# Finding the best cell lines across pan-cancer to use in pre-clinical research as a proxy for patient tumor samples considering immune cells, multi-omics, and cancer pathways

**DOI:** 10.1101/2022.12.18.520831

**Authors:** Banabithi Bose, Serdar Bozdag

## Abstract

In pre-clinical trials of anti-cancer drugs, cell lines are utilized as a model for patient tumor samples to understand the response of drugs. However, *in vitro* culture of cell lines, in general, alters the biology of the cell lines and likely gives rise to systematic differences from the tumor samples’ genomic profiles; hence the drug response of cell lines may deviate from actual patients’ drug response. In this study, we computed a similarity score for the selection of cell lines depicting the close and far resemblance to patient tumor samples in twenty-two different cancer types at genetic, genomic, and epigenetic levels integrating multi-omics datasets. We also considered the presence of immune cells in tumor samples and cancer-related biological pathways in this score which aids personalized medicine research in cancer. We showed that based on these similarity scores cell lines were able to recapitulate the drug response of patient tumor samples for several FDA-approved cancer drugs in multiple cancer types. Based on these scores, several of the high-rank cell lines were shown to have a close likeness to the corresponding tumor type in previously reported in vitro experiments.

## Introduction

Drug testing on human subjects in cancer research creates major logistical and ethical concerns. Multiple anti-cancer drug trials on a real cancer patient are extremely dangerous to the patient’s health and survival. As a result, scientists commonly use human tumor-derived cell lines as models for cancer tissues in pre-clinical trials to better understand drug efficacy [1]. The fundamental benefit of employing a cell line for research is that it is eternal. Cell lines are inexpensive, and they can be cloned and cultured for long periods of time, allowing an experiment to be repeated several times. *In vitro* culture of cancer cell lines, on the other hand, frequently results in genetic alterations, resulting in systematic phenotypic variations from patient tumor samples [2], [3]. Hence, pre-clinical drug response models based on cell lines must be carefully evaluated before being translated into a clinical setting [1], [2].

Owing to the rapid technological advancements in high throughput technology, massive quantities of genome, proteome, transcriptome, and epigenome datasets, known as multiple “omes” or multi-omics, are now available to researchers all over the world. Several studies generated multi-omics datasets, such as gene expression, mutation, copy number aberration (CNA), DNA methylation, and microRNA (miRNA) expression from the large set of biospecimens [3]–[5]. Among these, multi-omics profiles of tumor samples and cancer cell lines in the Cancer Genome Atlas (TCGA) [3] and Cancer Cell Line Encyclopedia (CCLE) [4] databases pave the way for a comparative study of the patient tumor samples with cancer cell lines. To this end, using gene expression data, Liu *et al*. [6] used a Transcriptome Correlation (TC) analysis to correlate cell lines and tumor samples. Sandberg *et al*. computed similarity scores, namely, Tissue Similarity Index (TSI) between a cell line and a patient cohort using Singular Value Decomposition (SVD) with gene expression of tumor samples and cell lines. Recently, Warren *et al*. developed an unsupervised alignment method called Celligner that mapped the gene expression of tumor samples to the gene expression of the cell lines. Their study found an alignment of the majority of cell lines with tumor samples of the same cancer type while revealing systematic differences in others.

All these studies relied on a genome-wide correlation-based approach between cell lines and bulk tumor samples without considering the heterogeneous immune cell population present in bulk tumors [9], [10] that play roles in targeted therapies in cancer [11]–[13]. Also, these studies considered single omics profiles, *i*.*e*., only gene expression data to relate patient tumor samples and cell lines, whereas several studies have shown that cell lines mirror many aspects of the multi-omics signatures identified in patient tumors [14], [15]. To date, studies leveraging the multi-omics profiles to find systematic differences and similarities between patient tumor samples and cancer cell lines are limited. Furthermore, no previous computational approach was developed to consider the critical role of the biological pathway activities in computing cell line tumor similarity scores [16] despite the knowledge that pathway activation status in cancer cell lines might connect tumor samples [17], [18]. To address these gaps, our group developed a computational pipeline, *CTDPathSim*, a biological pathway activity-based approach to computing the similarity between patient tumor samples and cell lines in breast and ovarian cancer [19]. In *CTDPathSim*, we integrated DNA methylation and gene expression datasets and computed similarity scores between patient tumor samples and cell lines. We showed that the similarity scores computed by *CTDPathSim* depicted the drug response phenotype between breast and ovarian tumor samples and cell lines. However, establishing these ratings in a study that covers the entire cancer spectrum is important for choosing the best cell lines for a variety of tumor types. Additionally, it is essential to take into account CNA data when calculating these scores in order to portray the intricacy of cancer and patient-specific treatment response patterns when using cancer cell lines as a stand-in for tumor samples.

Recent studies show that CNAs can serve as prognostic and predictive biomarkers of drug response among cancer patients [20]. Since cell lines and tumor samples can have different copy number profiles [20], [21], it is important to consider the copy number effect while establishing a more accurate similarity index between a patient tumor sample and a cell line.

To this end, in the current study, we propose cancer-related biological-pathway-motivated integration of CNA data with DNA methylation and gene expression in pan-cancer to provide a resource for selecting suitable cell lines as a surrogate for primary tumor samples to be used in *in vitro* experiments. We present *CTDPathSim2*.*0*, a novel and comprehensive computational pipeline that combines the three forms of omics data discussed above to compute tumor sample-cell line similarity scores throughout the whole cancer spectrum. These scores are more reliable than other existing methods of identifying cell lines that have similarities and differences with real tumor samples. We performed an extensive analysis of our pipeline on 22 different cancer types and computed similarity scores between 7,228 patient tumor samples from TCGA and 1,018 cancer cell lines from CCLE capturing a wide range of tumor types. Based on these scores, several of the high-rank cell lines were shown to closely resemble the correct tumor type in previously published *in vitro* tests. Our results show that *CTDPathSim2*.*0* outperformed the previously published tool, *CTDPathSim* (hereafter, *CTDPathSim1*.*0*), and other three state-of-the-art methods in capturing the drug response concordance between patient tumor samples and cell lines for several FDA-approved cancer drugs in several cancer types. Also, *CTDPathSim2*.*0* outperformed all these tools in the selection of cell lines belonging to the same tissue type in several cancer types. Furthermore, *CTDPathSim2*.*0* computed similarity scores between cell lines and patient tumor samples that could capture known cancer types and subtypes in patient tumor samples. For the use and replication of our technology, we presented *CTDPathSim2*.*0* as an R software package. In this study, our approach was tested on cancers, however, a similar methodology to this could be used to uncover cell lines replicating other diseases.

## Results

### Computing cell line-tumor similarity score using multi-omics data

Considering the current gap in the understanding of similarities and differences between primary tumors i.e., patient tumor samples and cancer cell lines, in this study, we integrated DNA methylation, gene expression, and CNA datasets to compute multi-omics-based similarity scores between patient tumor samples and cell lines. Specifically, we utilized 7,228 patient tumor samples in 22 different cancer types from TCGA and 1,018 cancer cell lines from CCLE and computed their pairwise similarity score. For TCGA, we provided the cohort sizes of each data modality of the 22 cancer types in Table 1. For CCLE, we retrieved gene expression of 1,019 cell lines, DNA methylation of 831 cell lines, and CNA of 916 cell lines belonging to these 22 cancer types. We followed a systematic data processing approach to ensure the quality of the TCGA multi-omics datasets (see Methods). To address the fact that the gene expression and DNA methylation signal from bulk tumors could be confounded by the presence of the heterogeneous cell population [9], [10], in the first two steps of *CTDPathSim2*.*0* (Fig.1), we applied a deconvolution algorithm using quadratic programming approach (see Methods) utilizing different immune cell types, namely, B cell (B), natural killer (NK), CD4T, CD8T, monocytes (MC), adipocytes (AC), cortical neurons (CN), and vascular endothelial (VE) and computed deconvoluted DNA methylation and expression profiles of each patient tumor sample in each cancer type (Table 1). We anticipated that these deconvoluted omics profiles of patient tumor samples would be able to capture the true biological signals. In the third and fourth steps of *CTDPathSim2*.*0* (Fig.1), considering the critical role of the biological pathway activities in cancer [16] and the previous studies that suggested that pathway activation status in cancer cell lines could connect tumor samples [17], [18], we computed enriched biological pathways for the patient tumor samples and cancer cell lines utilizing patient-specific and cell line-specific differentially expressed (DE), differentially methylated (DM) and differentially aberrated (DA) genes (see Methods). We excluded one cell line that had no enriched pathway from this analysis. In the final step (Fig.1), utilizing DE, DM, and DA genes, we computed Spearman correlation coefficients to get gene expression-, DNA methylation-, and CNA-based similarity scores for each tumor sample-cell line pair (see Methods). We scaled these three similarity measures to the range of 0 to 1 using min-max normalization and computed an average similarity score for each sample-cell line pair in each cancer type. Since we did not have DNA methylation and copy number data for all the cell lines, there were sample-cell line pairs that had only expression-based scores. We also computed scores for *CTDPathSim1*.*0* to compare with our new pipeline. For *CTDPathSim1*.*0*, we considered expression-based similarity and DNA methylation-based similarity to compute an average similarity score for each cell line-sample pair. We applied z-normalization on the final scores to set the mean score to 0 and the standard deviation to 1 in each TCGA cancer type. For running each of the steps of our pipeline, we provided our tool as an R software package. For the detailed pipeline with algorithms and R functions, please follow the Methods section of the manuscript.

**Fig. 1.**
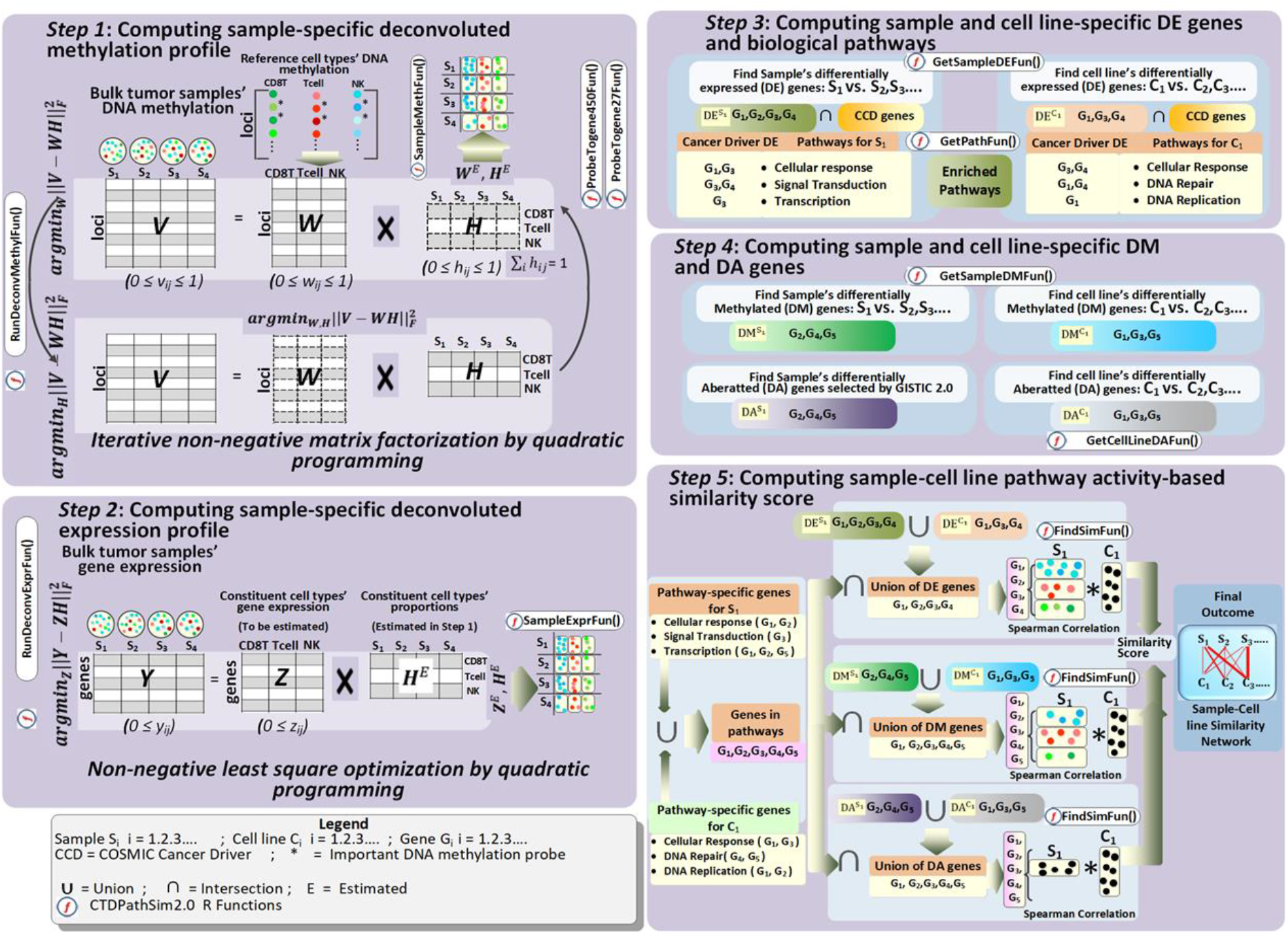
Flowchart of the CTDPathSim2.0 pipeline.

**Table 1.** Cohort sizes in different data modalities and cell types after deconvolution in 22 TCGA cancer types in the study.

| TCGA Cancer Type Name (Abbreviation) | Expr | Meth | CNA | Cell Types |
| --- | --- | --- | --- | --- |
| Adrenocortical carcinoma (ACC) | 79 | 79 | 77 | MC, VE, AC |
| Bladder Urothelial Carcinoma (BLCA) | 411 | 411 | 408 | CD4T, VE, AC |
| Breast invasive carcinoma (BRCA) | 1073 | 1073 | 1064 | CD4T, VE, MC |
| Cervical squamous cell carcinoma and endocervical adenocarcinoma (CESC) | 302 | 302 | 290 | CD4T, CN, VE, AC |
| Lymphoid Neoplasm Diffuse Large B-cell Lymphoma (DLBC) | 47 | 47 | 46 | CD4T, NK, B |
| Esophageal carcinoma (ESCA) | 160 | 160 | 159 | VE, CN, MC, AC, CD4T |
| Glioblastoma multiforme (GBM) | 120 | 120 | 144 | CN, AC, MC |
| Head and Neck squamous cell carcinoma (HNSC) | 495 | 495 | 490 | AC, CD4T, VE |
| Kidney renal clear cell carcinoma (KIRC) | 490 | 293 | 486 | CD4T, VE, AC |
| Acute Myeloid Leukemia (LAML) | 133 | 133 | 131 | NK, NP, MC, B |
| Brain Lower Grade Glioma (LGG) | 492 | 492 | 491 | VE, CN, MC, AC |
| Liver hepatocellular carcinoma (LIHC) | 371 | 371 | 369 | CD4T, CN, AC, VE |
| Lung adenocarcinoma (LUAD) | 508 | 508 | 509 | AC, VE, CD4T, CN |
| Lung squamous cell carcinoma (LUSC) | 491 | 491 | 490 | AC, CD4T, CN, VE |
| Mesothelioma (MESO) | 85 | 85 | 85 | VE, AC, CD4T |
| Ovarian serous cystadenocarcinoma (OV) | 368 | 594 | 358 | CN, VE, AC, MC |
| Pancreatic adenocarcinoma (PAAD) | 146 | 146 | 146 | VE, CD4T, CN |
| Prostate adenocarcinoma (PRAD) | 400 | 400 | 397 | CD4T, VE, AC |
| Skin Cutaneous Melanoma (SKCM) | 103 | 103 | 103 | CD4T, AC, CE |
| Stomach adenocarcinoma (STAD) | 369 | 369 | 368 | VE, AC, CD4T |
| Thyroid carcinoma (THCA) | 468 | 468 | 466 | CN, CD4T, AC |
| Uterine Corpus Endometrial Carcinoma (UCEC) | 546 | 546 | 538 | CD4T, CN, AC |

Note: 'Expr' stands for gene expression; 'Meth' stands for DNA methylation; 'CNA' stands for copy number alteration. Cell types are the different cell populations such as B cell (B), natural killer (NK), CD4T, monocytes (MC), adipocytes (AC), cortical neurons (CN), and vascular endothelial (VE) found to be present in bulk tumor DNA methylation data after applying deconvolution algorithm in step 1 of CTDPathSim2.0.

### CTDPathSim2.0 outperformed the existing methods for computing similarity scores that recapitulate the known drug responses between the tumor samples and the cell lines

The scores computed by *CTDPathSim2*.*0* were compared to the scores computed by *CTDPathSim1*.*0* and three state-of-the-art methods, namely TSI, TC analysis, and Celligner (see Methods). The drug response concordance between TCGA samples and CCLE cell lines was compared to see if the sample-cell line similarity scores obtained by these five approaches were concordant with drug response. We examined only those drugs and similarity scores that fulfilled the four filtering criteria given in Methods, using z-normalized scores for all the methods (Fig. 2). Except for Celligner (which had no filtered drug for ACC), we retrieved 24 total drugs in 19 distinct cancer types common to TCGA samples and CCLE cell lines using these filters on the generated scores by the various approaches (see Supplemental Table S1). Fig. 3 displays the conditional density plots for known drug responses vs. similarity scores of five different drugs from five methodologies in five cancer types. *CTDPathSim2*.*0* shows the drug response concordance (i.e., the matching percentage of the response of a drug increases as the similarity score increases) for all these five drugs in all five cancer types. The conditional density plots for all other drugs in 19 cancer types for the five methods with drug response trend summary are in Supplemental Figure S1. There were 86 different conditional density plots for 24 different drugs in 19 different cancer types, with the concordance trend obvious in 42, 32, 23, 19, and 22 plots for *CTDPathSim2*.*0, CTDPathSim1*.*0*, TSI method, TC analysis, and Celligner, respectively. These results indicate that similarity scores computed by our method align with known drug responses between the tumor samples and the cell lines.

**Fig. 2.**
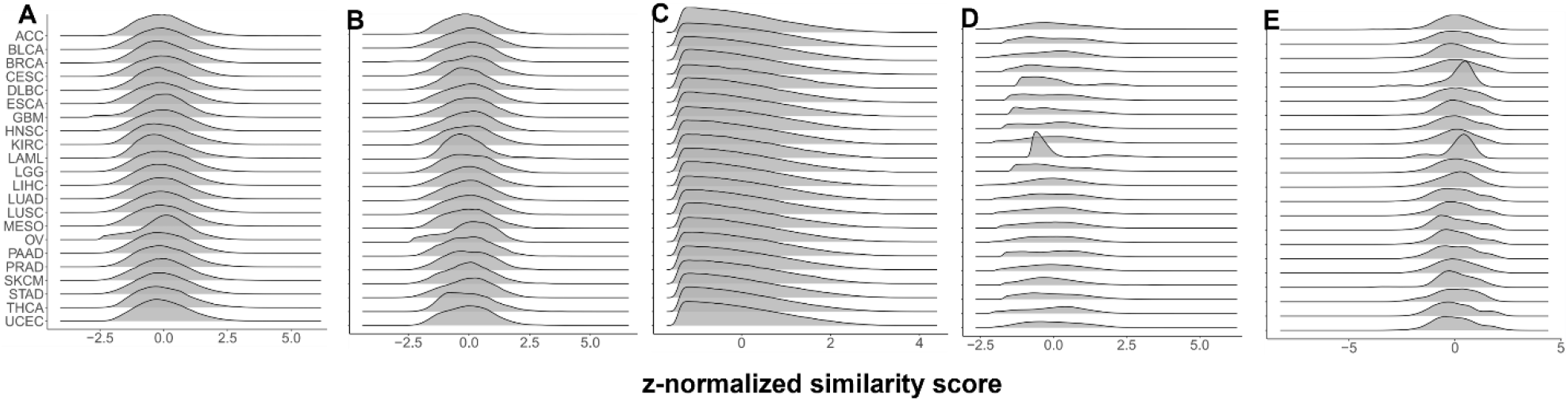
Computed similarity scores in different cancer types by different methods. Z-normalized scores between CCLE cell lines and TCGA samples of 22 different cancer types computed by A) CTDPathSim2.0, B) CTDPathSim1.0, C) TSI, D)TC analysis and E) Celligner.

**Fig. 3.**
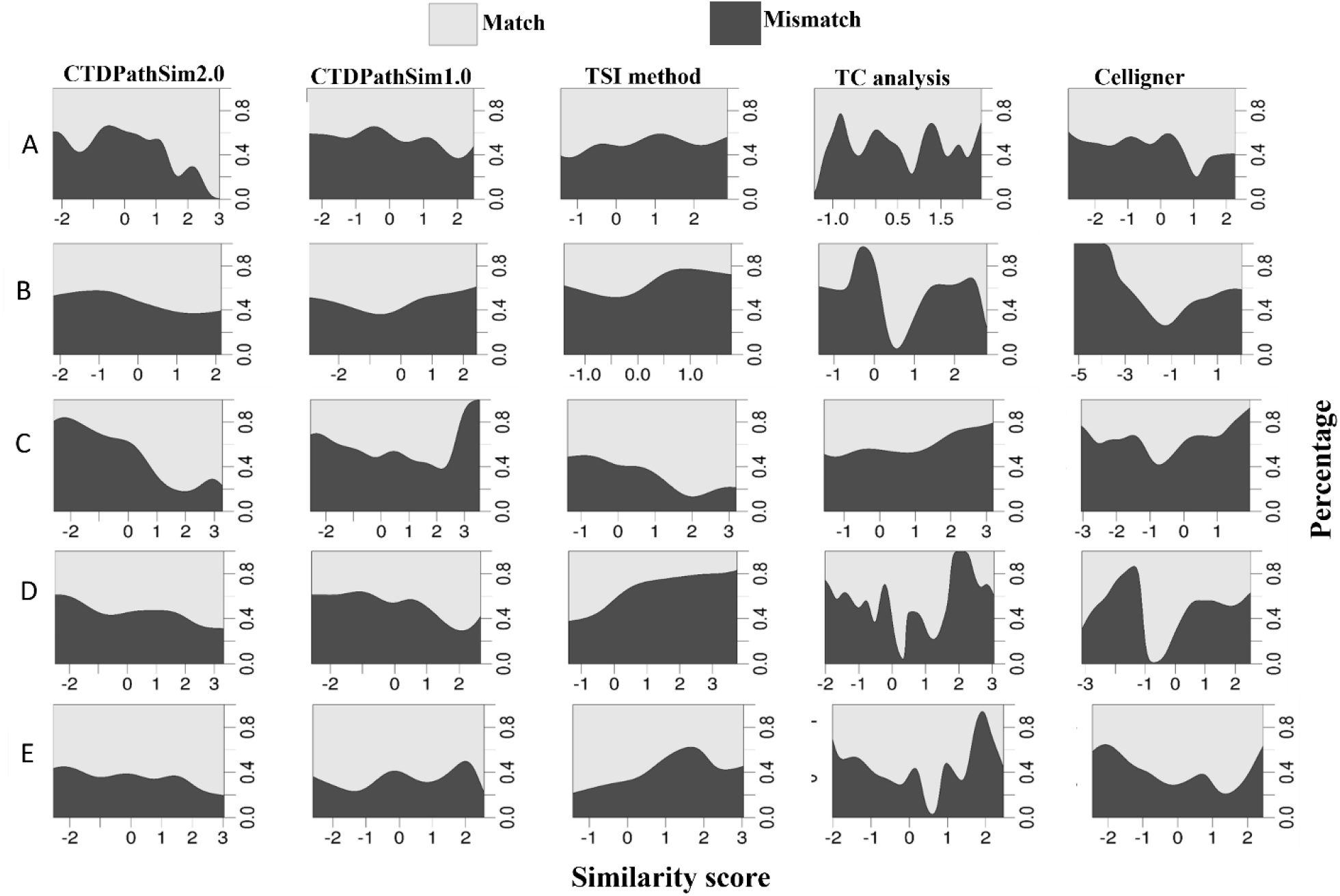
Drug response concordance with sample-cell line similarity scores. Conditional density plots of match and mismatch rate of drug response between TCGA samples and CCLE cell lines with computed z-normalized similarity scores by five different methods: A) Gemcitabine in BLCA, B) Doxorubicin in GBM, C) 5-Fluoruoracil in ESCA, D) Paclitaxel in LUSC and E) Erlotinib in LUAD. The matching percentage of the response of a drug increases as the similarity score increases in CTDPathSim2.0 in all these cancer types

Based on the aforementioned conditional density plots, we derived a drug response concordance score in each cancer type (i.e., micro concordance score) for each method to summarize its performance. We examined drug response concordance using computed similarity scores in high and low similarities, using the top 20 and bottom 20 percentiles of the scores, respectively. We discovered that these thresholds were better at depicting conditional density plots of drug response matching and mismatch than other thresholds (see Supplemental File S1). Because of the skewed distribution of TSI scores (see Fig. 2B), to allow enough data points for our comparison, we used high and low similarity thresholds of above 0 and below 0, respectively. The drugs were labeled as “concordant” if the difference between response matching percentage in the top and bottom percentiles was ≥ 0.10, “discordant” if that was ≤ − 0.10, and “undecided” otherwise. In each cancer type, we computed two different concordance scores (i.e., micro concordance scores): (1) 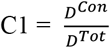 and (2) 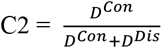; where *D*^*Tot*^ denotes the total number of drugs, *D*^*Con*^ denotes the number of concordant drugs and *D*^*Dis*^ denotes the number of discordant drugs. Table 2 displays the micro concordant scores for five different methods in 19 different cancer types. For *CTDPathSim2*.*0, CTDPathSim1*.*0*, TSI method, TC analysis, and Celligner, the number of cancer types with the highest micro concordant scores with C1 was eight, six, three, five, and four, respectively. Similarly, for *CTDPathSim2*.*0, CTDPathSim1*.*0*, TSI method, TC analysis, and Celligner, the number of cancer types with the highest micro concordant scores with C2 was eight, five, three, five, and four, respectively. We computed macro-averaged and weighted macro-averaged concordance scores using C1 and C2 independently to measure the overall effectiveness of the five different methods on pan-cancer (Table 3). We considered eighteen distinct cancer types omitting ACC in these scorings because there were no filtered drugs with computed Celligner scores in ACC. For each method, the macro-averaged concordance 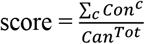, where *c* denotes the cancer type, *Con*^*c*^ denotes the micro concordance score (i.e., C1 or C2) in cancer type *c* and *Can*^*Tot*^ denotes the number of cancer types; the weighted macro-averaged concordance 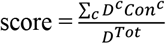, where *c* denotes the cancer type, *Con*^*c*^ denotes the micro concordance score (i.e., C1 or C2) in cancer type *c, D*^*Tot*^ denotes the total number of drugs across all cancer types and *D*^*c*^ denotes the number of drugs in cancer type *c*. In comparison to all other methods, *CTDPathSim2*.*0* achieved the greatest macro-averaged and weighted macro-averaged concordant scores with both C1 and C2. *CTDPathSim1*.*0* achieved the second-highest weighted macro-averaged concordance score with C1 and C2. *CTDPathSim1*.*0* outperformed TSI and TC analysis for macro-averaged with C1 and C2 as well. However, in terms of macro-averaged scores, Celligner slightly outscored *CTDPathSim1*.*0*. Overall, *CTDPathSim2*.*0* outperformed *CTDPathSim1*.*0* in terms of reproducing known drug response concordance between sample-cell line pairs; *CTDPathSim1*.*0* outperformed the TSI technique, TC analysis, and Celligner, which were all in the same ballpark.

**Table 2.**
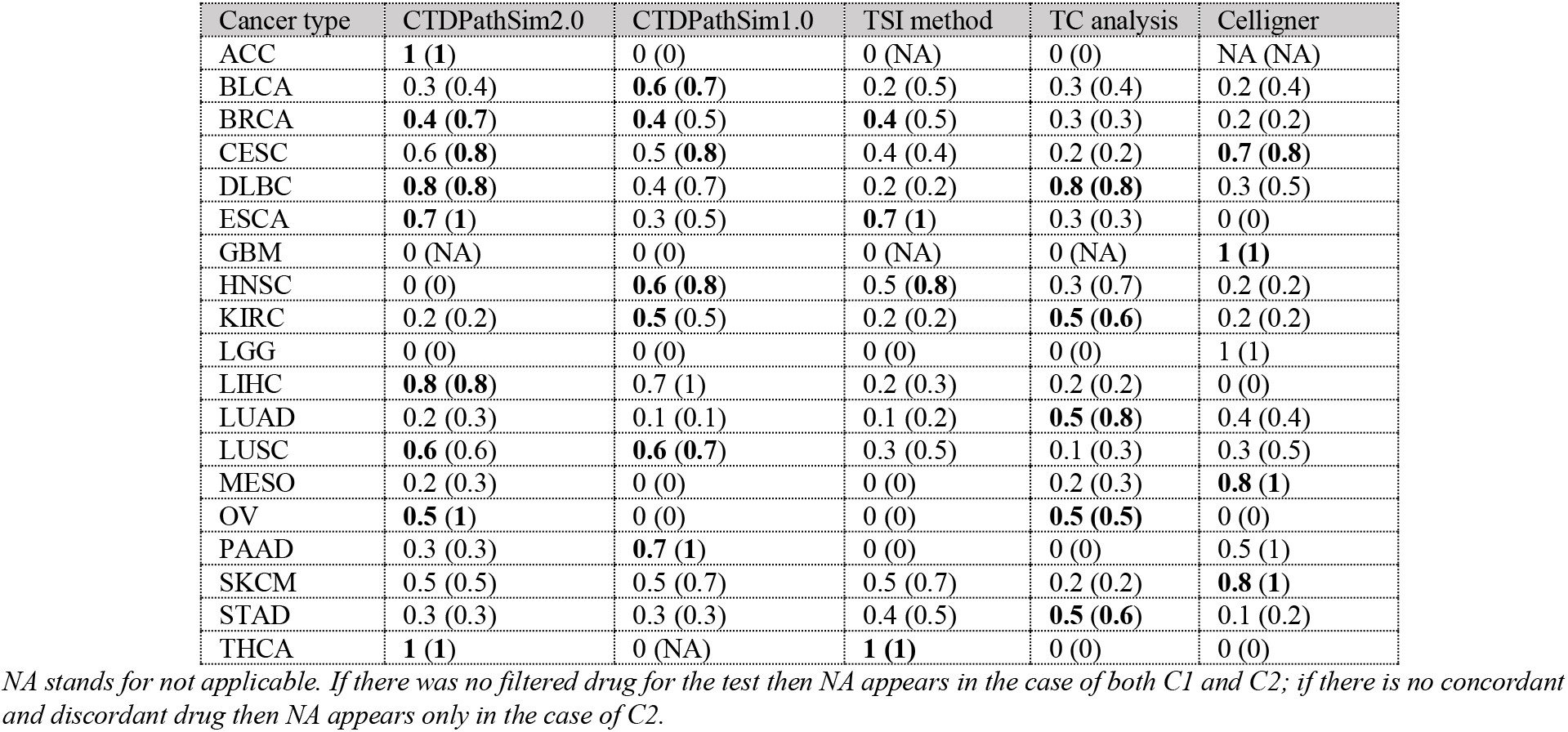
The micro concordant scores for five different methods in 19 different cancer types with C1 outside of the parenthesis and C2 inside the parenthesis. The highest values for each cancer type are shown in bold.

**Table 3.**
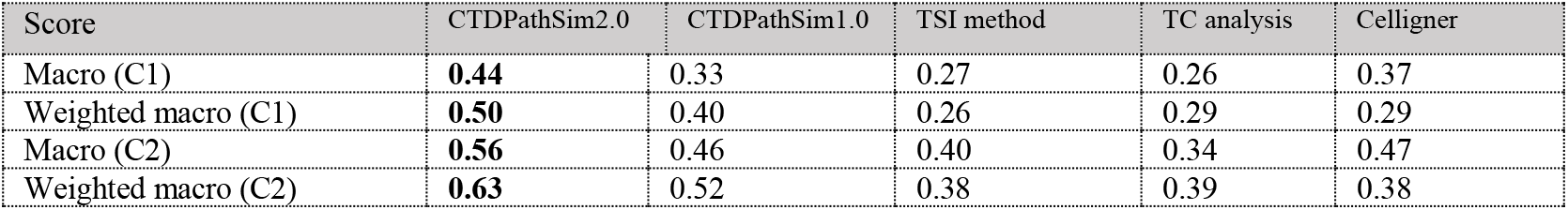
The macro-averaged and weighted macro-averaged concordant scores with C1 and C2. The highest values in each score are shown in bold.

### CTDPathSim2.0 outperformed state-of-the-art methods in the selection of significant cell lines belonging to the same tissue type in several cancers

We expected that the cell lines that have high similarity to a primary tumor sample would be significantly enriched in the cell lines derived from that tissue type, whereas low similarity cell lines would not have such enrichment. To test which similarity calculation method met this expectation, we picked CCLE cell lines that had high and low similarity to a primary tumor sample and performed enrichment of these CCLE cell lines in the cell lines derived from that tissue type in 21 different TCGA cancer types using all five methods. Since there were no Adrenocortical Carcinoma-derived cell lines in CCLE, we removed ACC samples from this analysis. We performed this evaluation using a high similarity threshold ≥ 1 and a low similarity threshold ≤ -1 for the sample-cell line pairs for all the methods, except for TSI method. These thresholds were chosen to include a sufficient number of sample-cell line pairs for all different methods to run this analysis. Because of the skewed distribution of TSI scores (see Fig. 2B), to allow enough data points for our comparison, we used high and low similarity thresholds of ≥0 and ≤0, respectively. In these thresholds, we computed highly and lowly similar cell lines for each patient tumor sample and checked if the cell lines for each patient tumor sample had highly overlapping tissue-specific cell lines by running hypergeometric test for each patient tumor sample. For instance, we checked if the highly similar cell lines to LUSC samples had significantly overlapping lung cancer cell lines. We found that highly similar cell lines to primary tissue samples computed by *CTDPathSim2*.*0* had significantly lower hypergeometric p-values than hypergeometric p-values for lowly similar cell lines. Fig. 4 shows the distribution of hypergeometric p-values (i.e., enrichment density plot) for corresponding tissue-specific cancer cell lines in high and low similarity thresholds for all five methods in GBM, UCEC, KIRC, LUAD, and LUSC samples in panels A-E, respectively. In *CTDPathSim2*.*0*, for all five cancer types (Fig. 4.; Column 1), the tissue-specific cell lines were picked with significant hypergeometric p-values (< 0.05) in high similarity thresholds compared to low similarity thresholds and hypergeometric p-values were separated within two groups. *CTDPathSim1*.*0* was also able to separate the hypergeometric p-values within two groups in four cancer types (Fig. 4.; Column 2) except GBM. The TSI method (Fig. 4.; Column 3) produced highly overlapping p-values in all five cancer types. For the Celligner (Fig. 4.; Column 5) significant p-values appeared to be present in low similarity groups with higher frequencies than in high similarity groups. TC analysis (Fig. 4.; Column 4) performed well in separating the two groups in four cancer types except GBM, however, there are several insignificant hypergeometric p-values (> 0.05) in high similarity group in KIRC. The enrichment density plots for all cancer types with different thresholds can be accessed from Supplemental Figure S2.

**Fig. 4.**
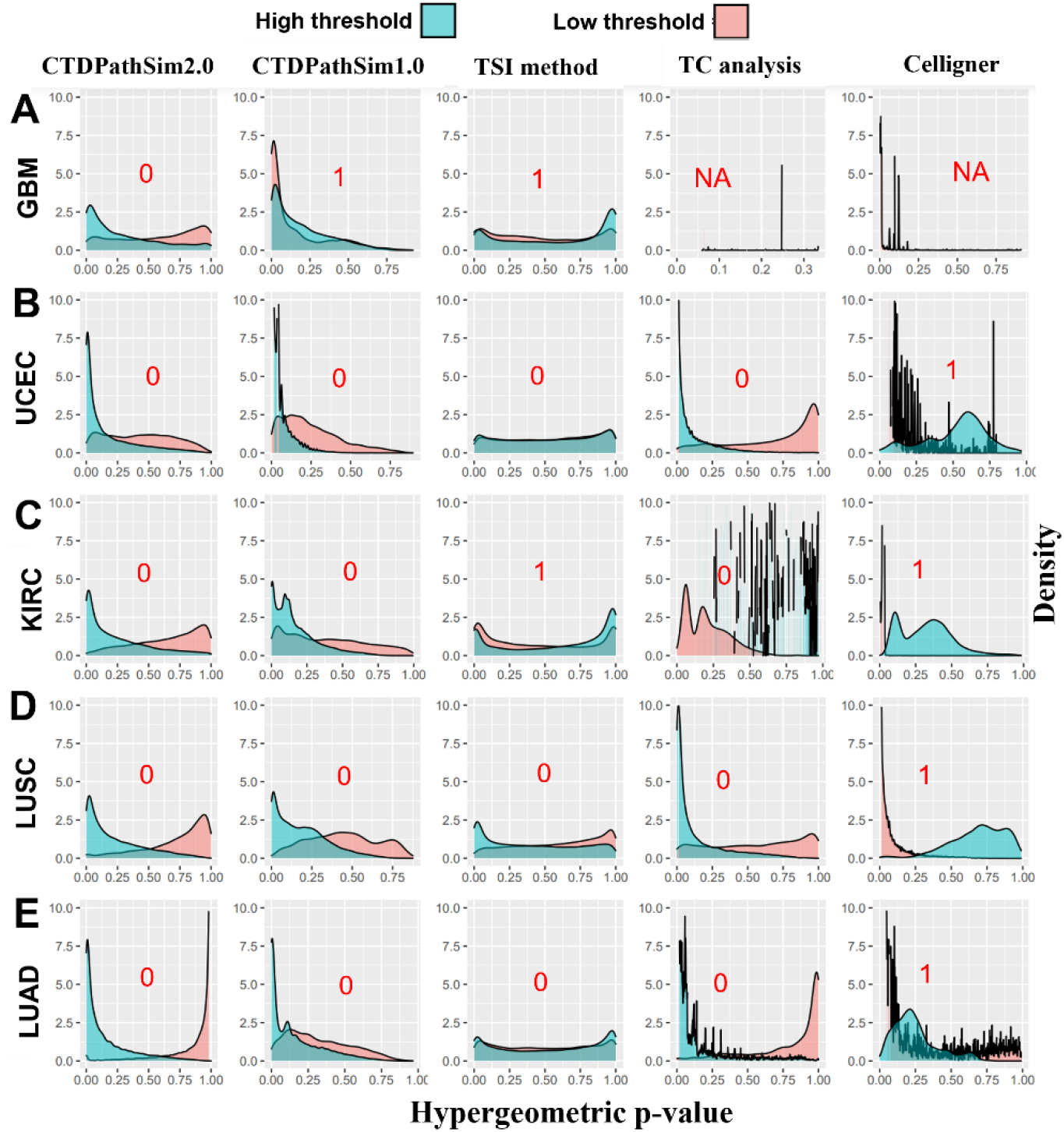
Density plots of hypergeometric p-values of enrichment for known tissue-specific cancer cell lines for the five methods. Enrichment of highly (cyan) and lowly (red) similar cell lines in A) GBM-specific cell lines; B) UCEC-specific cell lines; C) KIRC-specific cell lines; D) LUAD-specific cell lines and E) LUSC-specific cell lines. The t-test (alternative = “less”) p-values comparing each pair of the density plots in each cancer type are shown in red. NA stands for not applicable in cases where a t-test could not be run due to insufficient data points between two groups.

We compared the density plots of hypergeometric p-values for the enrichment of known tissue-specific cancer cell lines in 21 cancer types for five different methods using a one-sided t-test (Table 4). The significant t-test (alternative = “less”) p-value (p-value < 0.05) shows that the enrichment density plot in the high similarity group has a lower mean compared to the low similarity group. There were 20, 17, 14, 16, and 16 distinct cancer types with significant t-test p-values for *CTDPathSim2*.*0, CTDPathSim1*.*0*, TSI method, TC analysis, and Celligner, respectively. Overall, *CTDPathSim2*.*0* outscored all other methods in this test, with *CTDPathSim1*.*0* coming in second, the TC analysis and Celligner being comparable, and the TSI approach coming in last.

**Table 4.** One-sided t-test p-values for comparing the density plots of hypergeometric p-values for the enrichment of known tissue-specific cancer cell lines in 21 cancer types for five different methods.

| Cancer type | CTDPathSim2.0 | CTDPathSim1.0 | TSI method | TC analysis | Celligner |
| --- | --- | --- | --- | --- | --- |
| BLCA | 0 | 0 | 0 | 0 | 1 |
| BRCA | 0 | 0 | 0 | 0 | 1 |
| CESC | NA | NA | 6.7e-75 | NA | NA |
| DLBC | 6.2e-202 | 0 | 1 | NA | NA |
| ESCA | 8.9e-248 | 0 | 1 | 2.3e-02 | NA |
| GBM | 0 | 1 | 1 | NA | NA |
| HNSC | 0 | 0 | 0 | 6.1e-06 | NA |
| KIRC | 0 | 0 | 1 | 0 | 1 |
| LAML | 3.5e-18 | NA | 1 | NA | NA |
| LGG | 0 | 0 | 9.9e-01 | NA | 1 |
| LIHC | 1.1e-123 | 0 | 1 | 0 | 1 |
| LUAD | 0 | 0 | 2.6e-28 | 0 | 1 |
| LUSC | 0 | 0 | 0 | 0 | 1 |
| MESO | 4.8e-81 | 7.3e-166 | 3.8e-21 | 4.1e-17 | NA |
| OV | 0 | 0 | 0 | 0 | 1 |
| PAAD | 0 | 0 | 0 | 0 | 0.98 |
| PRAD | 4.1e-52 | 0 | 0 | 1.8e-258 | 1 |
| SKCM | 2.3e-115 | 1 | 3.1e-01 | 1.2e-86 | 1 |
| STAD | 1.5e-225 | 0 | 6.4e-116 | 0 | 1 |
| THCA | 1.3e-243 | 2.6e-06 | 3.9e-19 | 2.9e-22 | 1 |
| UCEC | 0 | 0 | 5.6e-47 | 0 | 1 |

Note: The significant t-test (alternative = “less”) p-value (p-value $< 0.05$ ) shows that the enrichment density plot (Fig. 4) in the high similarity group has a lower mean compared to the low similarity group. NA stands for not applicable in cases where a t-test could not be run due to insufficient data points between two groups.

### CTDPathSim2.0 tended to assign a high rank to tissue-specific cell lines among all the other cell lines in CCLE

In this analysis, for *CTDPathSim2*.*0*, we checked if the tissue-specific cell lines in each cancer type were in higher ranks among 1,018 cell lines from CCLE. We utilized the similarity scores computed in 21 different cancer types for which there was at least one tissue-specific cell line among 1,018 CCLE cell lines and computed the rank of 1,018 cell lines based on the median similarity values in each cancer type. In each cancer type, we additionally assigned an empirical p-value to these median rankings. For example, in BLCA, we had 25 BLCA-specific cell lines with a median rank of 710, hence, we picked 25 cell lines at random and computed their median rank to see if they had a better (i.e., higher) median rank than 710. We repeated this procedure 1000 and computed the empirical p-value as the percentage of cases where random cell lines had a better median rank. The number of tissue-specific cancer cell lines and their median rank in each cancer type are shown in Table 5 (here, the higher the rank better the result). Tissue-specific cell lines were shown to have a high rank (median of median rank in 21 cancer types was 738) among 1,018 CCLE cell lines for all of these cancer types. There were ten cancer types with an empirical p-value < 0.05 and sixteen cancer types with an empirical p-value ≤ 0.1. Fig. 5 displays several of the tissue-specific cell lines to be in higher rank among all the other cell lines in eight cancer types. The plots for all other cancer types can be accessed from Supplemental Figure S3. The rank of each tissue-specific cell line in 21 cancer types can be accessed from Supplemental Table S2.

**Table 5.** Number of tissue-specific cancer cell lines in 1,018 CCLE cell lines and their median rank with an empirical p-value in the parenthesis based on the median similarity values to the samples in each cancer type. Here, the higher the rank better the result.

| Cancer type | Cell line | Median Rank (p-value) | Cancer type | Cell line | Median Rank (p-value) |
| --- | --- | --- | --- | --- | --- |
| BLCA | 25 | 710 (0.02) | LIHC | 25 | 601 (0.16) |
| BRCA | 51 | 608 (0.08) | LUAD | 76 | 728 (0.00) |
| CESC | 3 | 827 (0.08) | LUSC | 23 | 663 (0.06) |
| DLBC | 39 | 985 (0.00) | MESO | 9 | 749 (0.06) |
| ESCA | 27 | 629 (0.11) | OV | 44 | 506 (0.53) |
| GBM | 33 | 655 (0.03) | PAAD | 40 | 684 (0.00) |
| HNSC | 33 | 744 (0.00) | PRAD | 8 | 680 (0.11) |
| KIRC | 33 | 785 (0.00) | SKCM | 54 | 677 (0.00) |
| LAML | 35 | 993 (0.00) | STAD | 37 | 573 (0.21) |
| LGG | 10 | 646 (0.17) | THCA | 11 | 577 (0.33) |
|  |  |  | UCEC | 28 | 807 (0.00) |

**Fig. 5.**
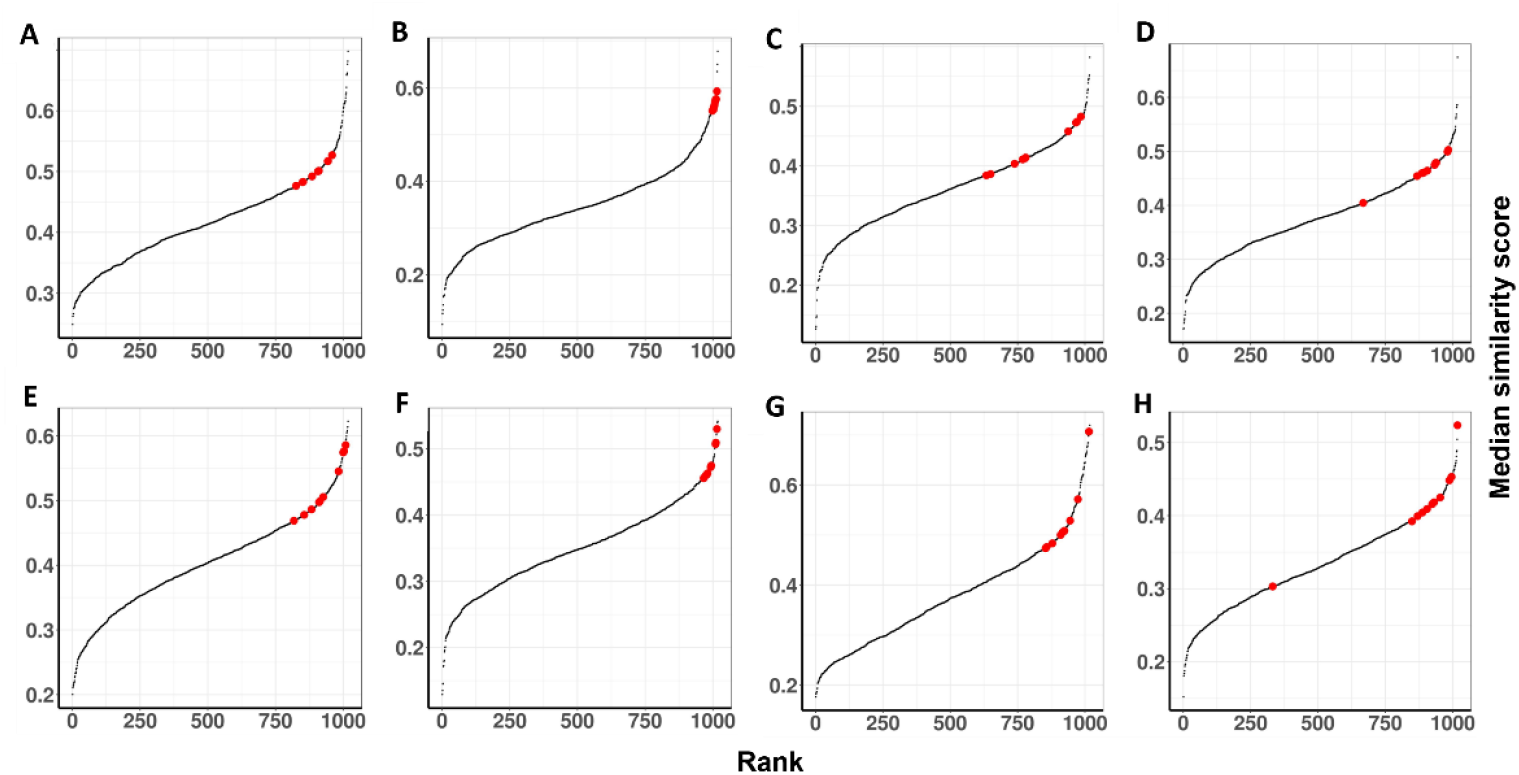
The rank of all 1,018 CCLE cell lines based on median similarity scores with all samples in the corresponding cancer type computed by CTDPathSim2.0 in eight cancer types. The rank of all cancer cell lines in A) BRCA; B) DLBC; C) GBM; D) LUAD; E) LIHC; F) LUSC; G) OV; H) UCEC. Each dot is a CCLE cell line and the few representatives of highly-ranked tissue-specific cell lines are marked by red.

### CTDPathSim2.0 computed similarity scores between cell lines and tumor samples were able to capture known cancer types and subtypes

To further evaluate *CTDPathSim2*.*0*, we tested whether the computed similarity scores between CCLE cell lines and primary tumor samples could detect known cancer types and subtypes present in the tumor samples. We used the cancer type and subtype annotation data from [8] focusing on situations where there were matched cancer type and subtype annotations across cell lines and tumor samples. We filtered annotation labels for which at least ten tumor samples were available in TCGA. To display the clusters of patient tumor samples based on their similarity scores with 1,018 cell lines, we used 7,223 tumor samples with 19 distinct cancer type labels and 26 distinct cancer subtype labels (Fig. 6A, 6C). We performed k-means clustering of the patient tumor samples using cell line-specific similarity scores and compared these clusters (Fig. 6B) with annotated cancer type groups (Fig. 6A) using Rand Index (RI) and Uniform Manifold Approximation and Projection (UMAP) [22] plots. To perform k-means, we used R base package Stats with k = 19 and all other parameters as default. The UMAP plot in Fig. 6B shows the clear separation between different cancer type-specific clusters. The computed RI between annotated cancer-type labels with 19 predicted clusters was 0.89. Similarly, for subtypes, we performed unsupervised k-means clustering of the tumor samples using cell line-specific similarity scores with k = 26 and compared these clusters (Fig. 6D) with annotated subtype clusters (Fig. 6C) using RI and UMAP. The UMAP plot in Fig. 6D shows the clear separation between different subtype-specific clusters. The computed RI between annotated subtype labels with 26 predicted clusters was 0.92 Furthermore, we ran the Random Forest classifier using cell lines as features with the computed similarity matrix to classify the patient tumor samples and assessed this classification based on the known cancer type and subtype labels. For this purpose, we used the R package *keras* [23] implementation of a Random Forest classifier with 75% of tumor samples with computed similarity scores for training and testing with 10-fold cross-validation. We utilized 25% of tumor samples with computed similarity scores to check the performance of the model. We achieved high classification accuracy as 0.97 for cancer type labels and 0.90 for subtype labels. The macro-averaged F1 scores for cancer type and subtype classification were 0.96 and 0.88, respectively. This analysis shows that the tumor samples belonging to the same cancer type and subtype were having similar score distribution across the cell lines. For the HER2-enriched subtype, 67% of samples were identified as “luminal A” and 22% as “basal” leading to “NA” as the F1 score for this subtype. A recent study [24] found that in HER2-enriched breast cancer, we can find all of the transcriptional subtypes of breast cancer, including the luminal A and basal-like subtypes, and the Her2-enriched subtype can have a distinctive transcriptional landscape independent of HER2 amplification. Hence, this “NA” value is not surprising. Also, 70% of “melanocytic” subtype samples were classified as “transitory” samples leading to “NA” as an F1 score for this subtype. It has been reported that the “transitory” and “melanocytic” subtypes are a refinement of the similar differentiated proliferative phenotype [25]. These findings establish the fact that *CTDPathSim2*.*0* computed the similarity scores which were able to find the phenotypic similarity among patient tumor samples while establishing their association with cell lines accurately. The computed similarity scores between patient tumor samples and cell lines can guide the refinement of the cancer type and subtype annotations. The cancer type and subtype classification evaluation metrics such as precision, recall, and F1 scores can be accessed from Supplemental Table S3.

**Fig. 6.**
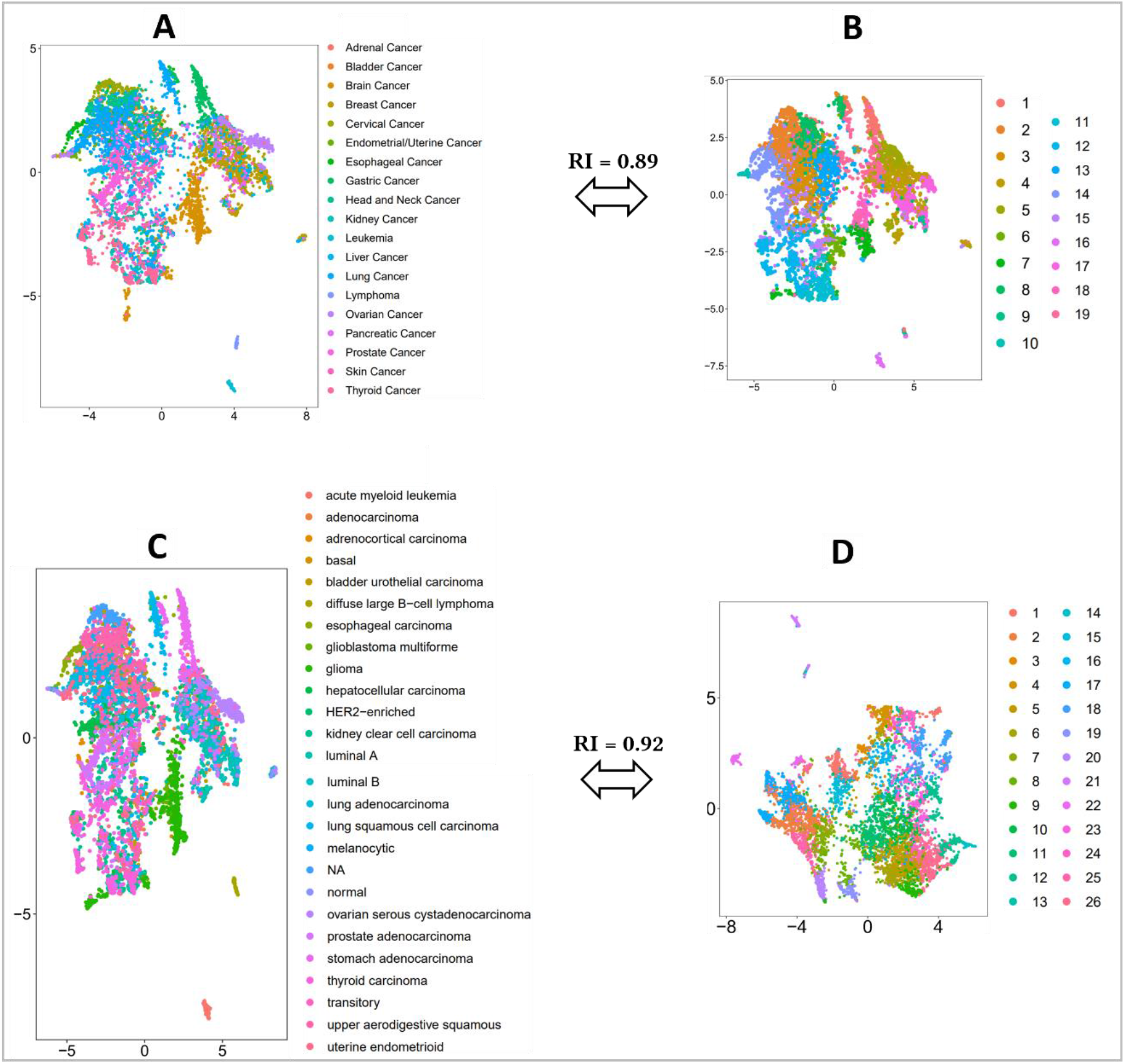
UMAP plots for visualizing annotated cancer type and subtype labels obtained from [30], k-means clustering for cancer type and subtype of TCGA samples based on CTDPathSim2.0 cell line-centric similarity scores. A) Cluster colors are showing 19 annotated cancer type labels; B) cluster colors are showing 19 clusters computed by k-means; C) Cluster colors show 26 annotated cancer subtype labels; D) cluster colors show 26 clusters computed by k-means.

## Discussion

We developed a computational pipeline, *CTDPathSim2*.*0*, that utilizes a pathway activity-based approach in computing the similarity between primary tumors and cell lines at genetic, genomic, and epigenetic levels. We applied *CTDPathSim2*.*0* on patient tumor samples from TCGA and cancer cell lines from CCLE for 22 different cancer types. We performed several tests to assess the similarity scores of sample-cell line pairs computed by *CTDPathSim2*.*0*, the previous pipeline, *CTDPathSim1*.*0*, and three other state-of-the-art methods. The computed similarity scores by *CTDPathSim2*.*0* were shown to be more successful than *CTDPathSim1*.*0* and the three state-of-the-art methods in capturing the drug response concordance between tumor samples and cell lines for several FDA-approved cancer drugs in several cancer types. Furthermore, in an enrichment test using CCLE cell lines from the same tissue type as TCGA tumor samples, *CTDPathSim2*.*0* outperformed all the methods.

Using a median similarity score computed by *CTDPathSim2*.*0* across the same tissue-specific tumor samples, we calculated the rank of tissue-specific CCLE cell lines. For example, each lung tissue-derived cell line with a median similarity score was taken for TCGA lung cancer (LUSC) samples. Table 6 shows the top five tissue-specific cell lines in fifteen different cancer types having more than ten tissue-specific cell lines. Several of the high-rank cell lines were shown to have a close likeness to the corresponding tumor type in previous *in vitro* experiments. For instance, the HCC1599 cell line was reported among highly resembling cell lines for the breast cancer samples with estrogen-positive, progesterone-positive, and human epidermal growth factor receptor 2-positive (HER2+) group [21]. A study reported CAOV3 as one of the most resembling cell lines for ovarian cancer [26]. DKMG cell line was found to have EGFR gene amplification, rearrangement, and expression, similar to GBM cancer samples [27]. In several studies, RAJI was a popular choice as a proxy for DLBC tumors [28], [29]. We also checked the tissue-specific cell lines that were low-ranked in the corresponding tumor samples. For DLBC, the REC-1 cell line had the lowest rank. An *in vitro* study found that the CD44 gene was hypermethylated and transcriptionally silenced in all DLBC cell lines, however, the same gene was found to be unmethylated and expressed in REC-1 cell lines [31]. We found that for BRCA, the UACC-893 cell line had the lowest rank. This cell line was reported to resemble the brain “metastasis” than the primary tumor in HER2+ breast cancer [32]. A study analyzed 57 breast cancer cell lines and found that the UACC-893 cell line was the only cell line that had the NAT2 gene’s expression higher than the NAT1 gene’s expression compared to other cell lines [33]. The same study reported the KPL-1 breast cancer cell line to be an MCF-7 derivative. Interestingly, our method ranks MCF7 and KPL-1 as 18 and 17, respectively among all 51 breast cancer cell lines used in the study. These findings point toward the ability of *CTDPathSim2*.*0* for ranking the cell lines based on their genomic profiles concerning primary tumors. *CTDPathSim2*.*0* also generated similarity scores between cell lines and patient tumor samples, which were able to capture known cancer types and subtypes, and these similarity scores can be used to refine current cancer and subtype classifications in tumor samples. Our tool enables the discovery of cell line models that best replicate the multi-omics properties of a tumor type, or even a specific tumor sample of interest, using biological pathway activation status. We anticipate that a deeper knowledge of the commonalities between cancer cell lines and tumors will enable better model selection and translation of the understanding of drug response from preclinical models to clinical samples. Finally, we provided *CTDPathSim2*.*0* as an R package to run the program for ease of application and reproducibility.

**Table 6.**
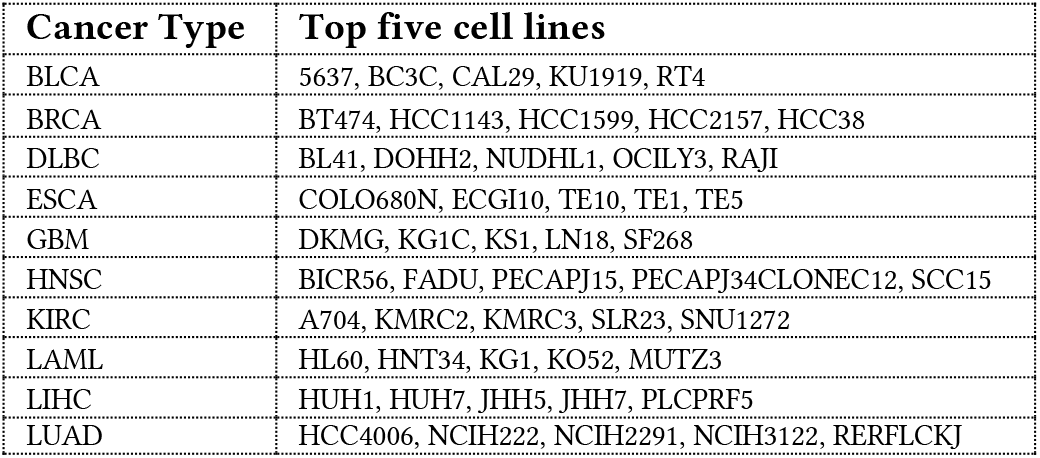
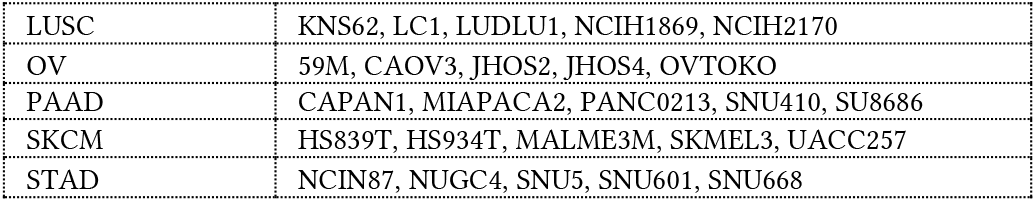
Five top-ranked tissue-specific cell lines for each cancer type.

## Methods

All the experiments were conducted in accordance with relevant guidelines and regulations.

### Datasets

#### Data preprocessing for running CTDPathSim2.0

In *CTDPathSim2*.*0*, we integrated DNA methylation, gene expression, and CNA datasets from 22 different cancer types. We downloaded the genomic data of cancer patient samples from the TCGA database. Using *TCGAbiolinks* [34], we downloaded RNA sequencing (RNA-Seq) data in HTSeq-FPKM format. We downloaded DNA methylation data from Infinium HumanMethylation27 Bead-Chip (27K) and Infinium HumanMethylation450 Bead-Chip (450K) platforms. We retrieved masked copy number variation (Affymetrix SNP Array 6.0) and computed the gene-centric copy number value compatible with hg38 genome assembly using R Bioconductor package *CNTools* [35]. From the COSMIC (Catalogue Of Somatic Mutation In Cancer) [36] cancer gene census project, we downloaded a list of 702 cancer-driving genes that are found to be frequently mutated in different cancer types.

We downloaded gene-centric copy number data for 1,040 cell lines from the CCLE database. We downloaded the CCLE_RNAseq_gene_rpkm _10180929.gct.gz file for gene expression for 1,019 cell lines and CCLE_RRBS_cgi_CpG_clusters_20181119 .txt.gz file for DNA methylation data of 831 cell lines. Using the R package *BioMethyl* [37], we removed CpG sites that had missing values in more than half of the samples and imputed the rest of the missing values by employing the *Enmix* [38] R package with default parameters.

To prepare the reference DNA methylation profiles for the deconvolution step, we processed raw methylation probes provided in *FlowSorted*.*Blood*.*450k* [39] R package that consists of DNA methylation probes from peripheral blood samples with five different cell types, namely, B cell (B), natural killer (NK), CD4T, CD8T, monocytes (MC), generated from adult men with replicates. We also processed raw methylation data available at NCBI Gene Expression Omnibus (GEO) database repository with the dataset identifier GSE122126 for three more cell types, namely, adipocytes (AC) (for 450K DNA methylation only), cortical neurons (CN), and vascular endothelial (VE) cells with replicates. We processed these raw methylation files using the R package *minfi* [40] to prepare our reference DNA methylation profiles of different cell types. Two different reference files, one for 450K probes and the other for 27K probes, with eight different cell types were prepared.

#### Data preprocessing for evaluating CTDPathSim2.0’s results

To compare the drug responses of patient tumor sample-cell line pairs, we compiled our ground truth drug response data for the patients from the TCGA database using Overall Survival (OS) and Progression-Free Survival (PFI) as clinical endpoints. These survival endpoints were considered representative of the drug response status of the patients. In OS, the patients who were dead from any cause were considered dead, otherwise censored. In PFI, the patients having a new tumor event whether it was a progression of the disease, local recurrence, distant metastasis, new primary tumor event, or died with cancer without a new tumor event, including cases with a new tumor event whose type is N/A were considered as the event occurred and all other patients were censored. We considered survival status 0 (i.e., censored), representative of sensitive drug response whereas survival status 1 (i.e., dead/event occurred), representative of resistant drug response as we mapped the patients’ survival status to drug response. In order to determine if PFI or OS status can be used as a drug response metric, we looked at the time intervals between drug applications. The following five rules were used to map the patients’ survival status to drug response (the details of these rules can be accessed from Supplemental File 1):

|  |  |
| --- | --- |
| *Drug response = OS status* | *; if PFI status = 0 and PFI.time = OS.time* |
| *Drug response = PFI status* | *; if PFI status = 1 and PFI.time = OS.time* |
| *Drug response = PFI status* | *; if drug application end days ≤ PFI days and PFI status = 1 with PFI.time < OS.time* |
| *Drug response = OS status* | *; if drug application end days ≤ PFI days and PFI status = 0 with PFI.time < OS.time* |
| *Drug response = OS status* | *; if drug application start days ≥ PFI days* |

For cell lines’ drug response data, we considered *IC*_50_ (the concentration of a drug that is required for 50% inhibition *in vitro*) values as drug response quantification. Furthermore, *ln* (*IC*_50_) < −2.0 was used to define positive (i.e., sensitive) anti-cancer drug response, and other *ln* (*IC*_50_) values were considered as negative (i.e., resistant) drug responses. Since common drugs between TCGA and CCLE were few (only twelve), we also added cancer cell lines drug response data with IC50 values from the Genomics of Drug Sensitivity in Cancer (GDSC) [41] database in our ground truth drug response data. We found 989 overlapping cell lines between the CCLE and the GDSC databases. We processed IC50 values of the GDSC drug similarly as we did for the cell lines in the CCLE. We retrieved 24 common drugs between TCGA patients of 19 different cancer types and cell lines from CCLE and GDSC after using the five filtering criteria mentioned below. A list of these drugs with the number of patient-cell line pairs, number of resistant and sensitive patients and cell lines can be accessed from Supplemental Table S1.

To compare the drug responses between sample-cell line pairs according to the computed similarity scores, we used z-normalized scores and considered only those drugs and similarity scores that passed the following filtering criteria to be eligible for the drug response concordance test between sample-cell line pairs.

##### 1. Filtering the drugs based on match-mismatch ratio

To check if a drug has a consistent response between sample-cell line pairs in the ground truth data, we computed a *match-mismatch* ratio, i.e., the ratio between the number of matched (i.e., same drug response) sample-cell line pairs to the number of mismatches (i.e., different drug response) sample-cell line pairs. The drugs that had a match-mismatch ratio < 0.2 were eliminated as even in the ground truth data, the majority of sample-cell line pairs have discordant drug responses.

##### 2. Filtering the drugs based on similarity thresholds

We considered the drugs for which the sample-cell line pairs had similarity thresholds at least above or below 1 and -1, respectively to allow enough data points in high and low similarity sample-cell line groups.

##### 3. Filtering the computed similarity scores based on hypergeometric p-value

Since we computed millions of sample-cell line similarity scores, we had discrete similarity scores, each with multiple sample-cell line pairs. For a specific drug, D, we computed the number of sample-cell line pairs for all similarity scores, namely, *Set 1*. Then, we computed the number of sample-cell line pairs for a specific discrete similarity score, K, namely, *Set 2*. We considered the discrete similarity scores for which there was a significant overlap between *Set 1* and *Set 2* (hypergeometric p-value < 0.05) to evaluate drug response match and mismatch between different sample-cell line pairs for drug D. The universe for this hypergeometric test was all the sample-cell lines pairs of the cancer type under study.

##### 4. Filtering the computed similarity scores based on match-mismatch difference

We filtered the similarity scores that showed at least a ten percent difference between match and mismatch drug response within sample-cell line pairs for considering these as confident scores for the drug response concordance test.

### Running CTDPathSim2.0

*CTDPathSim2*.*0* has five main computational steps. In the following paragraphs, we describe the *CTDPathSim2*.*0* R functions to run these steps. We performed these steps for each of 22 different cancer types separately. The entire pipeline of *CTDPathSim2*.*0* is illustrated in Fig. 1.

#### Step 1: Computing sample-specific deconvoluted DNA methylation profile

To capture the accurate DNA methylation signal of tumor samples, in the first step, we utilized a deconvolution-based algorithm [42] to infer the sample’s deconvoluted DNA methylation profile with proportions of different cell types in the sample’s bulk tumor tissue. The deconvolution algorithm requires a reference DNA methylation profile of different cell types. Hence, we compiled a list of reference DNA methylation profiles with eight different cell types, namely, B, NK, CD4T, CD8T, MC, AC, CN, and VE with replicates. We considered these cell types since a blood-based DNA methylation profile can depict the distribution of immune cell types [43], [44] and these cell types play roles in targeted therapies in cancer [11]–[13]. This algorithm computes a matrix with DNA methylation profiles of each of the constituent cell types (i.e., ***W*** = (*w*_*ij*_) where ***W*** ∈ ℝ^*l*×*c*^ ; *l* is the number of loci, *c* is the number of cell types, *i* = 1, 2,…., *l* and *j* = 1, 2,…., *c*.) and a matrix with the proportions of constituent cell types in each input tumor sample (i.e., ***H*** = (*h*_*ij*_) where ***H*** ∈ ℝ^*c*×*s*^ ; *c* is the number of cell types, *s* is the number of tumor samples, *i* = 1, 2,…., *c* and *j* = 1, 2,…., *s*.) minimizing the Euclidean distance between their linear combination (i.e., ***WH***) and the original matrix of samples’ DNA methylation profiles (i.e., ***V*** = (ν_*ij*_) where ***V*** ∈ ℝ^*l*×*s*^ ; *l* is the number of loci, *s* is the number of tumor samples, *i* = 1, 2,…., *l* and *j* = 1, 2,…., *s*.) (Fig. 1, Step1). We designed a function *RunDeconvMethylFun()* that ran a minimization algorithm in an iterative procedure that, in each round, alternates between estimating constituent cell-type proportions and DNA methylation profiles by solving constrained least-squares problems of 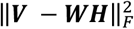 using quadratic programming method [42] with the constraints that 0 ≤ ν_*ij*_ ≤ 1, 0 ≤ *w*_*ij*_ ≤ 1 and ∑_*i*_ *h*_*ij*_ = 1. These three constraints for this problem were from the fact that DNA methylation measurements and cell type proportions were numbers in the range of 0 to 1 (hence, 0 ≤ ν_*ij*_ ≤ 1 and 0 ≤ *w*_*ij*_ ≤ 1), and cell type proportions within a sample added up to 1 (hence, ∑_*i*_ *h*_*ij*_ = 1) (Fig. 1, Step1). These constraints restrict the space of possible solutions, thus making it possible for the local iterative search to find a global minimum and an accurate solution reproducibly. To initiate the algorithm, we utilized the reference profiles as ***W*** with a set of selected 500 marker loci (probes) based on comparisons of each class of reference against all other samples using t-test. These loci were likely to exhibit variation in DNA methylation levels across different constituent cell types.

We designed a function, *SampleMethFun()* that utilized the estimated proportion of constituent cell types (i.e., ***H***^*E*^) and the estimated probe-centric DNA methylation profile of constituent cell types (i.e., ***W***^*E*^) from the output of *RunDeconvMethylFun()* to compute the sample-specific deconvoluted DNA methylation profile for each tumor sample (see Fig. 1, Step1). We ran this step for 450K DNA methylation probes (i.e., loci) and 27K DNA methylation probes separately. For each sample-specific deconvoluted DNA methylation profile, we computed gene-centric DNA methylation beta values from the probe-centric DNA methylation values with *ProbeToGeneFun()* function in our R package. For the 450K platform, the average beta value for promoter-specific probes was being considered due to their role in transcriptional silencing [45]. Given lower coverage in the 27K platform, the average beta value of all the probes of a gene was considered as the gene’s DNA methylation level.

#### Step 2: Computing sample-specific deconvoluted expression profile

In this step, we computed the deconvoluted expression profile for each tumor sample. We computed cell-type-specific expression profiles for the reference cell types utilized in Step 1 minimizing the objective function 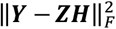 where ***Y*** *is* the original matrix of tumor samples’ expression profiles (i.e., ***Y*** = (*y*_*ij*_) where ***Y*** ∈ ℝ^*g*×*s*^, *g* is the number of genes, *s* is the number of samples, *i* = 1, 2,…., *g* and *j* = 1, 2,…., *s*.), and ***Z*** is the cell-type-specific expression profiles (i.e., ***Z*** = (*z*_*ij*_) where ***Z*** ∈ ℝ^*g*×*c*^, *g* is the number of genes, *c* is the number of cell types, *i* = 1, 2,…., *g* and *j* = 1, 2,…., *c*.) (Fig. 1, Step 2) to be estimated. We provided *RunDeconvExpr()* function in the package that utilized the estimated cell proportions (***H***^*E*^) from **Step 1** as a fixed input (i.e., ***H*** = ***H***^*E*^) and computed an average gene expression profiles of constituent cell types through a constrained least-squares of 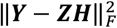 using quadratic programming with solutions constrained to [0,∞) for gene expression measurements [42]. Then, using estimated cell type-specific proportions in each sample (***H***^*E*^) and estimated cell type-specific expression (***Z***^*E*^) in *SampleExprFun()* function, we computed deconvoluted gene-centric expression profile of each tumor sample (Fig. 1, Step 2).

#### Step 3: Computing sample and cell line-specific DE genes and biological pathways

In this step, we computed enriched biological pathways listed in REACTOME [46] database for each sample and cell line using sample-specific DE genes. We designed *GetSampleDE()* function to compute the median expression of a gene across all samples and further computed the fold change of that gene to the computed median expression. A gene was considered up if the fold-change ≥ 4 and down if the fold-change ≤ −4. Likewise, using the *GetCellLineDEFun()* function and the similar thresholds, we computed cell line-specific DE genes. Considering the critical role of the biological pathway activities in cancer [16] and the previous studies that suggested that pathway activation status in cancer cell lines could connect tumor samples [17], [18], we identified enriched biological pathways for each sample and cell line using *GetPathFun()* function in our R package utilizing these DE genes. *GetPathFun()* found enriched biological pathways listed in the REACTOME database via a pathway enrichment tool *Pathfinder* [47]. To prioritize cancer-related biological pathways, specifically, we used the known frequently mutated genes that were considered cancer-driving genes from the DE gene list for this analysis. We obtained the frequently mutated cancer-driving genes from the COSMIC [36] database. *GetPath()* considered the pathways that were significantly enriched with FDR-corrected hypergeometric p-values < 0.05 in this DE cancer gene list.

#### Step 4: Computing sample and cell line-specific differentially methylated (DM) and differentially aberrated (DA) genes

In this step, we computed sample-specific and cell line-specific DM and DA genes. For sample-specific DM genes, we computed M-values [48] from gene-centric beta values and utilized *GetSampleDMFun()* function in the R package to compute the median of gene-centric M-values across all the samples in each cancer type. This function considered a gene hypermethylated if the M-value fold-change ≥ 4 and hypomethylated if fold-change ≤ -4. Likewise, using *GetCellLineDMFun()* with the same thresholds, we computed cell line-specific DM genes among all available cell lines in the study.

To compute sample-specific DA genes, we computed highly amplified and deleted genes utilizing chromosomal copy number profiles of each cancer type cohort as inputs in the GISTIC 2.0 [49] tool in the *GenePattern* [50] web server using a confidence interval of 0.90. We selected the genes that exceeded the high-level GISTIC thresholds for amplification and deletions as 2 and -2, respectively as DA genes.

For cell lines, we selected the highly amplified and deleted genes for each cell line compared with all other cell lines from various cancer types to capture the cancer-specific variation in copy number profiles in computing the similarity score between a patient sample and cell line. For this purpose, we utilized the *GetCellLineDA()* function, which selects cell line-specific DA genes based on Tukey’s mean-difference [51] curve based on gene-centric copy number data of cell lines. To determine whether a gene is DA for a specific cell line, we calculated the mean of that gene’s copy number values across all cell lines and the difference between the copy number value of that gene in the cell line under consideration and the mean of the copy number values of that gene in the other cell lines. For each cell line, the highly amplified genes were chosen based on mean ≥ 0.5 and difference ≤ 1, and highly deleted genes were chosen based on mean ≥ 0.5 and difference ≤ -1.

#### Step 5: Computing sample-cell line pathway activity-based similarity score

In this step, we computed Spearman rank correlation between each sample-cell line pair using the sample-specific deconvoluted expression, sample-specific deconvoluted DNA methylation, and sample’s gene-centric copy number values. All DM genes that occurred in an enriched pathway of a sample or cell line were used to compute the Spearman rank correlation between each sample-cell line pair via *FindSim()* function in the R package (Fig. 1, Step 5). Similarly, using DE and DA genes that were present in the gene list of the union of enriched pathways, we compute the Spearman rank correlation between each sample-cell line pair. Using DM and DE genes, we used the sample’s deconvoluted profiles to compute Spearman rank correlation, whereas for DA genes, we used the sample’s gene-centric copy number profile to compute Spearman rank correlation (Fig. 1, Step 5). We scaled the Spearman similarities to the range of 0 to 1 using min-max normalization and computed an average similarity score taking a mean of expression-based correlation, DNA methylation-based correlation, and copy number-based correlation for each sample-cell line pair. Since we did not have DNA methylation data and copy number data for all the cell lines, there were sample-cell line pairs that had only expression-based scores.

We presented the *CTDPathSim2*.*0* pipeline in this section, but our R package can be used to compute similarity scores without using CNA data to run the tool’s previous version, *CTDPathSim1*.*0*. In that case, the *FindSim*() function of the package can be used to run the pipeline without DA genes (please check the vignette of the *CTDPathSim2*.0 in the supplemental materials for these details).

### Running state-of-the-art-methods

We compared *CTDPathSim2*.*0* with our previous method *CTDPathSim1*.*0* and three other state-of-the-art methods, namely, TSI, TC analysis, and Celligner by running them on each of 22 different cancer type cohorts from TCGA and cancer cell line datasets from CCLE. For *CTDPathSim1*.*0*, we computed only DNA methylation-based and gene expression-based similarity scores and averaged them to get a unified similarity score for each sample-cell line pair in a cohort.

#### Running TSI method

For the TSI method, we performed z-normalization of RNA-Seq data for each gene in the samples and cell lines and performed all the required steps to compute tumor sample-cell line similarity scores based on the TSI paper [7]. We applied SVD on the normalized gene expression matrix of samples. Then each sample was projected into SVD space by measuring its correlation to the first 16 Eigenarrays (i.e., left singular vectors) that explain the most variance of the data. Similarly, we projected each cell line into SVD space. We computed a Pearson correlation score between each patient and cell line as a similarity score, namely, TSI score in the SVD space. We applied z-normalization on TSI scores computed between each cancer sample and cell line.

#### Running TC analysis

For TC analysis, we used 30,681 common genes between CCLE and TCGA RNA-Seq gene expression datasets with log2 transformation. Then, we rank-transformed gene RPKM (reads per kilobase of transcript per million reads mapped) values for each CCLE cell line and ranked all the genes according to their expression variation across all CCLE cell lines. The 1000 most variable genes were kept as “marker genes” to compute Spearman rank correlation, namely, TC scores, between each cell line and sample pair using the respective expression profiles.

#### Running Celligner

Considering Celligner’s global aligning approach of gene expression data between cell lines and tumor samples that depends on various cancer types and the composition of the dataset, we ran this pipeline as described in their paper utilizing the RNA-Seq data in TPM (transcript per million reads) of 12,236 tumor samples and 1,249 cell lines as log2(TPM + 1) transformation from Cancer Dependency Map [https://depmap.org] data portal for 37 different cancer types. We used the gene expression data without ‘non-coding RNA’ and ‘pseudogene’ as input in the Celligner pipeline. Celligner first performed unsupervised clustering of each dataset and removed the expression signatures with excess intra-cluster variance in the sample compared to cell line data. Then, using DE genes between clusters in the data it ran contrastive Principal Component Analysis (cPCA) followed by a batch correction method, MNN (mutual nearest neighbors) that aligned similar sample-cell line pairs to produce corrected gene expression data. We further filtered this gene expression matrix for 22 different TCGA cancer types utilized in the study and 1,015 CCLE cell lines that were common to *CTDPathSim2*.0. Then, we performed PCA on the Celligner-aligned data and took the Euclidean distance between each cell line and patient sample in PCA space (using the first 70 principal components following [30]) as a similarity measure.

## Supporting information

CTDPathSim2.0 R Vignette

README

## Data availability

The *CTDPathSim2*.*0* pipeline was developed as an R package. For installation, use *remotes::install_github(“boseb/CTDPathSim2*.*0”)*. The source codes of the package are available at *https://github.com/bozdaglab/CTDPathSim2.0* under Creative Commons Attribution Non-Commercial 4.0 International Public License. The vignette of the R package and the scripts for running the evaluation results can be accessed from the supplementary documents. The datasets can be accessed from Science Data Bank (https://doi.org/10.57760/sciencedb.01713).

## Acknowledgments

This work was supported by the National Institute of General Medical Sciences of the National Institutes of Health under Award Number R35GM133657.

## Author contributions

B.B and S.B conceived the study, B.B conducted the study, S.B supervised the study, B.B developed the software, B.B wrote the manuscript, B.B and S.B reviewed and edited the manuscript.

