## Supplemental Figure S1 for "Finding the best cell lines across pan-cancer to use in pre-clinical research as a proxy for patient tumor samples considering immune cells, multi-omics, and cancer pathways"

1(i.e., match)= 0(i.e., mismatch)=

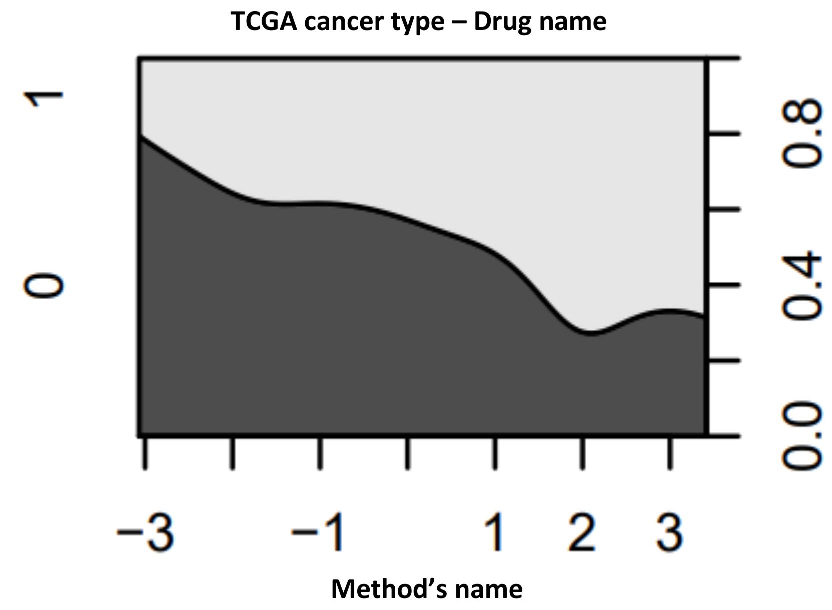


**Drug response concordance with sample-cell line similarity scores.** Conditional density plots of match and mismatch rate of drug response between TCGA samples and CCLE cell lines with computed z-normalized similarity scores by five different methods.

By visual inspection, we summed up the conditional density plots for known drug responses vs. similarity scores of five different methods in all cancer types. We label a drug as "concordant" if the matching percentage of the response increases as the similarity score increases; we label it as "discordant" if the matching percentage of the response decreases as the similarity score increases; if there is no increase or decrease, we label it as "linear," and all other trends are labeled as "undecided." We denoted a "concordant" trend for a drug in a method with a "Y," whereas all other trends were marked with a "N". There were 86 different conditional density plots for 24 different drugs in 19 different cancer types, with the concordance trend obvious in 42, 32, 23, 19, and 22 plots for CTDPathSim2.0, CTDPathSim1.0, TSI method, TC analysis, and Celligner, respectively.

**Table with visual summary of conditional density plots.** Summary of the conditional density plots for known drug responses vs. similarity scores of different drugs from five methodologies in all the cancer types by visual inspection. For each drug in the parenthesis, the "concordant" trend is denoted as "Y" and all other trends such as "discordant", "linear" and "undecided" are denoted as "N". NA stands for not applicable in case of missing conditional density plots for that drug due to filtering criteria provided in "Methods" section of the main manuscript.

| Cancer type (Drug name) | CTDPathSim2.0 | CTDPathSim1.0 | TSI method | TC analysis | Celligner |
|---|---|---|---|---|---|
| ACC (Doxorubicin) | Y | N | N | N | NA |
| BLCA (Cisplatin, Docetaxel,Doxorubicin,Etoposide,Gemcitabine,Methotrexate,Paclitaxel,Temsirolimus,Vinblastine) | N,Y,N,N,Y,N,N,Y,Y | N,N,N,NA,Y,N,N,Y,Y | N,Y,N,N,N,N,N,Y,N | N,N,N,N,N,N,N,N,N | N,NA,N,N,Y,N,N,N,N |
| BRCA (Doxorubicin, Gemcitabine,Lapatinib,Mitomycin-C,Vinorelbine) | Y,Y,N,Y,Y | N,N,N,N,Y | N,Y,Y,N,N | NA,N,NA,N,N | N,N,N,Y,N |
| CESC (5-Fluorouracil, Bleomycin, Doxorubicin, Gemcitabine, Mitomycin-C, Paclitaxel, Vinorelbine) | Y,N,Y,Y,Y,N,Y,Y | Y,Y,N,Y,NA,N,Y | N,N,N,N,NA,Y,NA | N,N,N,N,NA,Y,NA | Y,Y,N,N,N,Y,NA |
| DLBC (Bleomycin, Cisplatin, Doxorubicin, Gemcitabine, Methotrexate) | N,Y,Y,N,Y | Y,Y,Y,N,N | N,Y,Y,N,N | N,NA,Y,Y,Y | N,NA,NA,N,Y |
| ESCA(5- Fluorouracil, Gemcitabine, Paclitaxel) | Y,Y,N | N,Y,N | Y,N,Y | N,Y,N | N,Y,N |
| GBM (Doxorubicin) | Y | N | N | N | Y |
| HNSC (Cetuximab, Cisplatin,Docetaxel, Gemcitabine, Methotrexate, Paclitaxel | N,N,Y,N,N,N | NA,Y,N,Y,Y,N | Y,N,N,Y,Y,N | N,N,N,N,Y,N | N,Y,N,N,N,N |
| KIRC (Gemcitabine,Pazopanib,Rapamycin,Sorafinib,Sunitinib, Temsirolimus | Y,N,Y,N,N,N | Y,Y,Y,N,N,N | N,N,Y,NA,N,Y | N,Y,Y,N,N,Y | N,N,N,NA,,Y,N |
| LGG(Temozolomide) | N | N | N | N | Y |
| LIHC(Cisplatin, Doxorubicin, Gemcitabine, Mitomycin-C, Sorafenib, Temsirolimus) | Y,N,N,Y,Y,Y | Y,N,N,Y,Y,N | Y,N,N,NA,N,N | N.N.N.NA,N,N | N,NA,N,NA,N,NA |
| LUAD(Cisplatin, Docetaxel,Erlotinib, Etoposide,Gefitinib,Gemcitabine, Paclitaxel,Vinorelbine) | N,N,Y,Y,N,N,N | N,N,N,N,N,Y,N,N | N,Y,N,N,N,N,N,N | Y,N,N,Y,Y,N,Y,N | N,Y,N,N,Y,NN,Y |
| LUSC(Docetaxel, Doxorubicin, Erlotinib, Etoposide, Gemcitabine, Paclitaxel, Vinorelbine) | Y,N,Y,N,N,Y,Y | Y,Y,N,N,N,Y,Y | N,Y,N,N,N,N,Y | N,N,N,N,N,N,Y | Y,N,N,N,N,N,Y |
| MESO(Dasatinib, Doxorubicin, Gemcitabine, Temsirolimus) | Y,N,N,N | NA,N,N,N | N,N,N,N | Y,N,N,N | N,Y,Y,N |
| OV(Doxorubicin, Gemcitabine) | Y,N | N,N | N,N | Y,N | N,NA |
| PAAD(Doxorubicin, Gemcitabine, Paclitaxel) | N,N,Y | Y,N,Y | N,N,N | NA,N,N | NA,Y,N |
| SKCM(Cisplatin, Paclitaxel,Temozolomide,Trametinib) | Y,N,Y,N | Y,N,Y,N | Y,N,Y,N | Y,N,N,N | N,Y,N,Y |
| STAD(5- Fluorouracil, Cisplatin, Docetaxel, Doxorubicin, Etoposide, Mitomycin-C, Paclitaxel) | N,N,Y,N,N,N,Y | N,N,N,Y,N,N,Y | Y,N,N,Y,N,Y,N | N,N,Y,NA,N,Y,Y | Y,N,N,NA,N,N,N |
| THCA(Doxorubicin) | Y | N | Y | N | N |

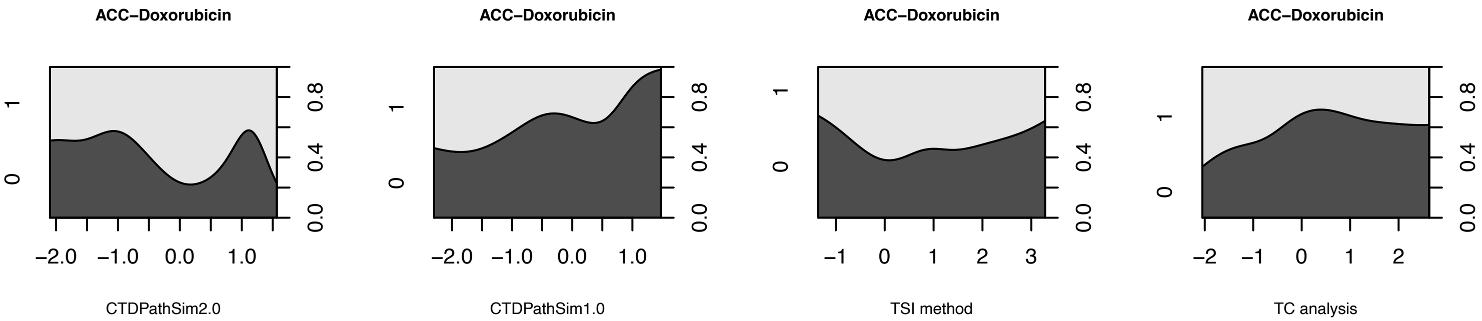
ACC−Doxorubicin
ACC−Doxorubicin
ACC−Doxorubicin
ACC−Doxorubicin
CTDPathSim2.0
CTDPathSim1.0
TSI method
TC analysis
0
1
0.0
0.4
0.8
−2.0
−1.0
0.0
1.0
−1
0
1
2
3
−2

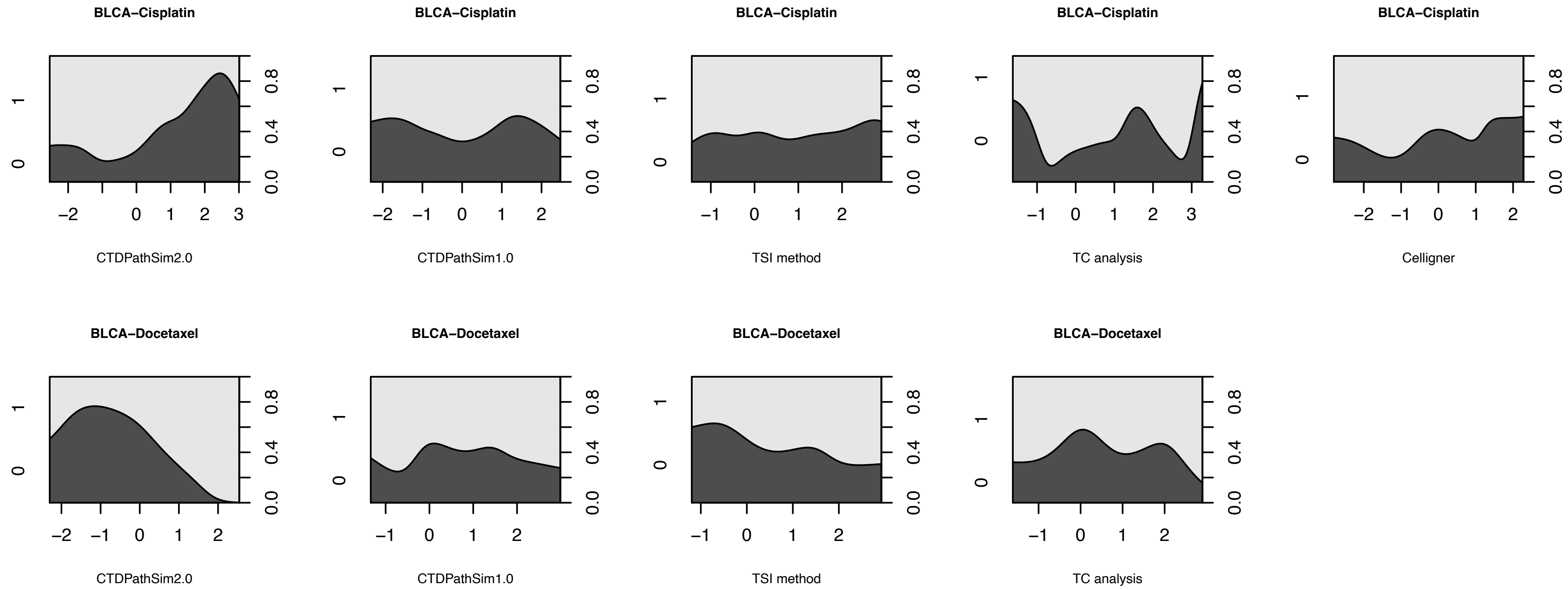
BLCA−Cisplatin
CTDPathSim2.0
BLCA−Cisplatin
CTDPathSim1.0
BLCA−Cisplatin
TSI method
BLCA−Cisplatin
TC analysis
BLCA−Cisplatin
Celligner
BLCA−Docetaxel
CTDPathSim2.0
BLCA−Docetaxel
CTDPathSim1.0
BLCA−Docetaxel
TSI method
BLCA−Docetaxel
TC analysis

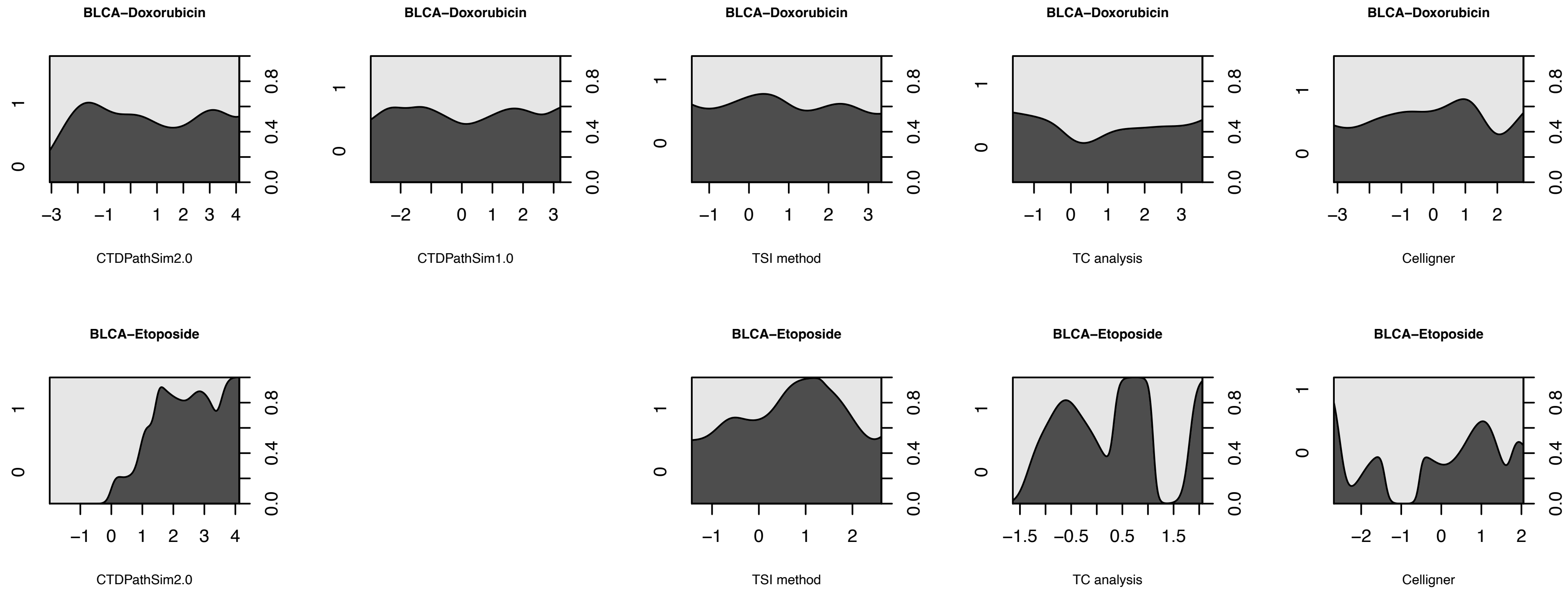
BLCA–Doxorubicin
CTDPathSim2.0
BLCA–Doxorubicin
CTDPathSim1.0
BLCA–Doxorubicin
TSI method
BLCA–Doxorubicin
TC analysis
BLCA–Doxorubicin
Celligner
BLCA–Etoposide
CTDPathSim2.0
BLCA–Etoposide
TSI method
BLCA–Etoposide
TC analysis
BLCA–Etoposide
Celligner

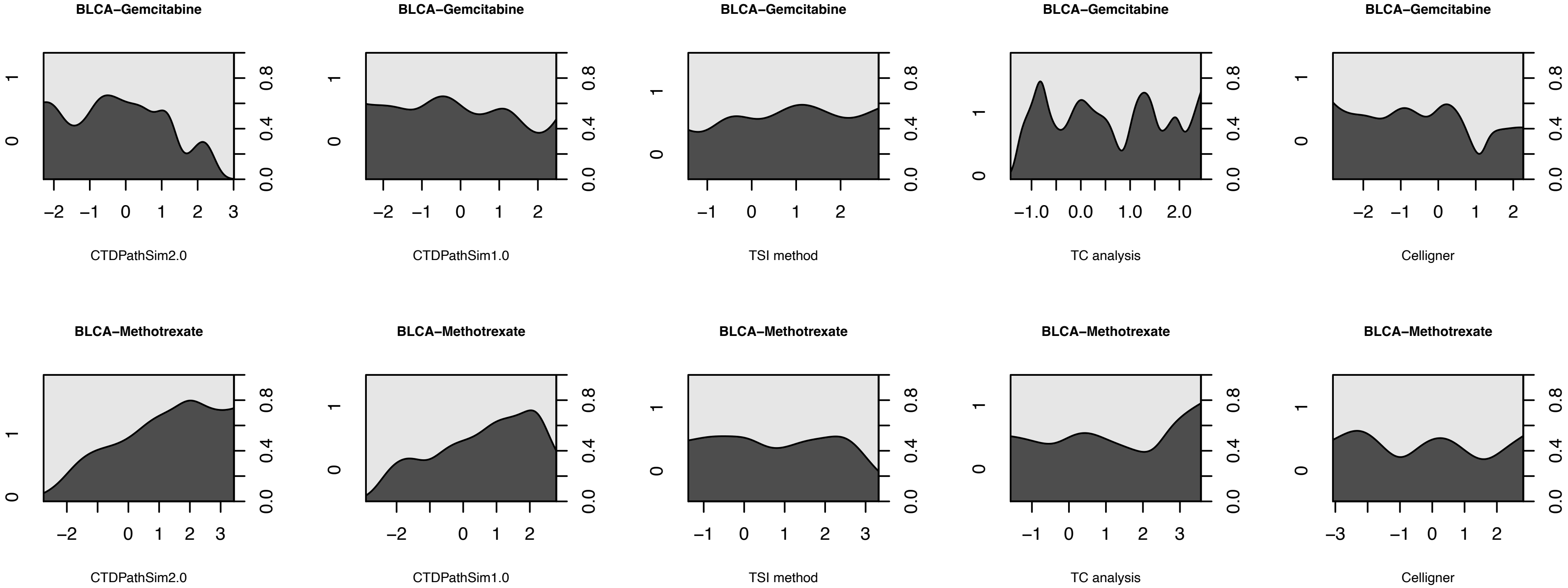
BLCA−Gemcitabine
CTDPathSim2.0
CTDPathSim1.0
TSI method
TC analysis
Celligner
BLCA−Methotrexate
CTDPathSim2.0
CTDPathSim1.0
TSI method
TC analysis
Celligner

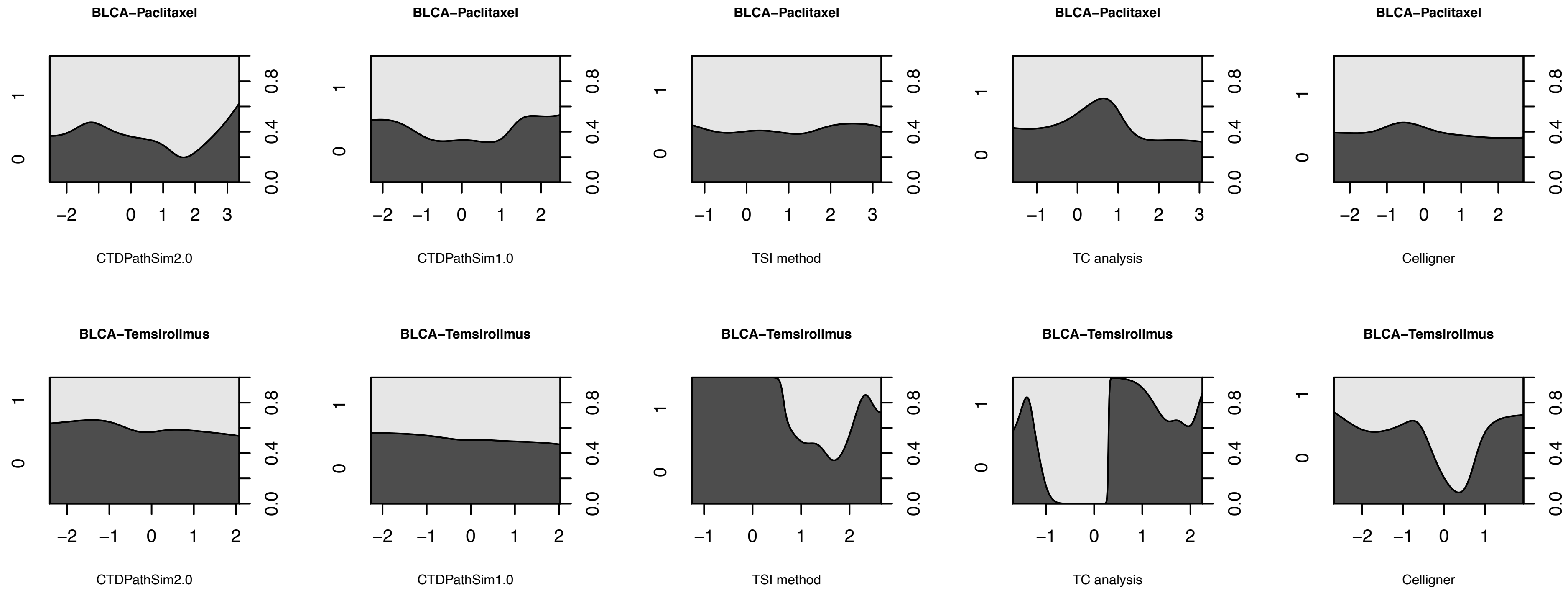

BLCA−Paclitaxel
CTDPathSim2.0
BLCA−Paclitaxel
CTDPathSim1.0
BLCA−Paclitaxel
TSI method
BLCA−Paclitaxel
TC analysis
BLCA−Paclitaxel
Celligner
BLCA−Temsirolimus
CTDPathSim2.0
BLCA−Temsirolimus
CTDPathSim1.0
BLCA−Temsirolimus
TSI method
BLCA−Temsirolimus
TC analysis
BLCA−Temsirolimus
Celligner

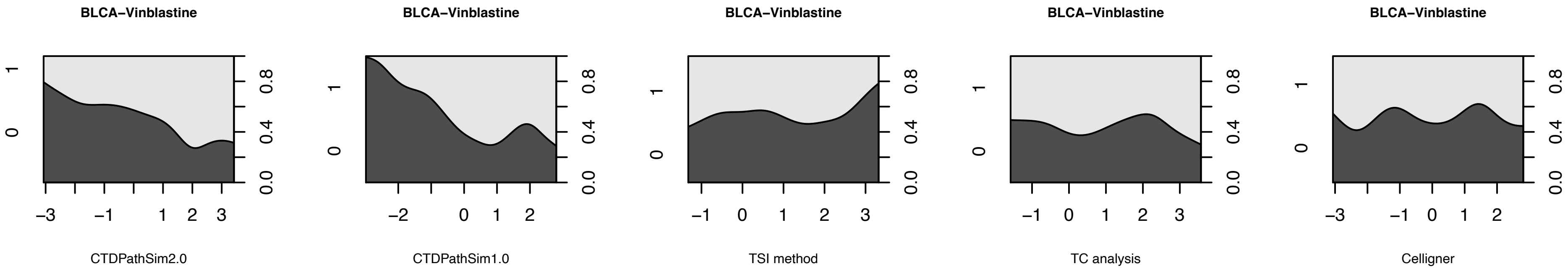
BLCA−Vinblastine
BLCA−Vinblastine
BLCA−Vinblastine
BLCA−Vinblastine
BLCA−Vinblastine
CTDPathSim2.0
CTDPathSim1.0
TSI method
TC analysis
Celligner

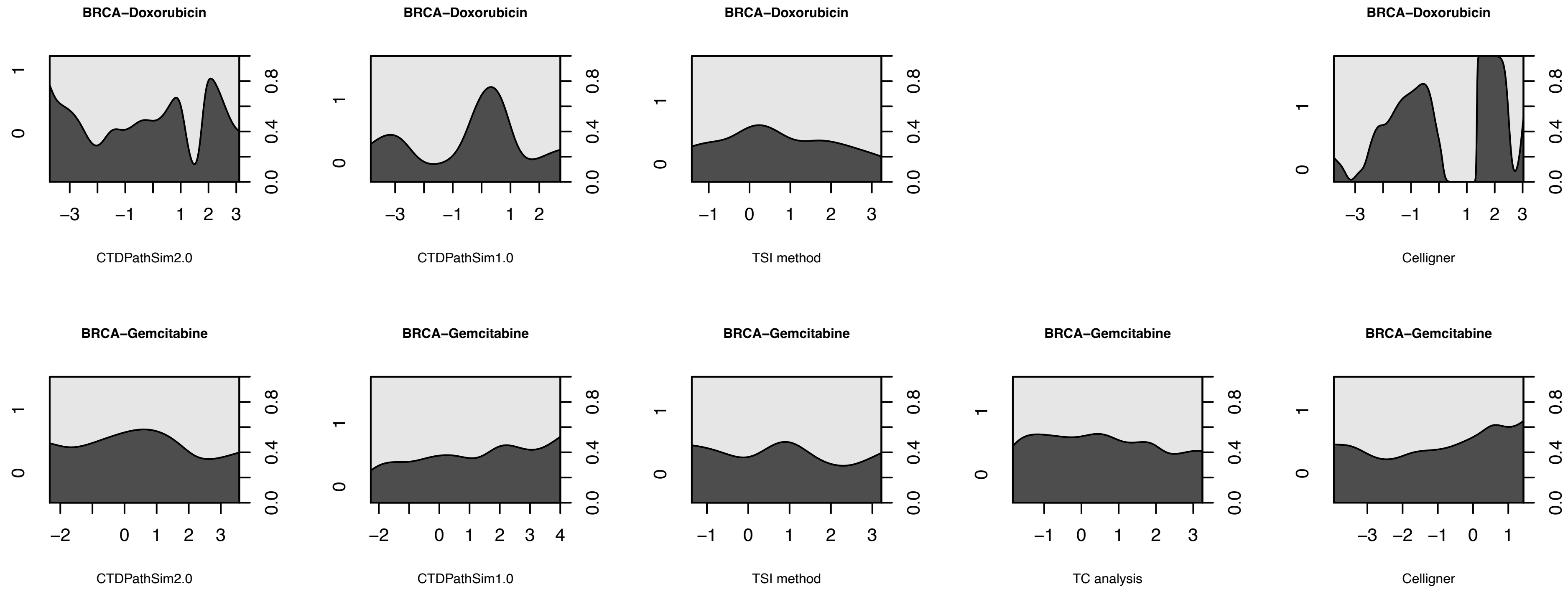
BRCA−Doxorubicin
CTDPathSim2.0
BRCA−Doxorubicin
CTDPathSim1.0
BRCA−Doxorubicin
TSI method
BRCA−Doxorubicin
Celligner
BRCA−Gemcitabine
CTDPathSim2.0
BRCA−Gemcitabine
CTDPathSim1.0
BRCA−Gemcitabine
TSI method
BRCA−Gemcitabine
TC analysis
BRCA−Gemcitabine
Celligner
0.0
0.4
0.8

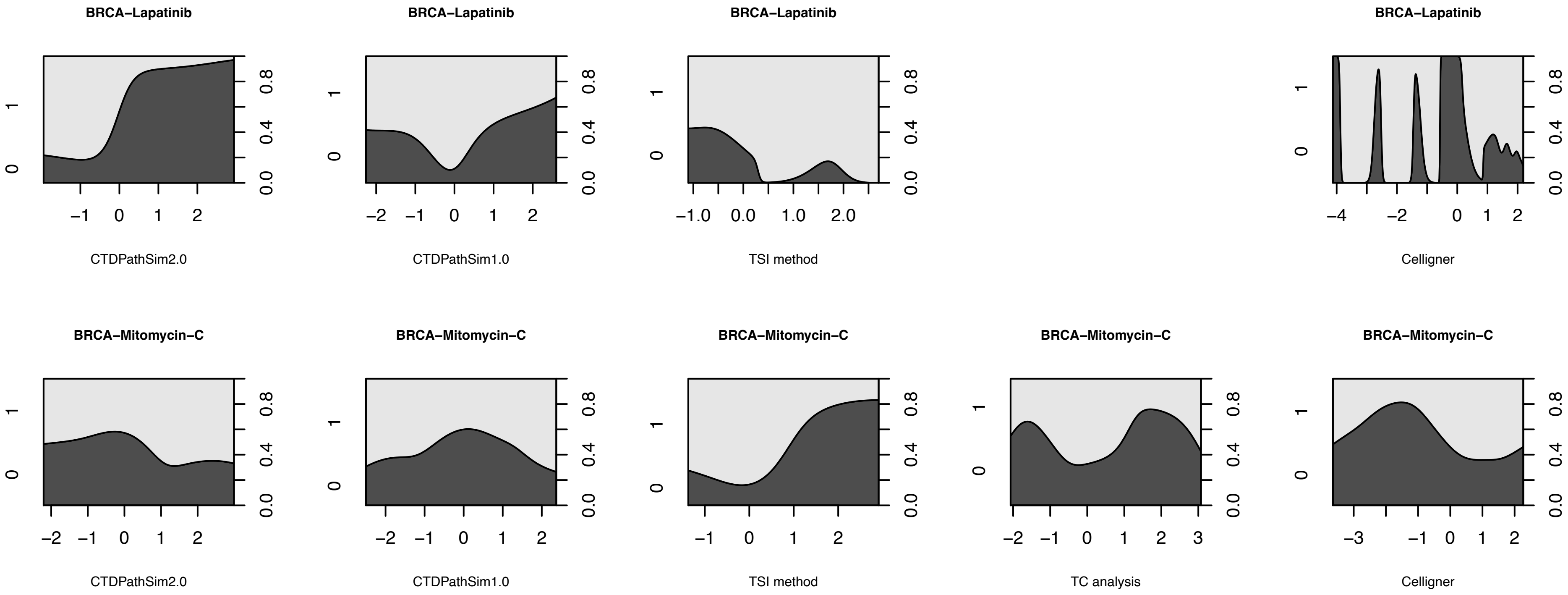
BRCA–Lapatinib
CTDPathSim2.0
BRCA–Lapatinib
CTDPathSim1.0
BRCA–Lapatinib
TSI method
BRCA–Lapatinib
Celligner
BRCA–Mitomycin–C
CTDPathSim2.0
BRCA–Mitomycin–C
CTDPathSim1.0
BRCA–Mitomycin–C
TSI method
BRCA–Mitomycin–C
TC analysis
BRCA–Mitomycin–C
Celligner

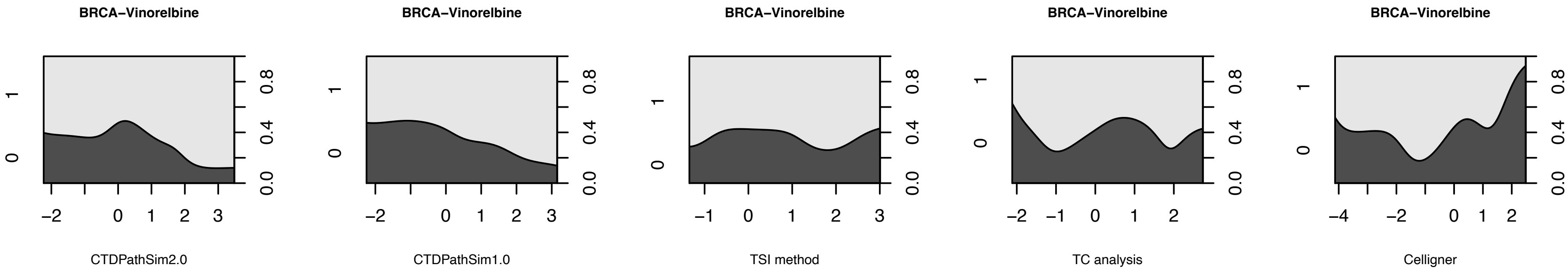

BRCA−Vinorelbine
CTDPathSim2.0
CTDPathSim1.0
TSI method
TC analysis
Celligner

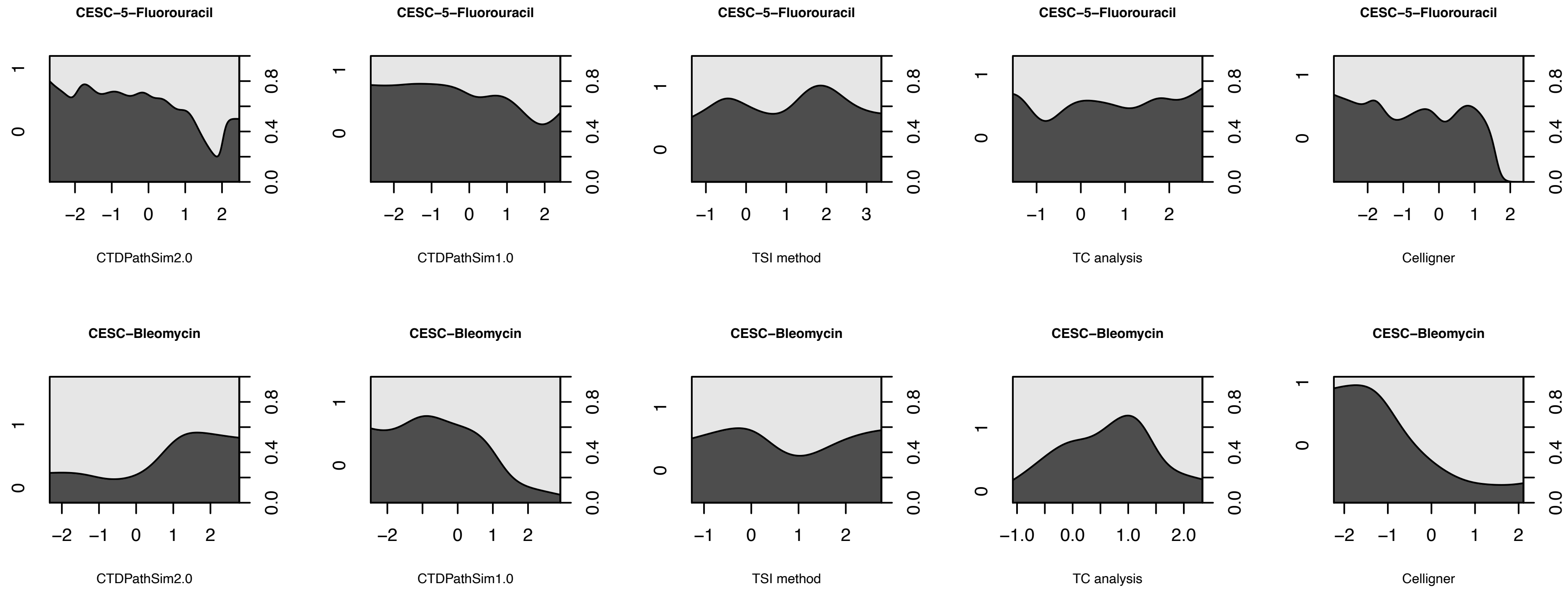
CESC−5−Fluorouracil
CTDPathSim2.0
CTDPathSim1.0
TSI method
TC analysis
Celligner
CESC−Bleomycin
CTDPathSim2.0
CTDPathSim1.0
TSI method
TC analysis
Celligner

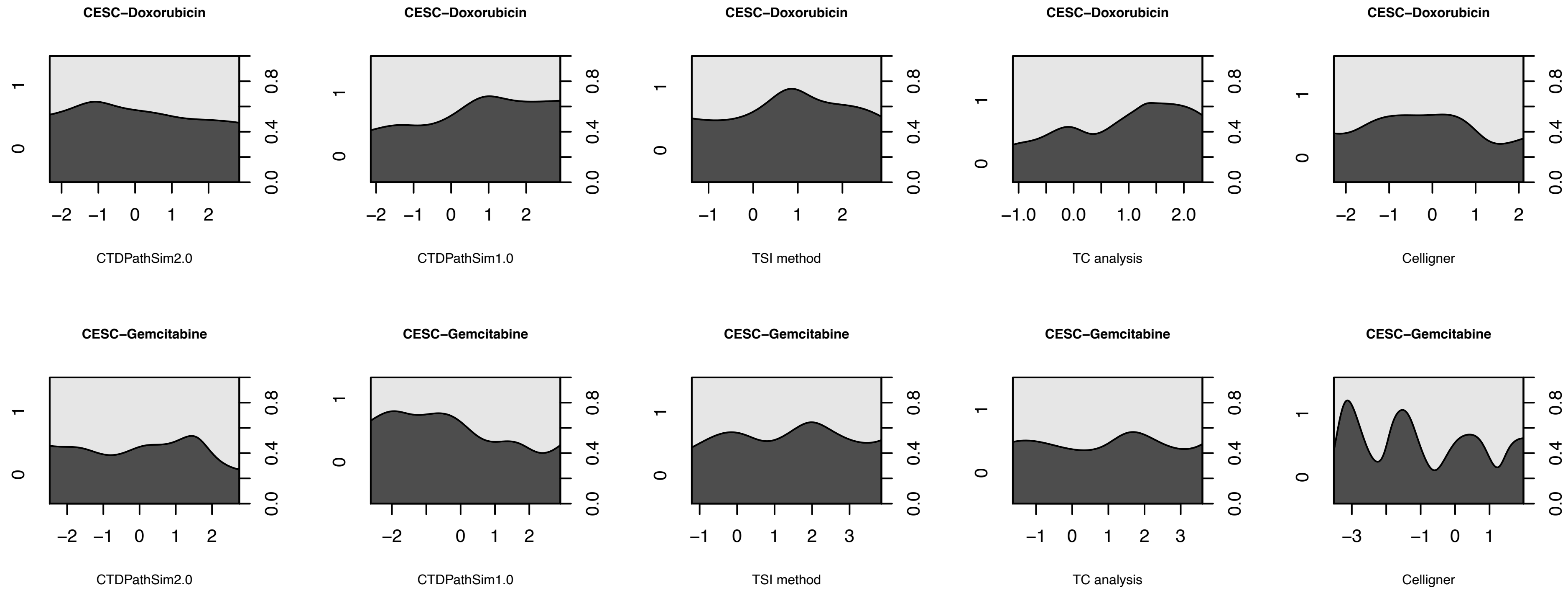

CESC−Doxorubicin
CTDPathSim2.0
CTDPathSim1.0
TSI method
TC analysis
Celligner
CESC−Gemcitabine
CTDPathSim2.0
CTDPathSim1.0
TSI method
TC analysis
Celligner

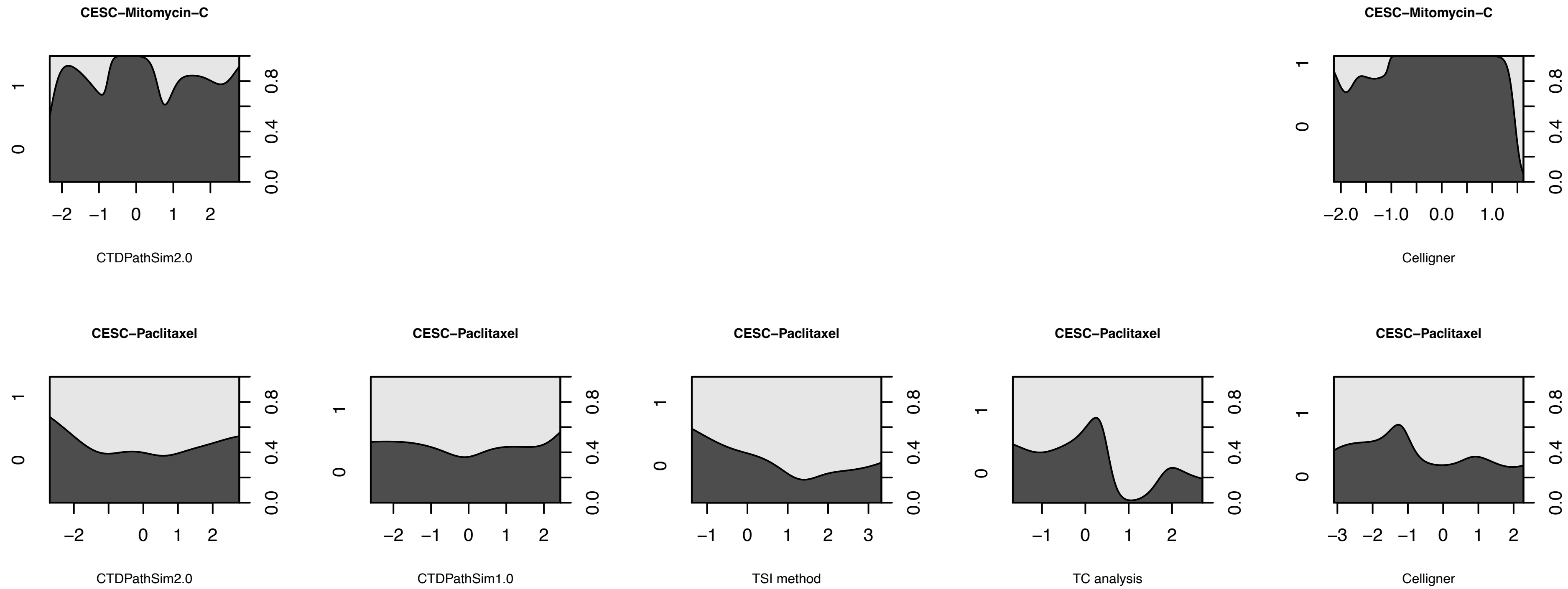
CESC−Mitomycin−C
CTDPathSim2.0
−2 −1 0 1 2
0 1
0.0 0.4 0.8
CESC−Mitomycin−C
Celligner
−2.0 −1.0 0.0 1.0
0 1
0.0 0.4 0.8
CESC−Paclitaxel
CTDPathSim2.0
−2 0 1 2
0 1
0.0 0.4 0.8
CESC−Paclitaxel
CTDPathSim1.0
−2 −1 0 1 2
0 1
0.0 0.4 0.8
CESC−Paclitaxel
TSI method
−1 0 1 2 3
0 1
0.0 0.4 0.8
CESC−Paclitaxel
TC analysis
−1 0 1 2
0 1
0.0 0.4 0.8
CESC−Paclitaxel
Celligner
−3 −2 −1 0 1 2
0 1
0.0 0.4 0.8

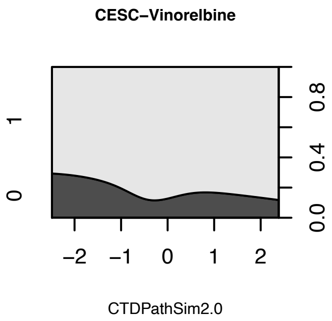
CESC−Vinorelbine
1
0
0.8
0.4
0.0
−2
−1
0
1
2
CTDPathSim2.0
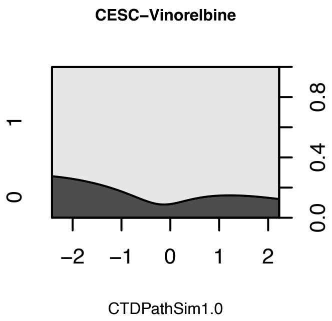
CESC−Vinorelbine
1
0
0.8
0.4
0.0
−2
−1
0
1
2
CTDPathSim1.0

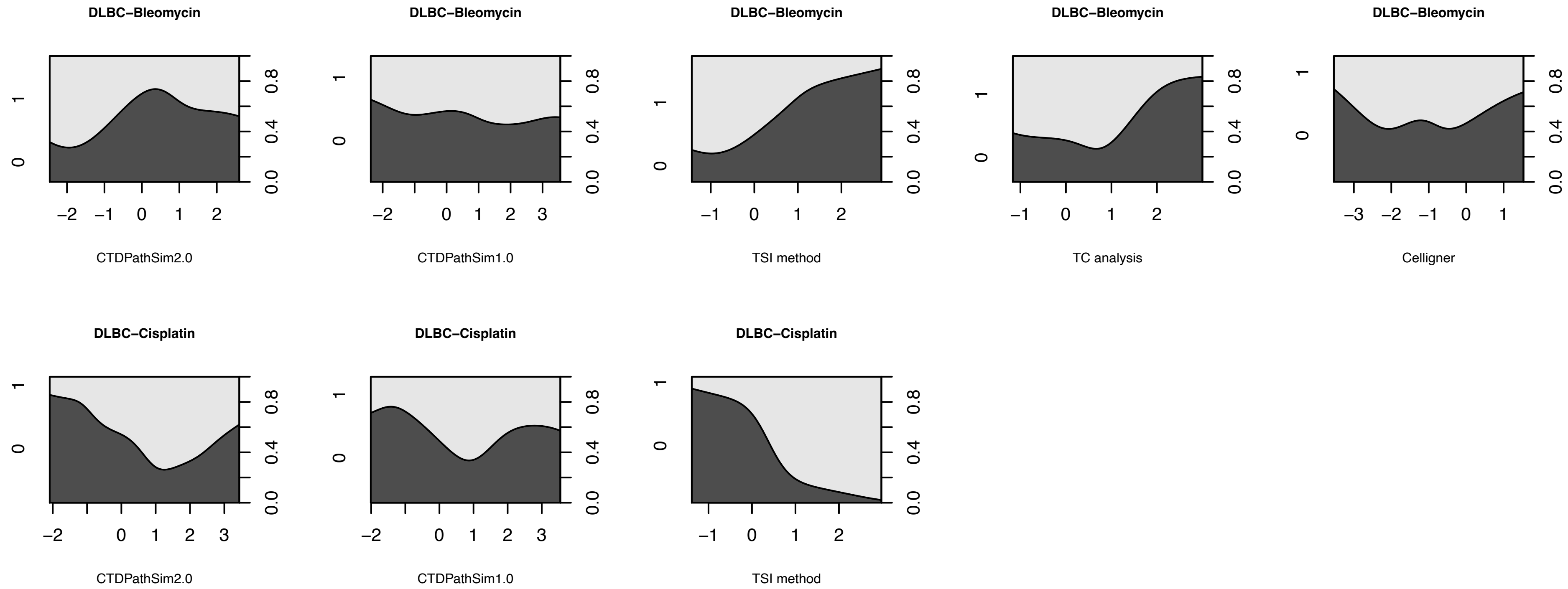
DLBC−Bleomycin
CTDPathSim2.0
DLBC−Bleomycin
CTDPathSim1.0
DLBC−Bleomycin
TSI method
DLBC−Bleomycin
TC analysis
DLBC−Bleomycin
Celligner
DLBC−Cisplatin
CTDPathSim2.0
DLBC−Cisplatin
CTDPathSim1.0
DLBC−Cisplatin
TSI method

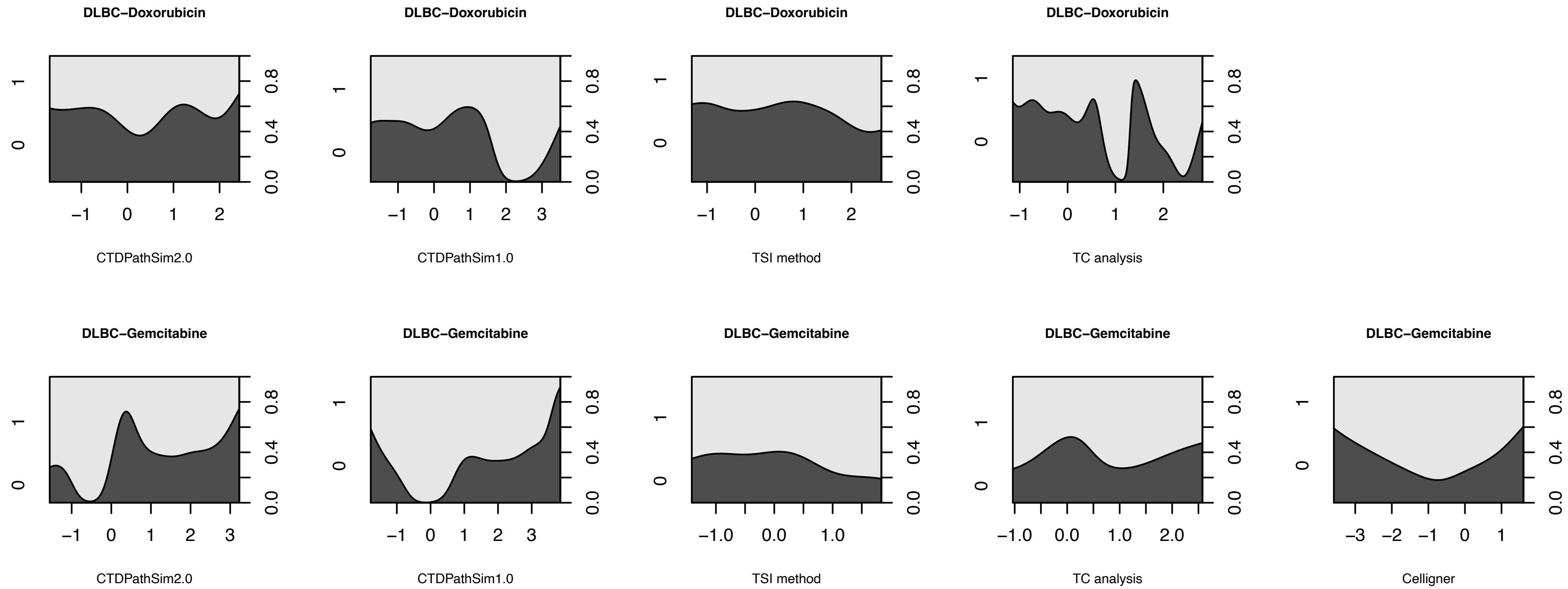
DLBC−Doxorubicin
CTDPathSim2.0
DLBC−Doxorubicin
CTDPathSim1.0
DLBC−Doxorubicin
TSI method
DLBC−Doxorubicin
TC analysis
DLBC−Gemcitabine
CTDPathSim2.0
DLBC−Gemcitabine
CTDPathSim1.0
DLBC−Gemcitabine
TSI method
DLBC−Gemcitabine
TC analysis
DLBC−Gemcitabine
Celligner

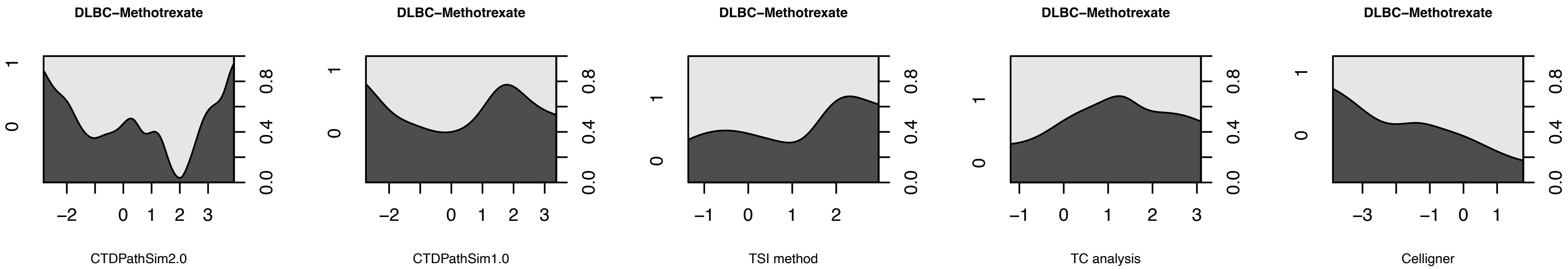
DLBC−Methotrexate
CTDPathSim2.0
DLBC−Methotrexate
CTDPathSim1.0
DLBC−Methotrexate
TSI method
DLBC−Methotrexate
TC analysis
DLBC−Methotrexate
Celligner

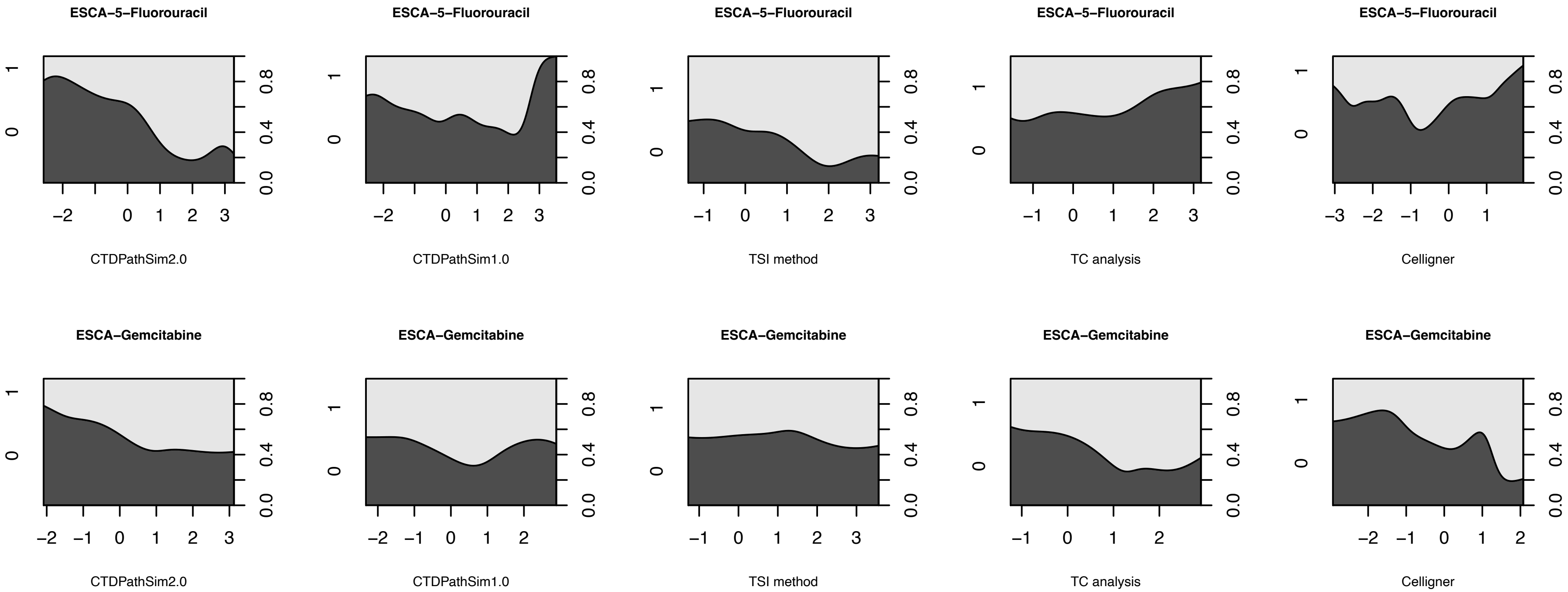
ESCA−5−Fluorouracil
CTDPathSim2.0
CTDPathSim1.0
TSI method
TC analysis
Celligner
ESCA−Gemcitabine
CTDPathSim2.0
CTDPathSim1.0
TSI method
TC analysis
Celligner

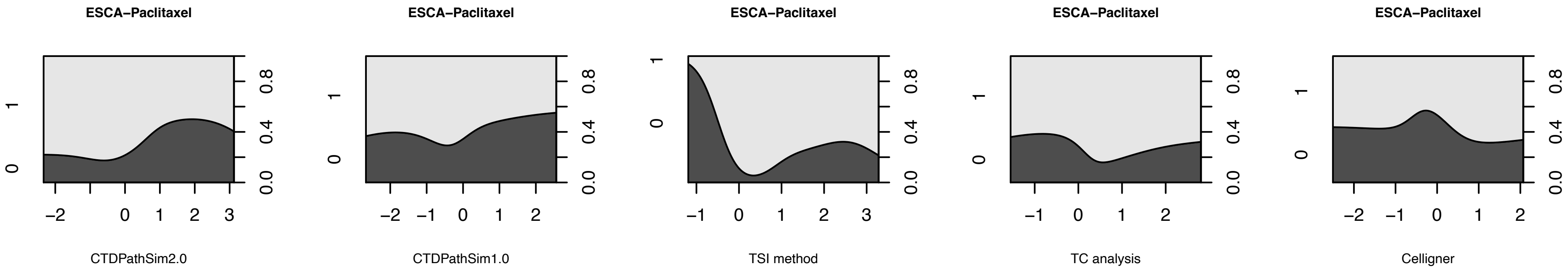
ESCA−Paclitaxel
CTDPathSim2.0
ESCA−Paclitaxel
CTDPathSim1.0
ESCA−Paclitaxel
TSI method
ESCA−Paclitaxel
TC analysis
ESCA−Paclitaxel
Celligner
0.0
0.4
0.8
−2
−1
0
1
2
3

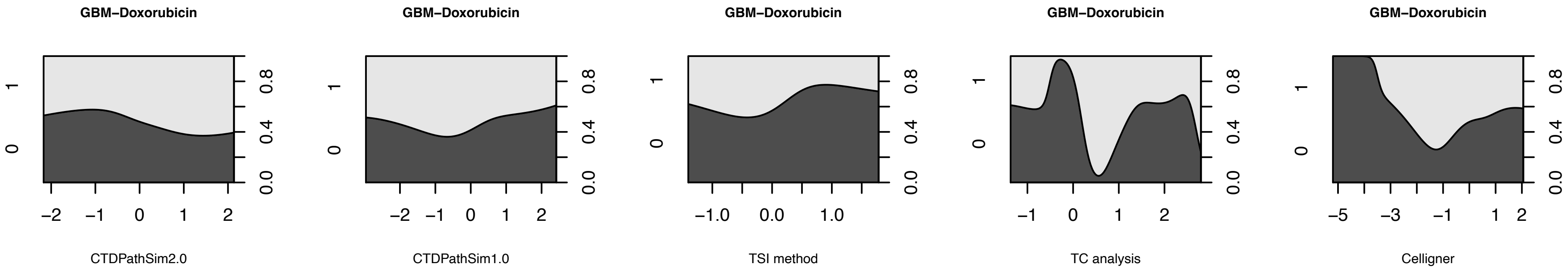
GBM−Doxorubicin
GBM−Doxorubicin
GBM−Doxorubicin
GBM−Doxorubicin
GBM−Doxorubicin
CTDPathSim2.0
CTDPathSim1.0
TSI method
TC analysis
Celligner

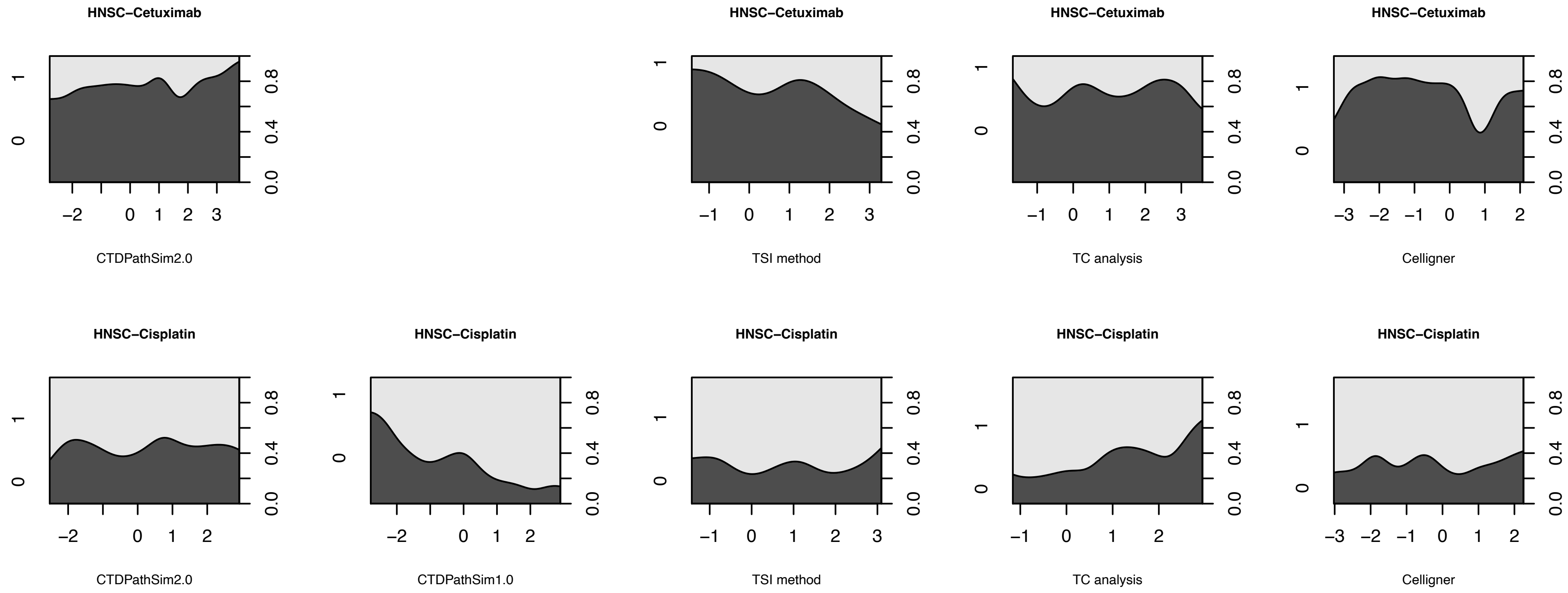
HNSC−Cetuximab
CTDPathSim2.0
HNSC−Cetuximab
TSI method
HNSC−Cetuximab
TC analysis
HNSC−Cetuximab
Celligner
HNSC−Cisplatin
CTDPathSim2.0
HNSC−Cisplatin
CTDPathSim1.0
HNSC−Cisplatin
TSI method
HNSC−Cisplatin
TC analysis
HNSC−Cisplatin
Celligner

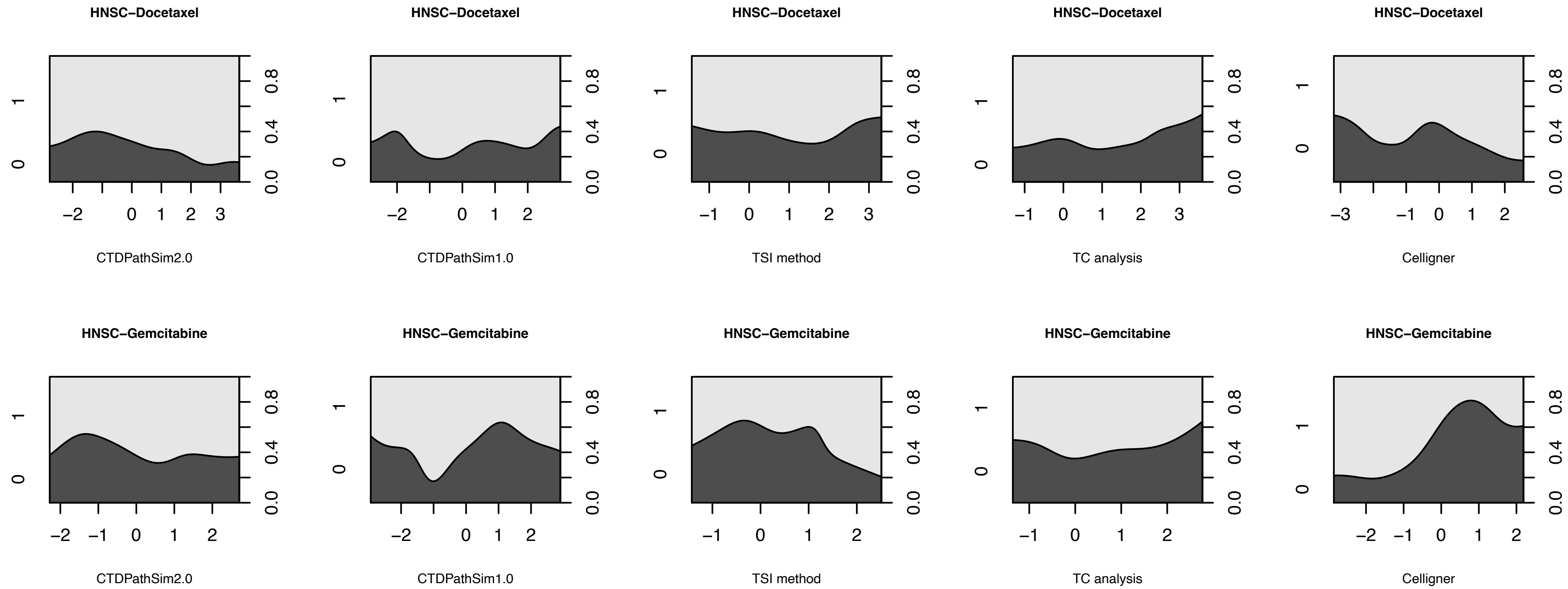

HNSC−Docetaxel
CTDPathSim2.0
CTDPathSim1.0
TSI method
TC analysis
Celligner
HNSC−Gemcitabine

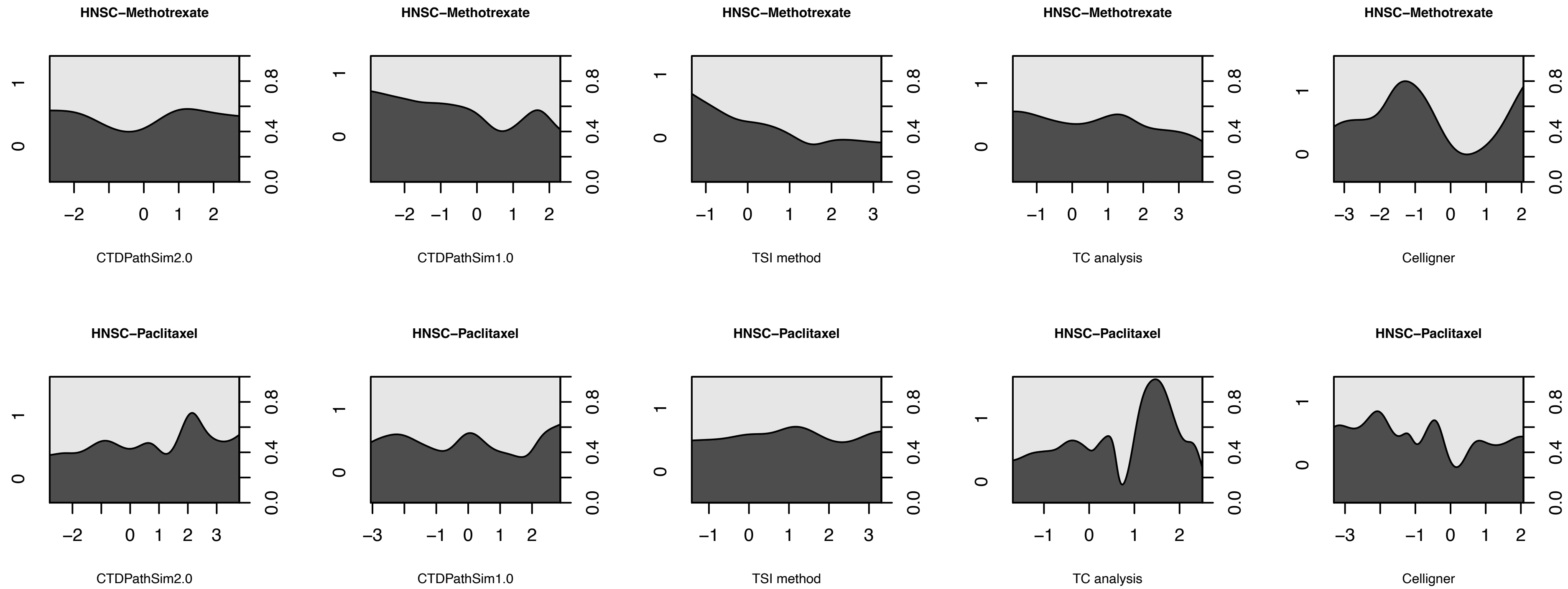

HNSC−Methotrexate
CTDPathSim2.0
CTDPathSim1.0
TSI method
TC analysis
Celligner
HNSC−Paclitaxel
CTDPathSim2.0
CTDPathSim1.0
TSI method
TC analysis
Celligner

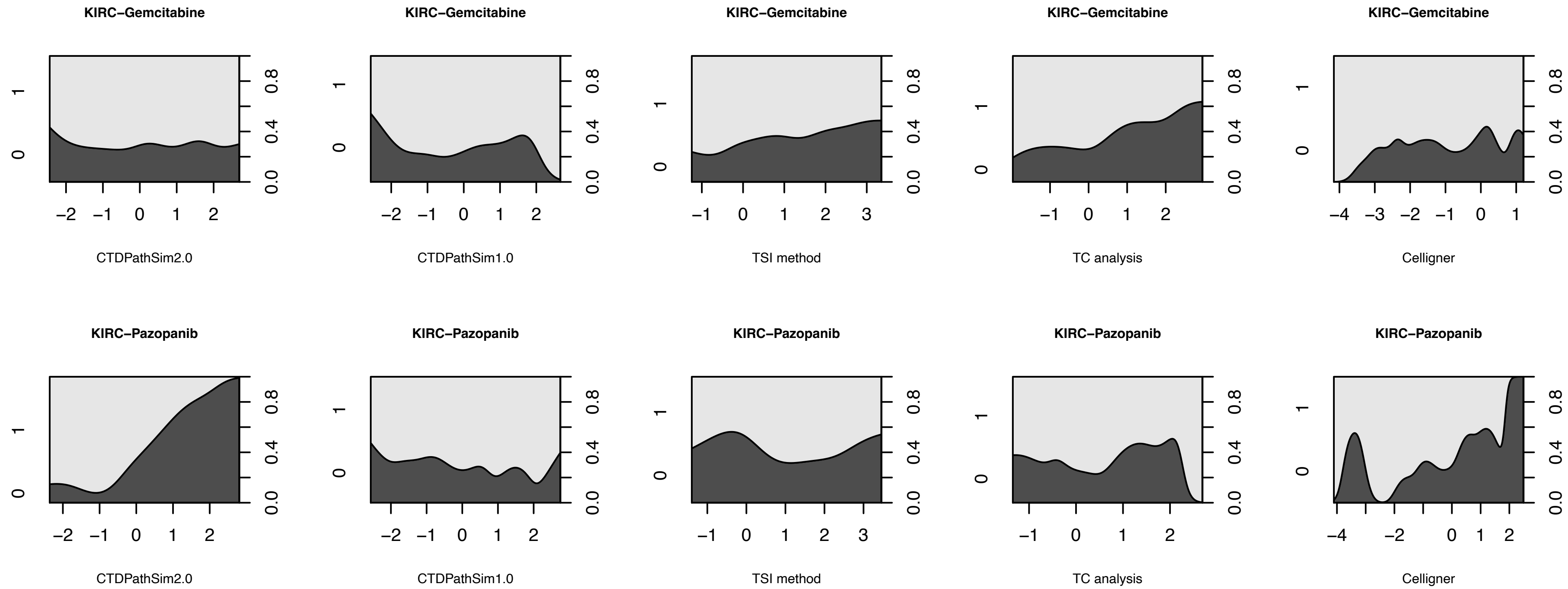

KIRC−Gemcitabine
CTDPathSim2.0
CTDPathSim1.0
TSI method
TC analysis
Celligner
KIRC−Pazopanib
CTDPathSim2.0
CTDPathSim1.0
TSI method
TC analysis
Celligner

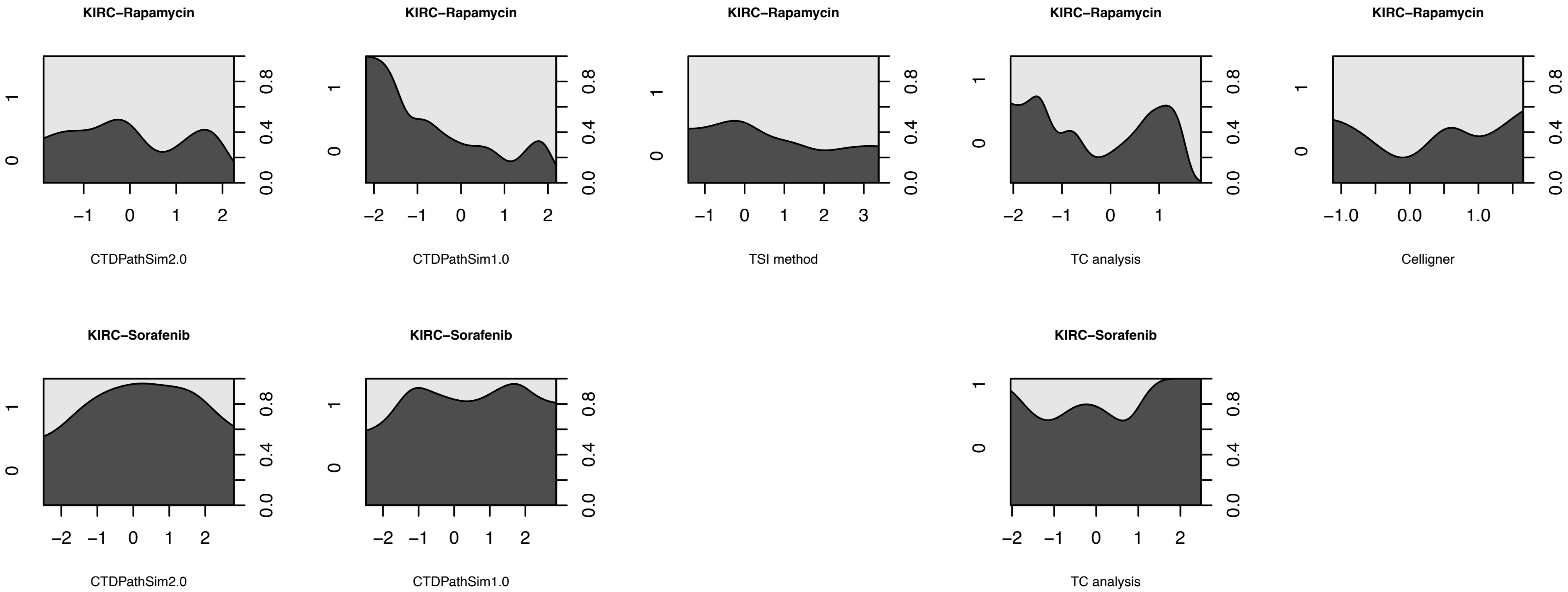

KIRC−Rapamycin
CTDPathSim2.0
KIRC−Rapamycin
CTDPathSim1.0
KIRC−Rapamycin
TSI method
KIRC−Rapamycin
TC analysis
KIRC−Rapamycin
Celligner
KIRC−Sorafenib
CTDPathSim2.0
KIRC−Sorafenib
CTDPathSim1.0
KIRC−Sorafenib
TC analysis

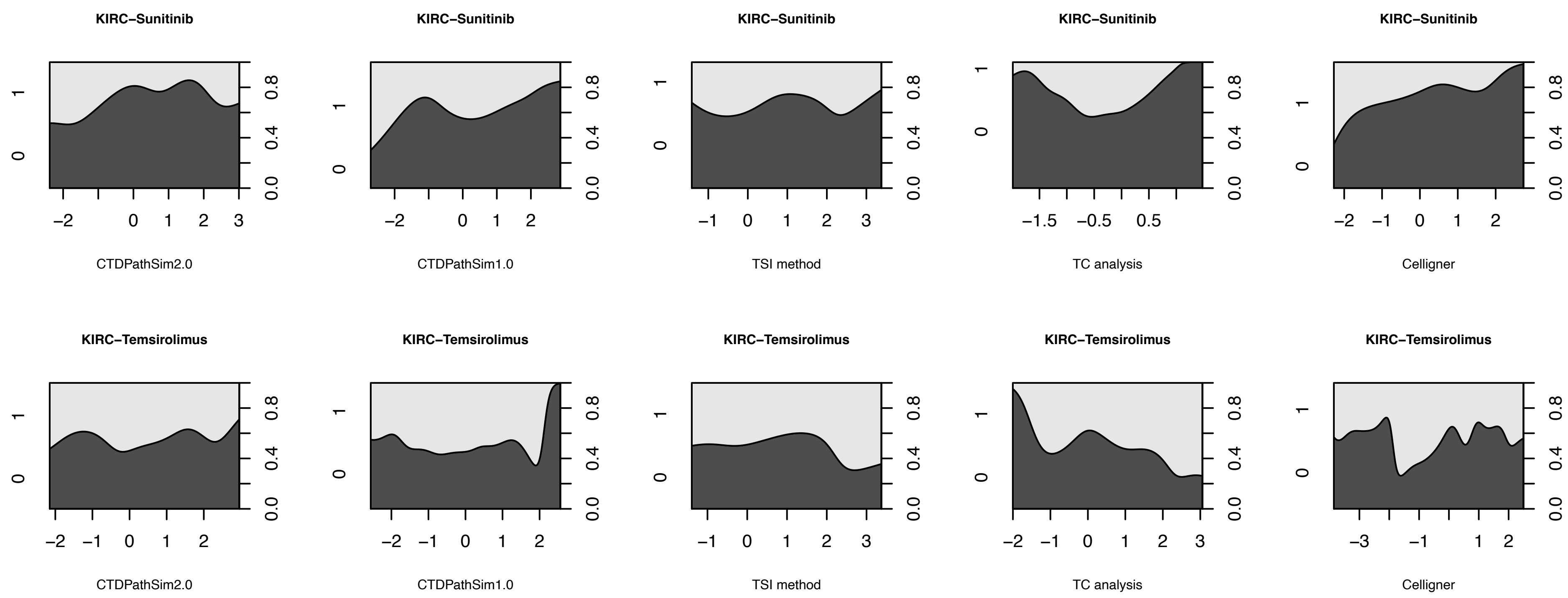

KIRC−Sunitinib
CTDPathSim2.0
CTDPathSim1.0
TSI method
TC analysis
Celligner
KIRC−Temsirolimus
CTDPathSim2.0
CTDPathSim1.0
TSI method
TC analysis
Celligner

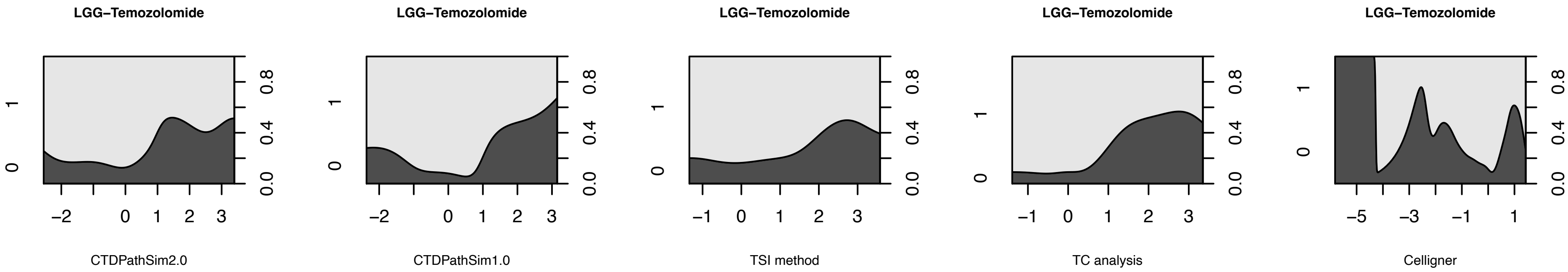
LGG−Temozolomide
CTDPathSim2.0
LGG−Temozolomide
CTDPathSim1.0
LGG−Temozolomide
TSI method
LGG−Temozolomide
TC analysis
LGG−Temozolomide
Celligner

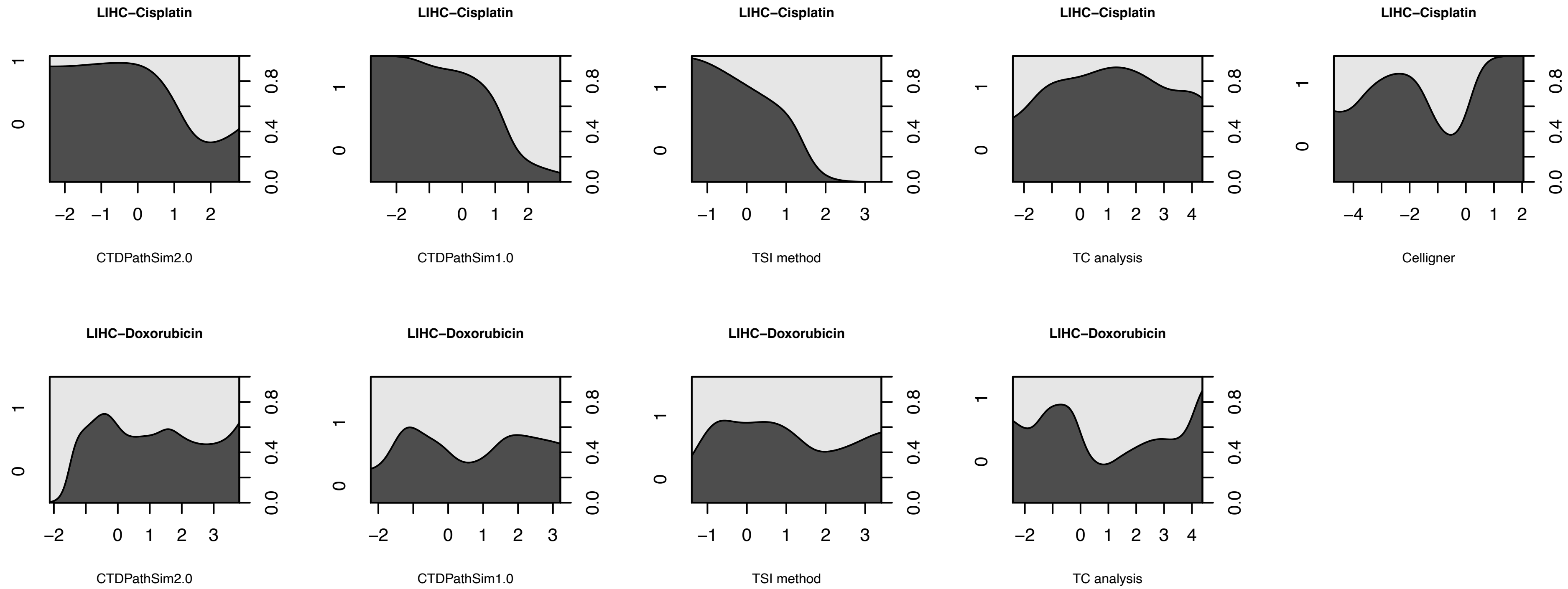

LIHC−Cisplatin
CTDPathSim2.0
LIHC−Cisplatin
CTDPathSim1.0
LIHC−Cisplatin
TSI method
LIHC−Cisplatin
TC analysis
LIHC−Cisplatin
Celligner
LIHC−Doxorubicin
CTDPathSim2.0
LIHC−Doxorubicin
CTDPathSim1.0
LIHC−Doxorubicin
TSI method
LIHC−Doxorubicin
TC analysis

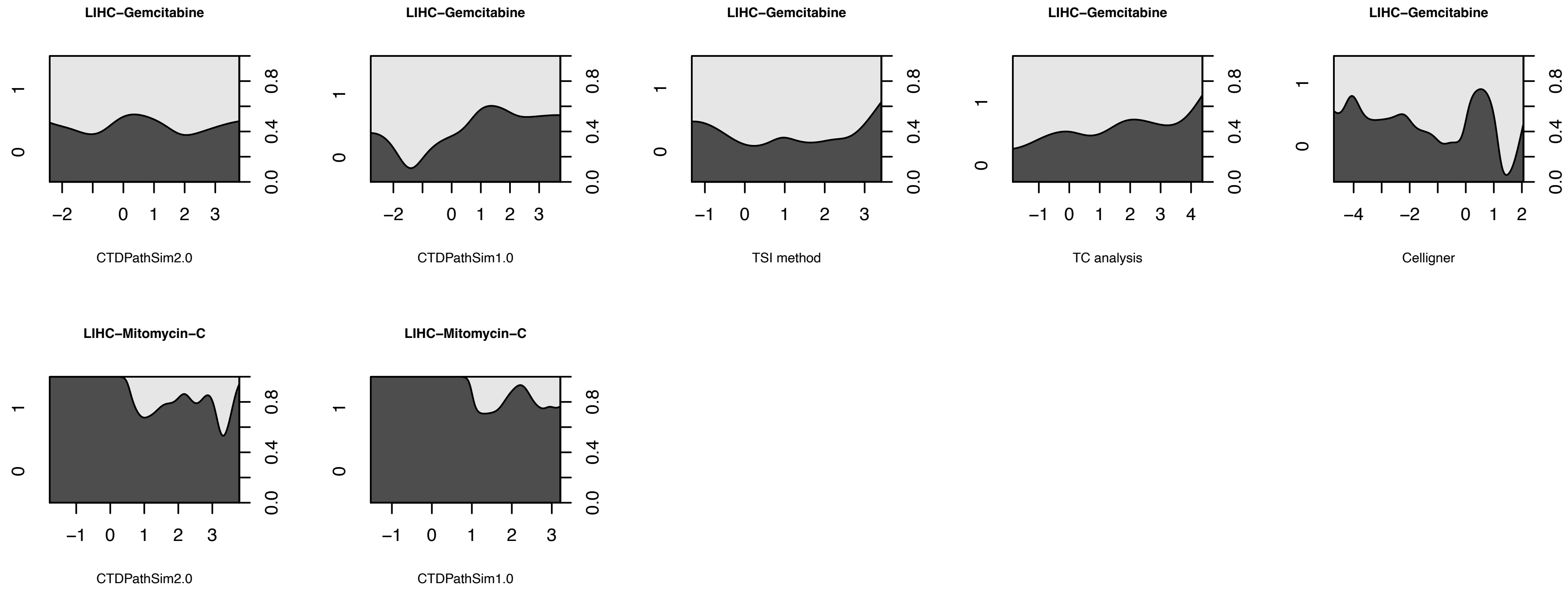

LIHC−Gemcitabine
CTDPathSim2.0
LIHC−Gemcitabine
CTDPathSim1.0
LIHC−Gemcitabine
TSI method
LIHC−Gemcitabine
TC analysis
LIHC−Gemcitabine
Celligner
LIHC−Mitomycin−C
CTDPathSim2.0
LIHC−Mitomycin−C
CTDPathSim1.0

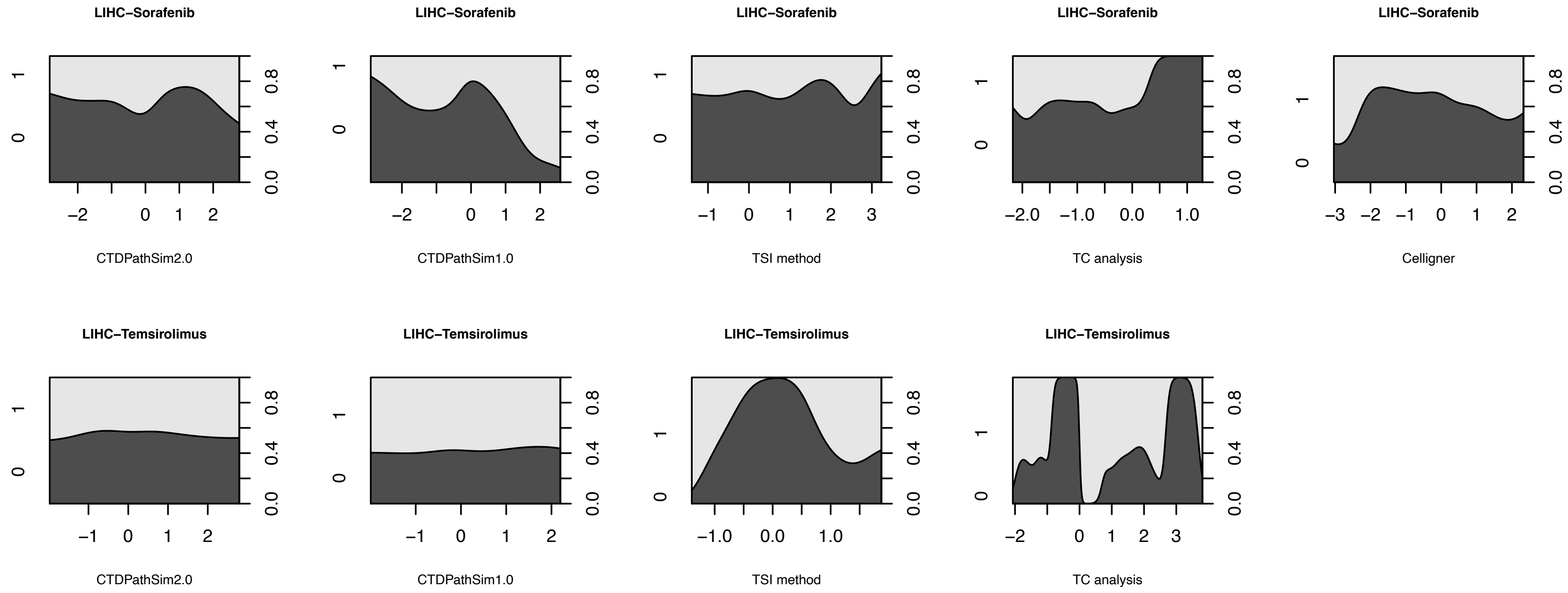
LIHC−Sorafenib
CTDPathSim2.0
LIHC−Sorafenib
CTDPathSim1.0
LIHC−Sorafenib
TSI method
LIHC−Sorafenib
TC analysis
LIHC−Sorafenib
Celligner
LIHC−Temsirolimus
CTDPathSim2.0
LIHC−Temsirolimus
CTDPathSim1.0
LIHC−Temsirolimus
TSI method
LIHC−Temsirolimus
TC analysis
0
1
0.0
0.4
0.8
−2
0
1
2
−1
3
−2.0
−1.0
0.0
1.0
−3

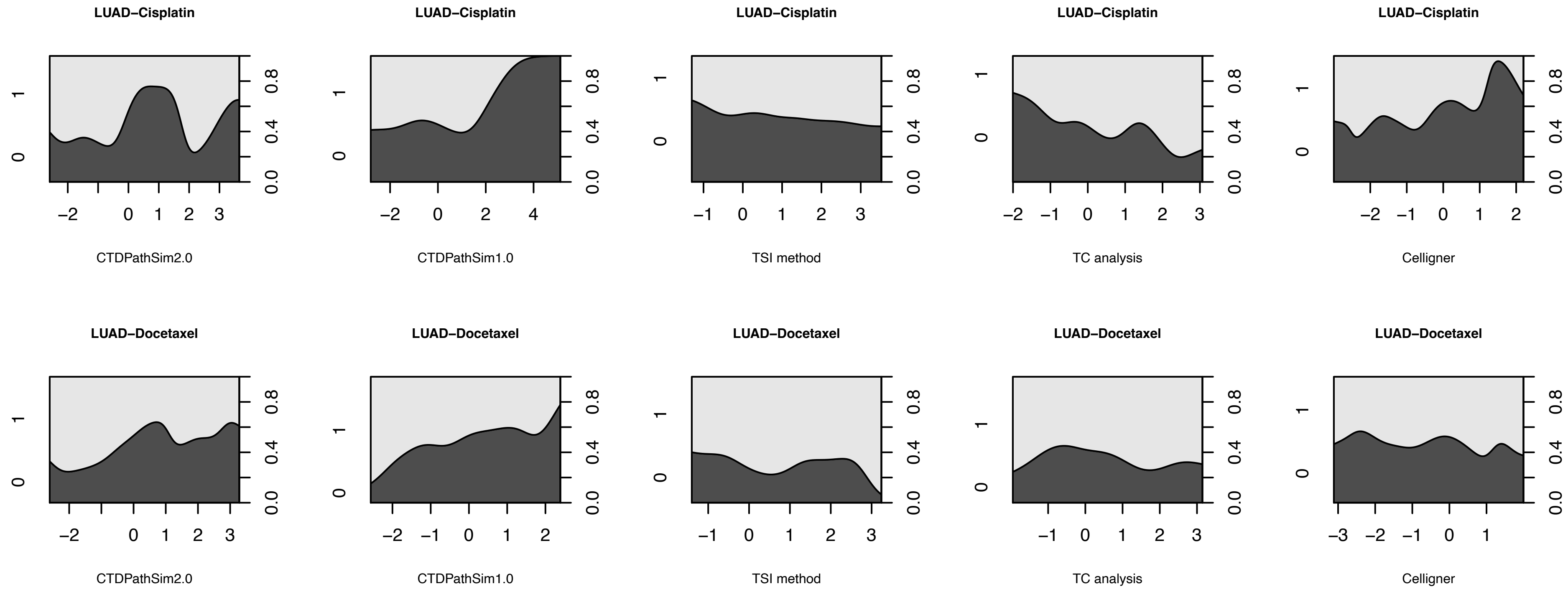

LUAD−Cisplatin
CTDPathSim2.0
CTDPathSim1.0
TSI method
TC analysis
Celligner
LUAD−Docetaxel
CTDPathSim2.0
CTDPathSim1.0
TSI method
TC analysis
Celligner

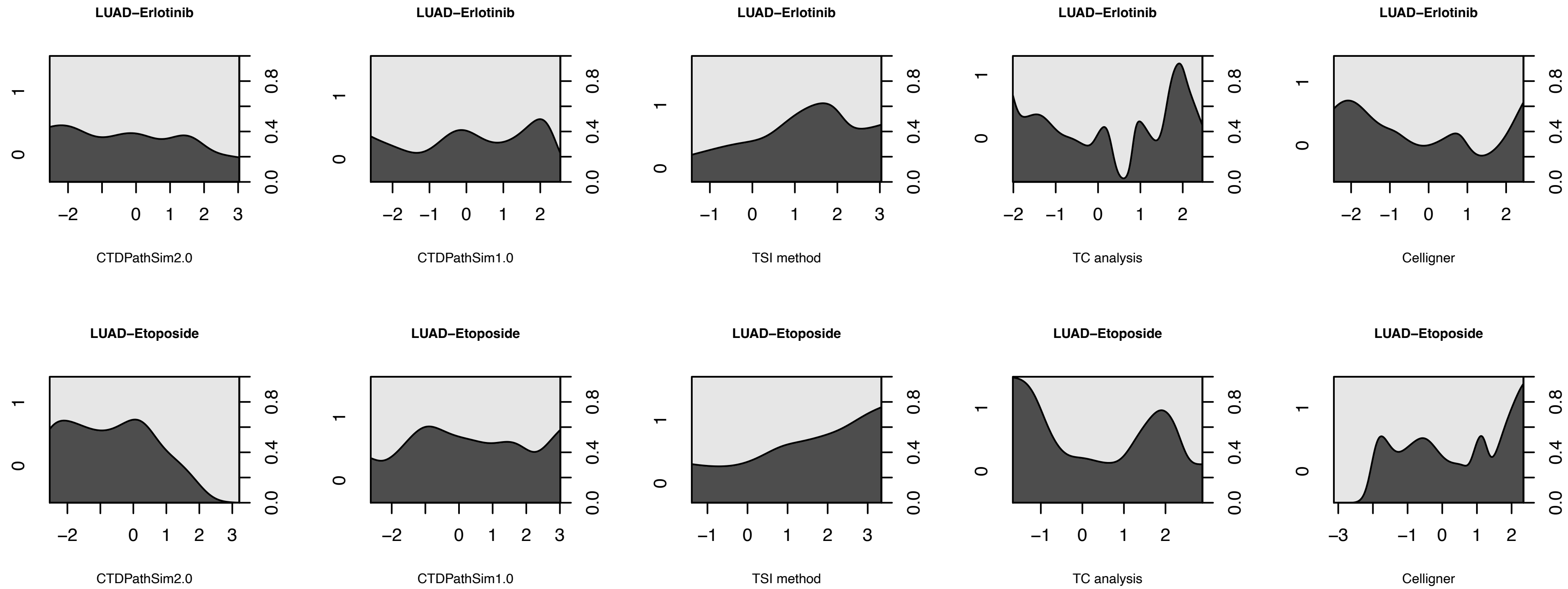

LUAD−Erlotinib
CTDPathSim2.0
CTDPathSim1.0
TSI method
TC analysis
Celligner
LUAD−Etoposide
CTDPathSim2.0
CTDPathSim1.0
TSI method
TC analysis
Celligner

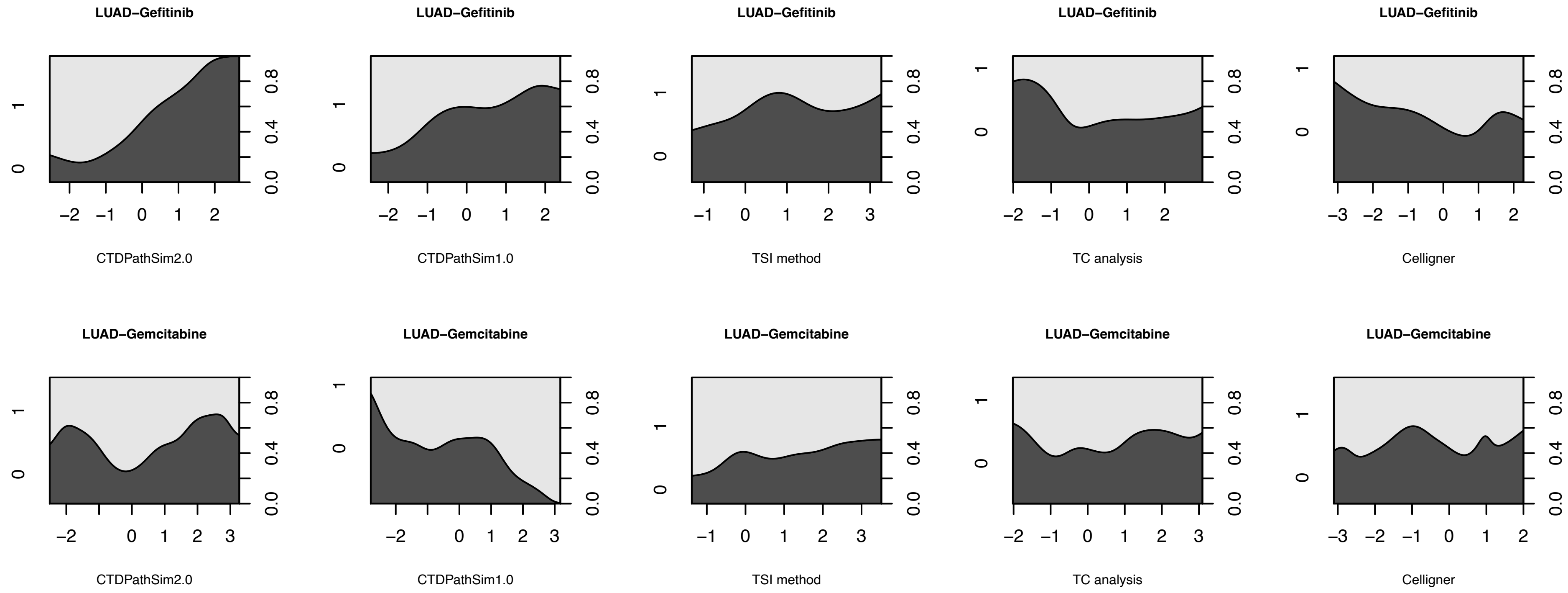

LUAD−Gefitinib
LUAD−Gefitinib
LUAD−Gefitinib
LUAD−Gefitinib
LUAD−Gefitinib
CTDPathSim2.0
CTDPathSim1.0
TSI method
TC analysis
Celligner
LUAD−Gemcitabine
LUAD−Gemcitabine
LUAD−Gemcitabine
LUAD−Gemcitabine
LUAD−Gemcitabine
CTDPathSim2.0
CTDPathSim1.0
TSI method
TC analysis
Celligner

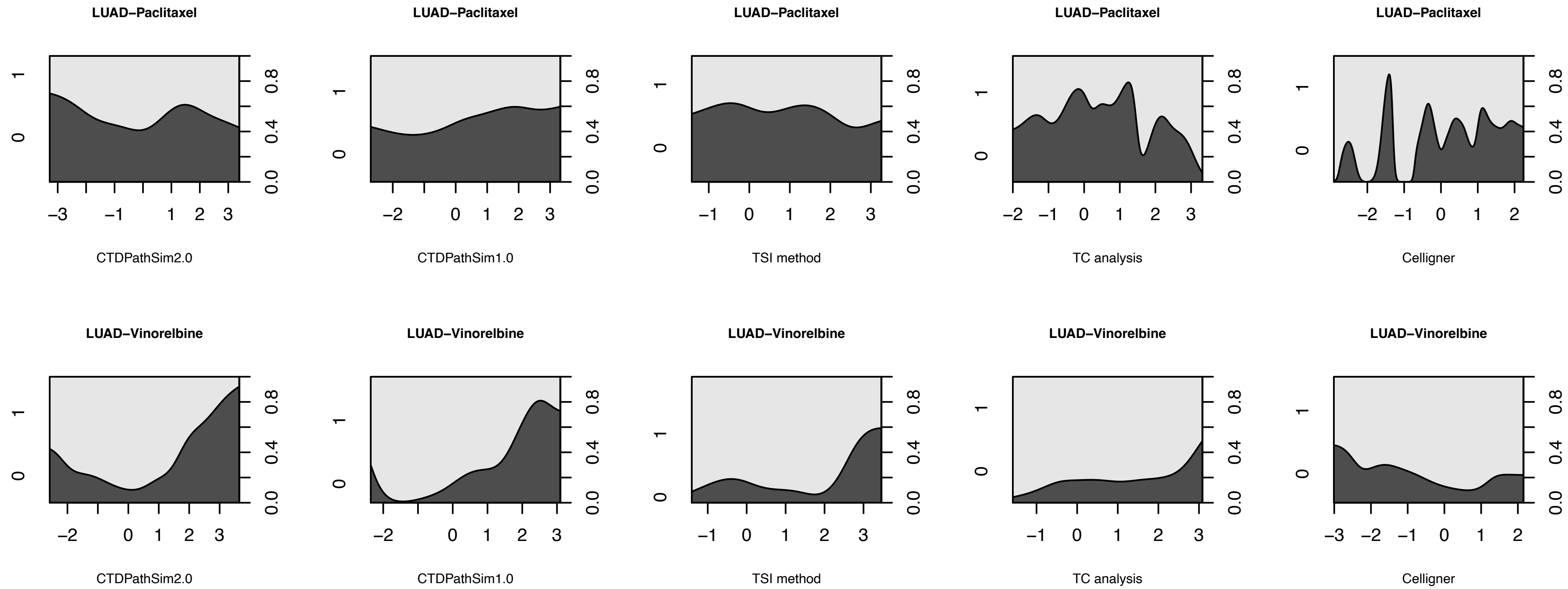
LUAD−Paclitaxel
CTDPathSim2.0
LUAD−Paclitaxel
CTDPathSim1.0
LUAD−Paclitaxel
TSI method
LUAD−Paclitaxel
TC analysis
LUAD−Paclitaxel
Celligner
LUAD−Vinorelbine
CTDPathSim2.0
LUAD−Vinorelbine
CTDPathSim1.0
LUAD−Vinorelbine
TSI method
LUAD−Vinorelbine
TC analysis
LUAD−Vinorelbine
Celligner
0.0
0.4
0.8
0
1

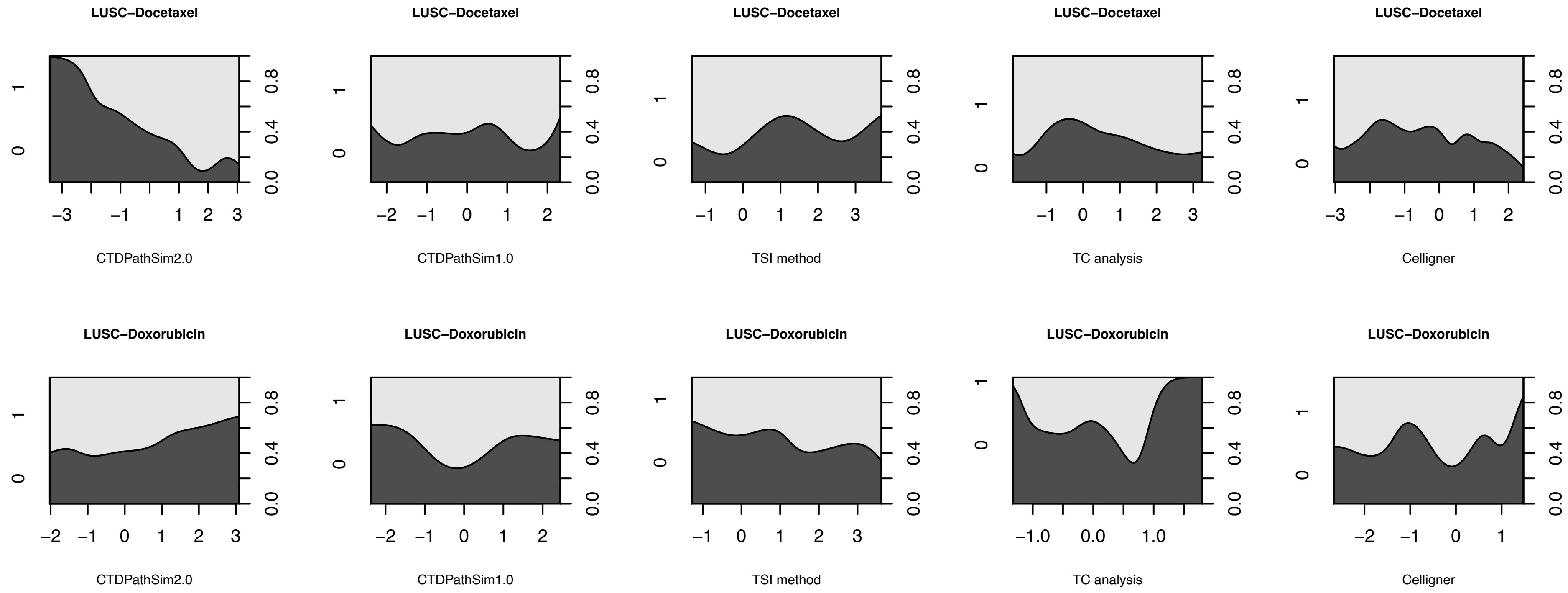

LUSC−Docetaxel
CTDPathSim2.0
CTDPathSim1.0
TSI method
TC analysis
Celligner
LUSC−Doxorubicin

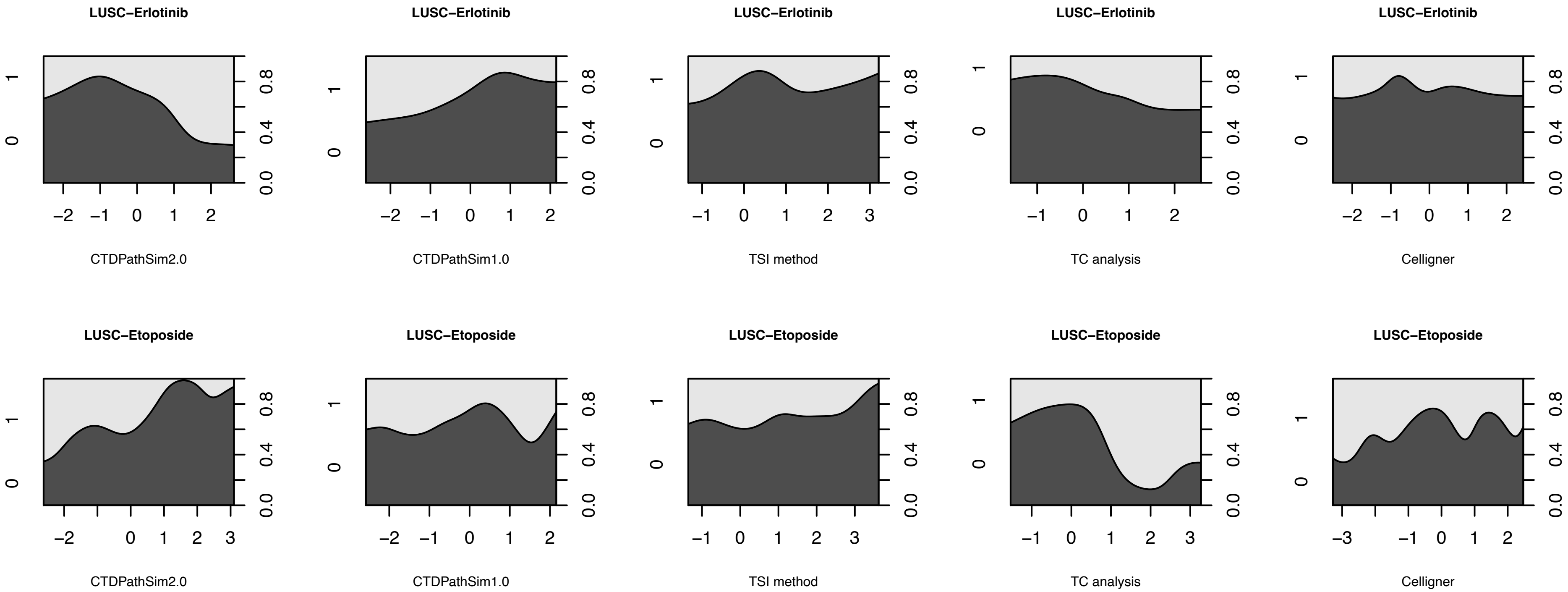
LUSC−Erlotinib
CTDPathSim2.0
LUSC−Erlotinib
CTDPathSim1.0
LUSC−Erlotinib
TSI method
LUSC−Erlotinib
TC analysis
LUSC−Erlotinib
Celligner
LUSC−Etoposide
CTDPathSim2.0
LUSC−Etoposide
CTDPathSim1.0
LUSC−Etoposide
TSI method
LUSC−Etoposide
TC analysis
LUSC−Etoposide
Celligner

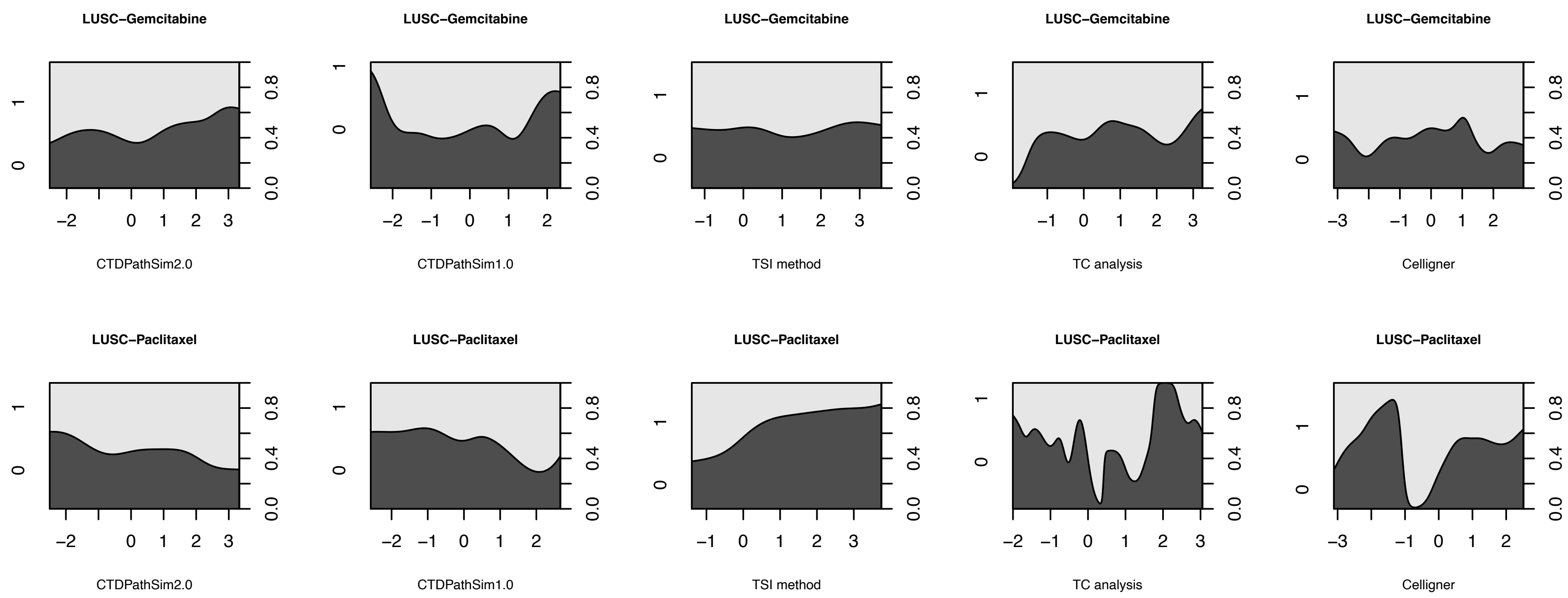
LUSC−Gemcitabine
CTDPathSim2.0
CTDPathSim1.0
TSI method
TC analysis
Celligner
LUSC−Paclitaxel

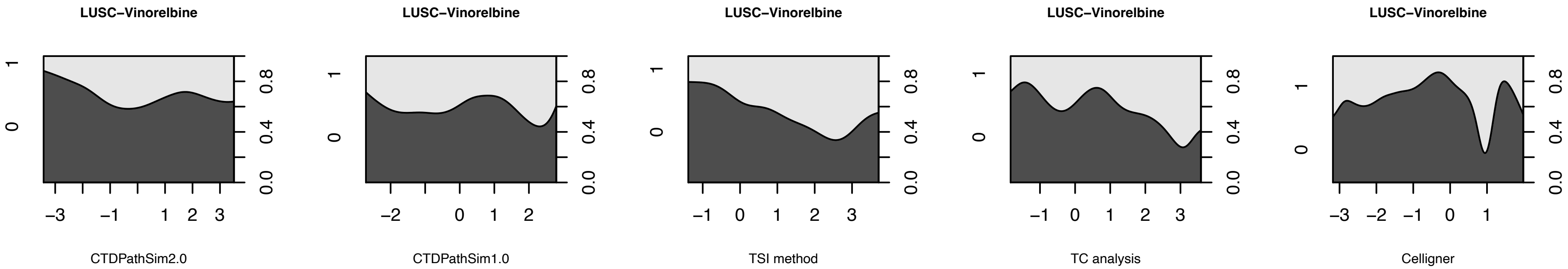

LUSC−Vinorelbine
LUSC−Vinorelbine
LUSC−Vinorelbine
LUSC−Vinorelbine
LUSC−Vinorelbine
CTDPathSim2.0
CTDPathSim1.0
TSI method
TC analysis
Celligner

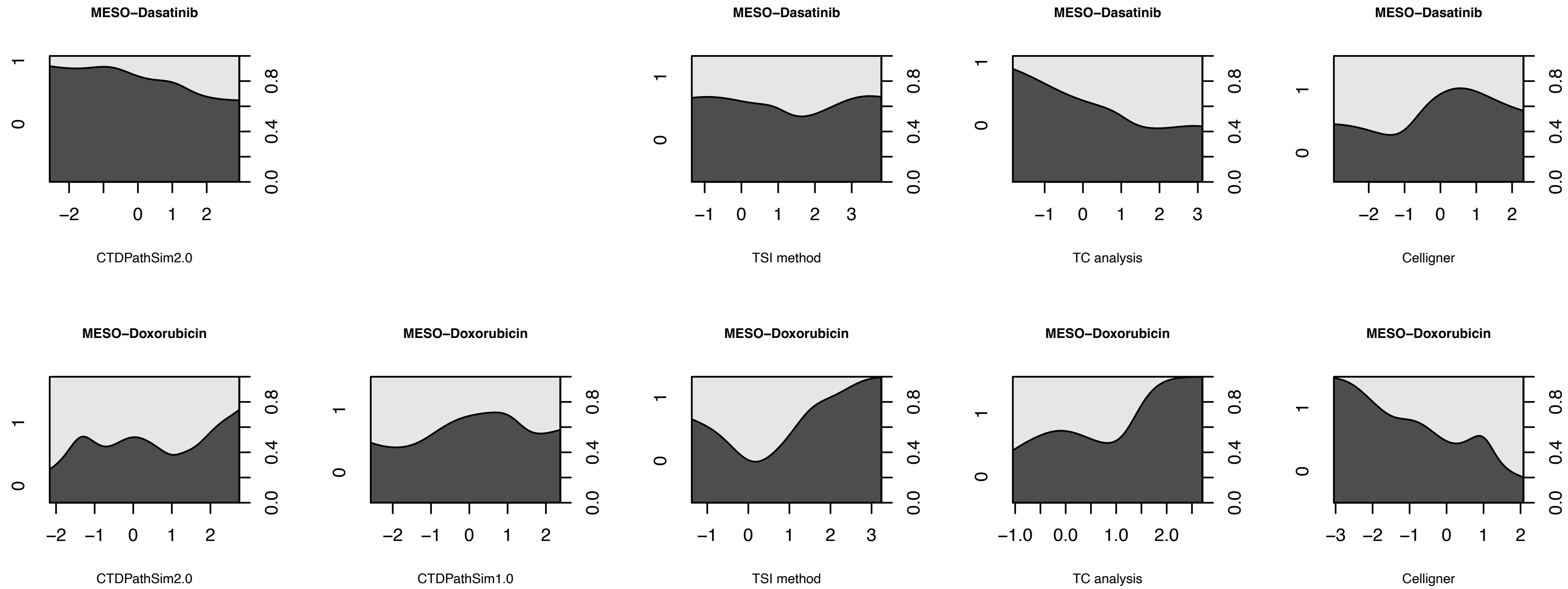
MESO−Dasatinib
CTDPathSim2.0
MESO−Dasatinib
TSI method
MESO−Dasatinib
TC analysis
MESO−Dasatinib
Celligner
MESO−Doxorubicin
CTDPathSim2.0
MESO−Doxorubicin
CTDPathSim1.0
MESO−Doxorubicin
TSI method
MESO−Doxorubicin
TC analysis
MESO−Doxorubicin
Celligner

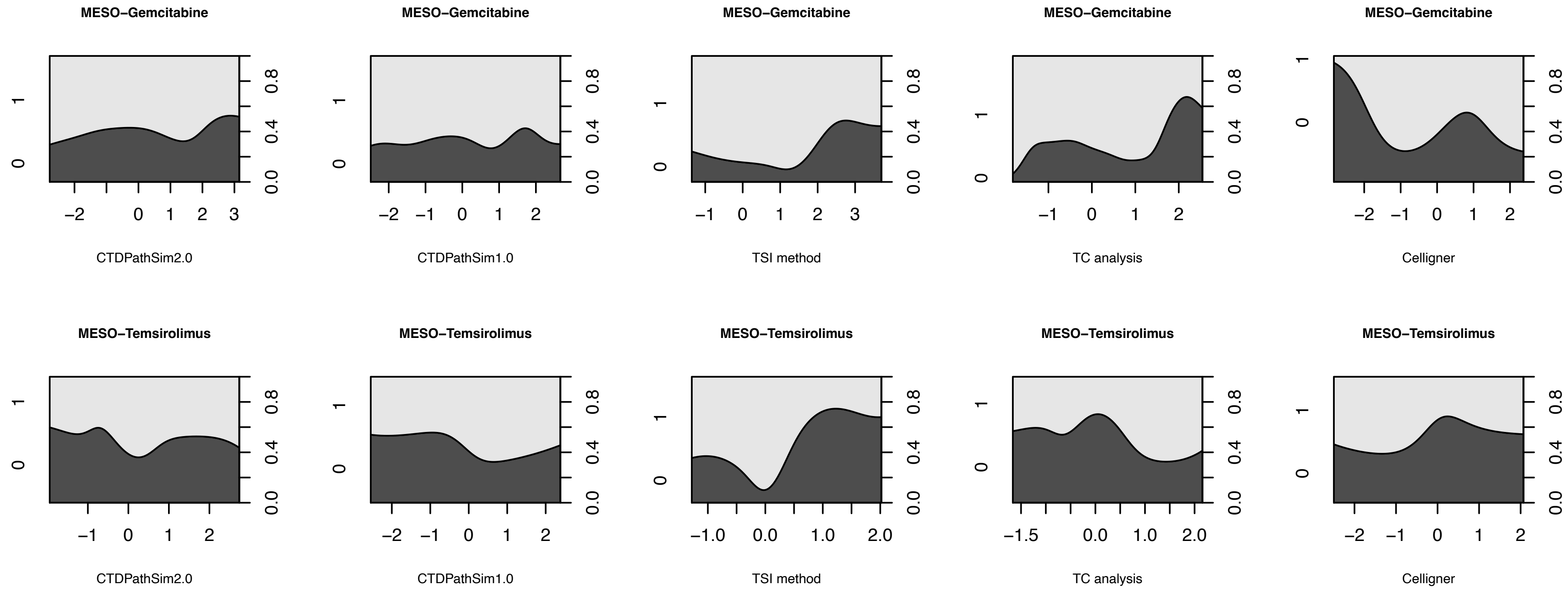
MESO−Gemcitabine
CTDPathSim2.0
CTDPathSim1.0
TSI method
TC analysis
Celligner
MESO−Temsirolimus
CTDPathSim2.0
CTDPathSim1.0
TSI method
TC analysis
Celligner

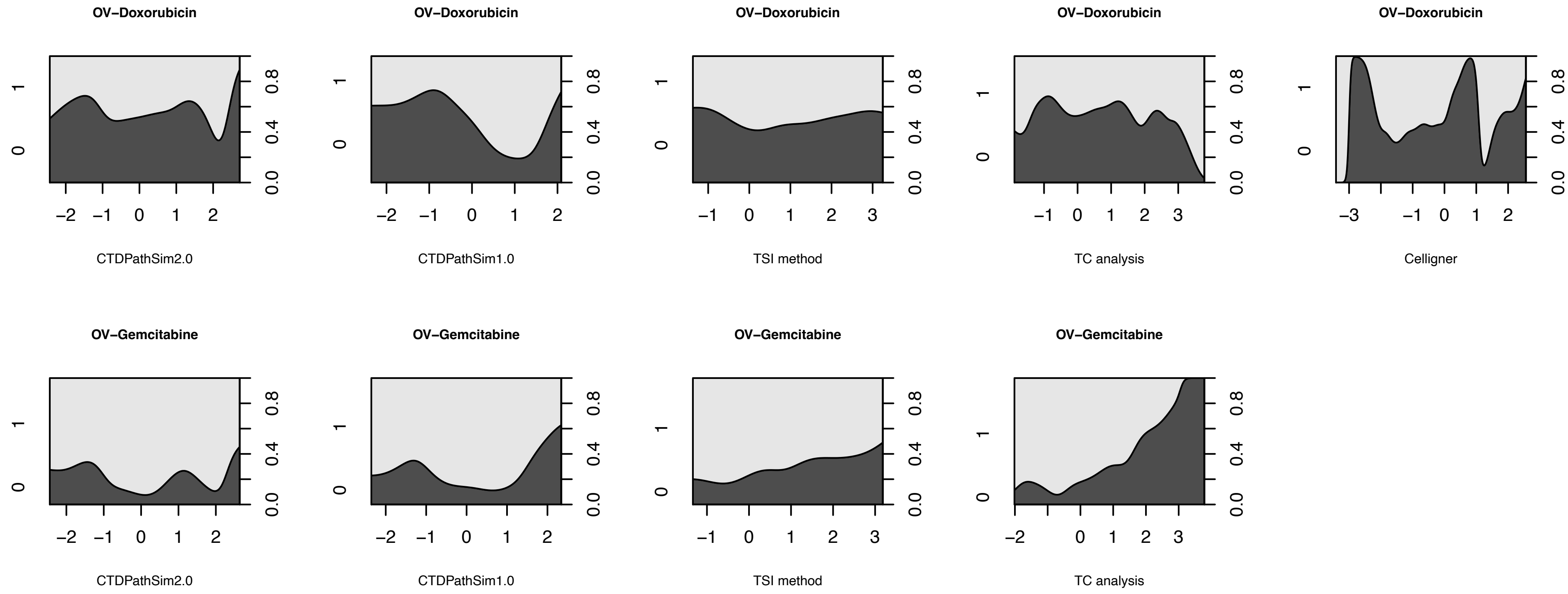
OV−Doxorubicin
CTDPathSim2.0
OV−Doxorubicin
CTDPathSim1.0
OV−Doxorubicin
TSI method
OV−Doxorubicin
TC analysis
OV−Doxorubicin
Celligner
OV−Gemcitabine
CTDPathSim2.0
OV−Gemcitabine
CTDPathSim1.0
OV−Gemcitabine
TSI method
OV−Gemcitabine
TC analysis

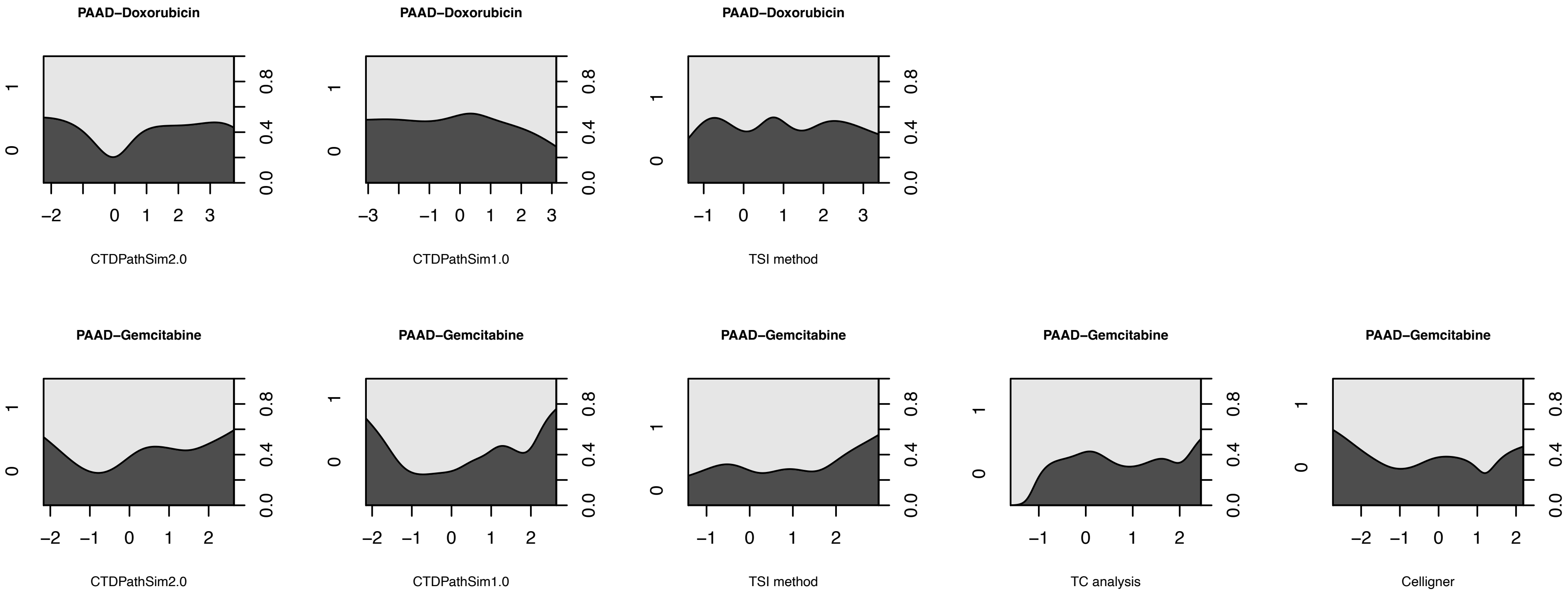
PAAD−Doxorubicin
CTDPathSim2.0
PAAD−Doxorubicin
CTDPathSim1.0
PAAD−Doxorubicin
TSI method
PAAD−Gemcitabine
CTDPathSim2.0
PAAD−Gemcitabine
CTDPathSim1.0
PAAD−Gemcitabine
TSI method
PAAD−Gemcitabine
TC analysis
PAAD−Gemcitabine
Celligner

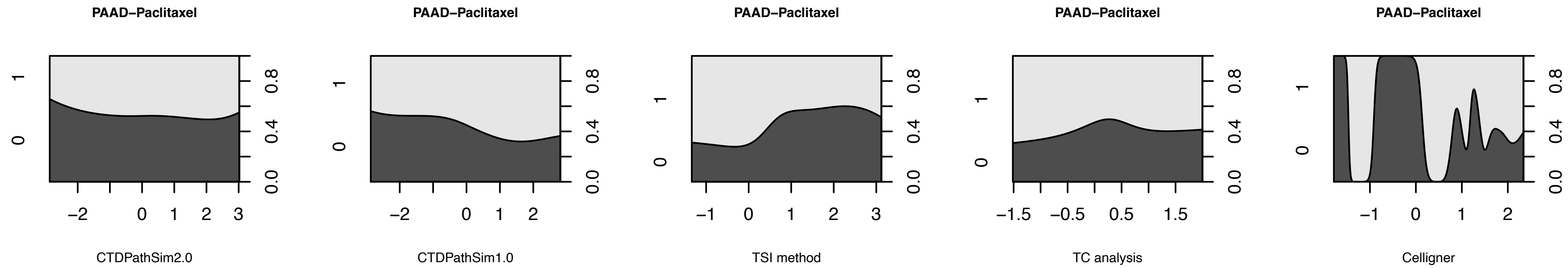

PAAD−Paclitaxel
CTDPathSim2.0
PAAD−Paclitaxel
CTDPathSim1.0
PAAD−Paclitaxel
TSI method
PAAD−Paclitaxel
TC analysis
PAAD−Paclitaxel
Celligner

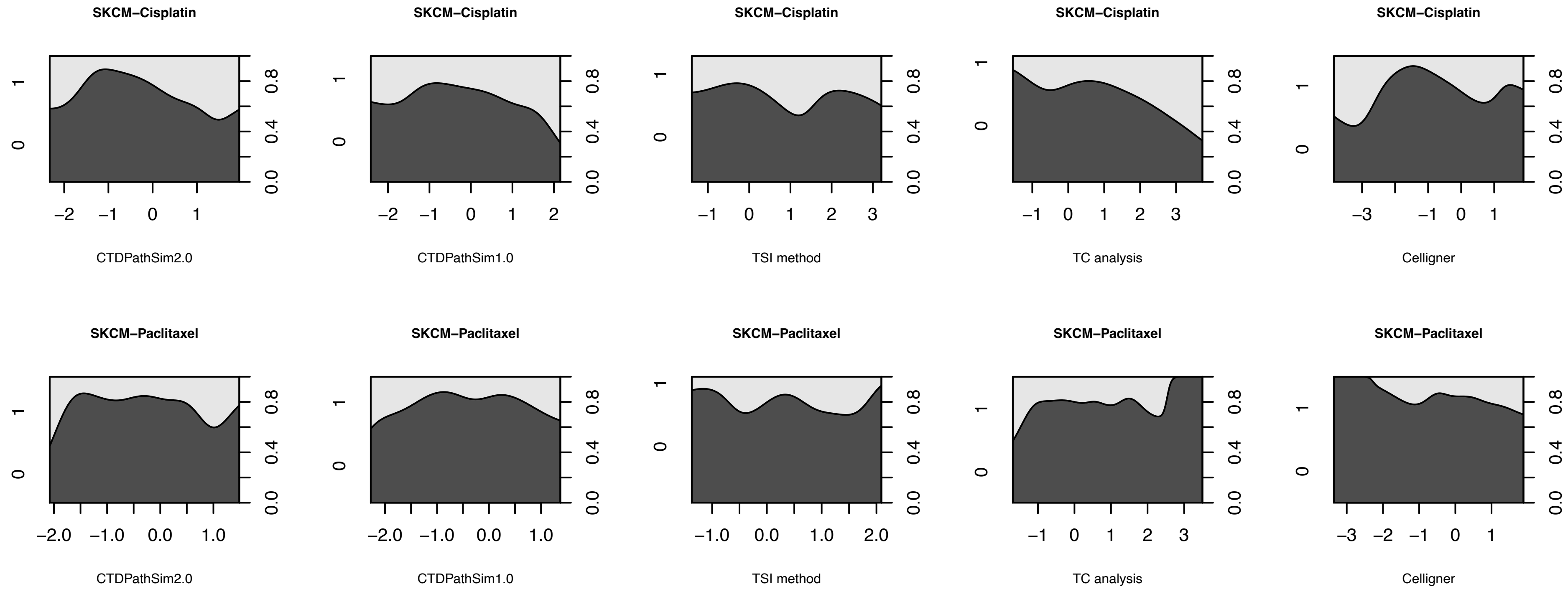
SKCM−Cisplatin
CTDPathSim2.0
SKCM−Cisplatin
CTDPathSim1.0
SKCM−Cisplatin
TSI method
SKCM−Cisplatin
TC analysis
SKCM−Cisplatin
Celligner
SKCM−Paclitaxel
CTDPathSim2.0
SKCM−Paclitaxel
CTDPathSim1.0
SKCM−Paclitaxel
TSI method
SKCM−Paclitaxel
TC analysis
SKCM−Paclitaxel
Celligner

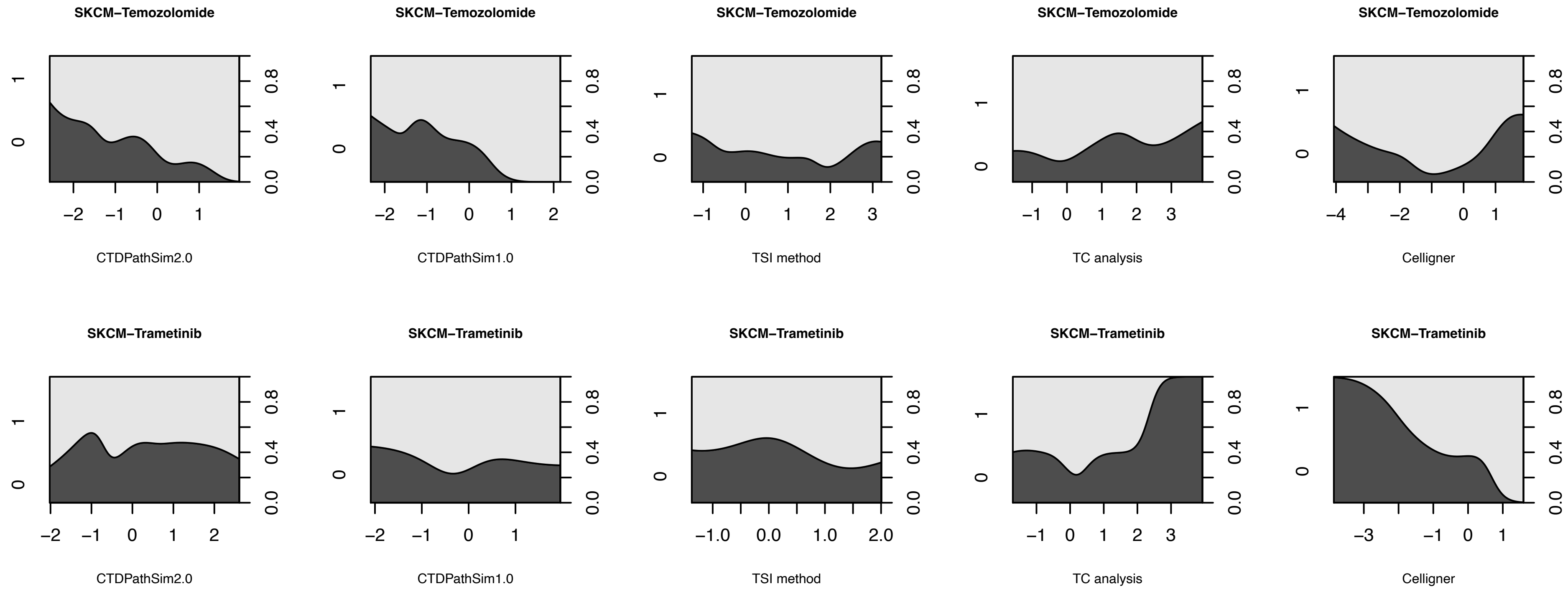
SKCM−Temozolomide
CTDPathSim2.0
CTDPathSim1.0
TSI method
TC analysis
Celligner
SKCM−Trametinib
CTDPathSim2.0
CTDPathSim1.0
TSI method
TC analysis
Celligner

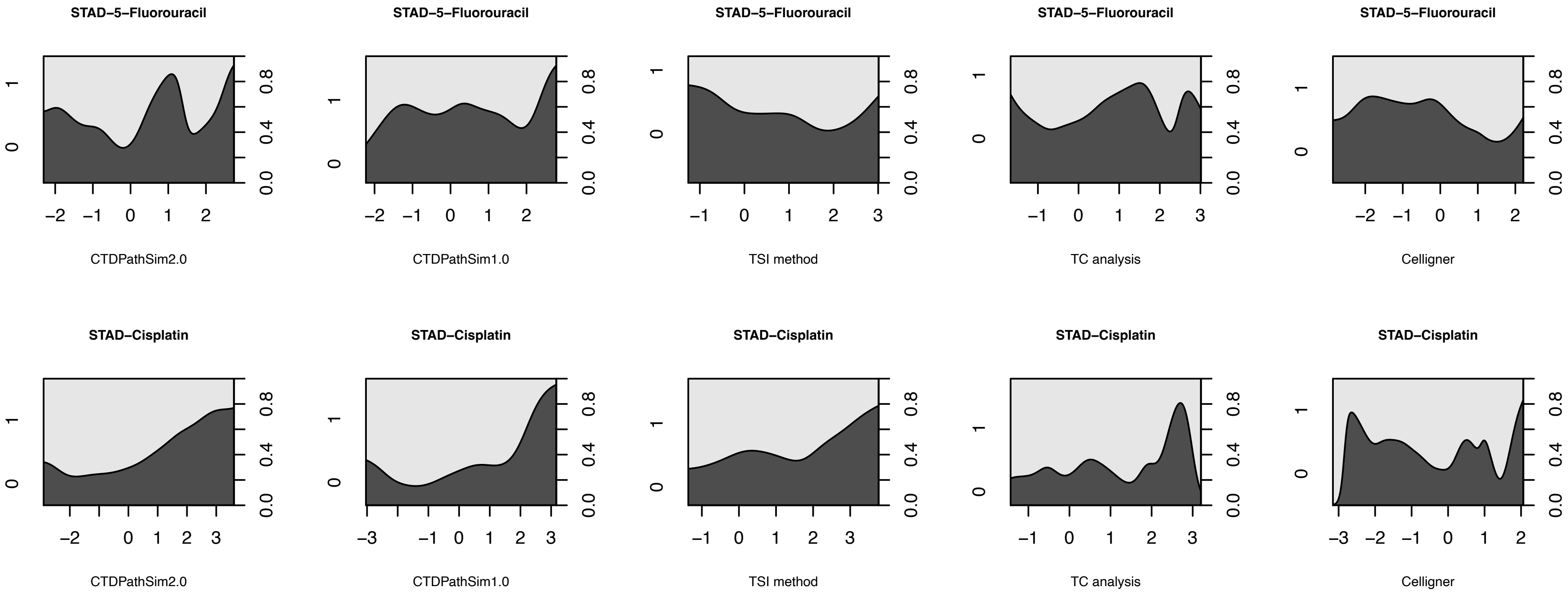

STAD−5−Fluorouracil
CTDPathSim2.0
CTDPathSim1.0
TSI method
TC analysis
Celligner
STAD−Cisplatin
CTDPathSim2.0
CTDPathSim1.0
TSI method
TC analysis
Celligner

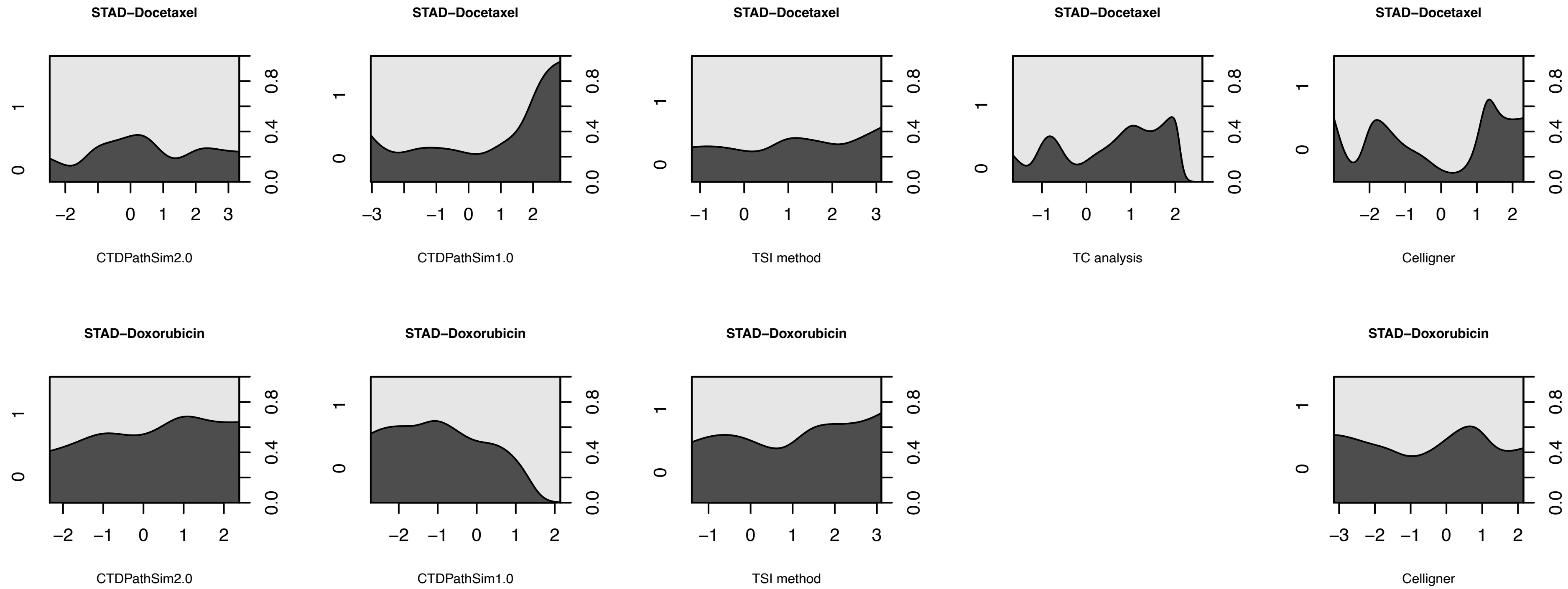
STAD−Docetaxel
CTDPathSim2.0
STAD−Docetaxel
CTDPathSim1.0
STAD−Docetaxel
TSI method
STAD−Docetaxel
TC analysis
STAD−Docetaxel
Celligner
STAD−Doxorubicin
CTDPathSim2.0
STAD−Doxorubicin
CTDPathSim1.0
STAD−Doxorubicin
TSI method
STAD−Doxorubicin
Celligner

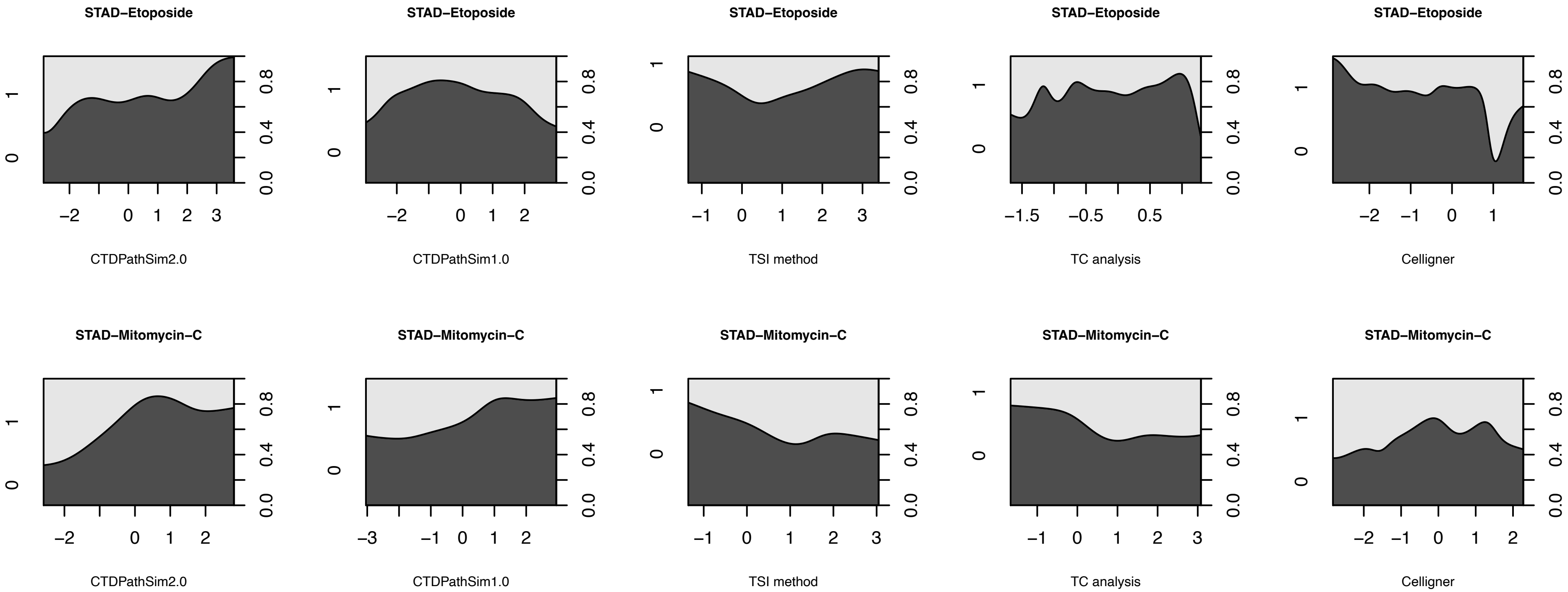

STAD−Etoposide
CTDPathSim2.0
CTDPathSim1.0
TSI method
TC analysis
Celligner
STAD−Mitomycin−C

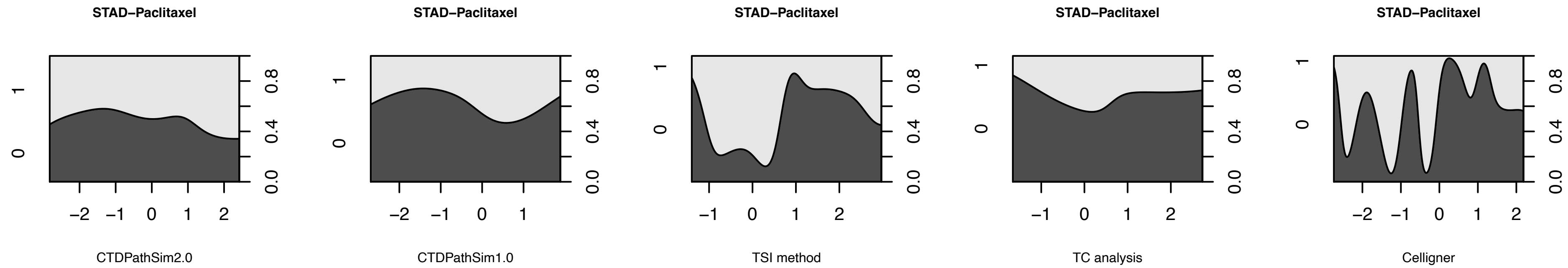
STAD−Paclitaxel
STAD−Paclitaxel
STAD−Paclitaxel
STAD−Paclitaxel
STAD−Paclitaxel
CTDPathSim2.0
CTDPathSim1.0
TSI method
TC analysis
Celligner

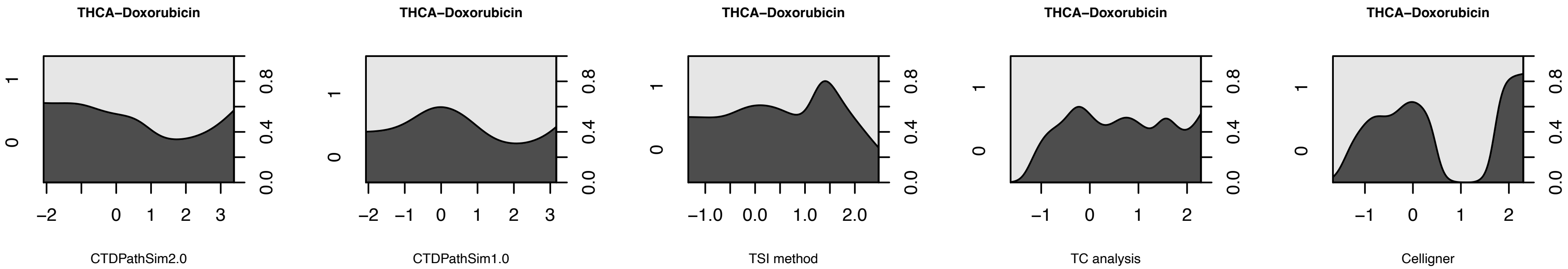
THCA–Doxorubicin
CTDPathSim2.0
THCA–Doxorubicin
CTDPathSim1.0
THCA–Doxorubicin
TSI method
THCA–Doxorubicin
TC analysis
THCA–Doxorubicin
Celligner
