## Supplemental Figure S2 for "Finding the best cell lines across pan-cancer to use in pre-clinical research as a proxy for patient tumor samples considering immune cells, multi-omics, and cancer pathways"

**TCGA cancer type**
**No. of tissue-specific cell lines in low similarity group vs high similarity group:10/15; t-test (alternative = “less”) p-value = 0.001**

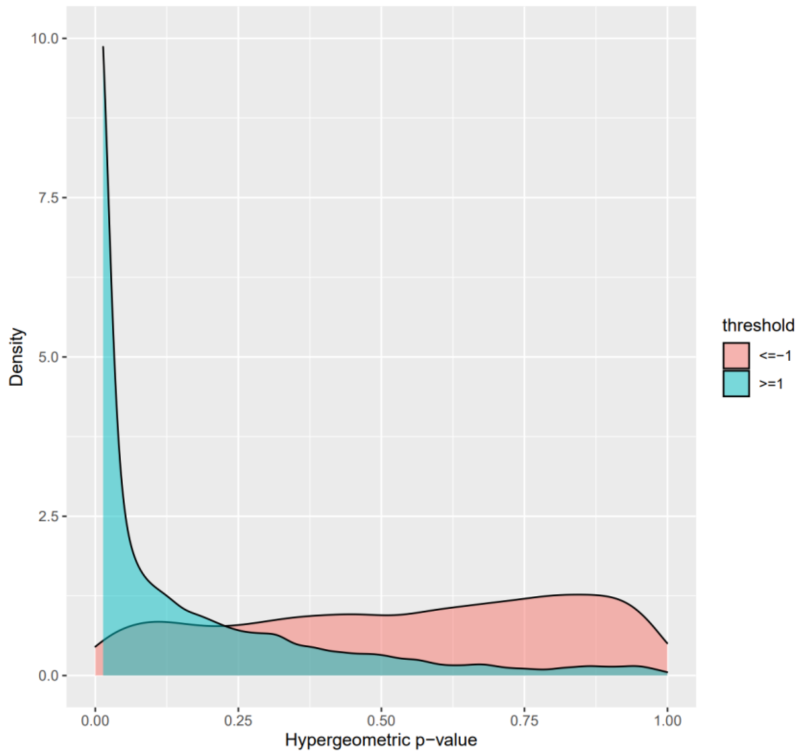


**Density plots of hypergeometric p-values of enrichment for known tissue-specific cancer cell lines for the five methods**. Enrichment of cancer tissue-specific cell lines in highly similar and lowly similar cell lines. The t-test (alternative = “less”) p-values comparing each pair of the density plots in each cancer type are in red. NA stands for not applicable in cases where t-test could not be run due to insufficient data points between two groups.

### CTDPathSim2.0

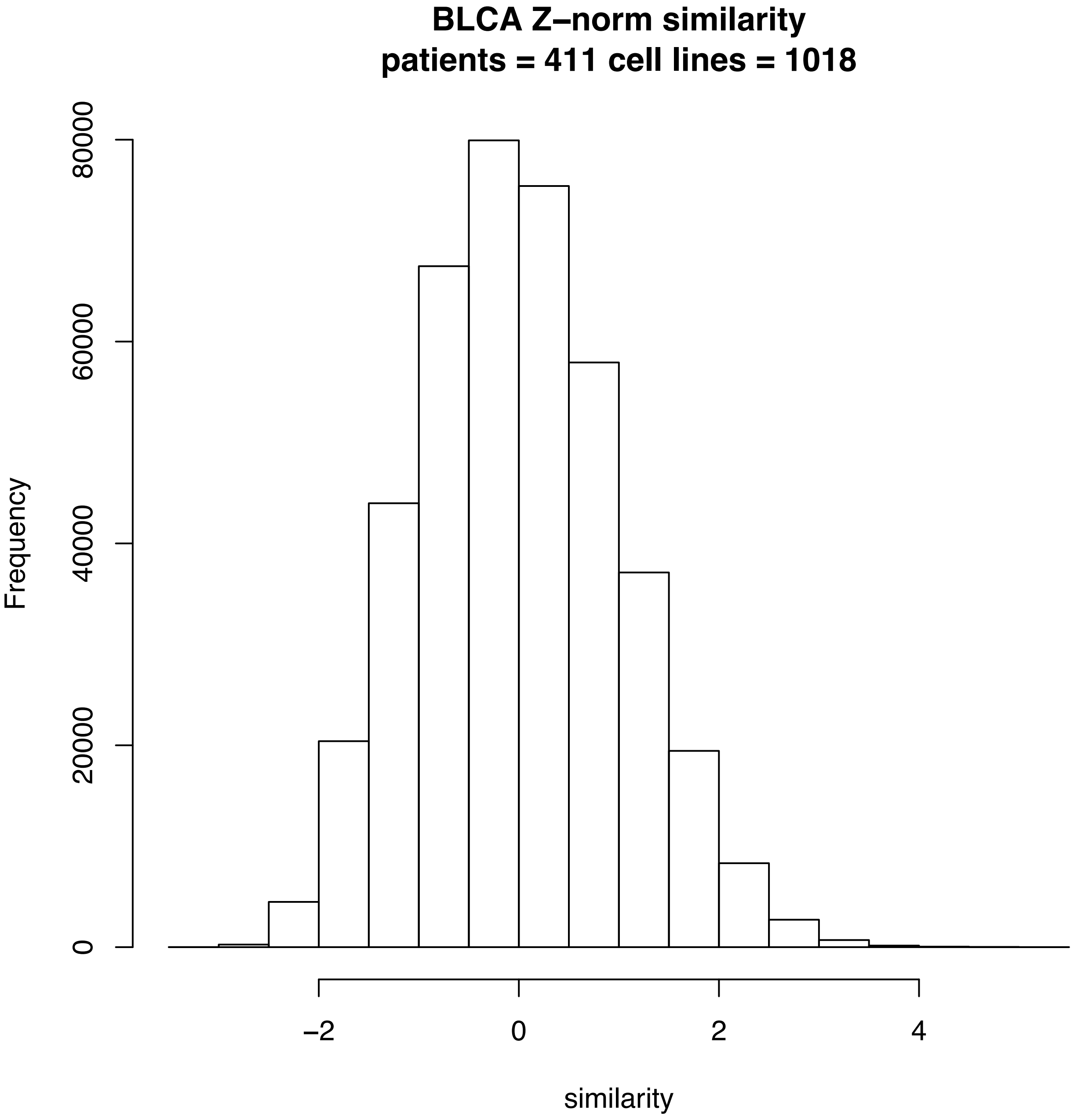
BLCA Z–norm similarity
patients = 411 cell lines = 1018
Frequency
0
20000
40000
60000
80000
–2
0
2
4
similarity

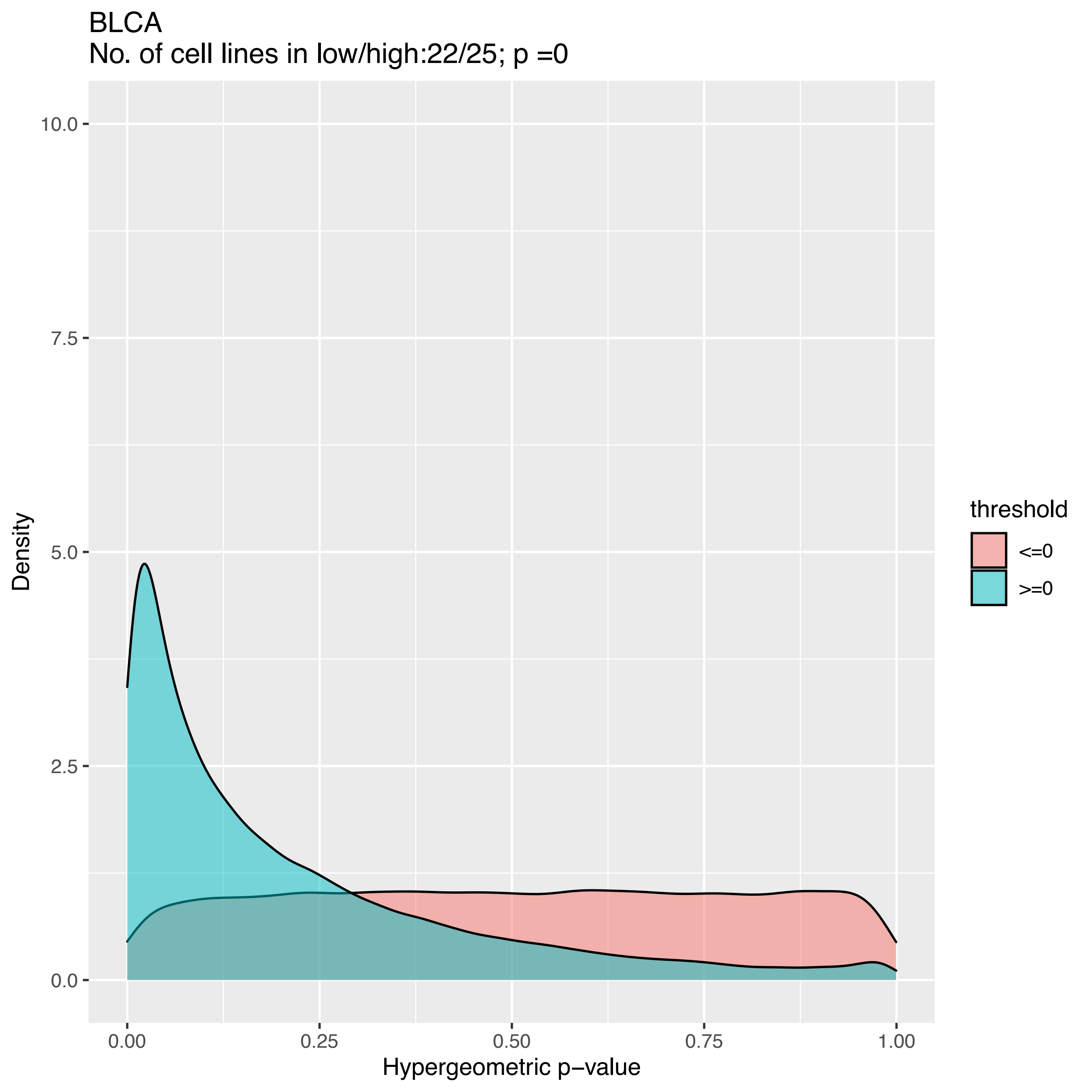
BLCA
No. of cell lines in low/high:22/25; p =0
Density
Hypergeometric p−value
threshold
<=0
>=0
0.0
2.5
5.0
7.5
10.0
0.00
0.25
0.50
0.75
1.00

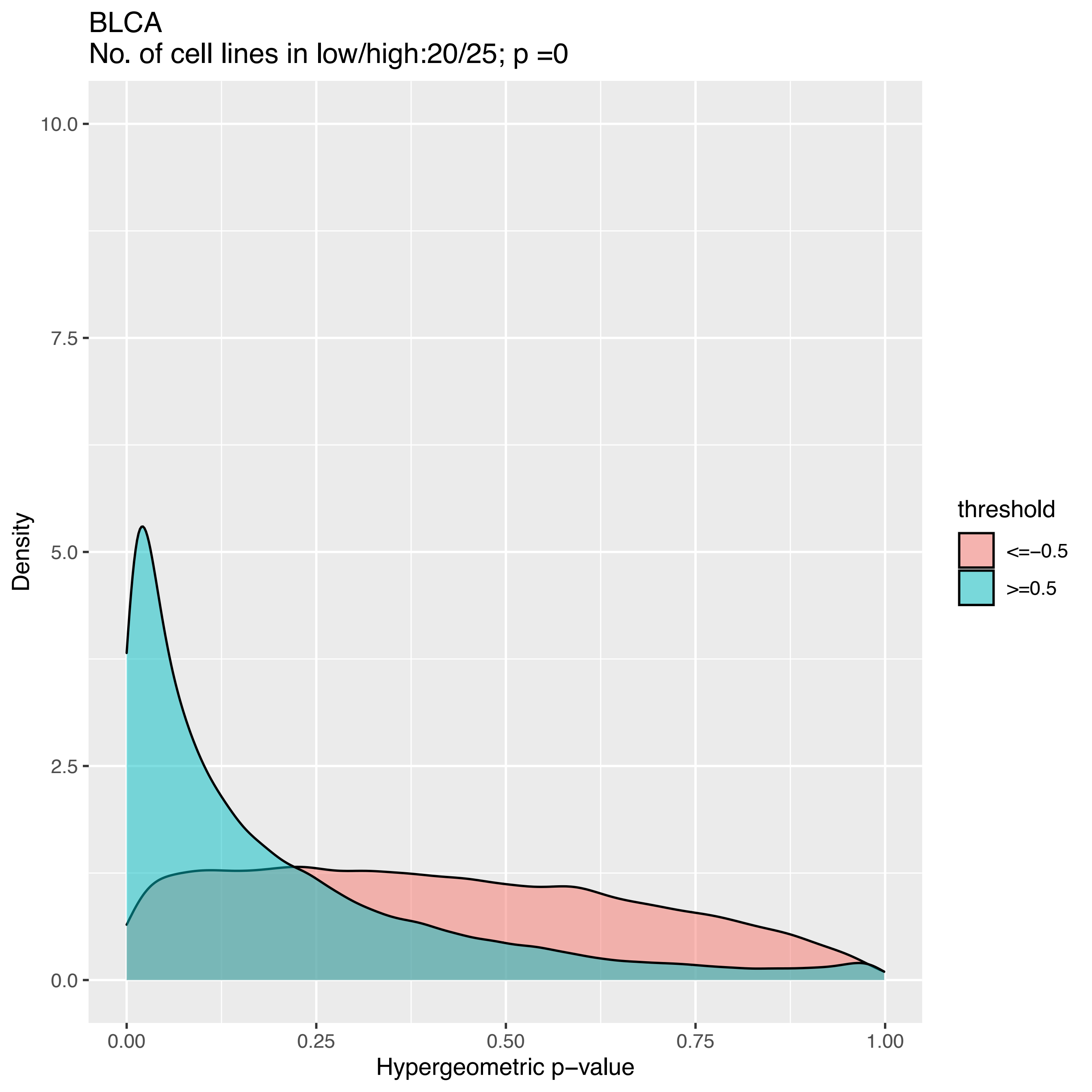

BLCA
No. of cell lines in low/high:20/25; p =0
10.0
7.5
5.0
2.5
0.0
Density
0.00
0.25
0.50
0.75
1.00
Hypergeometric p−value
threshold
<=−0.5
>=0.5

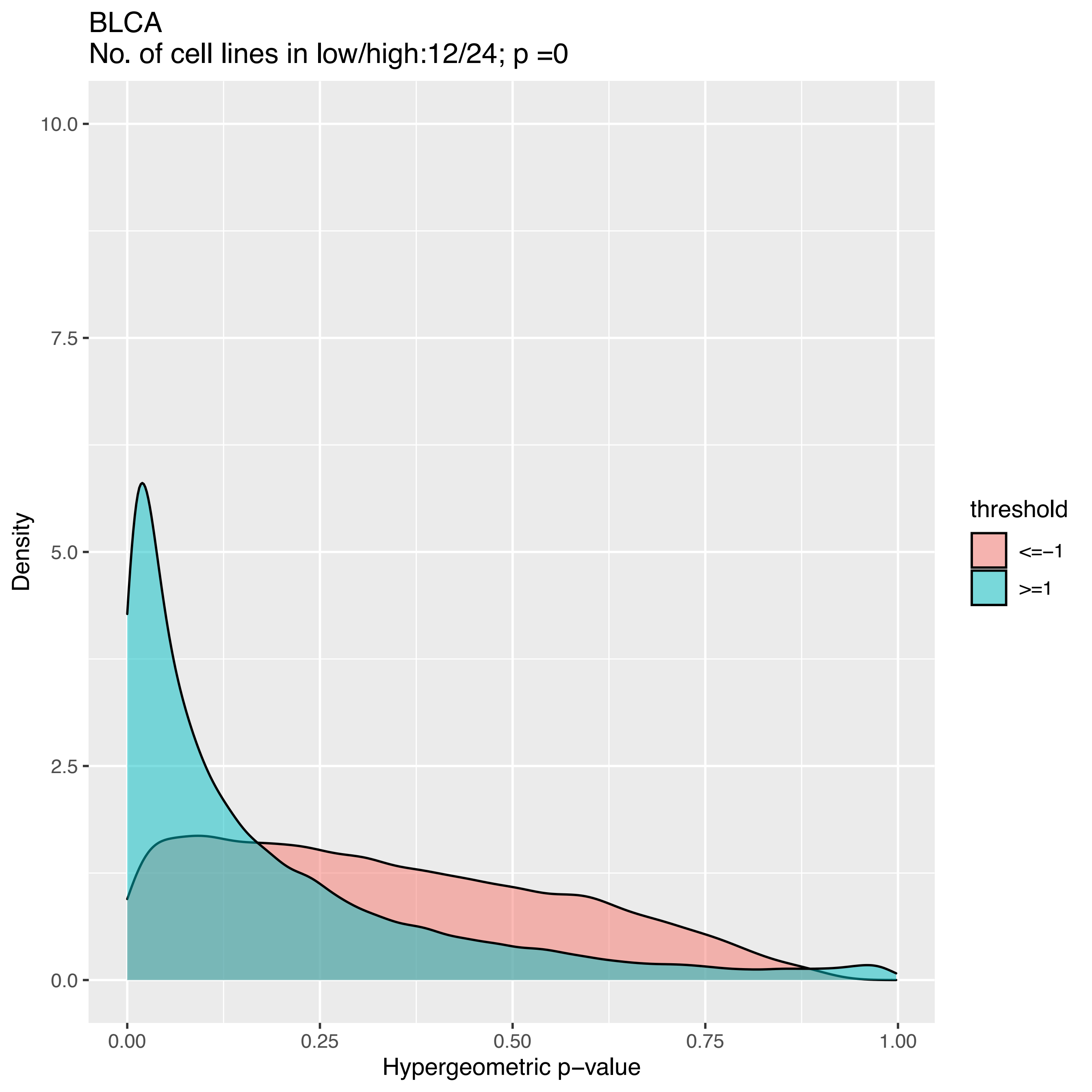
BLCA
No. of cell lines in low/high:12/24; p =0
Density
Hypergeometric p−value
10.0
7.5
5.0
2.5
0.0
0.00
0.25
0.50
0.75
1.00
threshold
<=−1
>=1

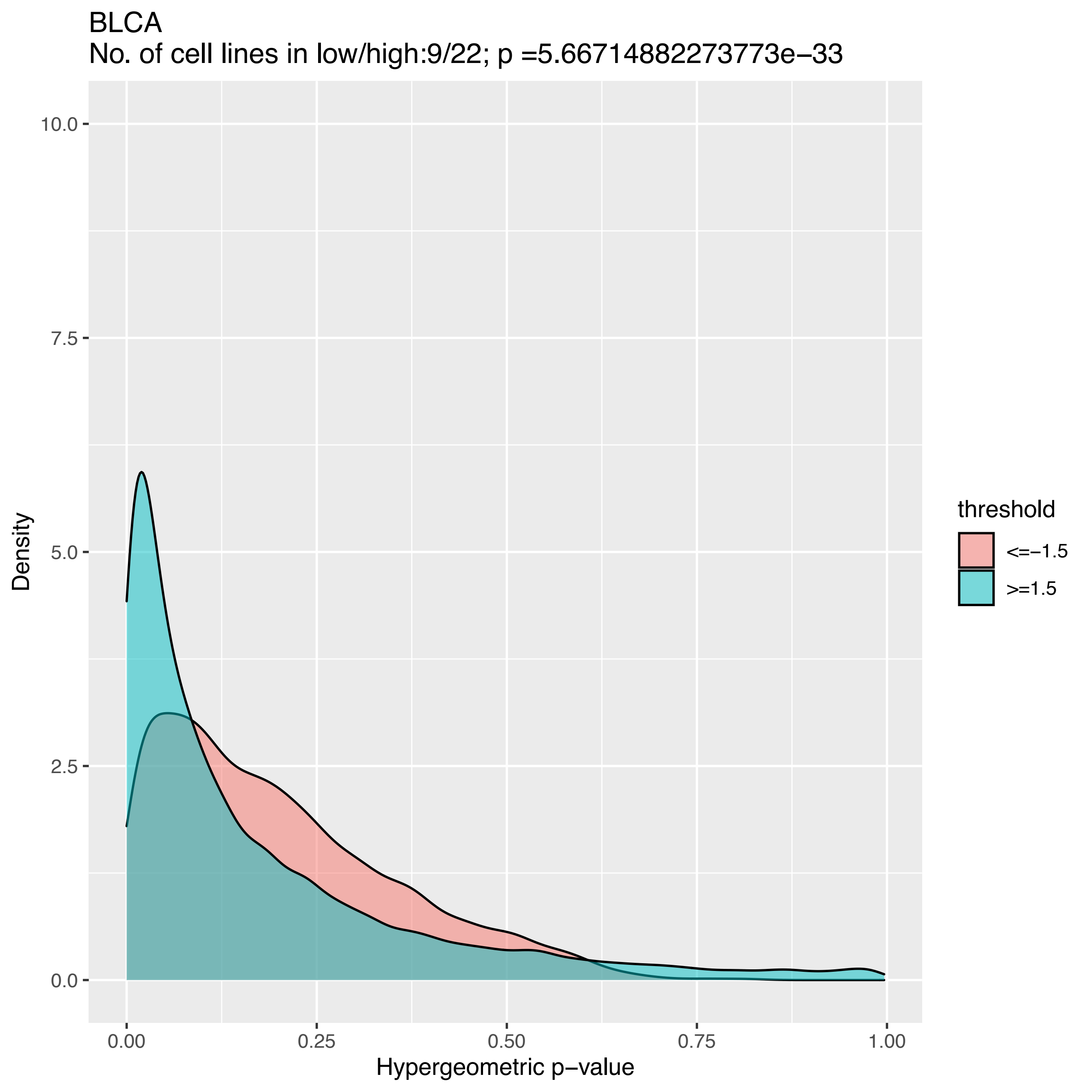
BLCA
No. of cell lines in low/high:9/22; p =5.66714882273773e−33
10.0
7.5
5.0
2.5
0.0
Density
0.00
0.25
0.50
0.75
1.00
Hypergeometric p−value
threshold
<=−1.5
>=1.5

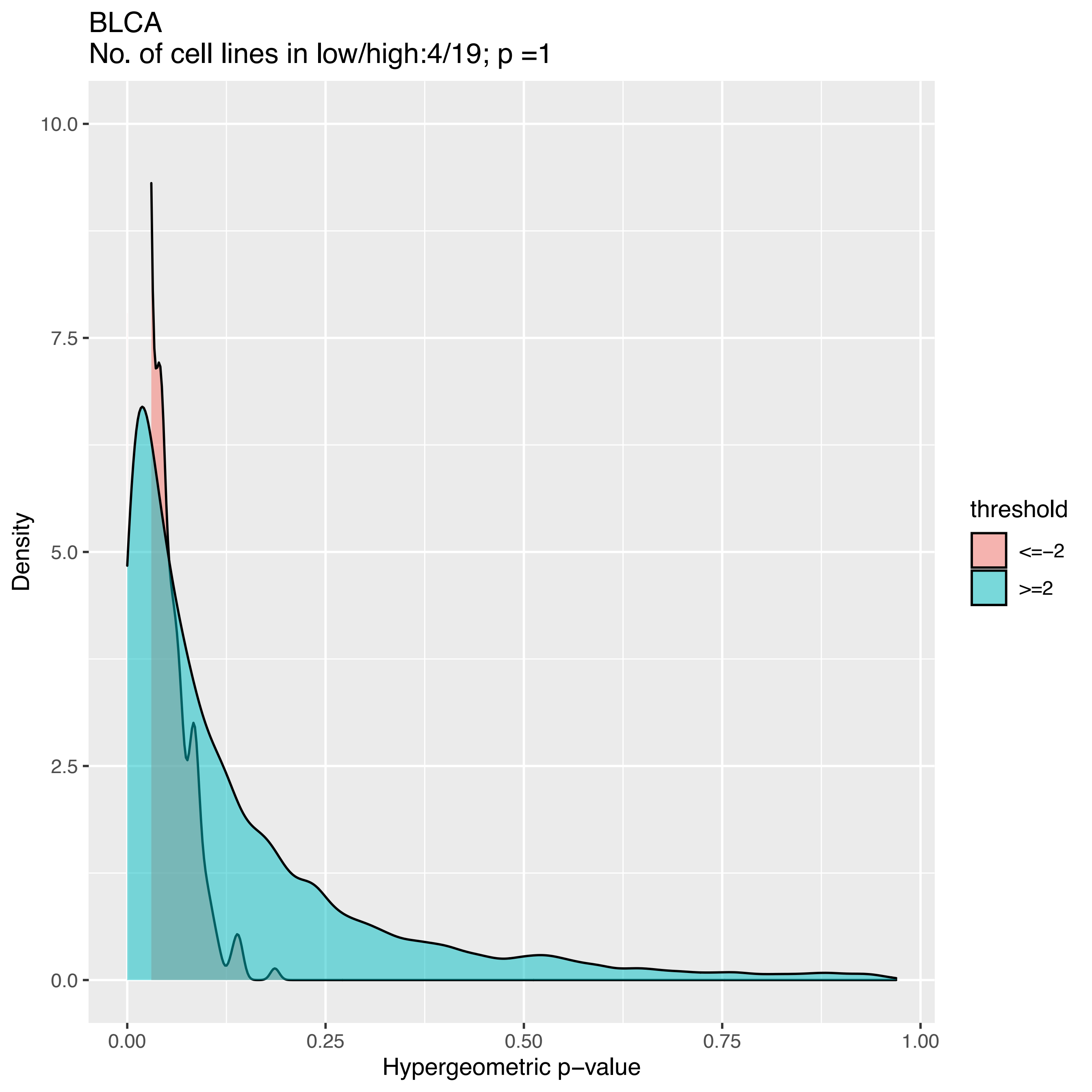
BLCA
No. of cell lines in low/high:4/19; p =1
Density
Hypergeometric p−value
10.0
7.5
5.0
2.5
0.0
0.00
0.25
0.50
0.75
1.00
threshold
<=−2
>=2

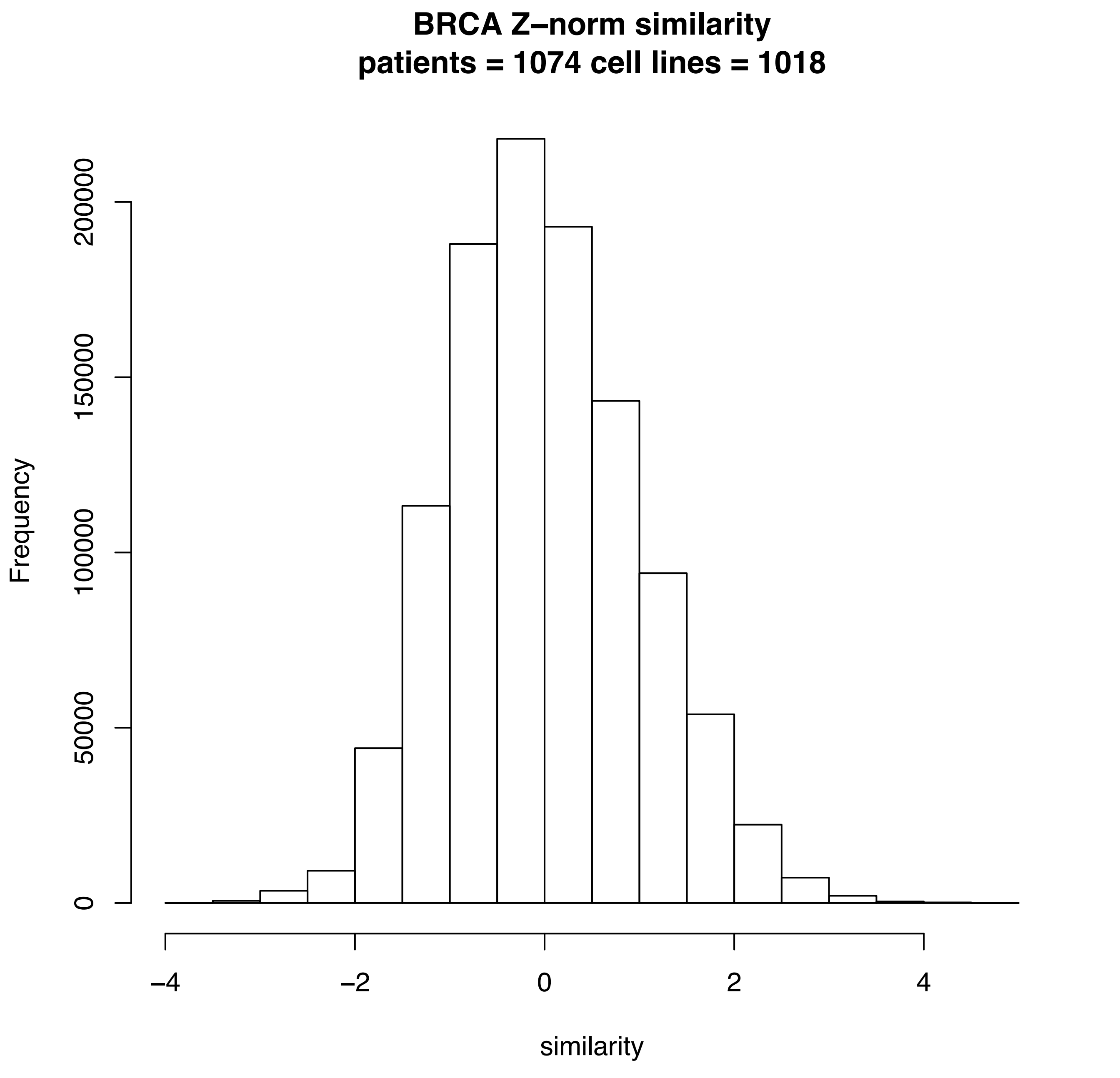
BRCA Z–norm similarity
patients = 1074 cell lines = 1018
Frequency
0
50000
100000
150000
200000
–4
–2
0
2
4
similarity

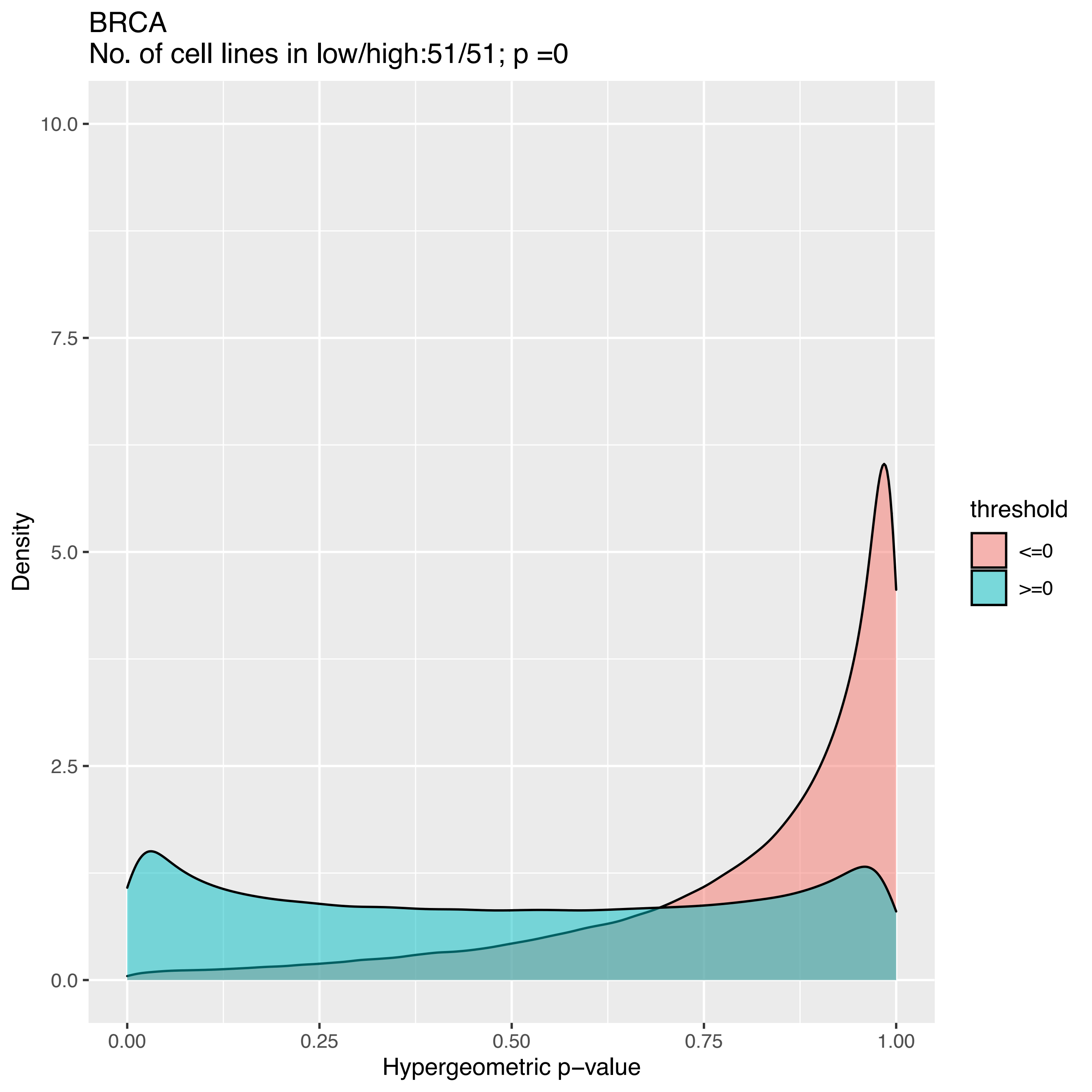

BRCA
No. of cell lines in low/high:51/51; p =0
Density
Hypergeometric p−value
threshold
<=0
>=0
10.0
7.5
5.0
2.5
0.0
0.00
0.25
0.50
0.75
1.00

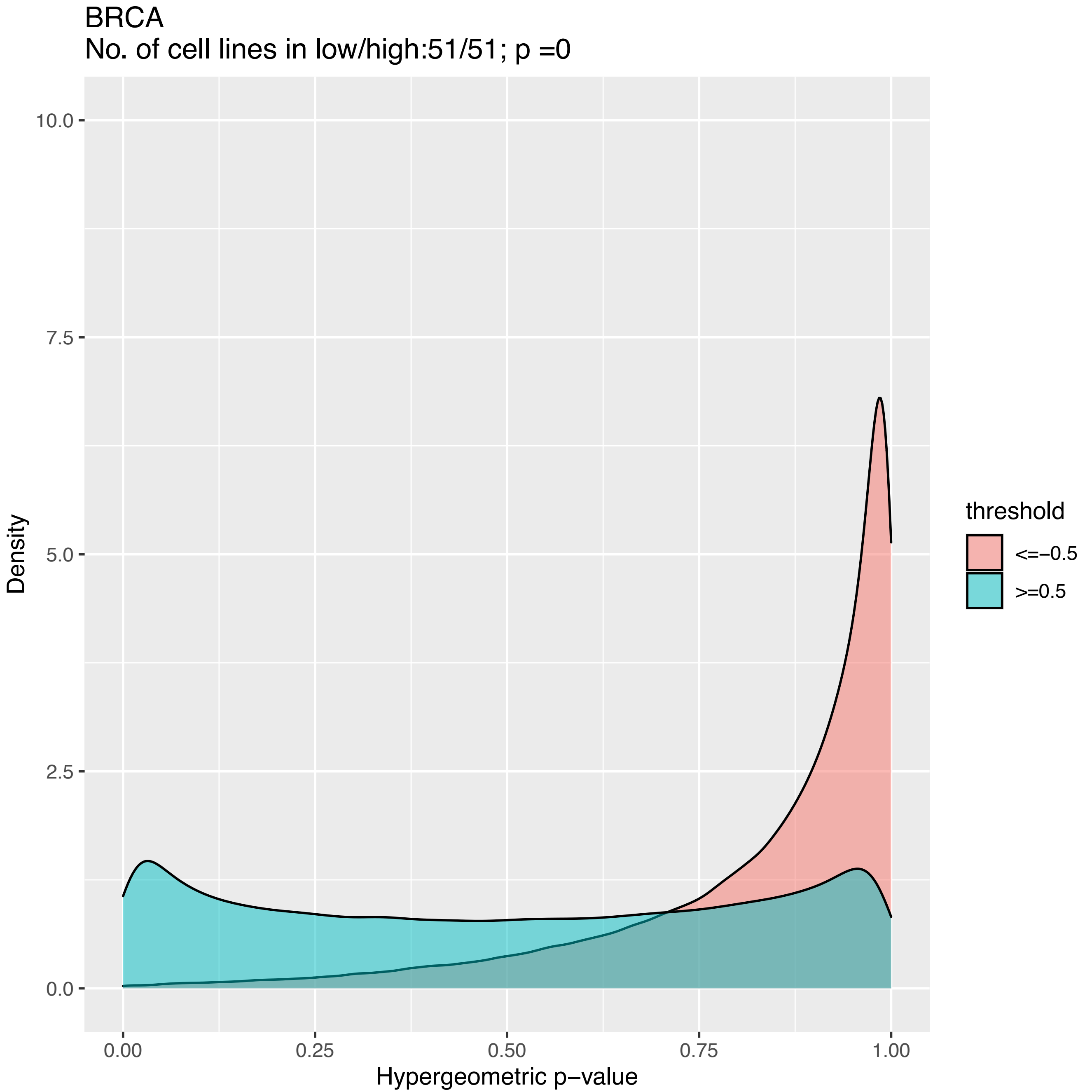
BRCA
No. of cell lines in low/high:51/51; p =0
Density
10.0
7.5
5.0
2.5
0.0
0.00
0.25
0.50
0.75
1.00
Hypergeometric p−value
threshold
<=−0.5
>=0.5

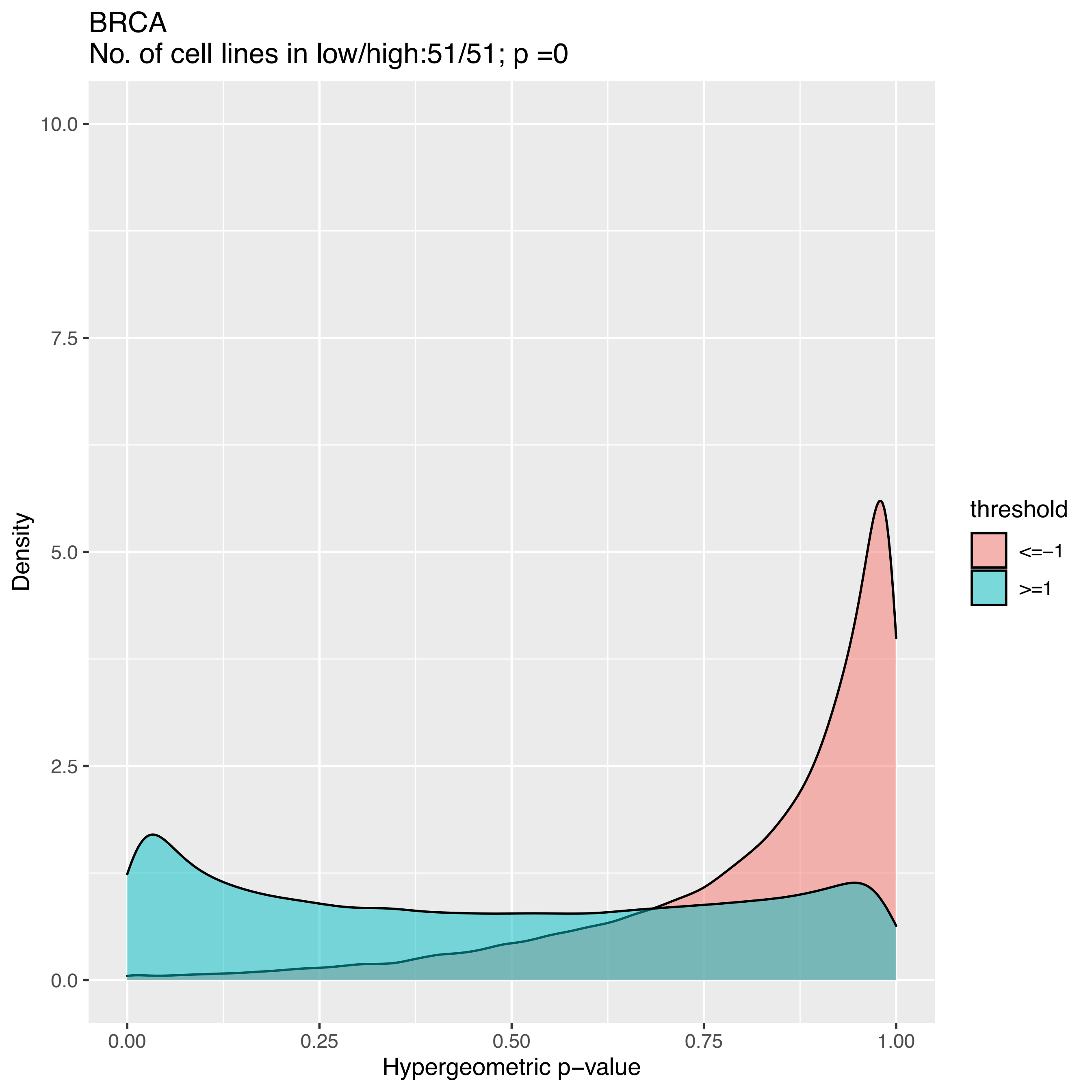

BRCA
No. of cell lines in low/high:51/51; p =0
Density
Hypergeometric p−value
10.0
7.5
5.0
2.5
0.0
0.00
0.25
0.50
0.75
1.00
threshold
<=−1
>=1

BRCA
No. of cell lines in low/high:51/48; p =0

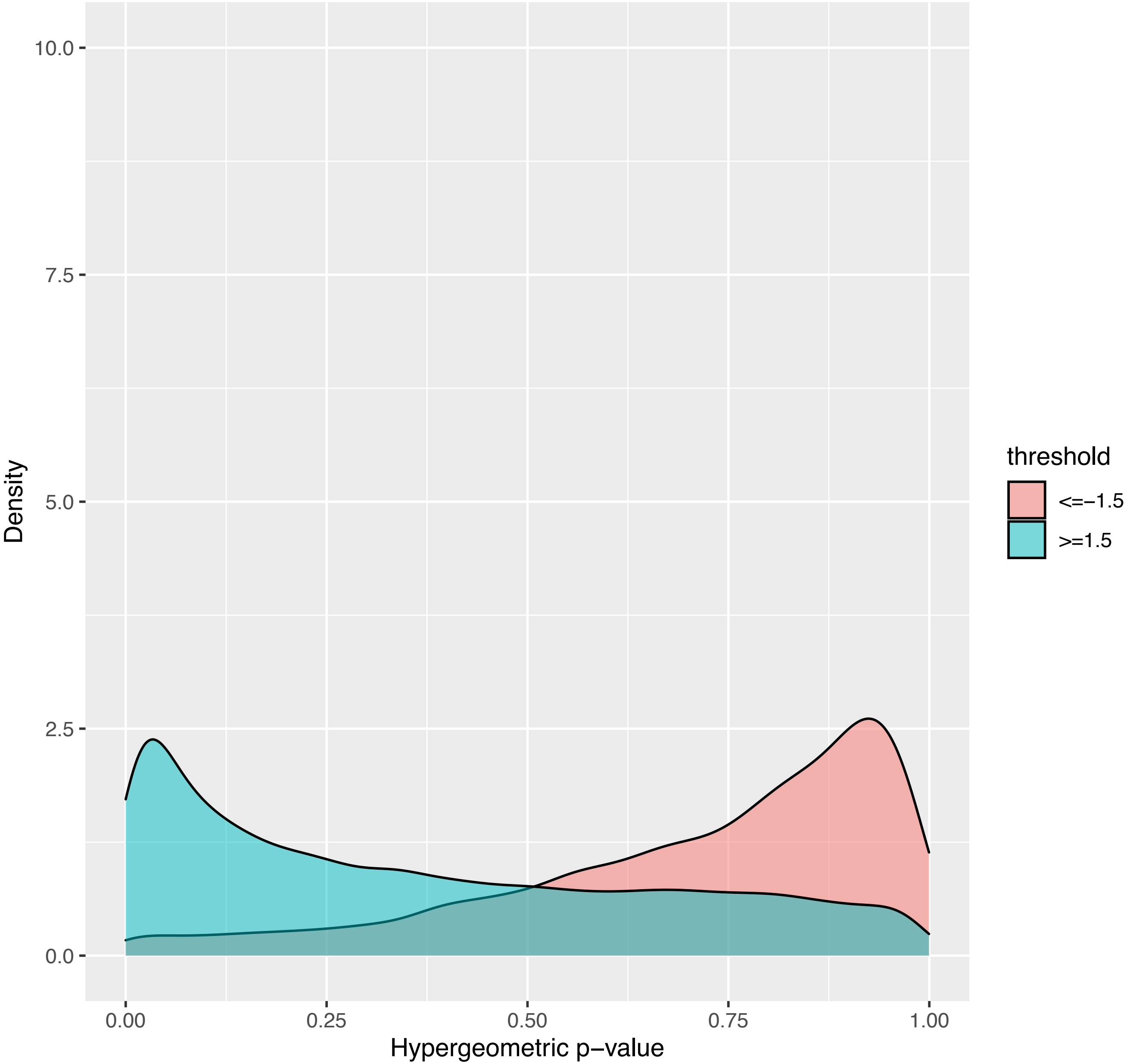

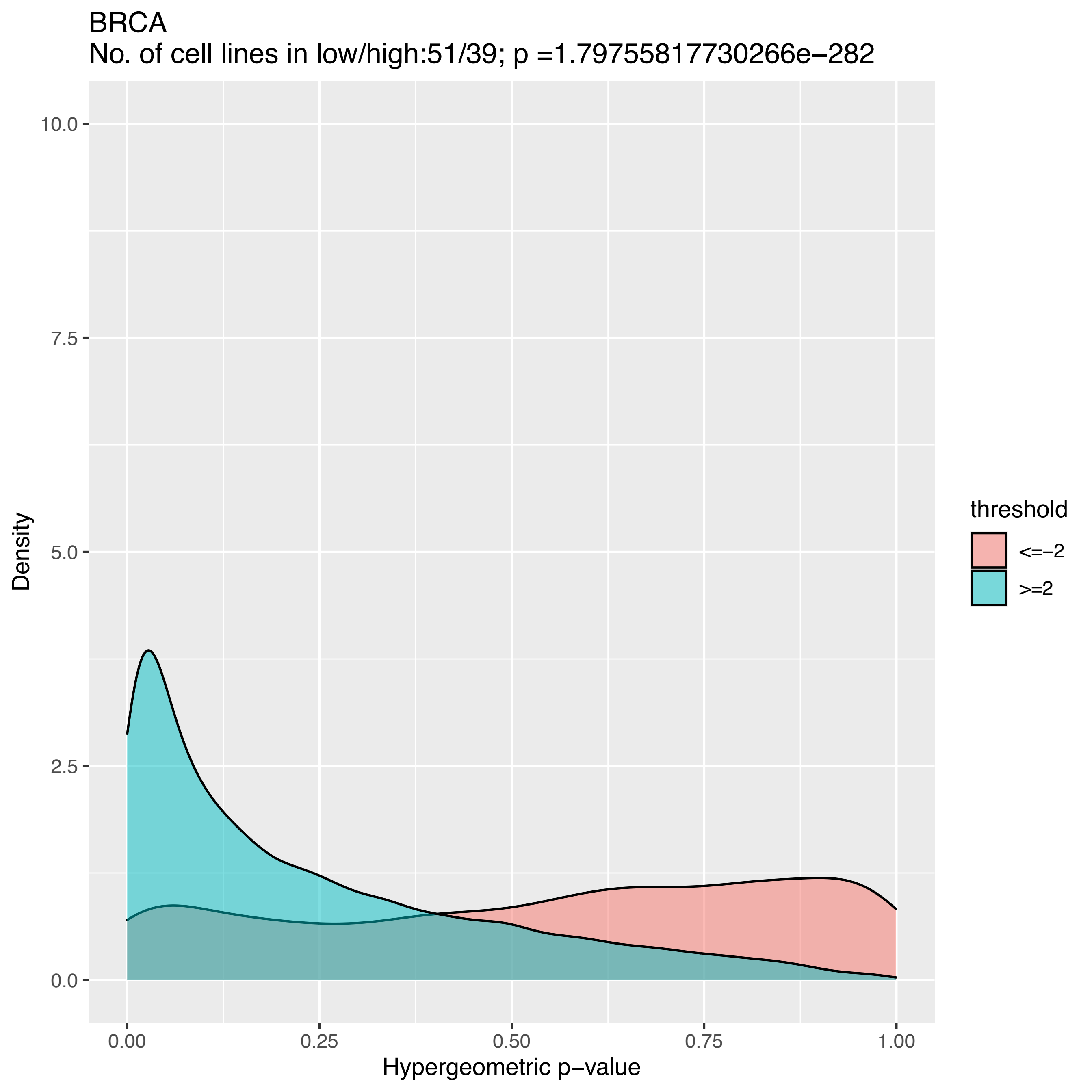
BRCA
No. of cell lines in low/high:51/39; p =1.79755817730266e−282
10.0
7.5
5.0
2.5
0.0
Density
0.00
0.25
0.50
0.75
1.00
Hypergeometric p−value
threshold
<=−2
>=2

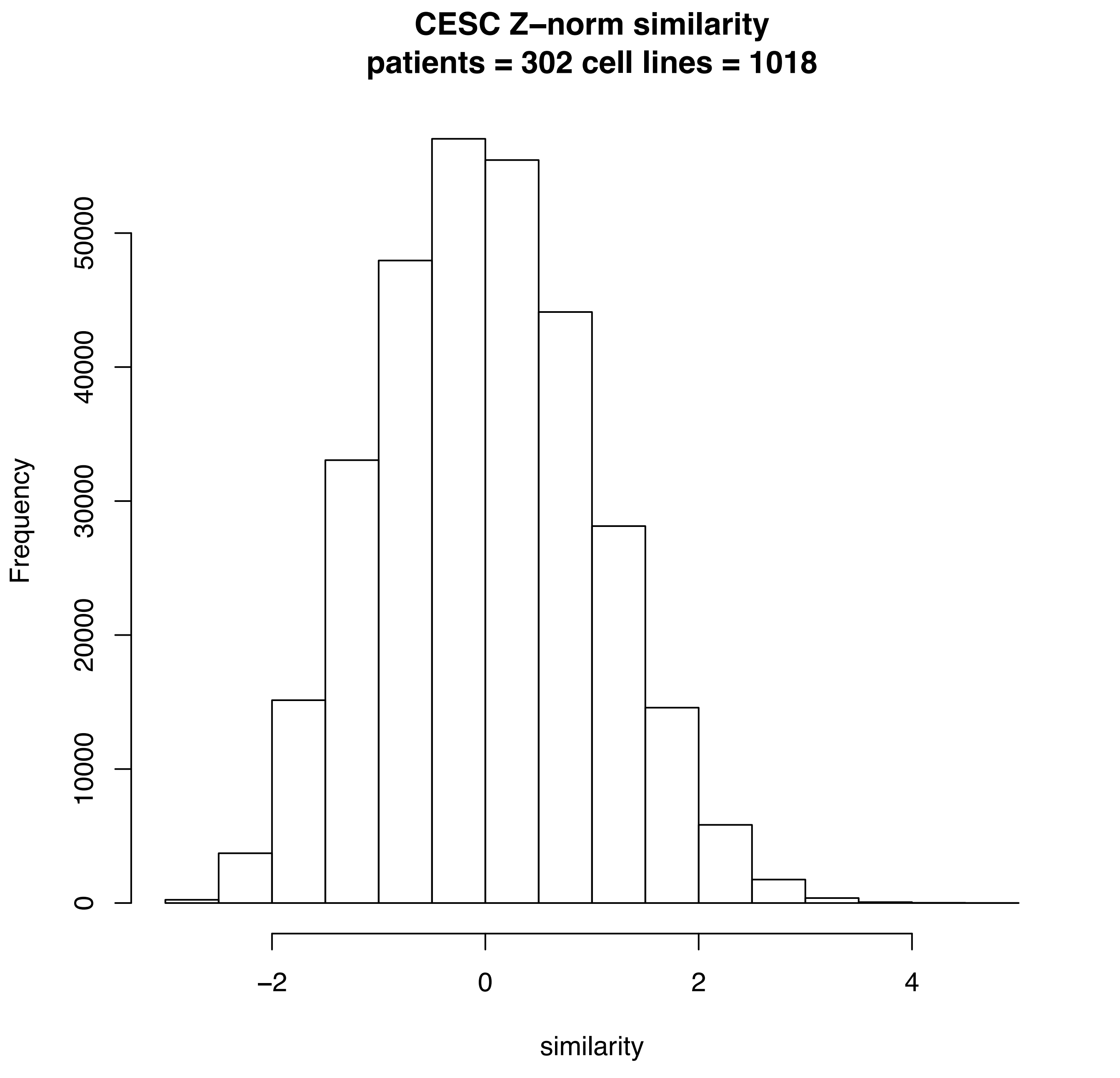
CESC Z–norm similarity
patients = 302 cell lines = 1018
Frequency
0
10000
20000
30000
40000
50000
–2
0
2
4
similarity

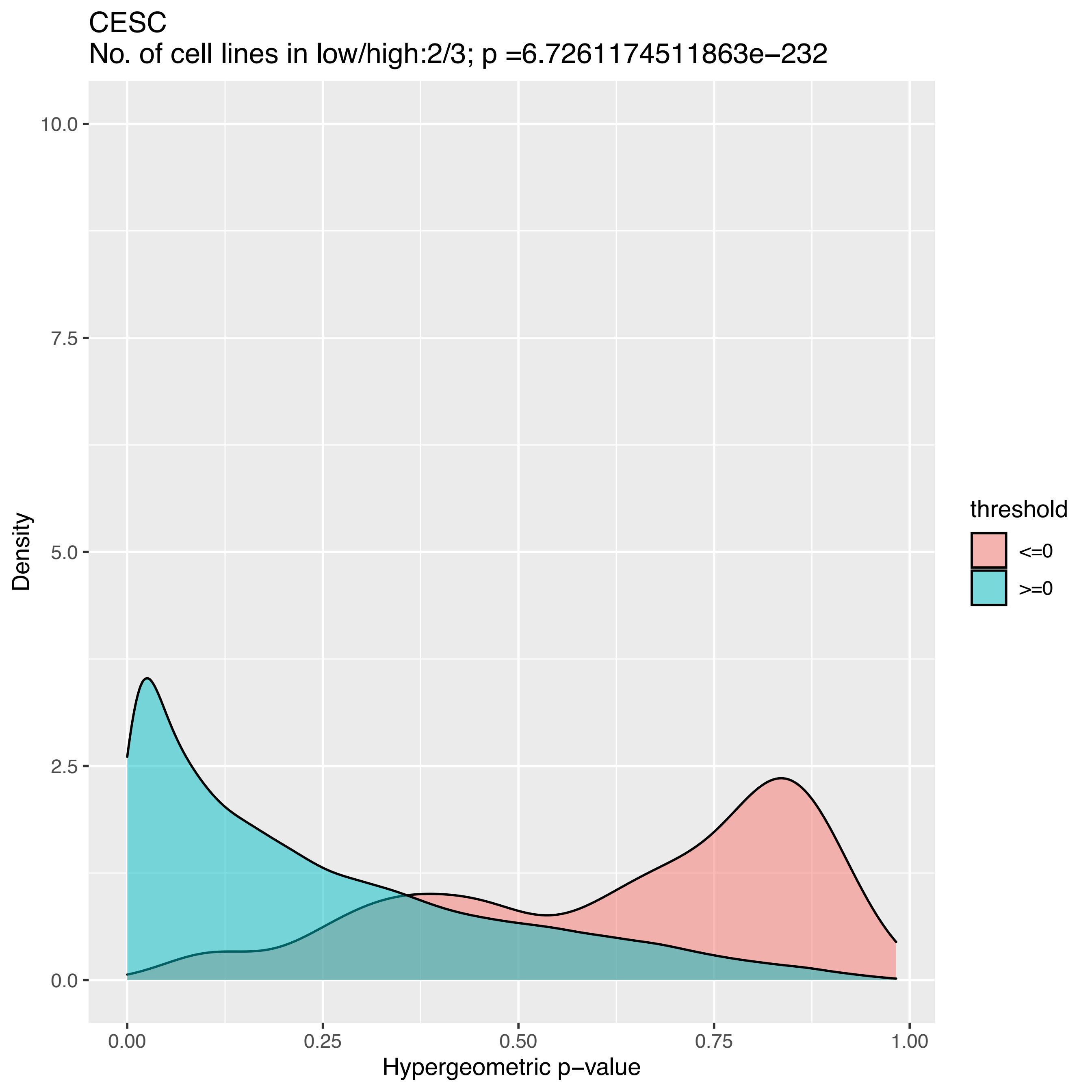
CESC
No. of cell lines in low/high:2/3; p =6.7261174511863e−232
Density
10.0
7.5
5.0
2.5
0.0
0.00
0.25
0.50
0.75
1.00
Hypergeometric p−value
threshold
<=0
>=0

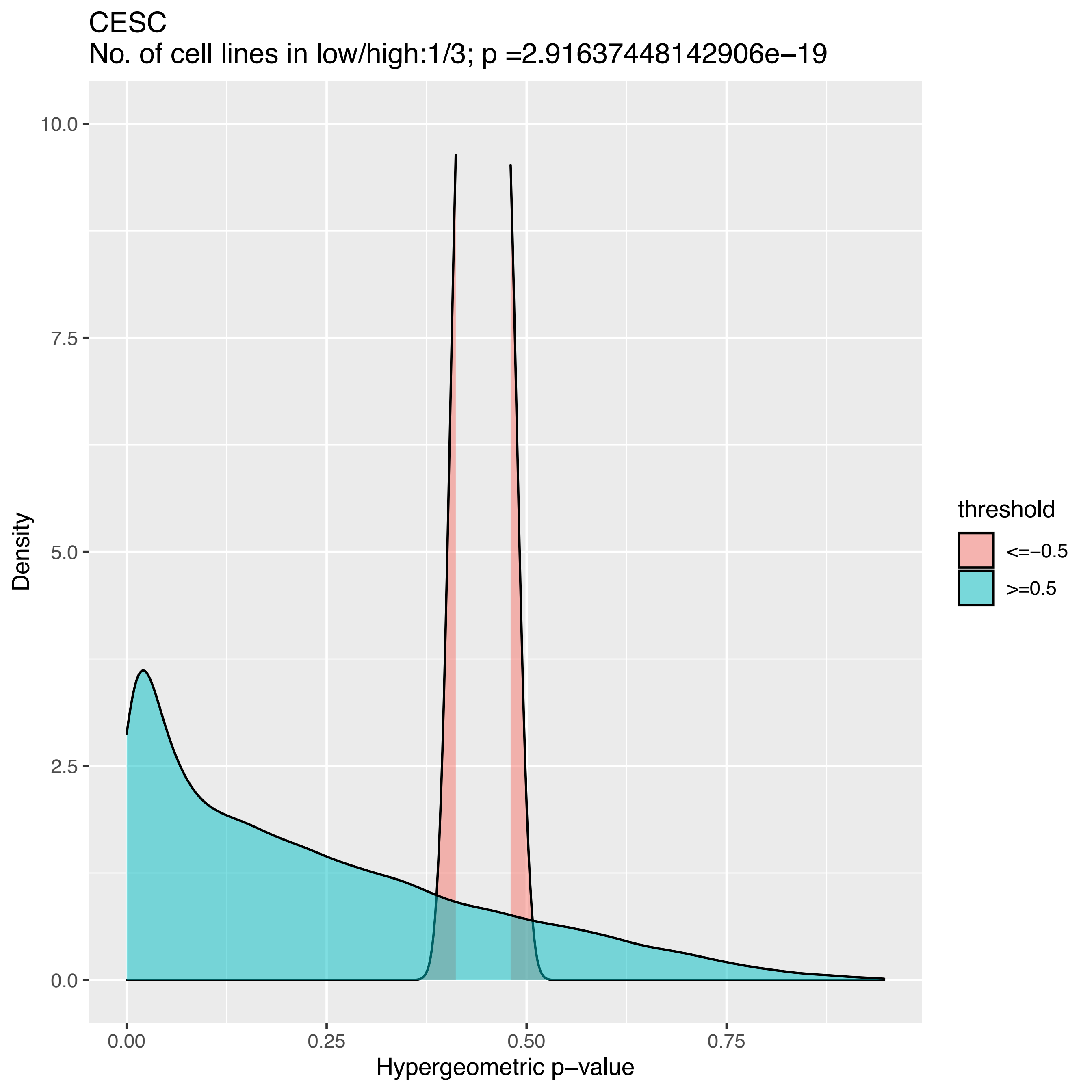
CESC
No. of cell lines in low/high:1/3; p =2.91637448142906e−19
Density
10.0
7.5
5.0
2.5
0.0
0.00
0.25
0.50
0.75
Hypergeometric p−value
threshold
<=−0.5
>=0.5

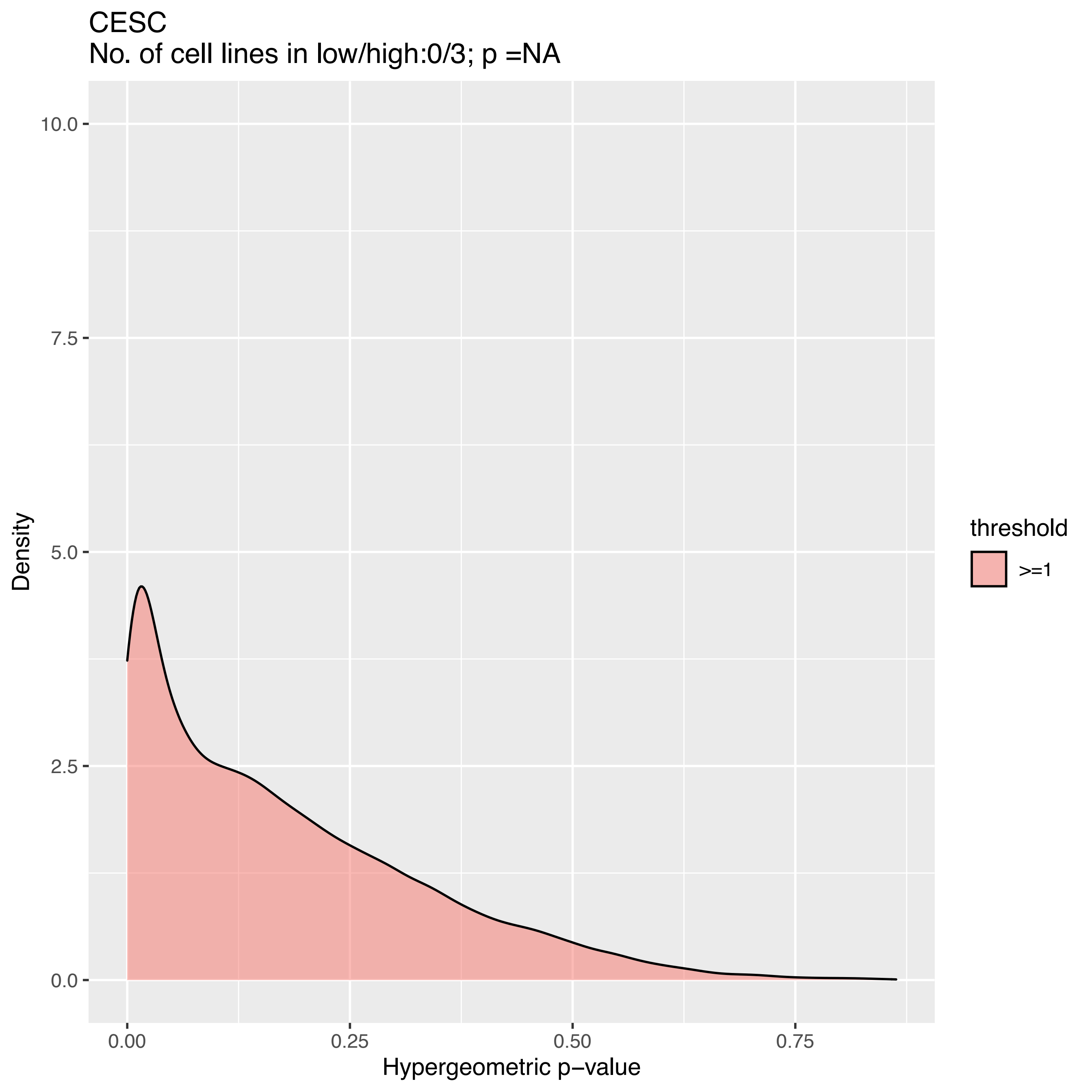
CESC
No. of cell lines in low/high:0/3; p =NA
10.0
7.5
5.0
2.5
0.0
Density
0.00
0.25
0.50
0.75
Hypergeometric p−value
threshold
>=1

CESC
No. of cell lines in low/high:0/3; p =NA

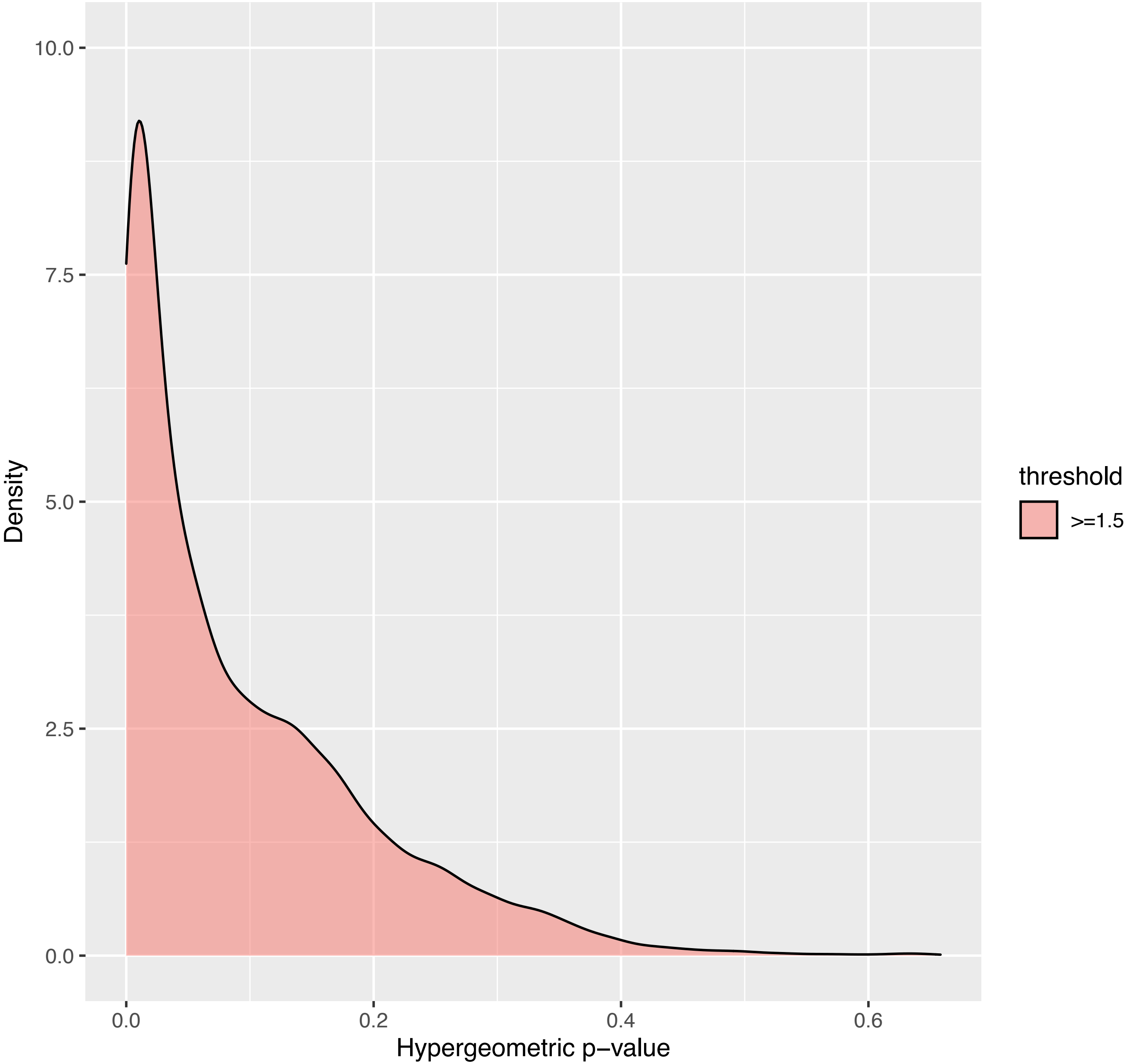

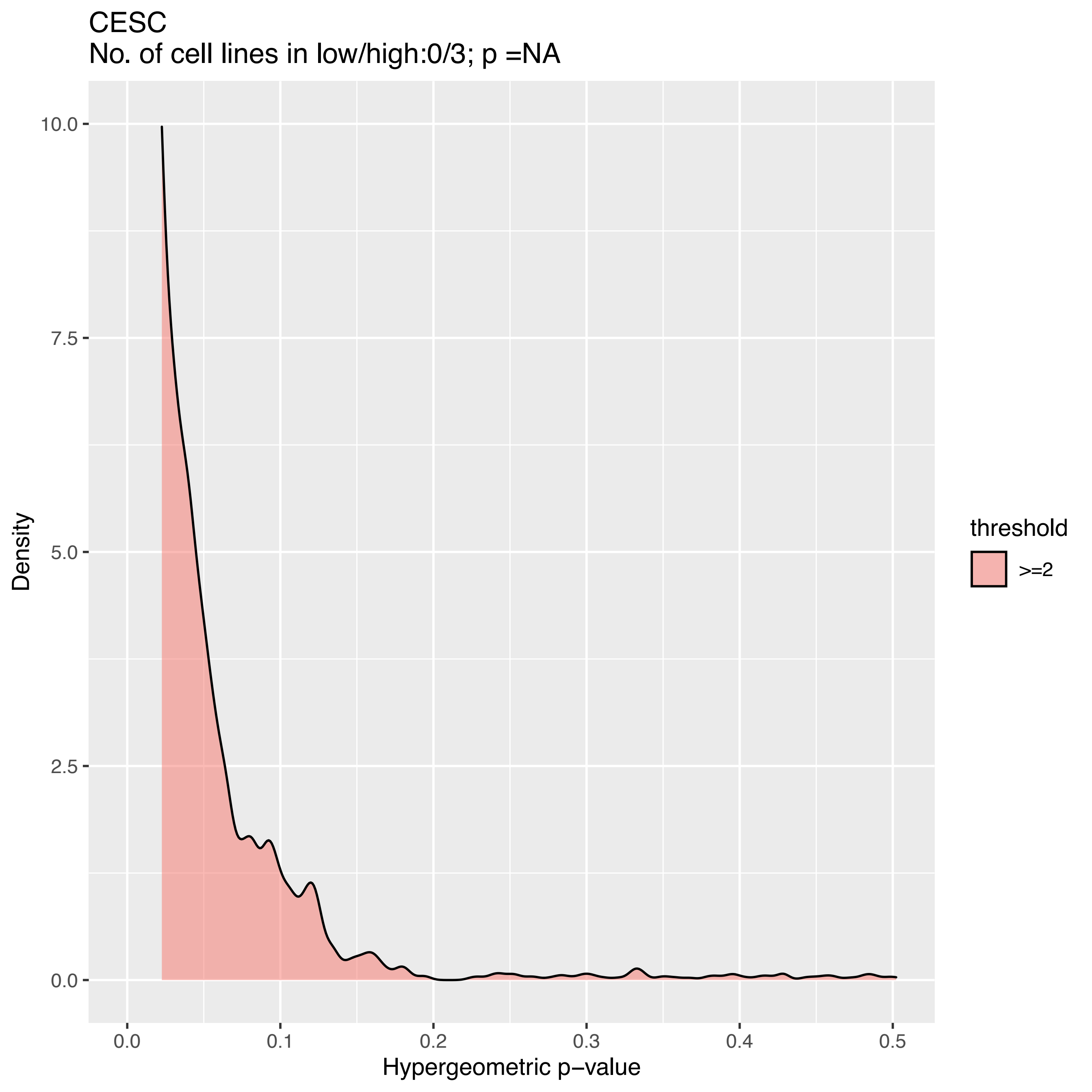
CESC
No. of cell lines in low/high:0/3; p =NA
Density
10.0
7.5
5.0
2.5
0.0
0.0
0.1
0.2
0.3
0.4
0.5
Hypergeometric p−value
threshold
>=2

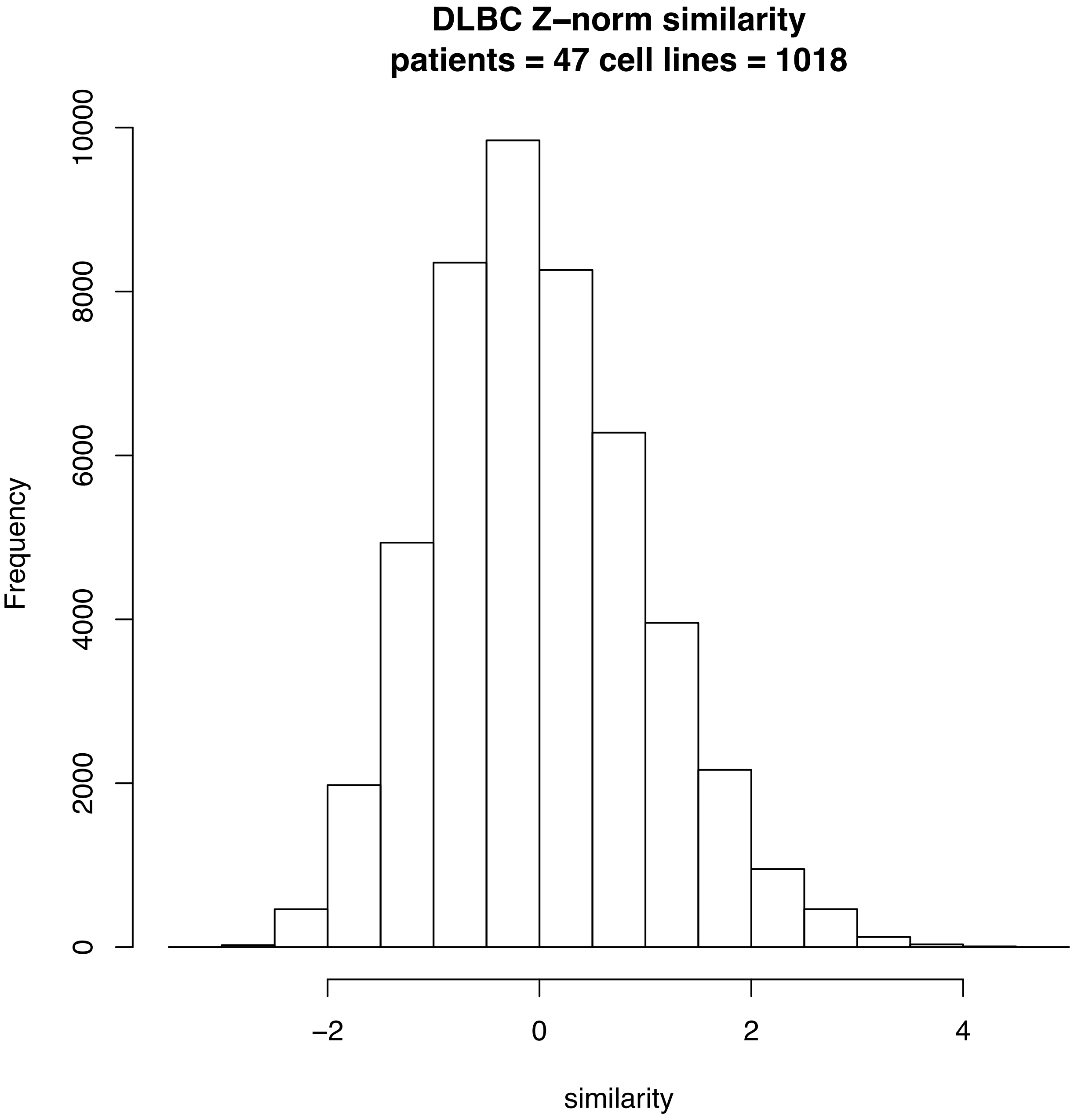
DLBC Z–norm similarity
patients = 47 cell lines = 1018
Frequency
0
2000
4000
6000
8000
10000
−2
0
2
4
similarity

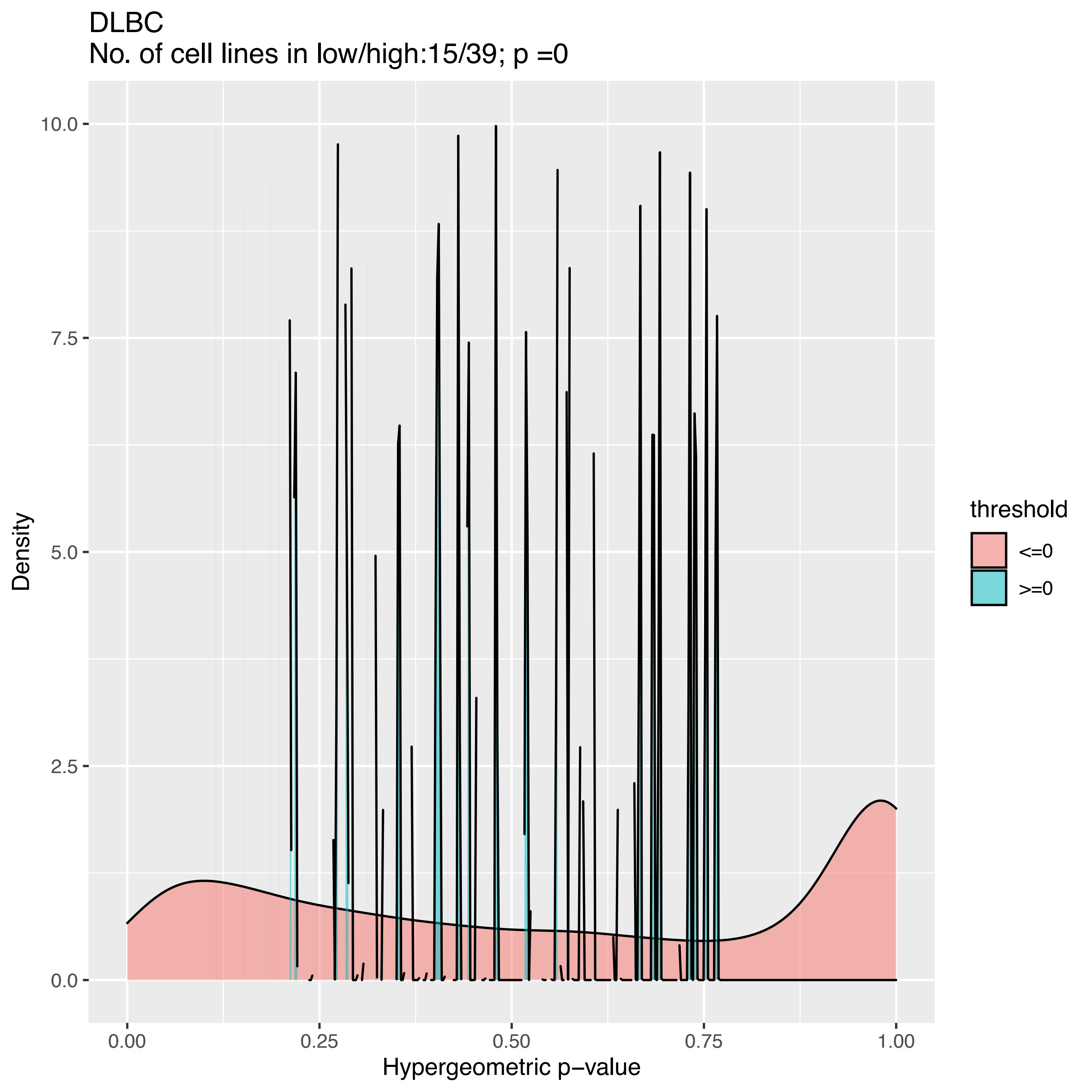
DLBC
No. of cell lines in low/high:15/39; p =0
Density
10.0
7.5
5.0
2.5
0.0
0.00
0.25
0.50
0.75
1.00
Hypergeometric p−value
threshold
<=0
>=0

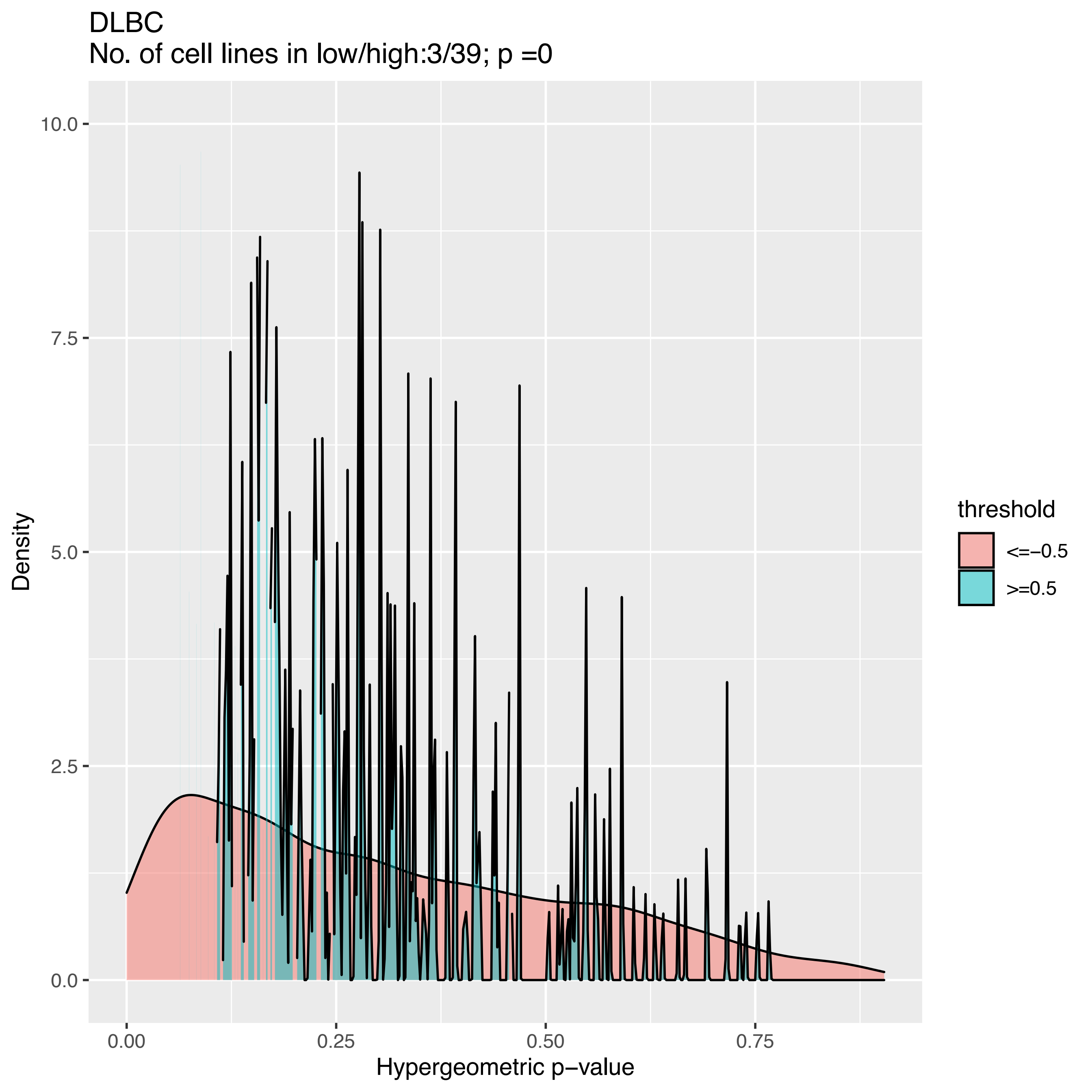
DLBC
No. of cell lines in low/high:3/39; p =0
Density
10.0
7.5
5.0
2.5
0.0
0.00
0.25
0.50
0.75
Hypergeometric p−value
threshold
<=−0.5
>=0.5

DLBC
No. of cell lines in low/high:3/38; p =6.16023595974476e−202

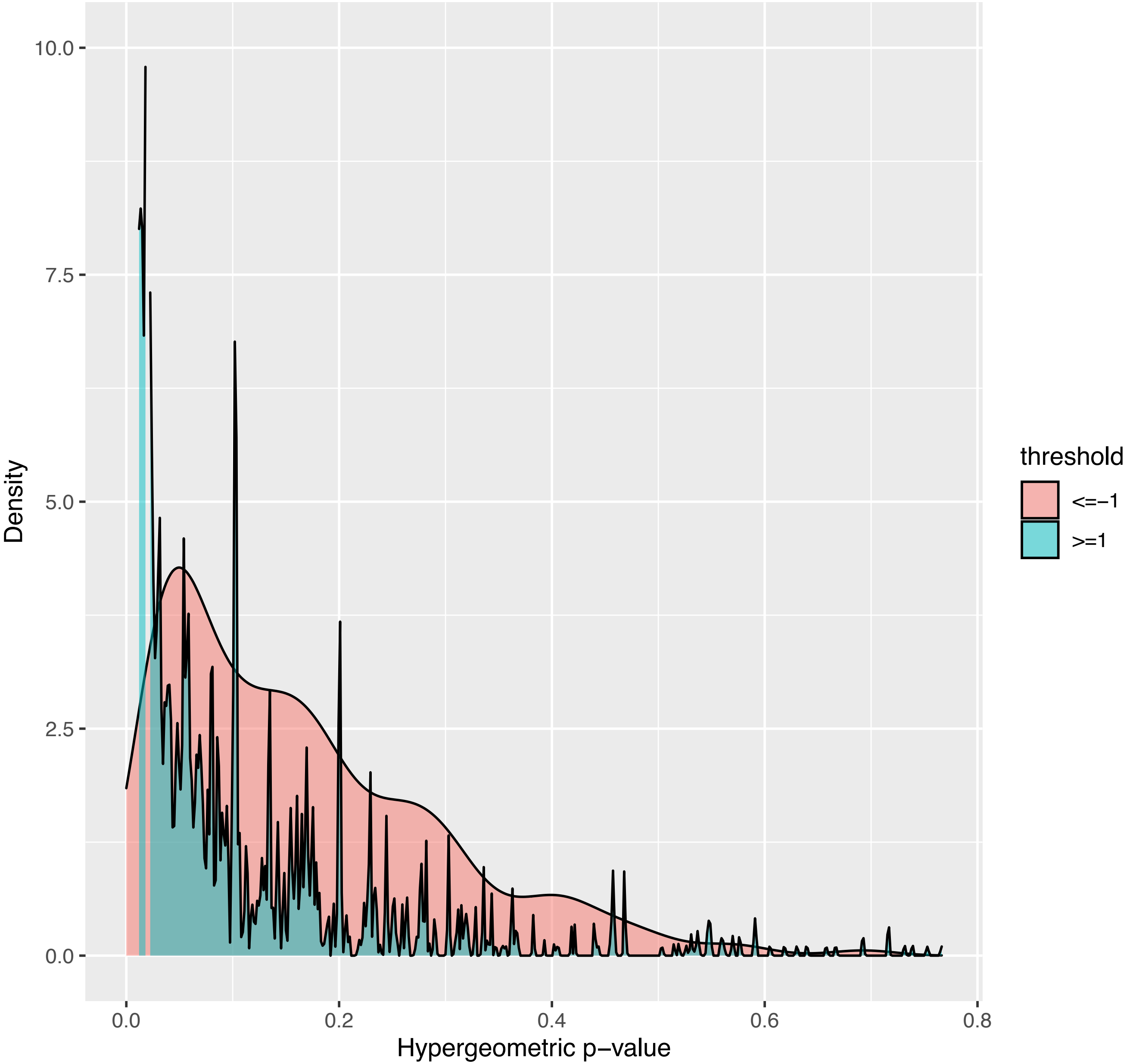

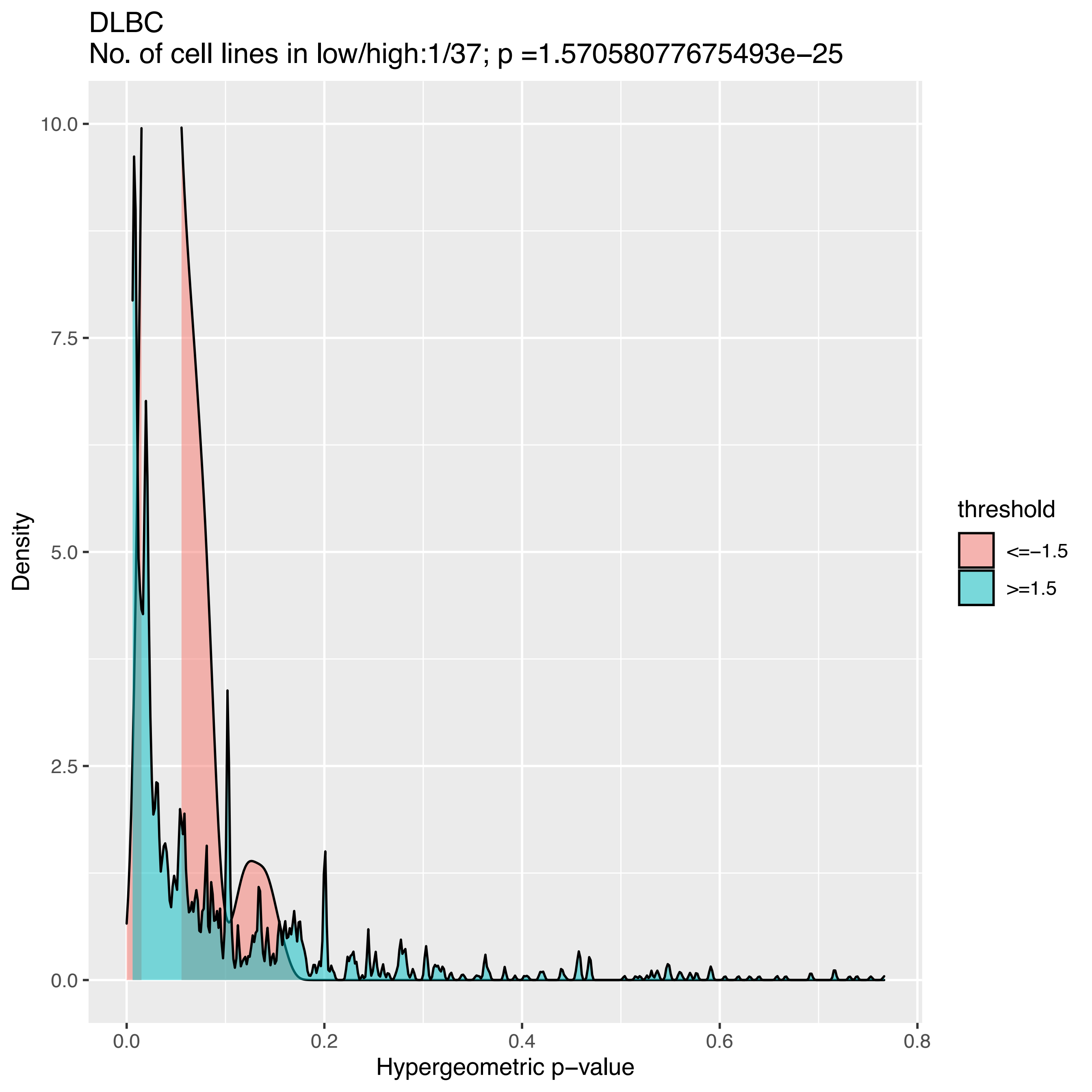
DLBC
No. of cell lines in low/high:1/37; p =1.57058077675493e−25
10.0
7.5
5.0
2.5
0.0
Density
0.0
0.2
0.4
0.6
0.8
Hypergeometric p−value
threshold
<=−1.5
>=1.5

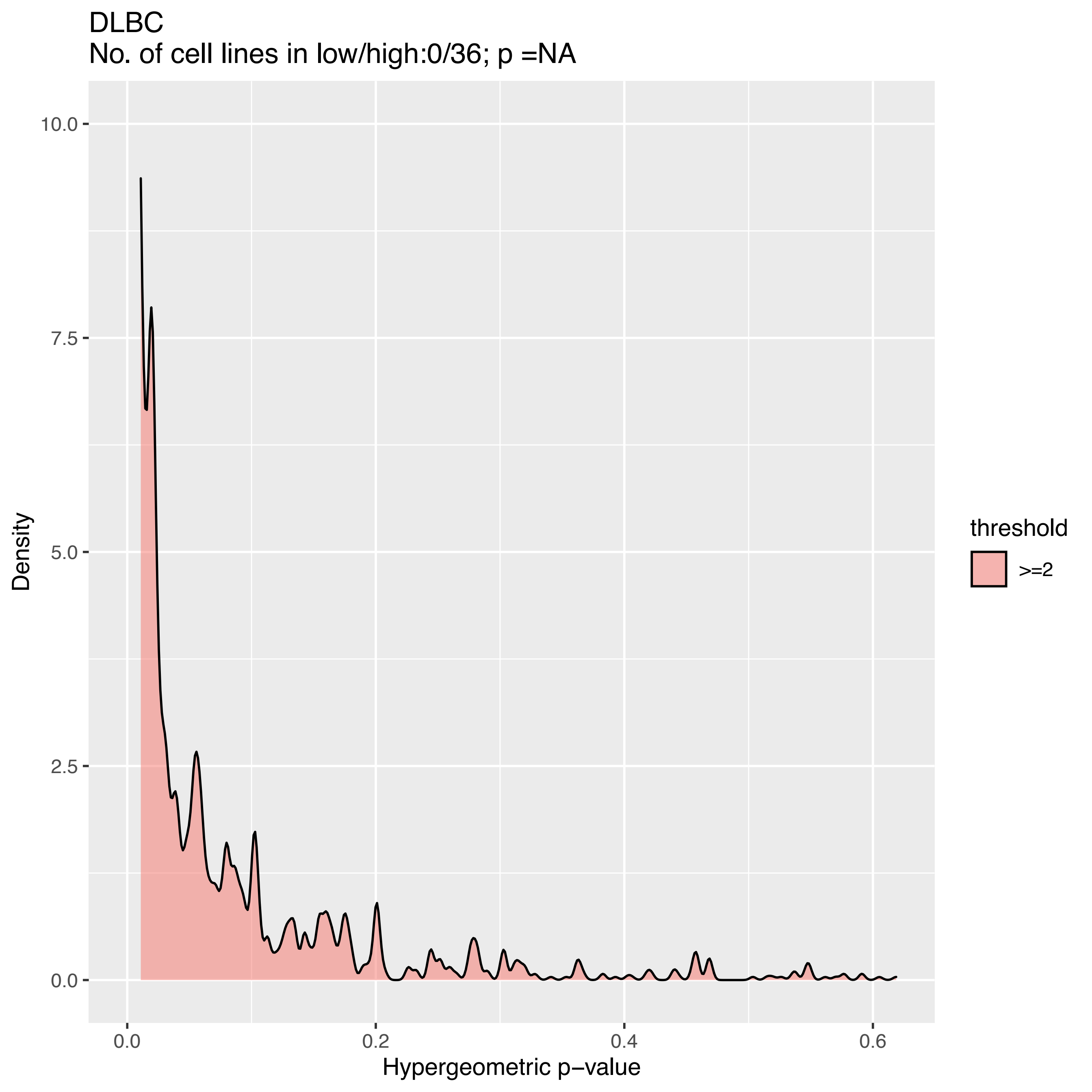
DLBC
No. of cell lines in low/high:0/36; p =NA
10.0
7.5
5.0
2.5
0.0
Density
0.0
0.2
0.4
0.6
Hypergeometric p−value
threshold
>=2

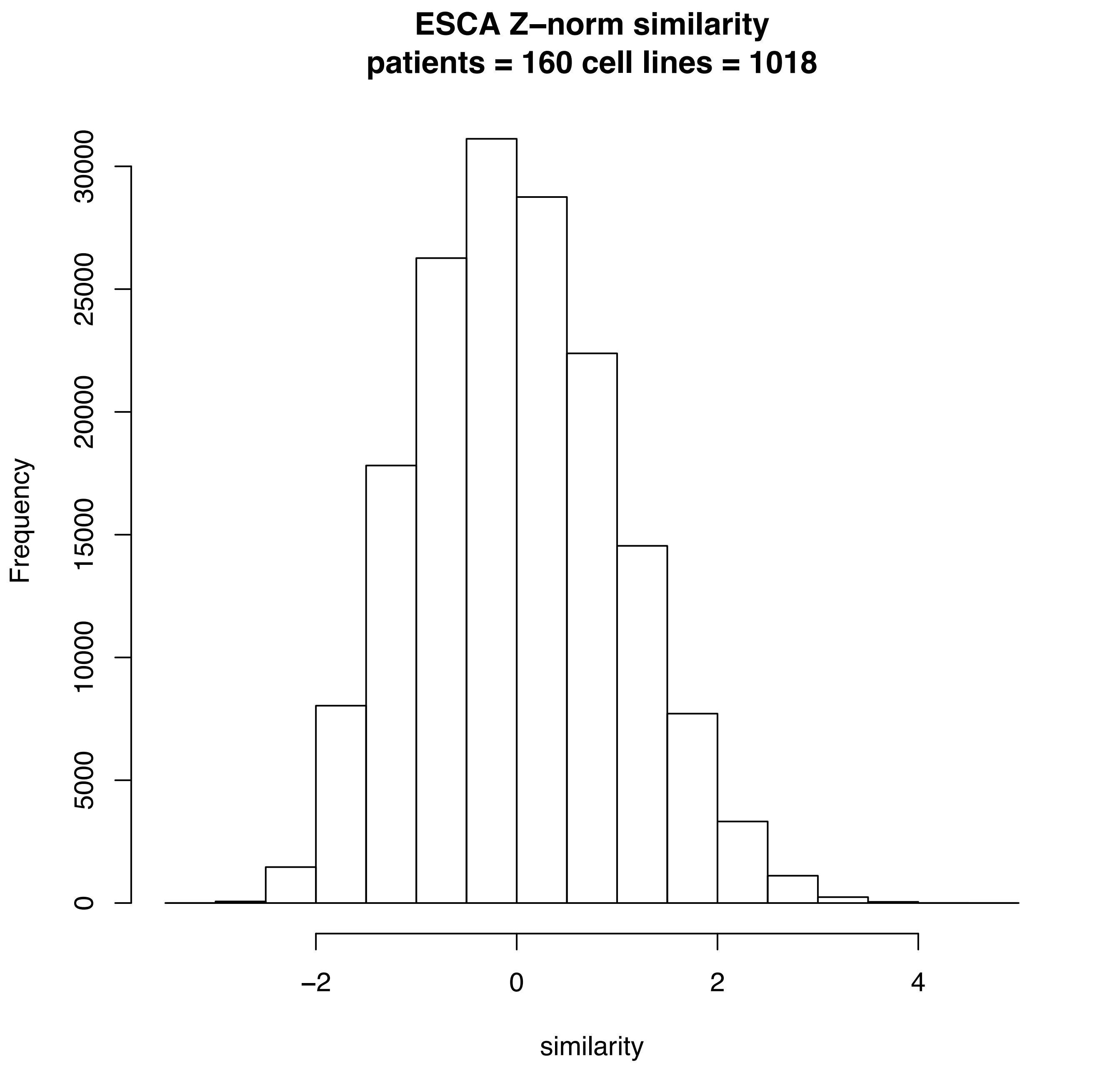

ESCA Z–norm similarity
patients = 160 cell lines = 1018
Frequency
0
5000
10000
15000
20000
25000
30000
−2
0
2
4
similarity

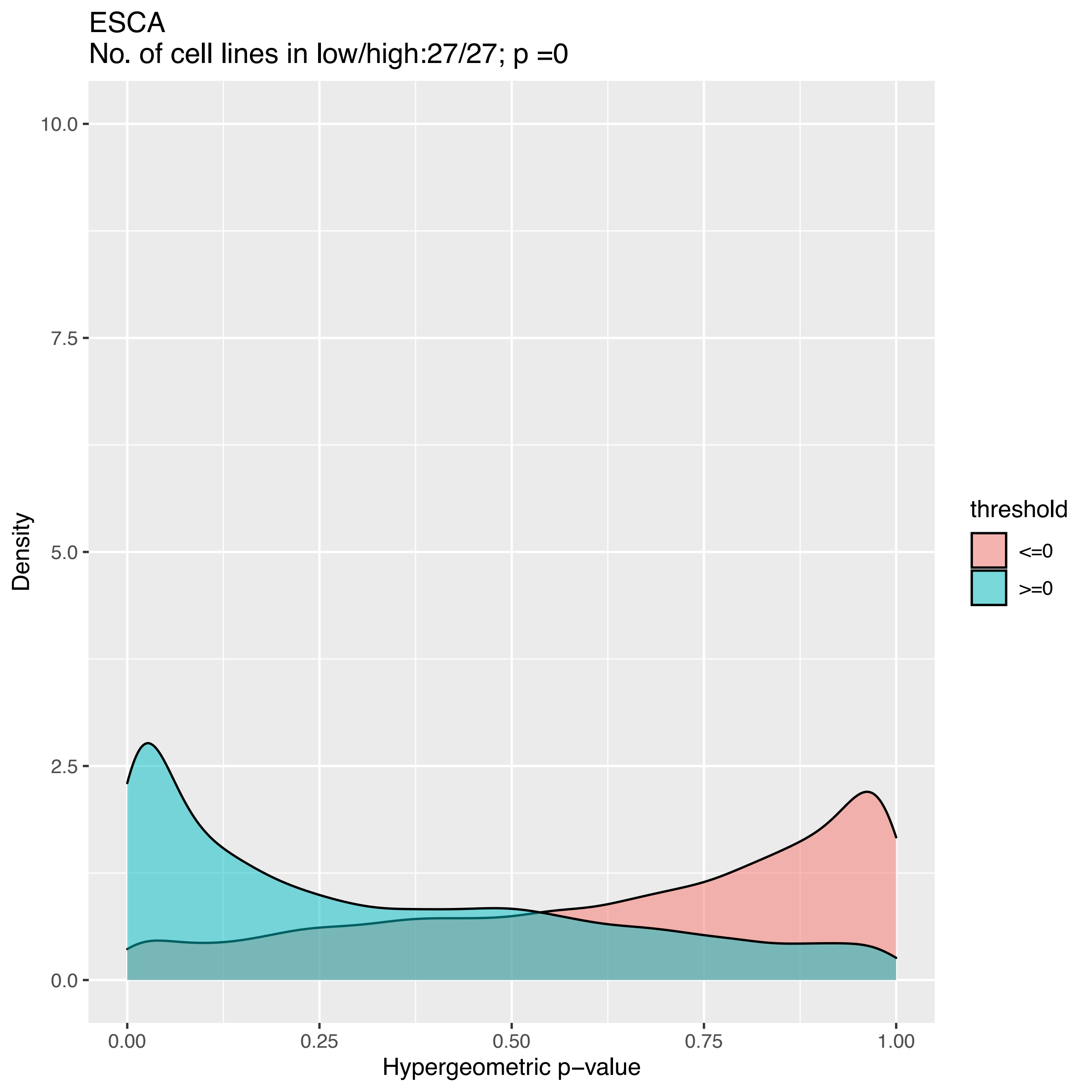
ESCA
No. of cell lines in low/high:27/27; p =0
Density
10.0
7.5
5.0
2.5
0.0
0.00
0.25
0.50
0.75
1.00
Hypergeometric p−value
threshold
<=0
>=0

ESCA
No. of cell lines in low/high:25/27; p =0

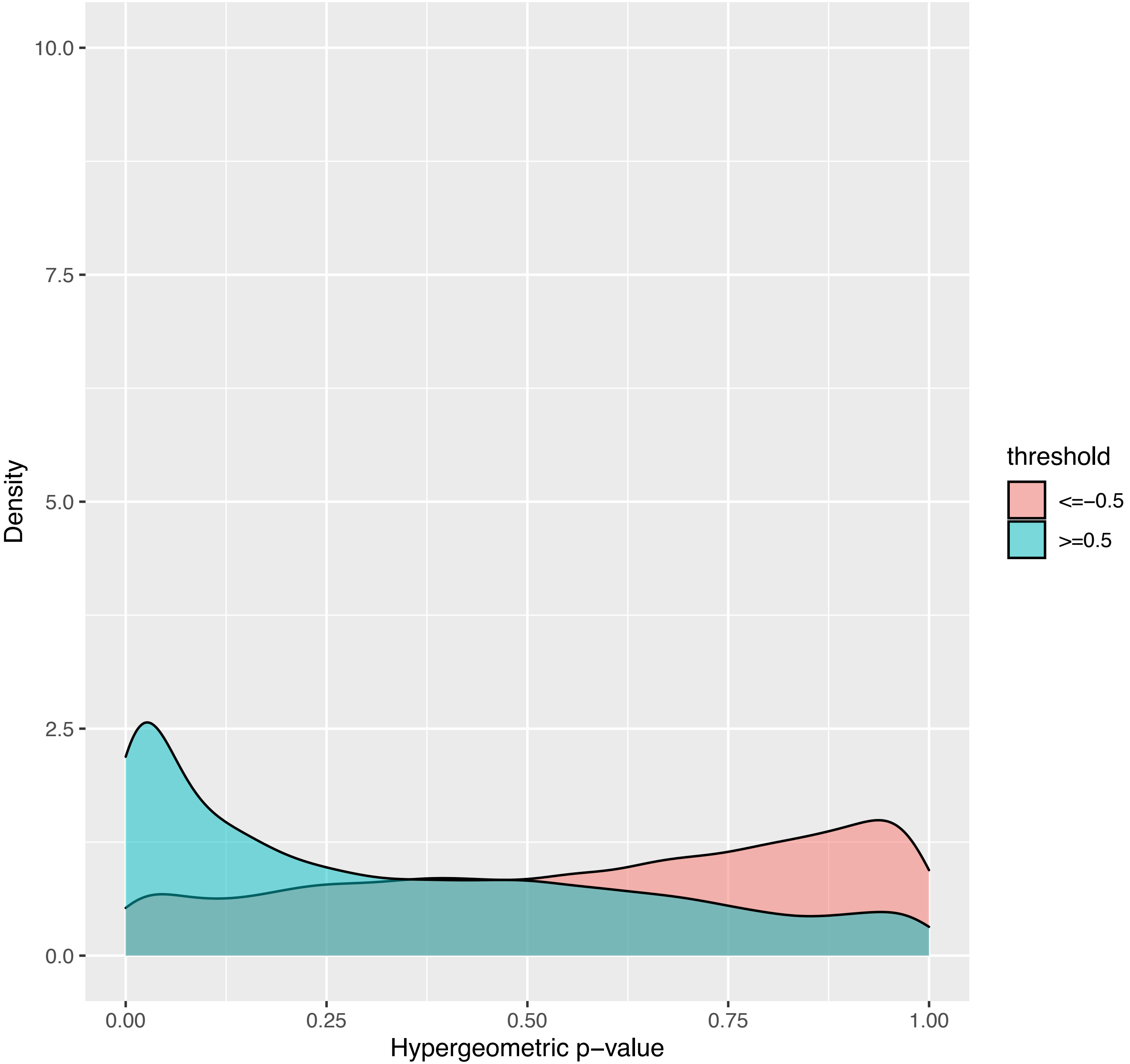

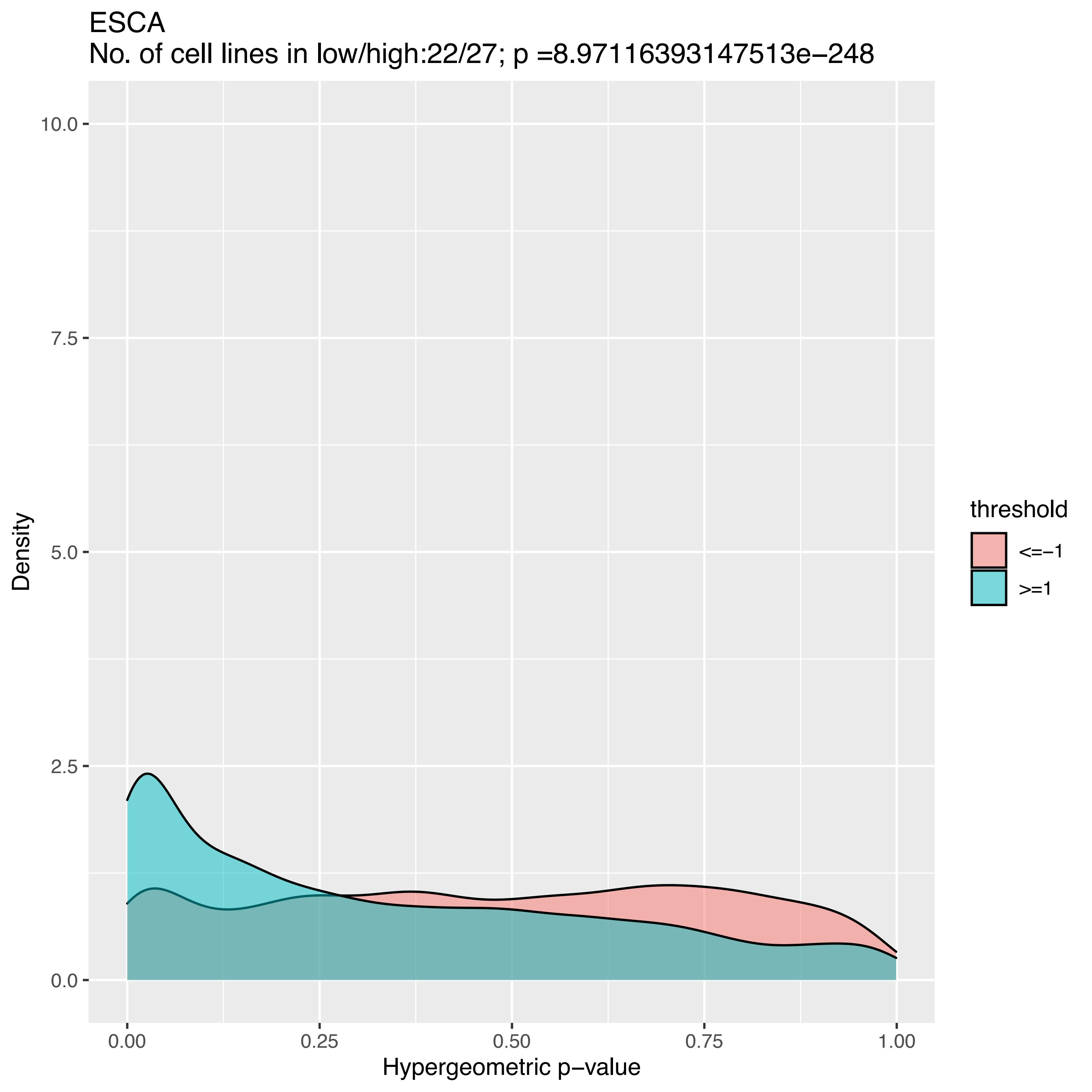

ESCA
No. of cell lines in low/high:22/27; p =8.97116393147513e−248
10.0
7.5
5.0
2.5
0.0
Density
0.00
0.25
0.50
0.75
1.00
Hypergeometric p−value
threshold
<=−1
>=1

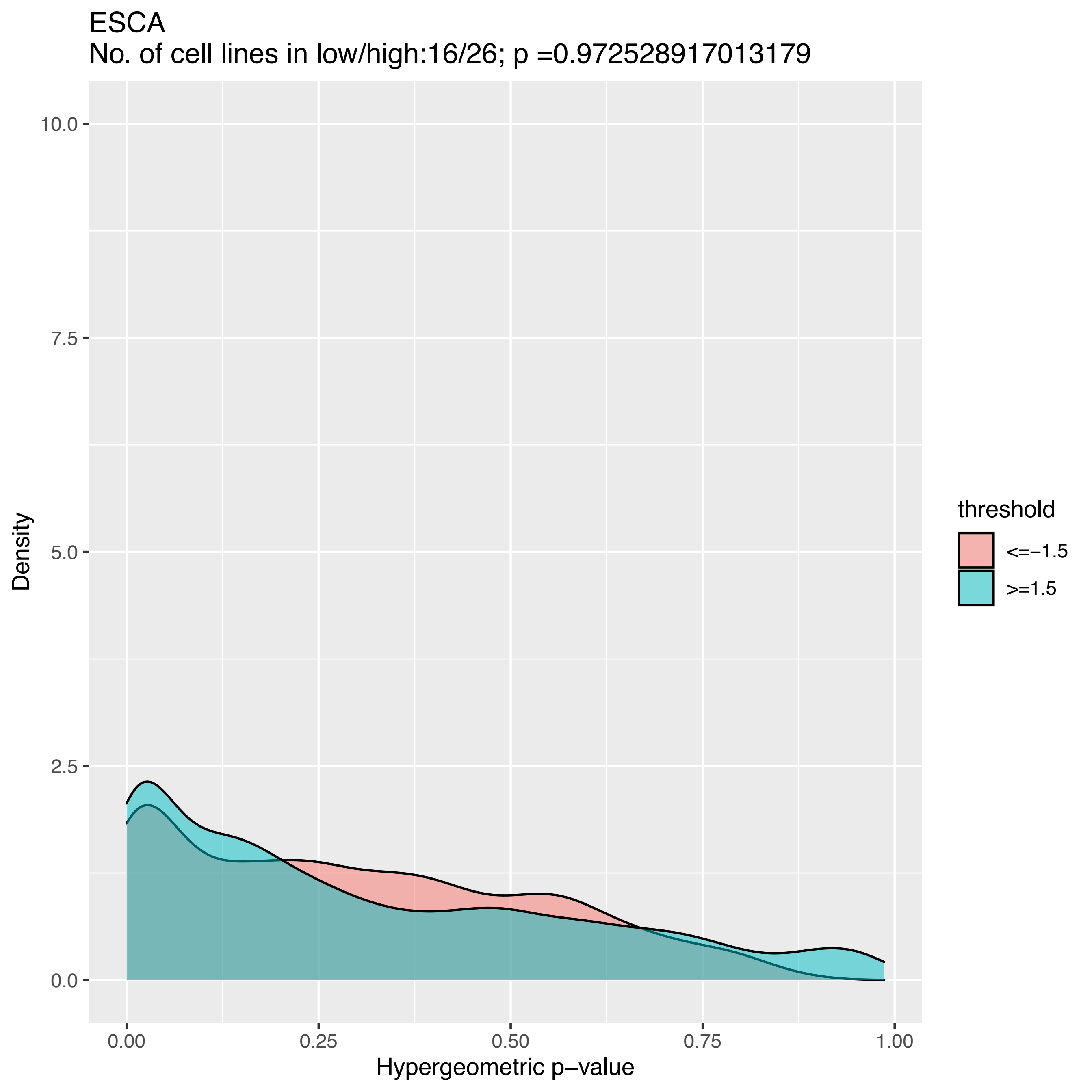

ESCA
No. of cell lines in low/high:16/26; p =0.972528917013179
10.0
7.5
5.0
2.5
0.0
Density
0.00
0.25
0.50
0.75
1.00
Hypergeometric p−value
threshold
<=−1.5
>=1.5

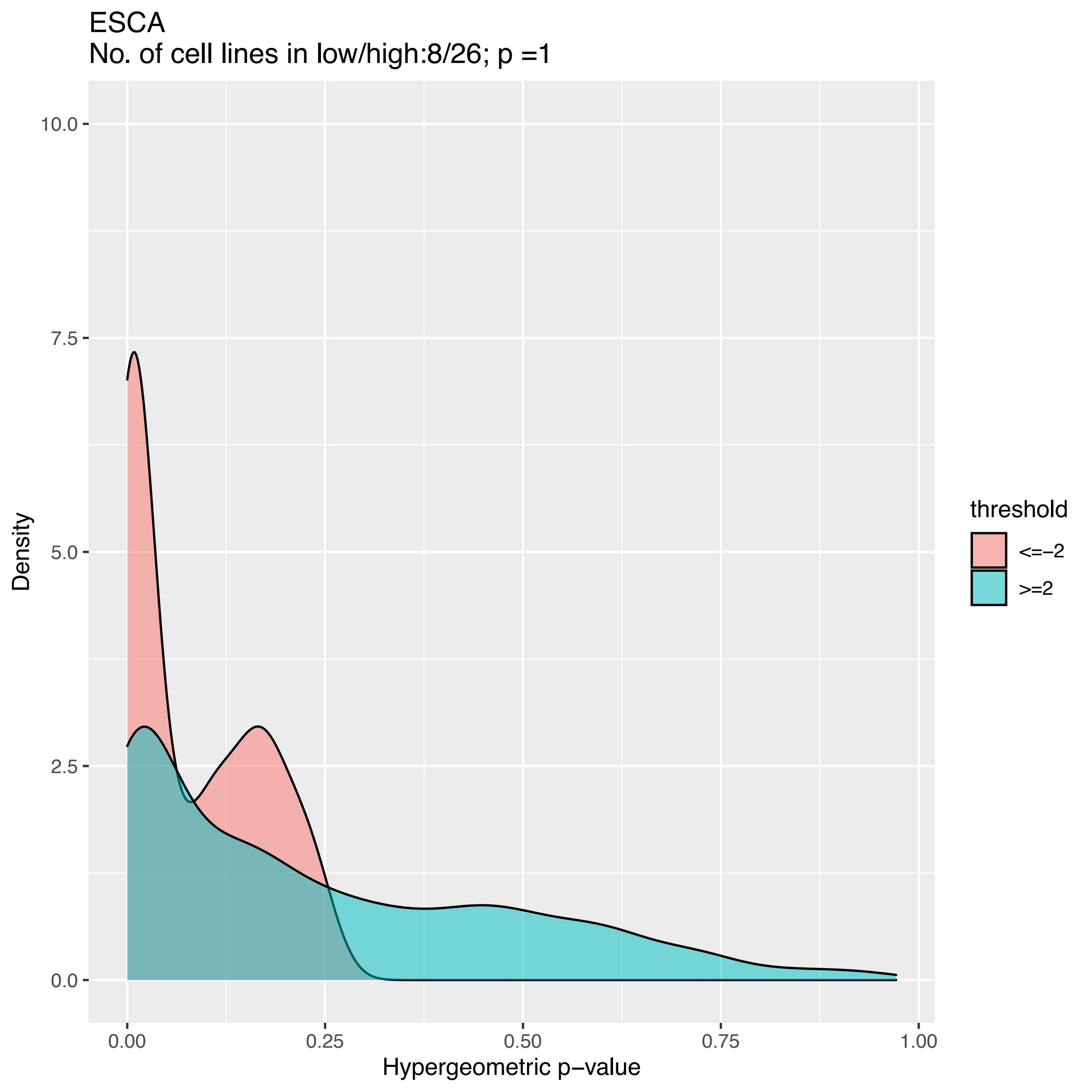
ESCA
No. of cell lines in low/high:8/26; p =1
Density
10.0
7.5
5.0
2.5
0.0
0.00
0.25
0.50
0.75
1.00
Hypergeometric p−value
threshold
<=−2
>=2

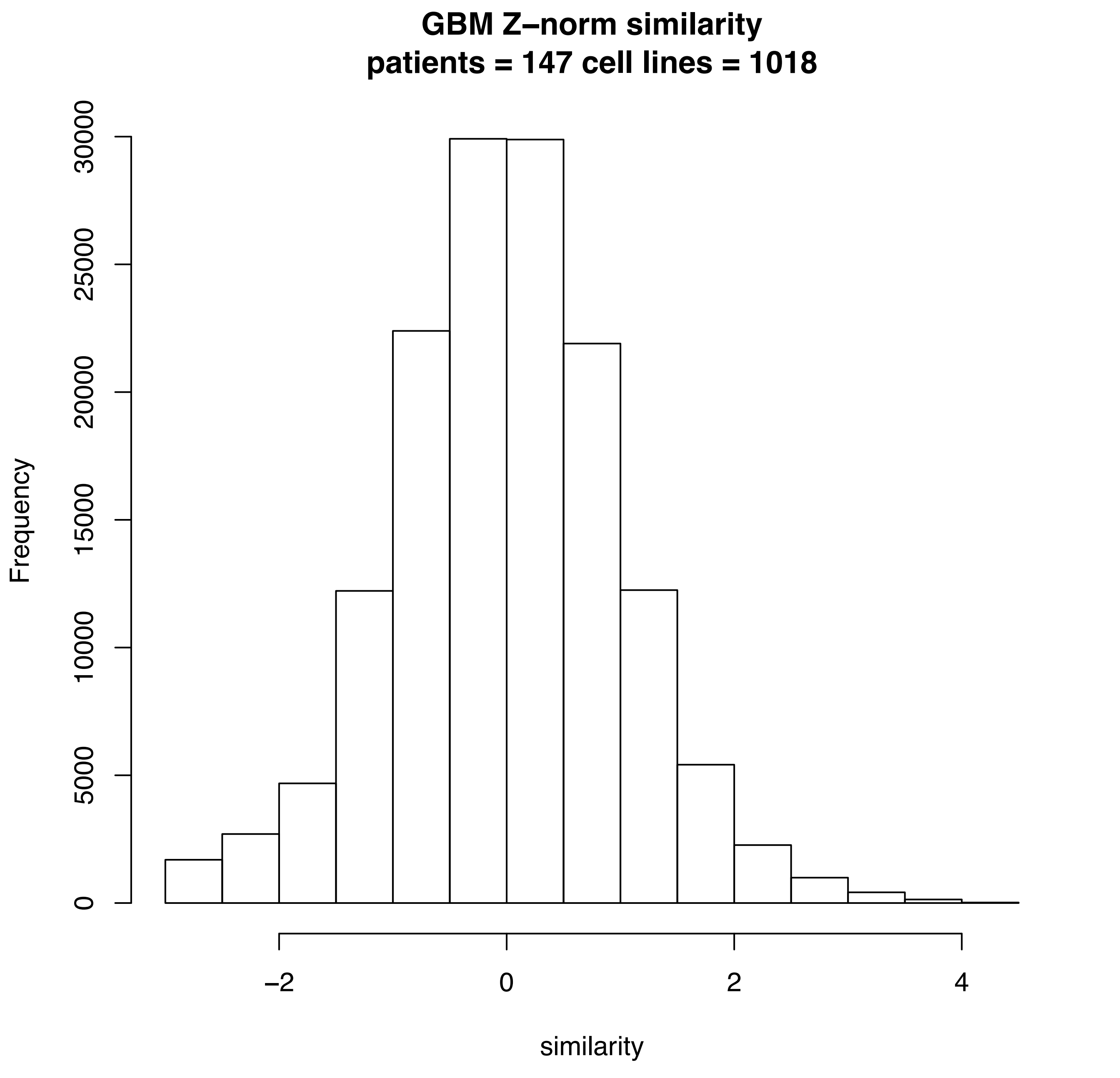

GBM Z–norm similarity
patients = 147 cell lines = 1018
Frequency
0
5000
10000
15000
20000
25000
30000
−2
0
2
4
similarity

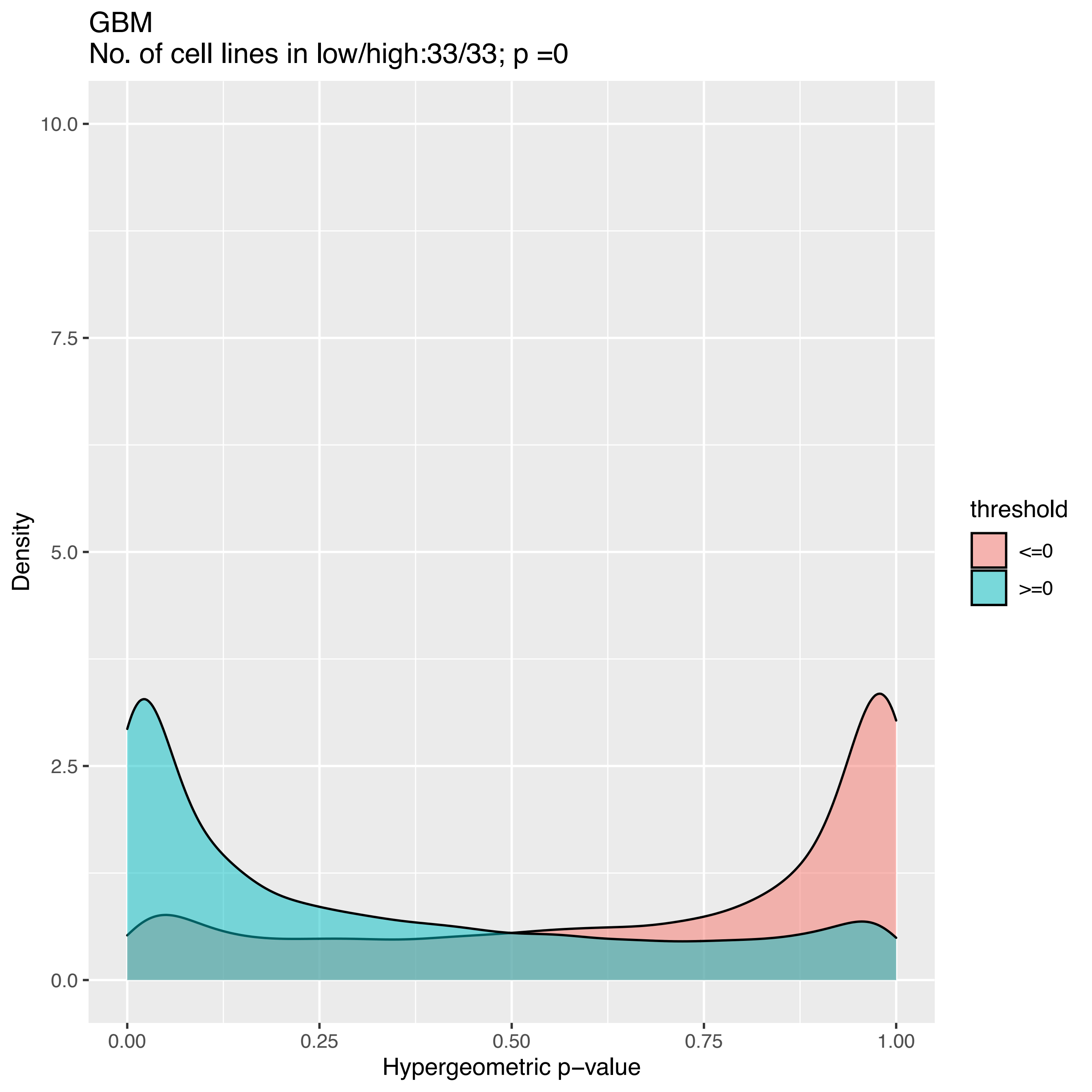
GBM
No. of cell lines in low/high:33/33; p =0
10.0
7.5
5.0
2.5
0.0
Density
0.00
0.25
0.50
0.75
1.00
Hypergeometric p−value
threshold
<=0
>=0

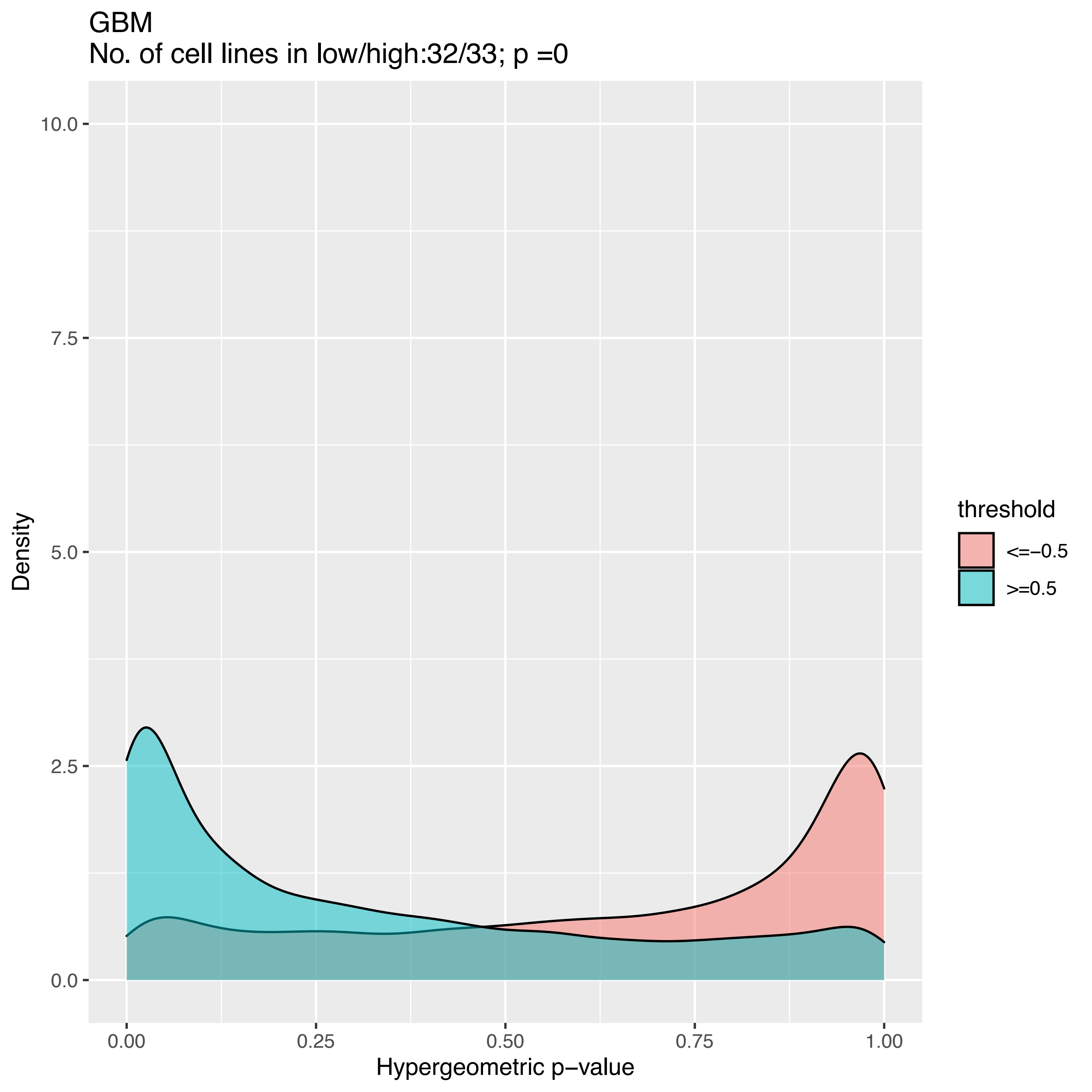
GBM
No. of cell lines in low/high:32/33; p =0
Density
10.0
7.5
5.0
2.5
0.0
0.00
0.25
0.50
0.75
1.00
Hypergeometric p−value
threshold
<=−0.5
>=0.5

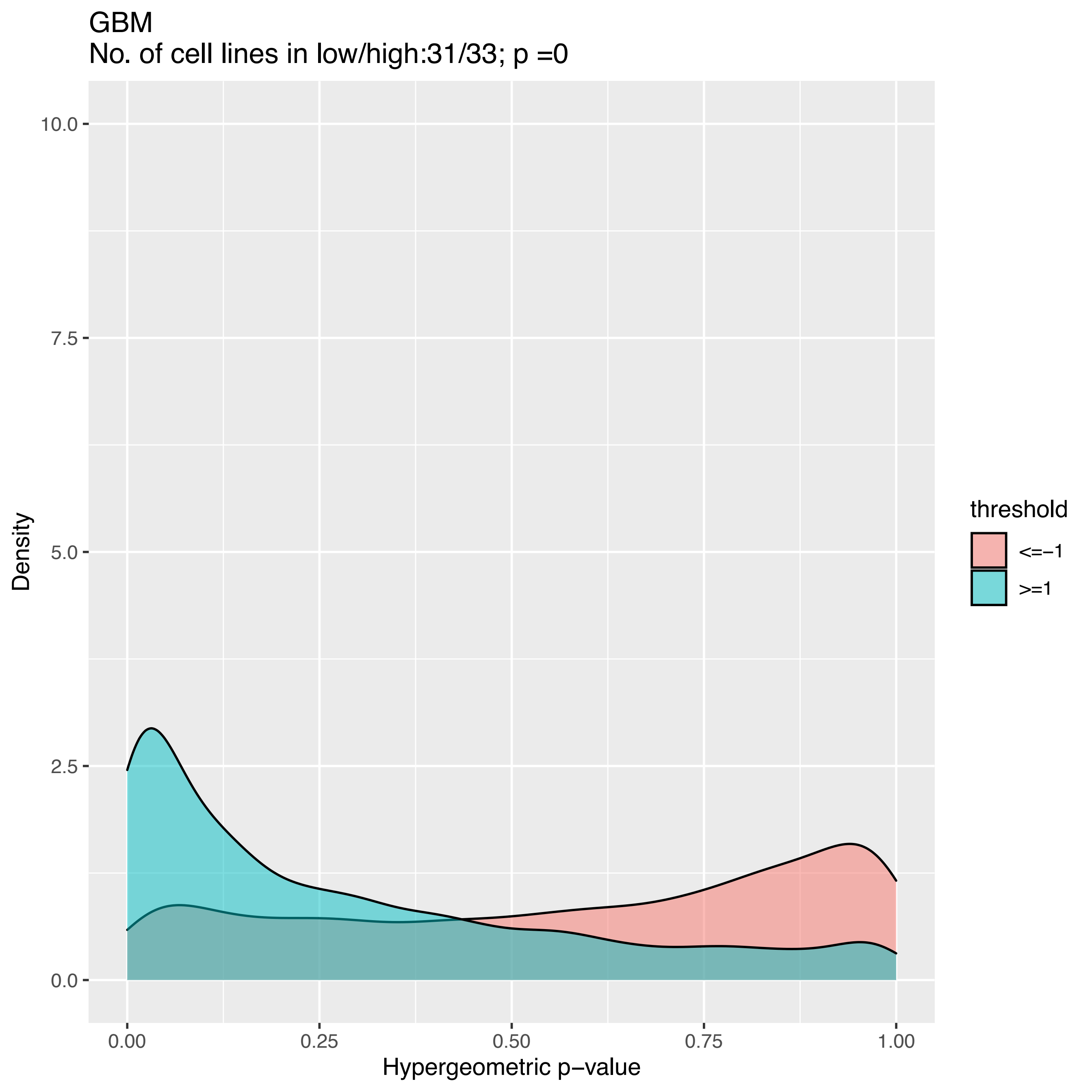

GBM
No. of cell lines in low/high:31/33; p =0
Density
10.0
7.5
5.0
2.5
0.0
0.00
0.25
0.50
0.75
1.00
Hypergeometric p−value
threshold
<=−1
>=1

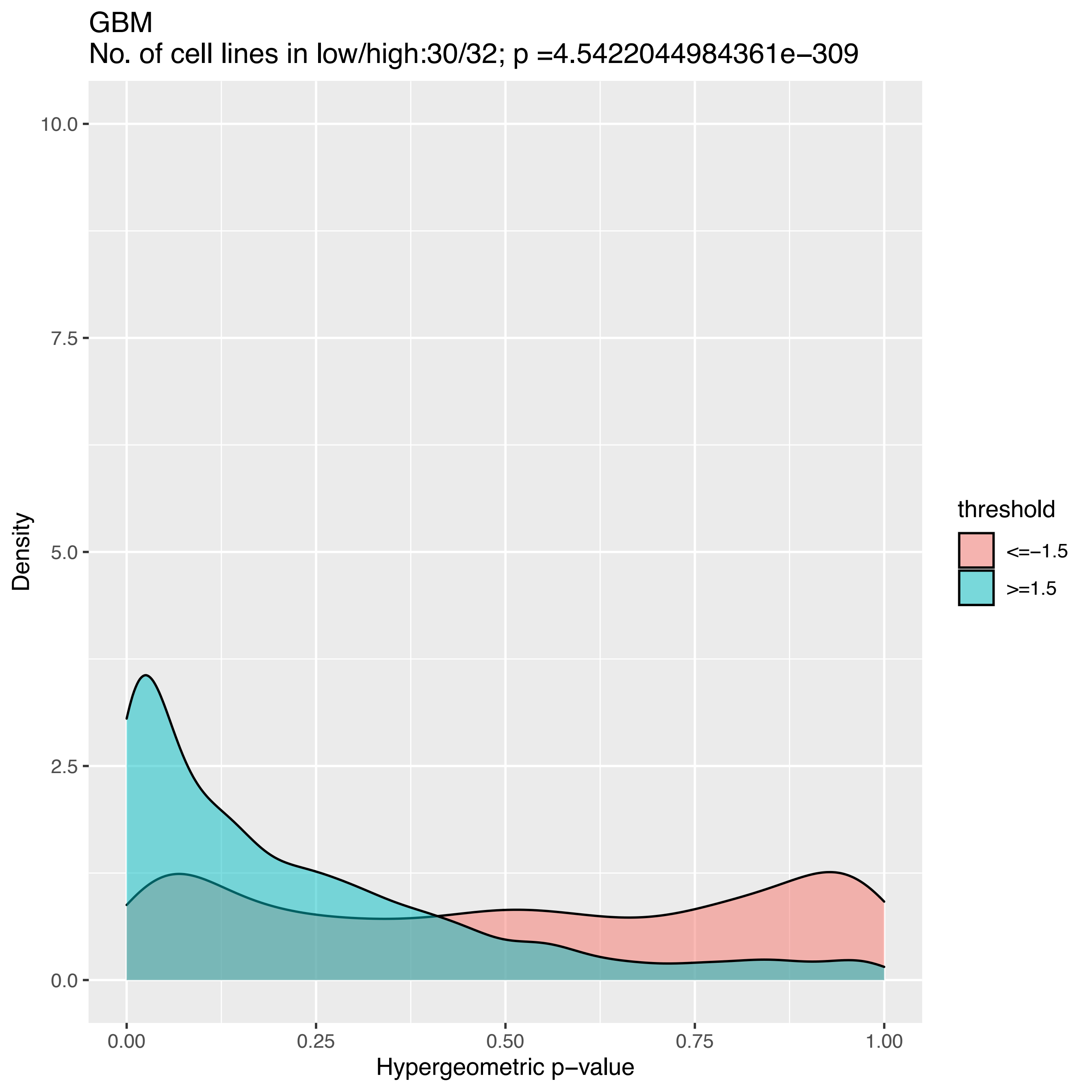
GBM
No. of cell lines in low/high:30/32; p =4.54220449843610e−309
Density
Hypergeometric p−value
10.0
7.5
5.0
2.5
0.0
0.00
0.25
0.50
0.75
1.00
threshold
<=−1.5
>=1.5

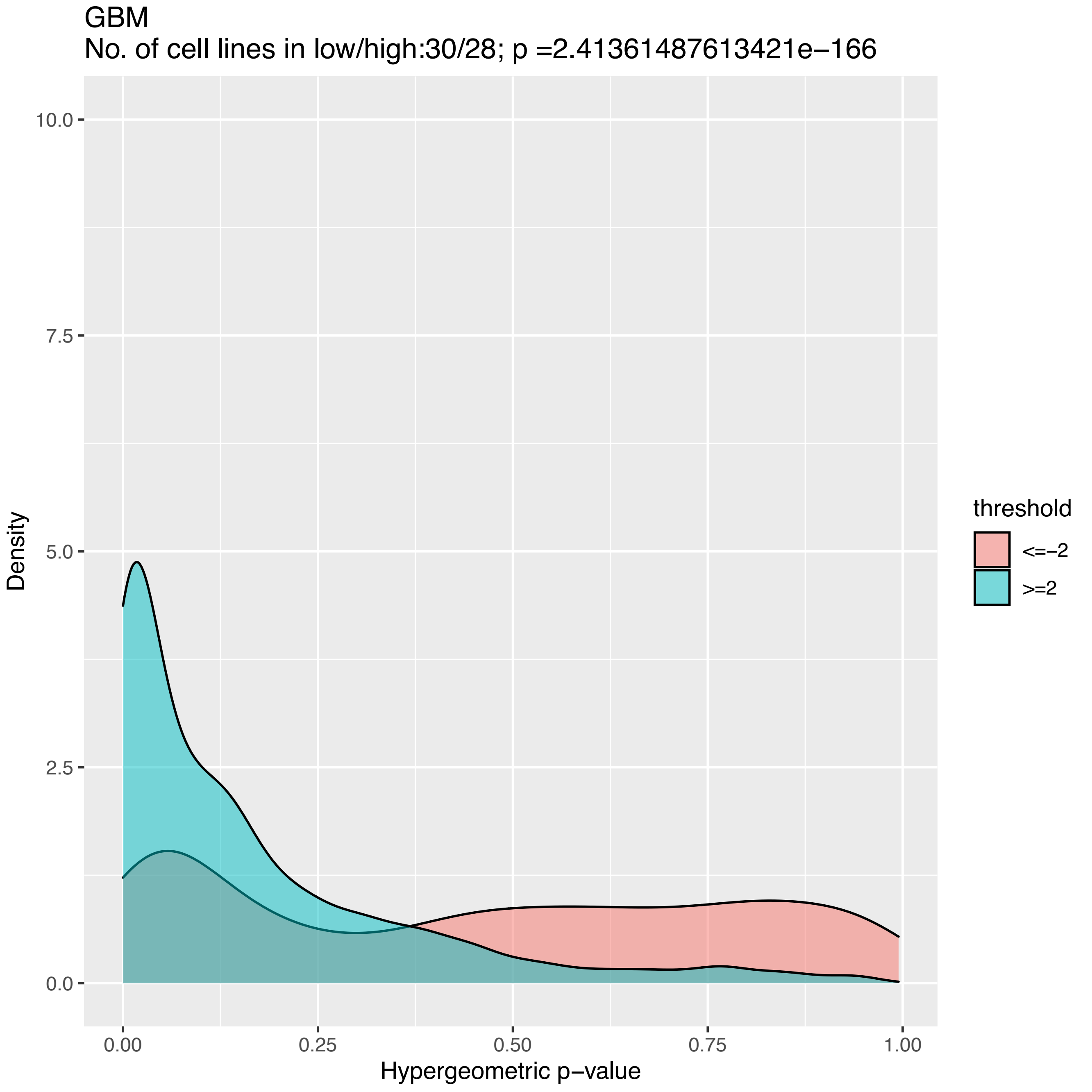
GBM
No. of cell lines in low/high:30/28; p =2.41361487613421e−166
Density
10.0
7.5
5.0
2.5
0.0
0.00
0.25
0.50
0.75
1.00
Hypergeometric p−value
threshold
<=−2
>=2

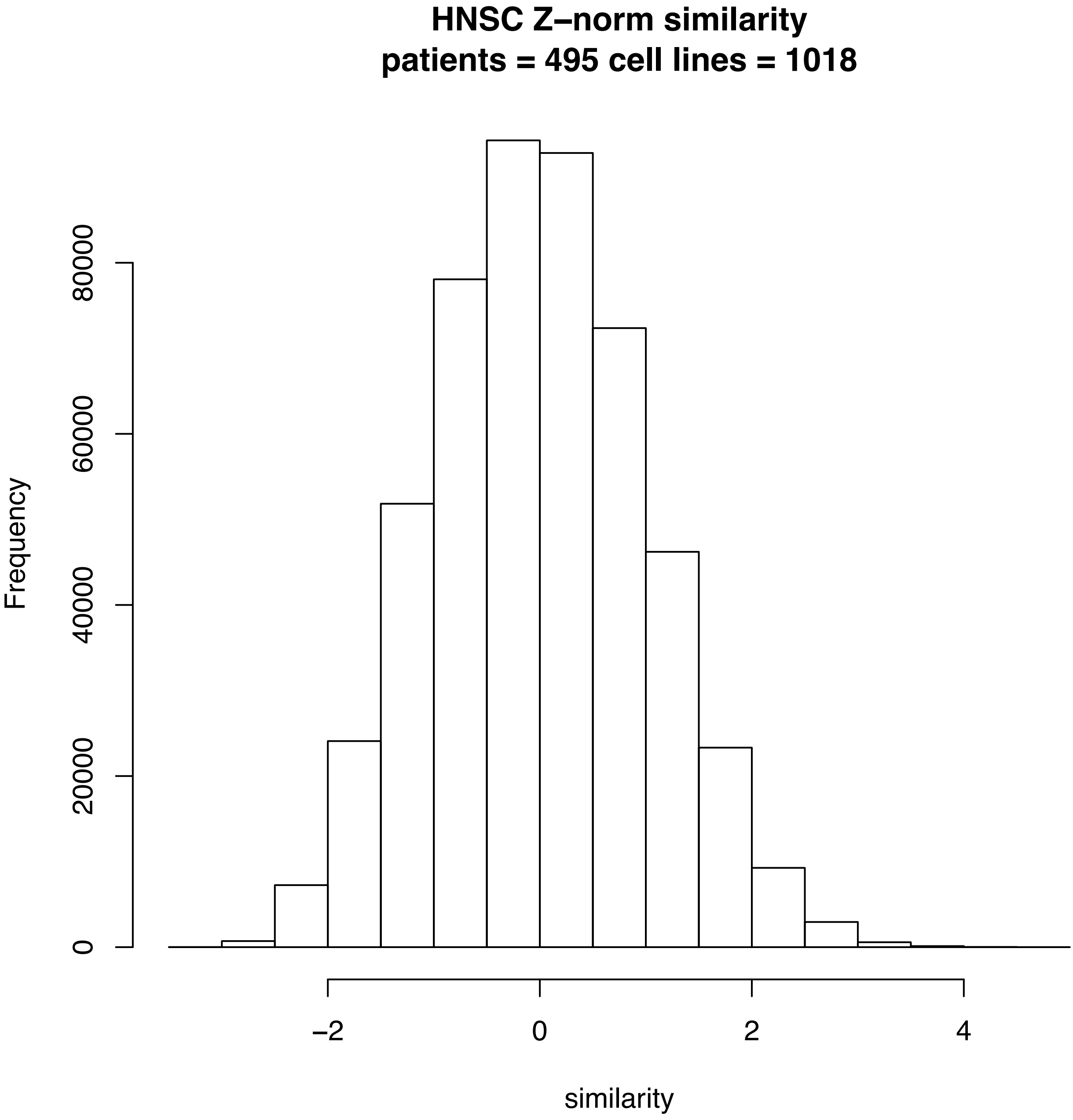
HNSC Z–norm similarity
patients = 495 cell lines = 1018
Frequency
0
20000
40000
60000
80000
–2
0
2
4
similarity

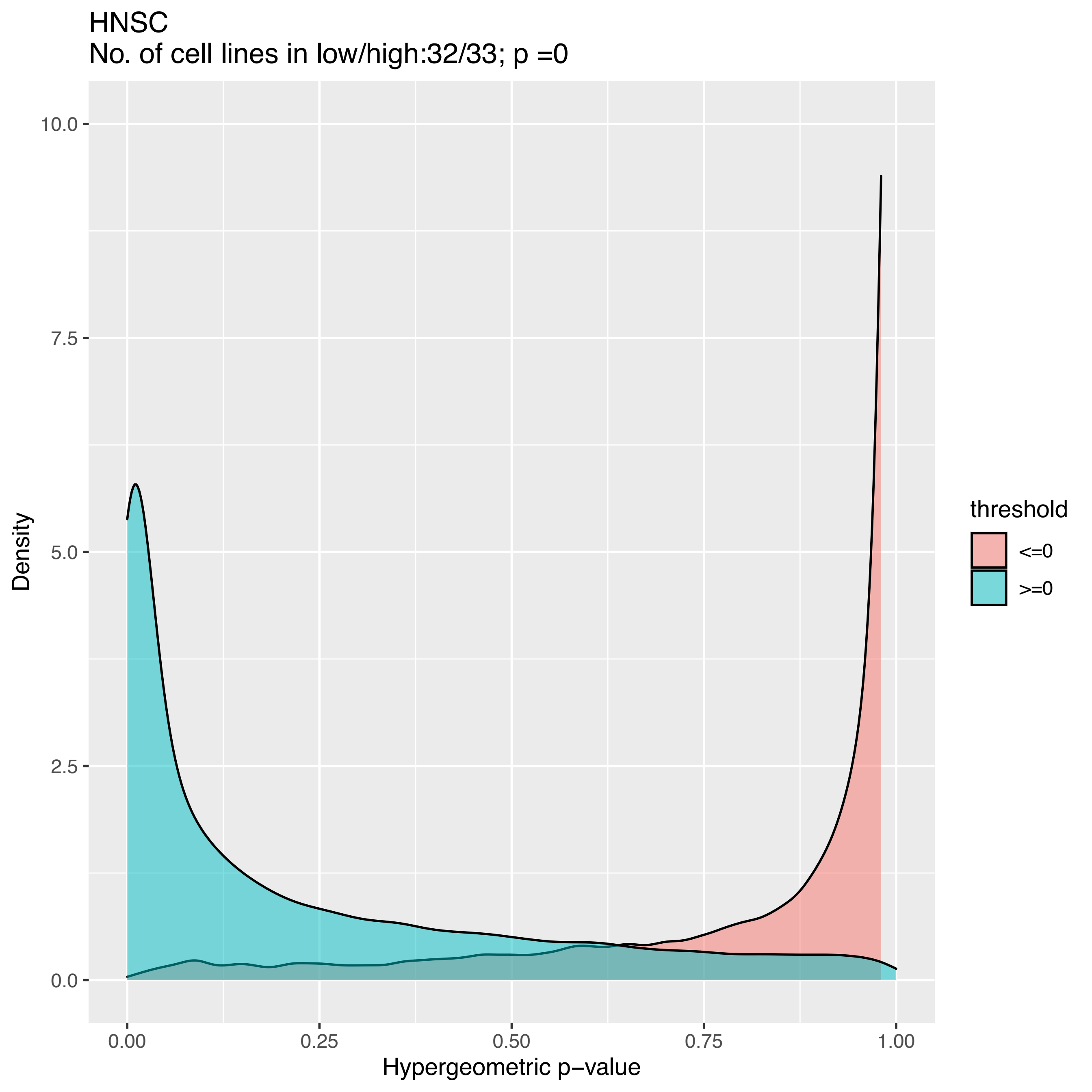

HNSC
No. of cell lines in low/high:32/33; p =0
10.0
7.5
5.0
2.5
0.0
Density
0.00
0.25
0.50
0.75
1.00
Hypergeometric p−value
threshold
<=0
>=0

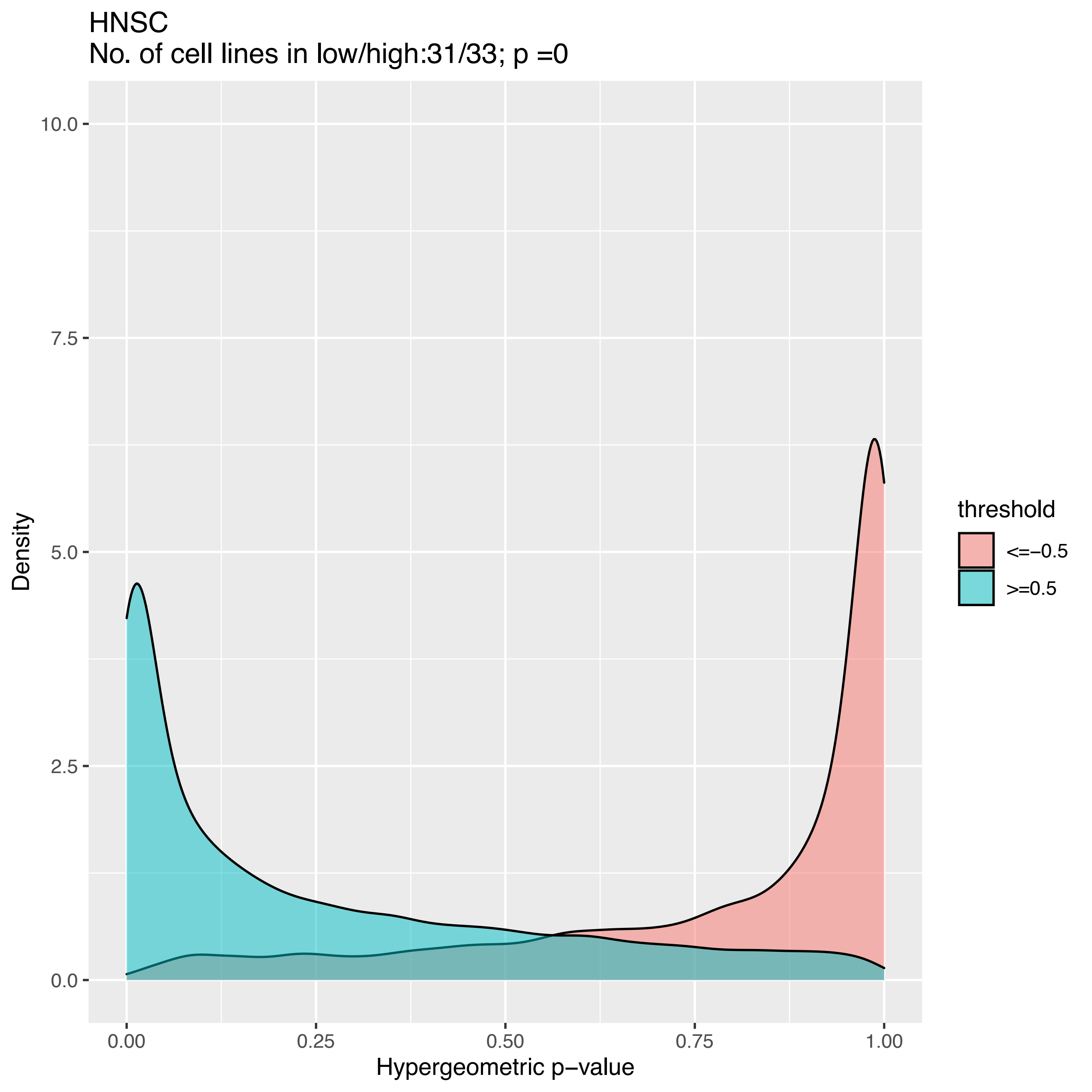
HNSC
No. of cell lines in low/high:31/33; p =0
Density
Hypergeometric p−value
0.00
0.25
0.50
0.75
1.00
0.0
2.5
5.0
7.5
10.0
threshold
<=−0.5
>=0.5

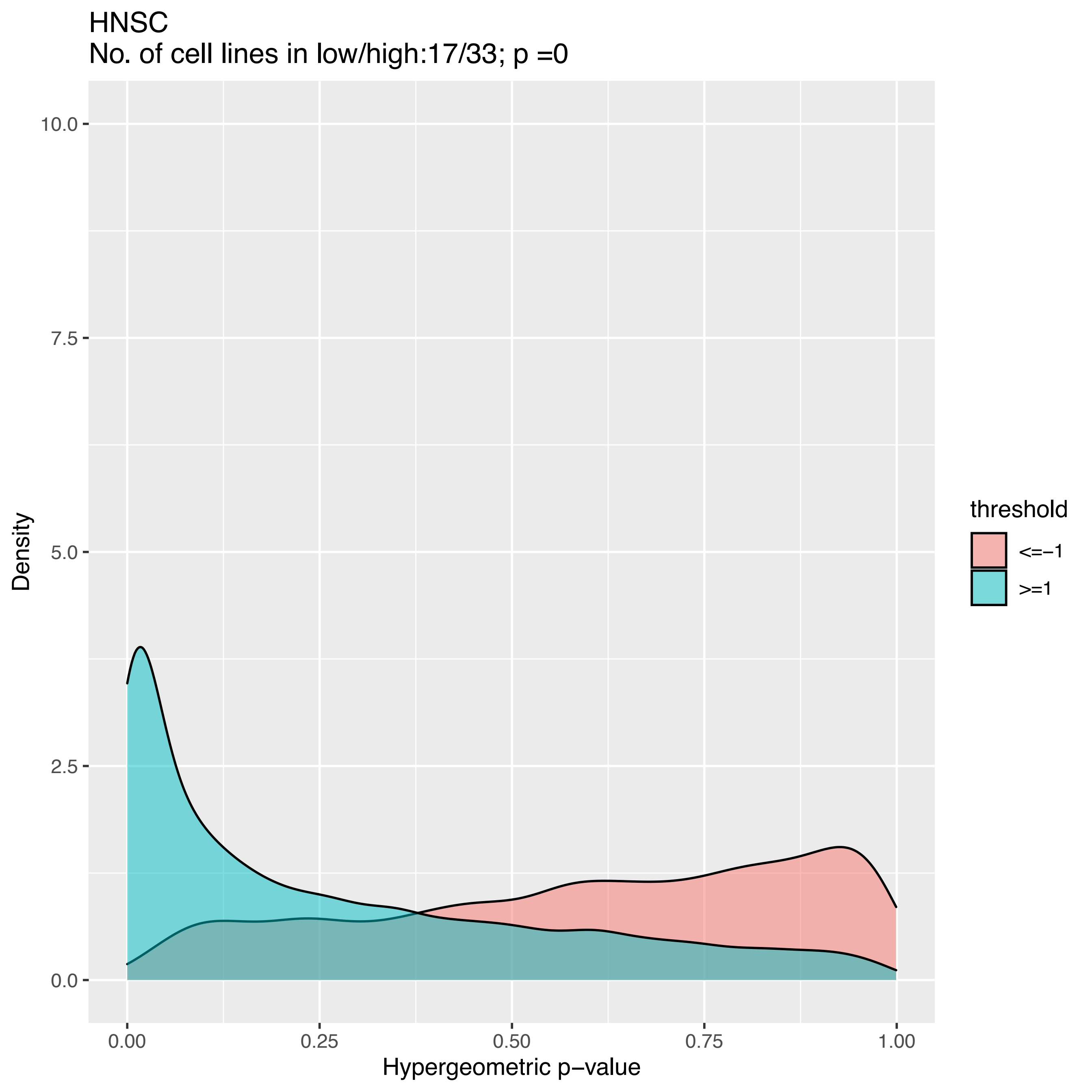

HNSC
No. of cell lines in low/high:17/33; p =0
10.0
7.5
5.0
2.5
0.0
Density
0.00
0.25
0.50
0.75
1.00
Hypergeometric p−value
threshold
<=−1
>=1

HNSC
No. of cell lines in low/high:7/33; p =0.99998362807959

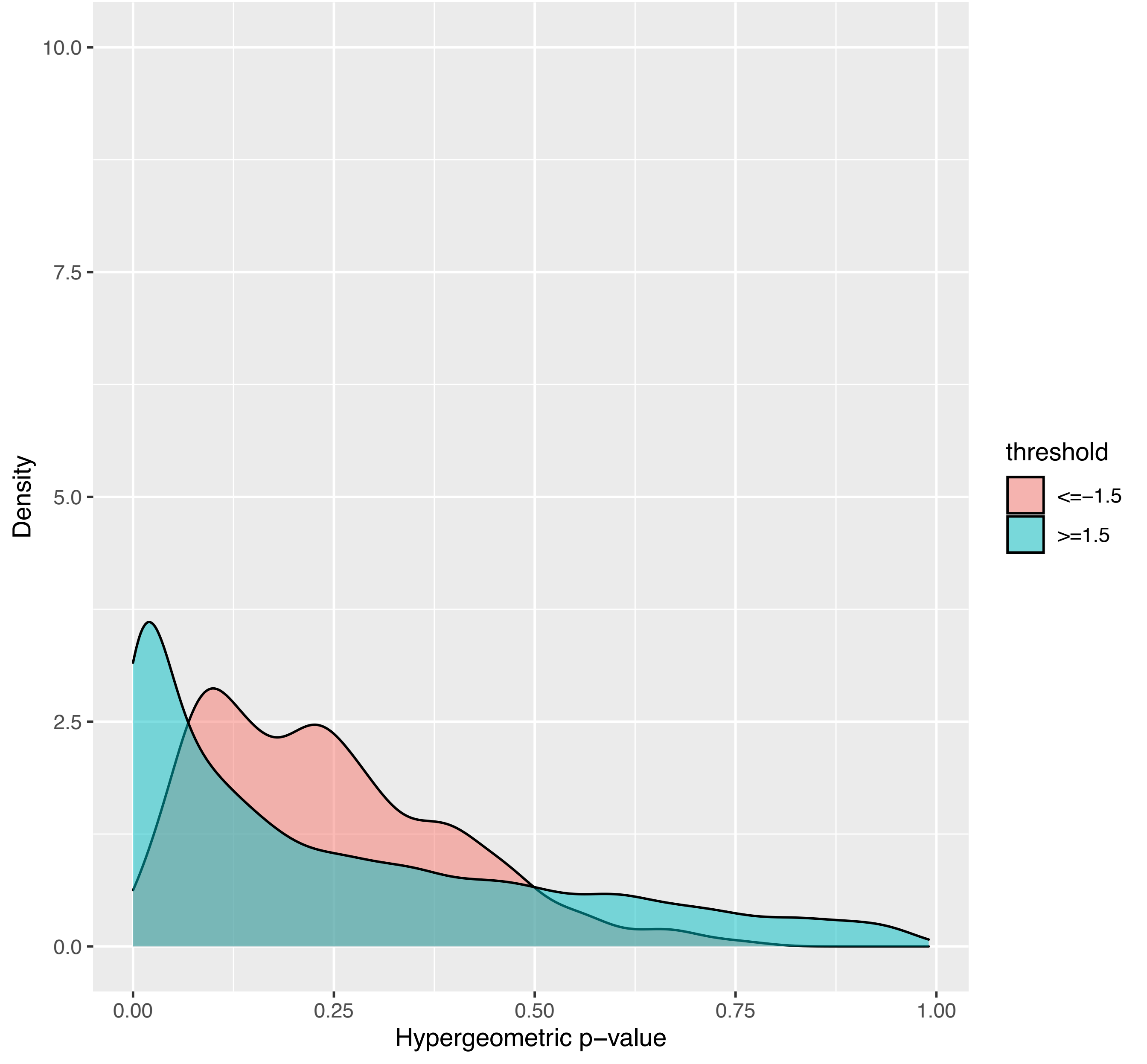

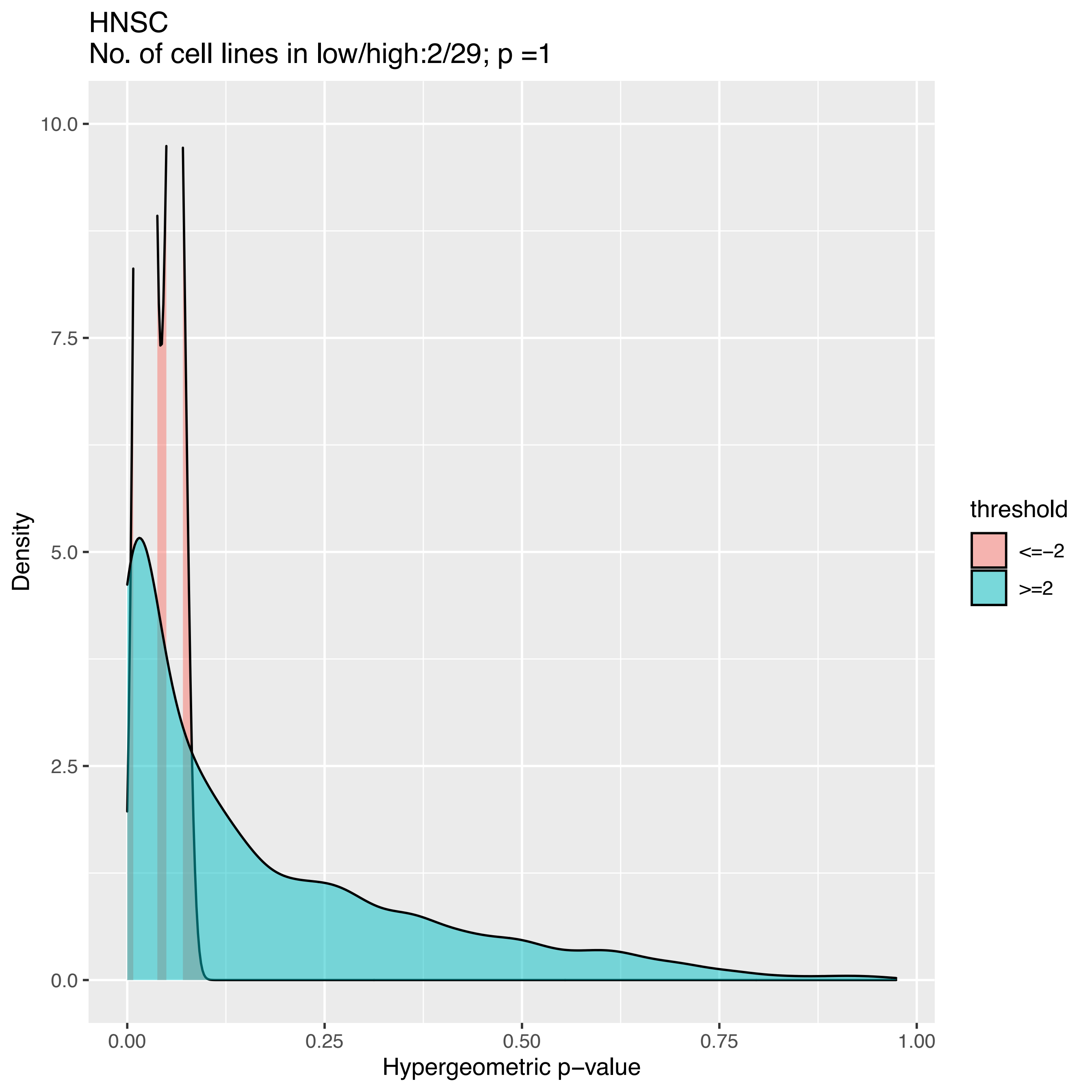
HNSC
No. of cell lines in low/high:2/29; p =1
Density
10.0
7.5
5.0
2.5
0.0
0.00
0.25
0.50
0.75
1.00
Hypergeometric p−value
threshold
<=−2
>=2

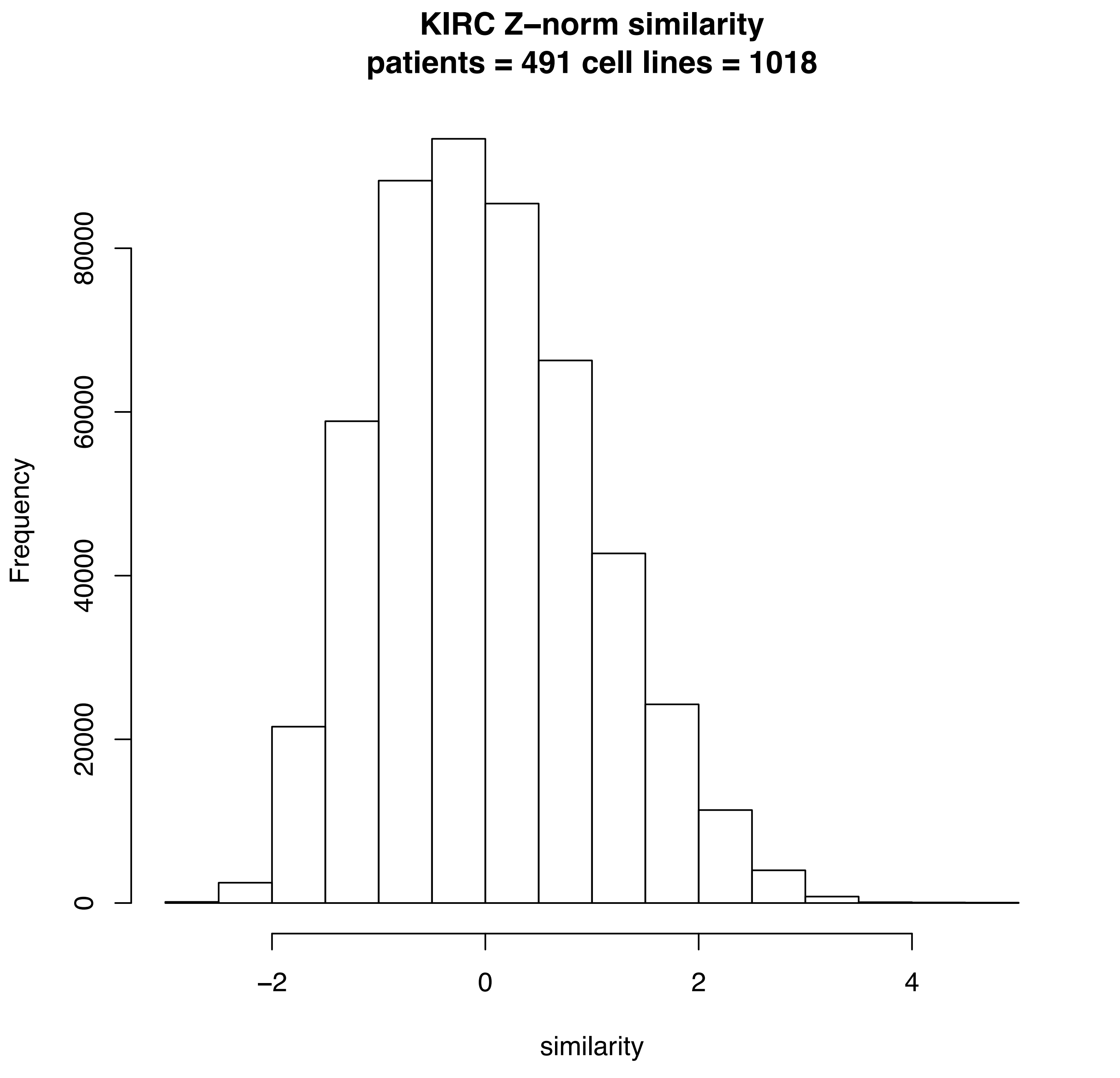

KIRC Z−norm similarity
patients = 491 cell lines = 1018
Frequency
0
20000
40000
60000
80000
−2
0
2
4
similarity

KIRC
No. of cell lines in low/high:32/33; p =0

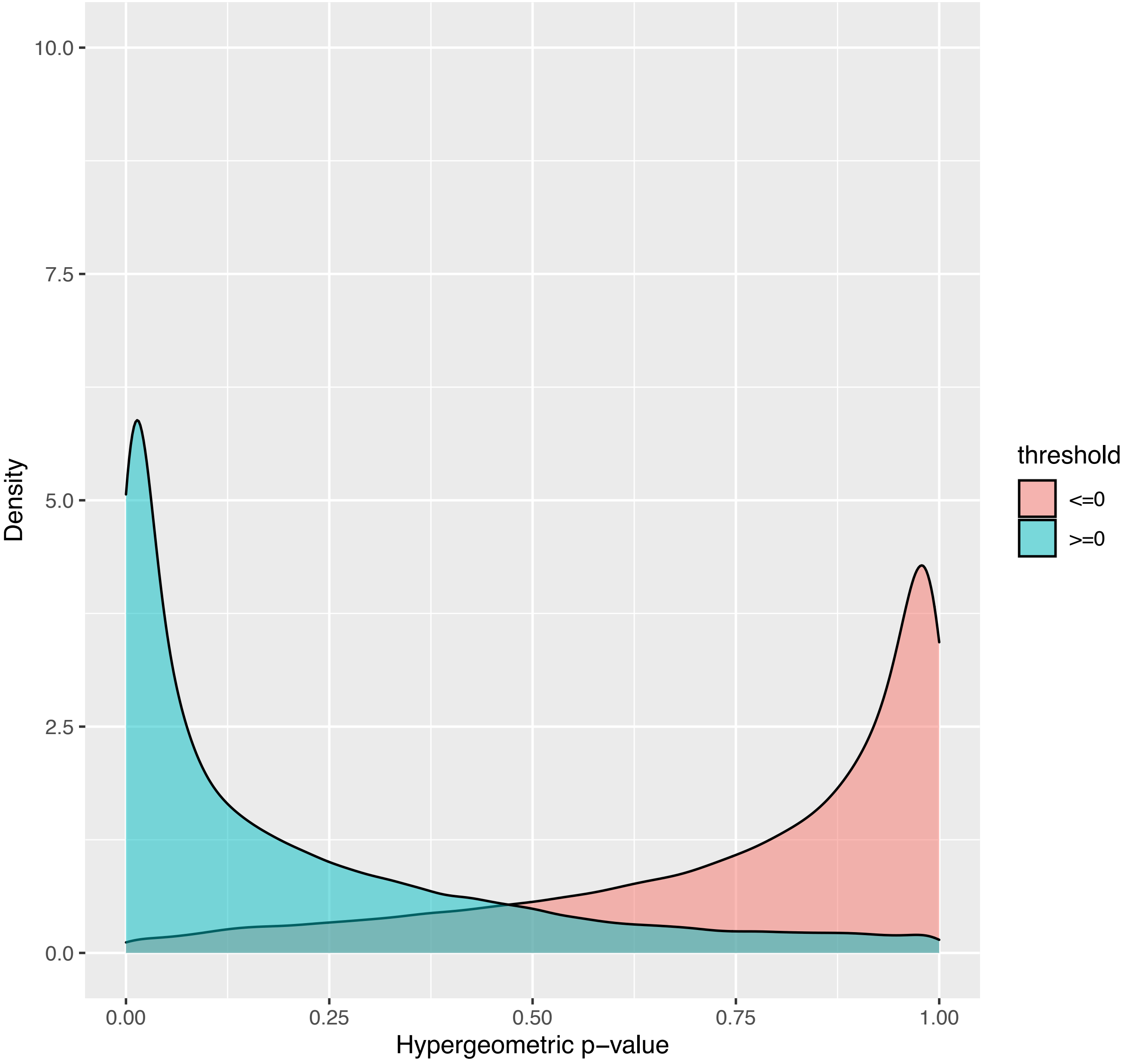

KIRC
No. of cell lines in low/high:32/33; p =0

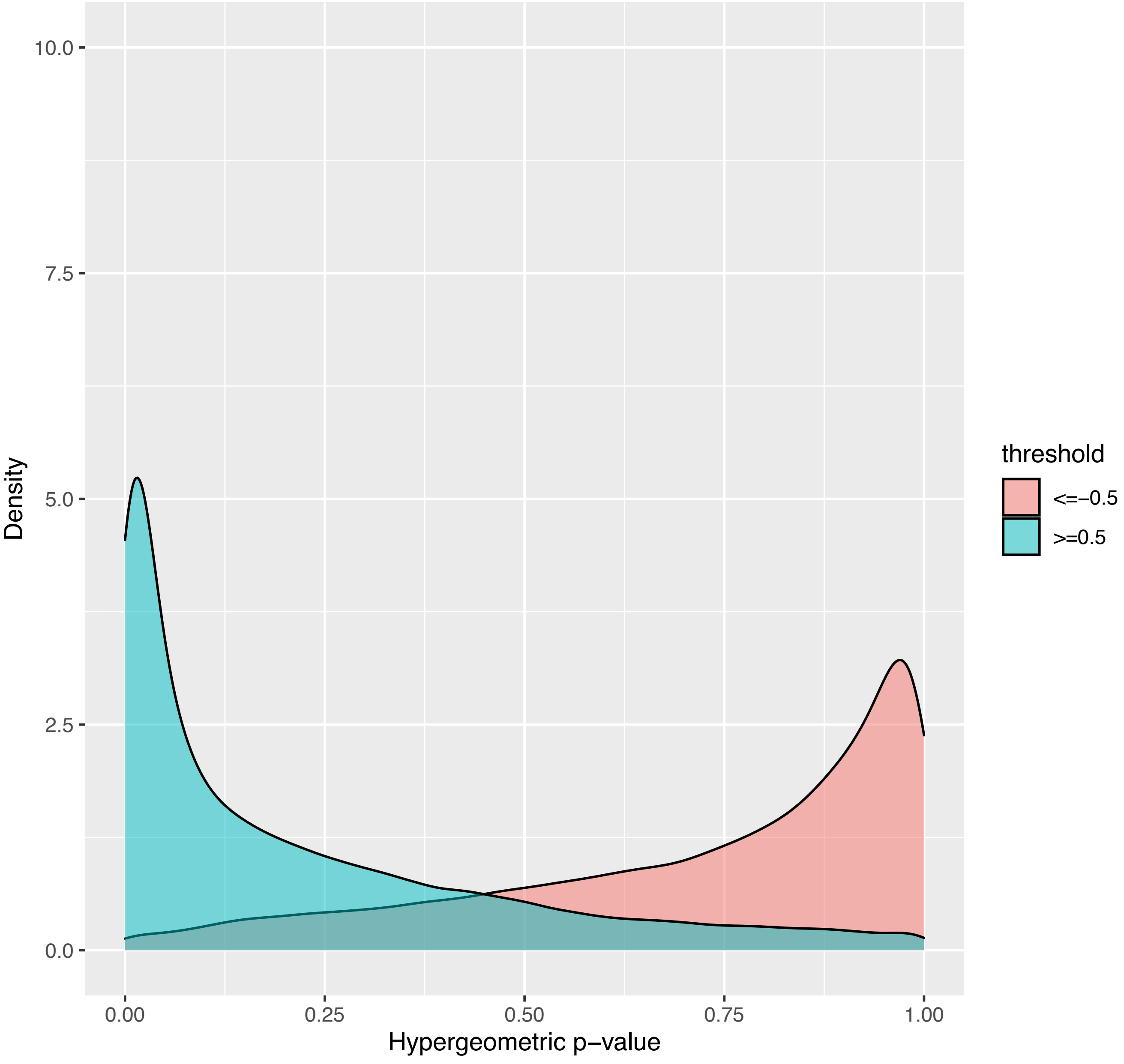

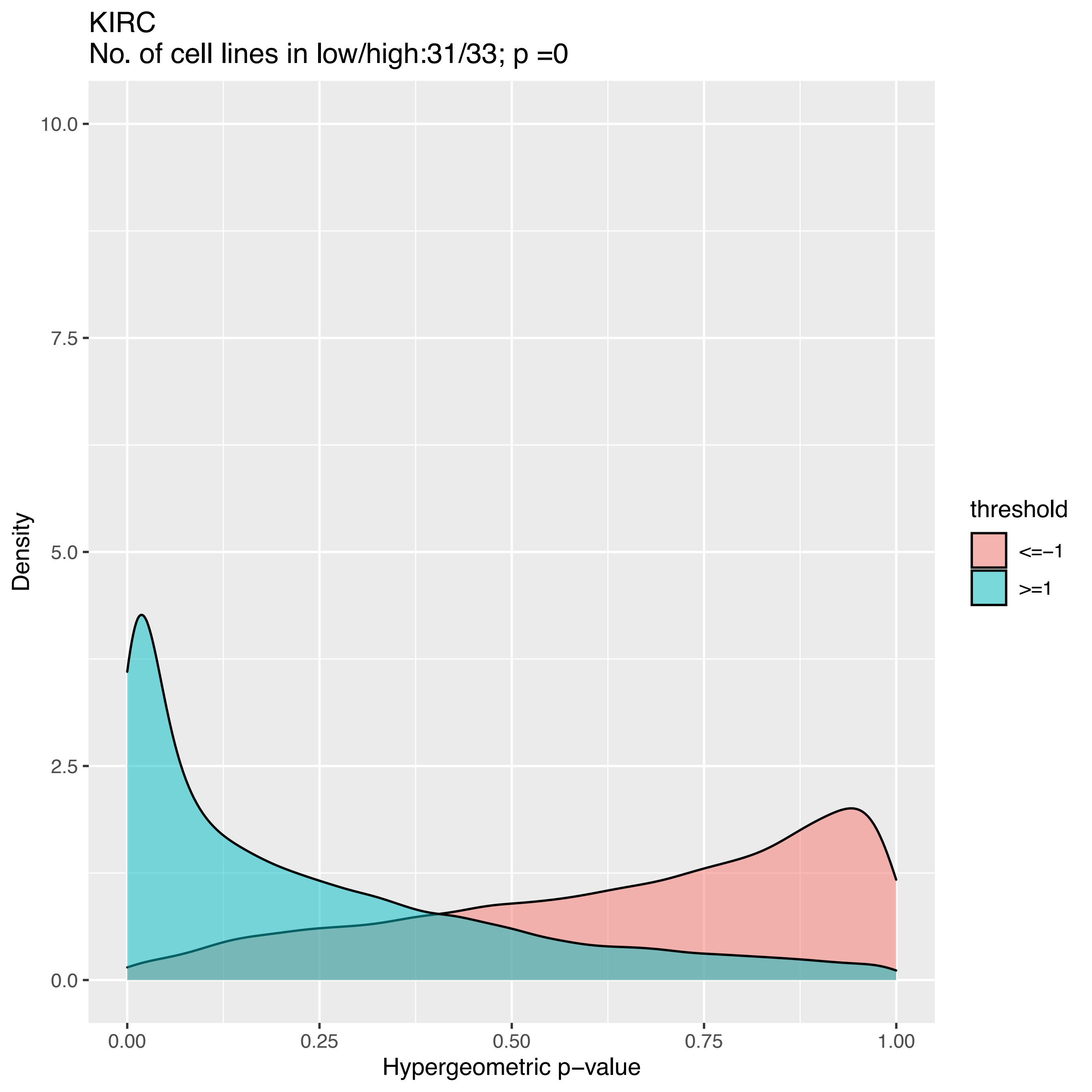

KIRC
No. of cell lines in low/high:31/33; p =0
Density
Hypergeometric p−value
10.0
7.5
5.0
2.5
0.0
0.00
0.25
0.50
0.75
1.00
threshold
<=−1
>=1

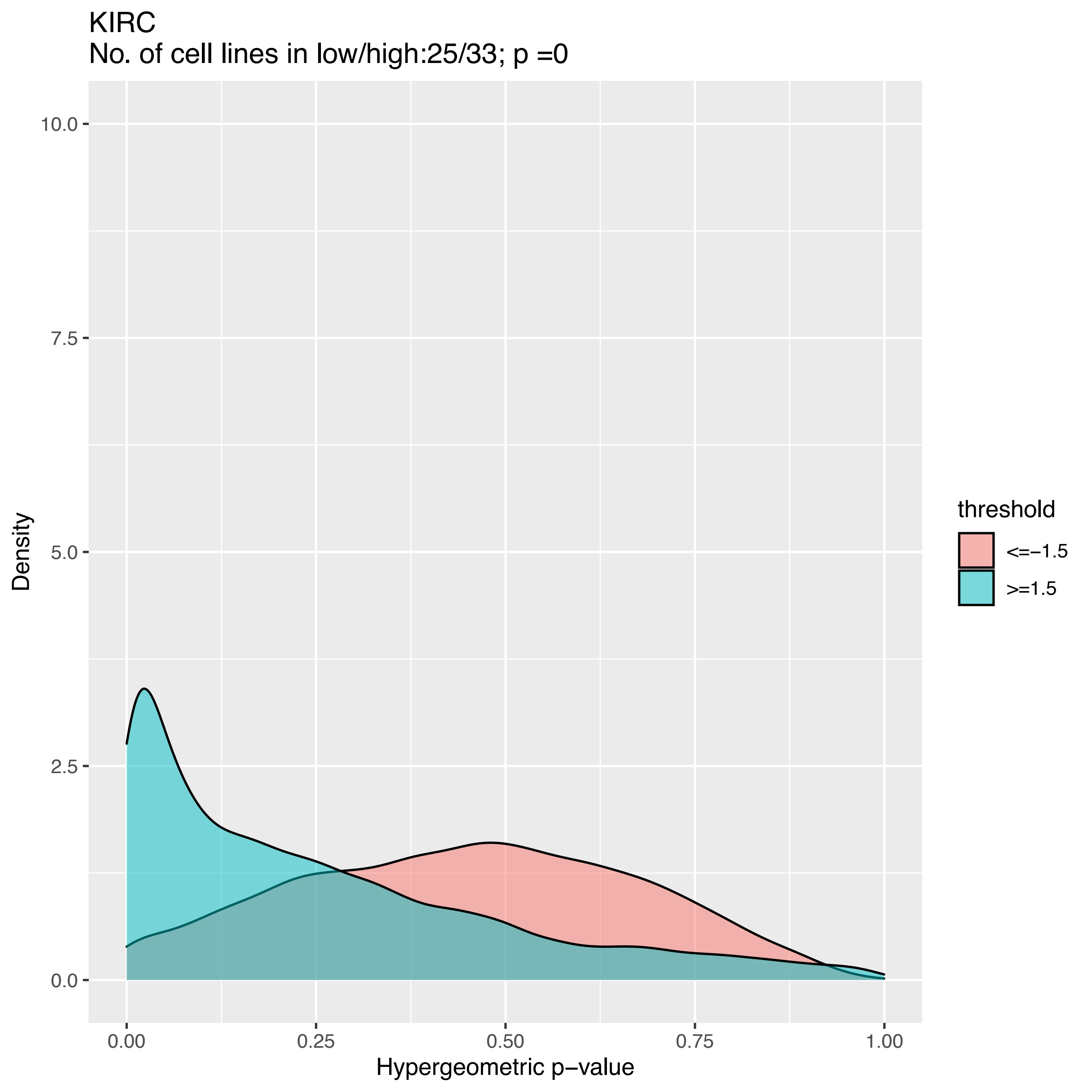

KIRC
No. of cell lines in low/high:25/33; p =0
10.0
7.5
5.0
2.5
0.0
Density
0.00
0.25
0.50
0.75
1.00
Hypergeometric p−value
threshold
<=−1.5
>=1.5

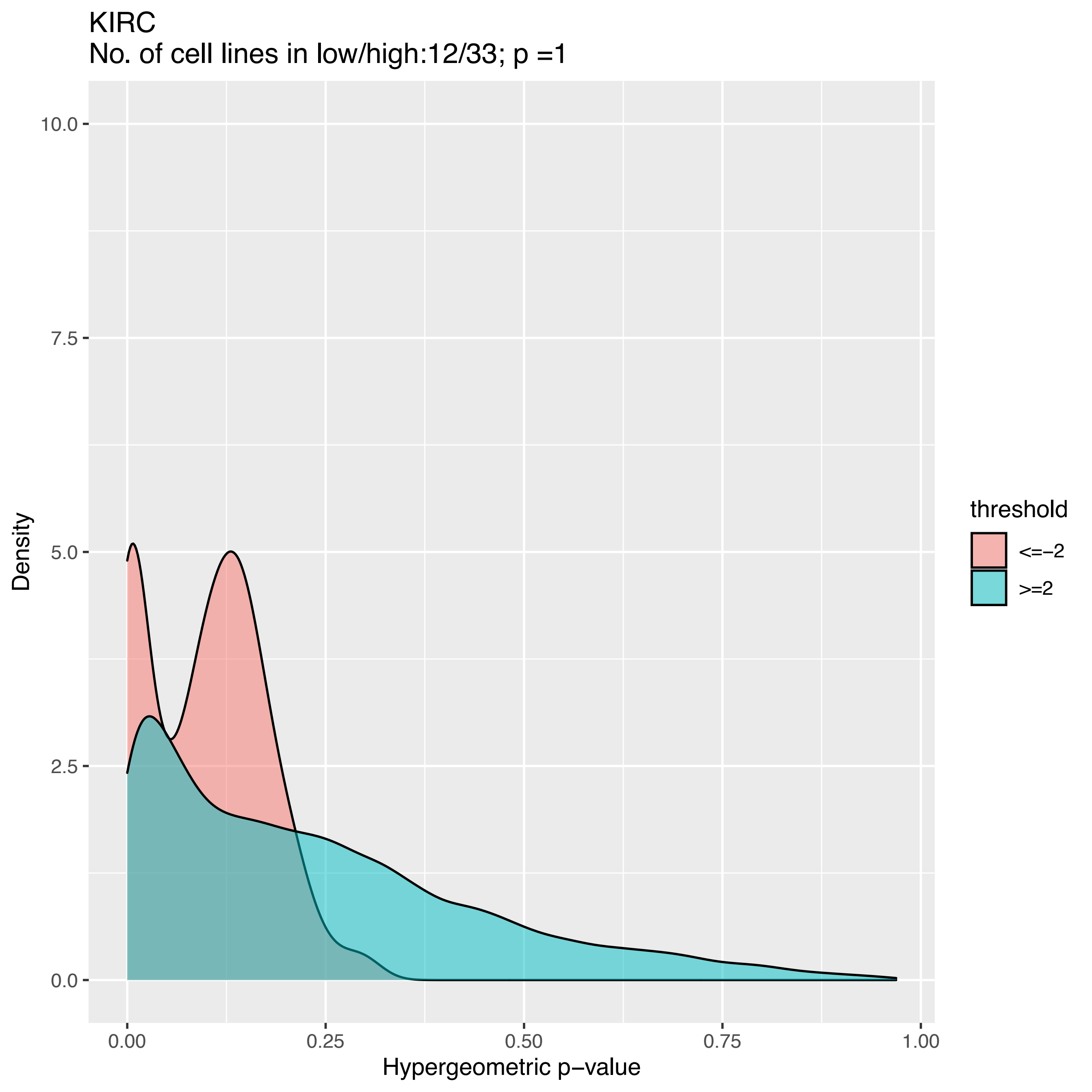

KIRC
No. of cell lines in low/high:12/33; p =1
Density
10.0
7.5
5.0
2.5
0.0
0.00
0.25
0.50
0.75
1.00
Hypergeometric p−value
threshold
<=−2
>=2

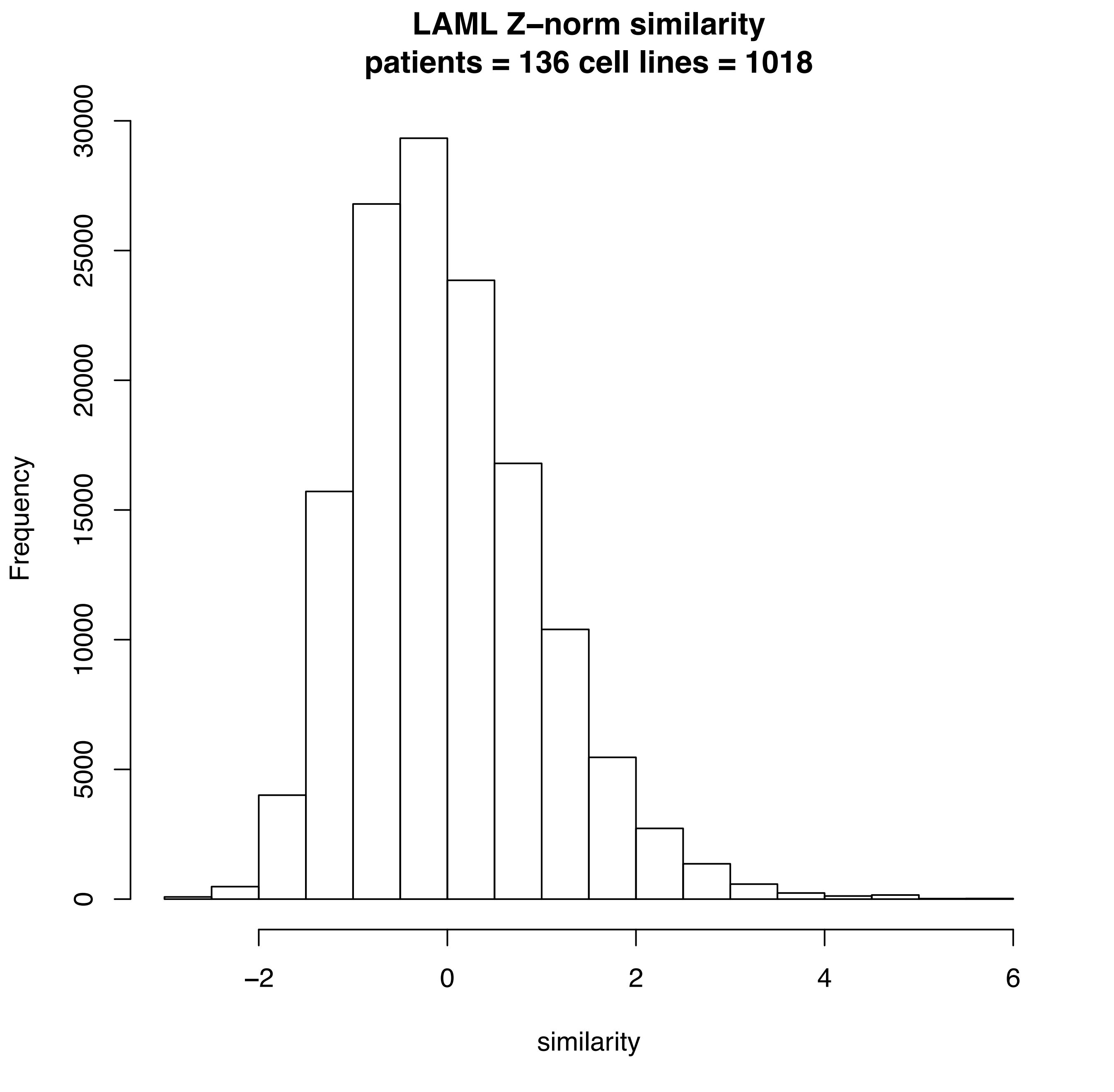

LAML Z–norm similarity
patients = 136 cell lines = 1018
Frequency
0
5000
10000
15000
20000
25000
30000
–2
0
2
4
6
similarity

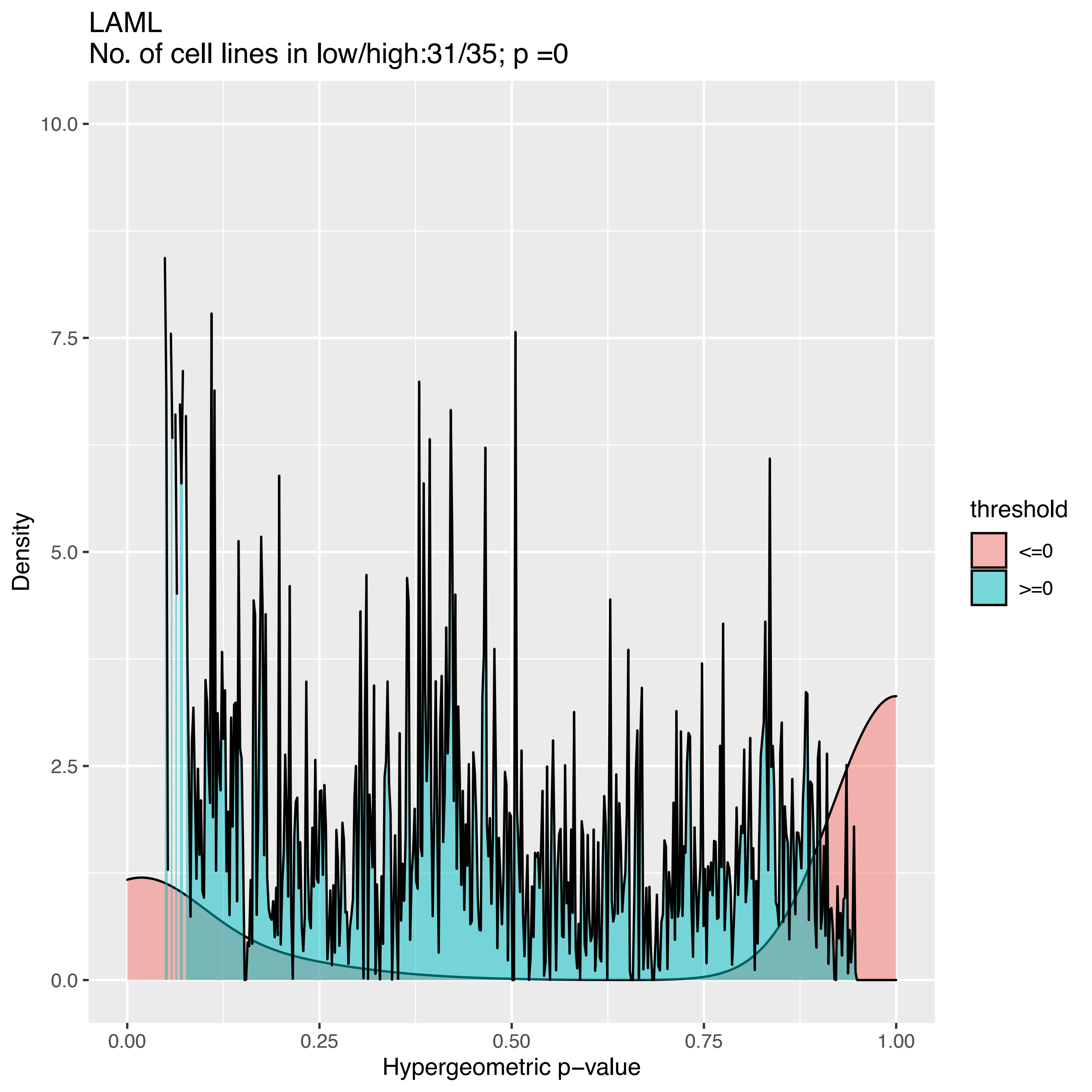

LAML
No. of cell lines in low/high:31/35; p =0
Density
10.0
7.5
5.0
2.5
0.0
0.00
0.25
0.50
0.75
1.00
Hypergeometric p−value
threshold
<=0
>=0

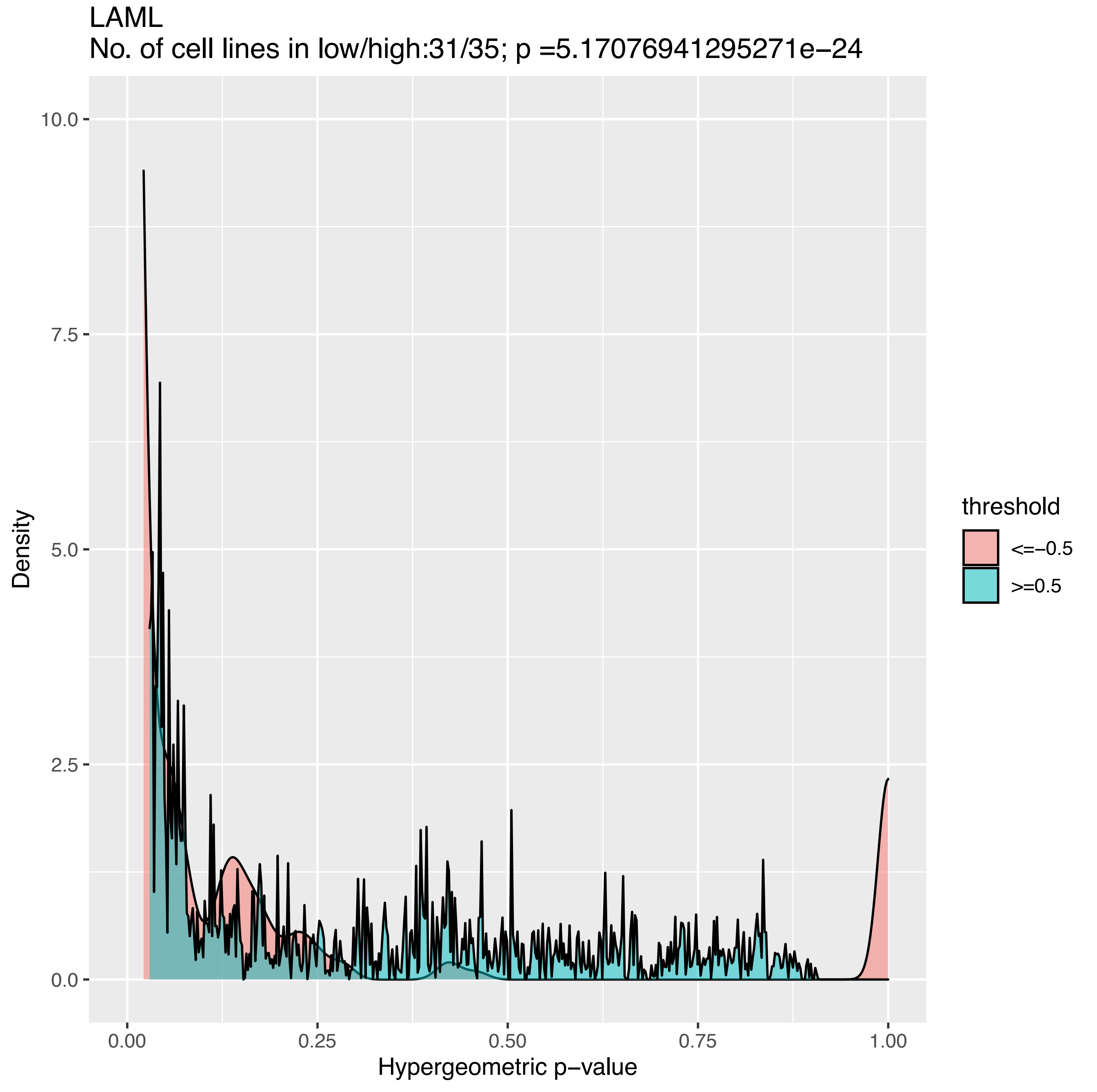
LAML
No. of cell lines in low/high:31/35; p =5.17076941295271e−24
Density
Hypergeometric p−value
10.0
7.5
5.0
2.5
0.0
0.00
0.25
0.50
0.75
1.00
threshold
<=−0.5
>=0.5

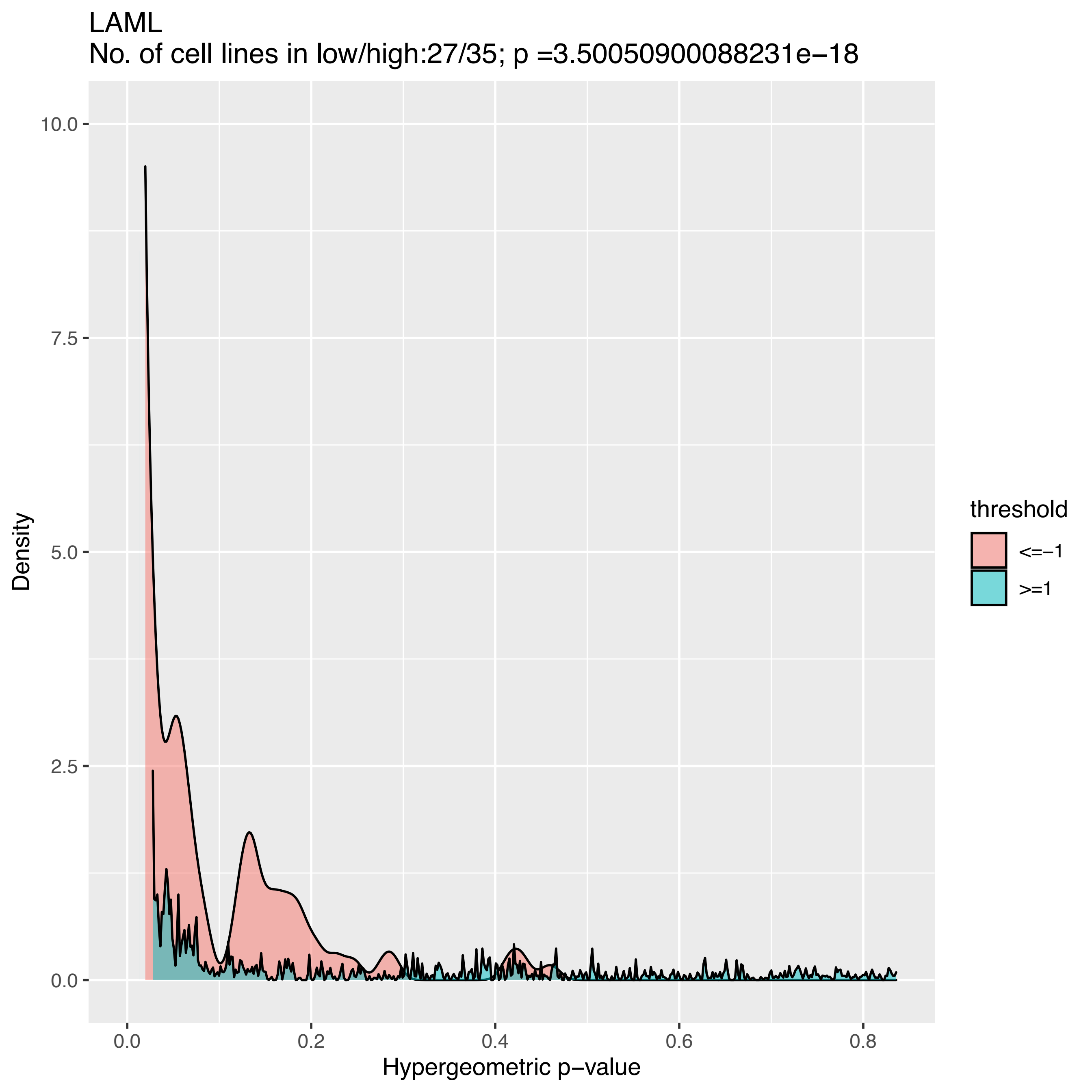
LAML
No. of cell lines in low/high:27/35; p =3.50050900088231e−18
Density
Hypergeometric p−value
threshold
<=−1
>=1
0.0
2.5
5.0
7.5
10.0
0.0
0.2
0.4
0.6
0.8

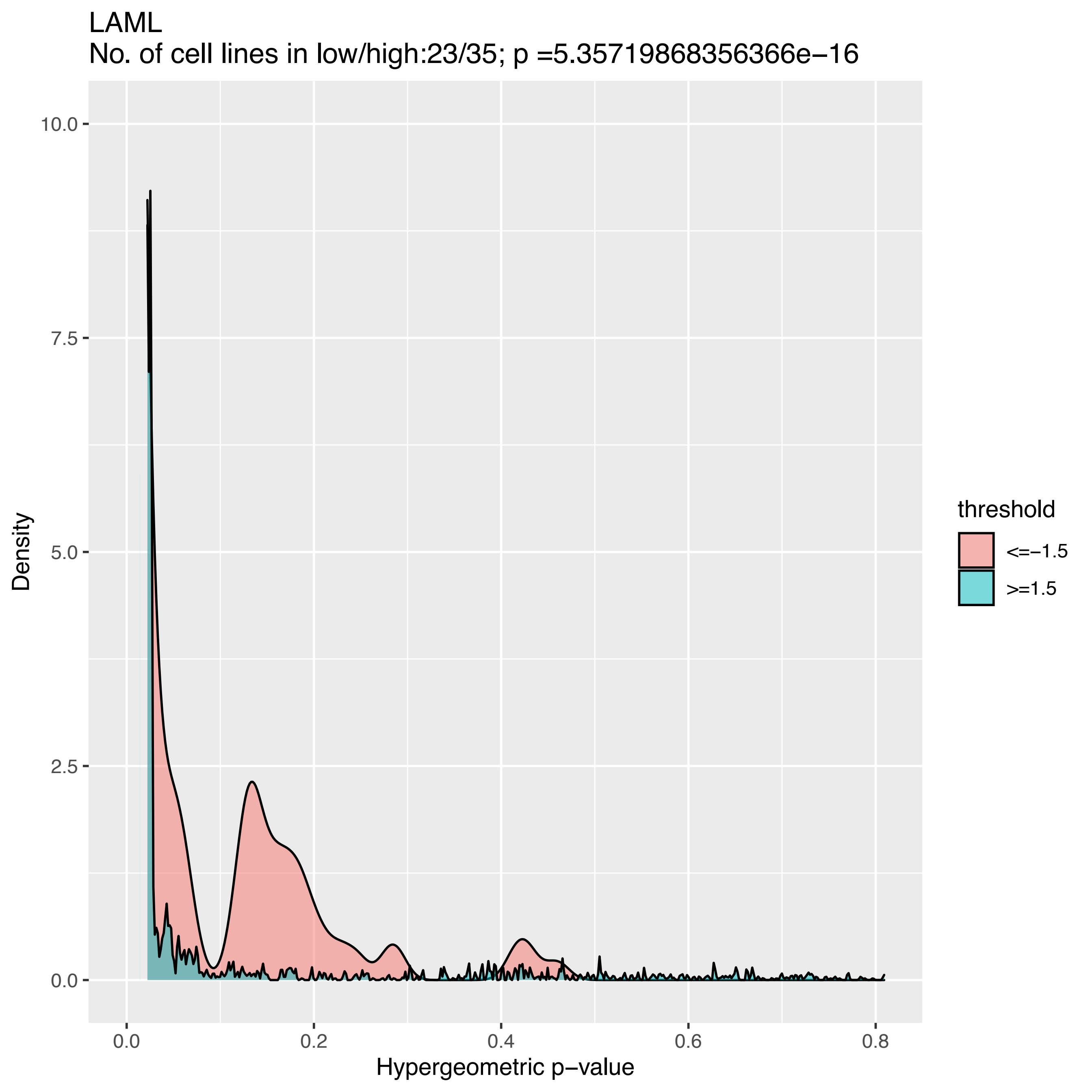
LAML
No. of cell lines in low/high:23/35; p =5.35719868356366e−16
Density
10.0
7.5
5.0
2.5
0.0
0.0
0.2
0.4
0.6
0.8
Hypergeometric p−value
threshold
<=−1.5
>=1.5

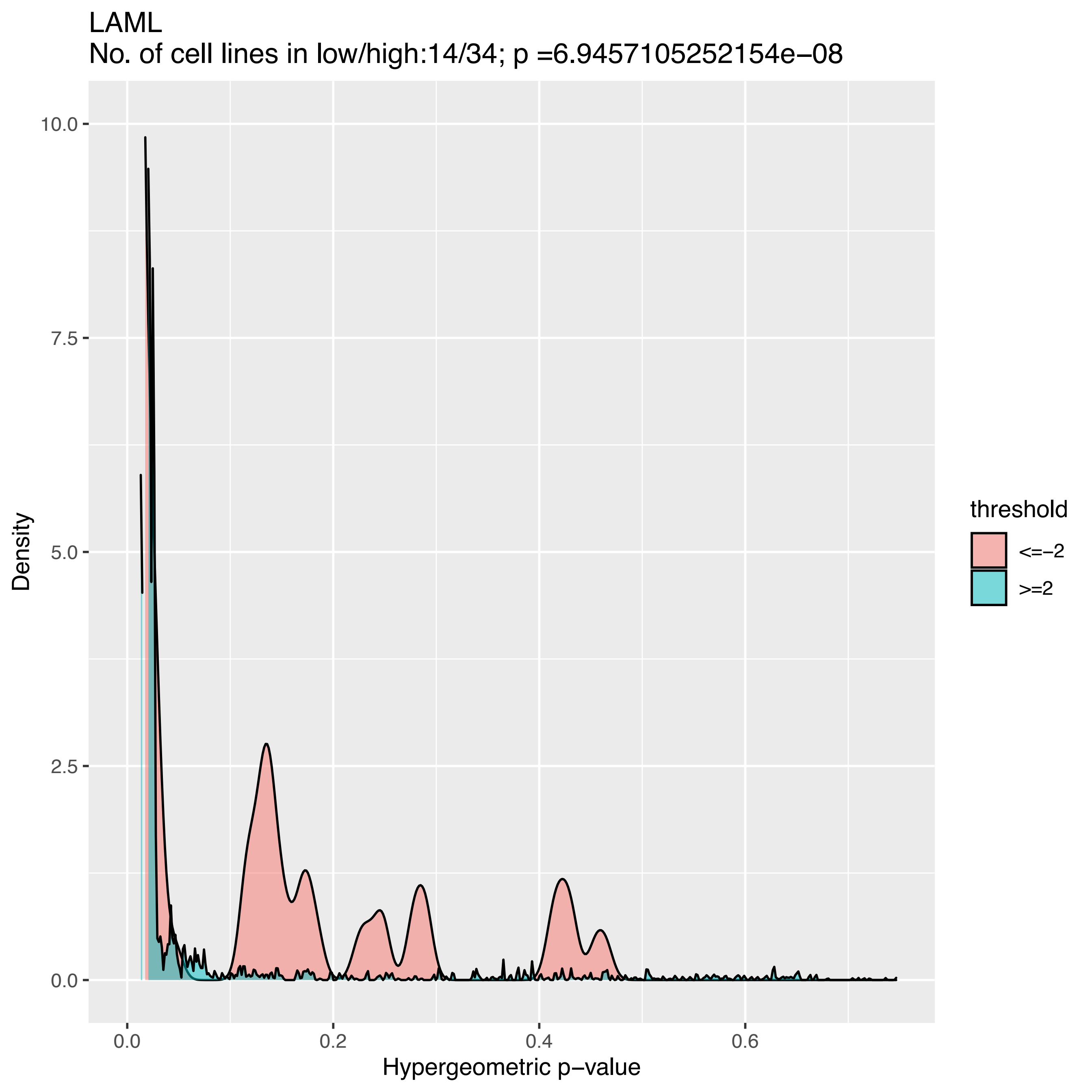

LAML
No. of cell lines in low/high:14/34; p =6.9457105252154e−08
10.0
7.5
5.0
2.5
0.0
Density
0.0
0.2
0.4
0.6
Hypergeometric p−value
threshold
<=−2
>=2

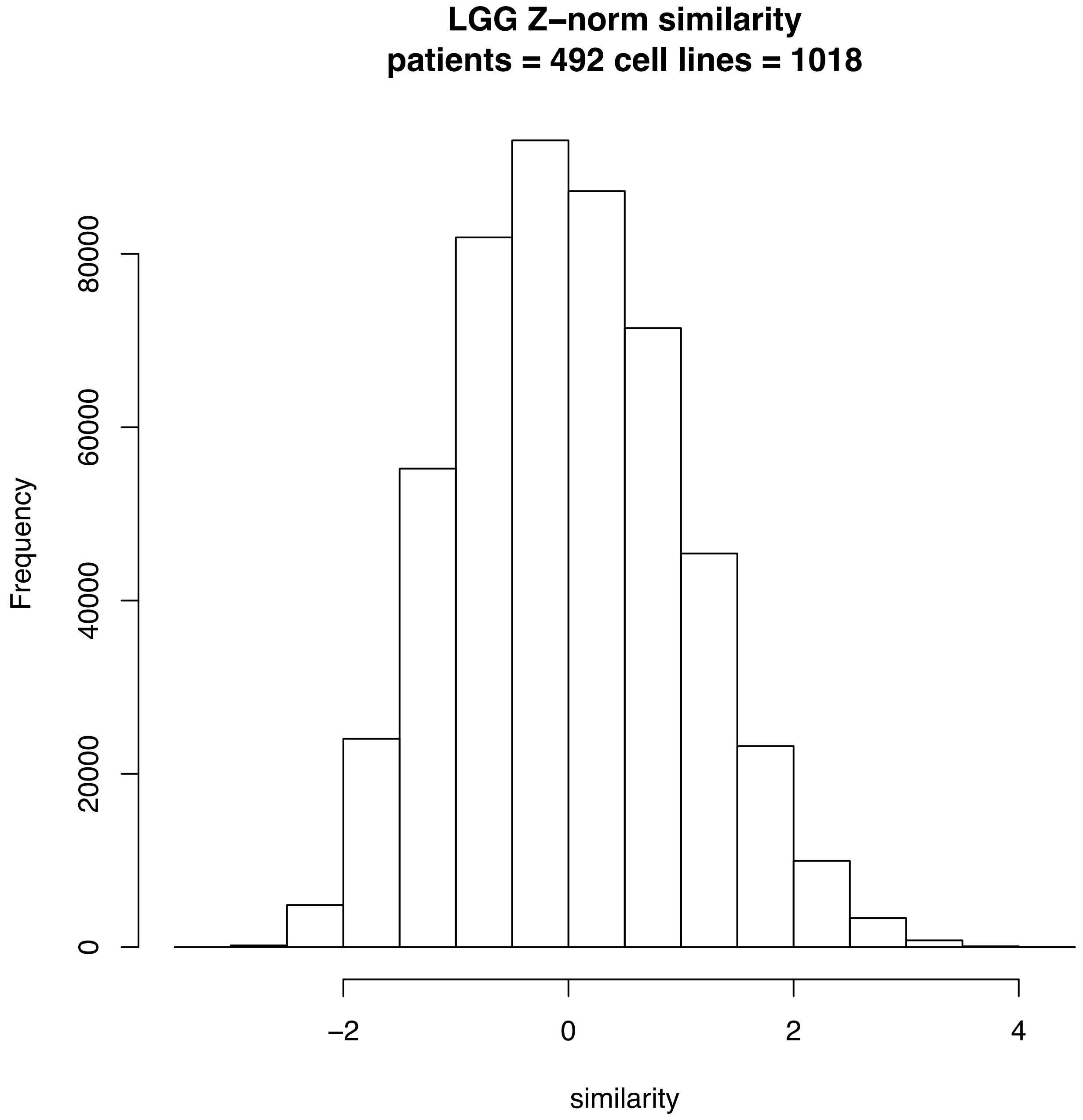
LGG Z–norm similarity
patients = 492 cell lines = 1018
Frequency
0
20000
40000
60000
80000
–2
0
2
4
similarity

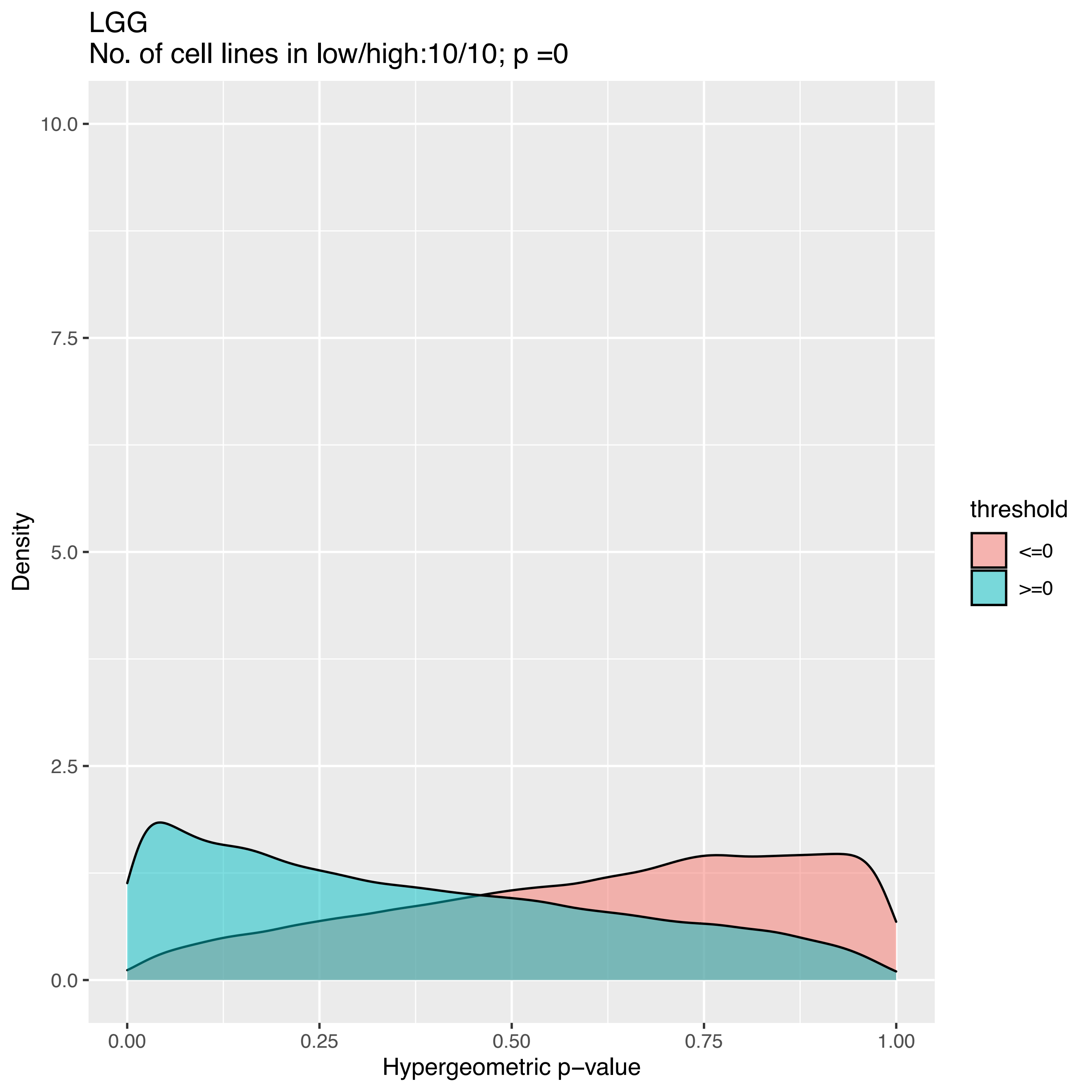
LGG
No. of cell lines in low/high:10/10; p =0
10.0
7.5
5.0
2.5
0.0
Density
0.00
0.25
0.50
0.75
1.00
Hypergeometric p−value
threshold
<=0
>=0

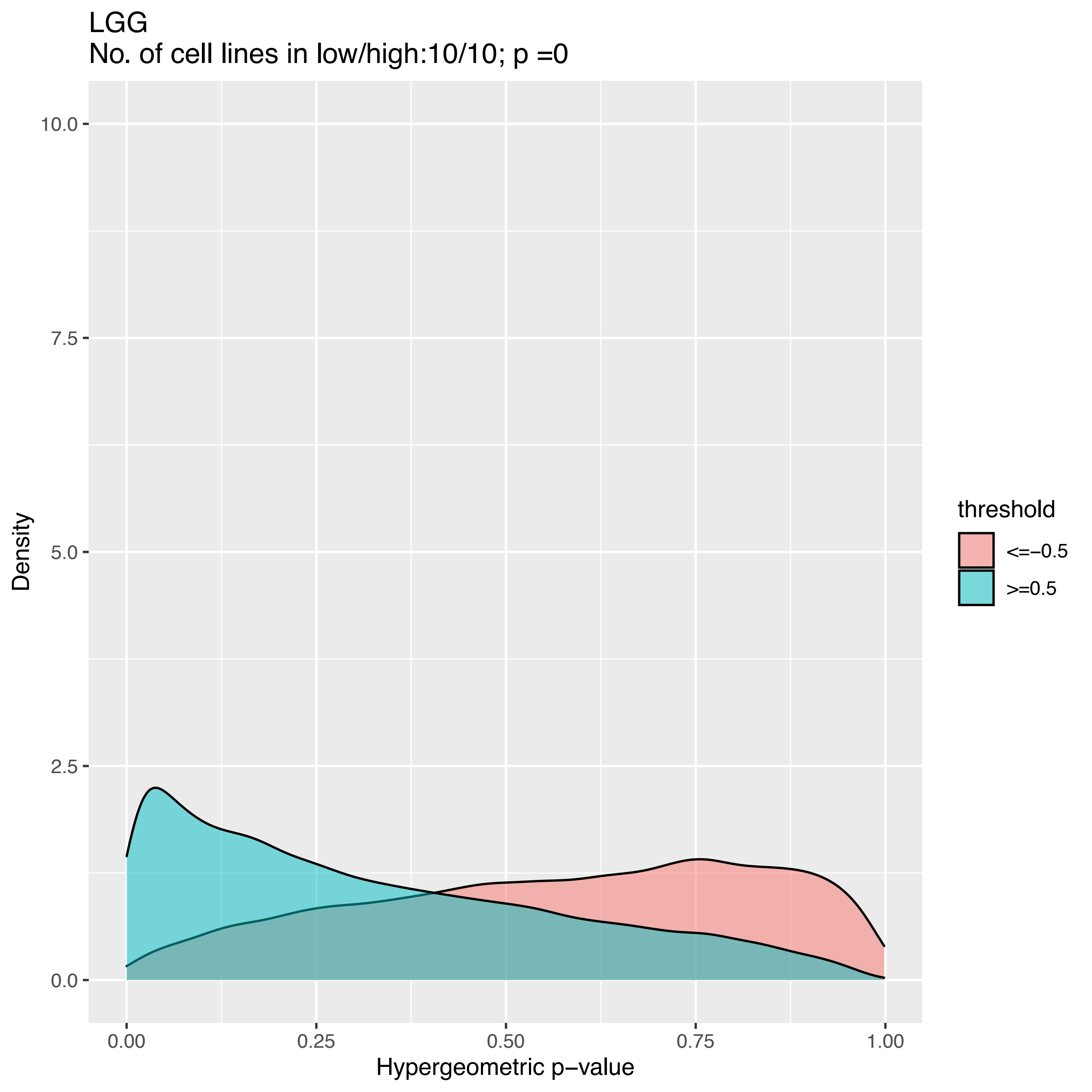

LGG
No. of cell lines in low/high:10/10; p =0
Density
10.0
7.5
5.0
2.5
0.0
0.00
0.25
0.50
0.75
1.00
Hypergeometric p−value
threshold
<=−0.5
>=0.5

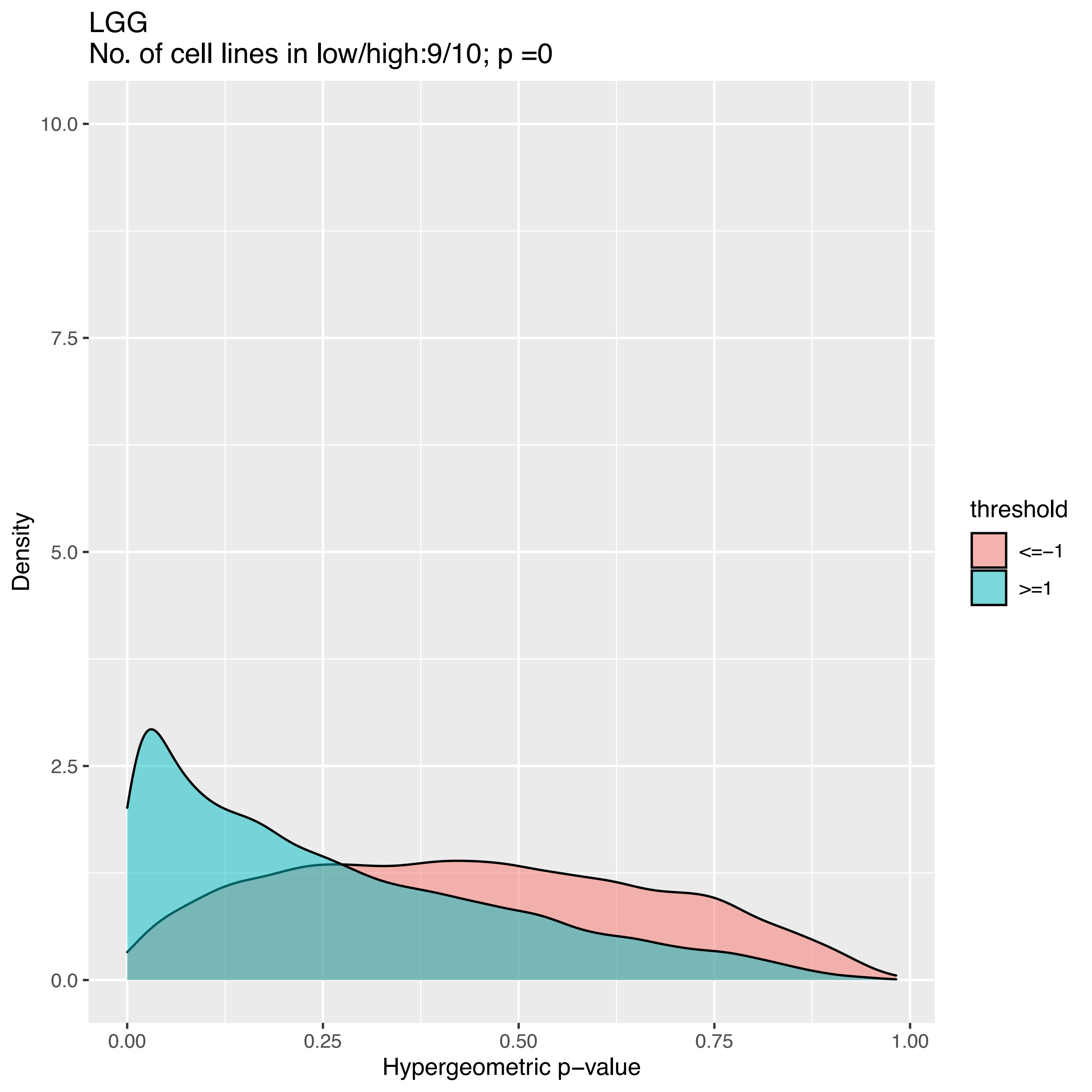
LGG
No. of cell lines in low/high:9/10; p =0
Density
Hypergeometric p−value
10.0
7.5
5.0
2.5
0.0
0.00
0.25
0.50
0.75
1.00
threshold
<=−1
>=1

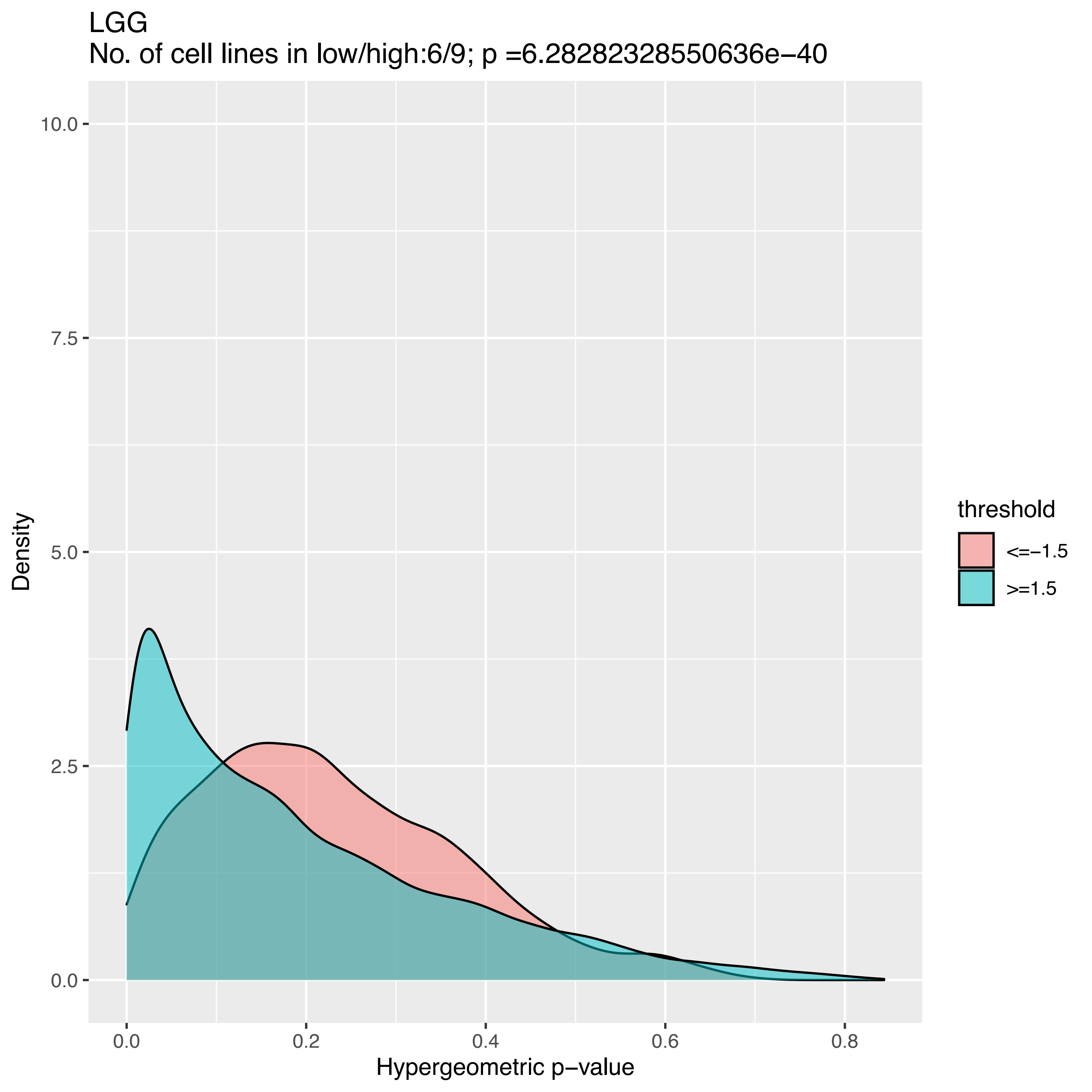
LGG
No. of cell lines in low/high:6/9; p =6.28282328550636e−40
10.0
7.5
5.0
2.5
0.0
Density
0.0
0.2
0.4
0.6
0.8
Hypergeometric p−value
threshold
<=−1.5
>=1.5

LGG
No. of cell lines in low/high:3/8; p =0.999999999834944

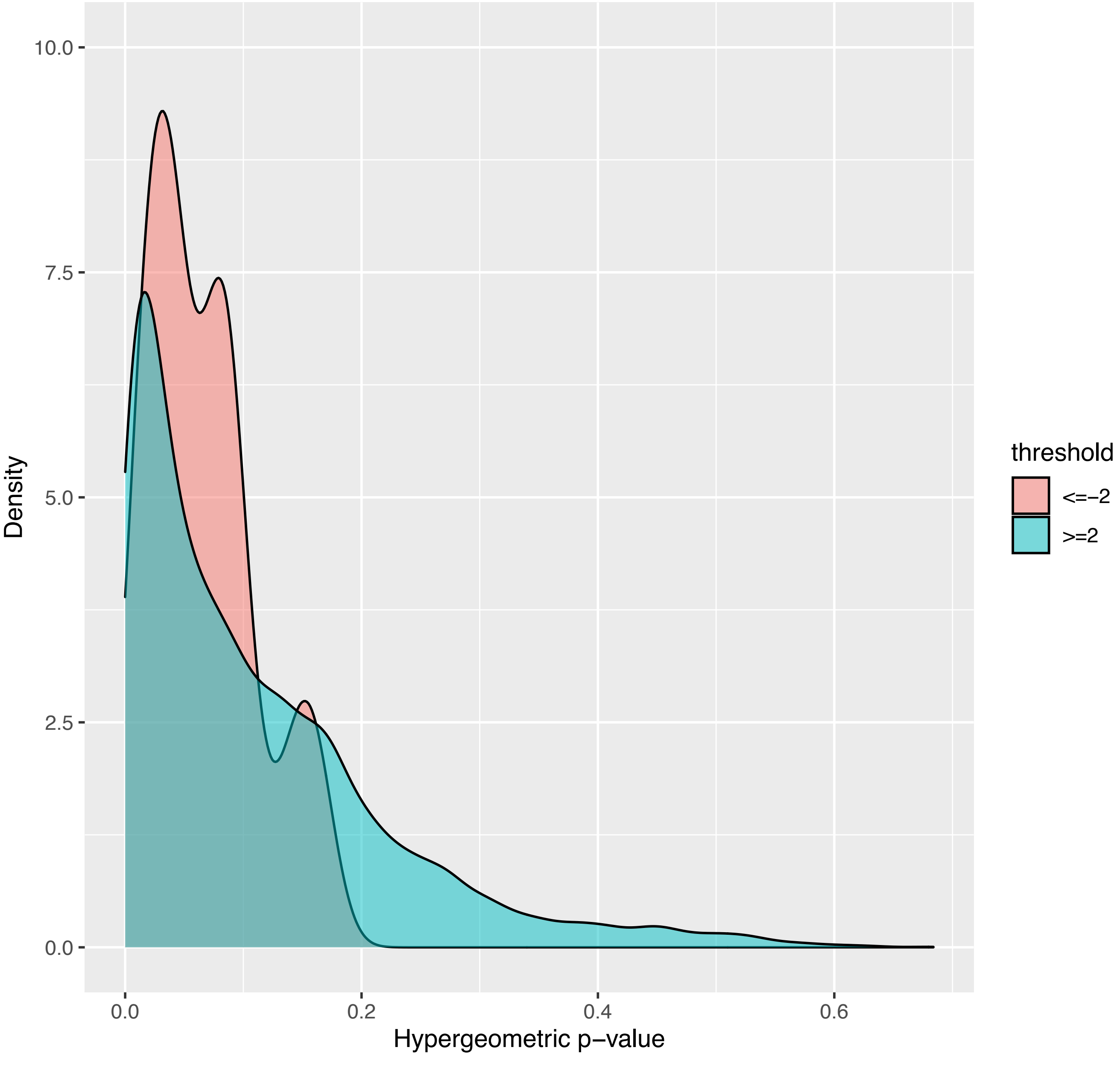

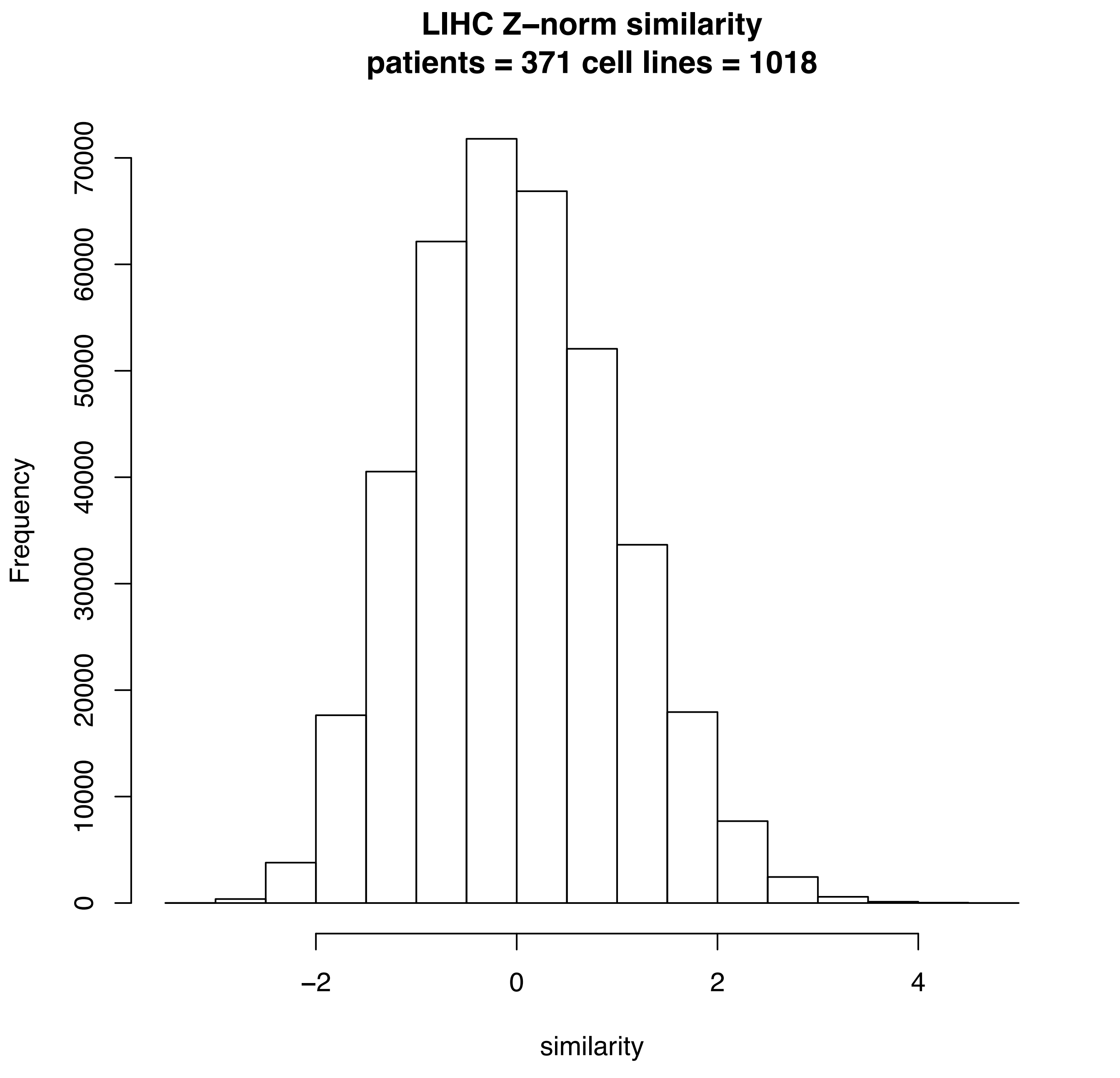
LIHC Z–norm similarity
patients = 371 cell lines = 1018
Frequency
0
10000
20000
30000
40000
50000
60000
70000
−2
0
2
4
similarity

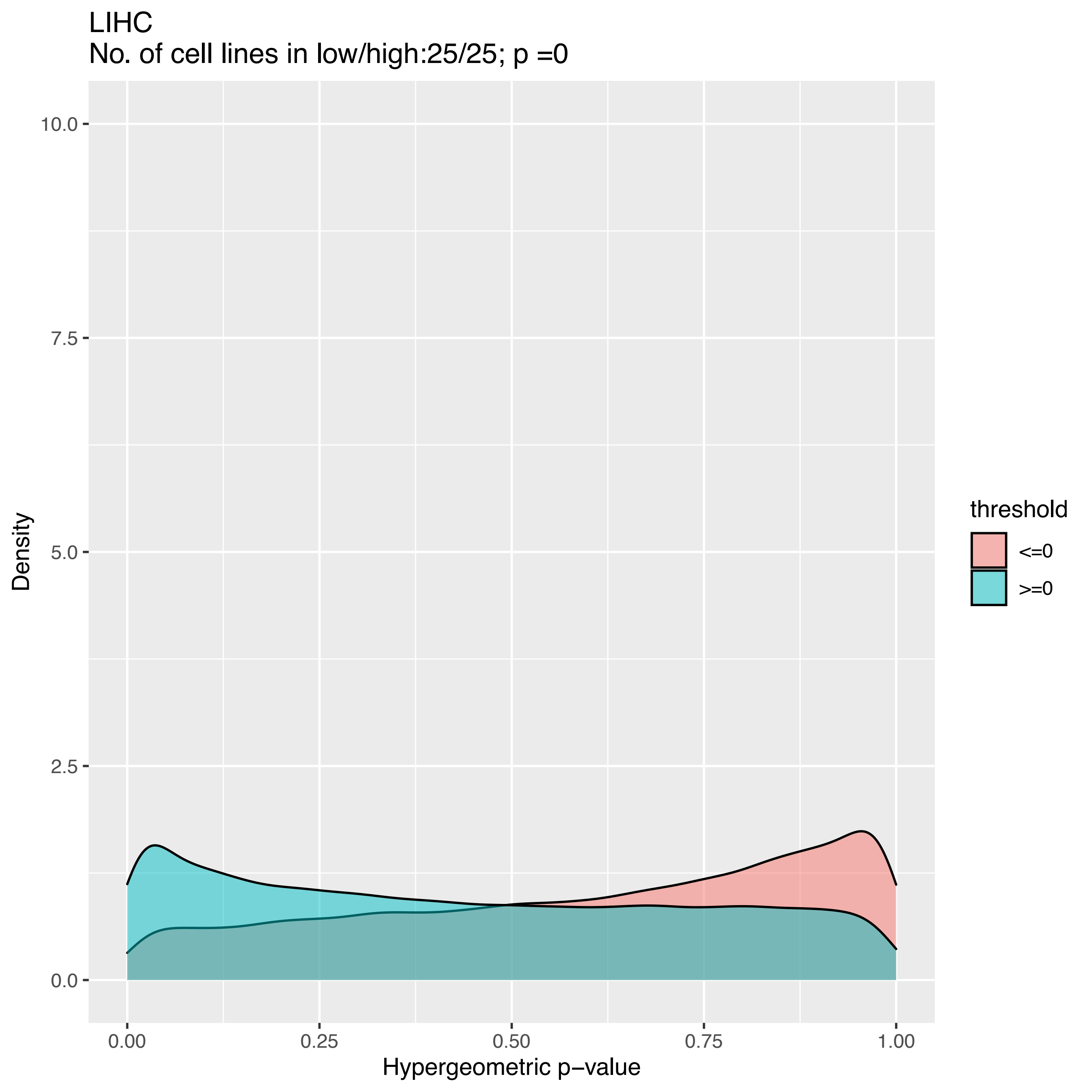
LIHC
No. of cell lines in low/high:25/25; p =0
Density
10.0
7.5
5.0
2.5
0.0
0.00
0.25
0.50
0.75
1.00
Hypergeometric p−value
threshold
<=0
>=0

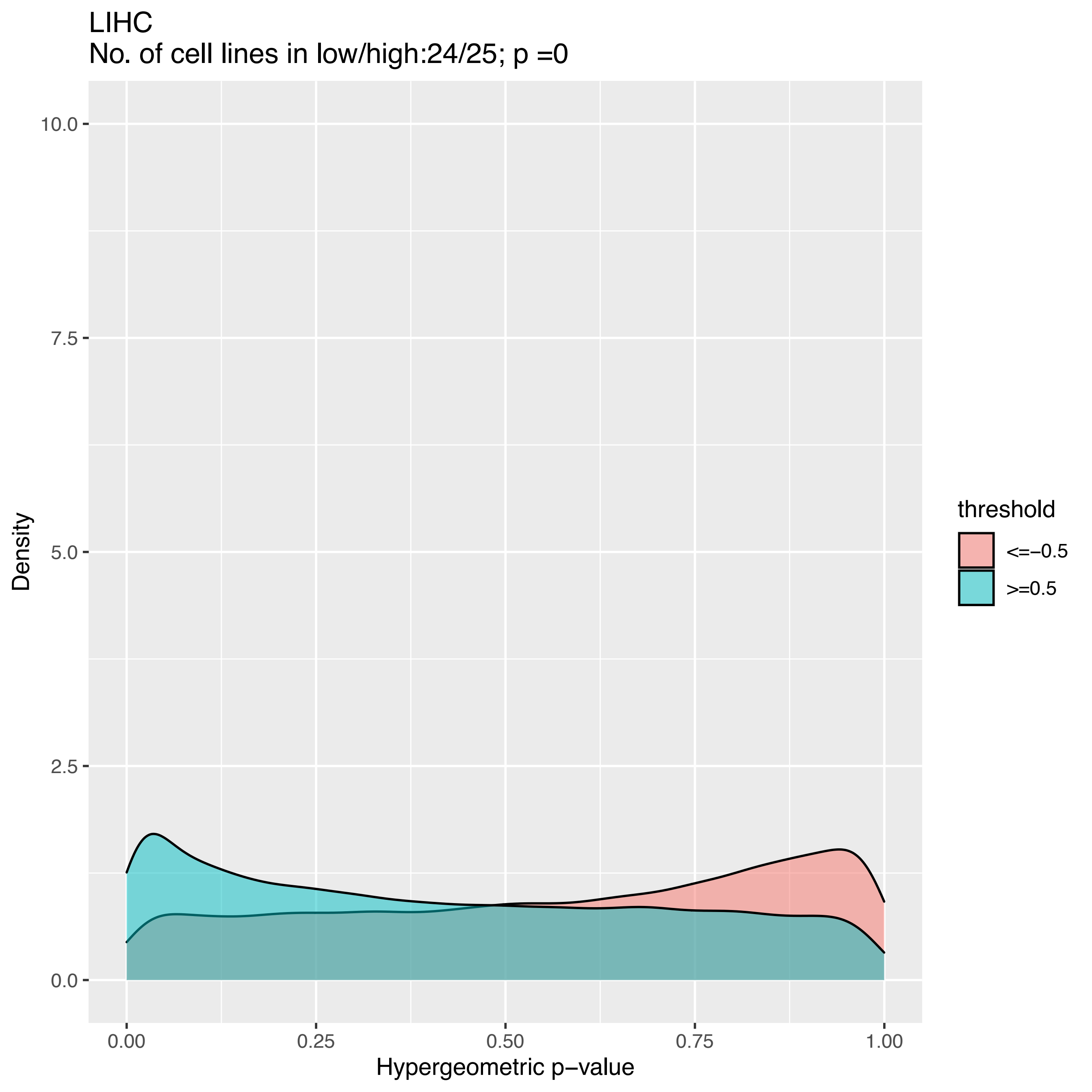
LIHC
No. of cell lines in low/high:24/25; p =0
10.0
7.5
5.0
2.5
0.0
Density
0.00
0.25
0.50
0.75
1.00
Hypergeometric p−value
threshold
<=−0.5
>=0.5

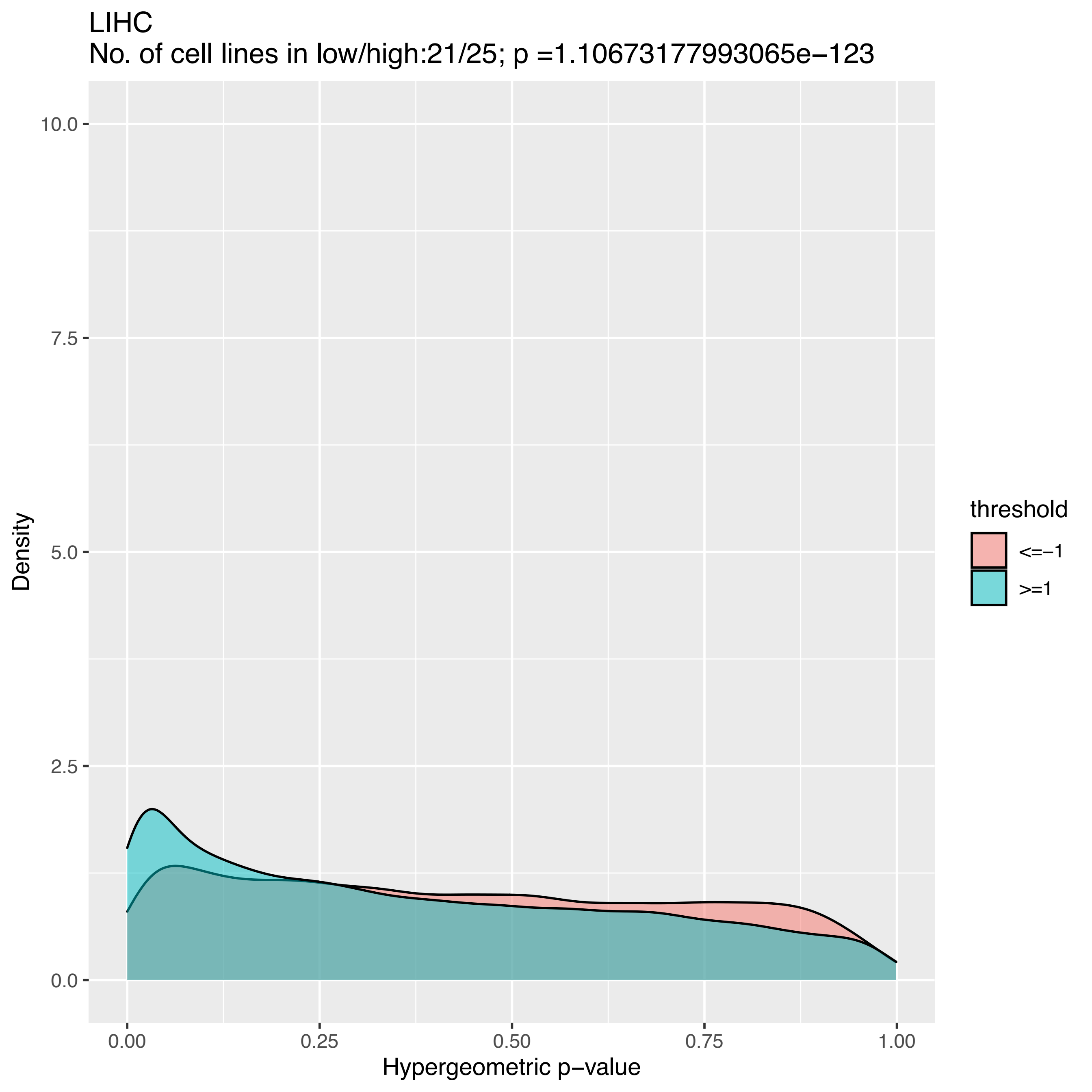

LIHC
No. of cell lines in low/high:21/25; p =1.10673177993065e−123
Density
10.0
7.5
5.0
2.5
0.0
0.00
0.25
0.50
0.75
1.00
Hypergeometric p−value
threshold
<=−1
>=1

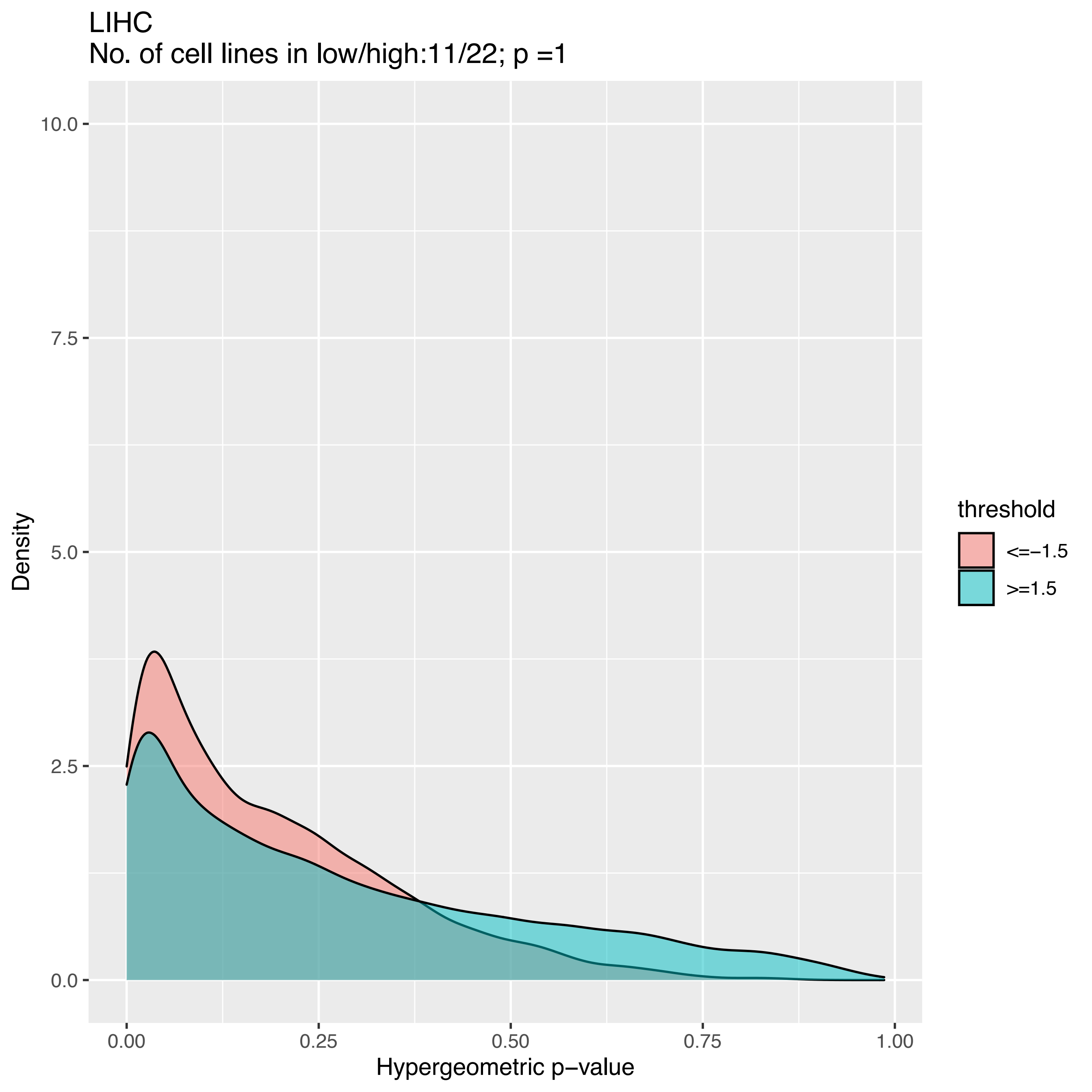

LIHC
No. of cell lines in low/high:11/22; p =1
10.0
7.5
5.0
2.5
0.0
Density
0.00
0.25
0.50
0.75
1.00
Hypergeometric p−value
threshold
<=−1.5
>=1.5

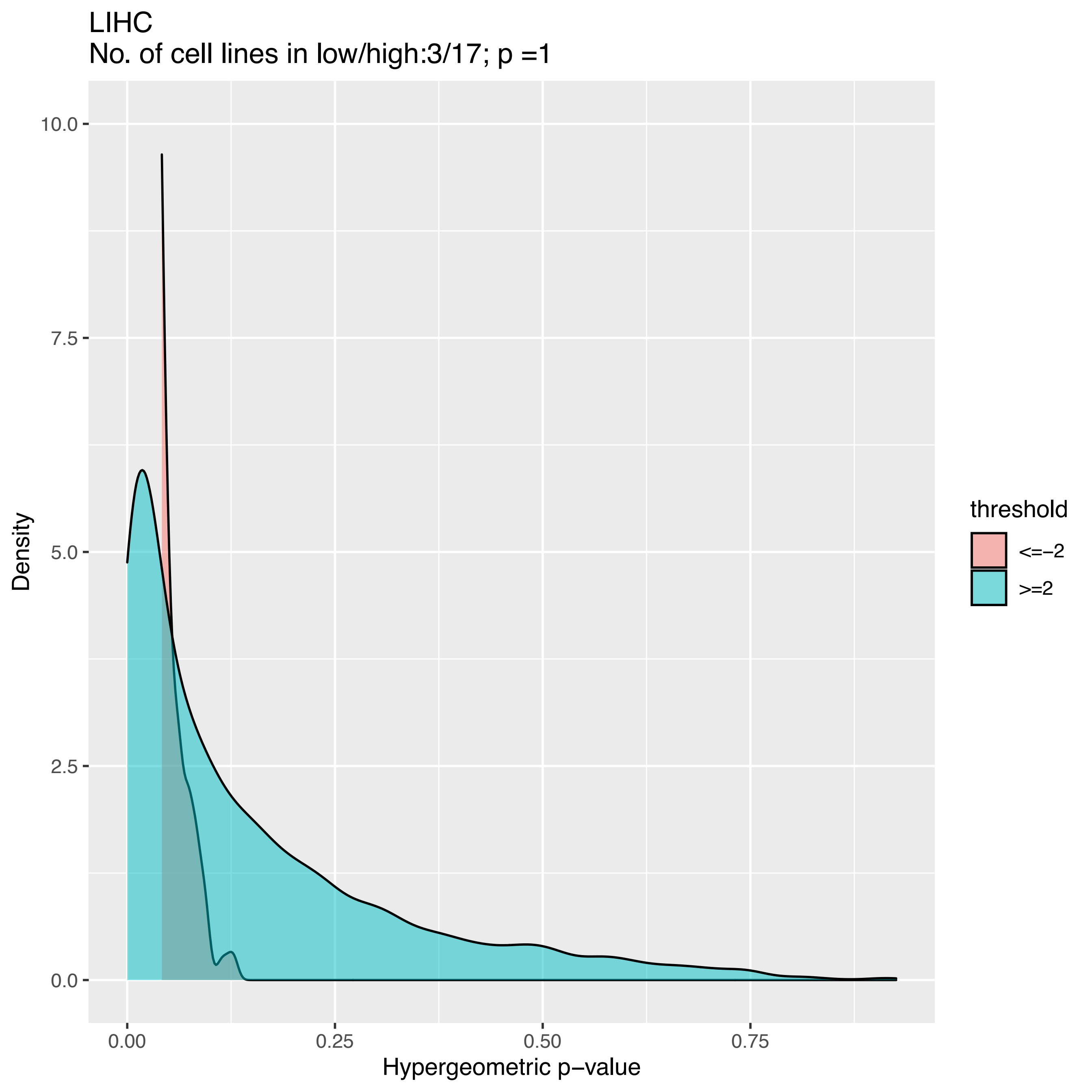

LIHC
No. of cell lines in low/high:3/17; p =1
Density
Hypergeometric p−value
10.0
7.5
5.0
2.5
0.0
0.00
0.25
0.50
0.75
threshold
<=−2
>=2

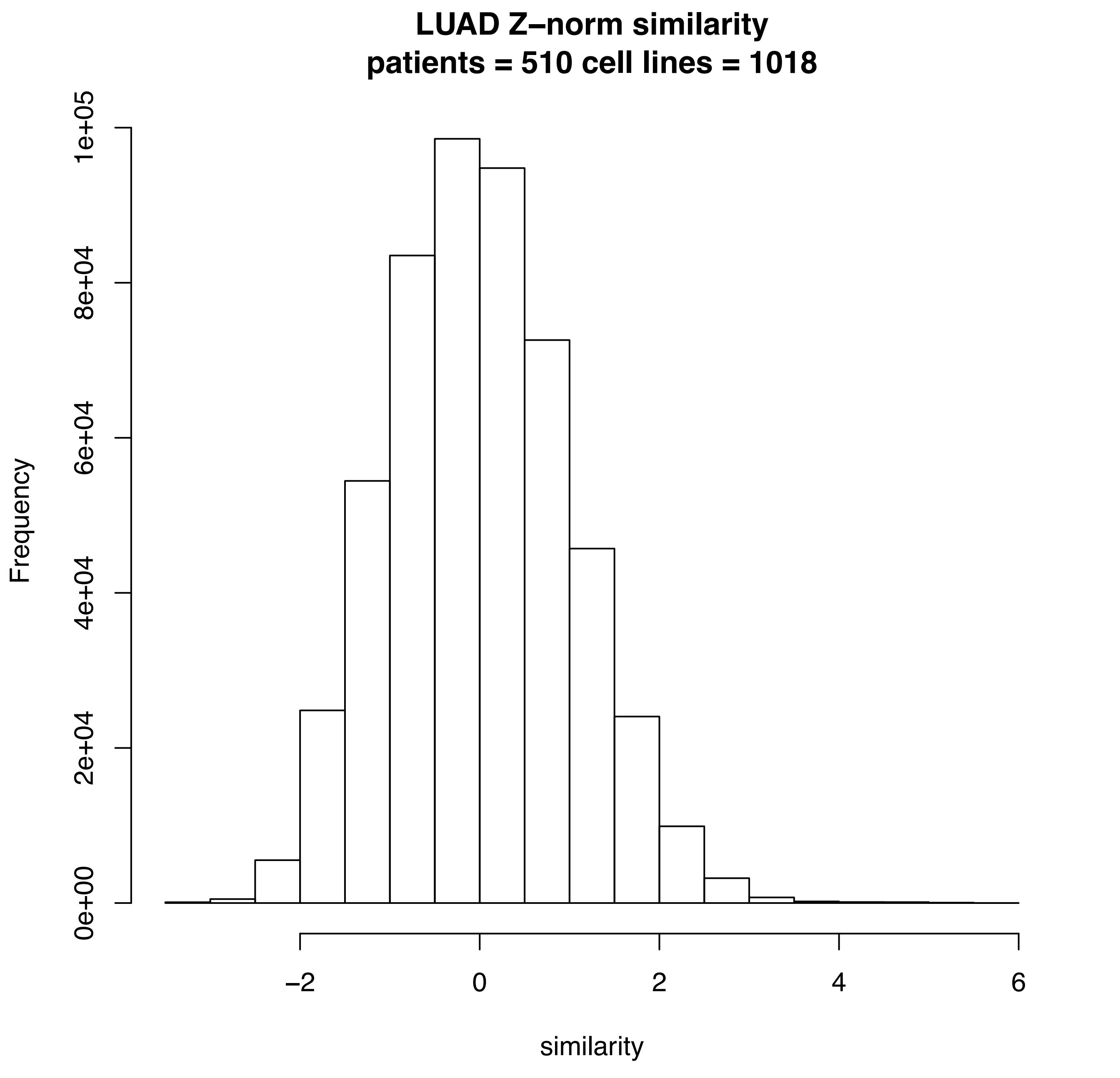
LUAD Z–norm similarity
patients = 510 cell lines = 1018
Frequency
0e+00
2e+04
4e+04
6e+04
8e+04
1e+05
similarity
–2
0
2
4
6

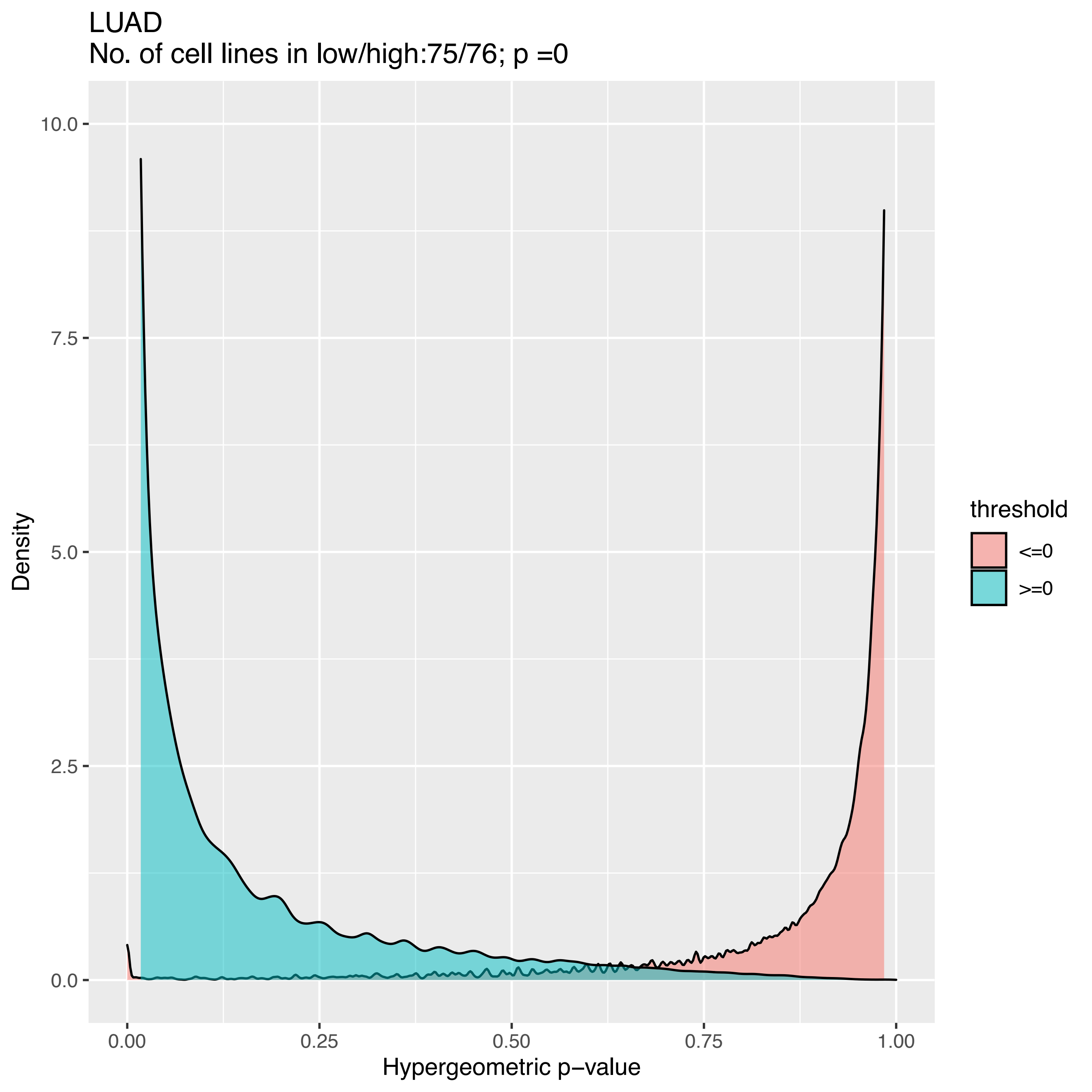
LUAD
No. of cell lines in low/high:75/76; p =0
10.0
7.5
5.0
2.5
0.0
0.00
0.25
0.50
0.75
1.00
Density
Hypergeometric p−value
threshold
<=0
>=0

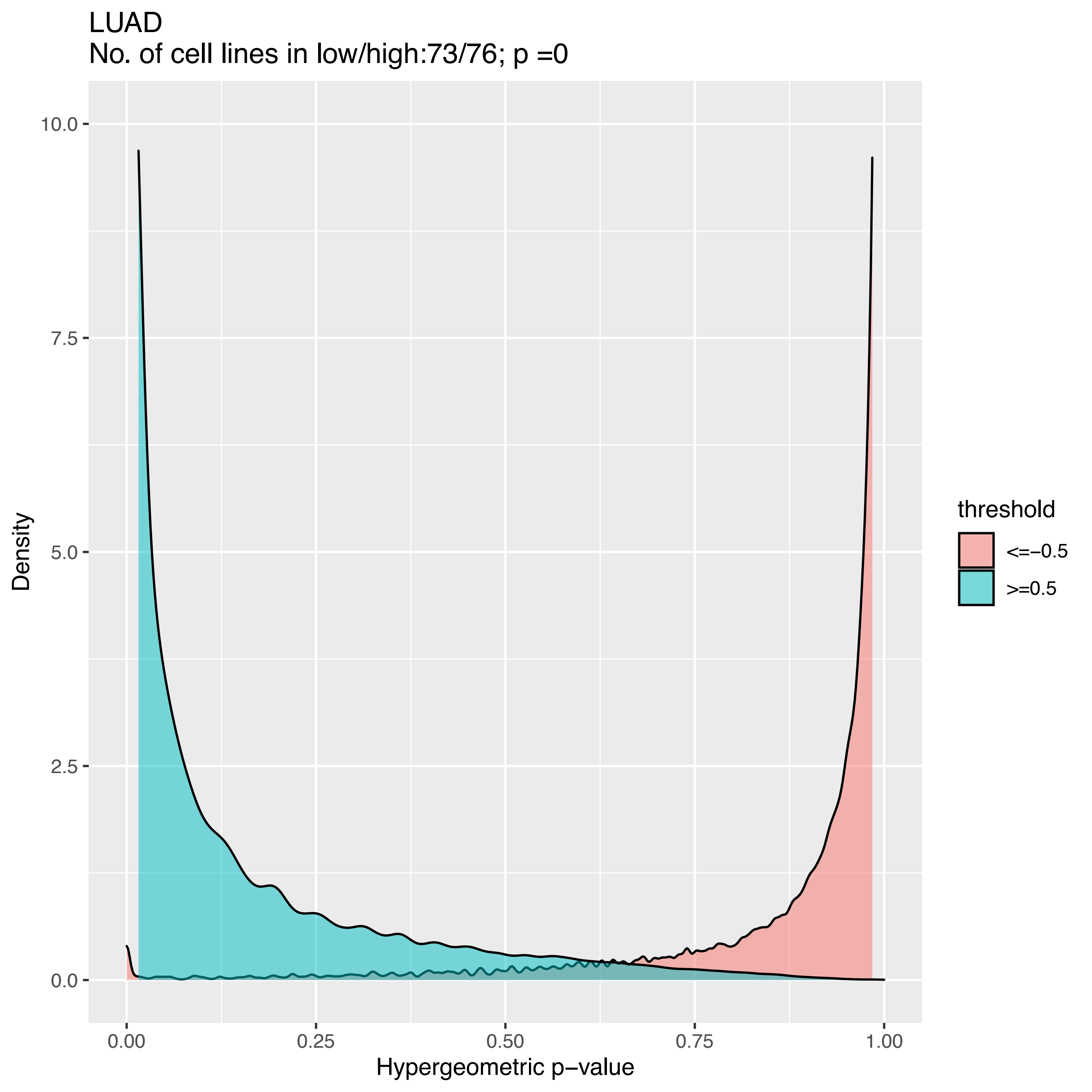

LUAD
No. of cell lines in low/high:73/76; p =0
Density
10.0
7.5
5.0
2.5
0.0
0.00
0.25
0.50
0.75
1.00
Hypergeometric p−value
threshold
<=−0.5
>=0.5

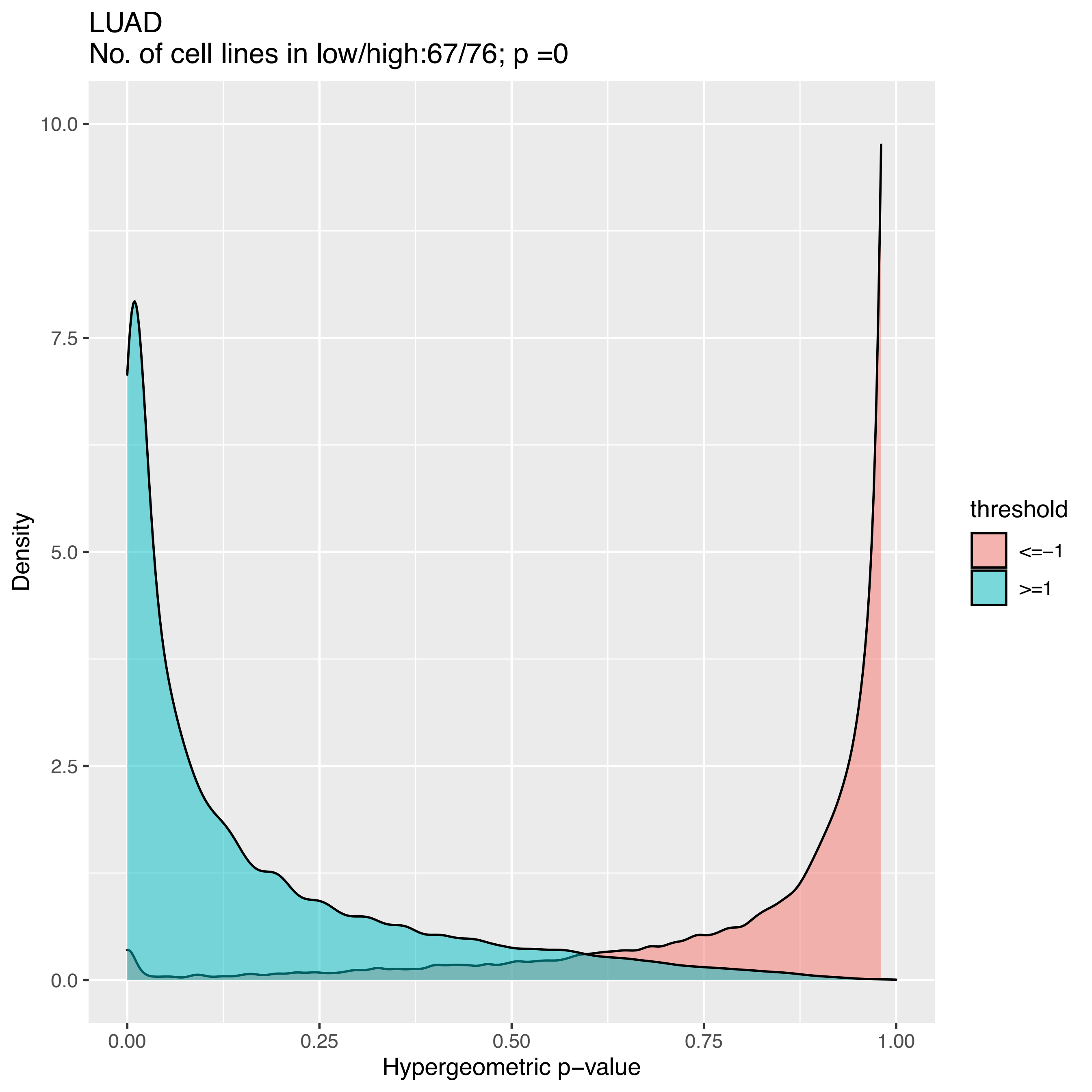

LUAD
No. of cell lines in low/high:67/76; p =0
Density
10.0
7.5
5.0
2.5
0.0
0.00
0.25
0.50
0.75
1.00
Hypergeometric p−value
threshold
<=−1
>=1

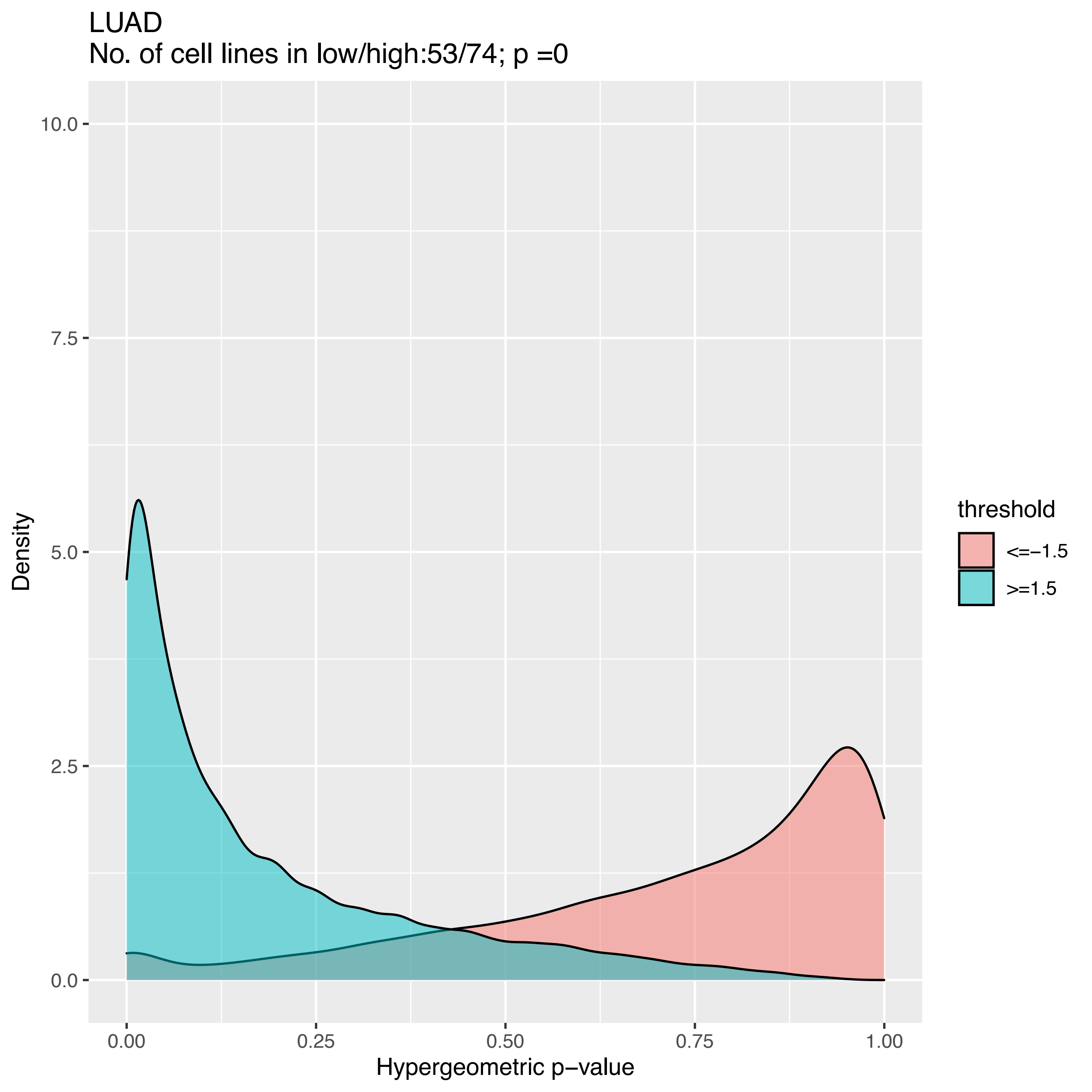
LUAD
No. of cell lines in low/high:53/74; p =0
Density
10.0
7.5
5.0
2.5
0.0
0.00
0.25
0.50
0.75
1.00
Hypergeometric p−value
threshold
<=−1.5
>=1.5

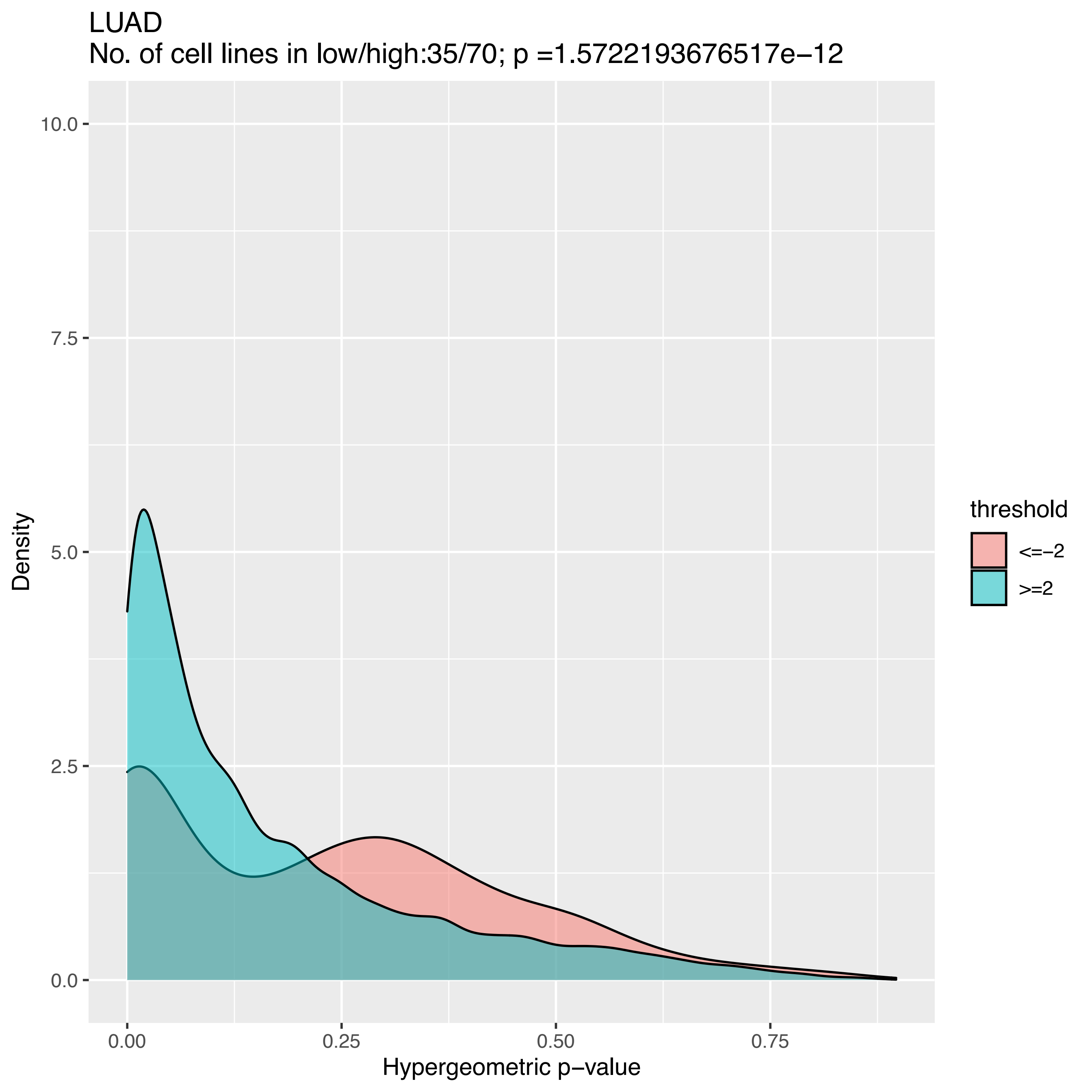
LUAD
No. of cell lines in low/high:35/70; p =1.5722193676517e−12
Density
10.0
7.5
5.0
2.5
0.0
0.00
0.25
0.50
0.75
Hypergeometric p−value
threshold
<=−2
>=2

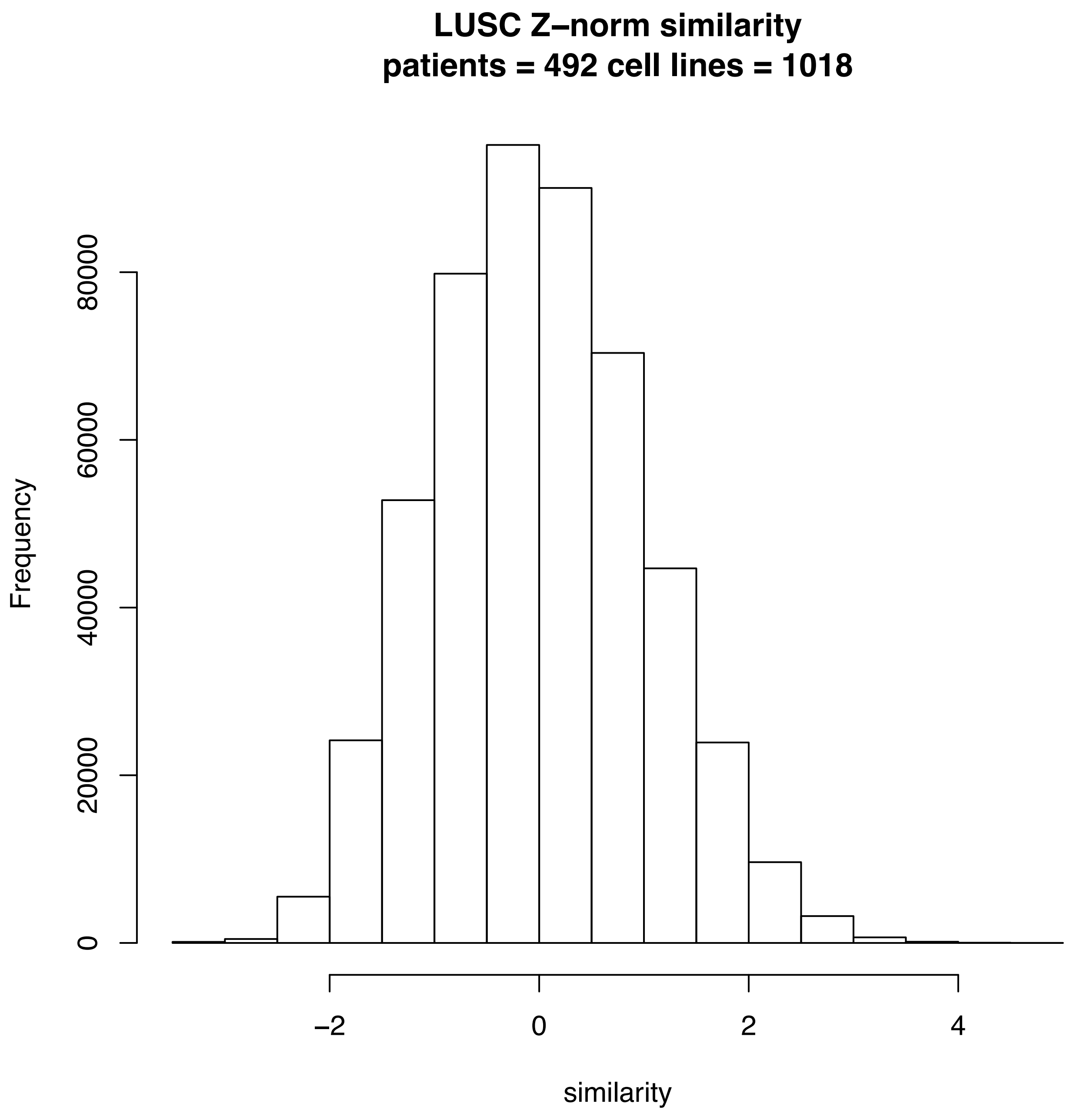

LUSC Z–norm similarity
patients = 492 cell lines = 1018
Frequency
0
20000
40000
60000
80000
–2
0
2
4
similarity

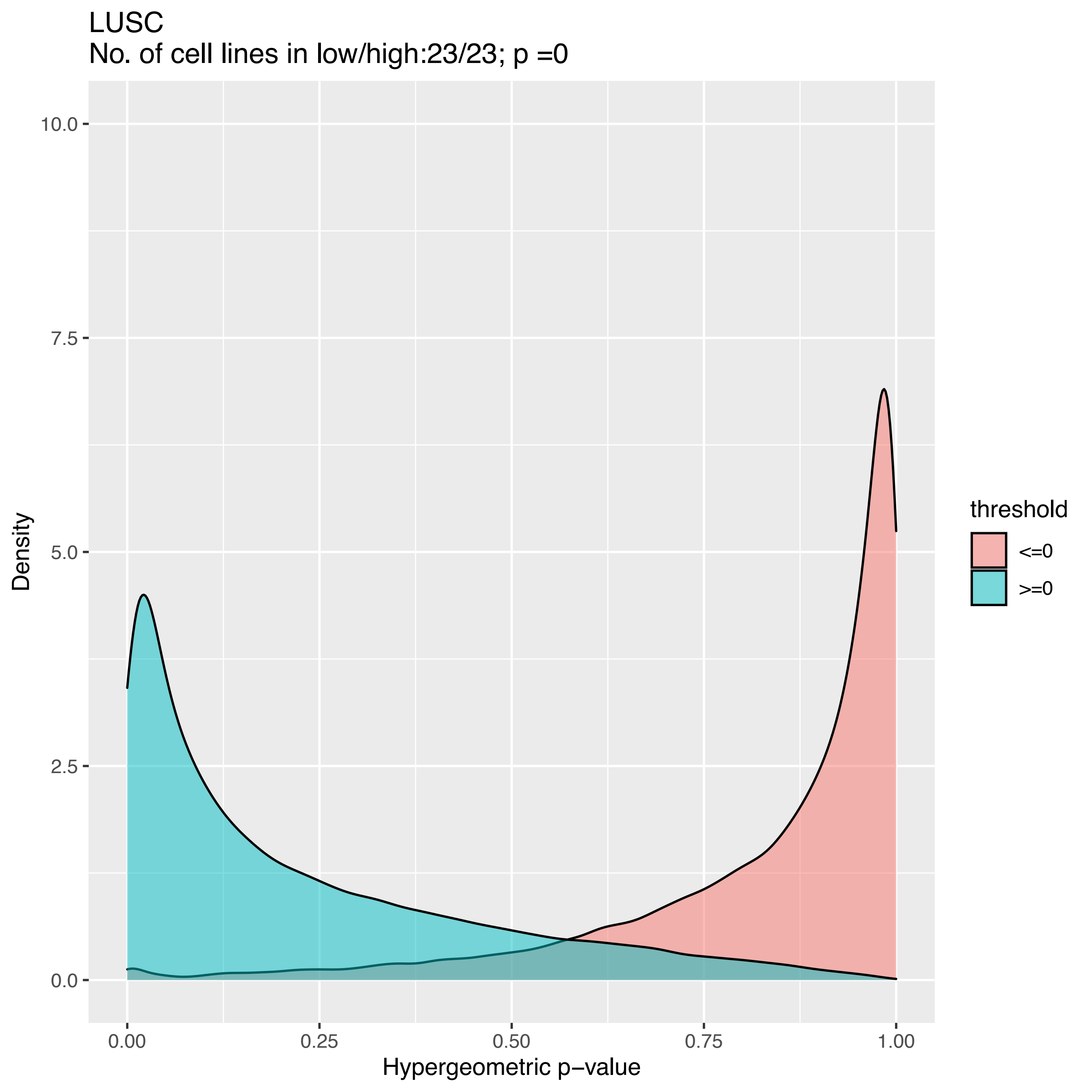
LUSC
No. of cell lines in low/high:23/23; p =0
Density
Hypergeometric p−value
0.0
2.5
5.0
7.5
10.0
0.00
0.25
0.50
0.75
1.00
threshold
<=0
>=0

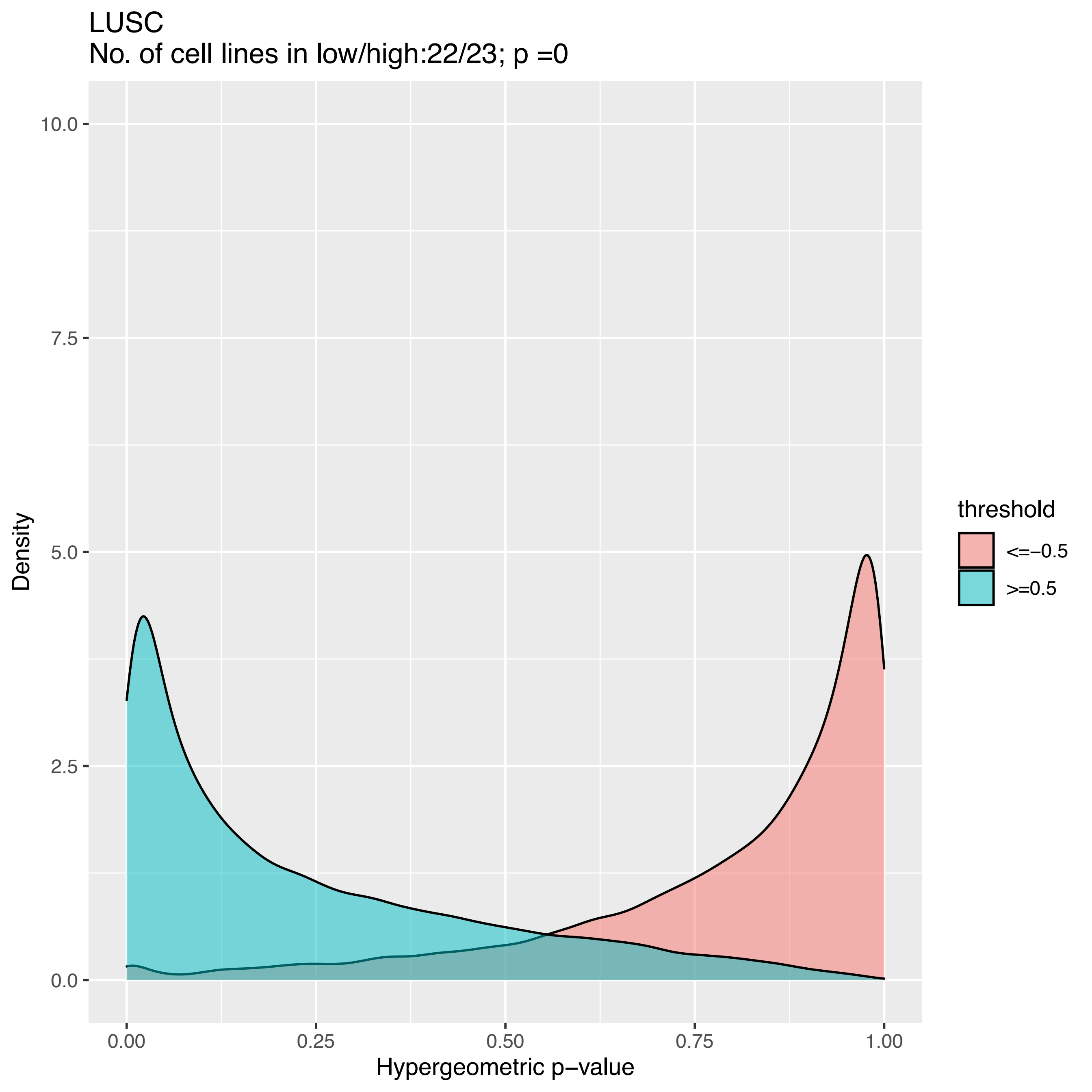
LUSC
No. of cell lines in low/high:22/23; p =0
10.0
7.5
5.0
2.5
0.0
0.00
0.25
0.50
0.75
1.00
Density
Hypergeometric p−value
threshold
<=−0.5
>=0.5

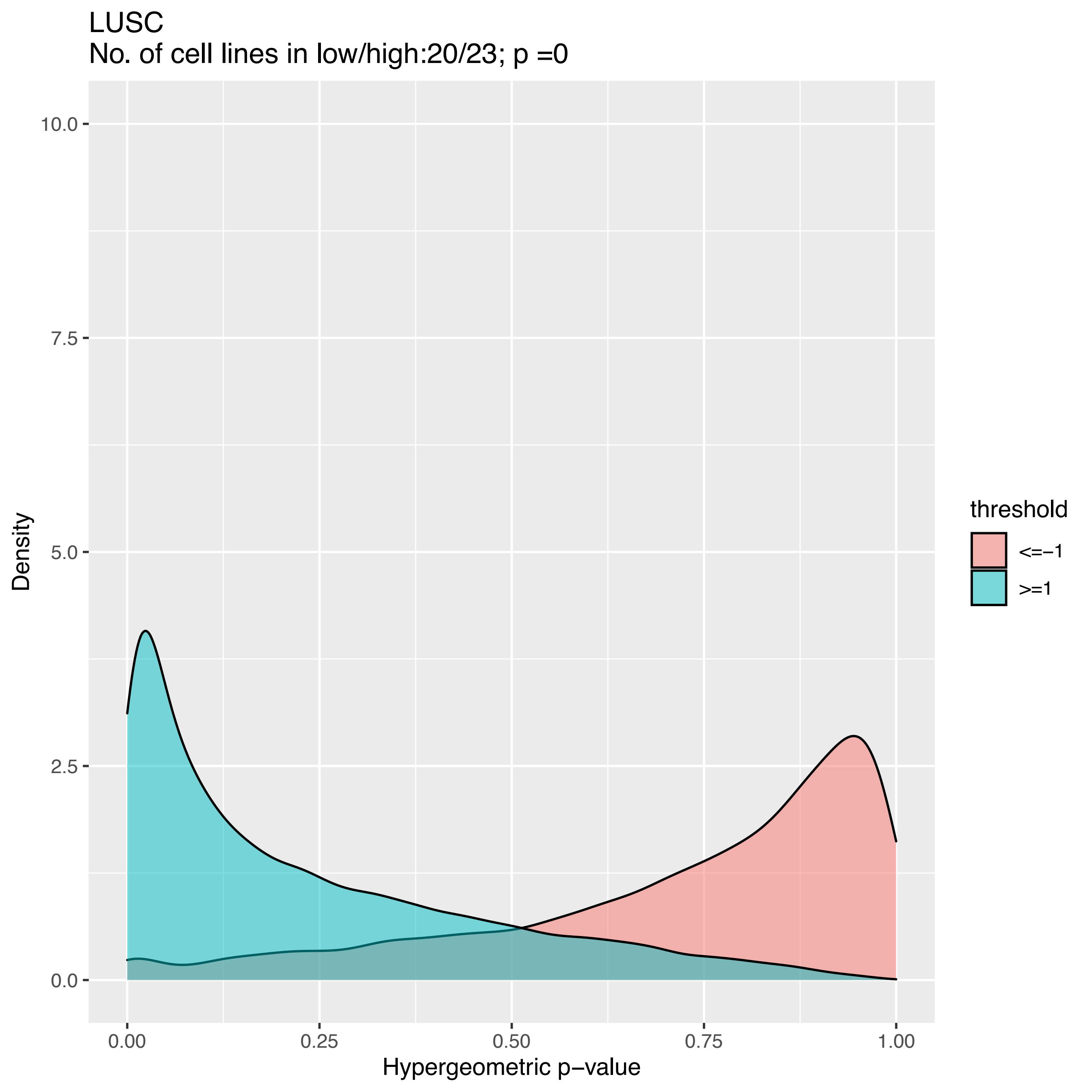
LUSC
No. of cell lines in low/high:20/23; p =0
Density
10.0
7.5
5.0
2.5
0.0
0.00
0.25
0.50
0.75
1.00
Hypergeometric p−value
threshold
<=−1
>=1

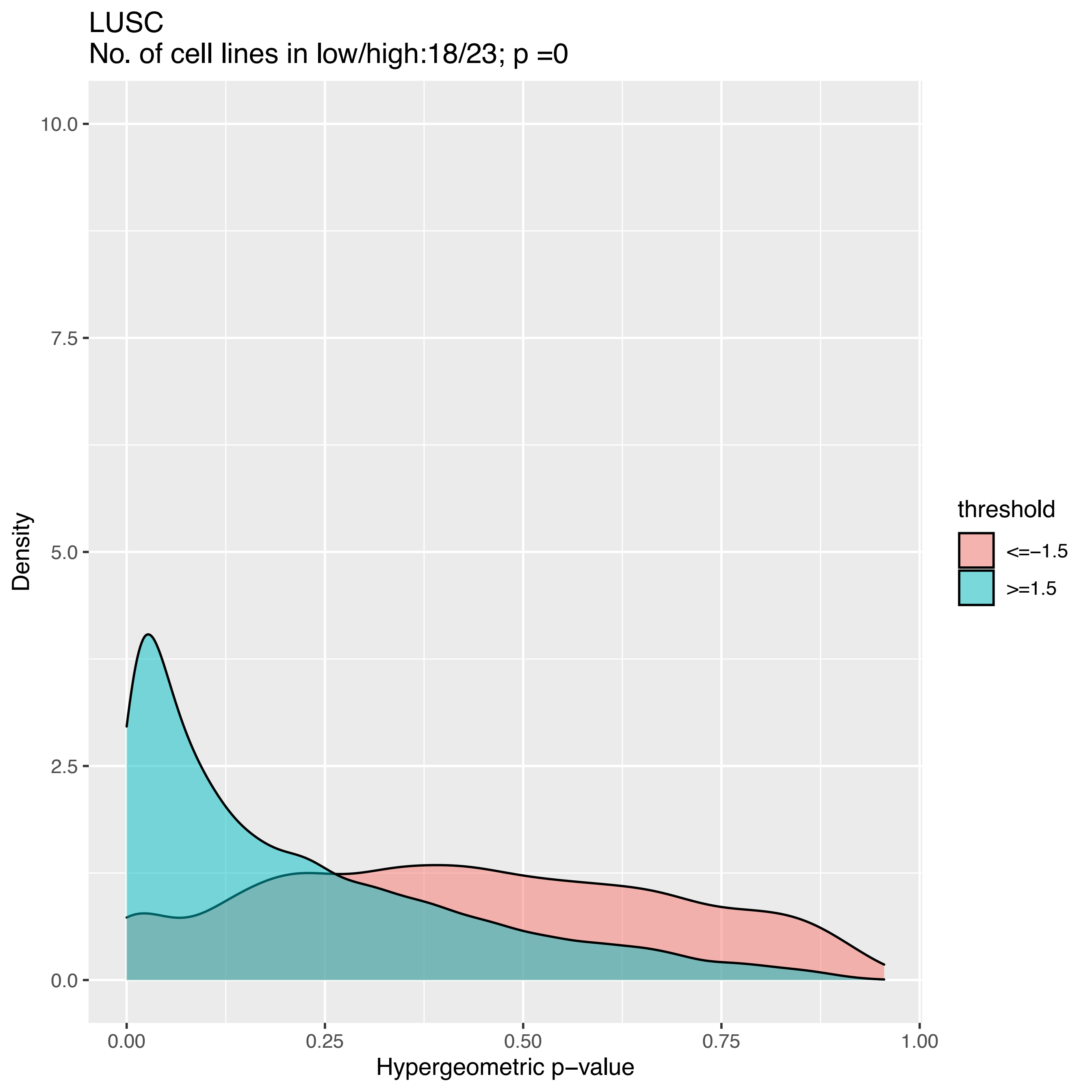
LUSC
No. of cell lines in low/high:18/23; p =0
10.0
7.5
5.0
2.5
0.0
0.00
0.25
0.50
0.75
1.00
Hypergeometric p−value
Density
threshold
<=−1.5
>=1.5

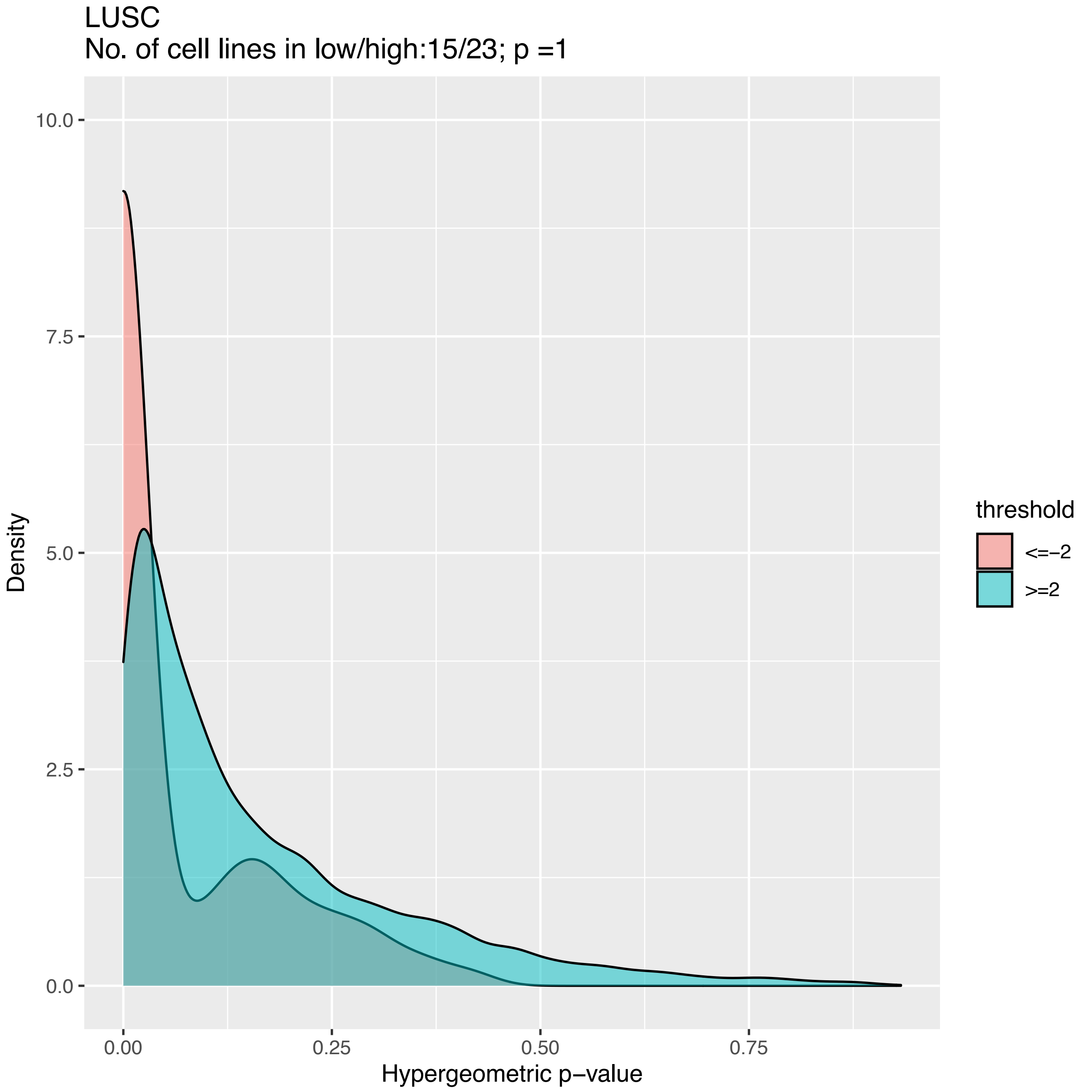

LUSC
No. of cell lines in low/high:15/23; p =1
10.0
7.5
5.0
2.5
0.0
Density
0.00
0.25
0.50
0.75
Hypergeometric p−value
threshold
<=−2
>=2

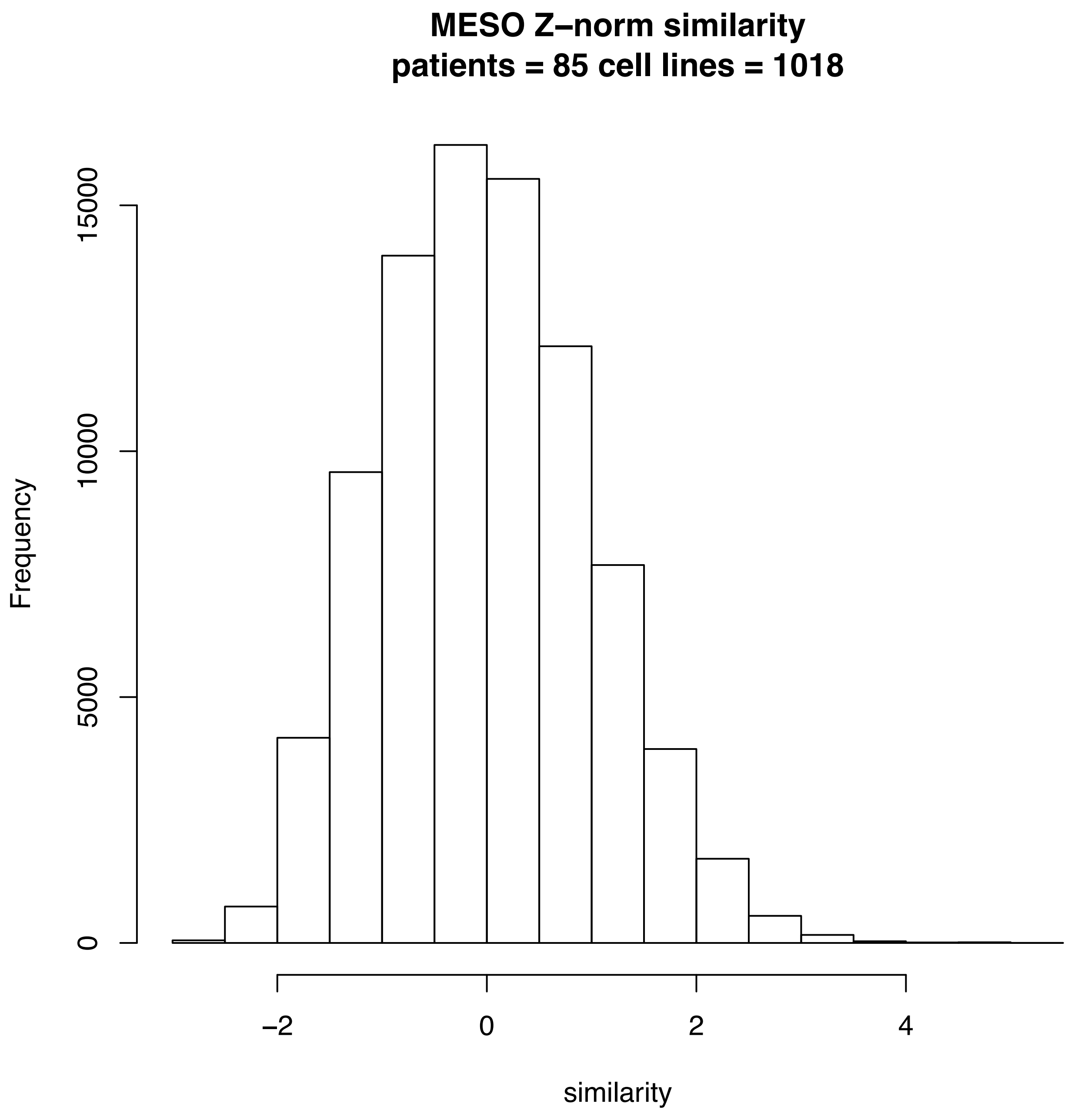
MESO Z−norm similarity
patients = 85 cell lines = 1018
Frequency
0
5000
10000
15000
−2
0
2
4
similarity

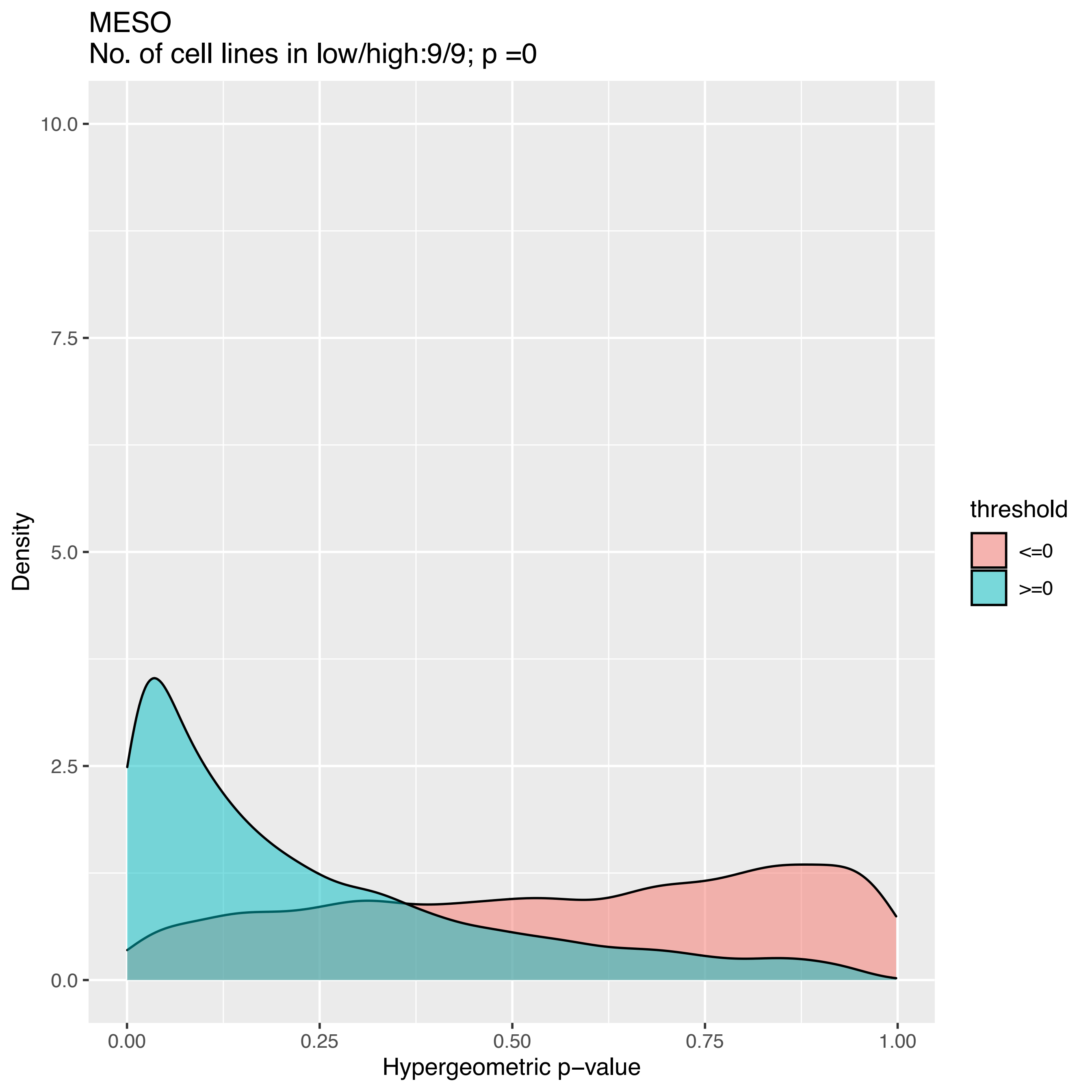

MESO
No. of cell lines in low/high:9/9; p =0
Density
10.0
7.5
5.0
2.5
0.0
0.00
0.25
0.50
0.75
1.00
Hypergeometric p−value
threshold
<=0
>=0

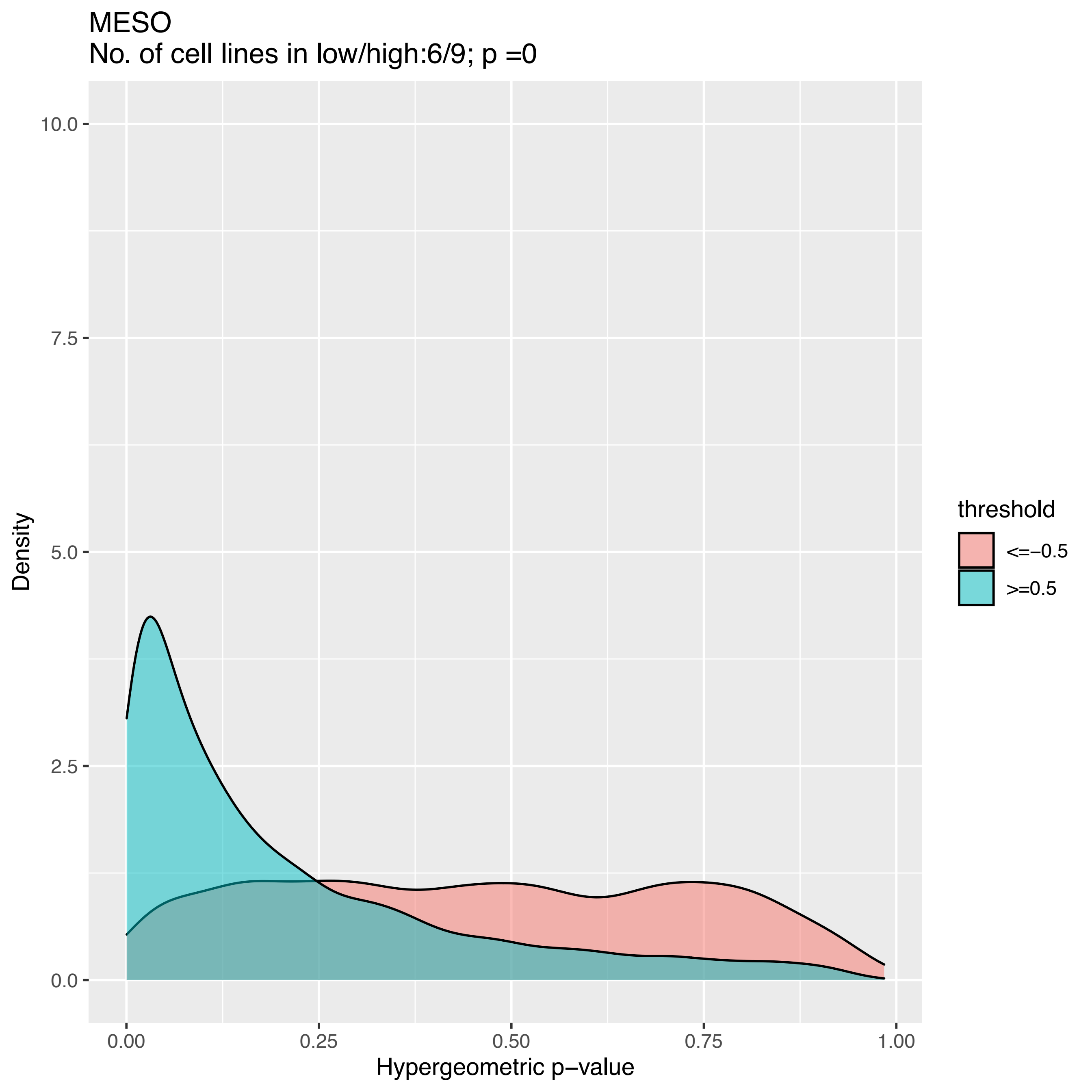
MESO
No. of cell lines in low/high:6/9; p =0
10.0
7.5
5.0
2.5
0.0
Density
0.00
0.25
0.50
0.75
1.00
Hypergeometric p−value
threshold
<=−0.5
>=0.5

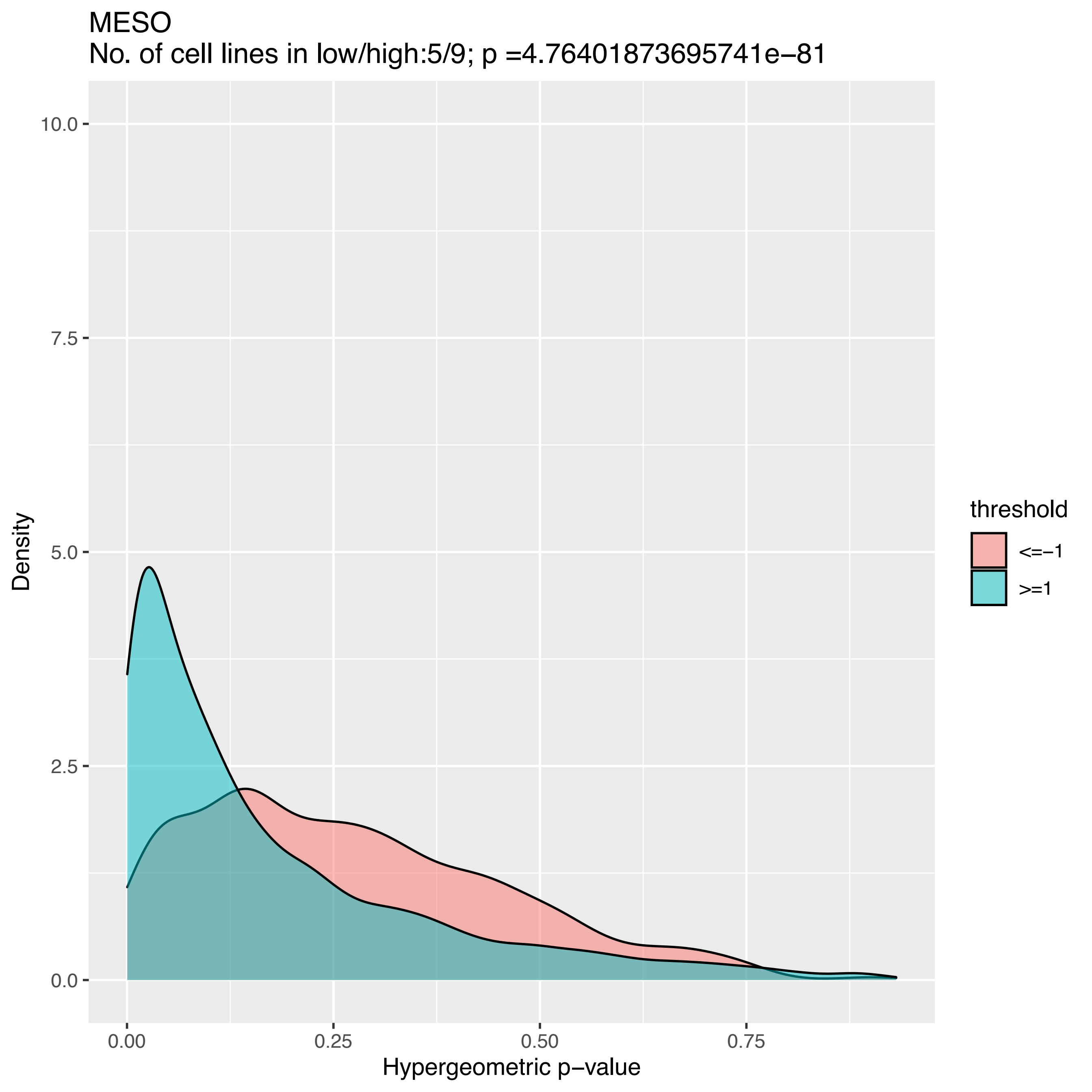

MESO
No. of cell lines in low/high:5/9; p =4.76401873695741e−81
Density
10.0
7.5
5.0
2.5
0.0
0.00
0.25
0.50
0.75
Hypergeometric p−value
threshold
<=−1
>=1

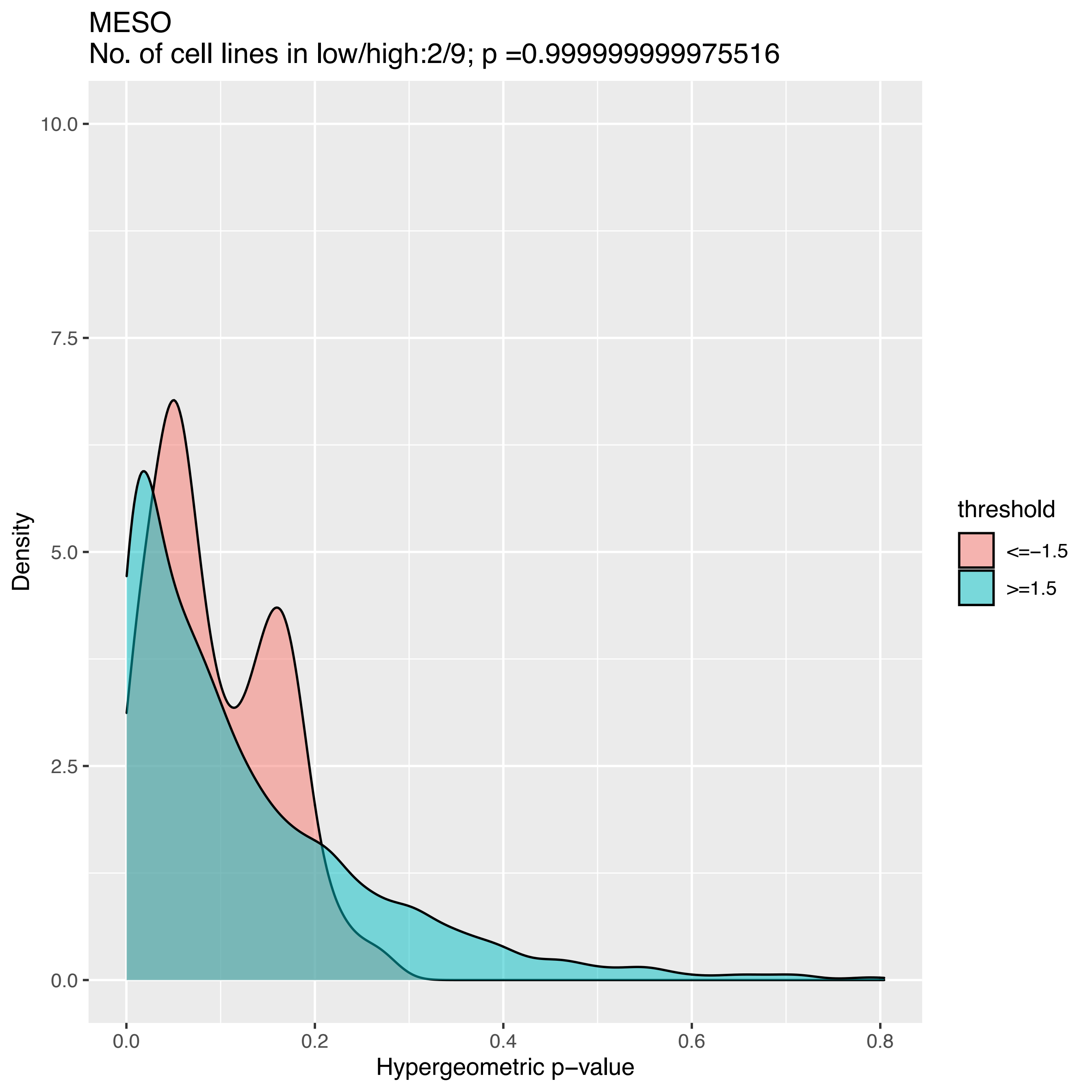
MESO
No. of cell lines in low/high:2/9; p =0.999999999975516
Density
10.0
7.5
5.0
2.5
0.0
0.0
0.2
0.4
0.6
0.8
Hypergeometric p−value
threshold
<=−1.5
>=1.5

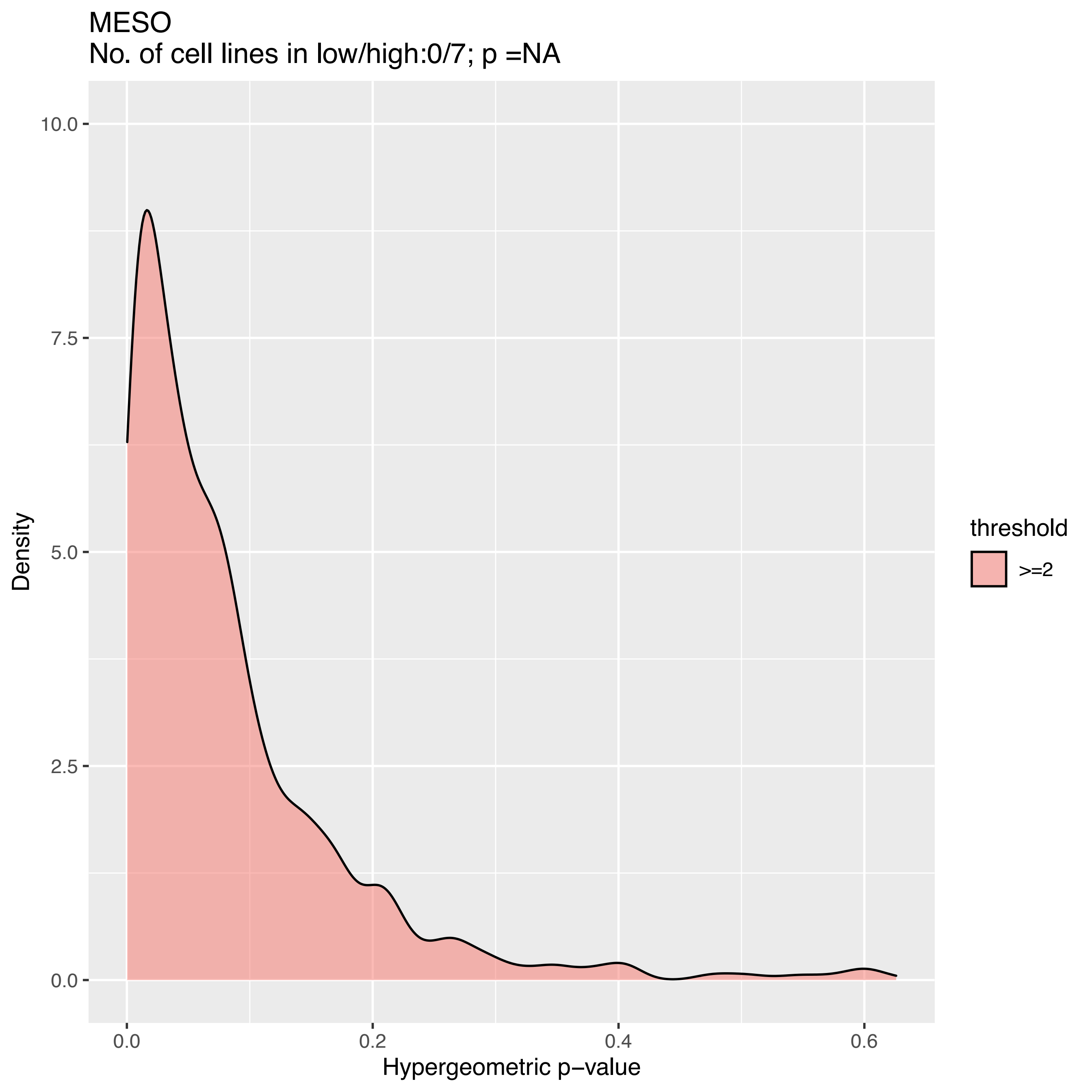
MESO
No. of cell lines in low/high:0/7; p =NA
Density
10.0
7.5
5.0
2.5
0.0
0.0
0.2
0.4
0.6
Hypergeometric p−value
threshold
>=2

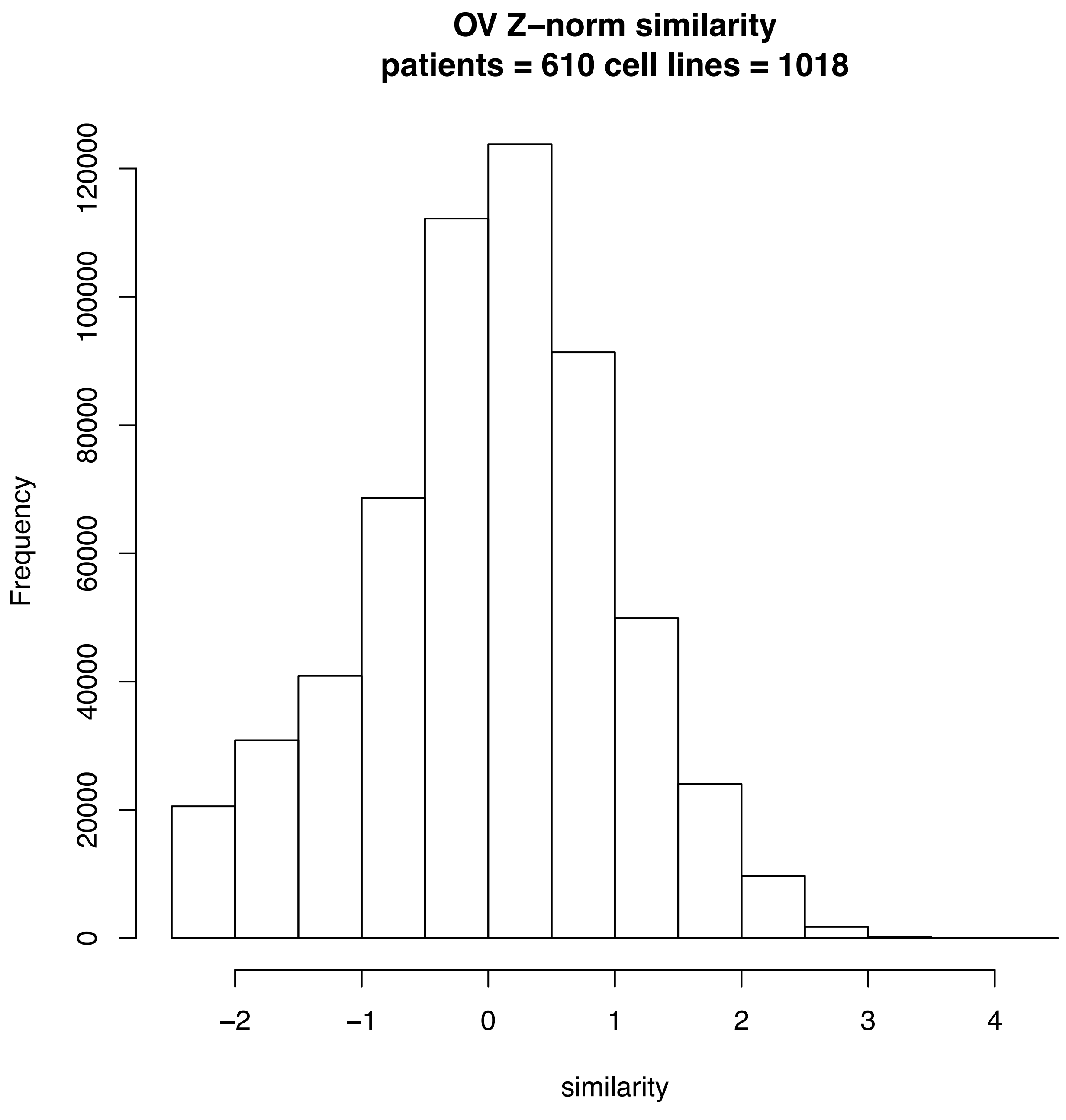
OV Z–norm similarity
patients = 610 cell lines = 1018
Frequency
0
20000
40000
60000
80000
100000
120000
−2
−1
0
1
2
3
4
similarity

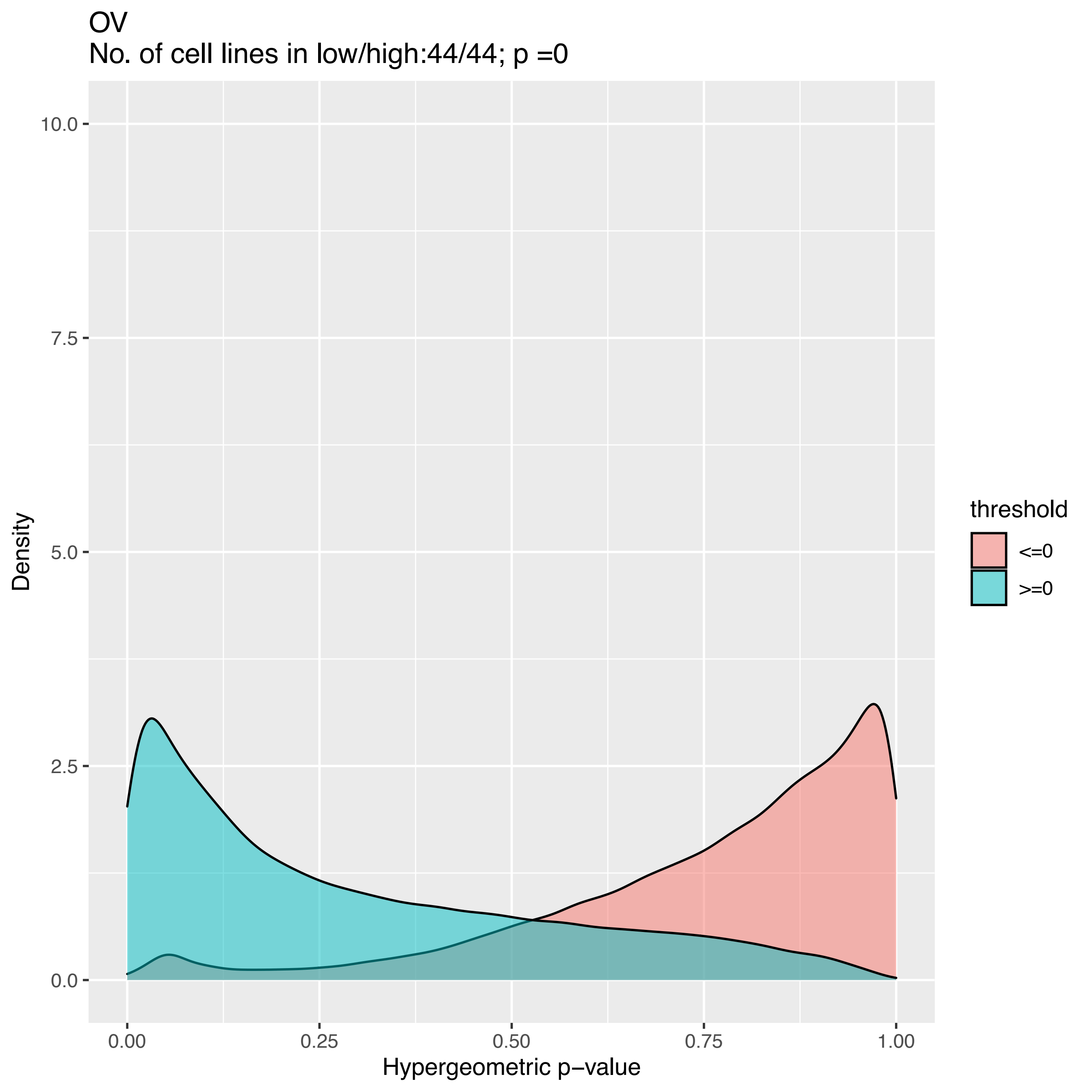
OV
No. of cell lines in low/high:44/44; p =0
Density
10.0
7.5
5.0
2.5
0.0
0.00
0.25
0.50
0.75
1.00
Hypergeometric p−value
threshold
<=0
>=0

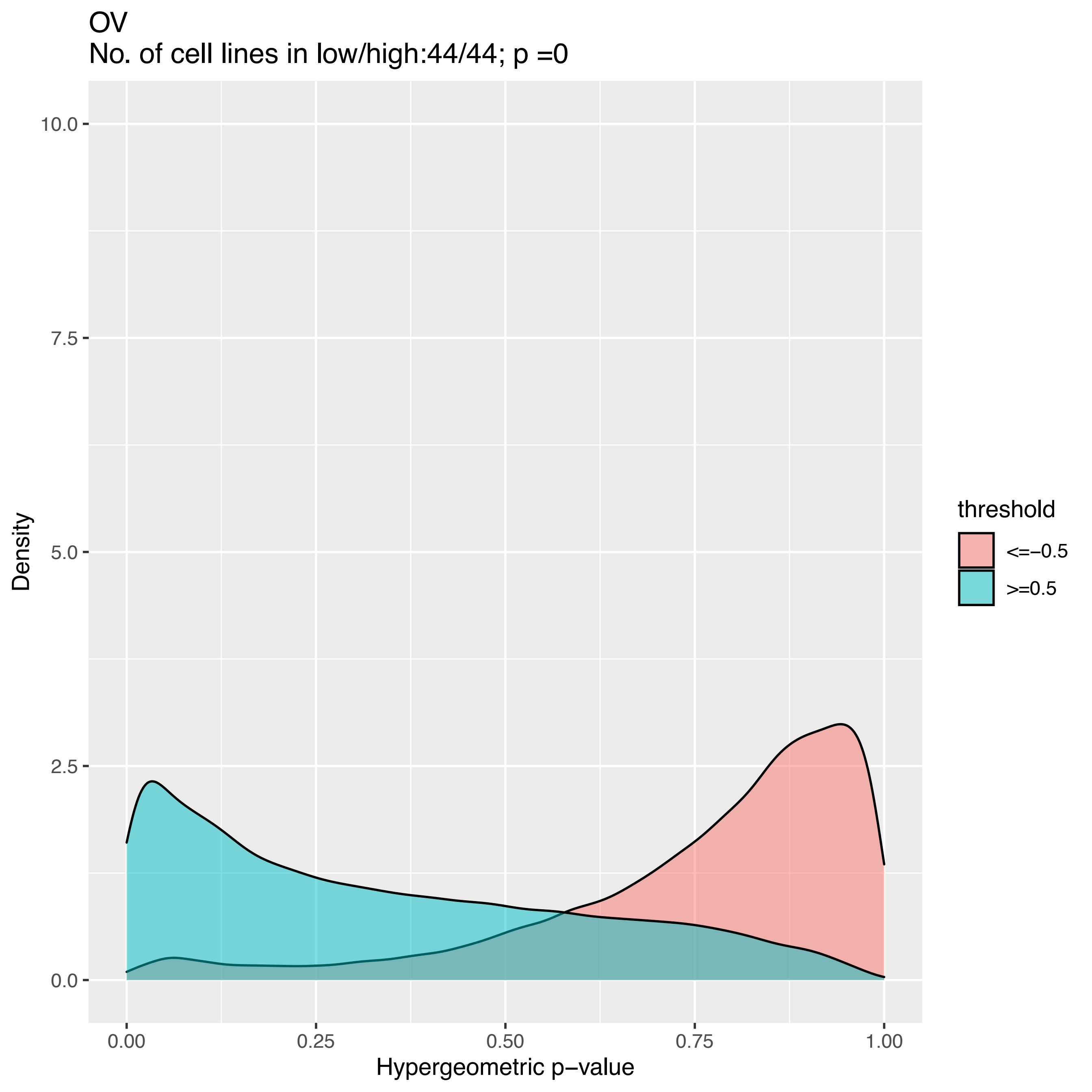

OV
No. of cell lines in low/high:44/44; p =0
10.0
7.5
5.0
2.5
0.0
Density
0.00
0.25
0.50
0.75
1.00
Hypergeometric p−value
threshold
<=−0.5
>=0.5

OV
No. of cell lines in low/high:44/44; p =0

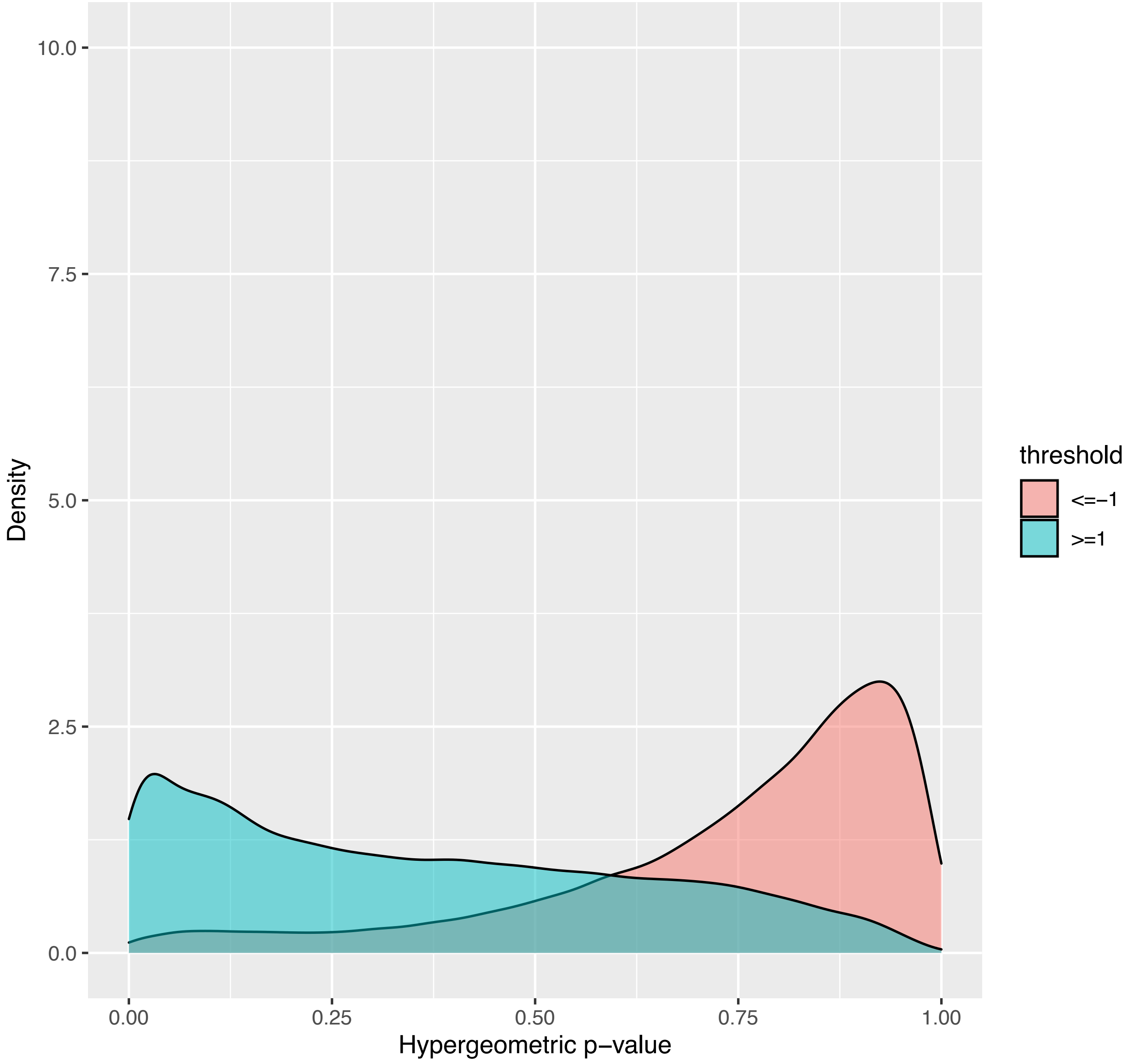

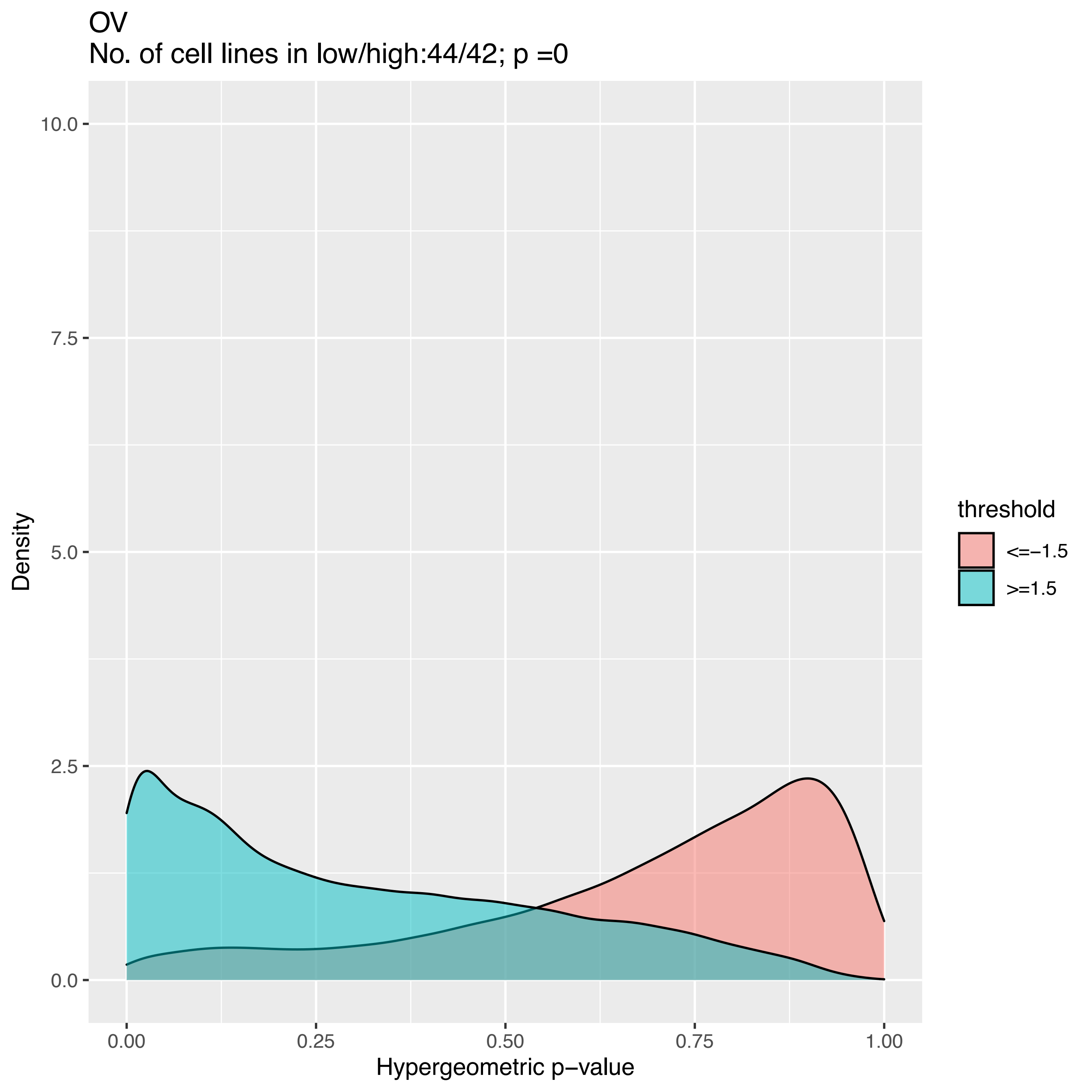
OV
No. of cell lines in low/high:44/42; p =0
Density
10.0
7.5
5.0
2.5
0.0
0.00
0.25
0.50
0.75
1.00
Hypergeometric p−value
threshold
<=−1.5
>=1.5

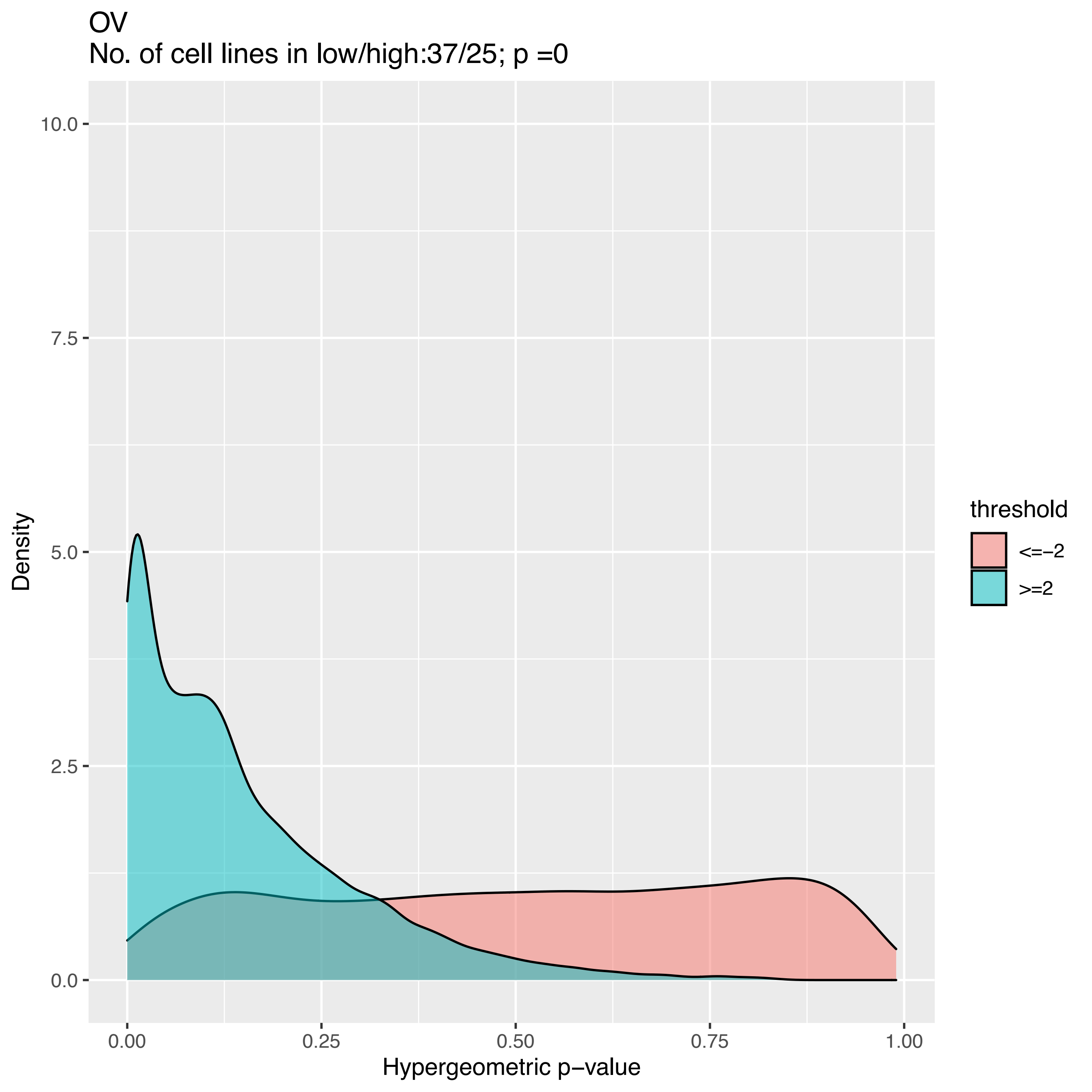

OV
No. of cell lines in low/high:37/25; p =0
Density
10.0
7.5
5.0
2.5
0.0
0.00
0.25
0.50
0.75
1.00
Hypergeometric p−value
threshold
<=−2
>=2

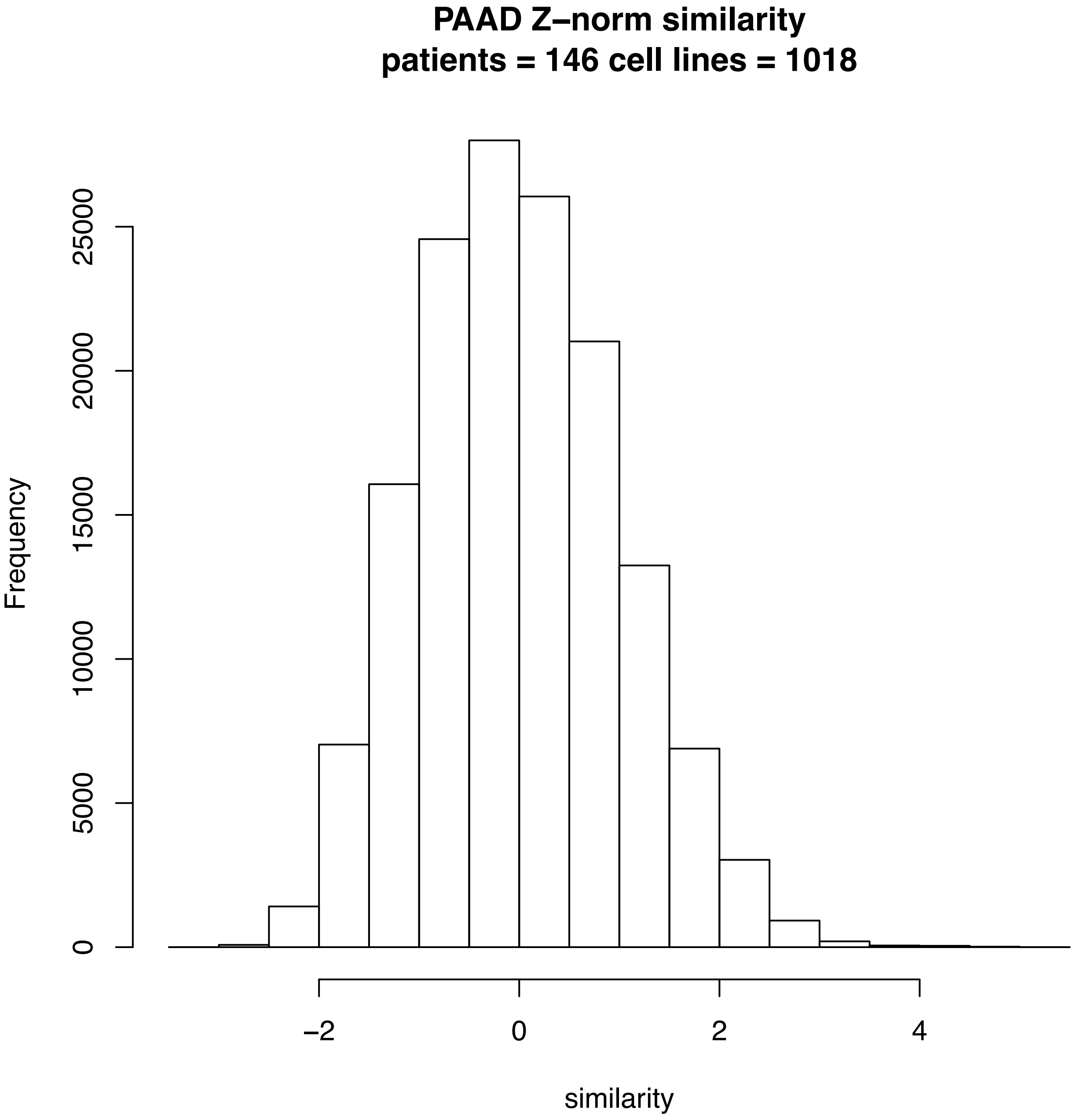
PAAD Z–norm similarity
patients = 146 cell lines = 1018
Frequency
0
5000
10000
15000
20000
25000
–2
0
2
4
similarity

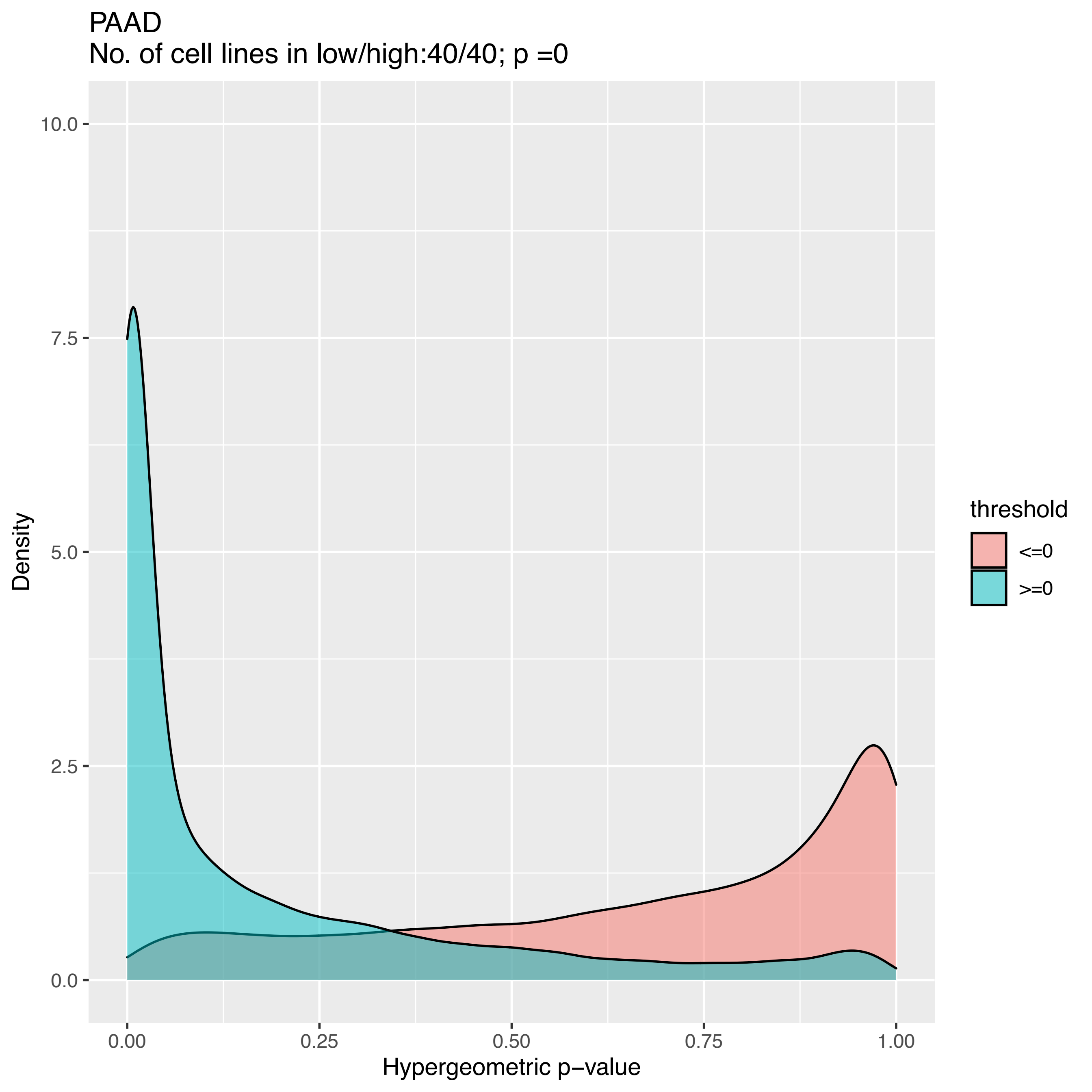
PAAD
No. of cell lines in low/high:40/40; p =0
Density
10.0
7.5
5.0
2.5
0.0
0.00
0.25
0.50
0.75
1.00
Hypergeometric p−value
threshold
<=0
>=0

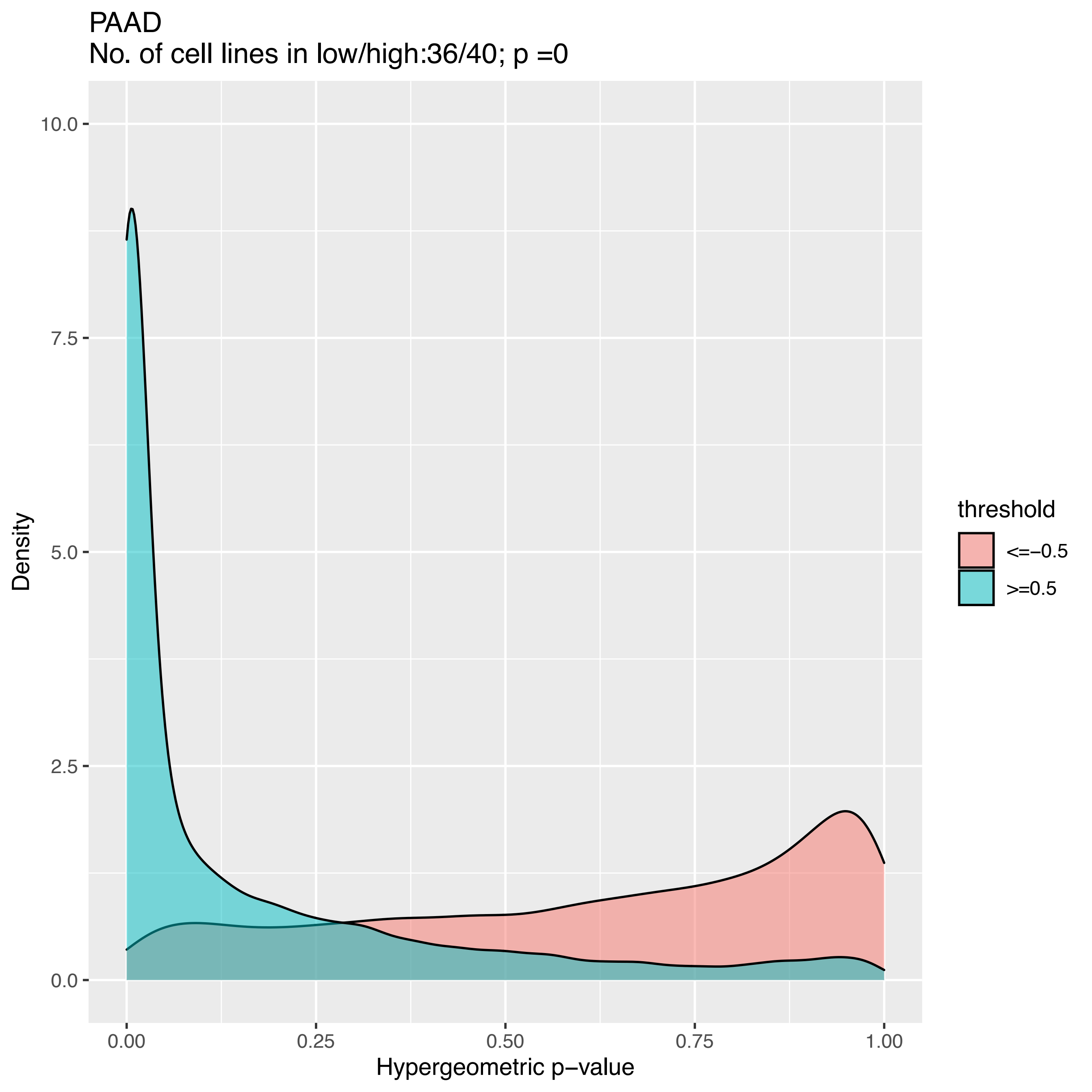

PAAD
No. of cell lines in low/high:36/40; p =0
10.0
7.5
5.0
2.5
0.0
Density
0.00
0.25
0.50
0.75
1.00
Hypergeometric p−value
threshold
<=−0.5
>=0.5

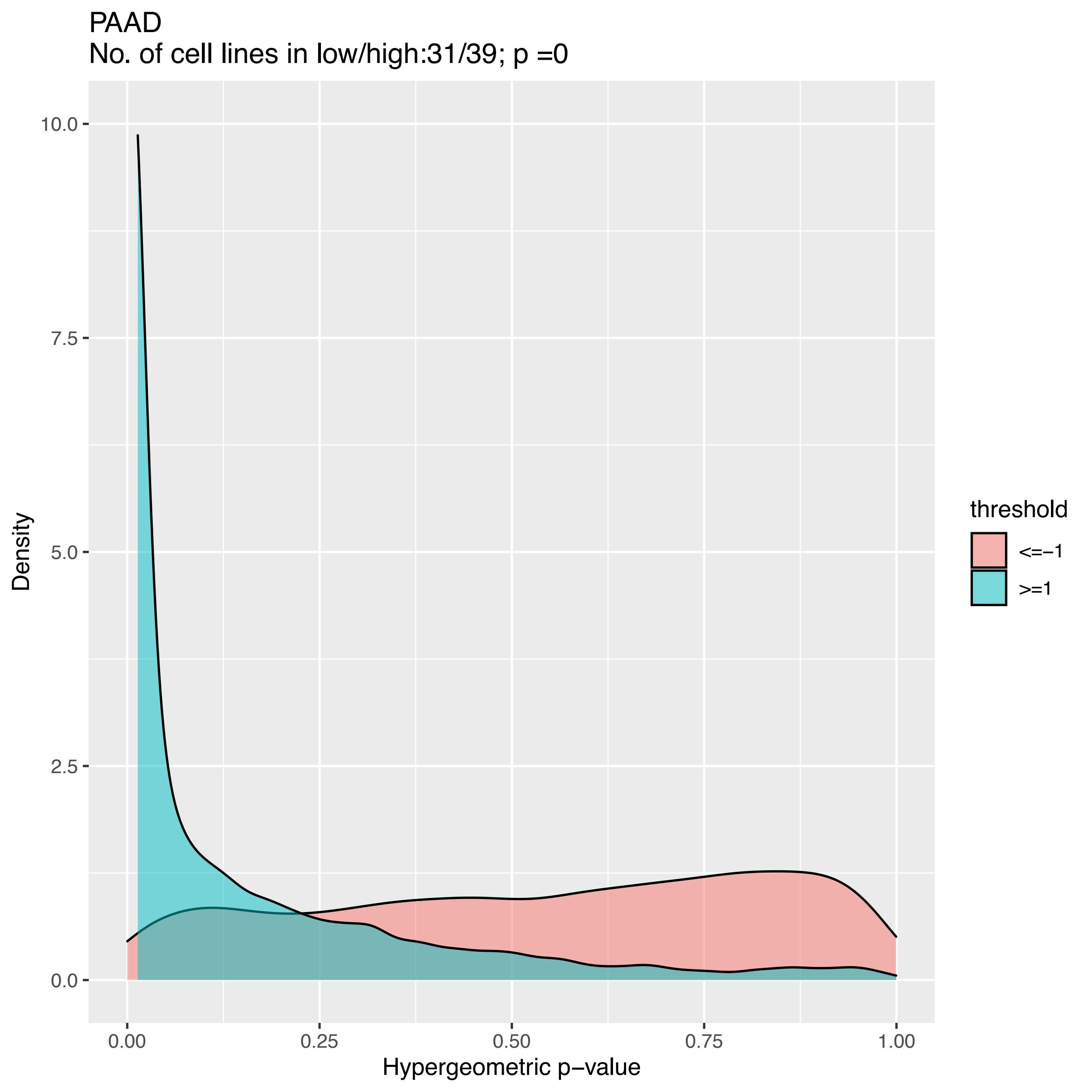
PAAD
No. of cell lines in low/high:31/39; p =0
Density
10.0
7.5
5.0
2.5
0.0
0.00
0.25
0.50
0.75
1.00
Hypergeometric p−value
threshold
<=−1
>=1

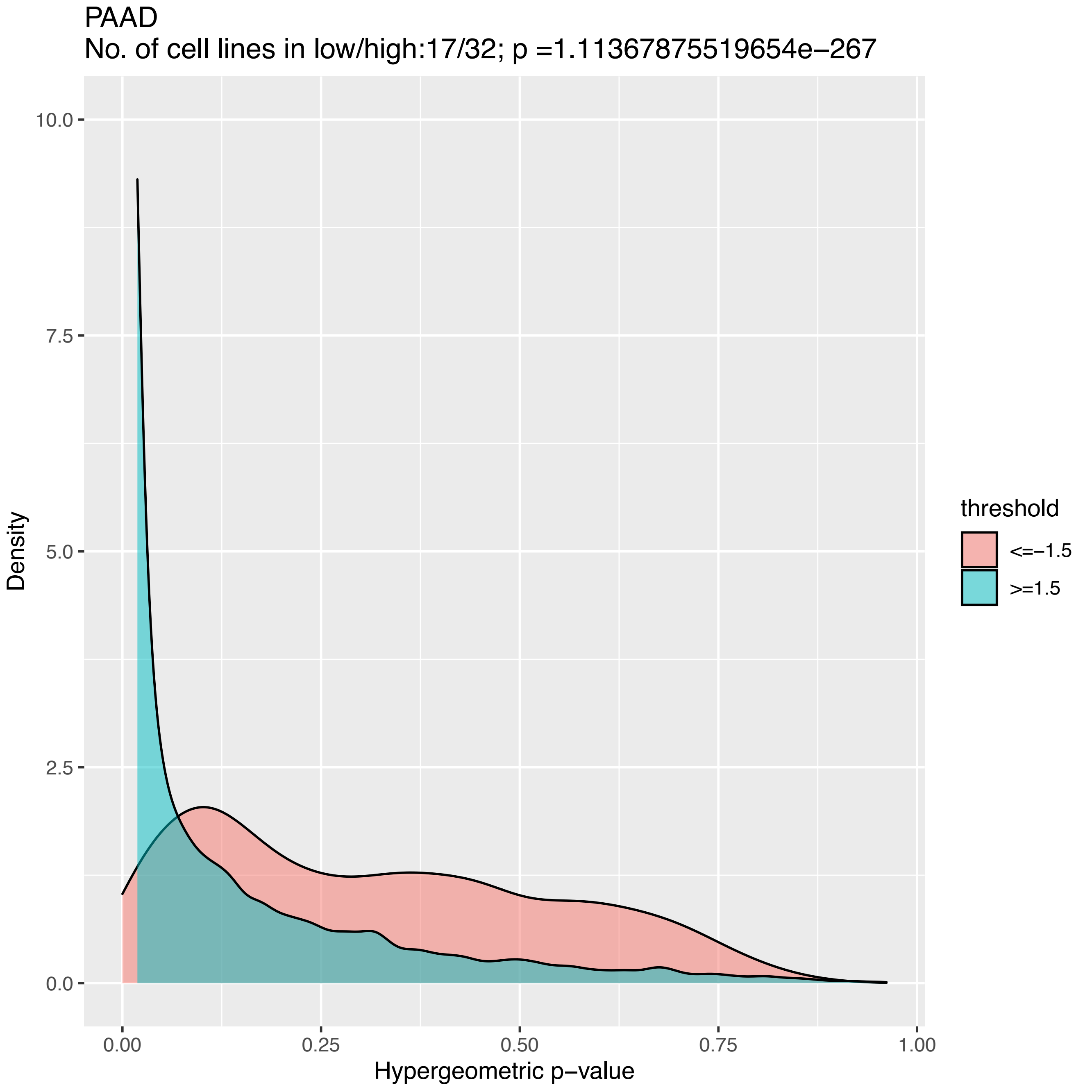
PAAD
No. of cell lines in low/high:17/32; p =1.11367875519654e−267
Density
10.0
7.5
5.0
2.5
0.0
0.00
0.25
0.50
0.75
1.00
Hypergeometric p−value
threshold
<=−1.5
>=1.5

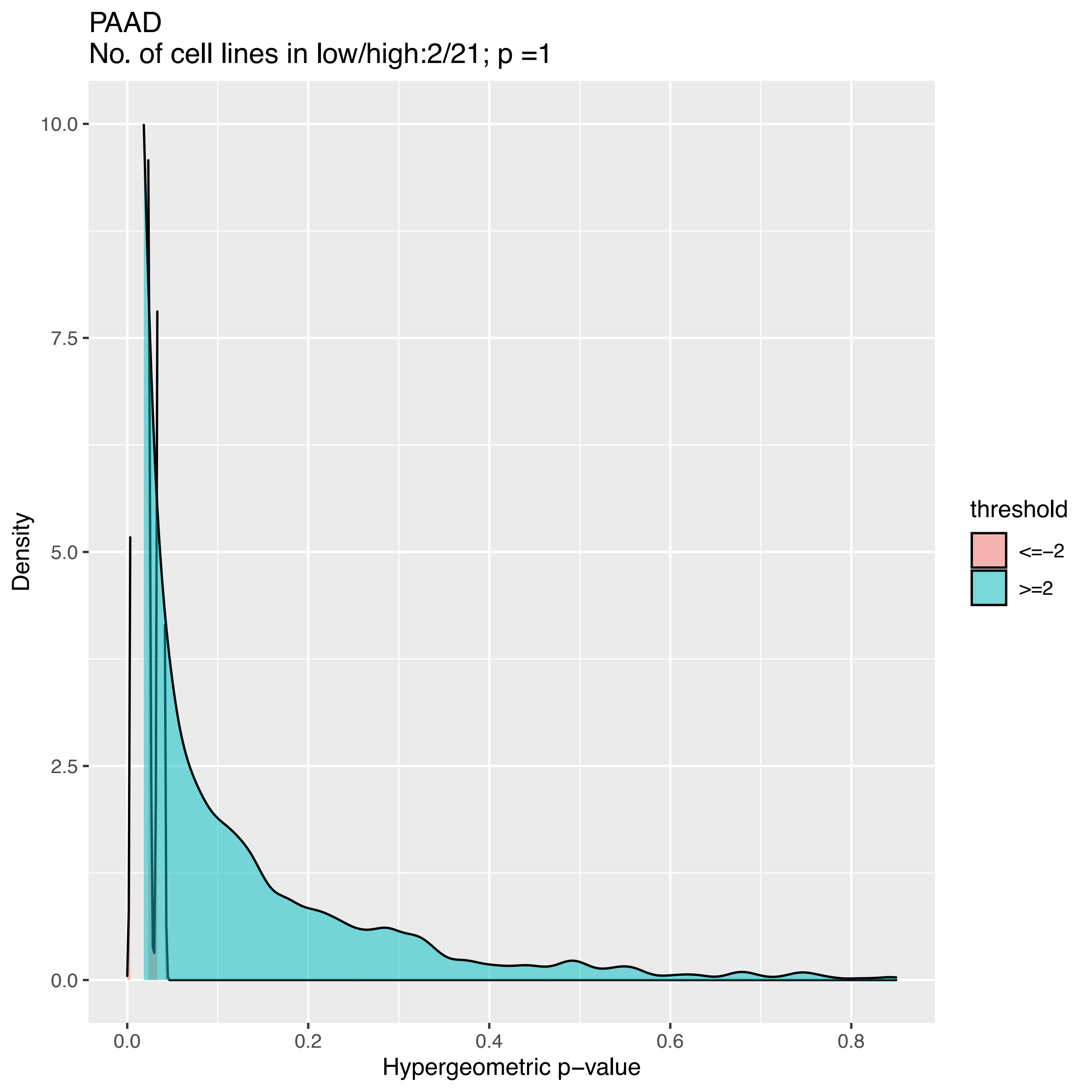
PAAD
No. of cell lines in low/high:2/21; p =1
Density
10.0
7.5
5.0
2.5
0.0
0.0
0.2
0.4
0.6
0.8
Hypergeometric p−value
threshold
<=−2
>=2

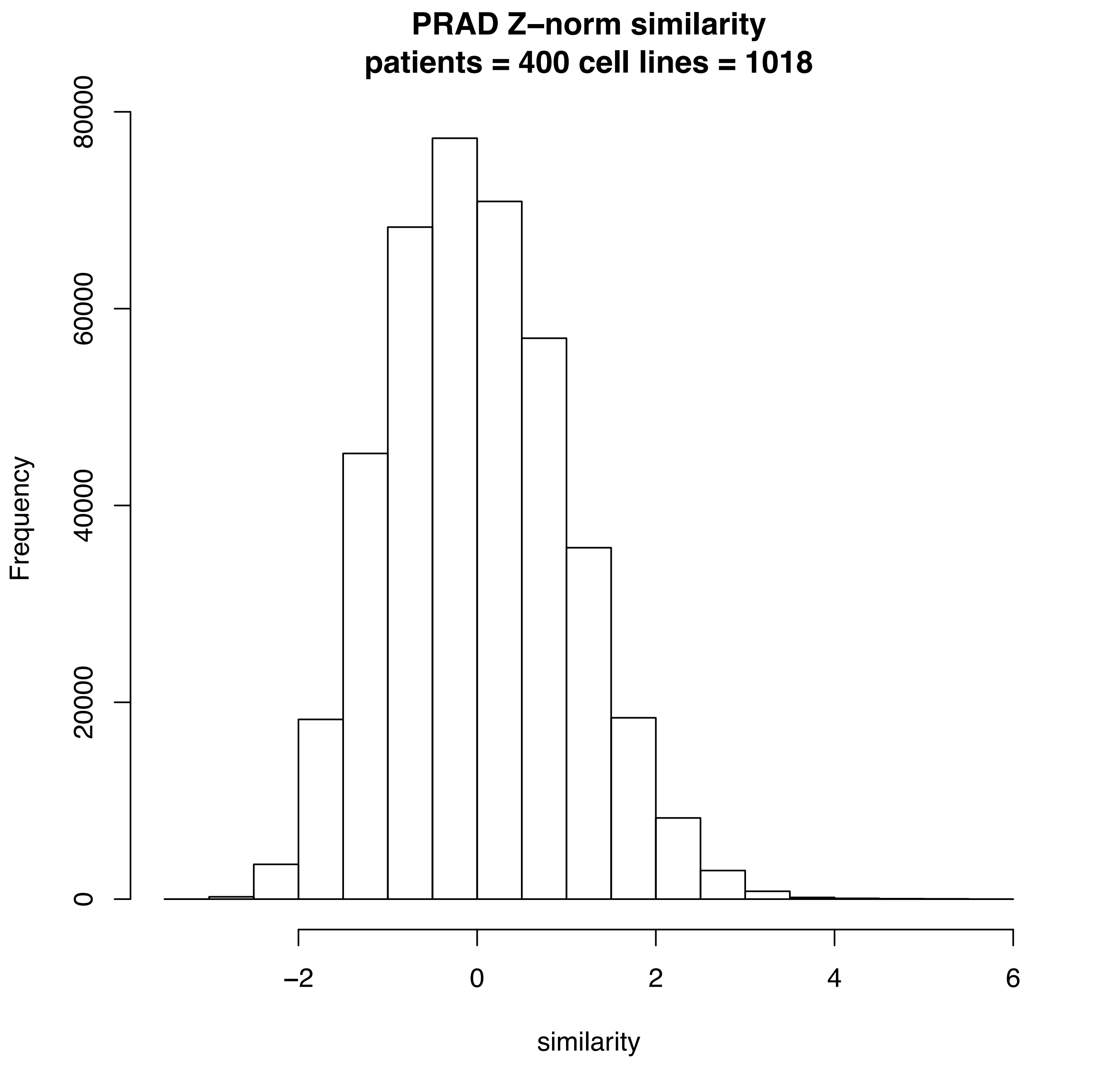
PRAD Z–norm similarity
patients = 400 cell lines = 1018
Frequency
0
20000
40000
60000
80000
–2
0
2
4
6
similarity

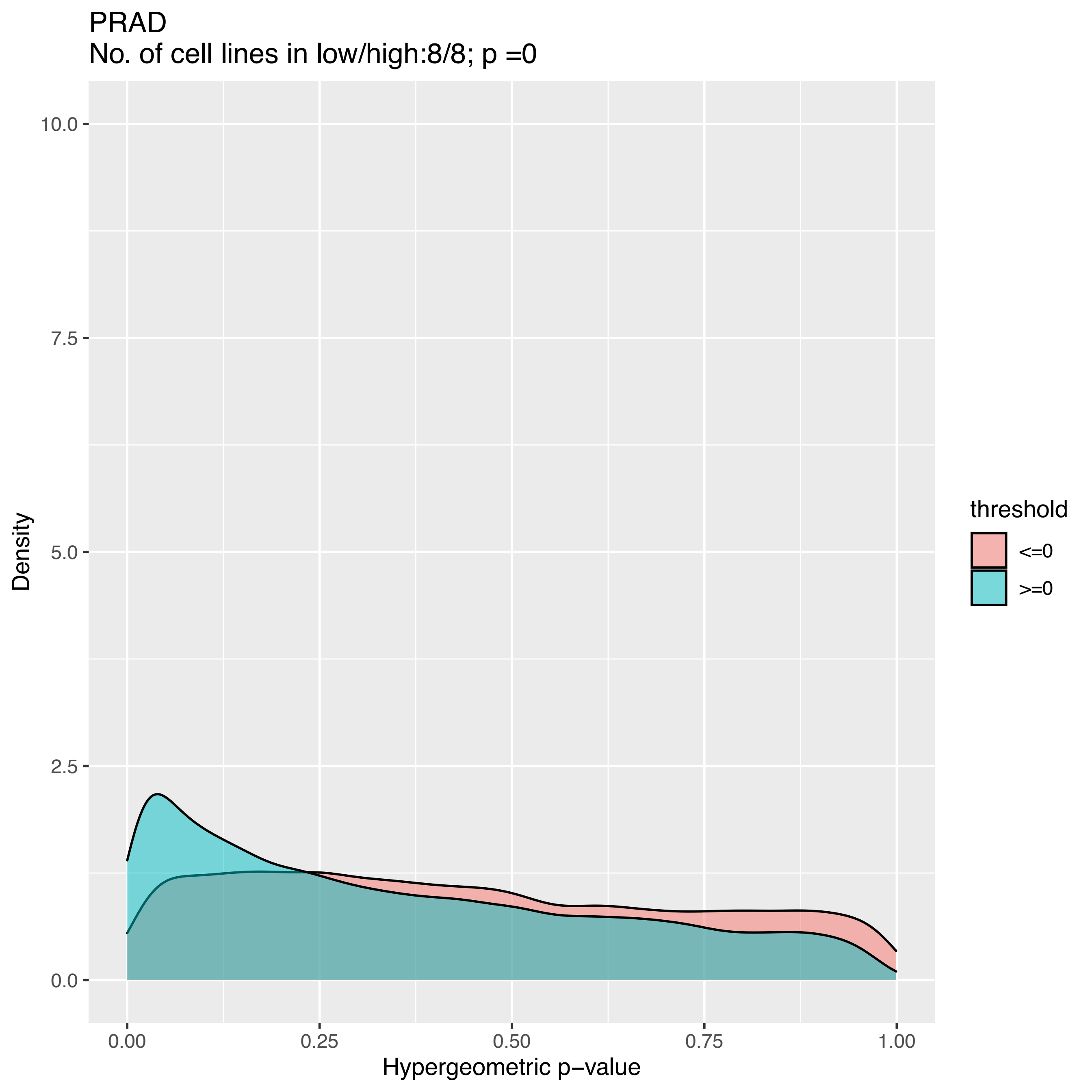

PRAD
No. of cell lines in low/high:8/8; p =0
Density
10.0
7.5
5.0
2.5
0.0
0.00
0.25
0.50
0.75
1.00
Hypergeometric p−value
threshold
<=0
>=0

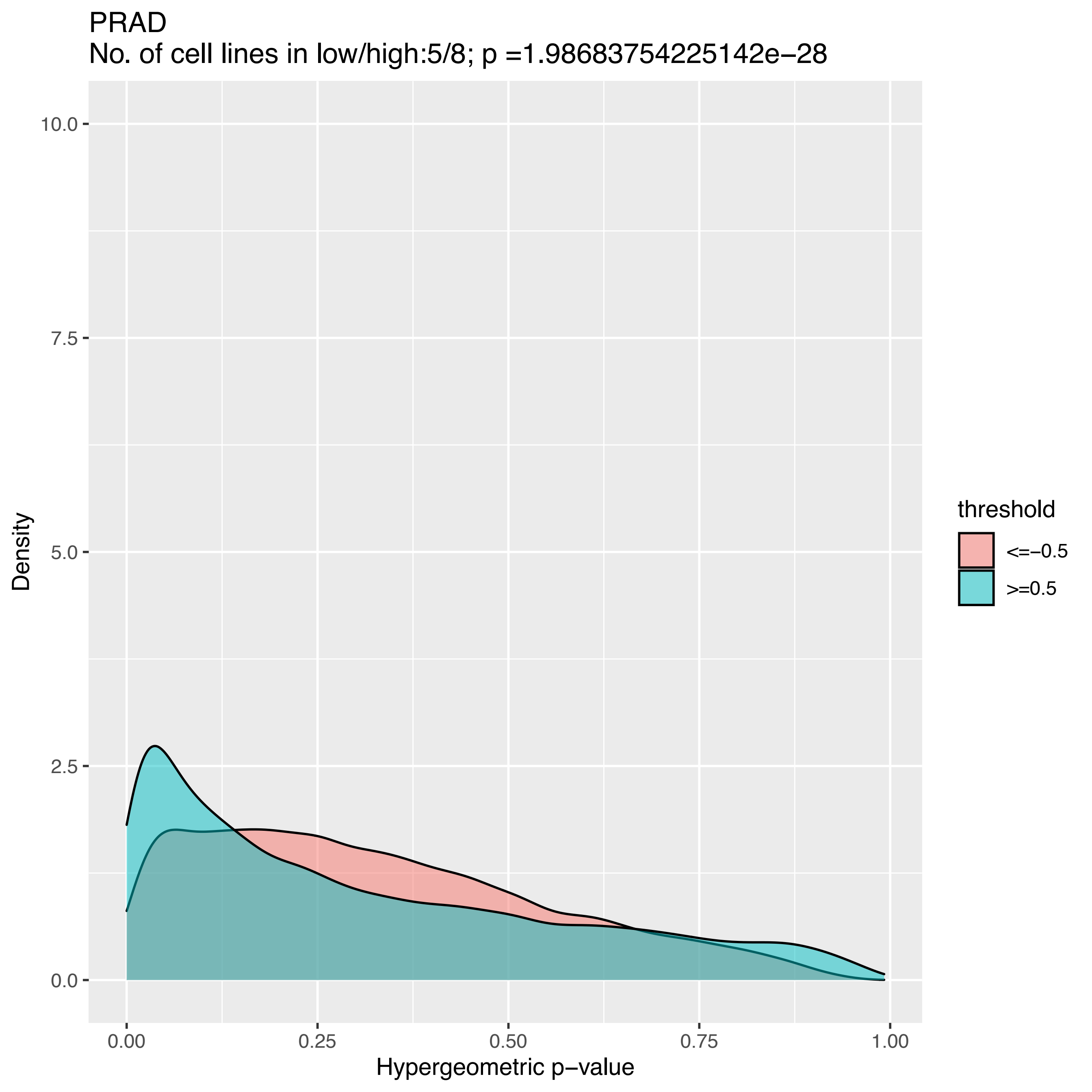
PRAD
No. of cell lines in low/high:5/8; p =1.98683754225142e−28
Density
Hypergeometric p−value
0.0
2.5
5.0
7.5
10.0
0.00
0.25
0.50
0.75
1.00
threshold
<=−0.5
>=0.5

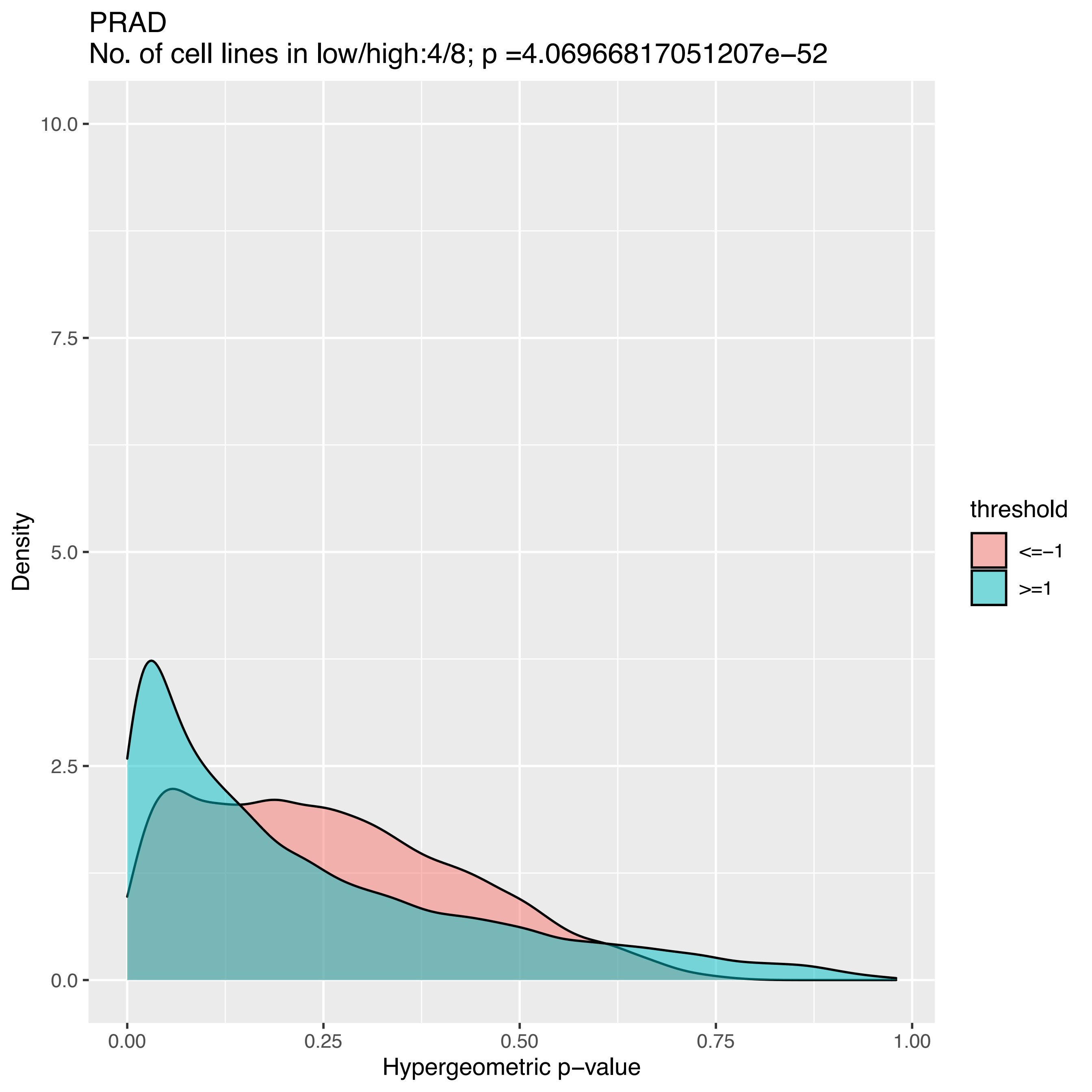
PRAD
No. of cell lines in low/high:4/8; p =4.06966817051207e−52
Density
10.0
7.5
5.0
2.5
0.0
0.00
0.25
0.50
0.75
1.00
Hypergeometric p−value
threshold
<=−1
>=1

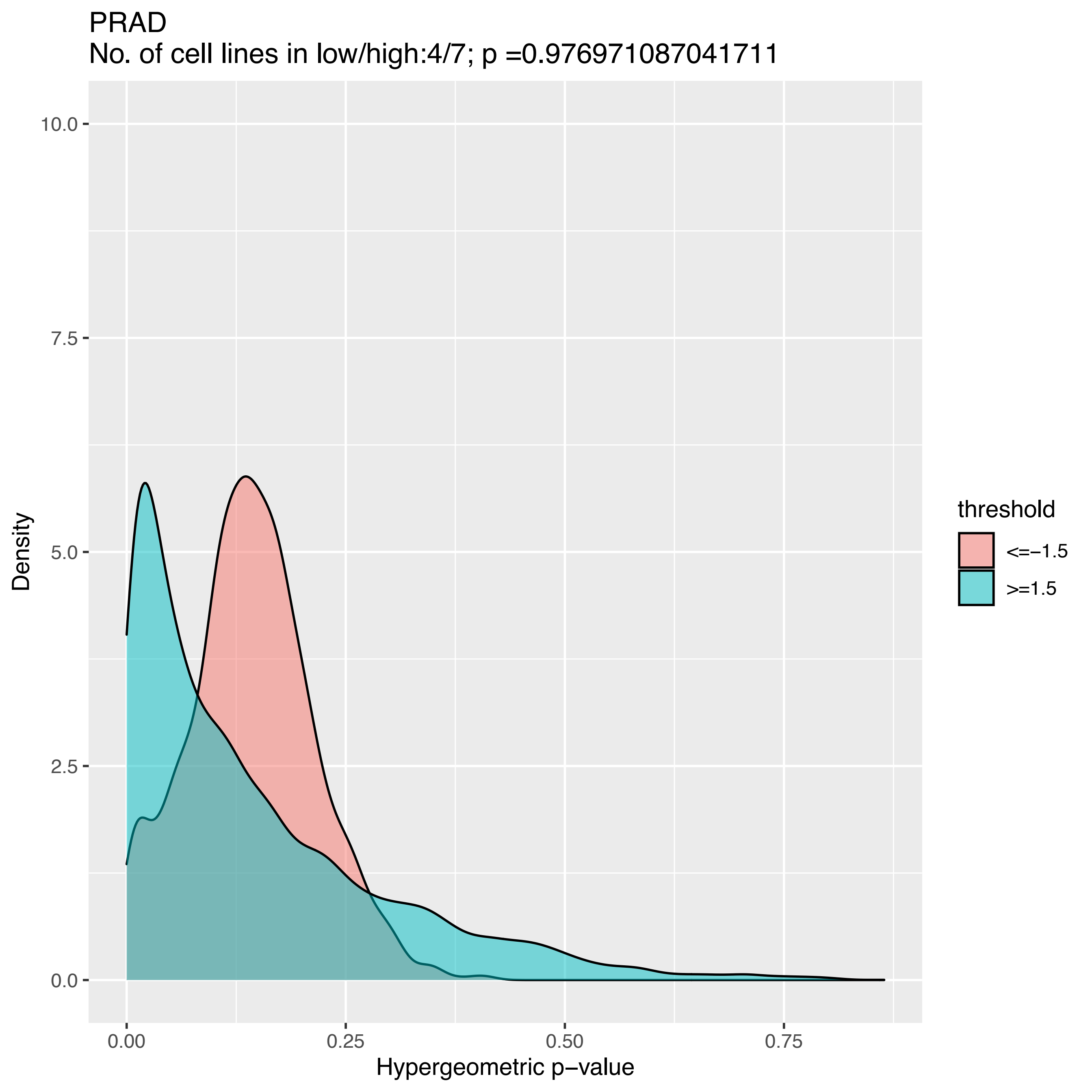
PRAD
No. of cell lines in low/high:4/7; p =0.976971087041711
10.0
7.5
5.0
2.5
0.0
Density
0.00
0.25
0.50
0.75
Hypergeometric p−value
threshold
<=−1.5
>=1.5

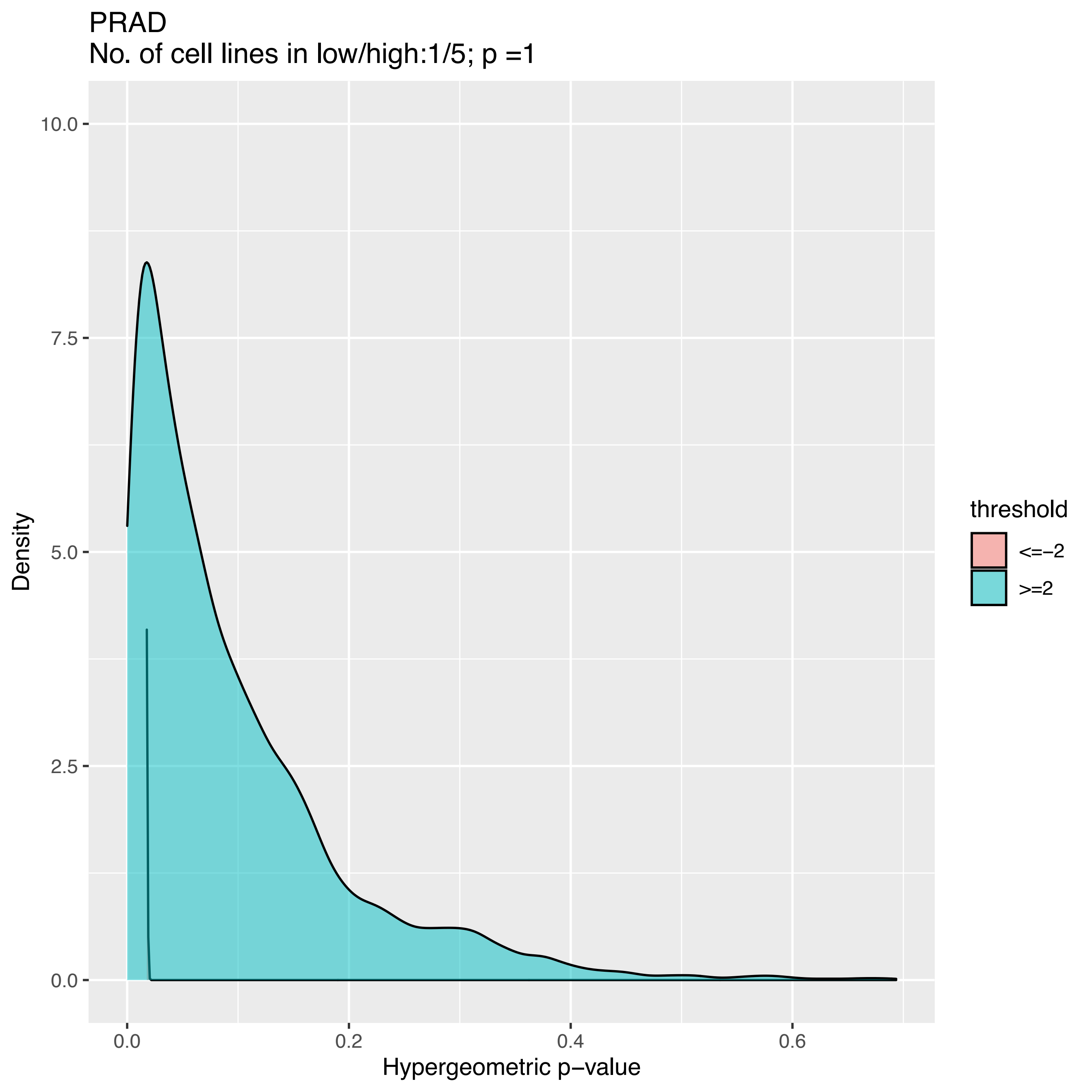
PRAD
No. of cell lines in low/high:1/5; p =1
Density
10.0
7.5
5.0
2.5
0.0
0.0
0.2
0.4
0.6
Hypergeometric p−value
threshold
<=−2
>=2

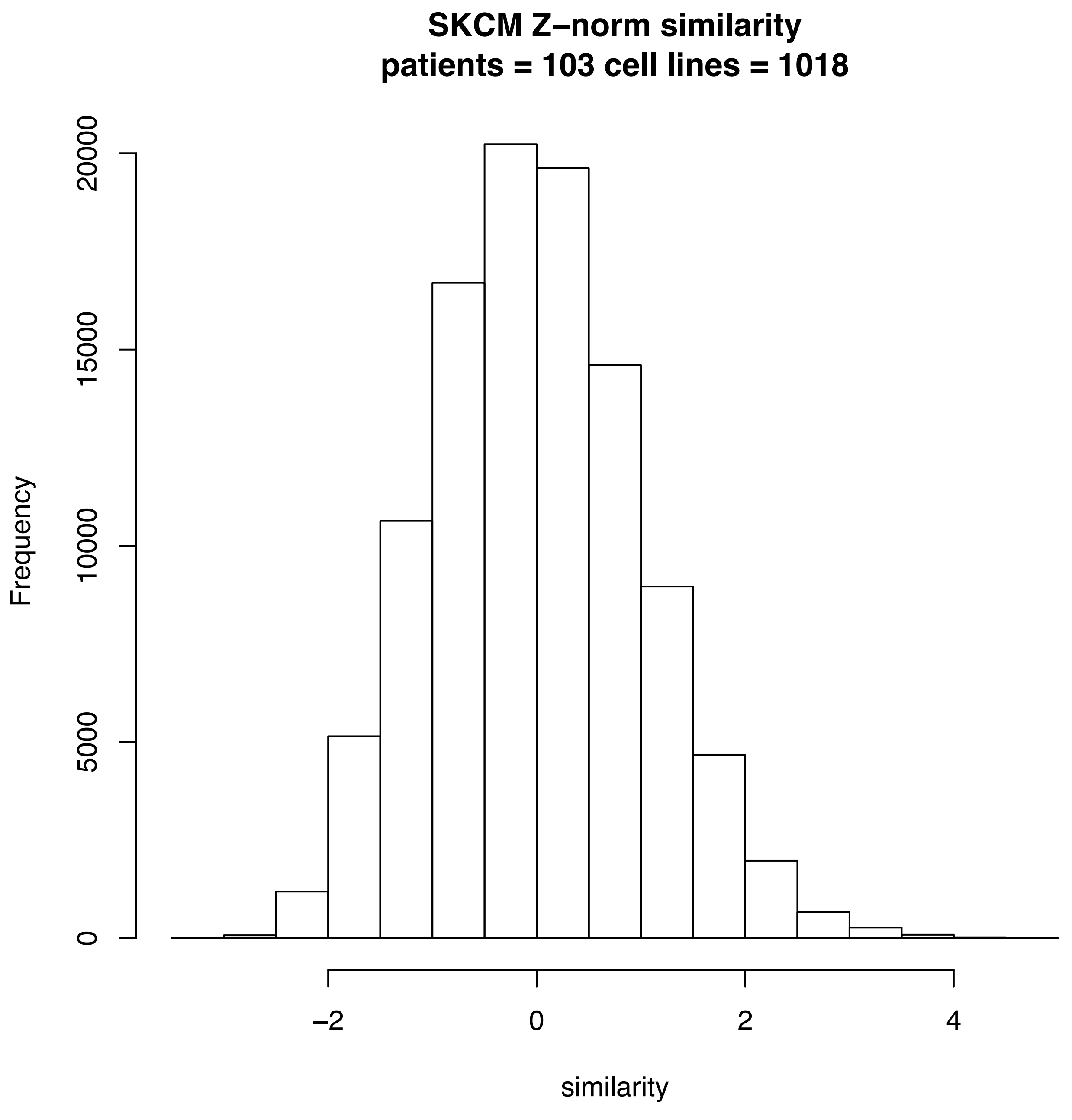
SKCM Z–norm similarity
patients = 103 cell lines = 1018
Frequency
0
5000
10000
15000
20000
−2
0
2
4
similarity

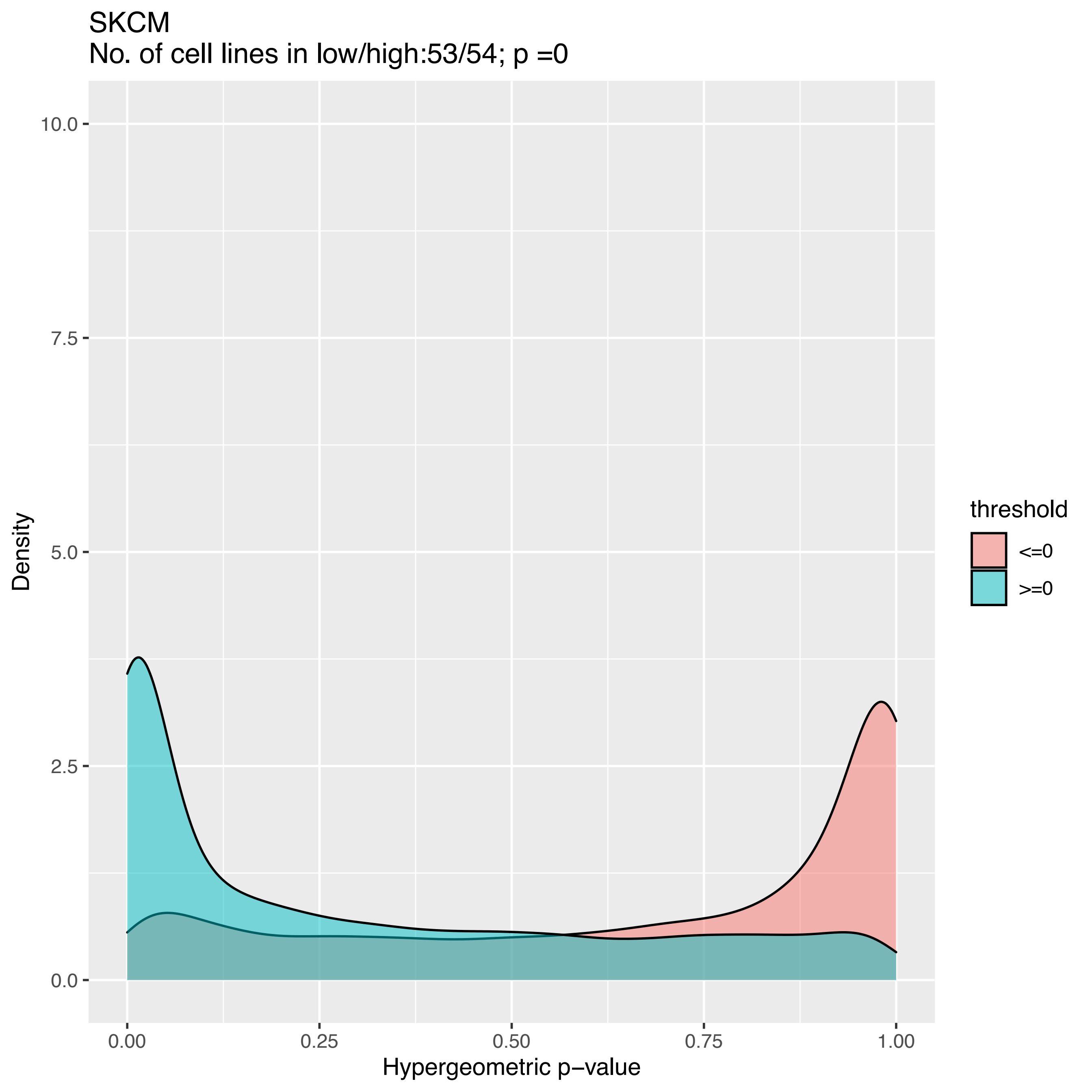
SKCM
No. of cell lines in low/high:53/54; p =0
10.0
7.5
5.0
2.5
0.0
Density
0.00
0.25
0.50
0.75
1.00
Hypergeometric p−value
threshold
<=0
>=0

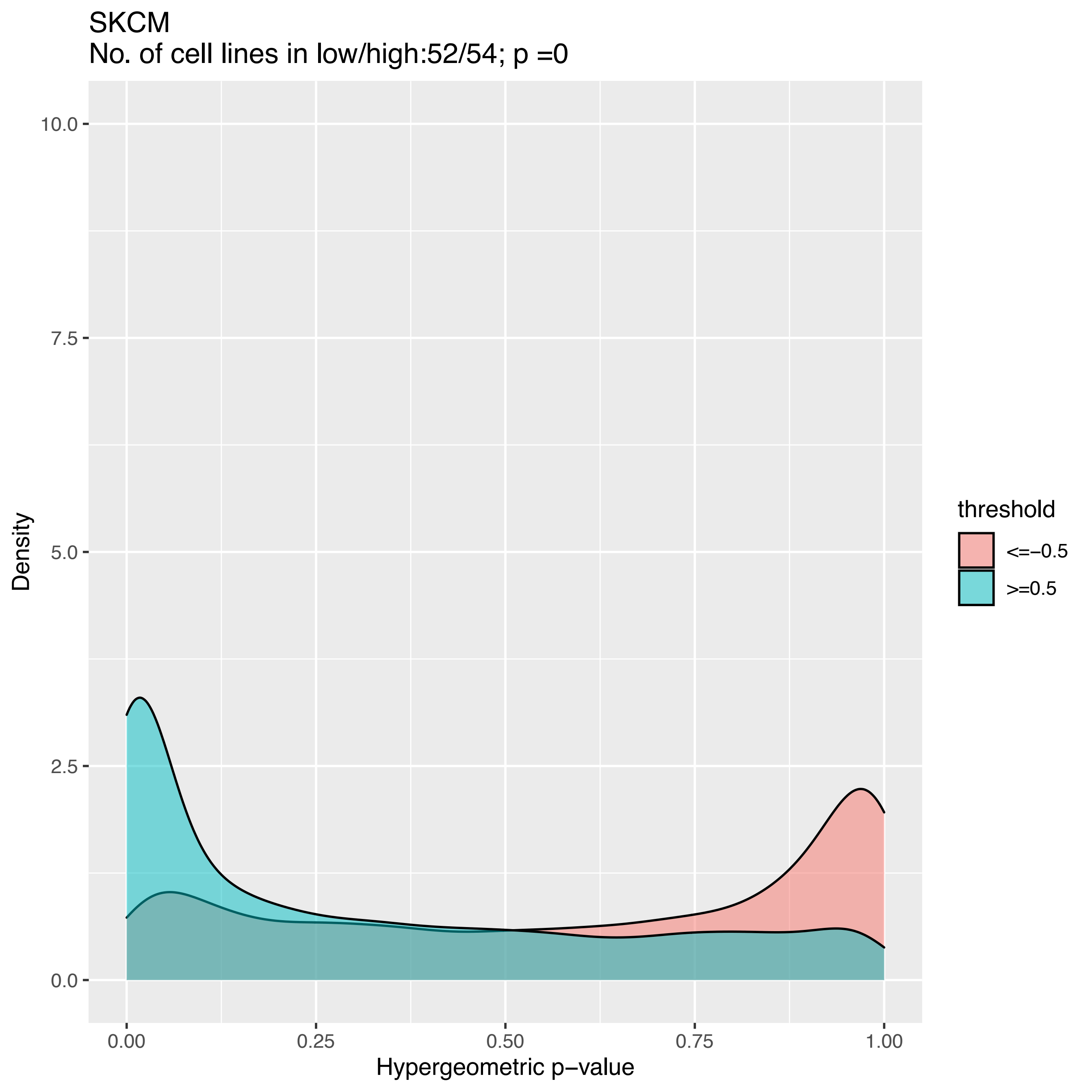
SKCM
No. of cell lines in low/high:52/54; p =0
10.0
7.5
5.0
2.5
0.0
Density
0.00
0.25
0.50
0.75
1.00
Hypergeometric p−value
threshold
<=−0.5
>=0.5

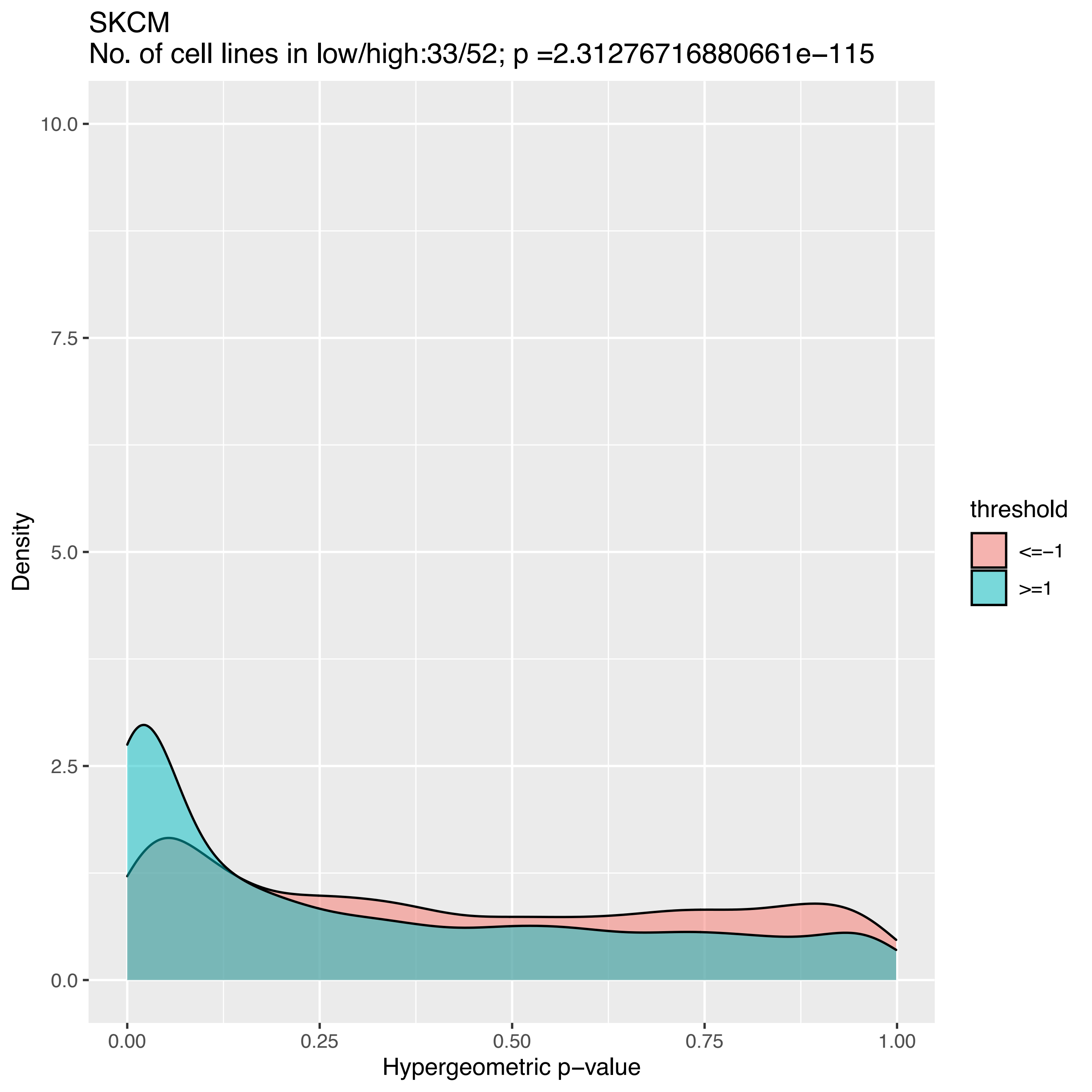
SKCM
No. of cell lines in low/high:33/52; p =2.31276716880661e−115
Density
Hypergeometric p−value
10.0
7.5
5.0
2.5
0.0
0.00
0.25
0.50
0.75
1.00
threshold
<=−1
>=1

SKCM
No. of cell lines in low/high:15/51; p =1

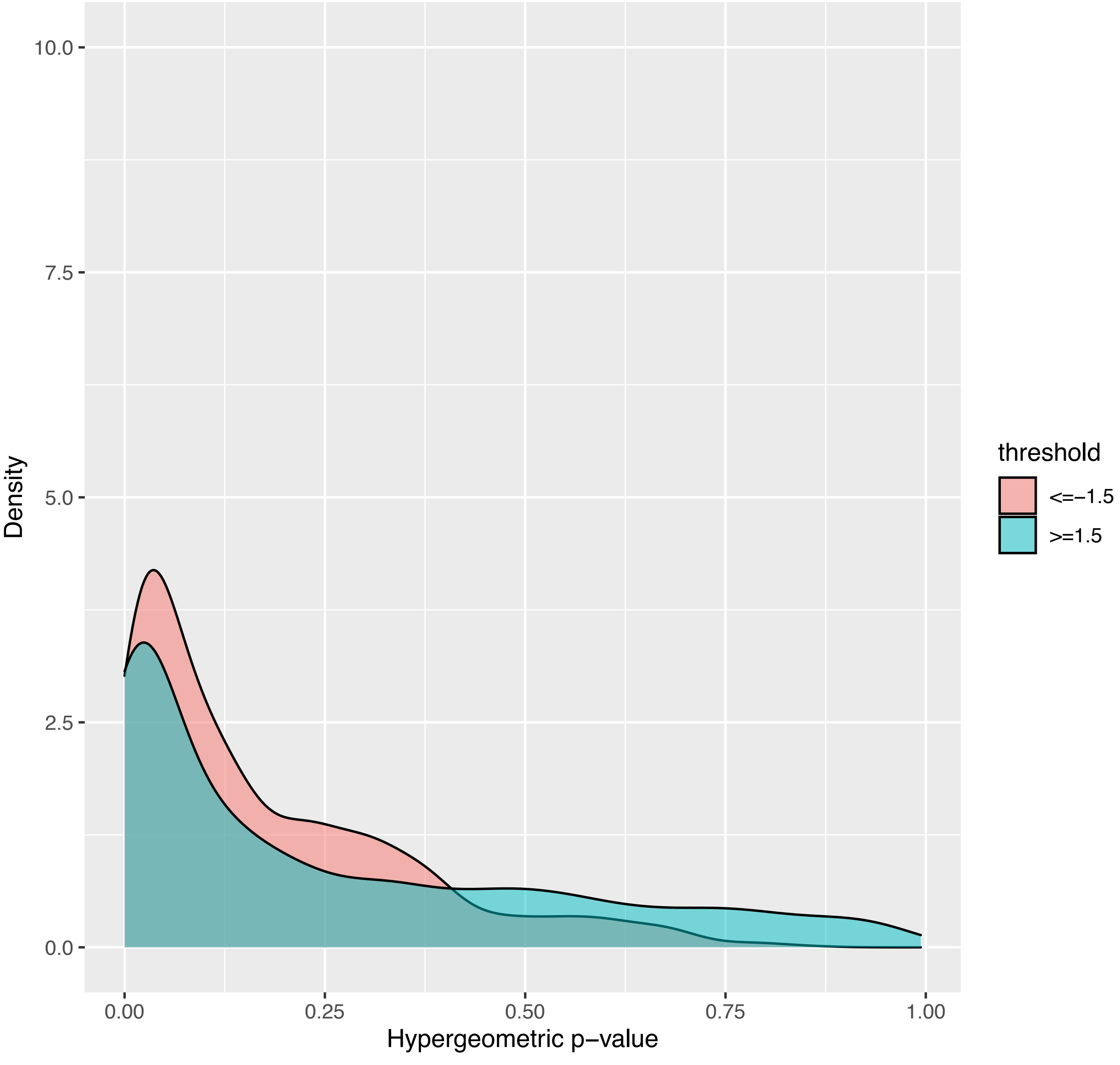

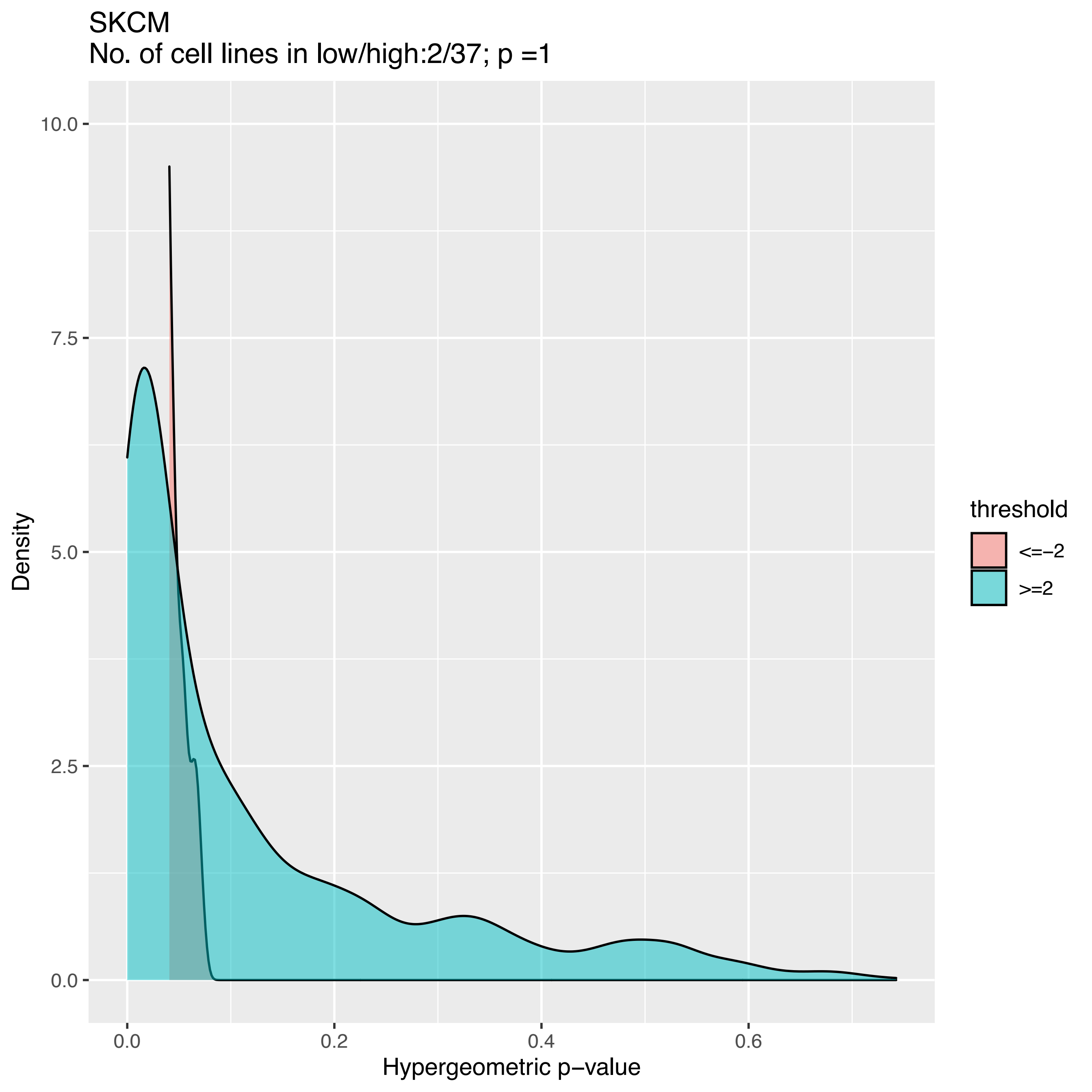
SKCM
No. of cell lines in low/high:2/37; p =1
Density
Hypergeometric p−value
10.0
7.5
5.0
2.5
0.0
0.0
0.2
0.4
0.6
threshold
<=−2
>=2

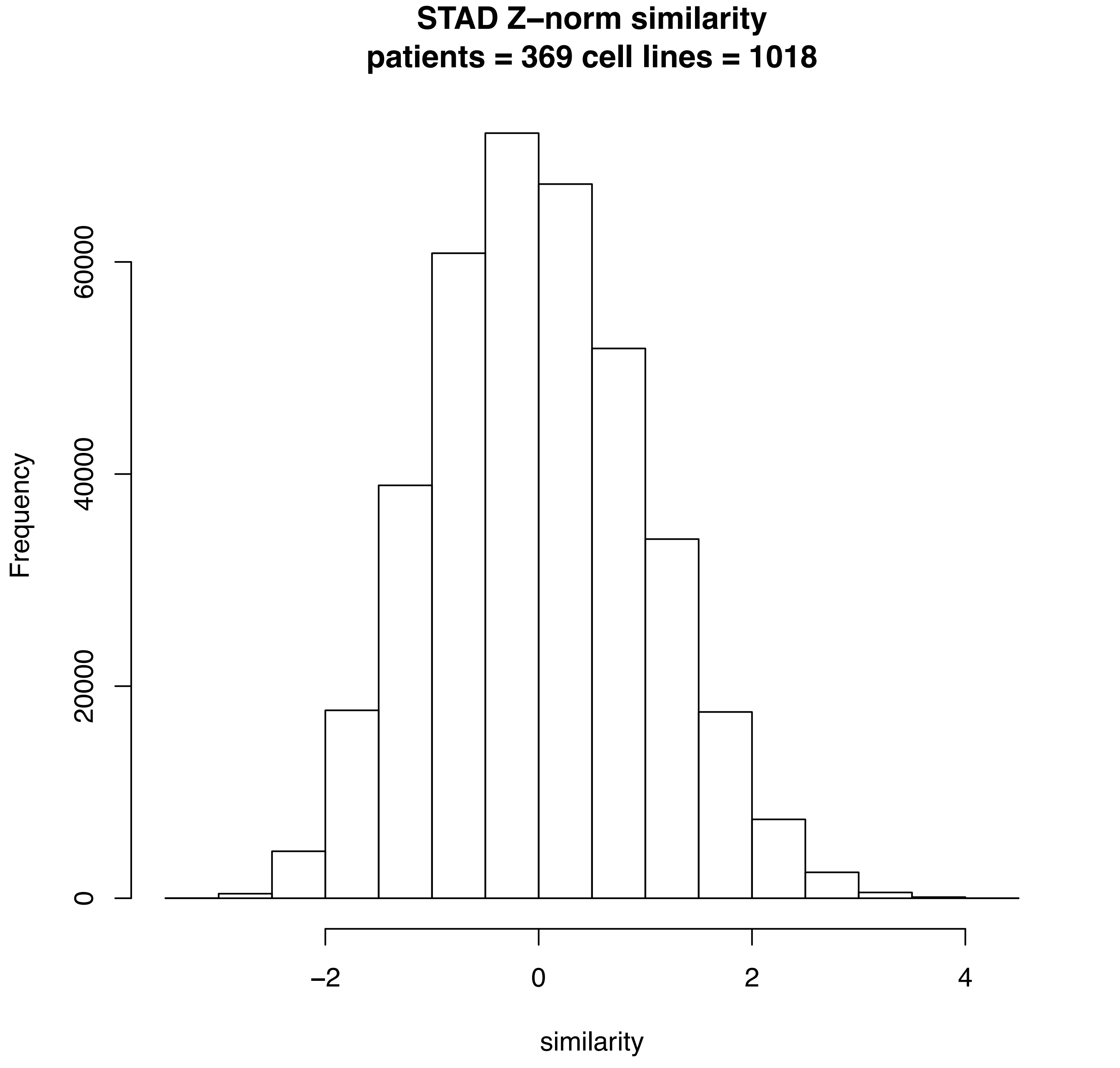
STAD Z−norm similarity
patients = 369 cell lines = 1018
Frequency
0
20000
40000
60000
−2
0
2
4
similarity

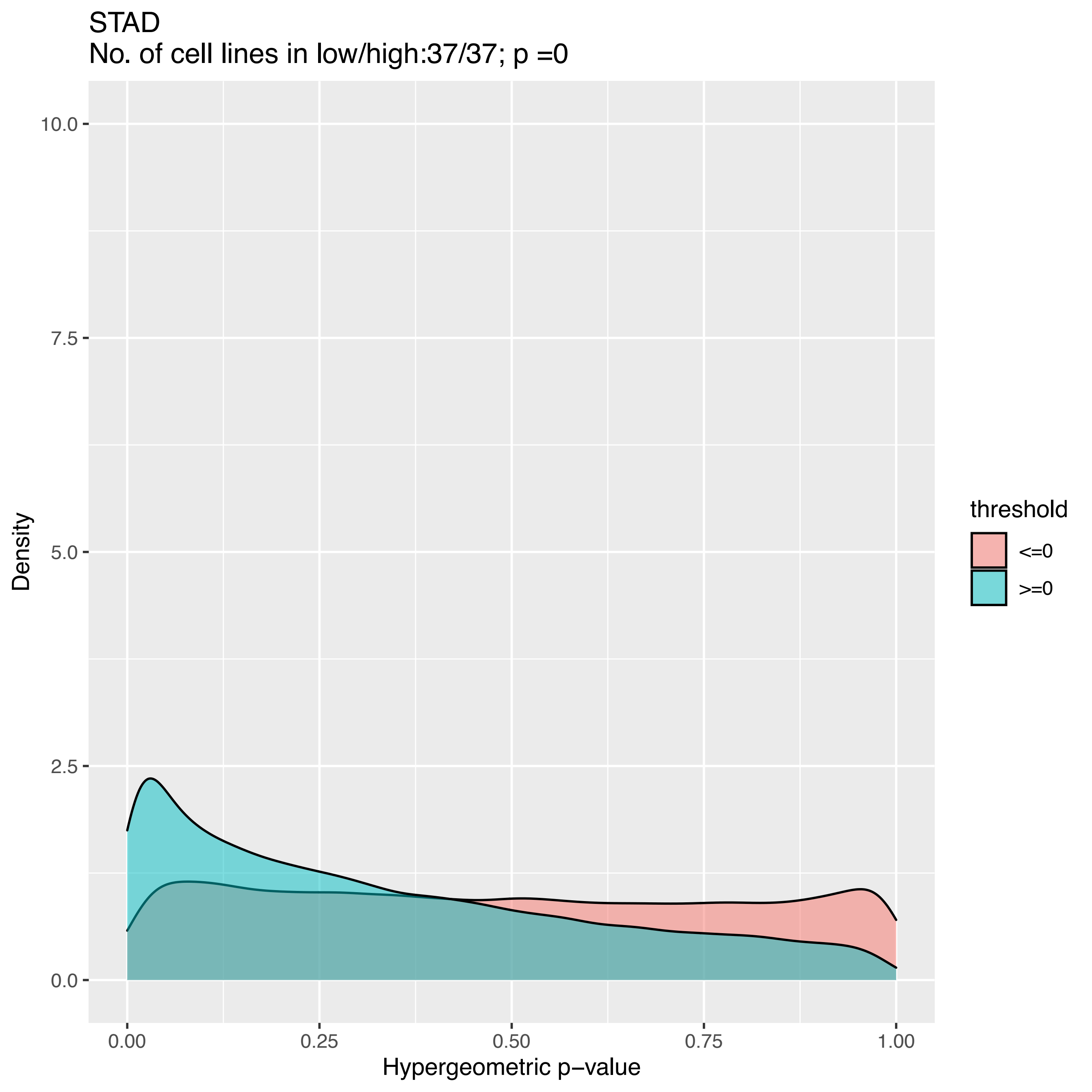
STAD
No. of cell lines in low/high:37/37; p =0
Density
10.0
7.5
5.0
2.5
0.0
0.00
0.25
0.50
0.75
1.00
Hypergeometric p−value
threshold
<=0
>=0

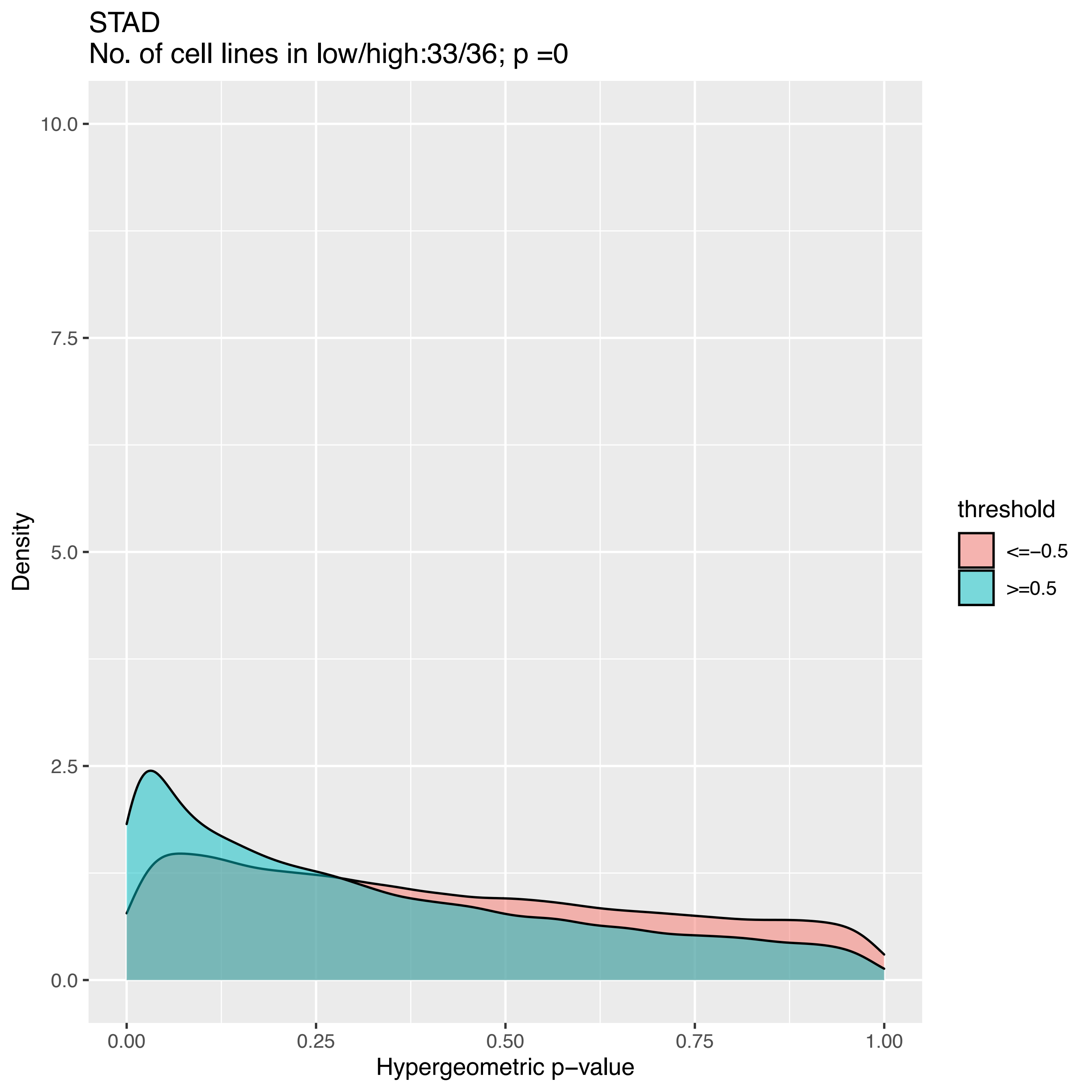

STAD
No. of cell lines in low/high:33/36; p =0
10.0
7.5
5.0
2.5
0.0
Density
0.00
0.25
0.50
0.75
1.00
Hypergeometric p−value
threshold
<=−0.5
>=0.5

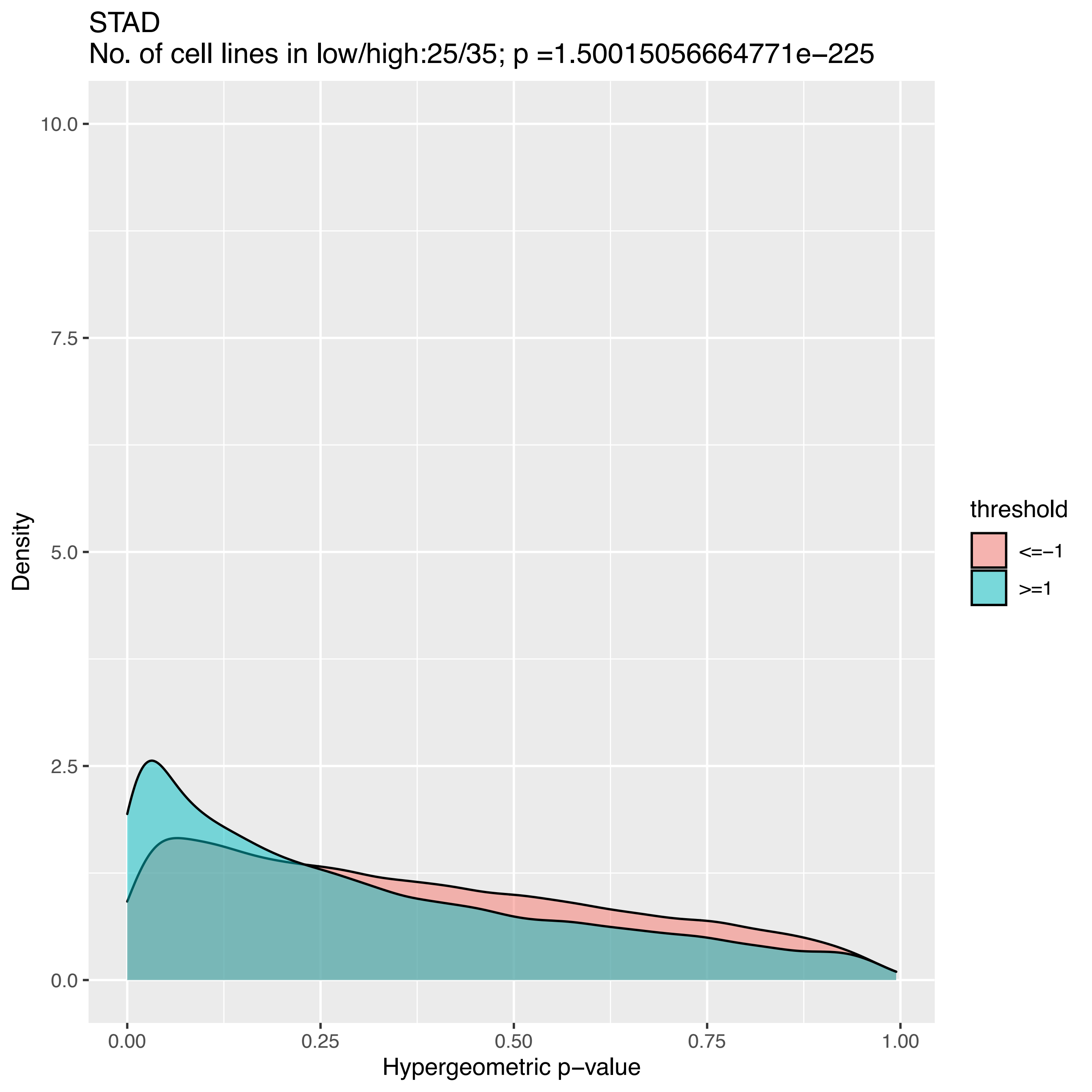

STAD
No. of cell lines in low/high:25/35; p =1.50015056664771e−225
Density
10.0
7.5
5.0
2.5
0.0
0.00
0.25
0.50
0.75
1.00
Hypergeometric p−value
threshold
<=−1
>=1

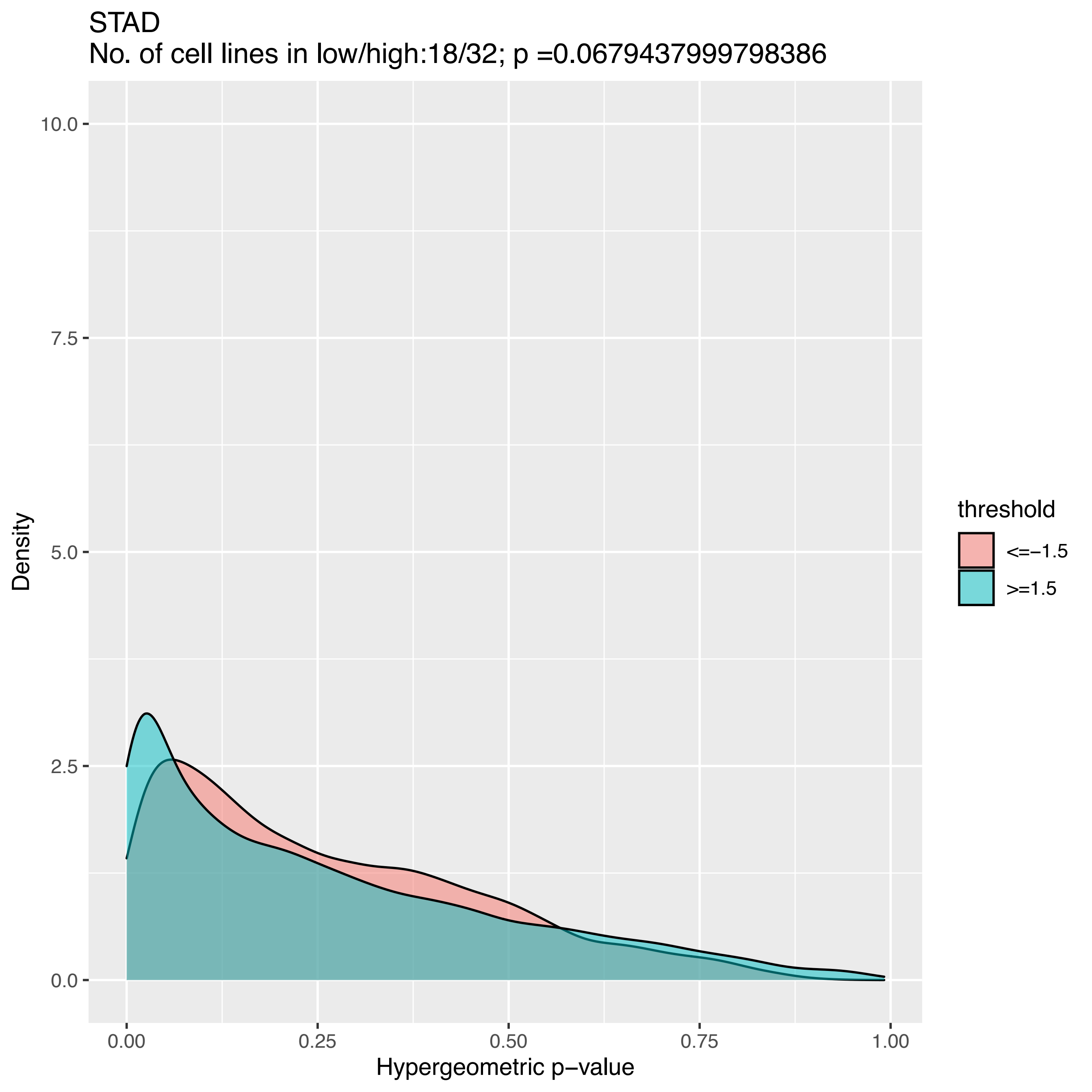
STAD
No. of cell lines in low/high:18/32; p =0.0679437999798386
Density
10.0
7.5
5.0
2.5
0.0
0.00
0.25
0.50
0.75
1.00
Hypergeometric p−value
threshold
<=−1.5
>=1.5

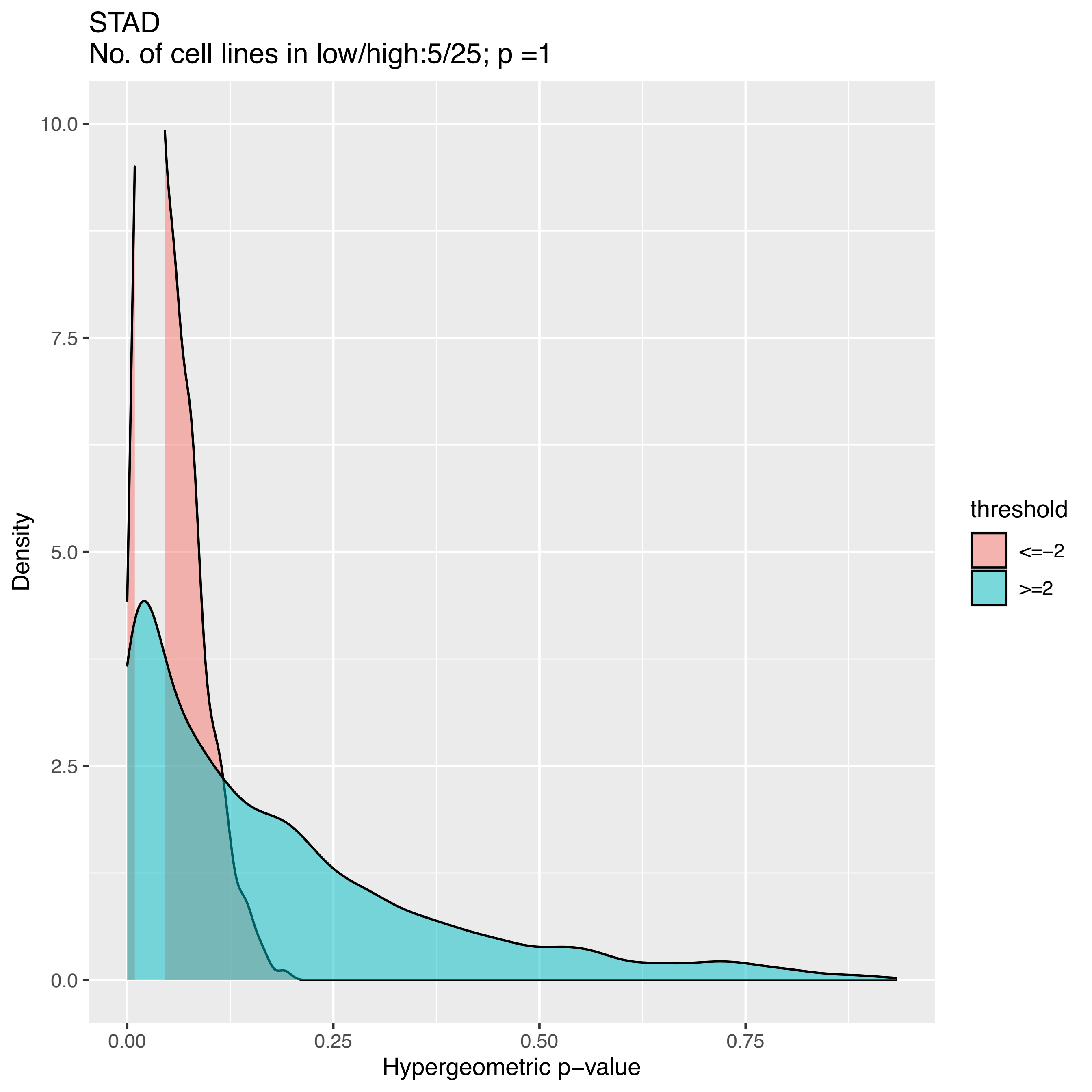

STAD
No. of cell lines in low/high:5/25; p =1
Density
10.0
7.5
5.0
2.5
0.0
0.00
0.25
0.50
0.75
Hypergeometric p−value
threshold
<=−2
>=2

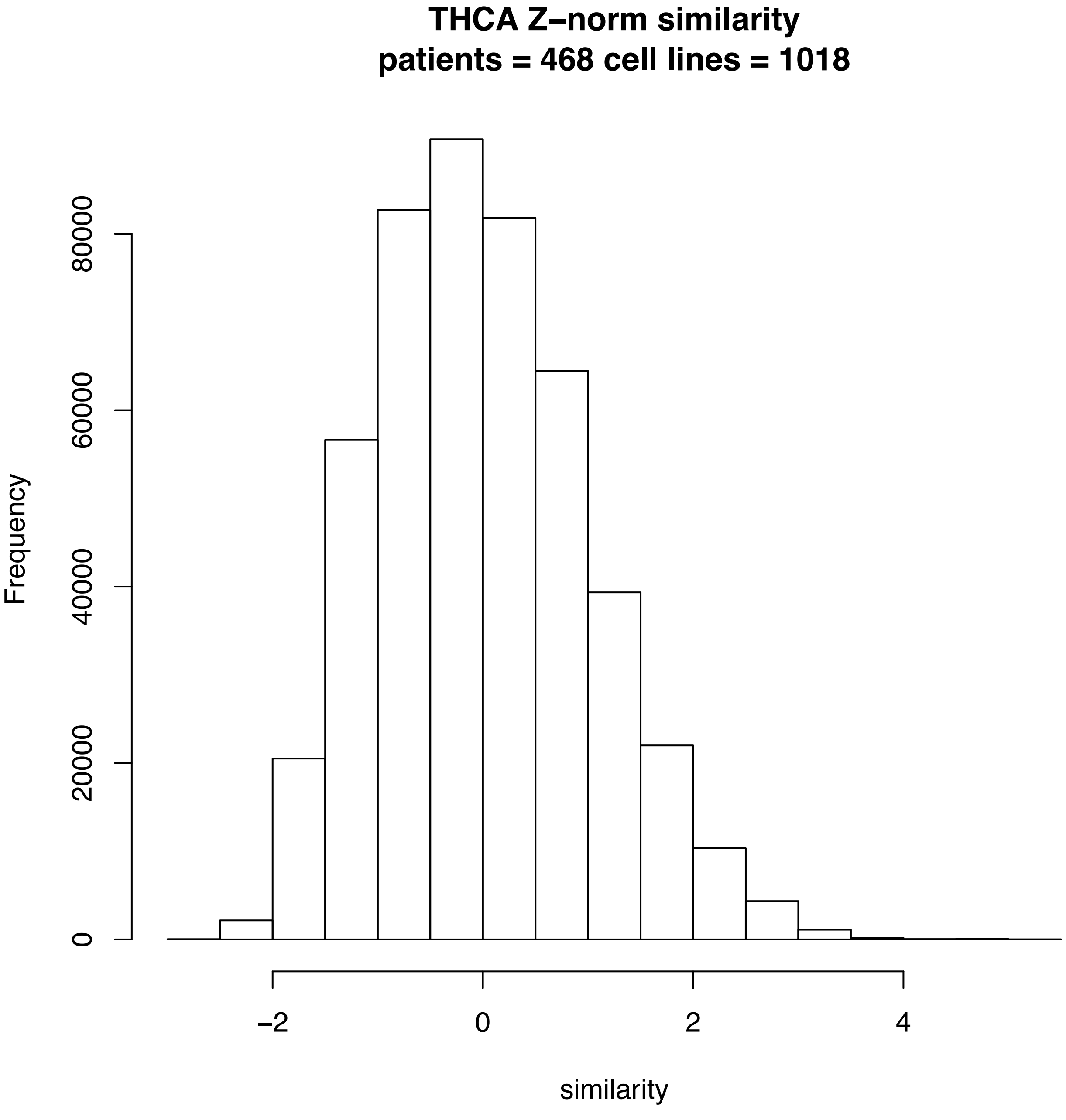
THCA Z–norm similarity
patients = 468 cell lines = 1018
Frequency
0
20000
40000
60000
80000
−2
0
2
4
similarity

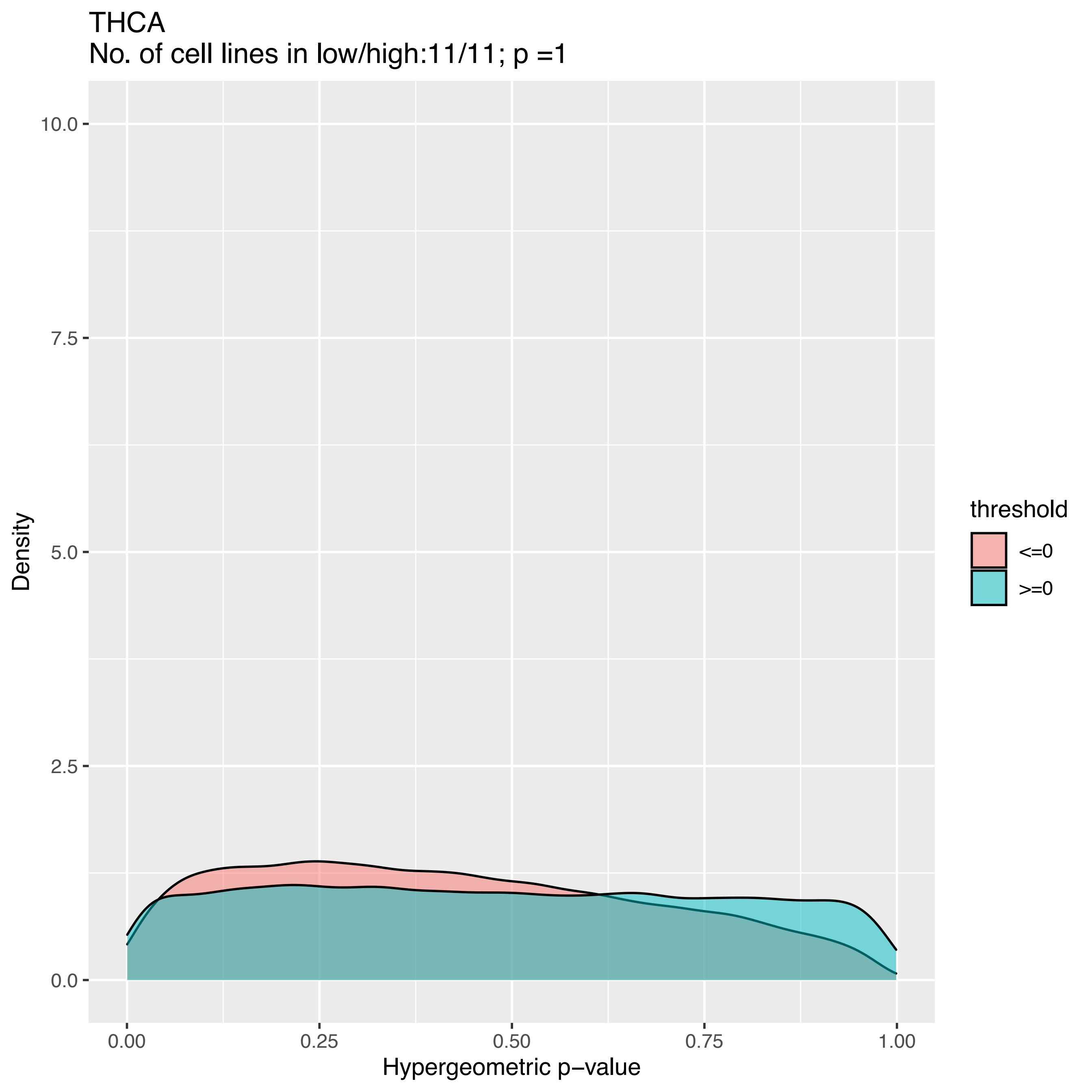
THCA
No. of cell lines in low/high:11/11; p =1
Density
10.0
7.5
5.0
2.5
0.0
0.00
0.25
0.50
0.75
1.00
Hypergeometric p−value
threshold
<=0
>=0

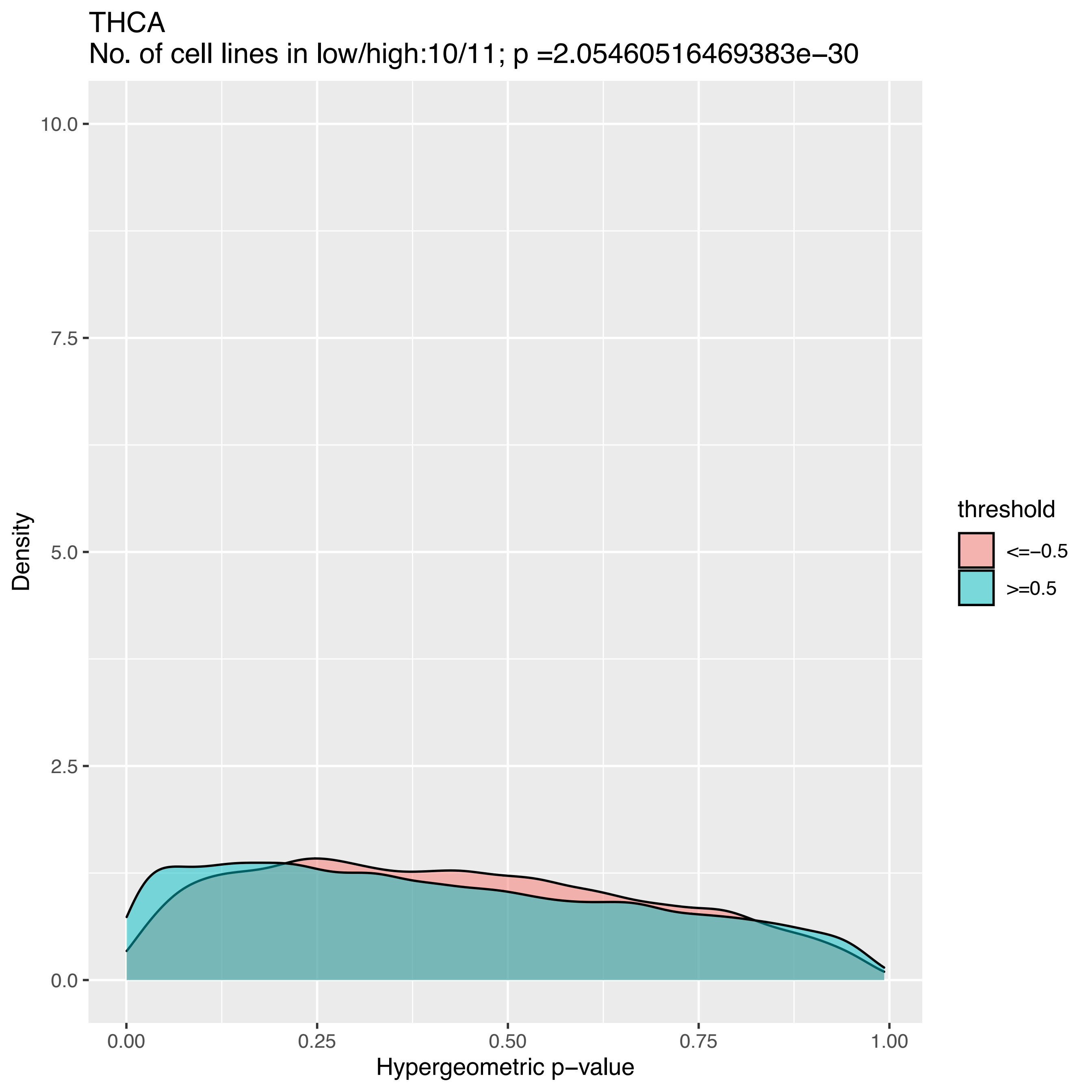
THCA
No. of cell lines in low/high:10/11; p =2.05460516469383e−30
Density
10.0
7.5
5.0
2.5
0.0
0.00
0.25
0.50
0.75
1.00
Hypergeometric p−value
threshold
<=−0.5
>=0.5

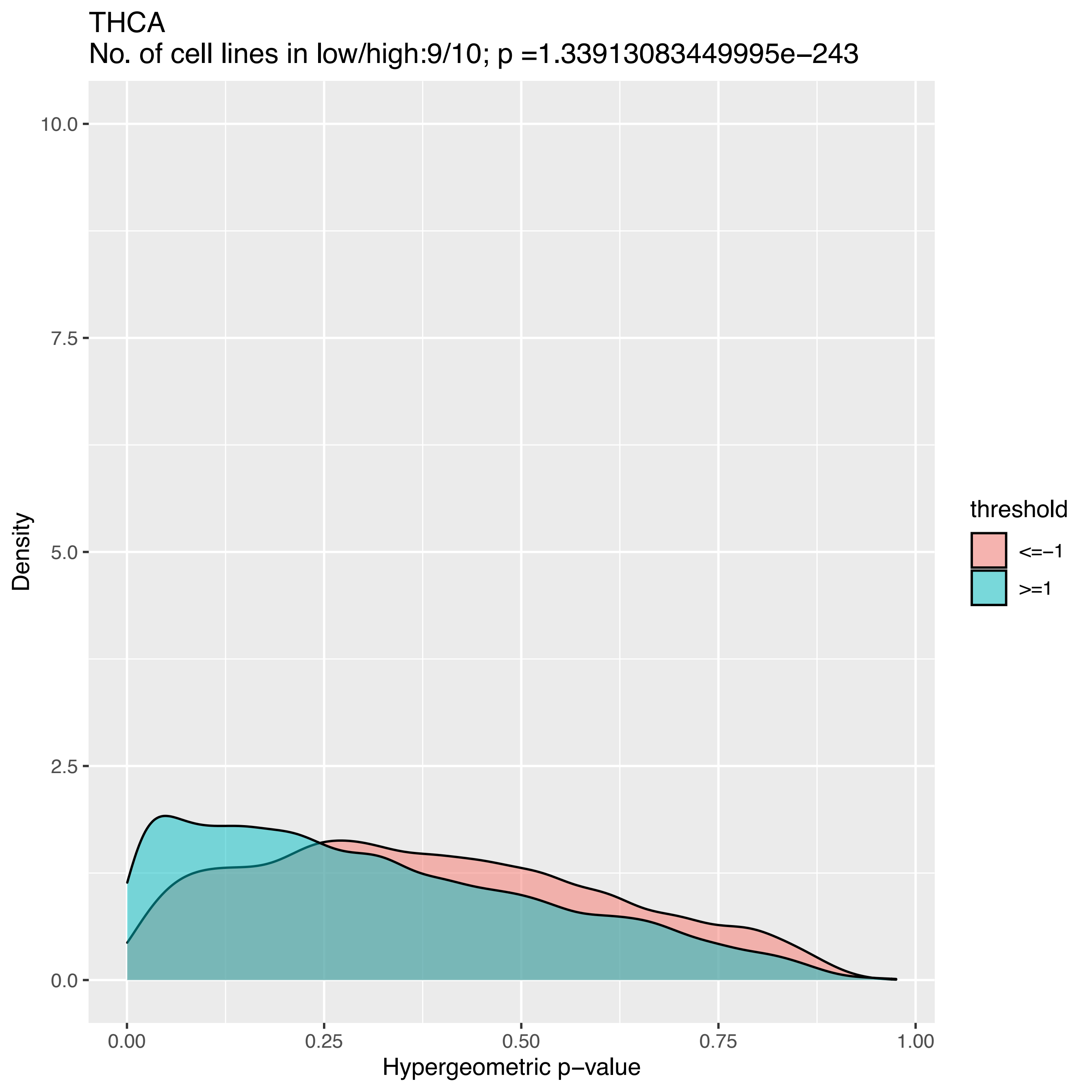
THCA
No. of cell lines in low/high:9/10; p =1.33913083449995e−243
10.0
7.5
5.0
2.5
0.0
Density
0.00
0.25
0.50
0.75
1.00
Hypergeometric p−value
threshold
<=−1
>=1

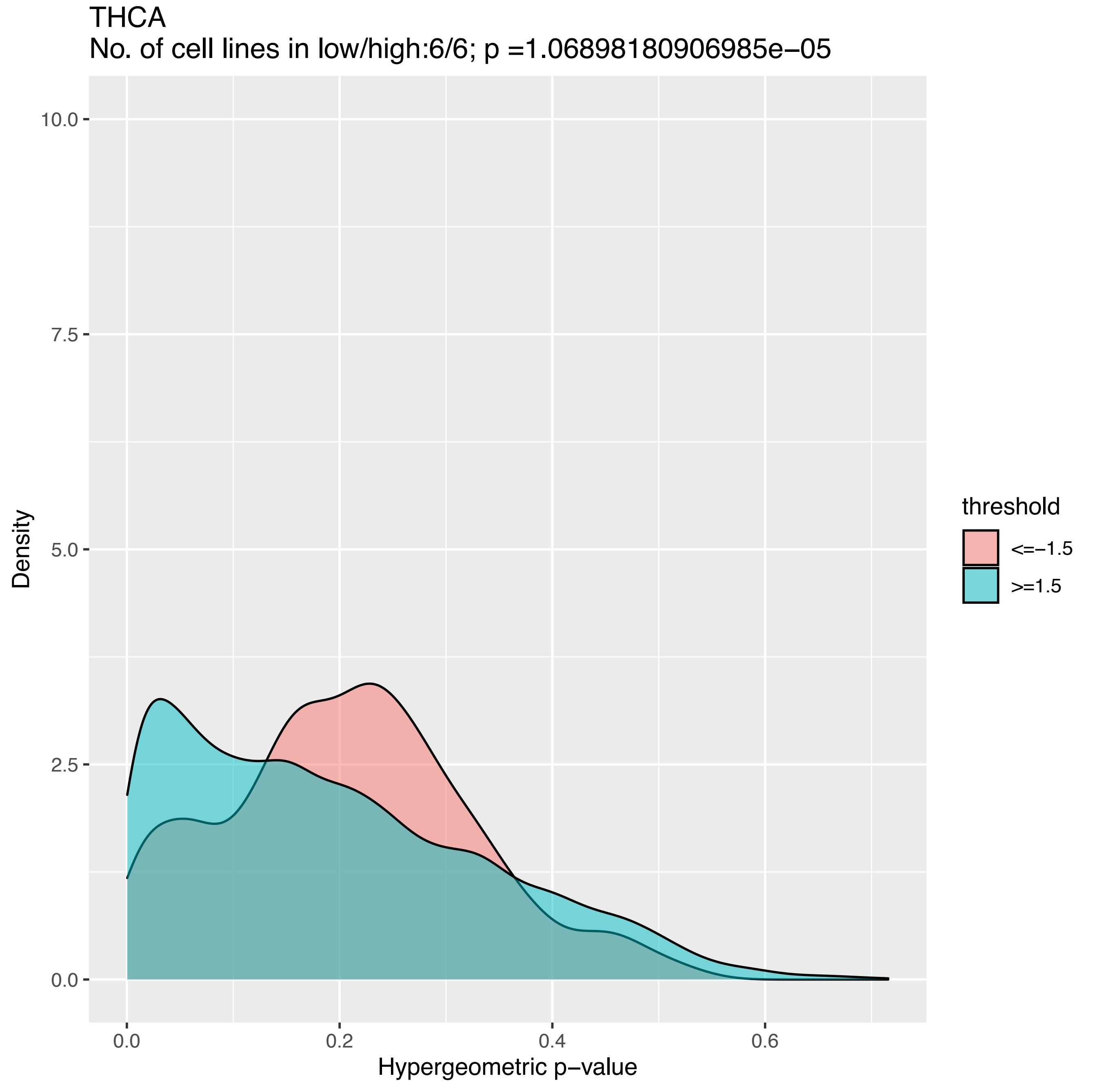
THCA
No. of cell lines in low/high:6/6; p =1.06898180906985e−05
Density
10.0
7.5
5.0
2.5
0.0
0.0
0.2
0.4
0.6
Hypergeometric p−value
threshold
<=−1.5
>=1.5

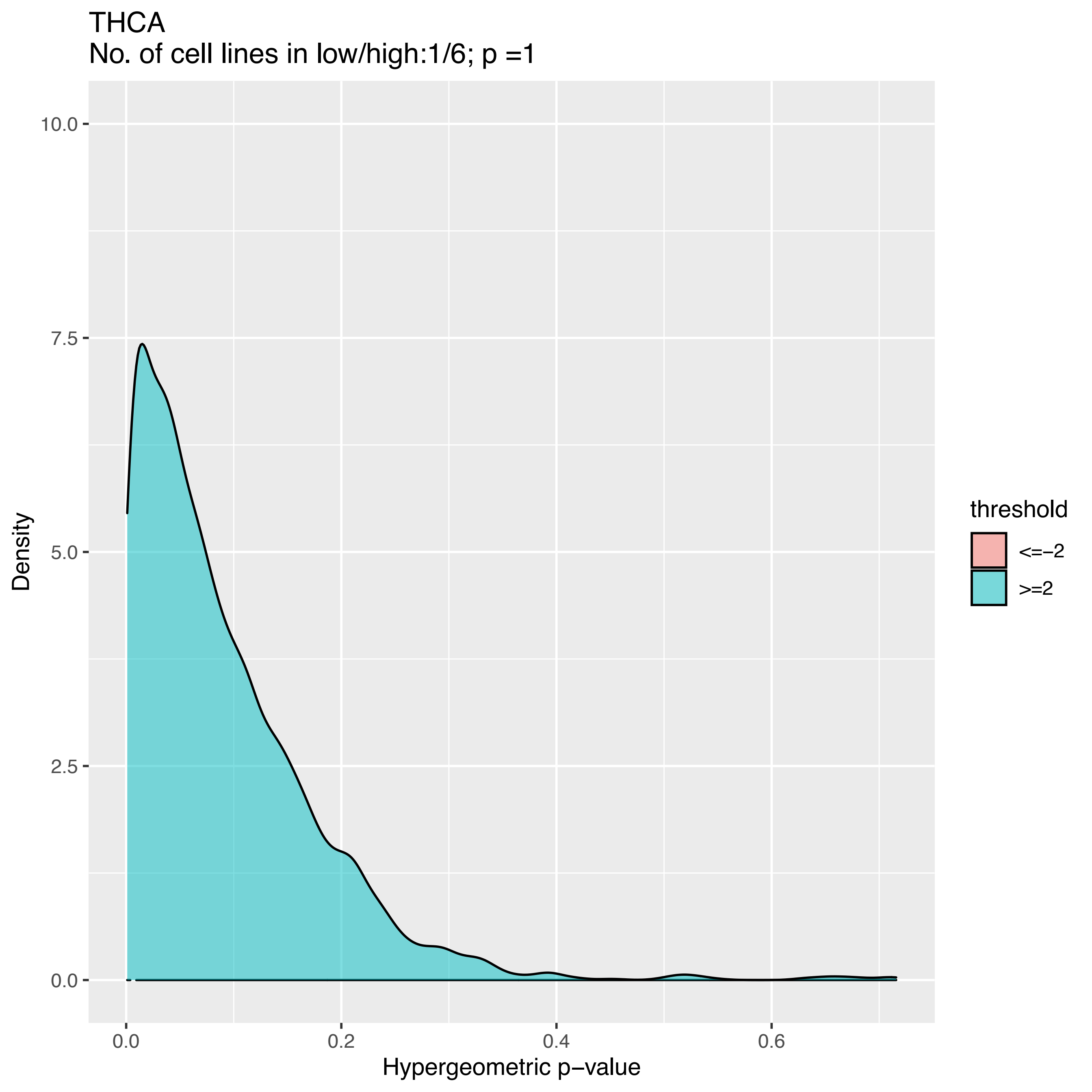
THCA
No. of cell lines in low/high:1/6; p =1
Density
10.0
7.5
5.0
2.5
0.0
0.0
0.2
0.4
0.6
Hypergeometric p−value
threshold
<=−2
>=2

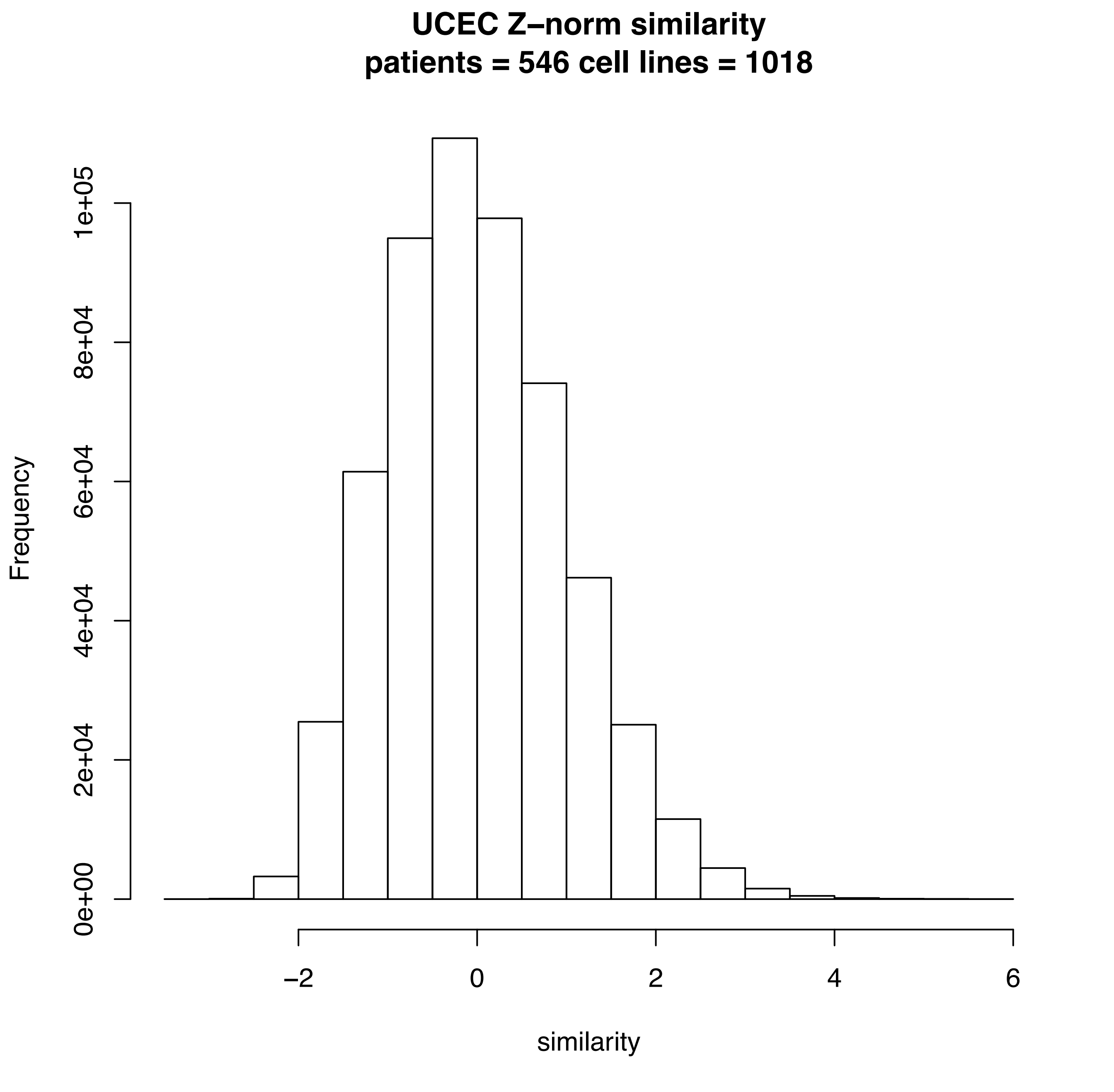
UCEC Z–norm similarity
patients = 546 cell lines = 1018
Frequency
0e+00
2e+04
4e+04
6e+04
8e+04
1e+05
–2
0
2
4
6
similarity

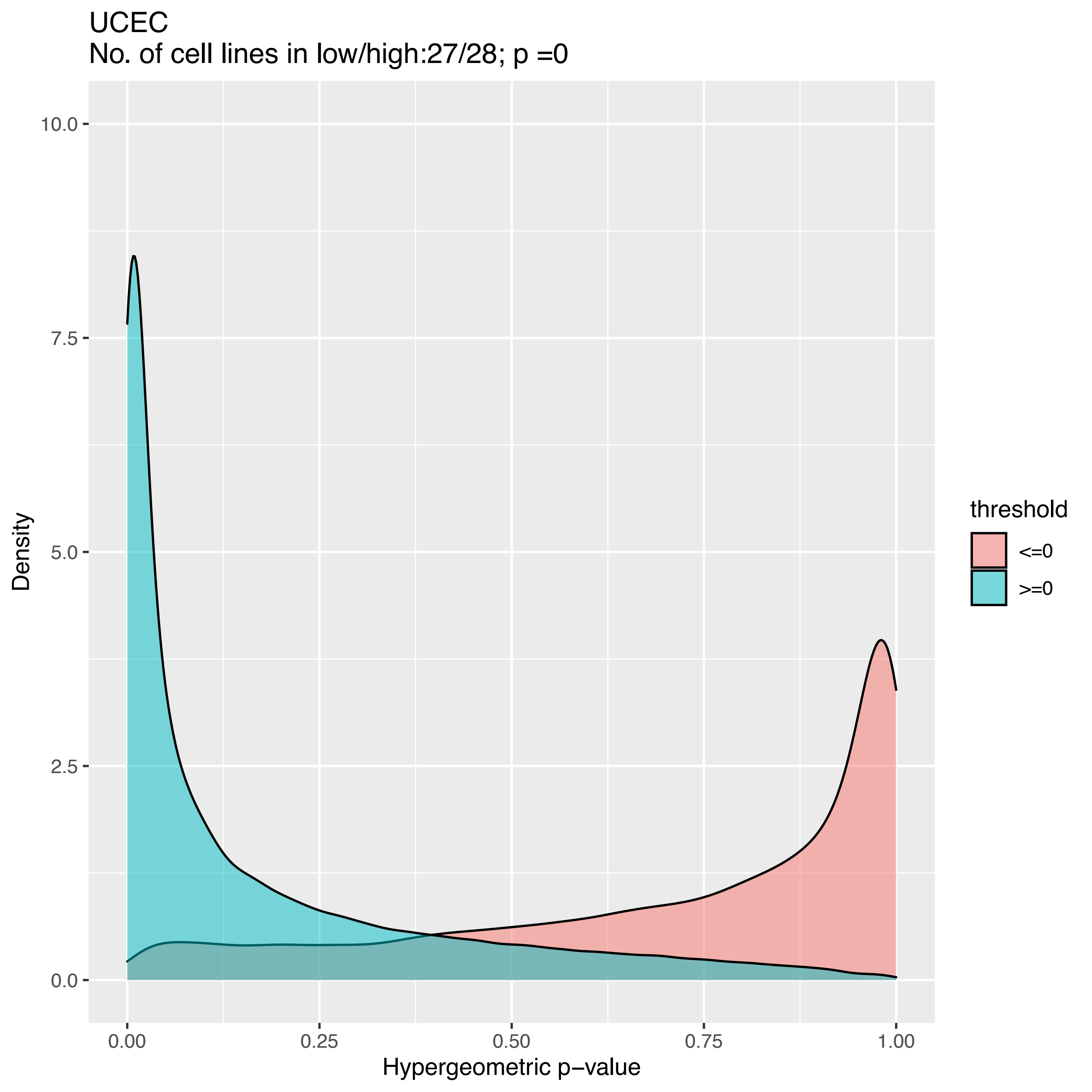
UCEC
No. of cell lines in low/high:27/28; p =0
Density
10.0
7.5
5.0
2.5
0.0
0.00
0.25
0.50
0.75
1.00
Hypergeometric p−value
threshold
<=0
>=0

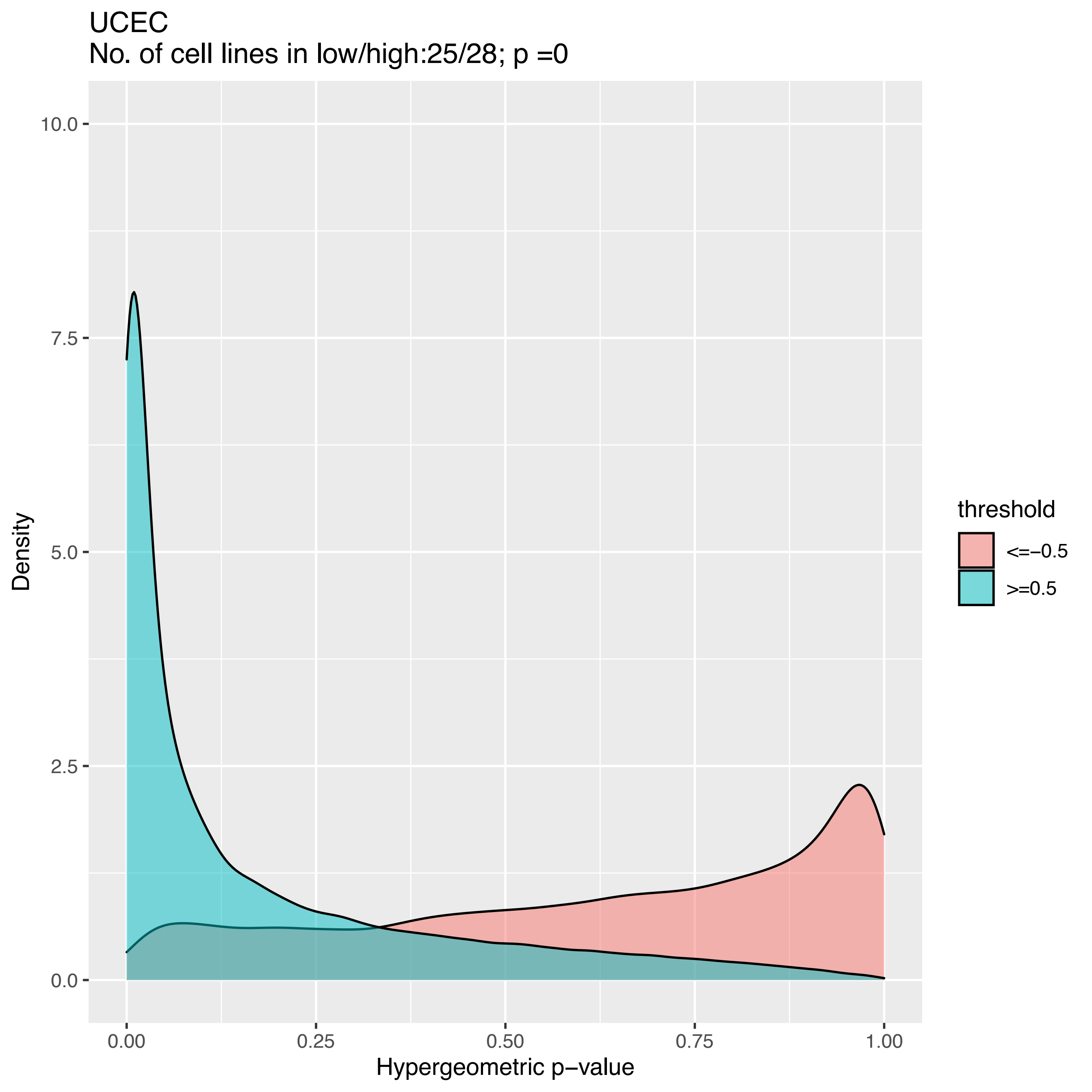
UCEC
No. of cell lines in low/high:25/28; p =0
10.0
7.5
5.0
2.5
0.0
Density
0.00
0.25
0.50
0.75
1.00
Hypergeometric p−value
threshold
<=−0.5
>=0.5

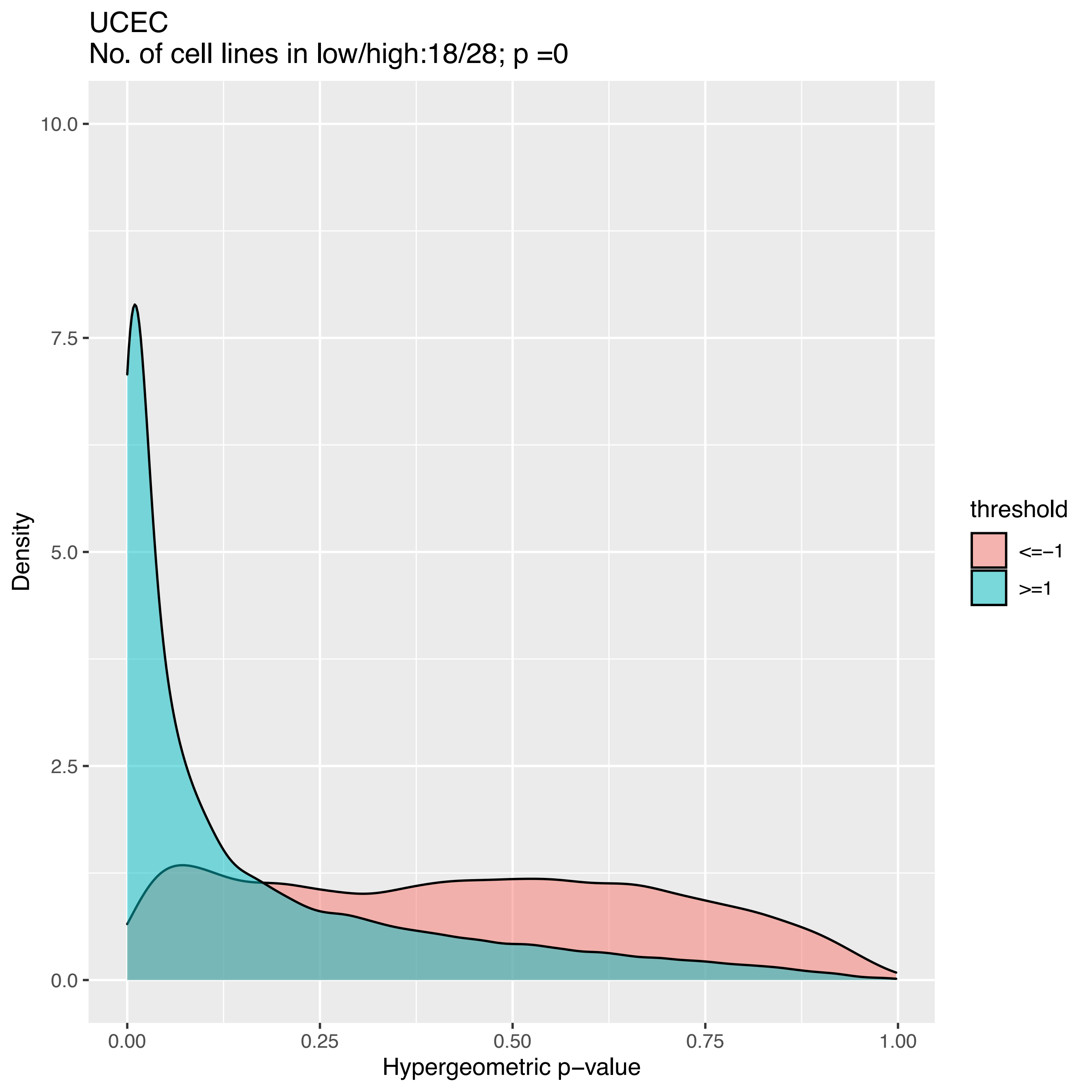

UCEC
No. of cell lines in low/high:18/28; p =0
Density
10.0
7.5
5.0
2.5
0.0
0.00
0.25
0.50
0.75
1.00
Hypergeometric p−value
threshold
<=−1
>=1

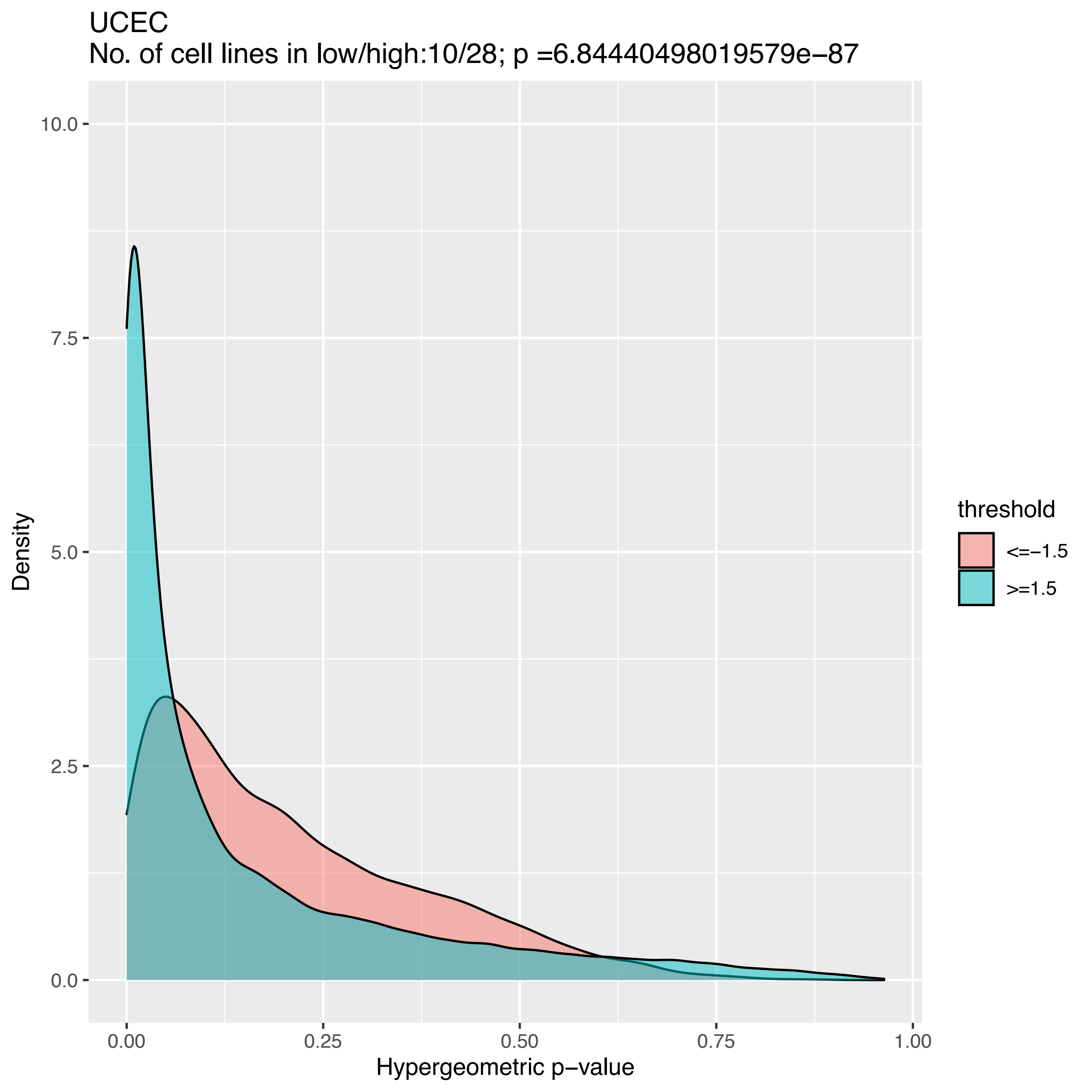
UCEC
No. of cell lines in low/high:10/28; p =6.84440498019579e−87
Density
10.0
7.5
5.0
2.5
0.0
0.00
0.25
0.50
0.75
1.00
Hypergeometric p−value
threshold
<=−1.5
>=1.5

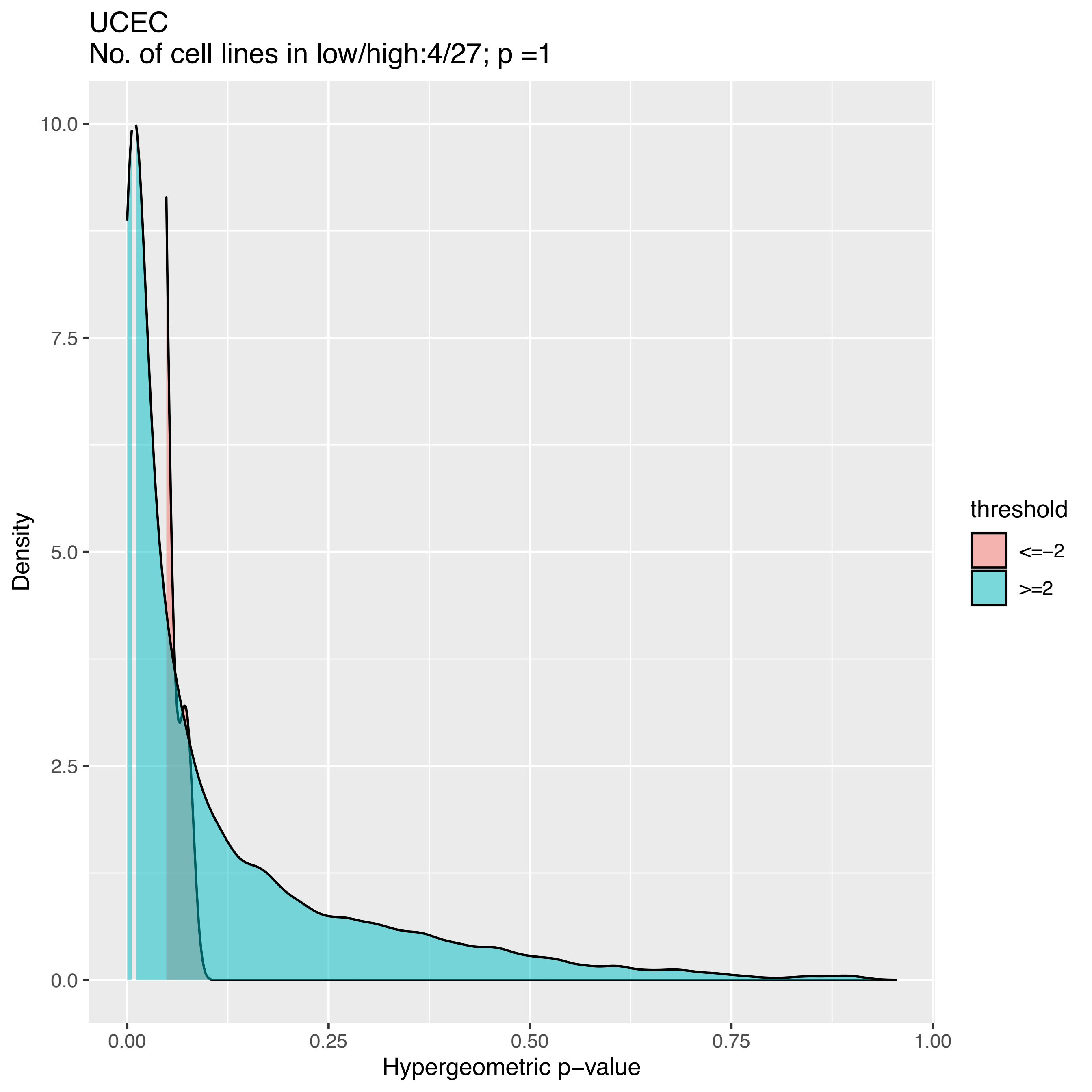

UCEC
No. of cell lines in low/high:4/27; p =1
Density
10.0
7.5
5.0
2.5
0.0
0.00
0.25
0.50
0.75
1.00
Hypergeometric p−value
threshold
<=−2
>=2

### CTDPathSim1.0

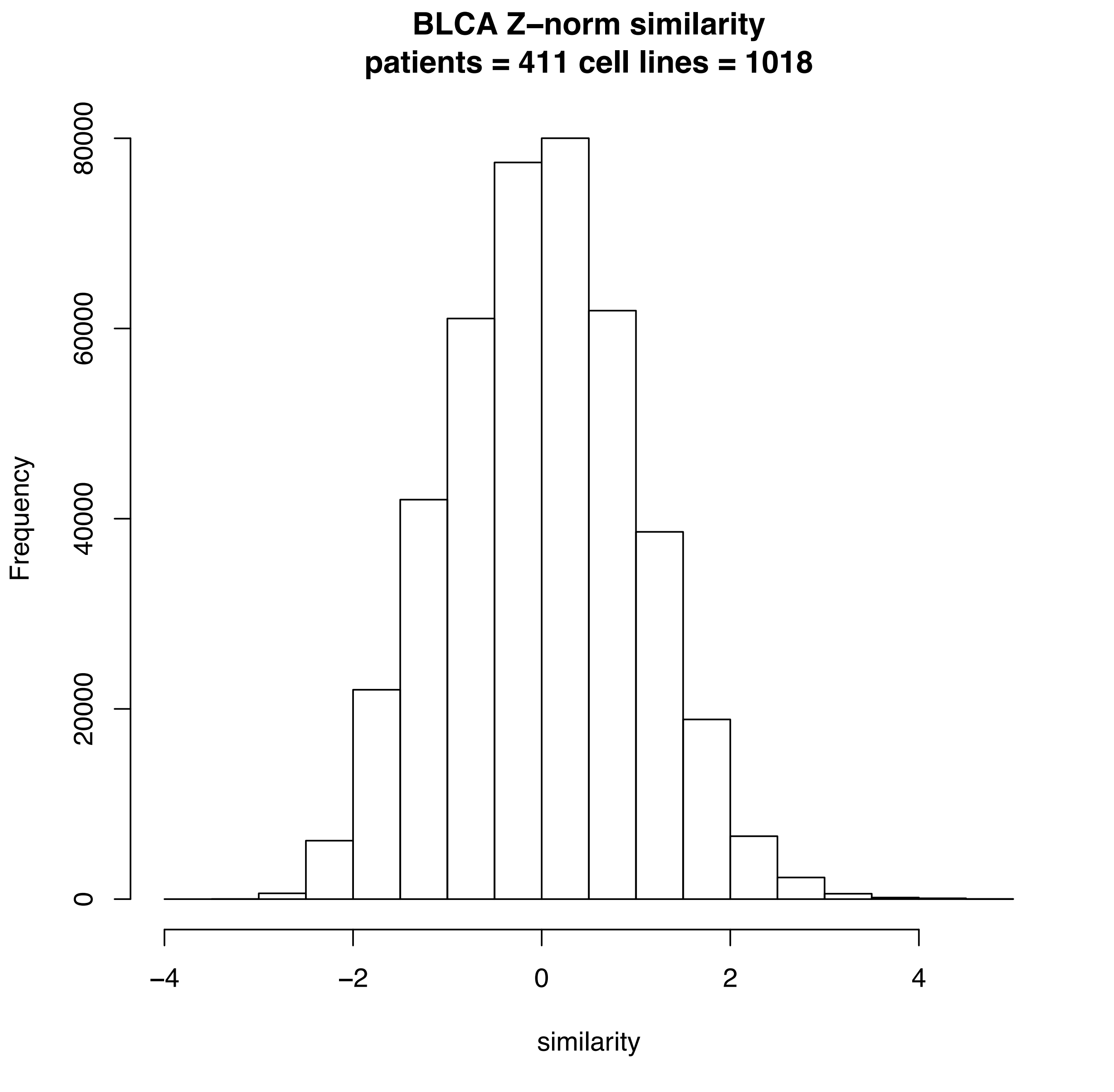
BLCA Z–norm similarity
patients = 411 cell lines = 1018
Frequency
0
20000
40000
60000
80000
–4
–2
0
2
4
similarity

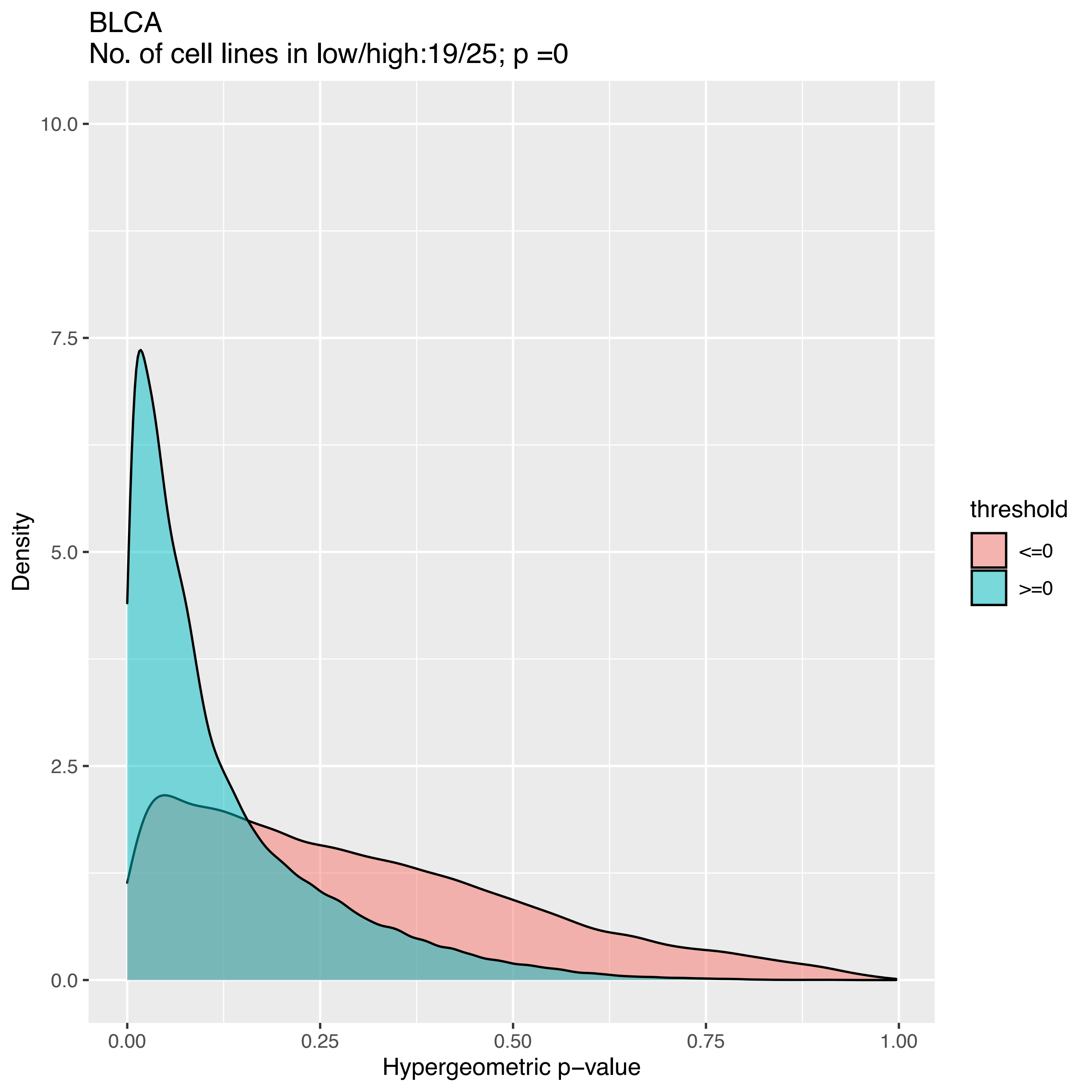

BLCA
No. of cell lines in low/high:19/25; p =0
Density
10.0
7.5
5.0
2.5
0.0
0.00
0.25
0.50
0.75
1.00
Hypergeometric p−value
threshold
<=0
>=0

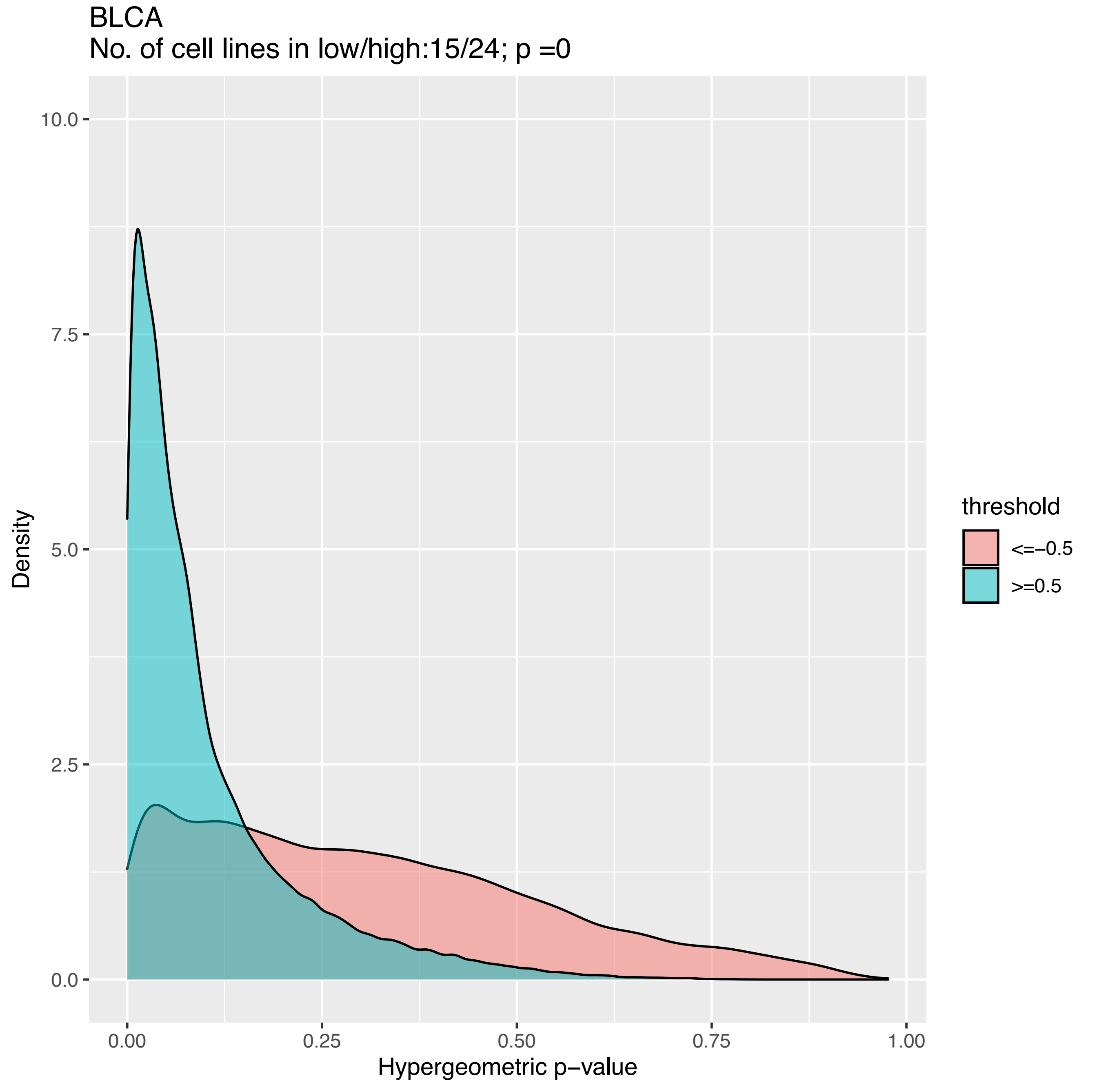
BLCA
No. of cell lines in low/high:15/24; p =0
Density
10.0
7.5
5.0
2.5
0.0
0.00
0.25
0.50
0.75
1.00
Hypergeometric p−value
threshold
<=−0.5
>=0.5

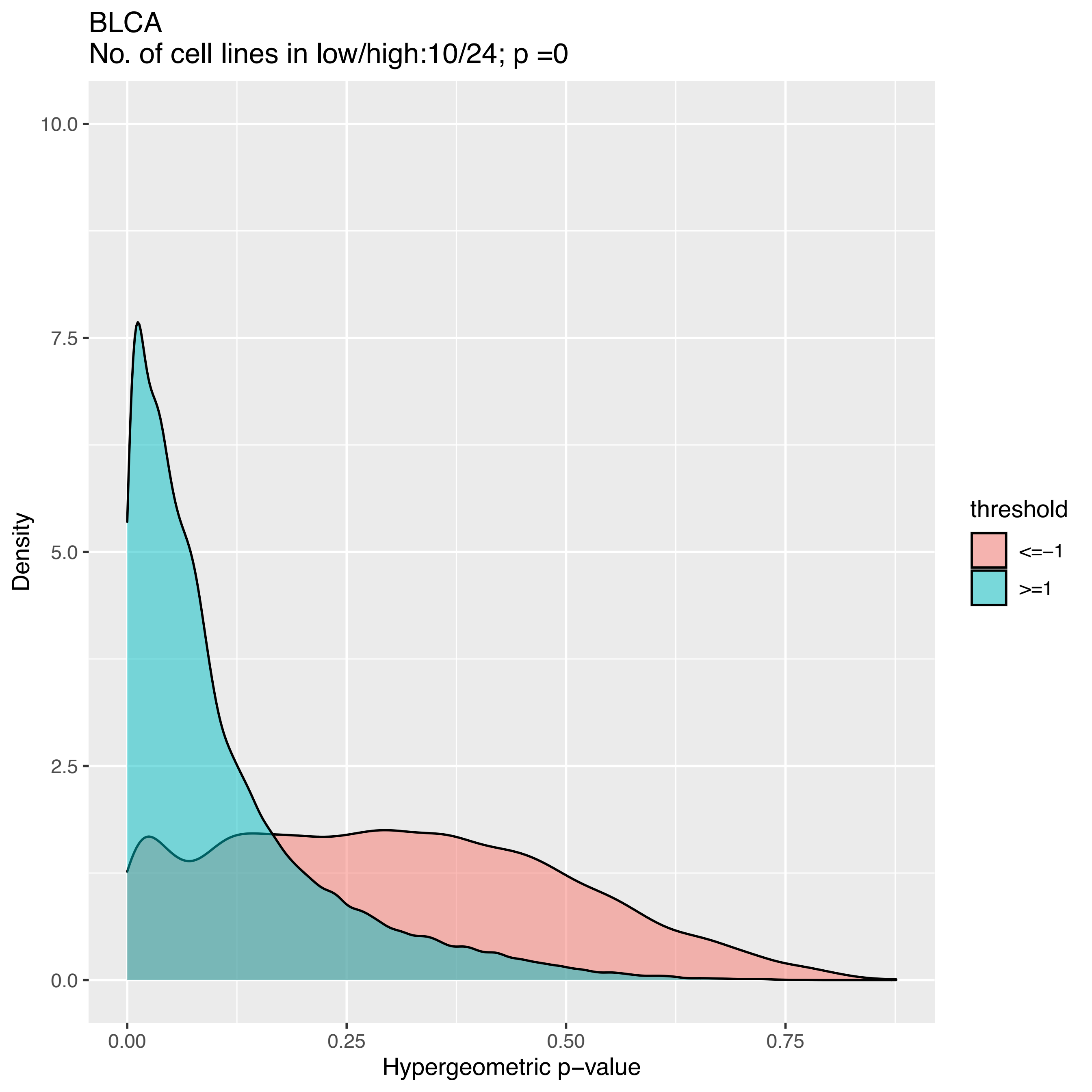
BLCA
No. of cell lines in low/high:10/24; p =0
Density
10.0
7.5
5.0
2.5
0.0
0.00
0.25
0.50
0.75
Hypergeometric p−value
threshold
<=−1
>=1

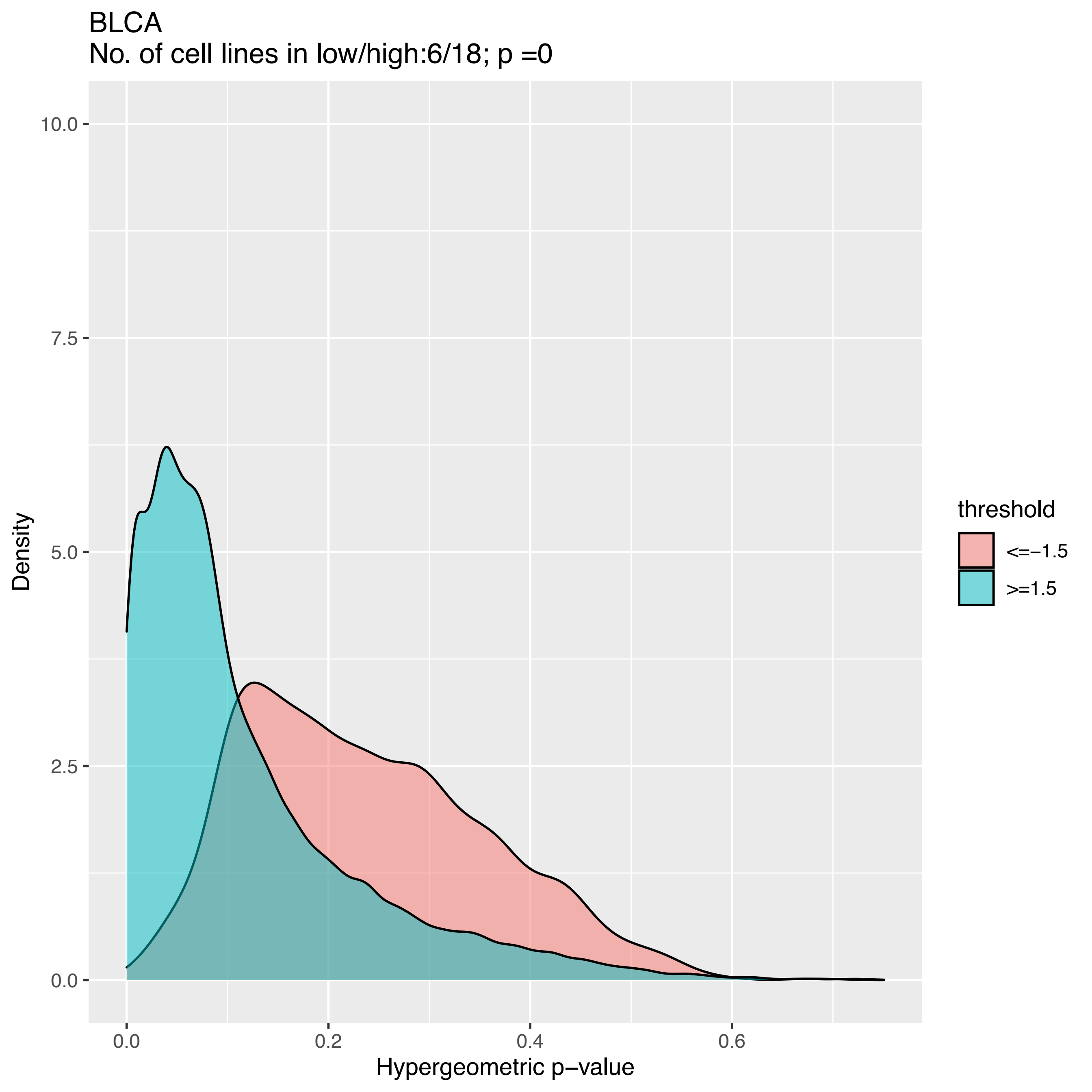
BLCA
No. of cell lines in low/high:6/18; p =0
Density
10.0
7.5
5.0
2.5
0.0
0.0
0.2
0.4
0.6
Hypergeometric p−value
threshold
<=−1.5
>=1.5

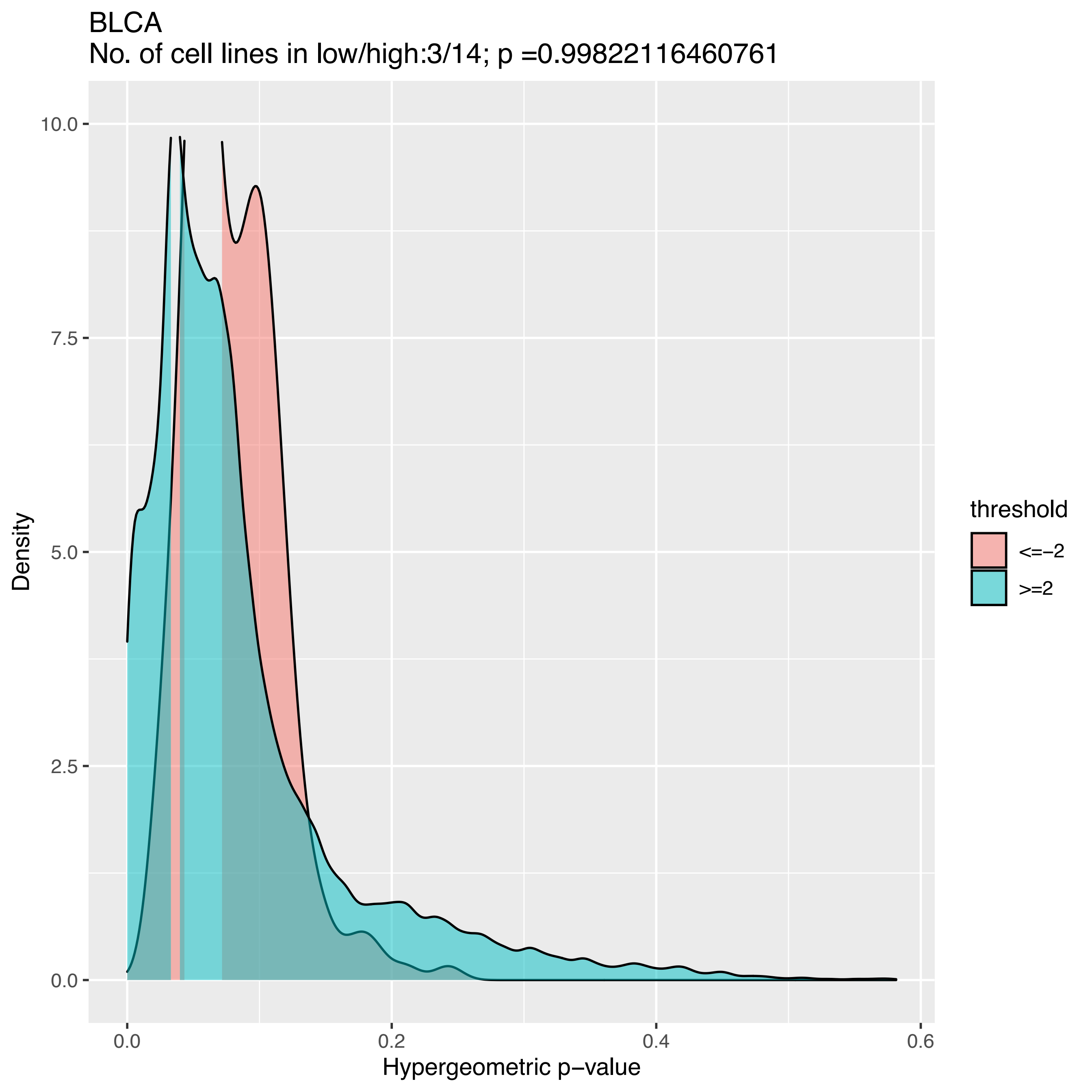

BLCA
No. of cell lines in low/high:3/14; p =0.99822116460761
Density
Hypergeometric p−value
10.0
7.5
5.0
2.5
0.0
0.0
0.2
0.4
0.6
threshold
<=−2
>=2

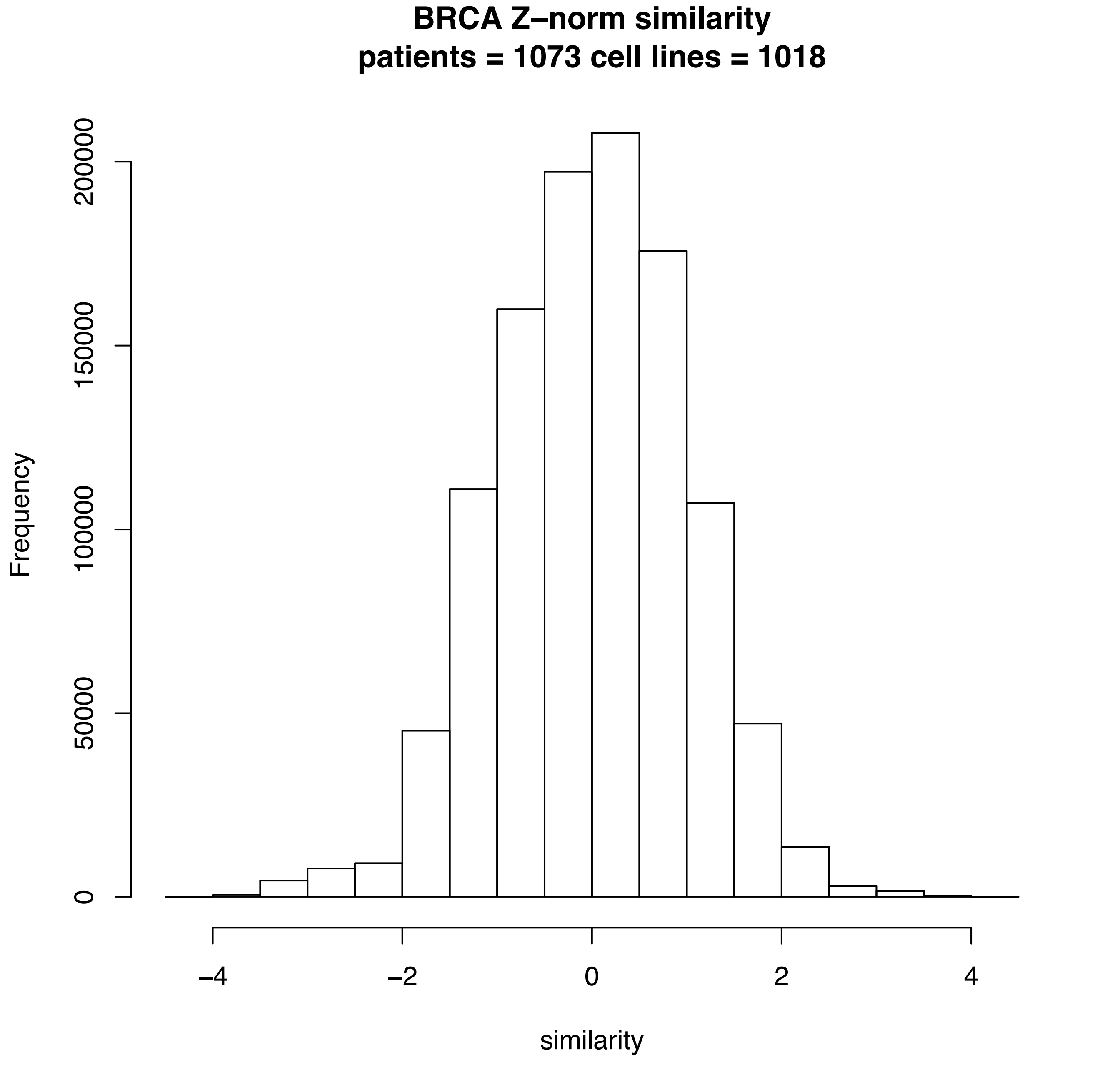
BRCA Z–norm similarity
patients = 1073 cell lines = 1018
Frequency
0
50000
100000
150000
200000
−4
−2
0
2
4
similarity

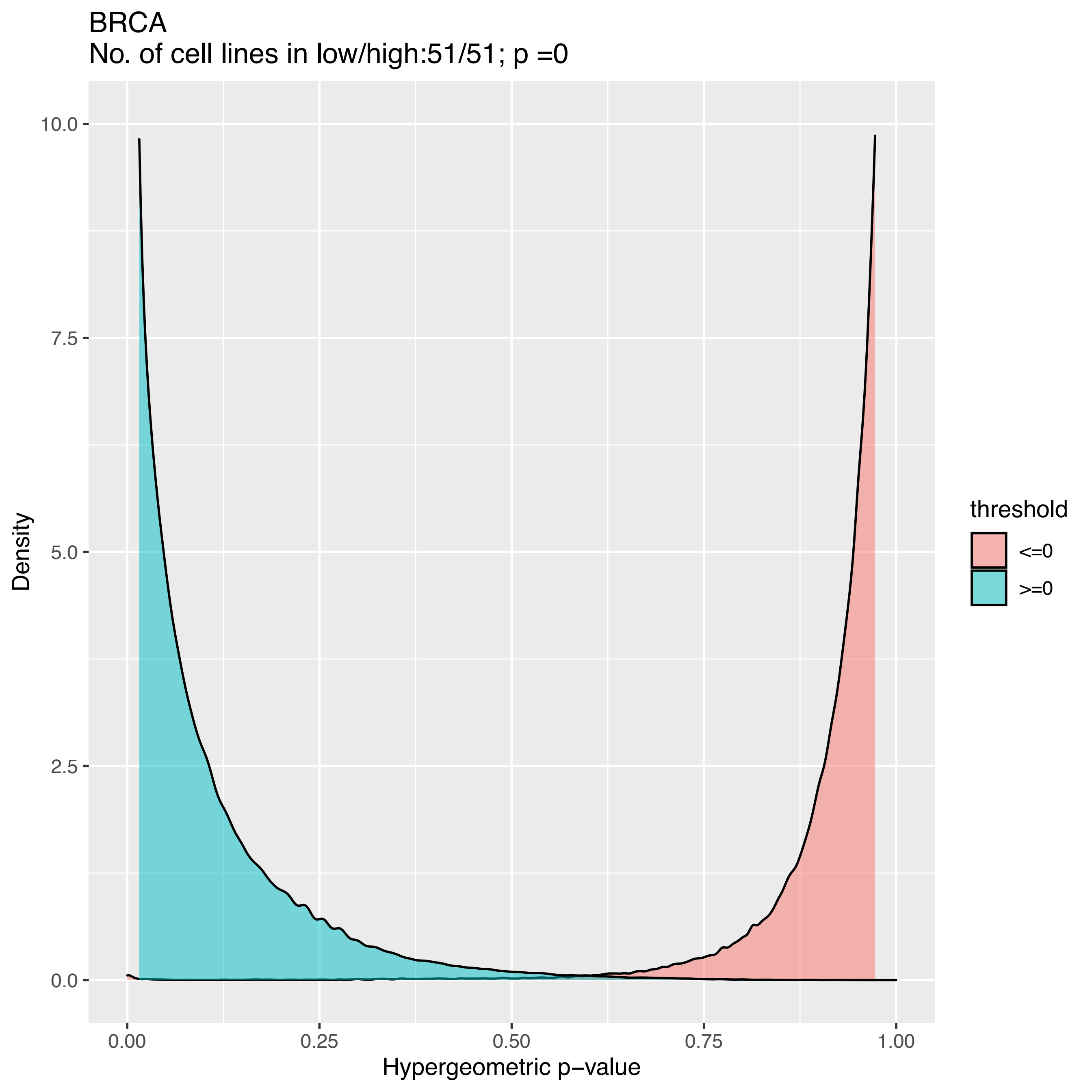
BRCA
No. of cell lines in low/high:51/51; p =0
Density
10.0
7.5
5.0
2.5
0.0
0.00
0.25
0.50
0.75
1.00
Hypergeometric p−value
threshold
<=0
>=0

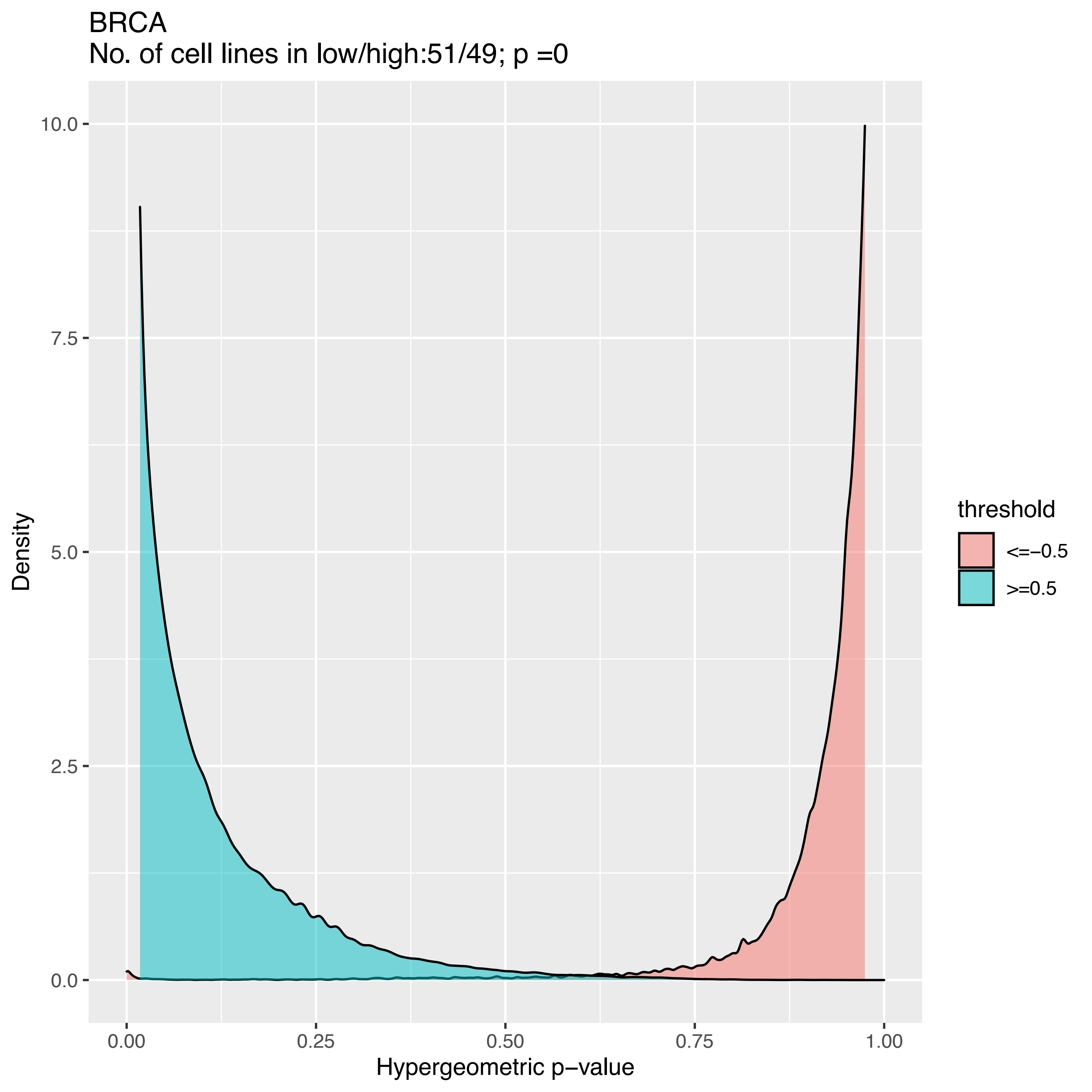
BRCA
No. of cell lines in low/high:51/49; p =0
10.0
7.5
5.0
2.5
0.0
Density
0.00
0.25
0.50
0.75
1.00
Hypergeometric p−value
threshold
<=−0.5
>=0.5

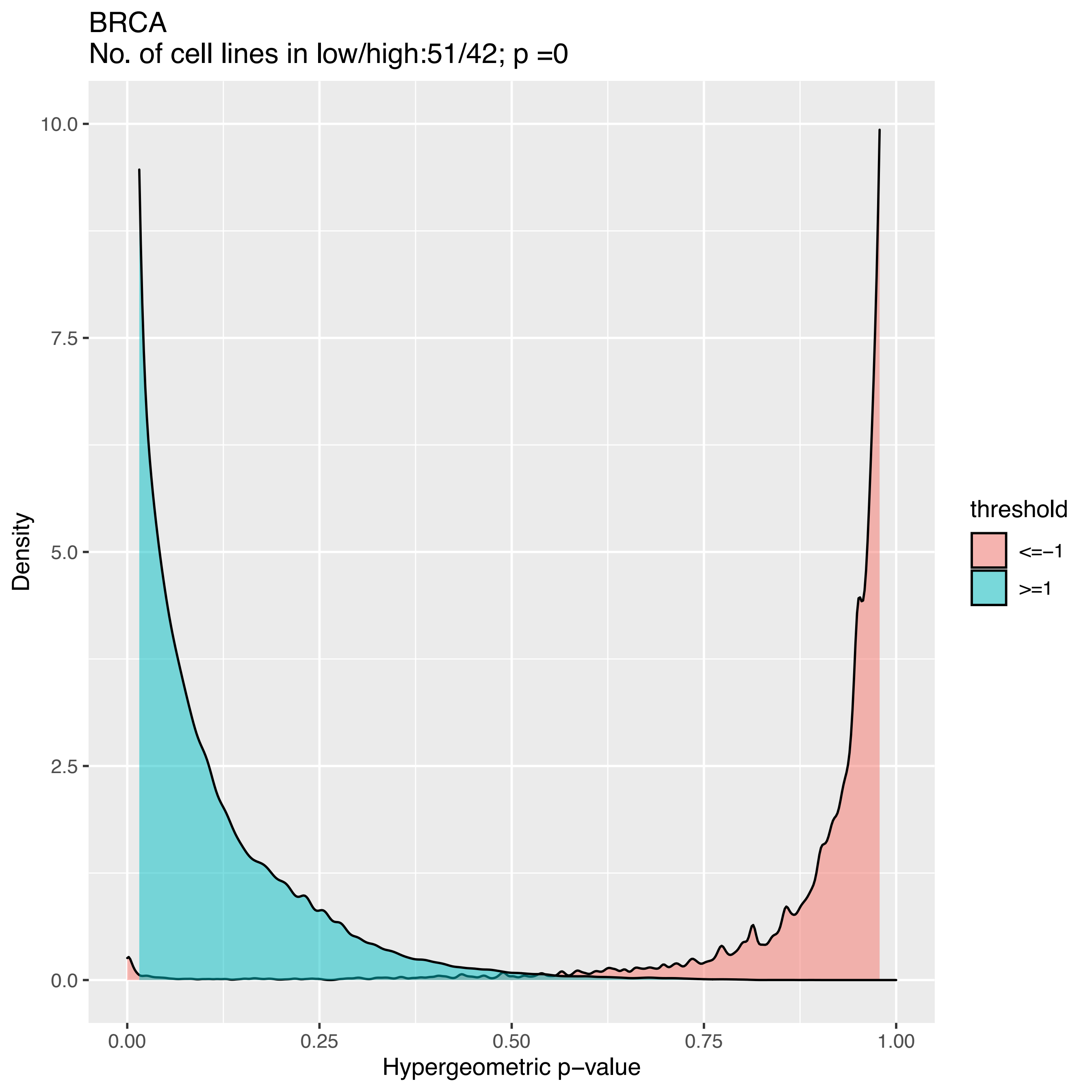
BRCA
No. of cell lines in low/high:51/42; p =0
Density
Hypergeometric p−value
0.0
2.5
5.0
7.5
10.0
0.00
0.25
0.50
0.75
1.00
threshold
<=−1
>=1

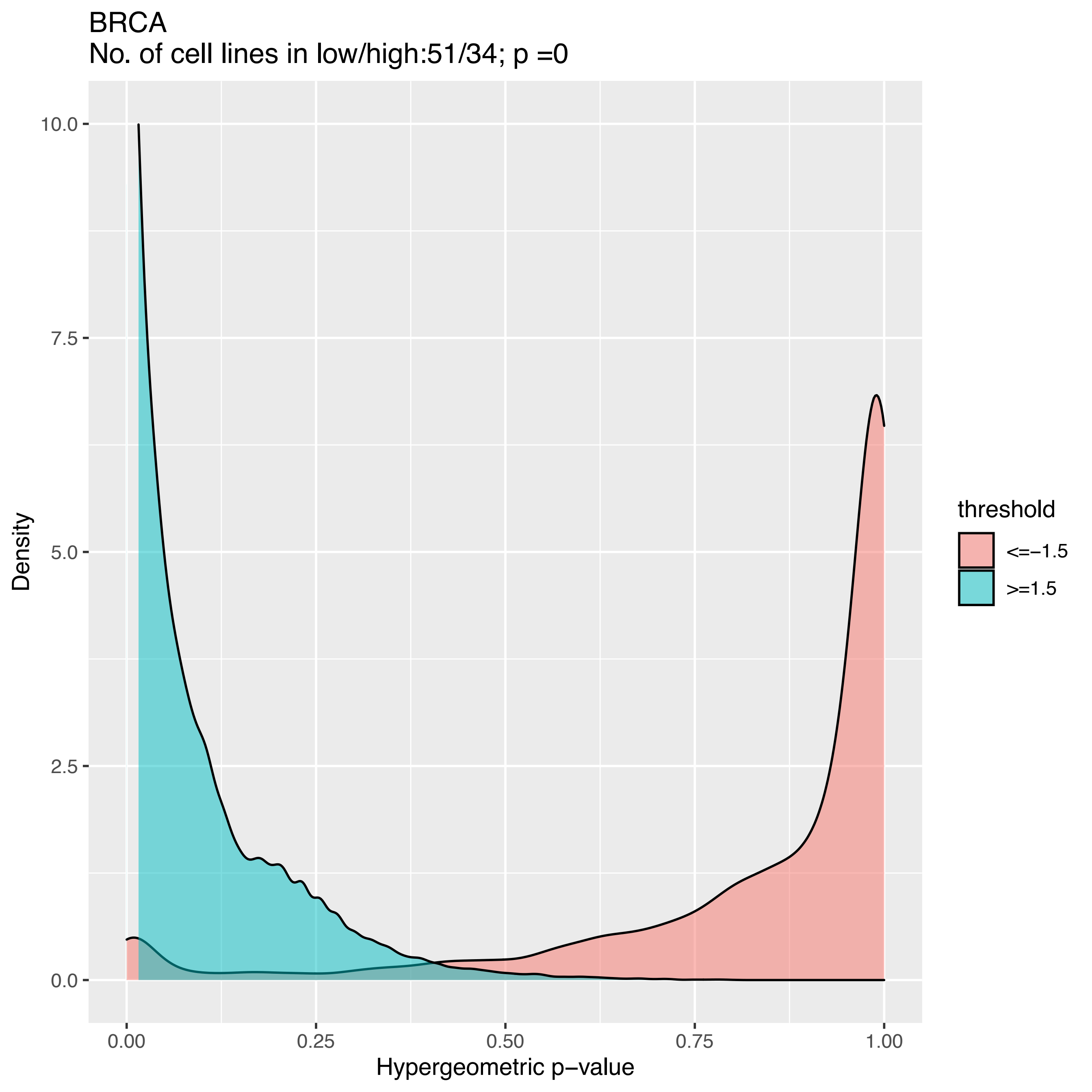
BRCA
No. of cell lines in low/high:51/34; p =0
10.0
7.5
5.0
2.5
0.0
Density
0.00
0.25
0.50
0.75
1.00
Hypergeometric p−value
threshold
<=−1.5
>=1.5

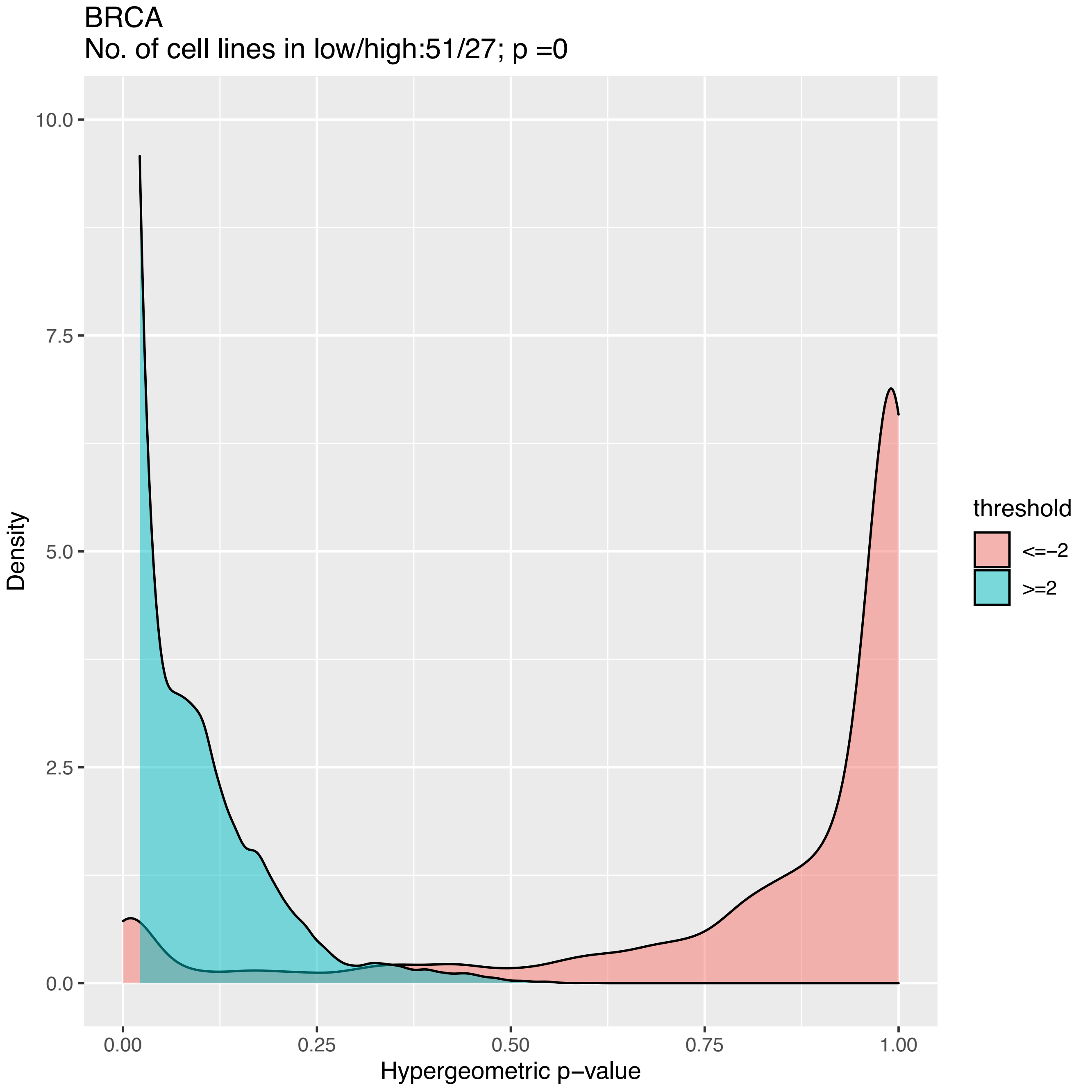
BRCA
No. of cell lines in low/high:51/27; p =0
10.0
7.5
5.0
2.5
0.0
Density
0.00
0.25
0.50
0.75
1.00
Hypergeometric p−value
threshold
<=−2
>=2

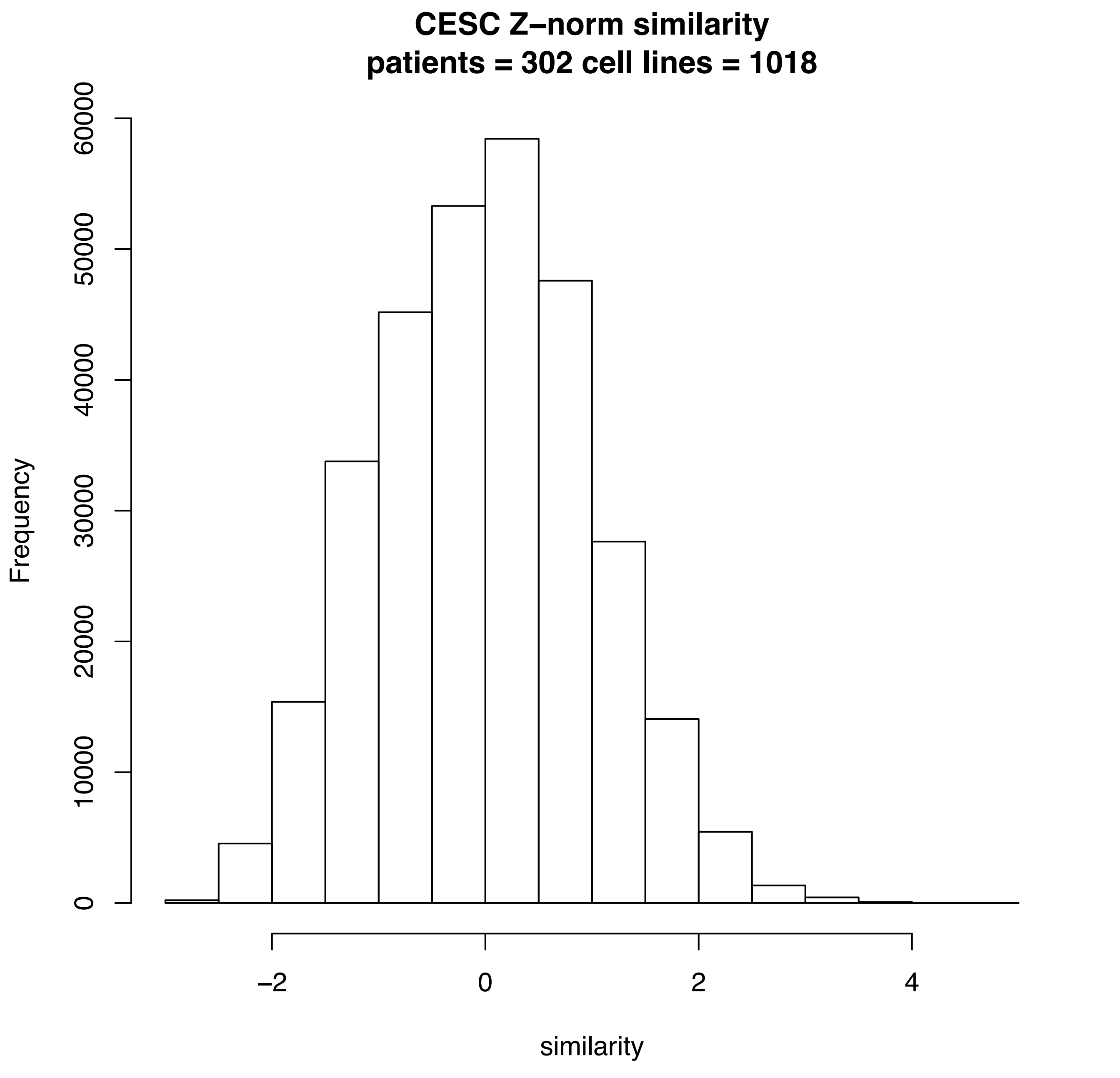
CESC Z–norm similarity
patients = 302 cell lines = 1018
Frequency
0
10000
20000
30000
40000
50000
60000
–2
0
2
4
similarity

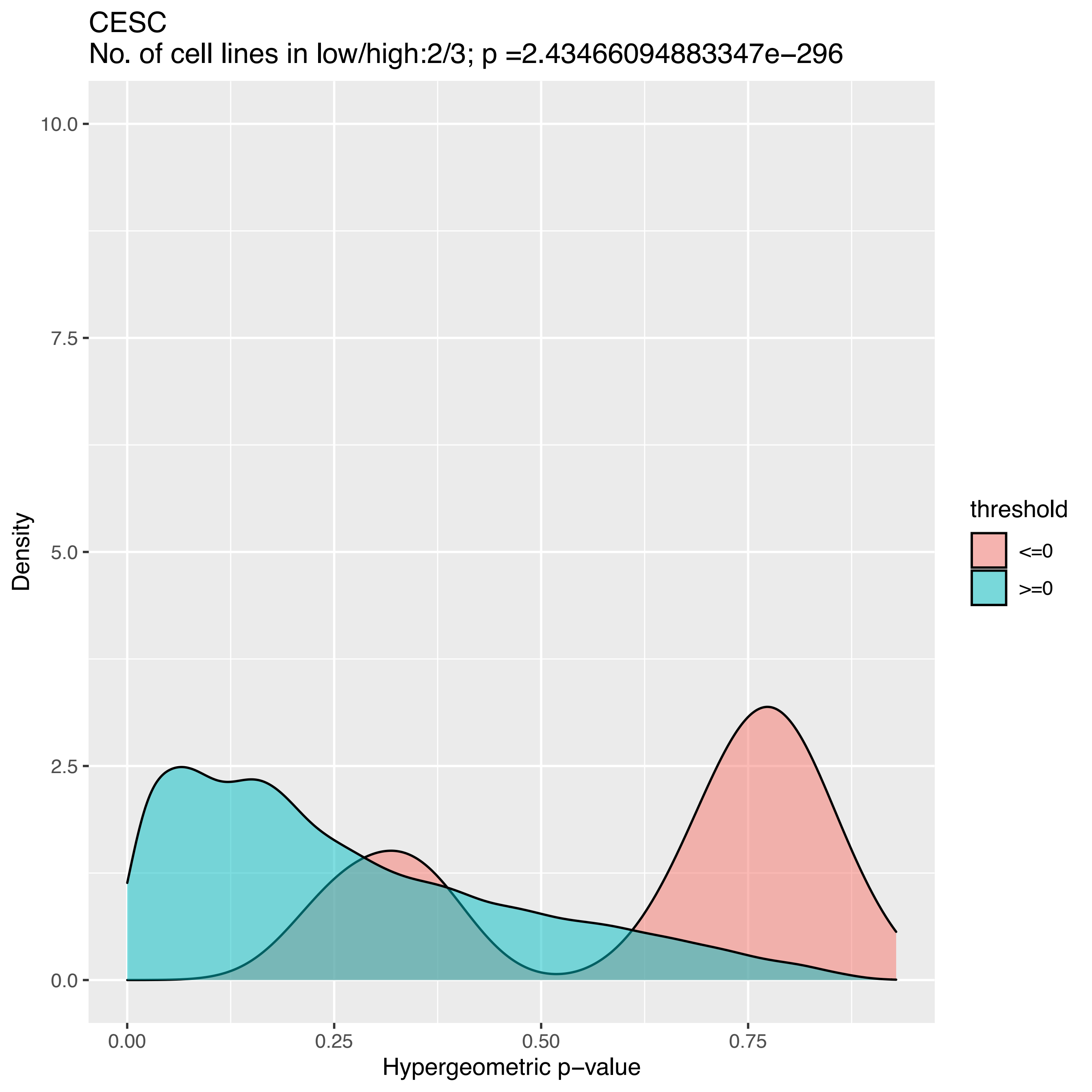
CESC
No. of cell lines in low/high:2/3; p =2.43466094883347e−296
10.0
7.5
5.0
2.5
0.0
Density
0.00
0.25
0.50
0.75
Hypergeometric p−value
threshold
<=0
>=0

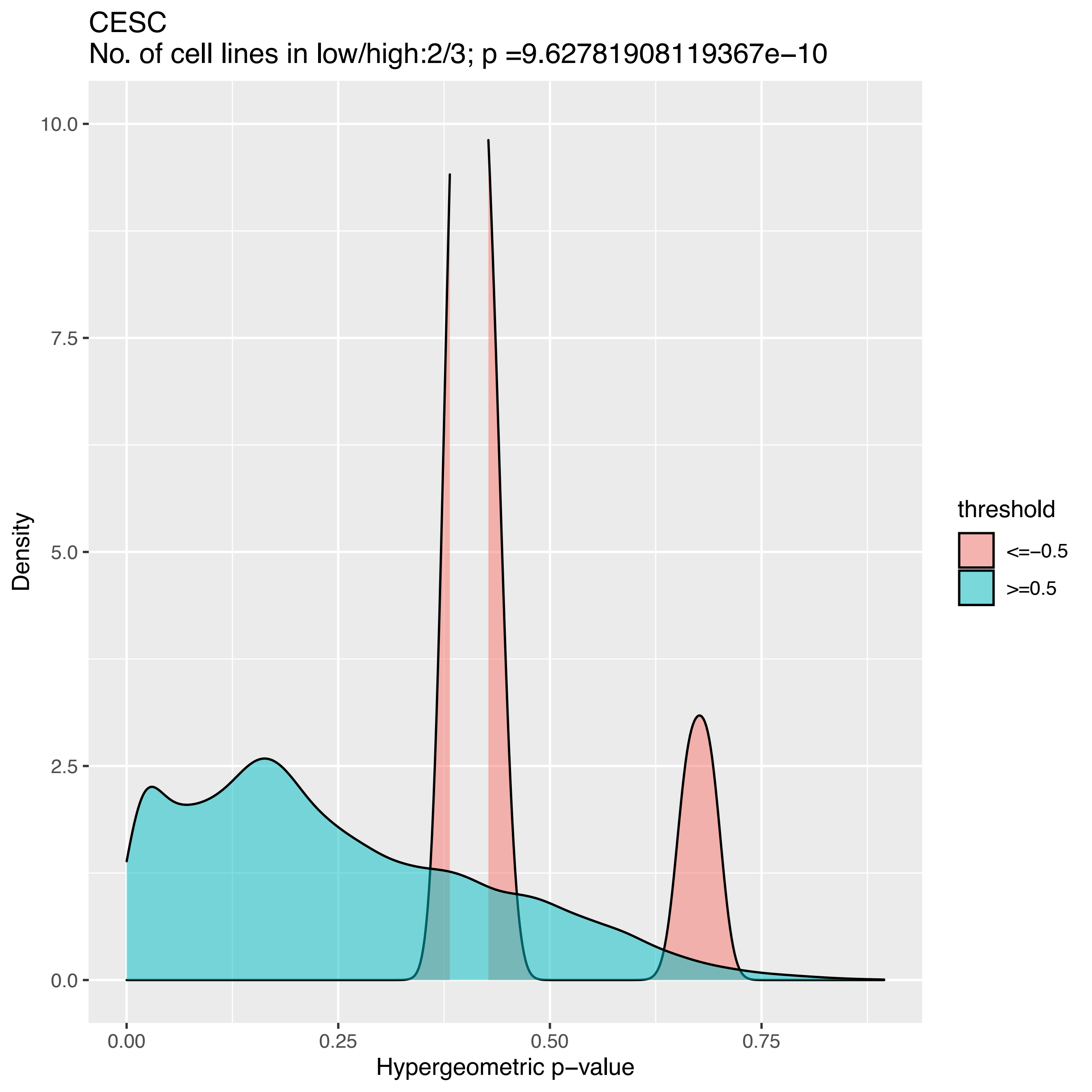
CESC
No. of cell lines in low/high:2/3; p =9.62781908119367e−10
Density
Hypergeometric p−value
10.0
7.5
5.0
2.5
0.0
0.00
0.25
0.50
0.75
threshold
<=−0.5
>=0.5

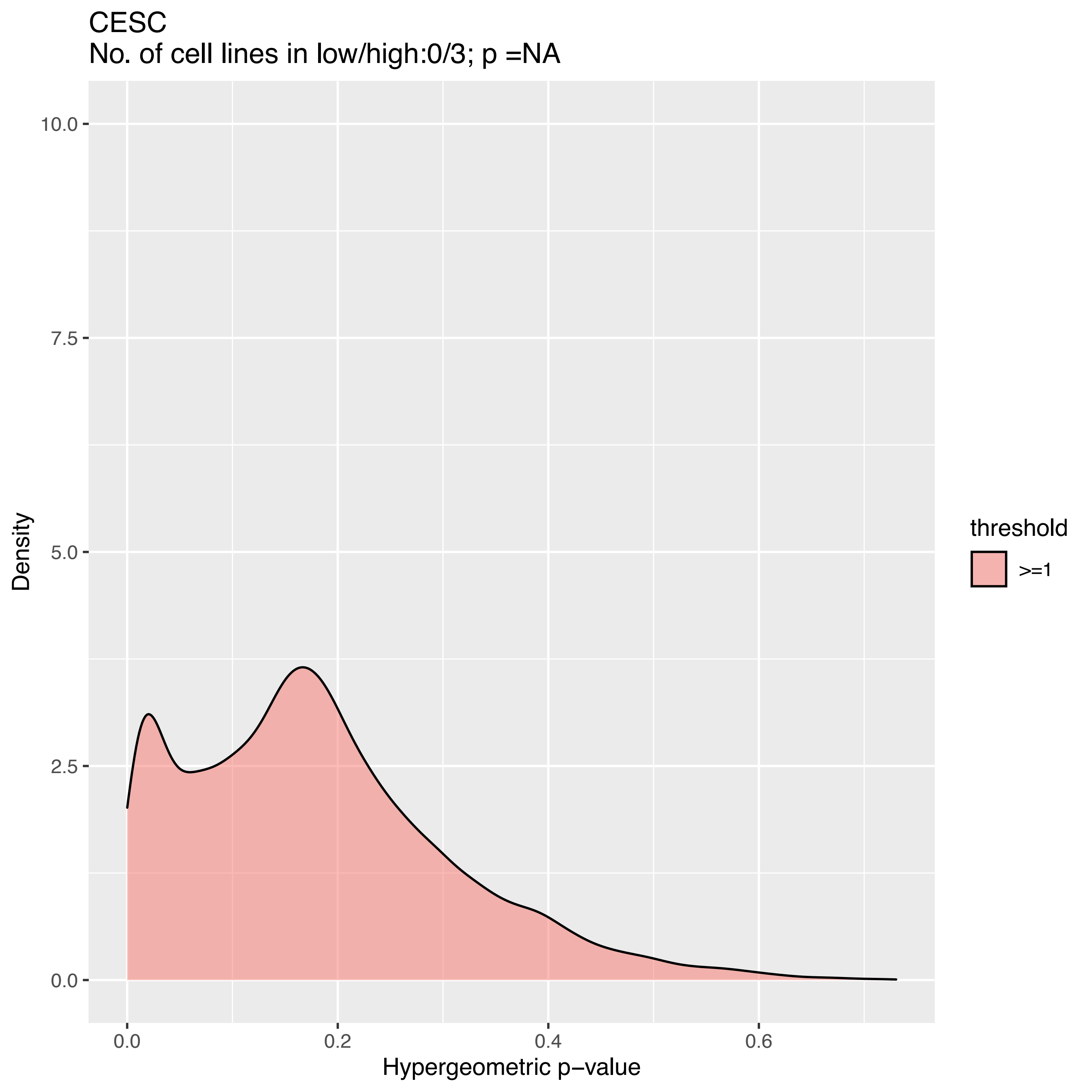
CESC
No. of cell lines in low/high:0/3; p =NA
Density
10.0
7.5
5.0
2.5
0.0
0.0
0.2
0.4
0.6
Hypergeometric p−value
threshold
>=1

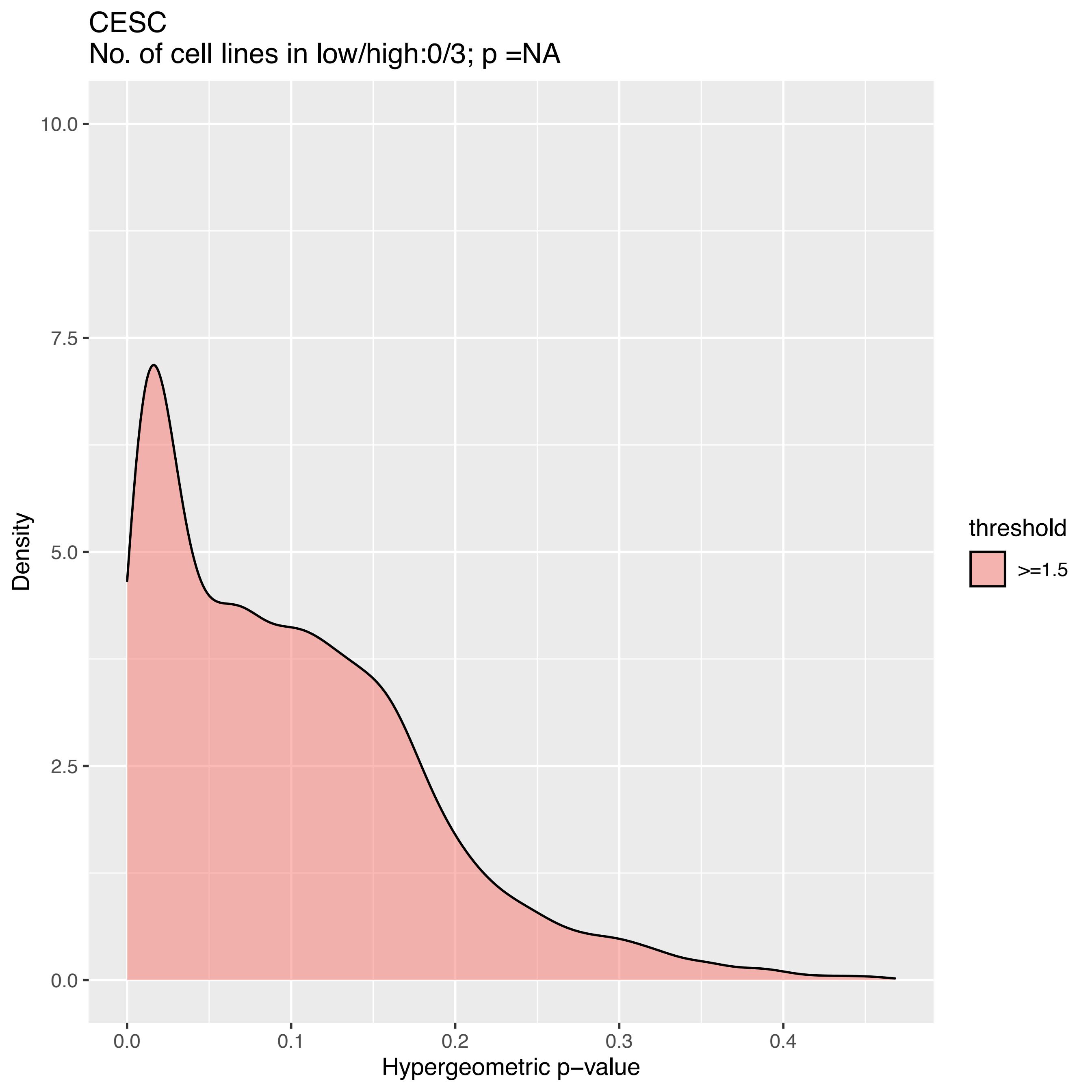
CESC
No. of cell lines in low/high:0/3; p =NA
10.0
7.5
5.0
2.5
0.0
Density
0.0
0.1
0.2
0.3
0.4
Hypergeometric p−value
threshold
>=1.5

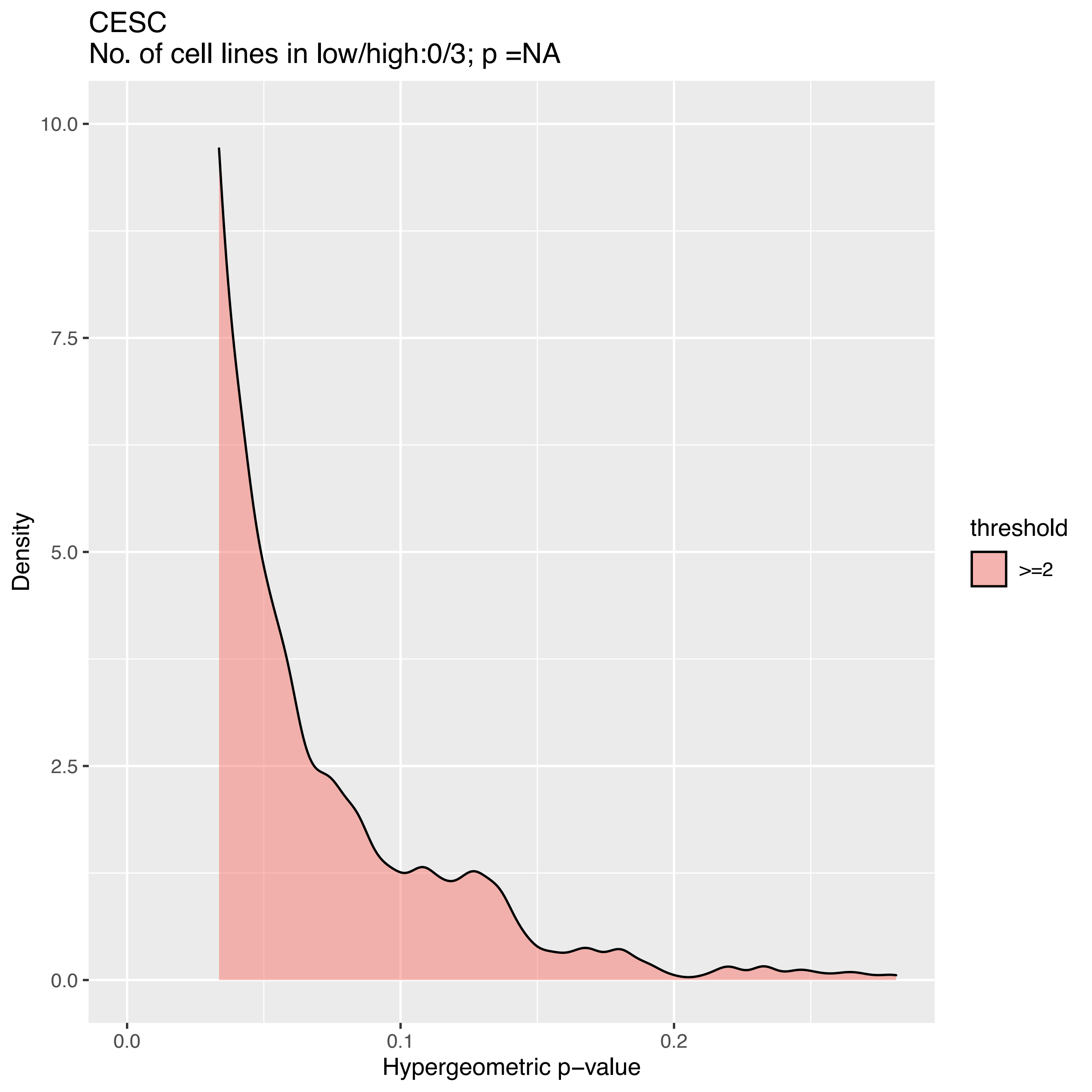

CESC
No. of cell lines in low/high:0/3; p =NA
10.0
7.5
5.0
2.5
0.0
Density
0.0
0.1
0.2
Hypergeometric p−value
threshold
>=2

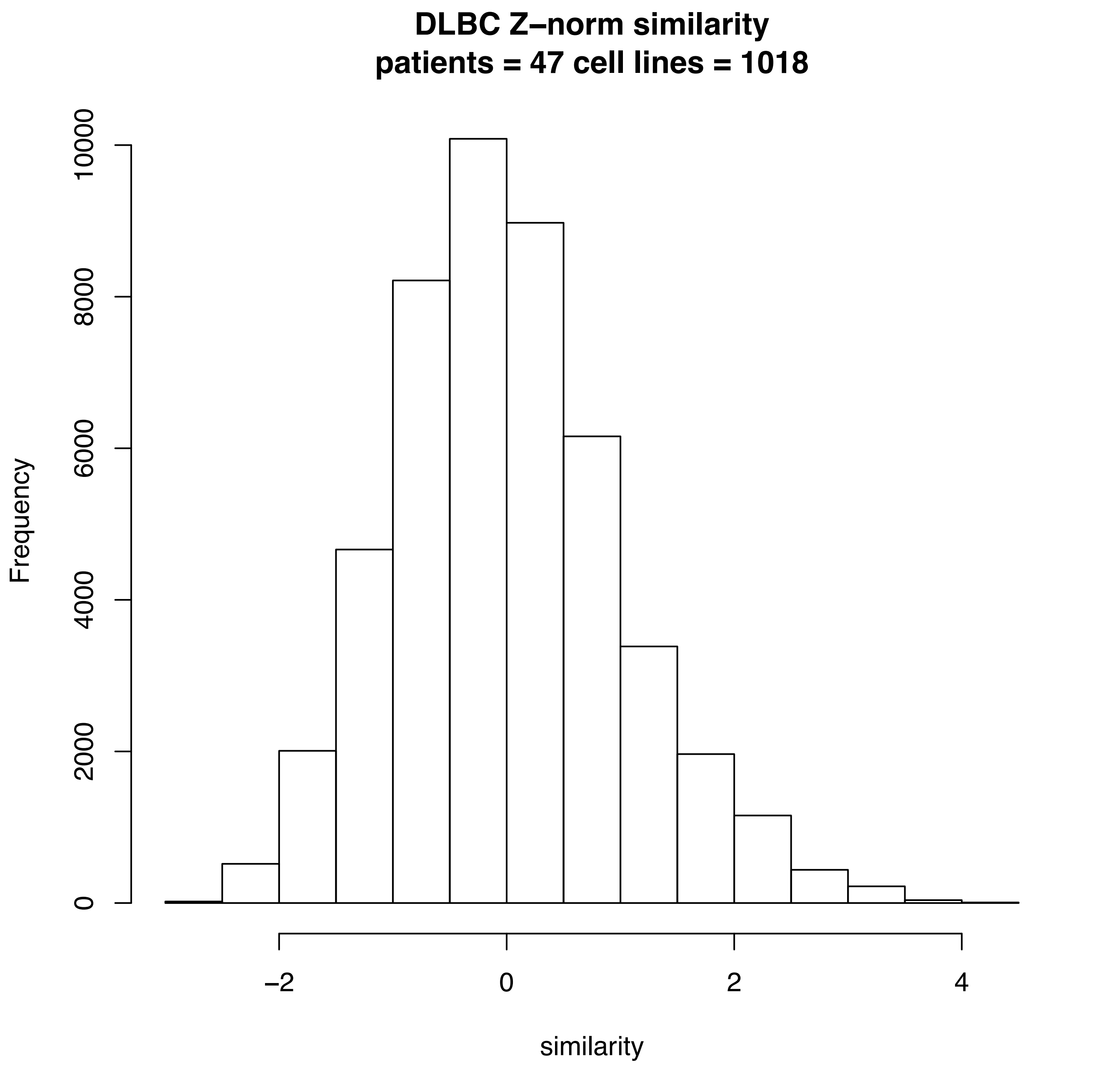
DLBC Z–norm similarity
patients = 47 cell lines = 1018
Frequency
0
2000
4000
6000
8000
10000
−2
0
2
4
similarity

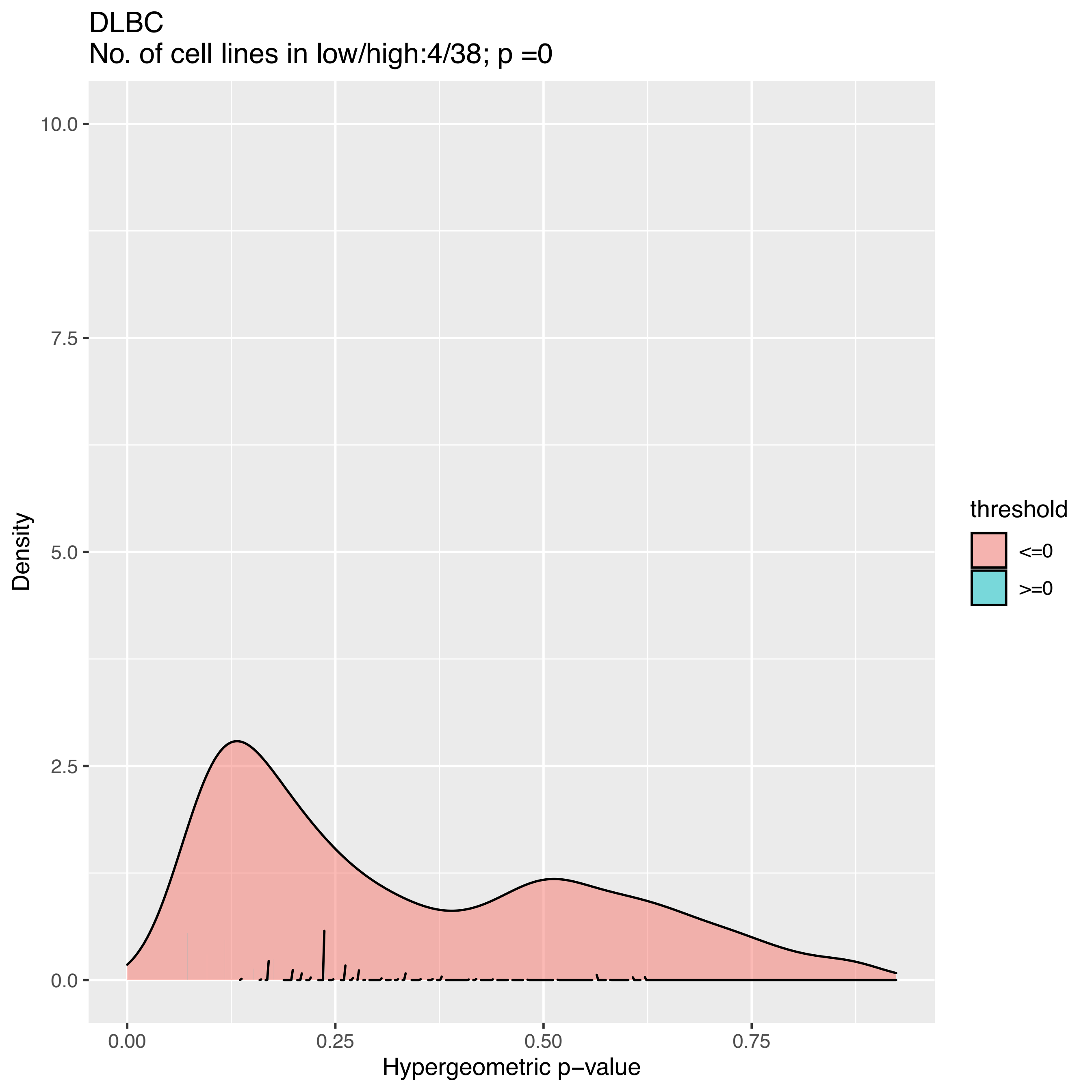

DLBC
No. of cell lines in low/high:4/38; p =0
Density
10.0
7.5
5.0
2.5
0.0
0.00
0.25
0.50
0.75
Hypergeometric p−value
threshold
<=0
>=0

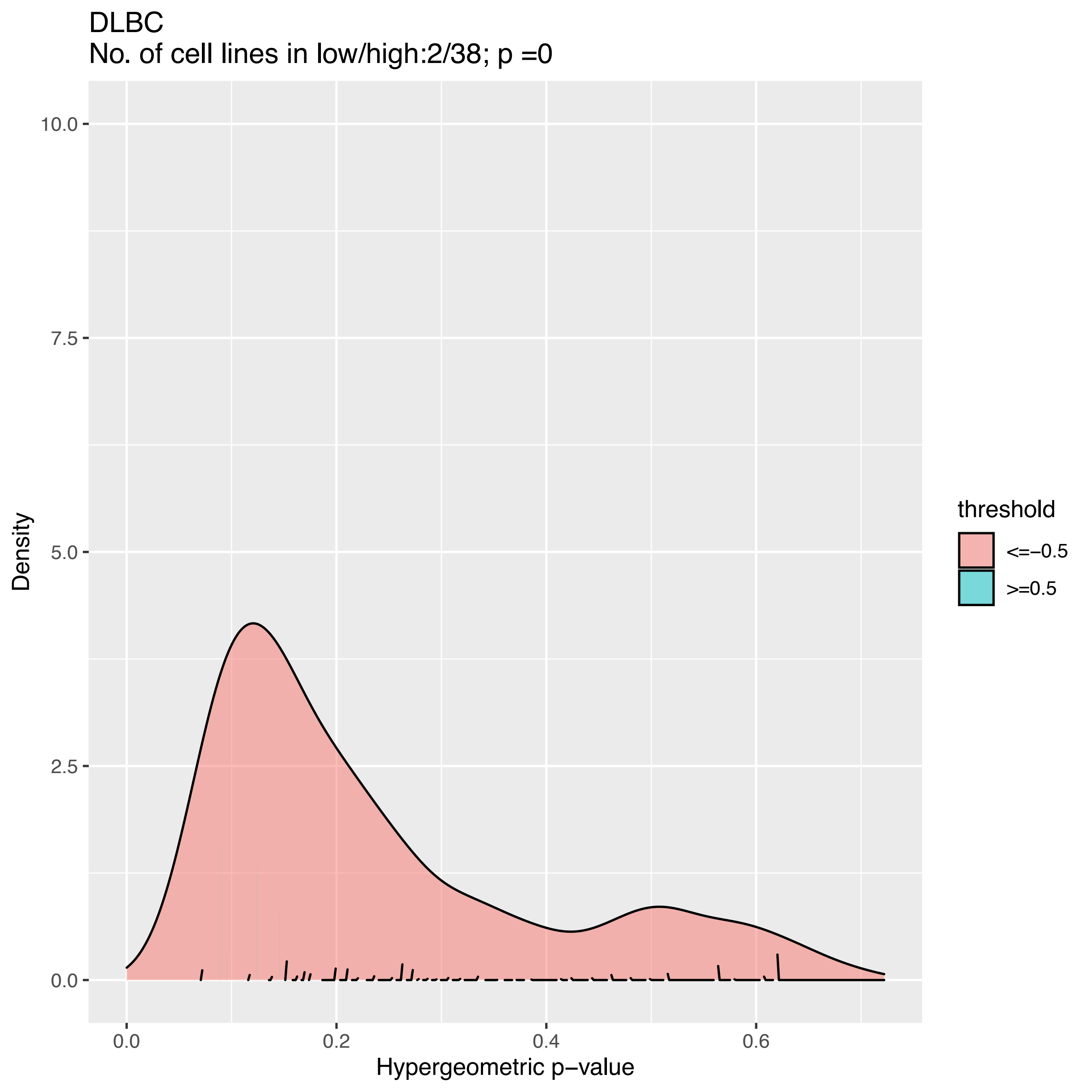
DLBC
No. of cell lines in low/high:2/38; p =0
Density
10.0
7.5
5.0
2.5
0.0
0.0
0.2
0.4
0.6
Hypergeometric p−value
threshold
<=−0.5
>=0.5

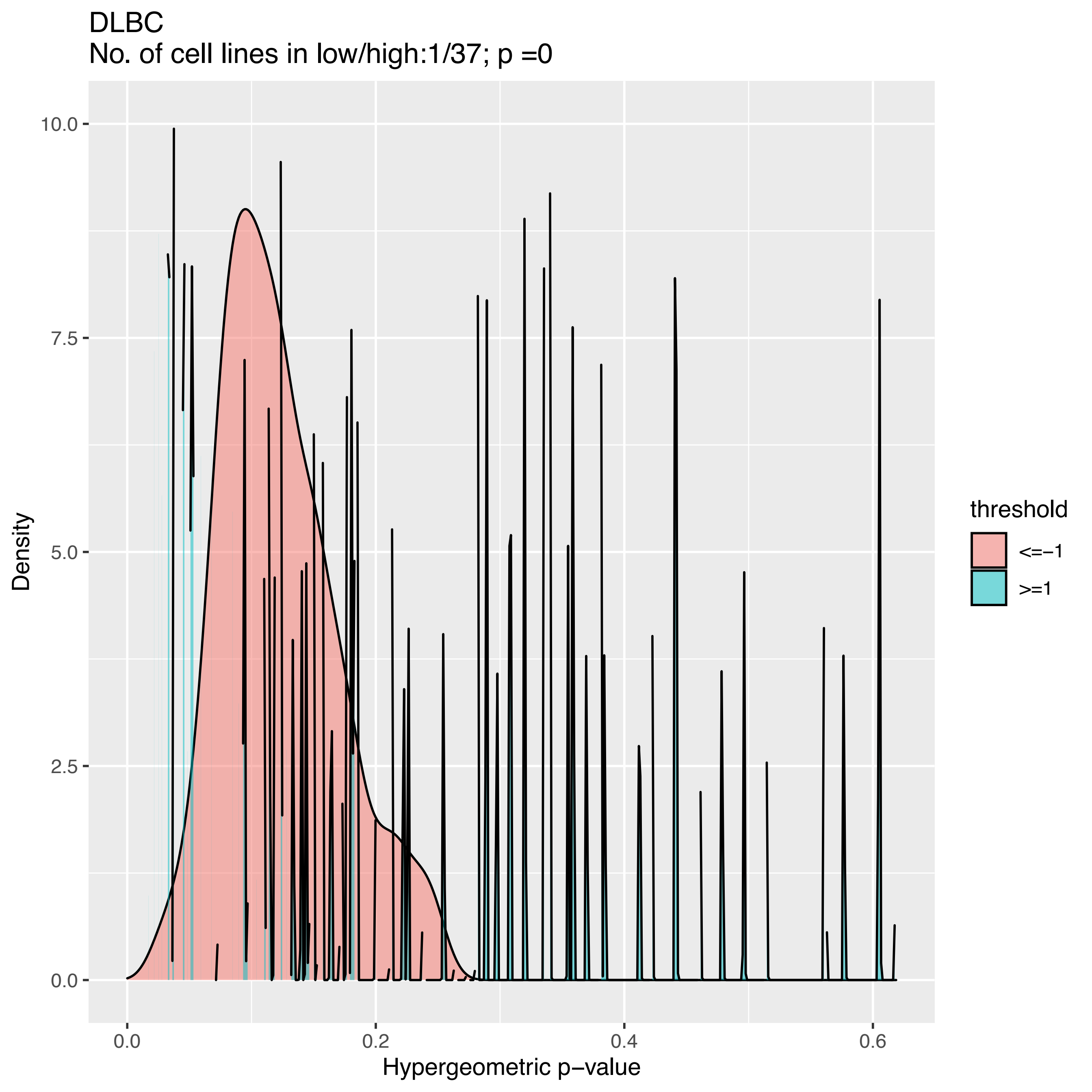
DLBC
No. of cell lines in low/high:1/37; p =0
Density
Hypergeometric p−value
threshold
<=−1
>=1
0.0
2.5
5.0
7.5
10.0
0.0
0.2
0.4
0.6

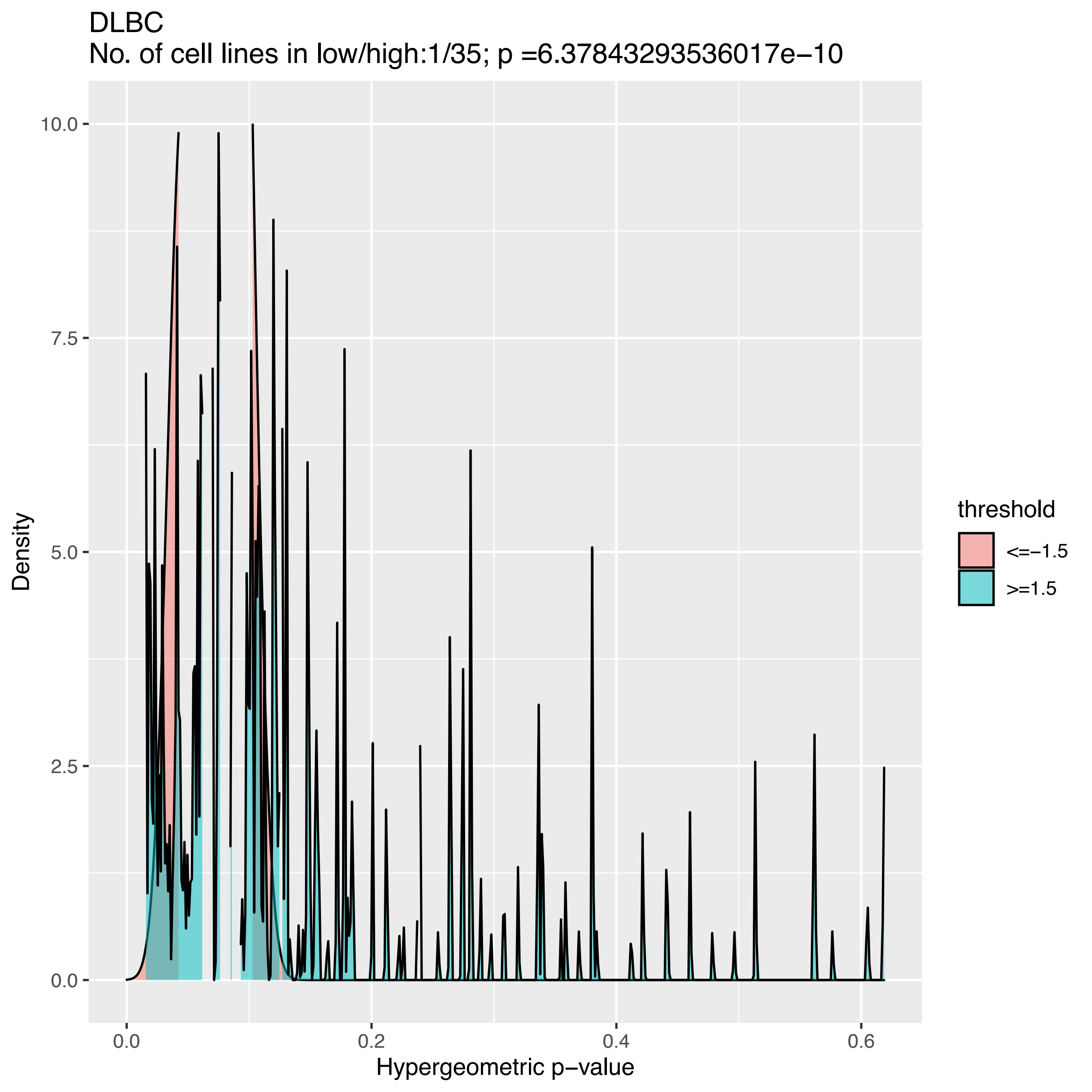

DLBC
No. of cell lines in low/high:1/35; p =6.37843293536017e−10
Density
Hypergeometric p−value
0.0
2.5
5.0
7.5
10.0
0.0
0.2
0.4
0.6
threshold
<=−1.5
>=1.5

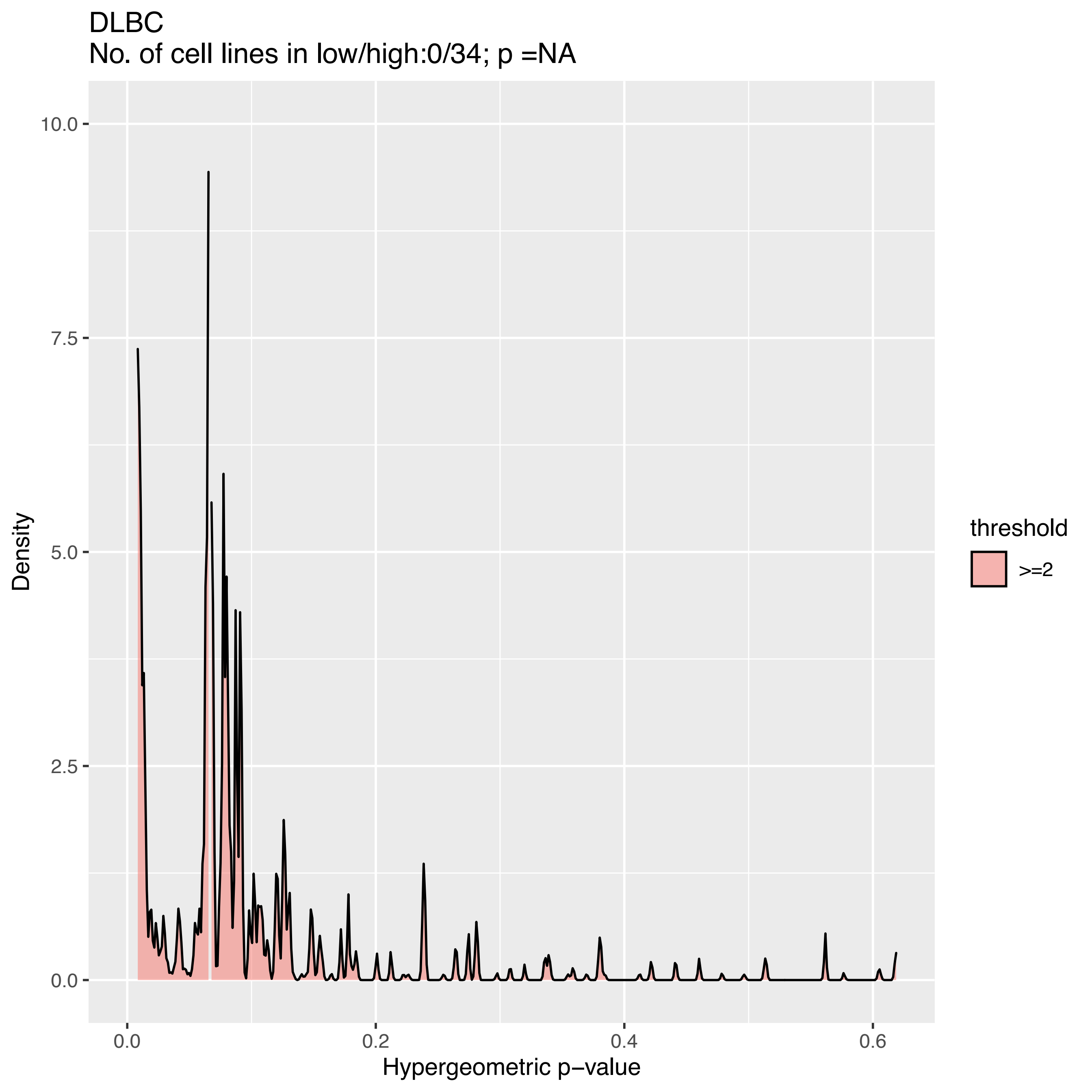
DLBC
No. of cell lines in low/high:0/34; p =NA
10.0
7.5
5.0
2.5
0.0
Density
0.0
0.2
0.4
0.6
Hypergeometric p−value
threshold
>=2

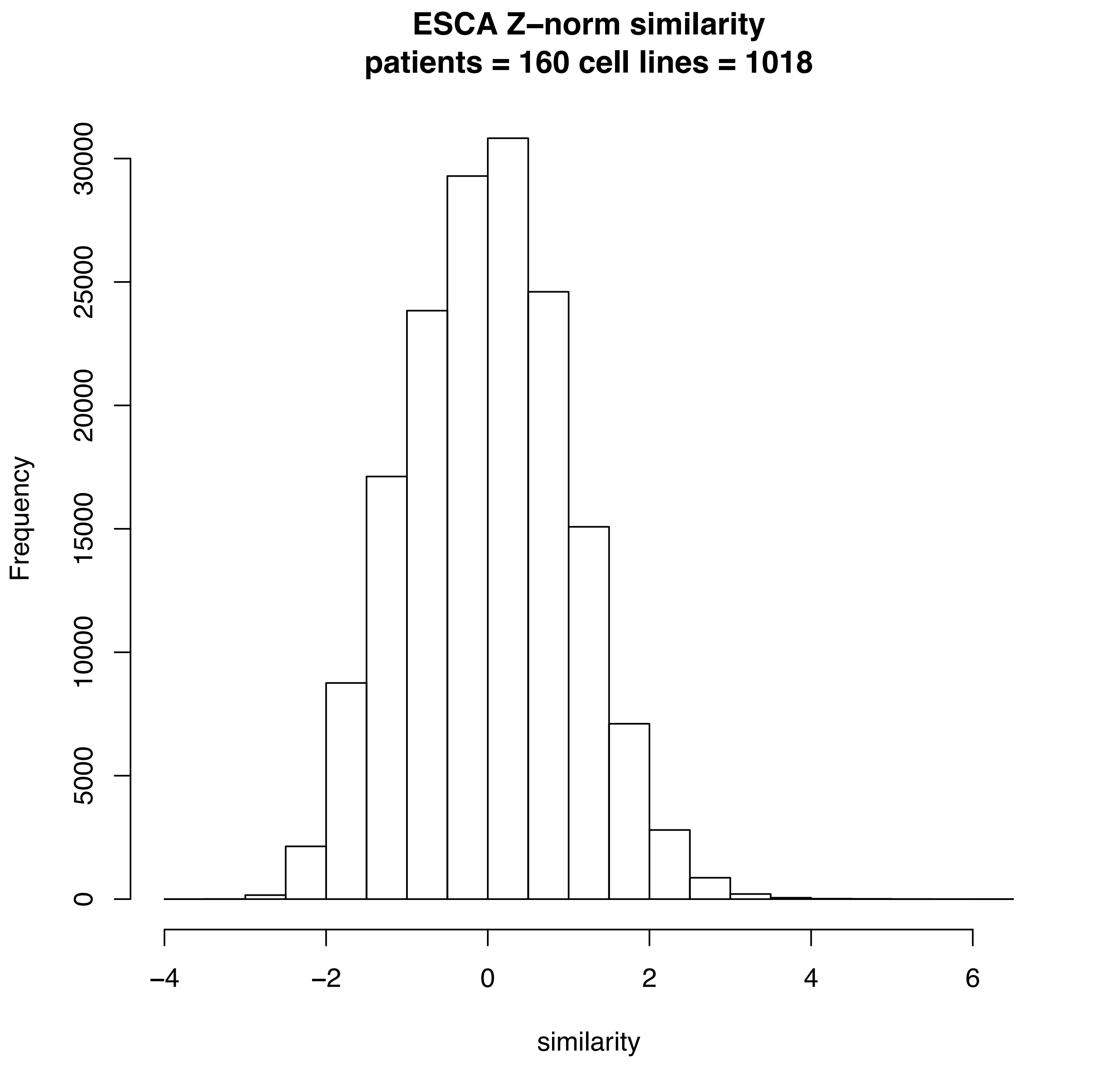
ESCA Z–norm similarity
patients = 160 cell lines = 1018
Frequency
0
5000
10000
15000
20000
25000
30000
–4
–2
0
2
4
6
similarity

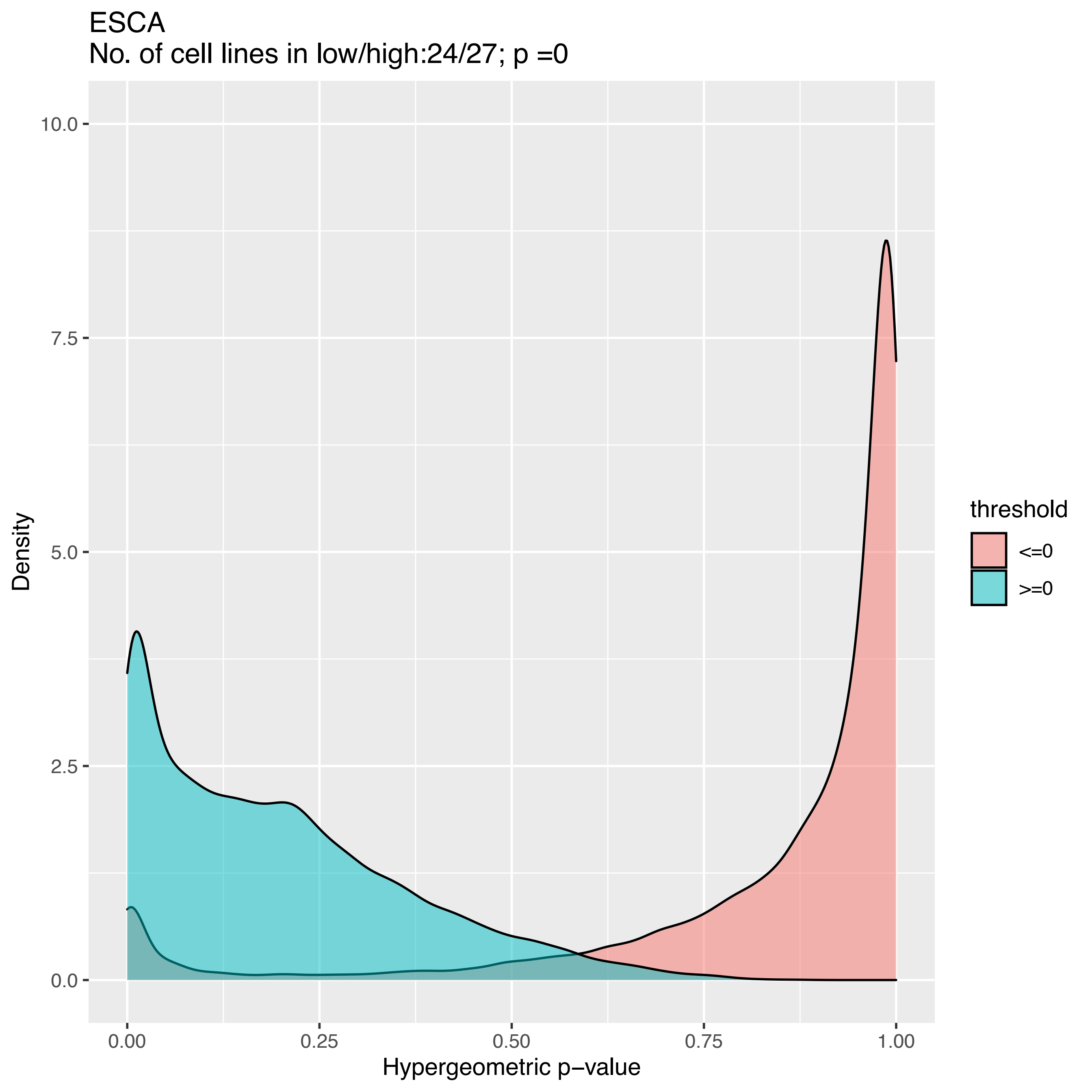

ESCA
No. of cell lines in low/high:24/27; p =0
10.0
7.5
5.0
2.5
0.0
Density
0.00
0.25
0.50
0.75
1.00
Hypergeometric p−value
threshold
<=0
>=0

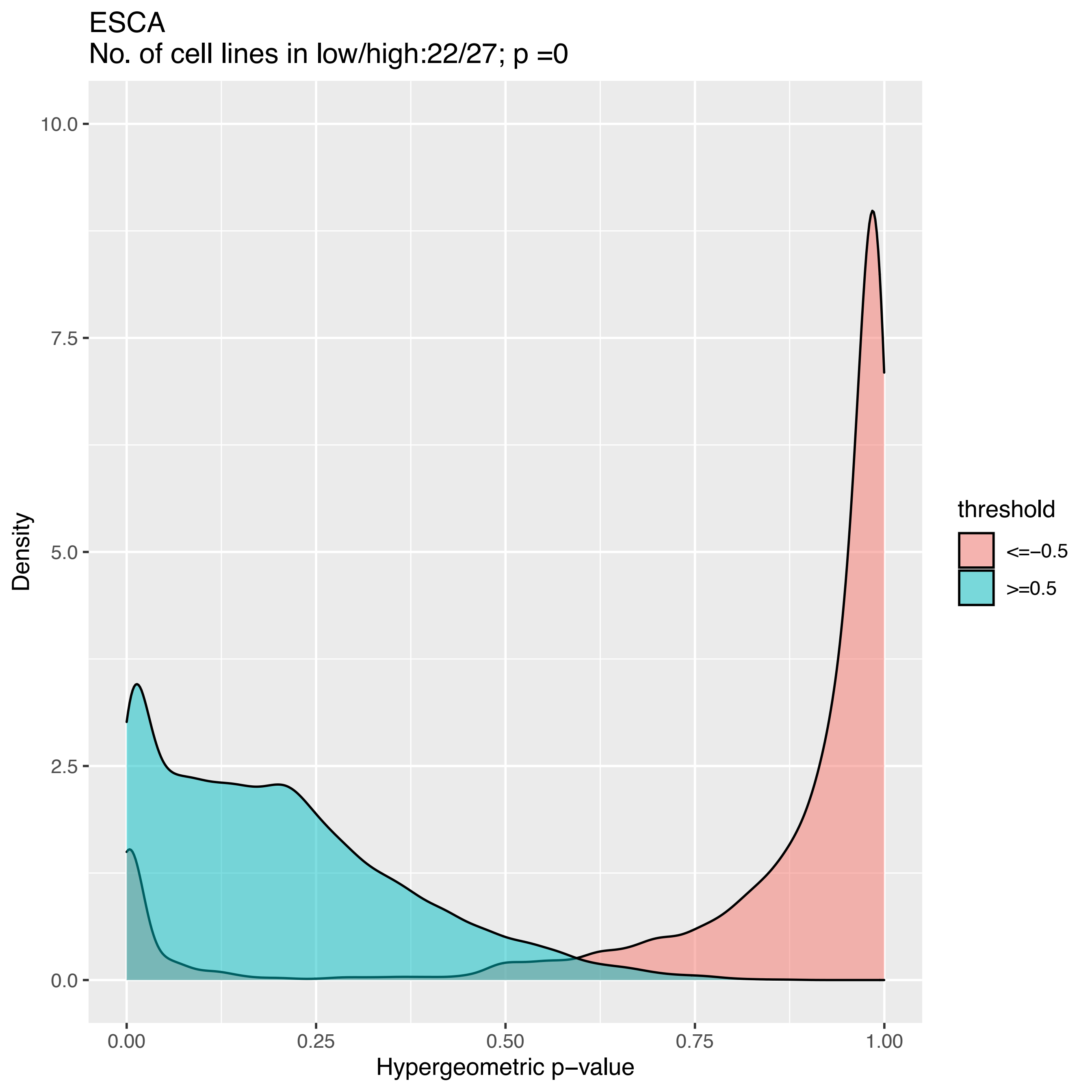

ESCA
No. of cell lines in low/high:22/27; p =0
Density
10.0
7.5
5.0
2.5
0.0
0.00
0.25
0.50
0.75
1.00
Hypergeometric p−value
threshold
<=−0.5
>=0.5

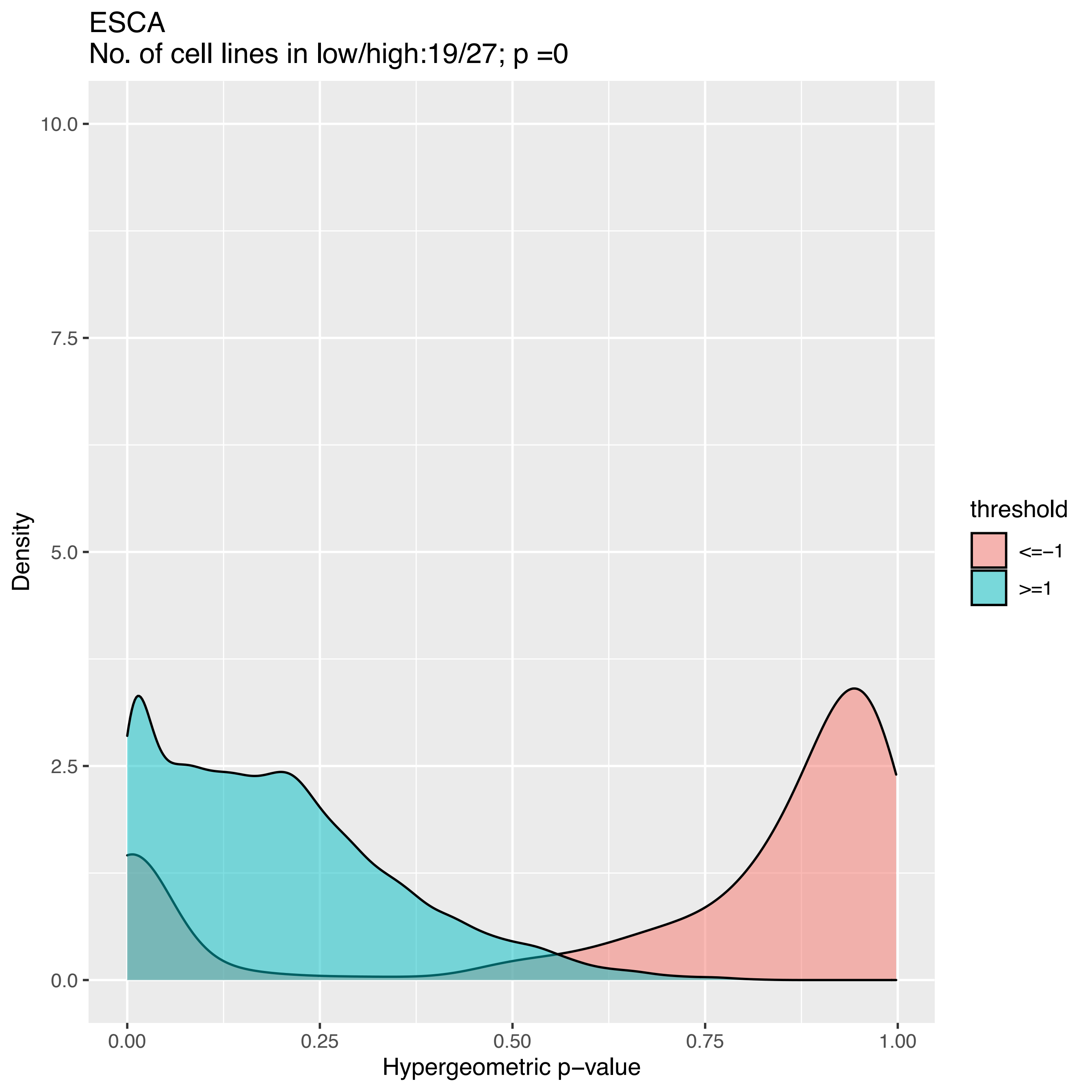

ESCA
No. of cell lines in low/high:19/27; p =0
10.0
7.5
5.0
2.5
0.0
Density
0.00
0.25
0.50
0.75
1.00
Hypergeometric p−value
threshold
<=−1
>=1

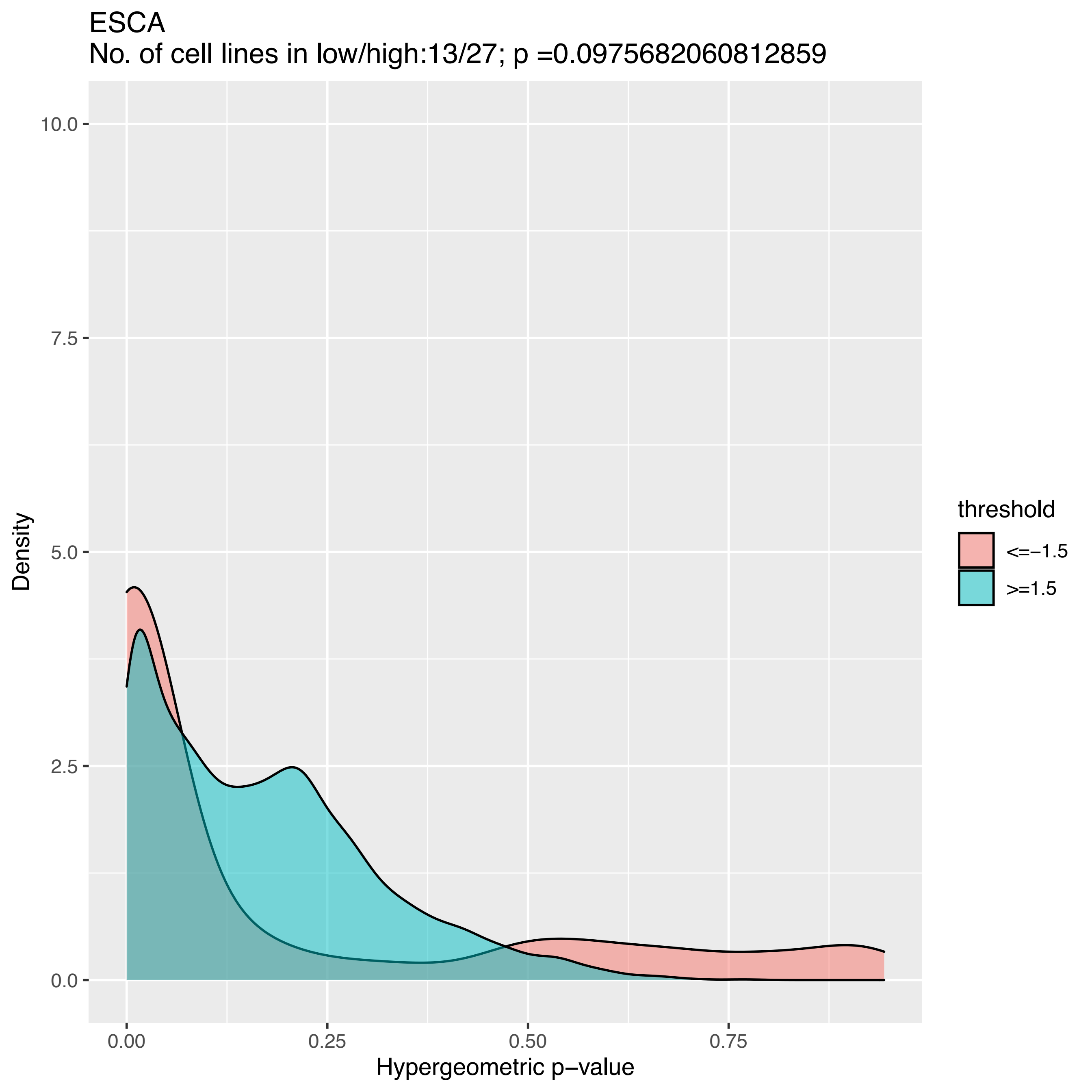

ESCA
No. of cell lines in low/high:13/27; p =0.0975682060812859
Density
10.0
7.5
5.0
2.5
0.0
0.00
0.25
0.50
0.75
Hypergeometric p−value
threshold
<=−1.5
>=1.5

ESCA
No. of cell lines in low/high:9/25; p =1

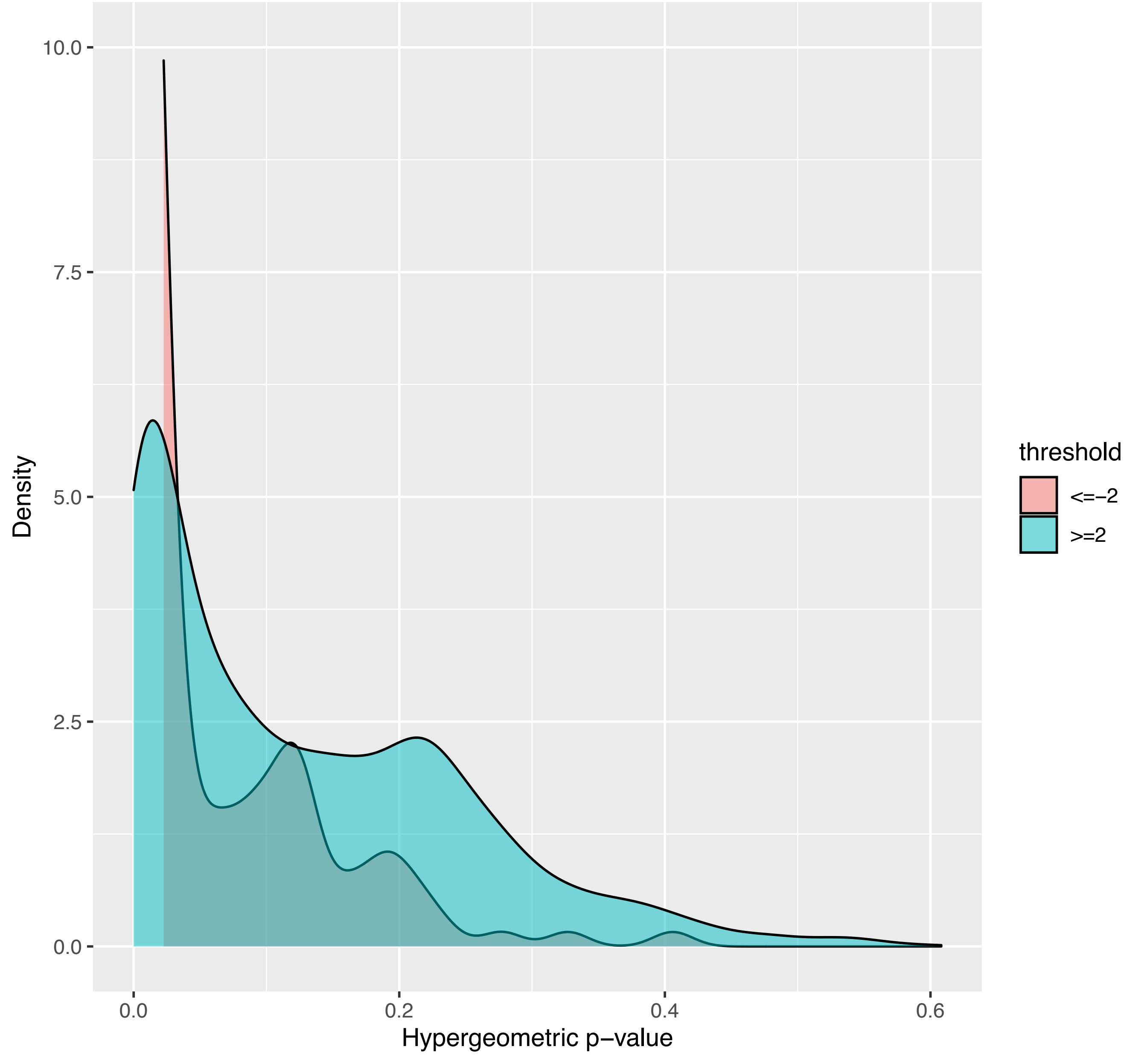

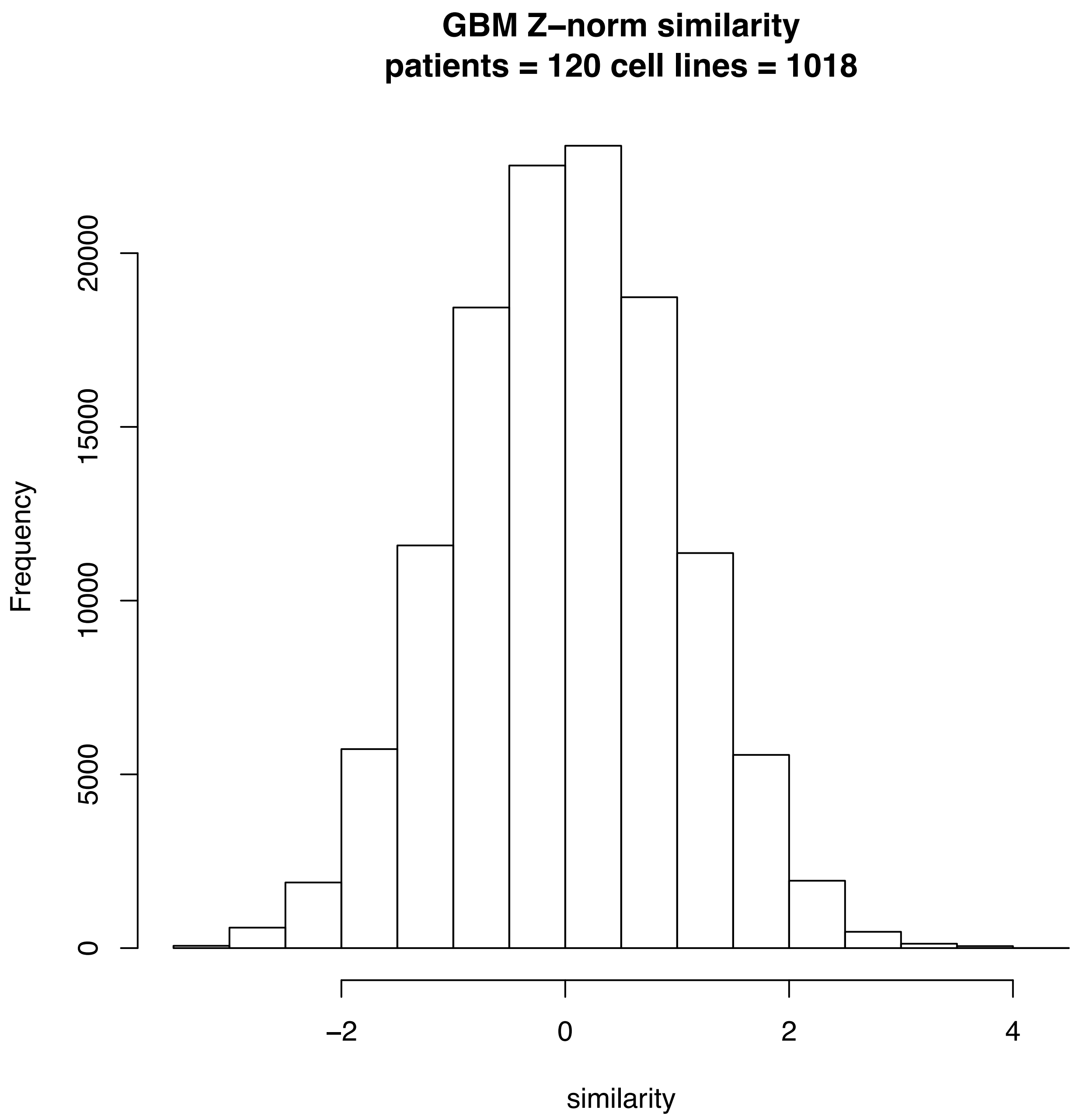
GBM Z–norm similarity
patients = 120 cell lines = 1018
Frequency
0
5000
10000
15000
20000
–2
0
2
4
similarity

GBM
No. of cell lines in low/high:26/31; p =1

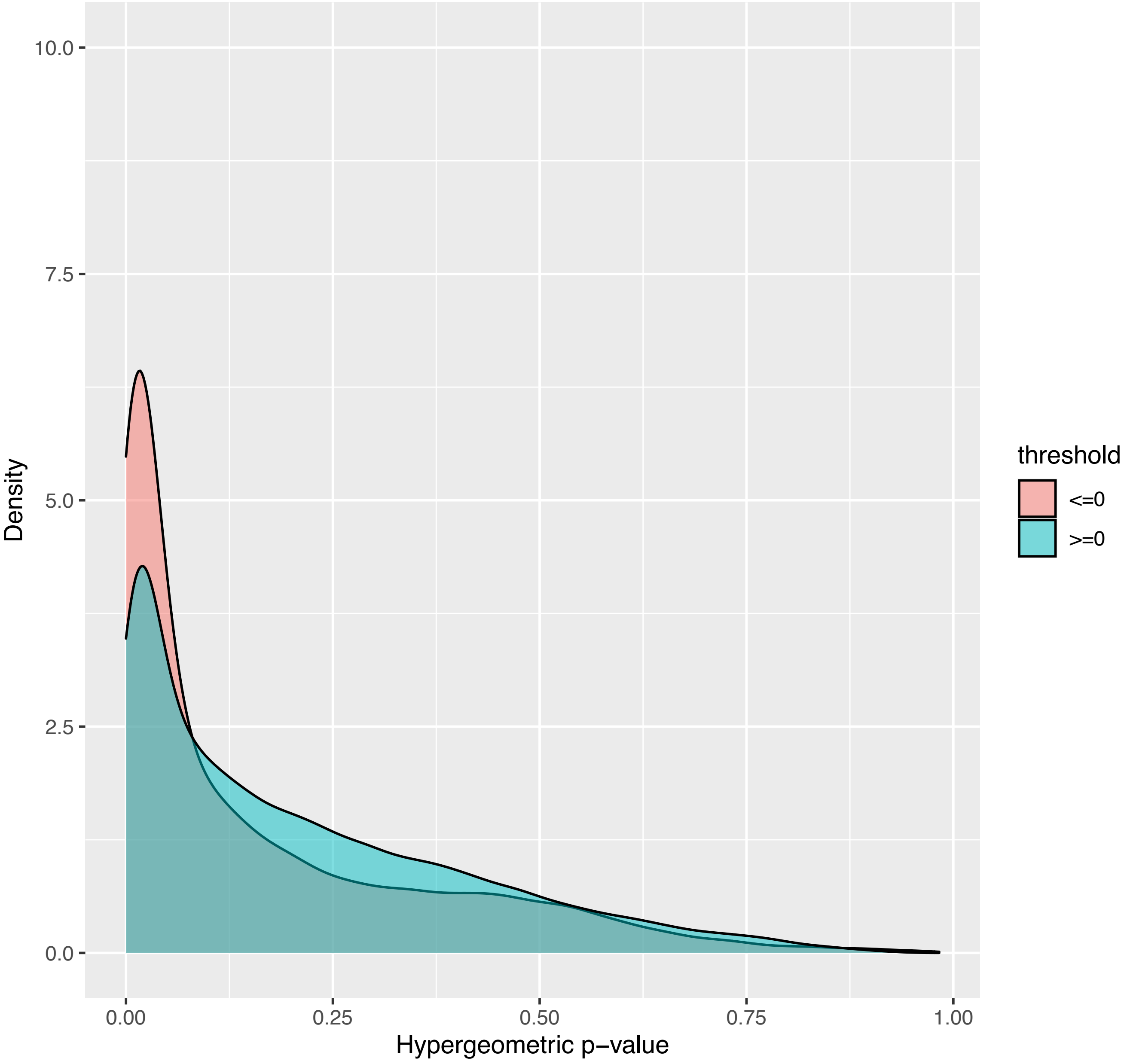

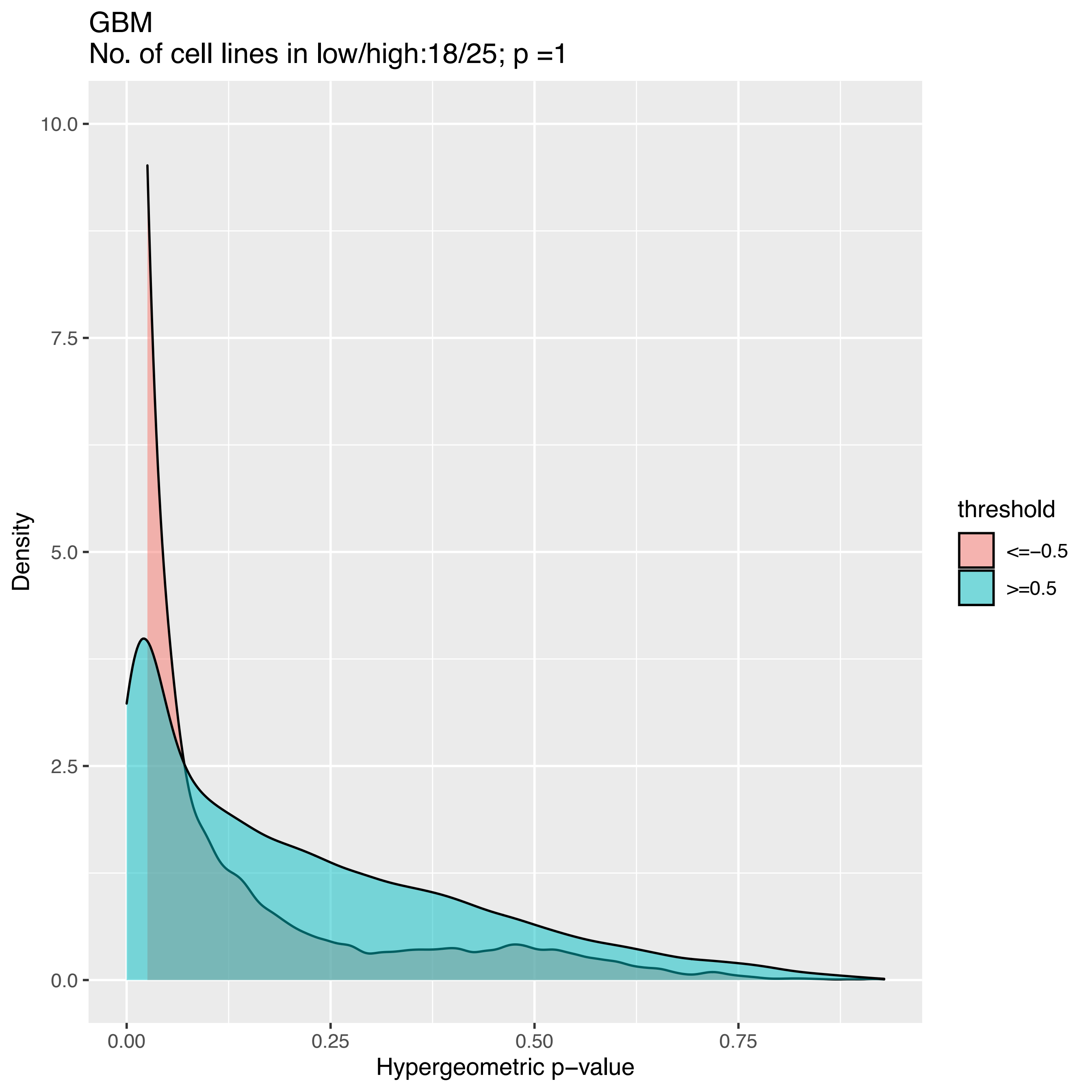
GBM
No. of cell lines in low/high:18/25; p =1
Density
10.0
7.5
5.0
2.5
0.0
0.00
0.25
0.50
0.75
Hypergeometric p−value
threshold
<=−0.5
>=0.5

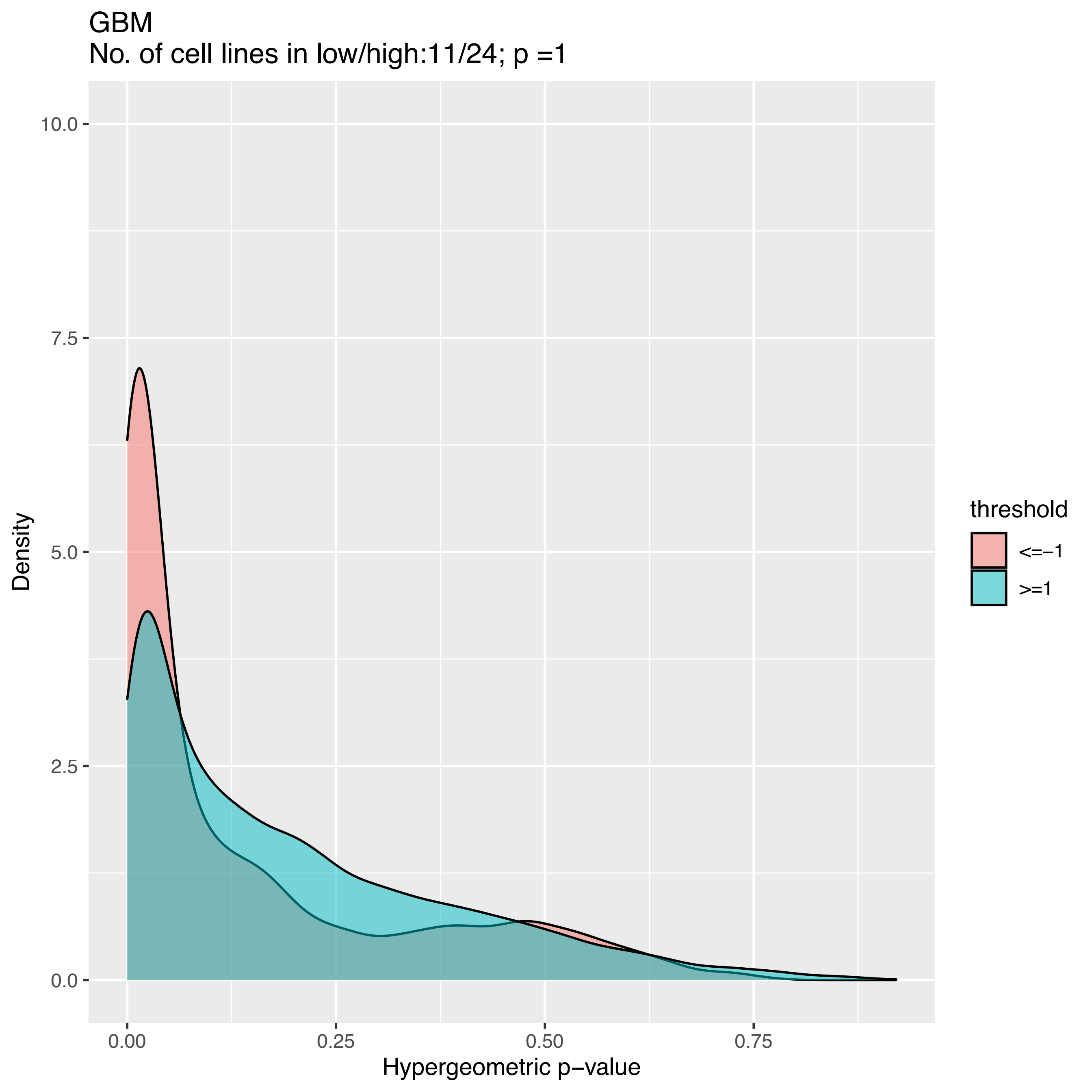
GBM
No. of cell lines in low/high:11/24; p =1
10.0
7.5
5.0
2.5
0.0
Density
0.00
0.25
0.50
0.75
Hypergeometric p−value
threshold
<=−1
>=1

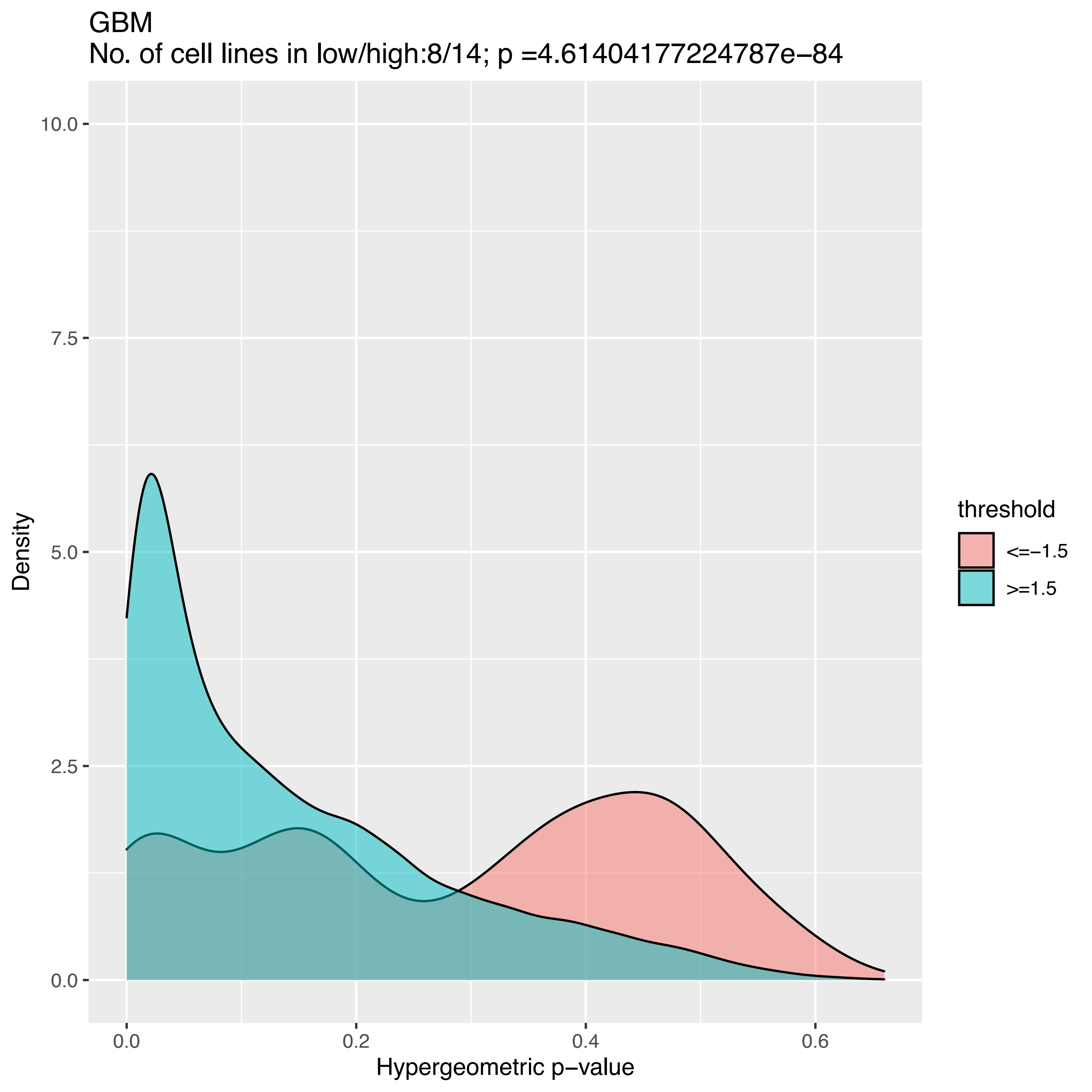
GBM
No. of cell lines in low/high:8/14; p =4.61404177224787e−84
10.0
7.5
5.0
2.5
0.0
Density
0.0
0.2
0.4
0.6
Hypergeometric p−value
threshold
<=−1.5
>=1.5

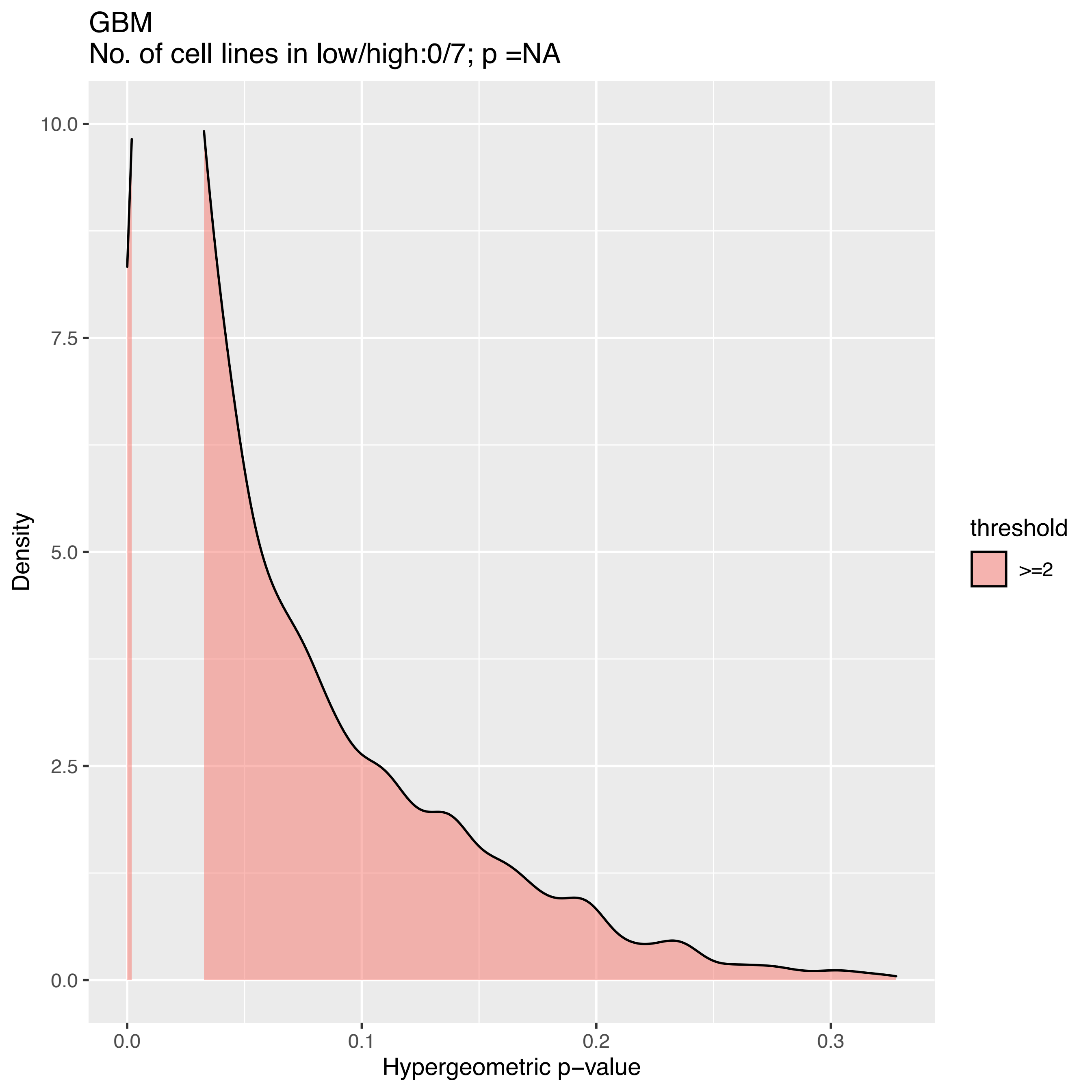
GBM
No. of cell lines in low/high:0/7; p =NA
Density
10.0
7.5
5.0
2.5
0.0
0.0
0.1
0.2
0.3
Hypergeometric p−value
threshold
>=2

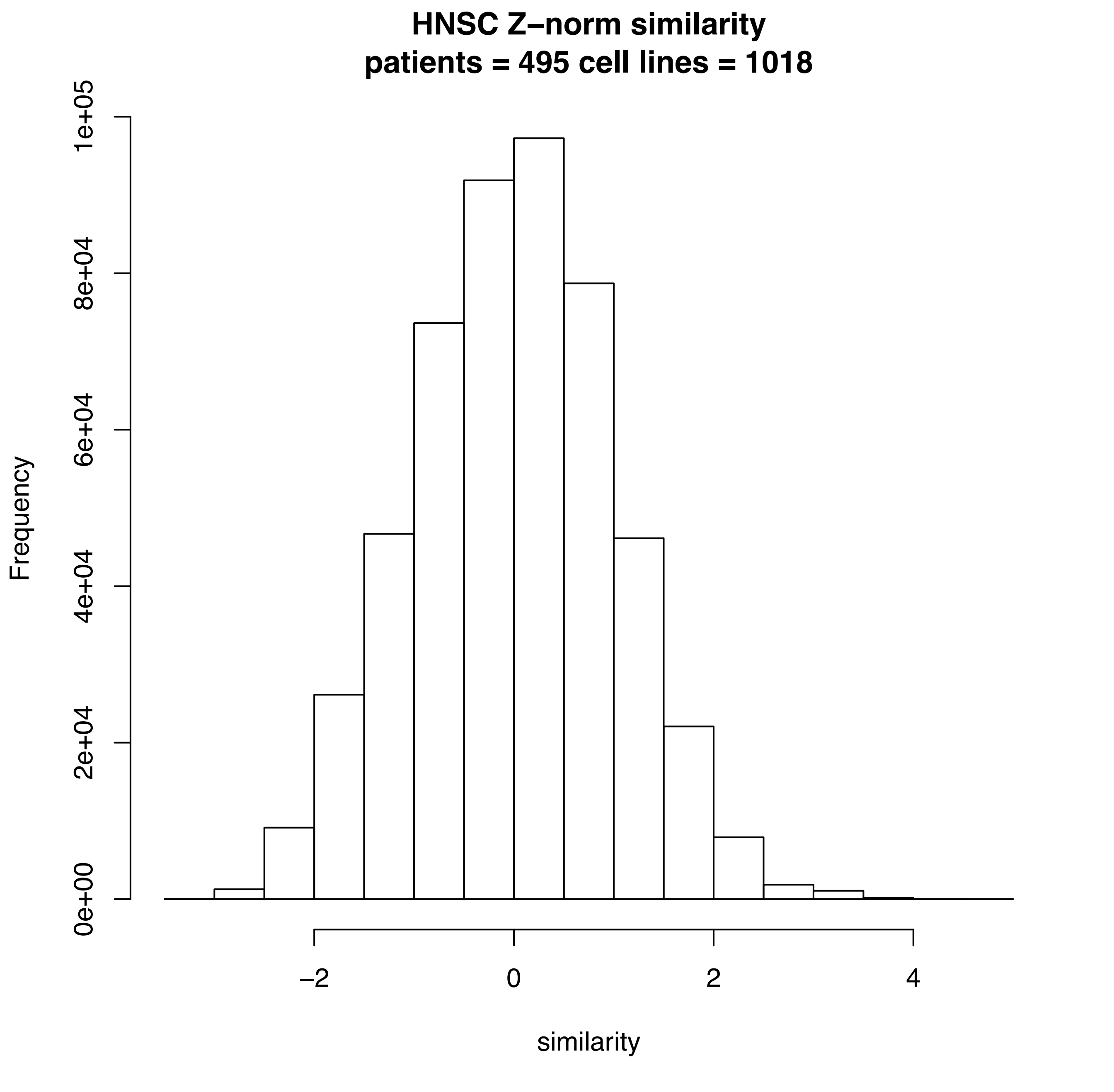
HNSC Z–norm similarity
patients = 495 cell lines = 1018
Frequency
0e+00
2e+04
4e+04
6e+04
8e+04
1e+05
–2
0
2
4
similarity

HNSC
No. of cell lines in low/high:28/33; p =0

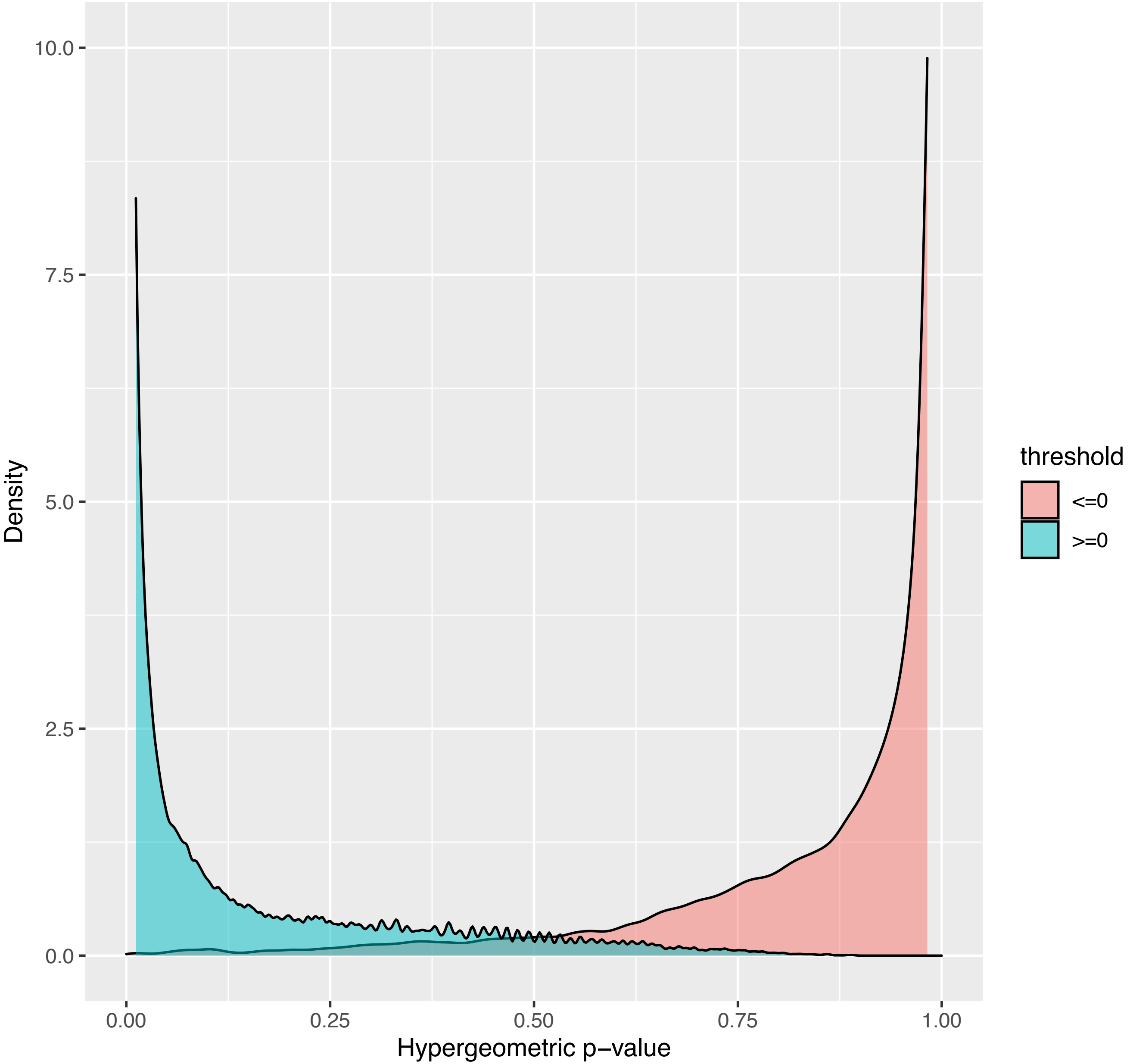

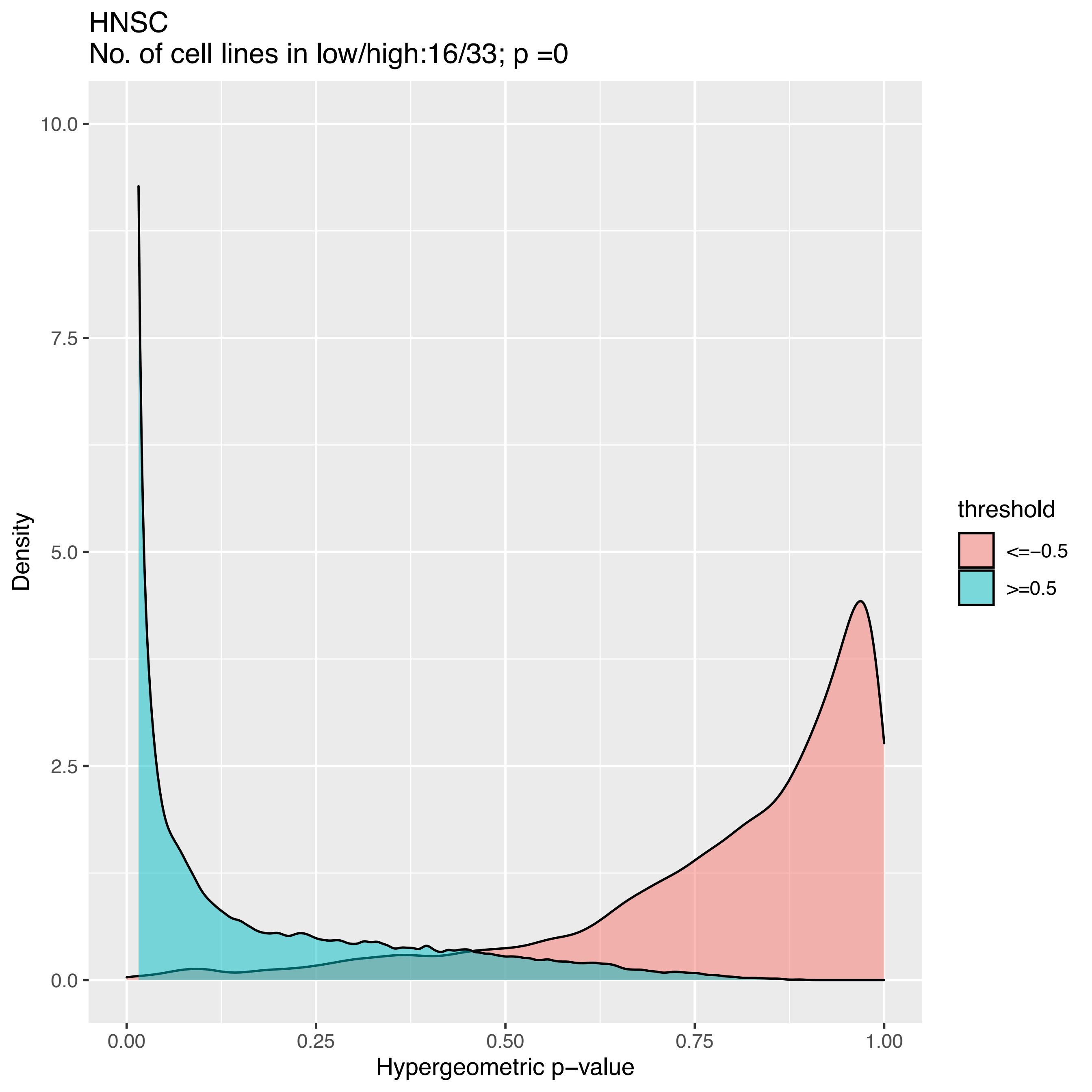
HNSC
No. of cell lines in low/high:16/33; p =0
Density
10.0
7.5
5.0
2.5
0.0
0.00
0.25
0.50
0.75
1.00
Hypergeometric p−value
threshold
<=−0.5
>=0.5

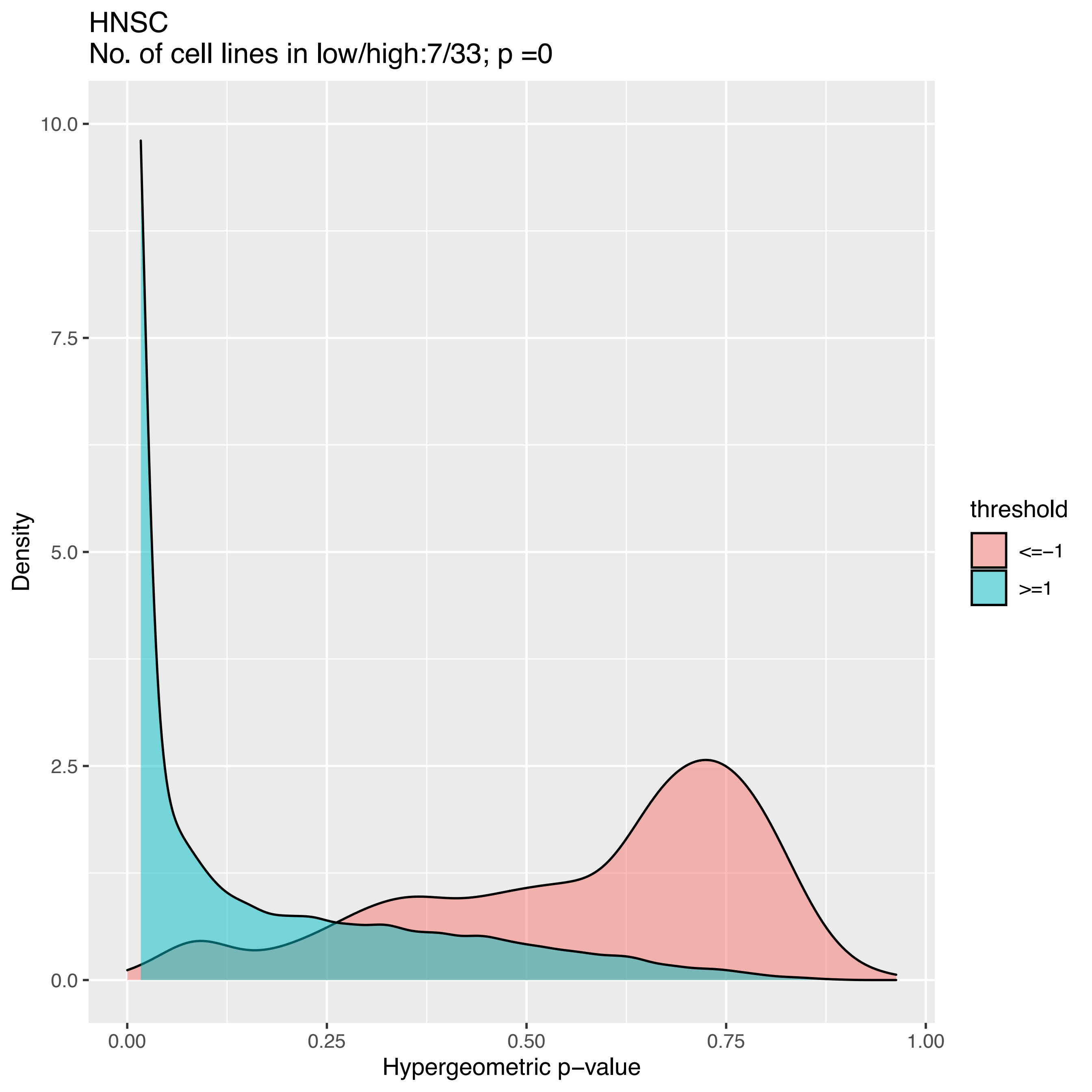

HNSC
No. of cell lines in low/high:7/33; p =0
Density
10.0
7.5
5.0
2.5
0.0
0.00
0.25
0.50
0.75
1.00
Hypergeometric p−value
threshold
<=−1
>=1

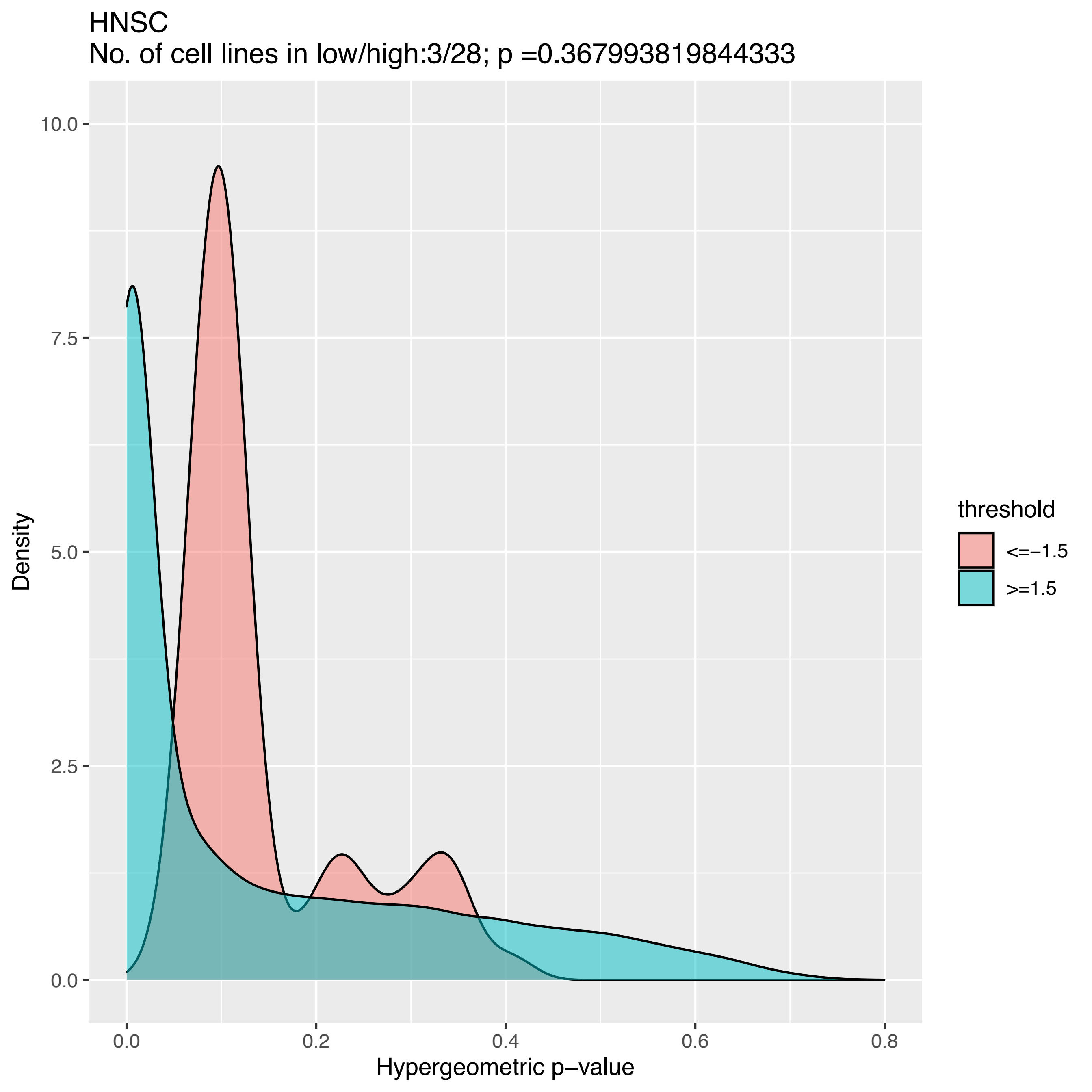
HNSC
No. of cell lines in low/high:3/28; p =0.367993819844333
10.0
7.5
5.0
2.5
0.0
Density
0.0
0.2
0.4
0.6
0.8
Hypergeometric p−value
threshold
<=−1.5
>=1.5

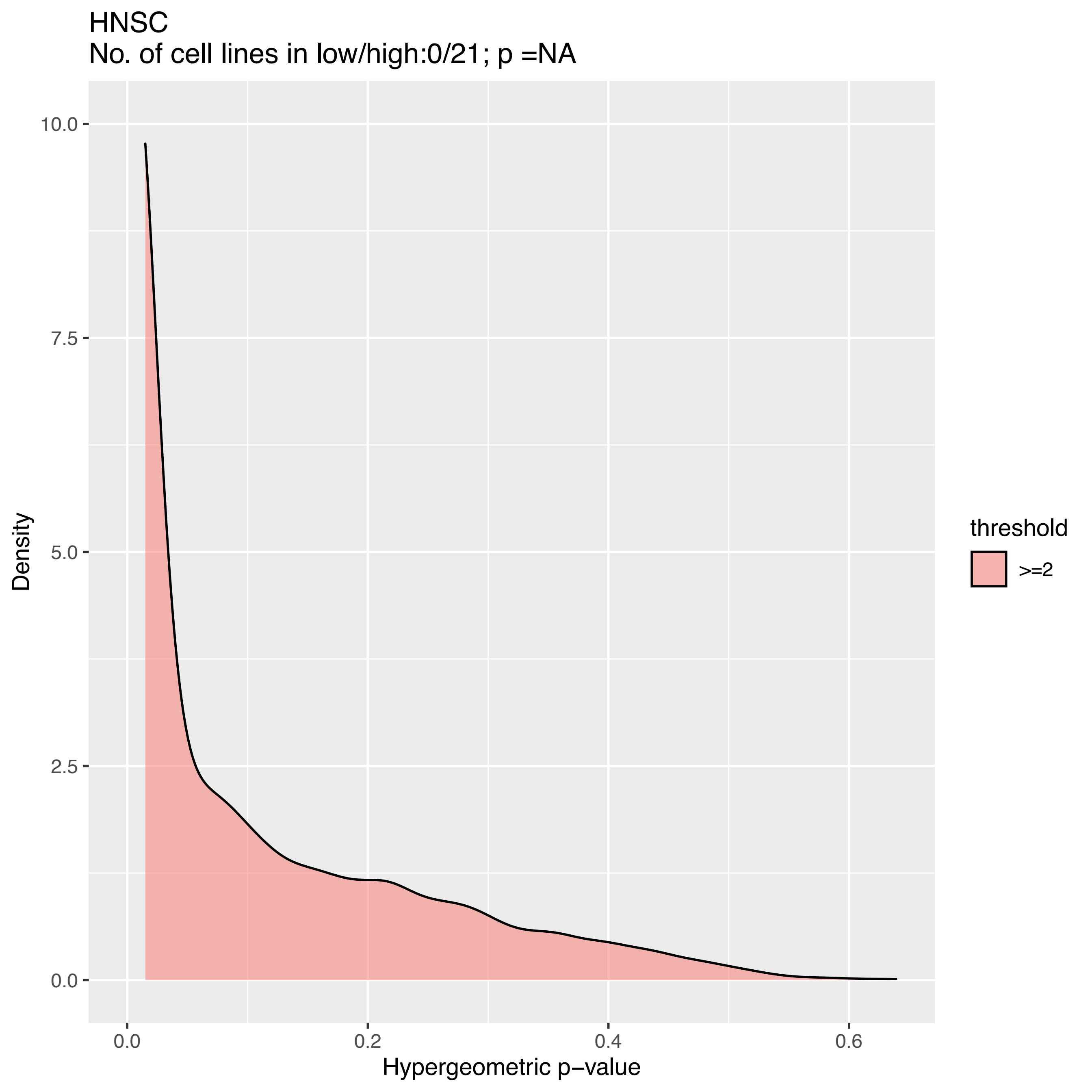
HNSC
No. of cell lines in low/high:0/21; p =NA
Density
10.0
7.5
5.0
2.5
0.0
0.0
0.2
0.4
0.6
Hypergeometric p−value
threshold
>=2

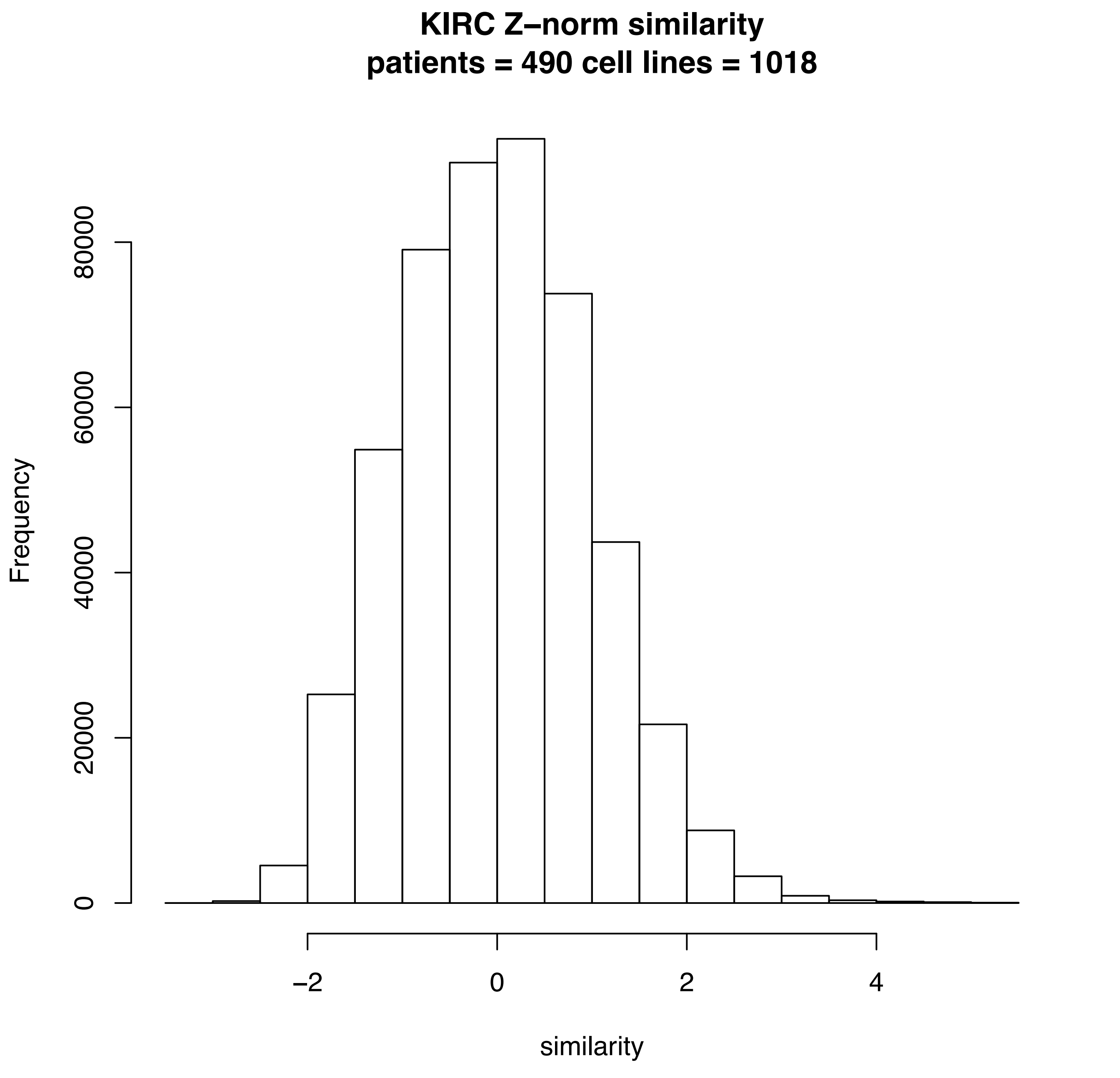
KIRC Z−norm similarity
patients = 490 cell lines = 1018
Frequency
0
20000
40000
60000
80000
−2
0
2
4
similarity

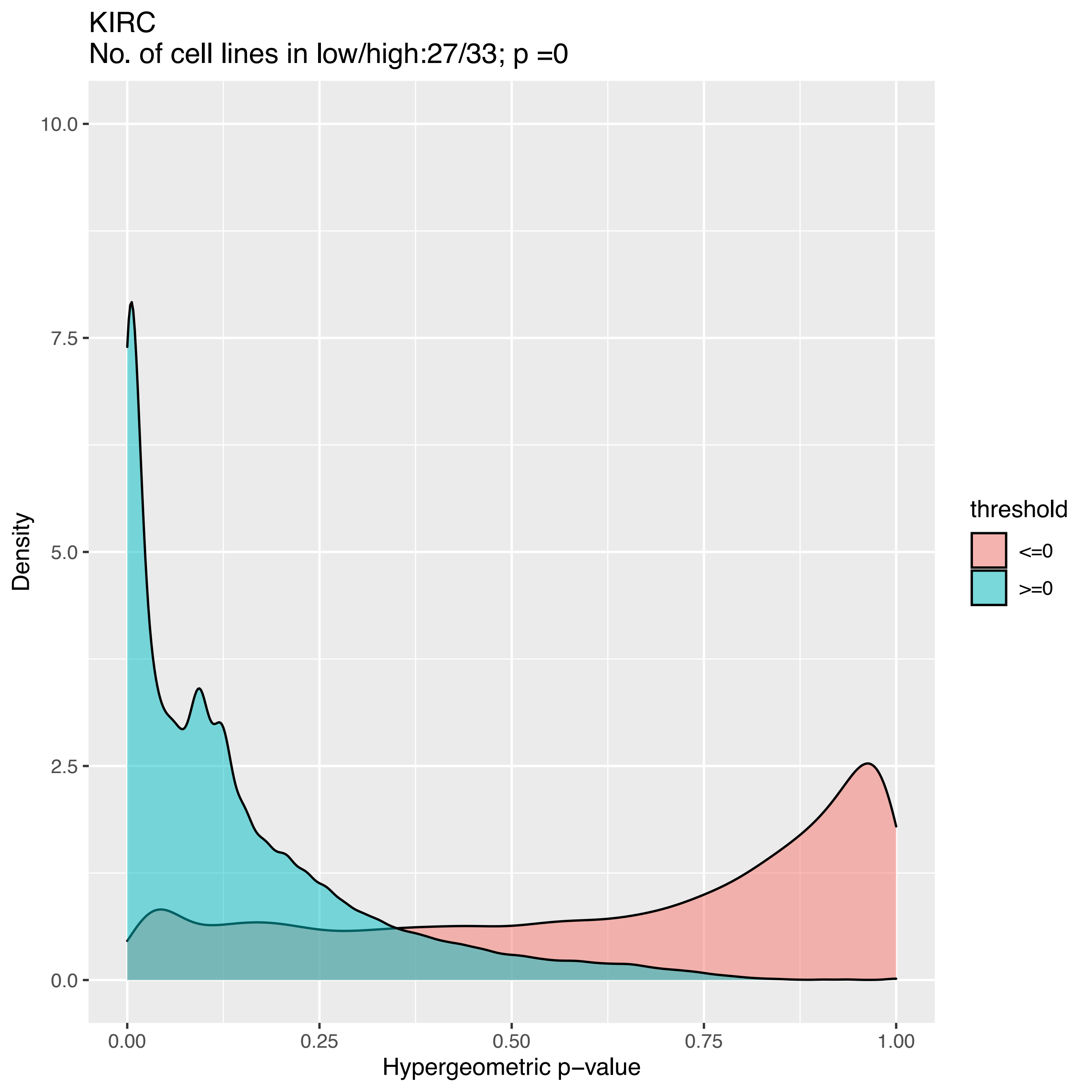
KIRC
No. of cell lines in low/high:27/33; p =0
10.0
7.5
5.0
2.5
0.0
Density
0.00
0.25
0.50
0.75
1.00
Hypergeometric p−value
threshold
<=0
>=0

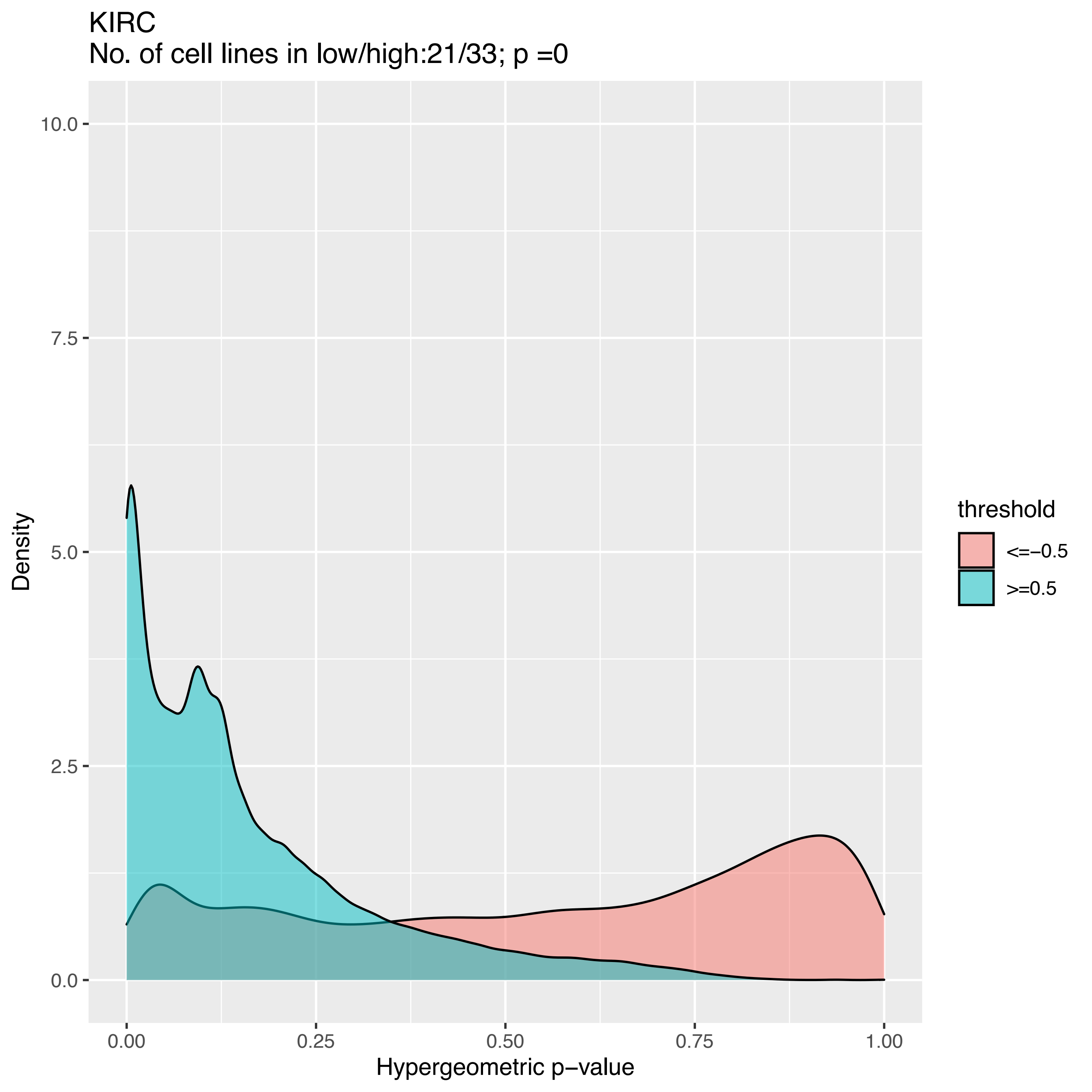
KIRC
No. of cell lines in low/high:21/33; p =0
10.0
7.5
5.0
2.5
0.0
Density
0.00
0.25
0.50
0.75
1.00
Hypergeometric p−value
threshold
<=−0.5
>=0.5

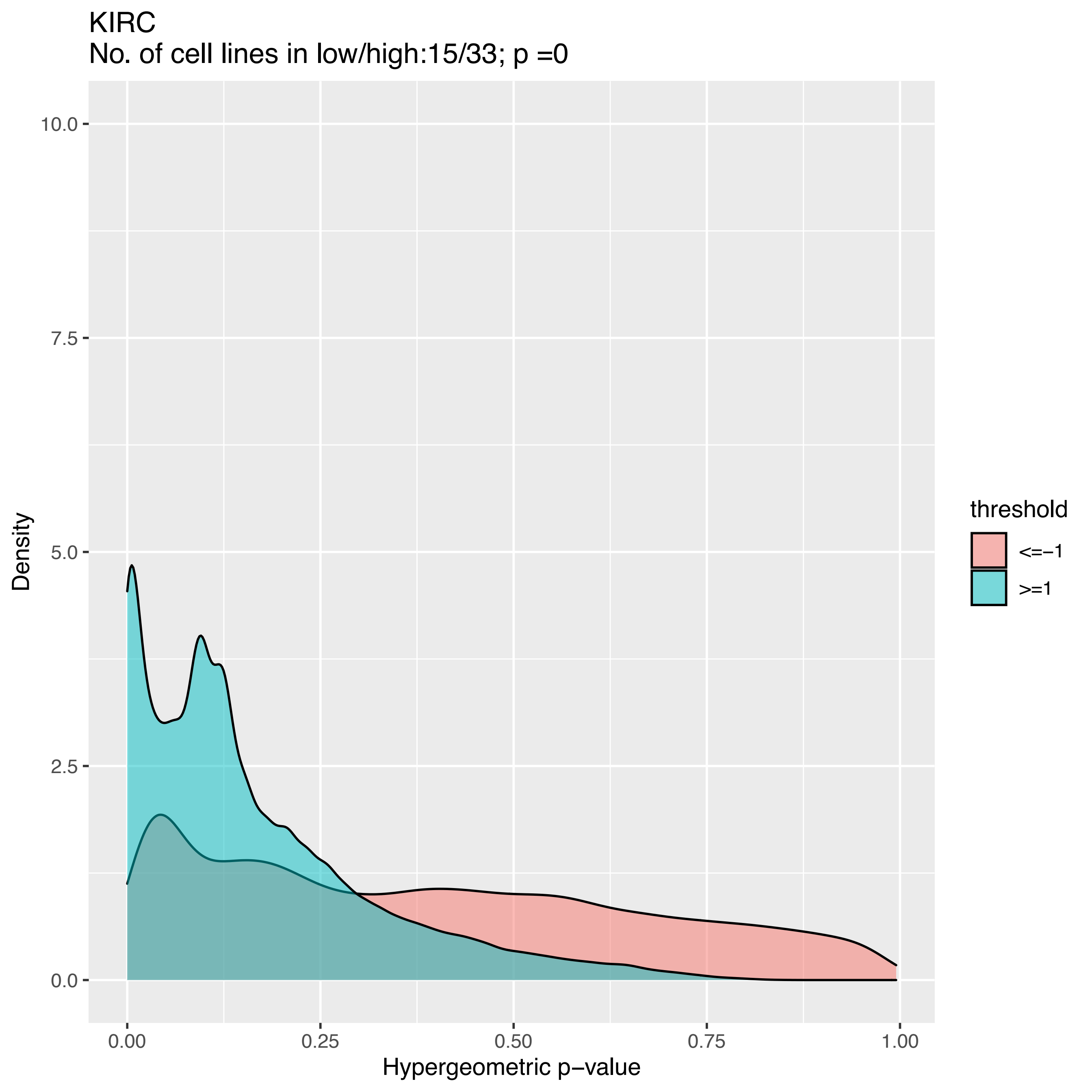
KIRC
No. of cell lines in low/high:15/33; p =0
Density
10.0
7.5
5.0
2.5
0.0
0.00
0.25
0.50
0.75
1.00
Hypergeometric p−value
threshold
<=−1
>=1

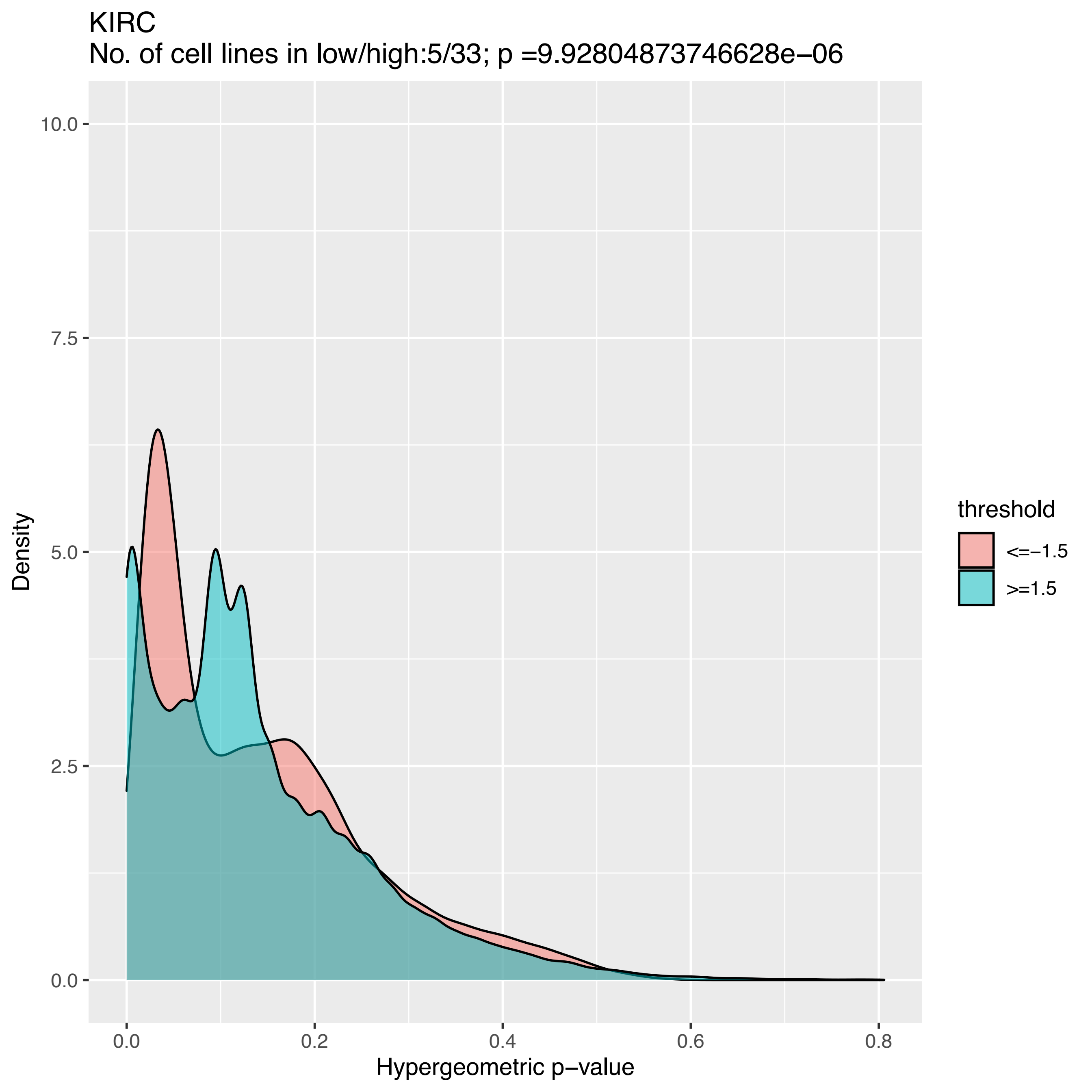

KIRC
No. of cell lines in low/high:5/33; p =9.92804873746628e−06
Density
10.0
7.5
5.0
2.5
0.0
0.0
0.2
0.4
0.6
0.8
Hypergeometric p−value
threshold
<=−1.5
>=1.5

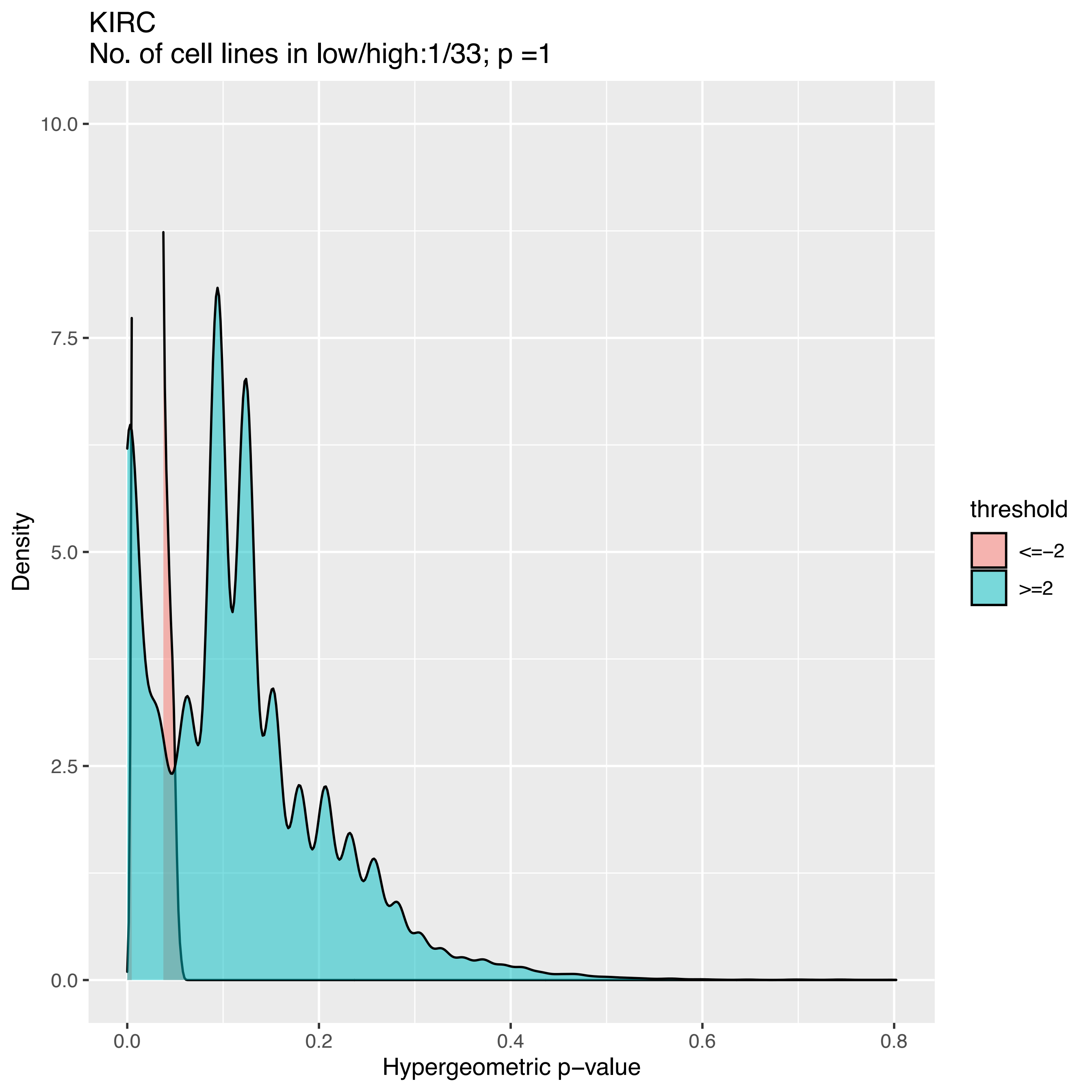
KIRC
No. of cell lines in low/high:1/33; p =1
Density
10.0
7.5
5.0
2.5
0.0
0.0
0.2
0.4
0.6
0.8
Hypergeometric p−value
threshold
<=−2
>=2

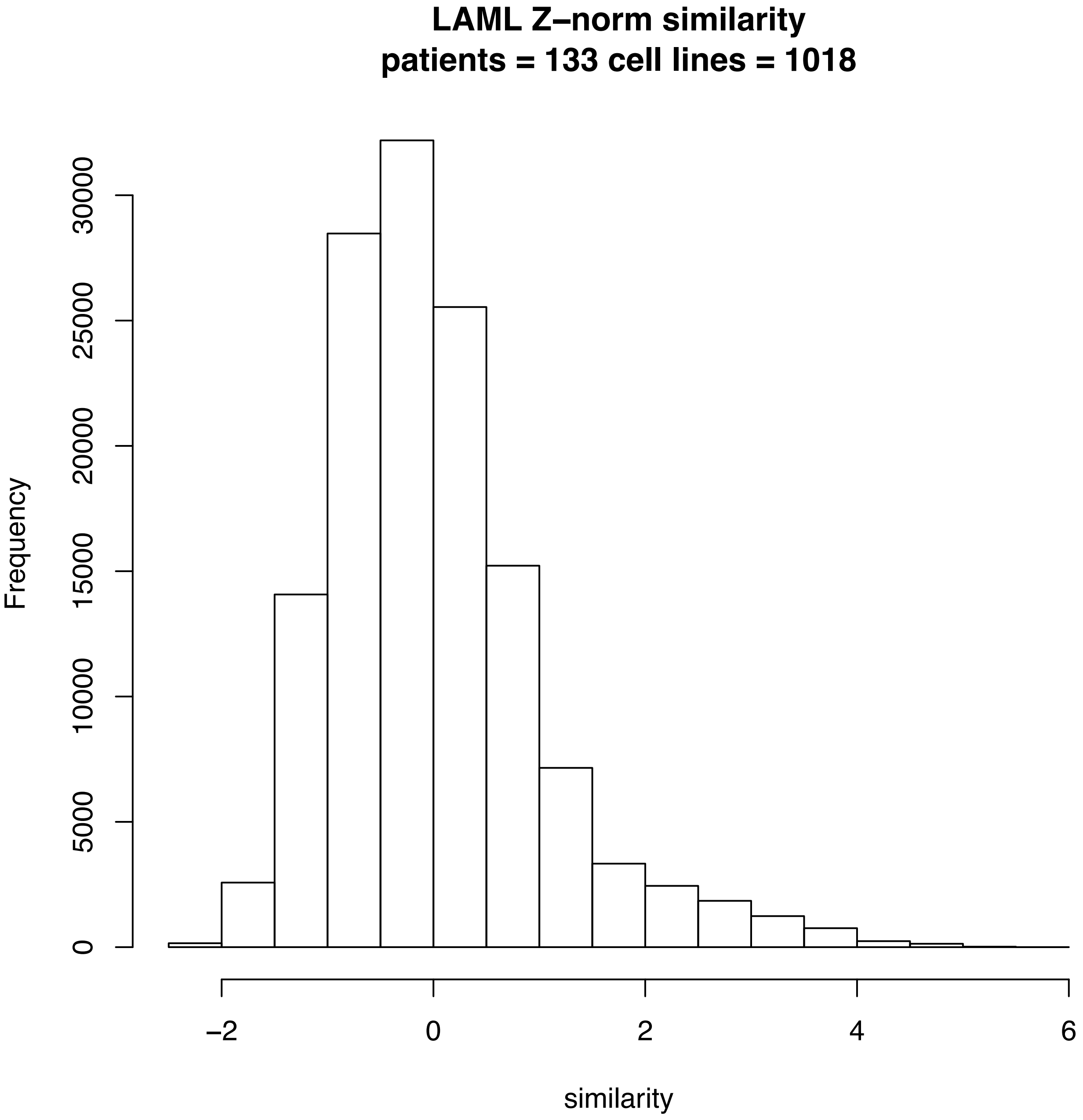
LAML Z–norm similarity
patients = 133 cell lines = 1018
Frequency
0
5000
10000
15000
20000
25000
30000
–2
0
2
4
6
similarity

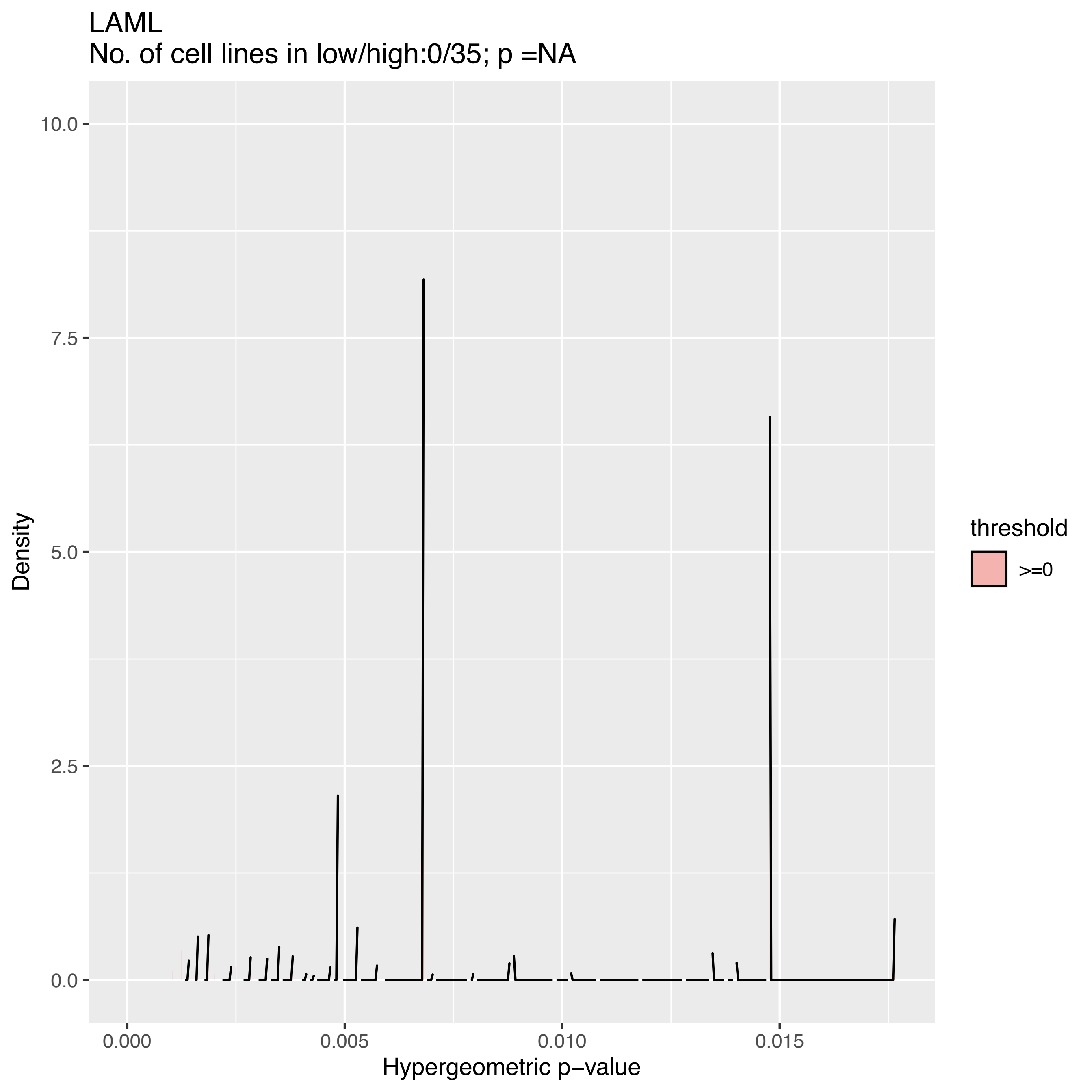

LAML
No. of cell lines in low/high:0/35; p =NA
10.0
7.5
5.0
2.5
0.0
Density
0.000
0.005
0.010
0.015
Hypergeometric p−value
threshold
>=0

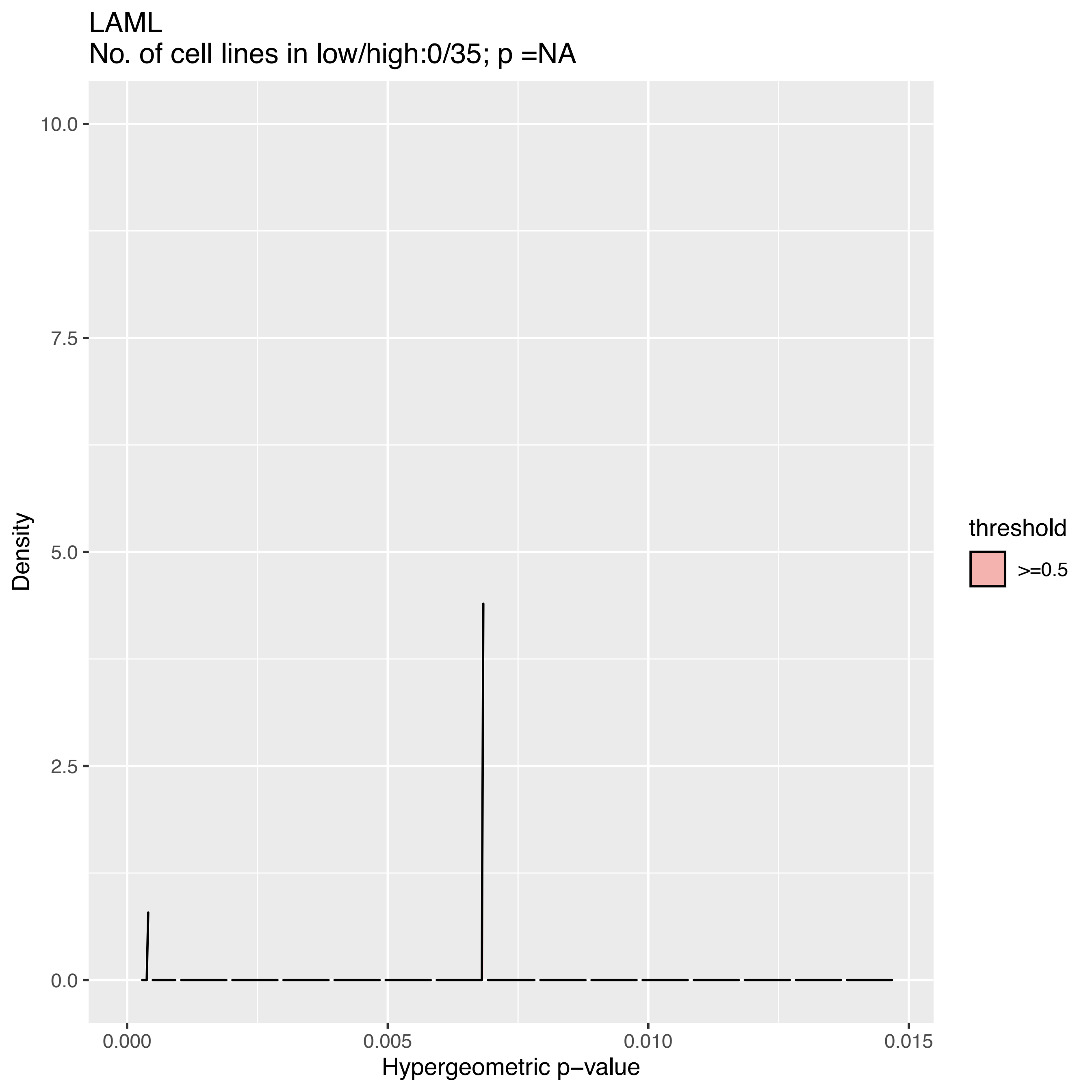

LAML
No. of cell lines in low/high:0/35; p =NA
10.0
7.5
5.0
2.5
0.0
0.000
0.005
0.010
0.015
Hypergeometric p−value
Density
threshold
>=0.5

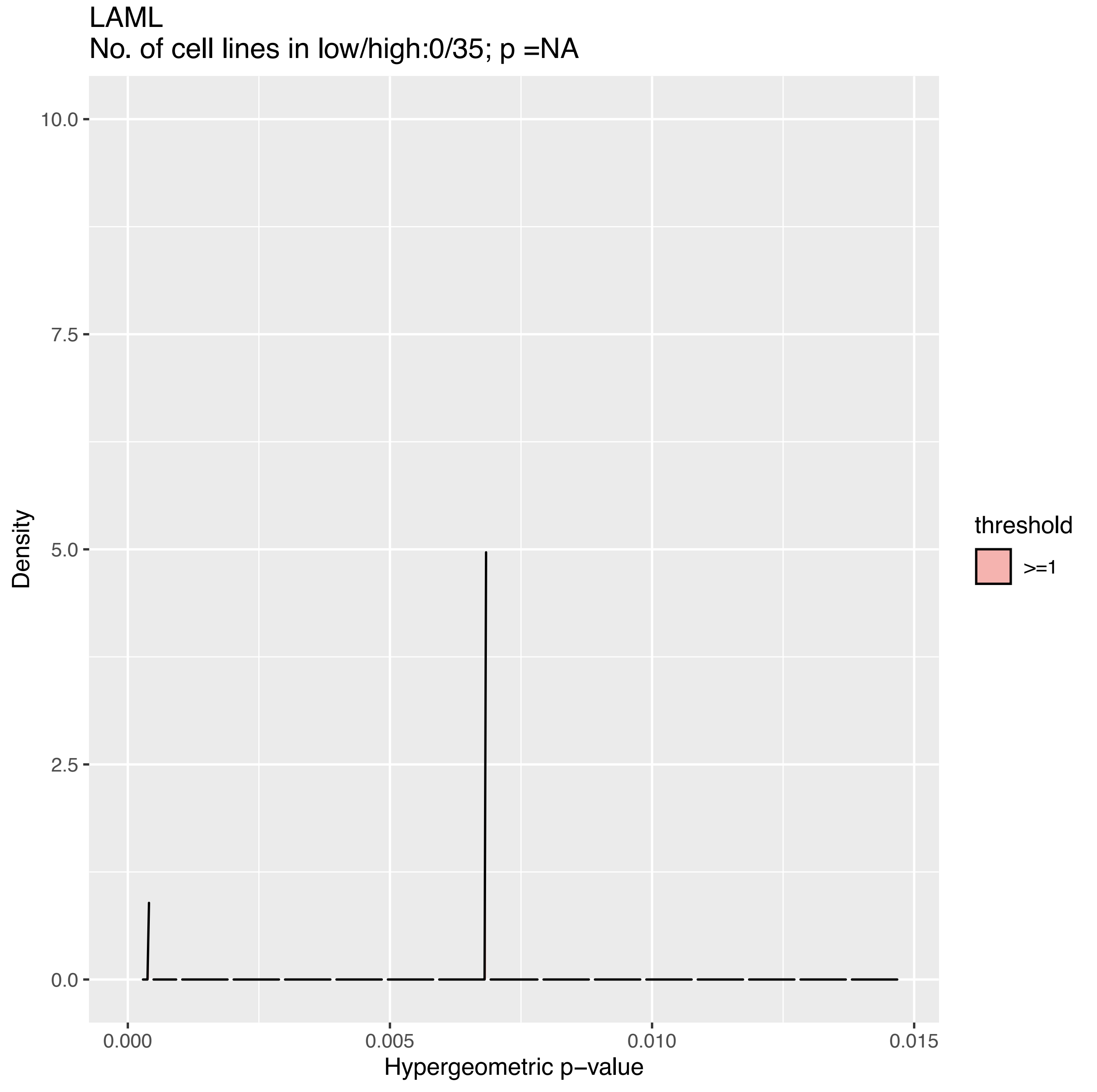

LAML
No. of cell lines in low/high:0/35; p =NA
Density
10.0
7.5
5.0
2.5
0.0
0.000
0.005
0.010
0.015
Hypergeometric p−value
threshold
>=1

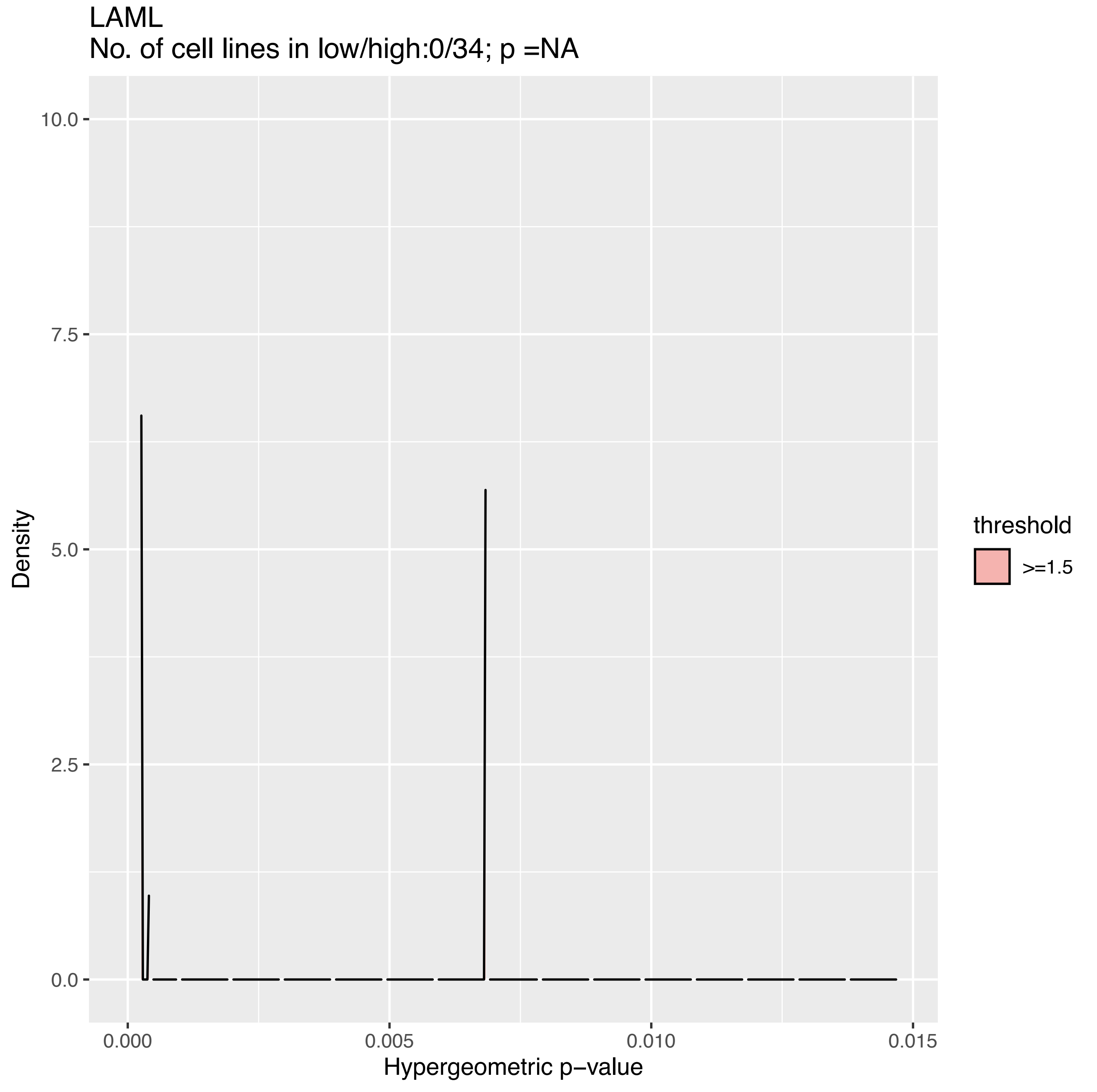
LAML
No. of cell lines in low/high:0/34; p =NA
10.0
7.5
5.0
2.5
0.0
Density
0.000
0.005
0.010
0.015
Hypergeometric p−value
threshold
>=1.5

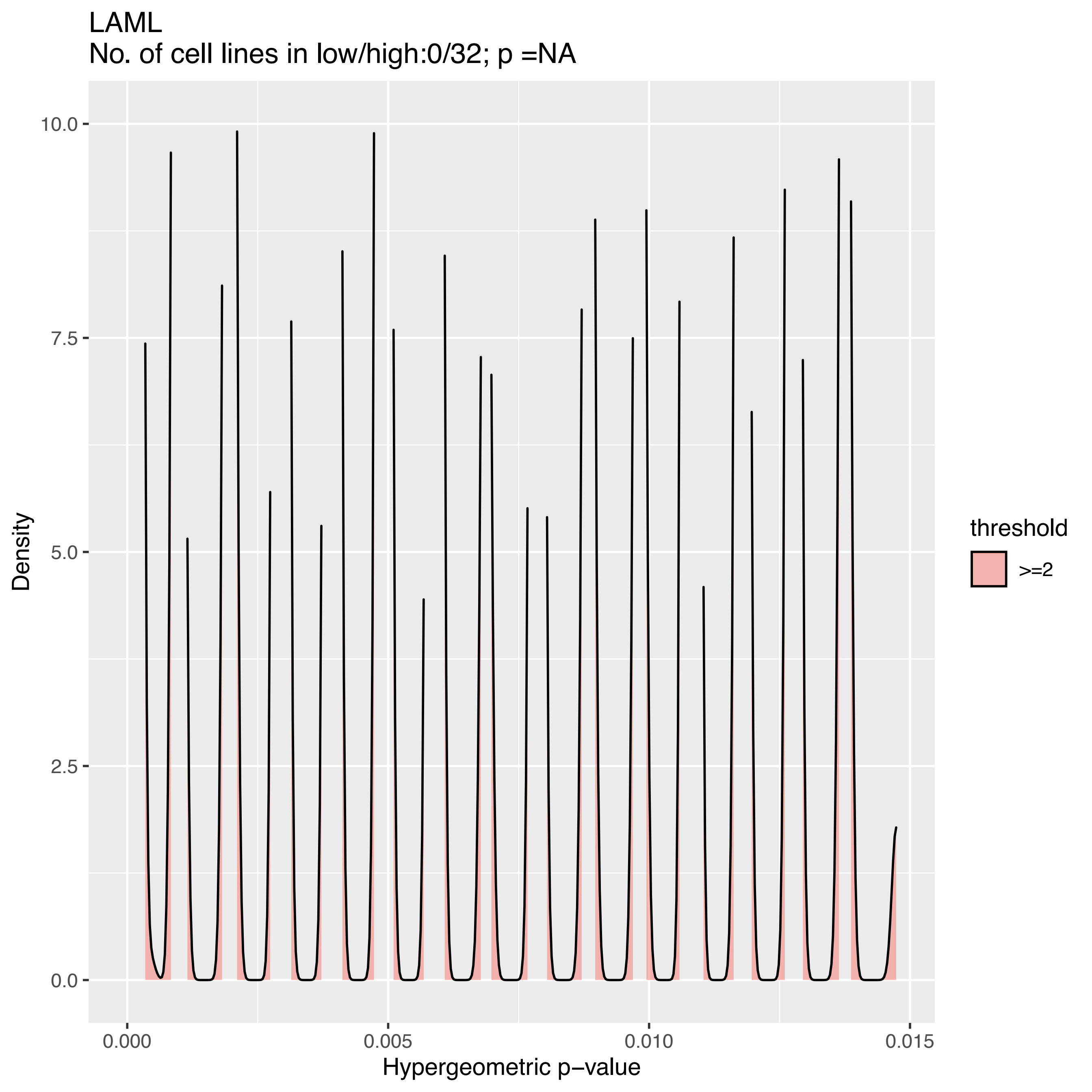

LAML
No. of cell lines in low/high:0/32; p =NA
Density
10.0
7.5
5.0
2.5
0.0
0.000
0.005
0.010
0.015
Hypergeometric p−value
threshold
>=2

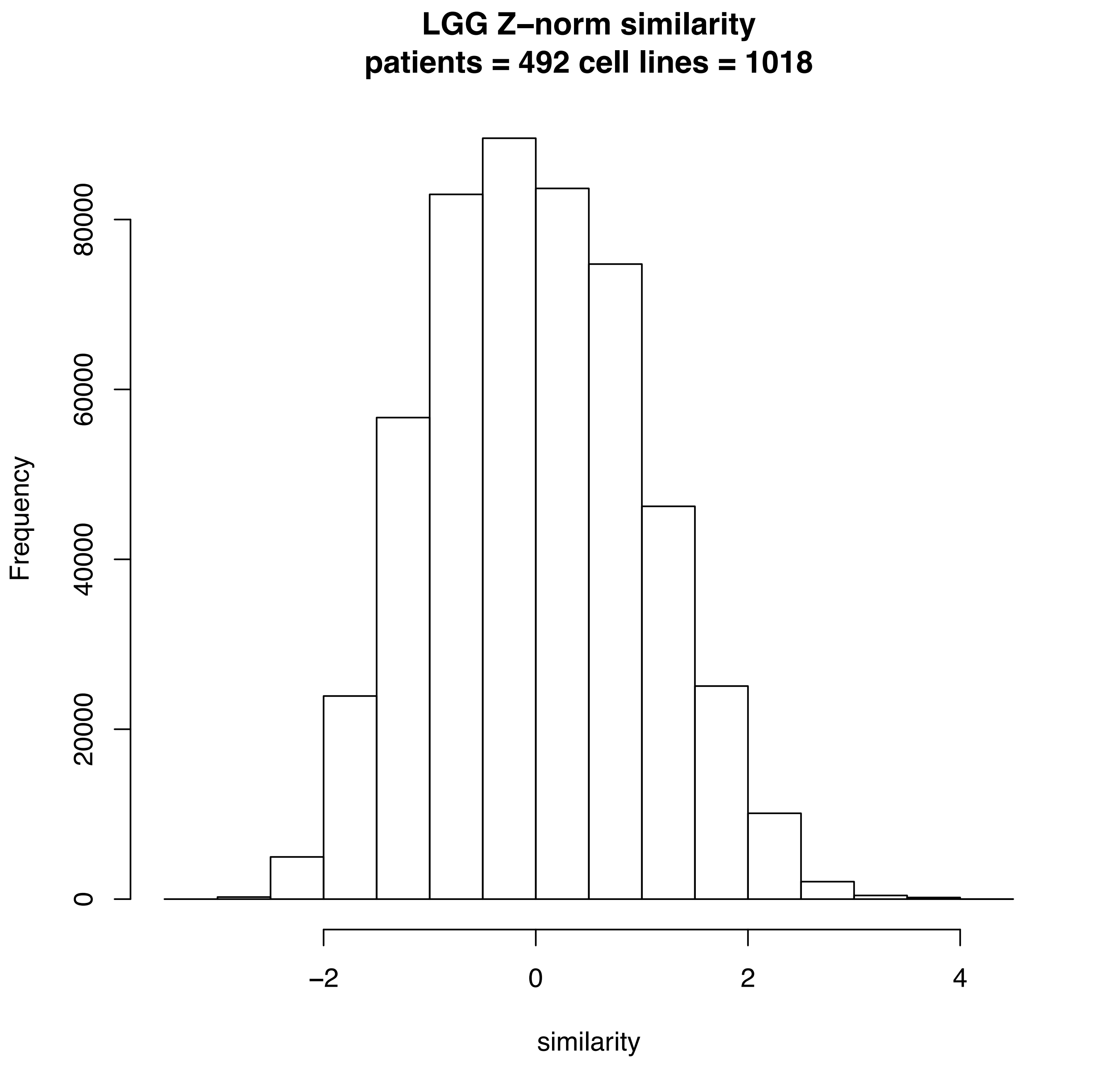
LGG Z–norm similarity
patients = 492 cell lines = 1018
Frequency
0
20000
40000
60000
80000
–2
0
2
4
similarity

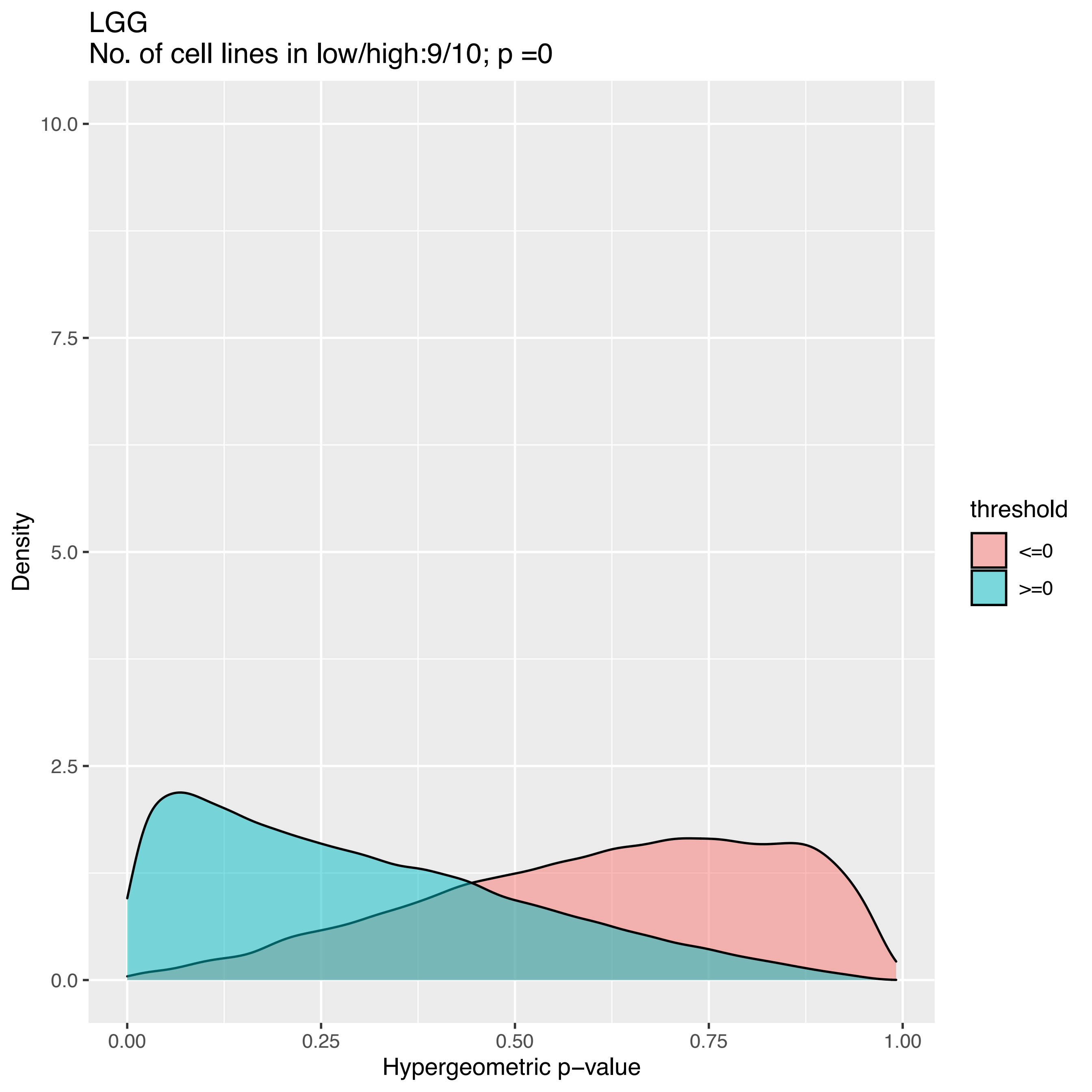
LGG
No. of cell lines in low/high:9/10; p =0
10.0
7.5
5.0
2.5
0.0
Density
0.00
0.25
0.50
0.75
1.00
Hypergeometric p−value
threshold
<=0
>=0

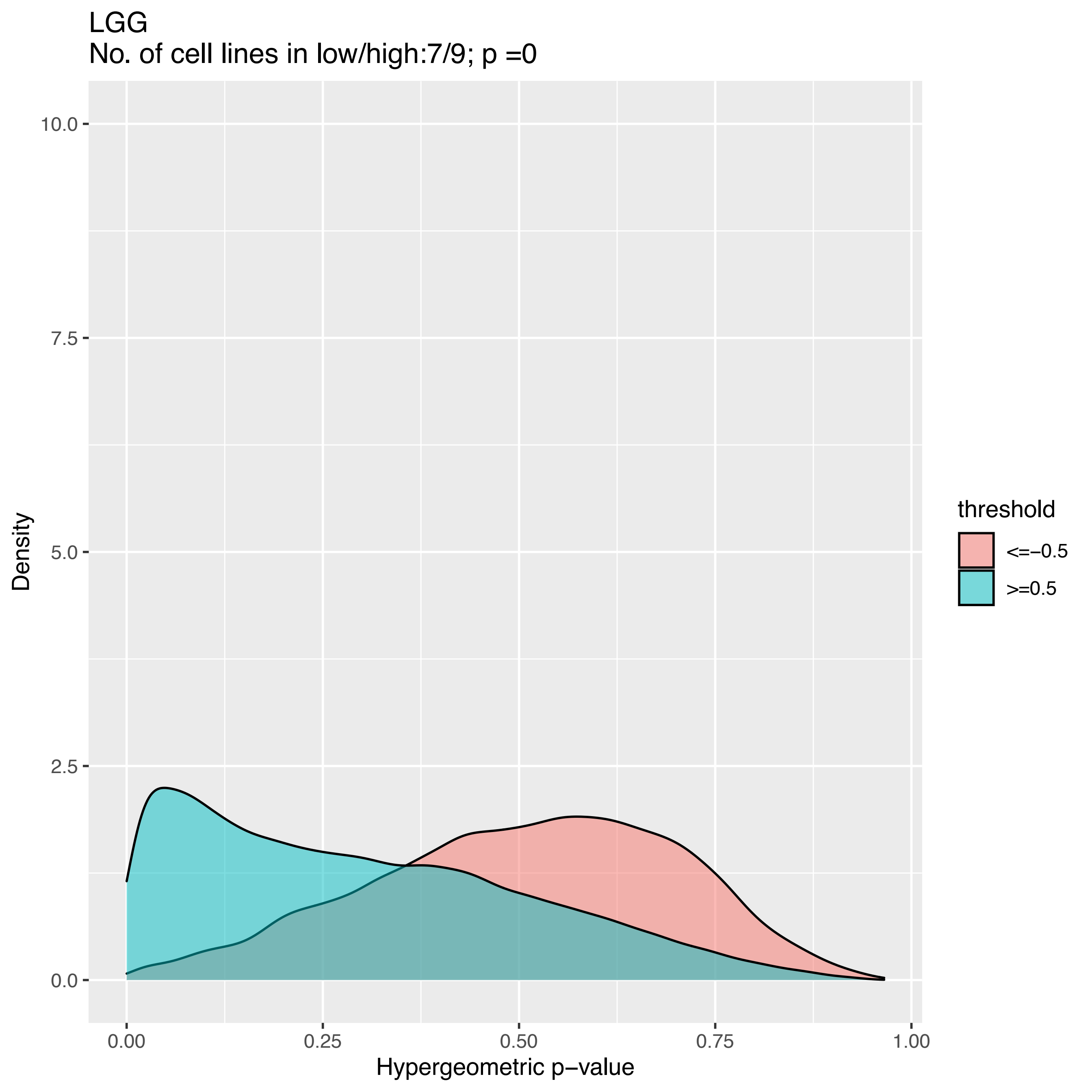
LGG
No. of cell lines in low/high:7/9; p =0
Density
10.0
7.5
5.0
2.5
0.0
0.00
0.25
0.50
0.75
1.00
Hypergeometric p−value
threshold
<=−0.5
>=0.5

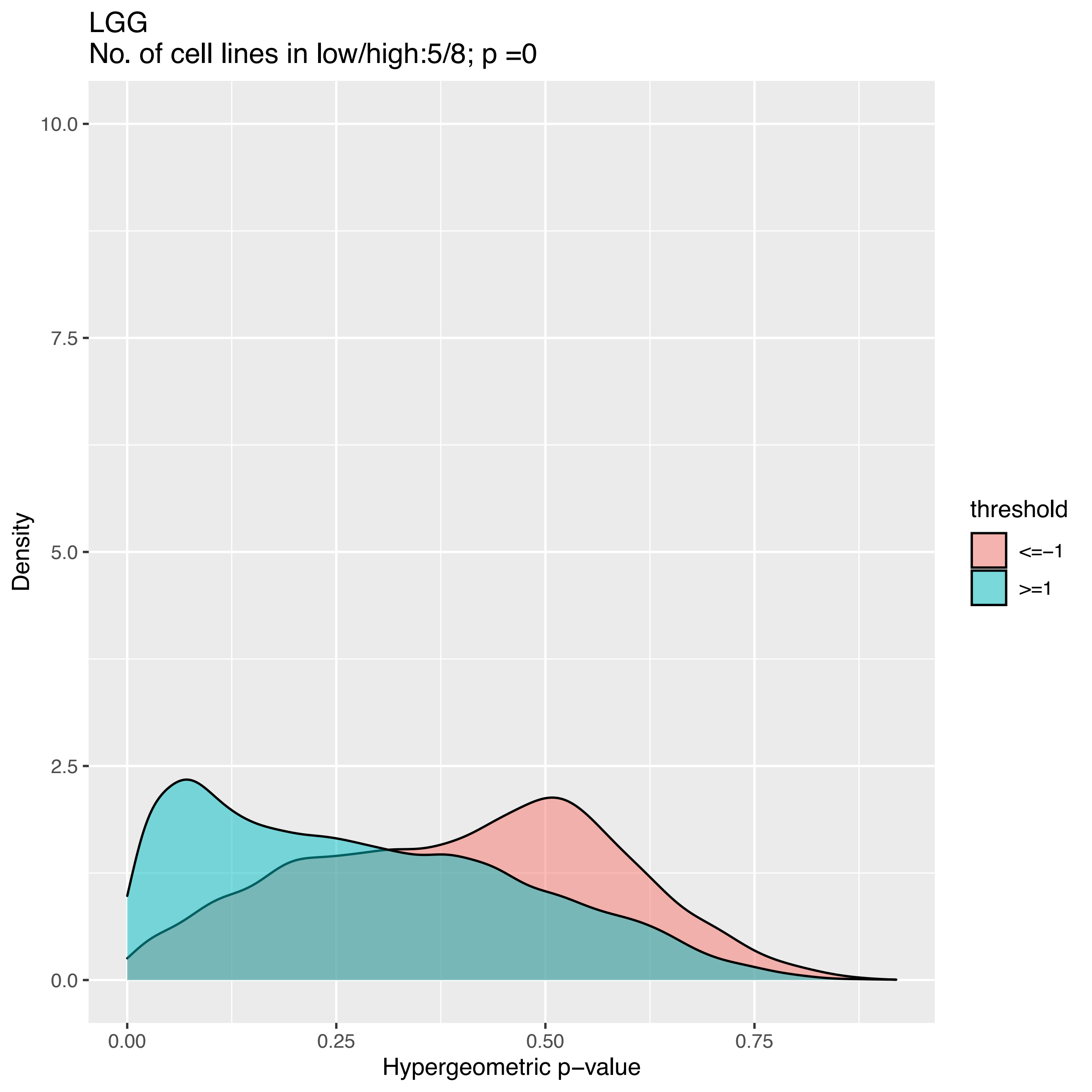

LGG
No. of cell lines in low/high:5/8; p =0
10.0
7.5
5.0
2.5
0.0
Density
0.00
0.25
0.50
0.75
Hypergeometric p−value
threshold
<=−1
>=1

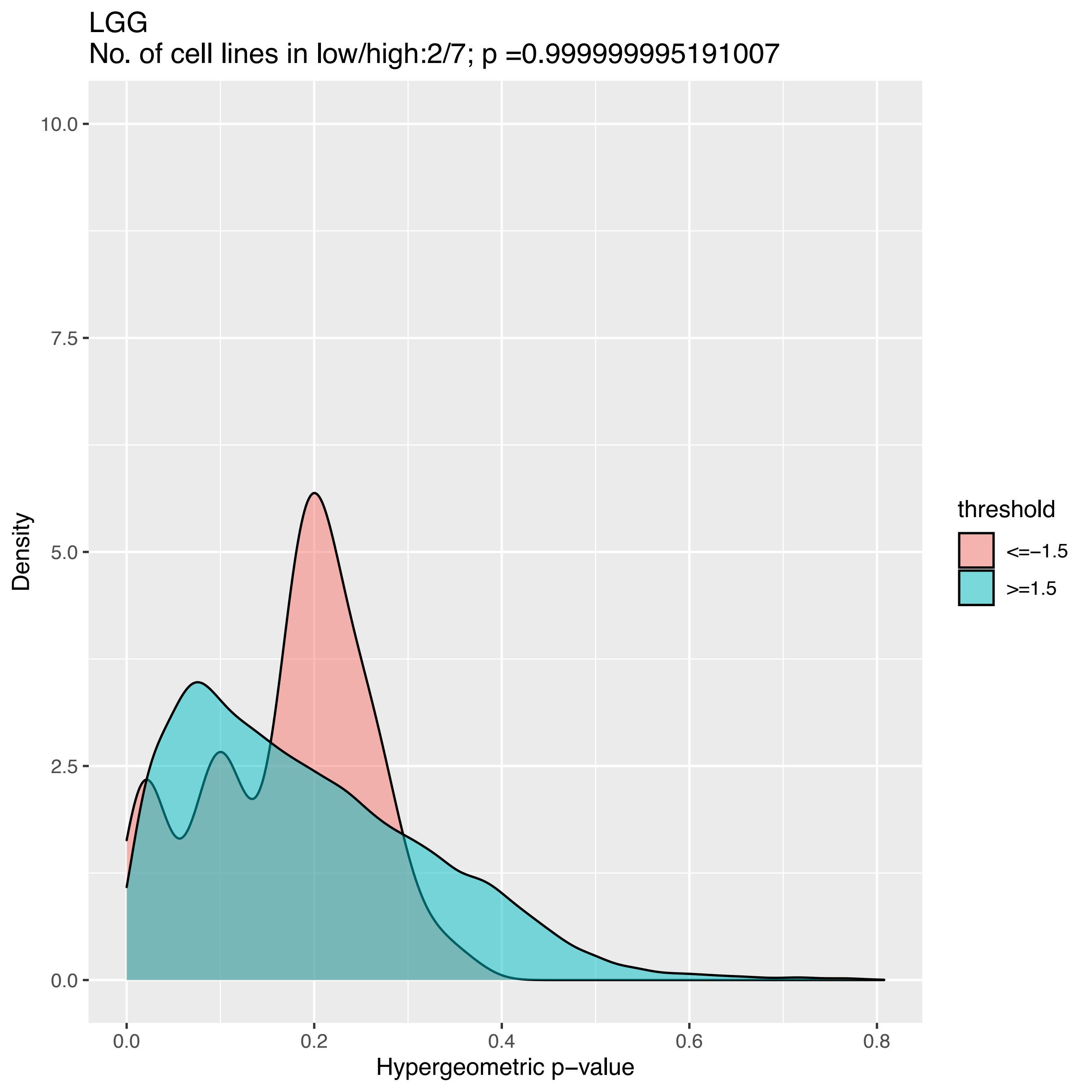
LGG
No. of cell lines in low/high:2/7; p =0.999999995191007
Density
10.0
7.5
5.0
2.5
0.0
0.0
0.2
0.4
0.6
0.8
Hypergeometric p−value
threshold
<=−1.5
>=1.5

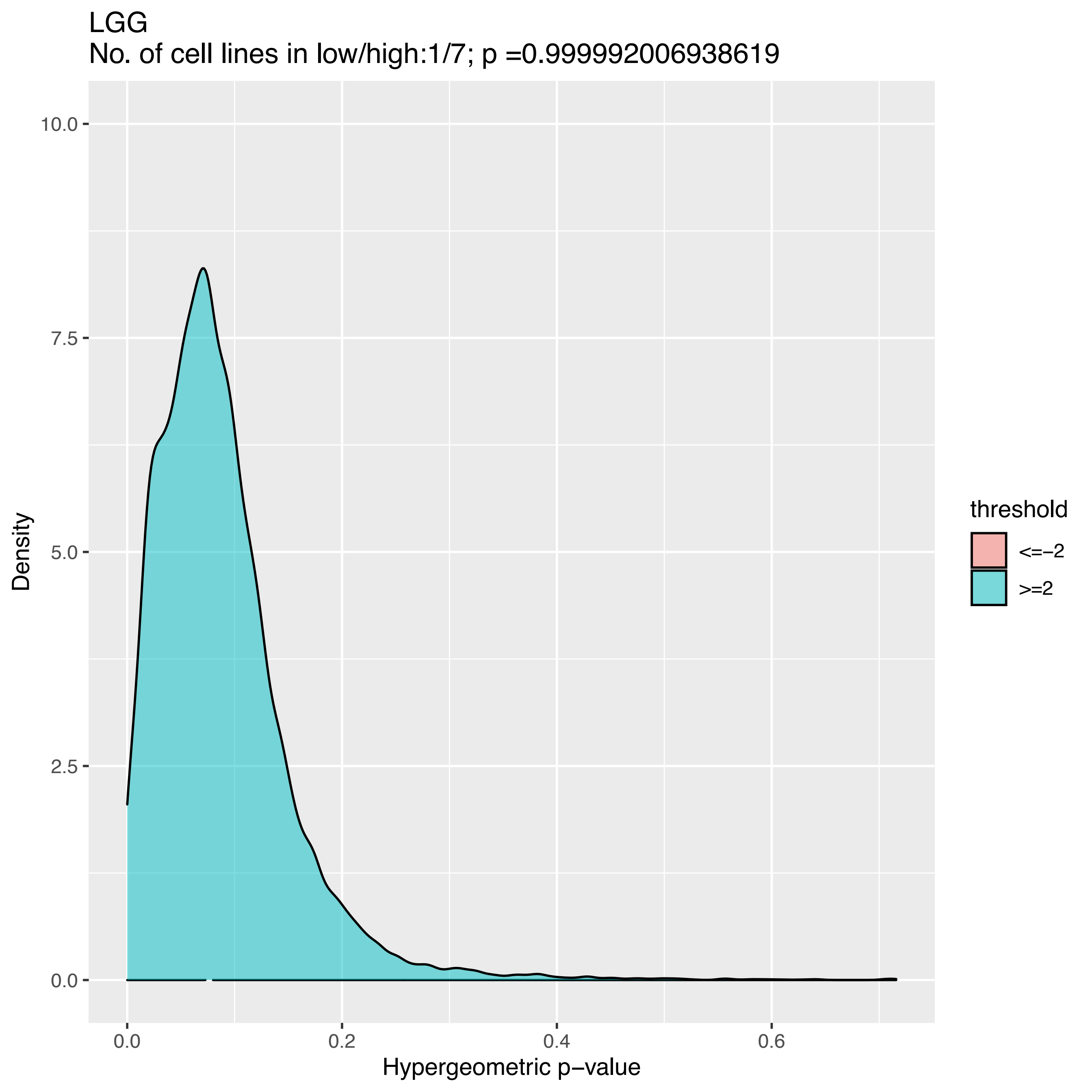
LGG
No. of cell lines in low/high:1/7; p =0.999992006938619
10.0
7.5
5.0
2.5
0.0
Density
0.0
0.2
0.4
0.6
Hypergeometric p−value
threshold
<=−2
>=2

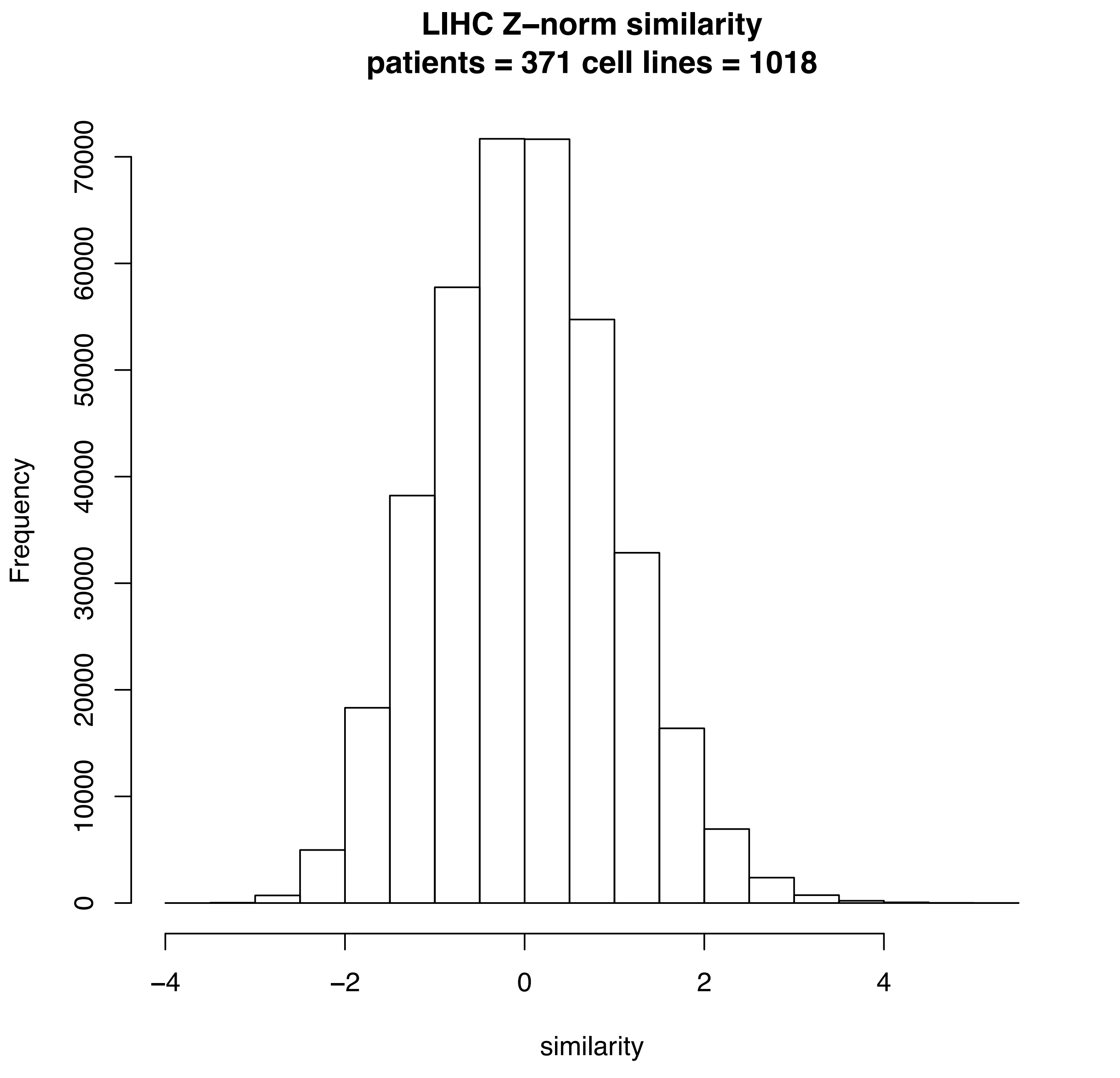
LIHC Z–norm similarity
patients = 371 cell lines = 1018
Frequency
0
10000
20000
30000
40000
50000
60000
70000
−4
−2
0
2
4
similarity

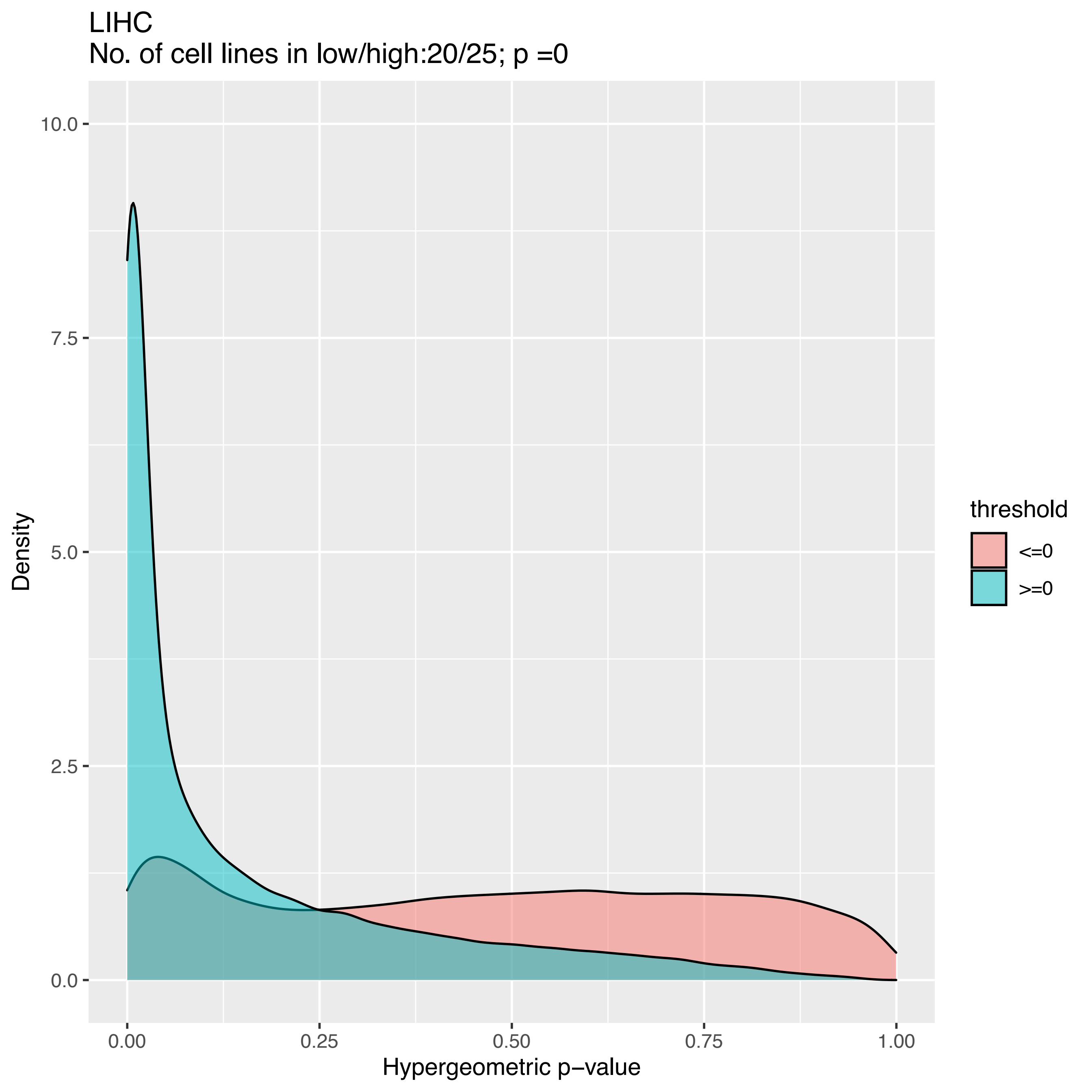
LIHC
No. of cell lines in low/high:20/25; p =0
Density
10.0
7.5
5.0
2.5
0.0
0.00
0.25
0.50
0.75
1.00
Hypergeometric p−value
threshold
<=0
>=0

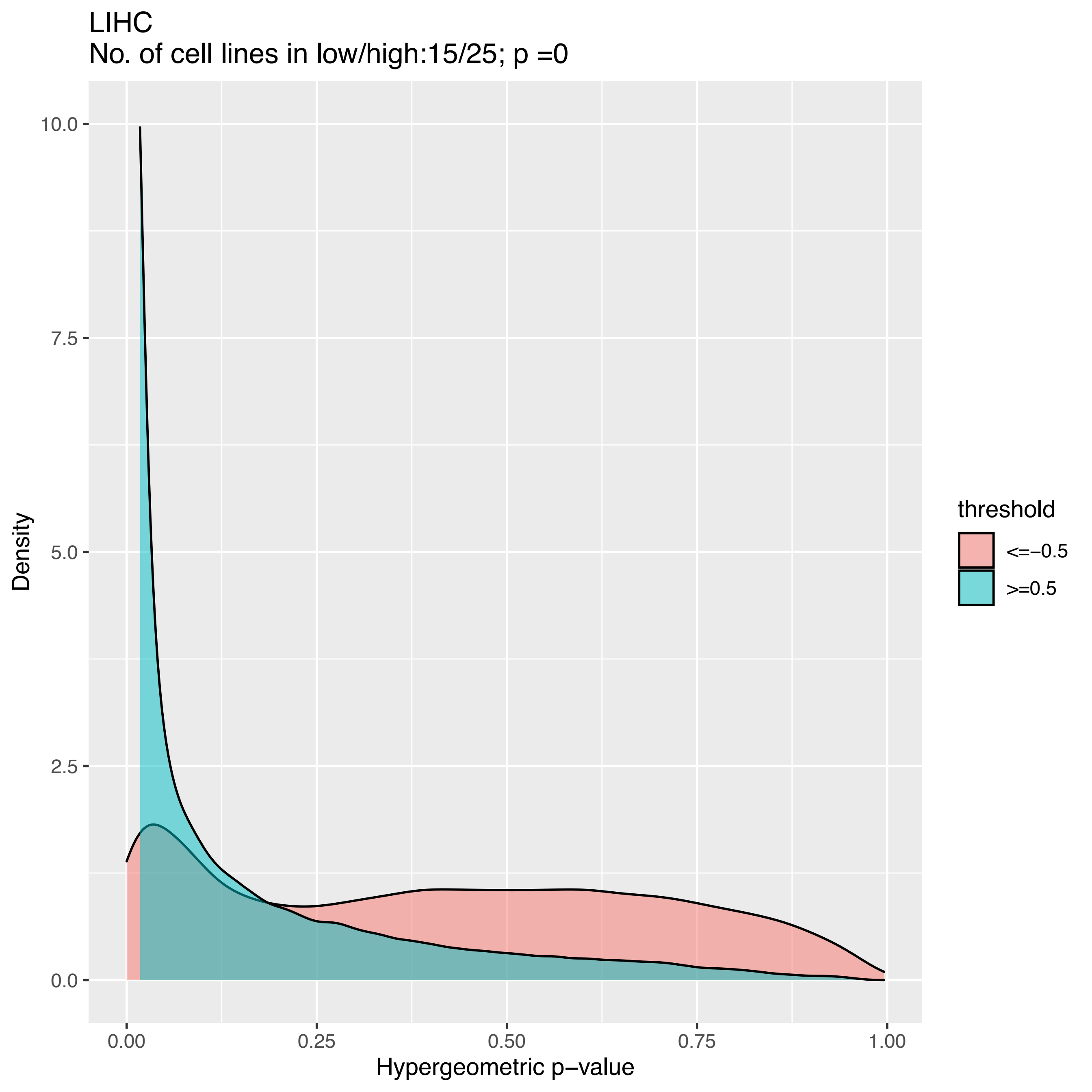

LIHC
No. of cell lines in low/high:15/25; p =0
Density
10.0
7.5
5.0
2.5
0.0
0.00
0.25
0.50
0.75
1.00
Hypergeometric p−value
threshold
<=−0.5
>=0.5

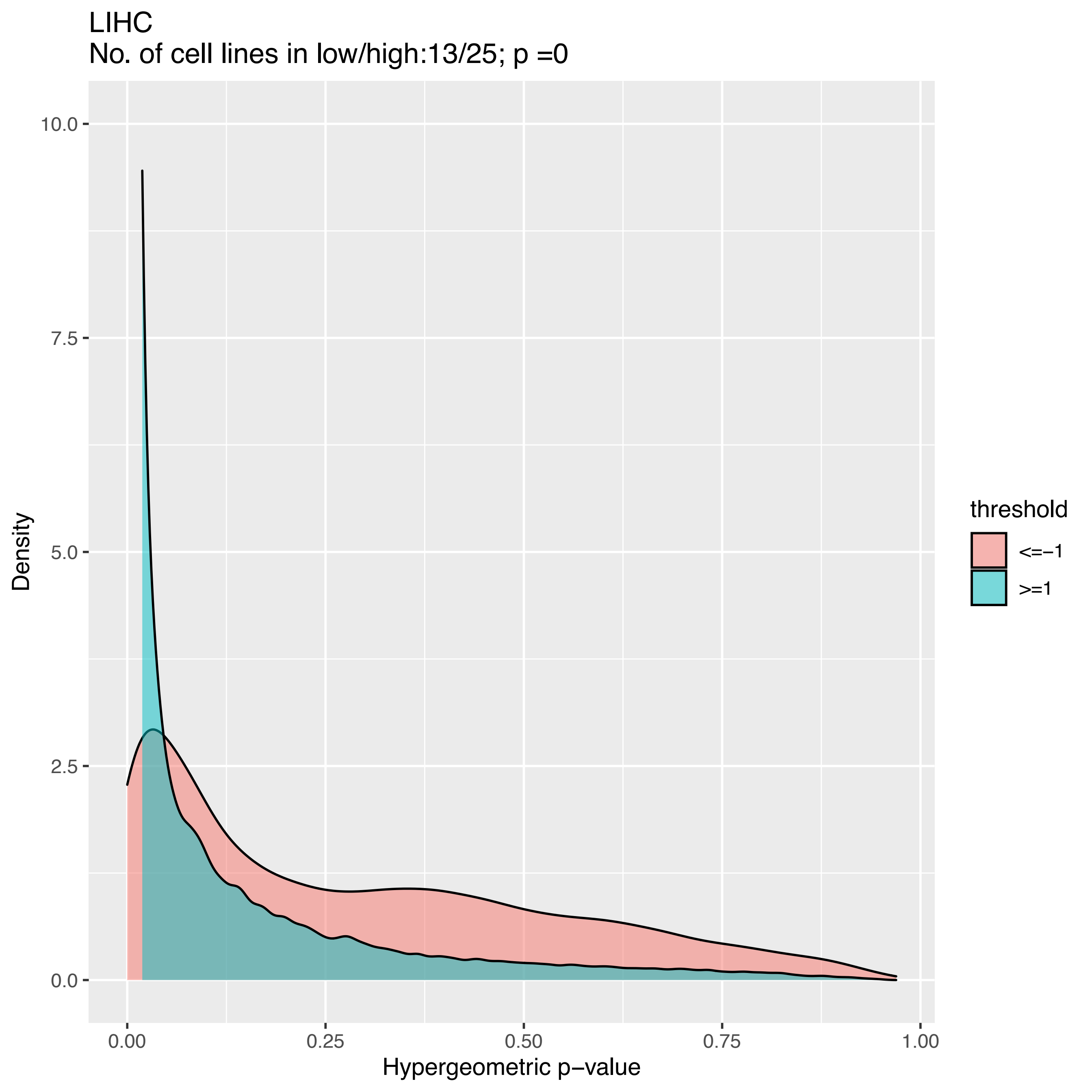
LIHC
No. of cell lines in low/high:13/25; p =0
Density
Hypergeometric p−value
0.0
2.5
5.0
7.5
10.0
0.00
0.25
0.50
0.75
1.00
threshold
<=−1
>=1

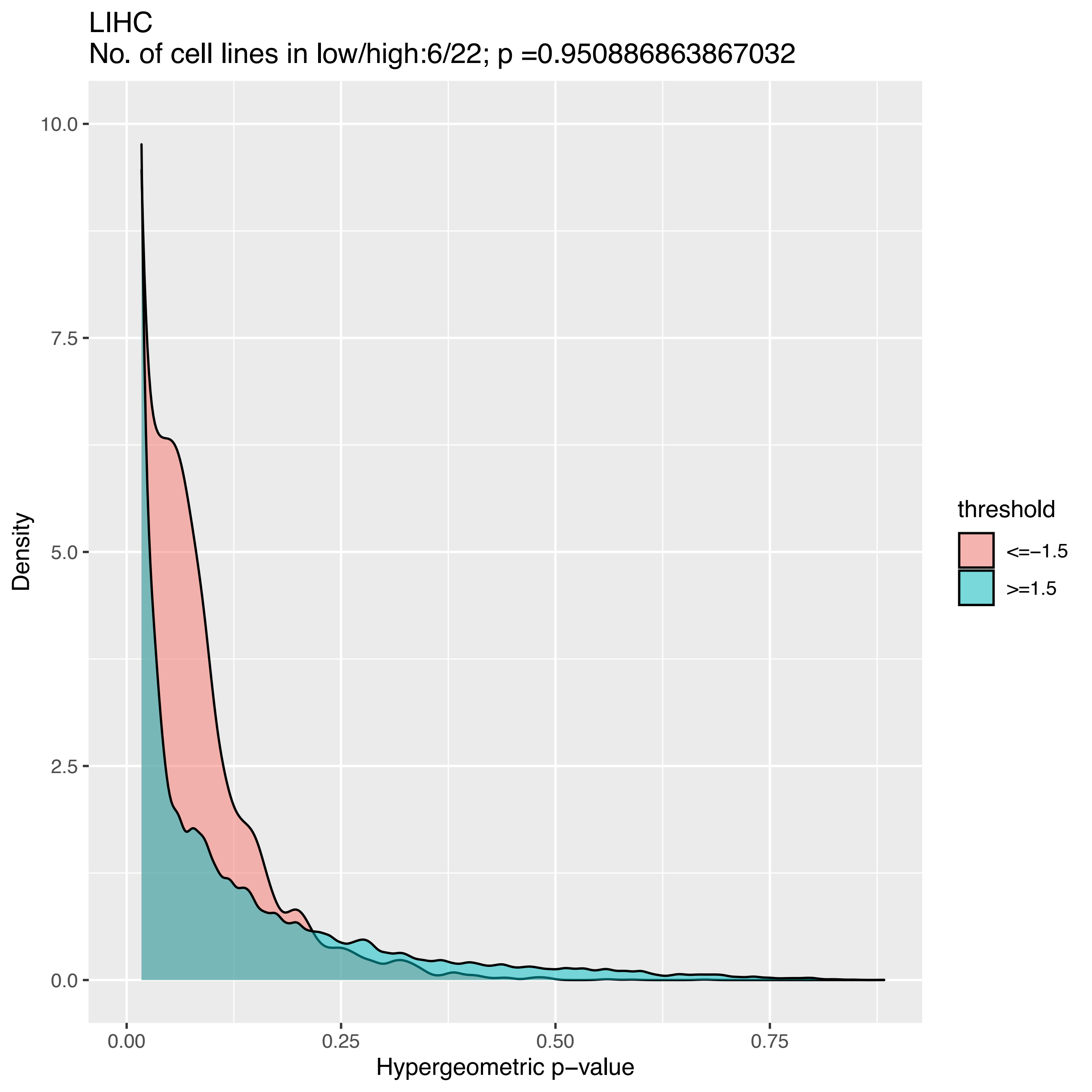

LIHC
No. of cell lines in low/high:6/22; p =0.950886863867032
10.0
7.5
5.0
2.5
0.0
Density
0.00
0.25
0.50
0.75
Hypergeometric p−value
threshold
<=−1.5
>=1.5

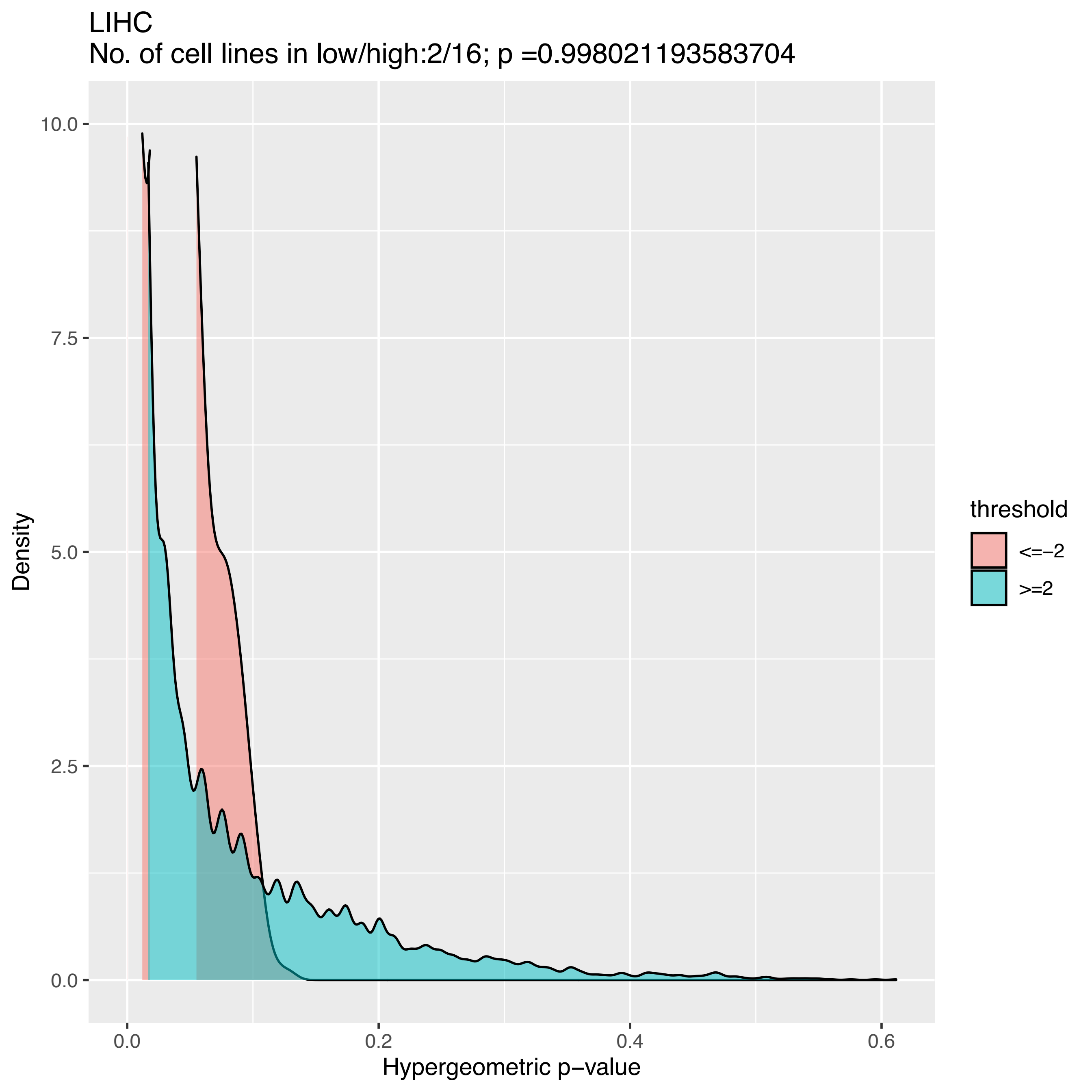

LIHC
No. of cell lines in low/high:2/16; p =0.998021193583704
10.0
7.5
5.0
2.5
0.0
Density
0.0
0.2
0.4
0.6
Hypergeometric p−value
threshold
<=−2
>=2

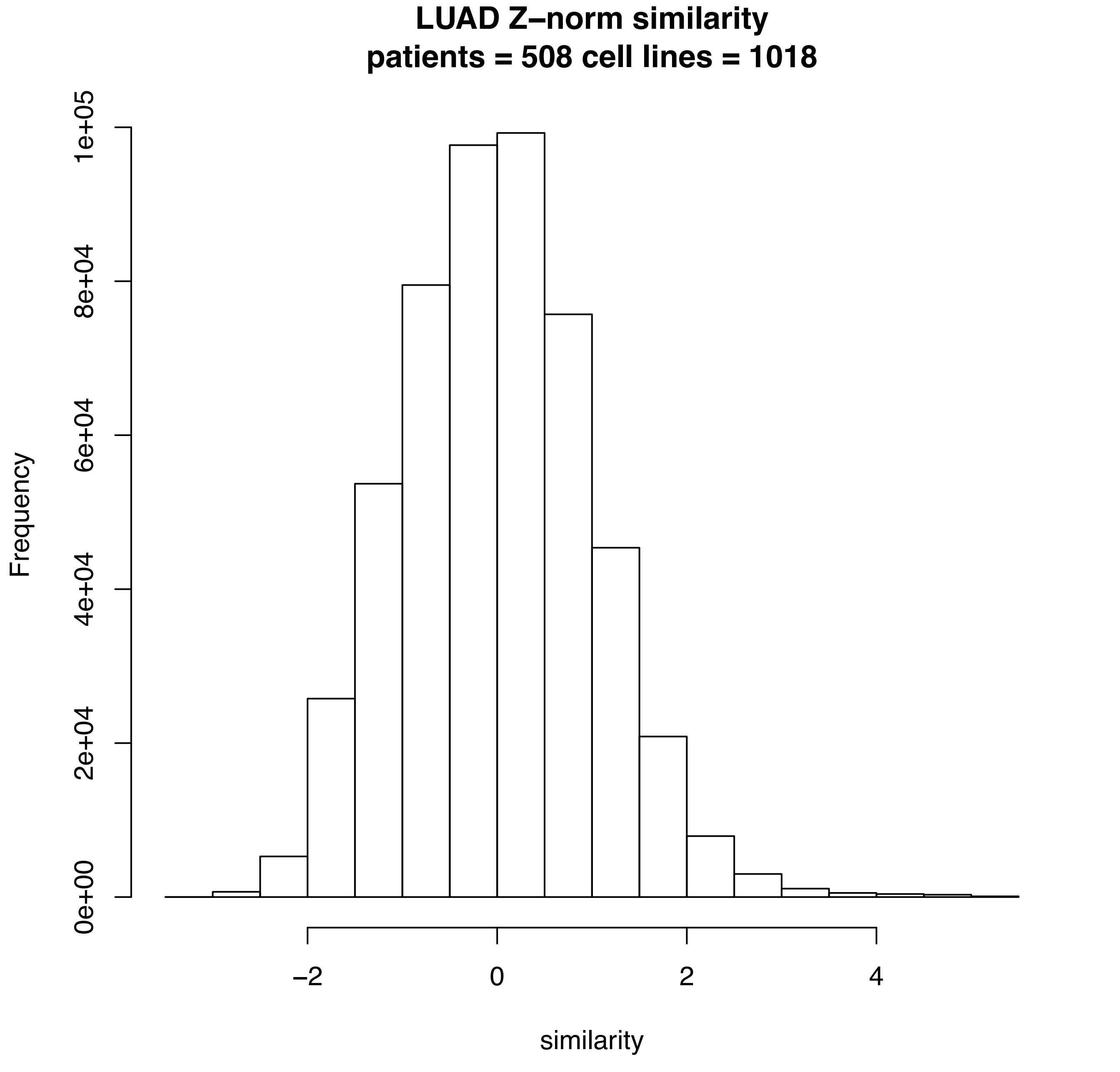
LUAD Z–norm similarity
patients = 508 cell lines = 1018
Frequency
0e+00
2e+04
4e+04
6e+04
8e+04
1e+05
−2
0
2
4
similarity

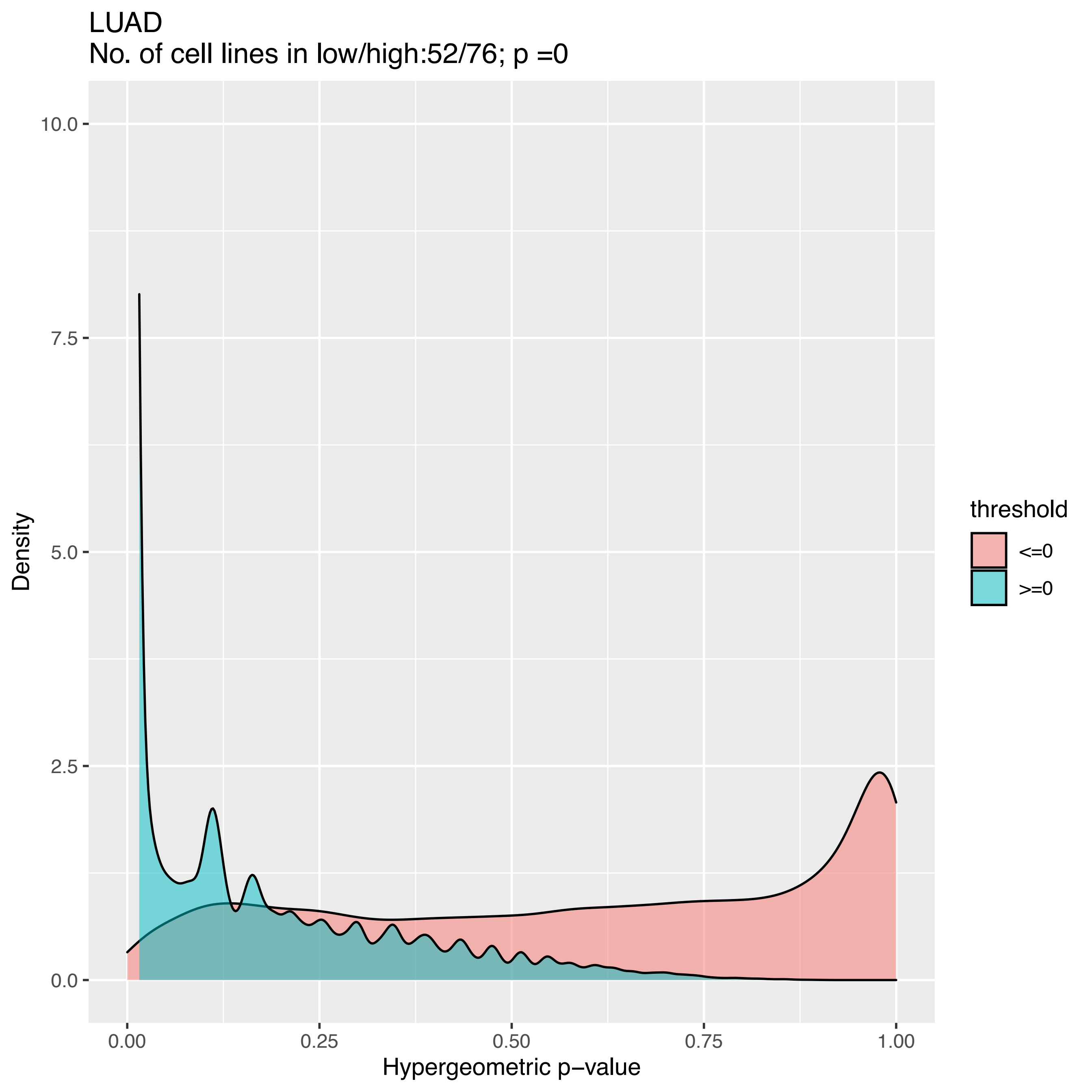
LUAD
No. of cell lines in low/high:52/76; p =0
Density
10.0
7.5
5.0
2.5
0.0
0.00
0.25
0.50
0.75
1.00
Hypergeometric p−value
threshold
<=0
>=0

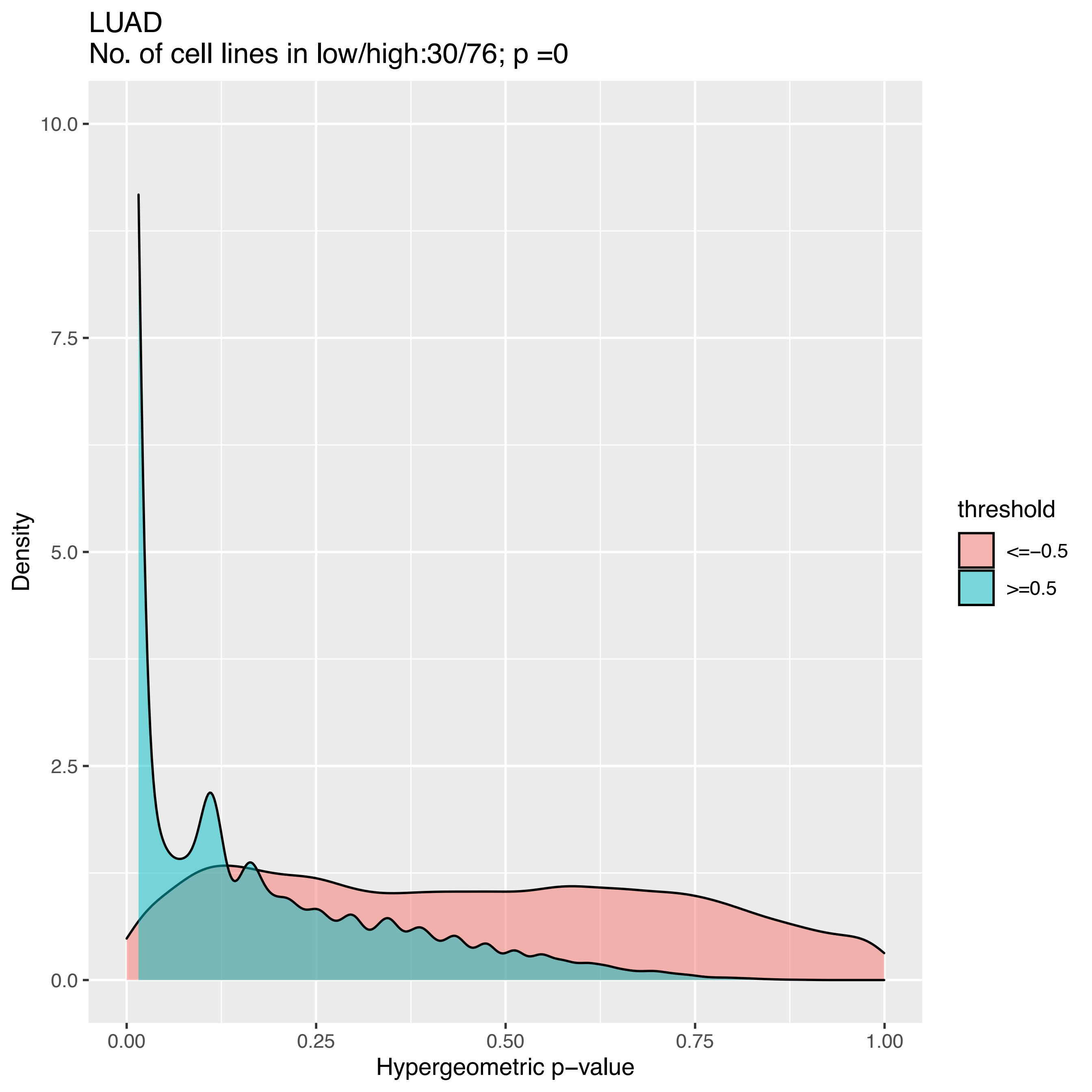

LUAD
No. of cell lines in low/high:30/76; p =0
Density
10.0
7.5
5.0
2.5
0.0
0.00
0.25
0.50
0.75
1.00
Hypergeometric p−value
threshold
<=−0.5
>=0.5

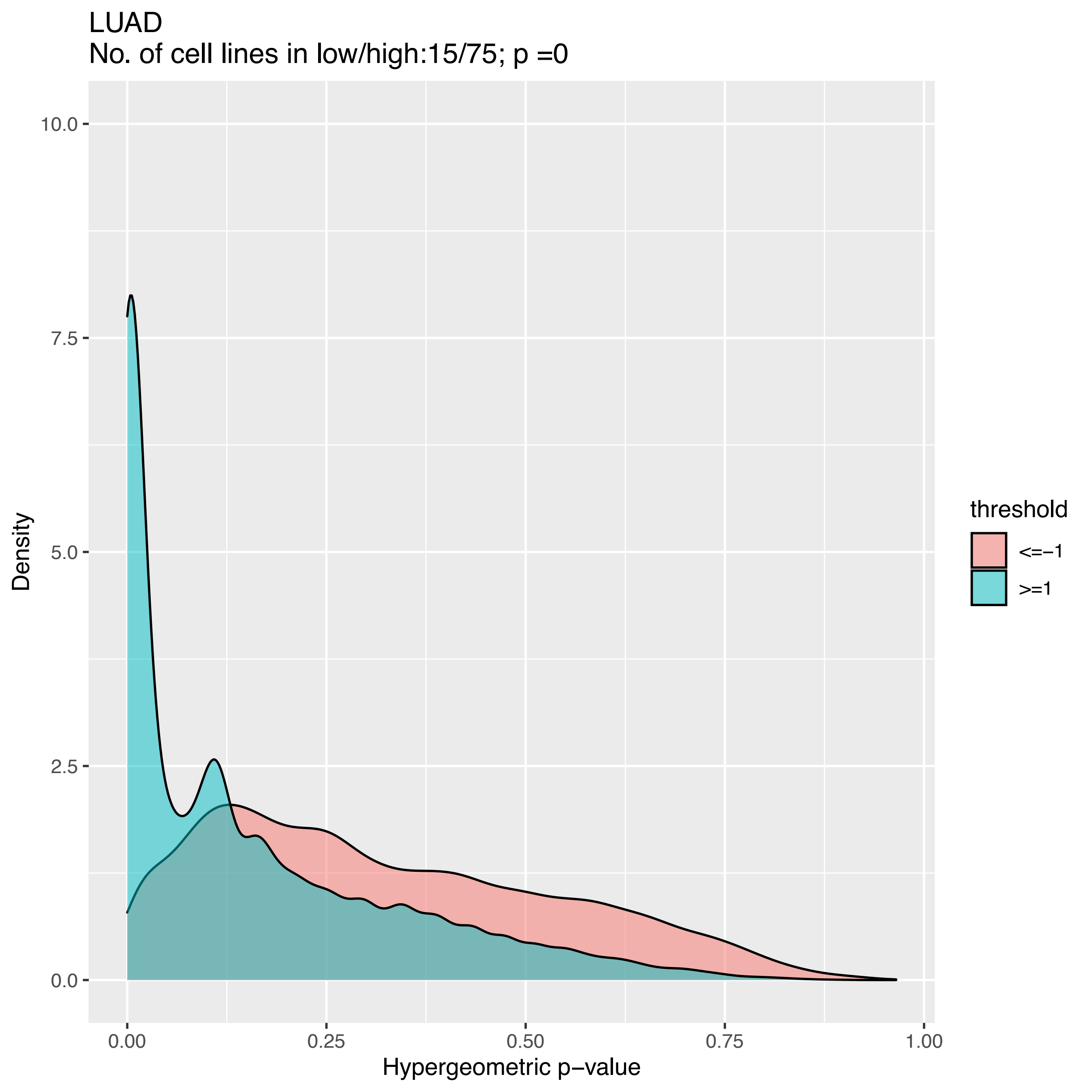
LUAD
No. of cell lines in low/high:15/75; p =0
Density
10.0
7.5
5.0
2.5
0.0
0.00
0.25
0.50
0.75
1.00
Hypergeometric p−value
threshold
<=−1
>=1

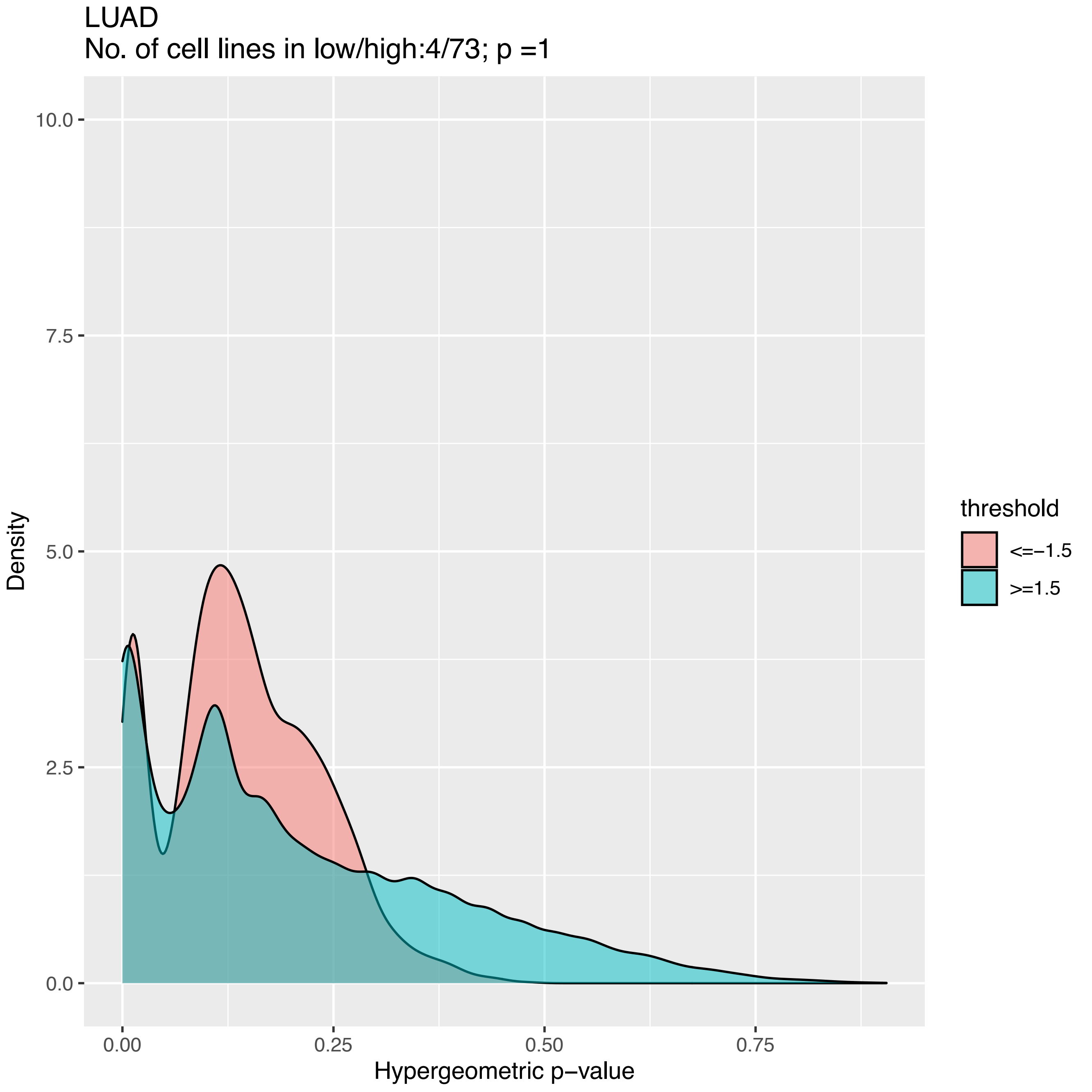
LUAD
No. of cell lines in low/high:4/73; p =1
Density
10.0
7.5
5.0
2.5
0.0
0.00
0.25
0.50
0.75
Hypergeometric p−value
threshold
<=−1.5
>=1.5

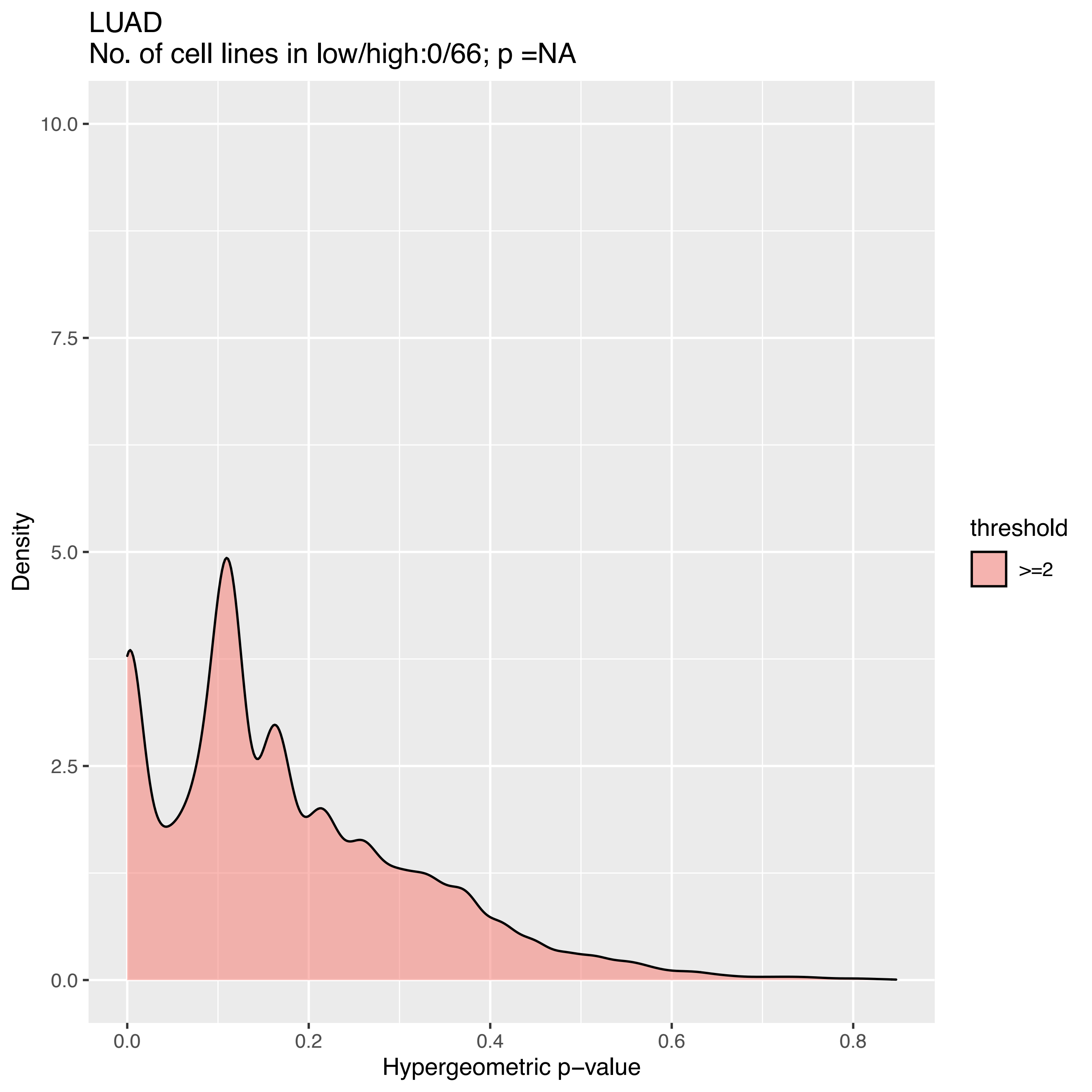
LUAD
No. of cell lines in low/high:0/66; p =NA
Density
10.0
7.5
5.0
2.5
0.0
0.0
0.2
0.4
0.6
0.8
Hypergeometric p−value
threshold
>=2

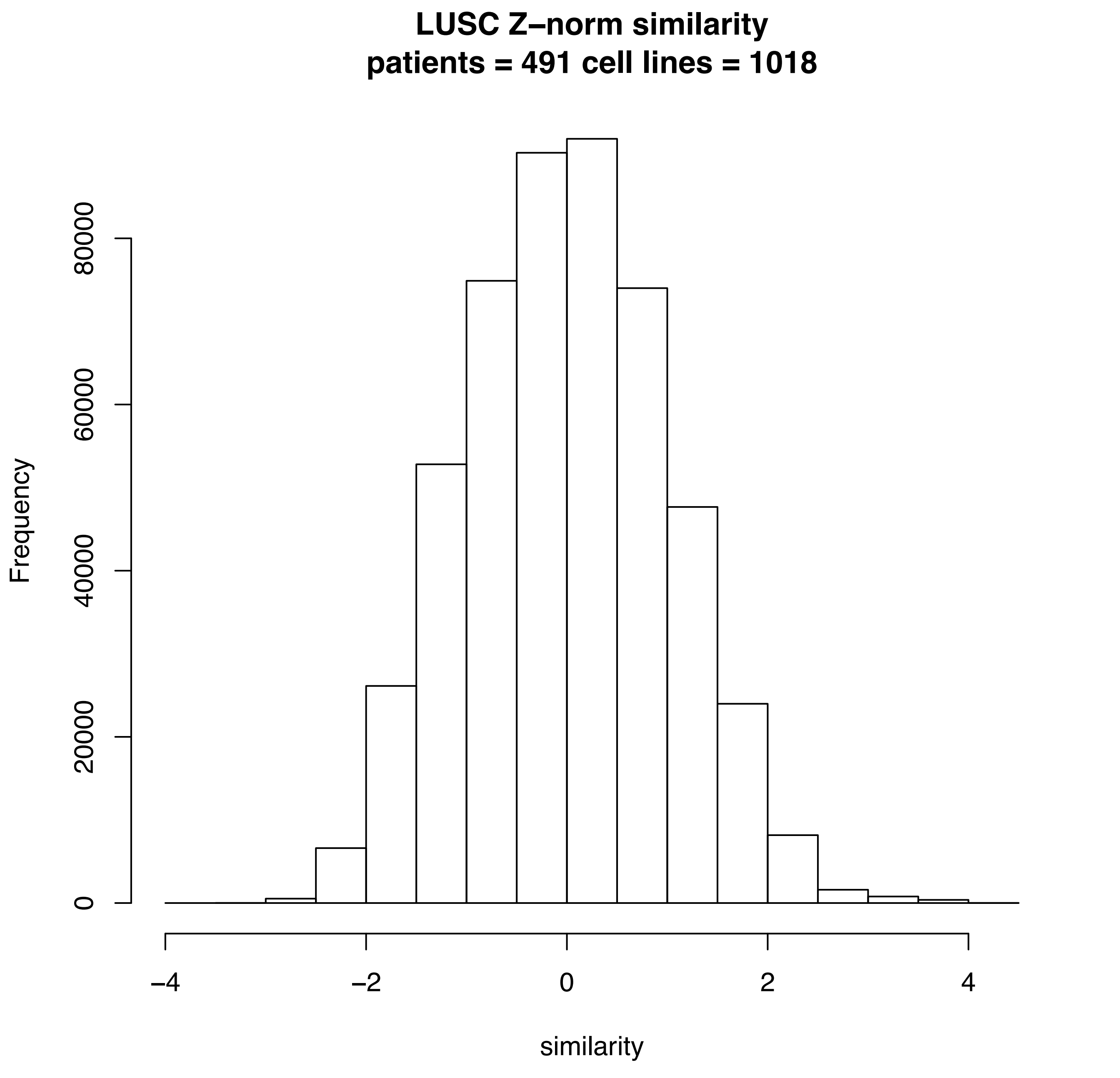
LUSC Z–norm similarity
patients = 491 cell lines = 1018
Frequency
0
20000
40000
60000
80000
–4
–2
0
2
4
similarity

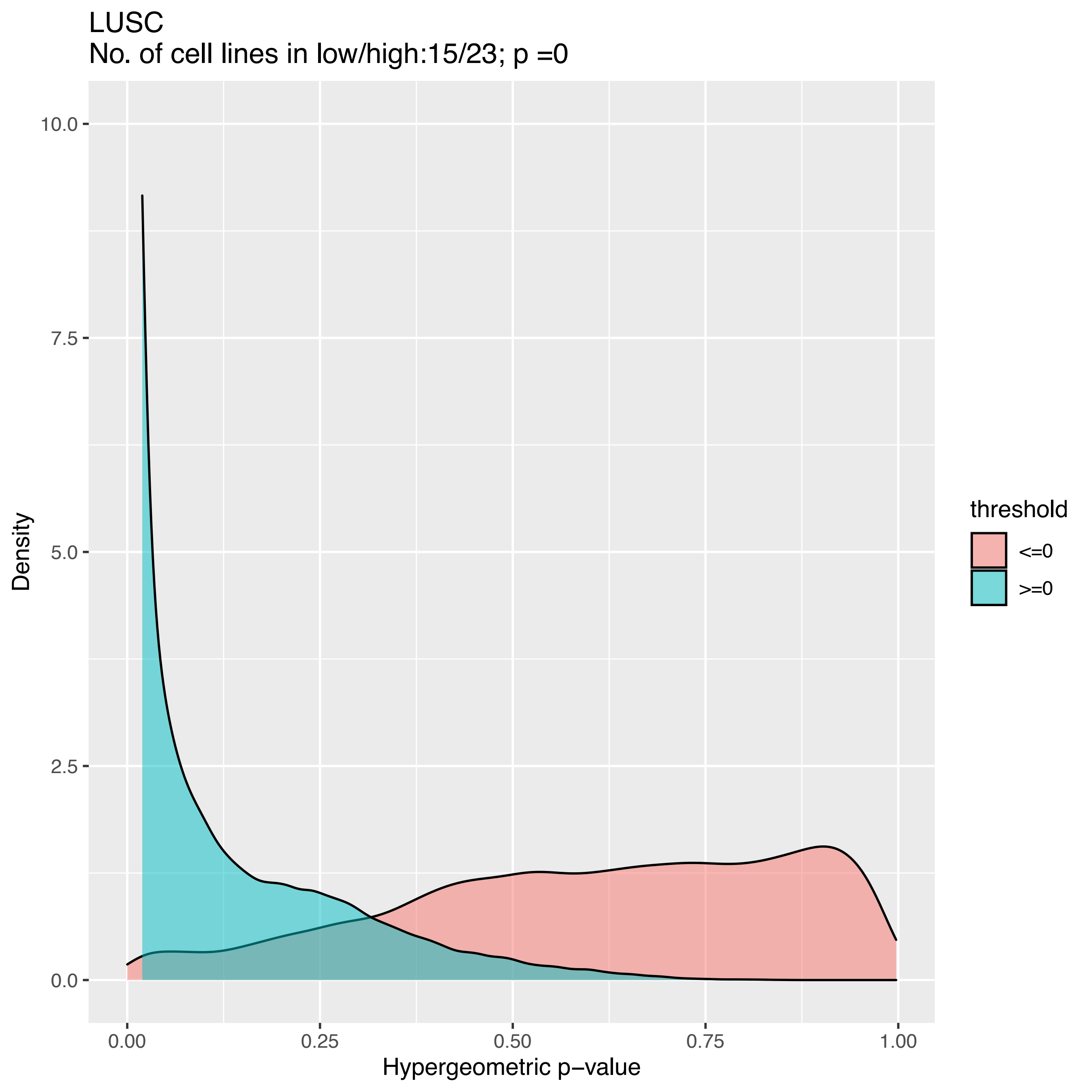
LUSC
No. of cell lines in low/high:15/23; p =0
Density
10.0
7.5
5.0
2.5
0.0
0.00
0.25
0.50
0.75
1.00
Hypergeometric p−value
threshold
<=0
>=0

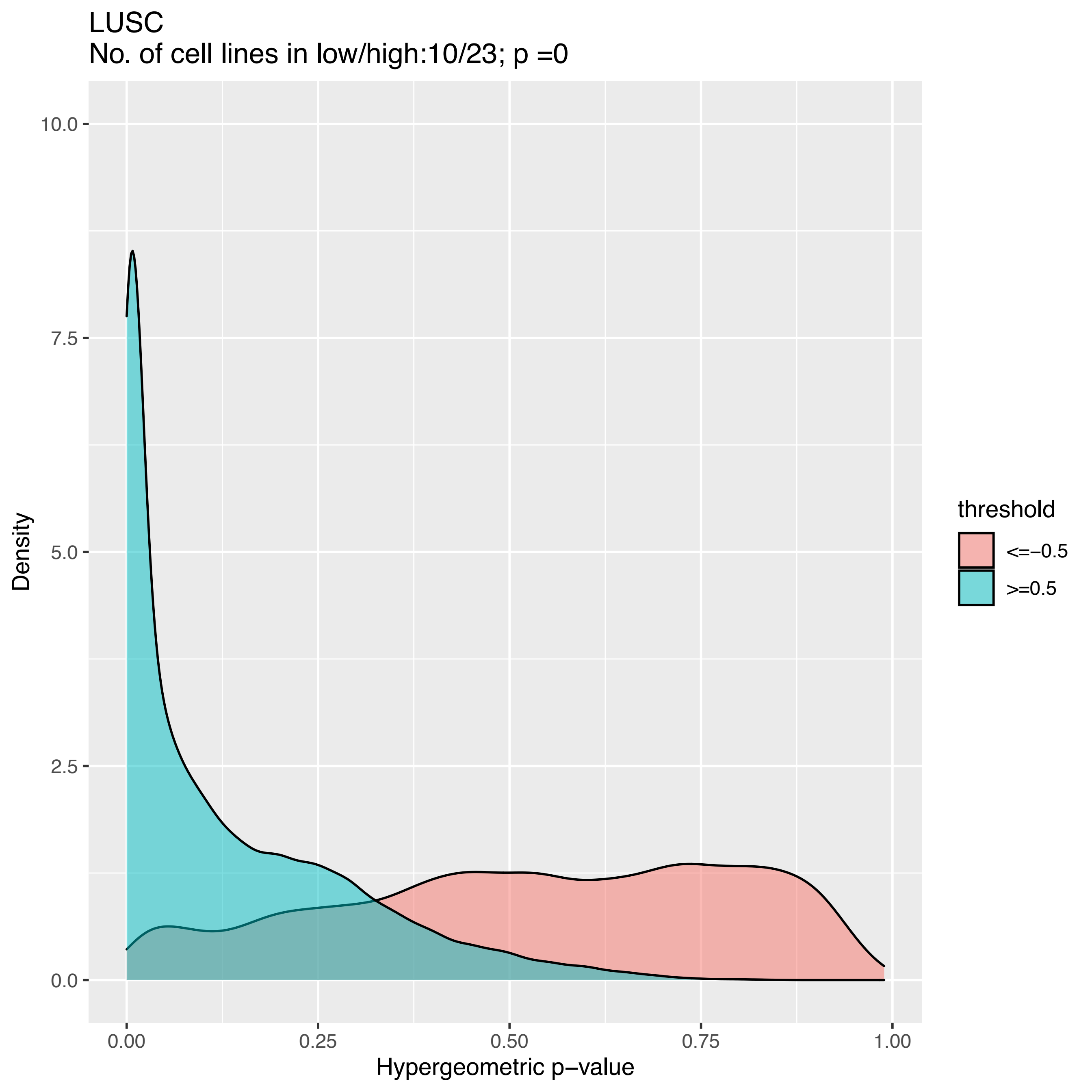
LUSC
No. of cell lines in low/high:10/23; p =0
Density
10.0
7.5
5.0
2.5
0.0
0.00
0.25
0.50
0.75
1.00
Hypergeometric p−value
threshold
<=−0.5
>=0.5

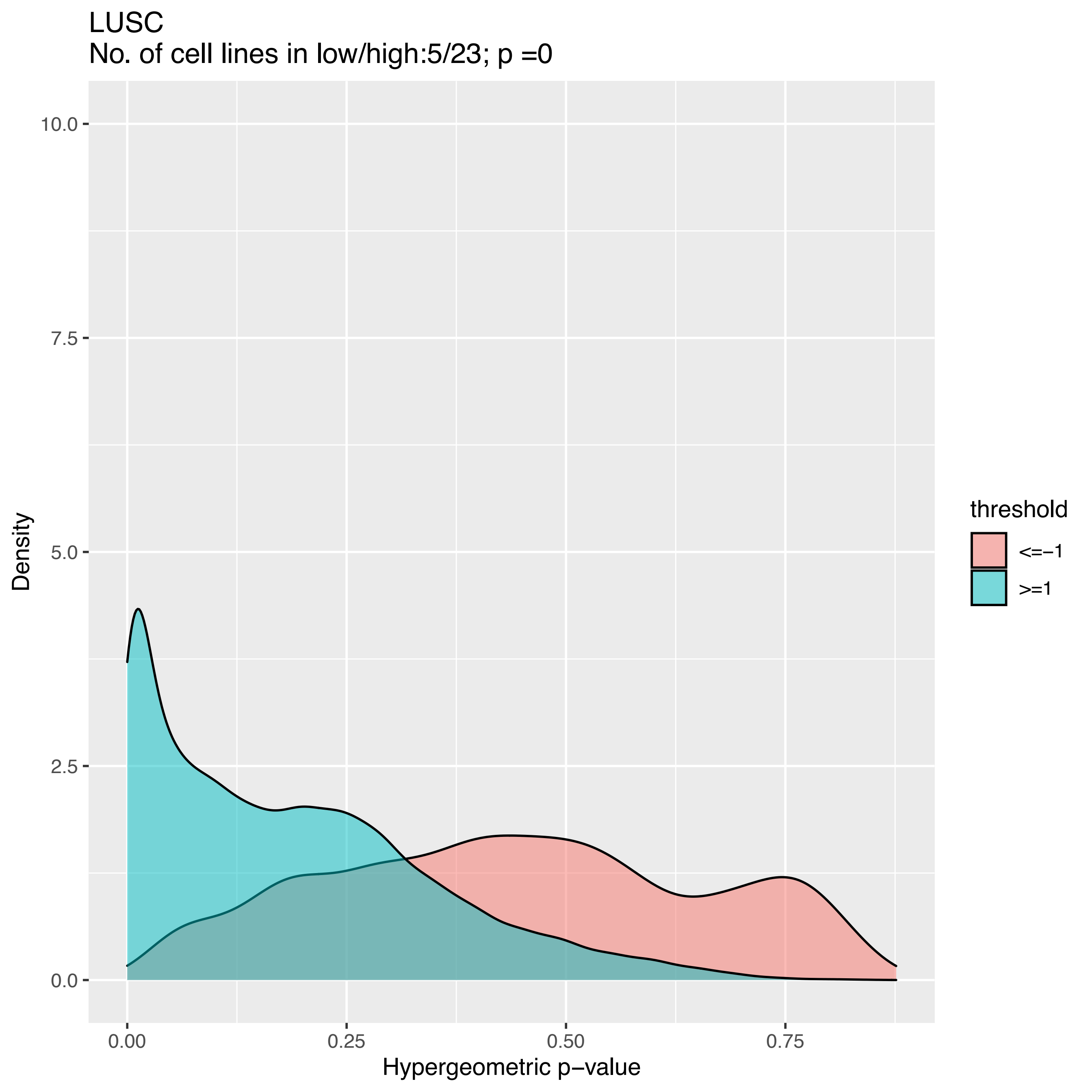
LUSC
No. of cell lines in low/high:5/23; p =0
Density
Hypergeometric p−value
threshold
<=−1
>=1
10.0
7.5
5.0
2.5
0.0
0.00
0.25
0.50
0.75

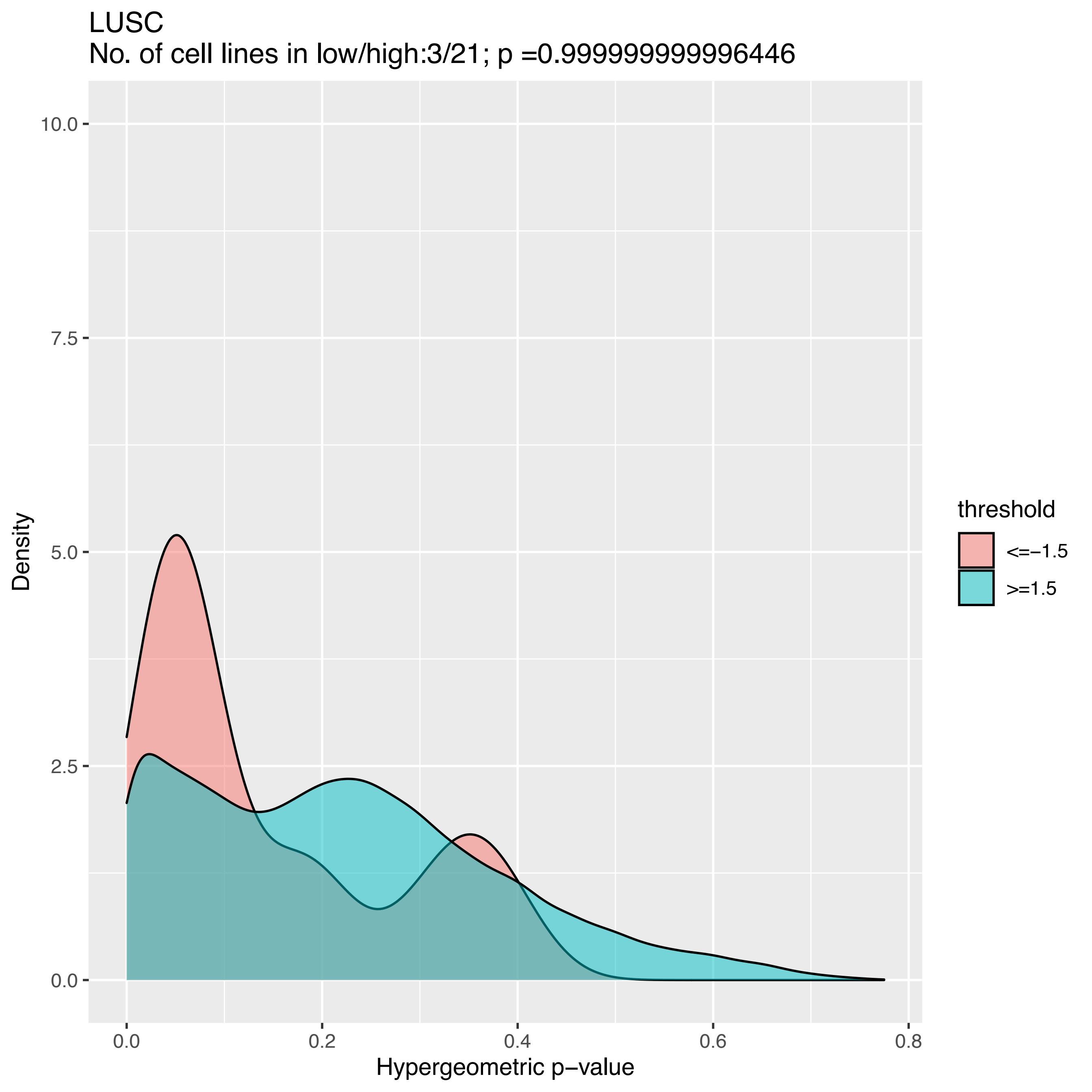

LUSC
No. of cell lines in low/high:3/21; p =0.999999999996446
10.0
7.5
5.0
2.5
0.0
Density
0.0
0.2
0.4
0.6
0.8
Hypergeometric p−value
threshold
<=−1.5
>=1.5

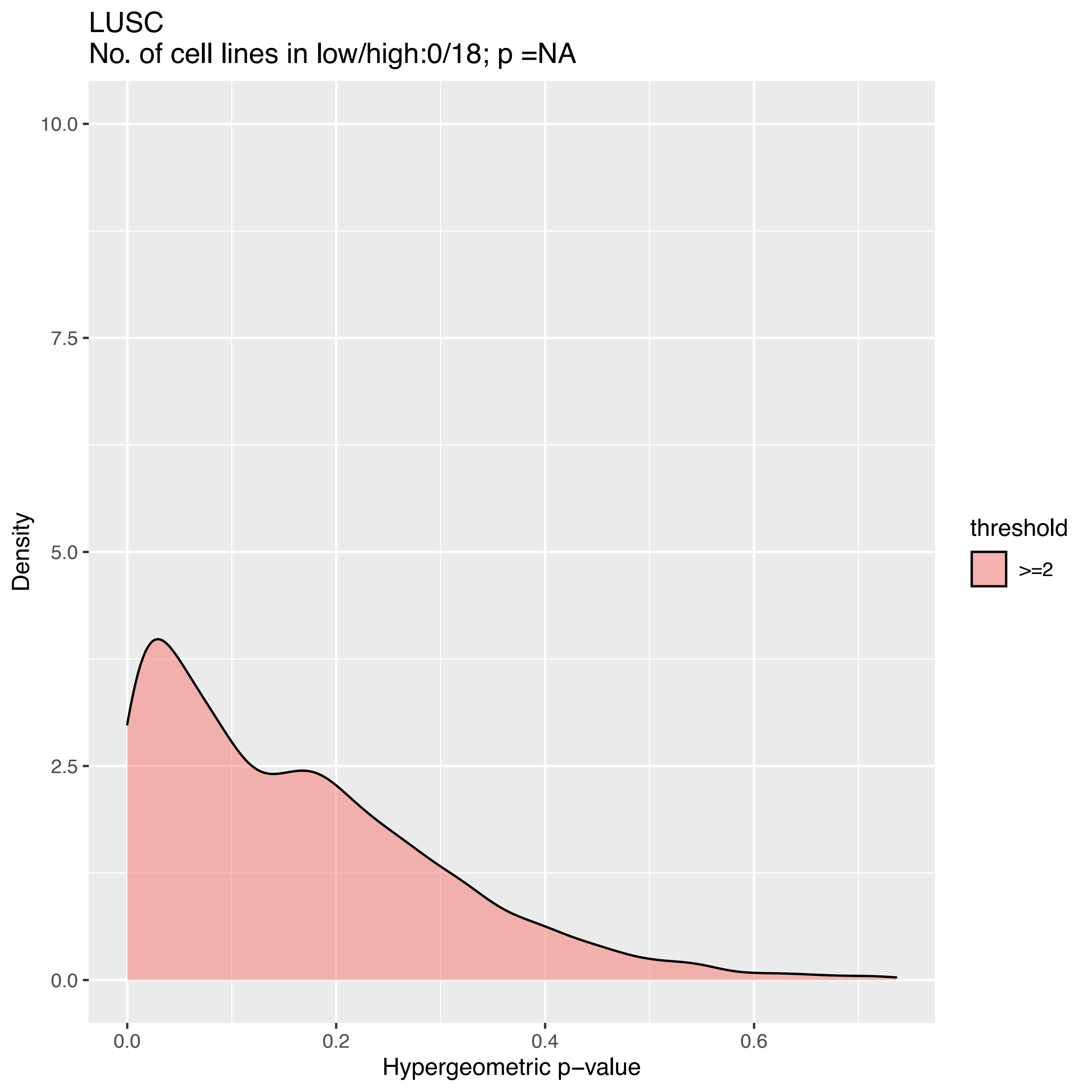
LUSC
No. of cell lines in low/high:0/18; p =NA
Density
10.0
7.5
5.0
2.5
0.0
0.0
0.2
0.4
0.6
Hypergeometric p−value
threshold
>=2

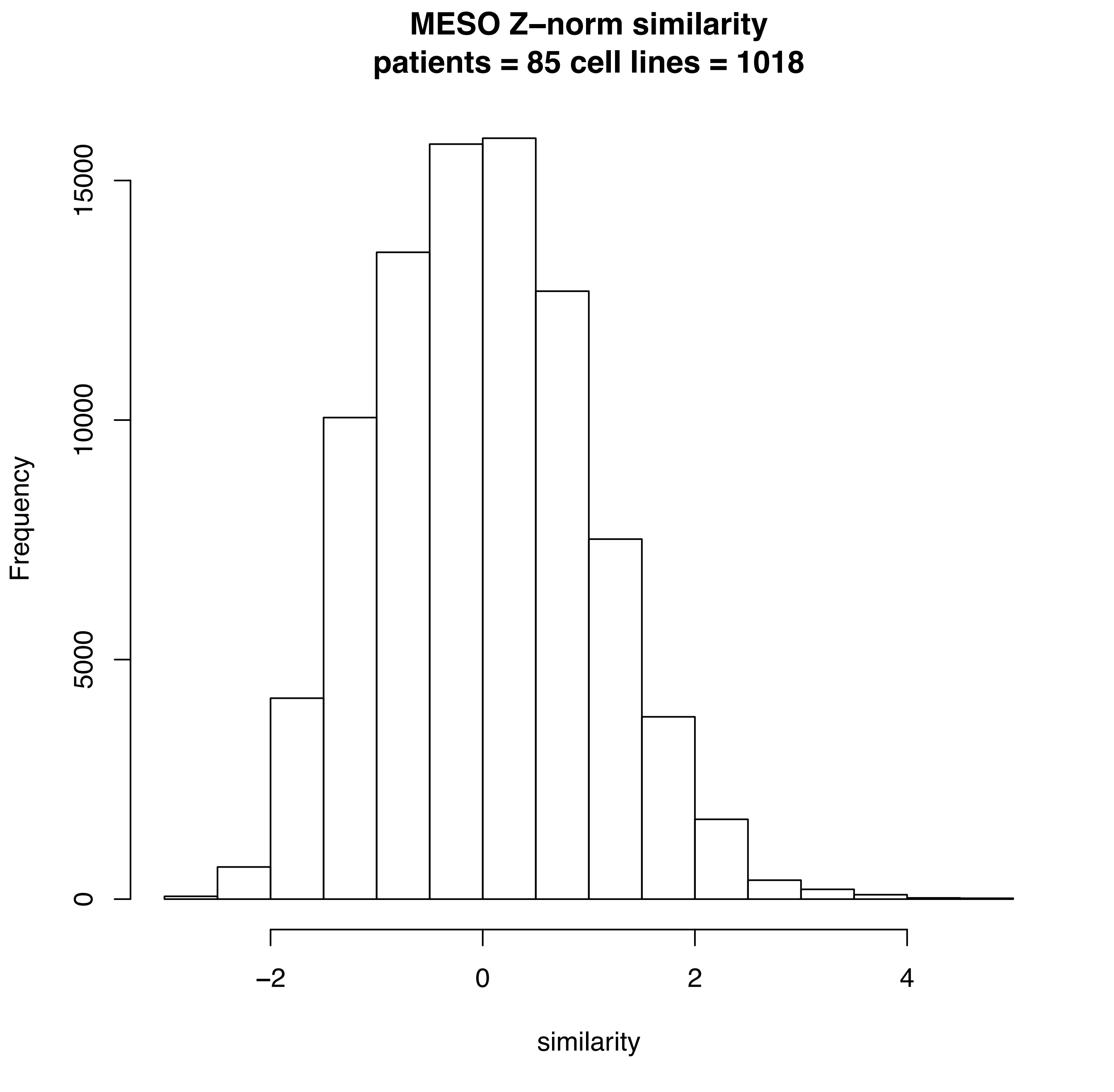
MESO Z–norm similarity
patients = 85 cell lines = 1018
Frequency
0
5000
10000
15000
−2
0
2
4
similarity

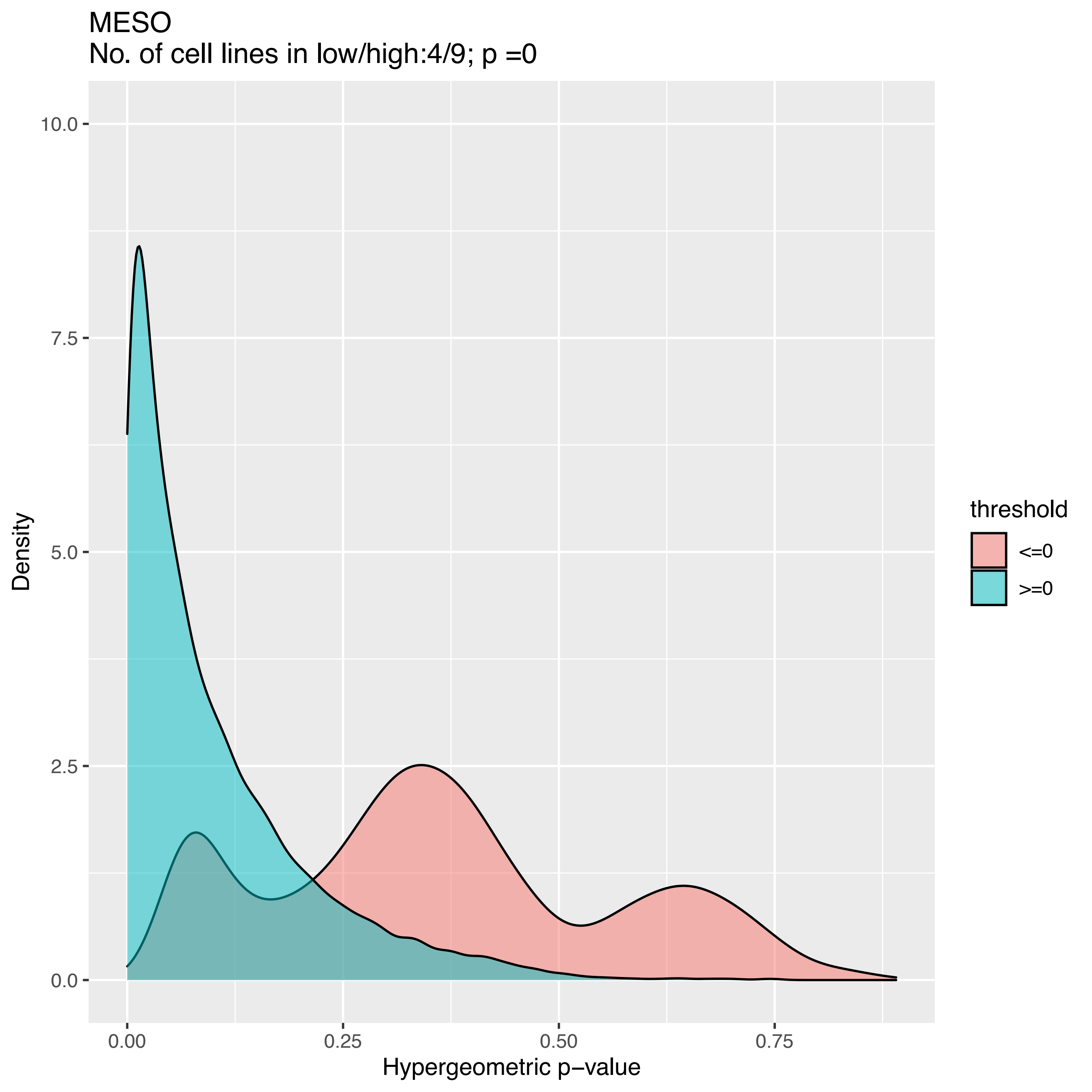

MESO
No. of cell lines in low/high:4/9; p =0
Density
10.0
7.5
5.0
2.5
0.0
0.00
0.25
0.50
0.75
Hypergeometric p−value
threshold
<=0
>=0

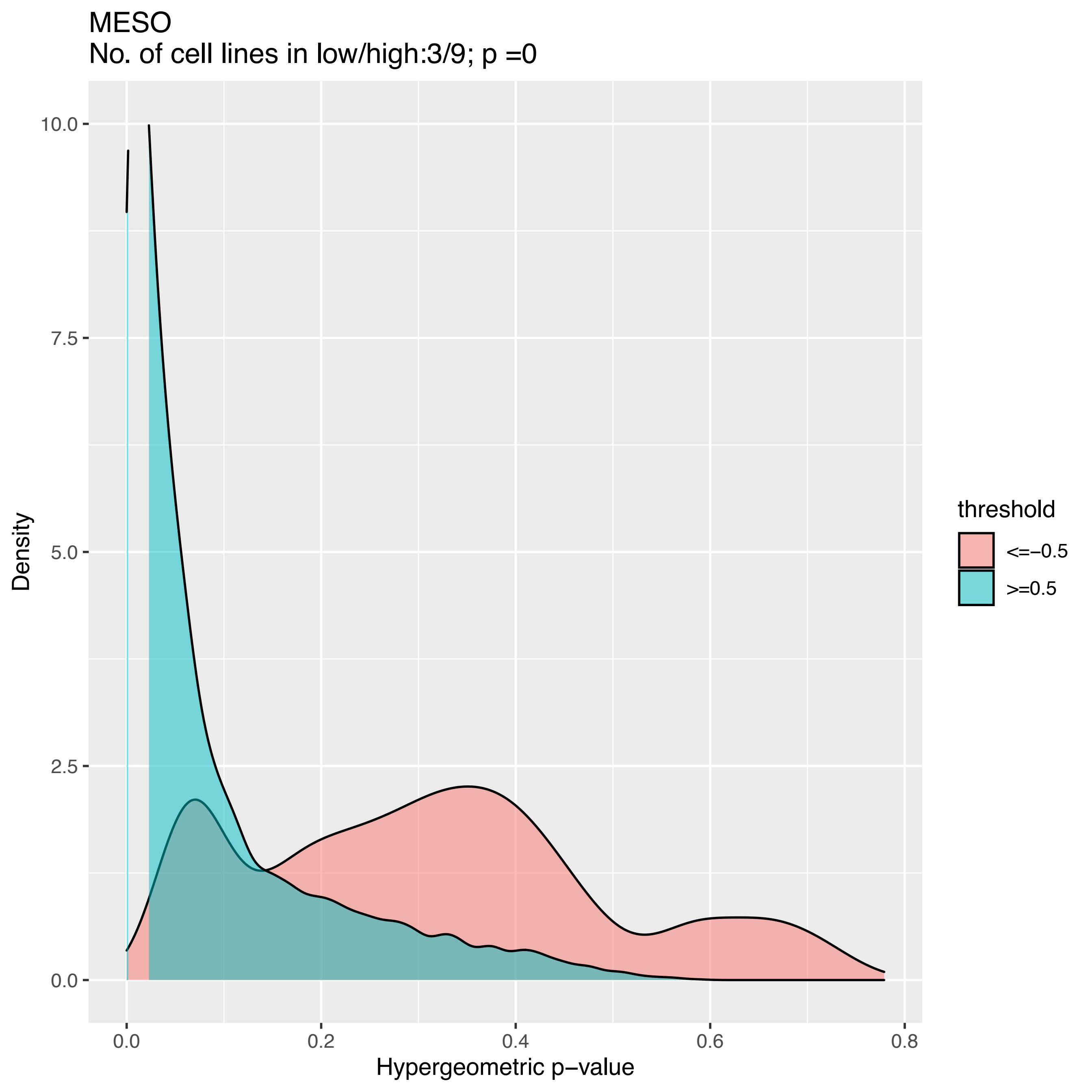
MESO
No. of cell lines in low/high:3/9; p =0
Density
10.0
7.5
5.0
2.5
0.0
0.0
0.2
0.4
0.6
0.8
Hypergeometric p−value
threshold
<=−0.5
>=0.5

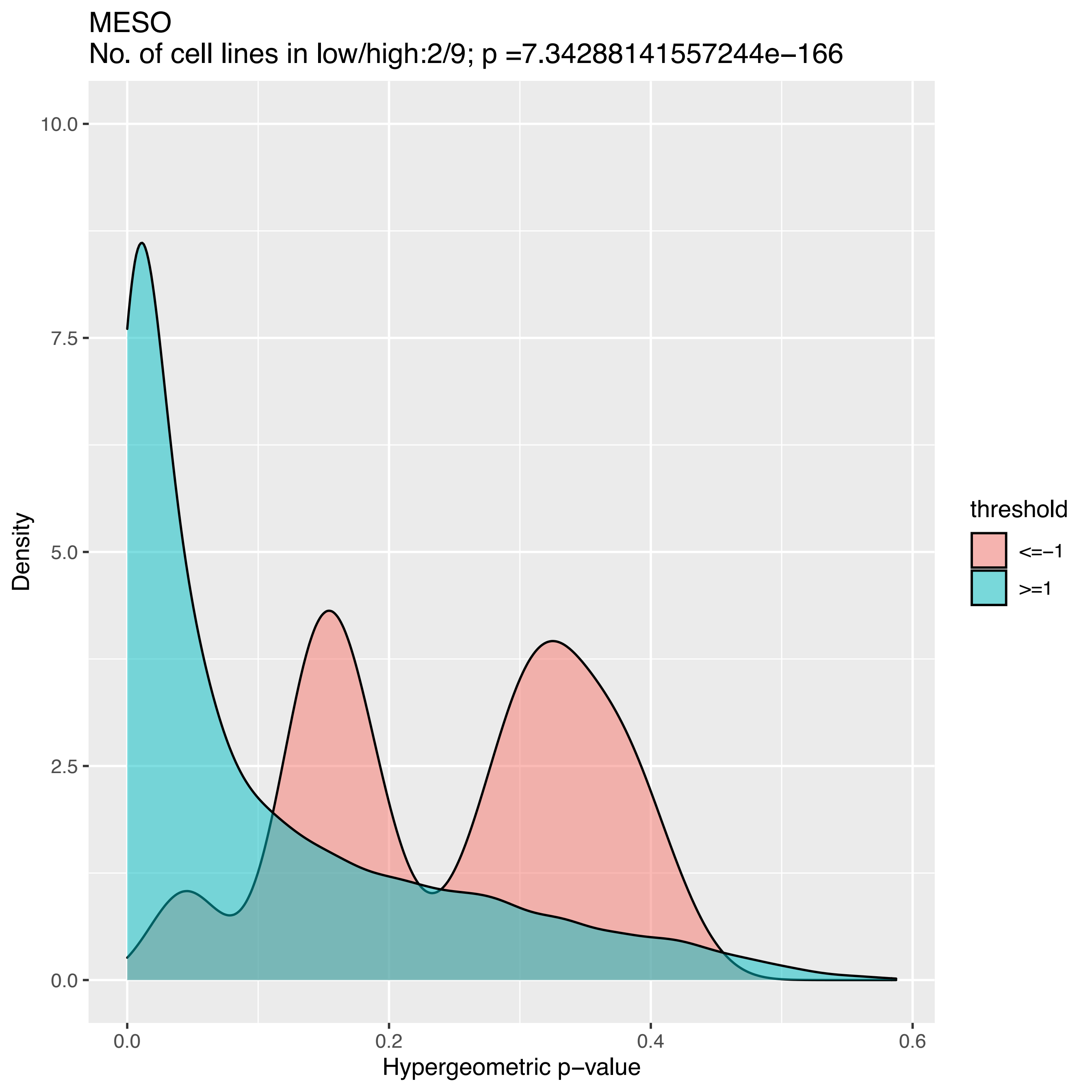

MESO
No. of cell lines in low/high:2/9; p =7.34288141557244e−166
Density
Hypergeometric p−value
0.0
2.5
5.0
7.5
10.0
0.0
0.2
0.4
0.6
threshold
<=−1
>=1

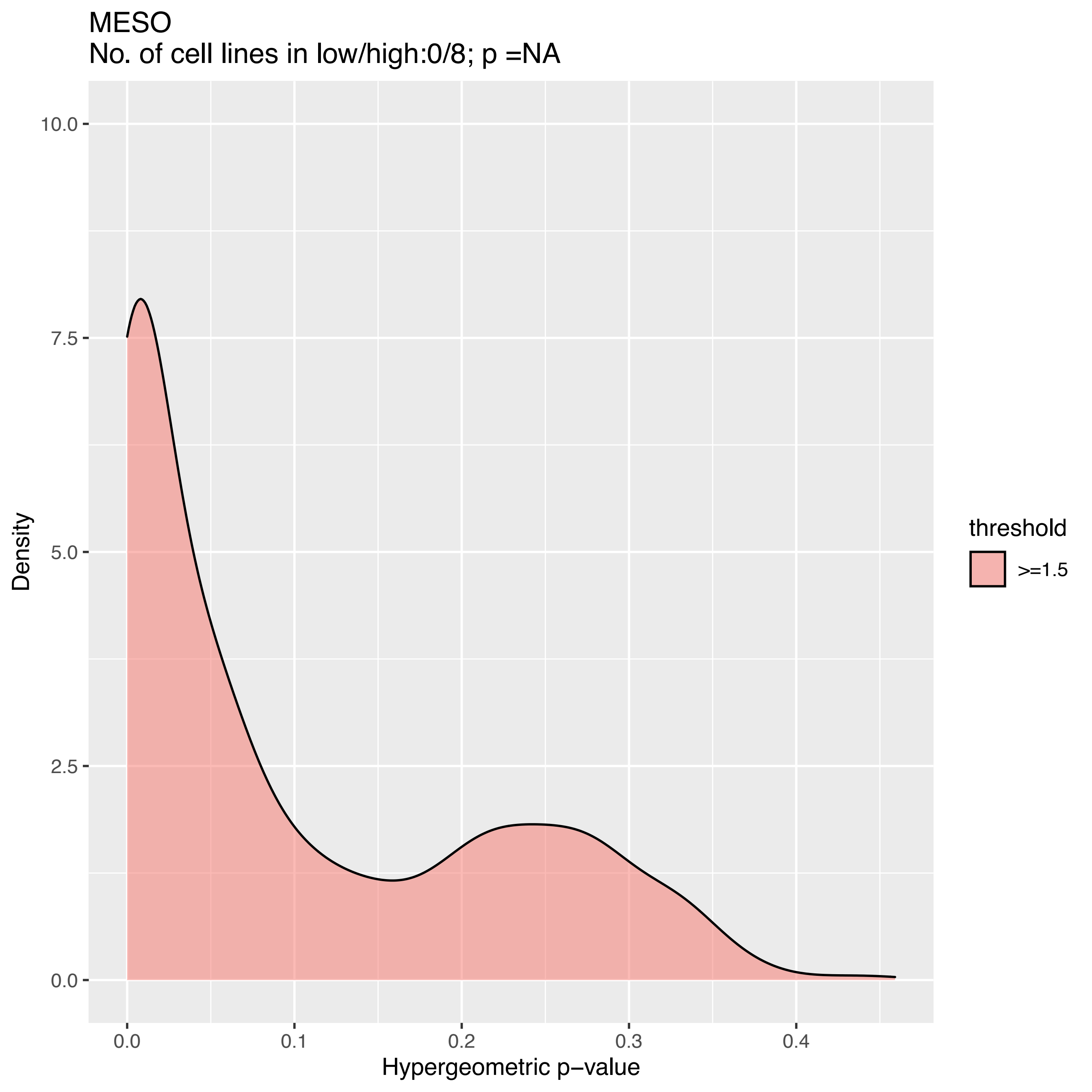

MESO
No. of cell lines in low/high:0/8; p =NA
Density
10.0
7.5
5.0
2.5
0.0
0.0
0.1
0.2
0.3
0.4
Hypergeometric p−value
threshold
>=1.5

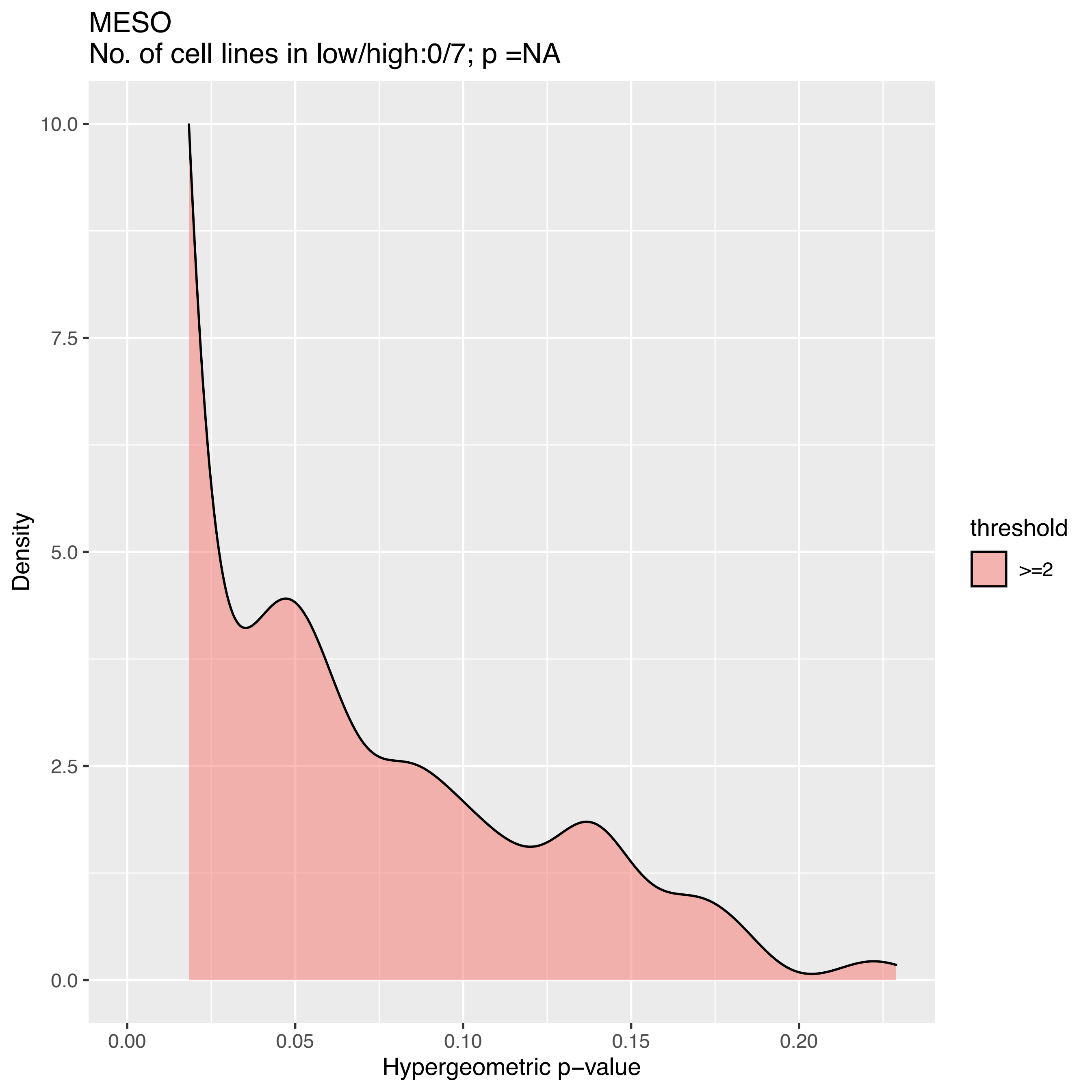

MESO
No. of cell lines in low/high:0/7; p =NA
10.0
7.5
5.0
2.5
0.0
Density
0.00
0.05
0.10
0.15
0.20
Hypergeometric p−value
threshold
>=2

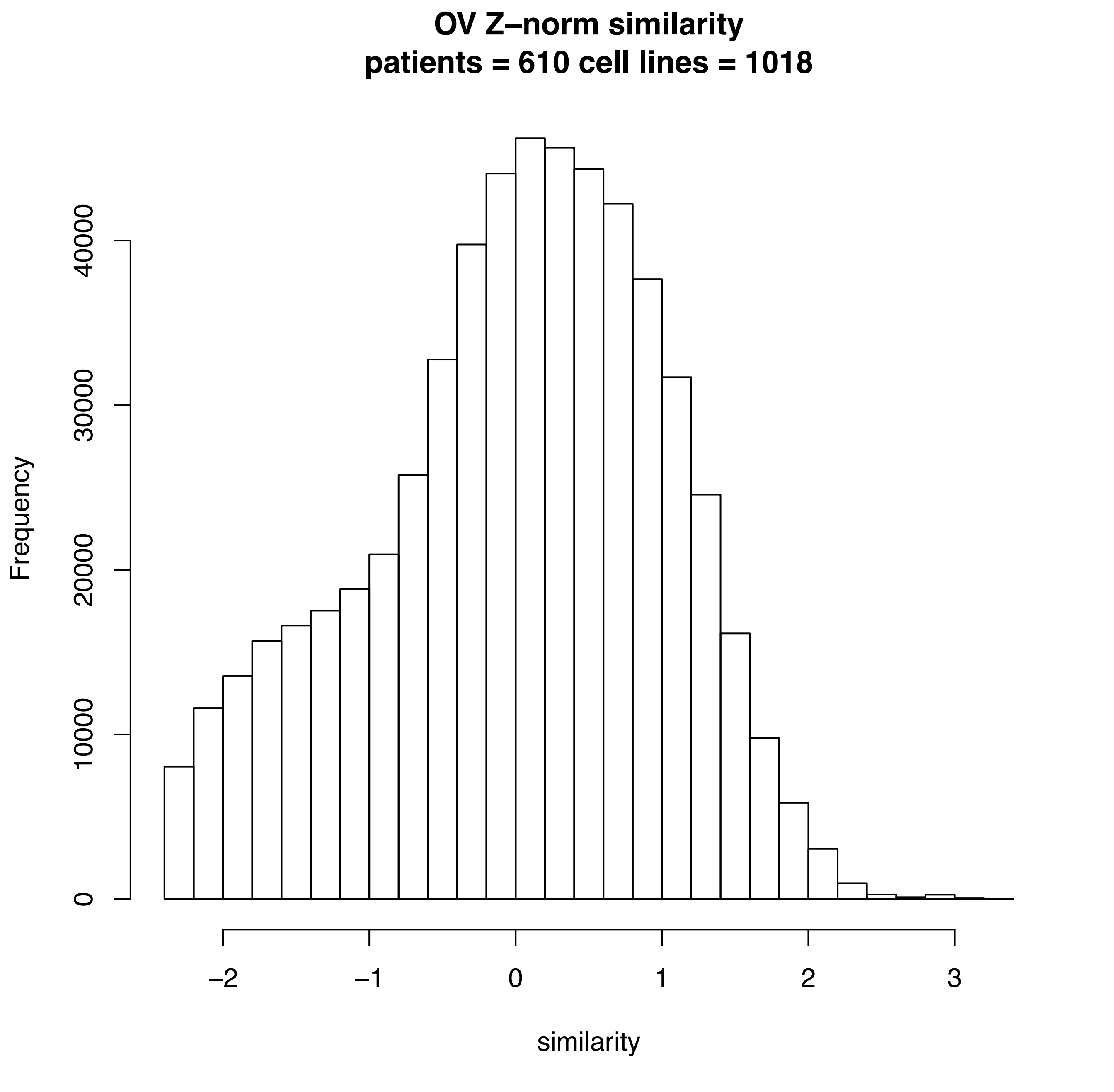
OV Z−norm similarity
patients = 610 cell lines = 1018
Frequency
0
10000
20000
30000
40000
−2
−1
0
1
2
3
similarity

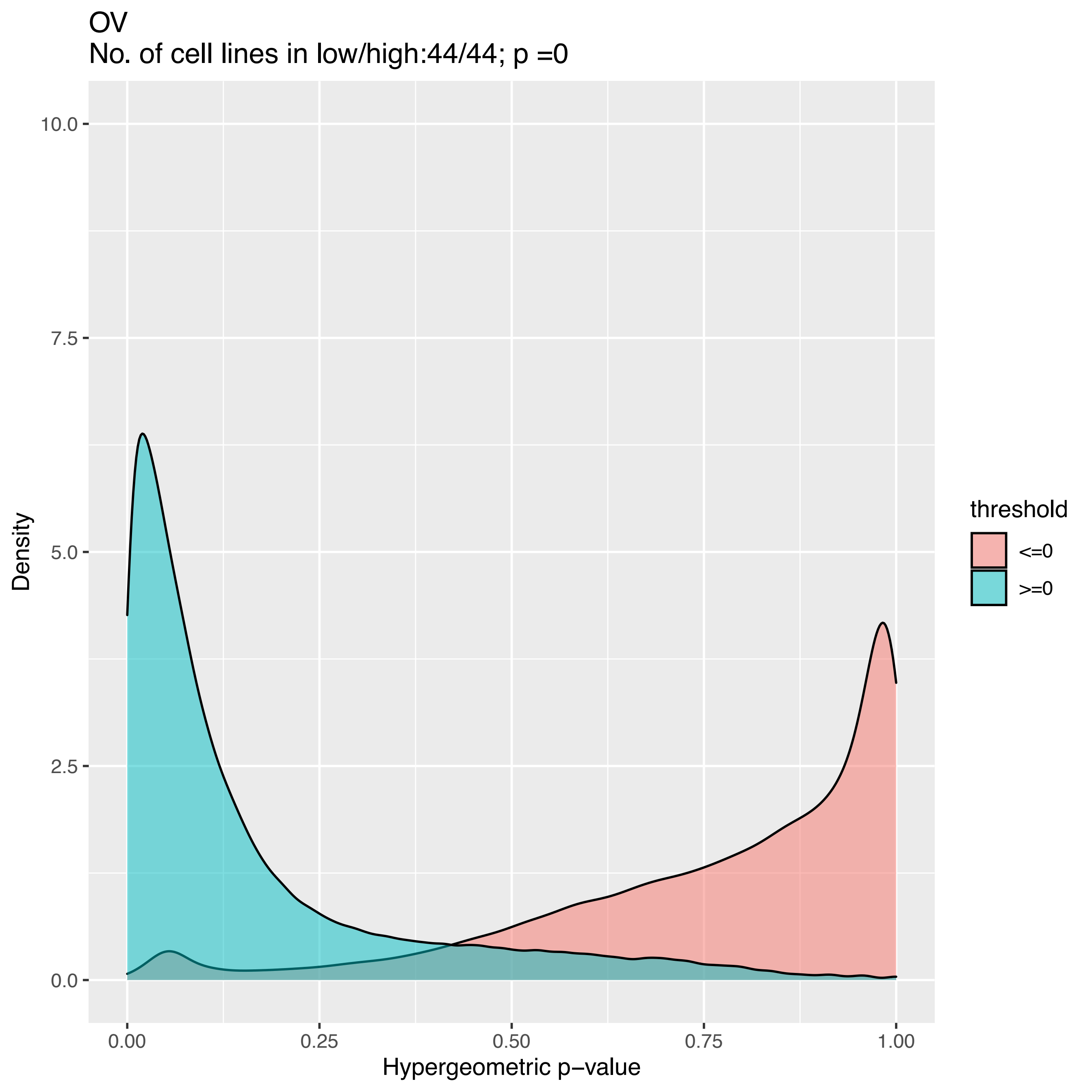

OV
No. of cell lines in low/high:44/44; p =0
10.0
7.5
5.0
2.5
0.0
Density
0.00
0.25
0.50
0.75
1.00
Hypergeometric p−value
threshold
<=0
>=0

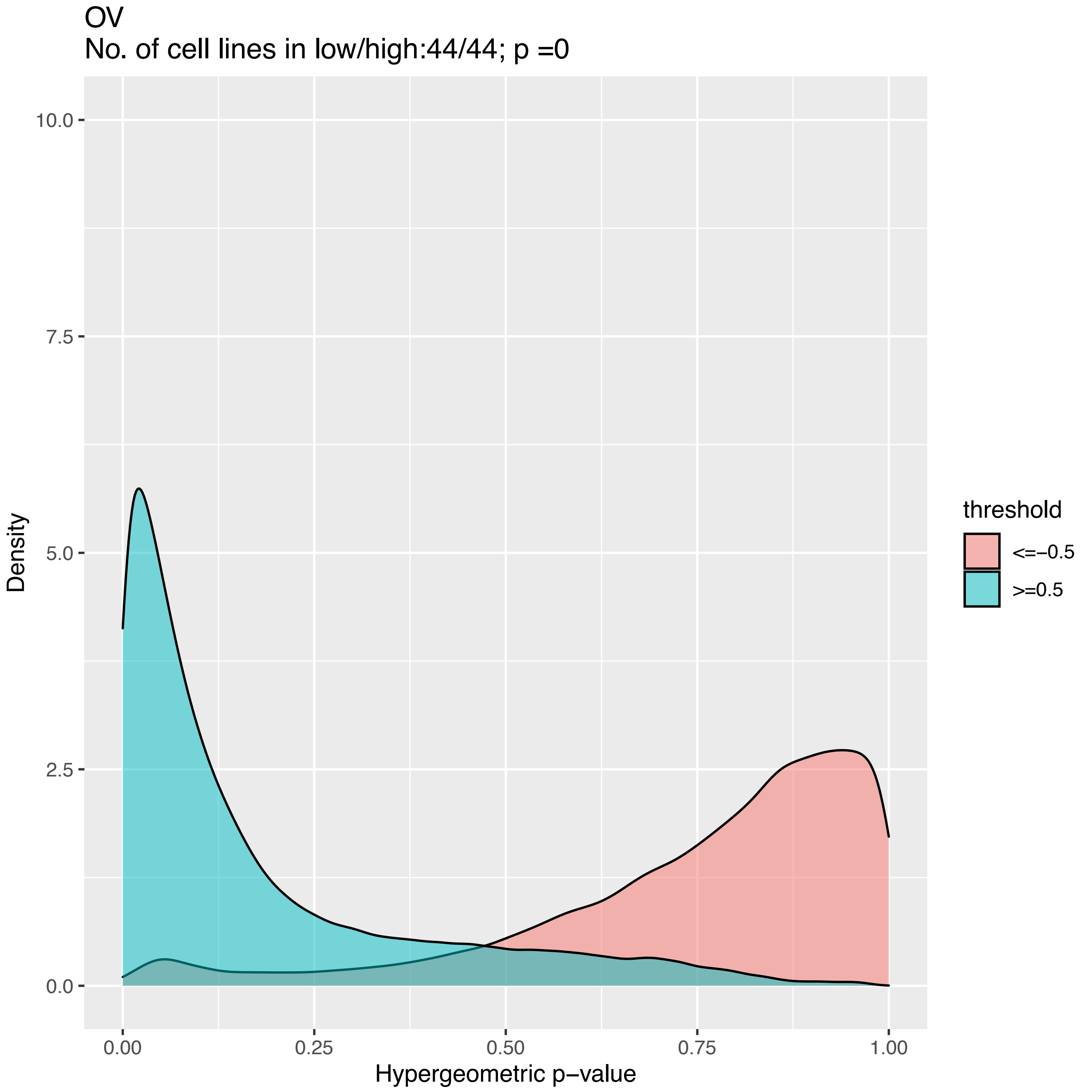

OV
No. of cell lines in low/high:44/44; p =0
Density
10.0
7.5
5.0
2.5
0.0
0.00
0.25
0.50
0.75
1.00
Hypergeometric p−value
threshold
<=−0.5
>=0.5

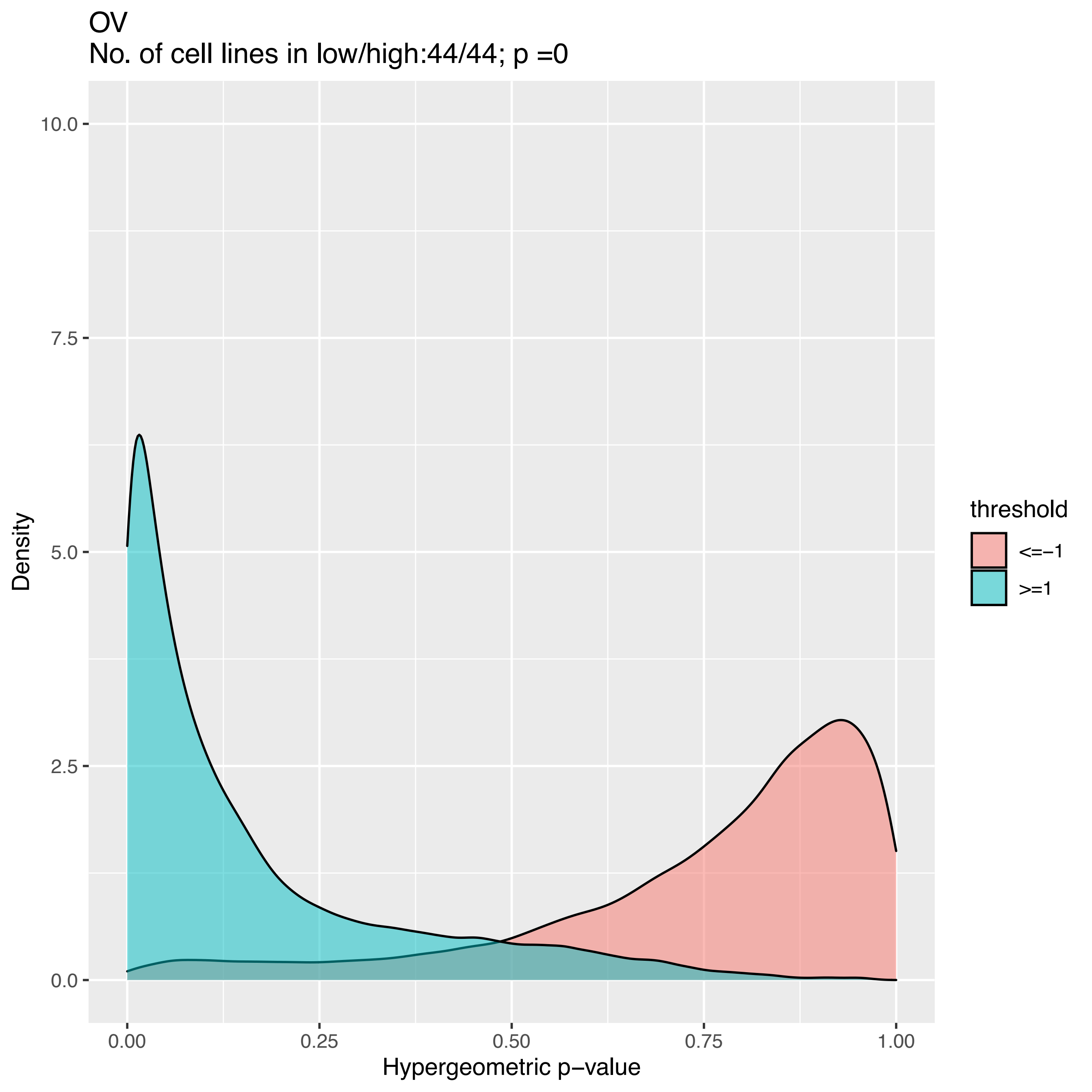

OV
No. of cell lines in low/high:44/44; p =0
10.0
7.5
5.0
2.5
0.0
Density
0.00
0.25
0.50
0.75
1.00
Hypergeometric p−value
threshold
<=−1
>=1

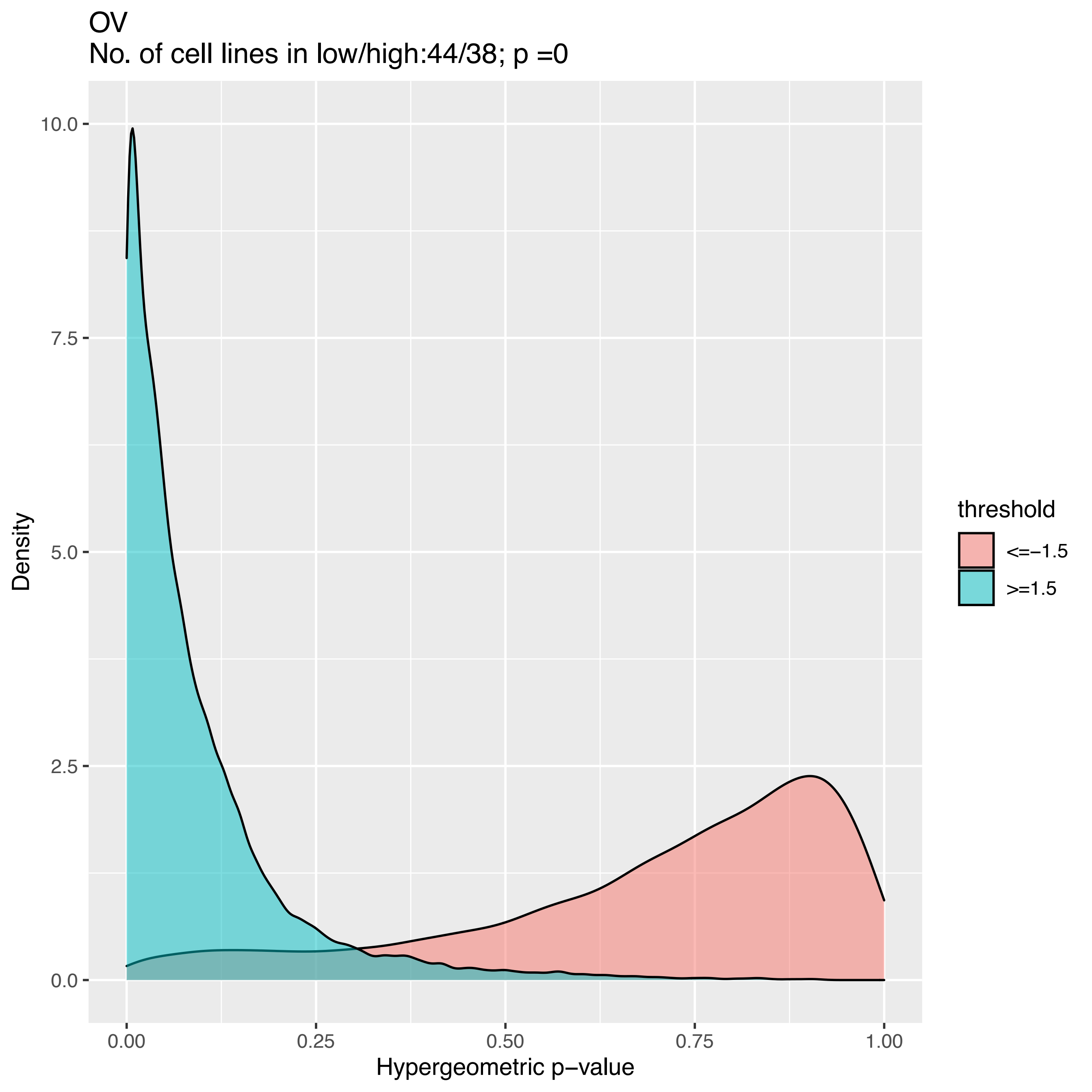
OV
No. of cell lines in low/high:44/38; p =0
10.0
7.5
5.0
2.5
0.0
Density
0.00
0.25
0.50
0.75
1.00
Hypergeometric p−value
threshold
<=−1.5
>=1.5

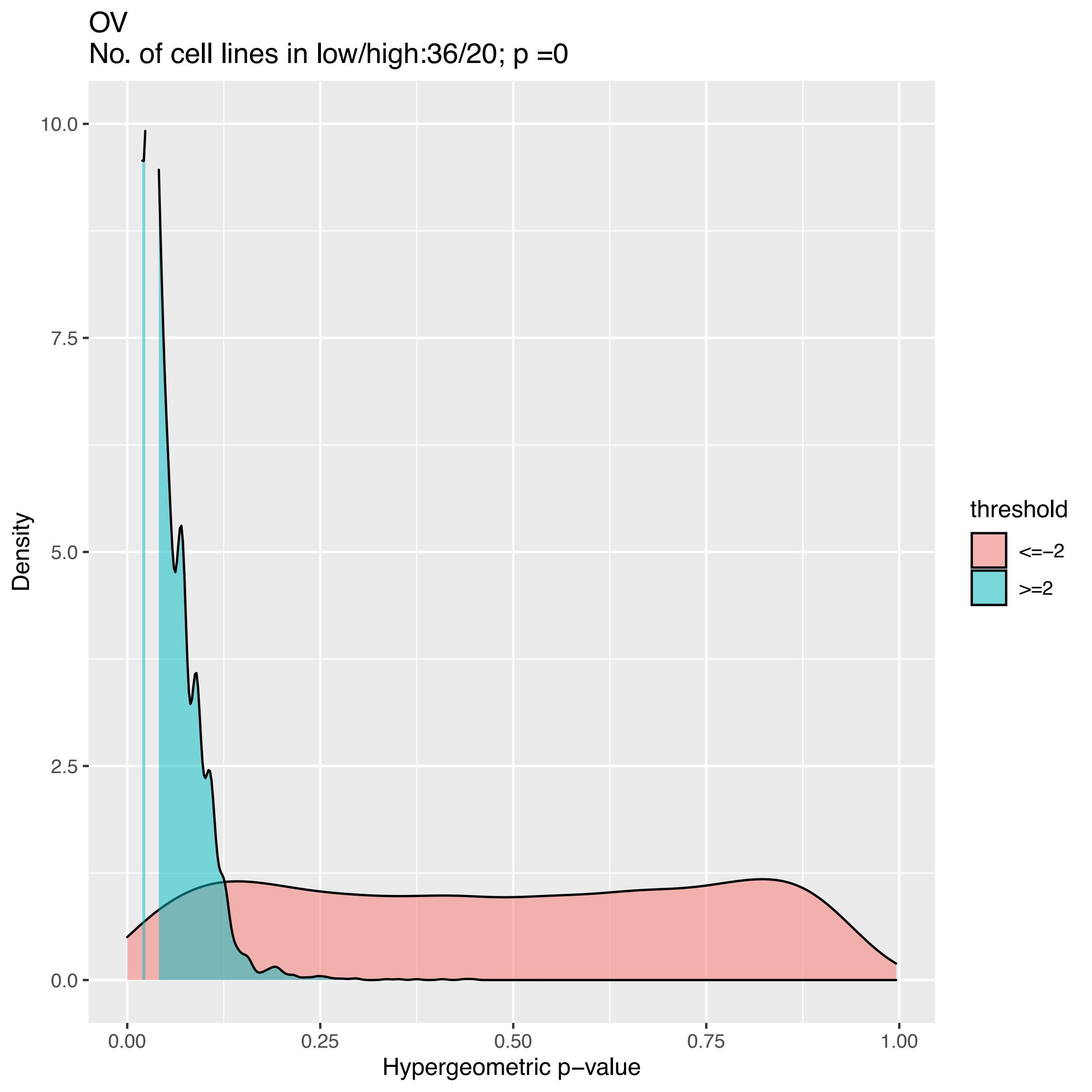
OV
No. of cell lines in low/high:36/20; p =0
10.0
7.5
5.0
2.5
0.0
Density
0.00
0.25
0.50
0.75
1.00
Hypergeometric p−value
threshold
<=−2
>=2

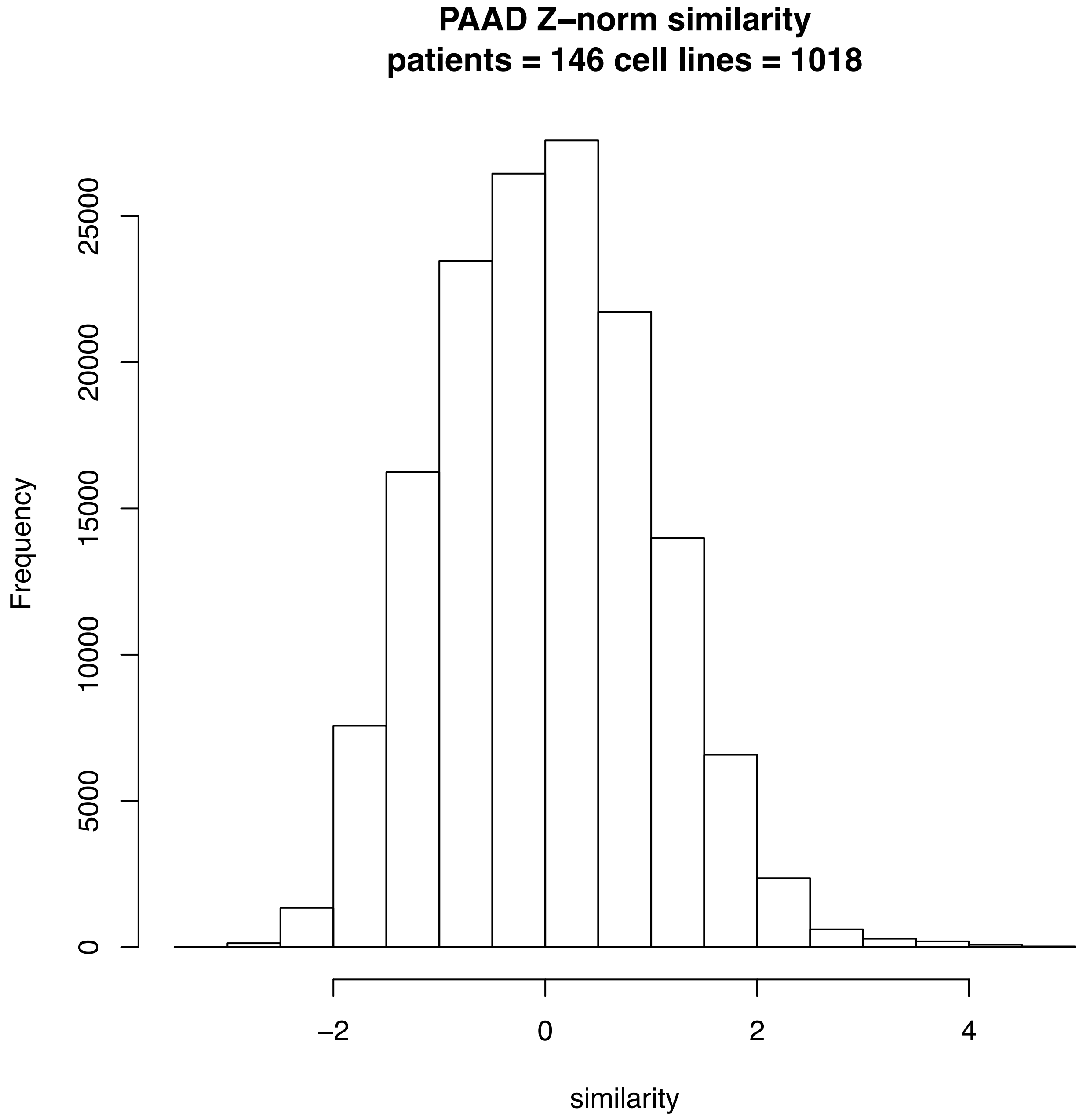
PAAD Z–norm similarity
patients = 146 cell lines = 1018
Frequency
0
5000
10000
15000
20000
25000
−2
0
2
4
similarity

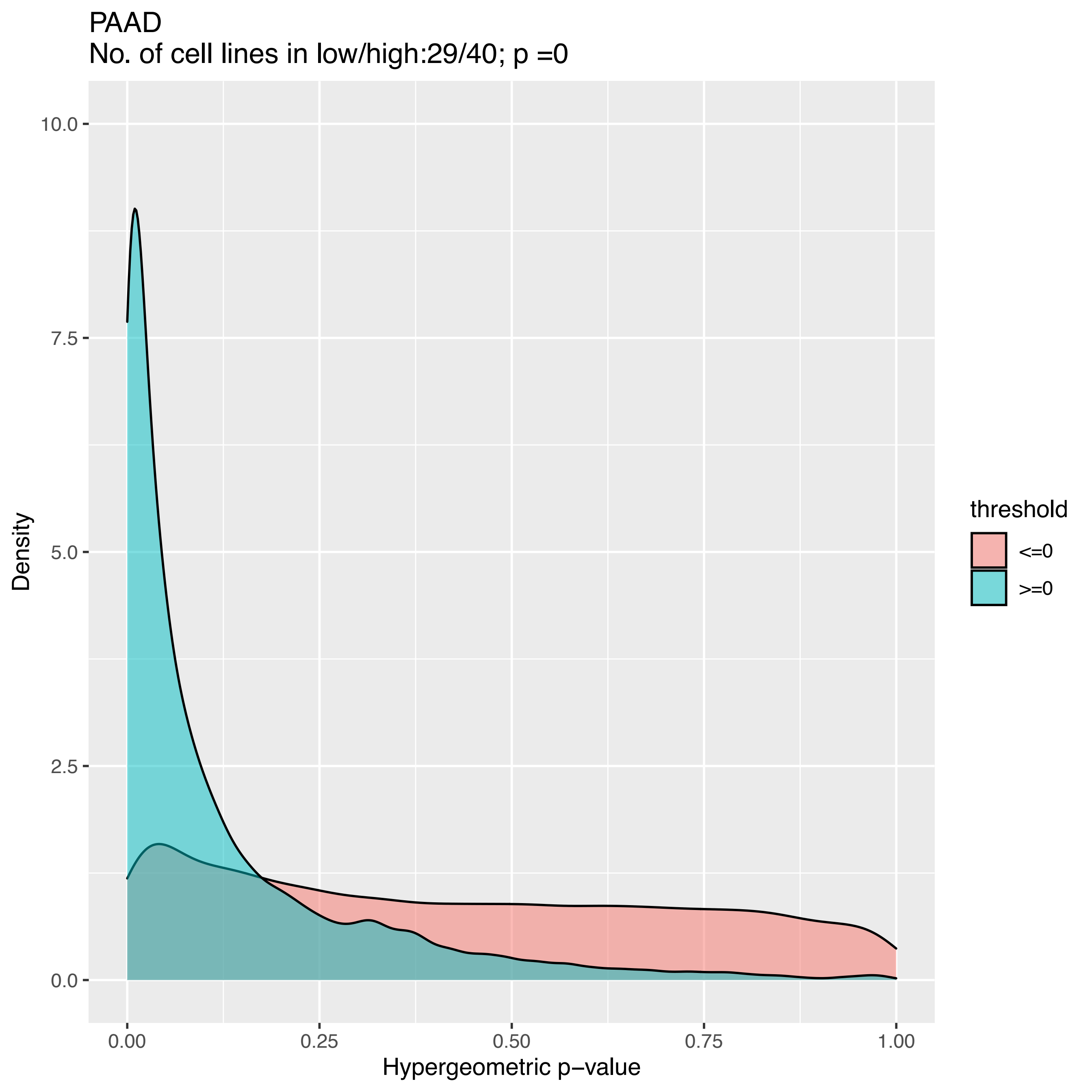

PAAD
No. of cell lines in low/high:29/40; p =0
Density
10.0
7.5
5.0
2.5
0.0
0.00
0.25
0.50
0.75
1.00
Hypergeometric p−value
threshold
<=0
>=0

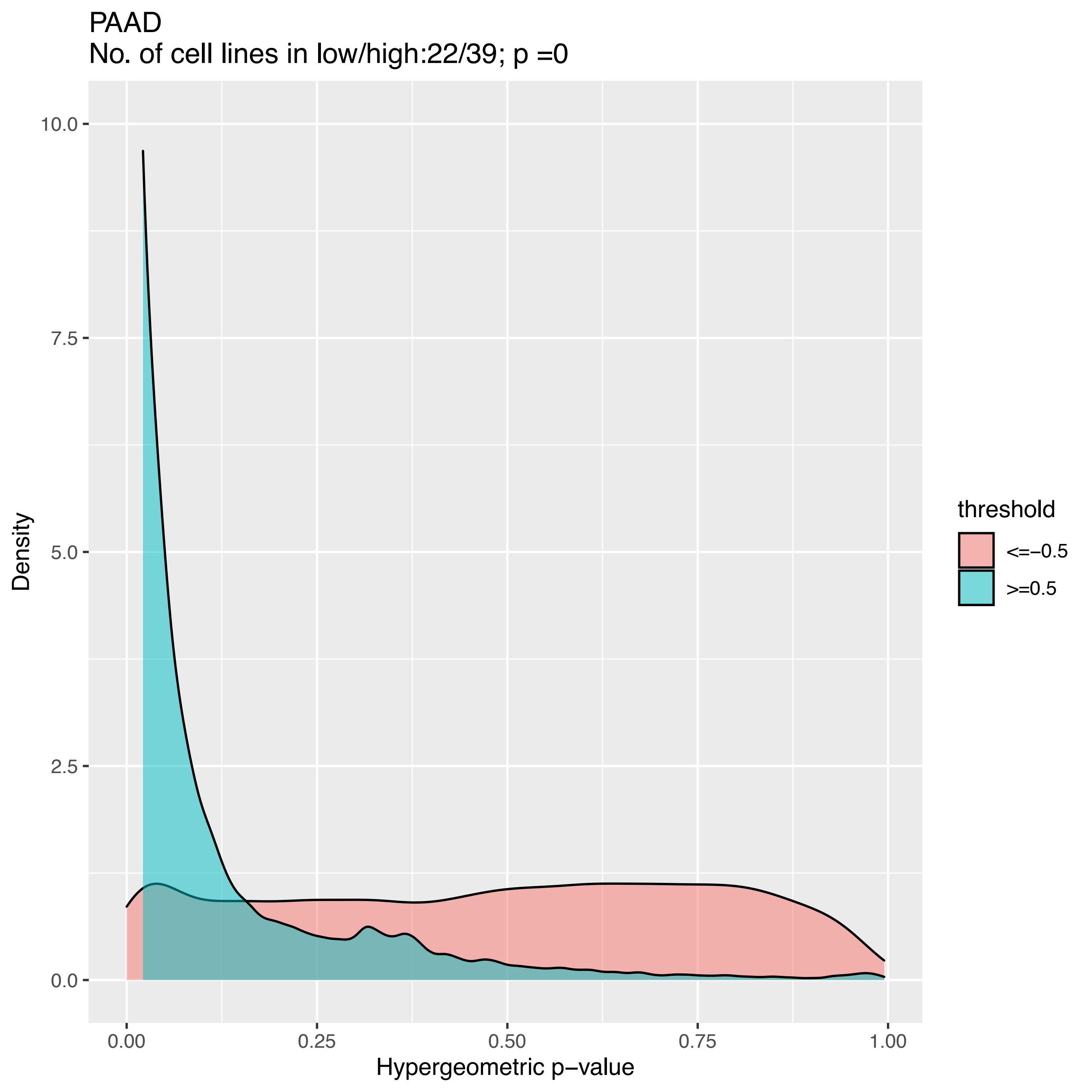

PAAD
No. of cell lines in low/high:22/39; p =0
Density
Hypergeometric p−value
10.0
7.5
5.0
2.5
0.0
0.00
0.25
0.50
0.75
1.00
threshold
<=−0.5
>=0.5

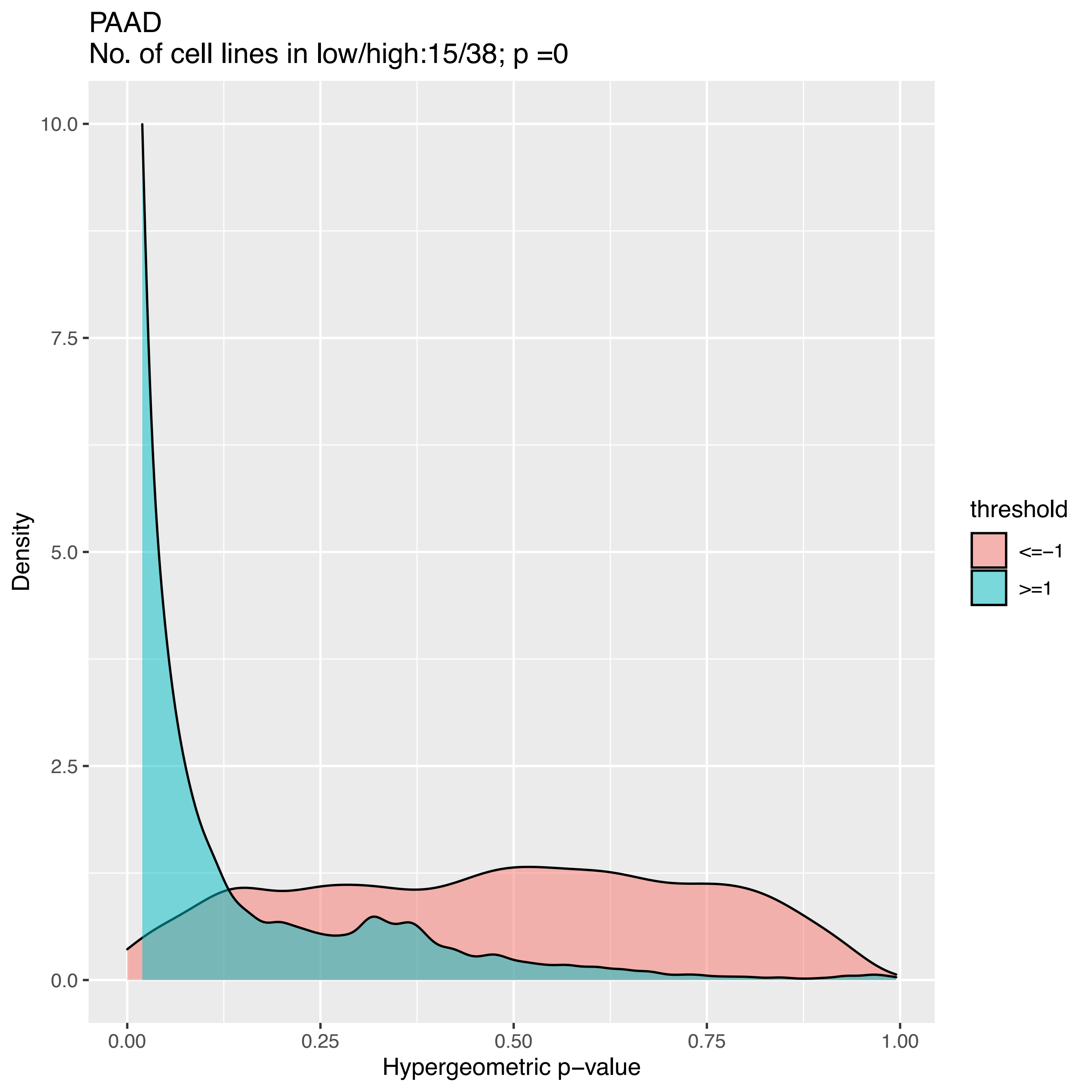
PAAD
No. of cell lines in low/high:15/38; p =0
Density
10.0
7.5
5.0
2.5
0.0
0.00
0.25
0.50
0.75
1.00
Hypergeometric p−value
threshold
<=−1
>=1

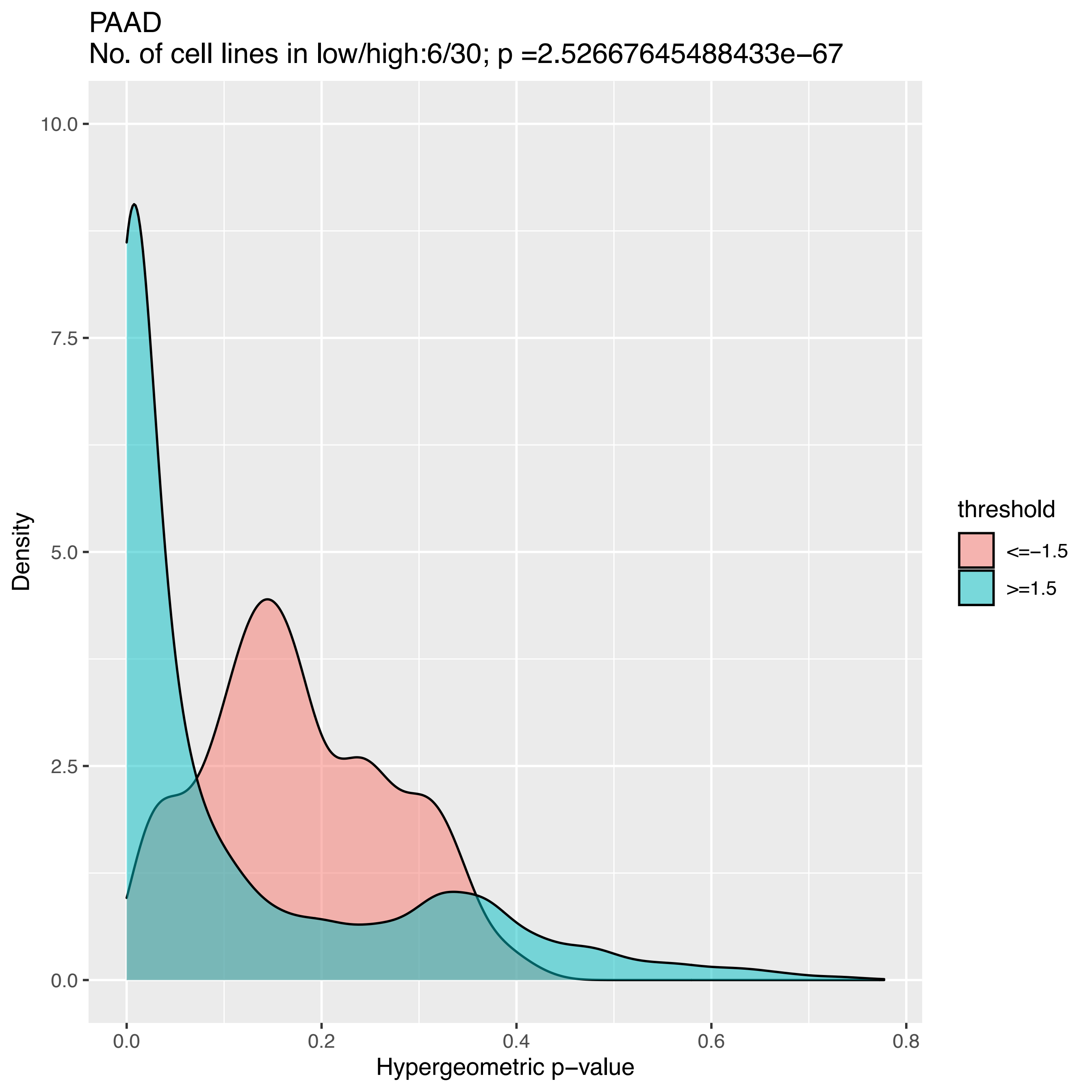
PAAD
No. of cell lines in low/high:6/30; p =2.52667645488433e−67
Density
10.0
7.5
5.0
2.5
0.0
0.0
0.2
0.4
0.6
0.8
Hypergeometric p−value
threshold
<=−1.5
>=1.5

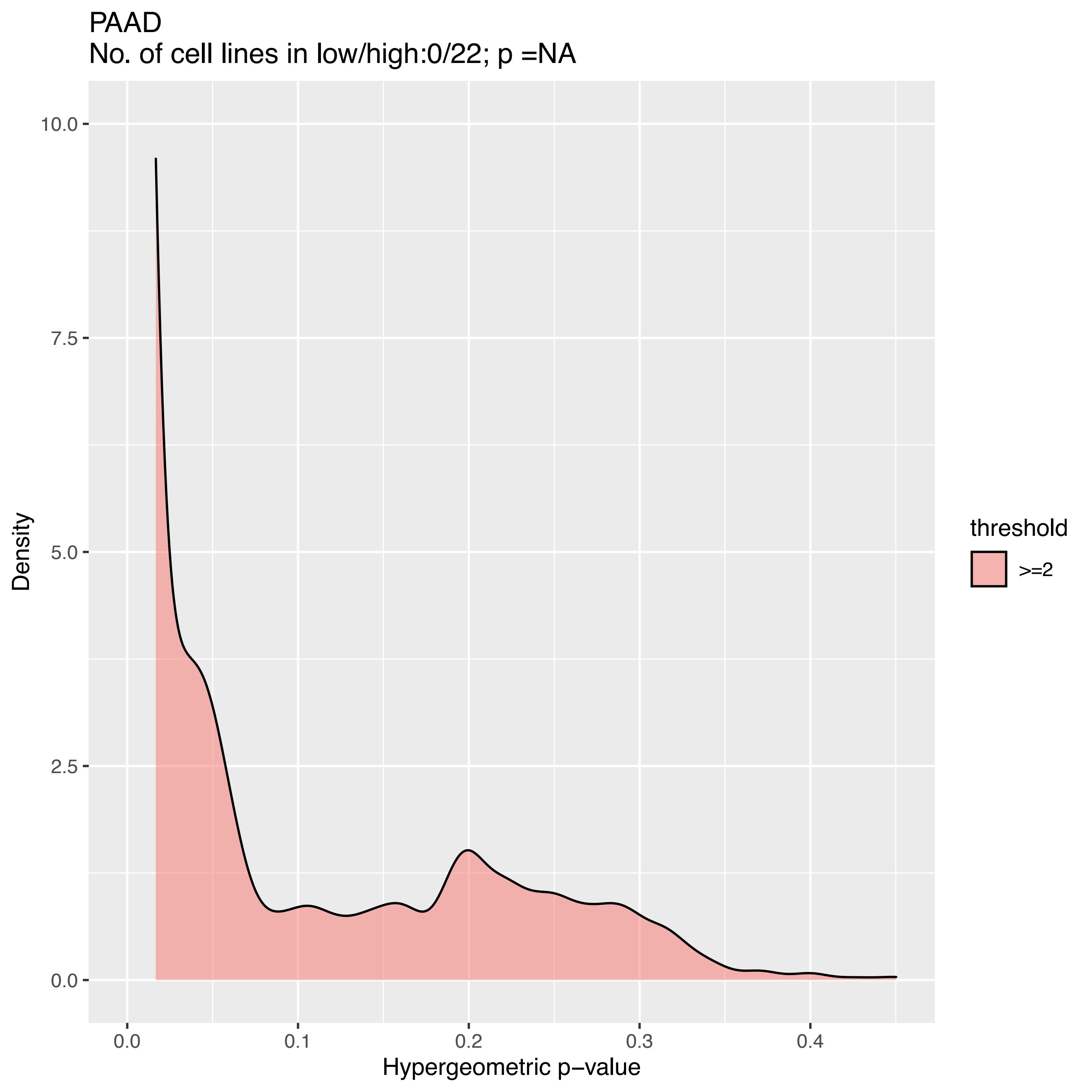

PAAD
No. of cell lines in low/high:0/22; p =NA
Density
10.0
7.5
5.0
2.5
0.0
0.0
0.1
0.2
0.3
0.4
Hypergeometric p−value
threshold
>=2

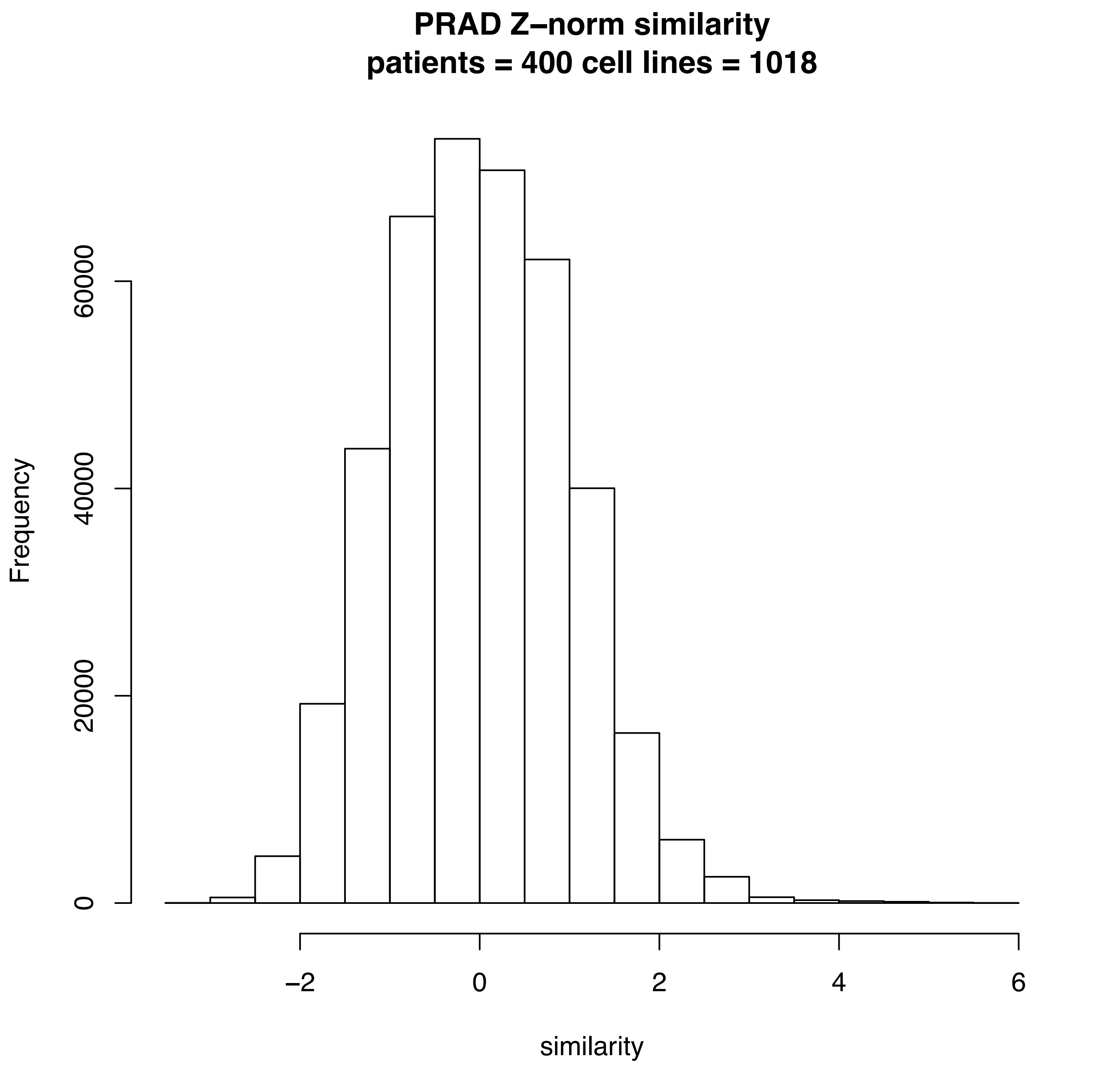
PRAD Z–norm similarity
patients = 400 cell lines = 1018
Frequency
0
20000
40000
60000
–2
0
2
4
6
similarity

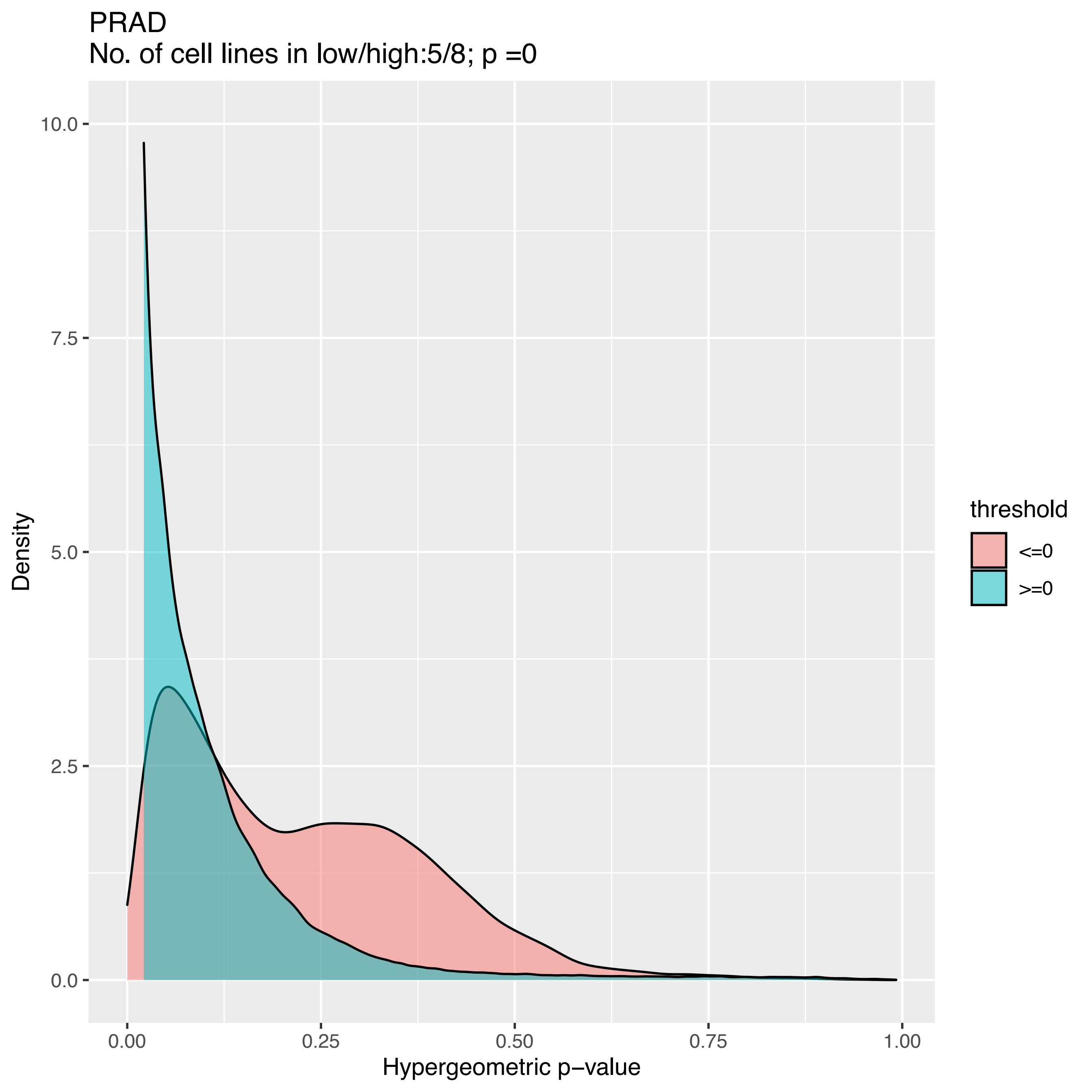
PRAD
No. of cell lines in low/high:5/8; p =0
Density
10.0
7.5
5.0
2.5
0.0
0.00
0.25
0.50
0.75
1.00
Hypergeometric p−value
threshold
<=0
>=0

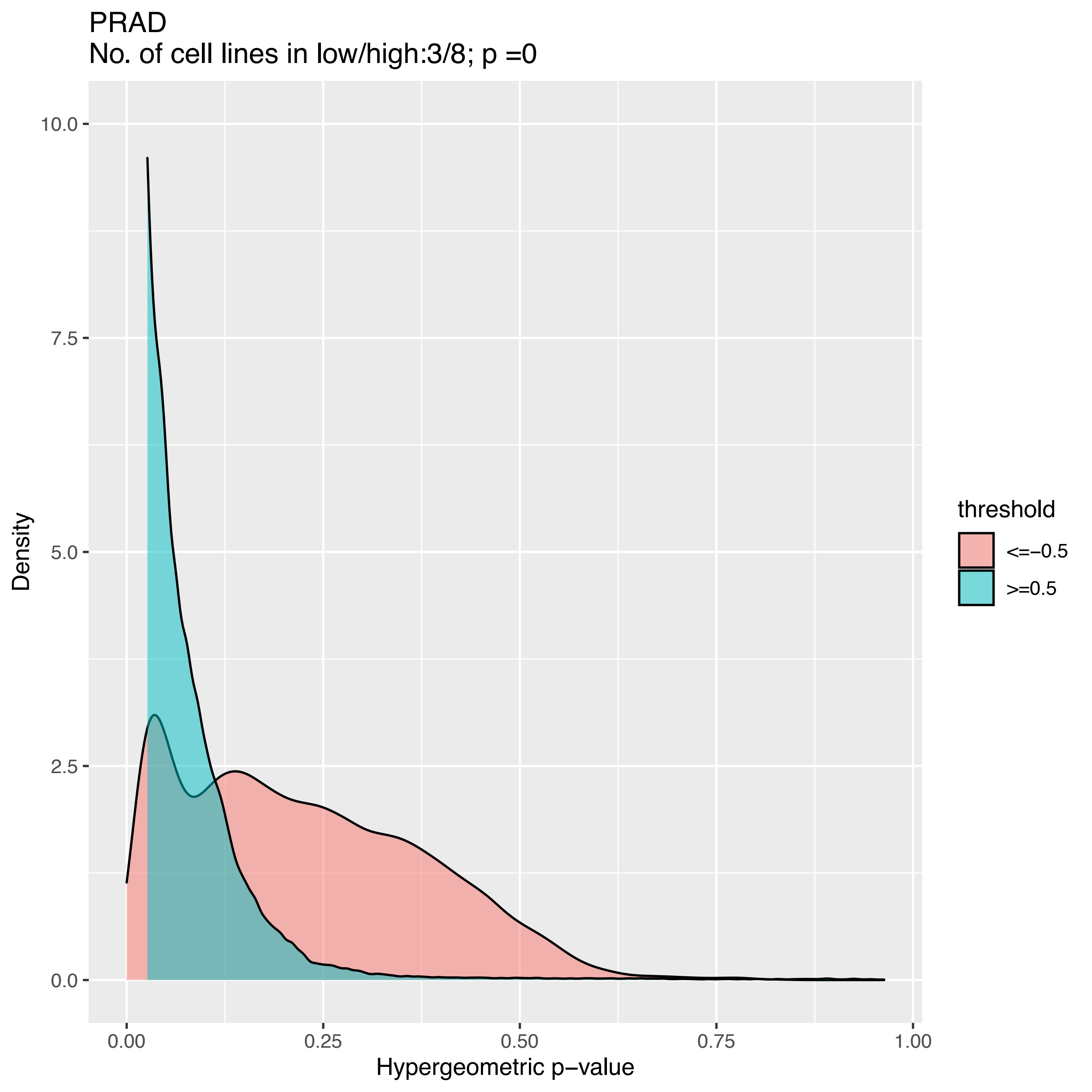
PRAD
No. of cell lines in low/high:3/8; p =0
Density
10.0
7.5
5.0
2.5
0.0
0.00
0.25
0.50
0.75
1.00
Hypergeometric p−value
threshold
<=−0.5
>=0.5

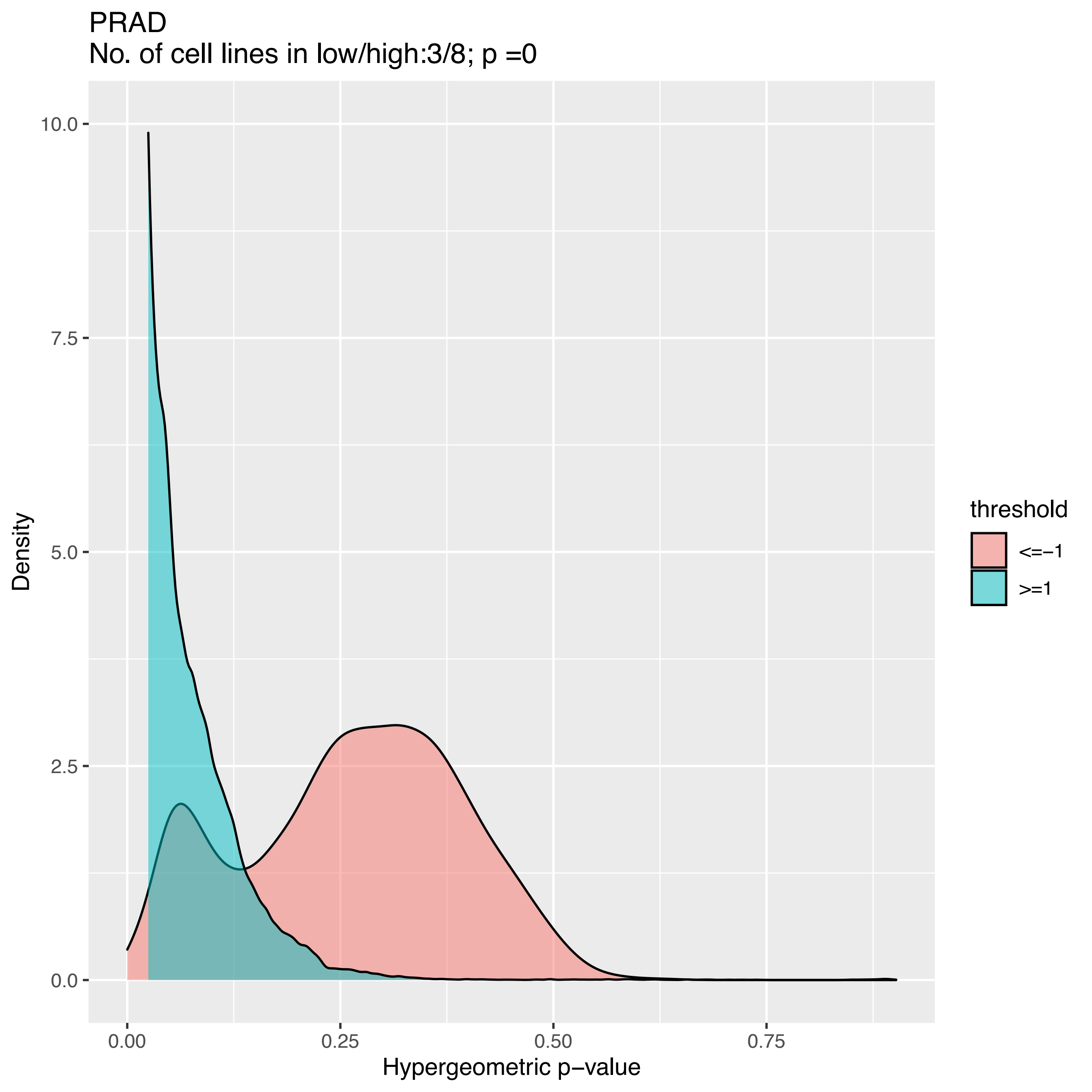

PRAD
No. of cell lines in low/high:3/8; p =0
Density
10.0
7.5
5.0
2.5
0.0
0.00
0.25
0.50
0.75
Hypergeometric p−value
threshold
<=−1
>=1

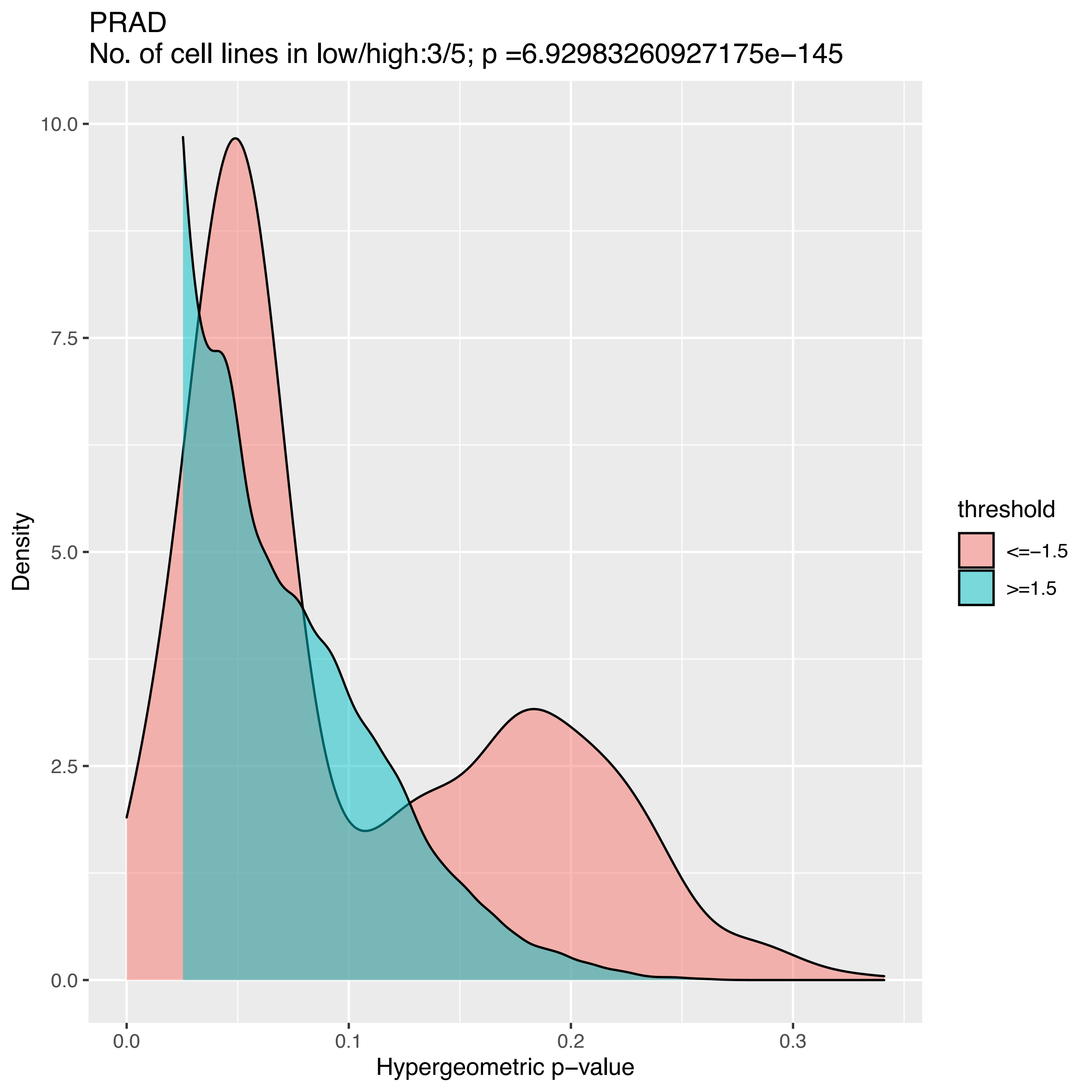

PRAD
No. of cell lines in low/high:3/5; p =6.92983260927175e−145
Density
10.0
7.5
5.0
2.5
0.0
0.0
0.1
0.2
0.3
Hypergeometric p−value
threshold
<=−1.5
>=1.5

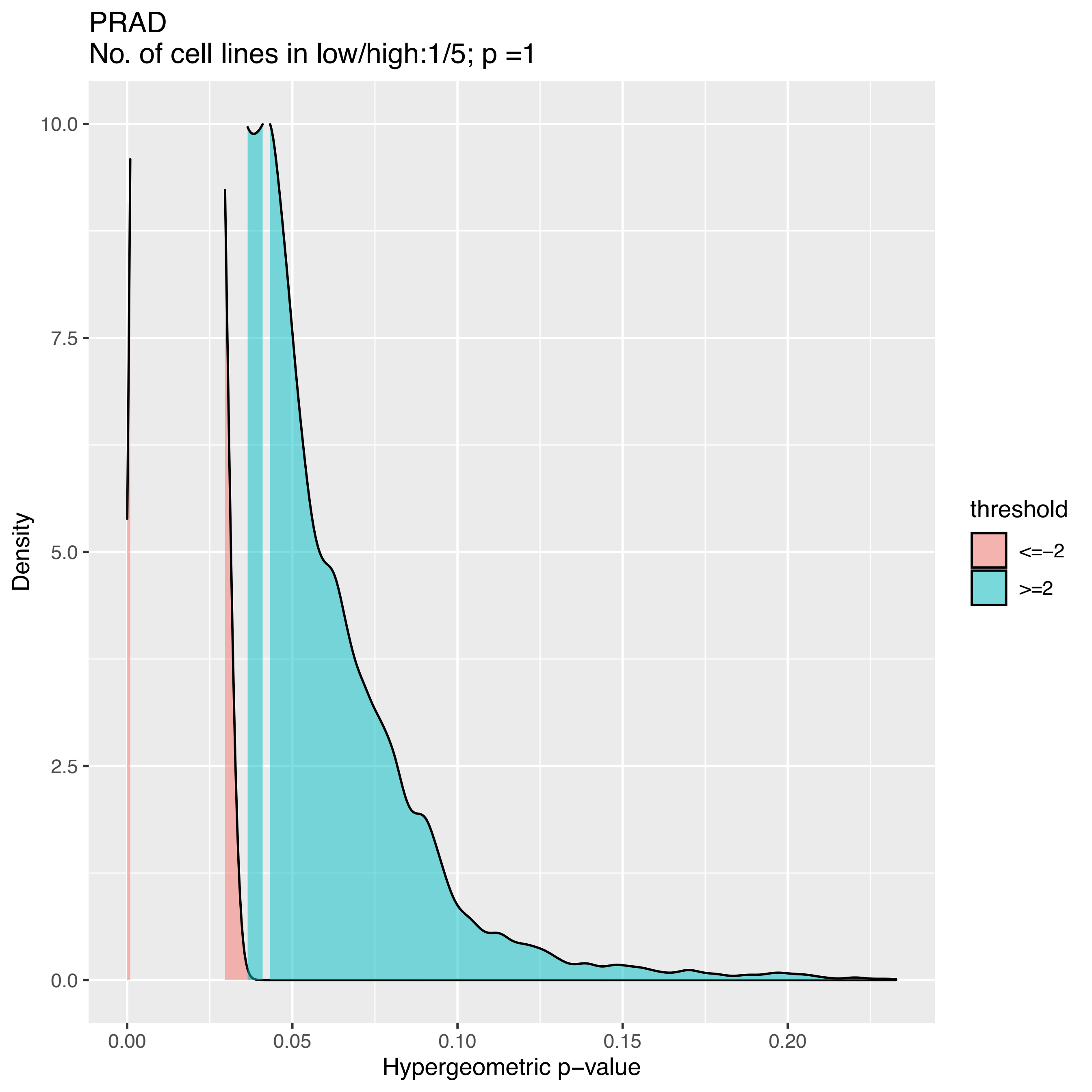
PRAD
No. of cell lines in low/high:1/5; p =1
Density
Hypergeometric p−value
10.0
7.5
5.0
2.5
0.0
0.00
0.05
0.10
0.15
0.20
threshold
<=−2
>=2

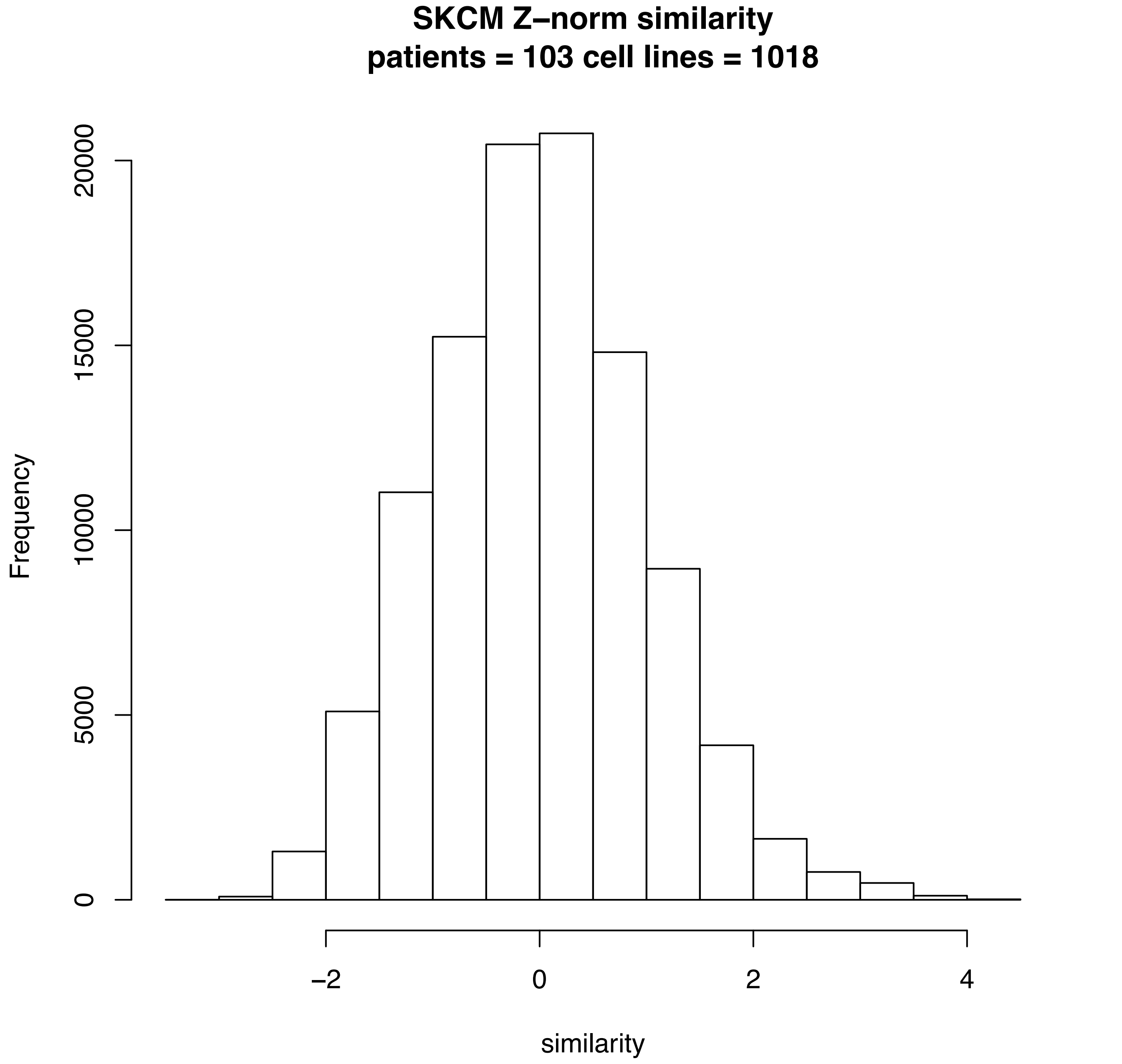
SKCM Z–norm similarity
patients = 103 cell lines = 1018
Frequency
0
5000
10000
15000
20000
–2
0
2
4
similarity

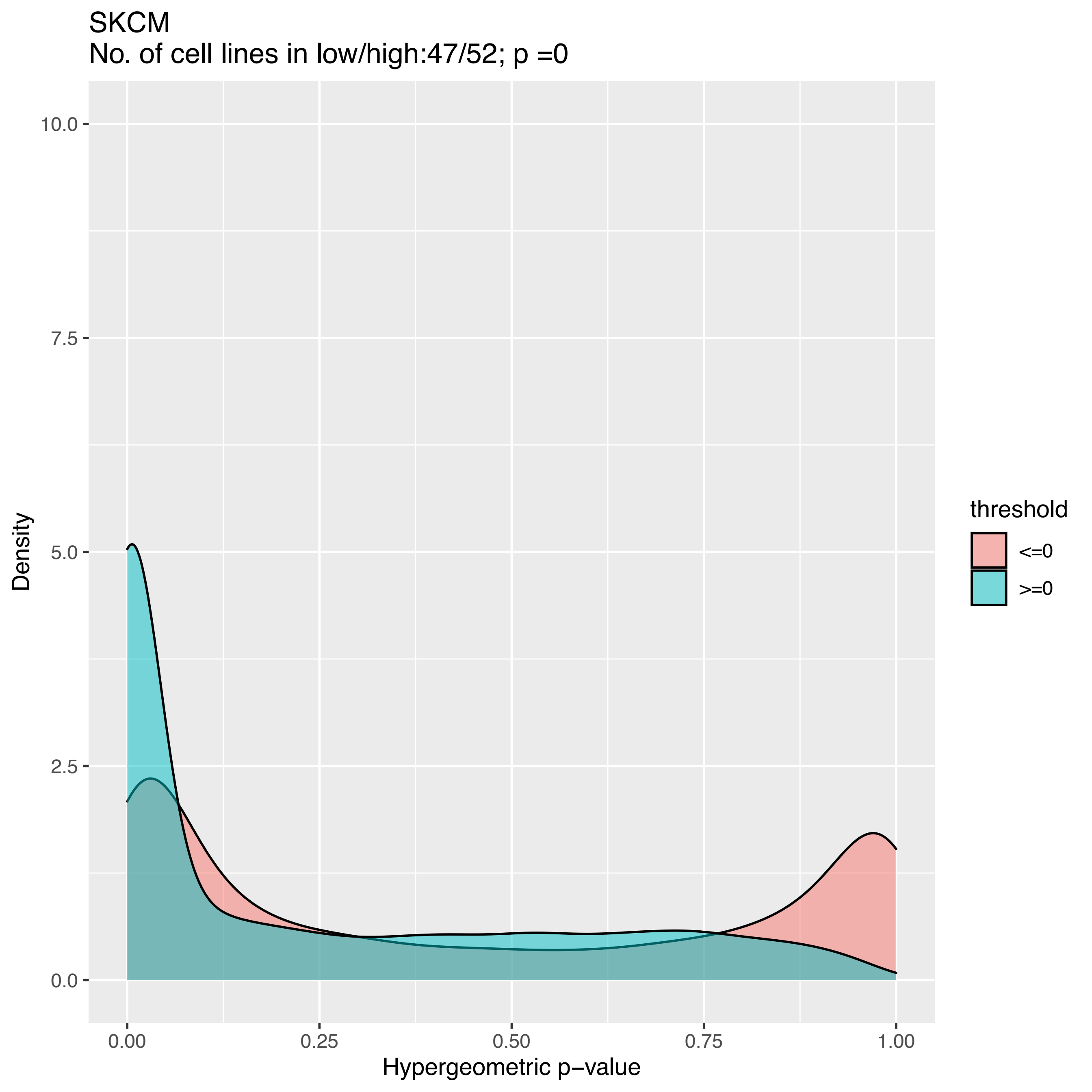

SKCM
No. of cell lines in low/high:47/52; p =0
Density
10.0
7.5
5.0
2.5
0.0
0.00
0.25
0.50
0.75
1.00
Hypergeometric p−value
threshold
<=0
>=0

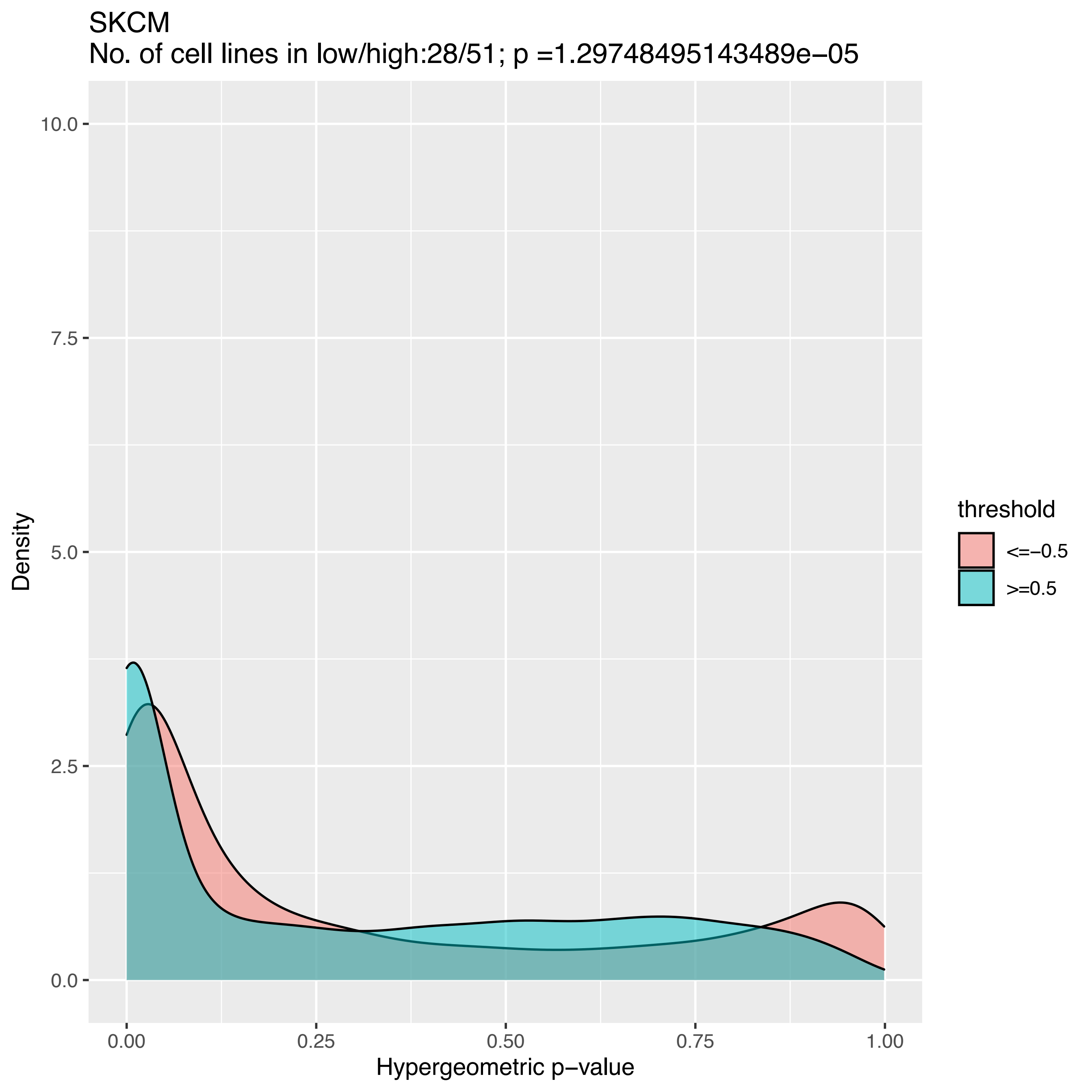
SKCM
No. of cell lines in low/high:28/51; p =1.29748495143489e−05
Density
Hypergeometric p−value
0.0
2.5
5.0
7.5
10.0
0.00
0.25
0.50
0.75
1.00
threshold
<=−0.5
>=0.5

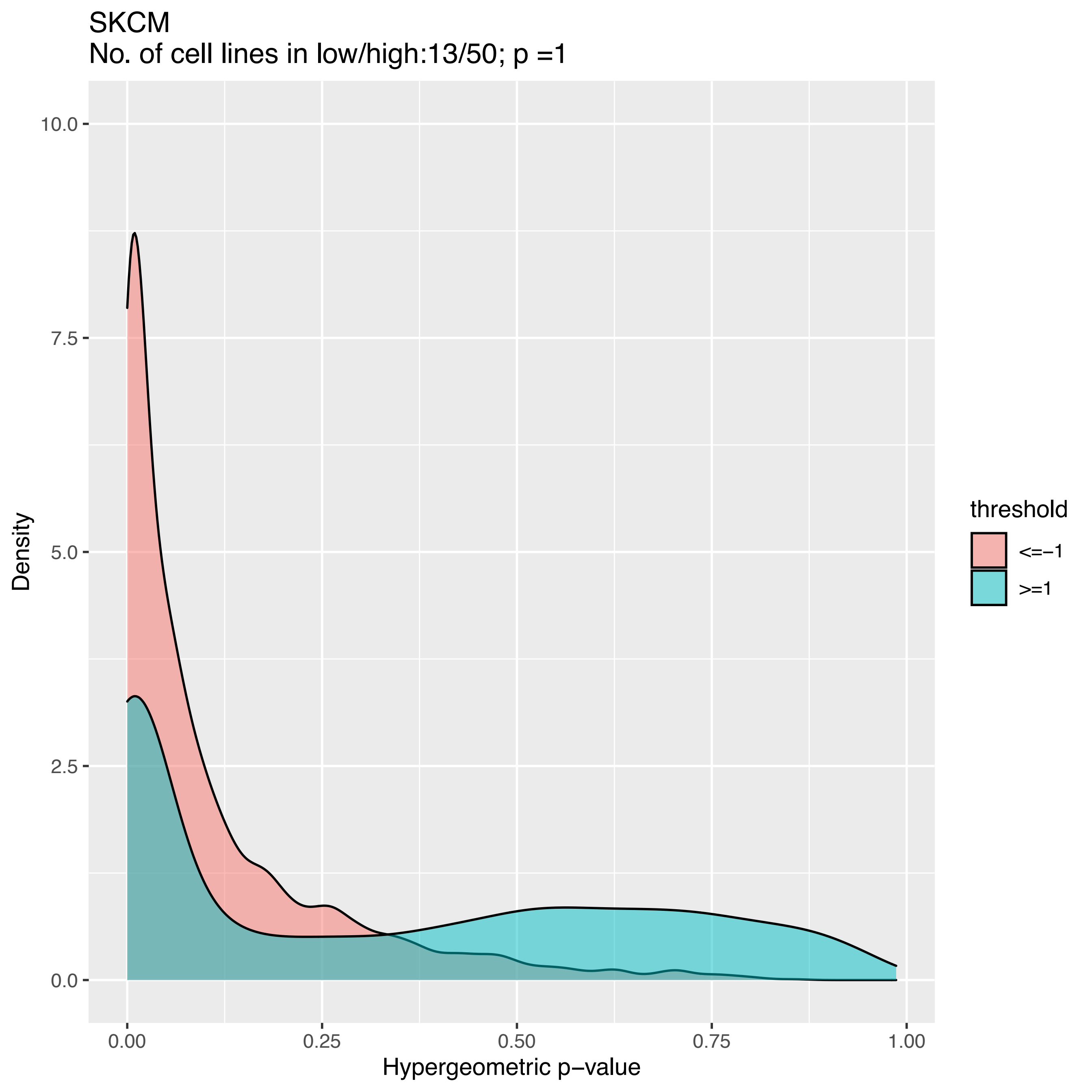

SKCM
No. of cell lines in low/high:13/50; p =1
Density
10.0
7.5
5.0
2.5
0.0
0.00
0.25
0.50
0.75
1.00
Hypergeometric p−value
threshold
<=−1
>=1

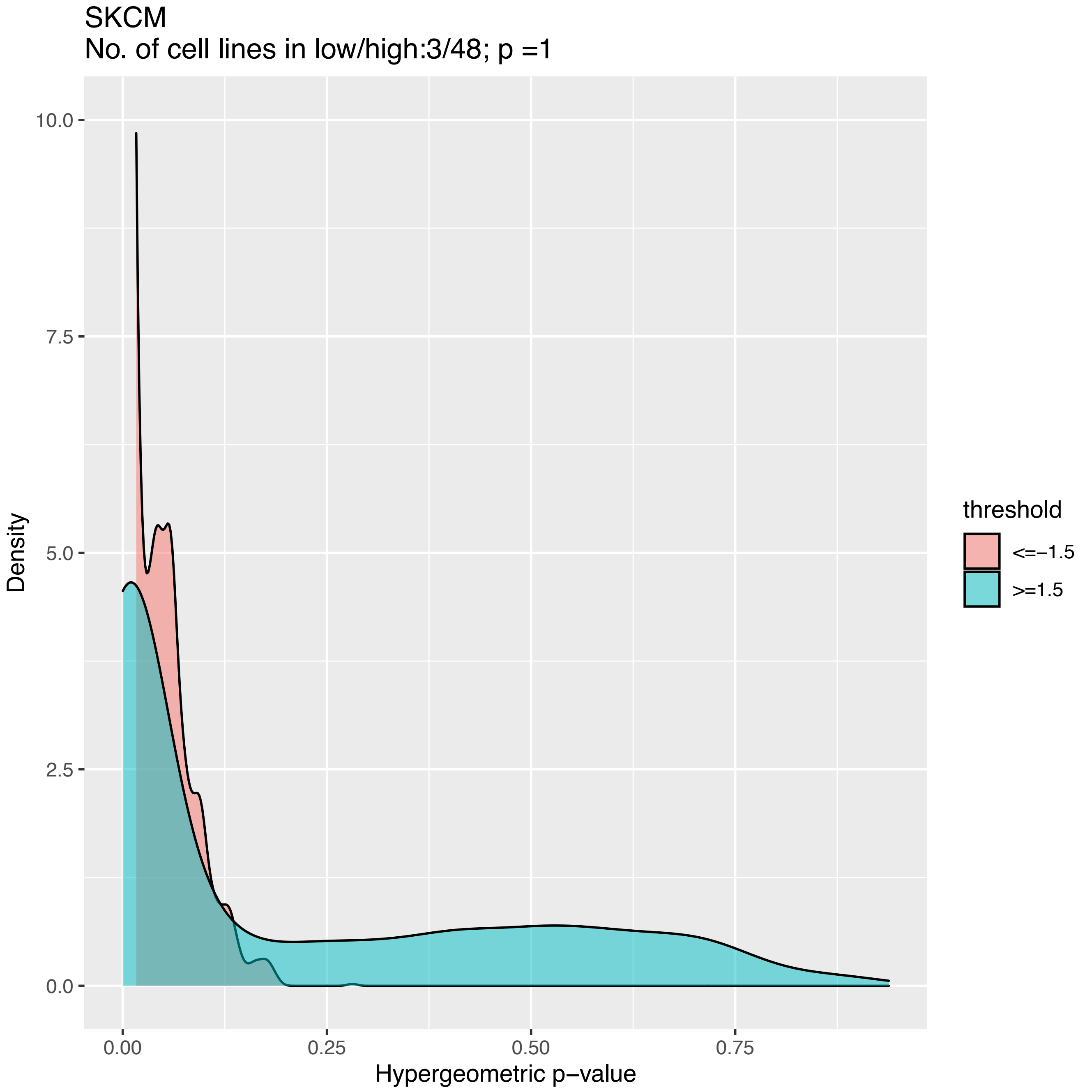

SKCM
No. of cell lines in low/high:3/48; p =1
Density
10.0
7.5
5.0
2.5
0.0
0.00
0.25
0.50
0.75
Hypergeometric p−value
threshold
<=−1.5
>=1.5

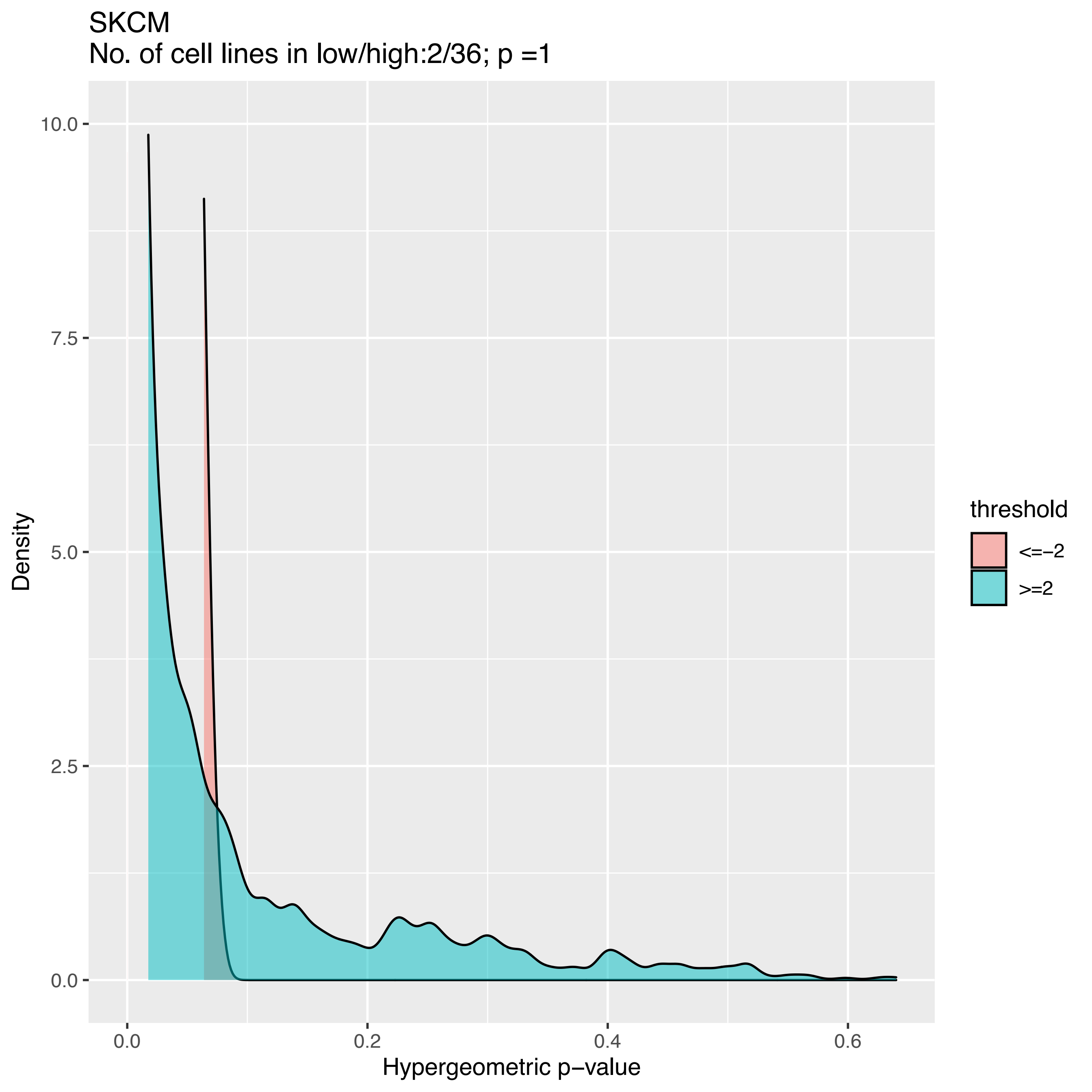
SKCM
No. of cell lines in low/high:2/36; p =1
Density
10.0
7.5
5.0
2.5
0.0
0.0
0.2
0.4
0.6
Hypergeometric p−value
threshold
<=−2
>=2

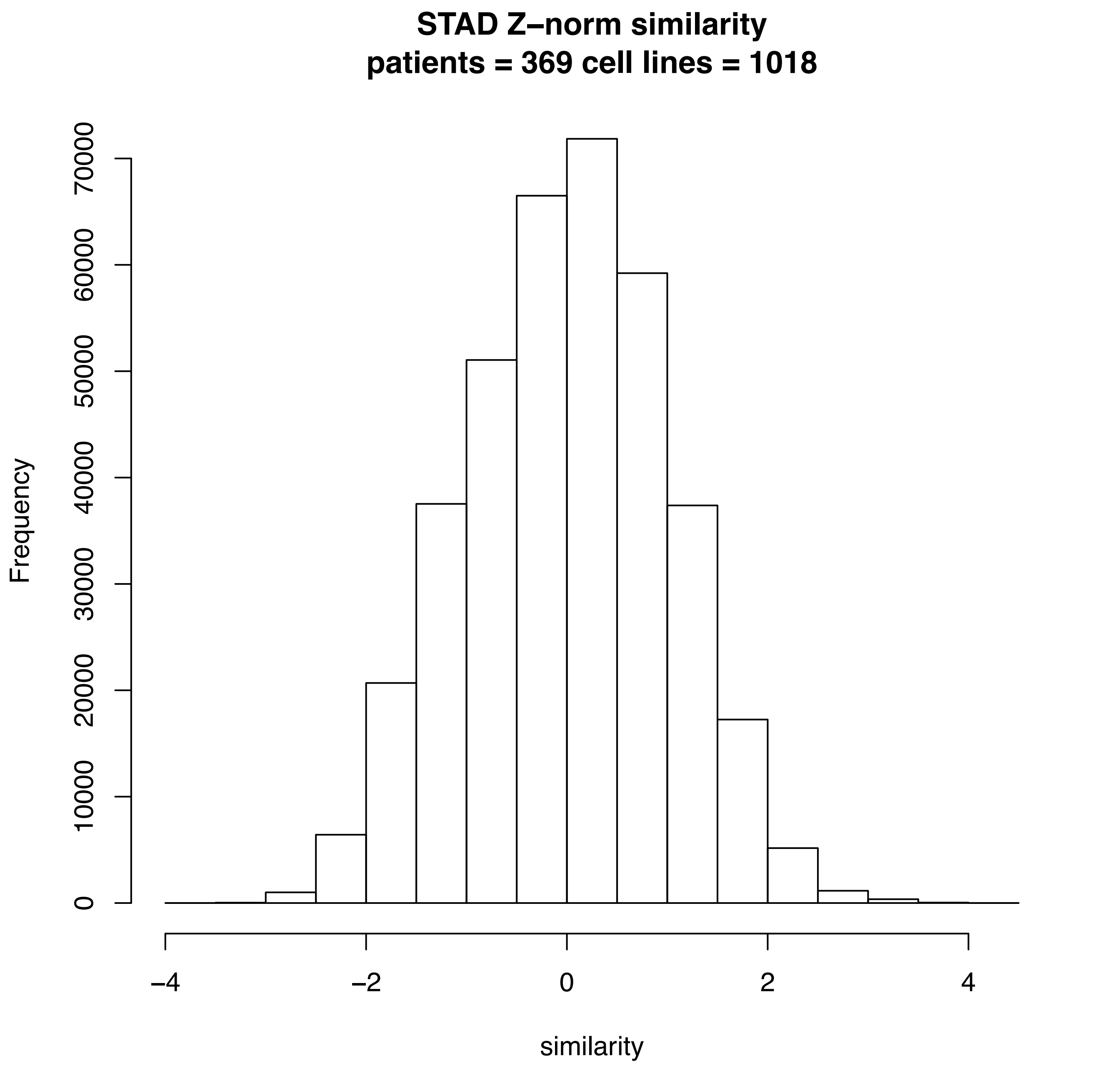

STAD Z–norm similarity
patients = 369 cell lines = 1018
Frequency
0
10000
20000
30000
40000
50000
60000
70000
–4
–2
0
2
4
similarity

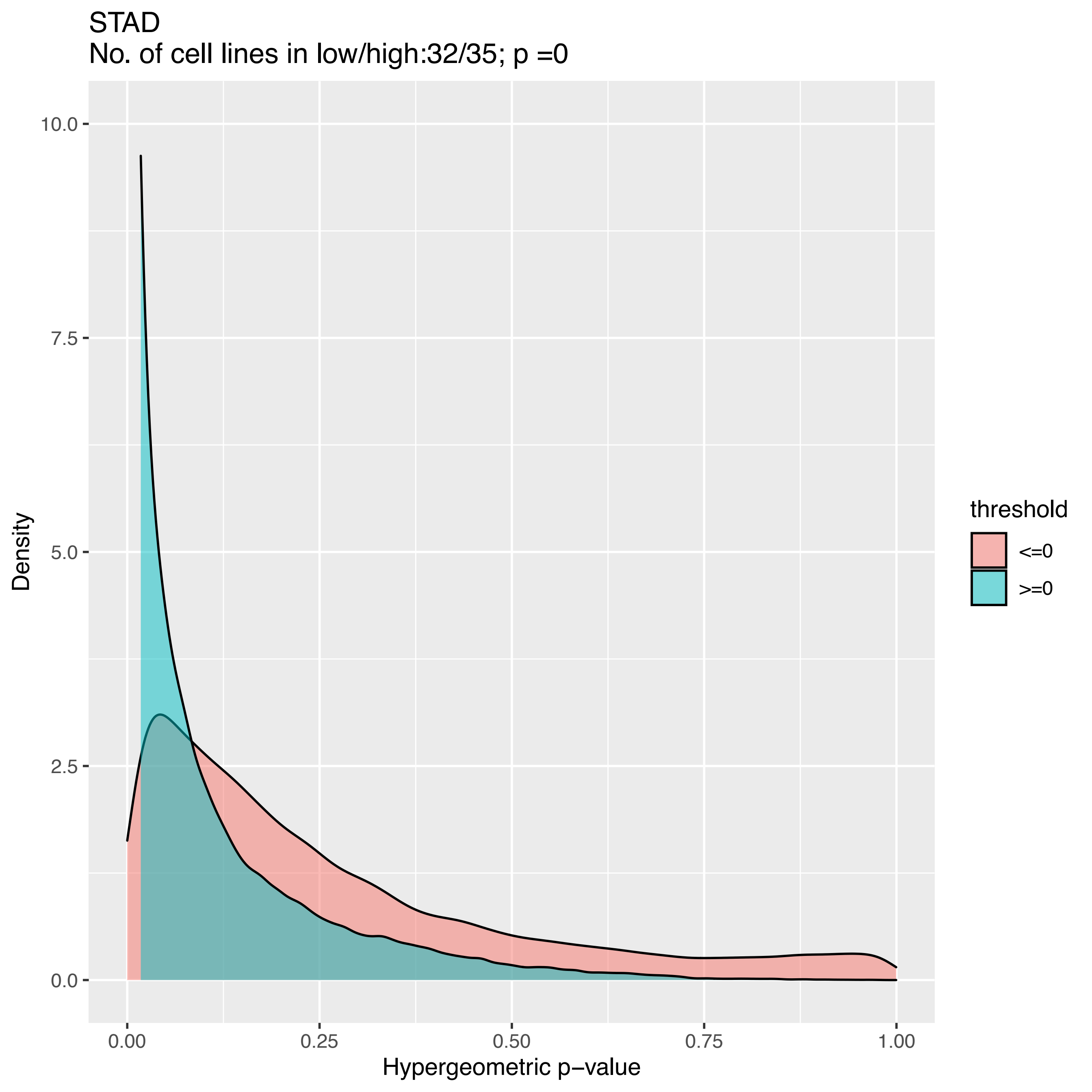

STAD
No. of cell lines in low/high:32/35; p =0
Density
10.0
7.5
5.0
2.5
0.0
0.00
0.25
0.50
0.75
1.00
Hypergeometric p−value
threshold
<=0
>=0

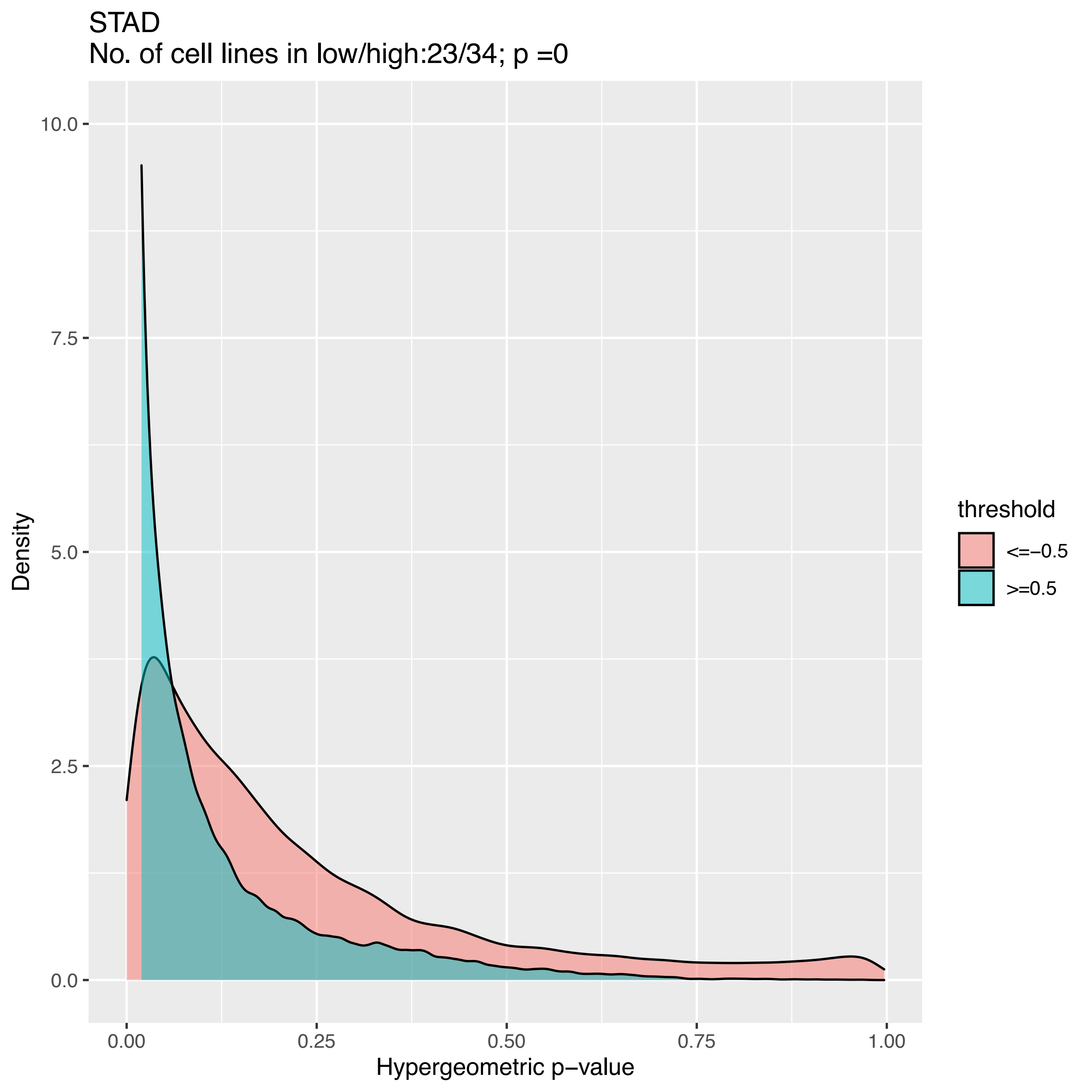
STAD
No. of cell lines in low/high:23/34; p =0
10.0
7.5
5.0
2.5
0.0
Density
0.00
0.25
0.50
0.75
1.00
Hypergeometric p−value
threshold
<=−0.5
>=0.5

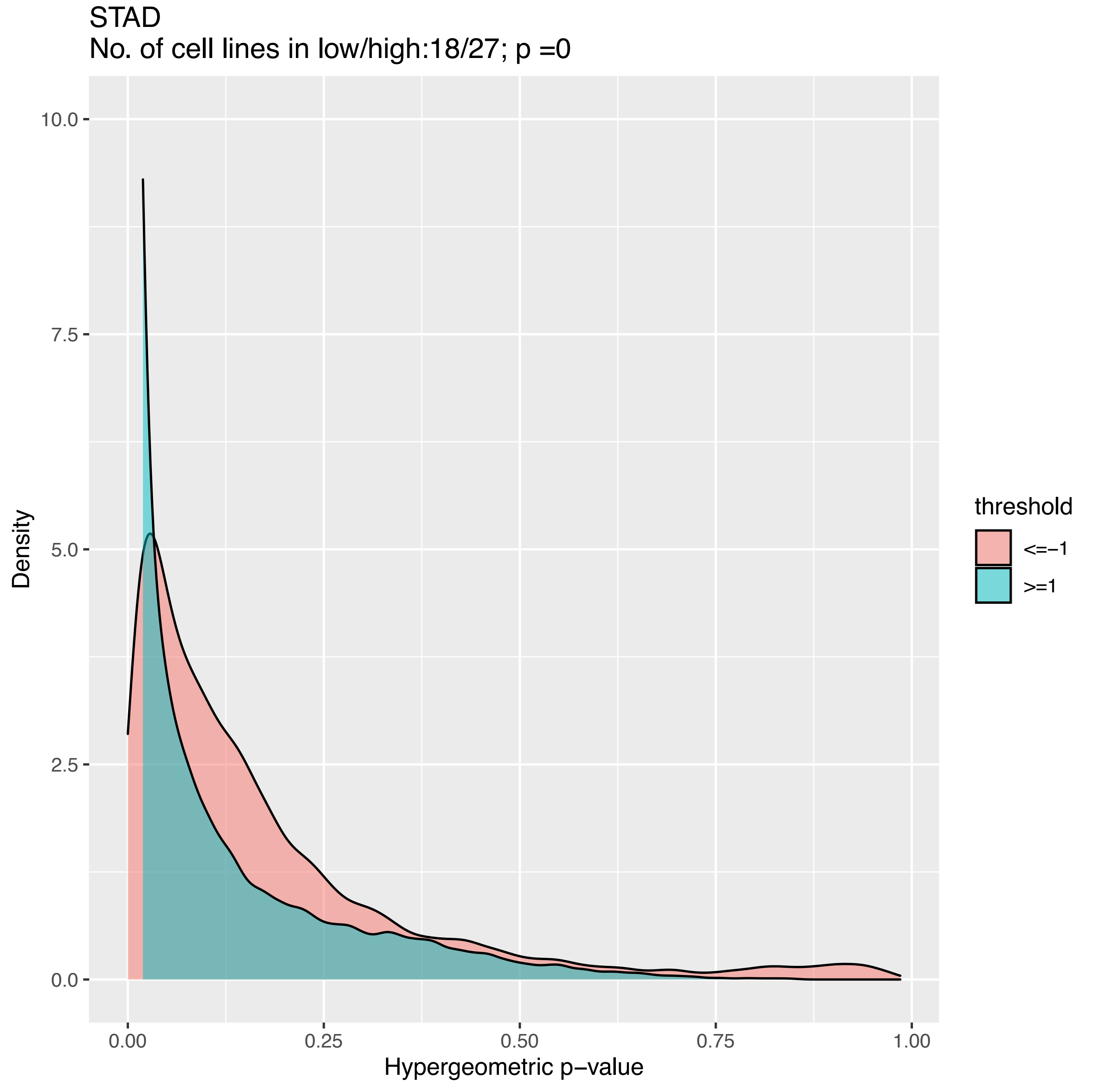
STAD
No. of cell lines in low/high:18/27; p =0
Density
10.0
7.5
5.0
2.5
0.0
0.00
0.25
0.50
0.75
1.00
Hypergeometric p−value
threshold
<=−1
>=1

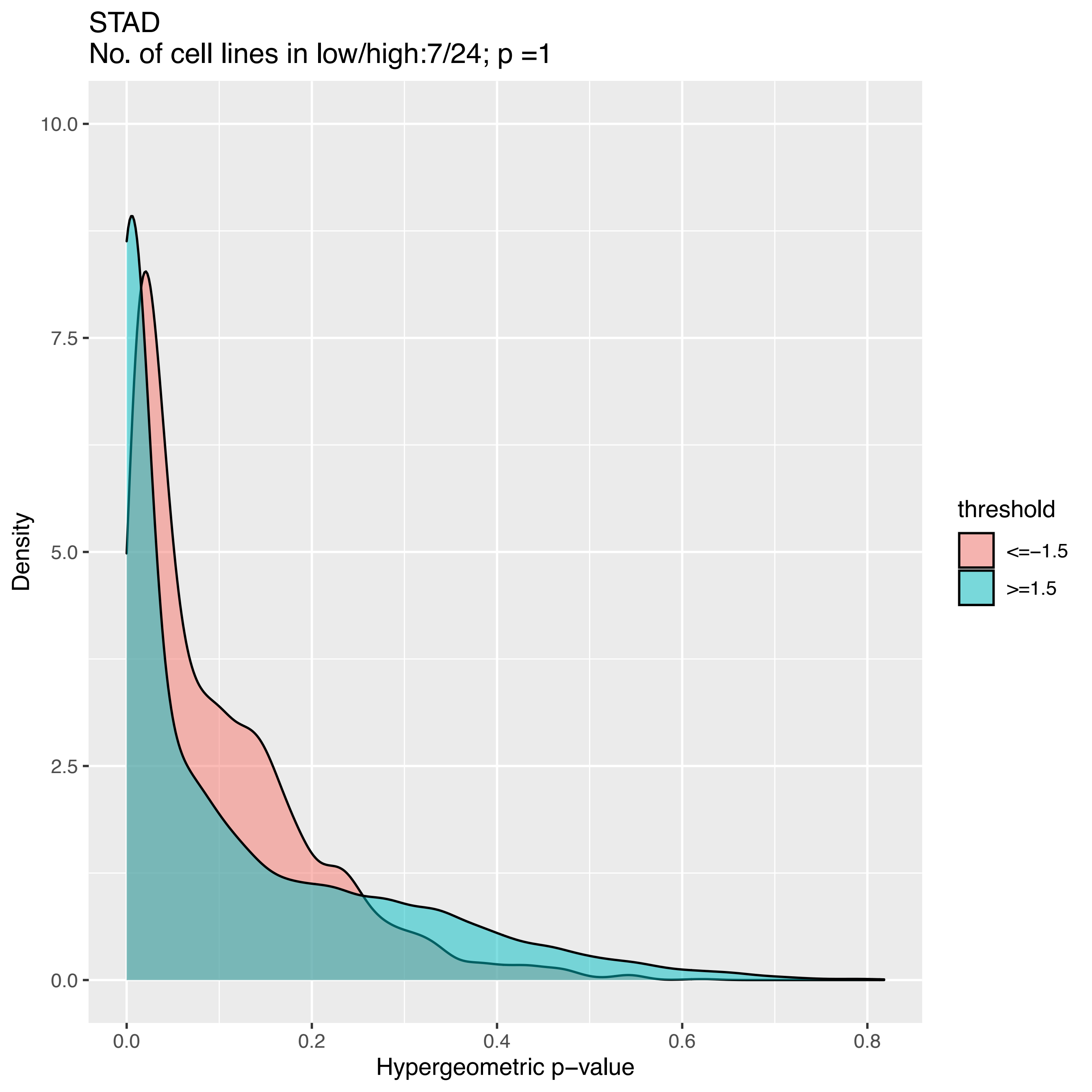
STAD
No. of cell lines in low/high:7/24; p =1
Density
10.0
7.5
5.0
2.5
0.0
0.0
0.2
0.4
0.6
0.8
Hypergeometric p−value
threshold
<=−1.5
>=1.5

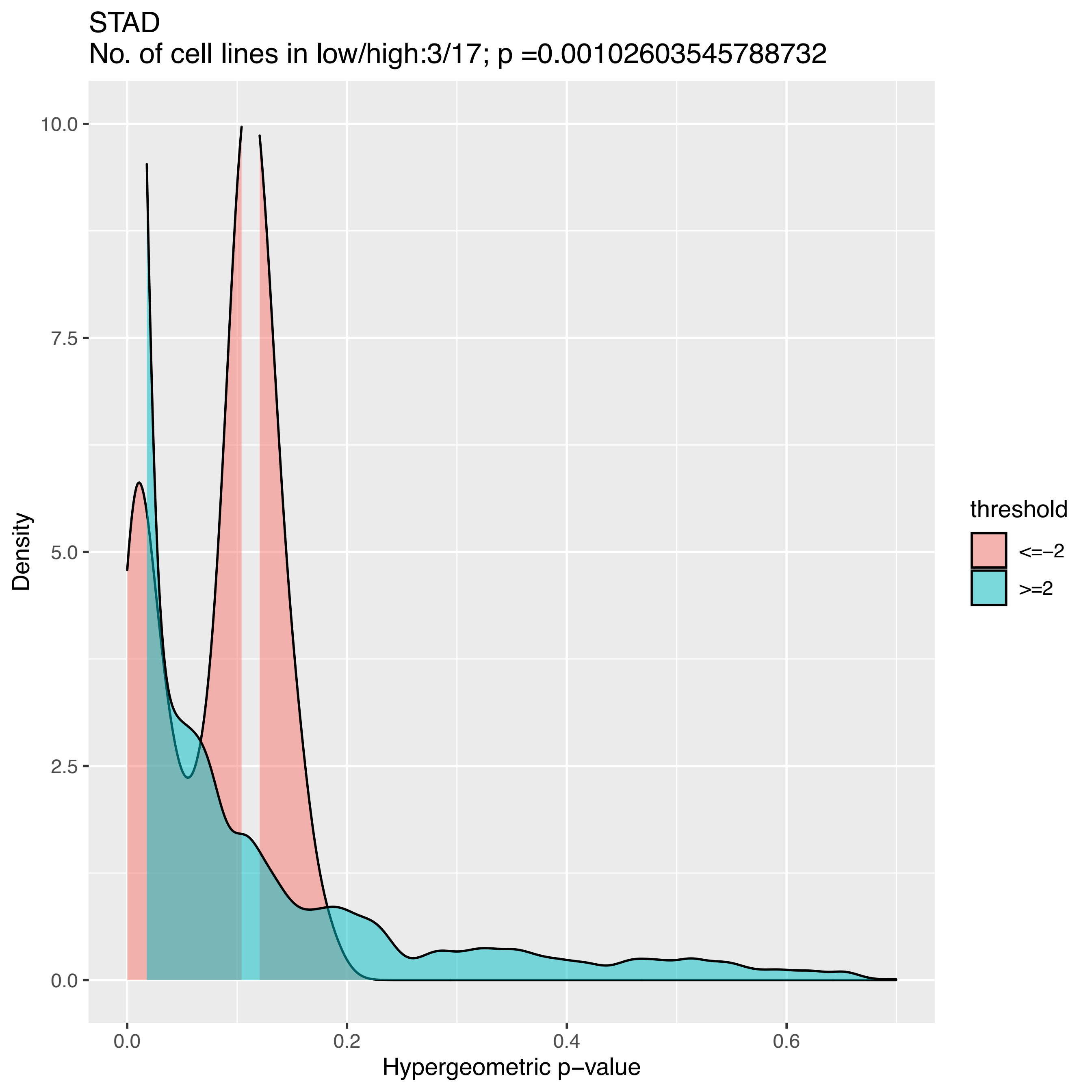
STAD
No. of cell lines in low/high:3/17; p =0.00102603545788732
Density
Hypergeometric p−value
threshold
<=−2
>=2
10.0
7.5
5.0
2.5
0.0
0.0
0.2
0.4
0.6

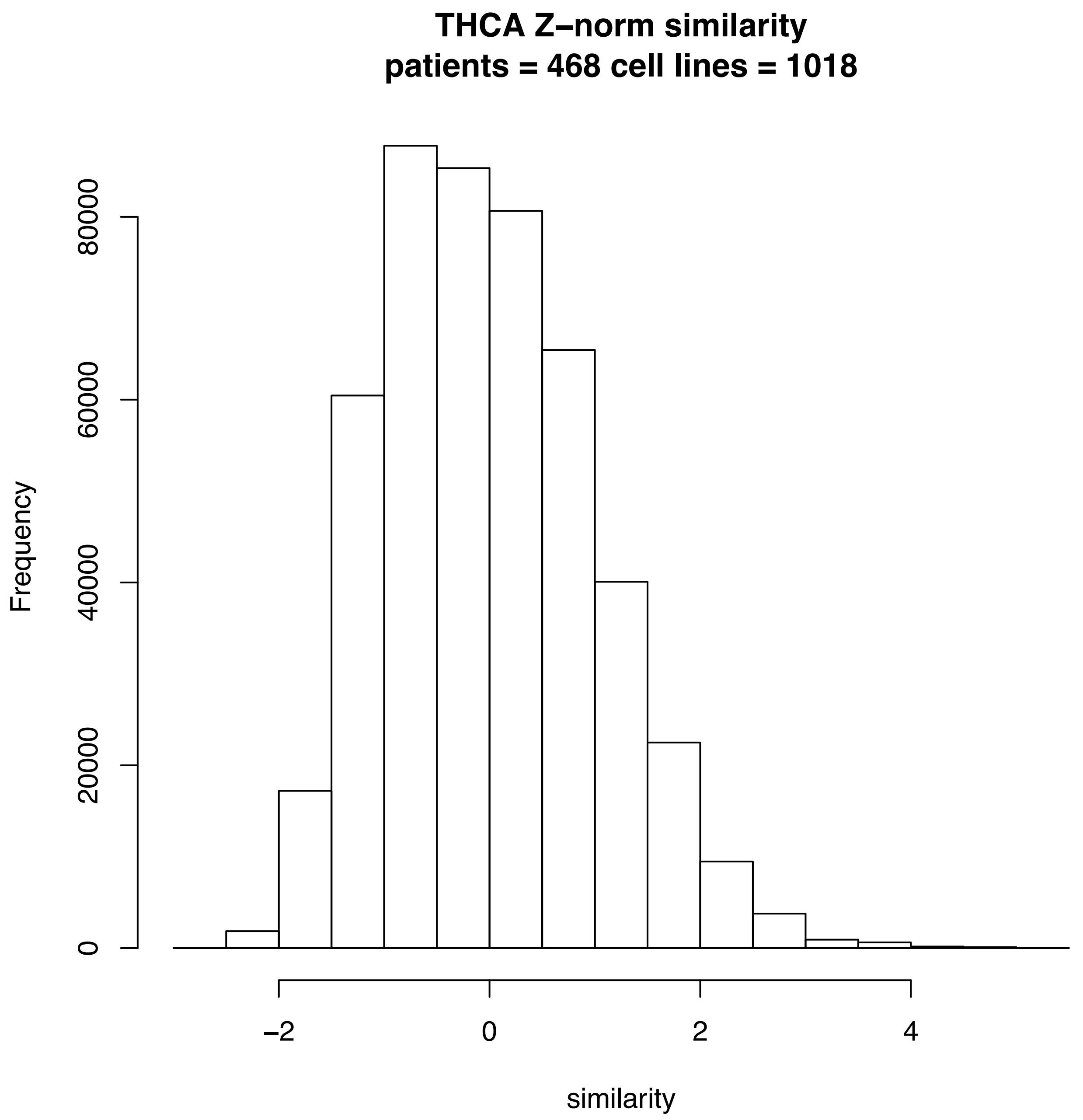
THCA Z–norm similarity
patients = 468 cell lines = 1018
Frequency
0
20000
40000
60000
80000
–2
0
2
4
similarity

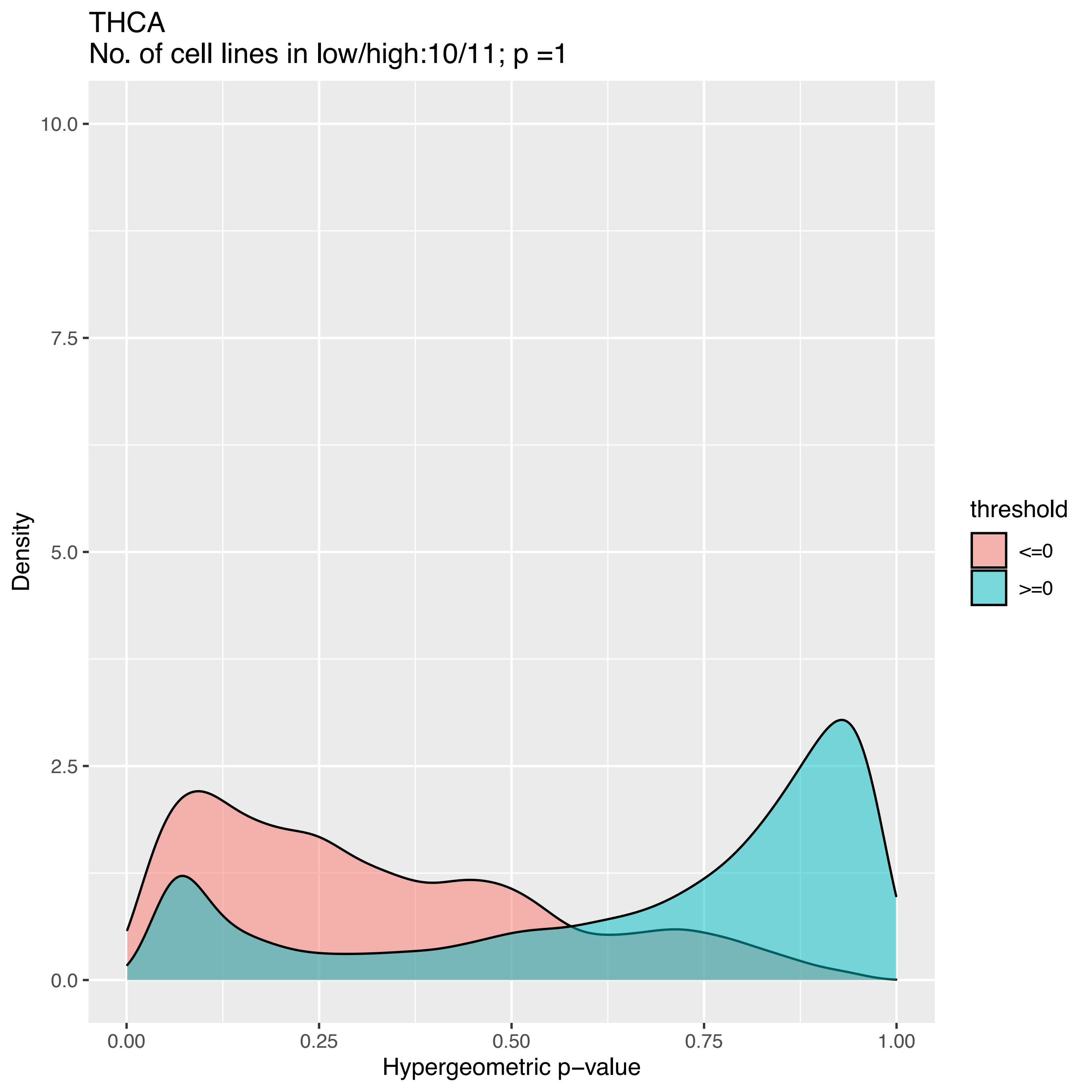
THCA
No. of cell lines in low/high:10/11; p =1
Density
10.0
7.5
5.0
2.5
0.0
0.00
0.25
0.50
0.75
1.00
Hypergeometric p−value
threshold
<=0
>=0

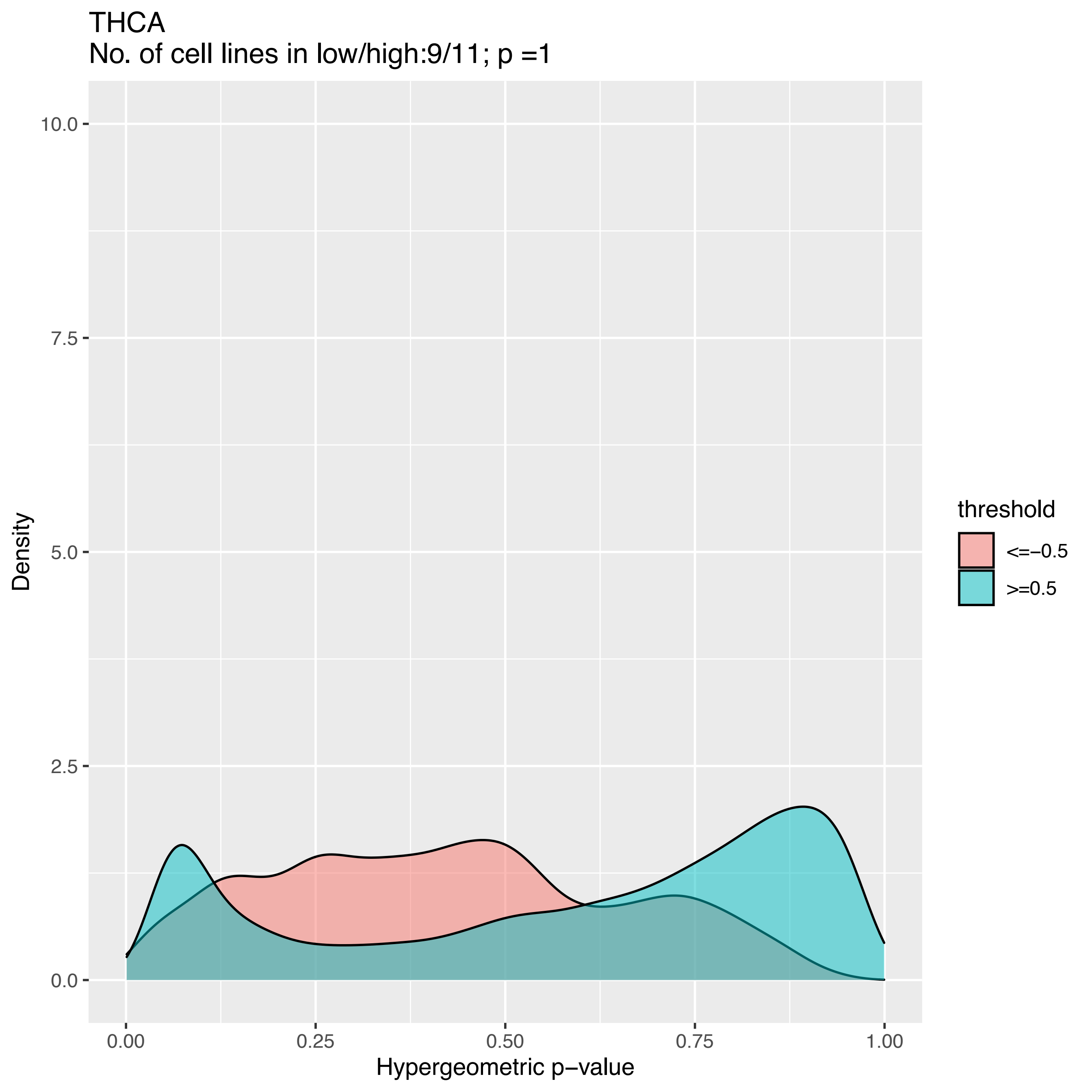
THCA
No. of cell lines in low/high:9/11; p =1
Density
10.0
7.5
5.0
2.5
0.0
0.00
0.25
0.50
0.75
1.00
Hypergeometric p−value
threshold
<=−0.5
>=0.5

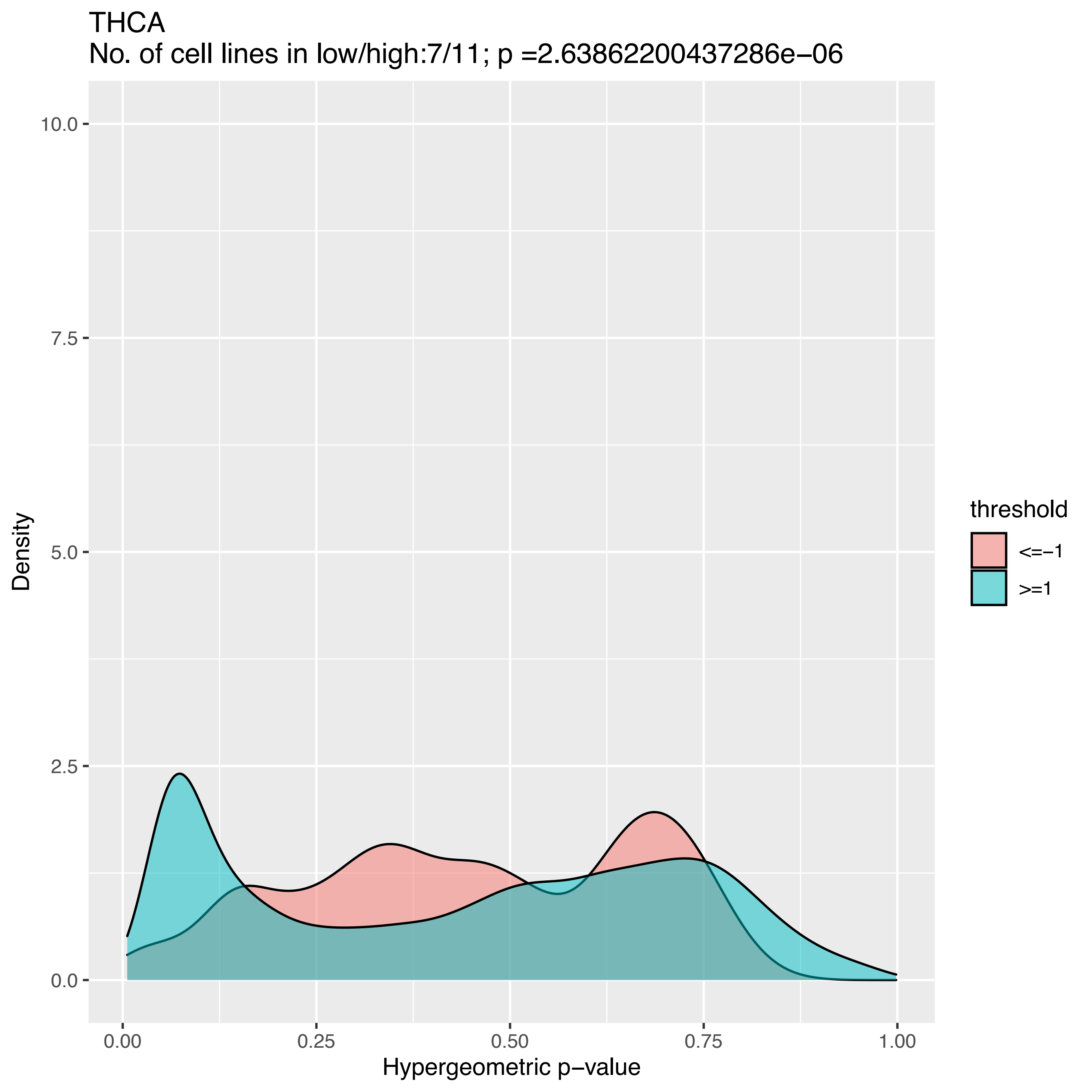
THCA
No. of cell lines in low/high:7/11; p =2.63862200437286e−06
Density
Hypergeometric p−value
0.0
2.5
5.0
7.5
10.0
0.00
0.25
0.50
0.75
1.00
threshold
<=−1
>=1

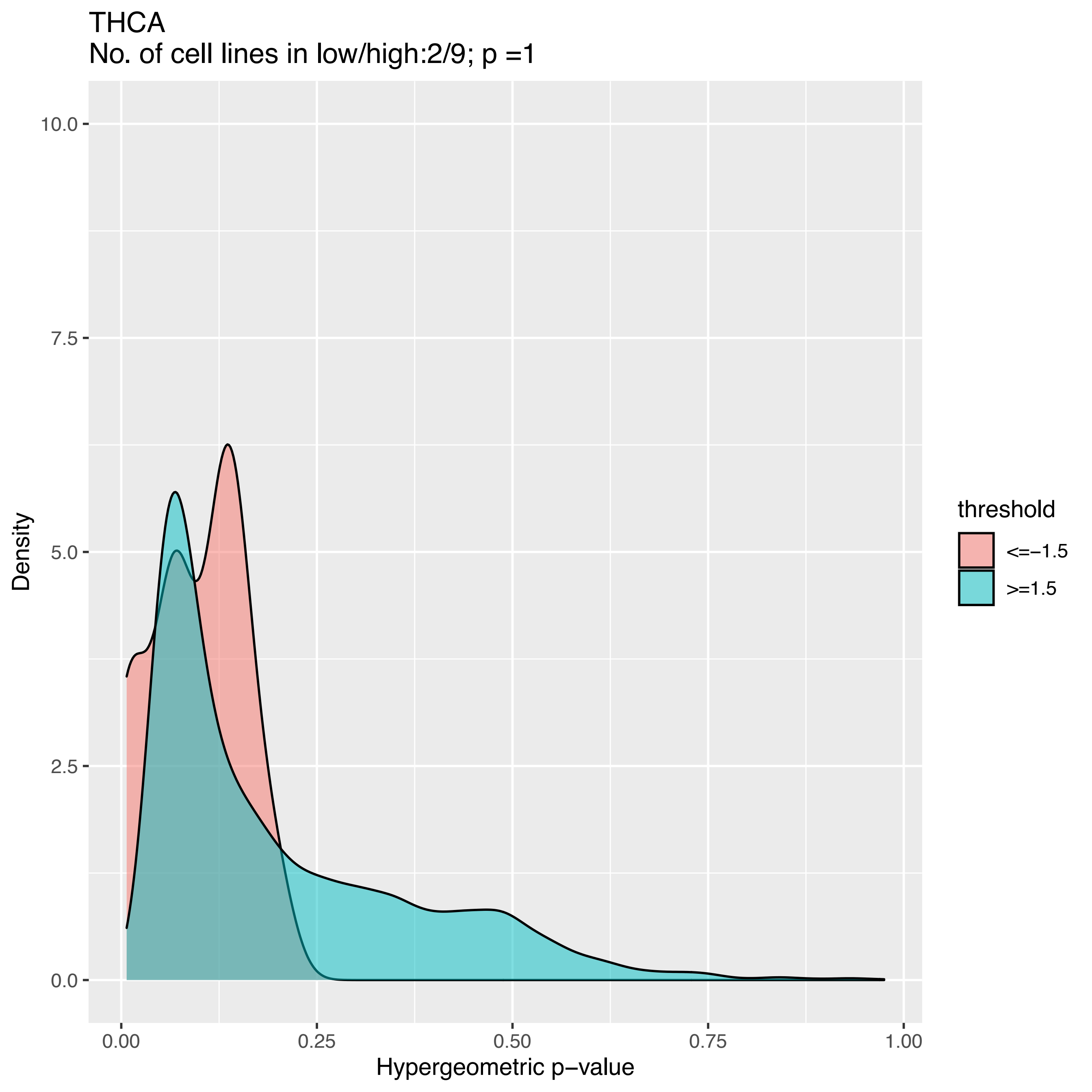
THCA
No. of cell lines in low/high:2/9; p =1
10.0
7.5
5.0
2.5
0.0
Density
0.00
0.25
0.50
0.75
1.00
Hypergeometric p−value
threshold
<=−1.5
>=1.5

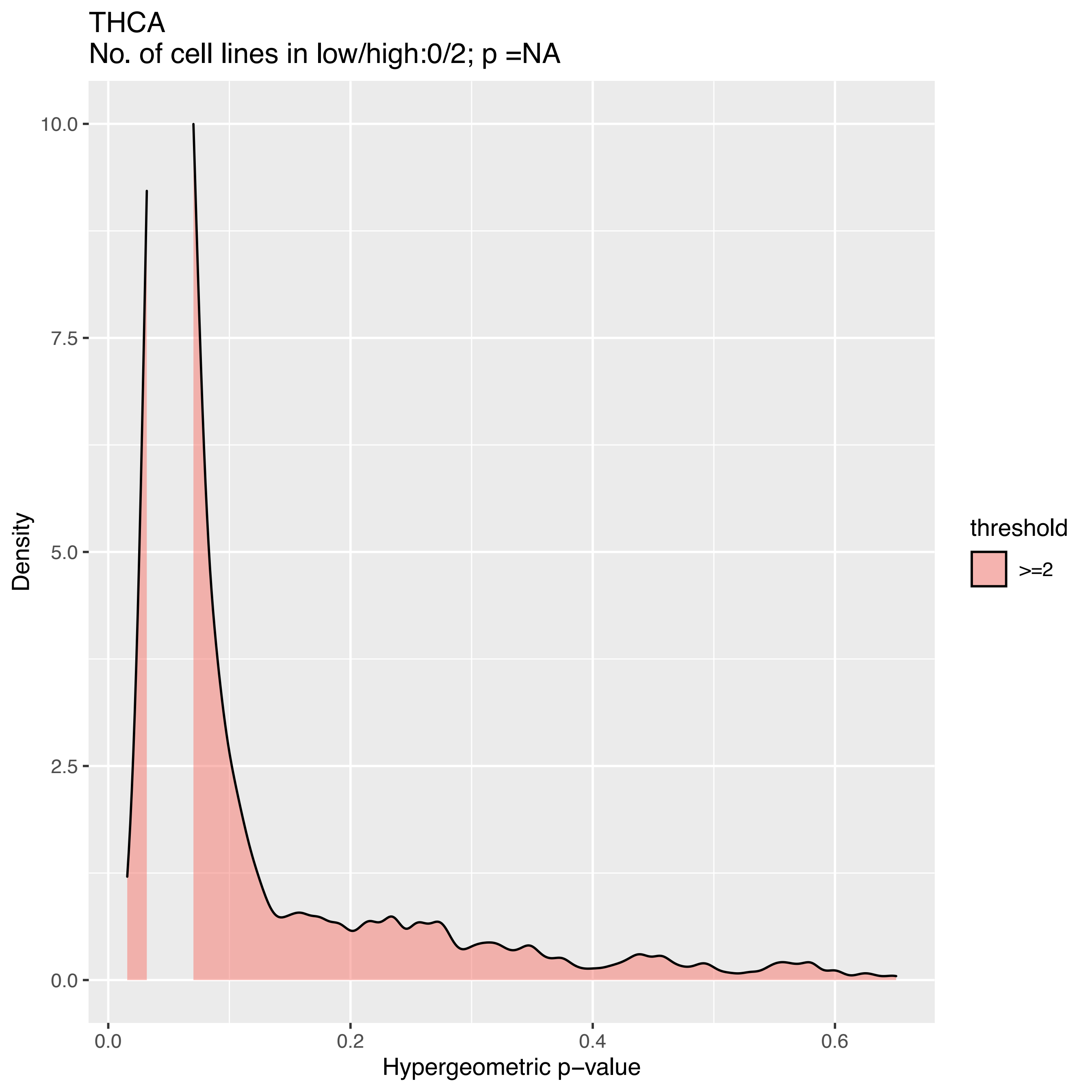
THCA
No. of cell lines in low/high:0/2; p =NA
Density
10.0
7.5
5.0
2.5
0.0
0.0
0.2
0.4
0.6
Hypergeometric p−value
threshold
>=2

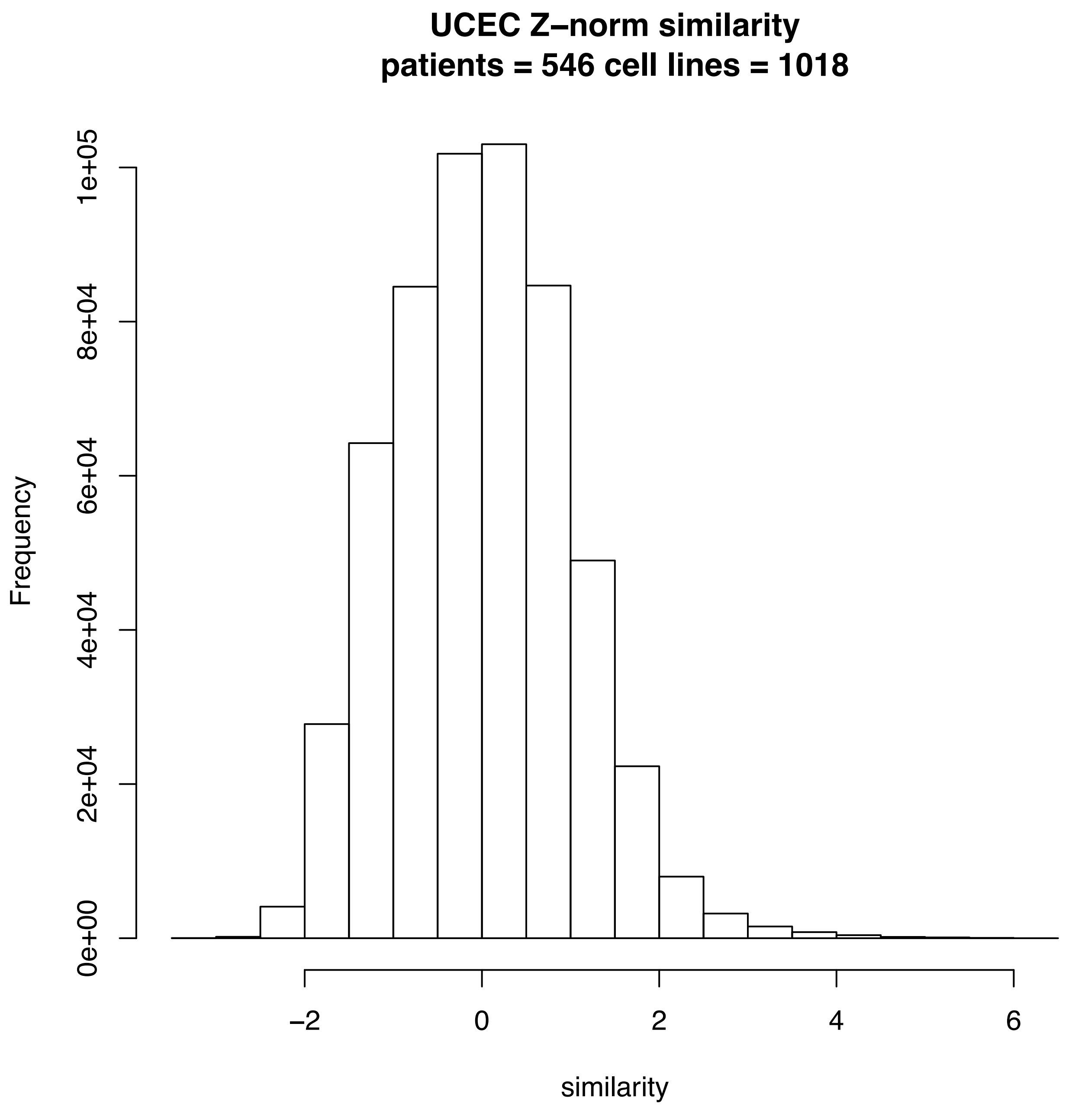
UCEC Z–norm similarity
patients = 546 cell lines = 1018
Frequency
0e+00
2e+04
4e+04
6e+04
8e+04
1e+05
–2
0
2
4
6
similarity

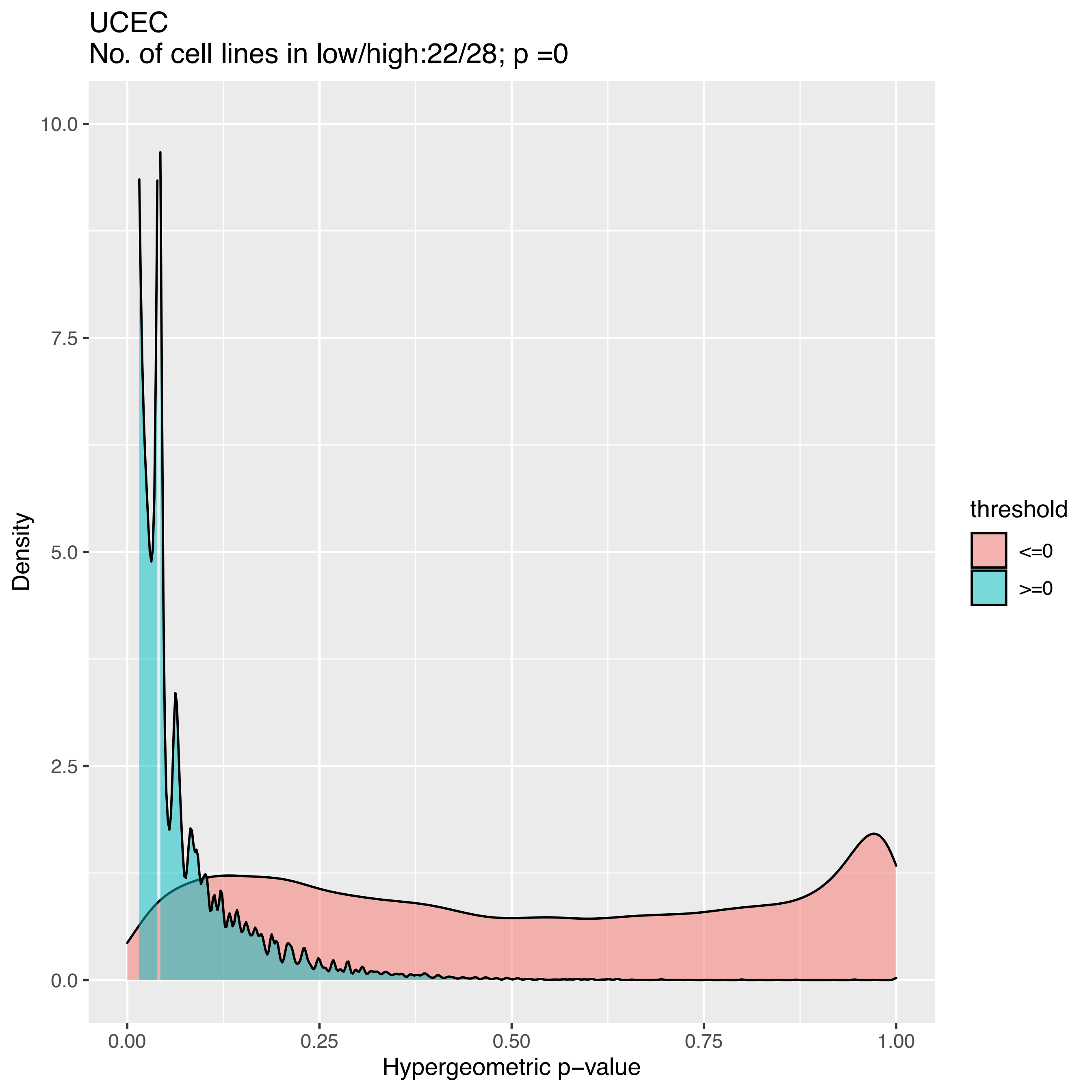
UCEC
No. of cell lines in low/high:22/28; p =0
Density
10.0
7.5
5.0
2.5
0.0
0.00
0.25
0.50
0.75
1.00
Hypergeometric p−value
threshold
<=0
>=0

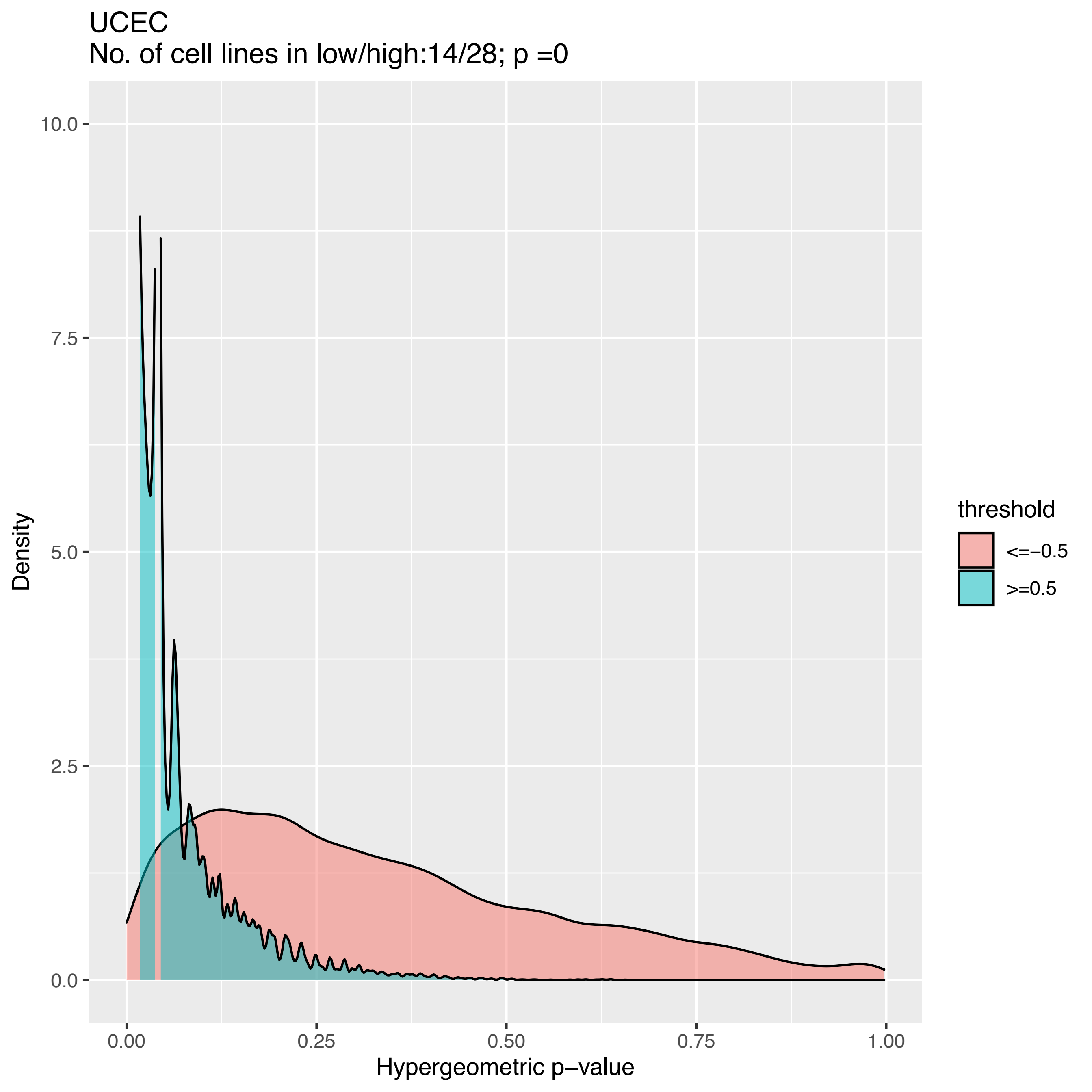
UCEC
No. of cell lines in low/high:14/28; p =0
Density
10.0
7.5
5.0
2.5
0.0
0.00
0.25
0.50
0.75
1.00
Hypergeometric p−value
threshold
<=−0.5
>=0.5

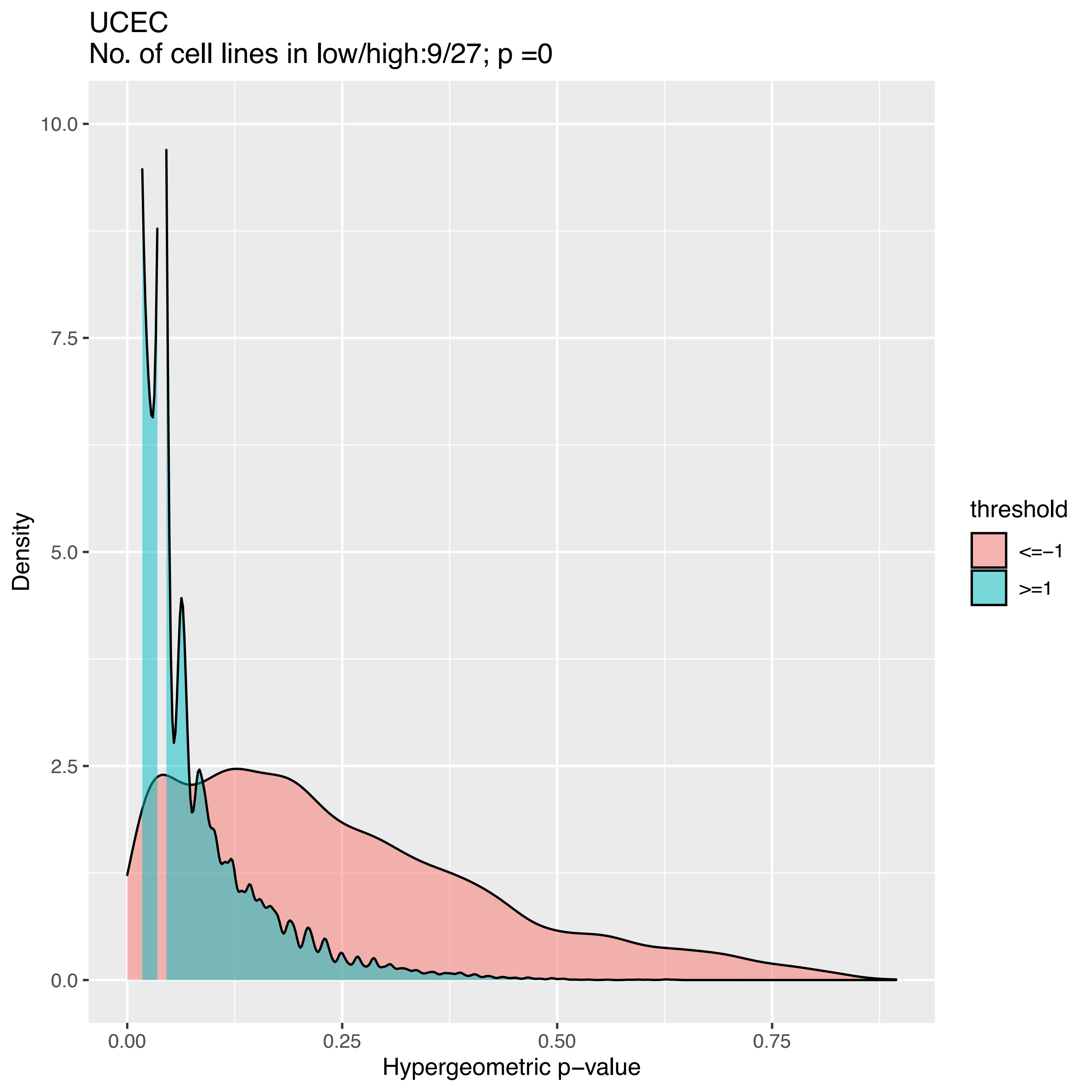
UCEC
No. of cell lines in low/high:9/27; p =0
Density
10.0
7.5
5.0
2.5
0.0
0.00
0.25
0.50
0.75
Hypergeometric p−value
threshold
<=−1
>=1

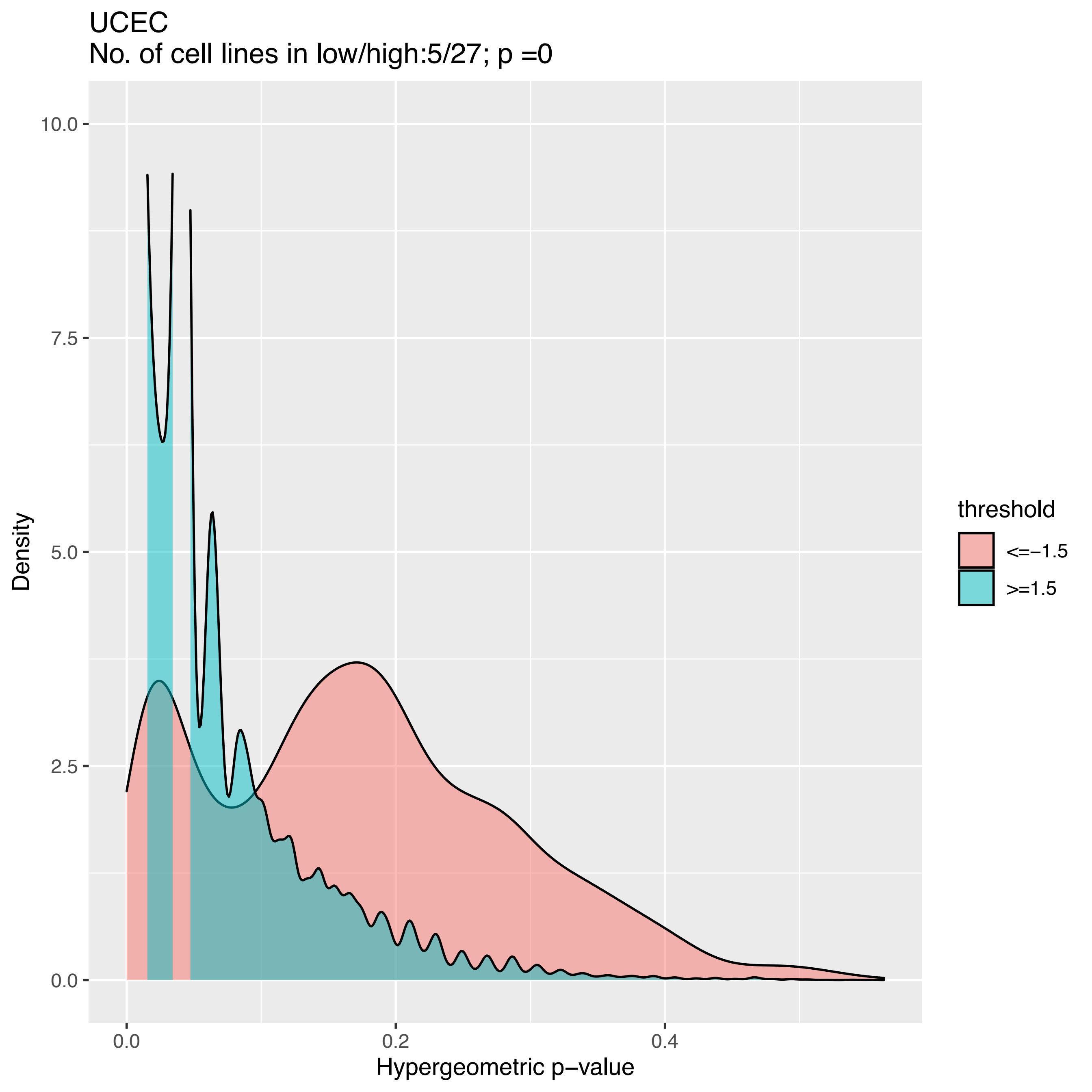
UCEC
No. of cell lines in low/high:5/27; p =0
Density
Hypergeometric p−value
10.0
7.5
5.0
2.5
0.0
0.0
0.2
0.4
threshold
<=−1.5
>=1.5

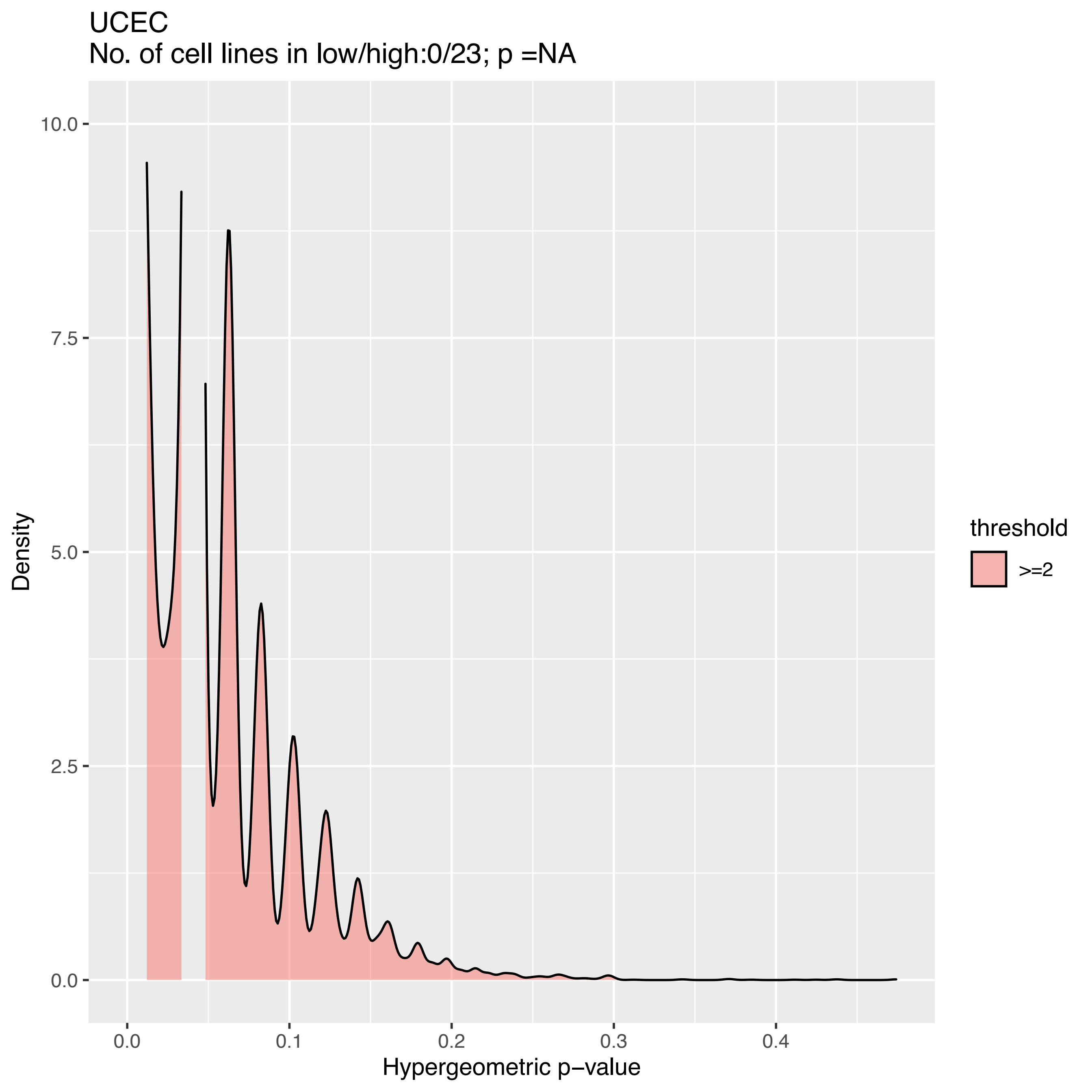
UCEC
No. of cell lines in low/high:0/23; p =NA
Density
10.0
7.5
5.0
2.5
0.0
0.0
0.1
0.2
0.3
0.4
Hypergeometric p−value
threshold
>=2

### TSI method

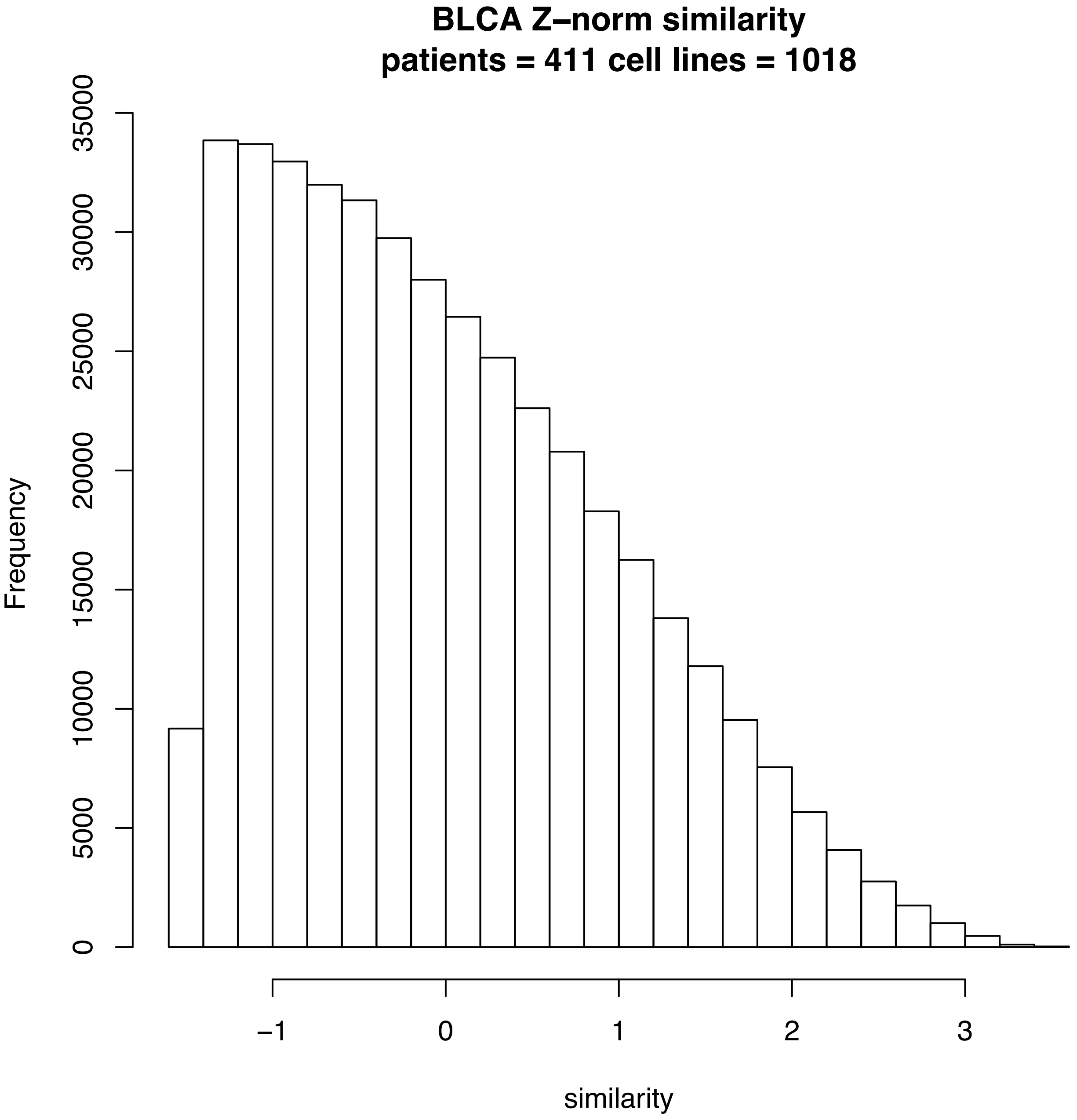
BLCA Z–norm similarity
patients = 411 cell lines = 1018
Frequency
0
5000
10000
15000
20000
25000
30000
35000
–1
0
1
2
3
similarity

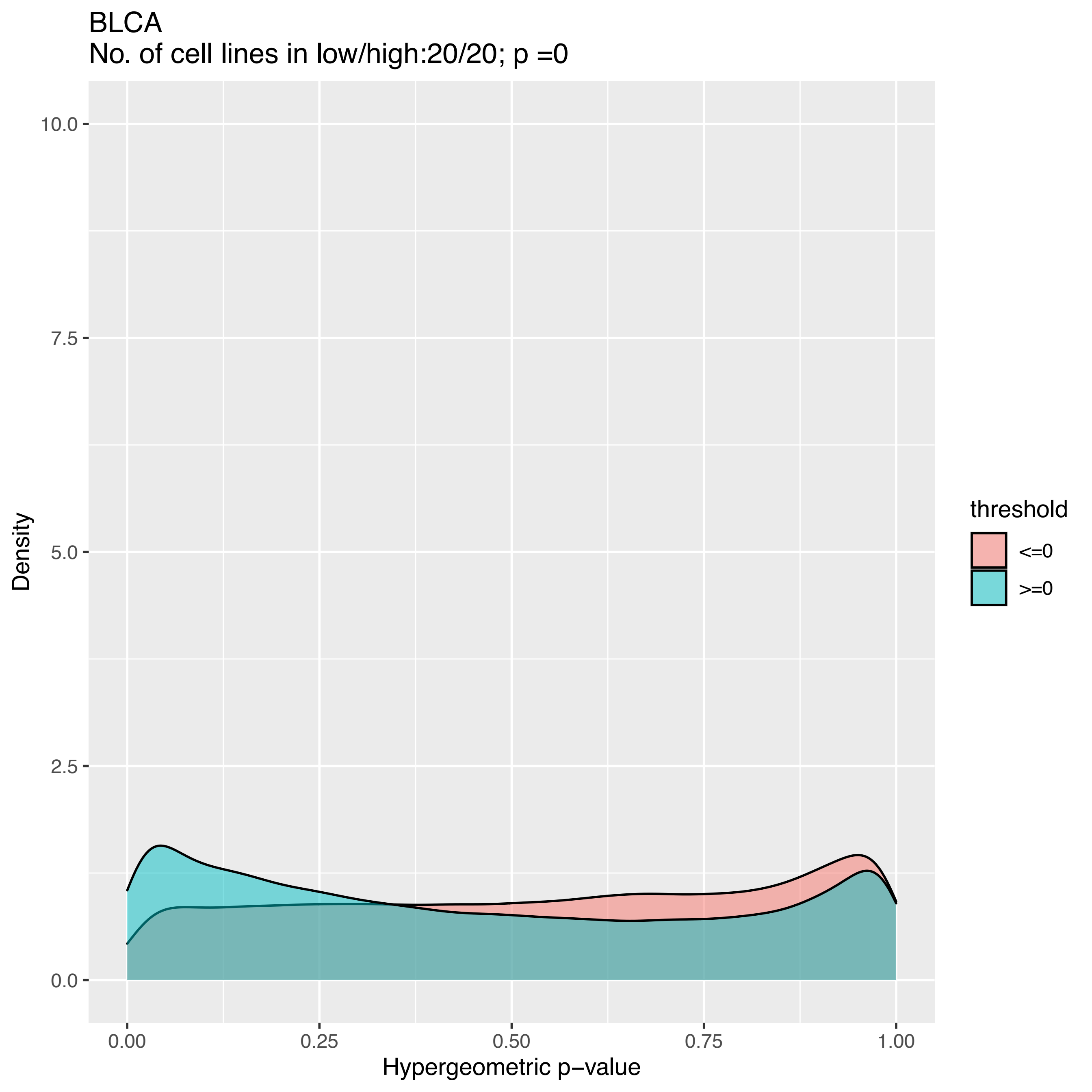
BLCA
No. of cell lines in low/high:20/20; p =0
Density
10.0
7.5
5.0
2.5
0.0
0.00
0.25
0.50
0.75
1.00
Hypergeometric p−value
threshold
<=0
>=0

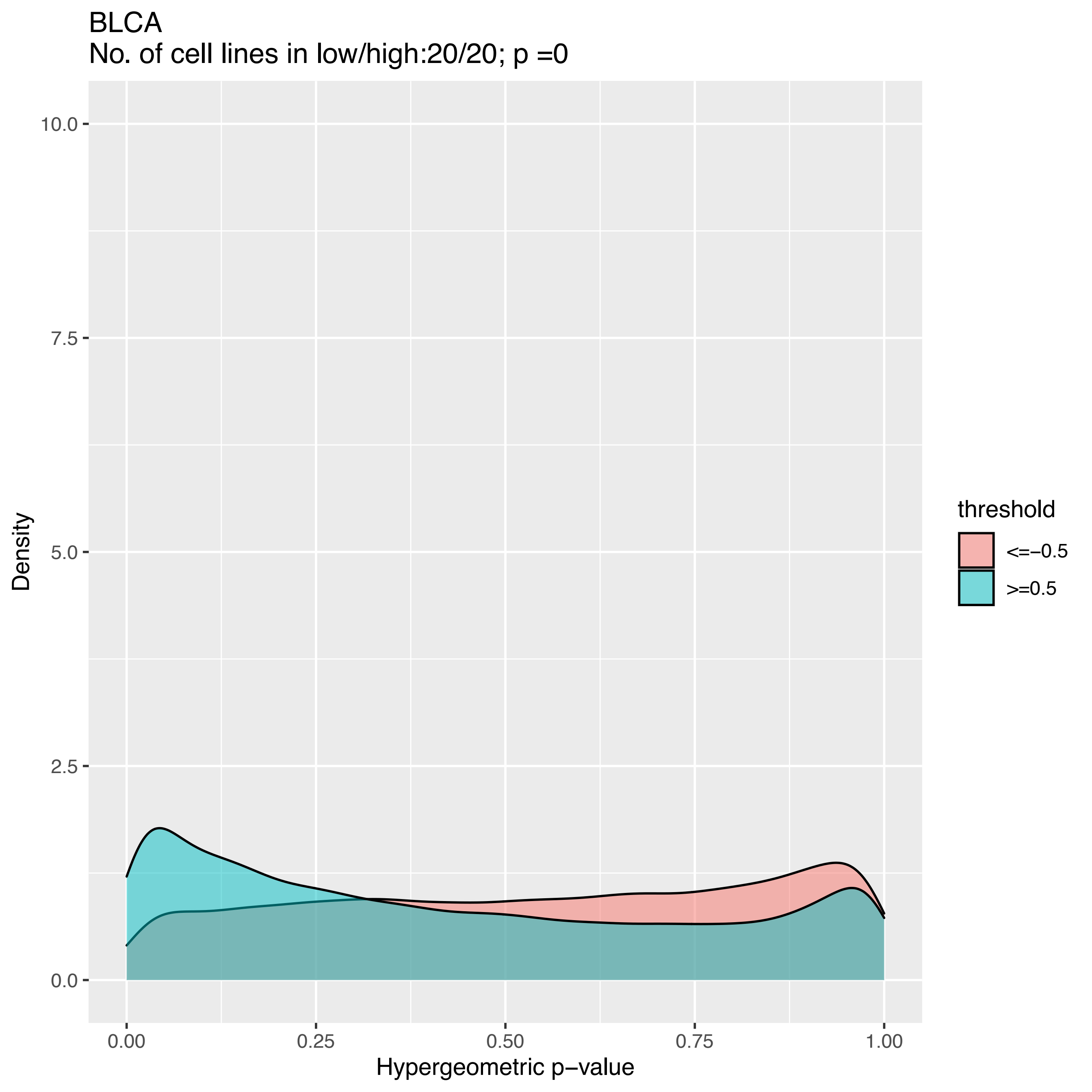
BLCA
No. of cell lines in low/high:20/20; p =0
10.0
7.5
5.0
2.5
0.0
Density
0.00
0.25
0.50
0.75
1.00
Hypergeometric p−value
threshold
<=−0.5
>=0.5

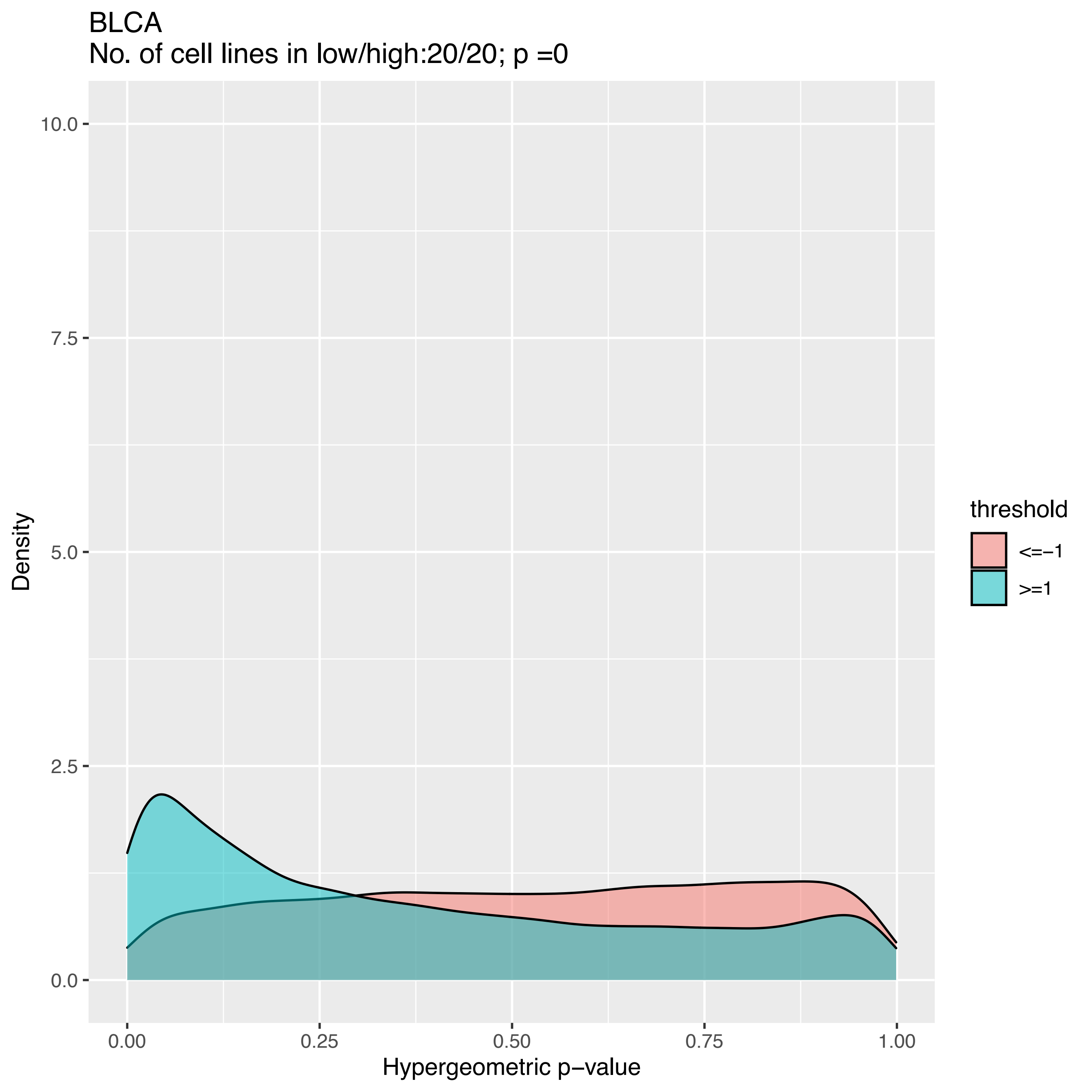
BLCA
No. of cell lines in low/high:20/20; p =0
Density
Hypergeometric p−value
threshold
<=−1
>=1
0.0
2.5
5.0
7.5
10.0
0.00
0.25
0.50
0.75
1.00

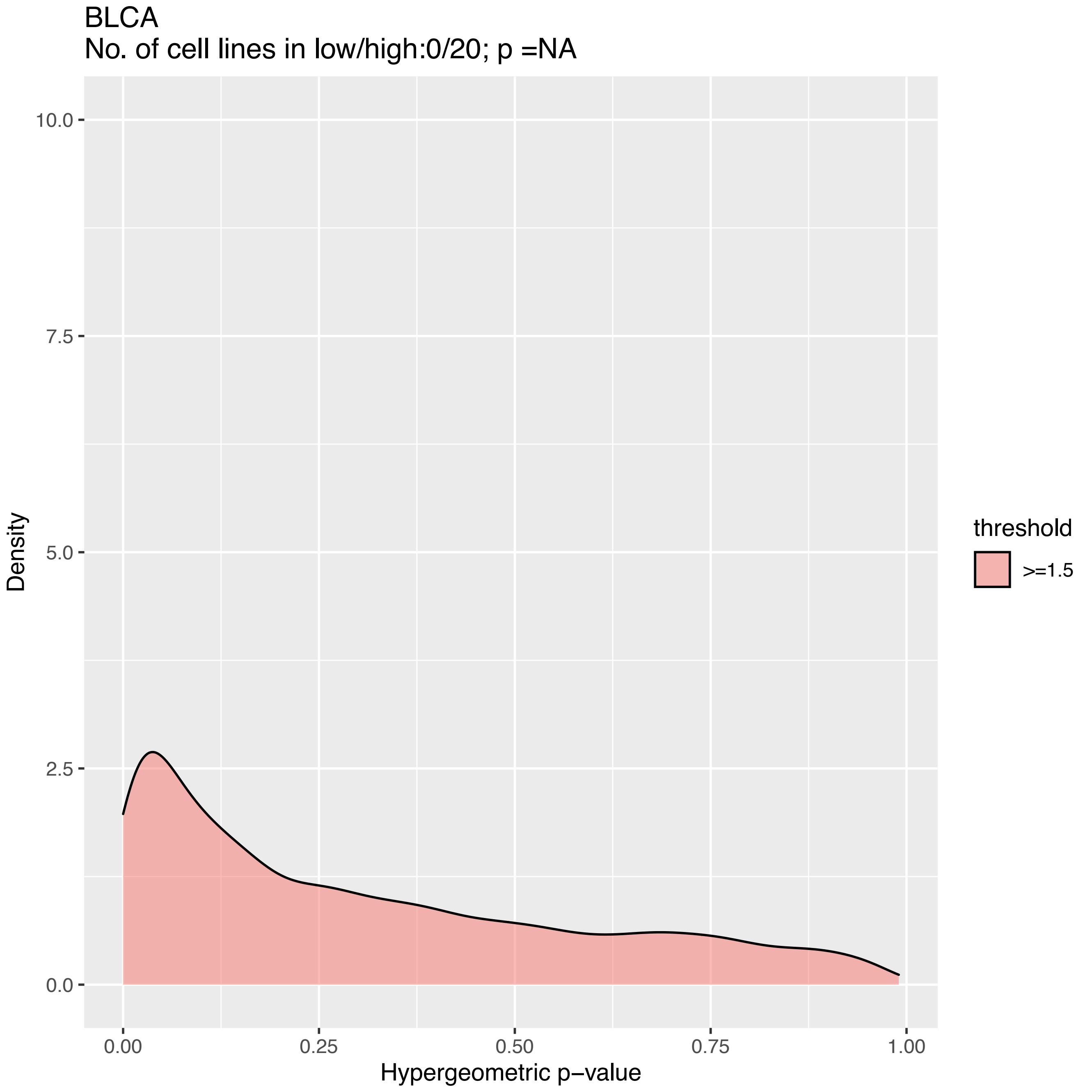

BLCA
No. of cell lines in low/high:0/20; p =NA
Density
10.0
7.5
5.0
2.5
0.0
0.00
0.25
0.50
0.75
1.00
Hypergeometric p−value
threshold
>=1.5

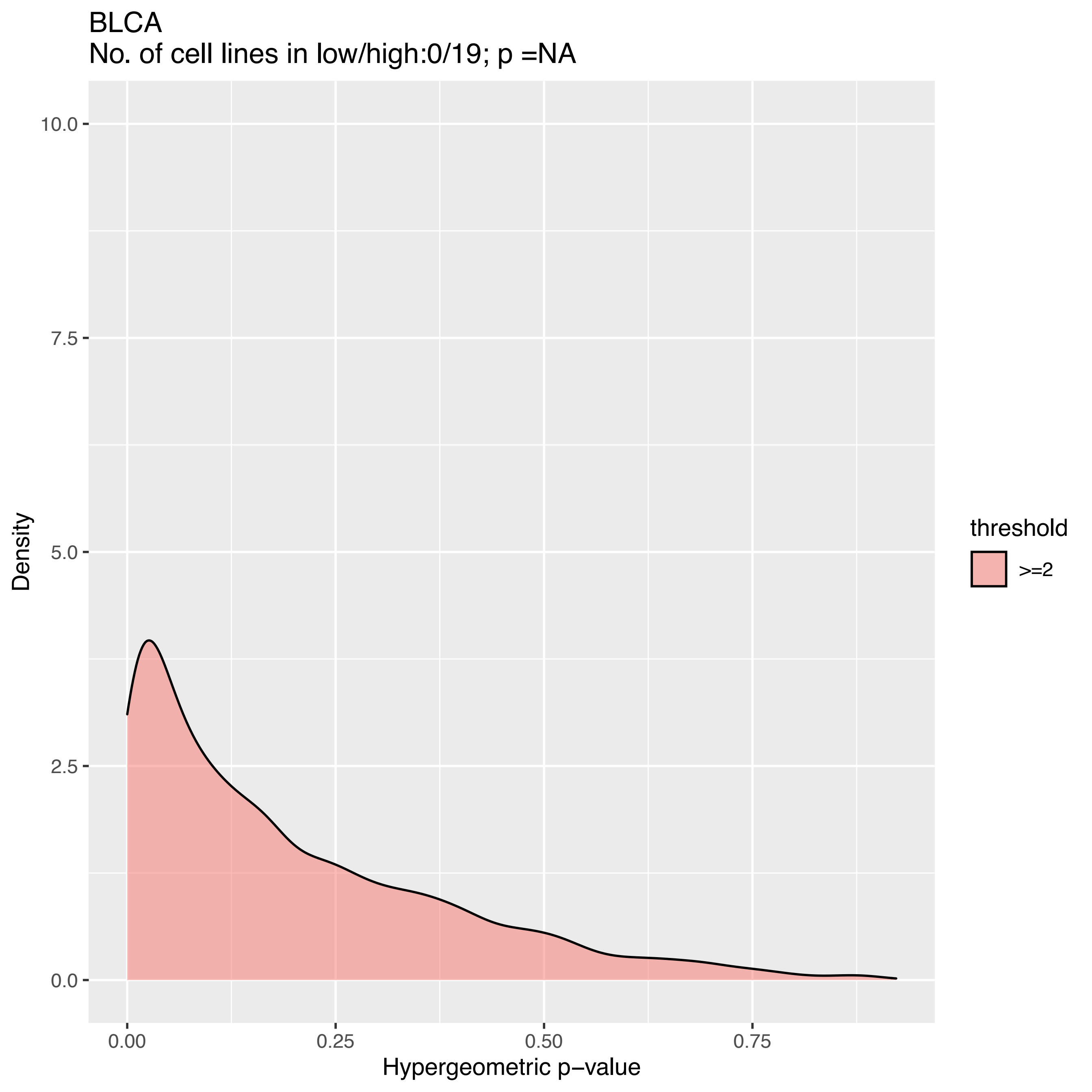

BLCA
No. of cell lines in low/high:0/19; p =NA
10.0
7.5
5.0
2.5
0.0
Density
0.00
0.25
0.50
0.75
Hypergeometric p−value
threshold
>=2

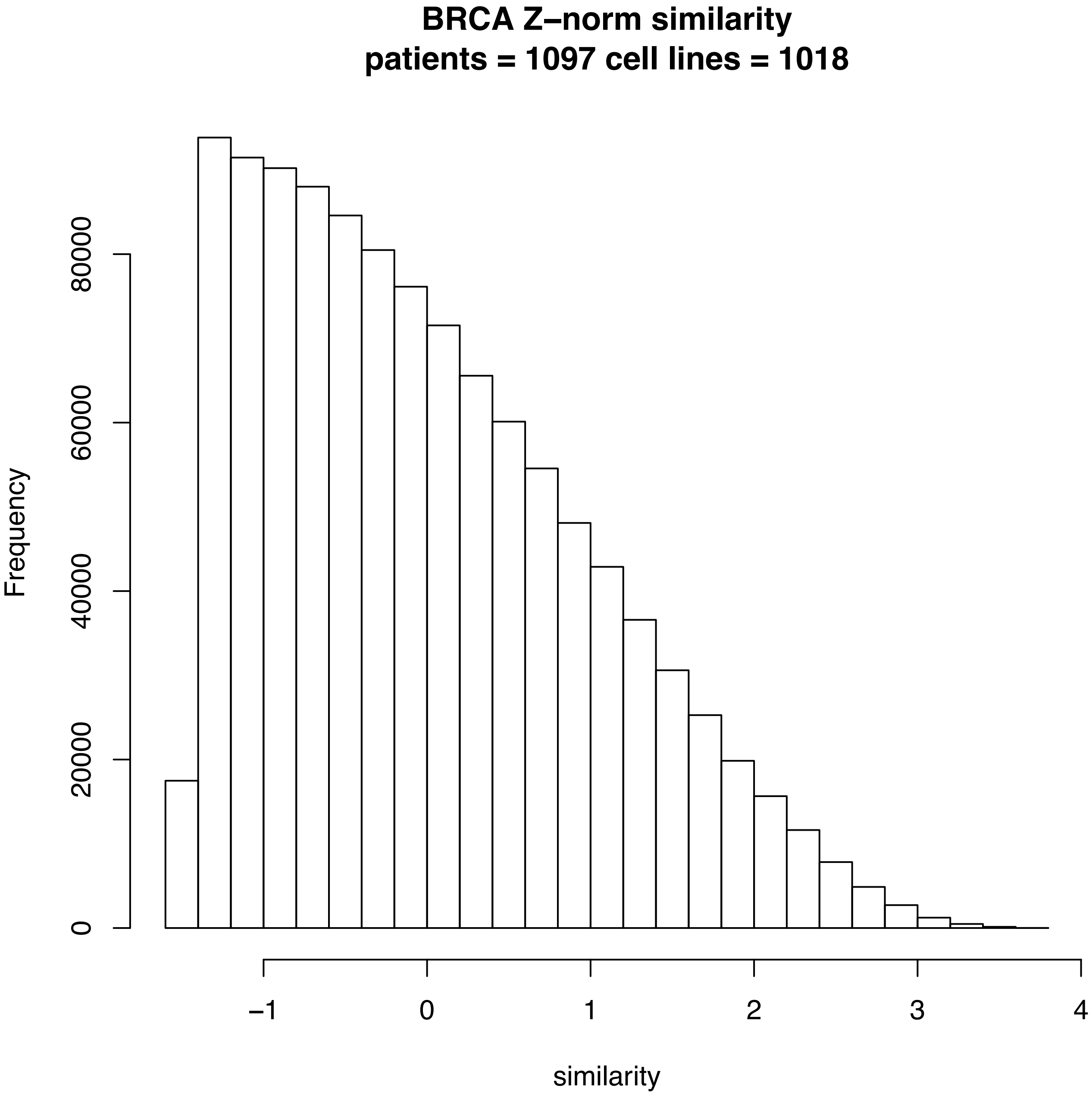
BRCA Z−norm similarity
patients = 1097 cell lines = 1018
Frequency
0
20000
40000
60000
80000
−1
0
1
2
3
4
similarity

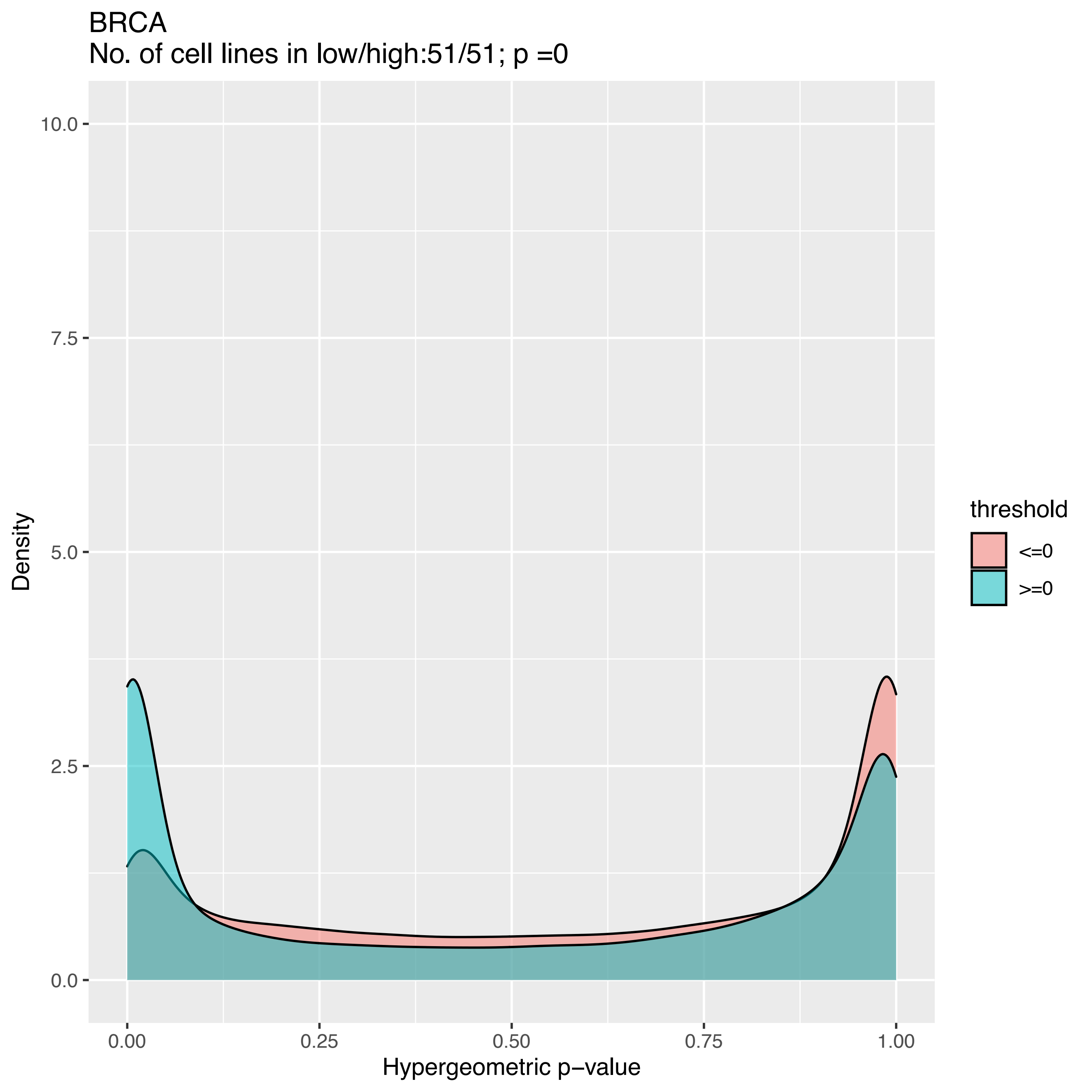

BRCA
No. of cell lines in low/high:51/51; p =0
Density
10.0
7.5
5.0
2.5
0.0
0.00
0.25
0.50
0.75
1.00
Hypergeometric p−value
threshold
<=0
>=0

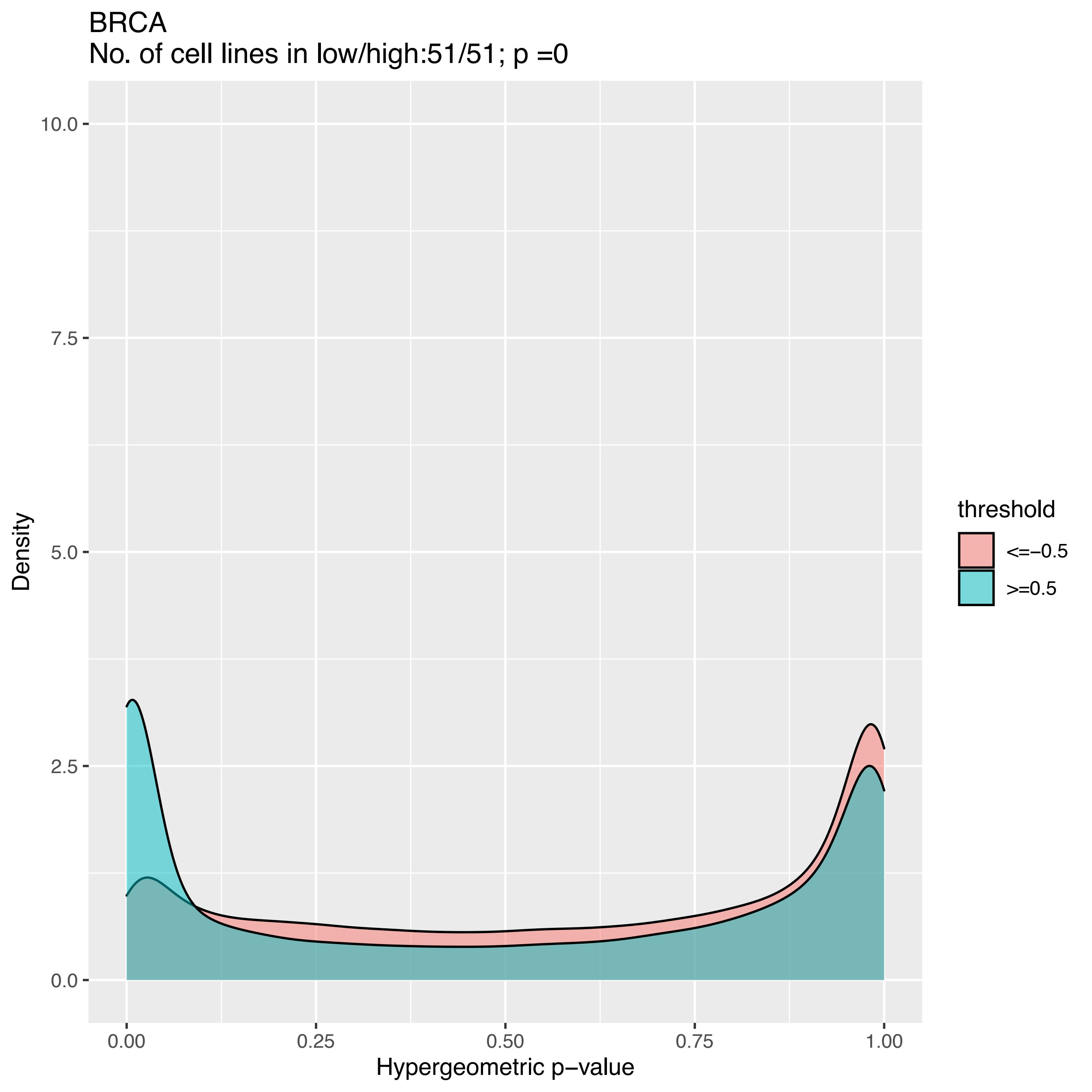

BRCA
No. of cell lines in low/high:51/51; p =0
Density
10.0
7.5
5.0
2.5
0.0
0.00
0.25
0.50
0.75
1.00
Hypergeometric p−value
threshold
<=−0.5
>=0.5

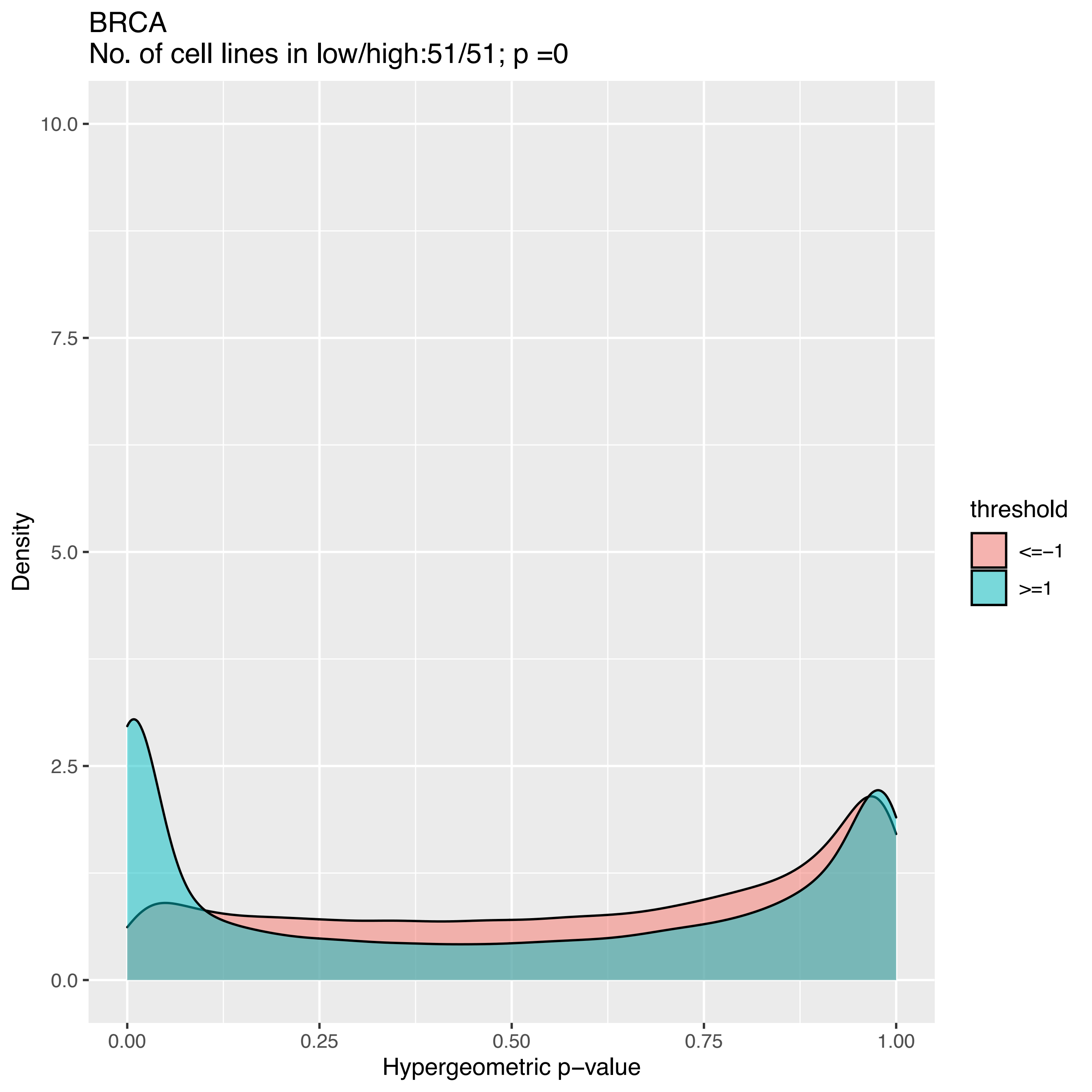
BRCA
No. of cell lines in low/high:51/51; p =0
10.0
7.5
5.0
2.5
0.0
0.00
0.25
0.50
0.75
1.00
Density
Hypergeometric p−value
threshold
<=−1
>=1

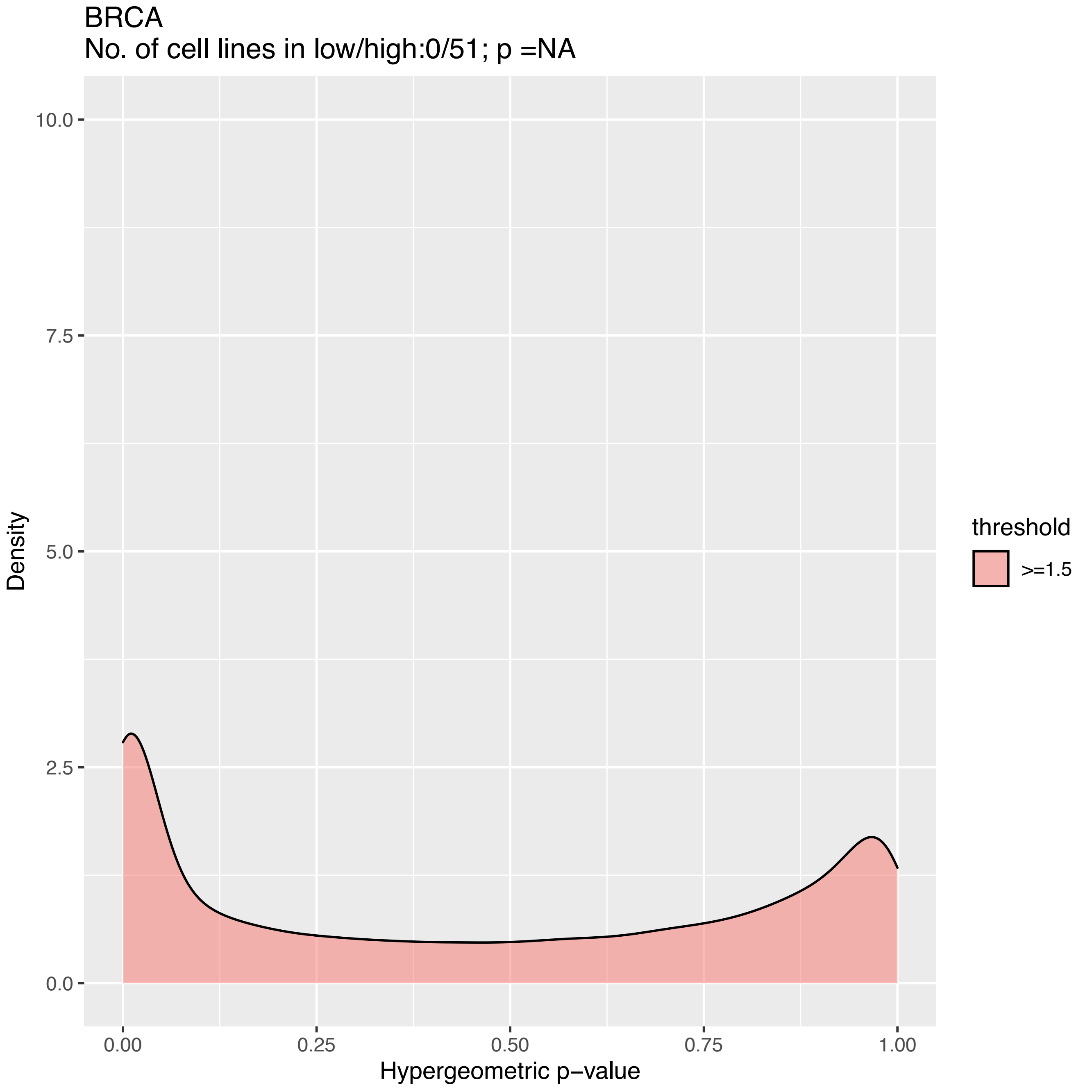
BRCA
No. of cell lines in low/high:0/51; p =NA
Density
10.0
7.5
5.0
2.5
0.0
0.00
0.25
0.50
0.75
1.00
Hypergeometric p−value
threshold
>=1.5

BRCA
No. of cell lines in low/high:0/51; p =NA

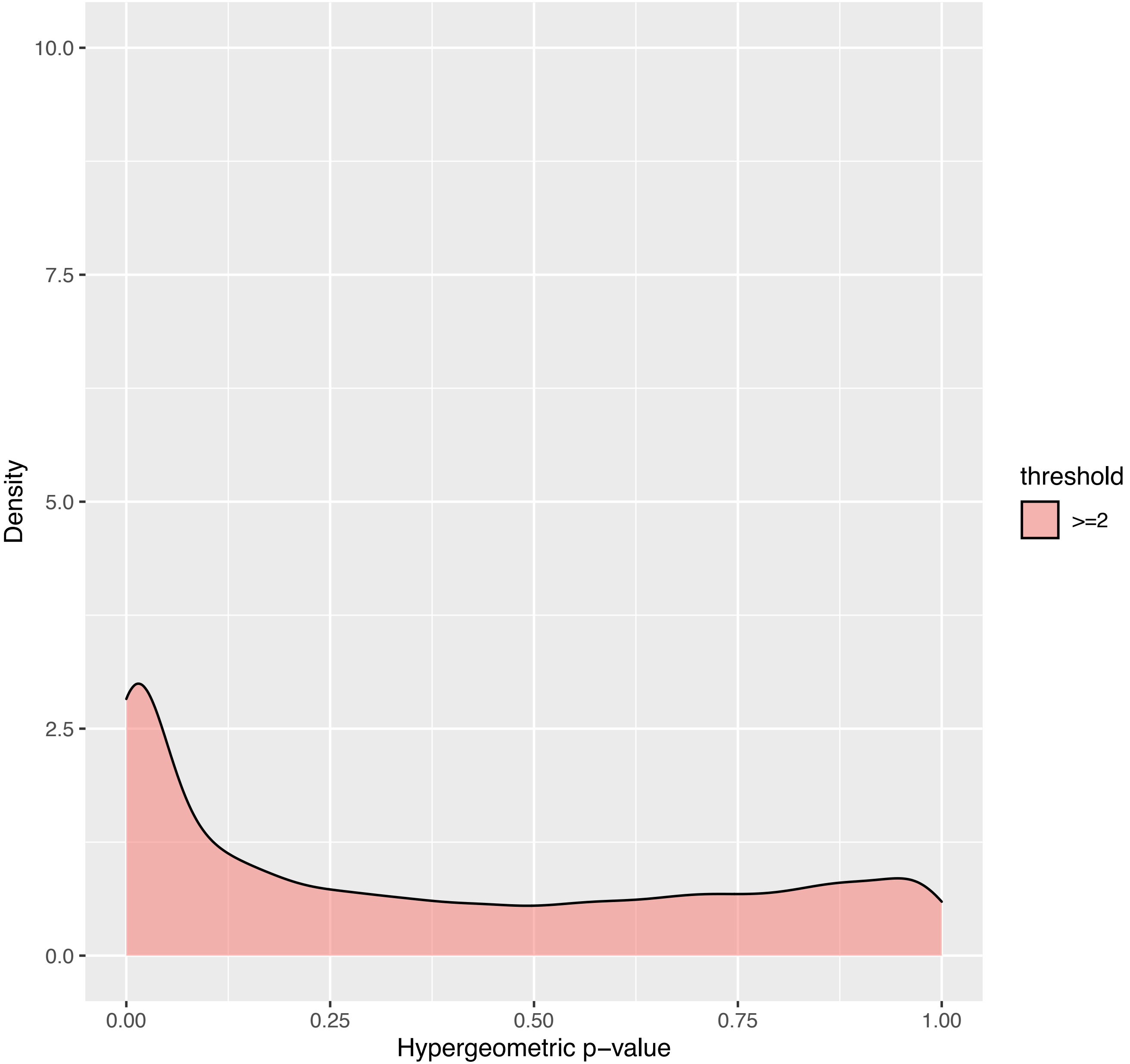

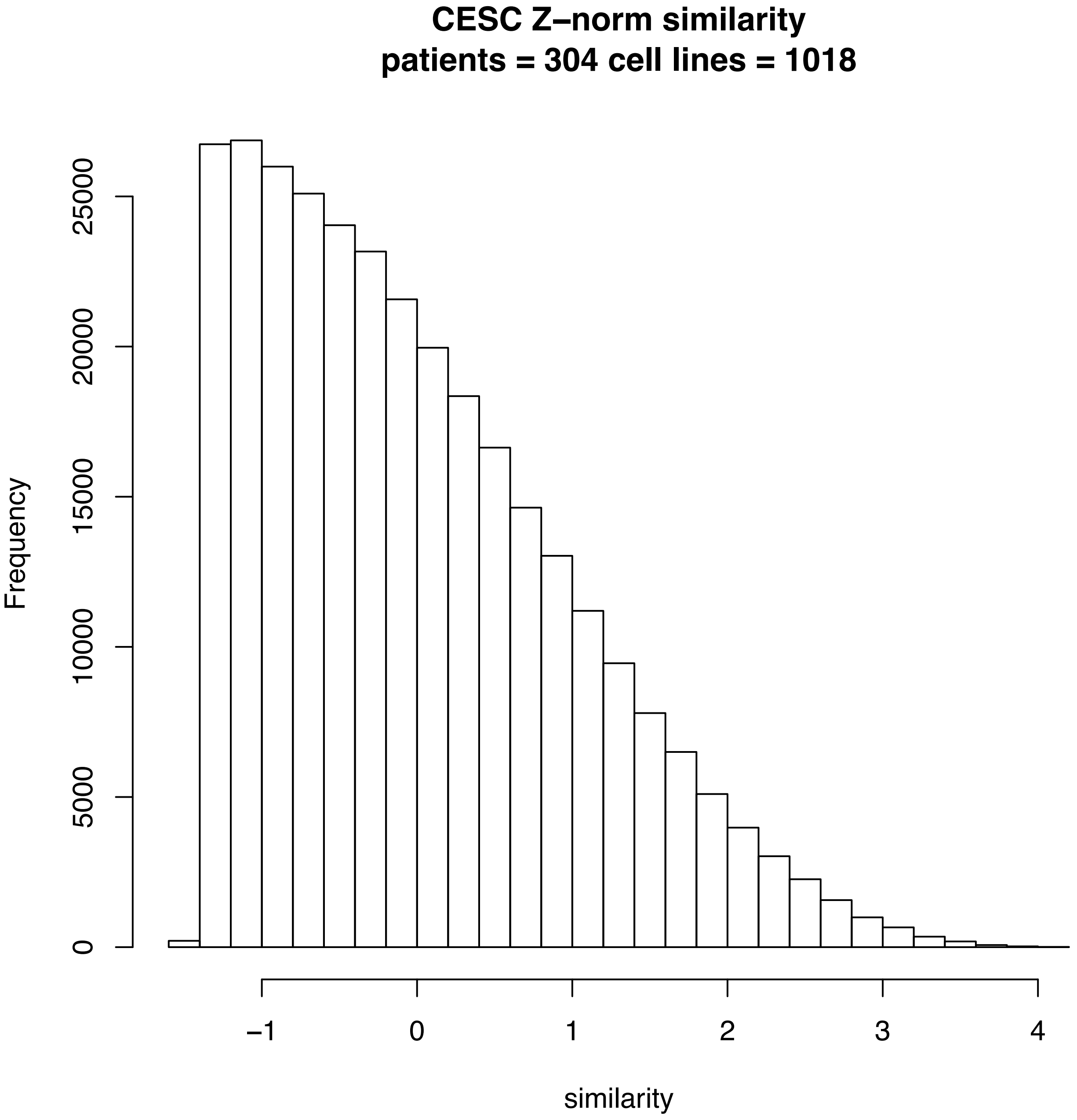
CESC Z–norm similarity
patients = 304 cell lines = 1018
Frequency
0
5000
10000
15000
20000
25000
−1
0
1
2
3
4
similarity

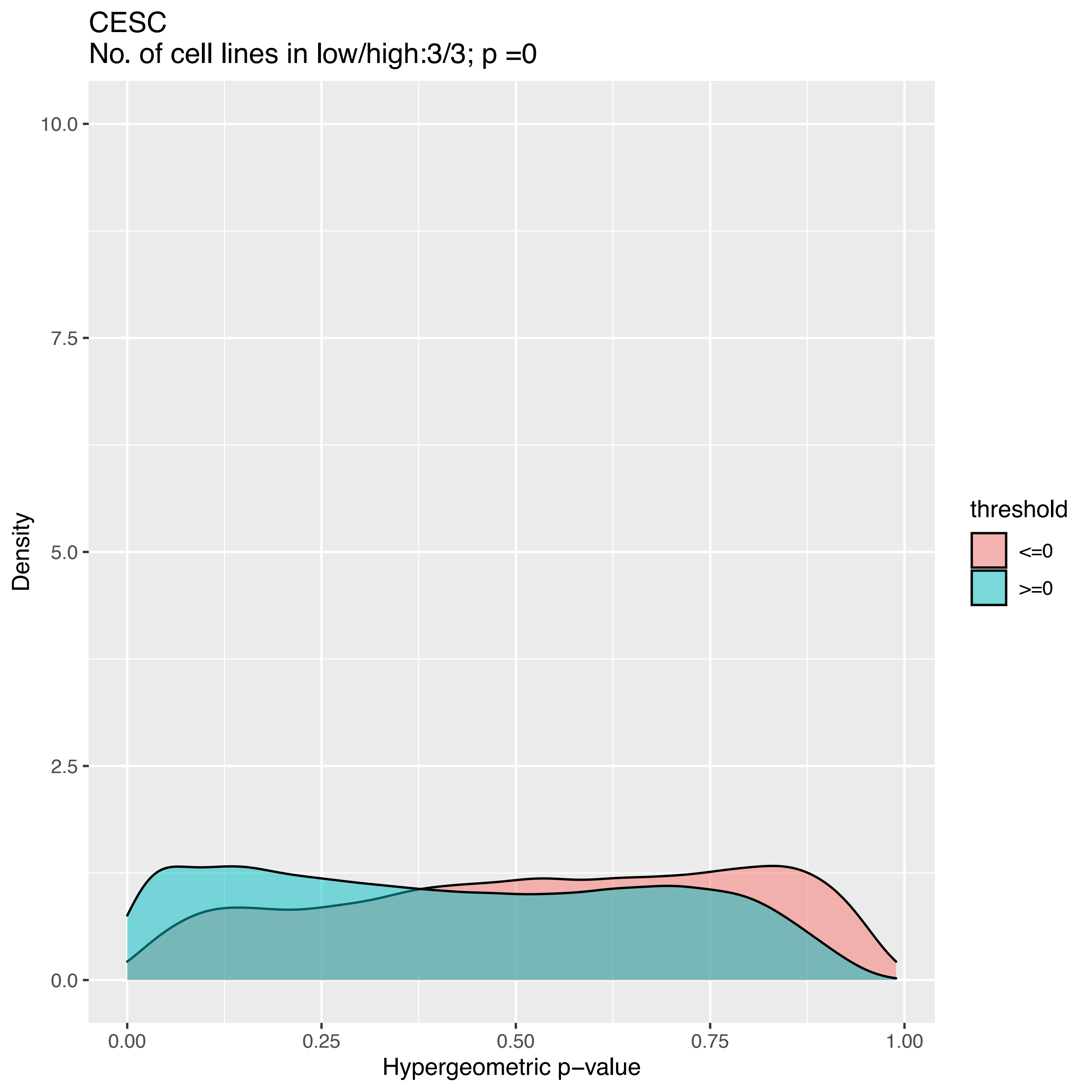

CESC
No. of cell lines in low/high:3/3; p =0
Density
10.0
7.5
5.0
2.5
0.0
0.00
0.25
0.50
0.75
1.00
Hypergeometric p−value
threshold
<=0
>=0

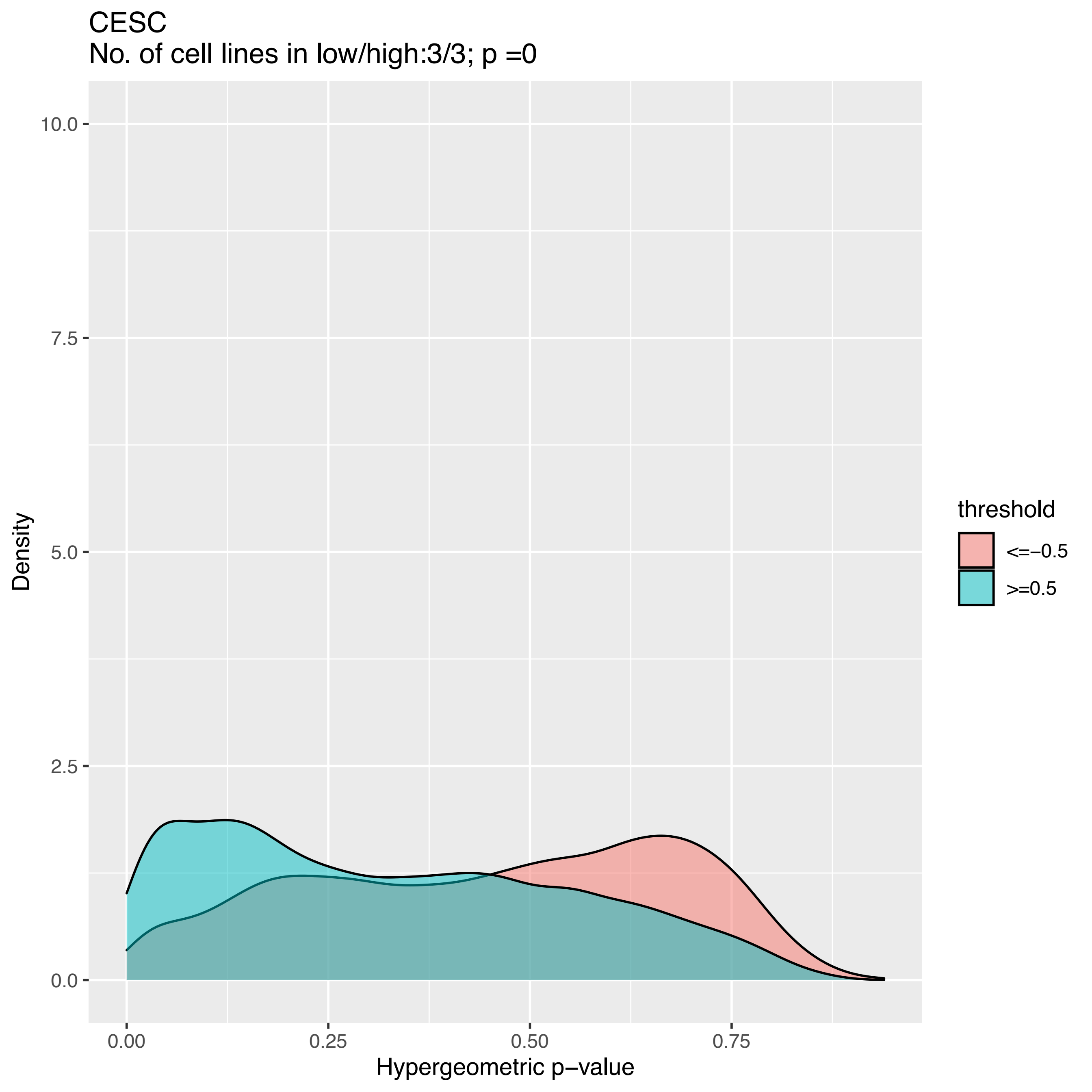
CESC
No. of cell lines in low/high:3/3; p =0
Density
Hypergeometric p−value
threshold
<=−0.5
>=0.5
0.0
2.5
5.0
7.5
10.0
0.00
0.25
0.50
0.75

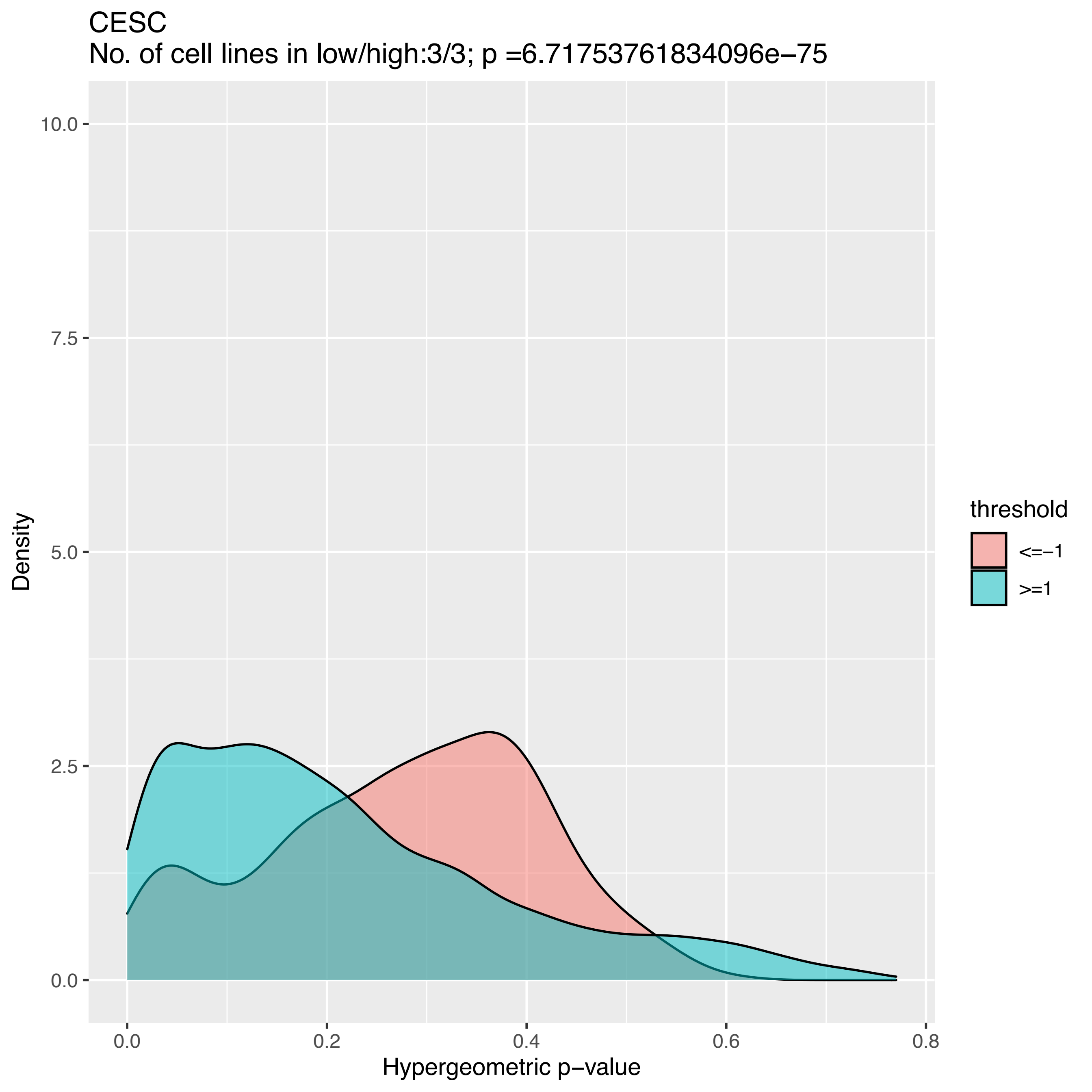

CESC
No. of cell lines in low/high:3/3; p =6.71753761834096e−75
Density
Hypergeometric p−value
10.0
7.5
5.0
2.5
0.0
0.0
0.2
0.4
0.6
0.8
threshold
<=−1
>=1

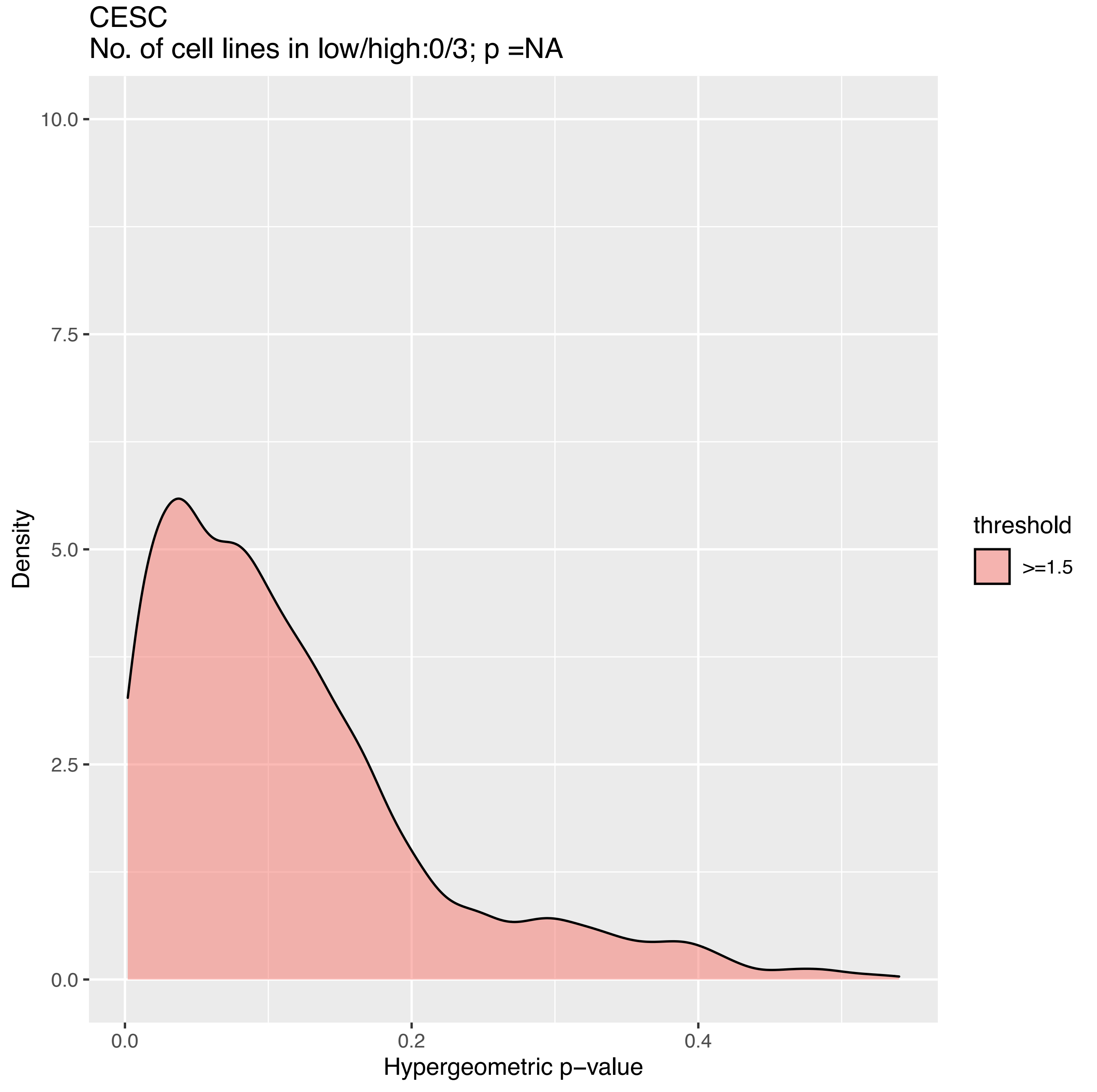
CESC
No. of cell lines in low/high:0/3; p =NA
10.0
7.5
5.0
2.5
0.0
Density
0.0
0.2
0.4
Hypergeometric p−value
threshold
>=1.5

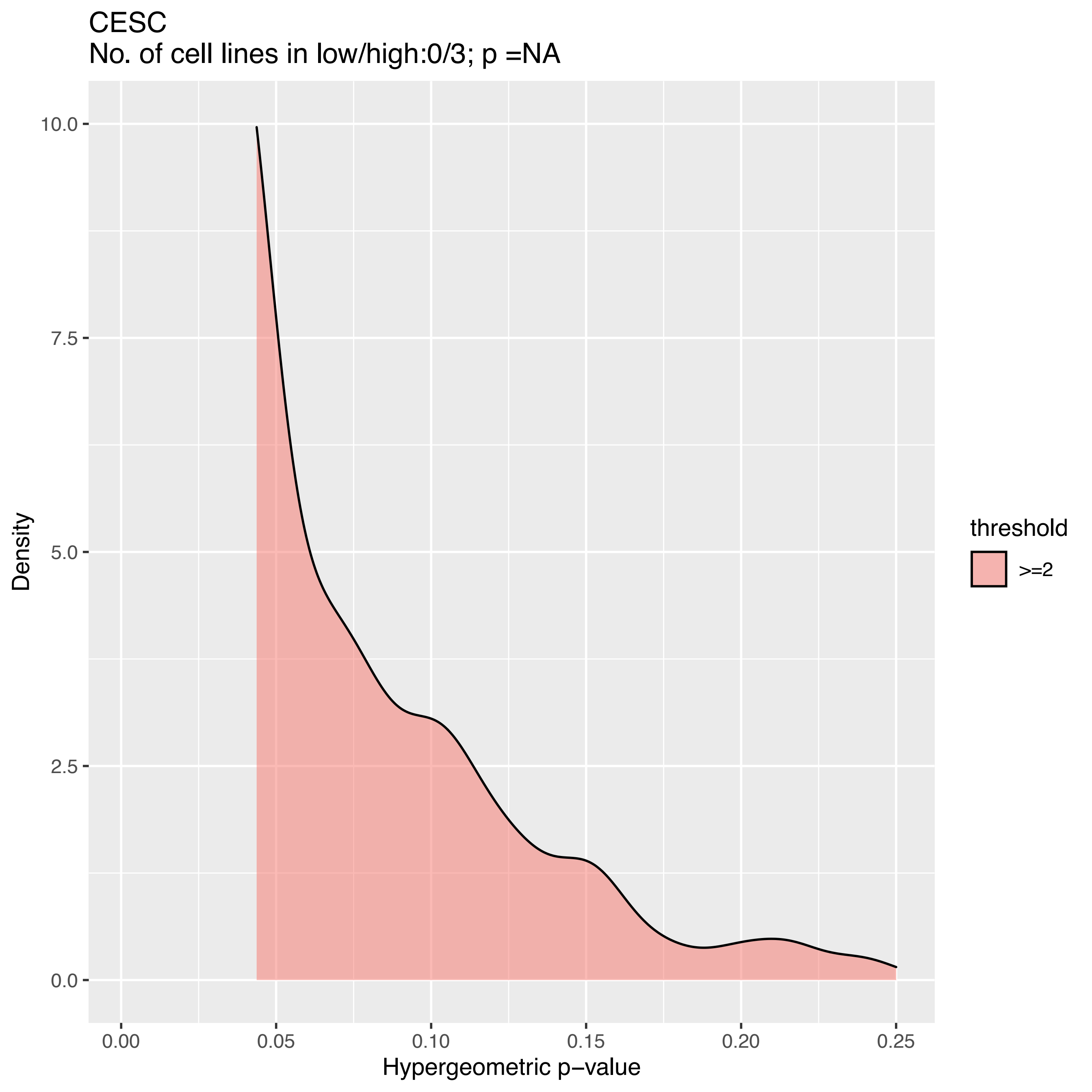
CESC
No. of cell lines in low/high:0/3; p =NA
10.0
7.5
5.0
2.5
0.0
Density
0.00
0.05
0.10
0.15
0.20
0.25
Hypergeometric p−value
threshold
>=2

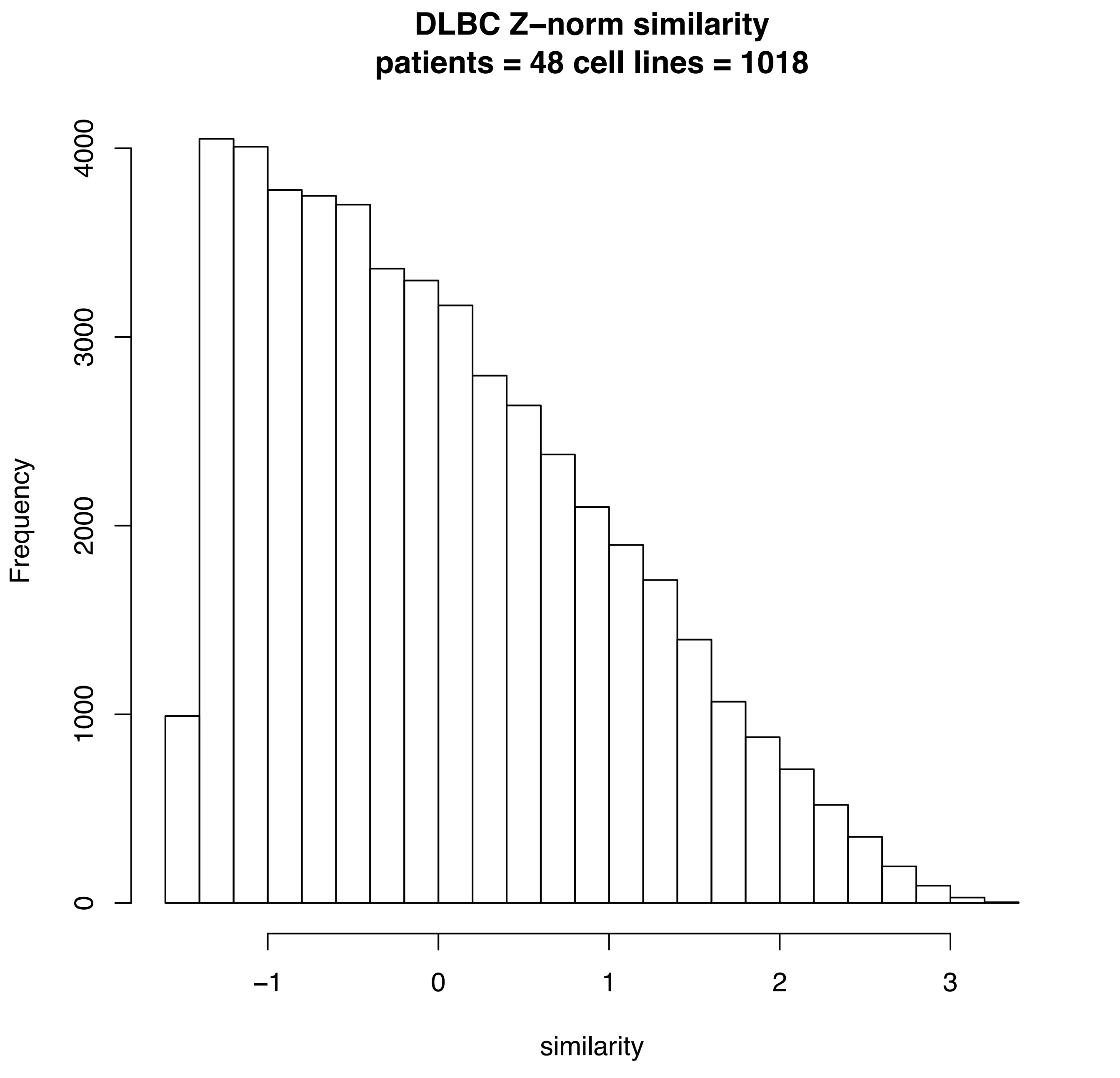
DLBC Z–norm similarity
patients = 48 cell lines = 1018
Frequency
0
1000
2000
3000
4000
−1
0
1
2
3
similarity

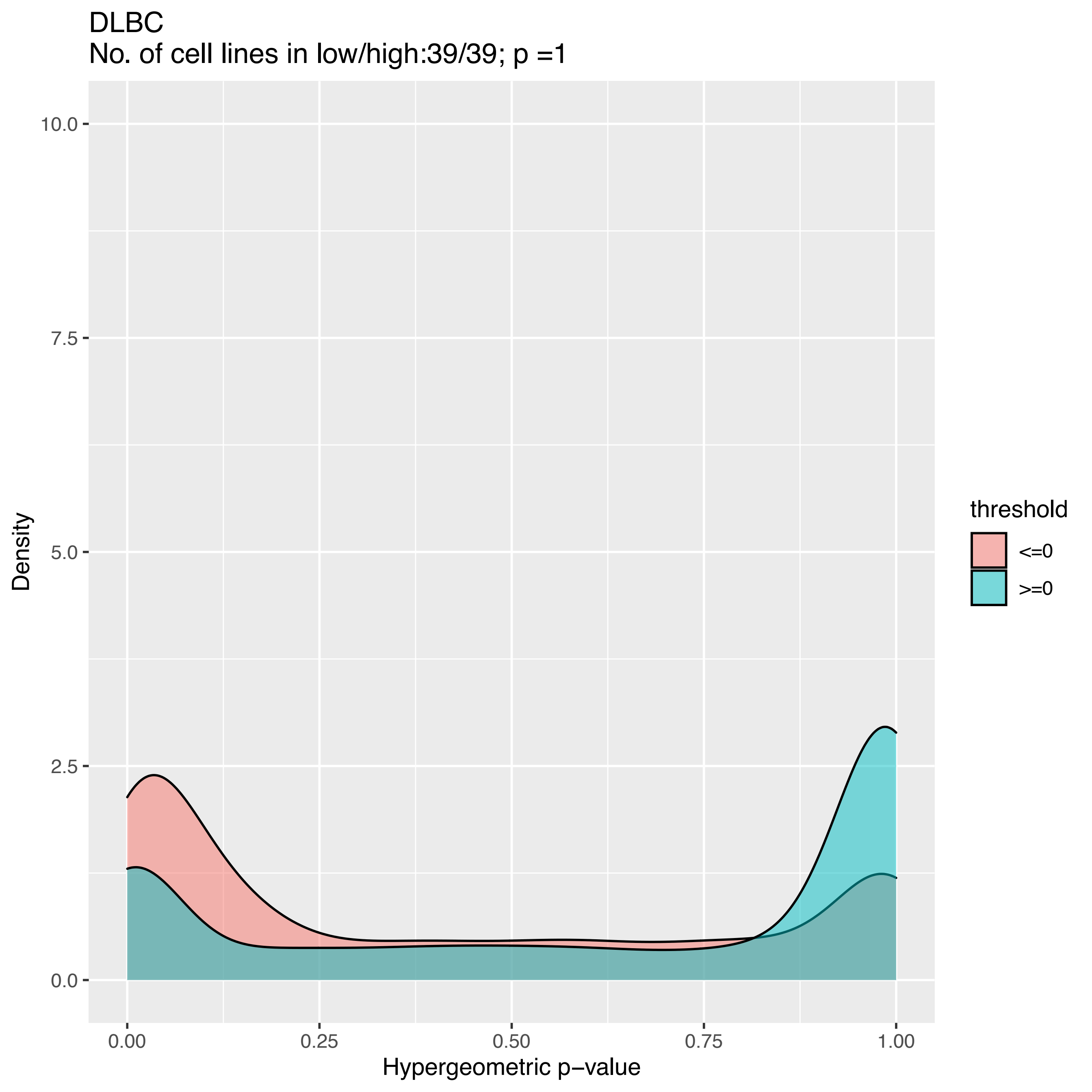
DLBC
No. of cell lines in low/high:39/39; p =1
10.0
7.5
5.0
2.5
0.0
Density
0.00
0.25
0.50
0.75
1.00
Hypergeometric p−value
threshold
<=0
>=0

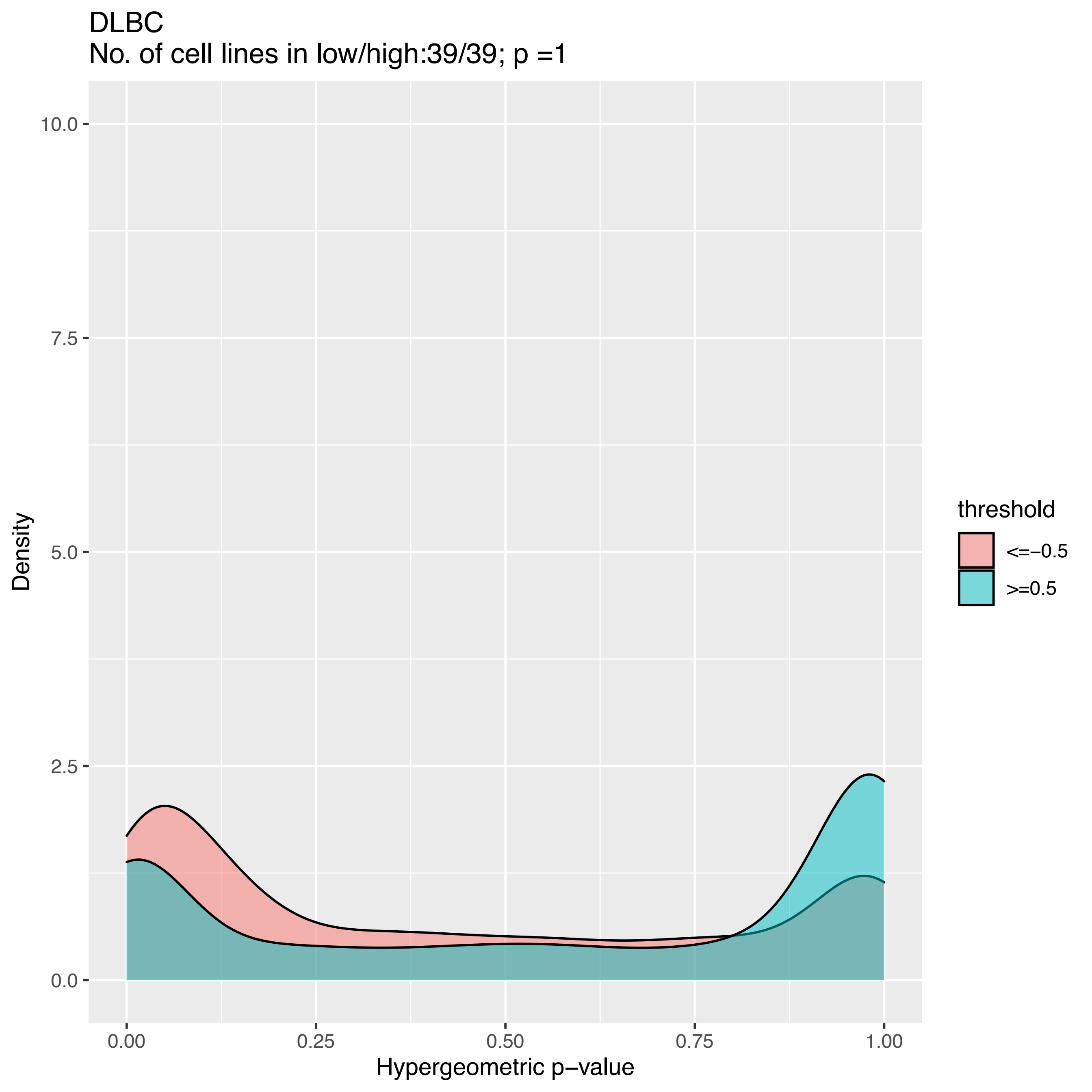
DLBC
No. of cell lines in low/high:39/39; p =1
Density
10.0
7.5
5.0
2.5
0.0
0.00
0.25
0.50
0.75
1.00
Hypergeometric p−value
threshold
<=−0.5
>=0.5

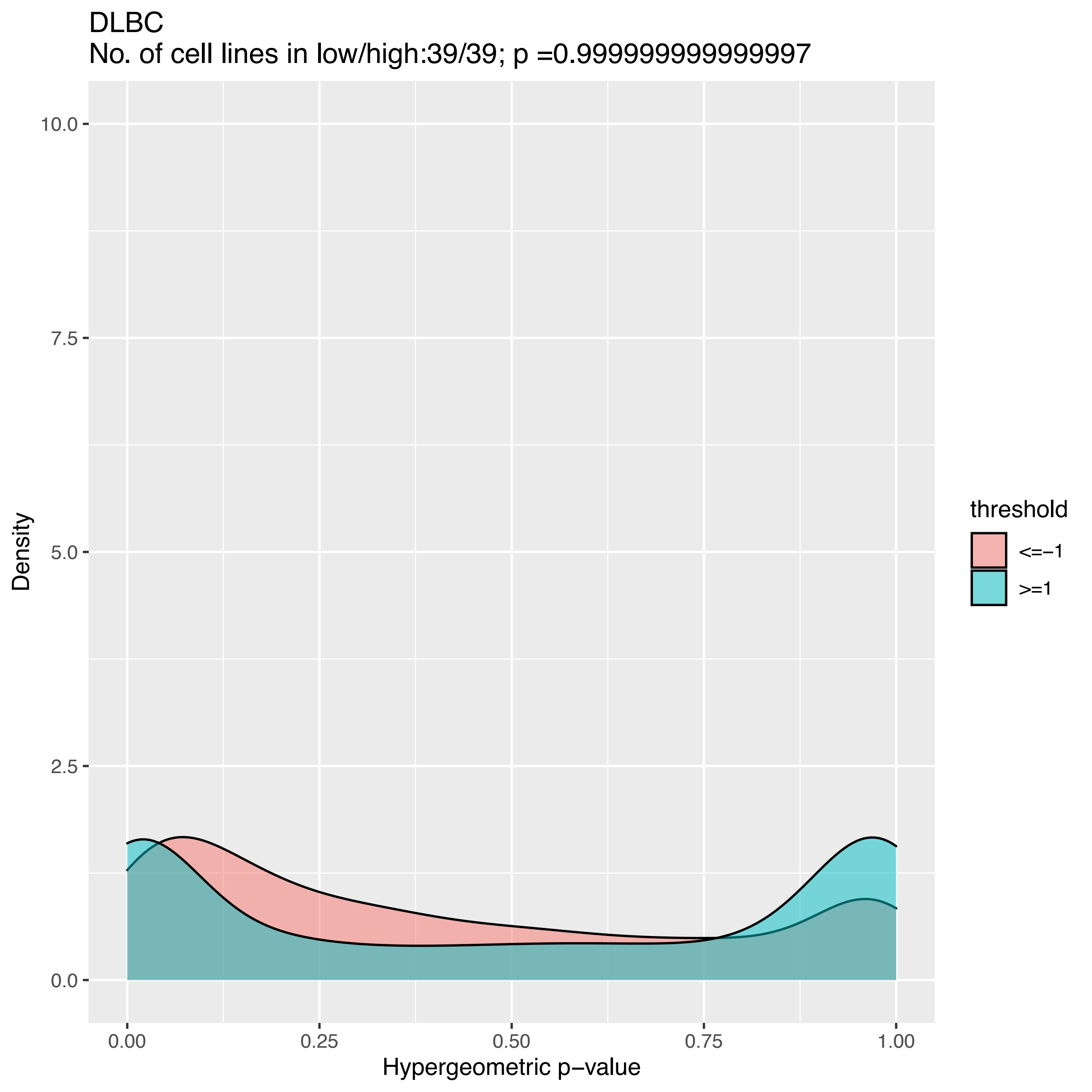
DLBC
No. of cell lines in low/high:39/39; p =0.9999999999999997
Density
10.0
7.5
5.0
2.5
0.0
0.00
0.25
0.50
0.75
1.00
Hypergeometric p−value
threshold
<=−1
>=1

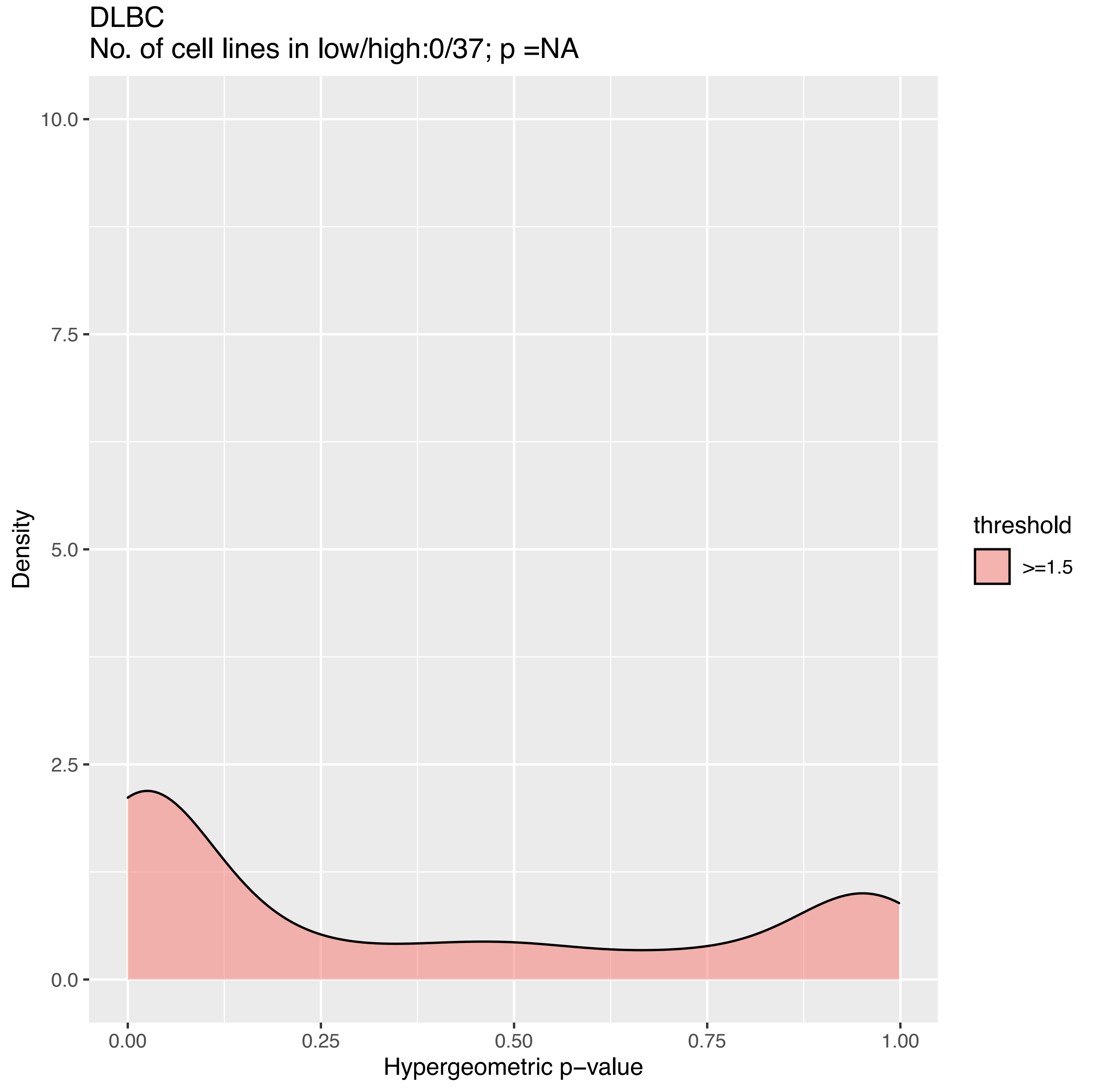
DLBC
No. of cell lines in low/high:0/37; p =NA
10.0
7.5
5.0
2.5
0.0
Density
0.00
0.25
0.50
0.75
1.00
Hypergeometric p−value
threshold
>=1.5

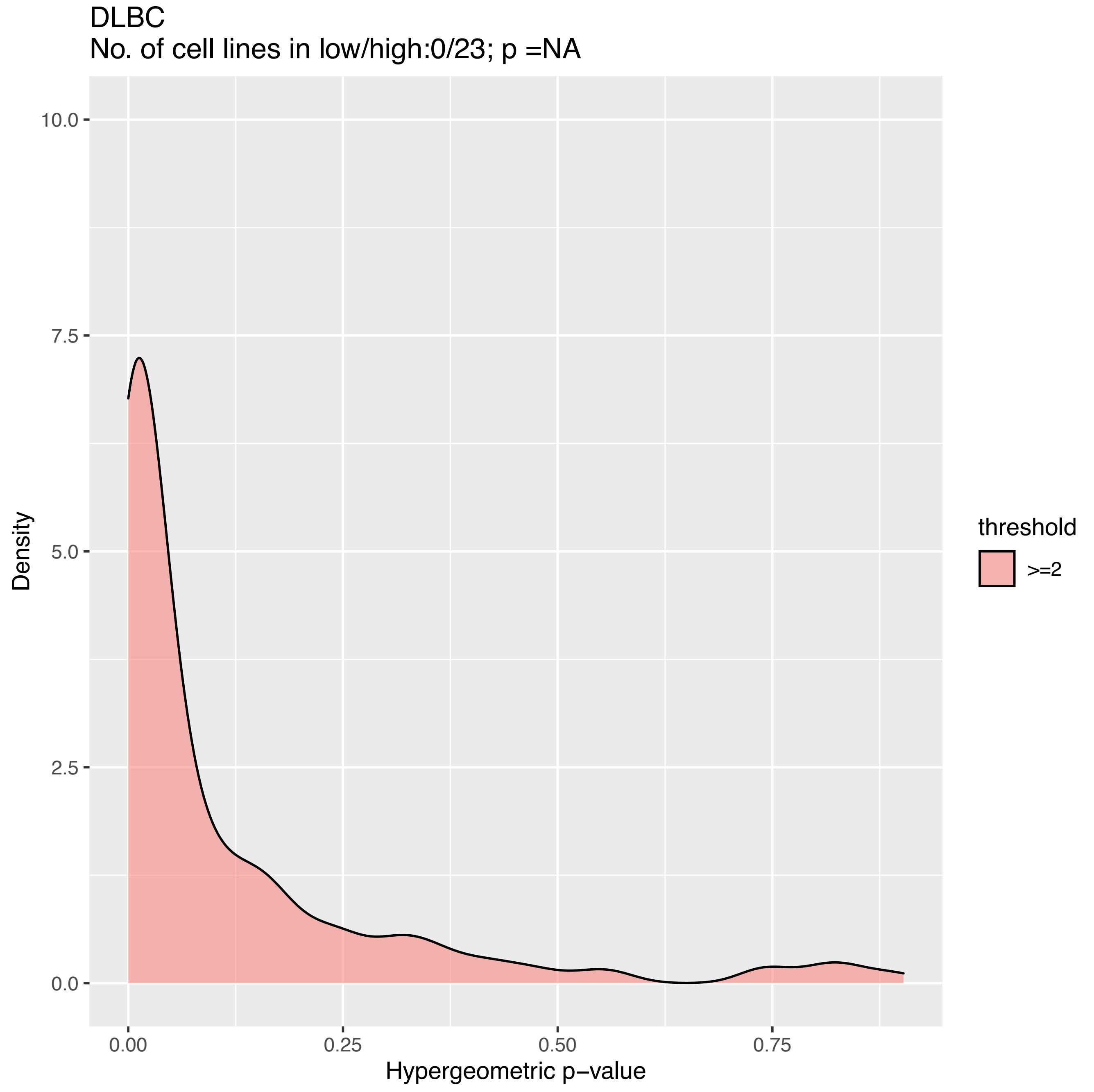

DLBC
No. of cell lines in low/high:0/23; p =NA
10.0
7.5
5.0
2.5
0.0
Density
0.00
0.25
0.50
0.75
Hypergeometric p−value
threshold
>=2

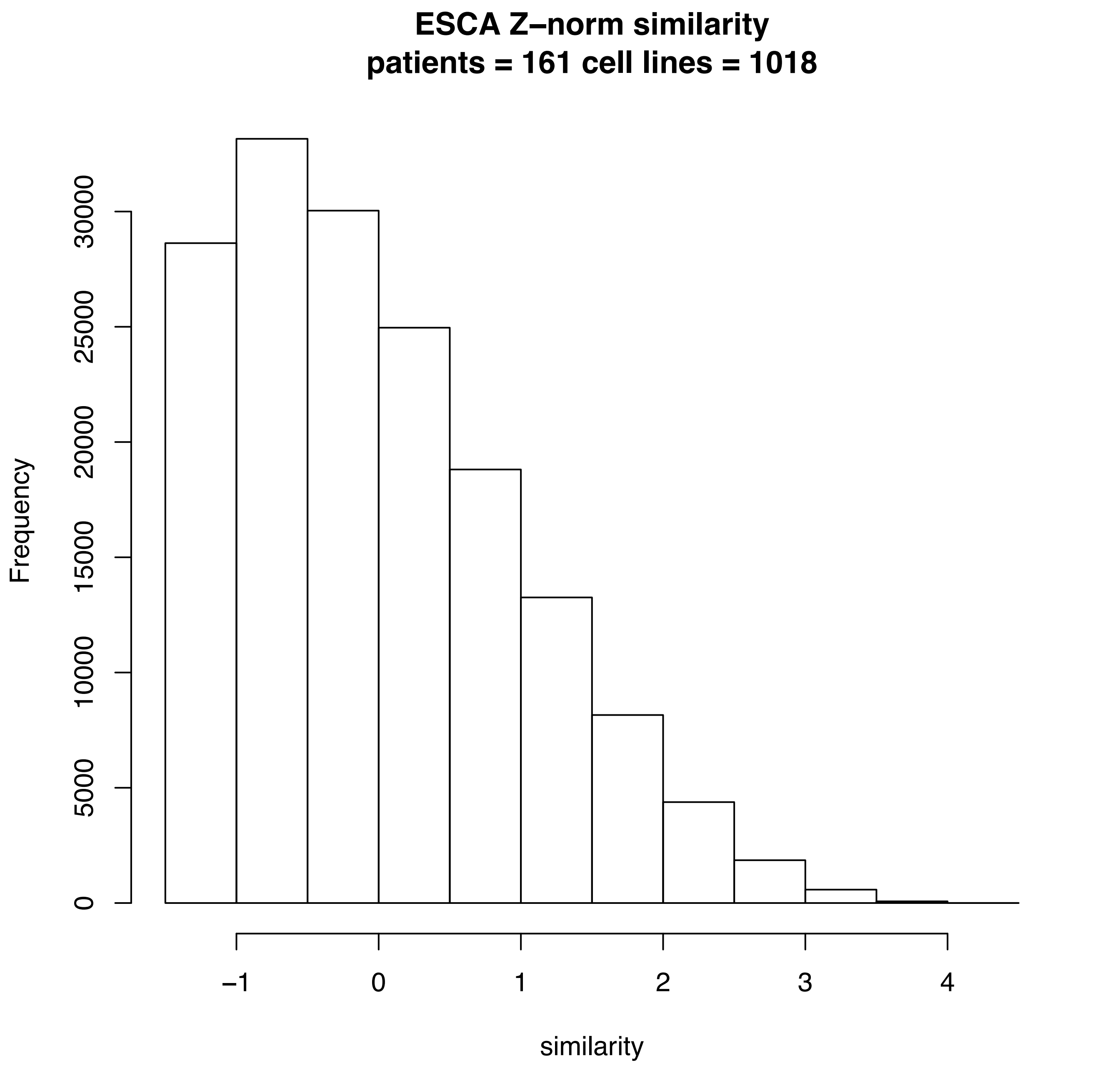
ESCA Z–norm similarity
patients = 161 cell lines = 1018
Frequency
0
5000
10000
15000
20000
25000
30000
–1
0
1
2
3
4
similarity

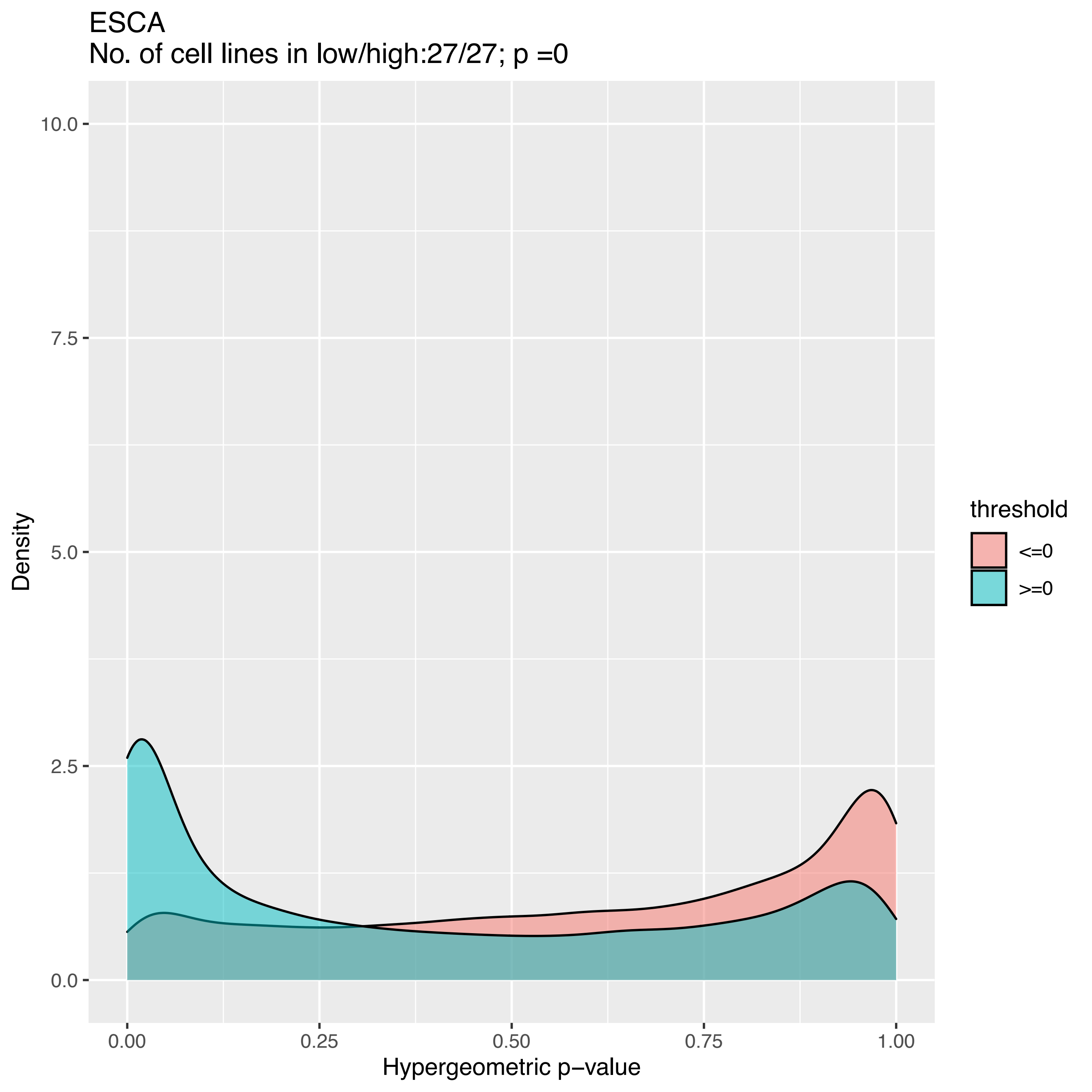
ESCA
No. of cell lines in low/high:27/27; p =0
Density
10.0
7.5
5.0
2.5
0.0
0.00
0.25
0.50
0.75
1.00
Hypergeometric p−value
threshold
<=0
>=0

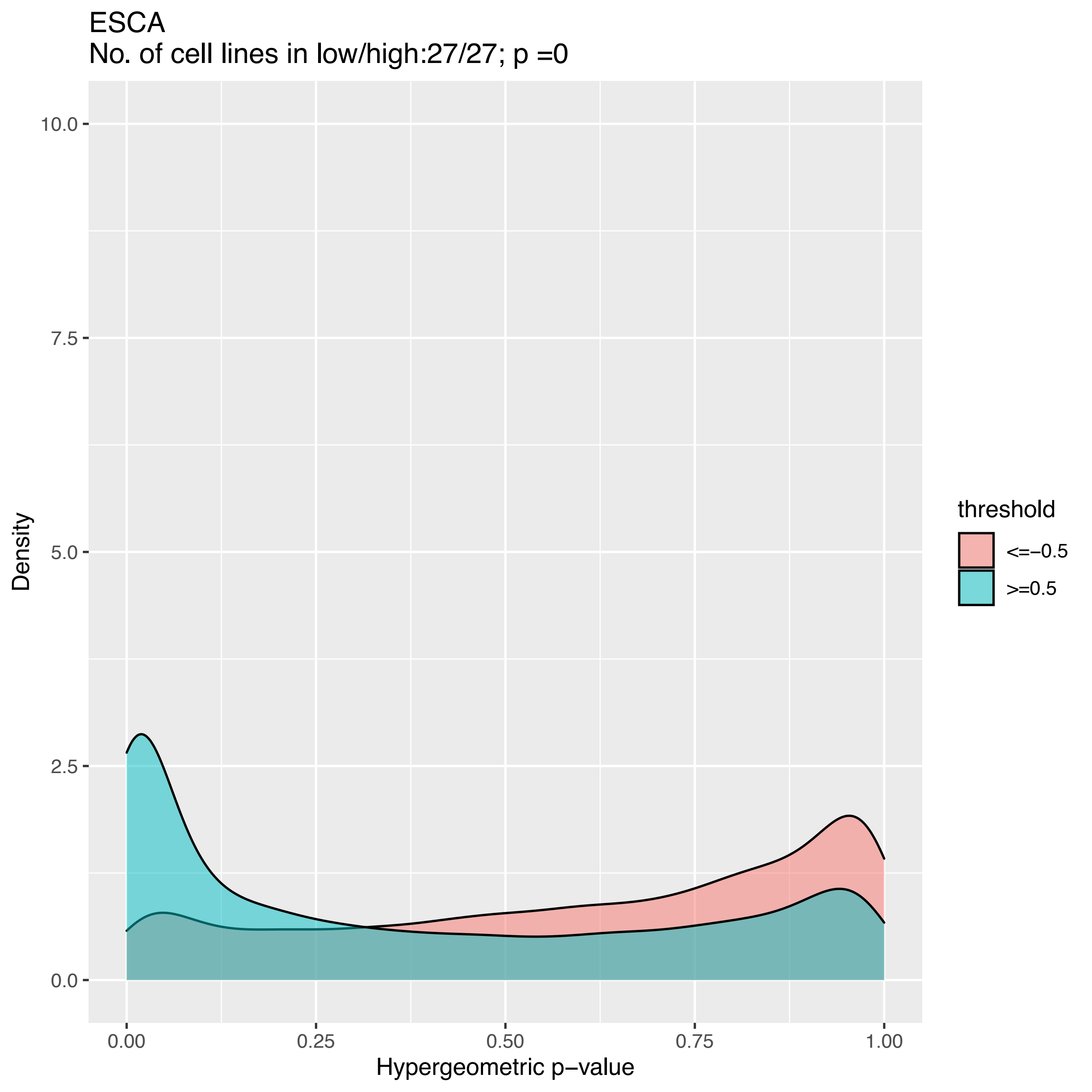
ESCA
No. of cell lines in low/high:27/27; p =0
Density
10.0
7.5
5.0
2.5
0.0
0.00
0.25
0.50
0.75
1.00
Hypergeometric p−value
threshold
<=−0.5
>=0.5

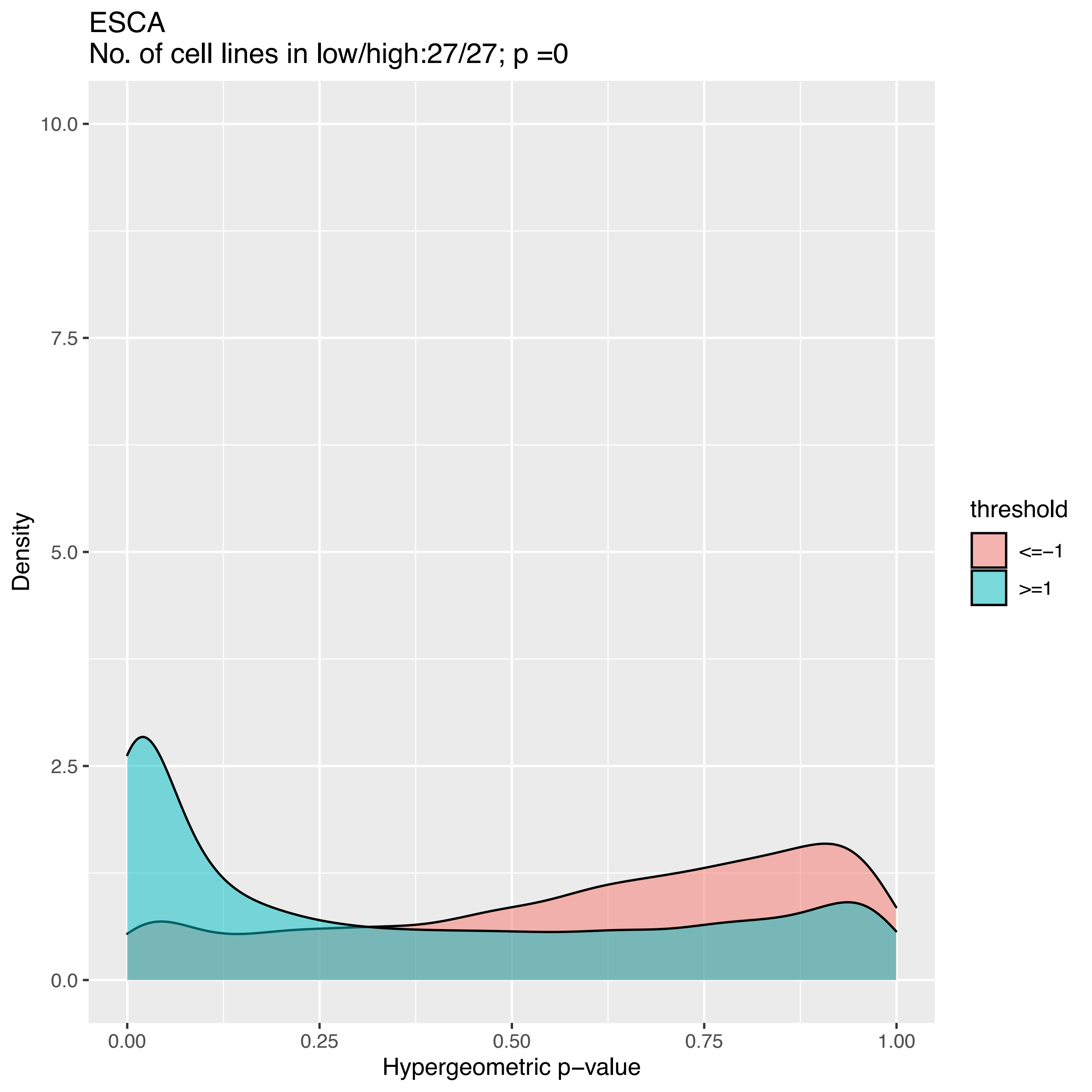
ESCA
No. of cell lines in low/high:27/27; p =0
10.0
7.5
5.0
2.5
0.0
Density
0.00
0.25
0.50
0.75
1.00
Hypergeometric p−value
threshold
<=−1
>=1

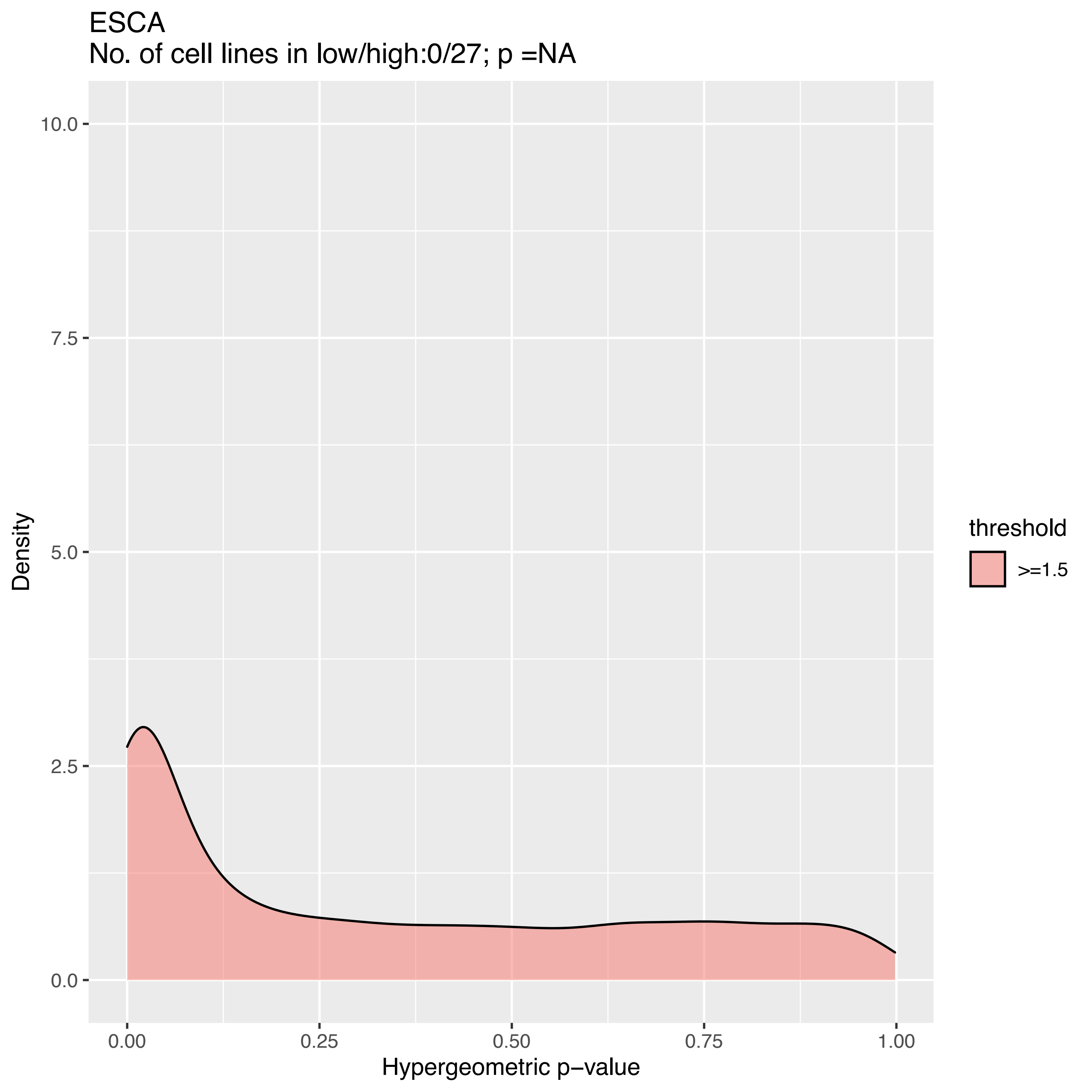

ESCA
No. of cell lines in low/high:0/27; p =NA
Density
10.0
7.5
5.0
2.5
0.0
0.00
0.25
0.50
0.75
1.00
Hypergeometric p−value
threshold
>=1.5

ESCA
No. of cell lines in low/high:0/27; p =NA

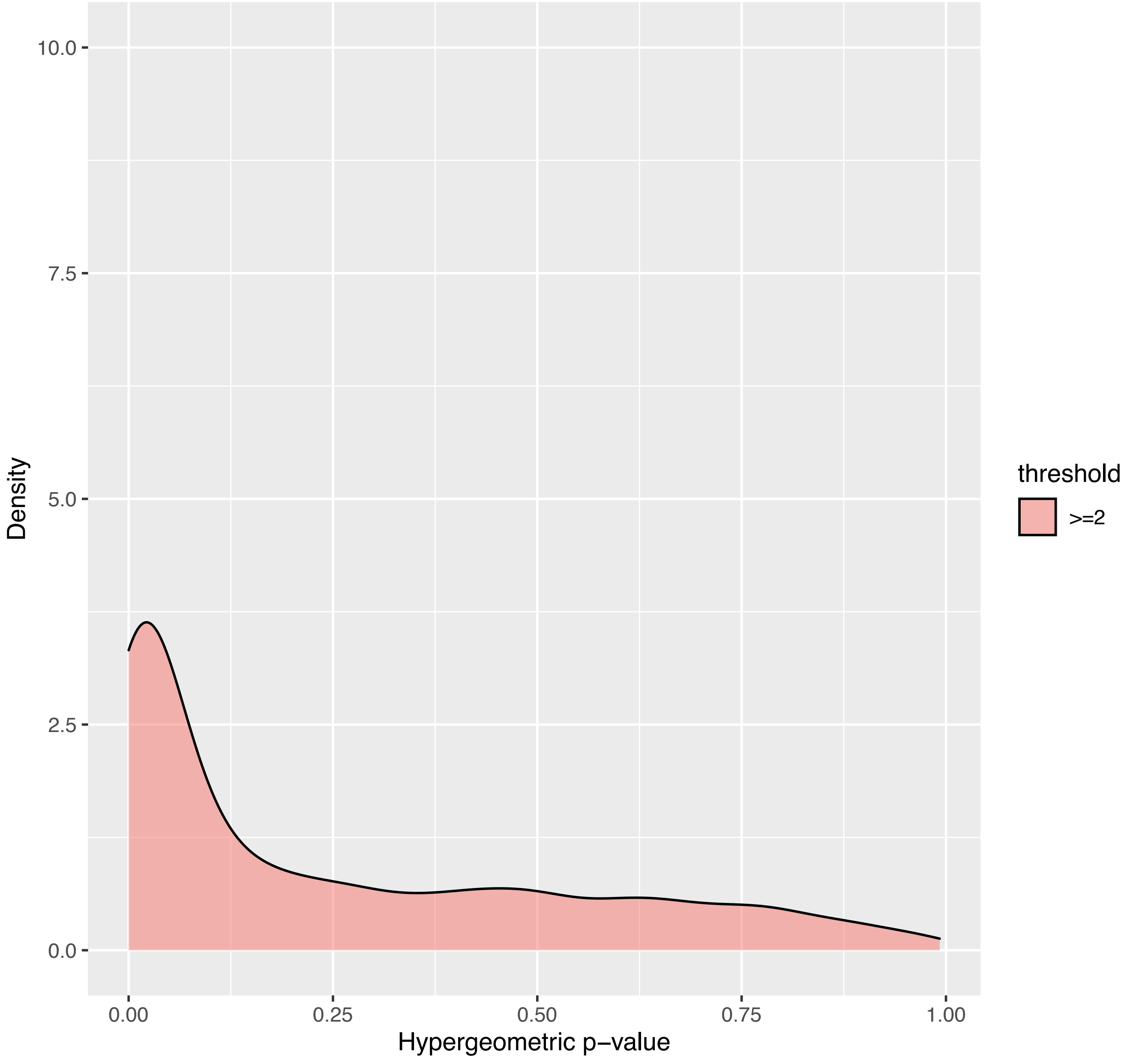

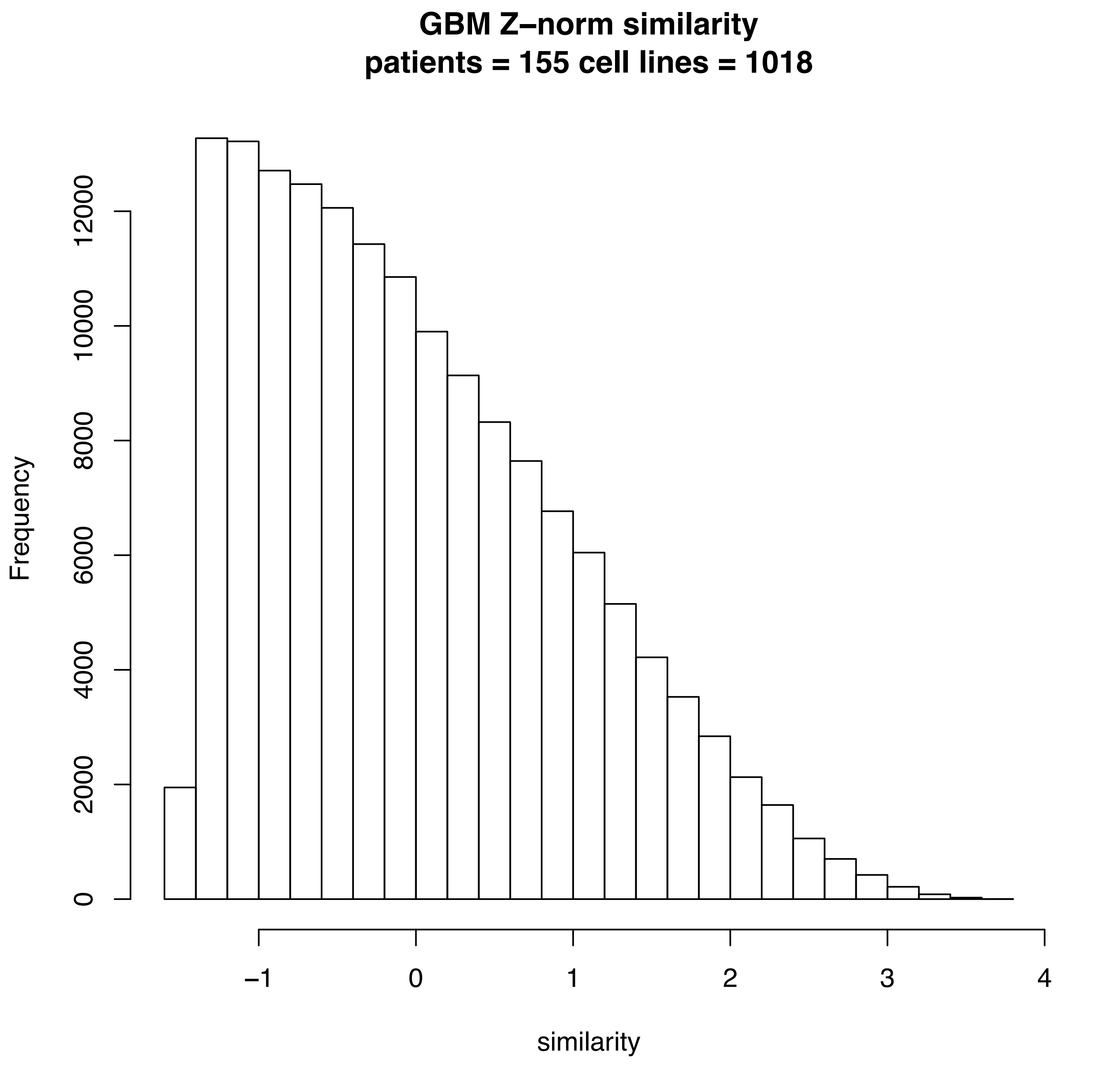
GBM Z–norm similarity
patients = 155 cell lines = 1018
Frequency
0
2000
4000
6000
8000
10000
12000
−1
0
1
2
3
4
similarity

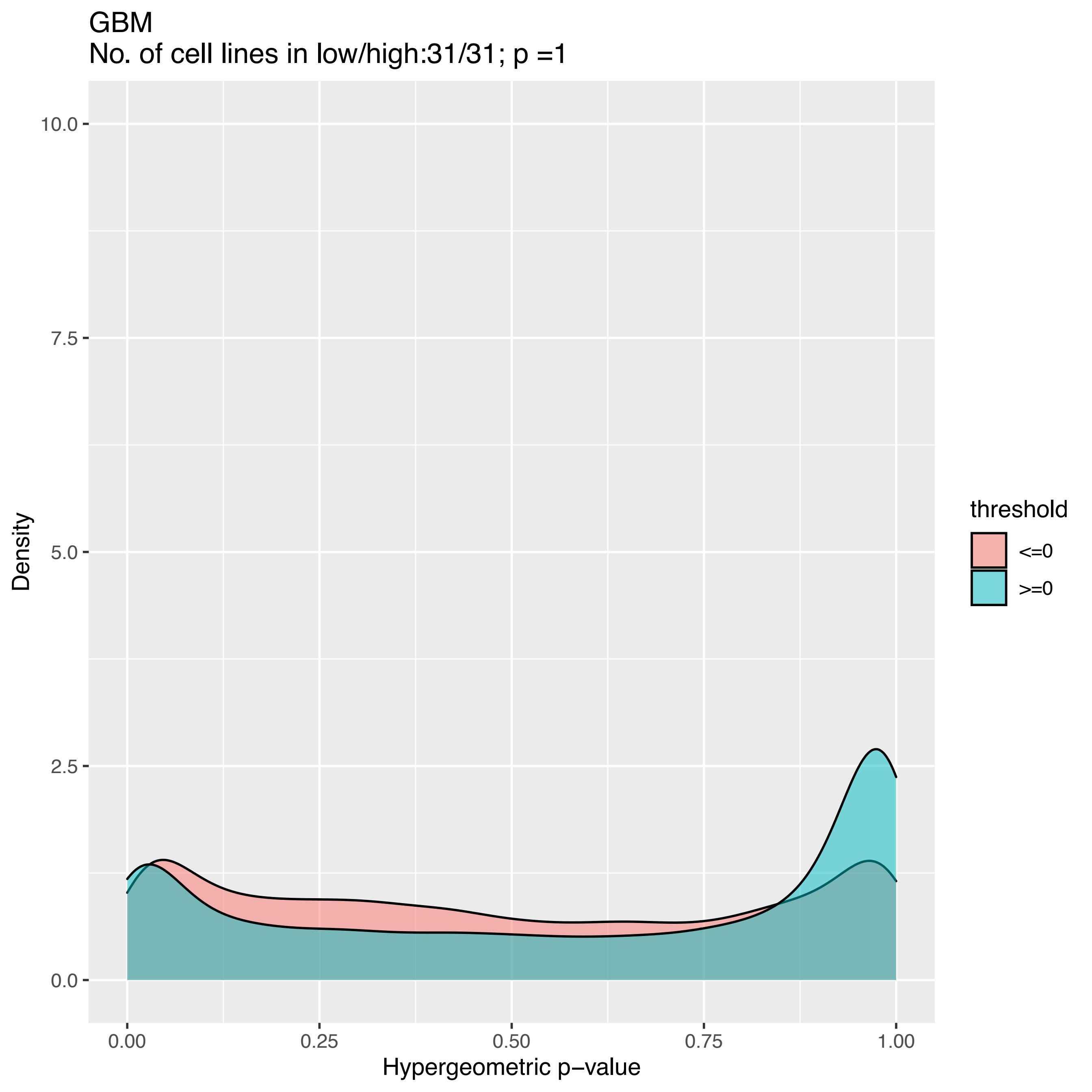

GBM
No. of cell lines in low/high:31/31; p =1
10.0
7.5
5.0
2.5
0.0
Density
0.00
0.25
0.50
0.75
1.00
Hypergeometric p−value
threshold
<=0
>=0

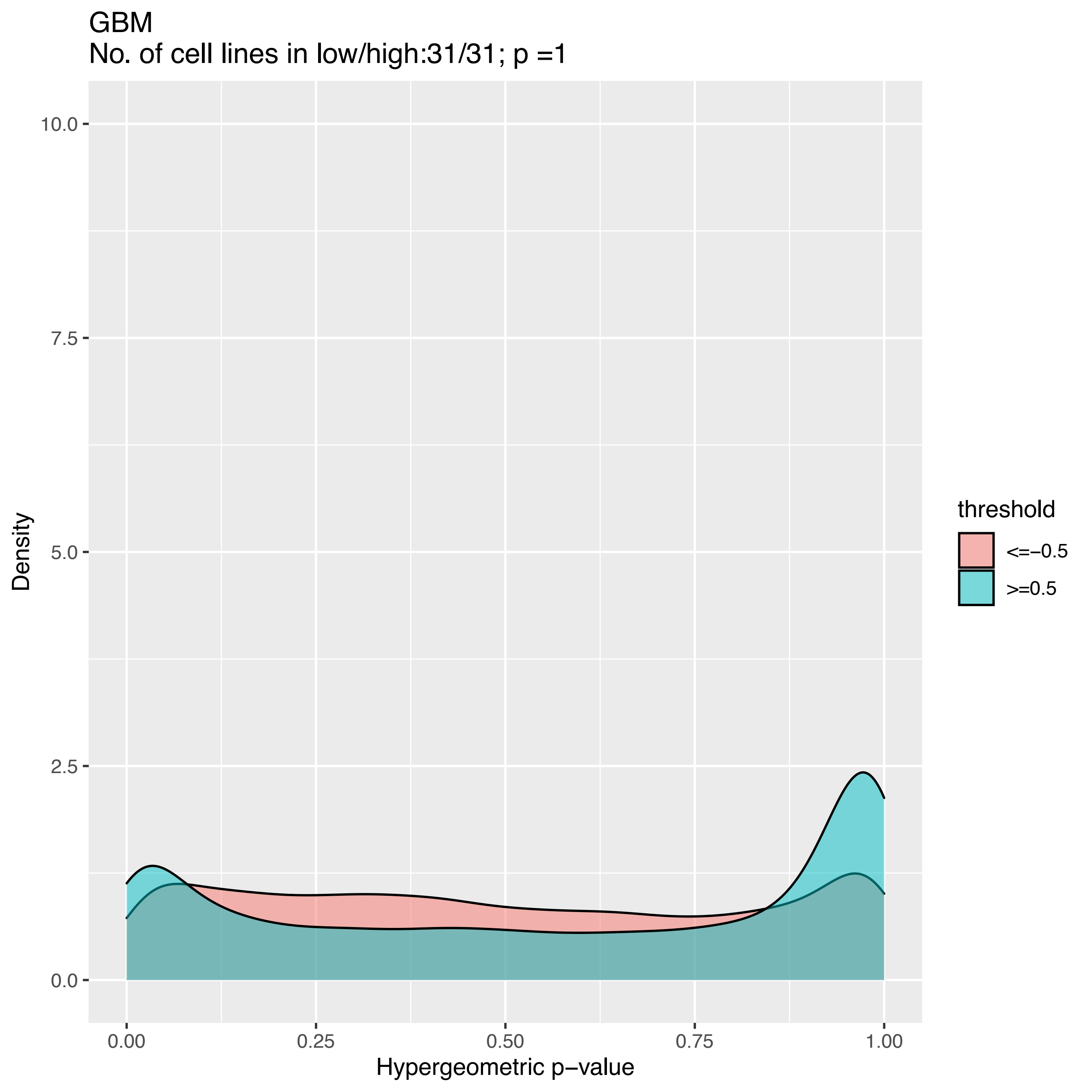
GBM
No. of cell lines in low/high:31/31; p =1
Density
10.0
7.5
5.0
2.5
0.0
0.00
0.25
0.50
0.75
1.00
Hypergeometric p−value
threshold
<=−0.5
>=0.5

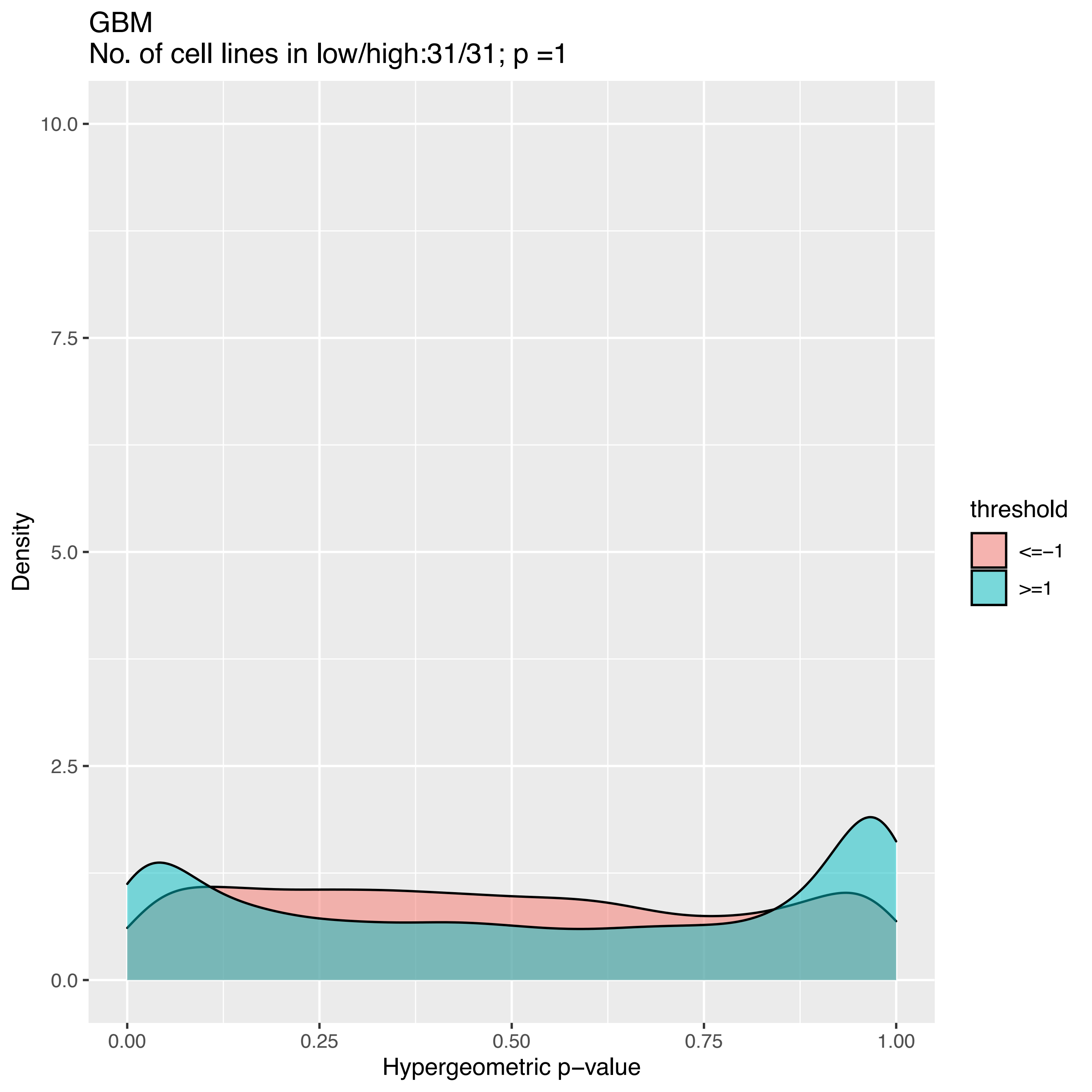
GBM
No. of cell lines in low/high:31/31; p =1
10.0
7.5
5.0
2.5
0.0
Density
0.00
0.25
0.50
0.75
1.00
Hypergeometric p−value
threshold
<=−1
>=1

GBM
No. of cell lines in low/high:0/31; p =NA

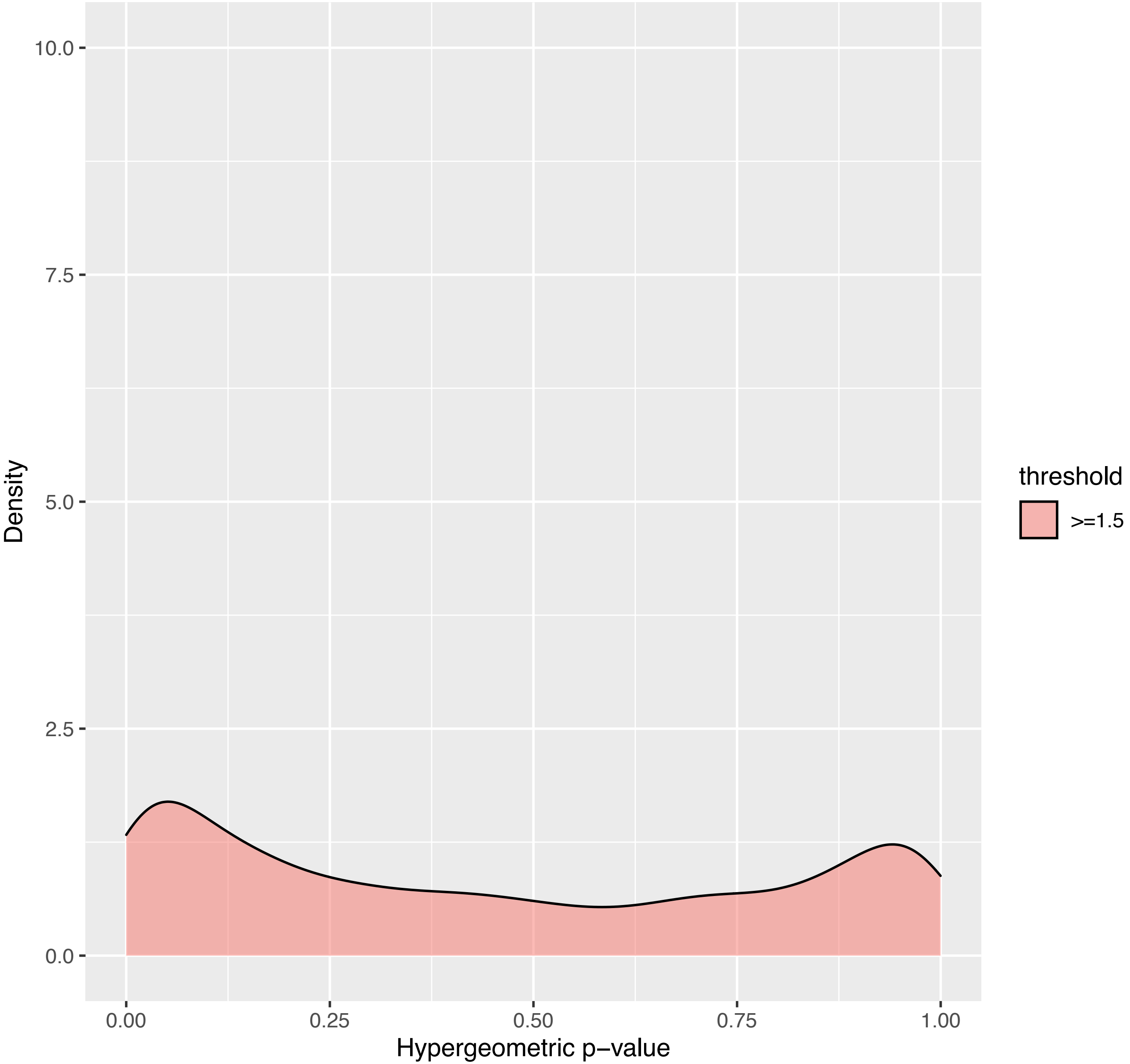

GBM
No. of cell lines in low/high:0/30; p =NA

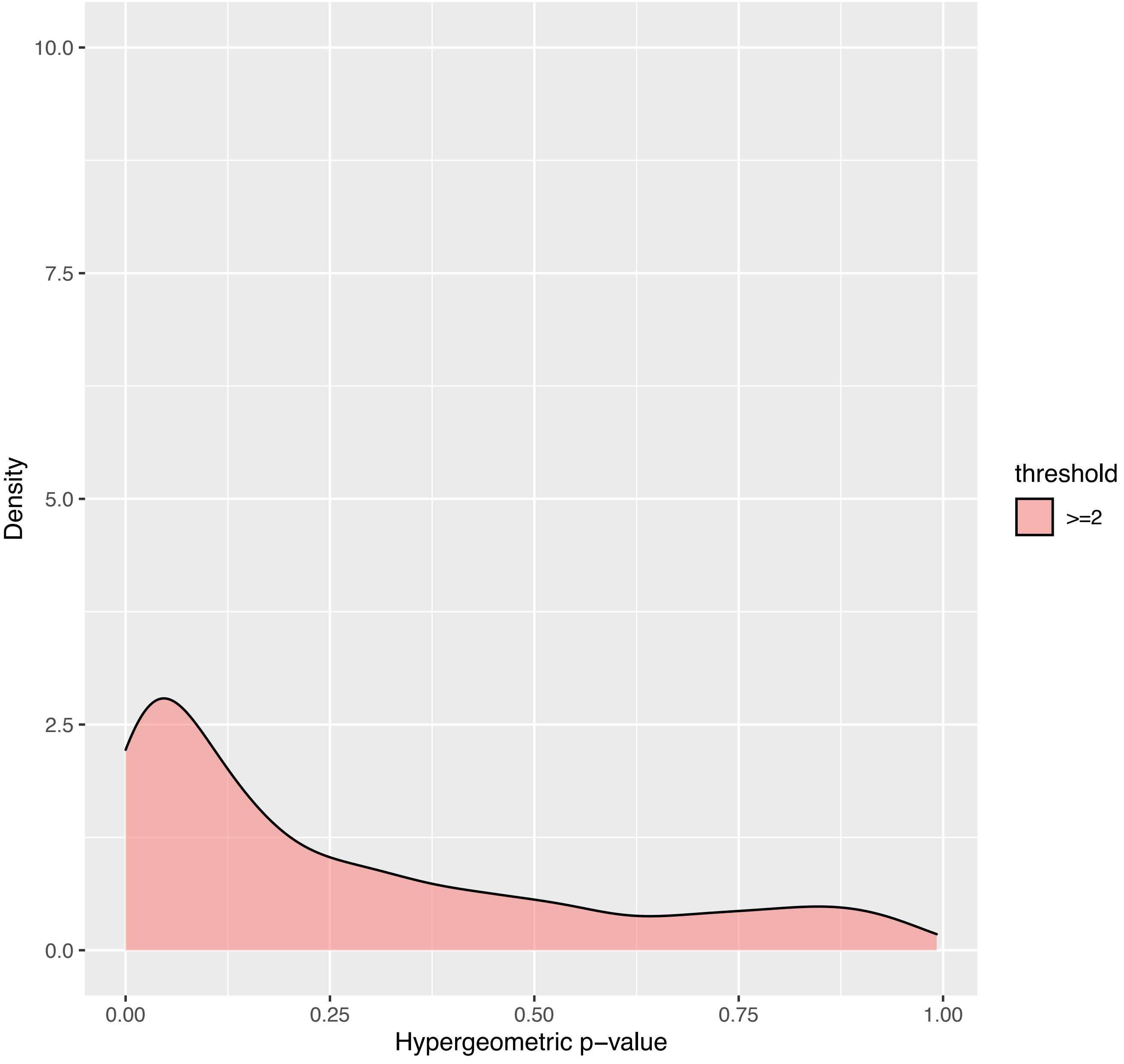

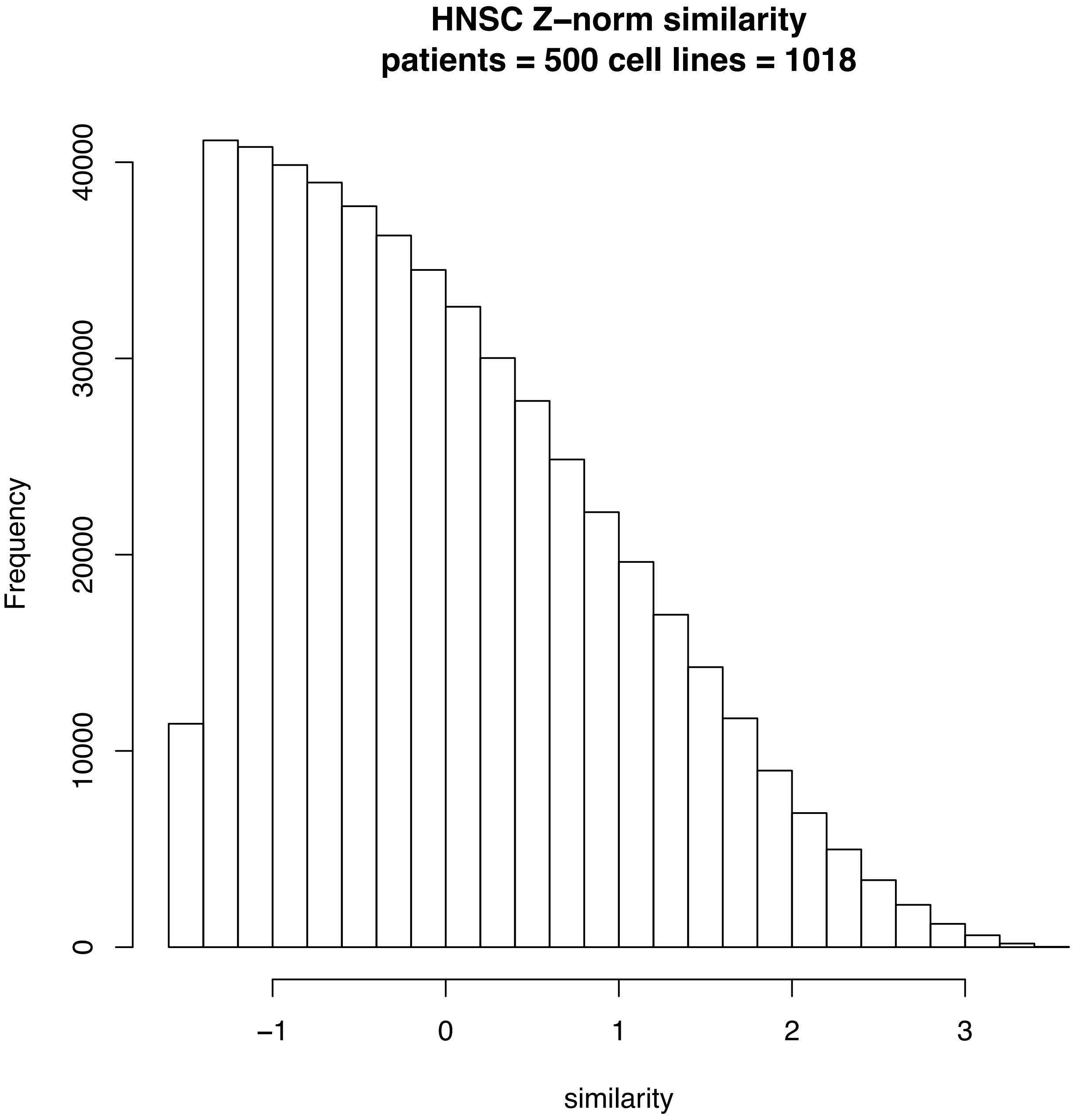

HNSC Z–norm similarity
patients = 500 cell lines = 1018
Frequency
0
10000
20000
30000
40000
−1
0
1
2
3
similarity

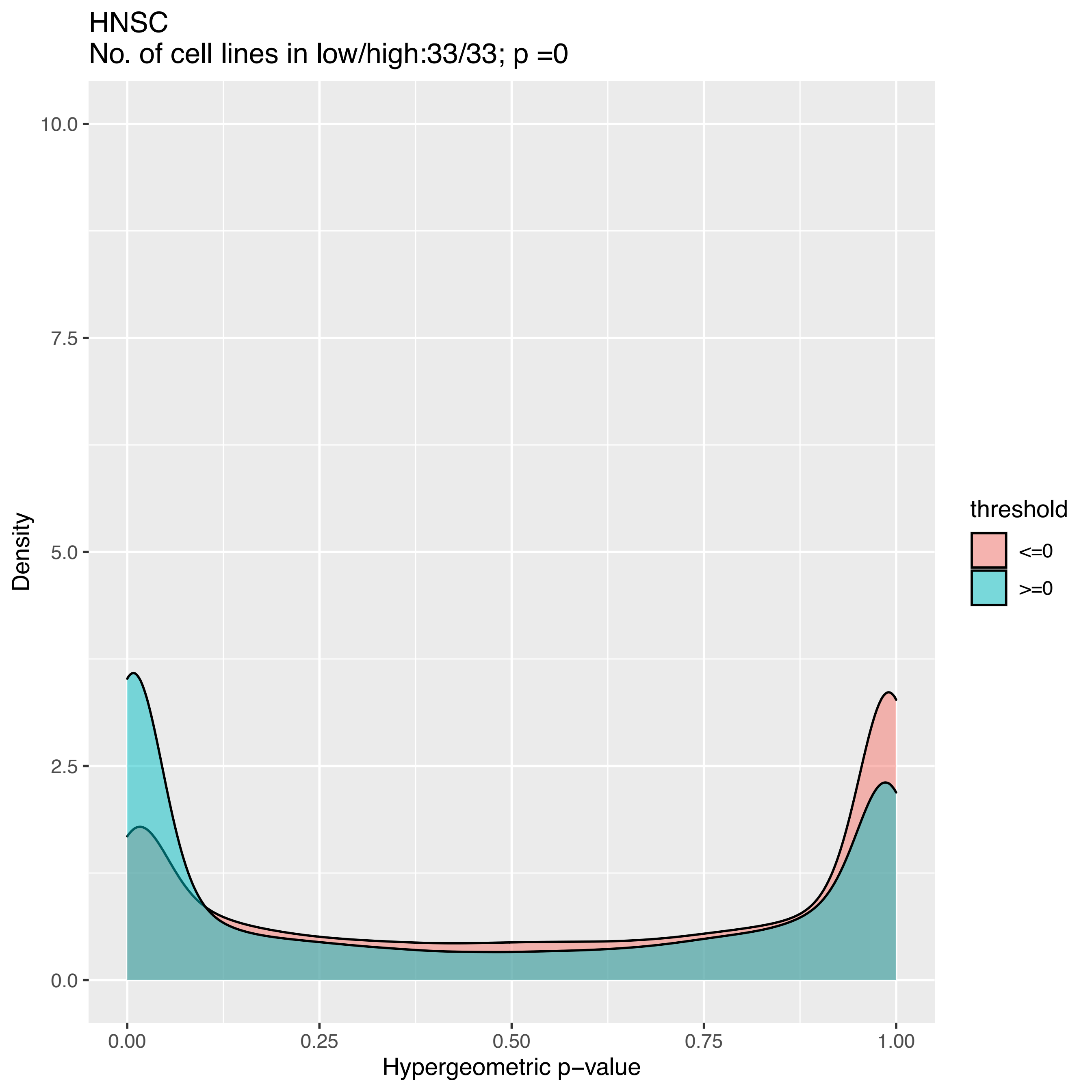

HNSC
No. of cell lines in low/high:33/33; p =0
Density
10.0
7.5
5.0
2.5
0.0
0.00
0.25
0.50
0.75
1.00
Hypergeometric p−value
threshold
<=0
>=0

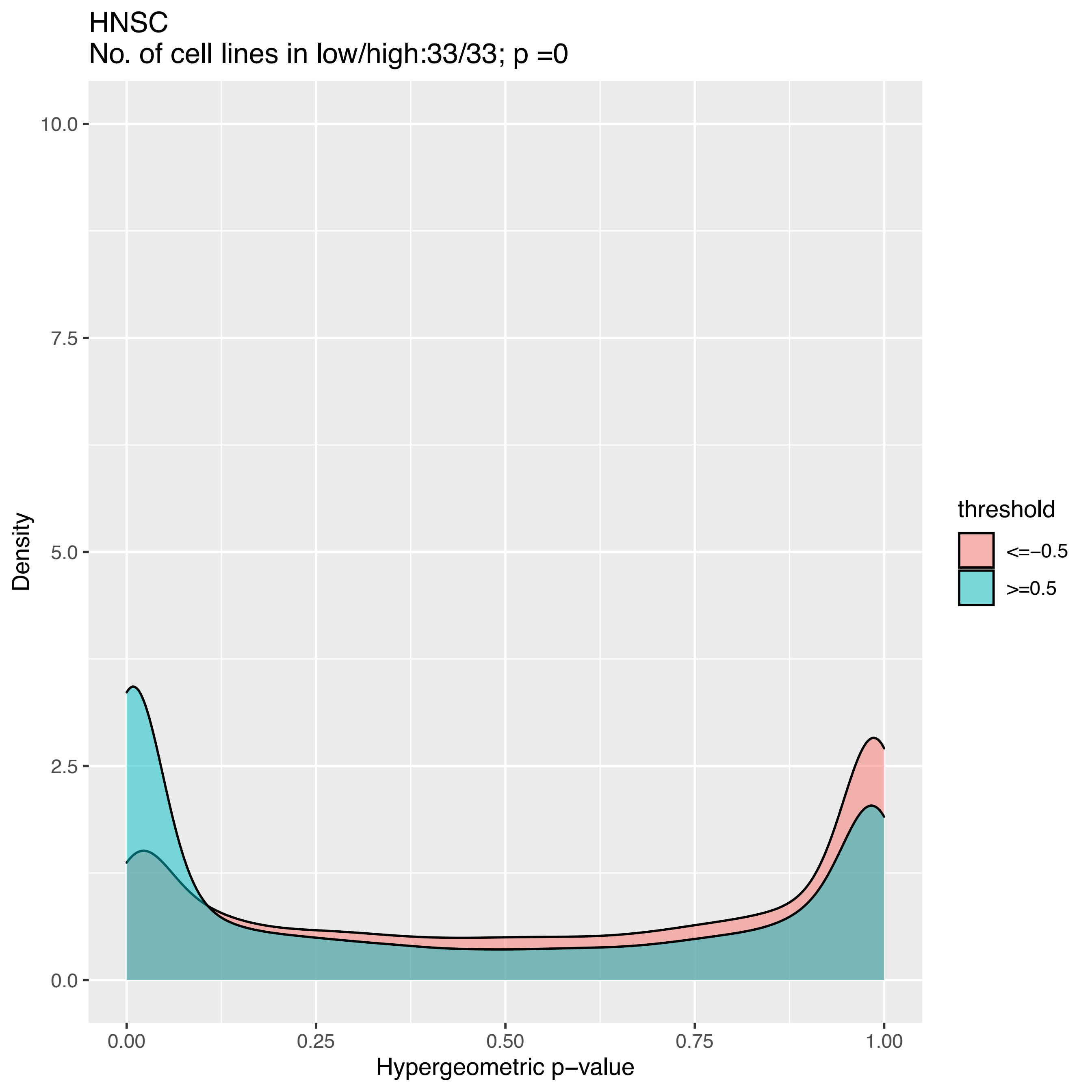
HNSC
No. of cell lines in low/high:33/33; p =0
Density
10.0
7.5
5.0
2.5
0.0
0.00
0.25
0.50
0.75
1.00
Hypergeometric p−value
threshold
<=−0.5
>=0.5

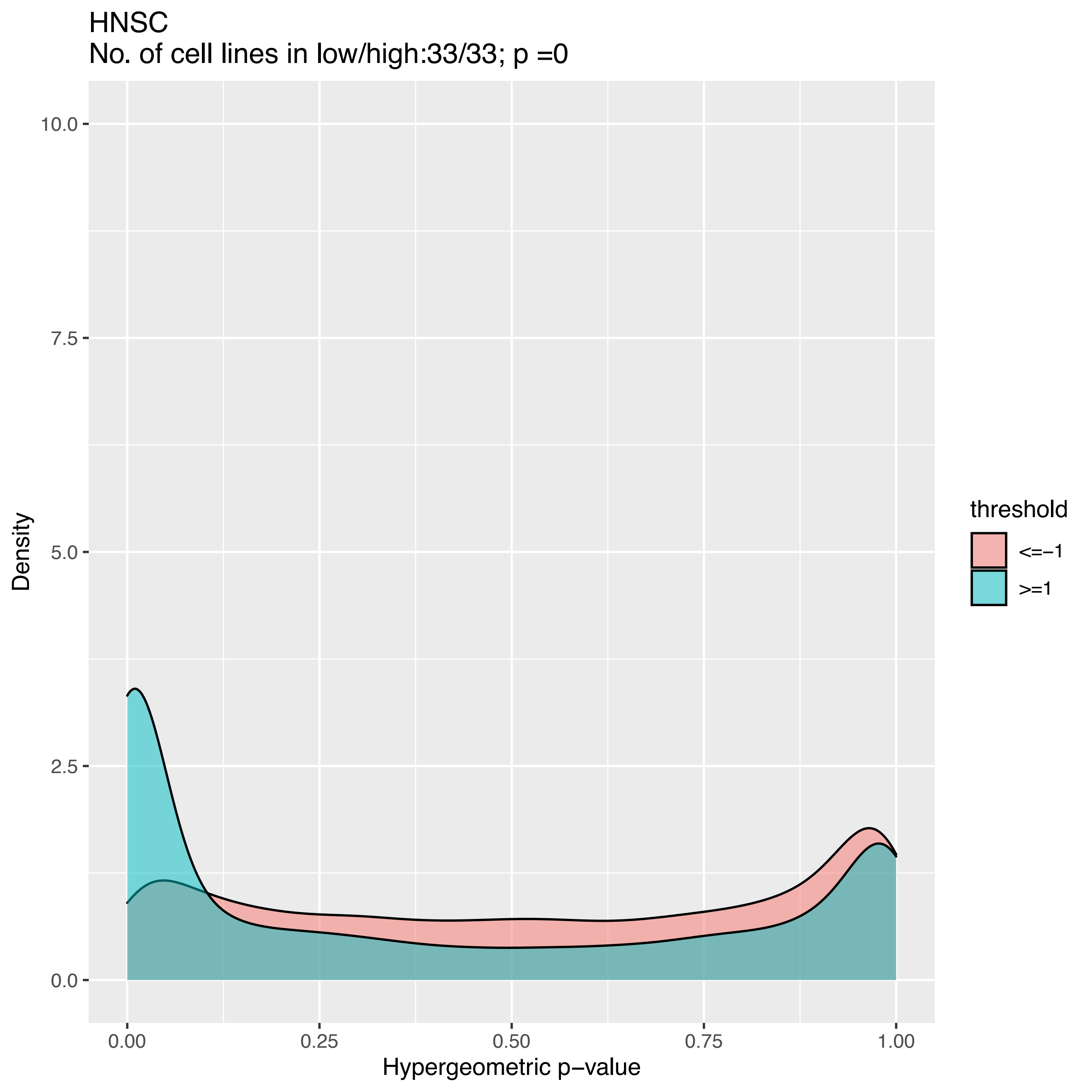
HNSC
No. of cell lines in low/high:33/33; p =0
10.0
7.5
5.0
2.5
0.0
Density
0.00
0.25
0.50
0.75
1.00
Hypergeometric p−value
threshold
<=−1
>=1

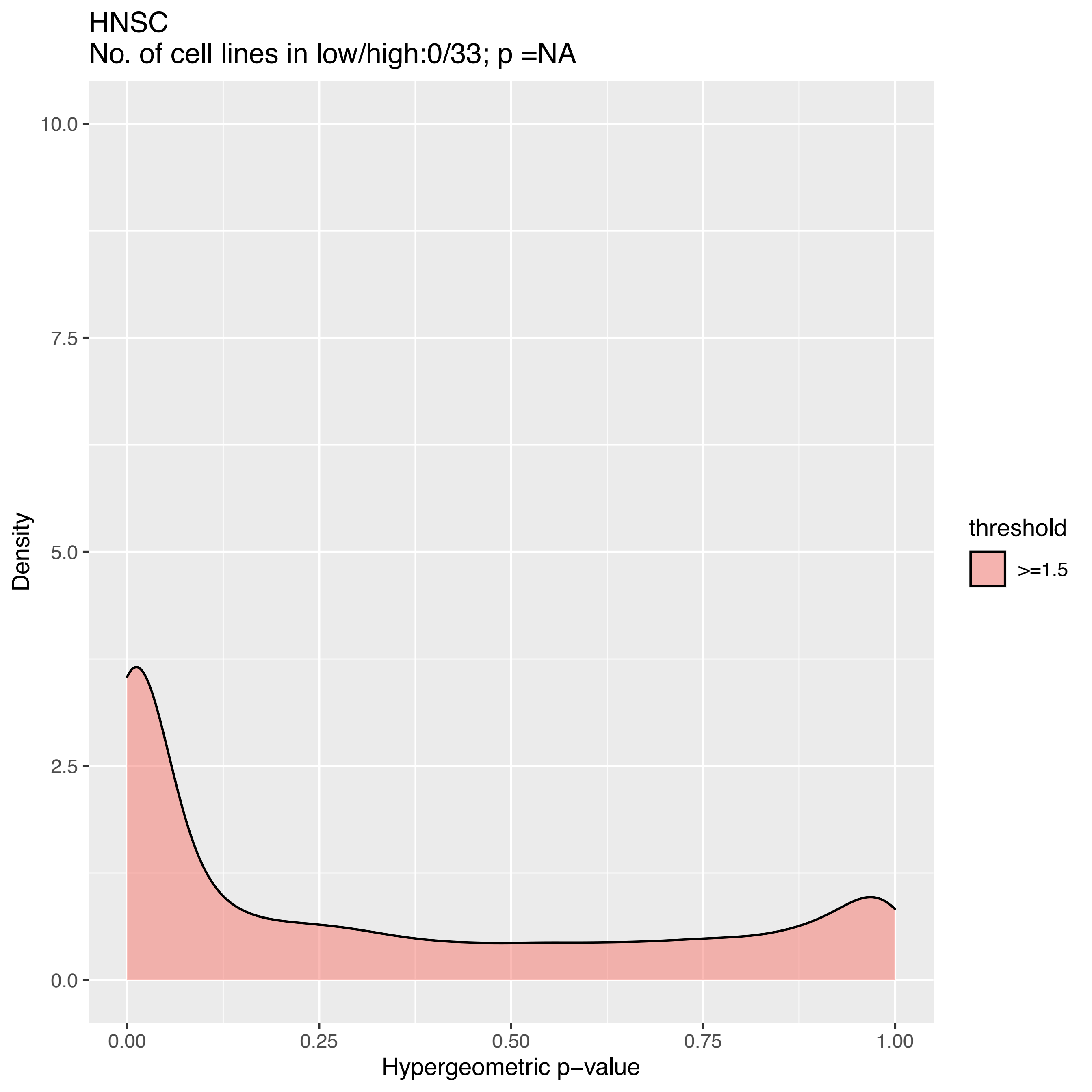

HNSC
No. of cell lines in low/high:0/33; p =NA
Density
10.0
7.5
5.0
2.5
0.0
0.00
0.25
0.50
0.75
1.00
Hypergeometric p−value
threshold
>=1.5

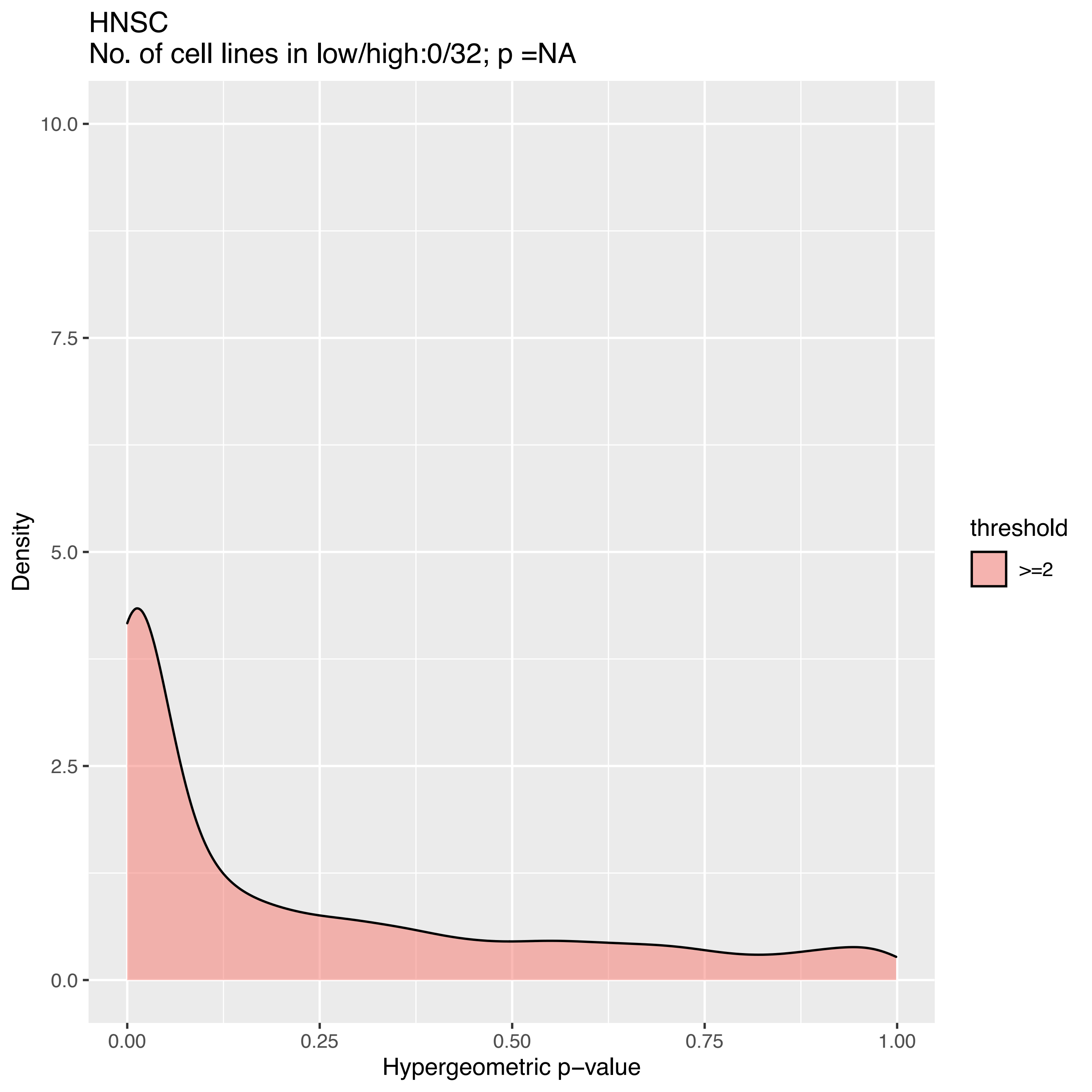
HNSC
No. of cell lines in low/high:0/32; p =NA
10.0
7.5
5.0
2.5
0.0
Density
0.00
0.25
0.50
0.75
1.00
Hypergeometric p−value
threshold
>=2

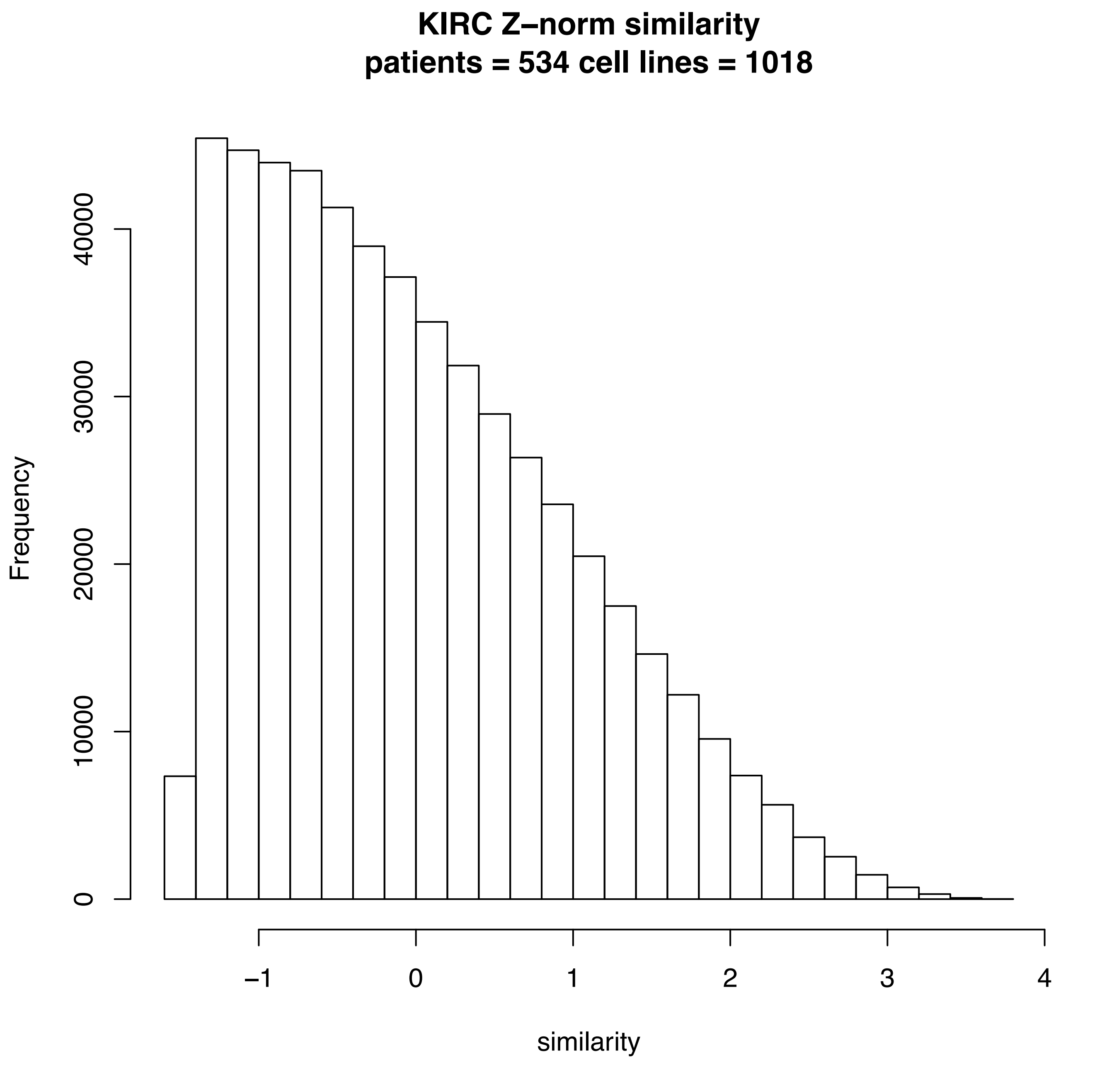
KIRC Z−norm similarity
patients = 534 cell lines = 1018
Frequency
0
10000
20000
30000
40000
−1
0
1
2
3
4
similarity

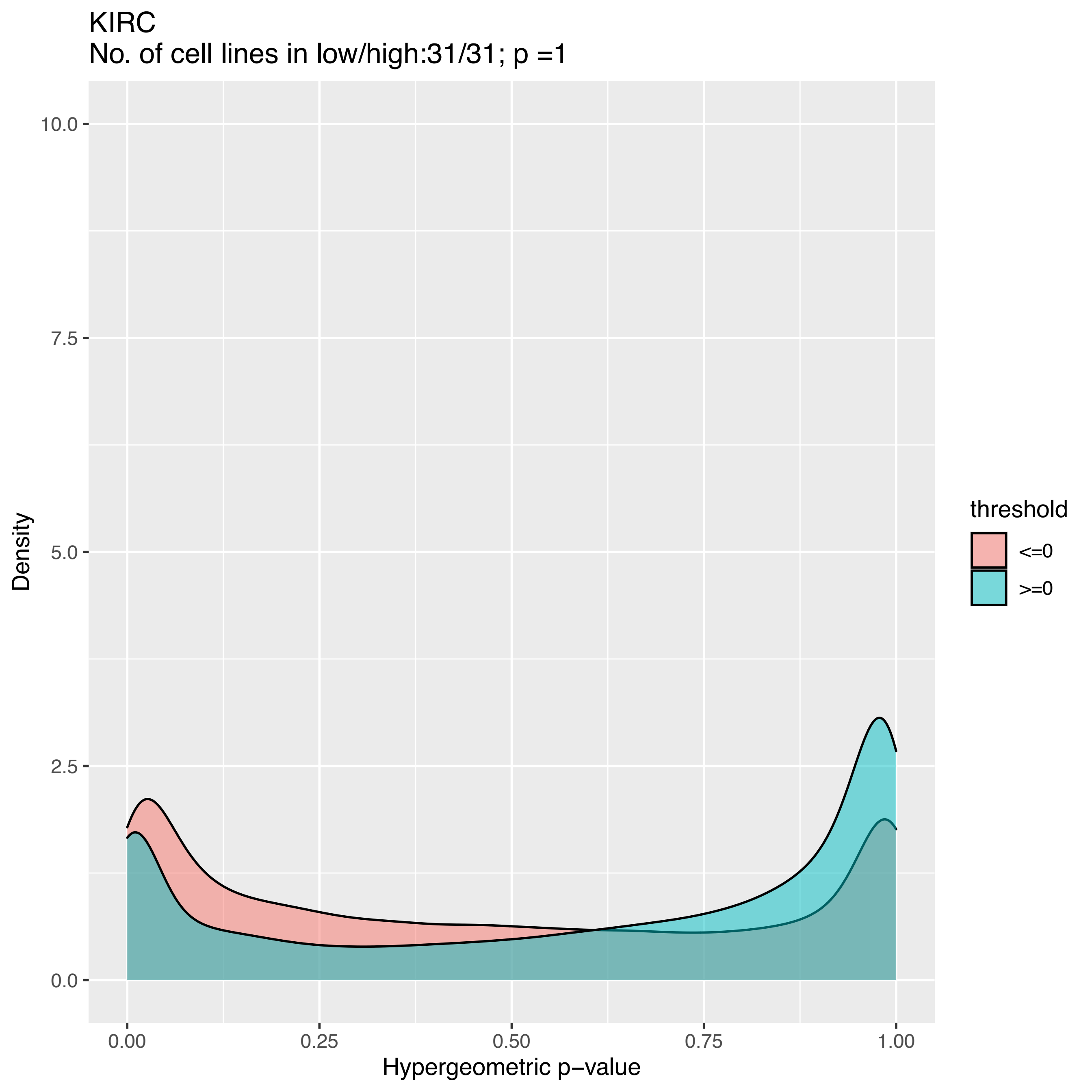

KIRC
No. of cell lines in low/high:31/31; p =1
Density
10.0
7.5
5.0
2.5
0.0
0.00
0.25
0.50
0.75
1.00
Hypergeometric p−value
threshold
<=0
>=0

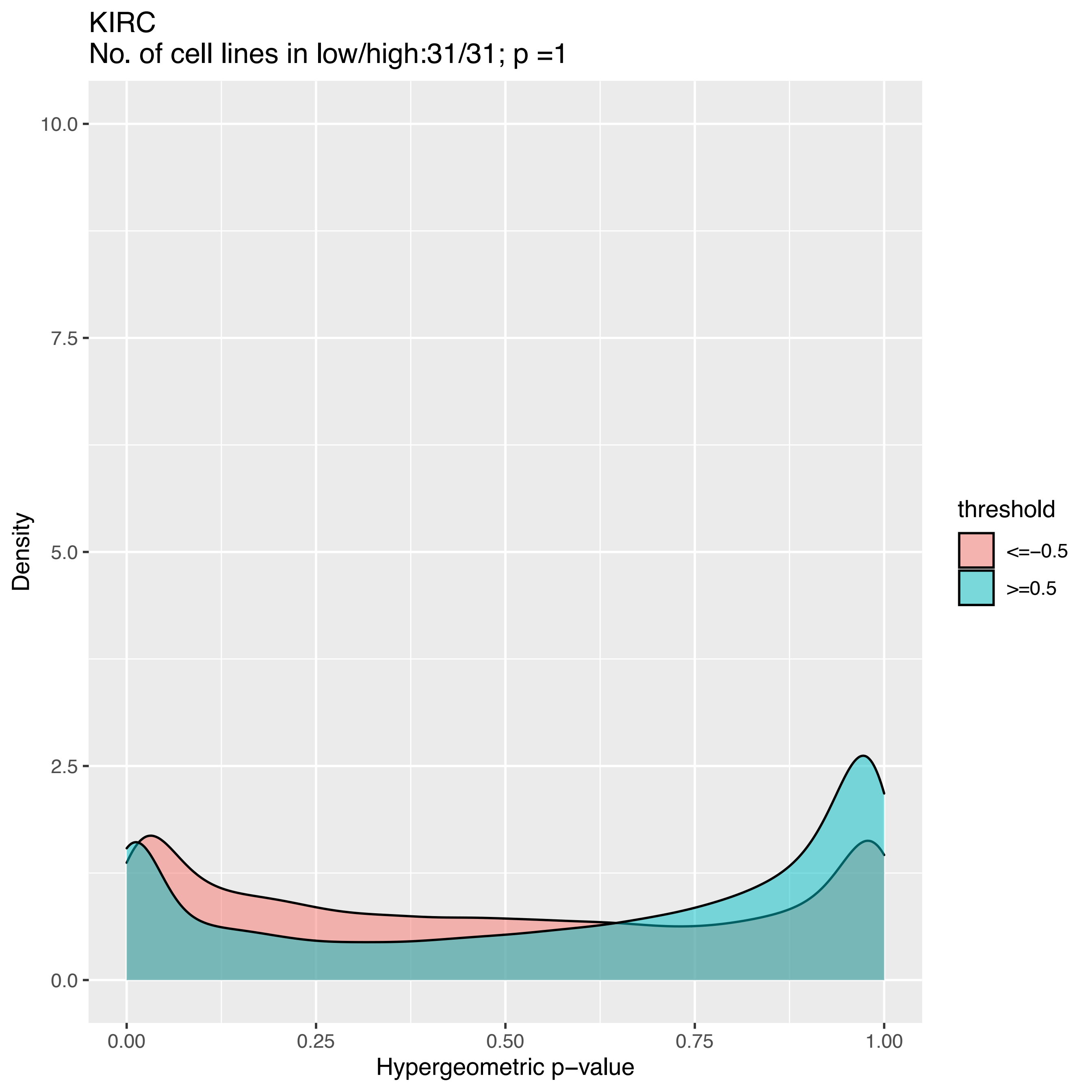
KIRC
No. of cell lines in low/high:31/31; p =1
Density
10.0
7.5
5.0
2.5
0.0
0.00
0.25
0.50
0.75
1.00
Hypergeometric p−value
threshold
<=−0.5
>=0.5

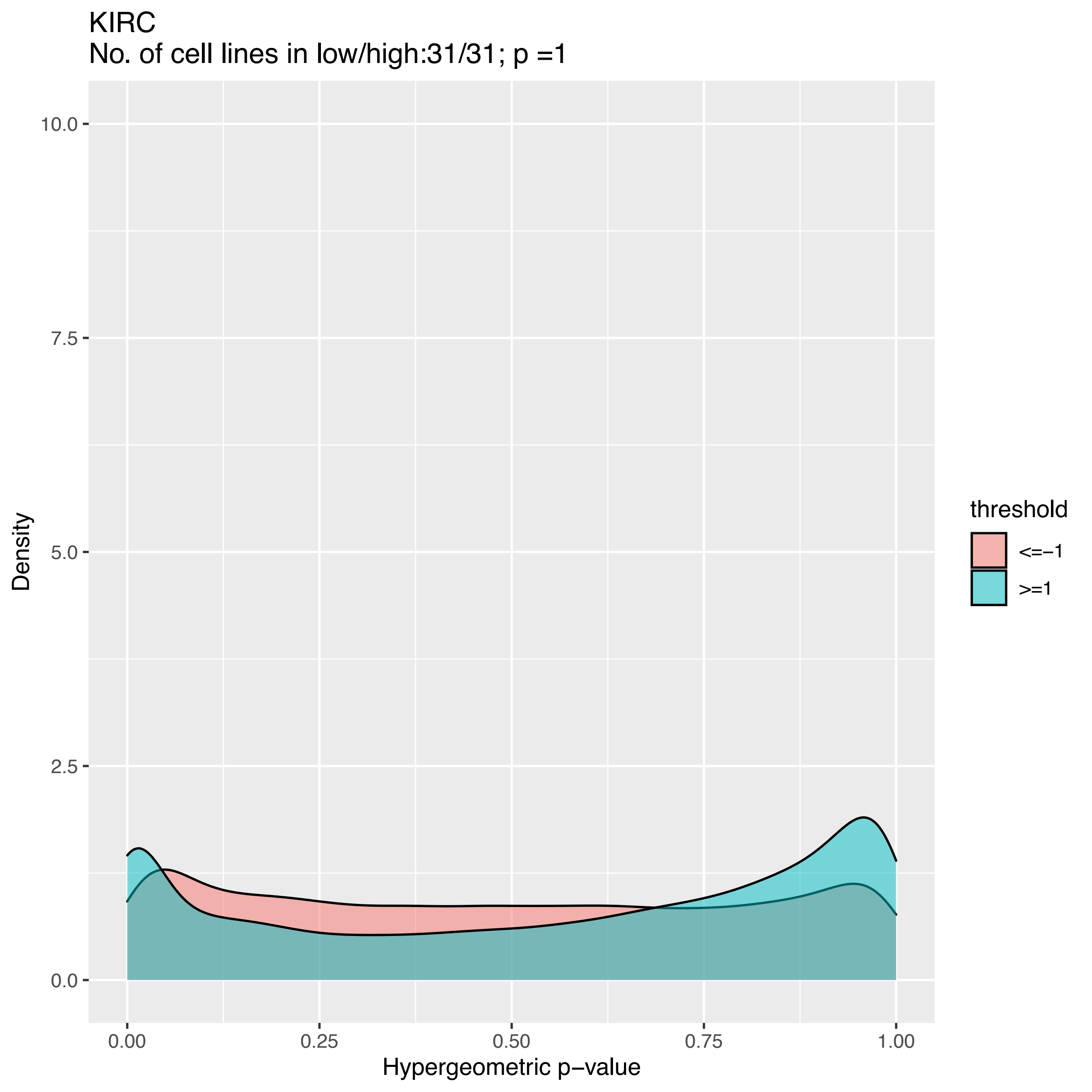

KIRC
No. of cell lines in low/high:31/31; p =1
threshold
<=−1
>=1
Density
Hypergeometric p−value
10.0
7.5
5.0
2.5
0.0
0.00
0.25
0.50
0.75
1.00

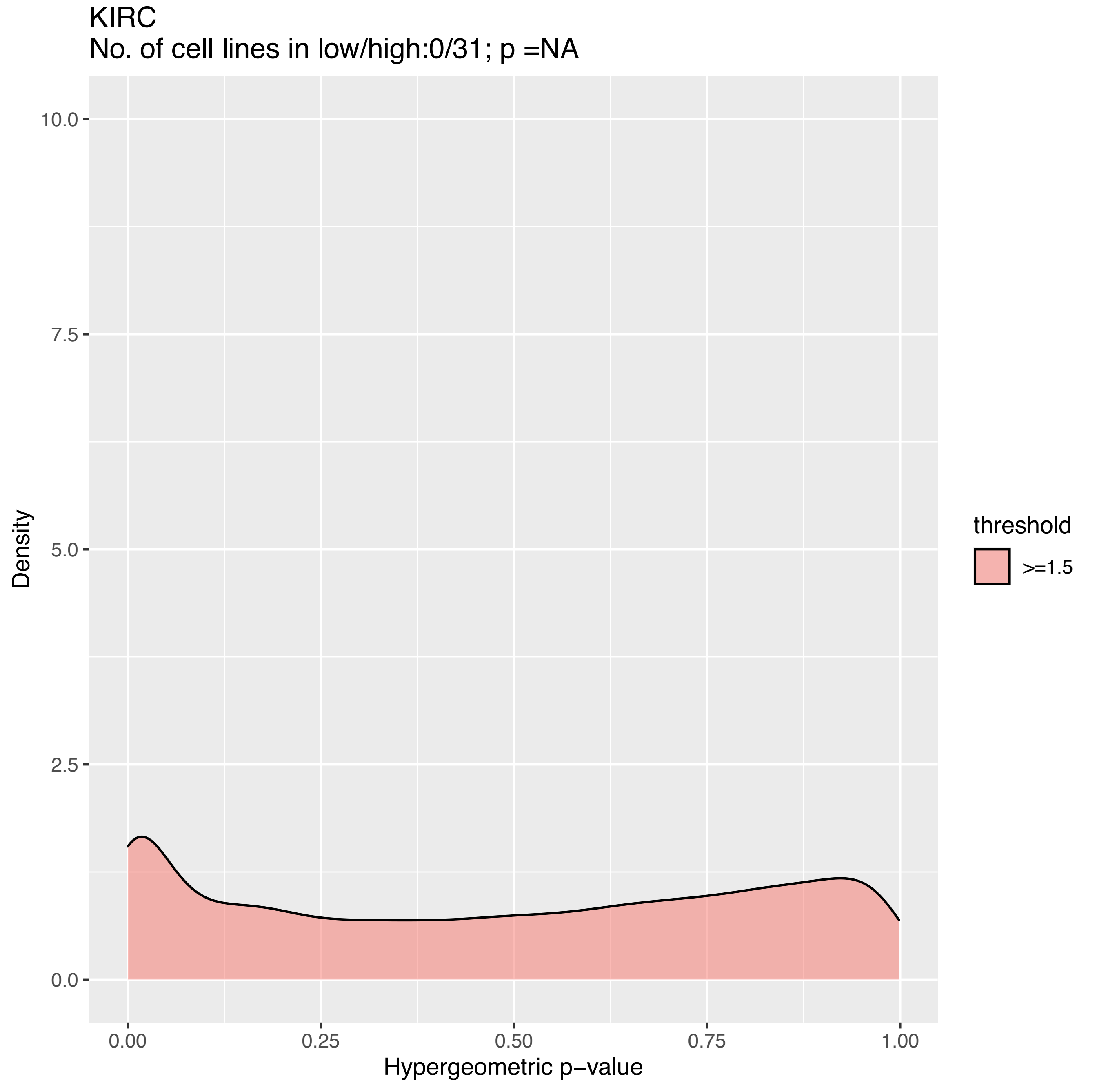

KIRC
No. of cell lines in low/high:0/31; p =NA
10.0
7.5
5.0
2.5
0.0
Density
0.00
0.25
0.50
0.75
1.00
Hypergeometric p−value
threshold
>=1.5

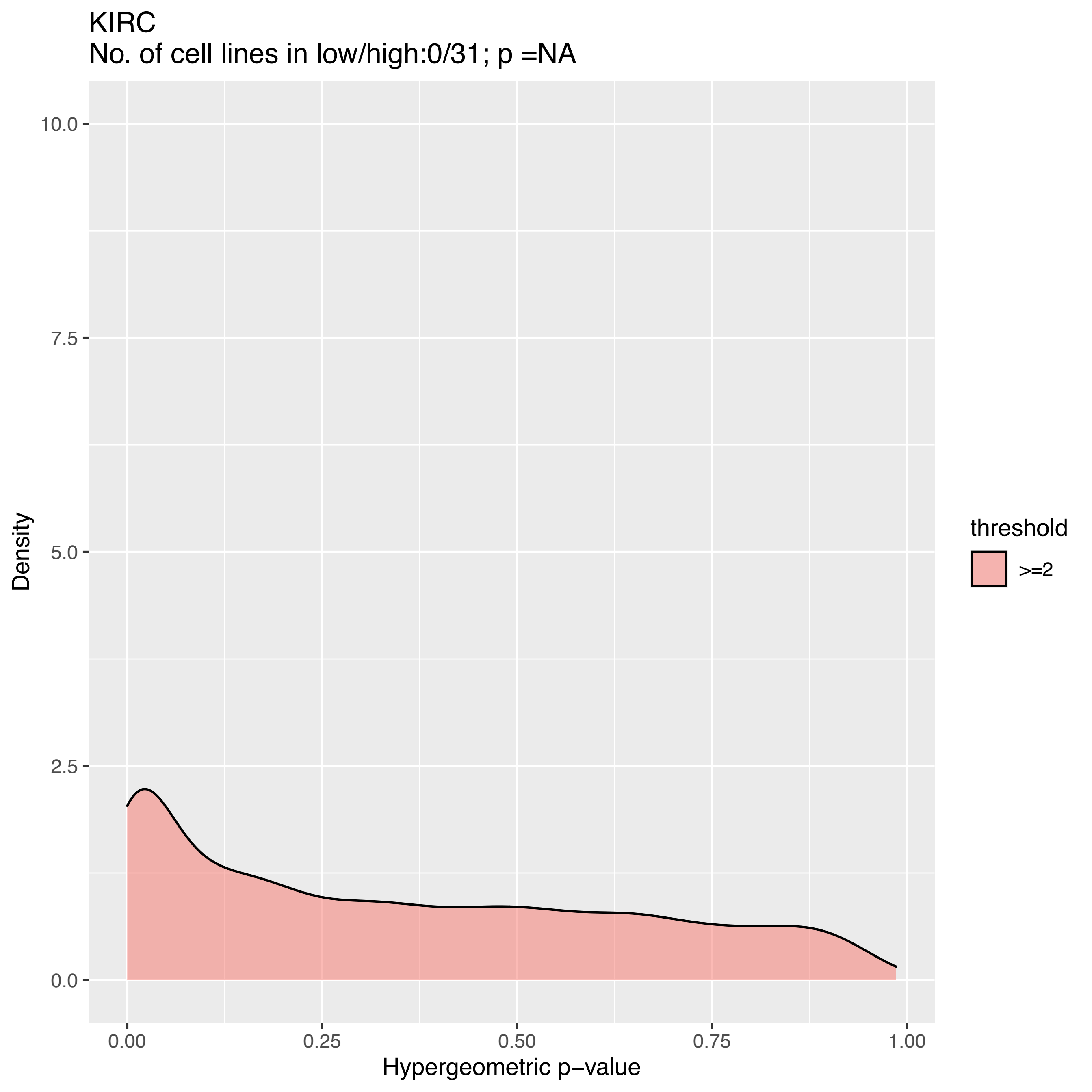
KIRC
No. of cell lines in low/high:0/31; p =NA
10.0
7.5
5.0
2.5
0.0
Density
0.00
0.25
0.50
0.75
1.00
Hypergeometric p−value
threshold
>=2

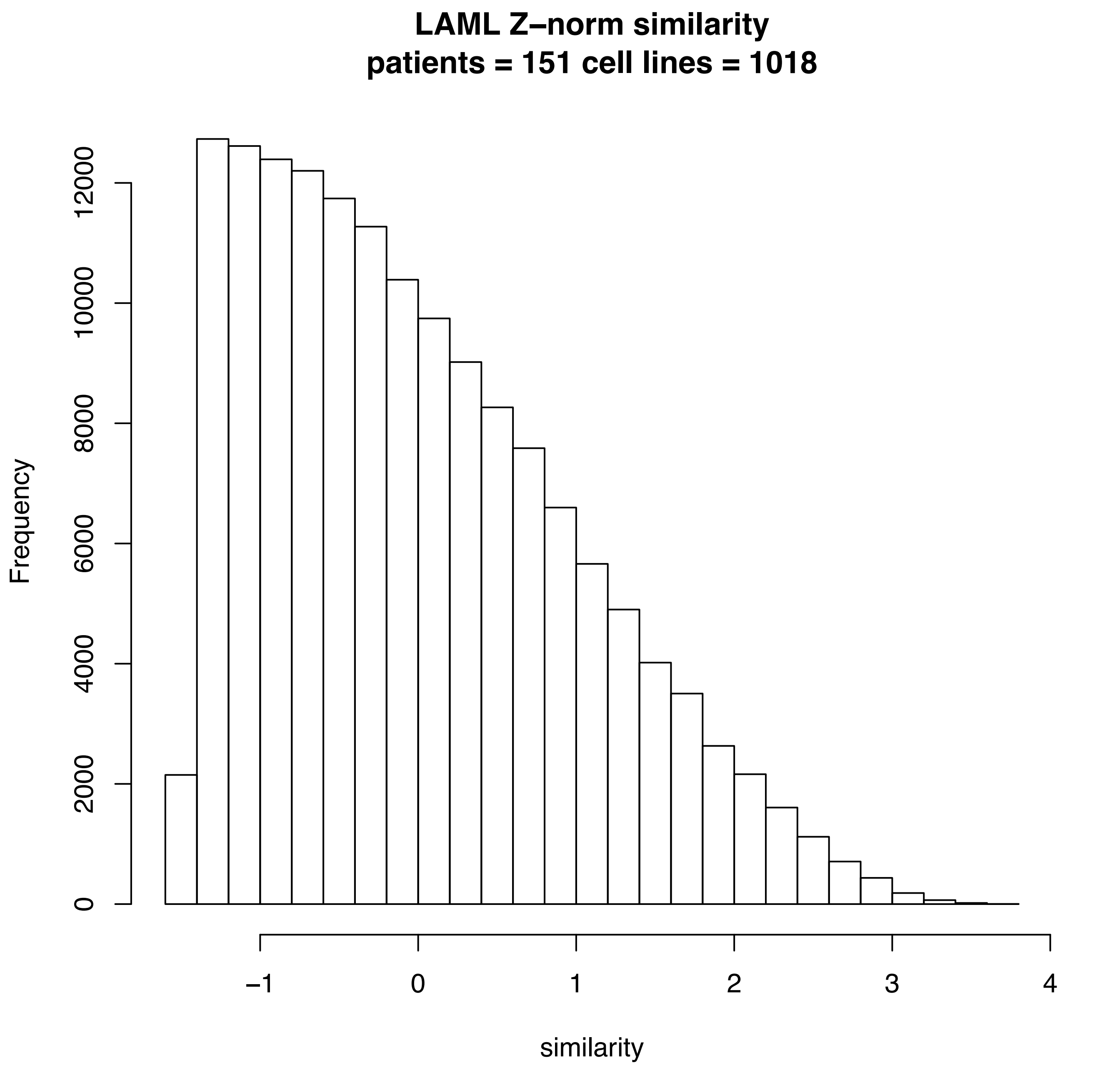
LAML Z–norm similarity
patients = 151 cell lines = 1018
Frequency
0
2000
4000
6000
8000
10000
12000
–1
0
1
2
3
4
similarity

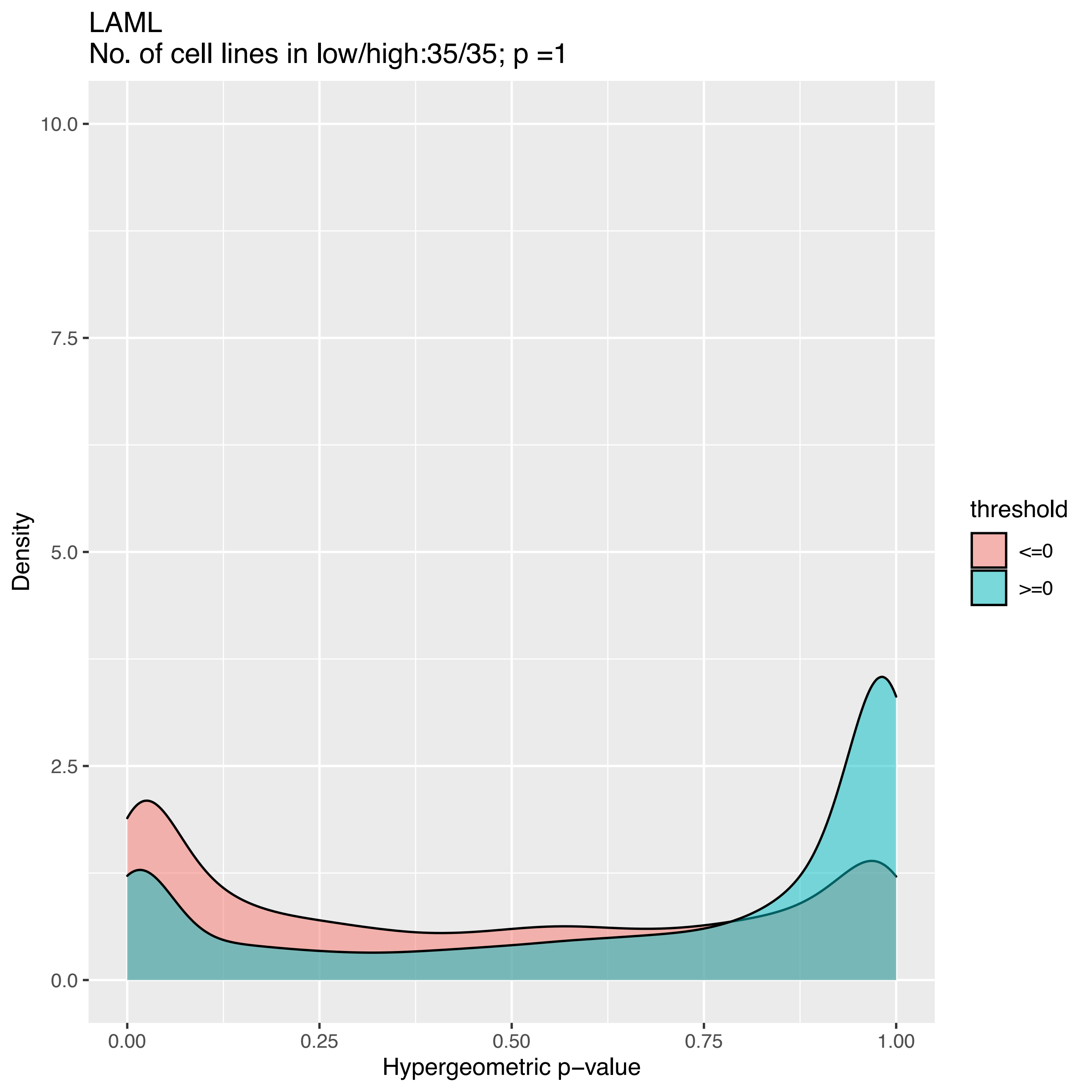

LAML
No. of cell lines in low/high:35/35; p =1
10.0
7.5
5.0
2.5
0.0
Density
0.00
0.25
0.50
0.75
1.00
Hypergeometric p−value
threshold
<=0
>=0

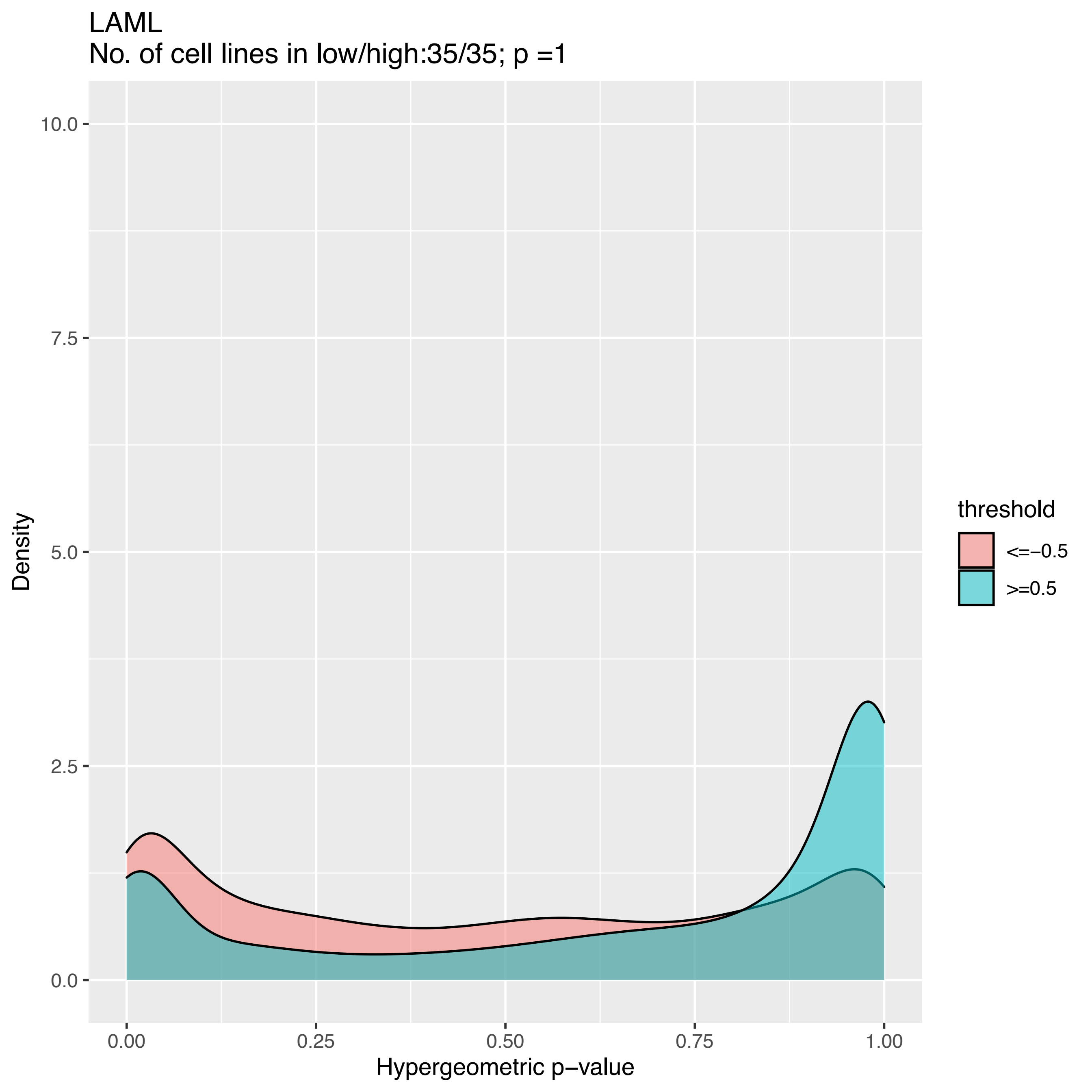
LAML
No. of cell lines in low/high:35/35; p =1
Density
10.0
7.5
5.0
2.5
0.0
0.00
0.25
0.50
0.75
1.00
Hypergeometric p−value
threshold
<=−0.5
>=0.5

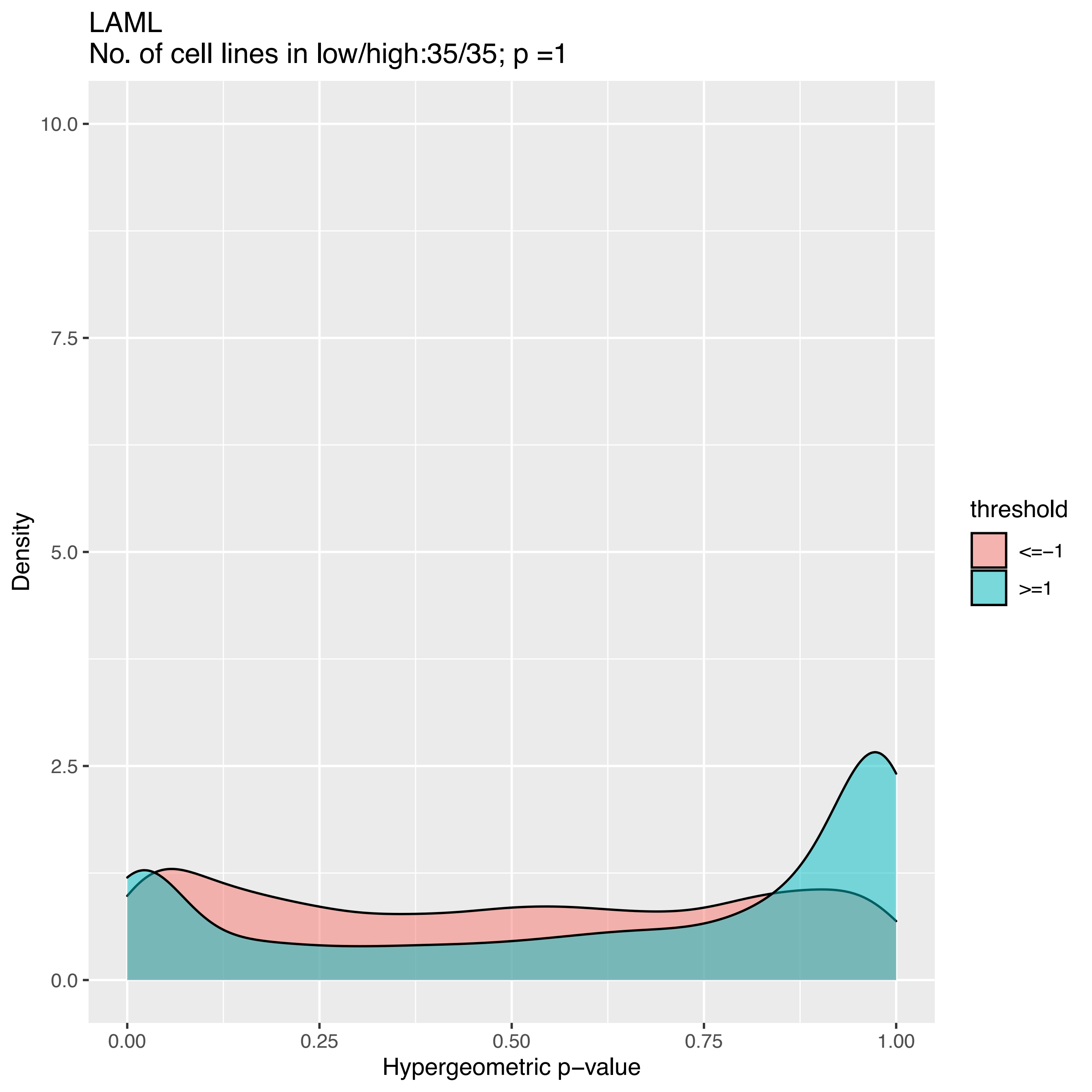
LAML
No. of cell lines in low/high:35/35; p =1
Density
10.0
7.5
5.0
2.5
0.0
0.00
0.25
0.50
0.75
1.00
Hypergeometric p−value
threshold
<=−1
>=1

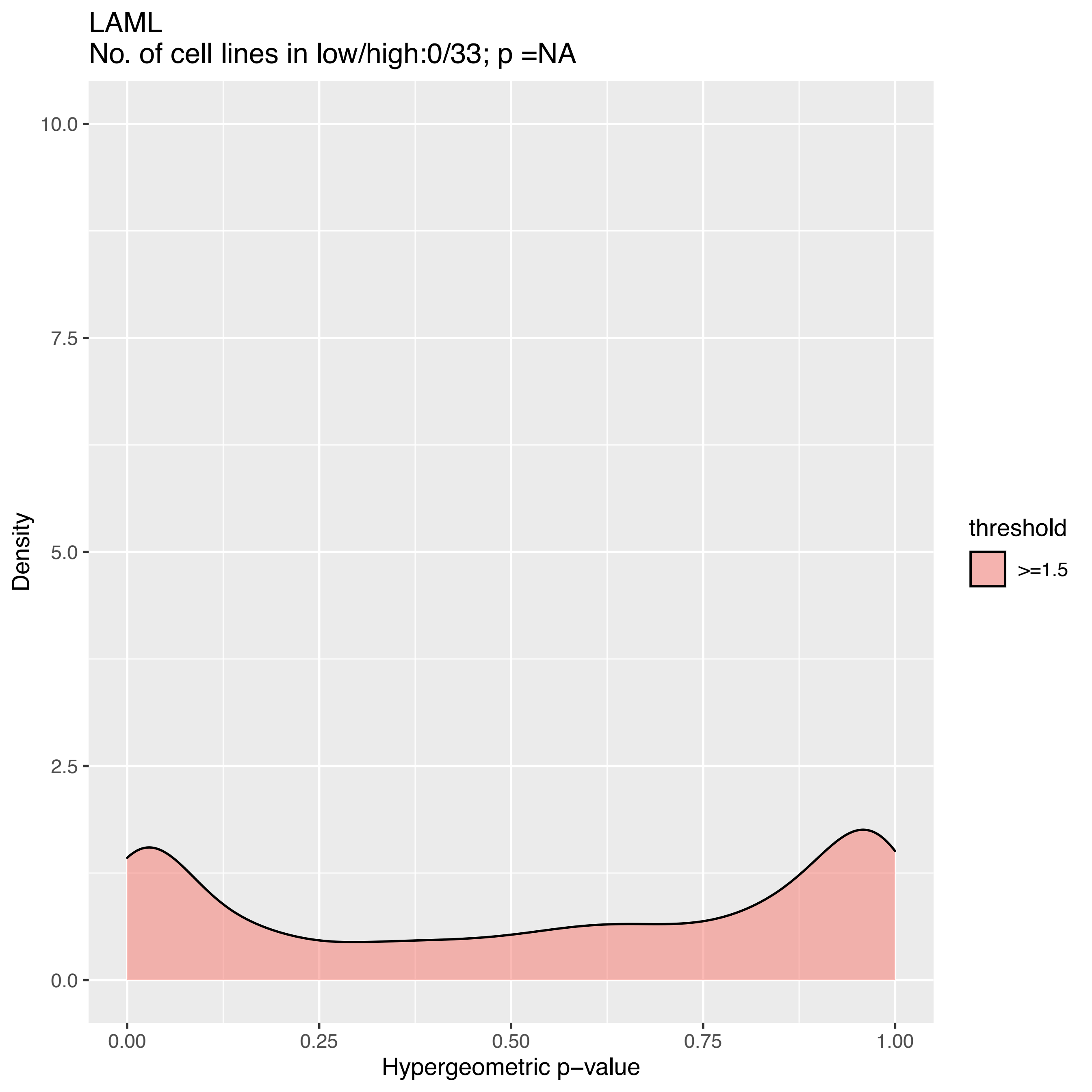
LAML
No. of cell lines in low/high:0/33; p =NA
Density
10.0
7.5
5.0
2.5
0.0
0.00
0.25
0.50
0.75
1.00
Hypergeometric p−value
threshold
>=1.5

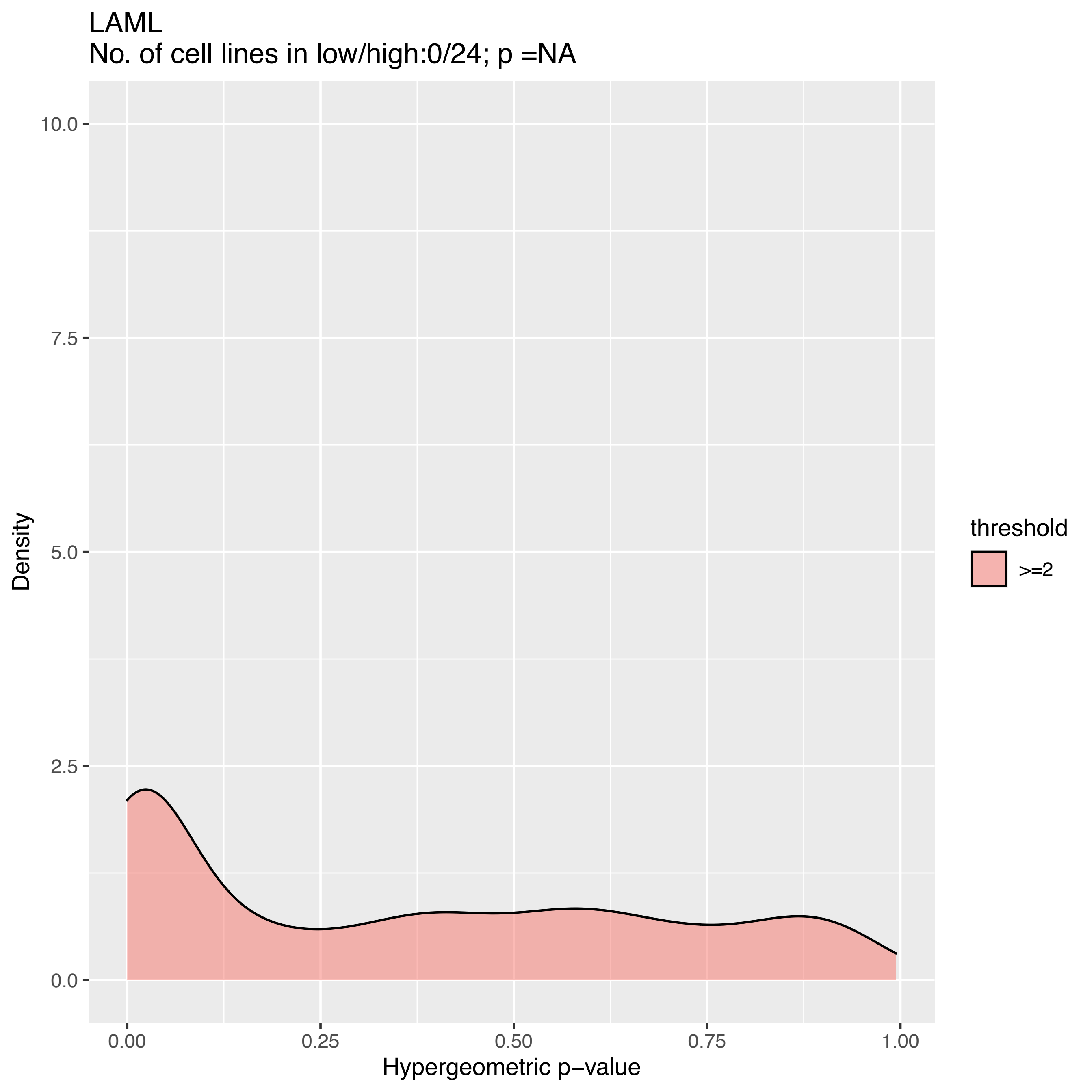

LAML
No. of cell lines in low/high:0/24; p =NA
10.0
7.5
5.0
2.5
0.0
Density
0.00
0.25
0.50
0.75
1.00
Hypergeometric p−value
threshold
>=2

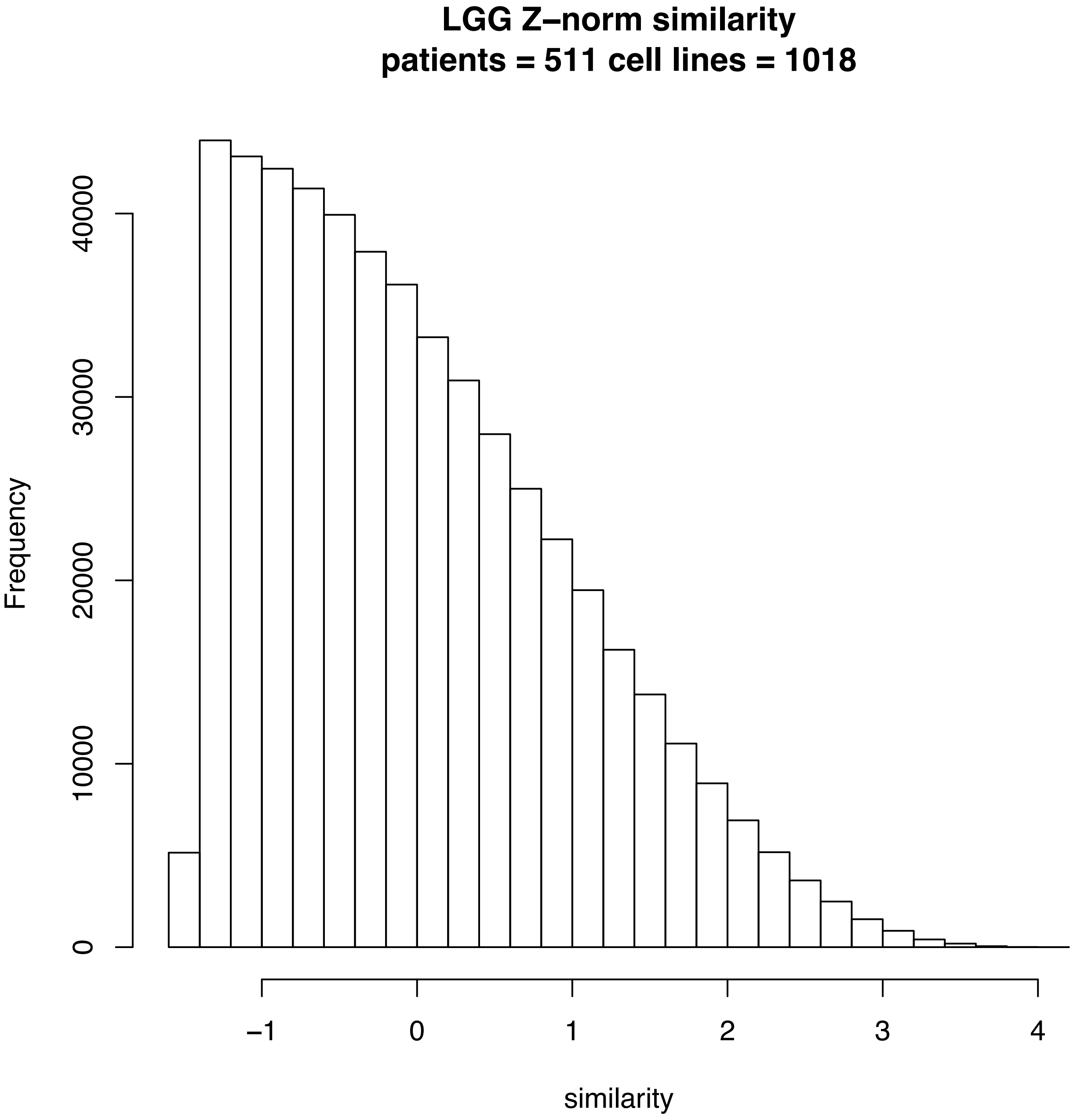
LGG Z–norm similarity
patients = 511 cell lines = 1018
Frequency
0
10000
20000
30000
40000
–1
0
1
2
3
4
similarity

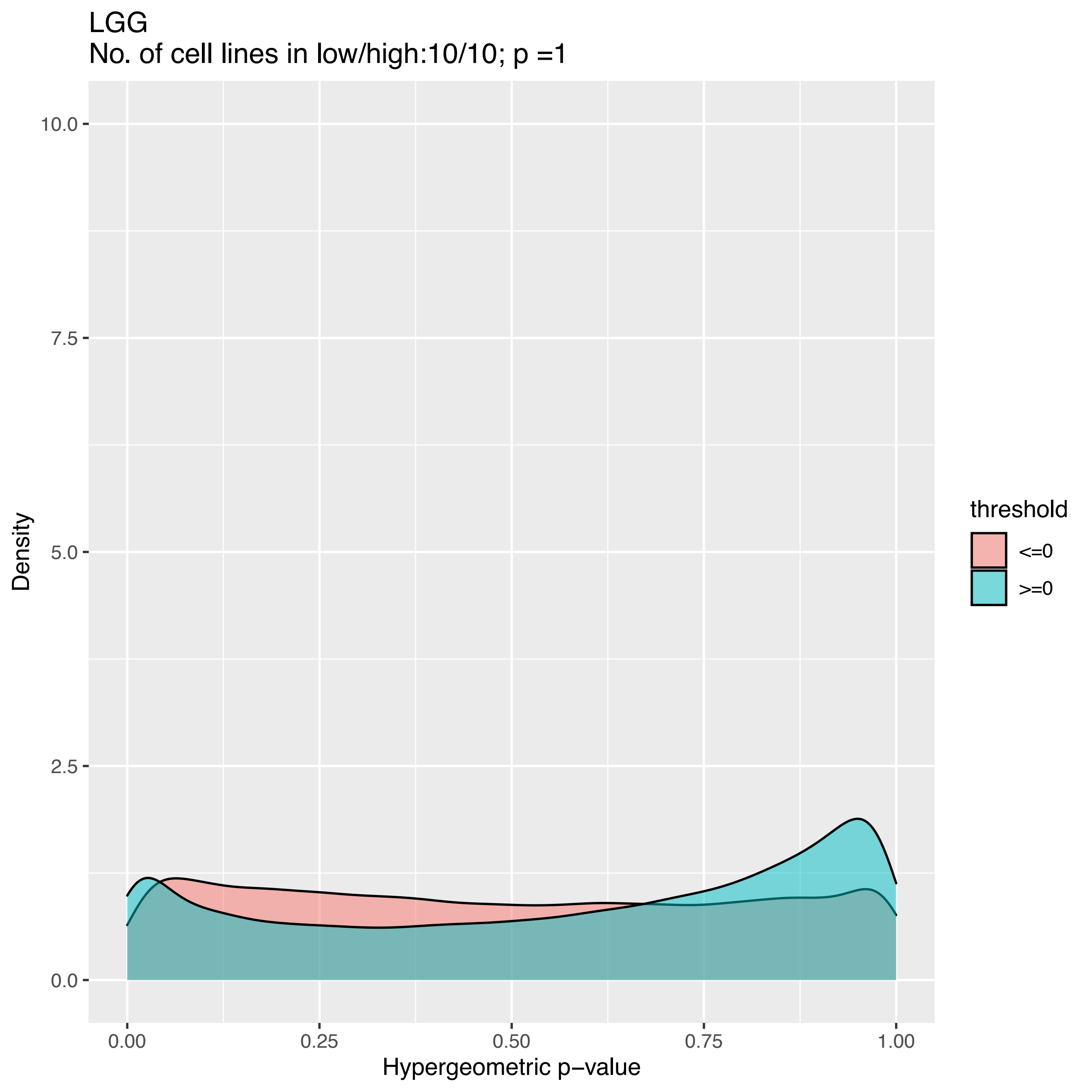
LGG
No. of cell lines in low/high:10/10; p =1
Density
Hypergeometric p−value
0.0
2.5
5.0
7.5
10.0
0.00
0.25
0.50
0.75
1.00
threshold
<=0
>=0

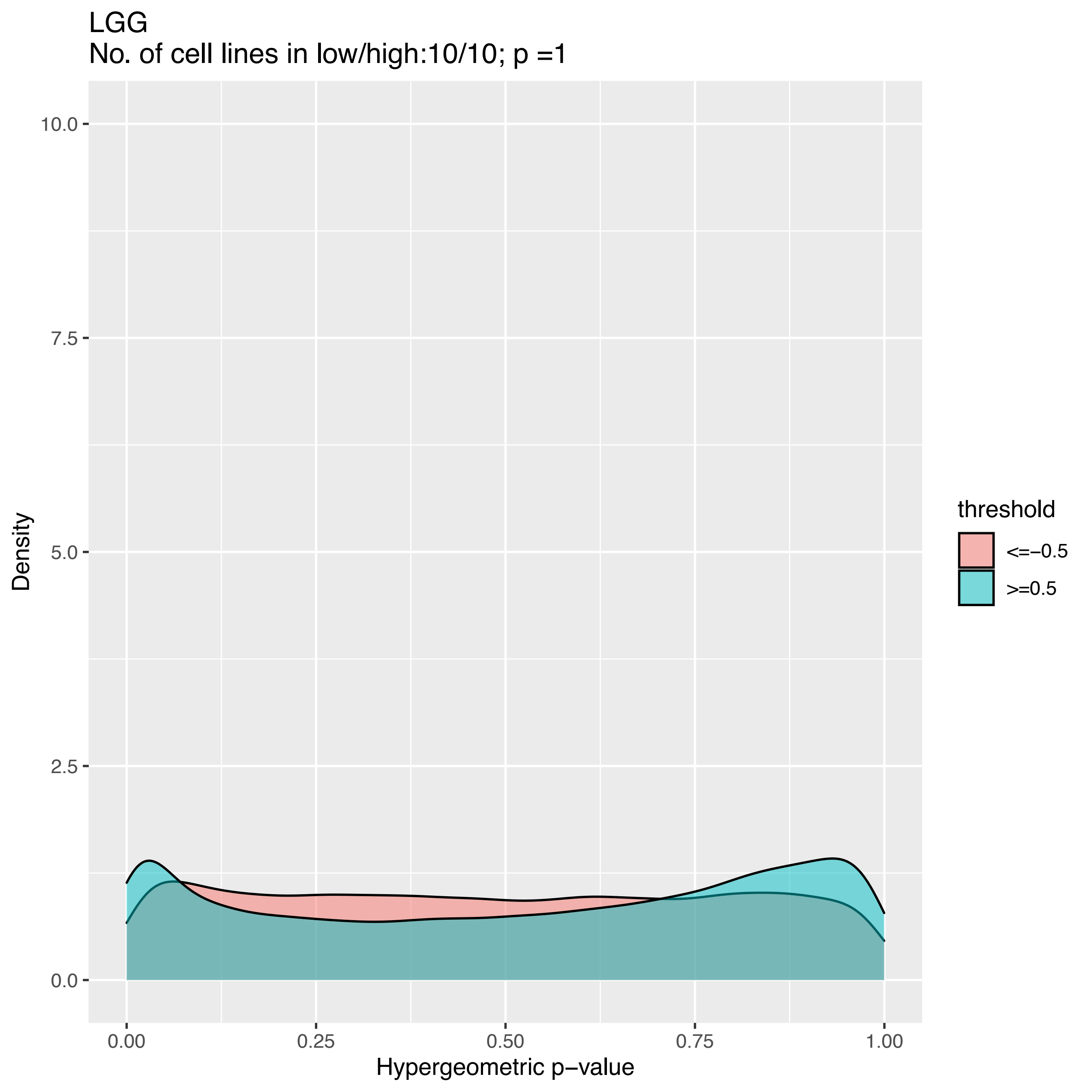

LGG
No. of cell lines in low/high:10/10; p =1
10.0
7.5
5.0
2.5
0.0
Density
0.00
0.25
0.50
0.75
1.00
Hypergeometric p−value
threshold
<=−0.5
>=0.5

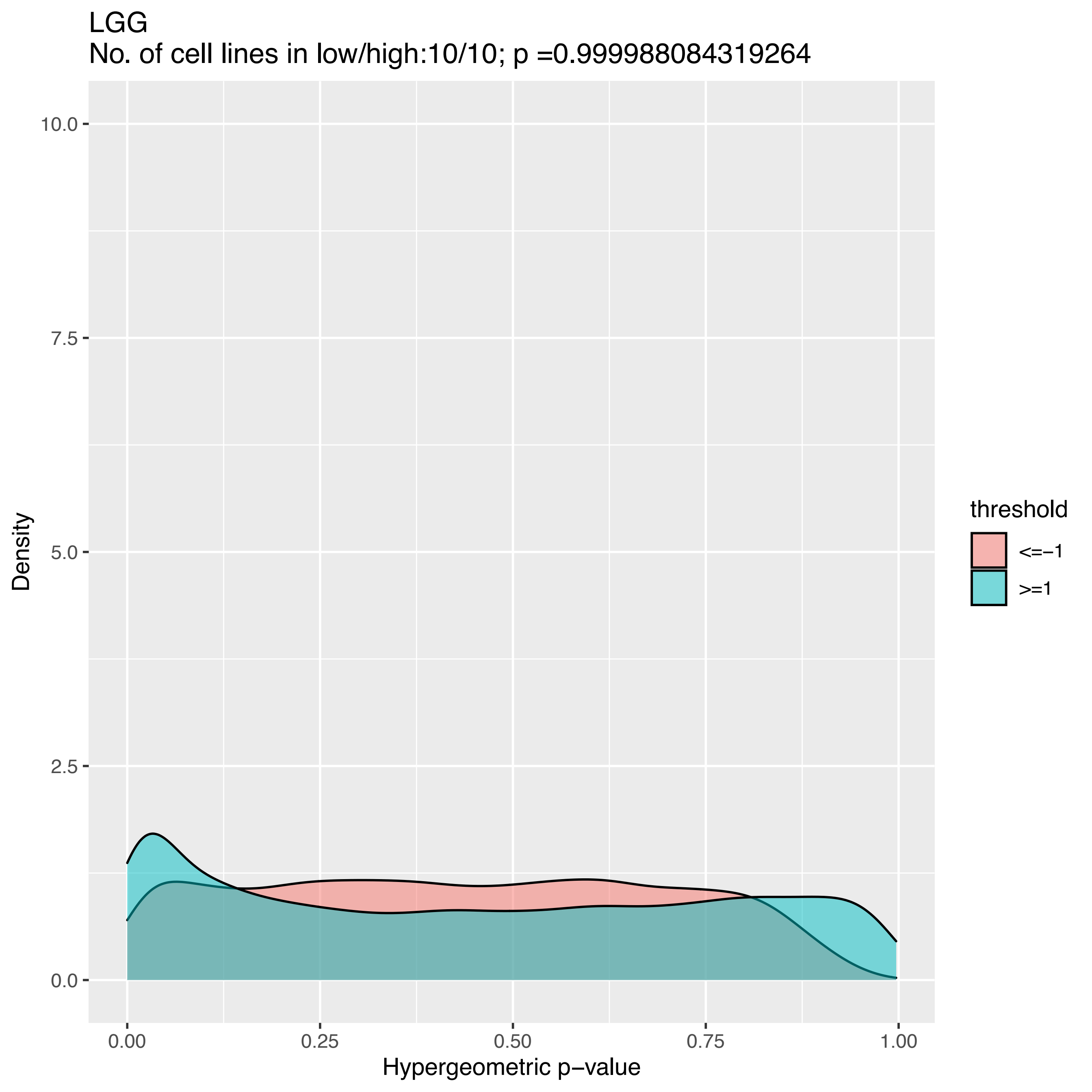

LGG
No. of cell lines in low/high:10/10; p =0.999988084319264
Density
10.0
7.5
5.0
2.5
0.0
0.00
0.25
0.50
0.75
1.00
Hypergeometric p−value
threshold
<=−1
>=1

LGG
No. of cell lines in low/high:0/10; p =NA

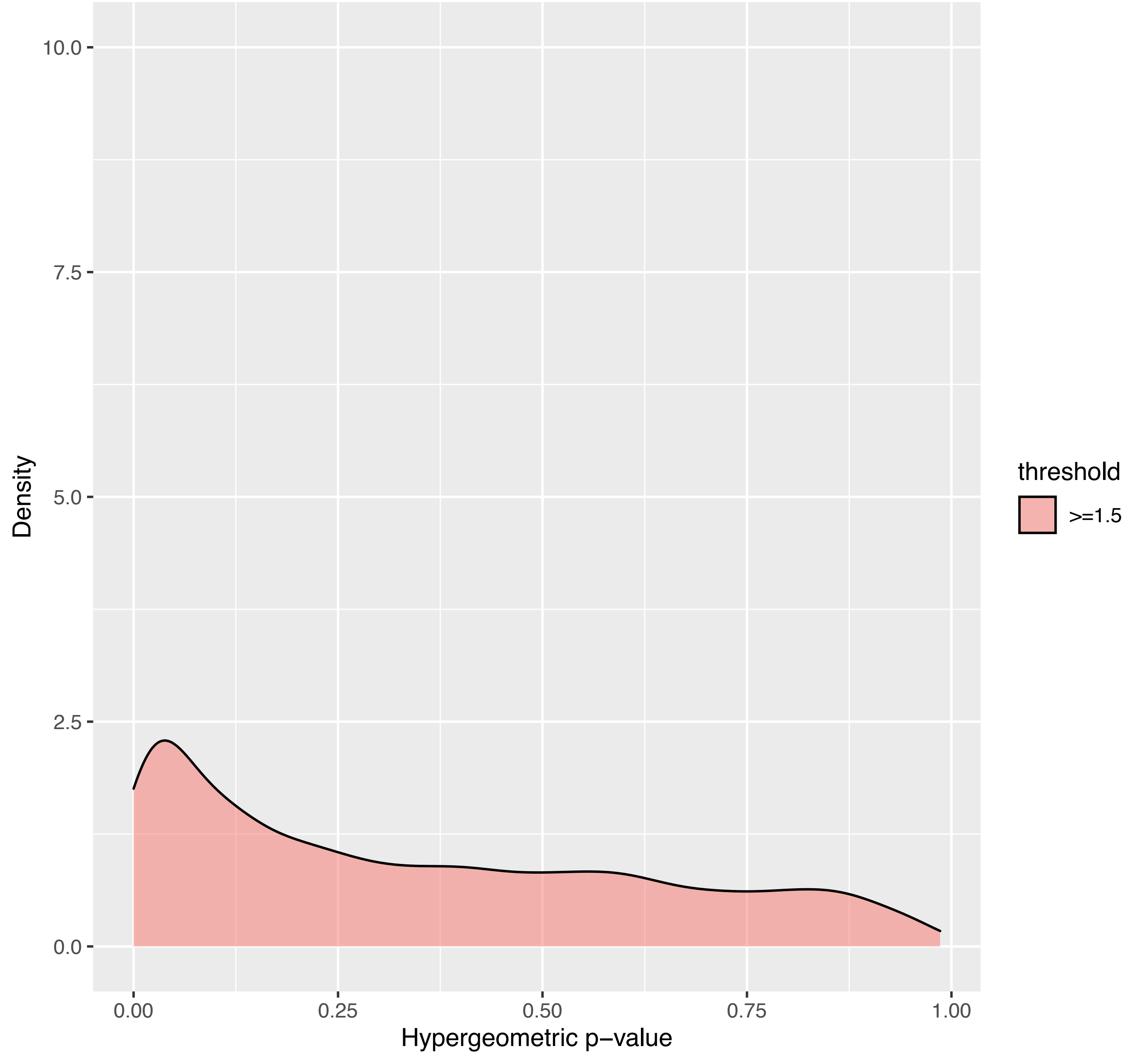

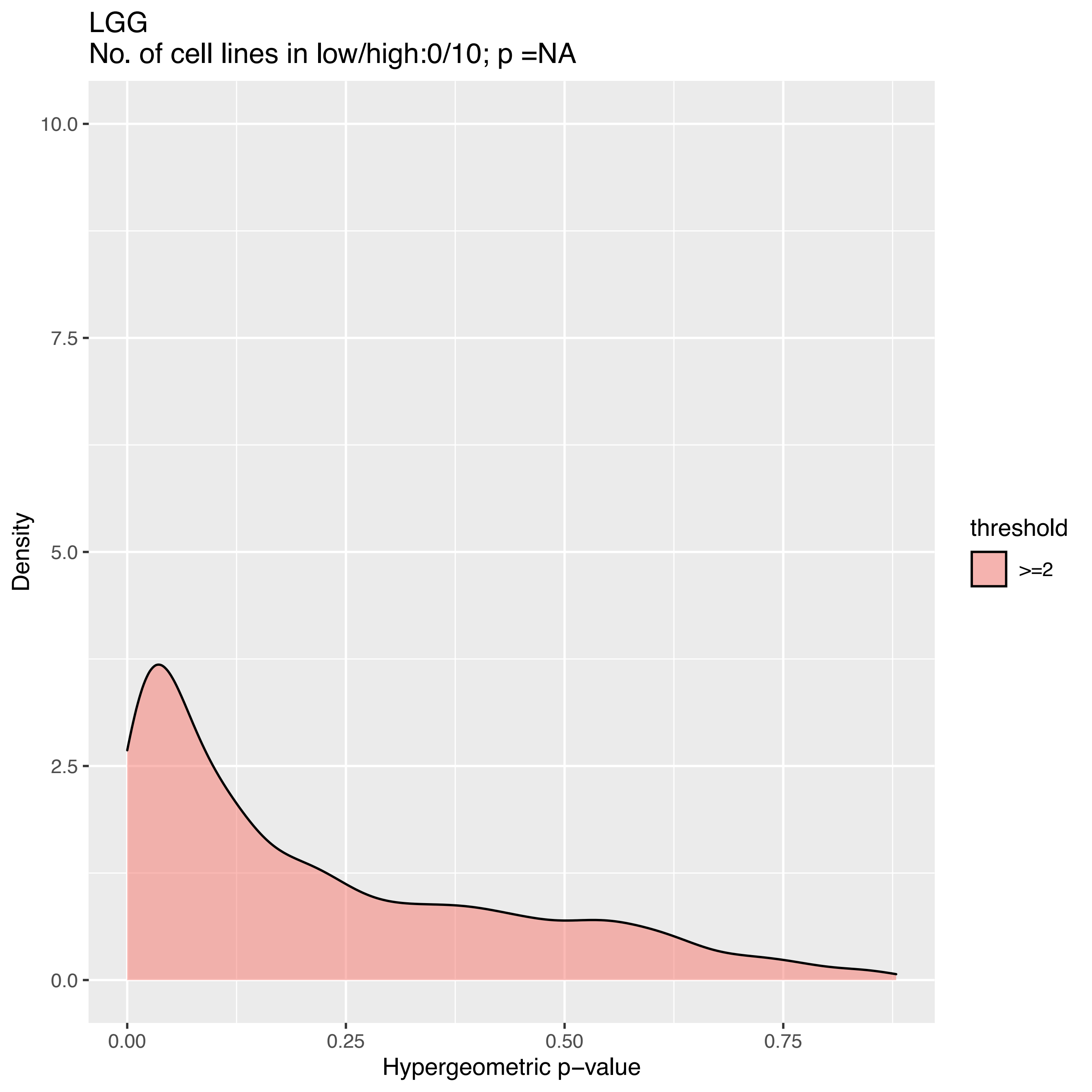
LGG
No. of cell lines in low/high:0/10; p =NA
Density
10.0
7.5
5.0
2.5
0.0
0.00
0.25
0.50
0.75
Hypergeometric p−value
threshold
>=2

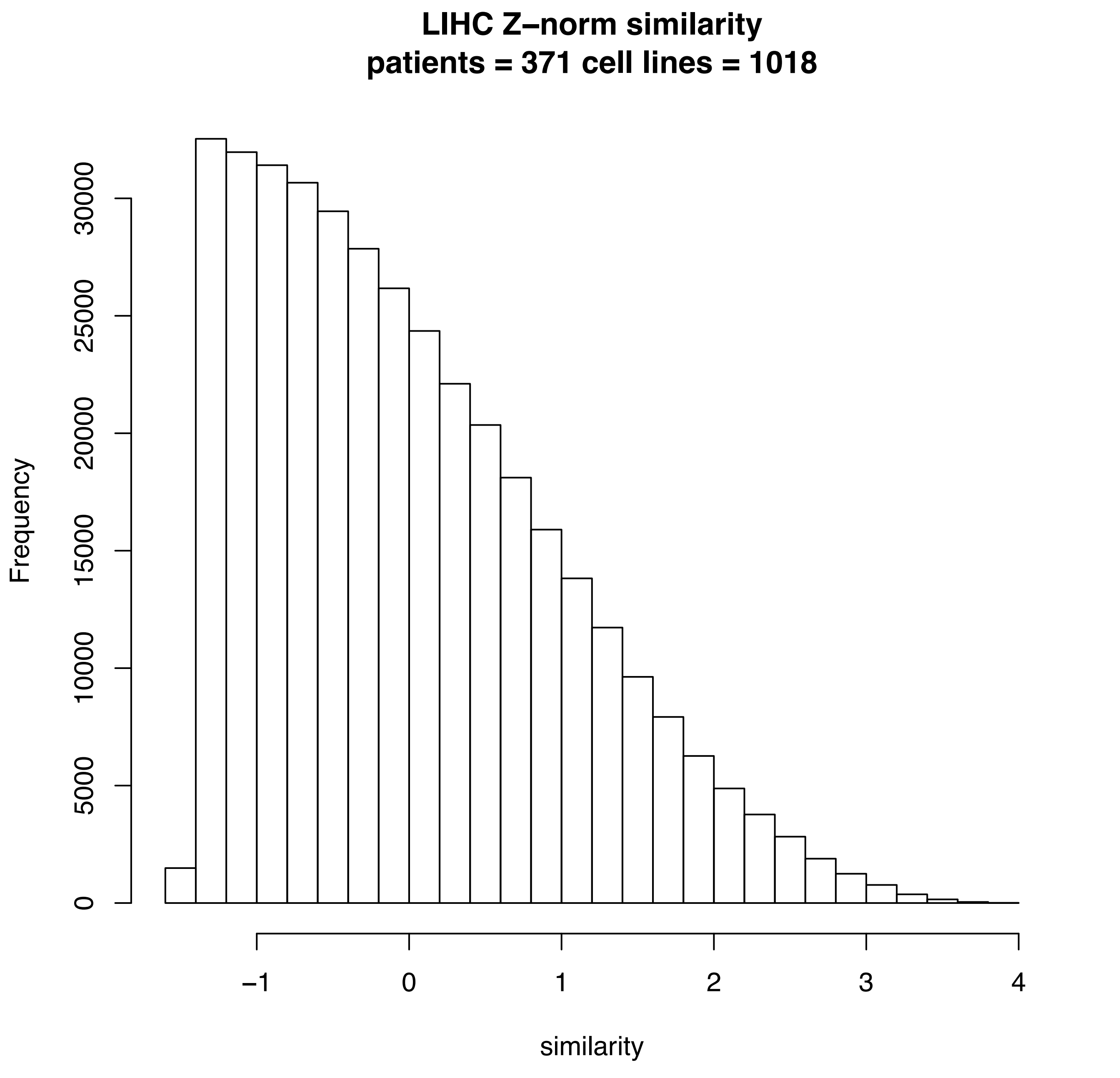

LIHC Z–norm similarity
patients = 371 cell lines = 1018
Frequency
0
5000
10000
15000
20000
25000
30000
–1
0
1
2
3
4
similarity

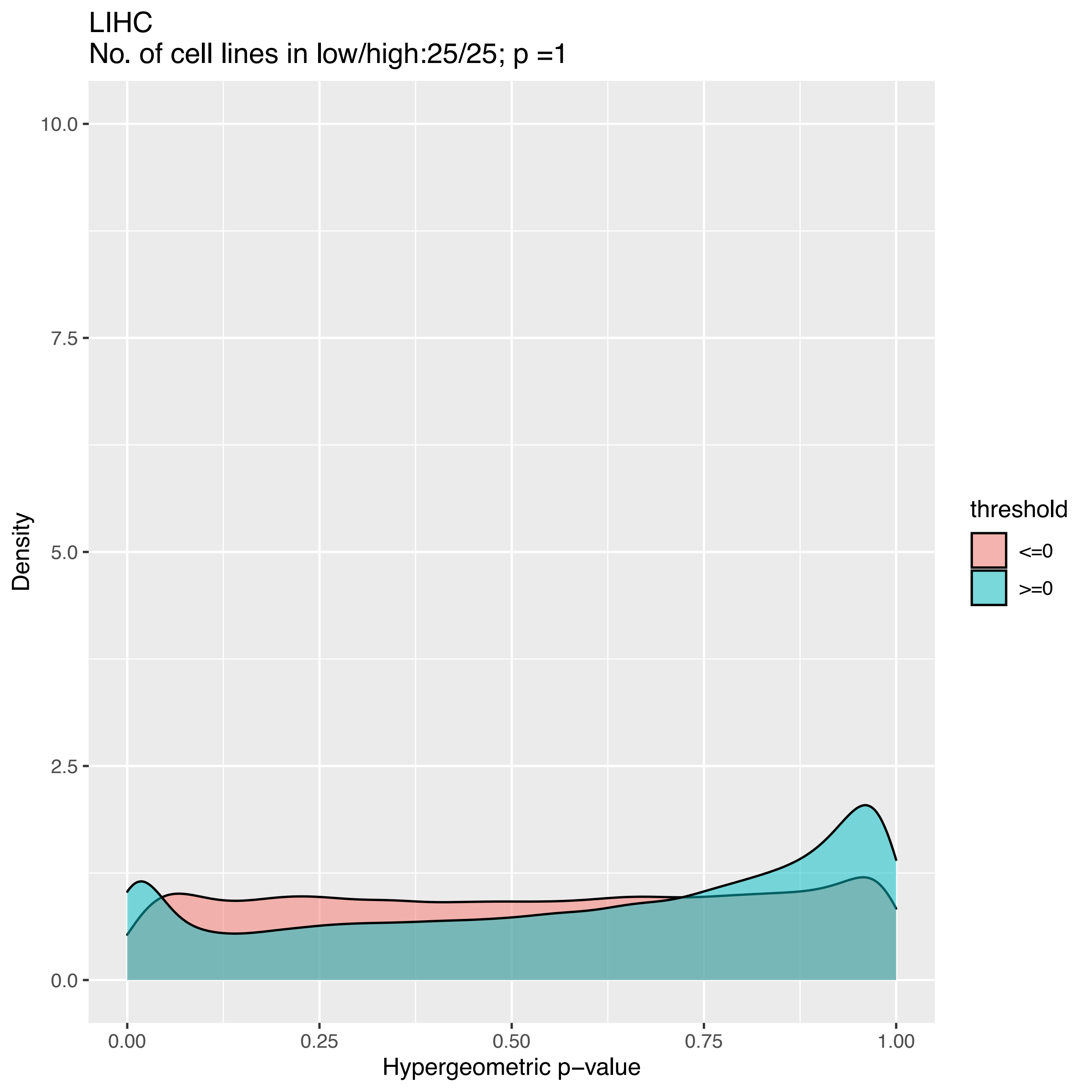

LIHC
No. of cell lines in low/high:25/25; p =1
Density
10.0
7.5
5.0
2.5
0.0
0.00
0.25
0.50
0.75
1.00
Hypergeometric p−value
threshold
<=0
>=0

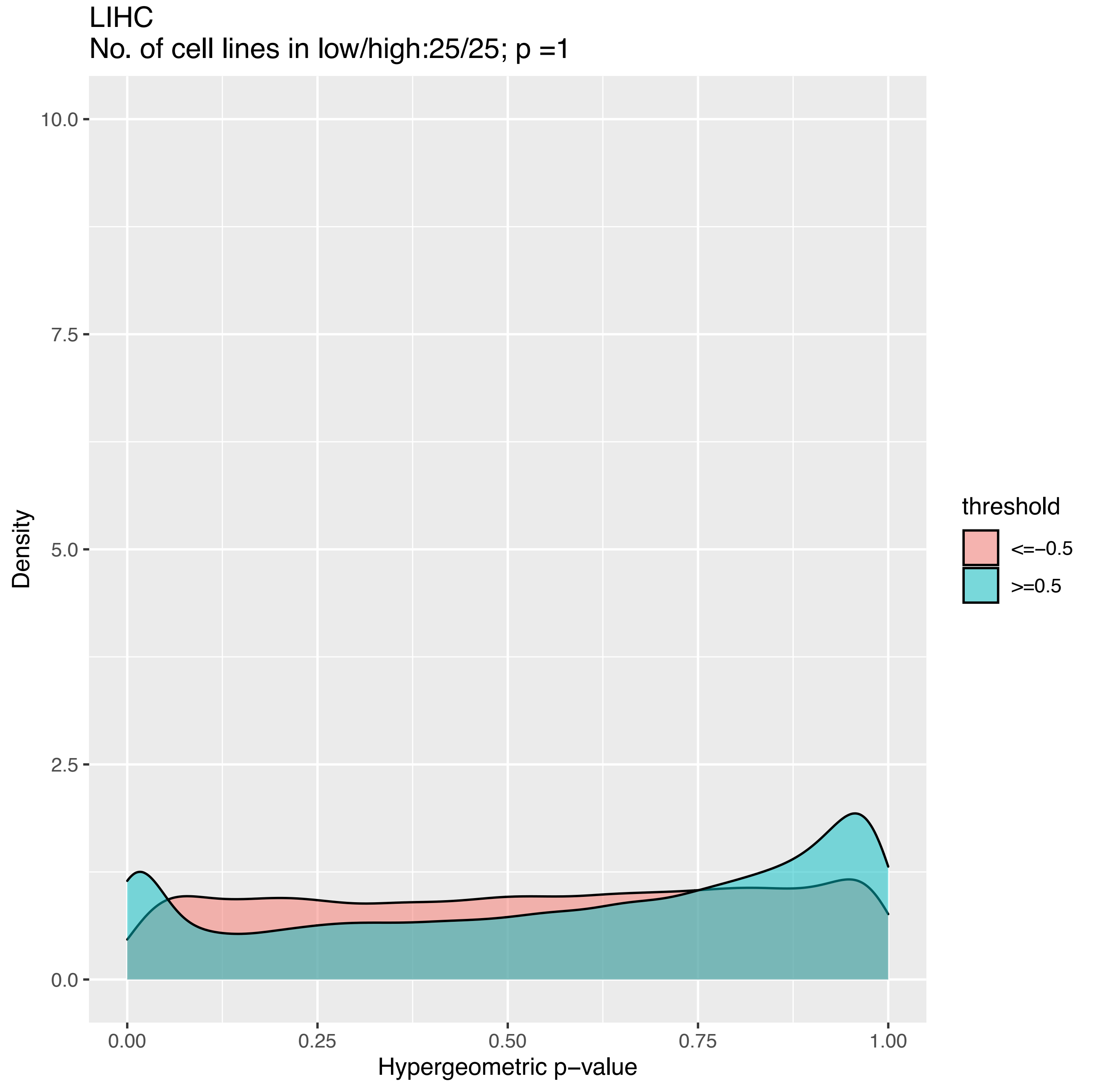
LIHC
No. of cell lines in low/high:25/25; p =1
Density
Hypergeometric p−value
0.0
2.5
5.0
7.5
10.0
0.00
0.25
0.50
0.75
1.00
threshold
<=−0.5
>=0.5

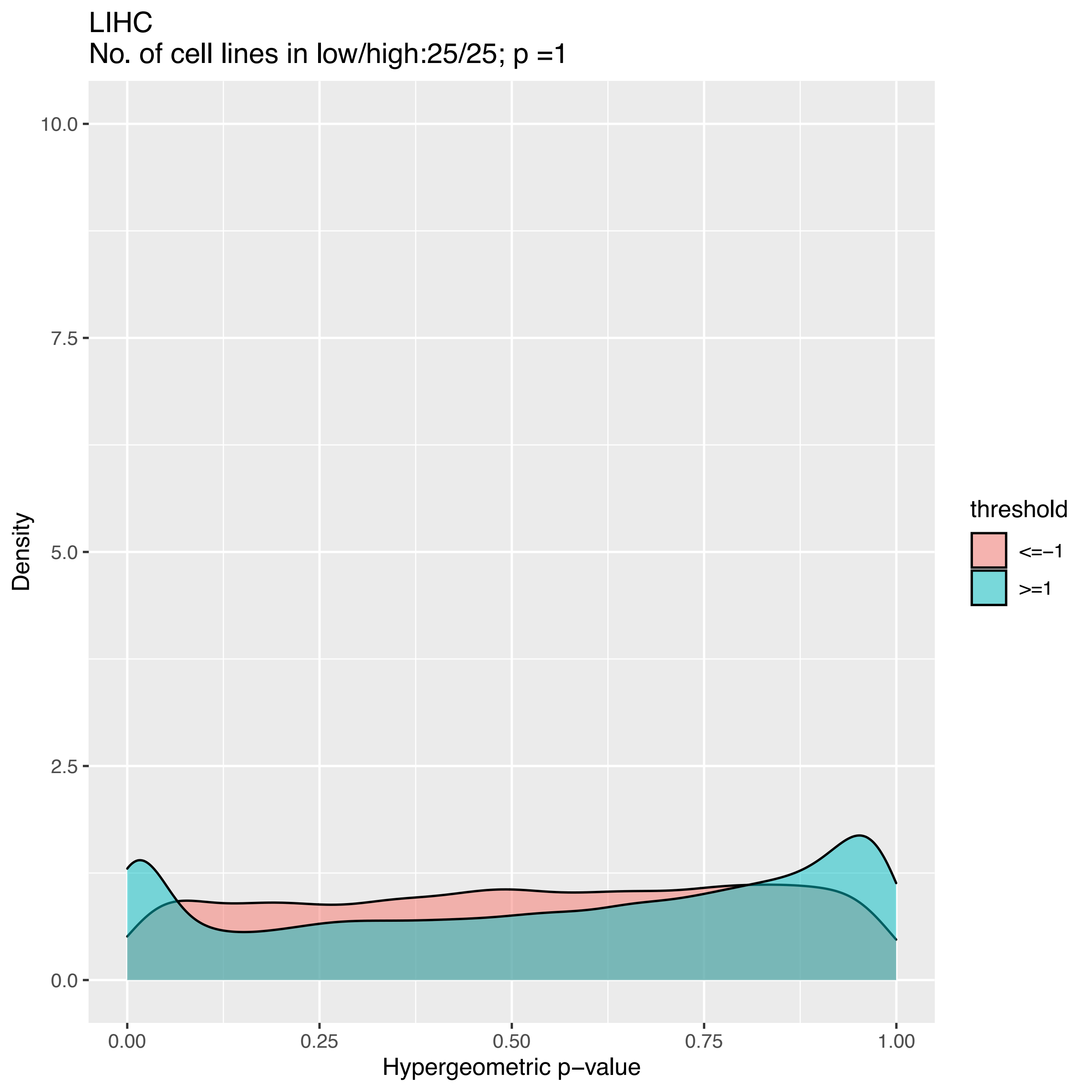
LIHC
No. of cell lines in low/high:25/25; p =1
Density
10.0
7.5
5.0
2.5
0.0
0.00
0.25
0.50
0.75
1.00
Hypergeometric p−value
threshold
<=−1
>=1

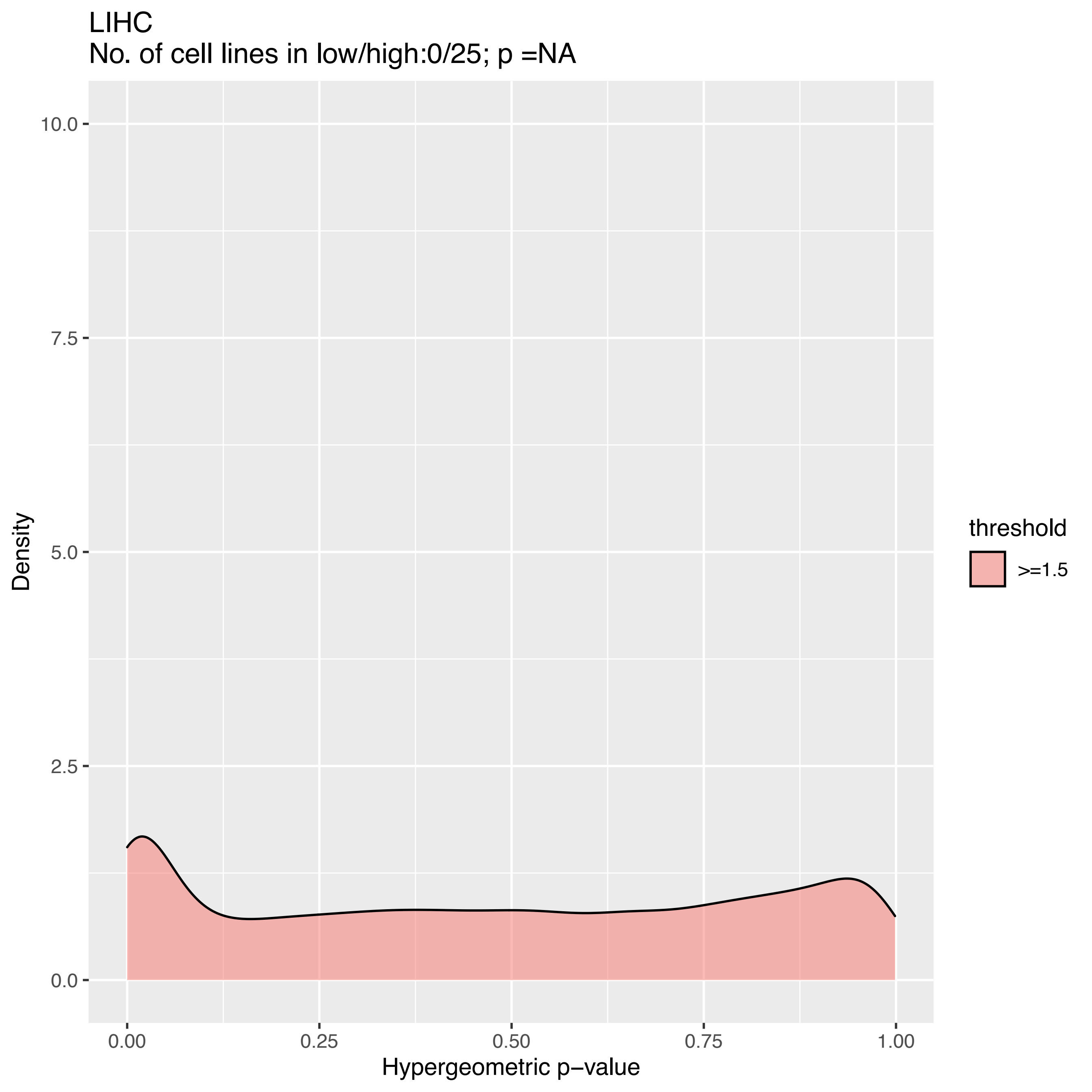

LIHC
No. of cell lines in low/high:0/25; p =NA
10.0
7.5
5.0
2.5
0.0
Density
0.00
0.25
0.50
0.75
1.00
Hypergeometric p−value
threshold
>=1.5

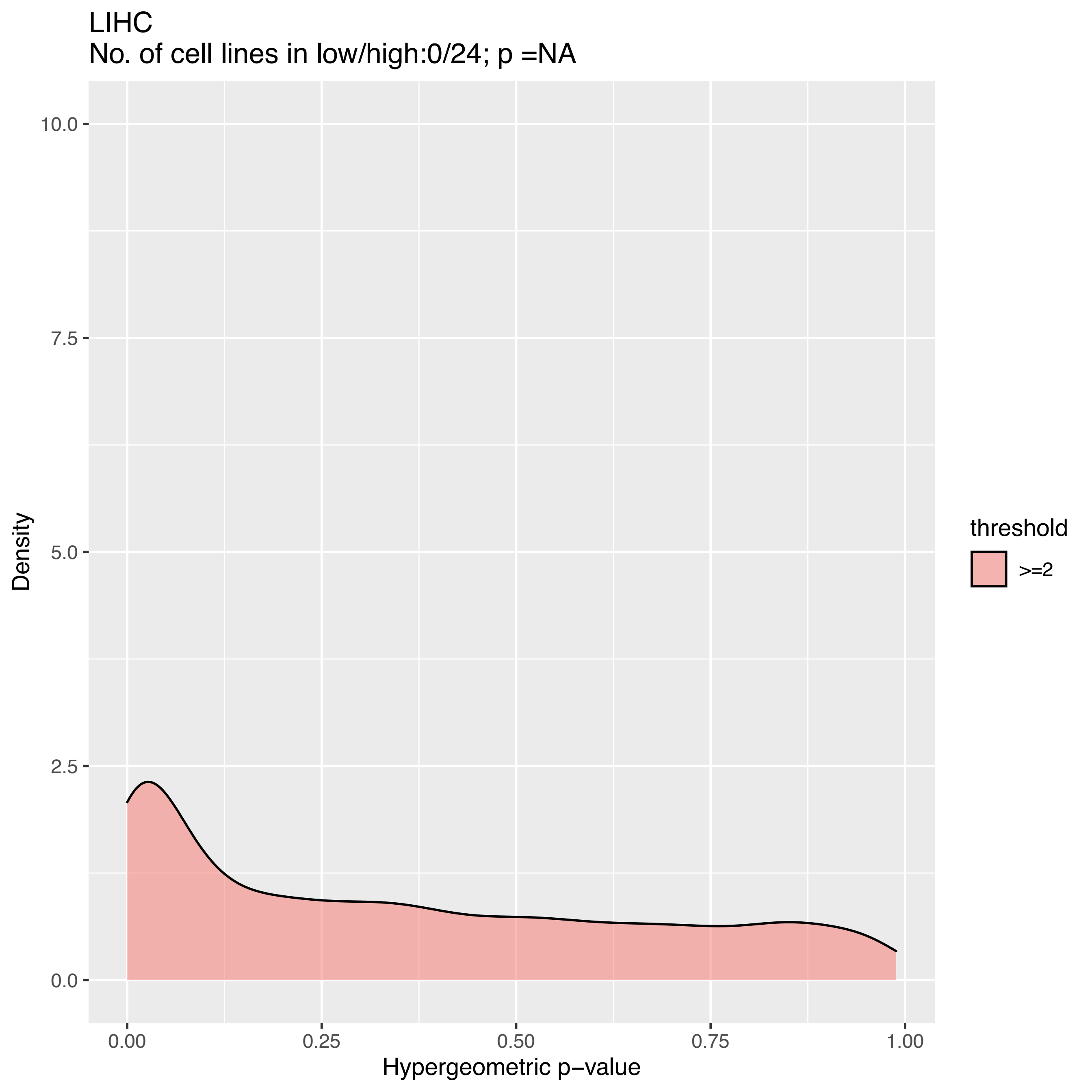
LIHC
No. of cell lines in low/high:0/24; p =NA
Density
10.0
7.5
5.0
2.5
0.0
0.00
0.25
0.50
0.75
1.00
Hypergeometric p−value
threshold
>=2

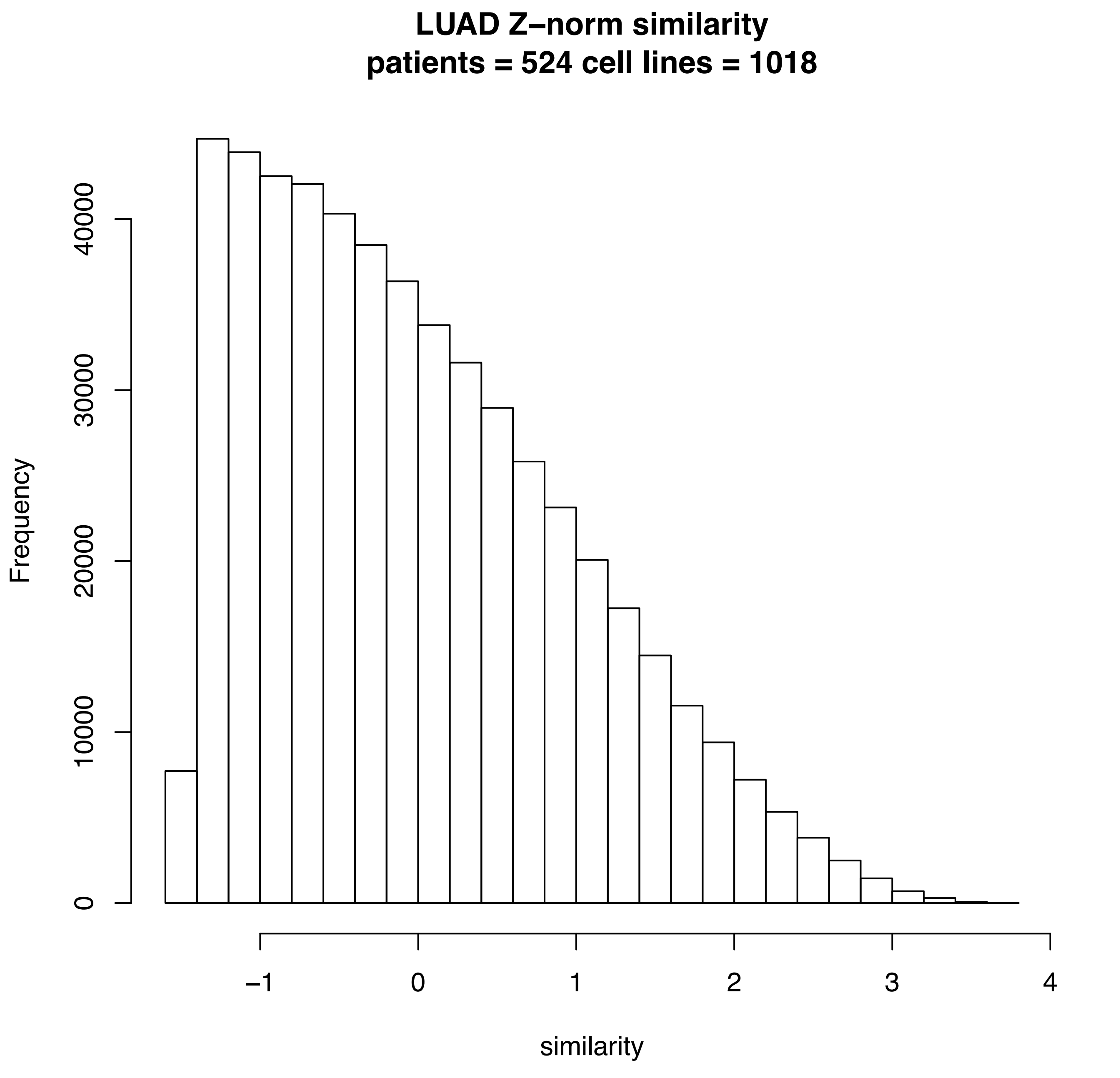
LUAD Z–norm similarity
patients = 524 cell lines = 1018
Frequency
0
10000
20000
30000
40000
–1
0
1
2
3
4
similarity

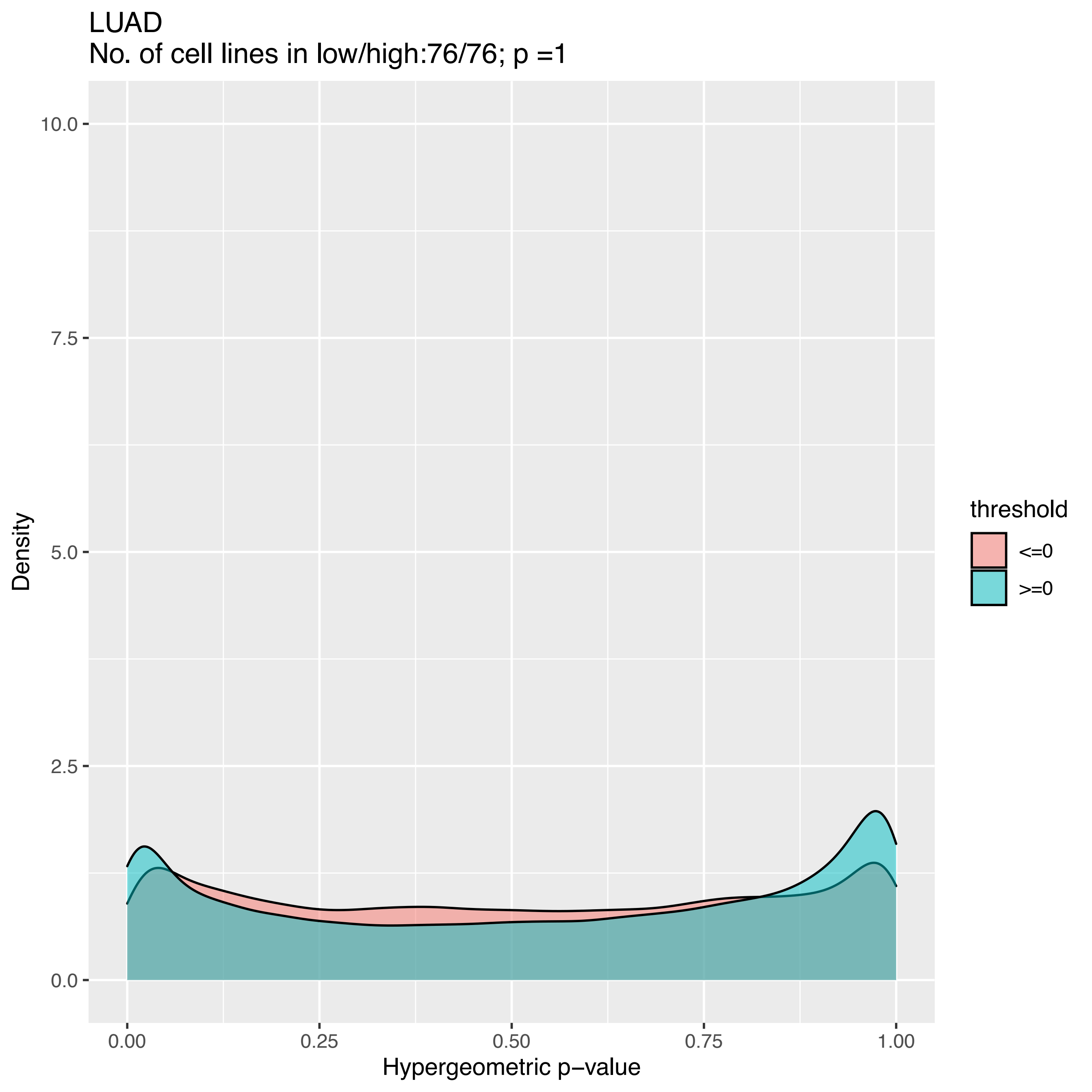
LUAD
No. of cell lines in low/high:76/76; p =1
10.0
7.5
5.0
2.5
0.0
Density
0.00
0.25
0.50
0.75
1.00
Hypergeometric p−value
threshold
<=0
>=0

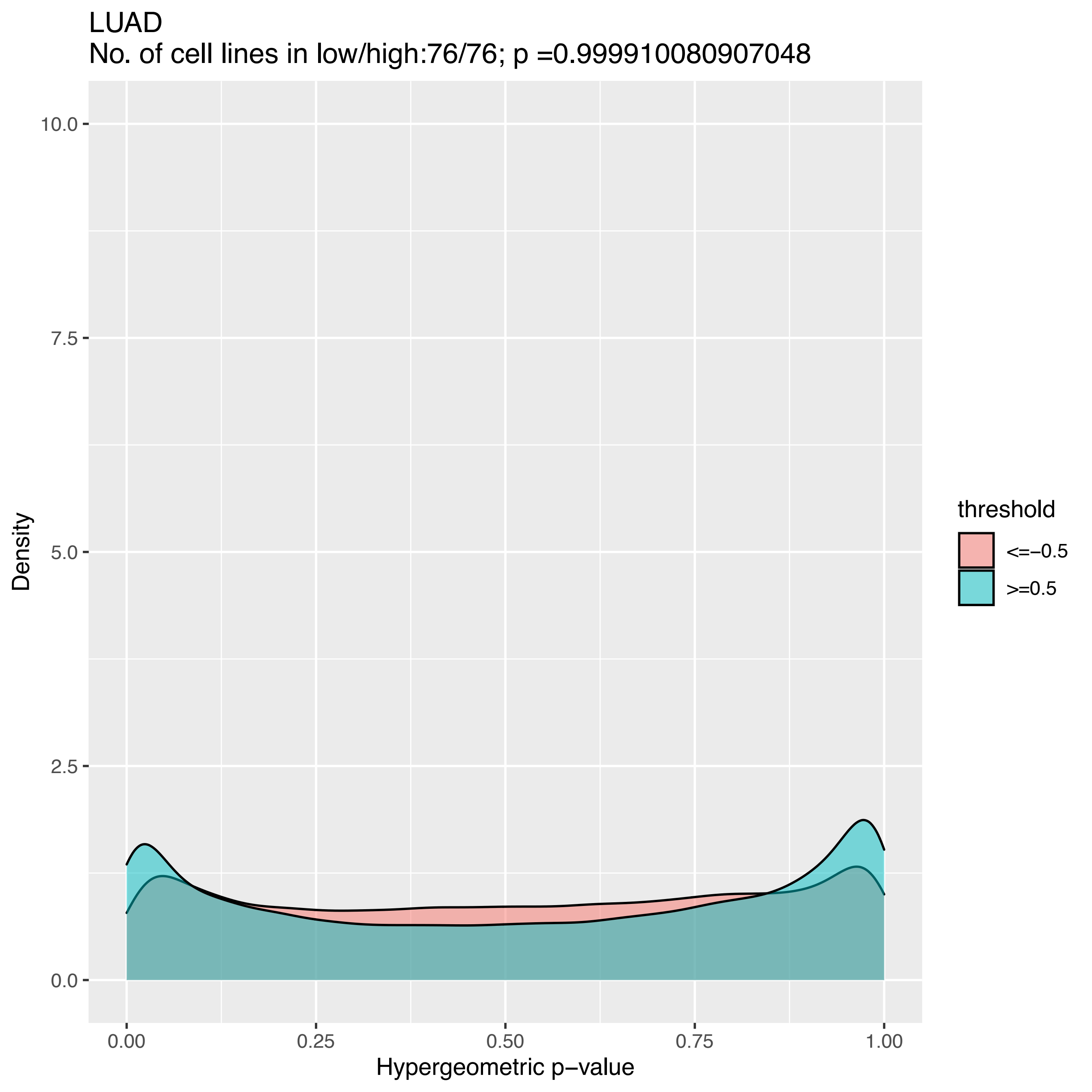
LUAD
No. of cell lines in low/high:76/76; p =0.999910080907048
Density
10.0
7.5
5.0
2.5
0.0
0.00
0.25
0.50
0.75
1.00
Hypergeometric p−value
threshold
<=−0.5
>=0.5

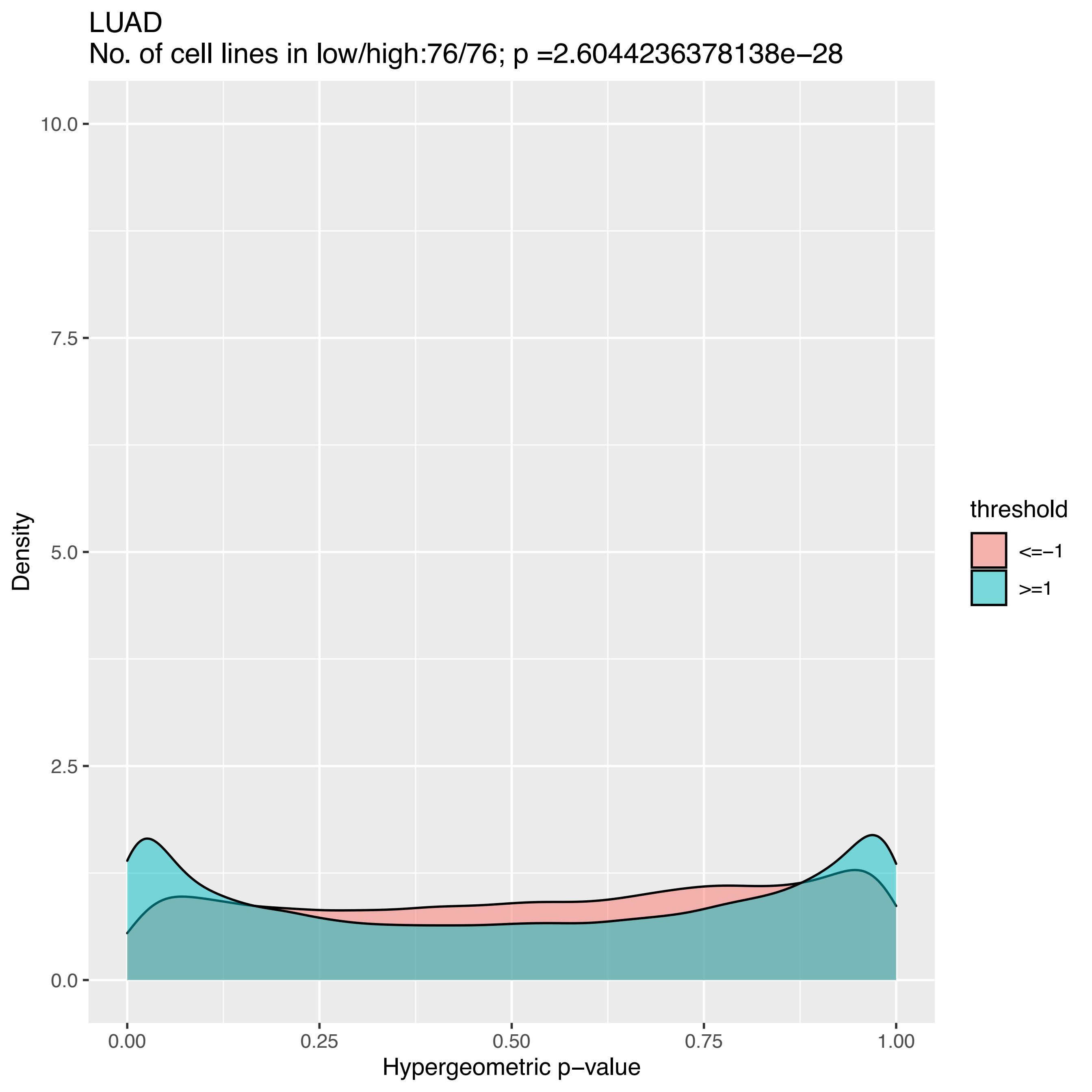
LUAD
No. of cell lines in low/high:76/76; p =2.6044236378138e−28
10.0
7.5
5.0
2.5
0.0
Density
0.00
0.25
0.50
0.75
1.00
Hypergeometric p−value
threshold
<=−1
>=1

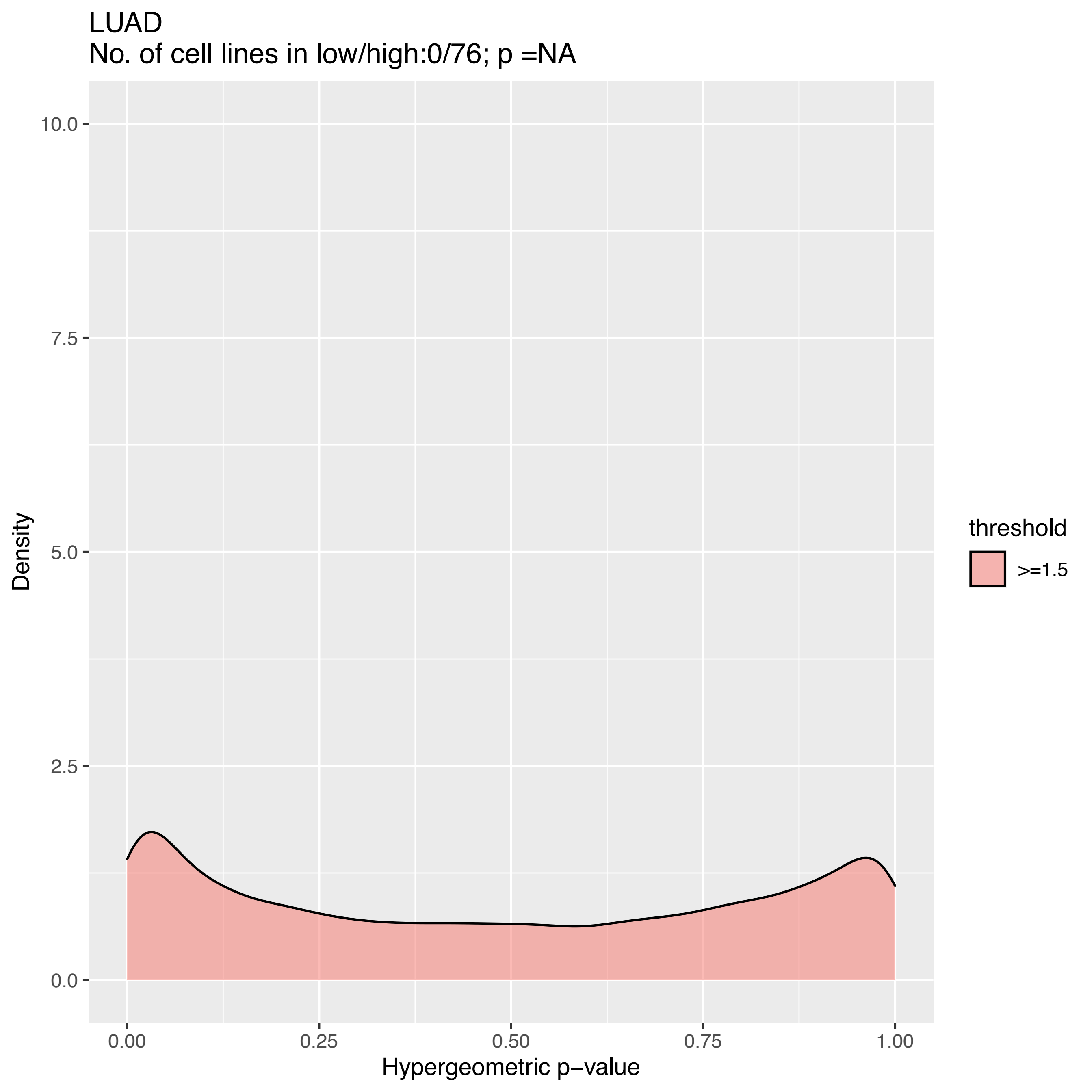

LUAD
No. of cell lines in low/high:0/76; p =NA
10.0
7.5
5.0
2.5
0.0
Density
0.00
0.25
0.50
0.75
1.00
Hypergeometric p−value
threshold
>=1.5

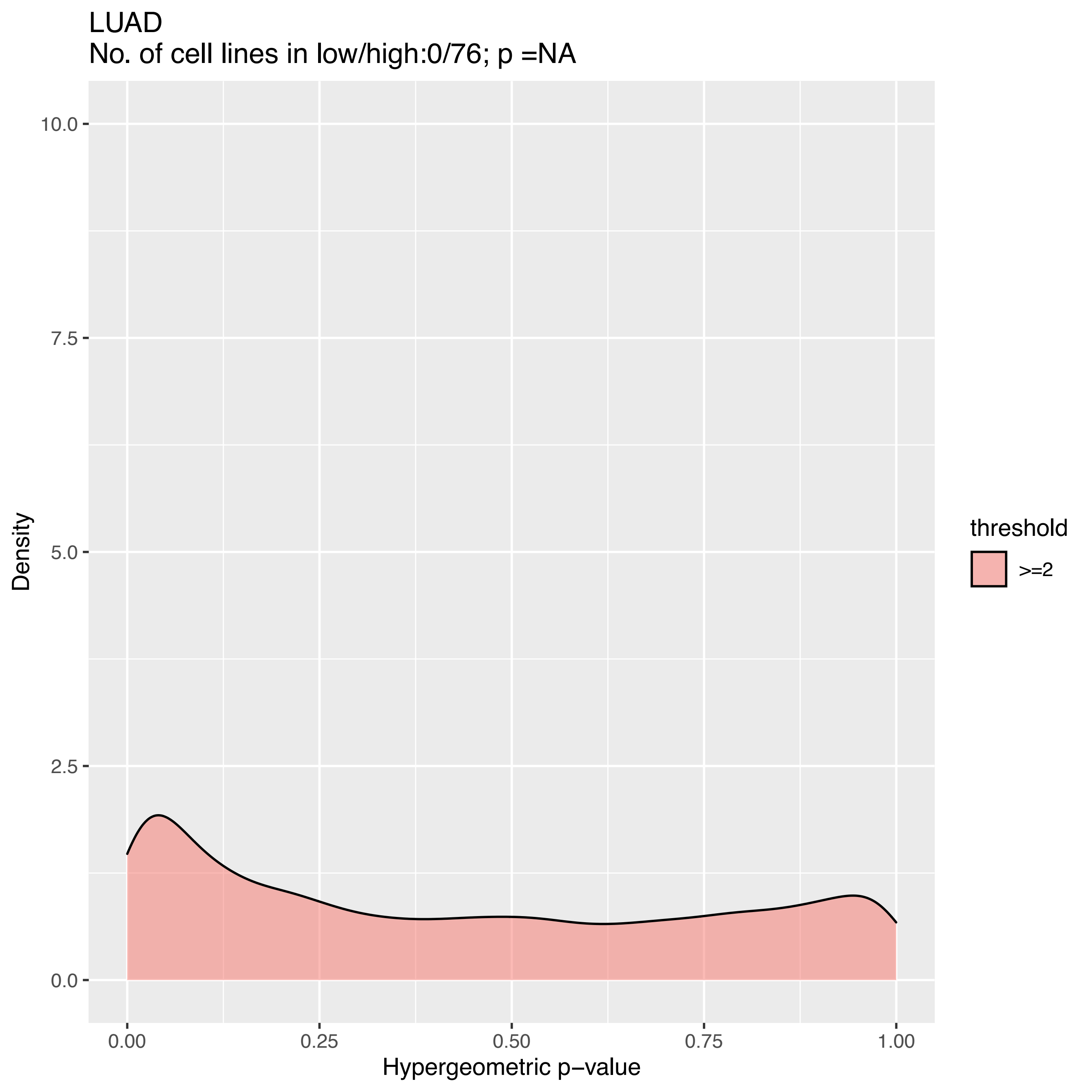

LUAD
No. of cell lines in low/high:0/76; p =NA
Density
10.0
7.5
5.0
2.5
0.0
0.00
0.25
0.50
0.75
1.00
Hypergeometric p−value
threshold
>=2

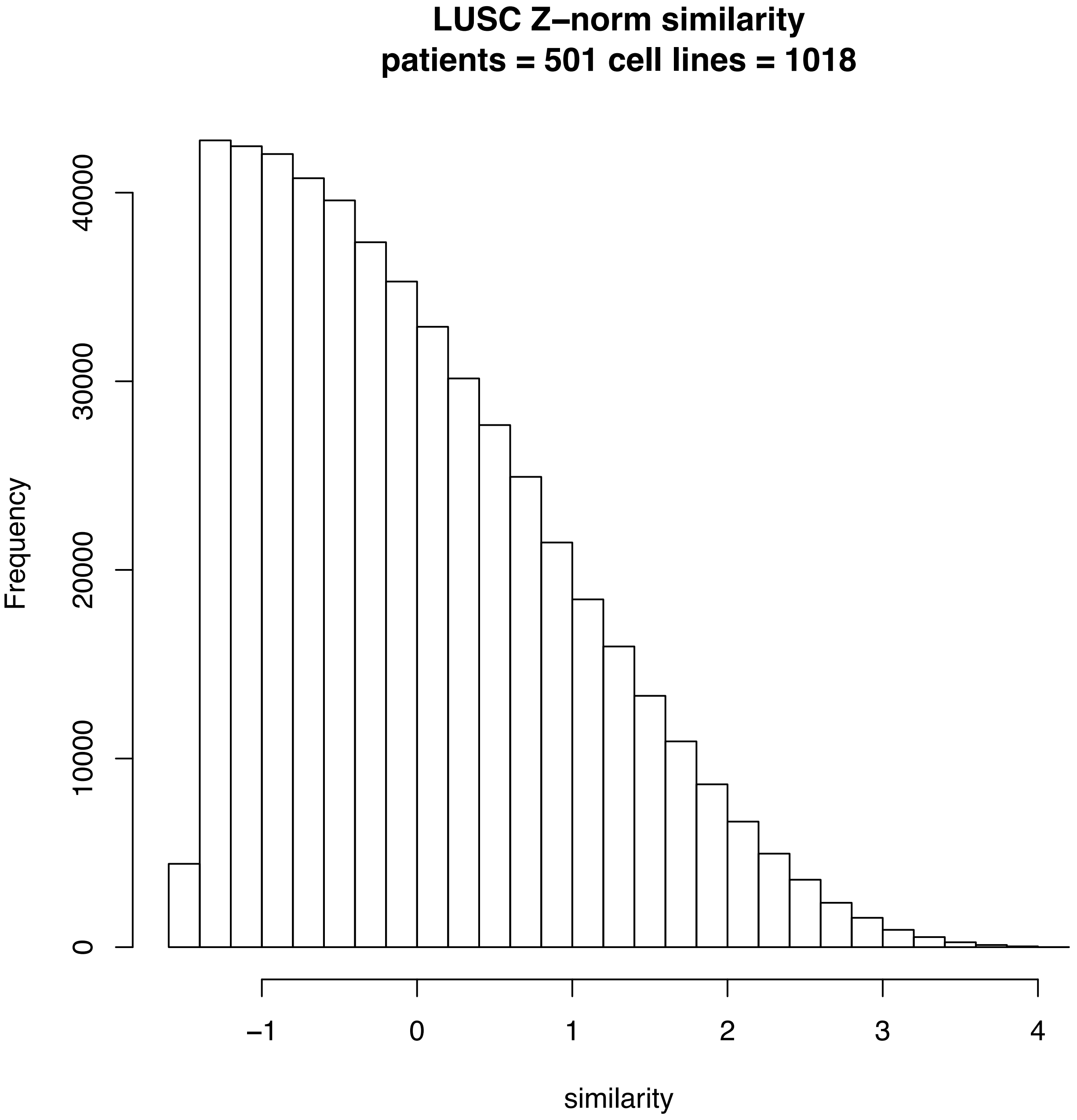

LUSC Z–norm similarity
patients = 501 cell lines = 1018
Frequency
0
10000
20000
30000
40000
−1
0
1
2
3
4
similarity

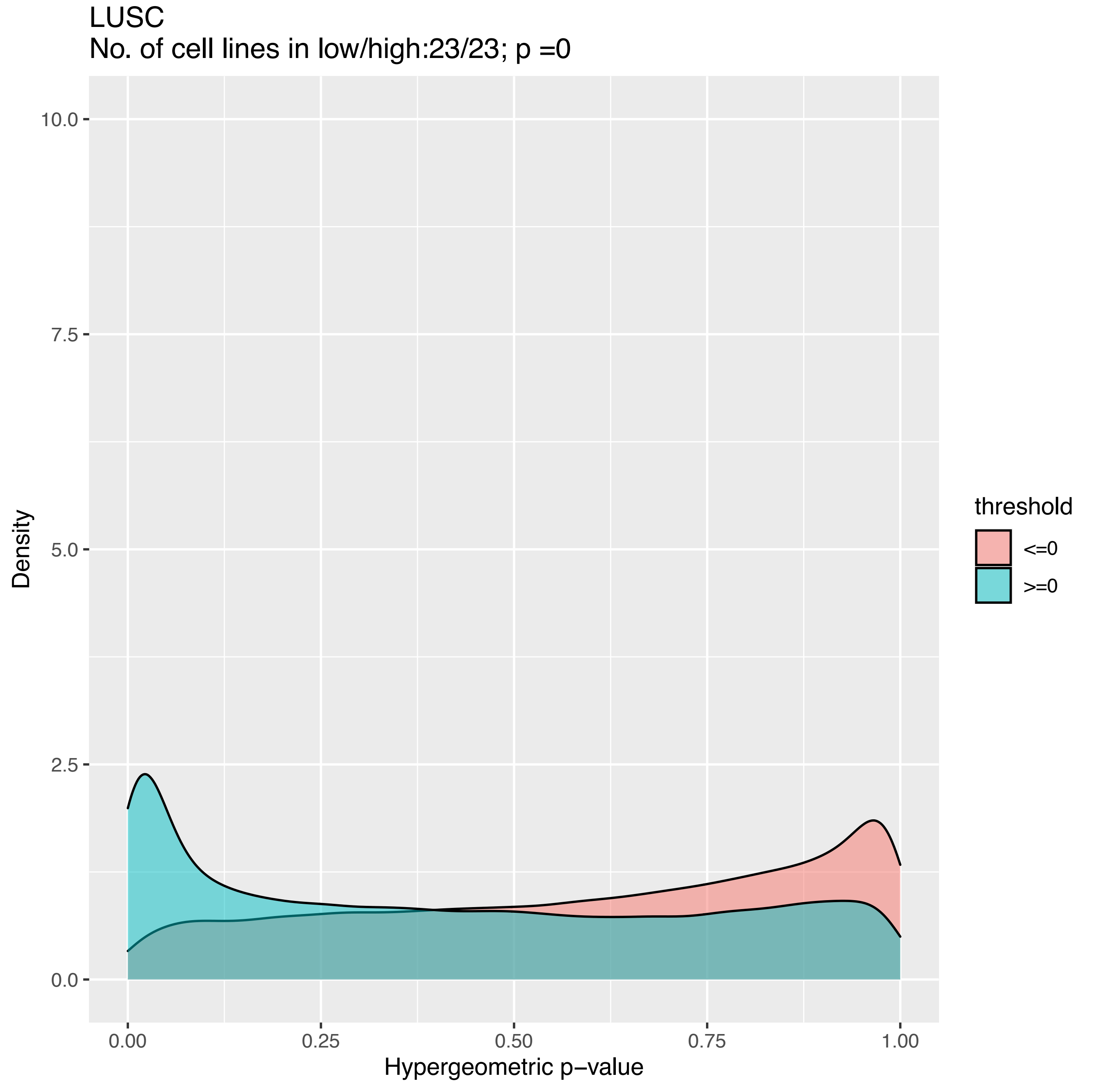

LUSC
No. of cell lines in low/high:23/23; p =0
10.0
7.5
5.0
2.5
0.0
Density
0.00
0.25
0.50
0.75
1.00
Hypergeometric p−value
threshold
<=0
>=0

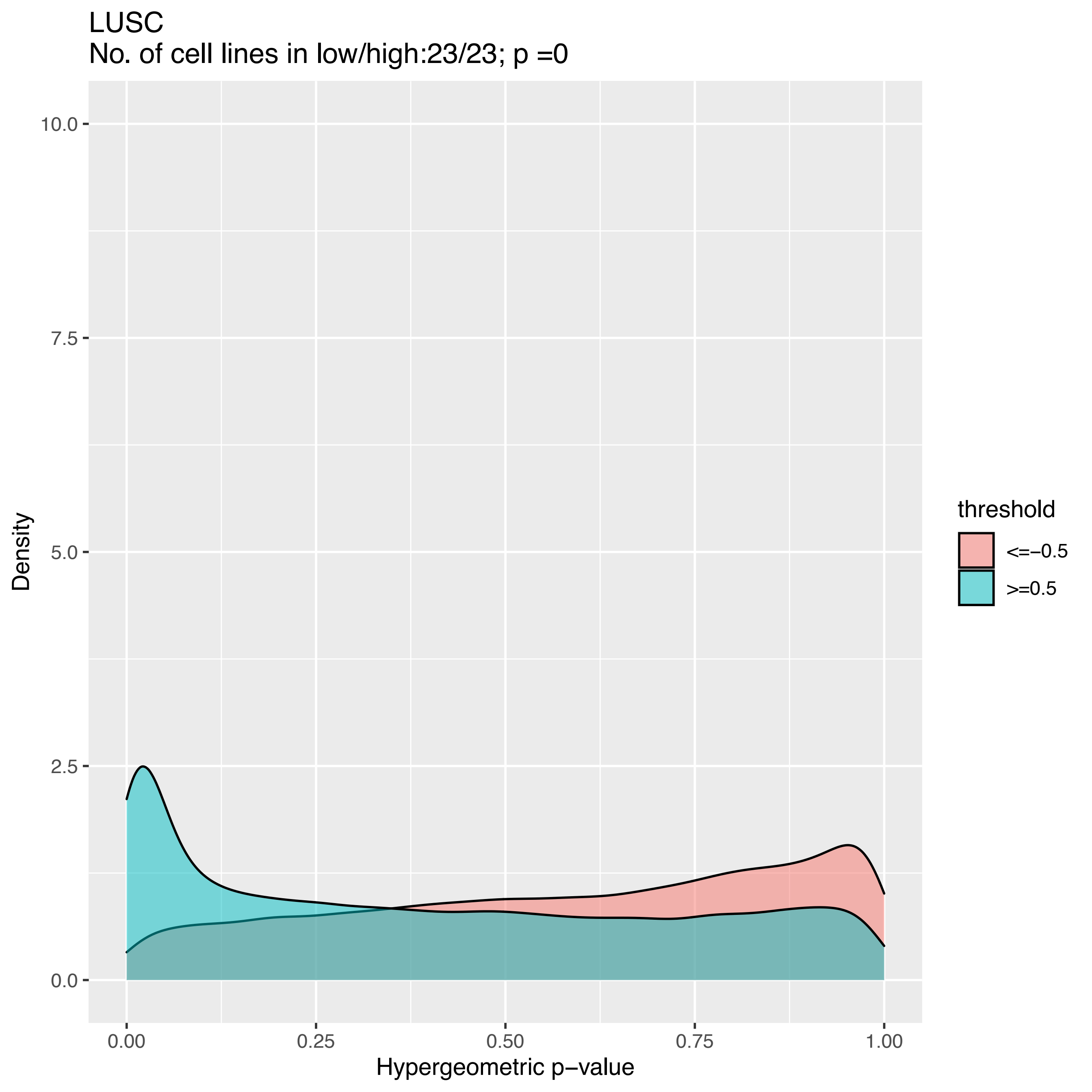
LUSC
No. of cell lines in low/high:23/23; p =0
10.0
7.5
5.0
2.5
0.0
Density
0.00
0.25
0.50
0.75
1.00
Hypergeometric p−value
threshold
<=−0.5
>=0.5

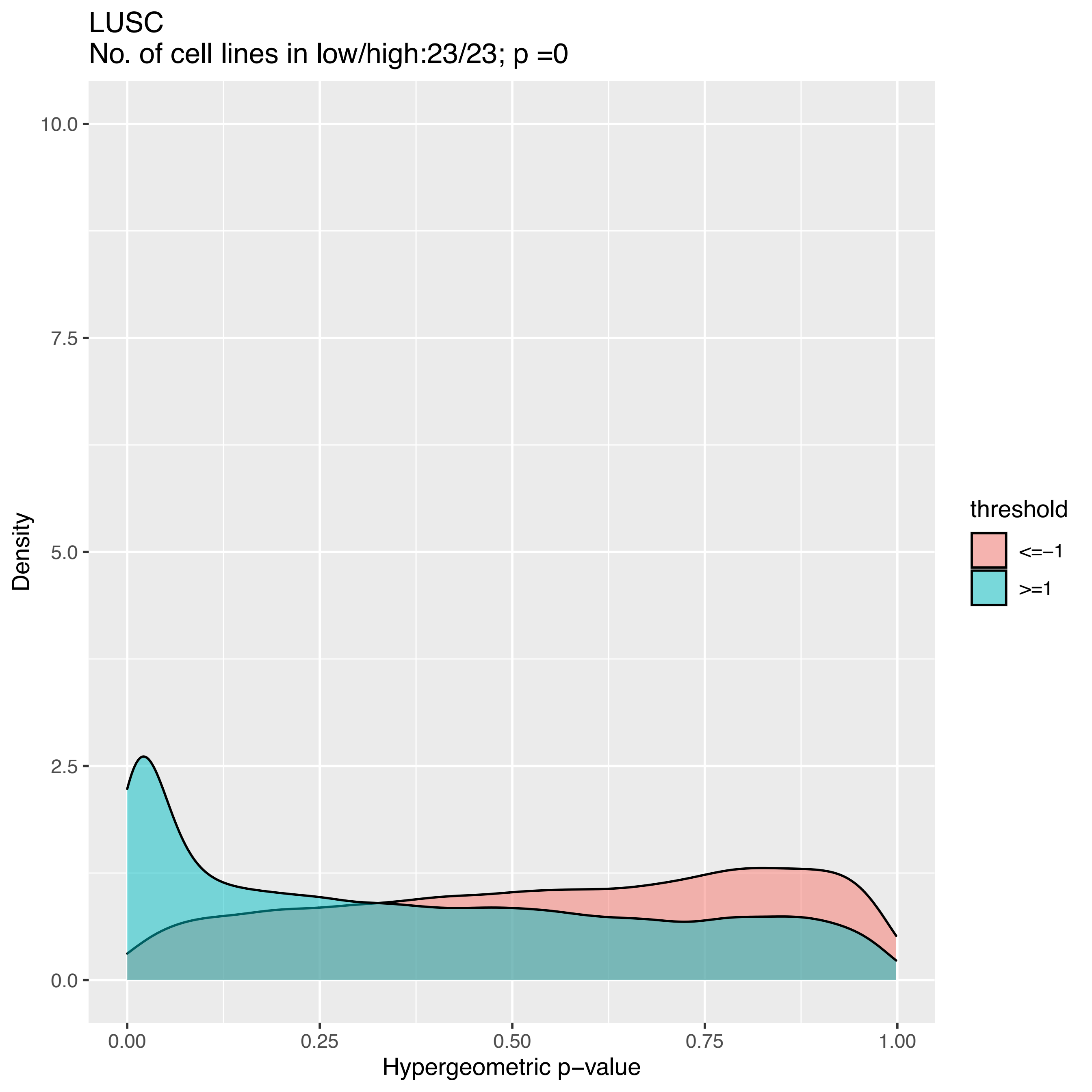

LUSC
No. of cell lines in low/high:23/23; p =0
10.0
7.5
5.0
2.5
0.0
Density
0.00
0.25
0.50
0.75
1.00
Hypergeometric p−value
threshold
<=−1
>=1

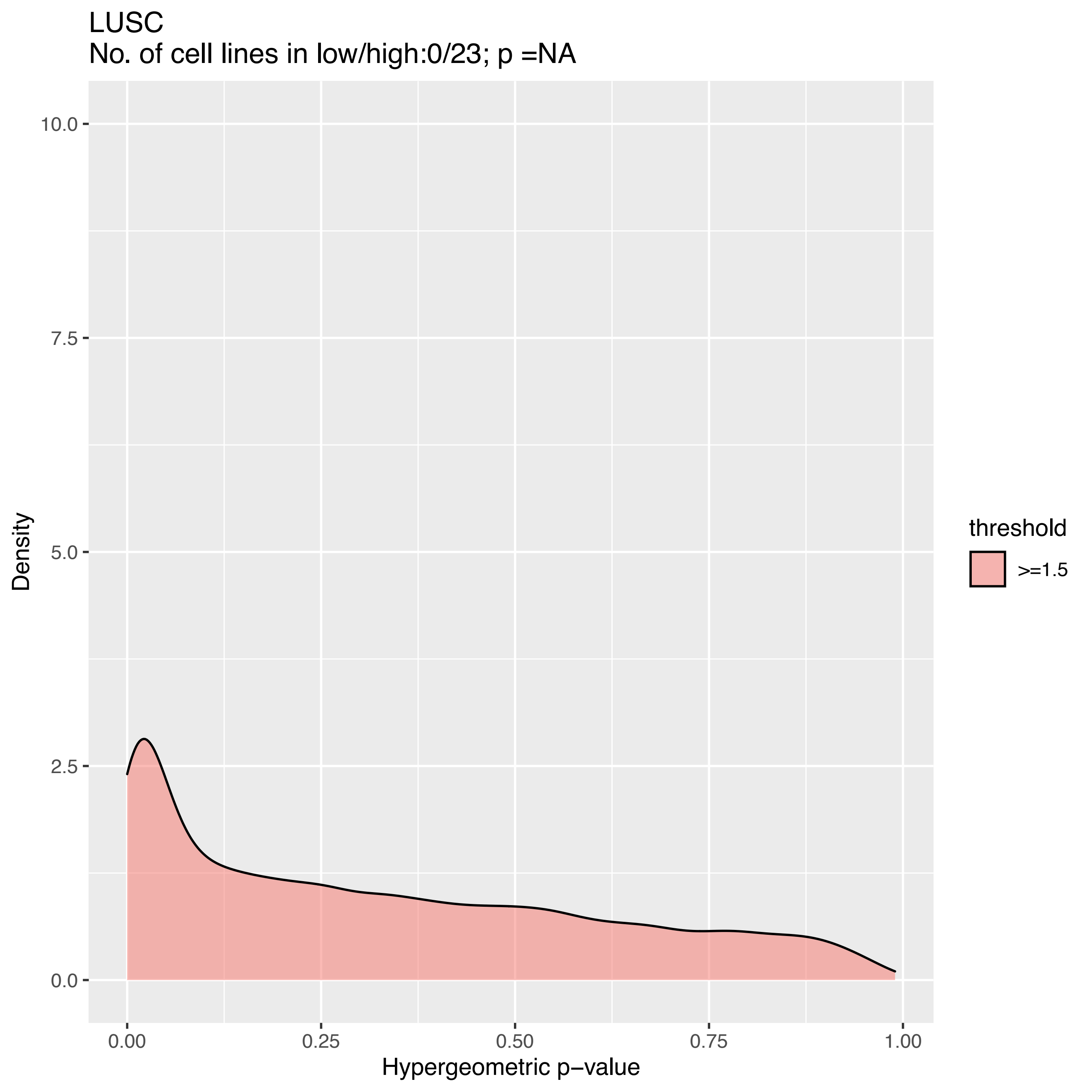

LUSC
No. of cell lines in low/high:0/23; p =NA
10.0
7.5
5.0
2.5
0.0
Density
0.00
0.25
0.50
0.75
1.00
Hypergeometric p−value
threshold
>=1.5

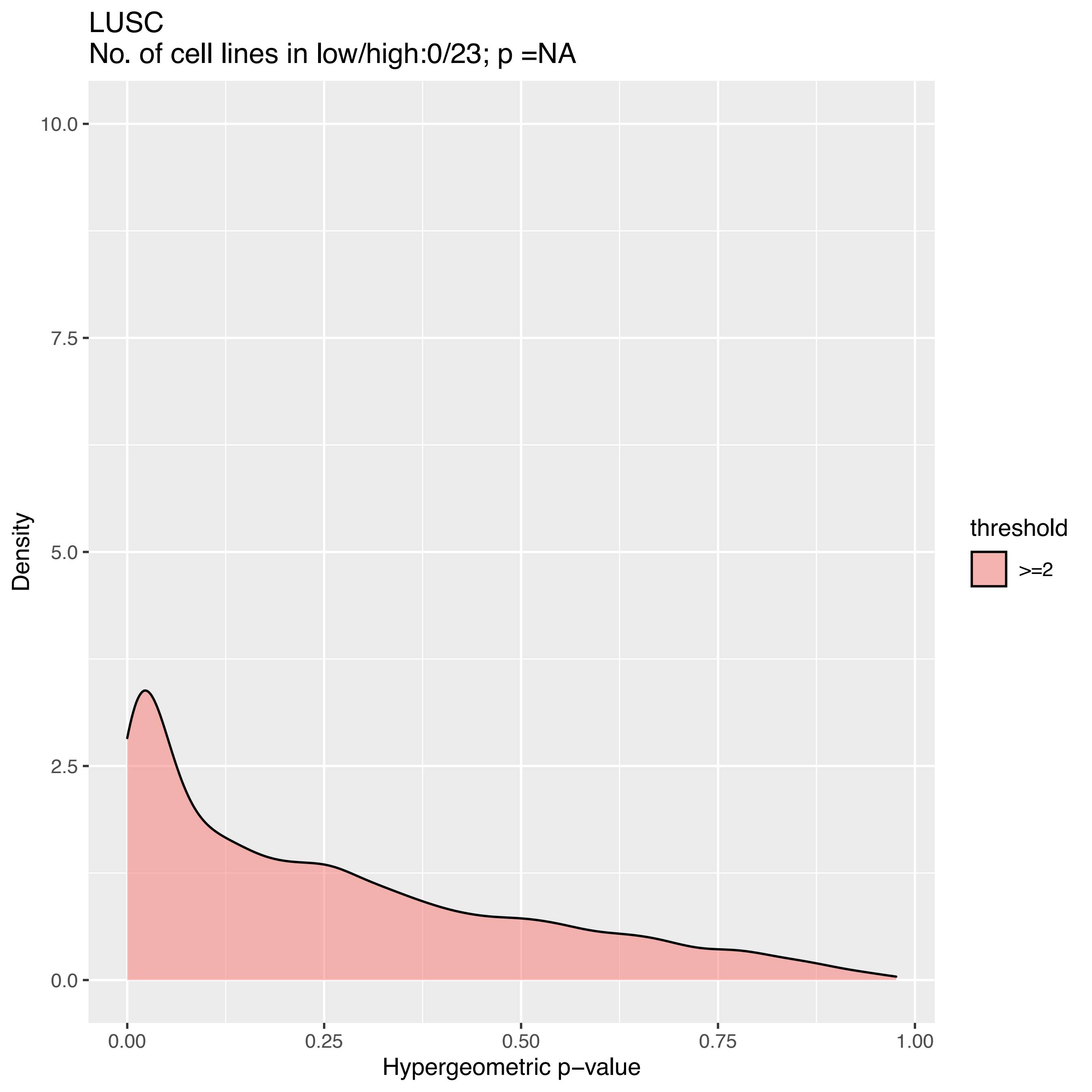
LUSC
No. of cell lines in low/high:0/23; p =NA
10.0
7.5
5.0
2.5
0.0
Density
0.00
0.25
0.50
0.75
1.00
Hypergeometric p−value
threshold
>=2

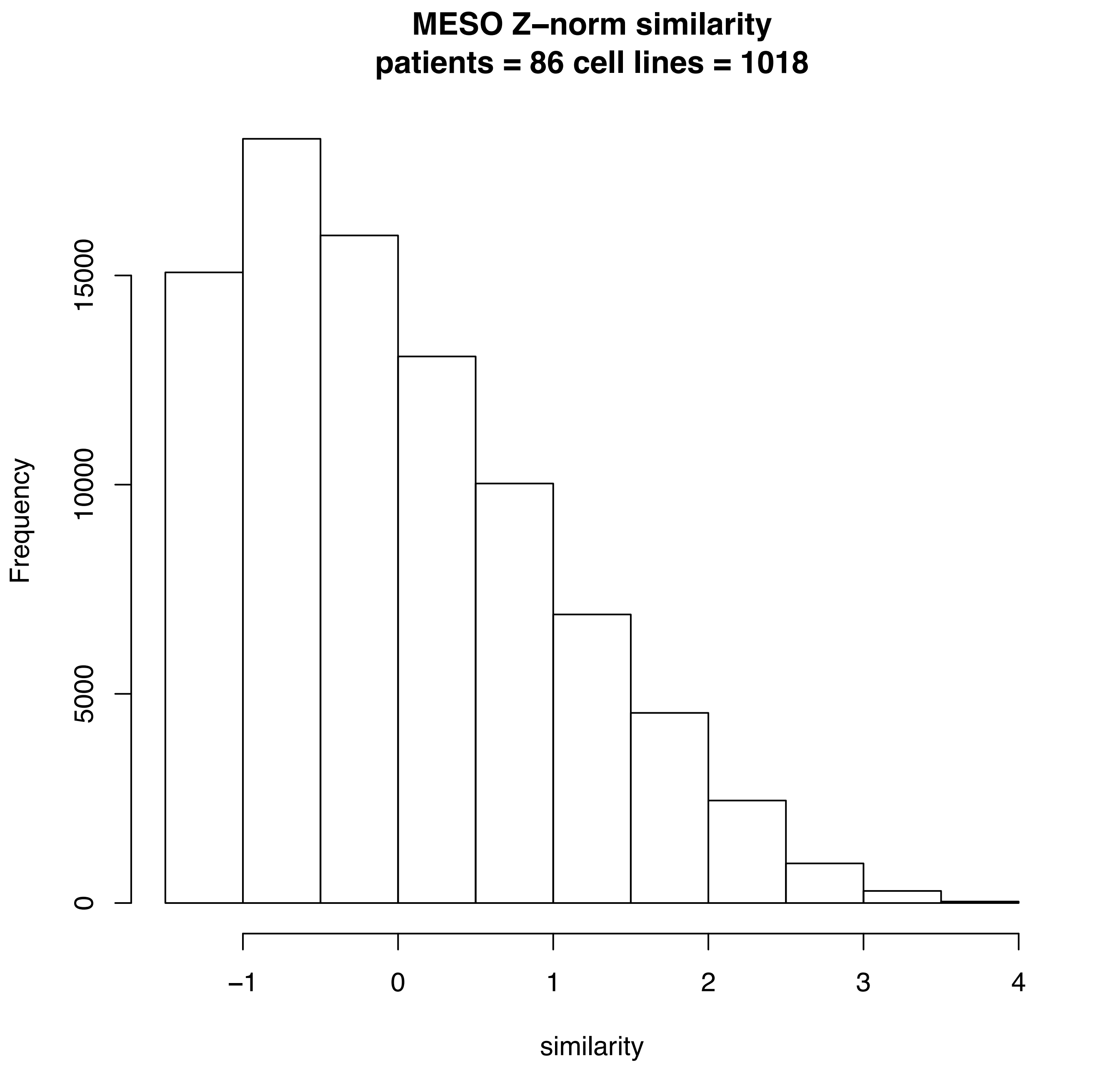
MESO Z–norm similarity
patients = 86 cell lines = 1018
Frequency
0
5000
10000
15000
−1
0
1
2
3
4
similarity

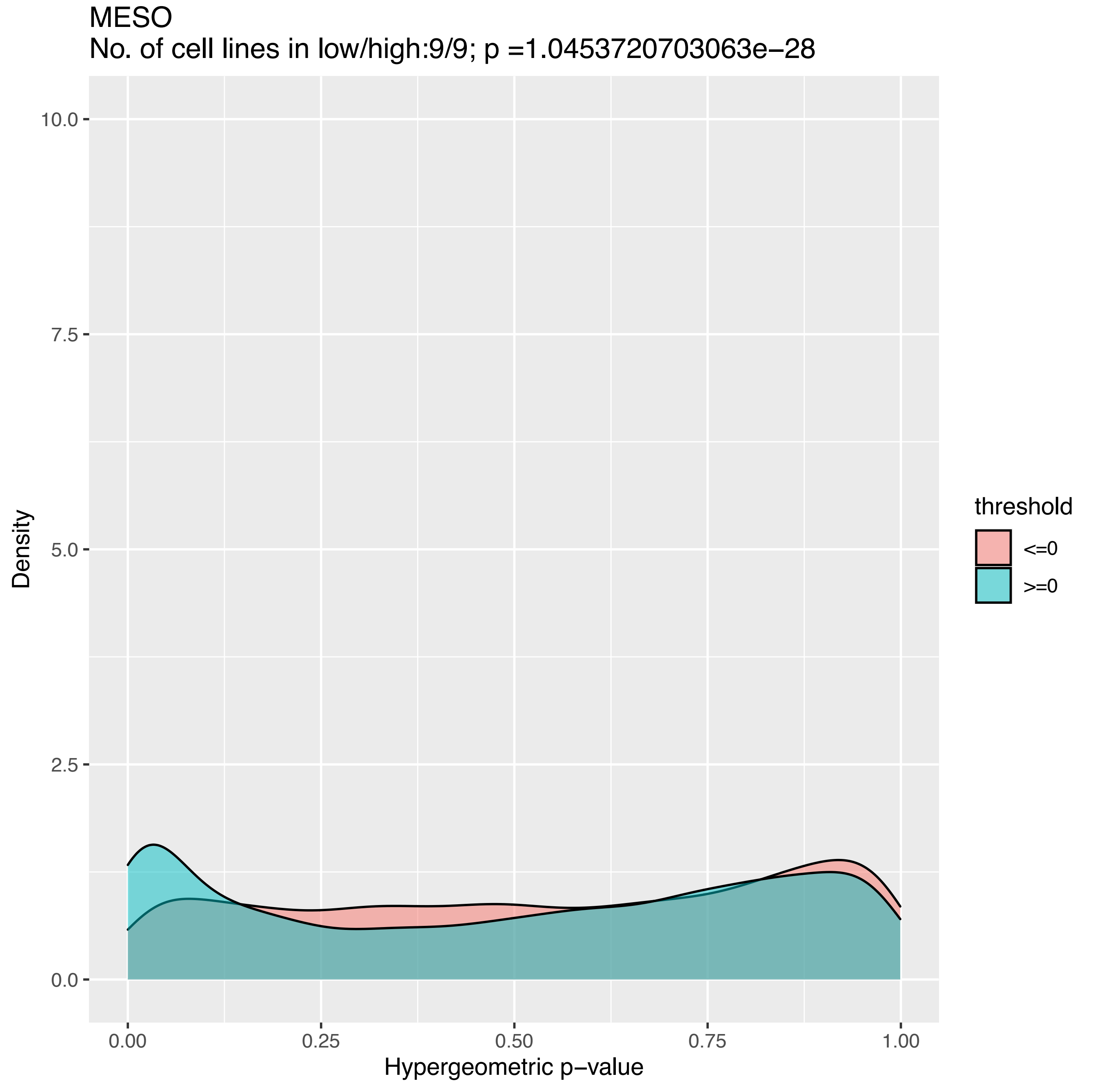
MESO
No. of cell lines in low/high:9/9; p =1.0453720703063e−28
10.0
7.5
5.0
2.5
0.0
Density
0.00
0.25
0.50
0.75
1.00
Hypergeometric p−value
threshold
<=0
>=0

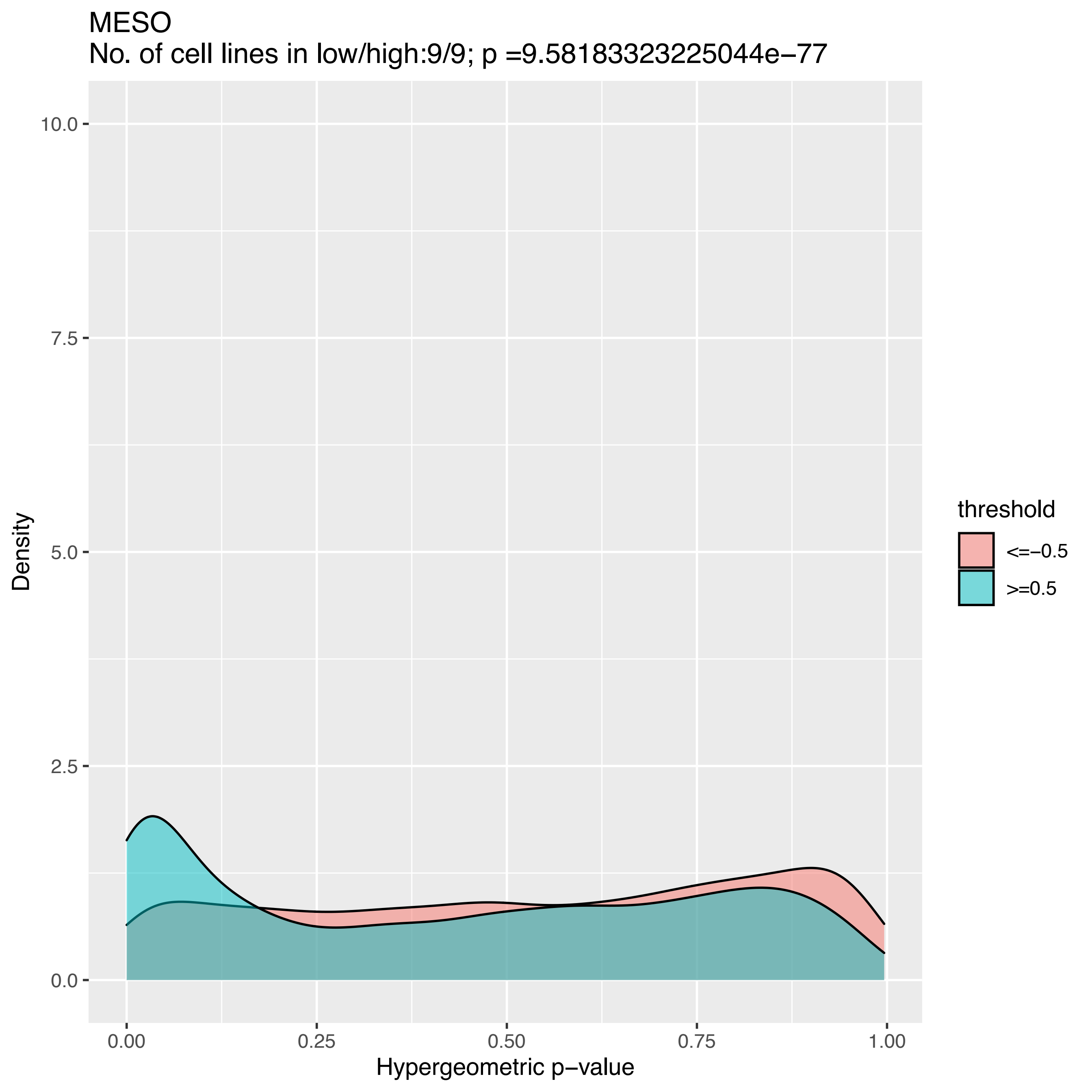

MESO
No. of cell lines in low/high:9/9; p =9.58183323225044e−77
10.0
7.5
5.0
2.5
0.0
Density
0.00
0.25
0.50
0.75
1.00
Hypergeometric p−value
threshold
<=−0.5
>=0.5

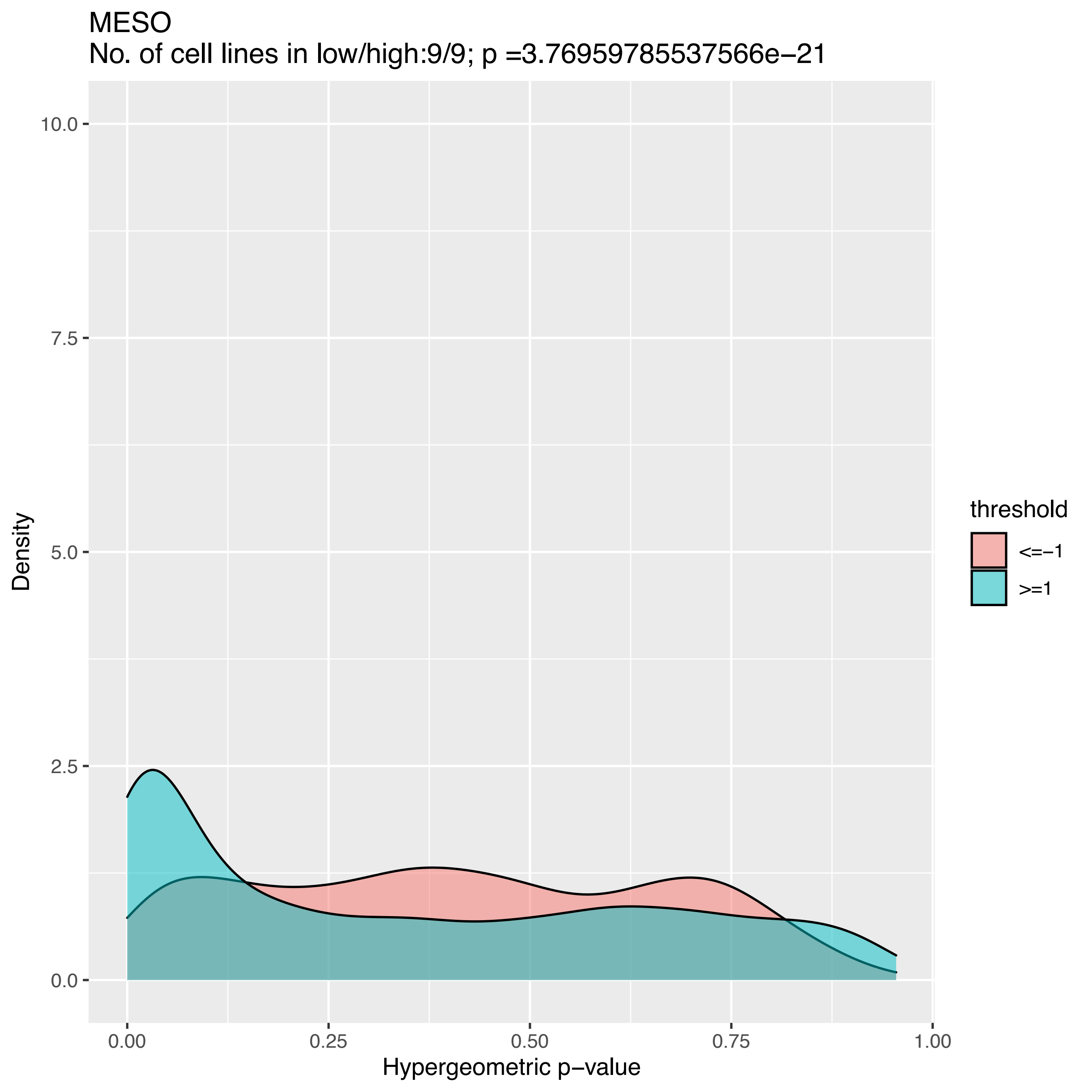

MESO
No. of cell lines in low/high:9/9; p =3.76959785537566e−21
10.0
7.5
5.0
2.5
0.0
Density
0.00
0.25
0.50
0.75
1.00
Hypergeometric p−value
threshold
<=−1
>=1

MESO
No. of cell lines in low/high:0/9; p =NA

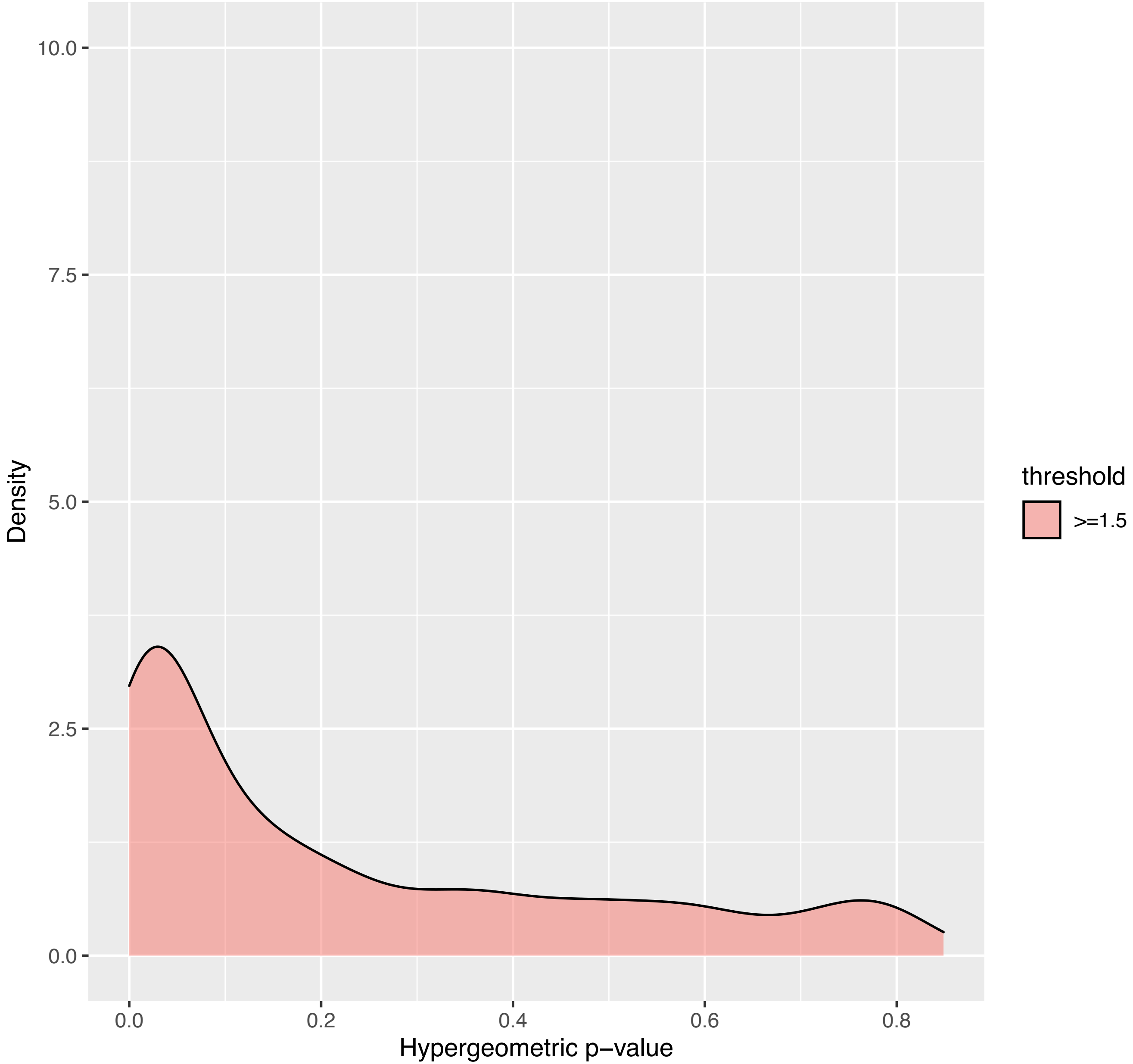

MESO
No. of cell lines in low/high:0/9; p =NA

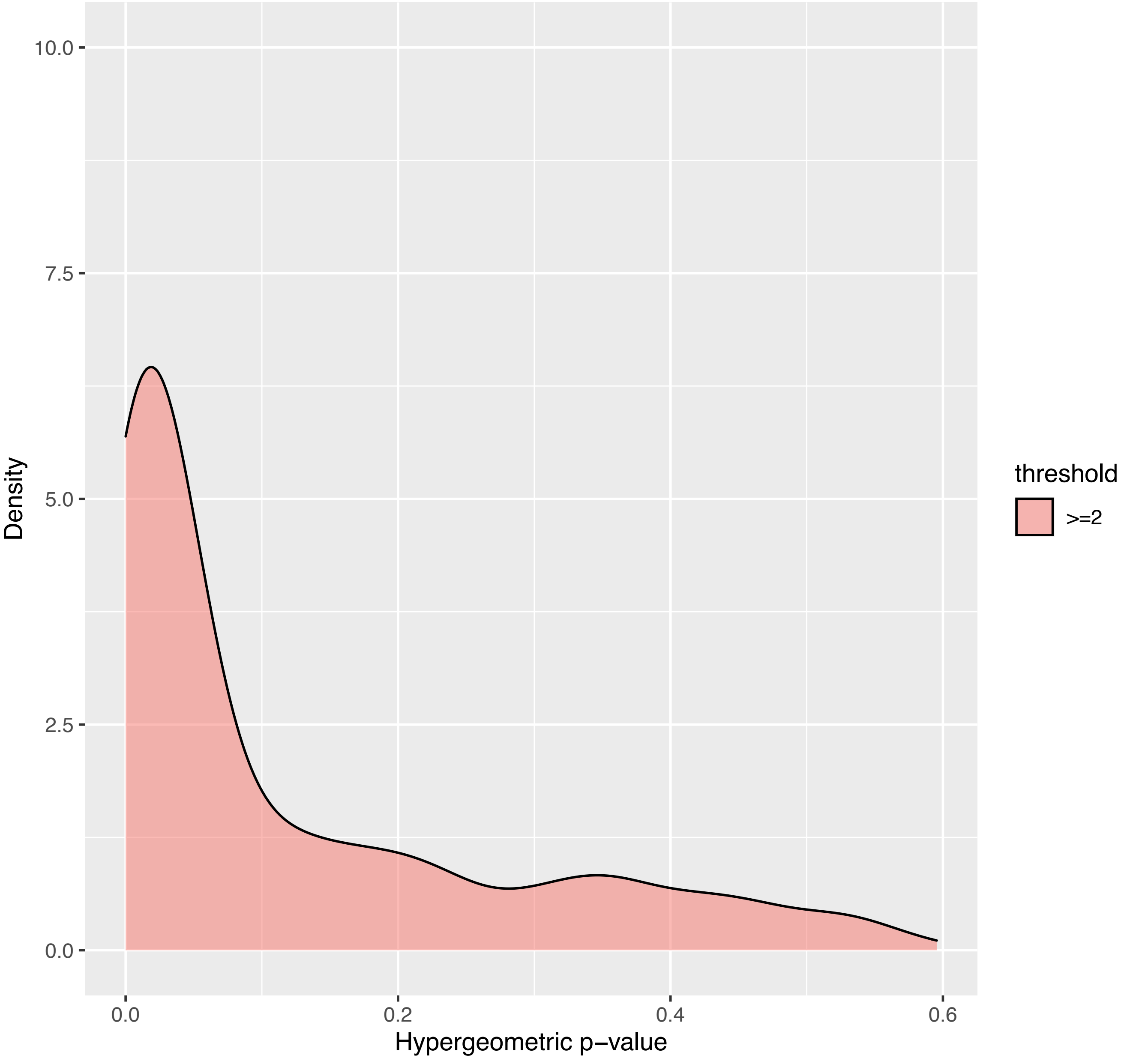

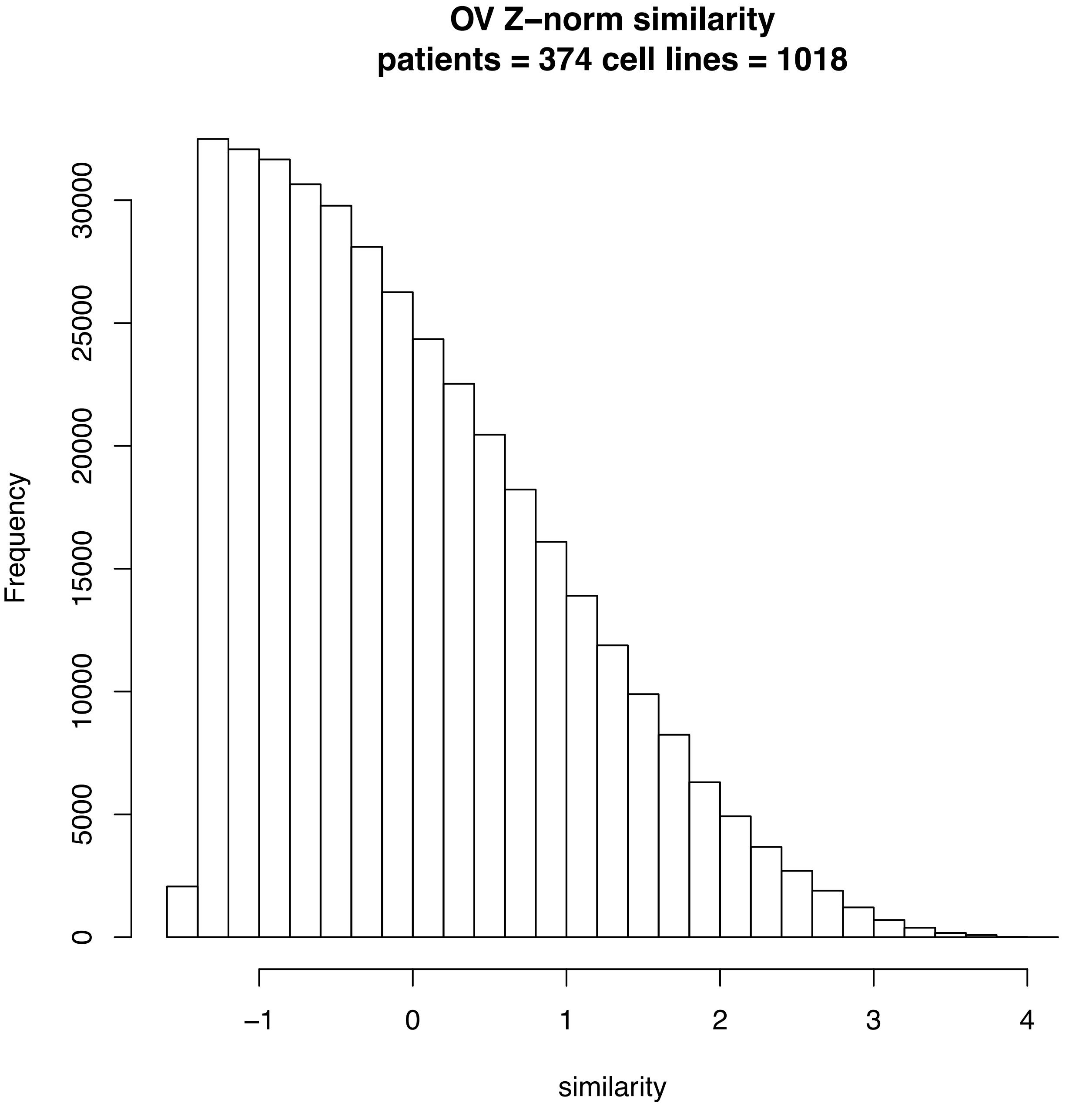
OV Z–norm similarity
patients = 374 cell lines = 1018
Frequency
0
5000
10000
15000
20000
25000
30000
–1
0
1
2
3
4
similarity

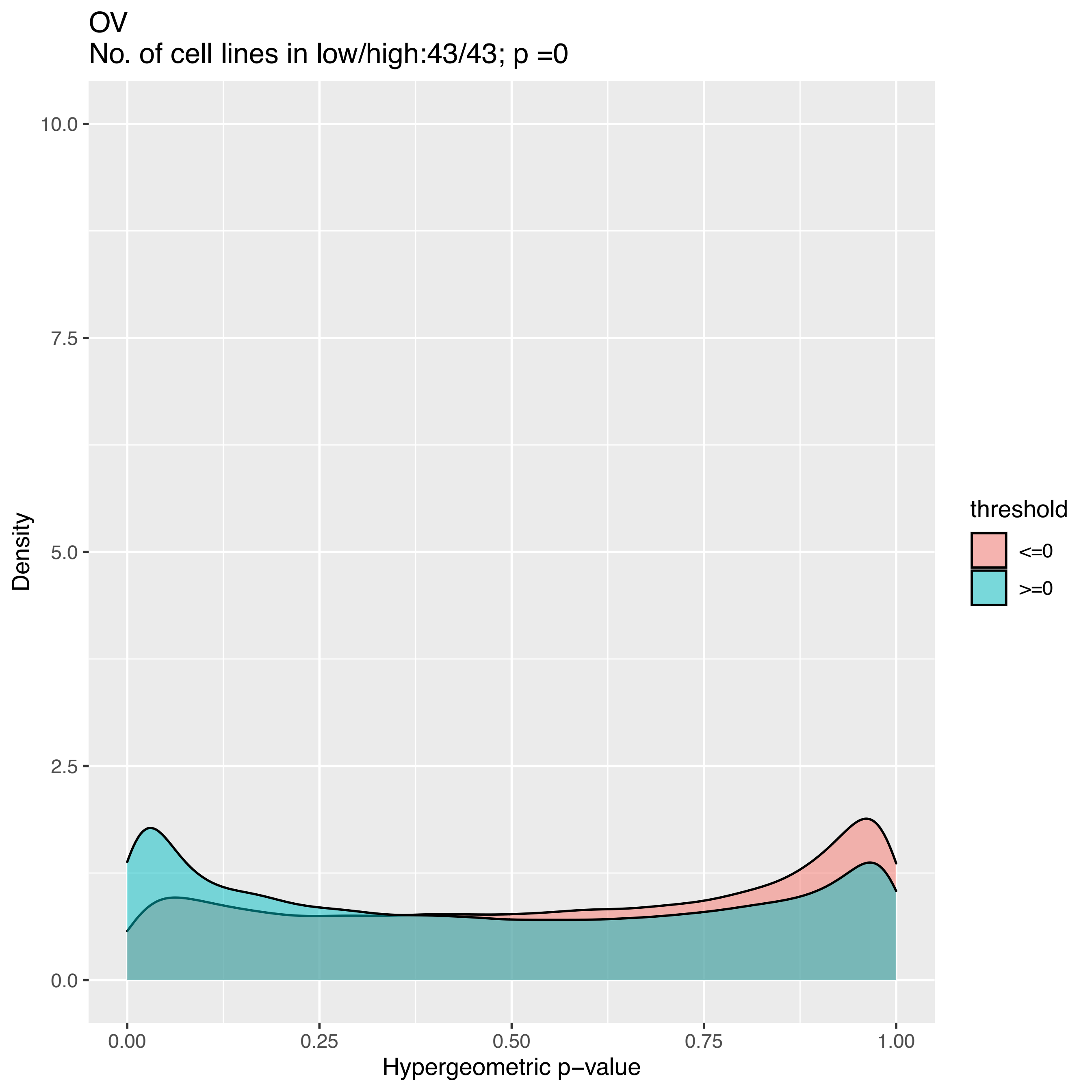
OV
No. of cell lines in low/high:43/43; p =0
Density
Hypergeometric p−value
threshold
<=0
>=0
0.0
2.5
5.0
7.5
10.0
0.00
0.25
0.50
0.75
1.00

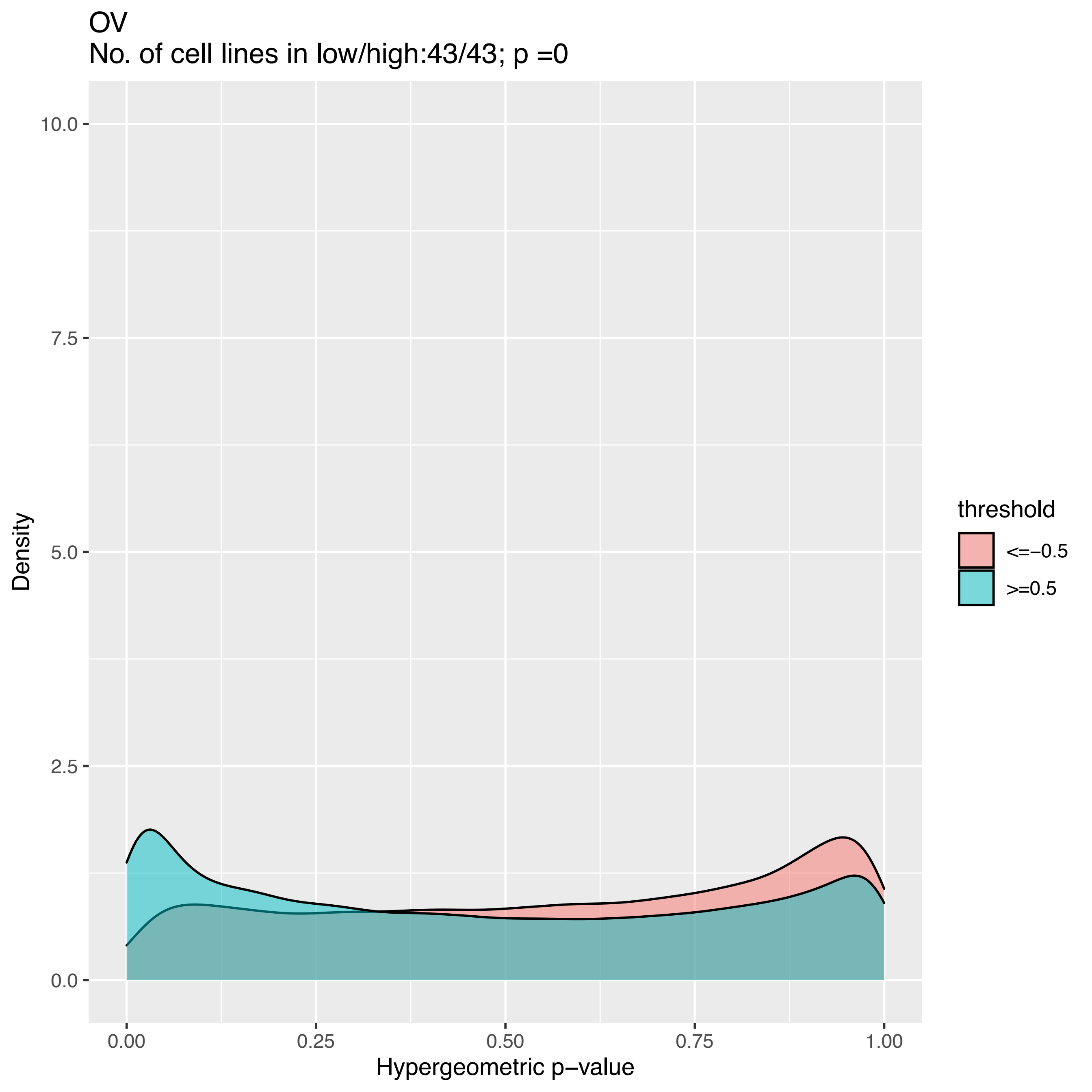
OV
No. of cell lines in low/high:43/43; p =0
Density
10.0
7.5
5.0
2.5
0.0
0.00
0.25
0.50
0.75
1.00
Hypergeometric p−value
threshold
<=−0.5
>=0.5

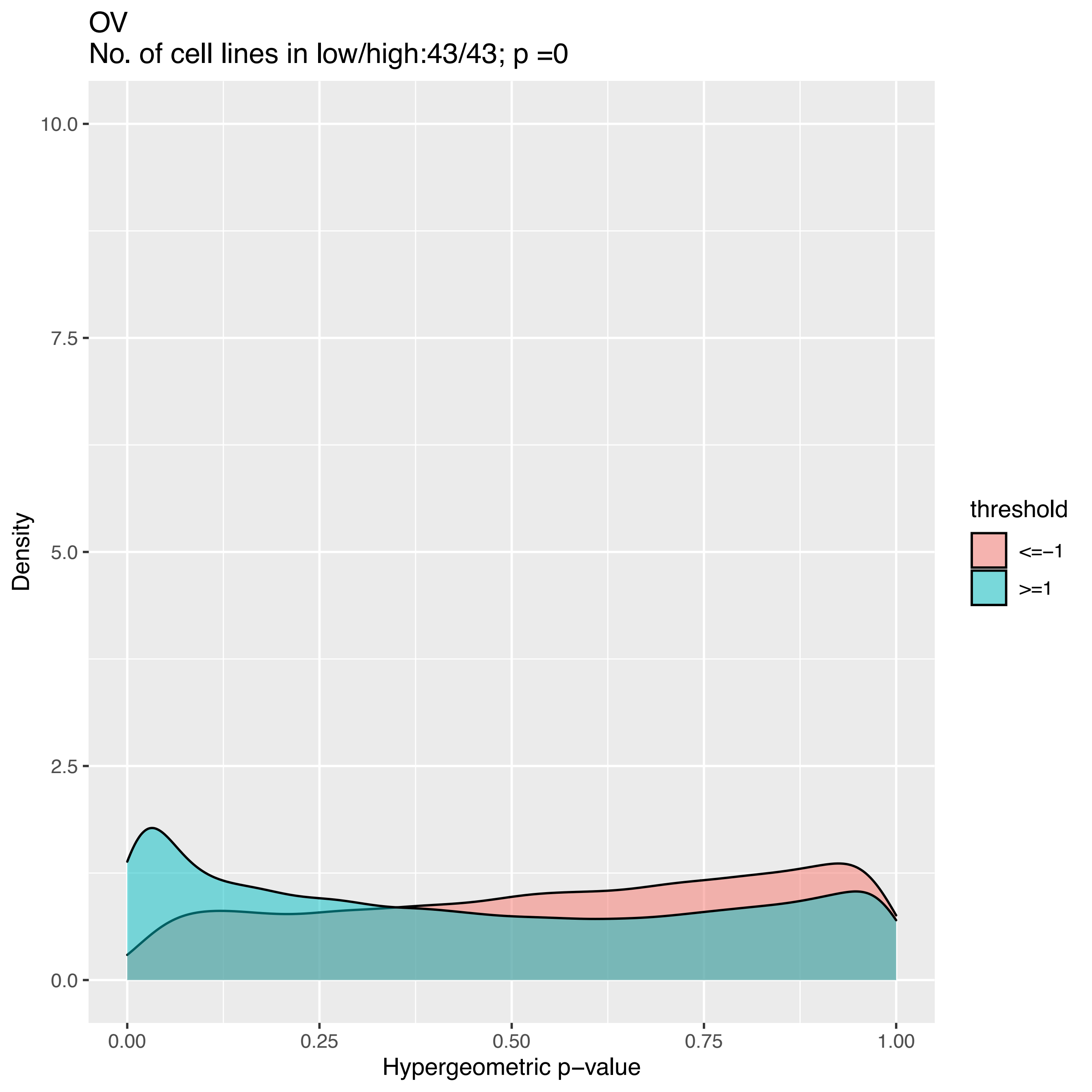
OV
No. of cell lines in low/high:43/43; p =0
10.0
7.5
5.0
2.5
0.0
Density
0.00
0.25
0.50
0.75
1.00
Hypergeometric p−value
threshold
<=−1
>=1

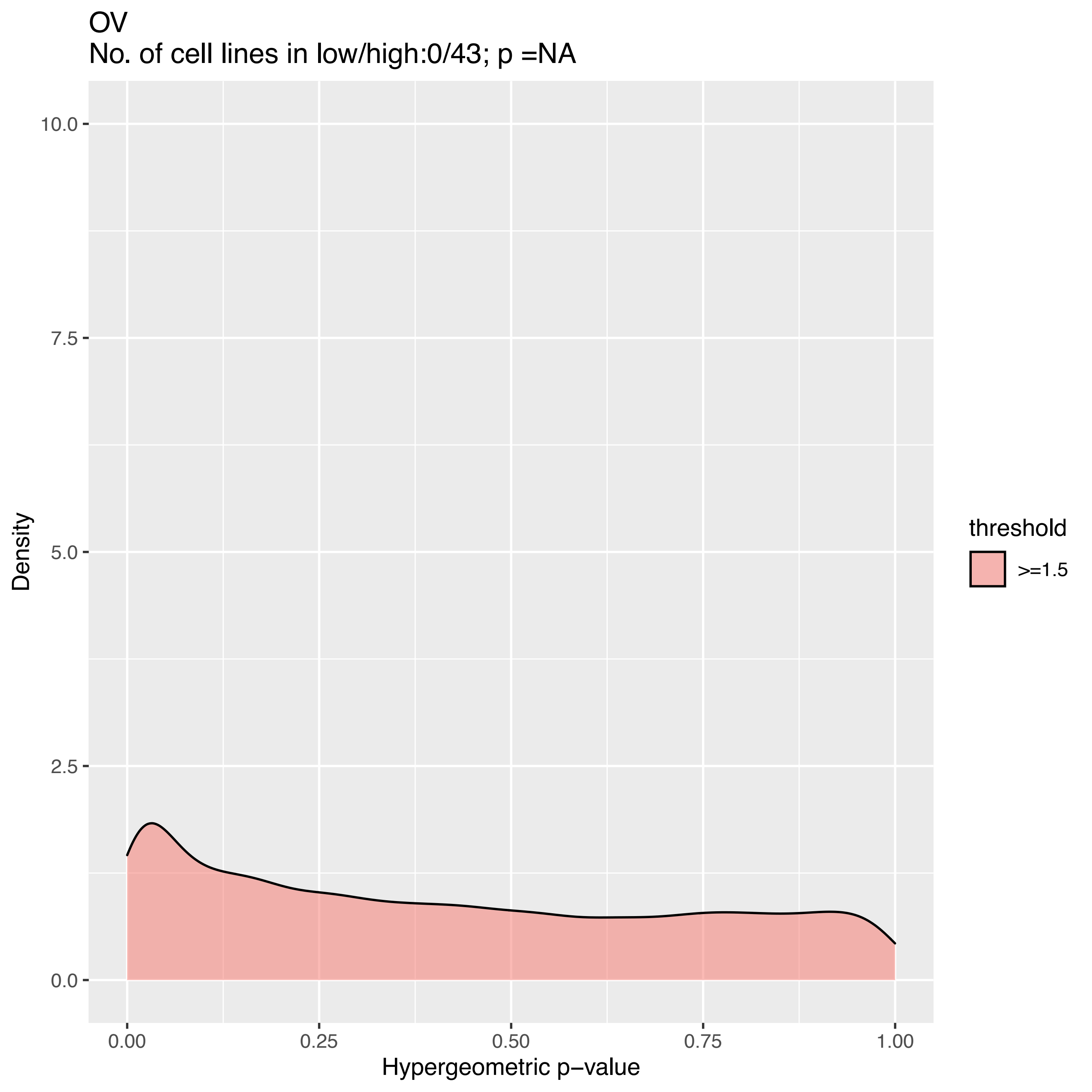
OV
No. of cell lines in low/high:0/43; p =NA
10.0
7.5
5.0
2.5
0.0
Density
0.00
0.25
0.50
0.75
1.00
Hypergeometric p−value
threshold
>=1.5

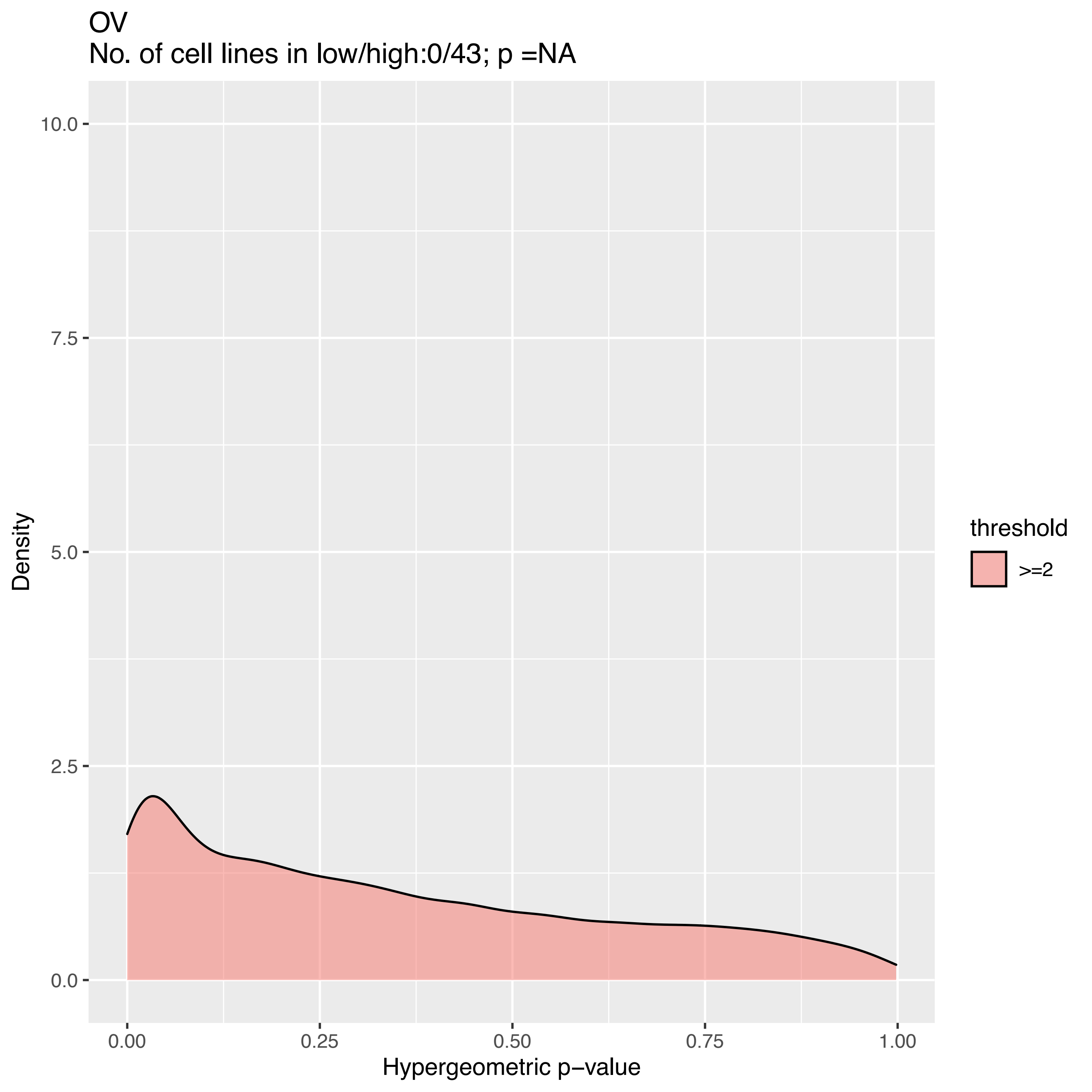
OV
No. of cell lines in low/high:0/43; p =NA
Density
10.0
7.5
5.0
2.5
0.0
0.00
0.25
0.50
0.75
1.00
Hypergeometric p−value
threshold
>=2

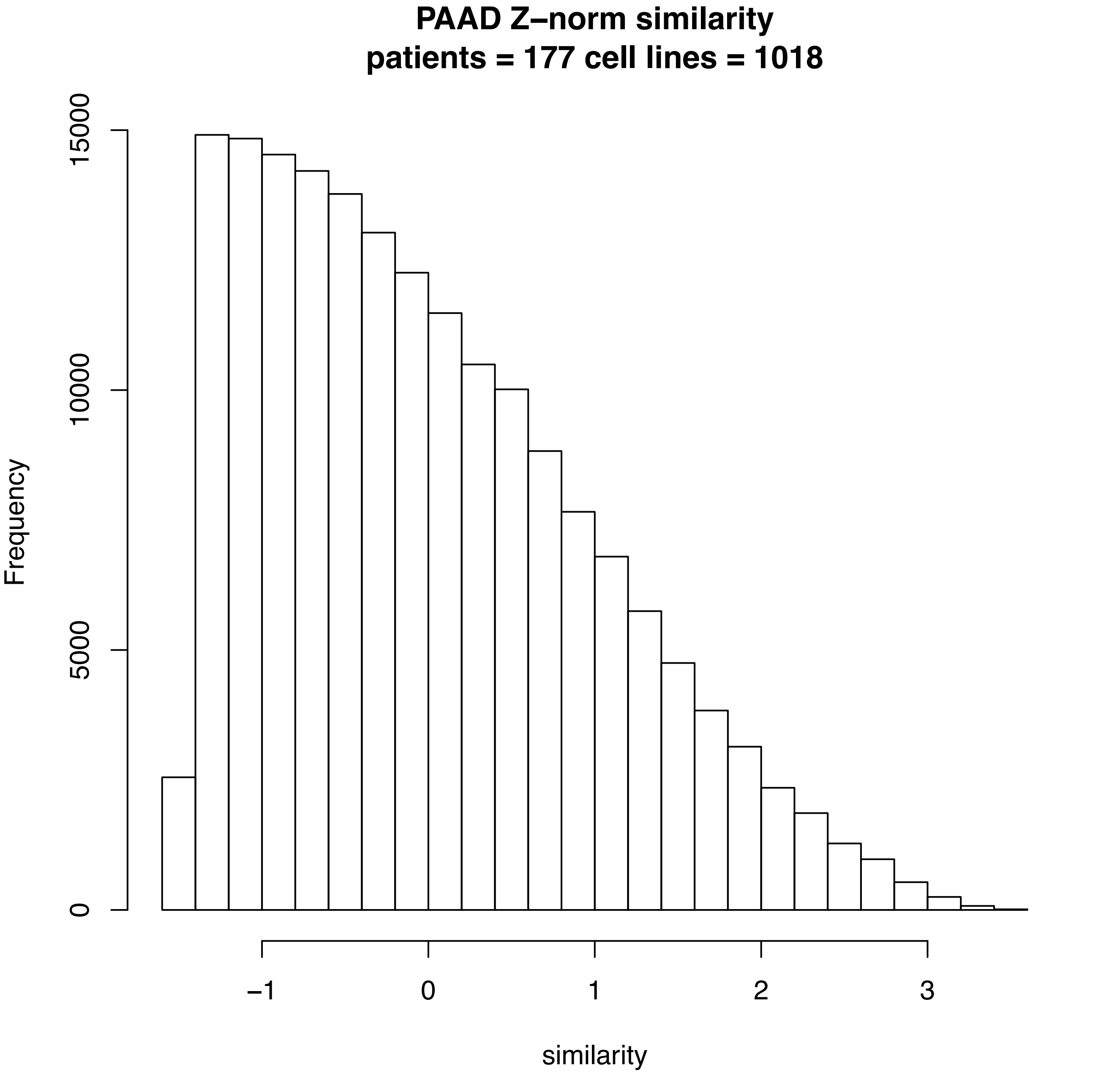
PAAD Z–norm similarity
patients = 177 cell lines = 1018
Frequency
0
5000
10000
15000
–1
0
1
2
3
similarity

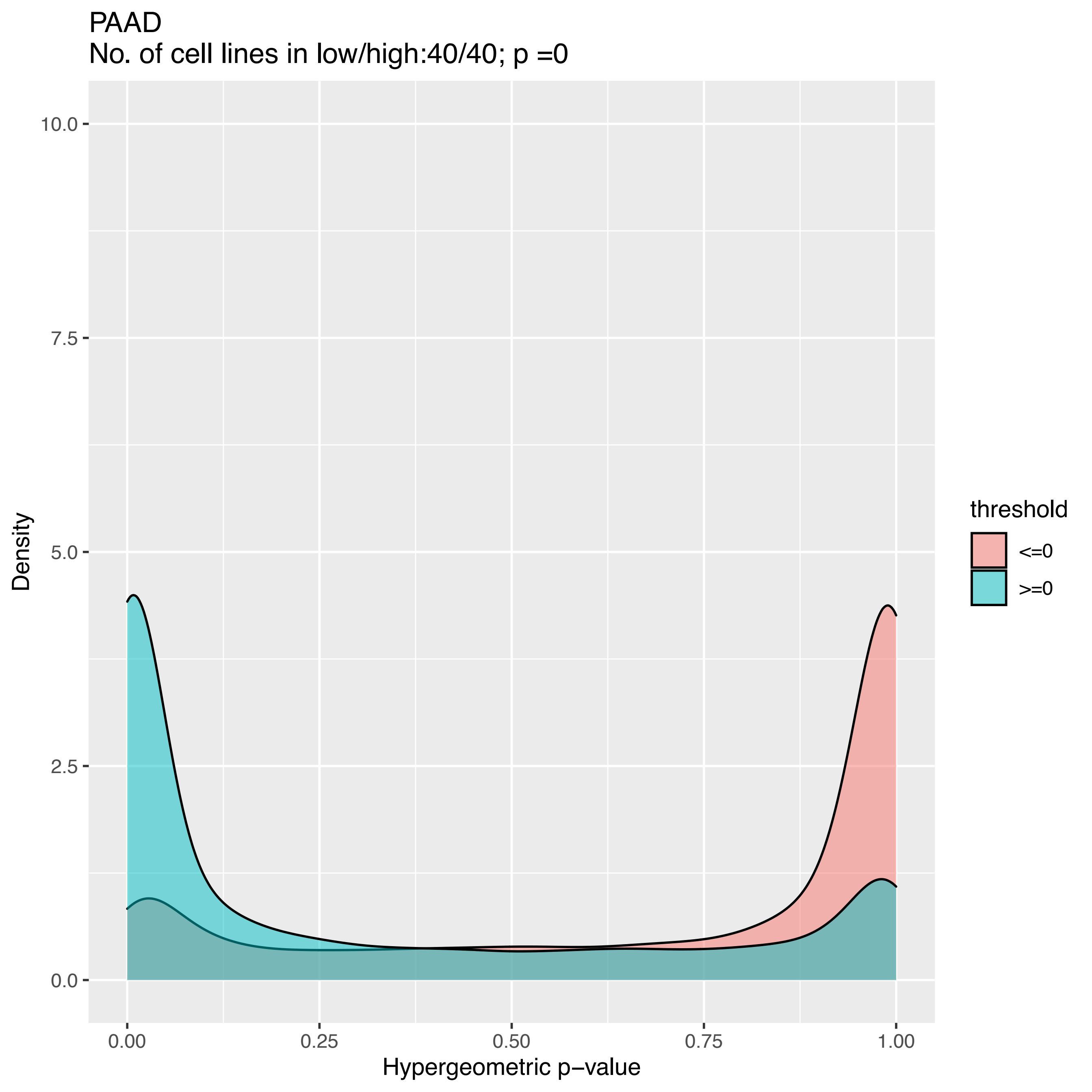

PAADBRK
No. of cell lines in low/high:40/40; p =0
Density
10.0
7.5
5.0
2.5
0.0
0.00
0.25
0.50
0.75
1.00
Hypergeometric p−value
threshold
<=0
>=0

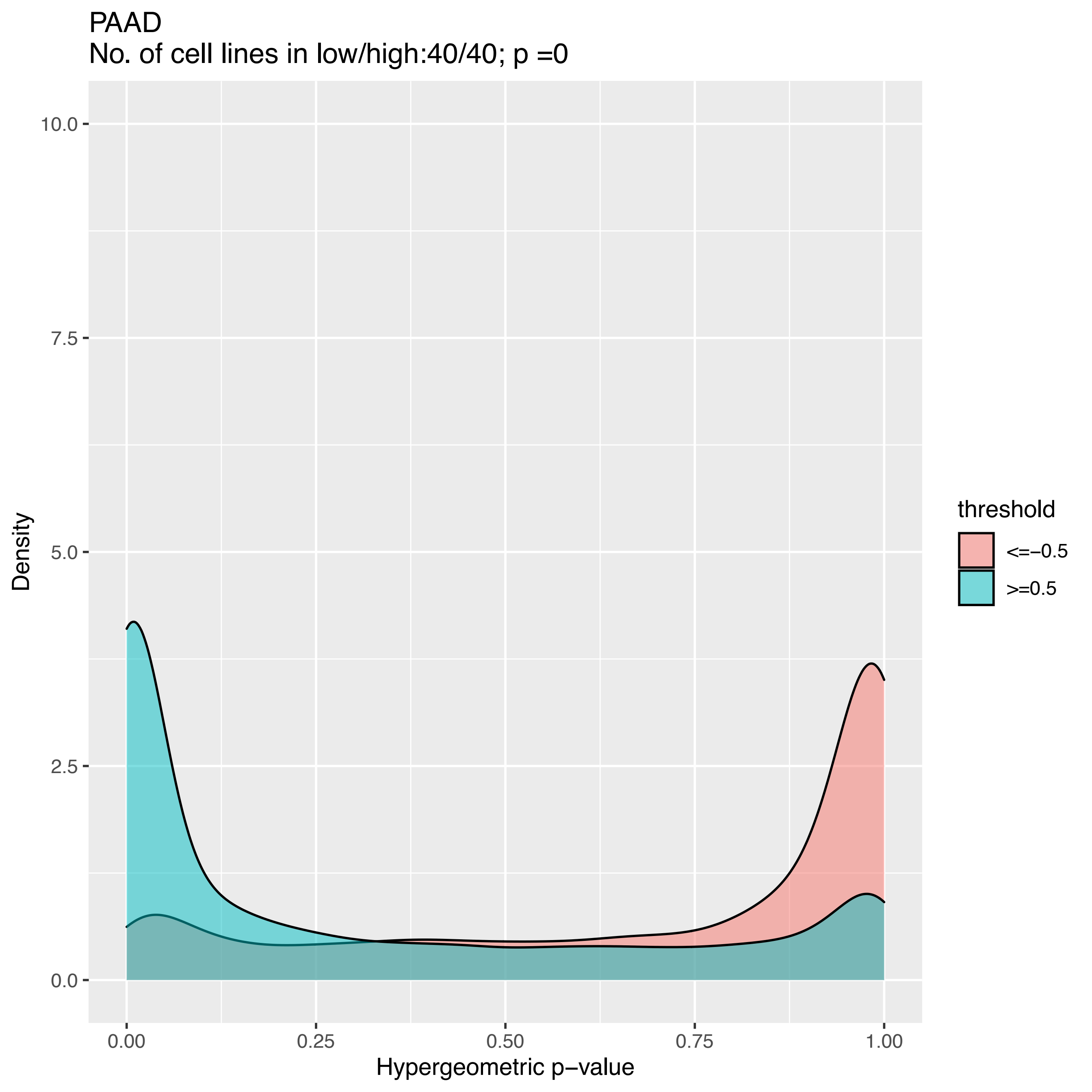
PAAD
No. of cell lines in low/high:40/40; p =0
Density
10.0
7.5
5.0
2.5
0.0
0.00
0.25
0.50
0.75
1.00
Hypergeometric p−value
threshold
<=−0.5
>=0.5

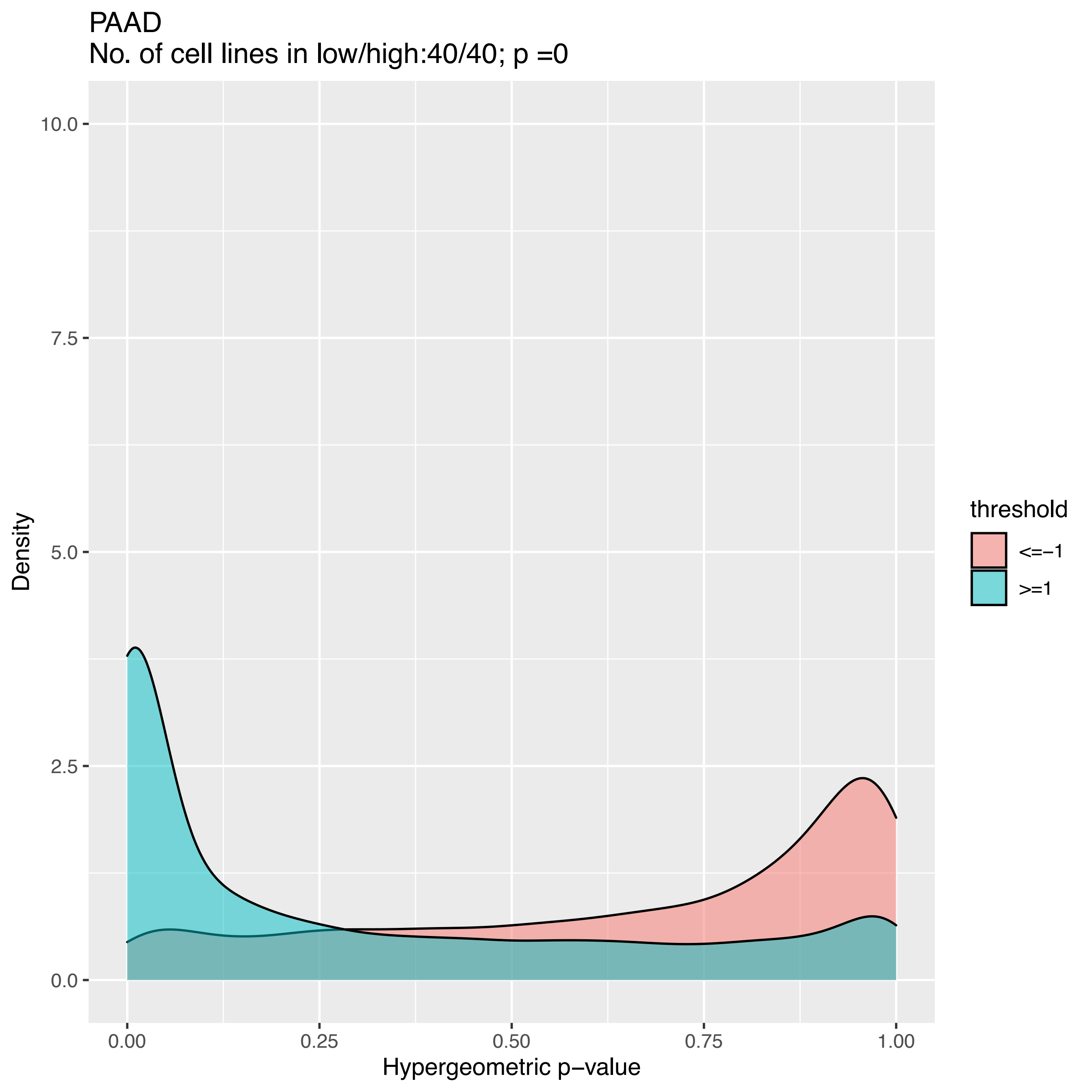
PAAD
No. of cell lines in low/high:40/40; p =0
Density
10.0
7.5
5.0
2.5
0.0
0.00
0.25
0.50
0.75
1.00
Hypergeometric p−value
threshold
<=−1
>=1

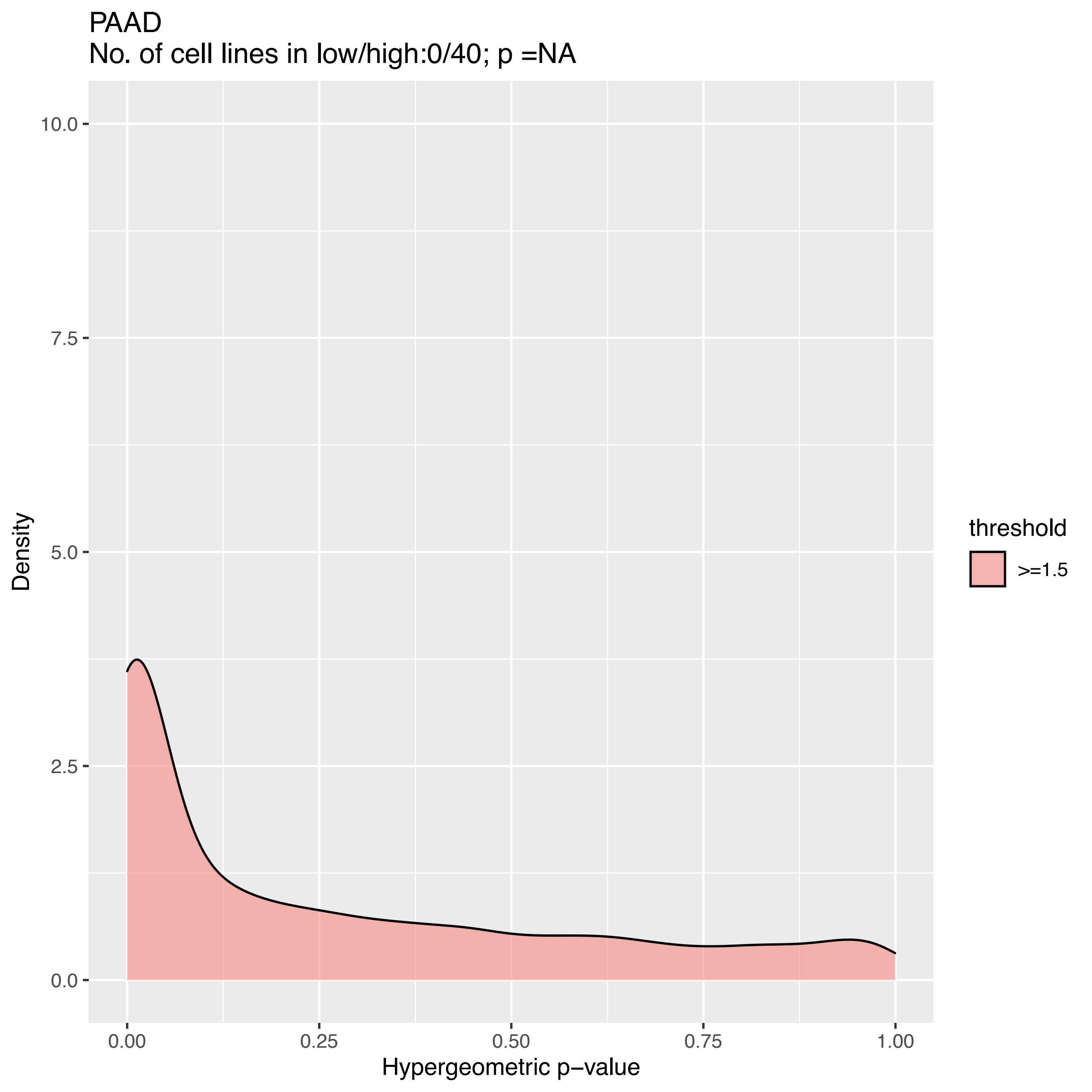

PAAD
No. of cell lines in low/high:0/40; p =NA
10.0
7.5
5.0
2.5
0.0
Density
0.00
0.25
0.50
0.75
1.00
Hypergeometric p−value
threshold
>=1.5

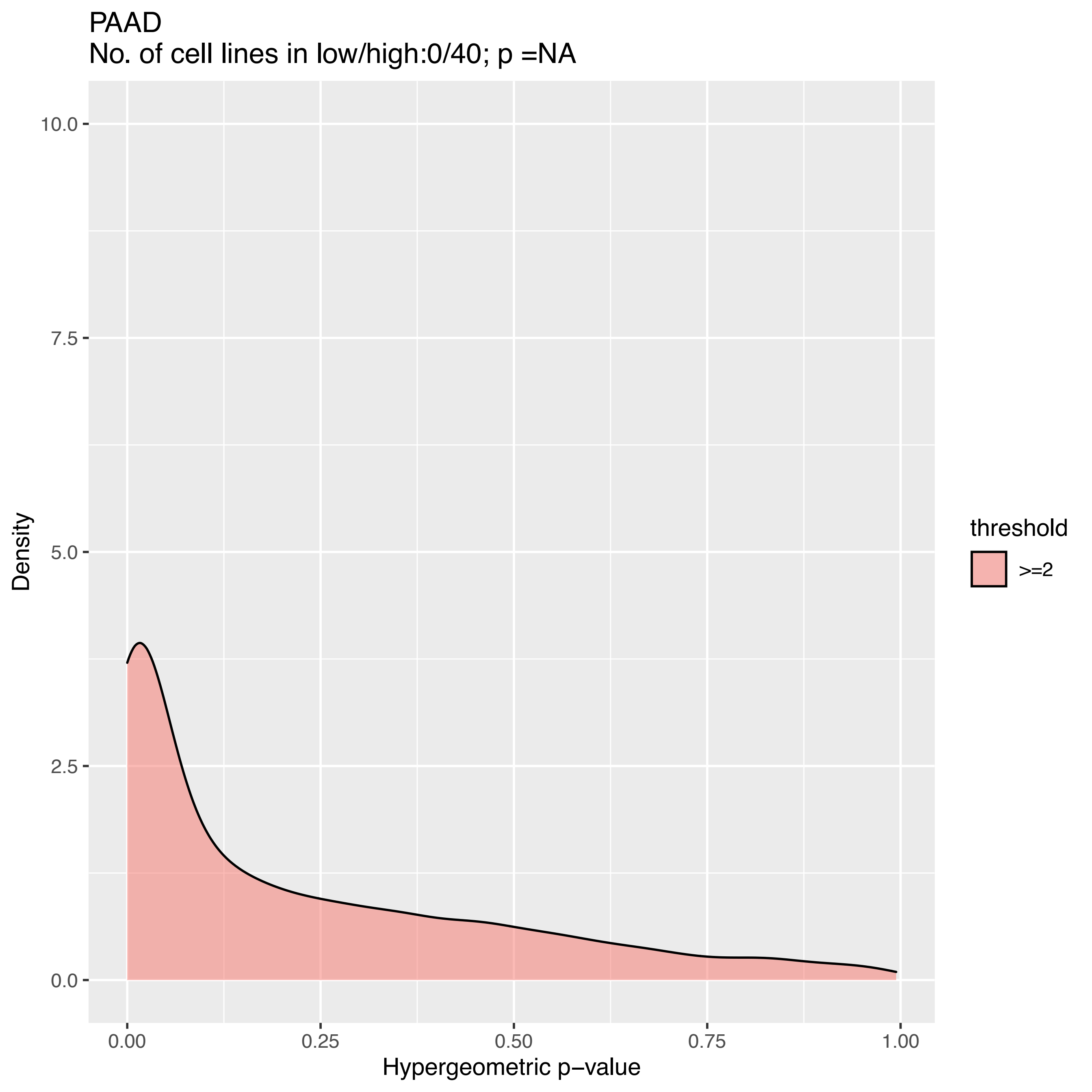
PAAD
No. of cell lines in low/high:0/40; p =NA
Density
10.0
7.5
5.0
2.5
0.0
0.00
0.25
0.50
0.75
1.00
Hypergeometric p−value
threshold
>=2

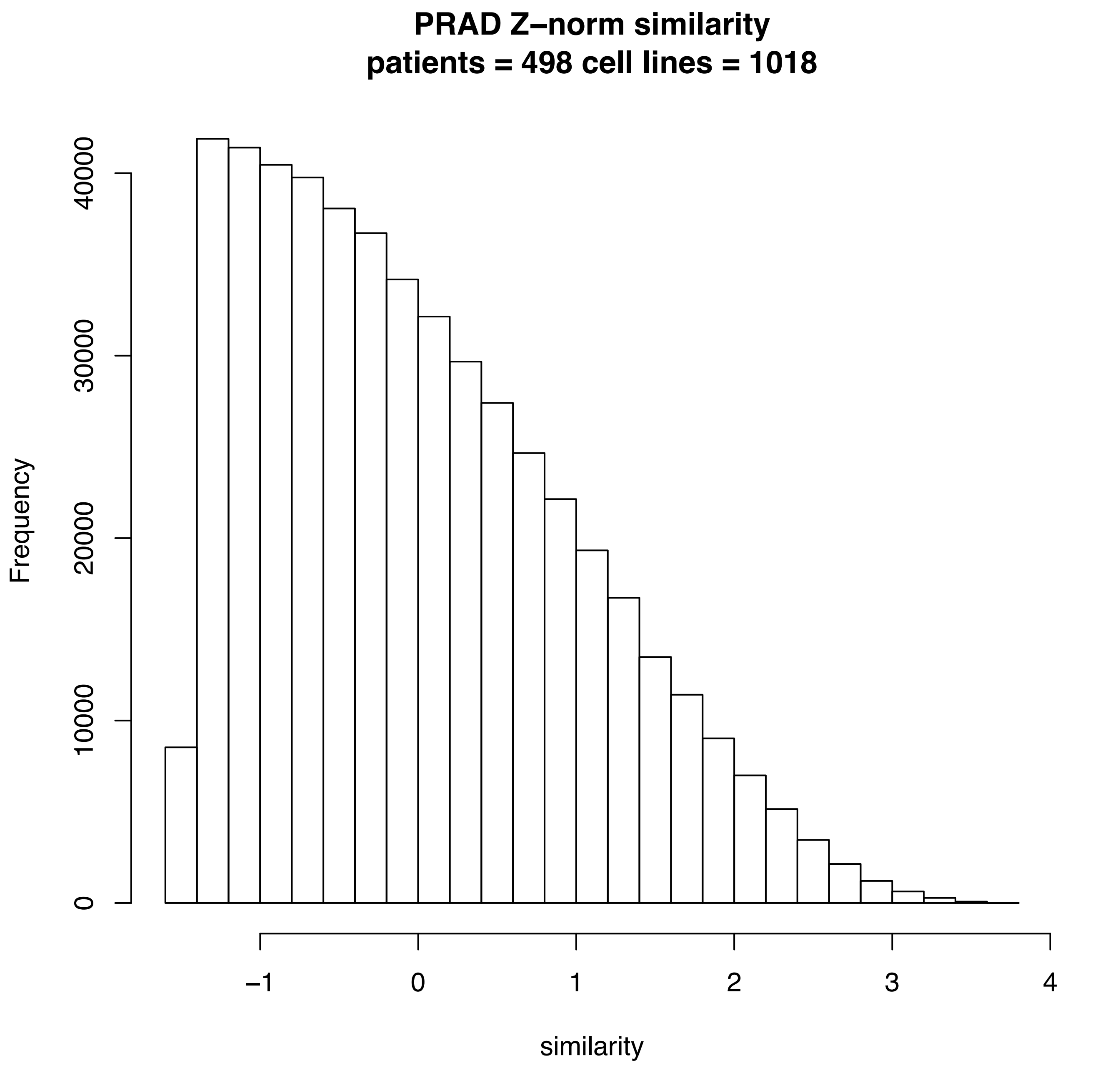
PRAD Z–norm similarity
patients = 498 cell lines = 1018
Frequency
0
10000
20000
30000
40000
−1
0
1
2
3
4
similarity

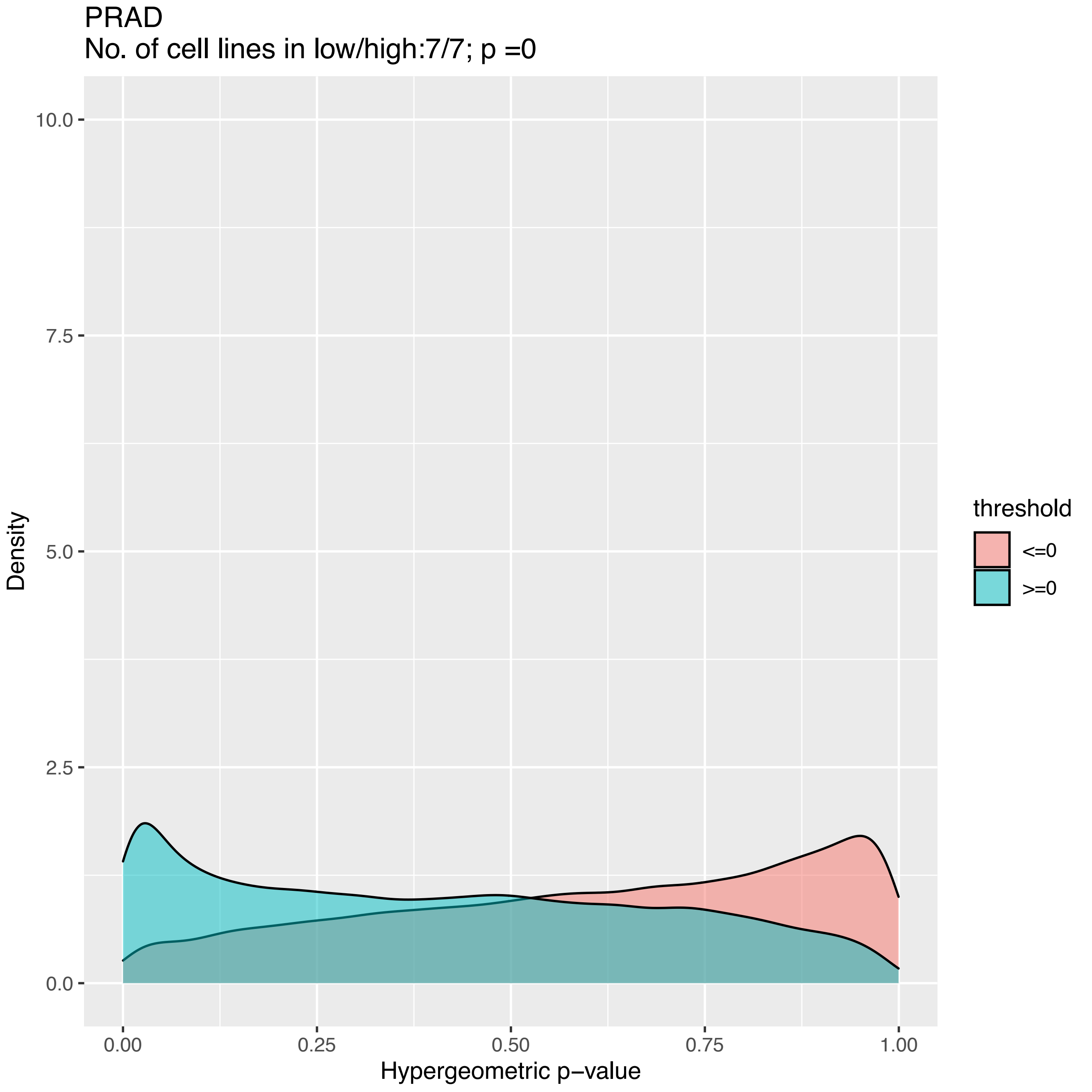
PRAD
No. of cell lines in low/high:7/7; p =0
Density
10.0
7.5
5.0
2.5
0.0
0.00
0.25
0.50
0.75
1.00
Hypergeometric p−value
threshold
<=0
>=0

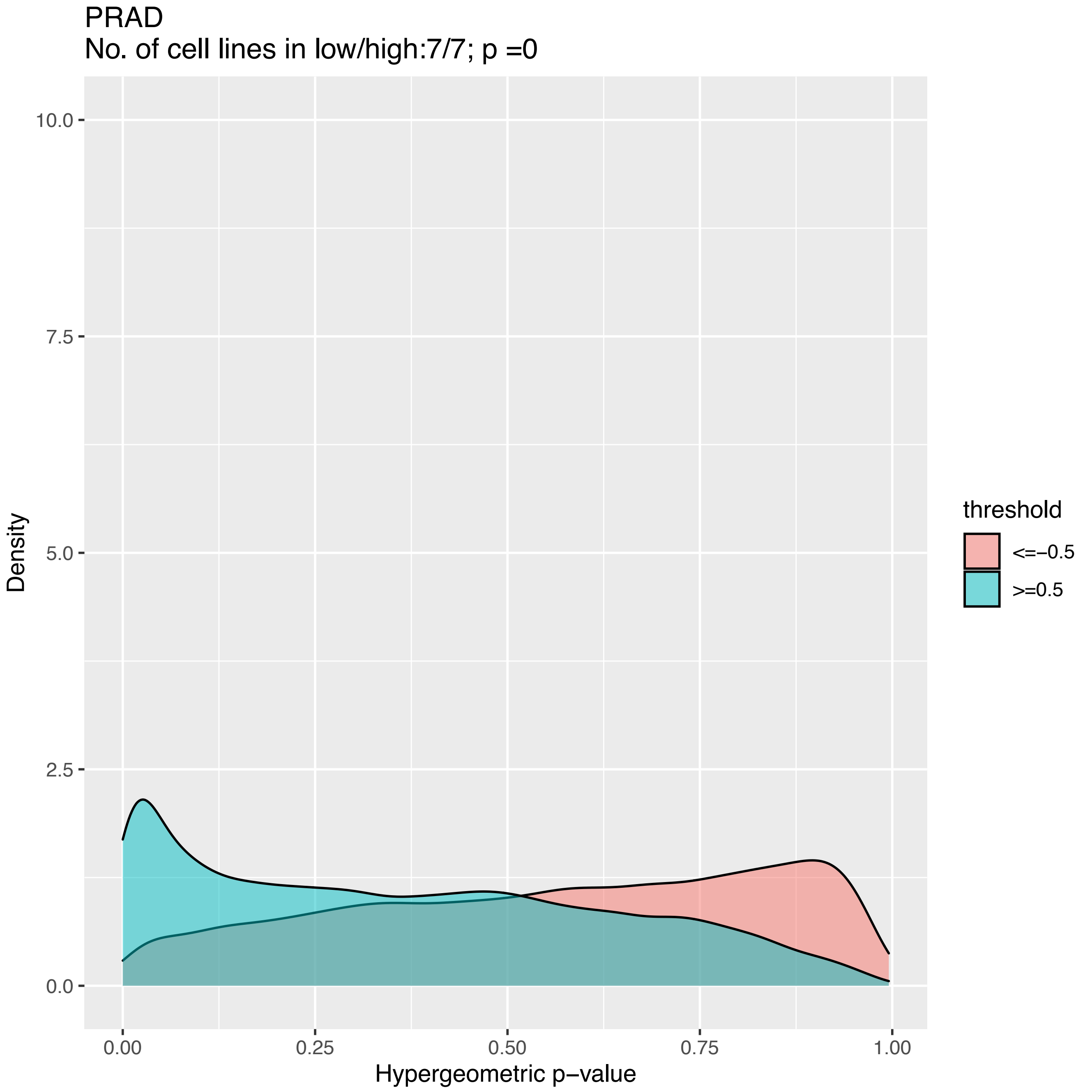
PRAD
No. of cell lines in low/high:7/7; p =0
10.0
7.5
5.0
2.5
0.0
Density
0.00
0.25
0.50
0.75
1.00
Hypergeometric p−value
threshold
<=−0.5
>=0.5

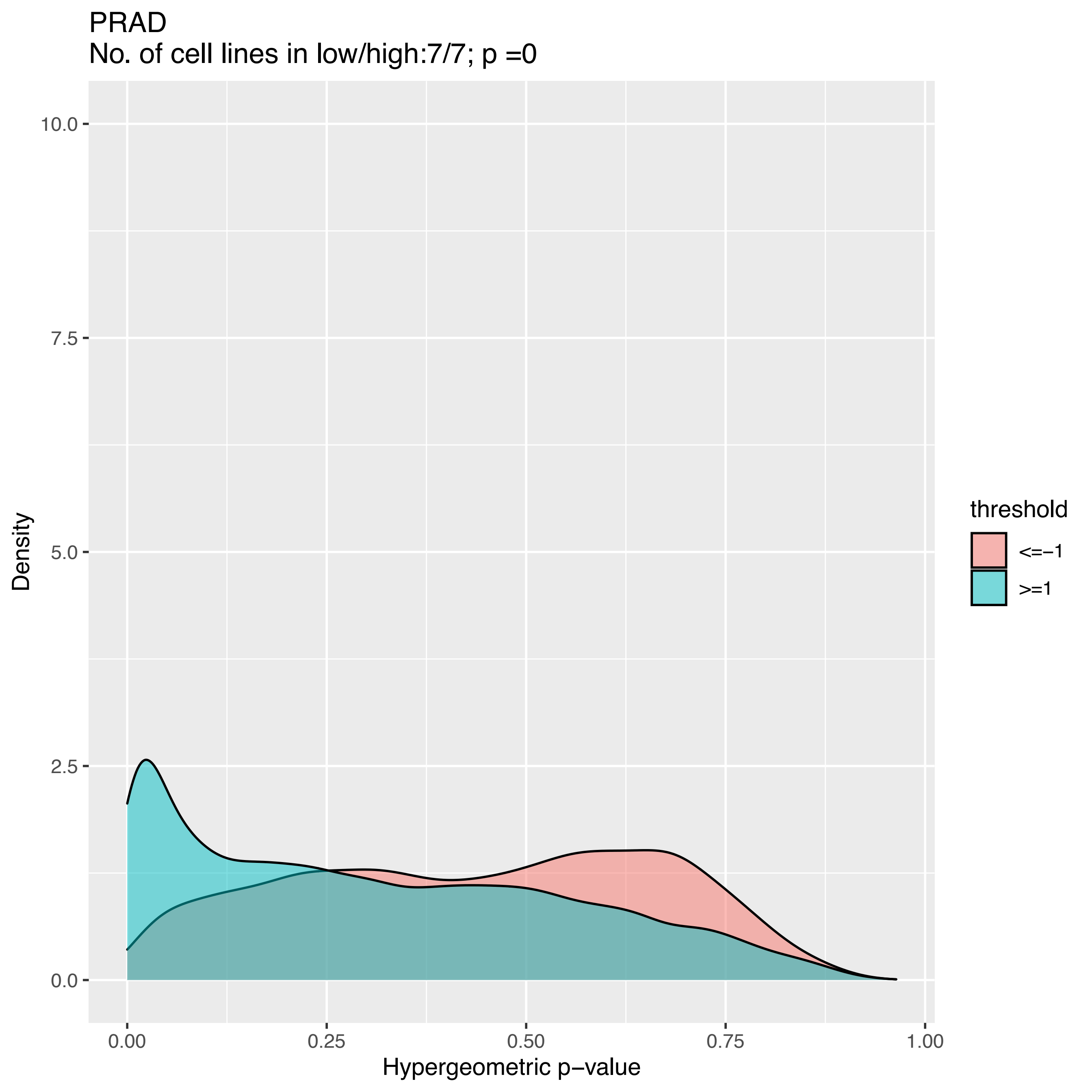
PRAD
No. of cell lines in low/high:7/7; p =0
Density
10.0
7.5
5.0
2.5
0.0
0.00
0.25
0.50
0.75
1.00
Hypergeometric p−value
threshold
<=−1
>=1

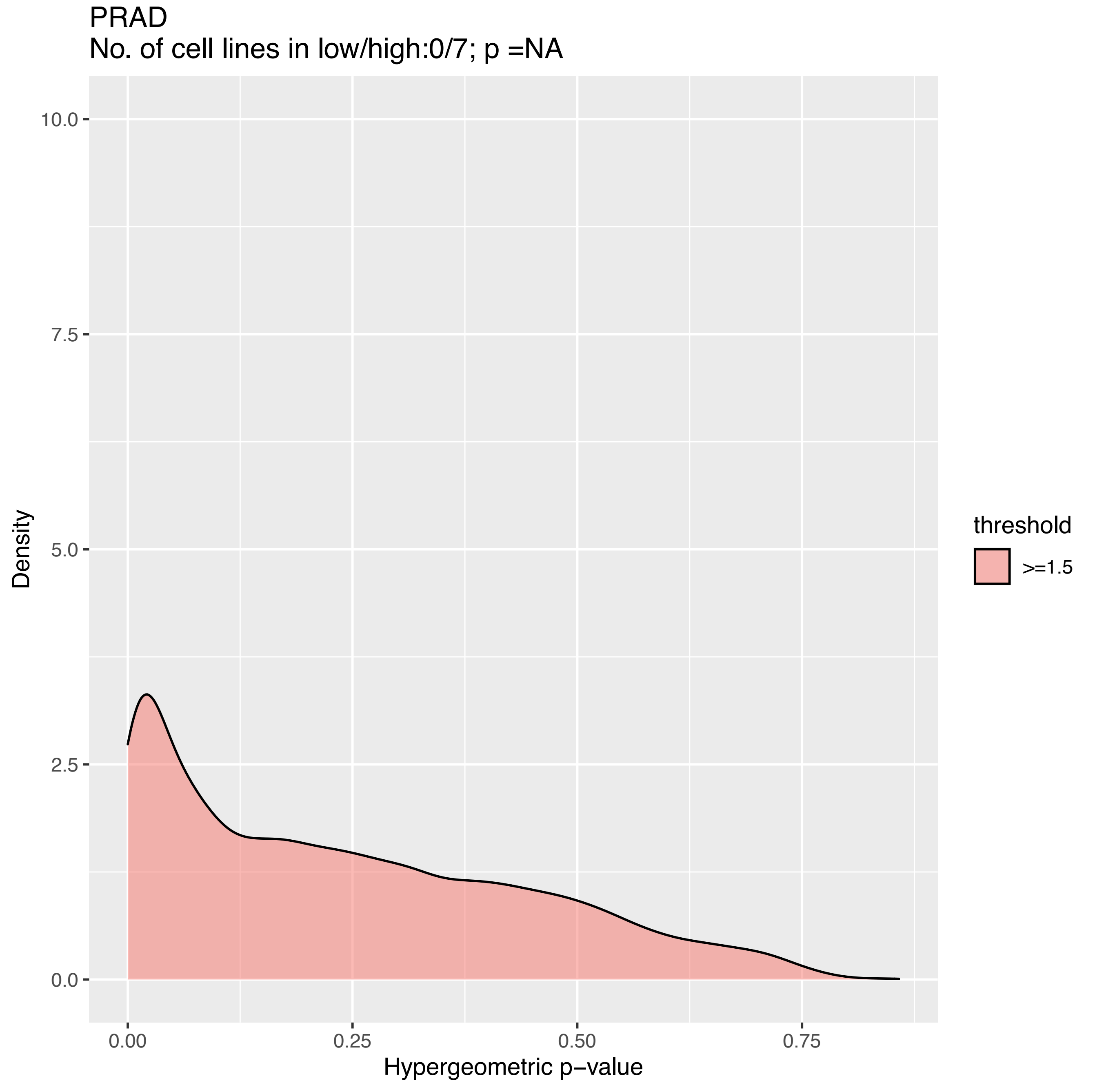
PRAD
No. of cell lines in low/high:0/7; p =NA
Density
10.0
7.5
5.0
2.5
0.0
0.00
0.25
0.50
0.75
Hypergeometric p−value
threshold
>=1.5

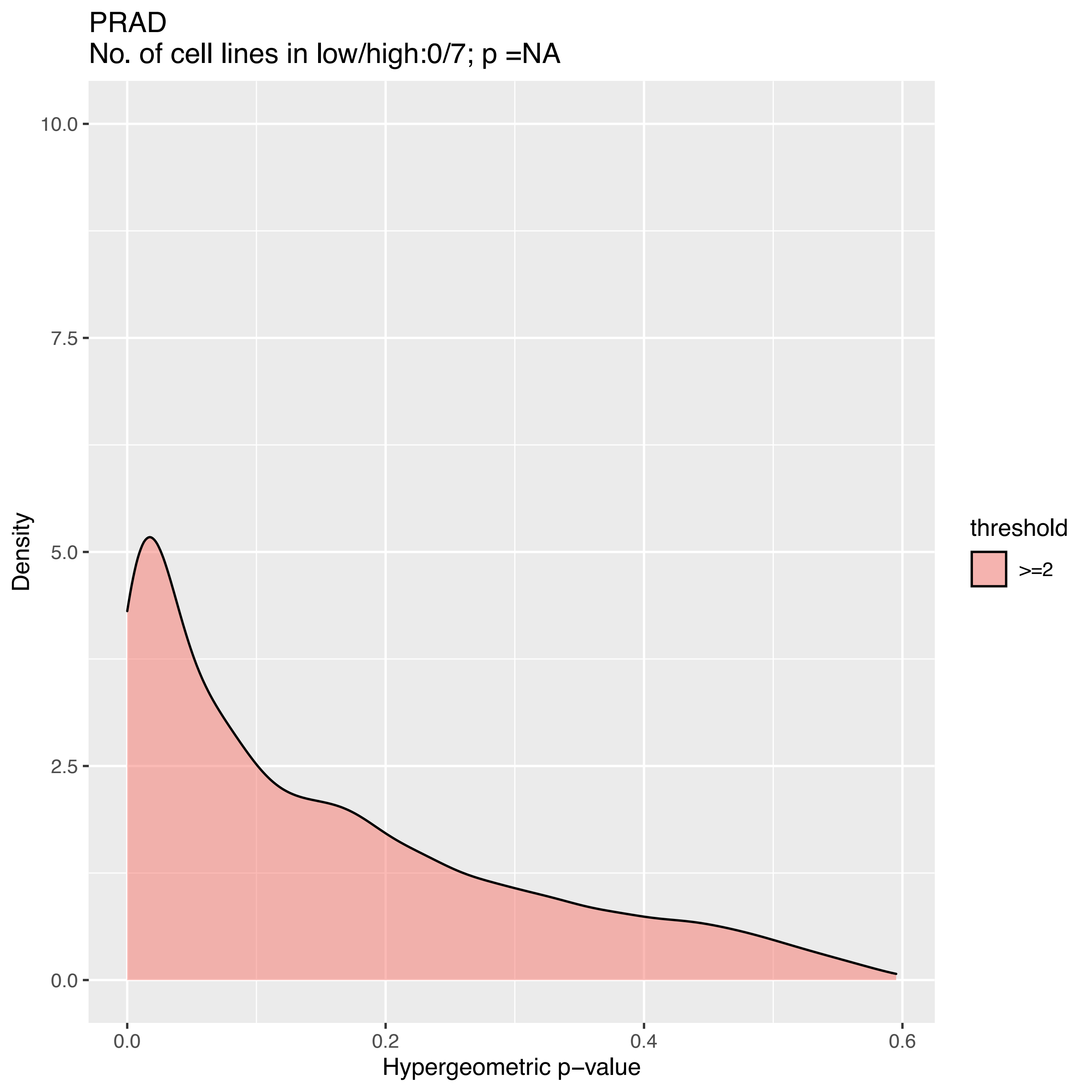

PRAD
No. of cell lines in low/high:0/7; p =NA
Density
10.0
7.5
5.0
2.5
0.0
0.0
0.2
0.4
0.6
Hypergeometric p−value
threshold
>=2

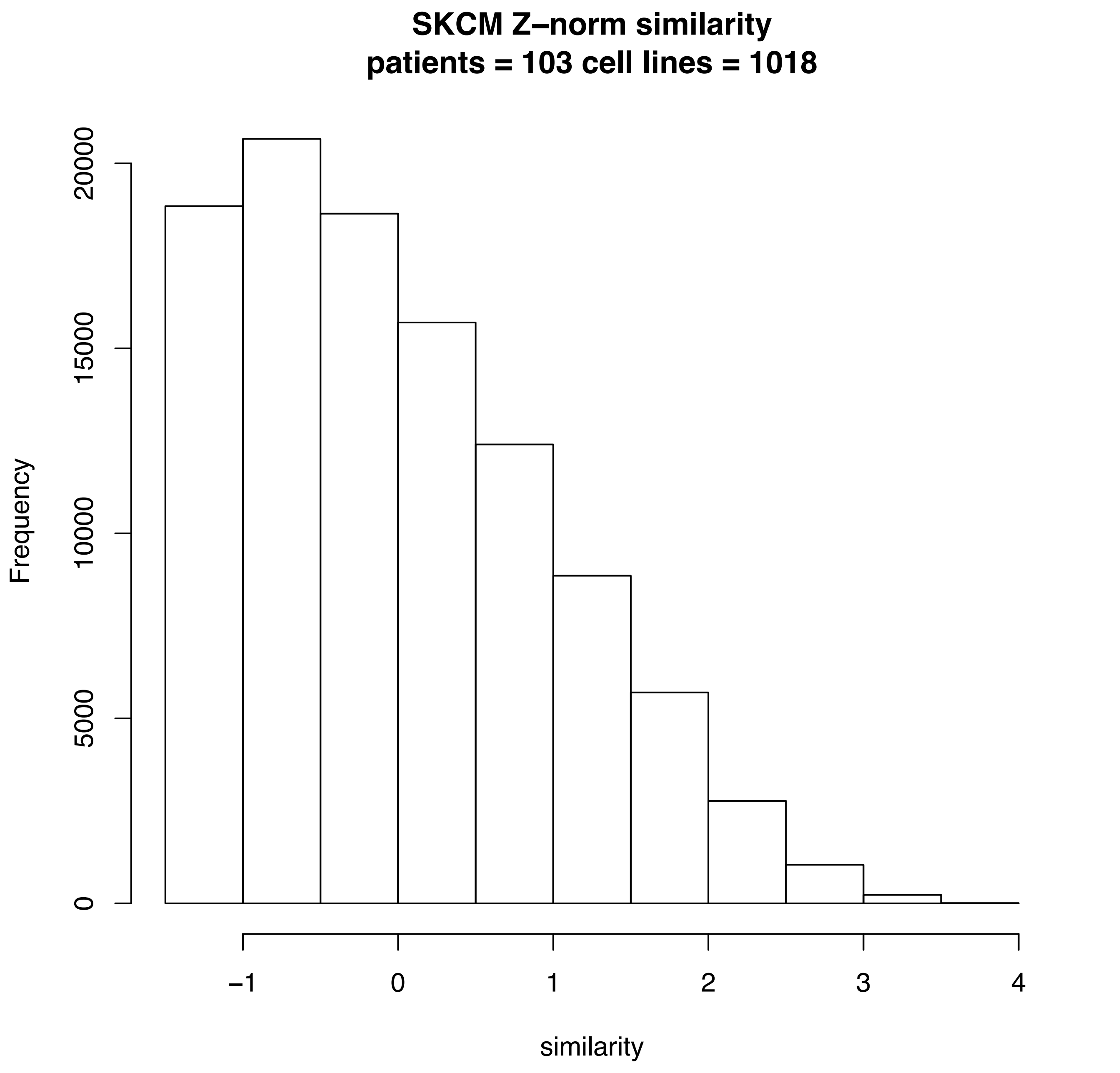
SKCM Z–norm similarity
patients = 103 cell lines = 1018
Frequency
0
5000
10000
15000
20000
−1
0
1
2
3
4
similarity

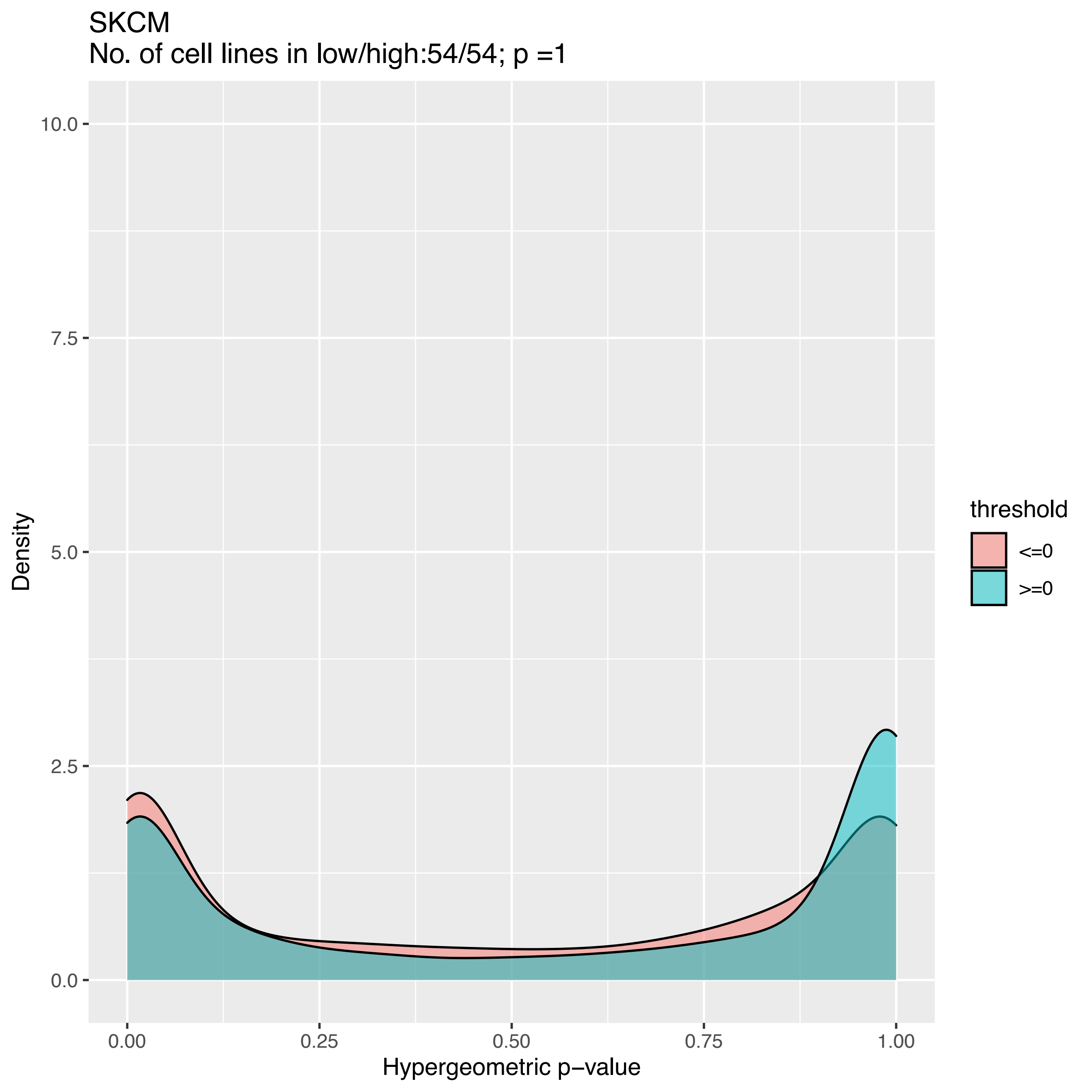

SKCM
No. of cell lines in low/high:54/54; p =1
10.0
7.5
5.0
2.5
0.0
Density
0.00
0.25
0.50
0.75
1.00
Hypergeometric p−value
threshold
<=0
>=0

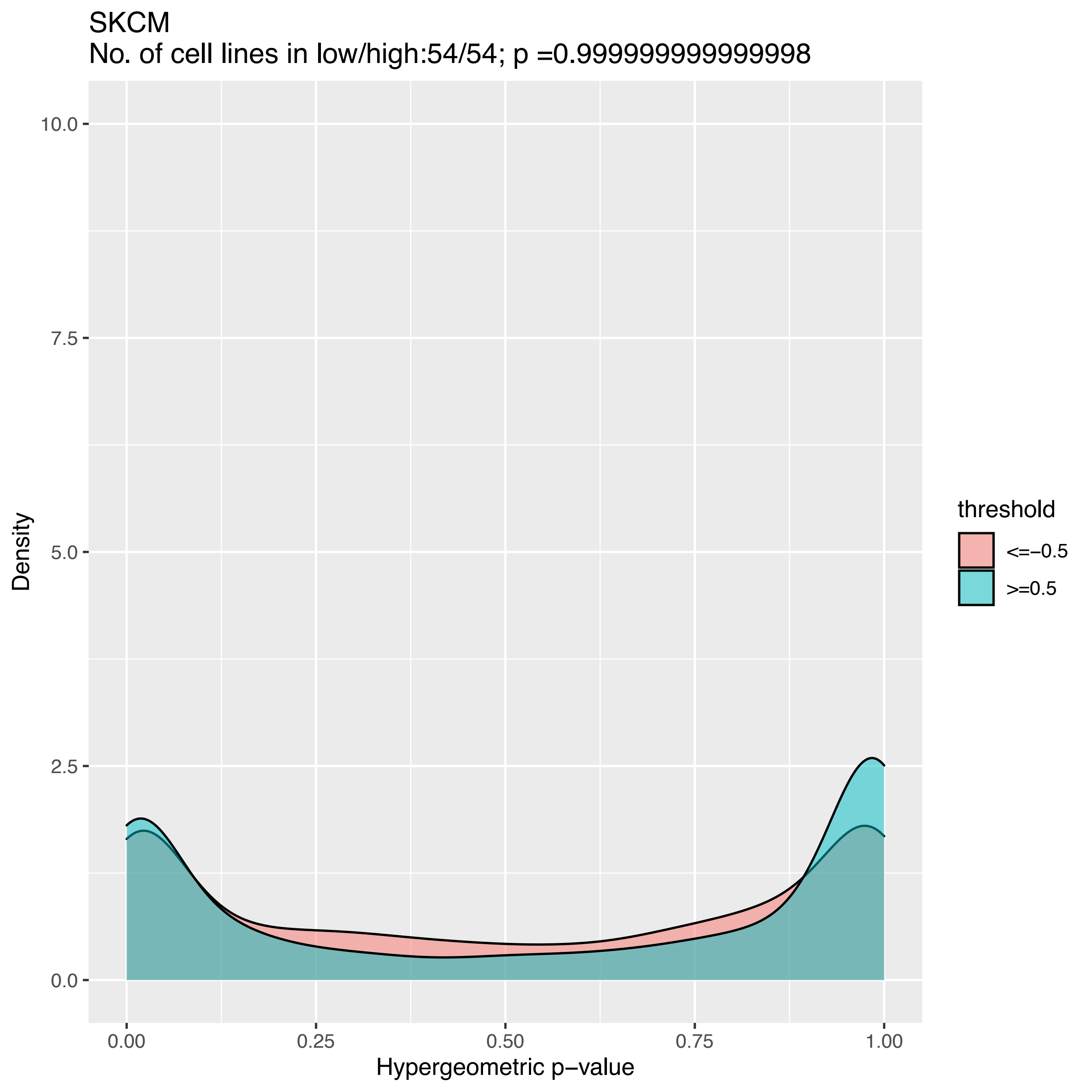

SKCM
No. of cell lines in low/high:54/54; p =0.9999999999999998
Density
10.0
7.5
5.0
2.5
0.0
0.00
0.25
0.50
0.75
1.00
Hypergeometric p−value
threshold
<=−0.5
>=0.5

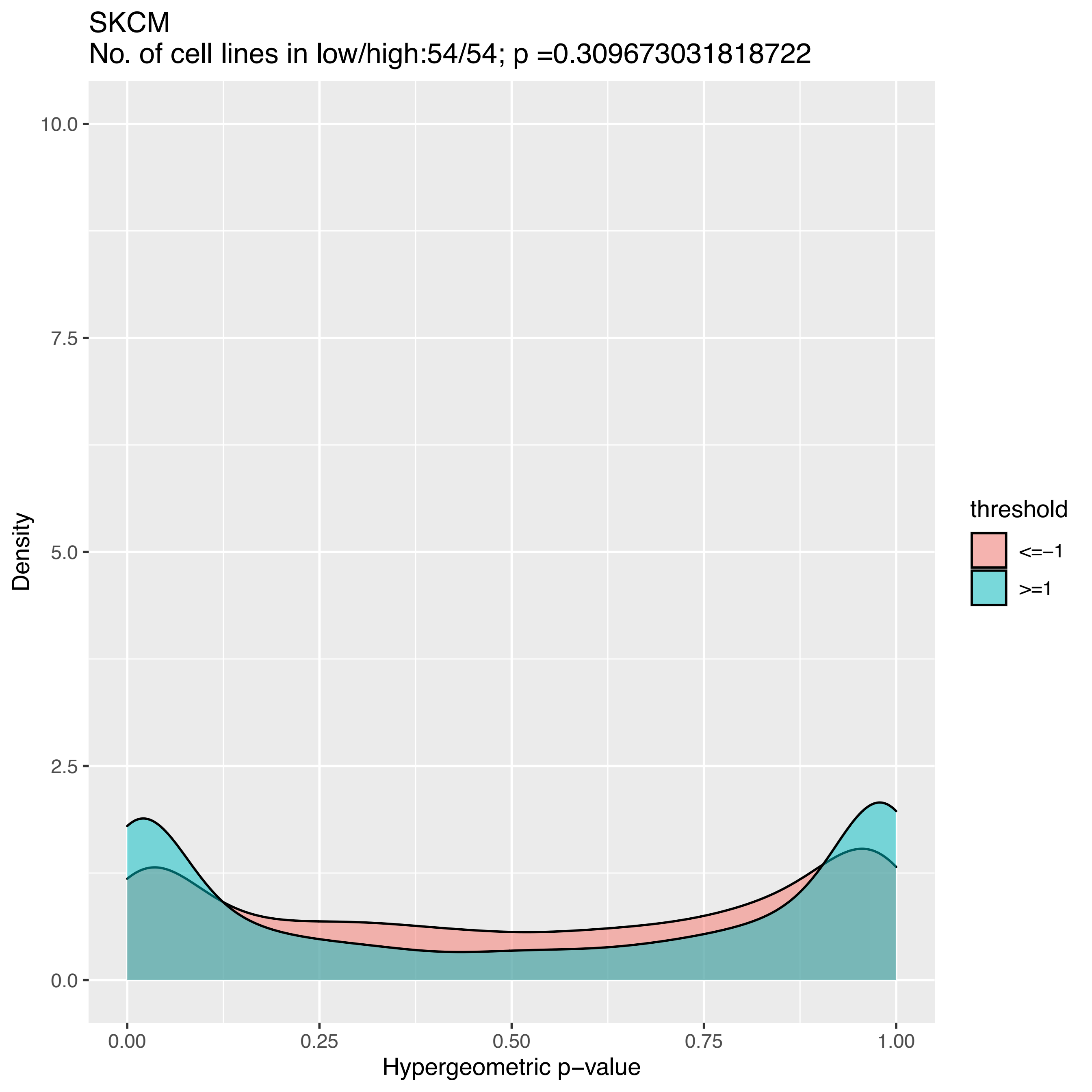
SKCM
No. of cell lines in low/high:54/54; p =0.309673031818722
Density
Hypergeometric p−value
threshold
<=−1
>=1
0.0
2.5
5.0
7.5
10.0
0.00
0.25
0.50
0.75
1.00

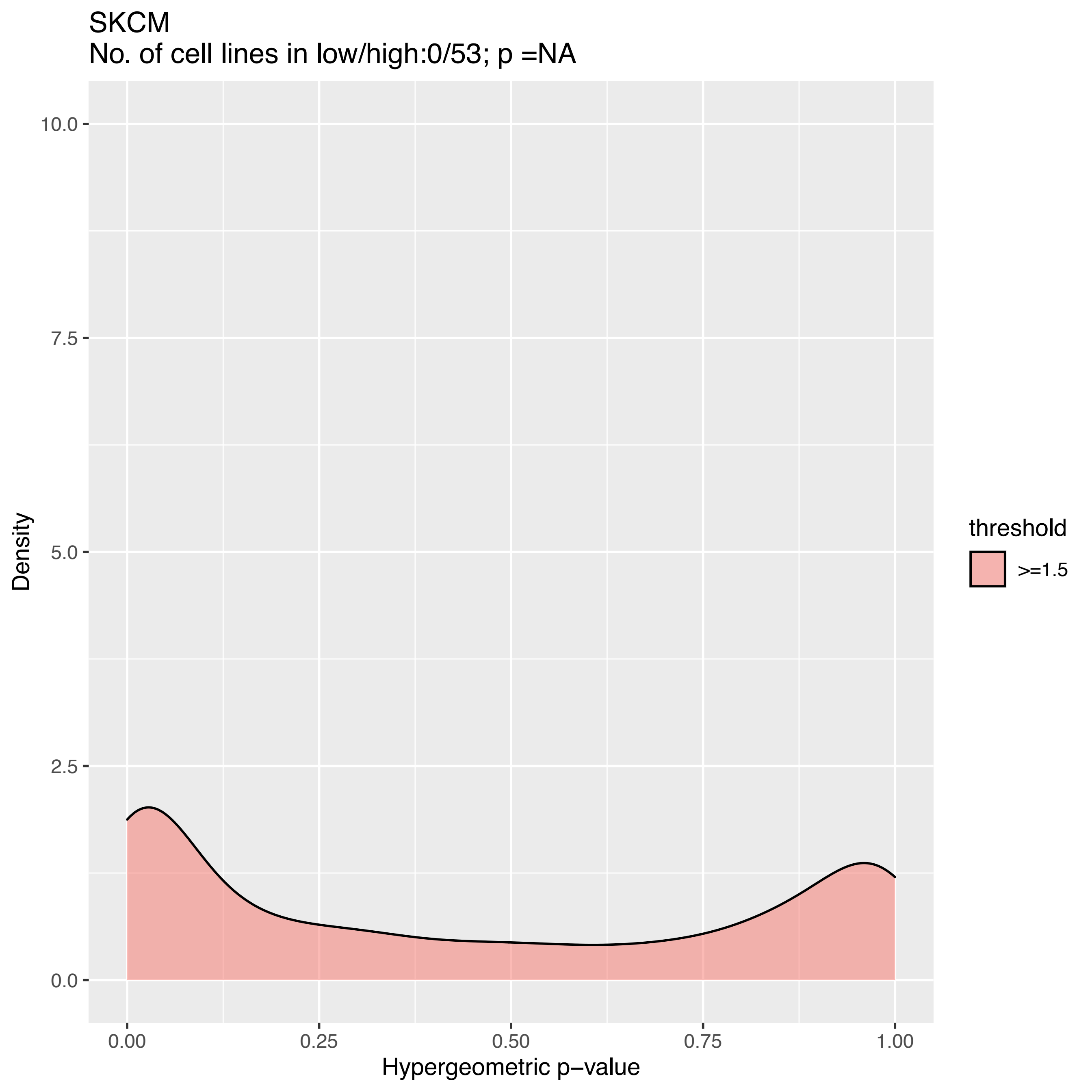
SKCM
No. of cell lines in low/high:0/53; p =NA
Density
10.0
7.5
5.0
2.5
0.0
0.00
0.25
0.50
0.75
1.00
Hypergeometric p−value
threshold
>=1.5

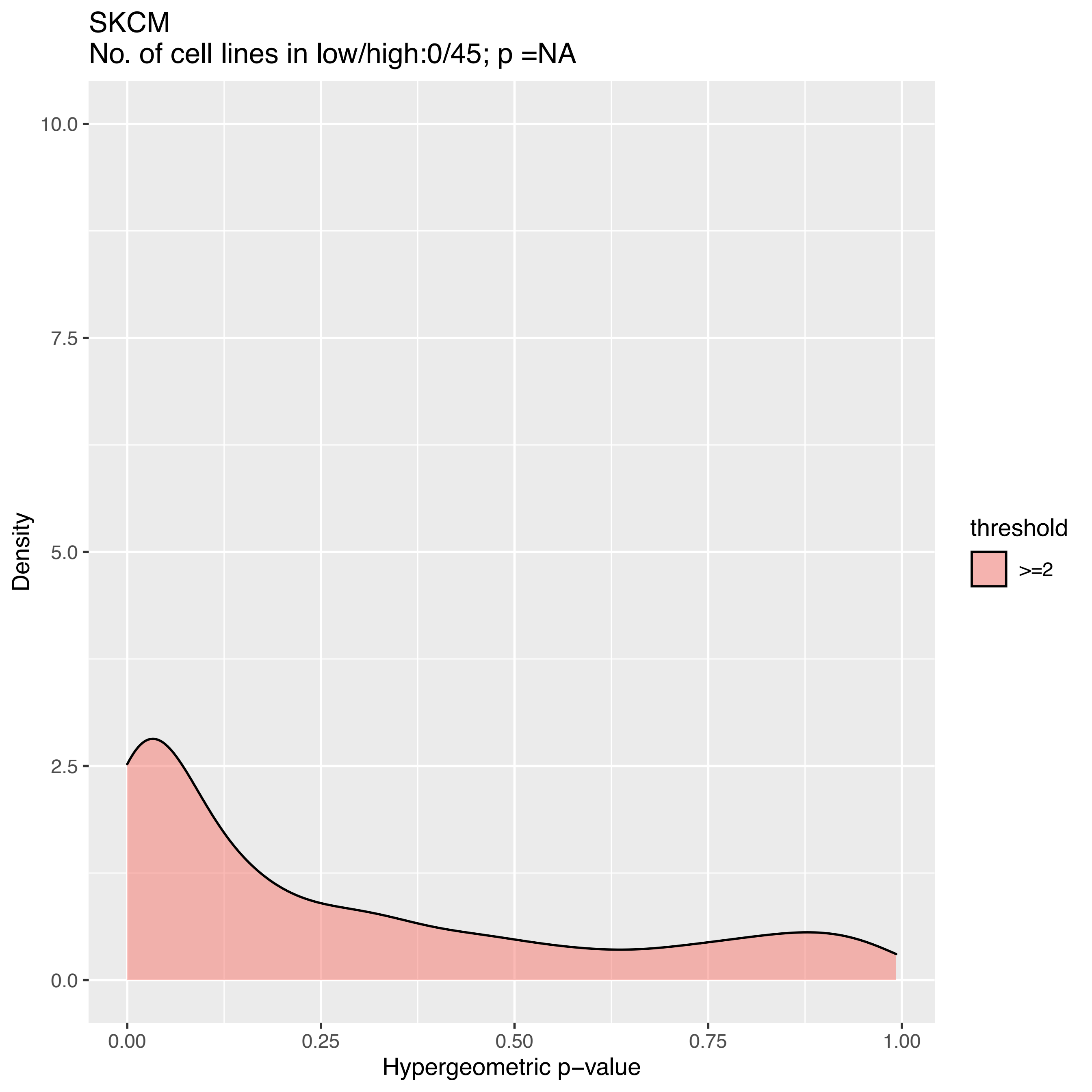
SKCM
No. of cell lines in low/high:0/45; p =NA
10.0
7.5
5.0
2.5
0.0
Density
0.00
0.25
0.50
0.75
1.00
Hypergeometric p−value
threshold
>=2

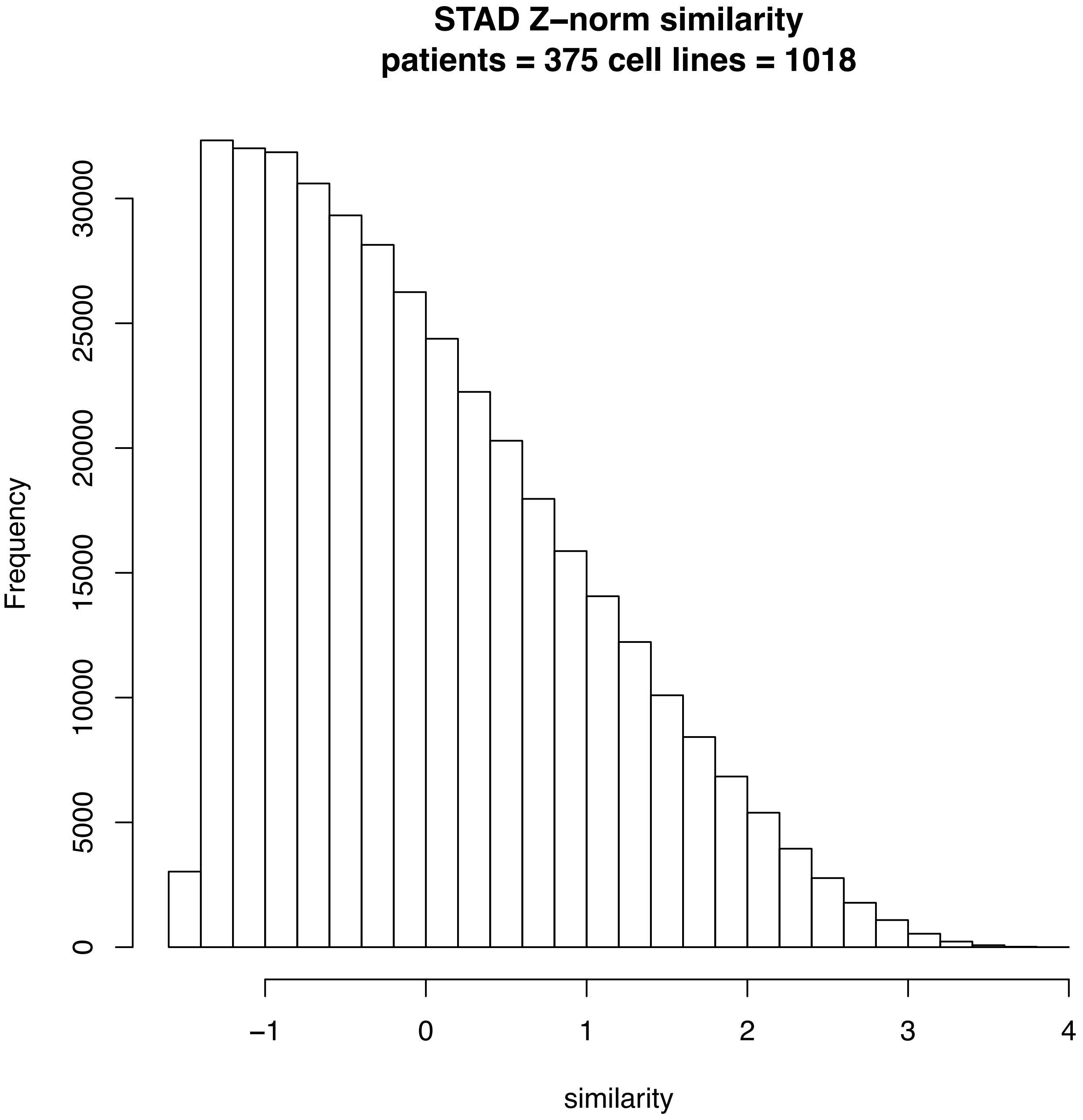
STAD Z–norm similarity
patients = 375 cell lines = 1018
Frequency
0
5000
10000
15000
20000
25000
30000
−1
0
1
2
3
4
similarity

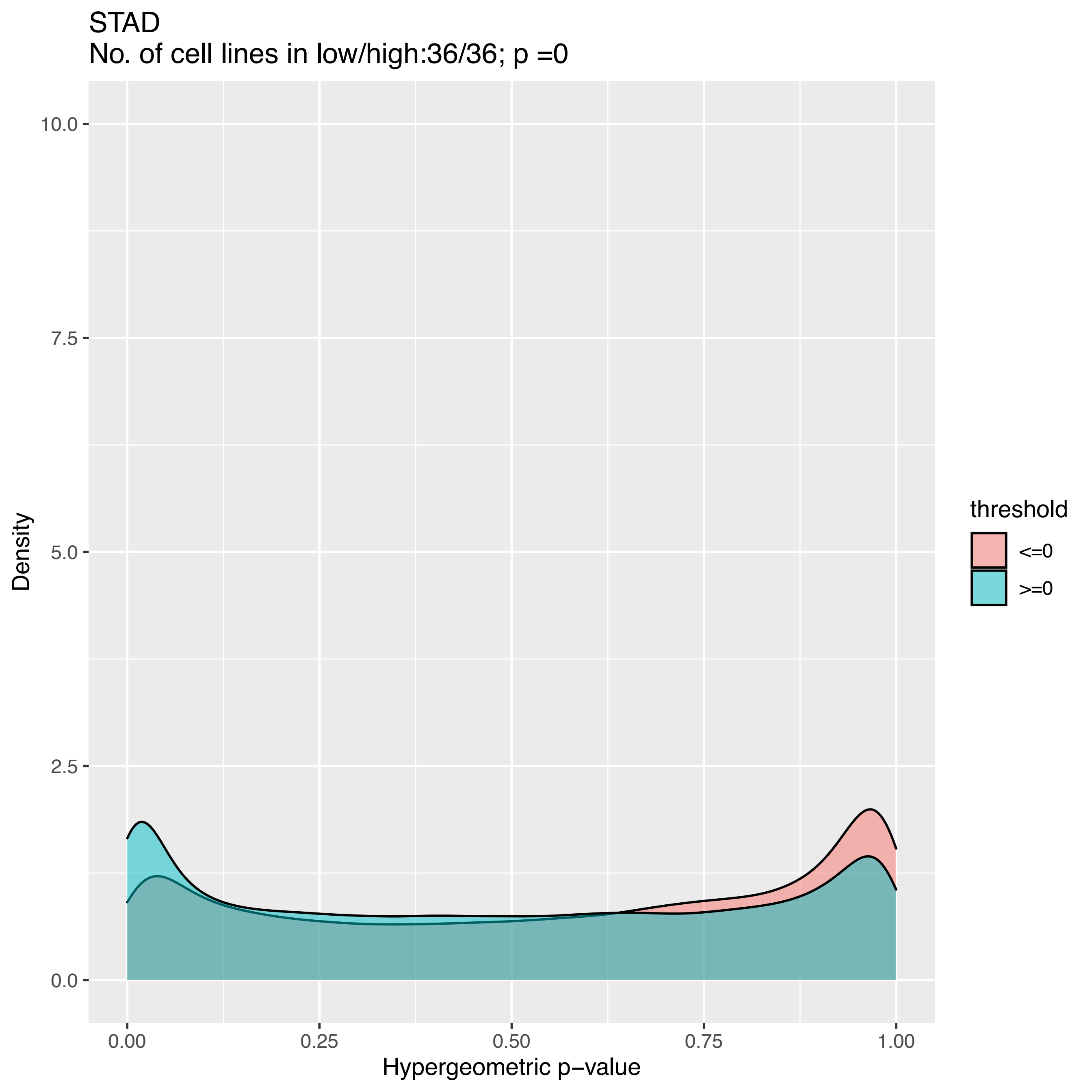
STAD
No. of cell lines in low/high:36/36; p =0
10.0
7.5
5.0
2.5
0.0
Density
0.00
0.25
0.50
0.75
1.00
Hypergeometric p−value
threshold
<=0
>=0

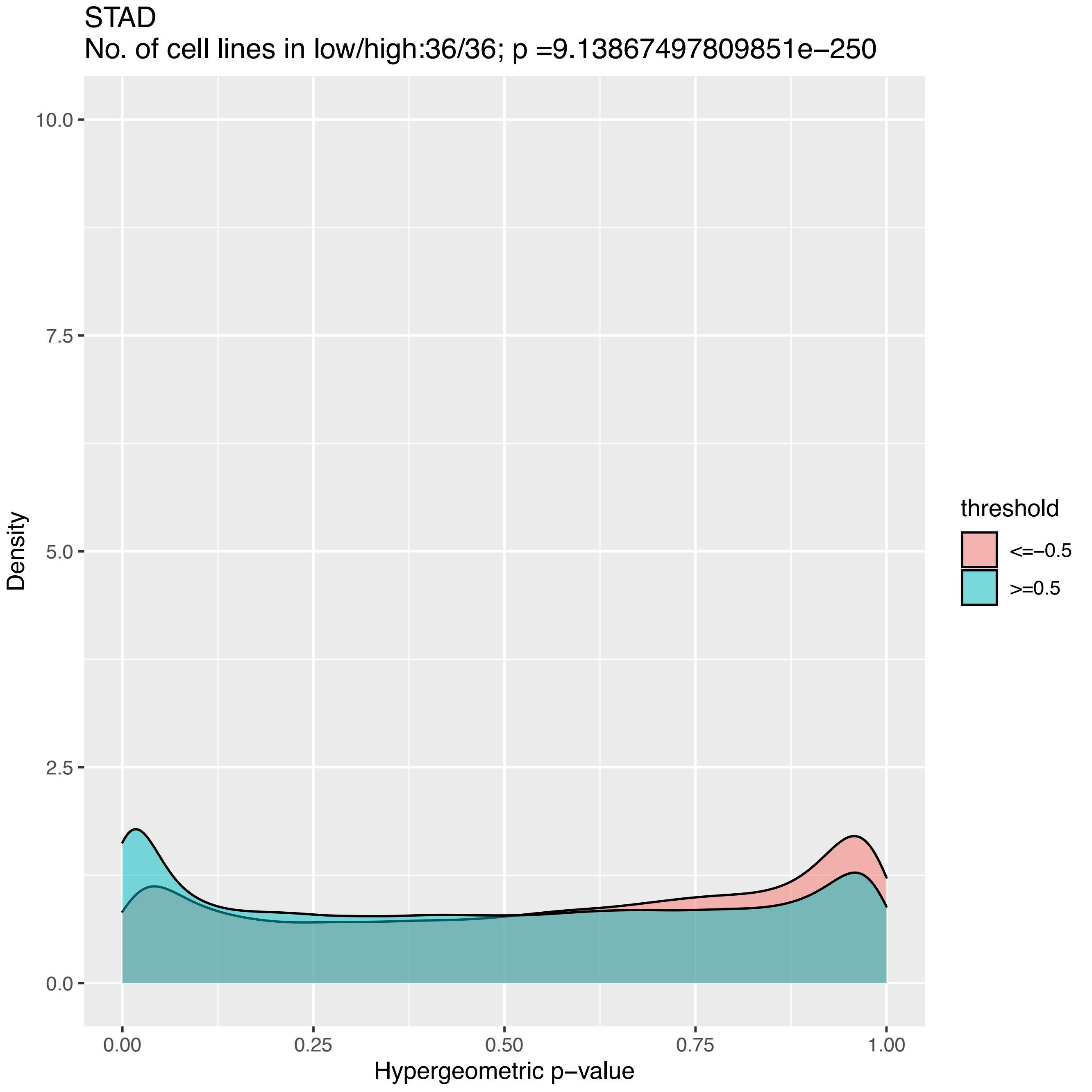
STAD
No. of cell lines in low/high:36/36; p =9.13867497809851e−250
Density
10.0
7.5
5.0
2.5
0.0
0.00
0.25
0.50
0.75
1.00
Hypergeometric p−value
threshold
<=−0.5
>=0.5

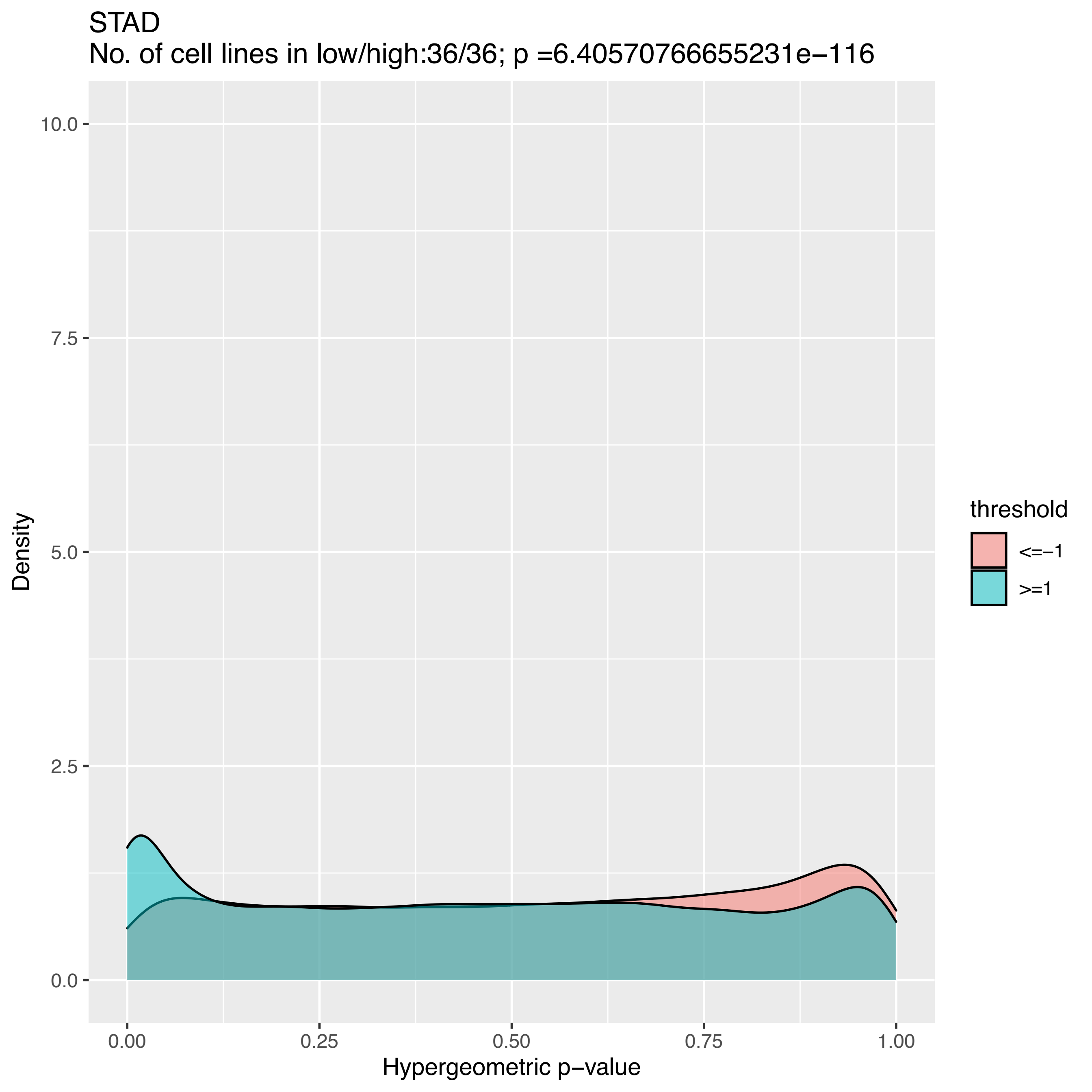

STAD
No. of cell lines in low/high:36/36; p =6.40570766655231e−116
10.0
7.5
5.0
2.5
0.0
Density
0.00
0.25
0.50
0.75
1.00
Hypergeometric p−value
threshold
<=−1
>=1

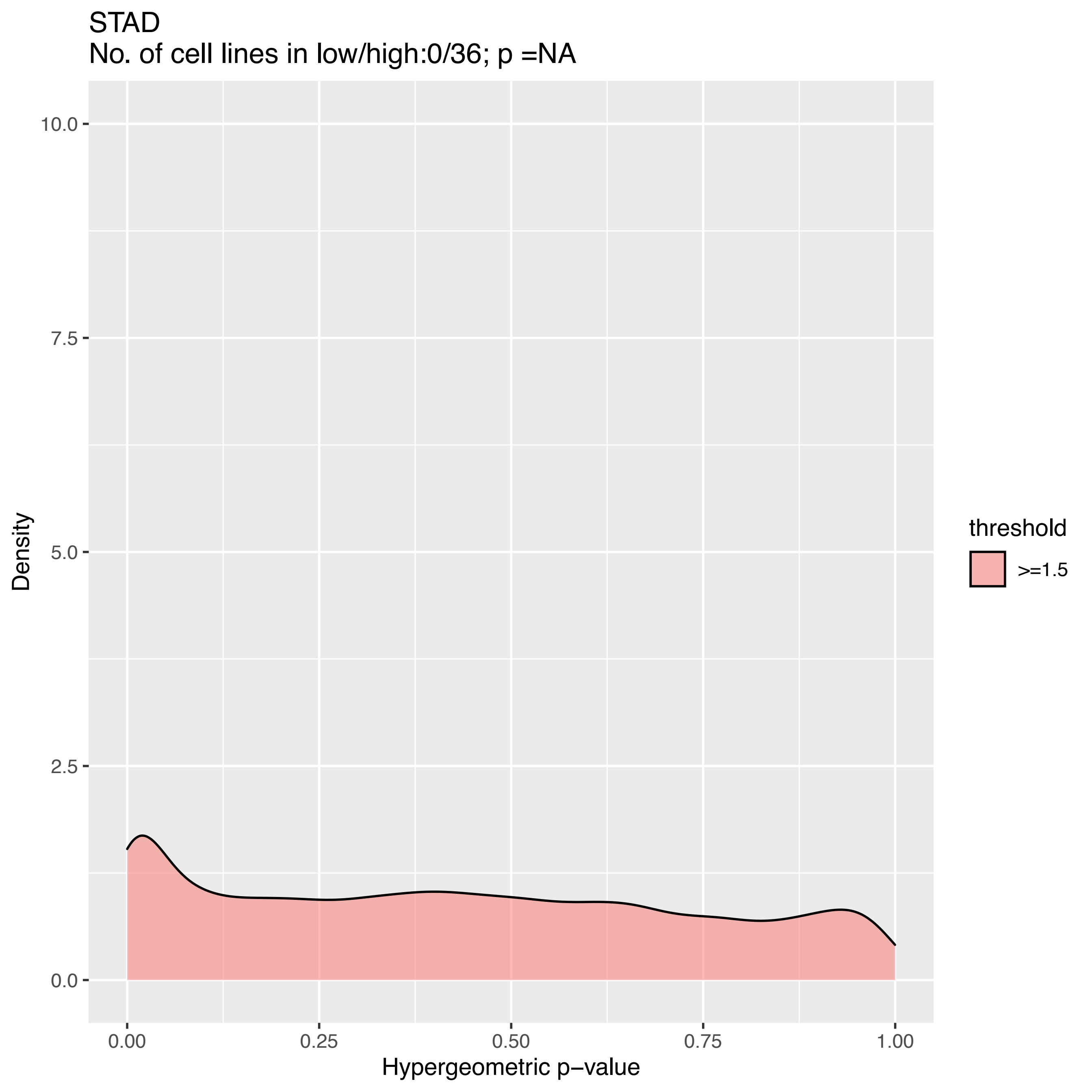

STAD
No. of cell lines in low/high:0/36; p =NA
Density
10.0
7.5
5.0
2.5
0.0
0.00
0.25
0.50
0.75
1.00
Hypergeometric p−value
threshold
>=1.5

STAD
No. of cell lines in low/high:0/36; p =NA

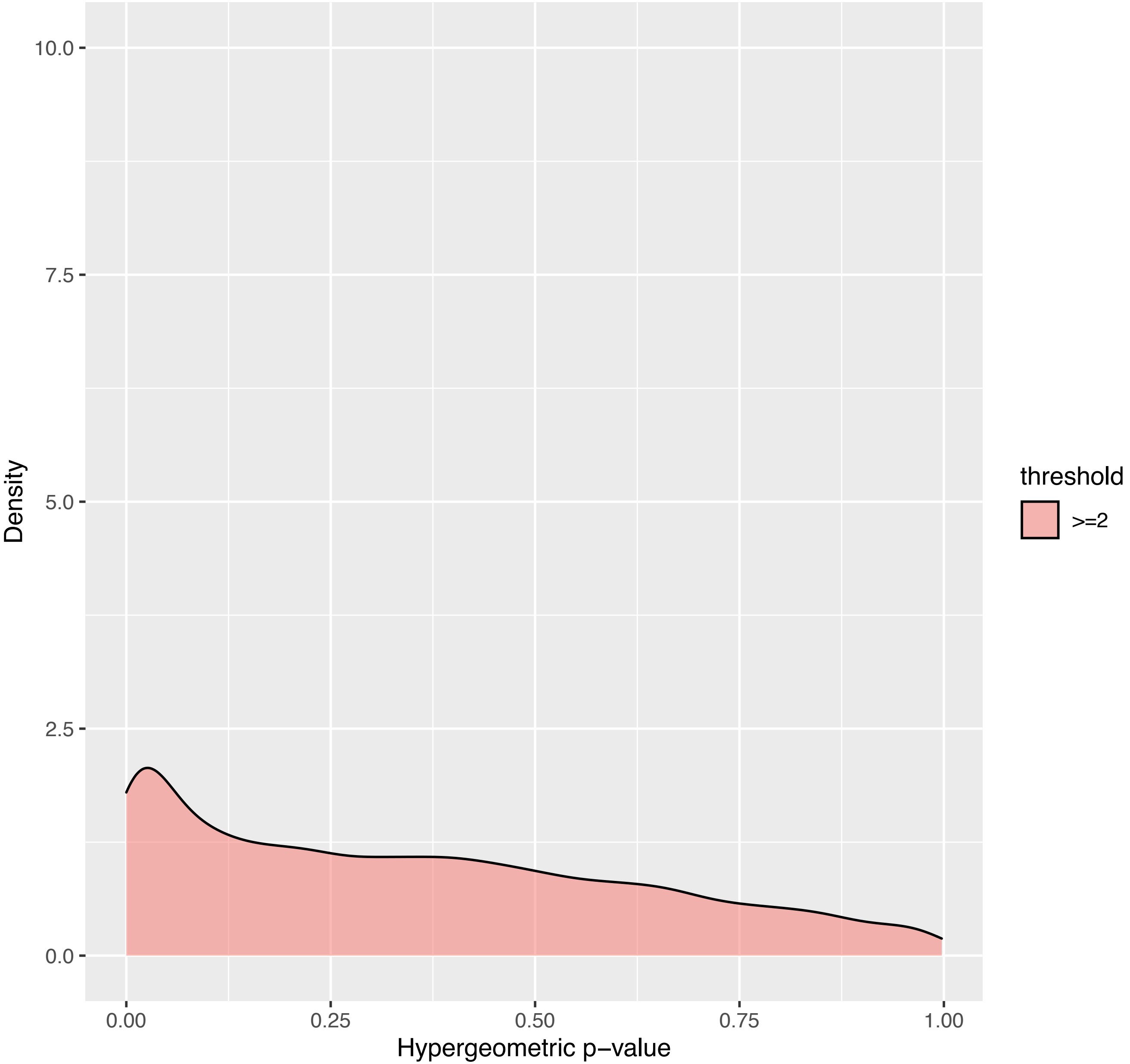

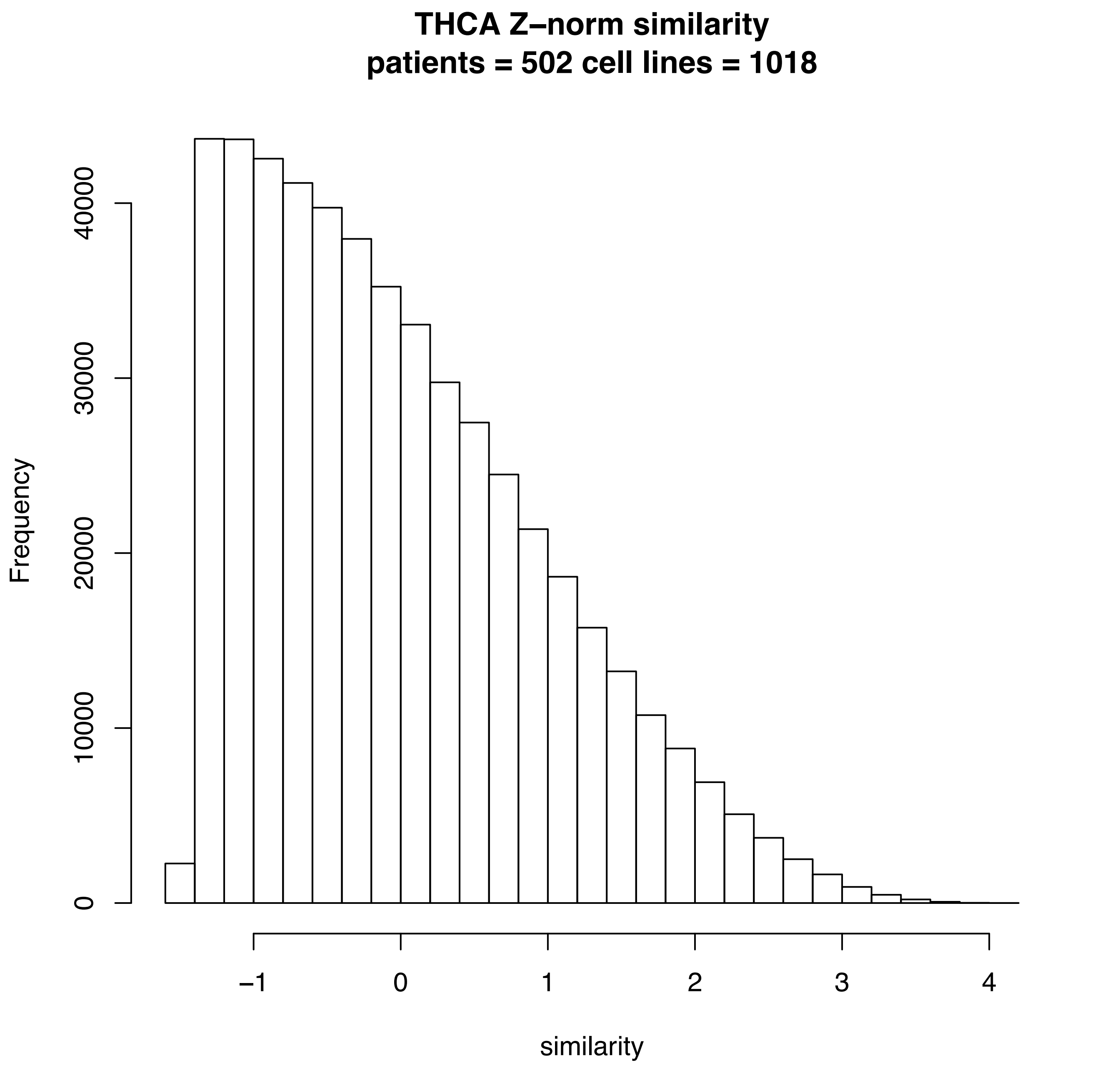
THCA Z–norm similarity
patients = 502 cell lines = 1018
Frequency
0
10000
20000
30000
40000
–1
0
1
2
3
4
similarity

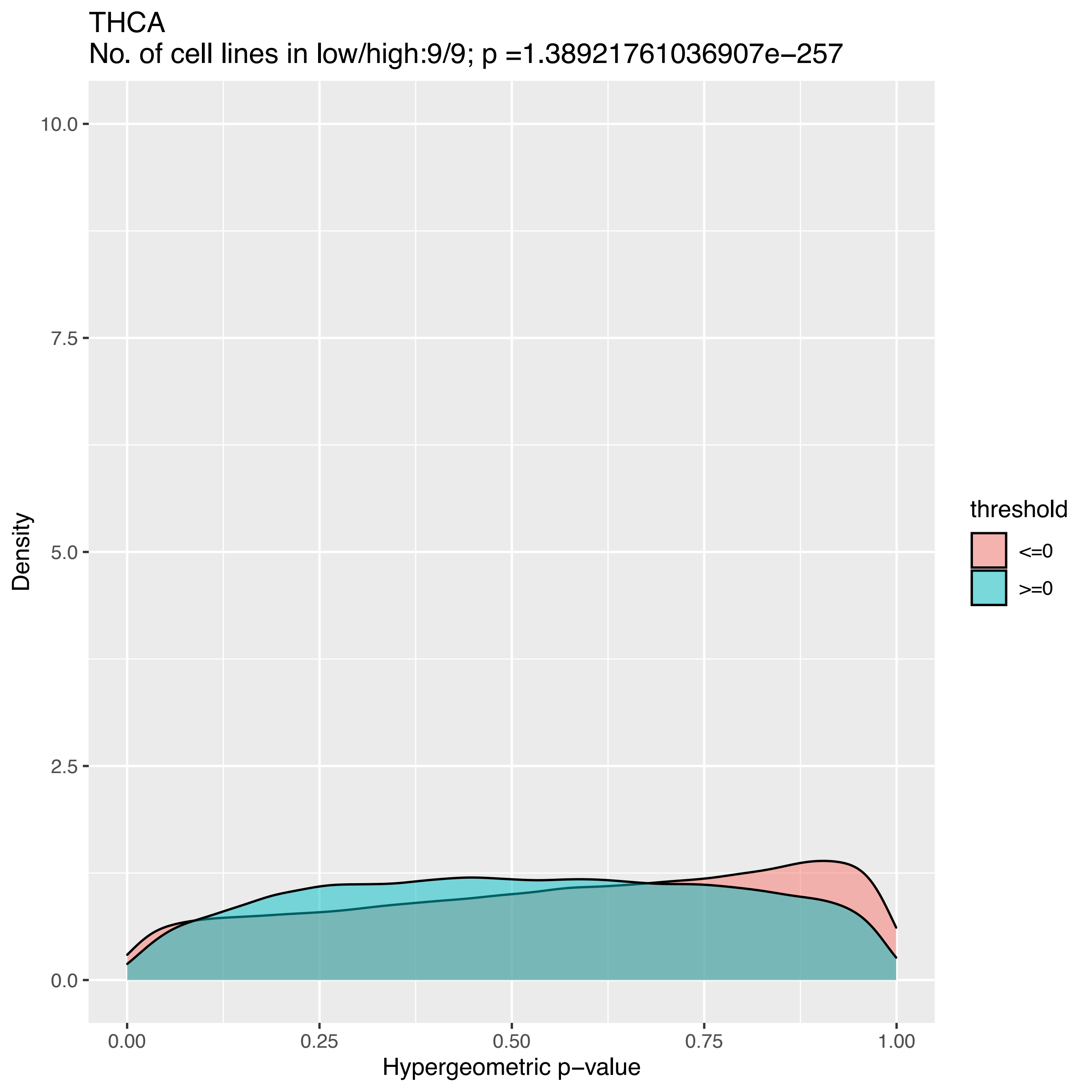
THCA
No. of cell lines in low/high:9/9; p =1.38921761036907e−257
10.0
7.5
5.0
2.5
0.0
Density
0.00
0.25
0.50
0.75
1.00
Hypergeometric p−value
threshold
<=0
>=0

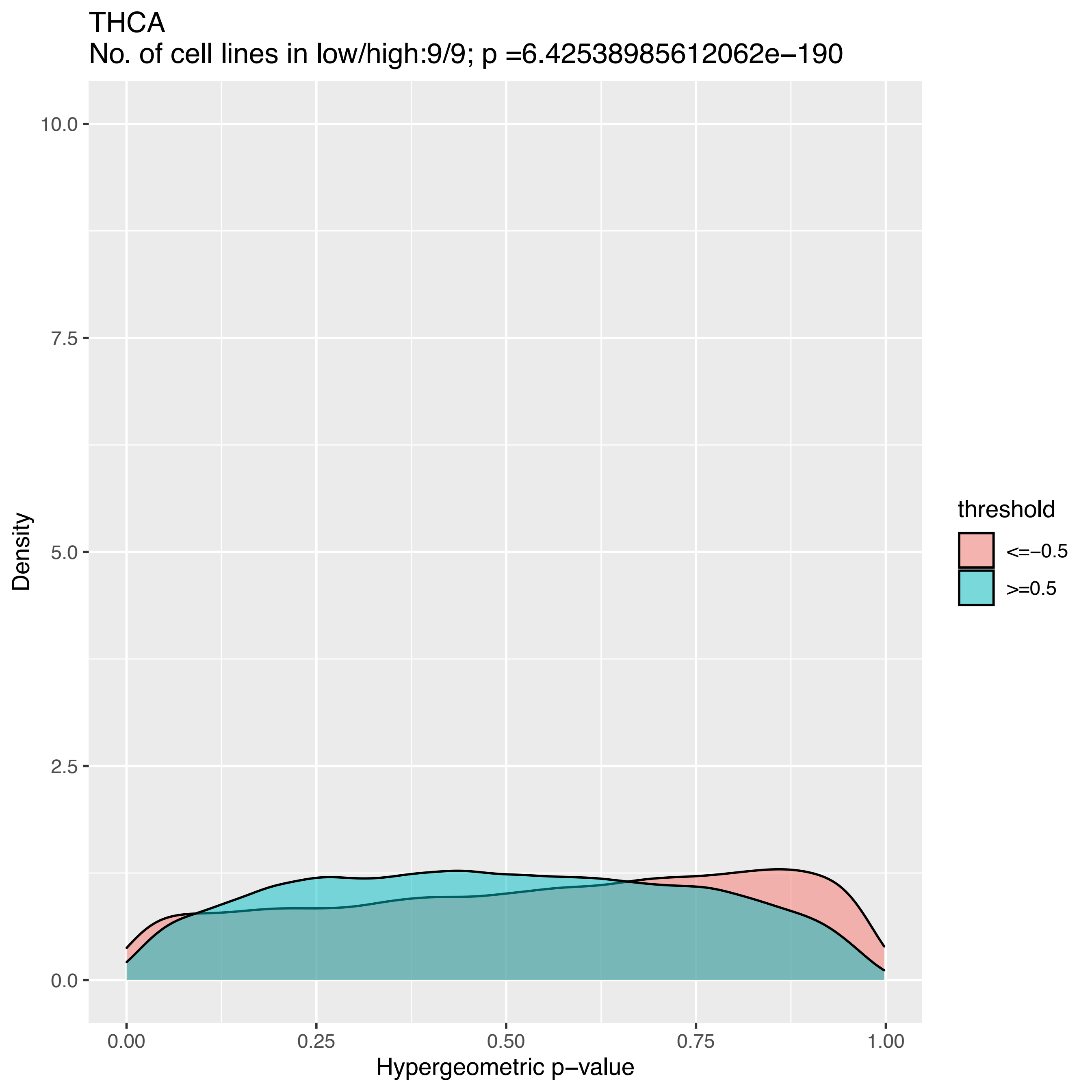
THCA
No. of cell lines in low/high:9/9; p =6.42538985612062e−190
Density
10.0
7.5
5.0
2.5
0.0
0.00
0.25
0.50
0.75
1.00
Hypergeometric p−value
threshold
<=−0.5
>=0.5

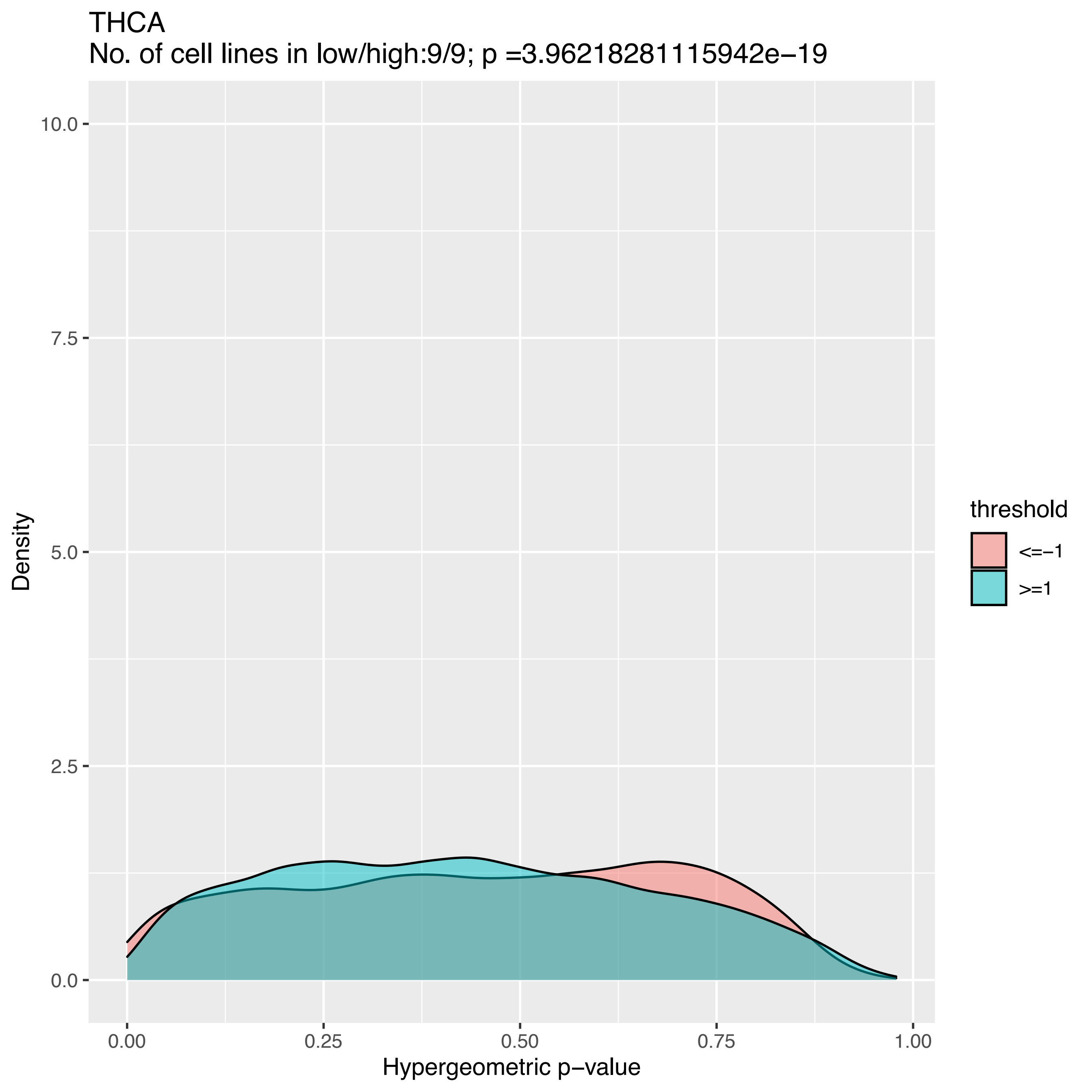
THCA
No. of cell lines in low/high:9/9; p =3.96218281115942e−19
Density
Hypergeometric p−value
0.0
2.5
5.0
7.5
10.0
0.00
0.25
0.50
0.75
1.00
threshold
<=−1
>=1

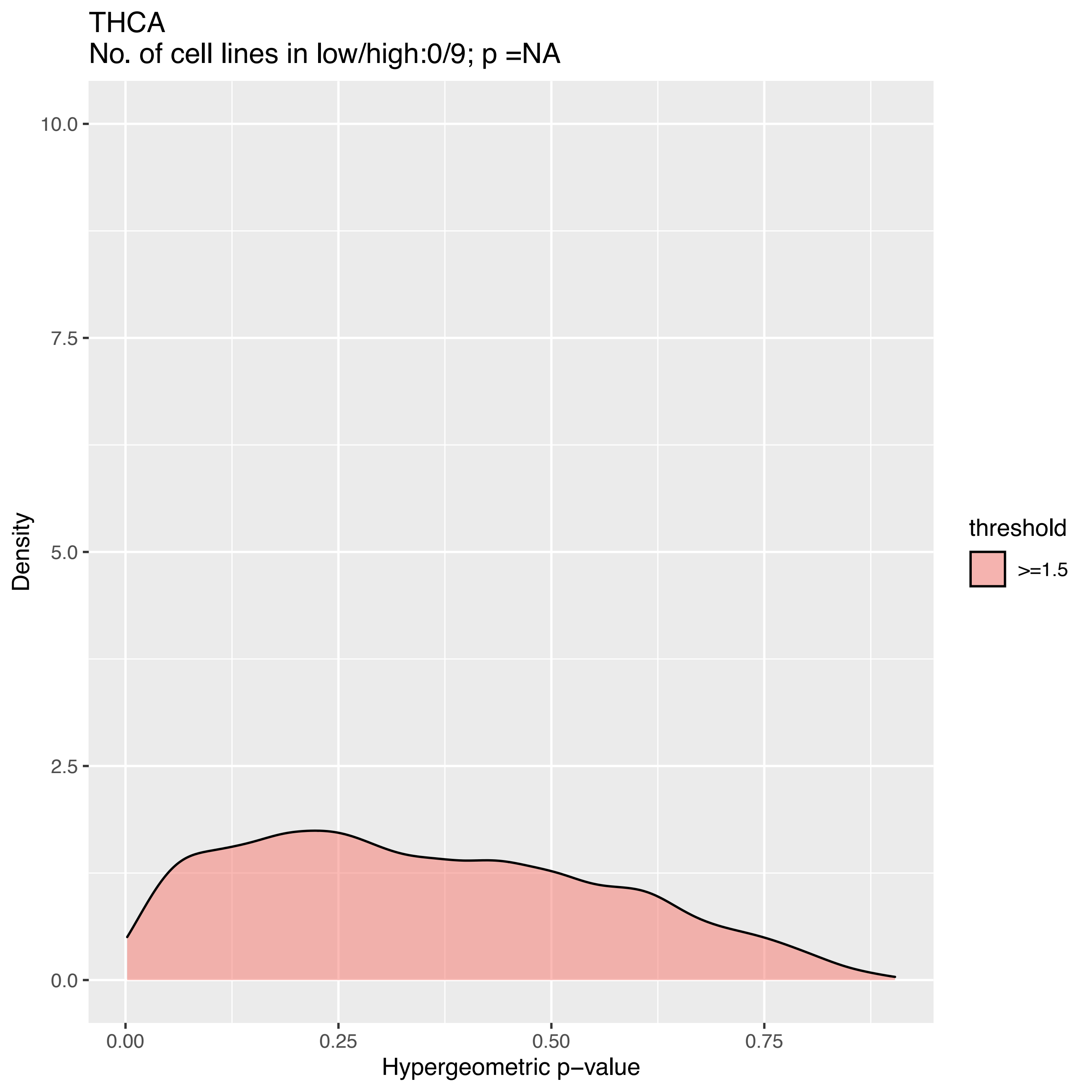
THCA
No. of cell lines in low/high:0/9; p =NA
10.0
7.5
5.0
2.5
0.0
Density
0.00
0.25
0.50
0.75
Hypergeometric p−value
threshold
>=1.5

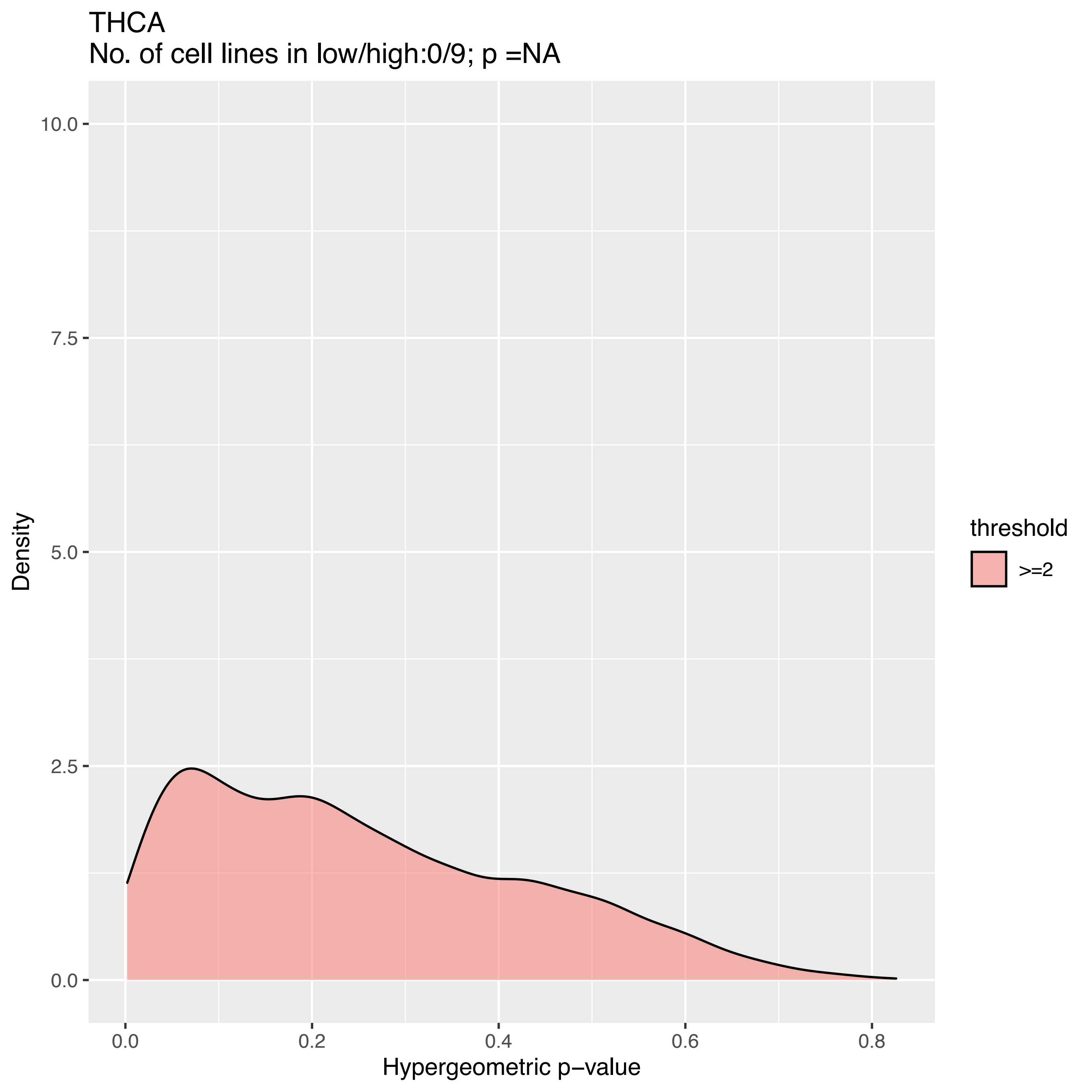
THCA
No. of cell lines in low/high:0/9; p =NA
10.0
7.5
5.0
2.5
0.0
Density
0.0
0.2
0.4
0.6
0.8
Hypergeometric p−value
threshold
>=2

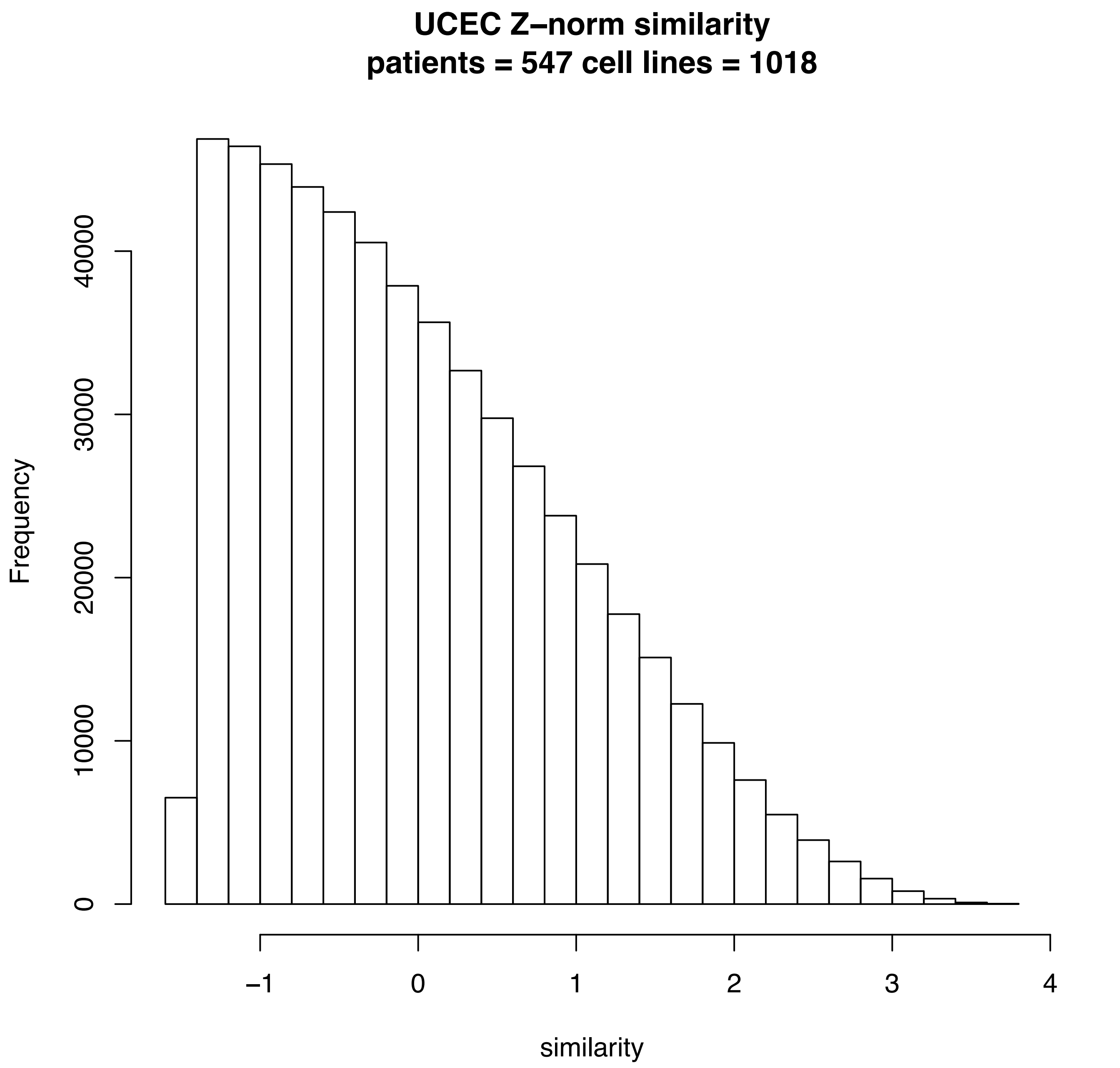
UCEC Z–norm similarity
patients = 547 cell lines = 1018
Frequency
0
10000
20000
30000
40000
−1
0
1
2
3
4
similarity

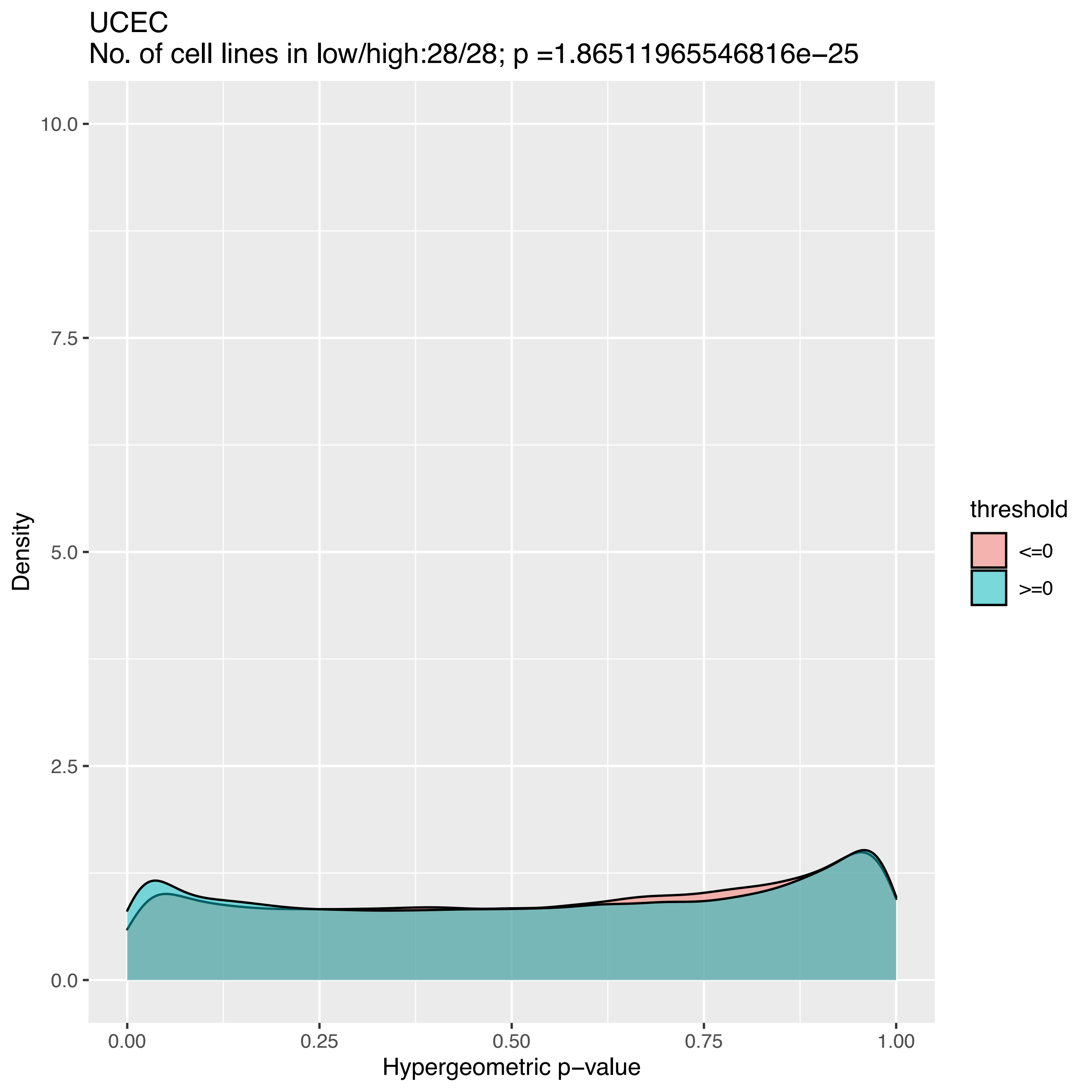

UCEC
No. of cell lines in low/high:28/28; p =1.86511965546816e−25
10.0
7.5
5.0
2.5
0.0
Density
0.00
0.25
0.50
0.75
1.00
Hypergeometric p−value
threshold
<=0
>=0

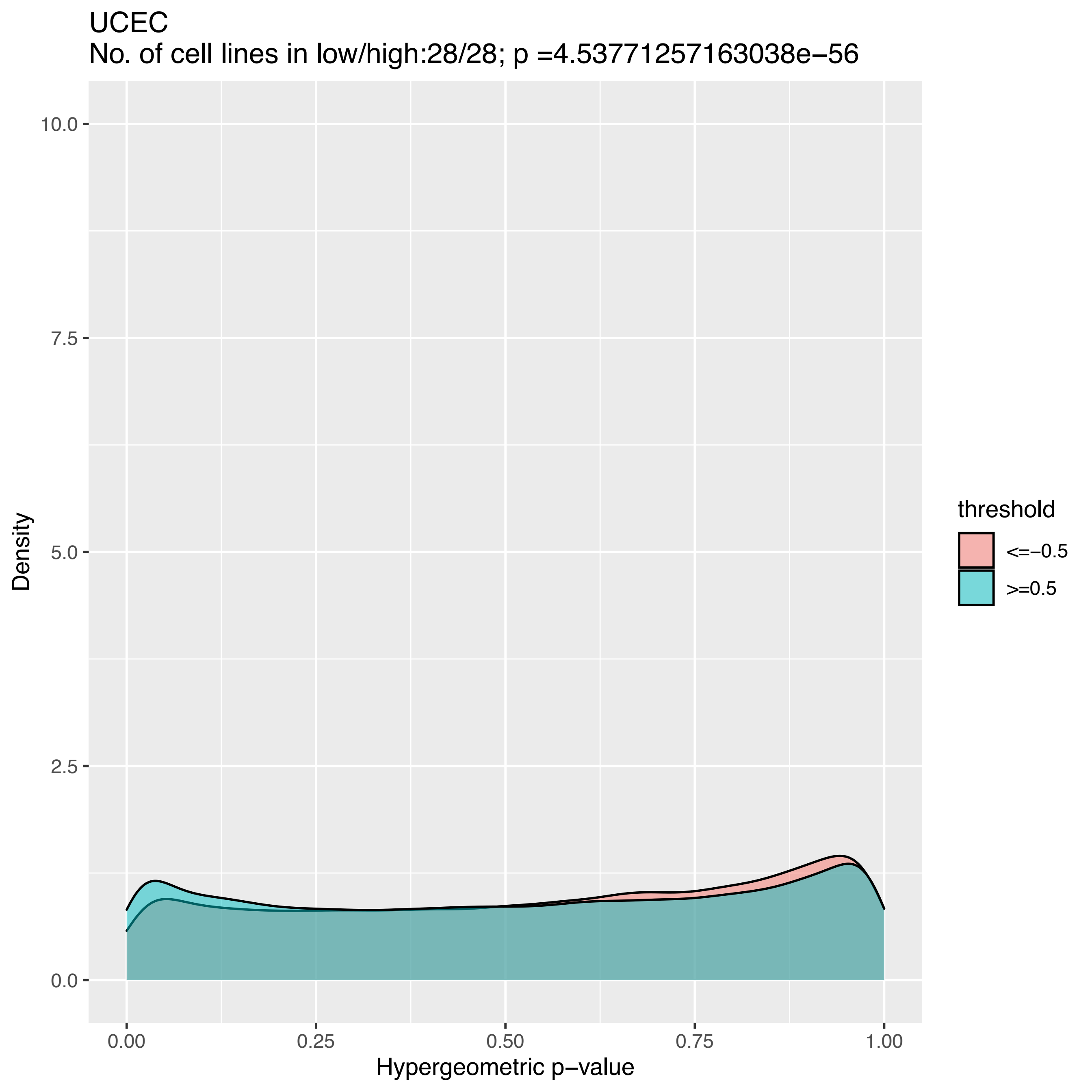
UCEC
No. of cell lines in low/high:28/28; p =4.53771257163038e−56
Density
10.0
7.5
5.0
2.5
0.0
0.00
0.25
0.50
0.75
1.00
Hypergeometric p−value
threshold
<=−0.5
>=0.5

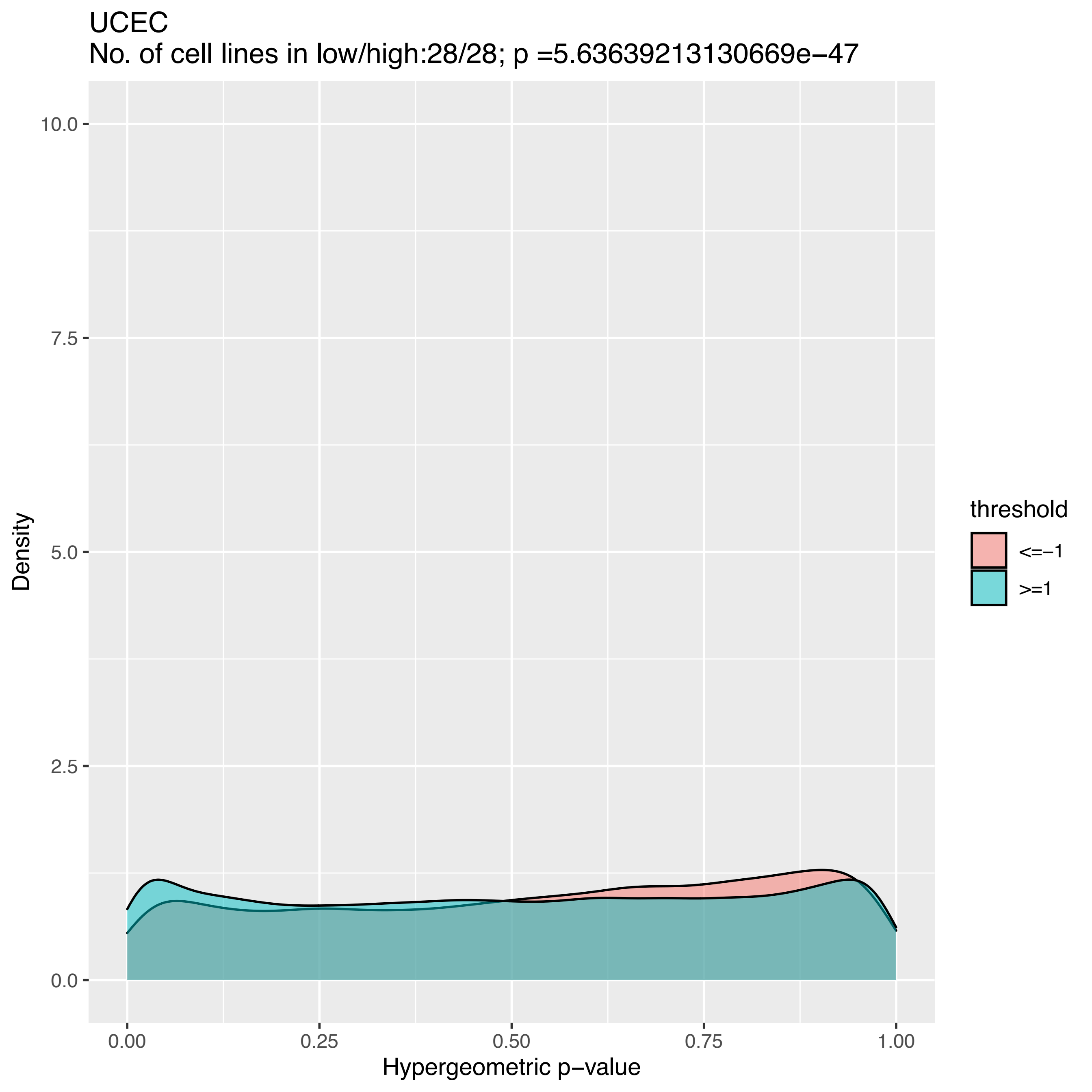
UCEC
No. of cell lines in low/high:28/28; p =5.63639213130669e−47
Density
10.0
7.5
5.0
2.5
0.0
0.00
0.25
0.50
0.75
1.00
Hypergeometric p−value
threshold
<=−1
>=1

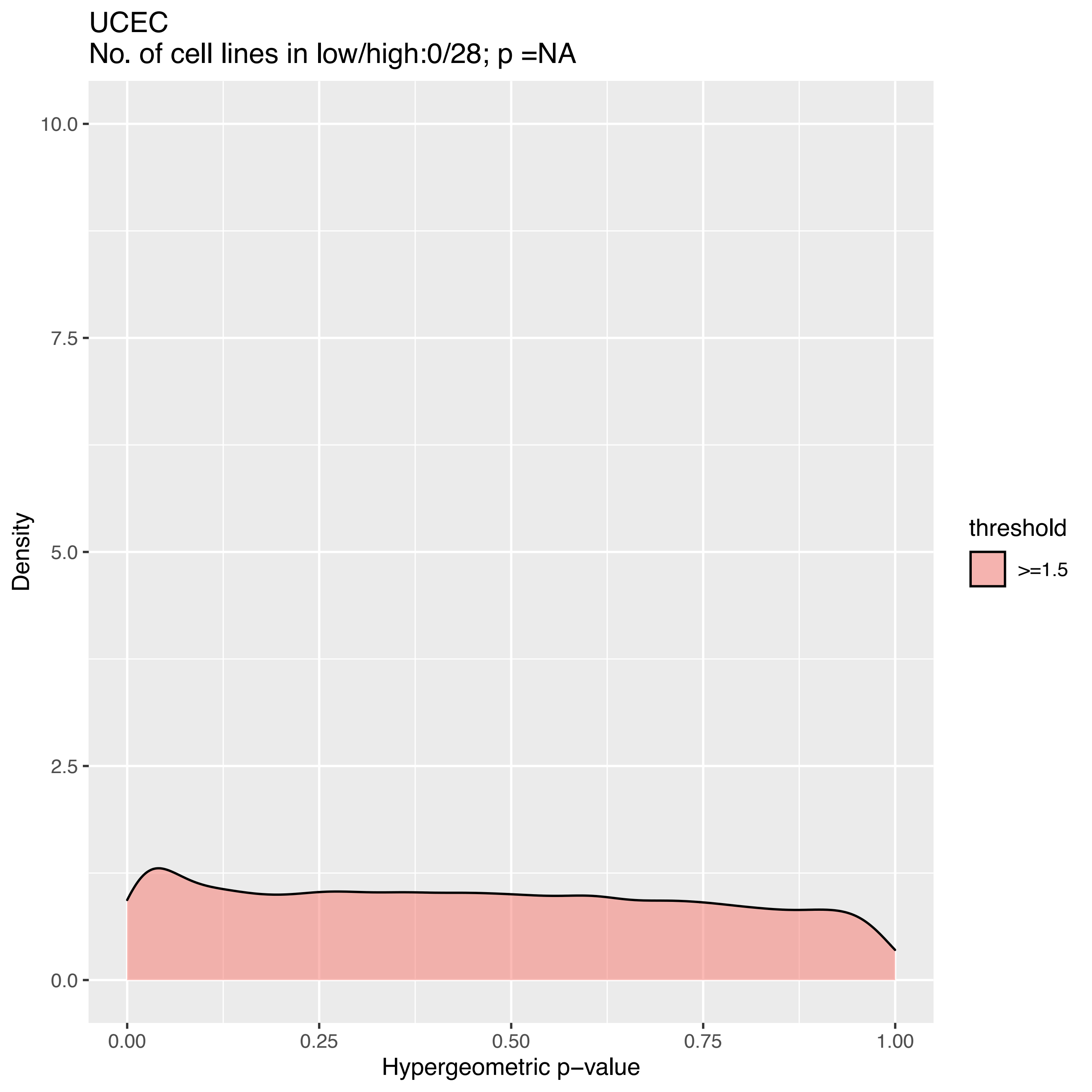
UCEC
No. of cell lines in low/high:0/28; p =NA
Density
10.0
7.5
5.0
2.5
0.0
0.00
0.25
0.50
0.75
1.00
Hypergeometric p−value
threshold
>=1.5

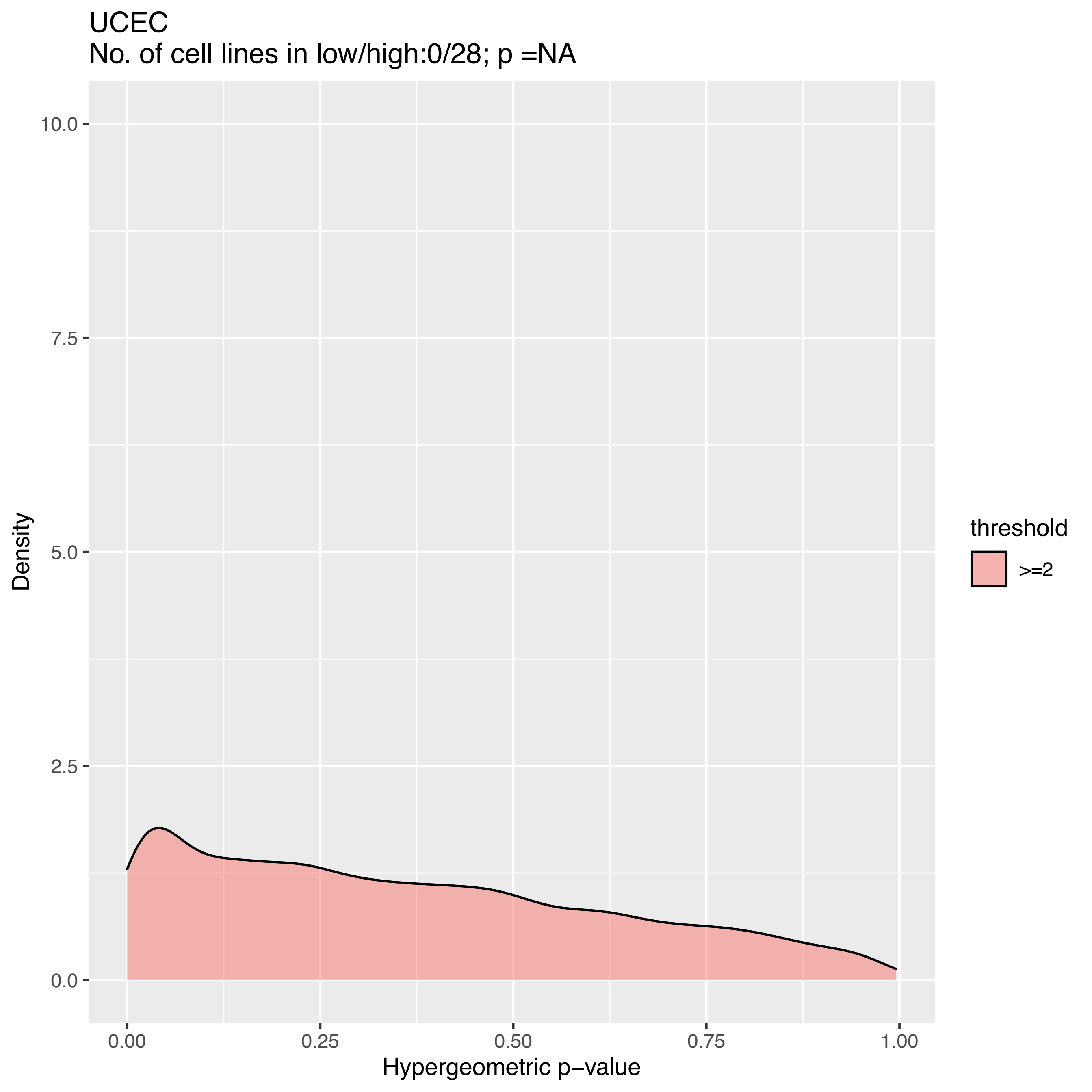

UCEC
No. of cell lines in low/high:0/28; p =NA
Density
10.0
7.5
5.0
2.5
0.0
0.00
0.25
0.50
0.75
1.00
Hypergeometric p−value
threshold
>=2

### TC analysis

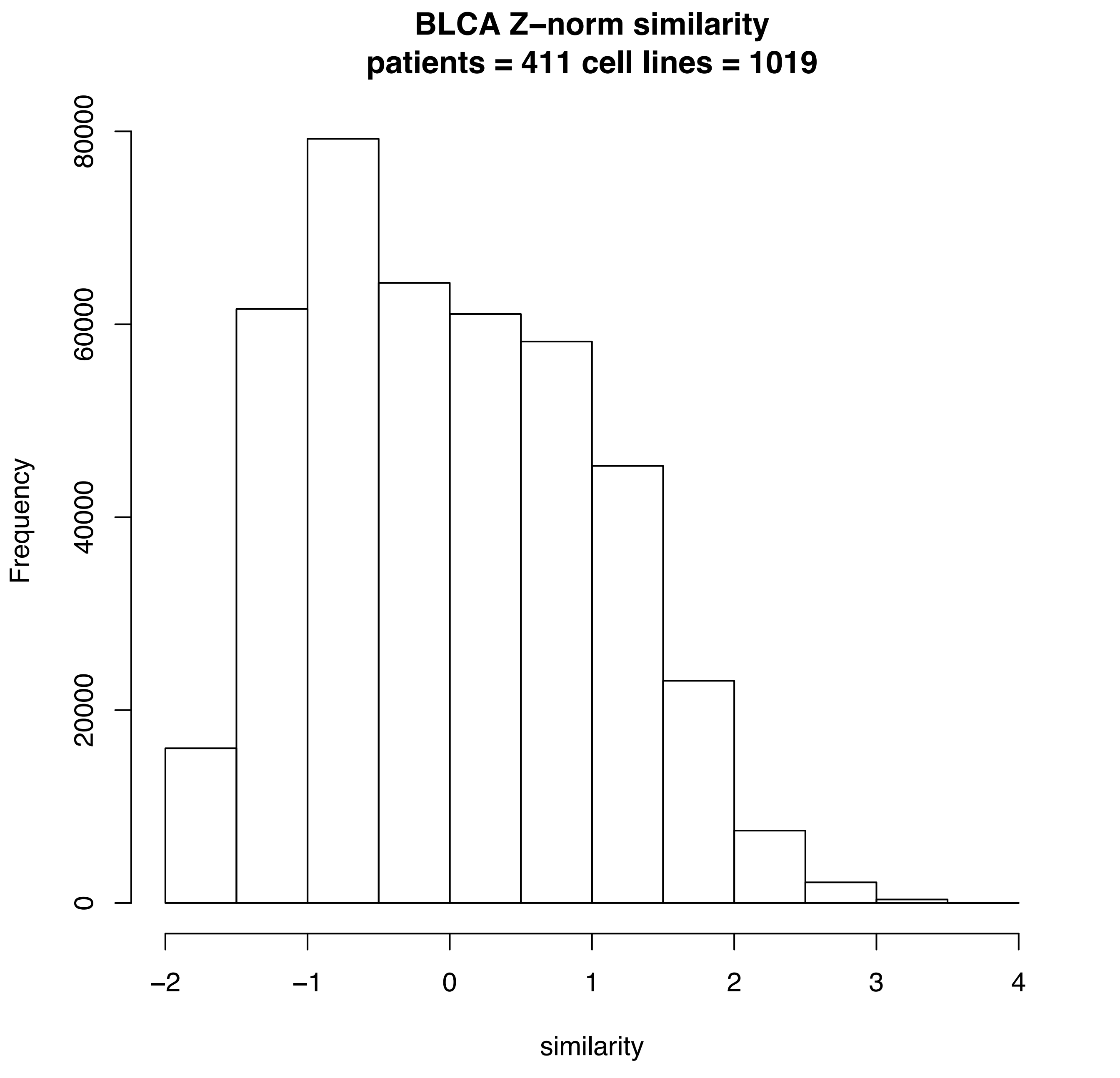
BLCA Z–norm similarity
patients = 411 cell lines = 1019
Frequency
0
20000
40000
60000
80000
–2
–1
0
1
2
3
4
similarity

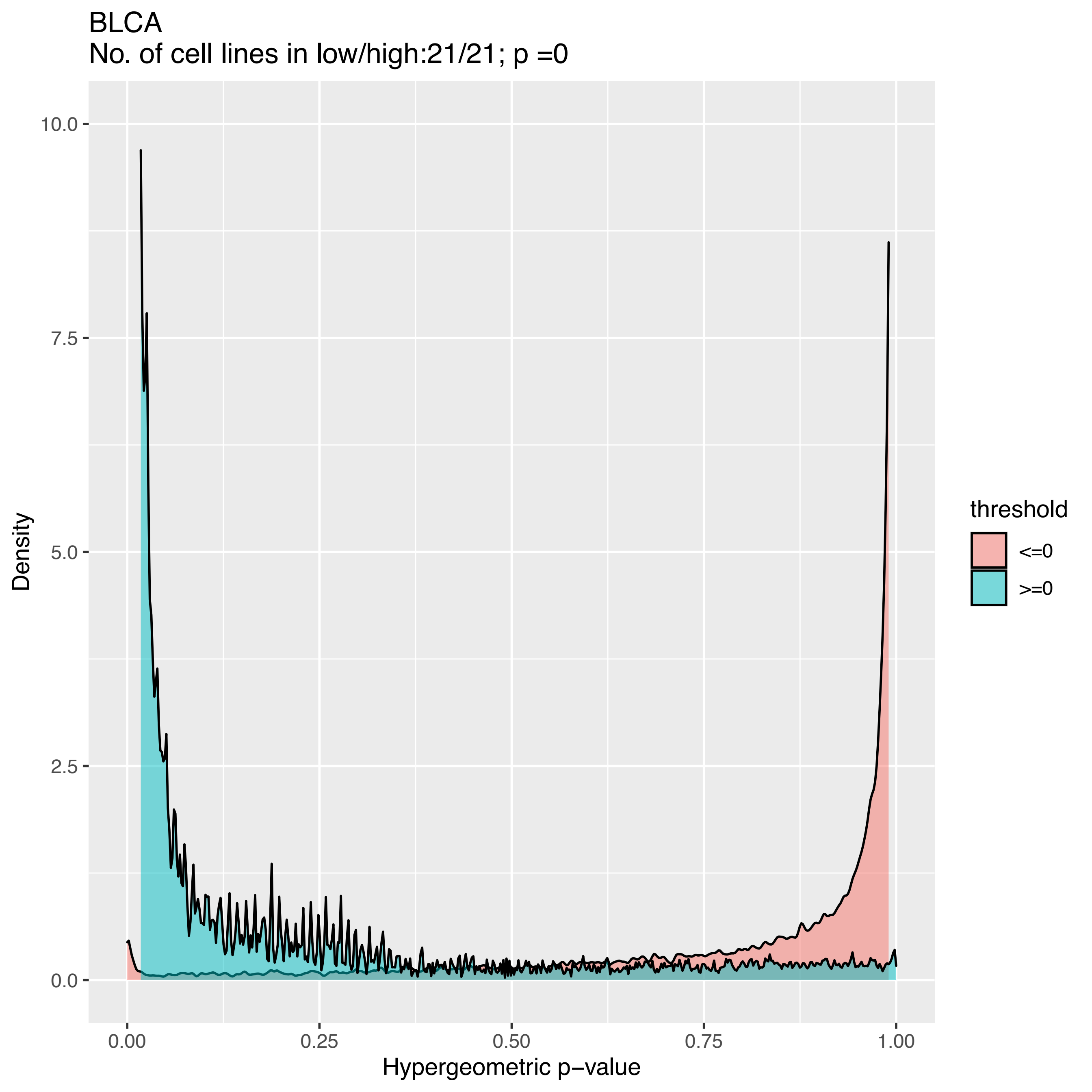

BLCA
No. of cell lines in low/high:21/21; p =0
Density
10.0
7.5
5.0
2.5
0.0
0.00
0.25
0.50
0.75
1.00
Hypergeometric p−value
threshold
<=0
>=0

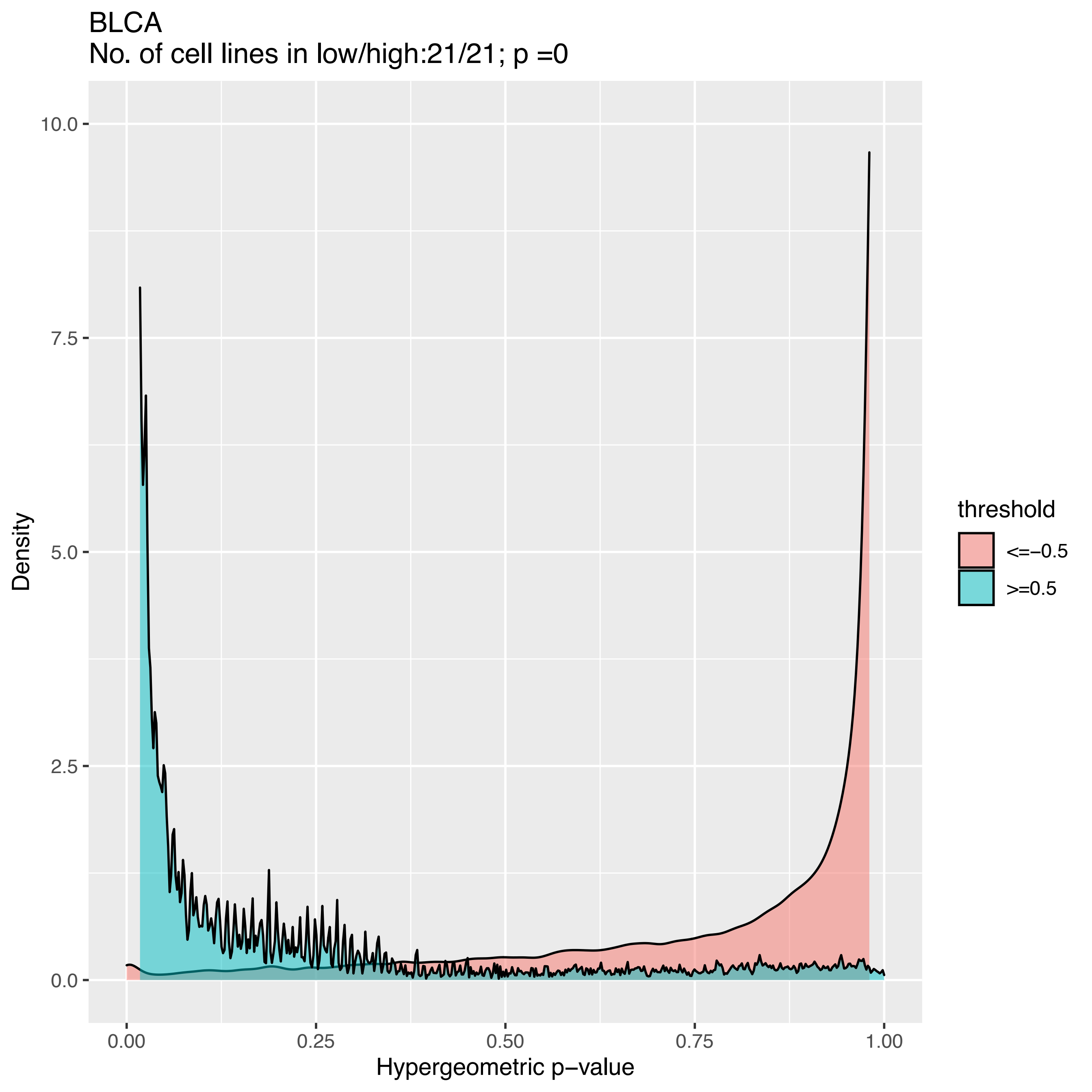
BLCA
No. of cell lines in low/high:21/21; p =0
10.0
7.5
5.0
2.5
0.0
Density
0.00
0.25
0.50
0.75
1.00
Hypergeometric p−value
threshold
<=−0.5
>=0.5

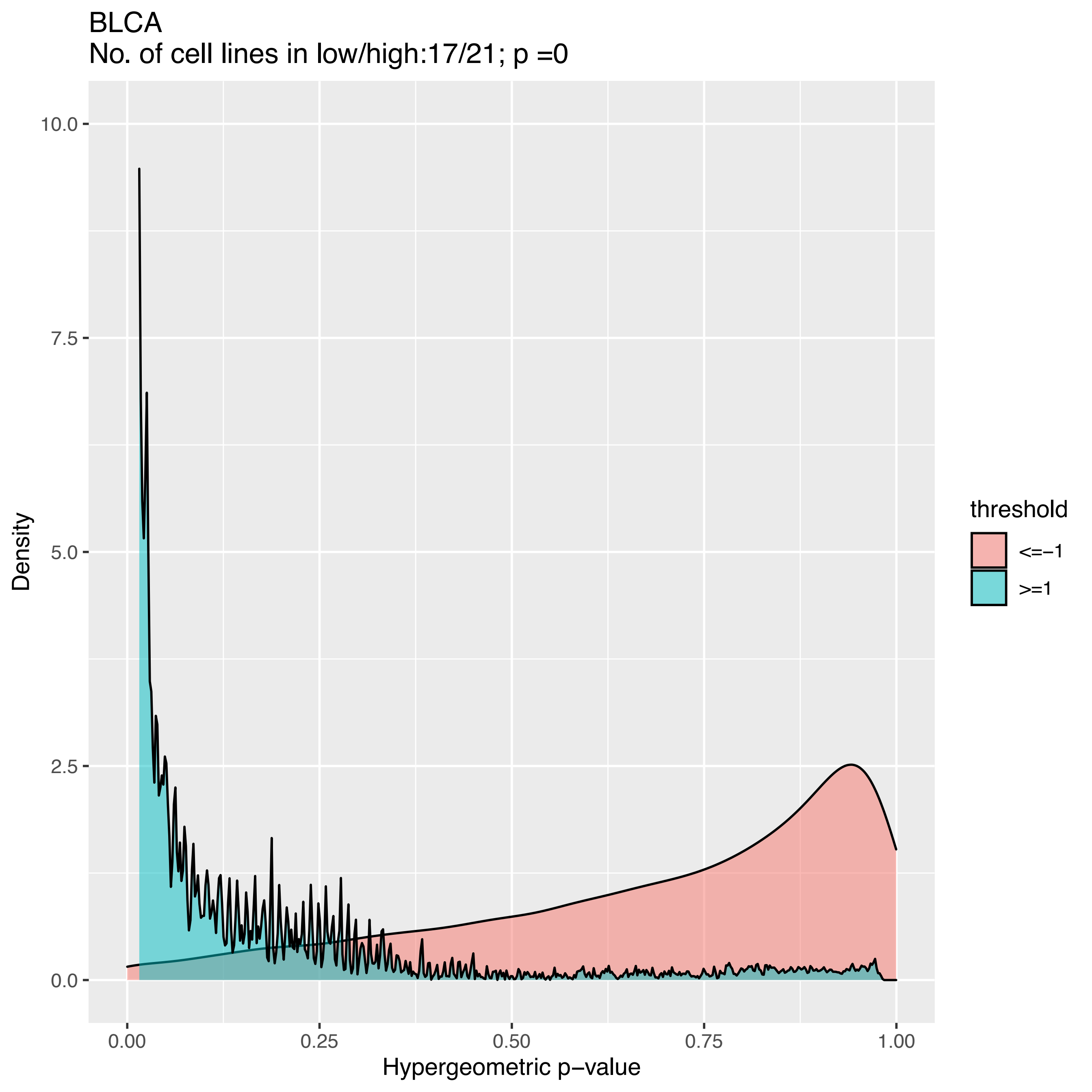
BLCA
No. of cell lines in low/high:17/21; p =0
Density
Hypergeometric p−value
10.0
7.5
5.0
2.5
0.0
0.00
0.25
0.50
0.75
1.00
threshold
<=−1
>=1

BLCA
No. of cell lines in low/high:8/18; p =3.2793858548964e−141

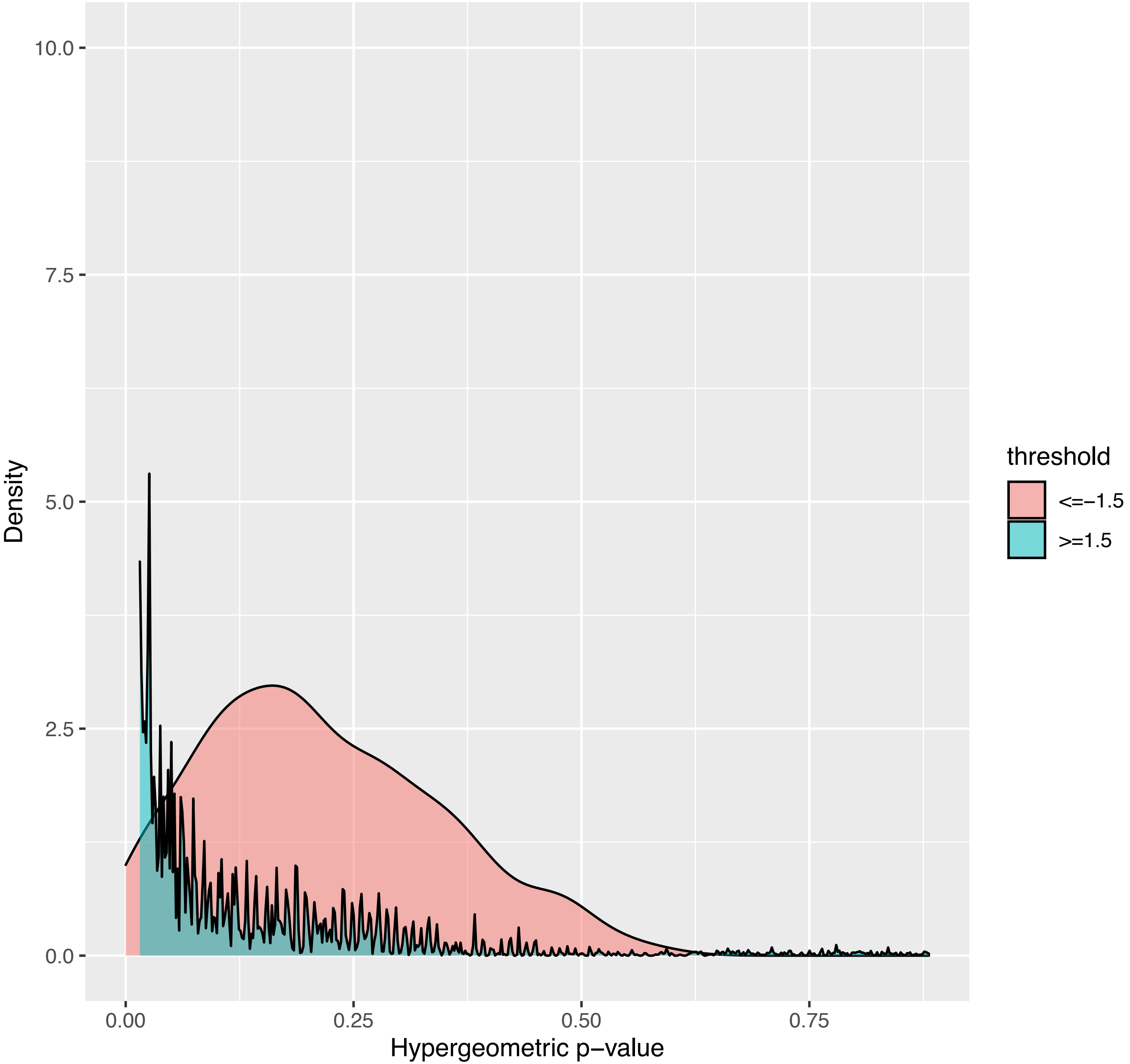

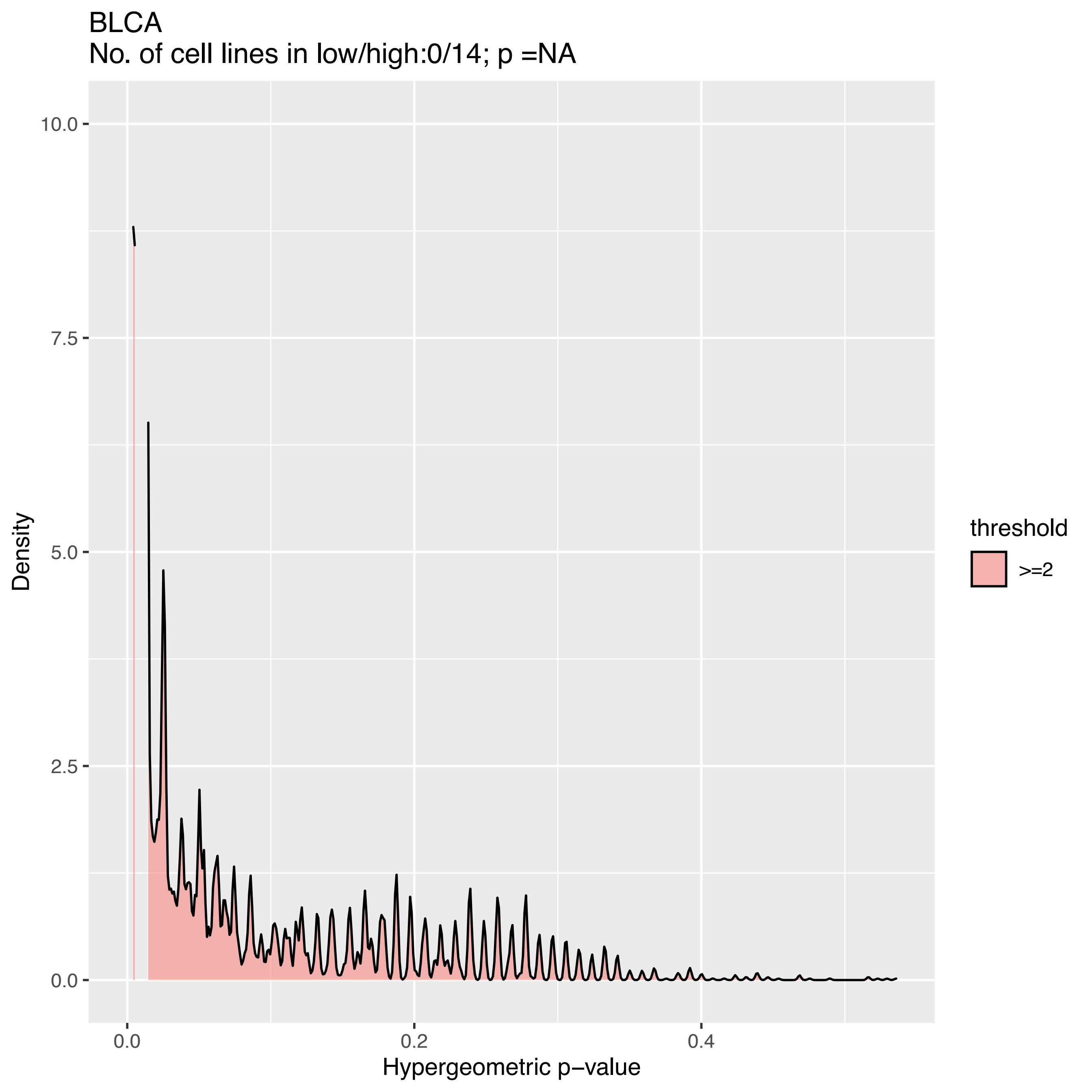
BLCA
No. of cell lines in low/high:0/14; p =NA
Density
10.0
7.5
5.0
2.5
0.0
0.0
0.2
0.4
Hypergeometric p−value
threshold
>=2

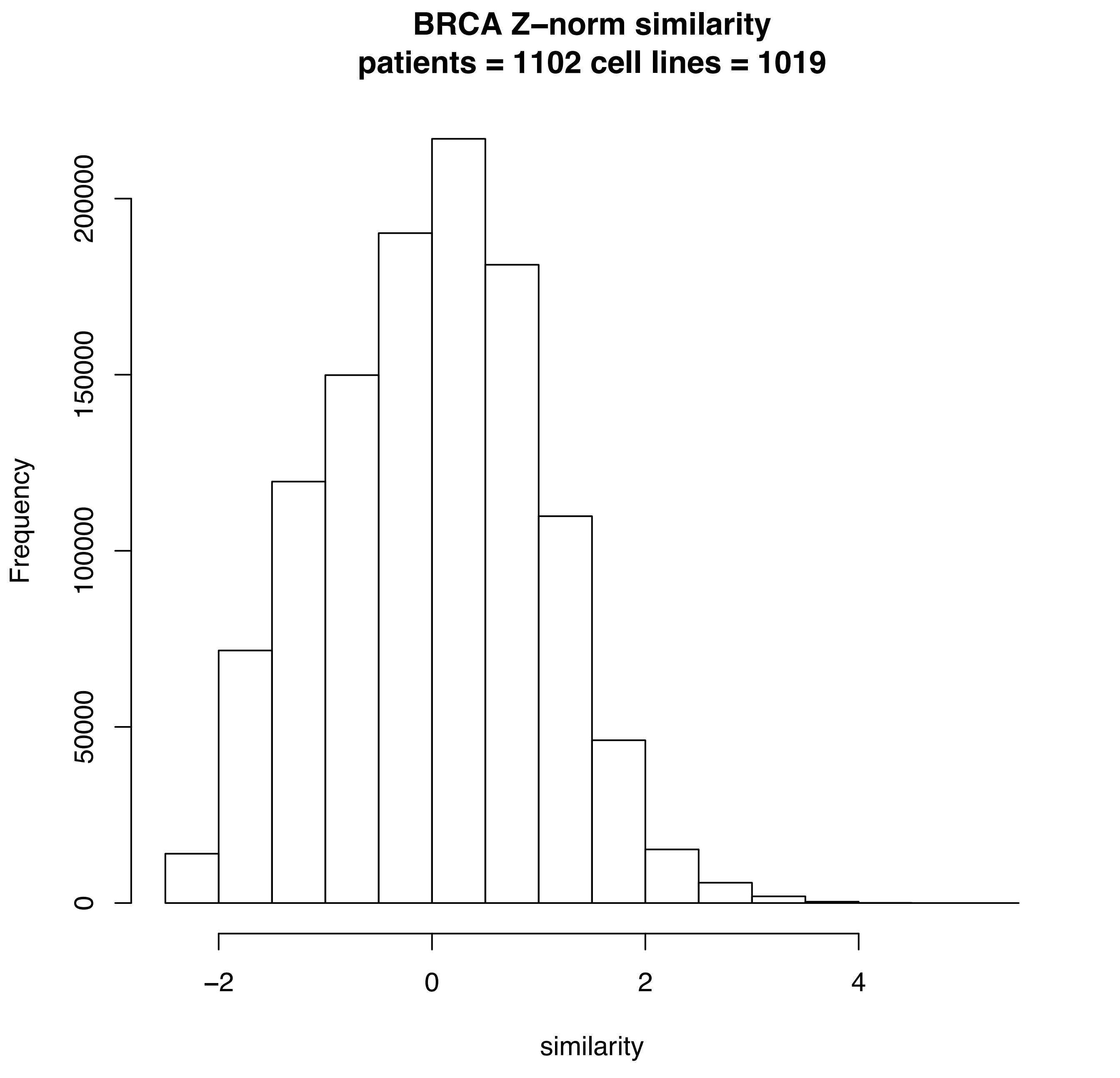
BRCA Z–norm similarity
patients = 1102 cell lines = 1019
Frequency
0
50000
100000
150000
200000
−2
0
2
4
similarity

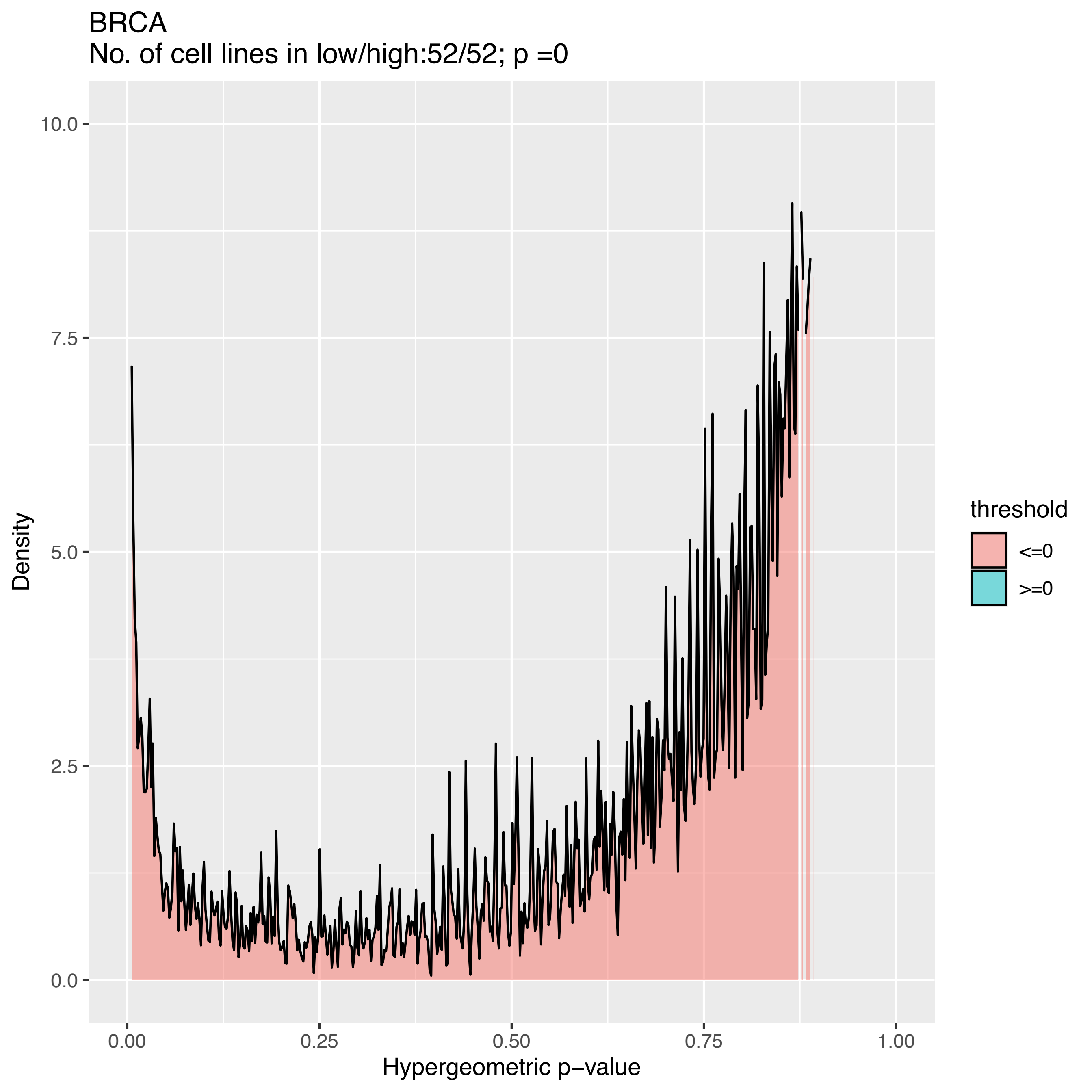
BRCA
No. of cell lines in low/high:52/52; p =0
Density
10.0
7.5
5.0
2.5
0.0
0.00
0.25
0.50
0.75
1.00
Hypergeometric p−value
threshold
<=0
>=0

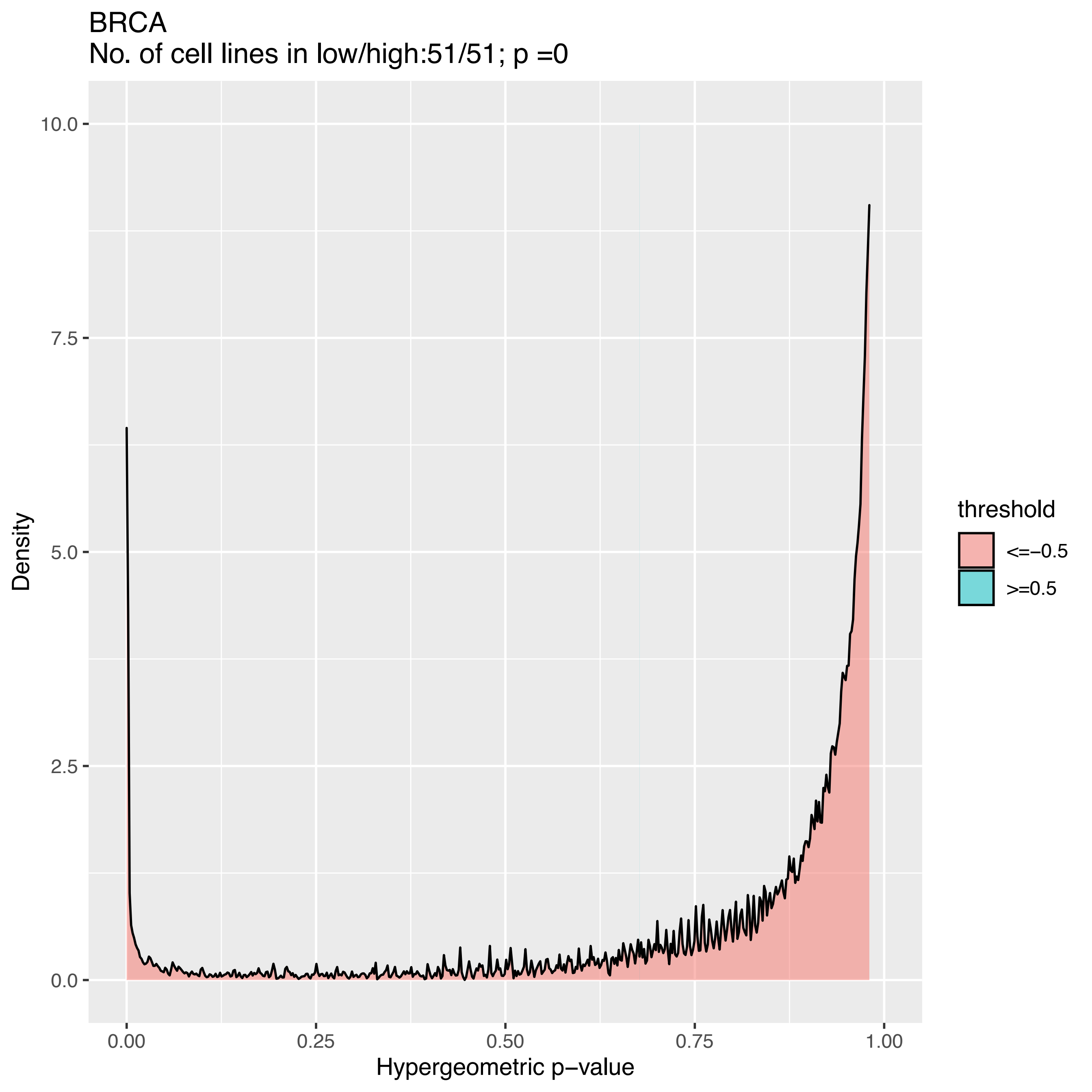
BRCA
No. of cell lines in low/high:51/51; p =0
Density
10.0
7.5
5.0
2.5
0.0
0.00
0.25
0.50
0.75
1.00
Hypergeometric p−value
threshold
<=−0.5
>=0.5

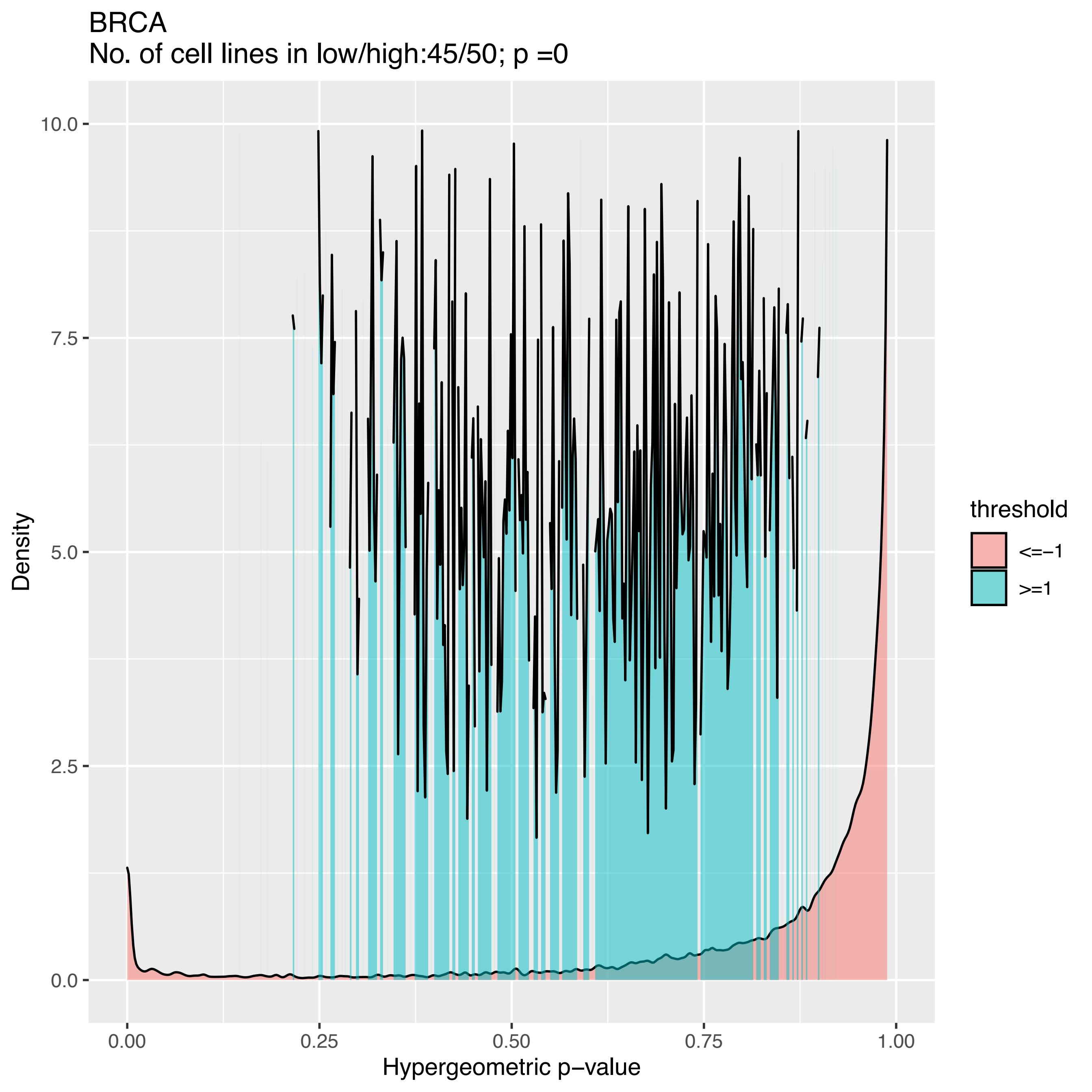
BRCA
No. of cell lines in low/high:45/50; p =0
Density
10.0
7.5
5.0
2.5
0.0
0.00
0.25
0.50
0.75
1.00
Hypergeometric p−value
threshold
<=−1
>=1

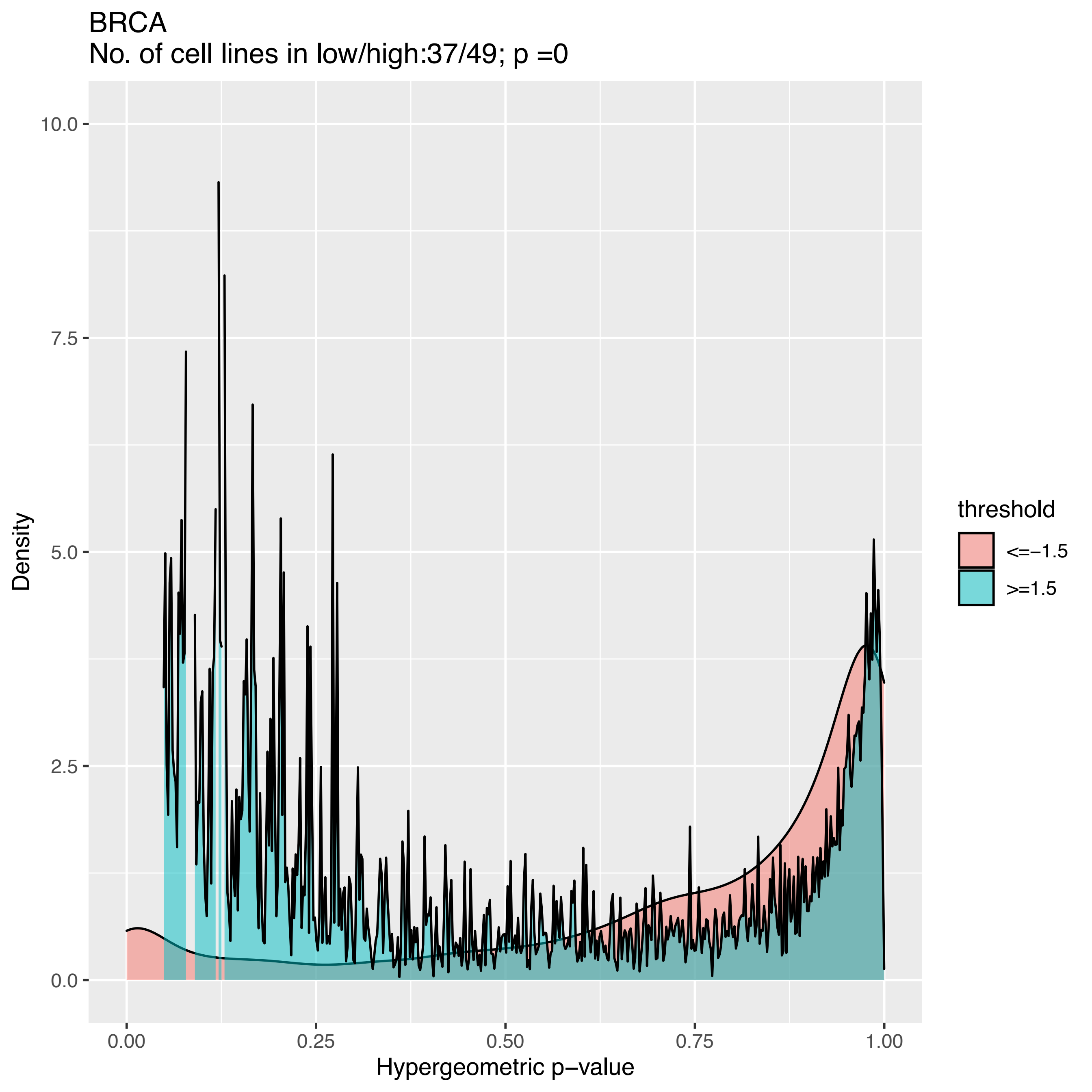

BRCA
No. of cell lines in low/high:37/49; p =0
Density
Hypergeometric p−value
0.0
2.5
5.0
7.5
10.0
0.00
0.25
0.50
0.75
1.00
threshold
<=−1.5
>=1.5

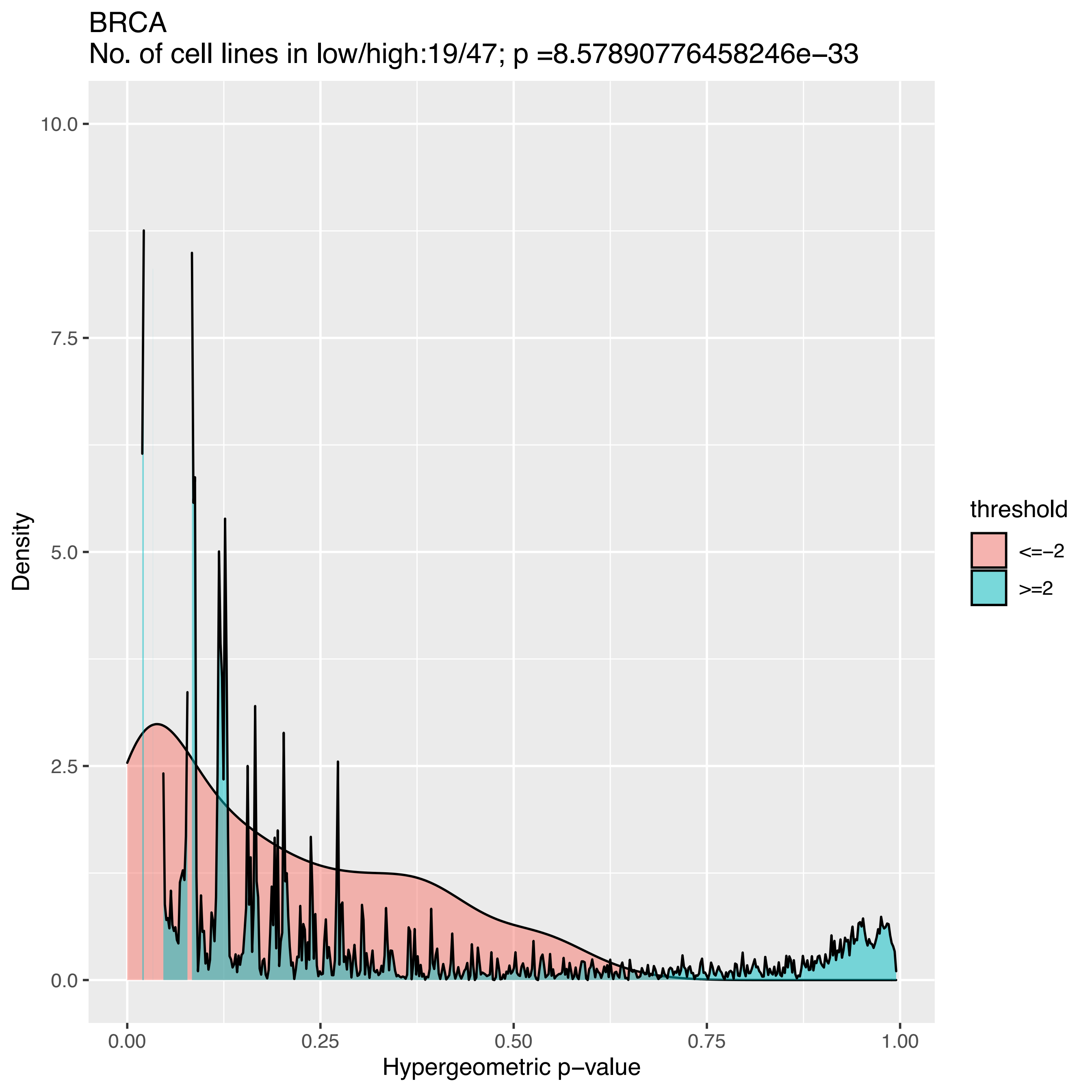

BRCA
No. of cell lines in low/high:19/47; p =8.57890776458246e−33
Density
10.0
7.5
5.0
2.5
0.0
0.00
0.25
0.50
0.75
1.00
Hypergeometric p−value
threshold
<=−2
>=2

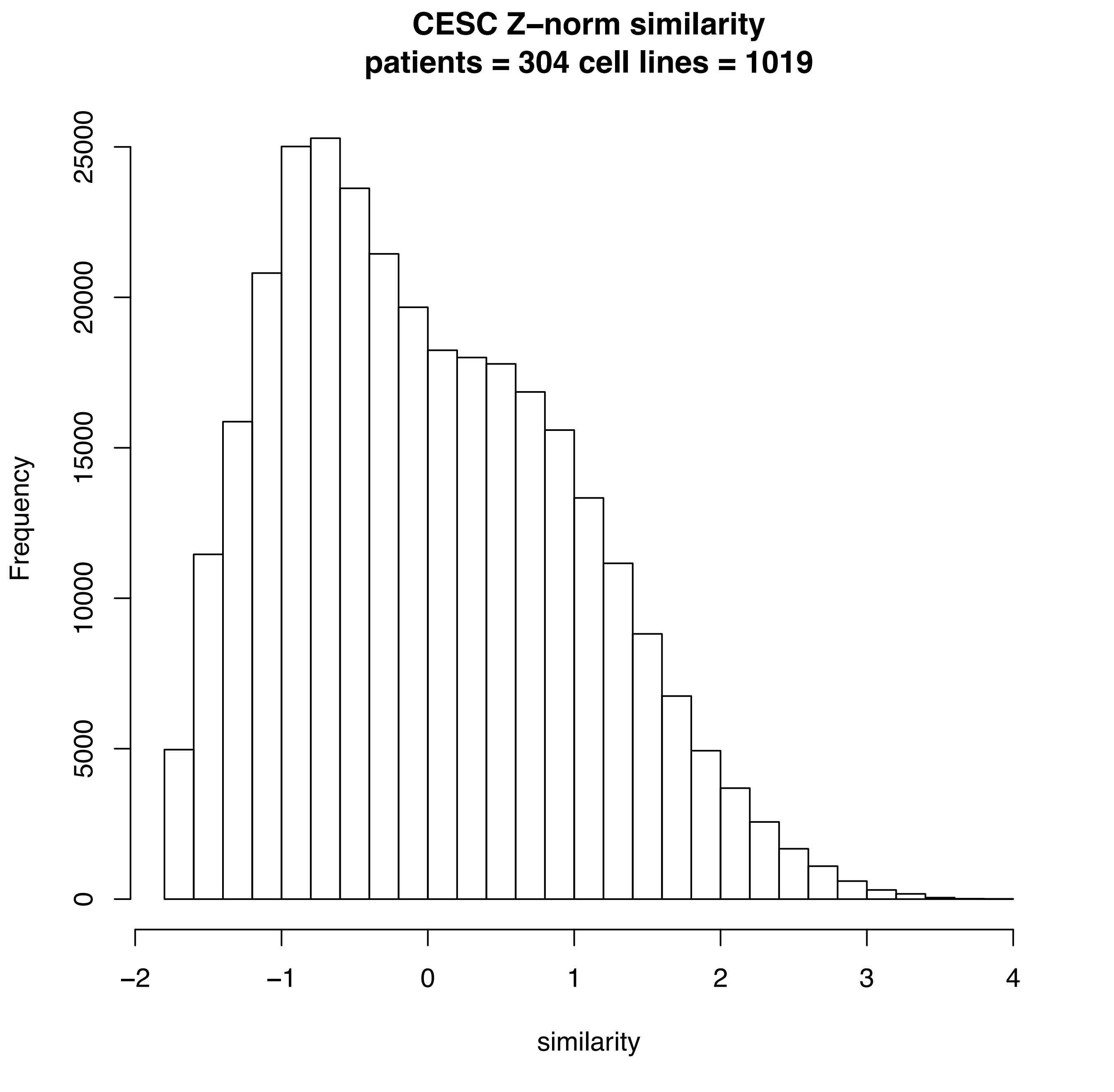
CESC Z–norm similarity
patients = 304 cell lines = 1019
Frequency
0
5000
10000
15000
20000
25000
–2
–1
0
1
2
3
4
similarity

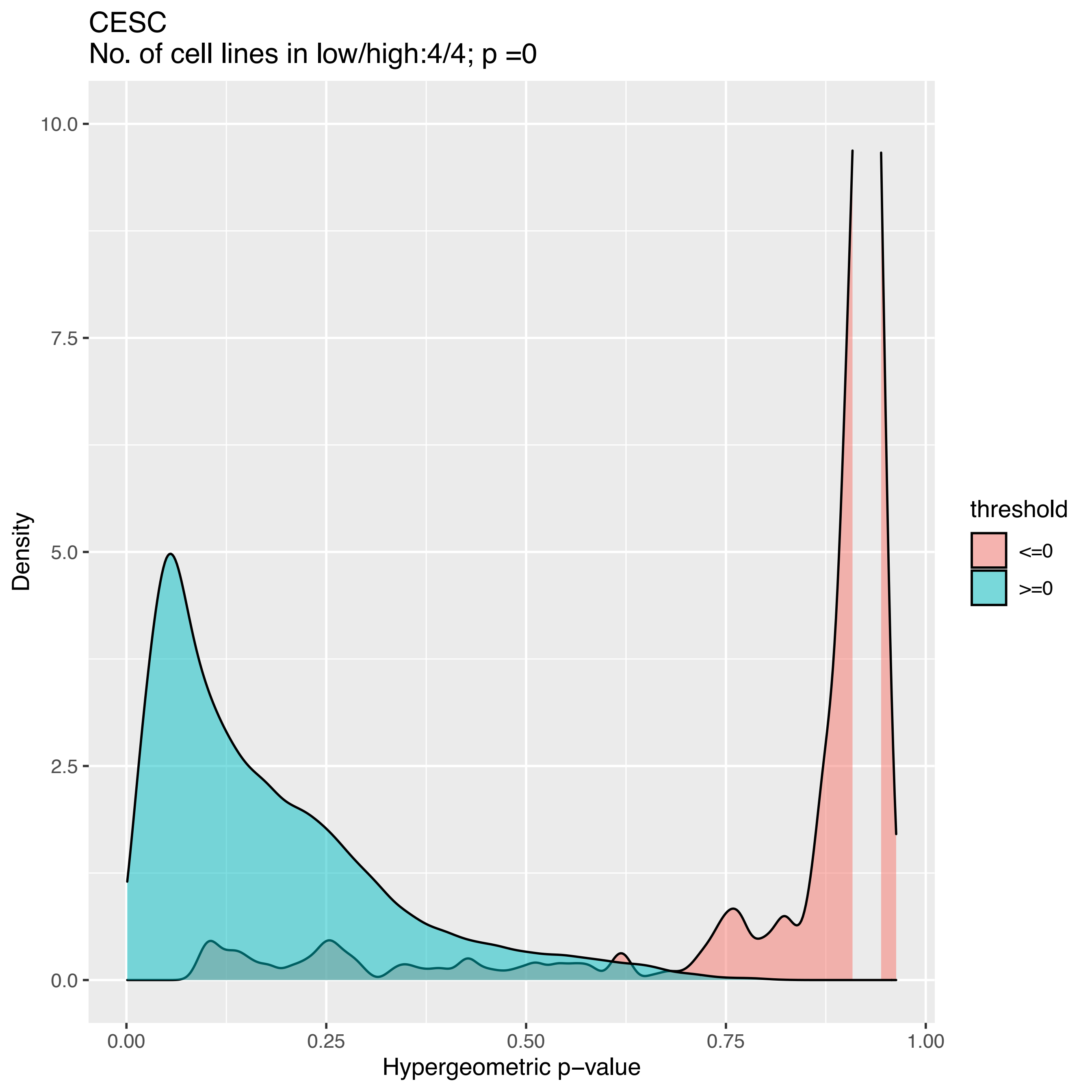
CESC
No. of cell lines in low/high:4/4; p =0
Density
Hypergeometric p−value
threshold
<=0
>=0
0.0
2.5
5.0
7.5
10.0
0.00
0.25
0.50
0.75
1.00

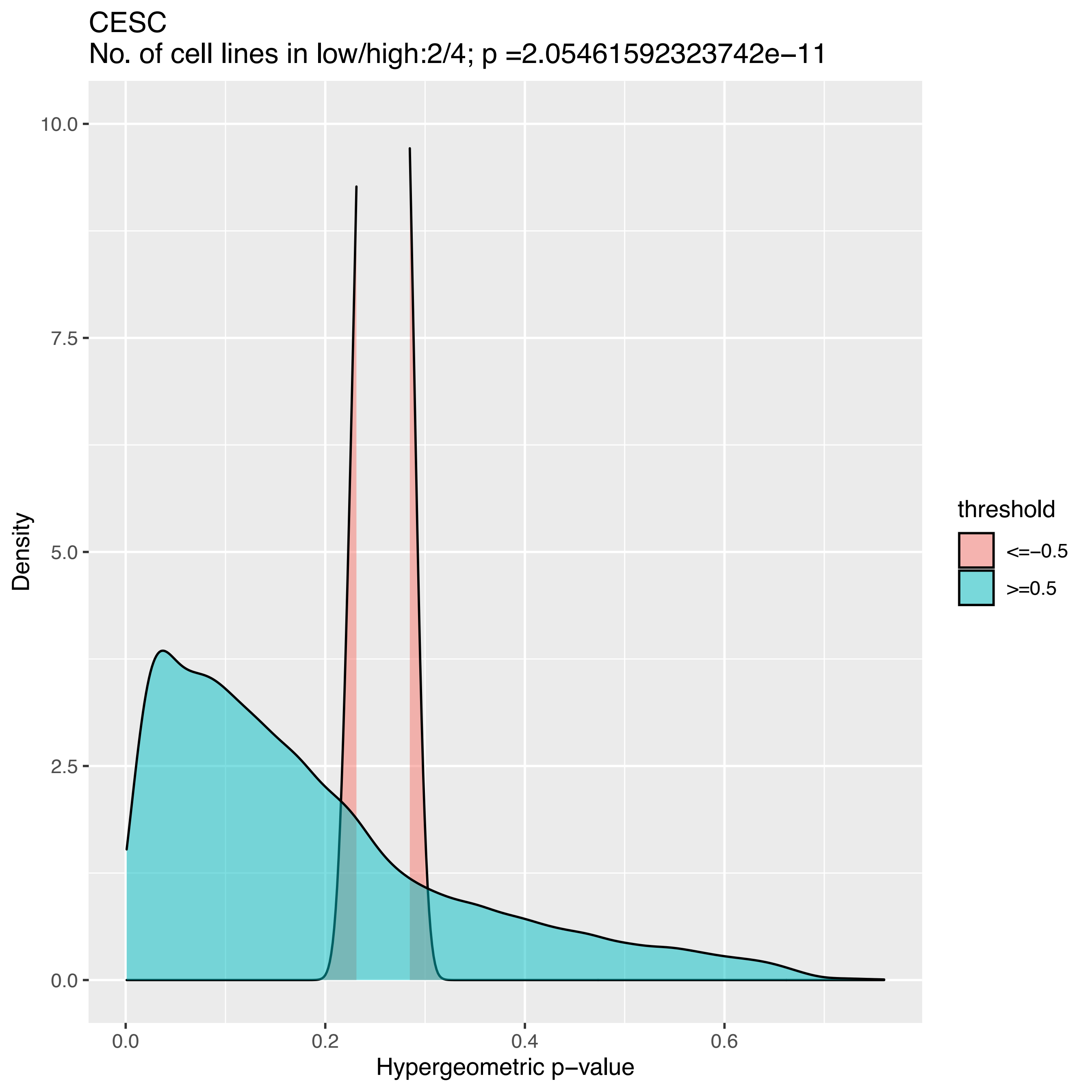
CESC
No. of cell lines in low/high:2/4; p =2.05461592323742e−11
Density
Hypergeometric p−value
10.0
7.5
5.0
2.5
0.0
0.0
0.2
0.4
0.6
threshold
<=−0.5
>=0.5

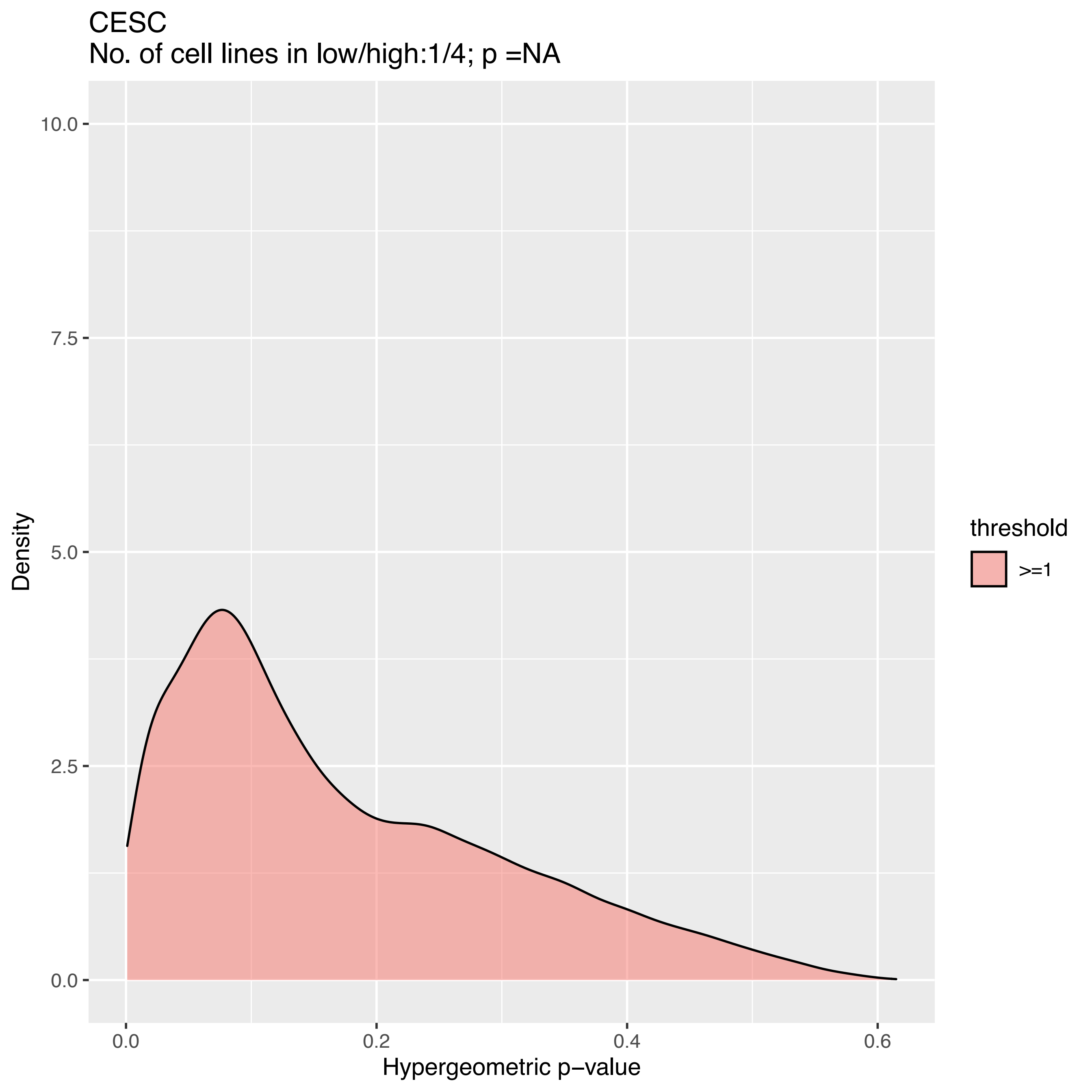
CESC
No. of cell lines in low/high:1/4; p =NA
10.0
7.5
5.0
2.5
0.0
Density
0.0
0.2
0.4
0.6
Hypergeometric p−value
threshold
>=1

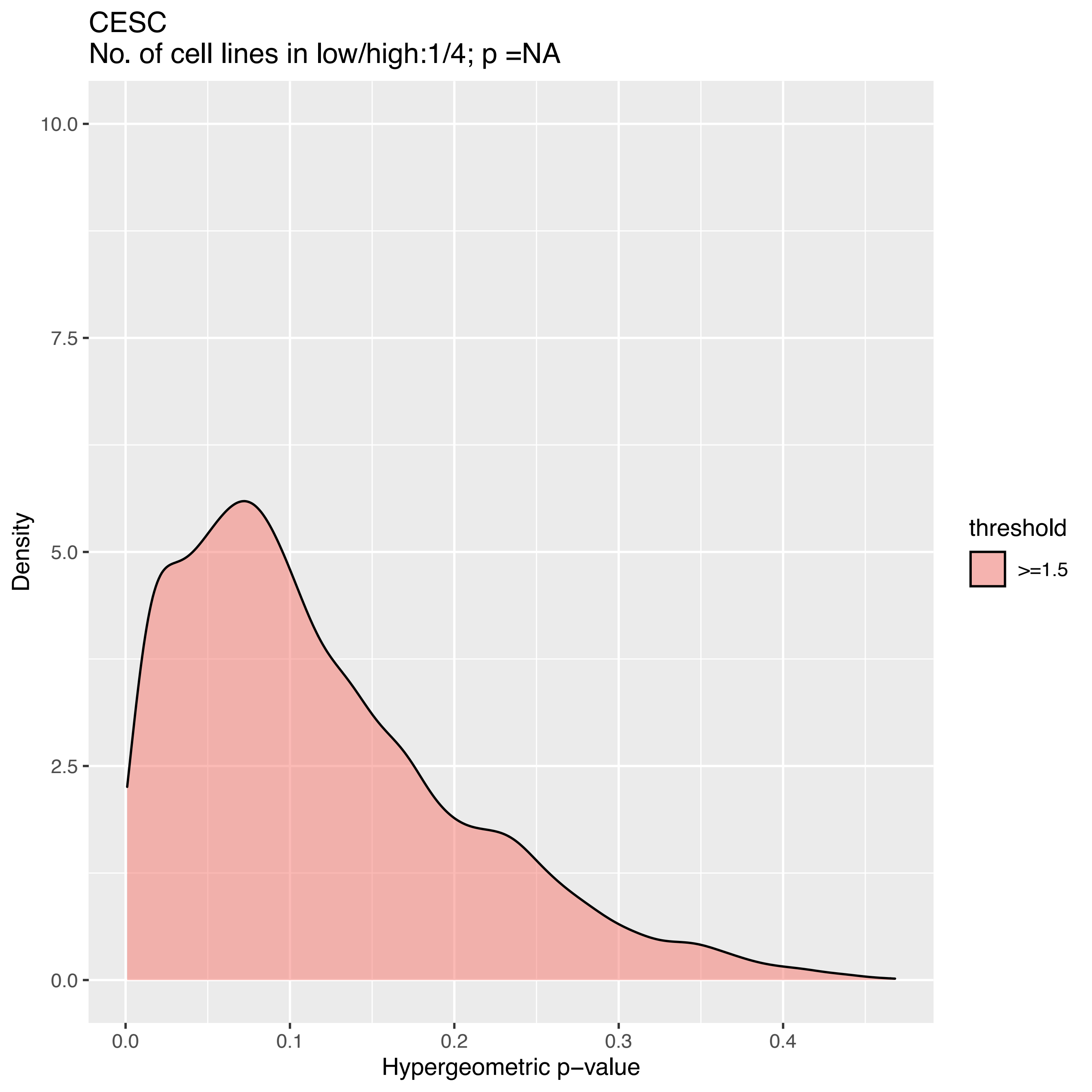

CESC
No. of cell lines in low/high:1/4; p =NA
Density
10.0
7.5
5.0
2.5
0.0
0.0
0.1
0.2
0.3
0.4
Hypergeometric p−value
threshold
>=1.5

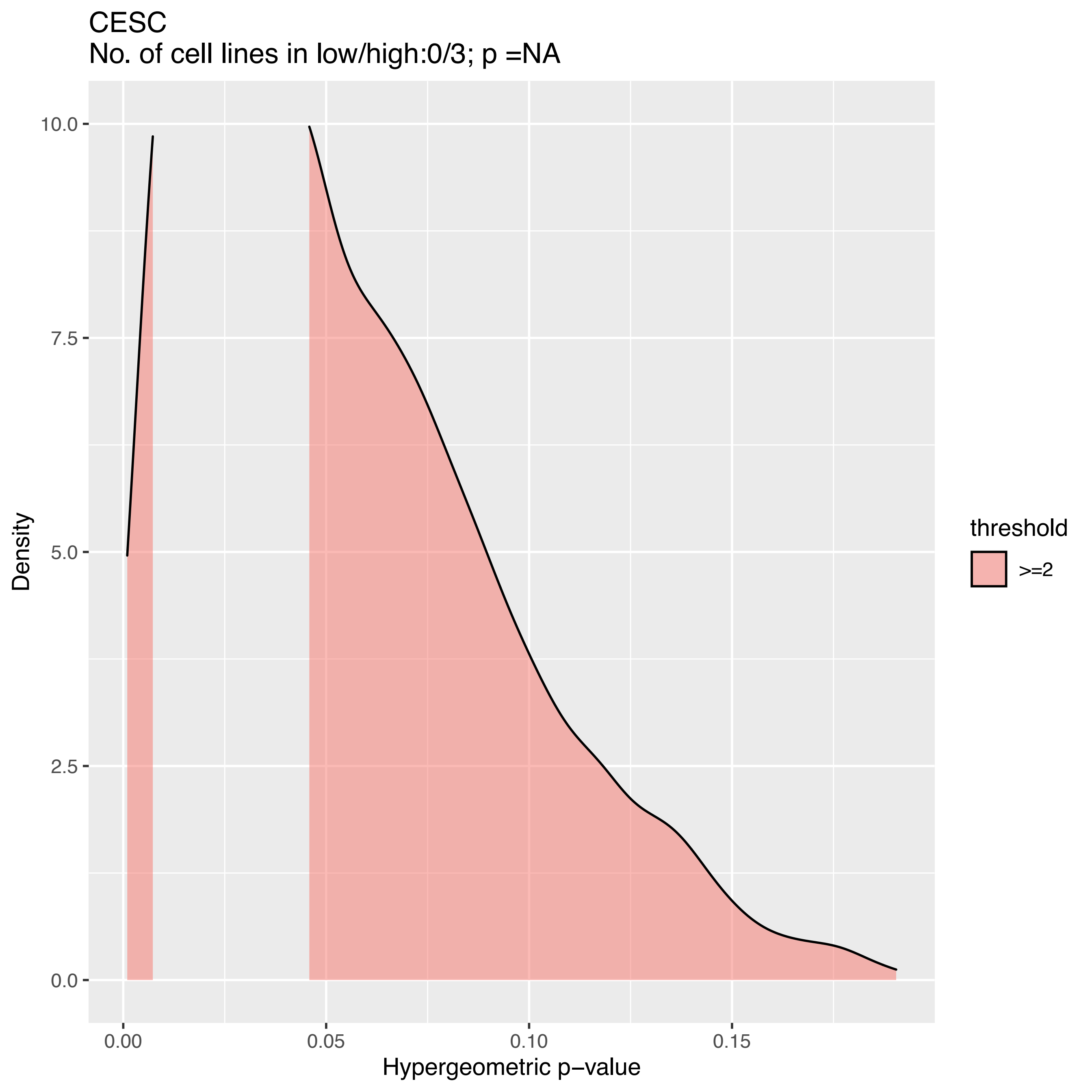

CESC
No. of cell lines in low/high:0/3; p =NA
Density
10.0
7.5
5.0
2.5
0.0
0.00
0.05
0.10
0.15
Hypergeometric p−value
threshold
>=2

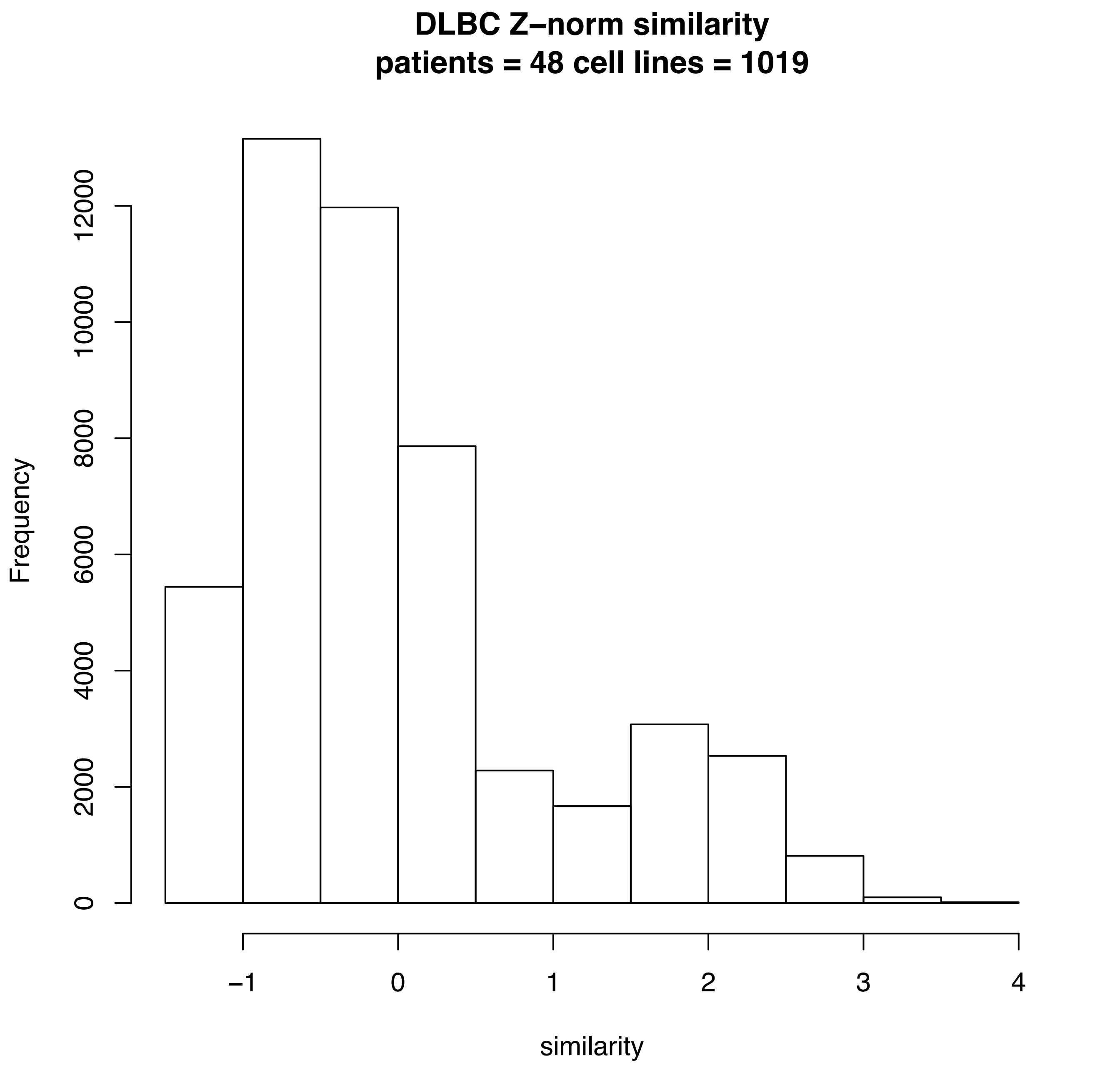
DLBC Z–norm similarity
patients = 48 cell lines = 1019
Frequency
0
2000
4000
6000
8000
10000
12000
–1
0
1
2
3
4
similarity

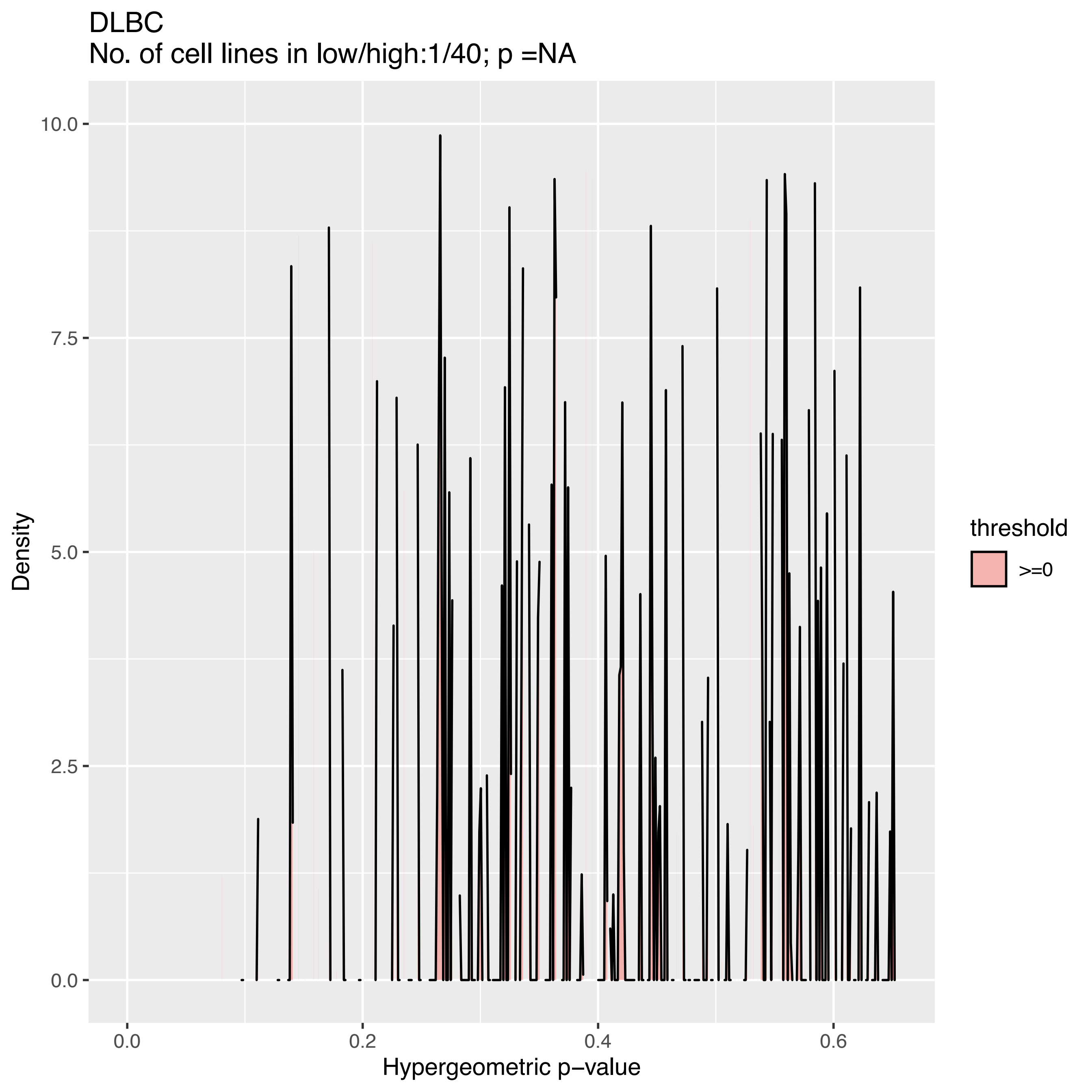

DLBC
No. of cell lines in low/high:1/40; p =NA
Density
10.0
7.5
5.0
2.5
0.0
0.0
0.2
0.4
0.6
Hypergeometric p−value
threshold
>=0

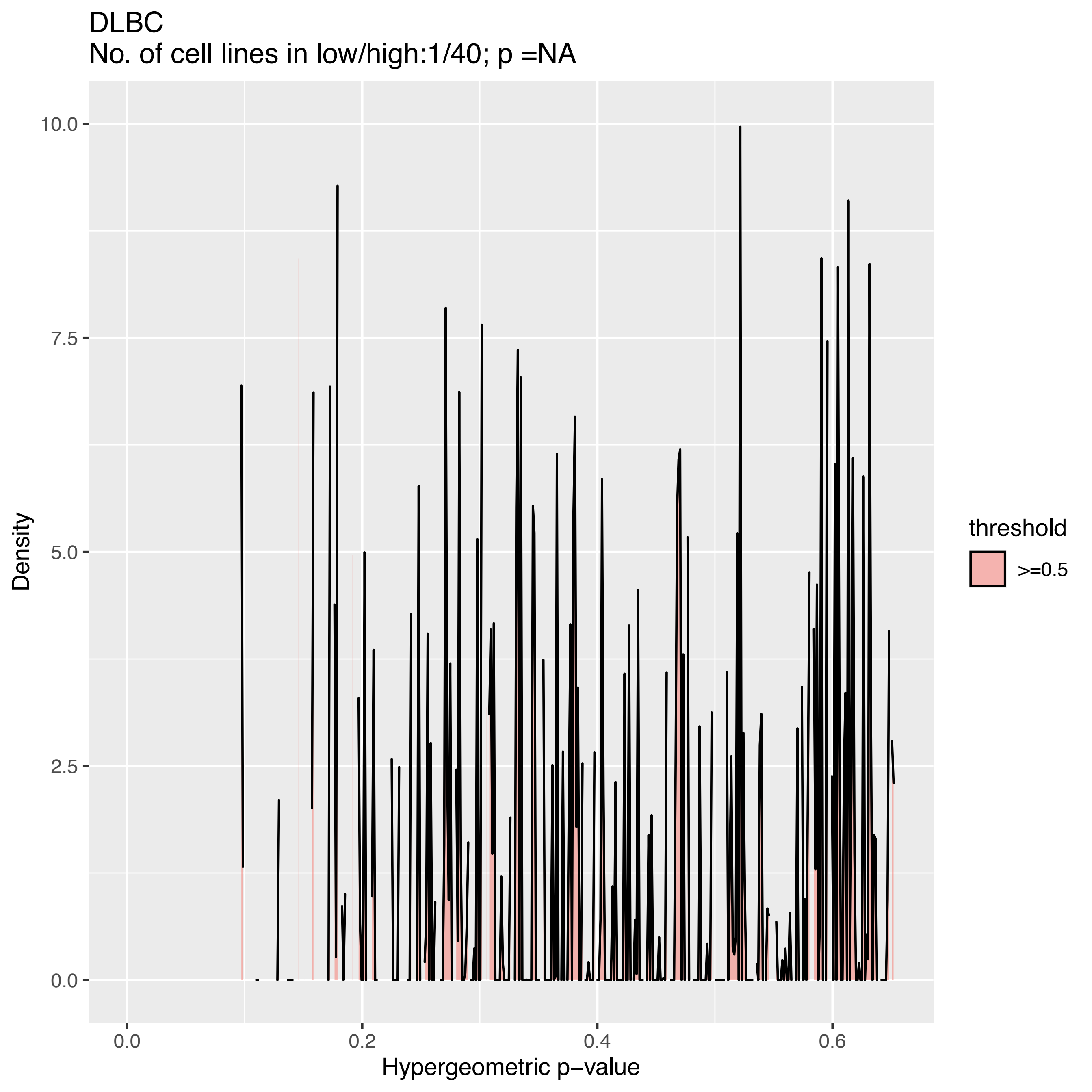

DLBC
No. of cell lines in low/high:1/40; p =NA
Density
Hypergeometric p−value
0.0
2.5
5.0
7.5
10.0
0.0
0.2
0.4
0.6
threshold
>=0.5

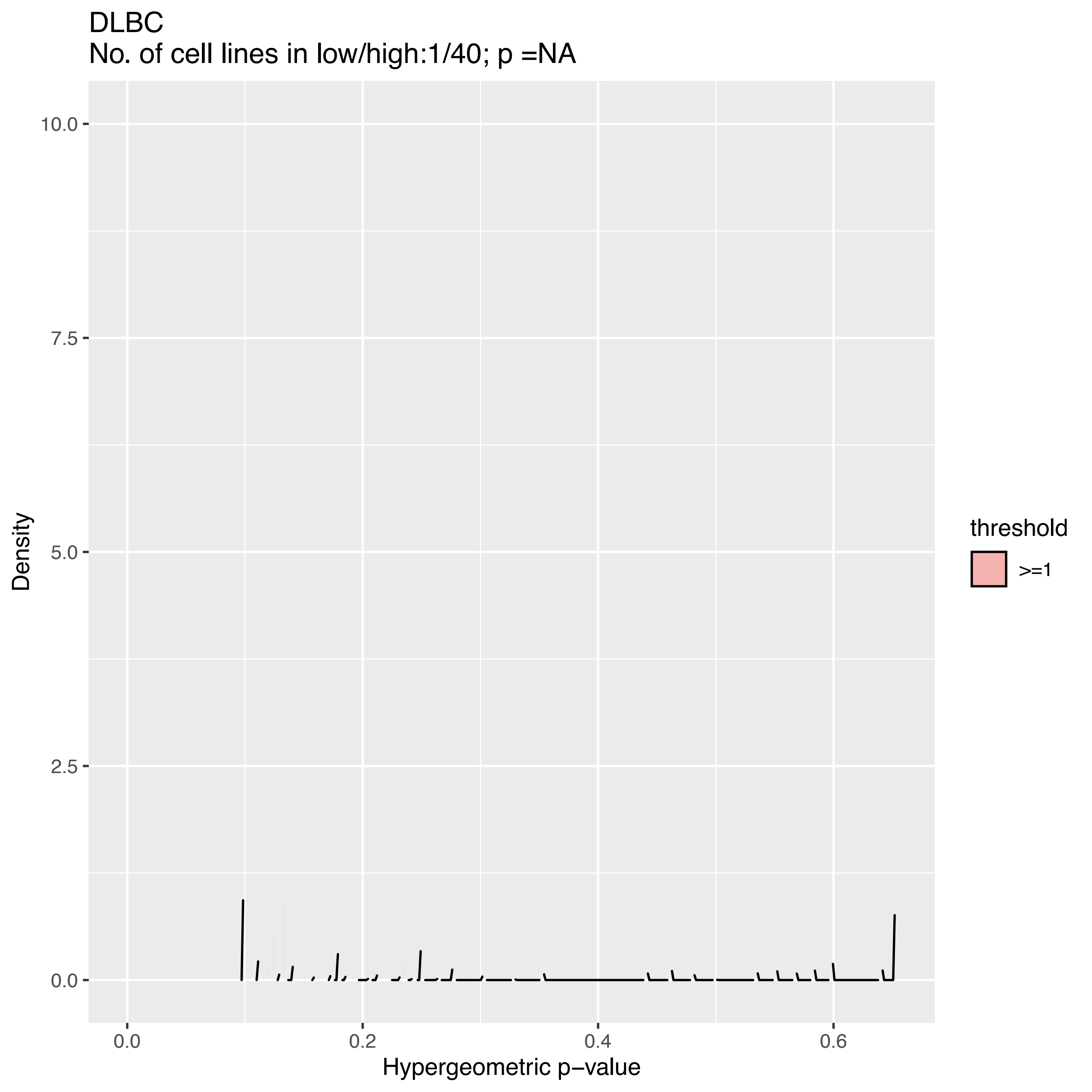
DLBC
No. of cell lines in low/high:1/40; p =NA
Density
10.0
7.5
5.0
2.5
0.0
0.0
0.2
0.4
0.6
Hypergeometric p−value
threshold
>=1

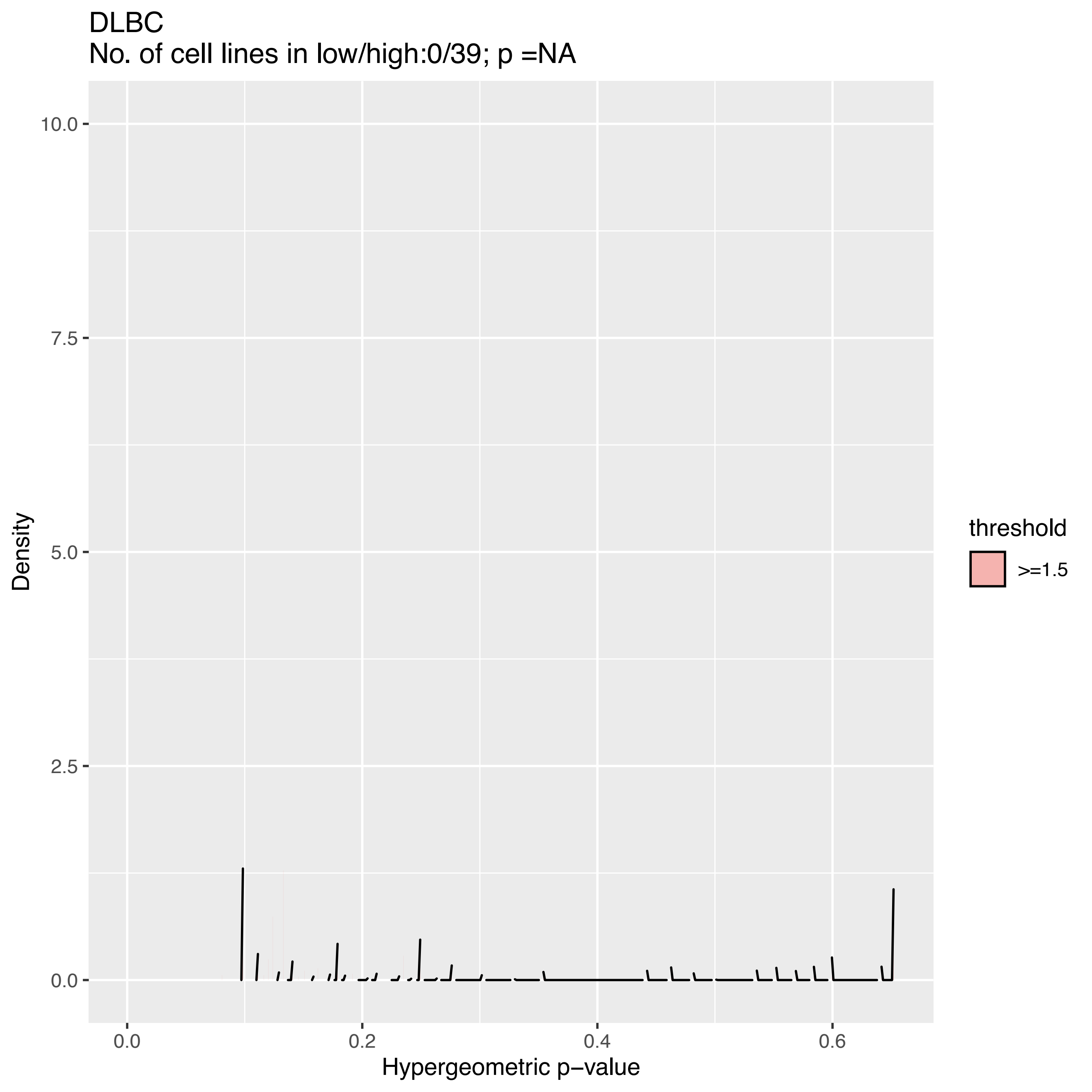

DLBC
No. of cell lines in low/high:0/39; p =NA
Density
10.0
7.5
5.0
2.5
0.0
0.0
0.2
0.4
0.6
Hypergeometric p−value
threshold
>=1.5

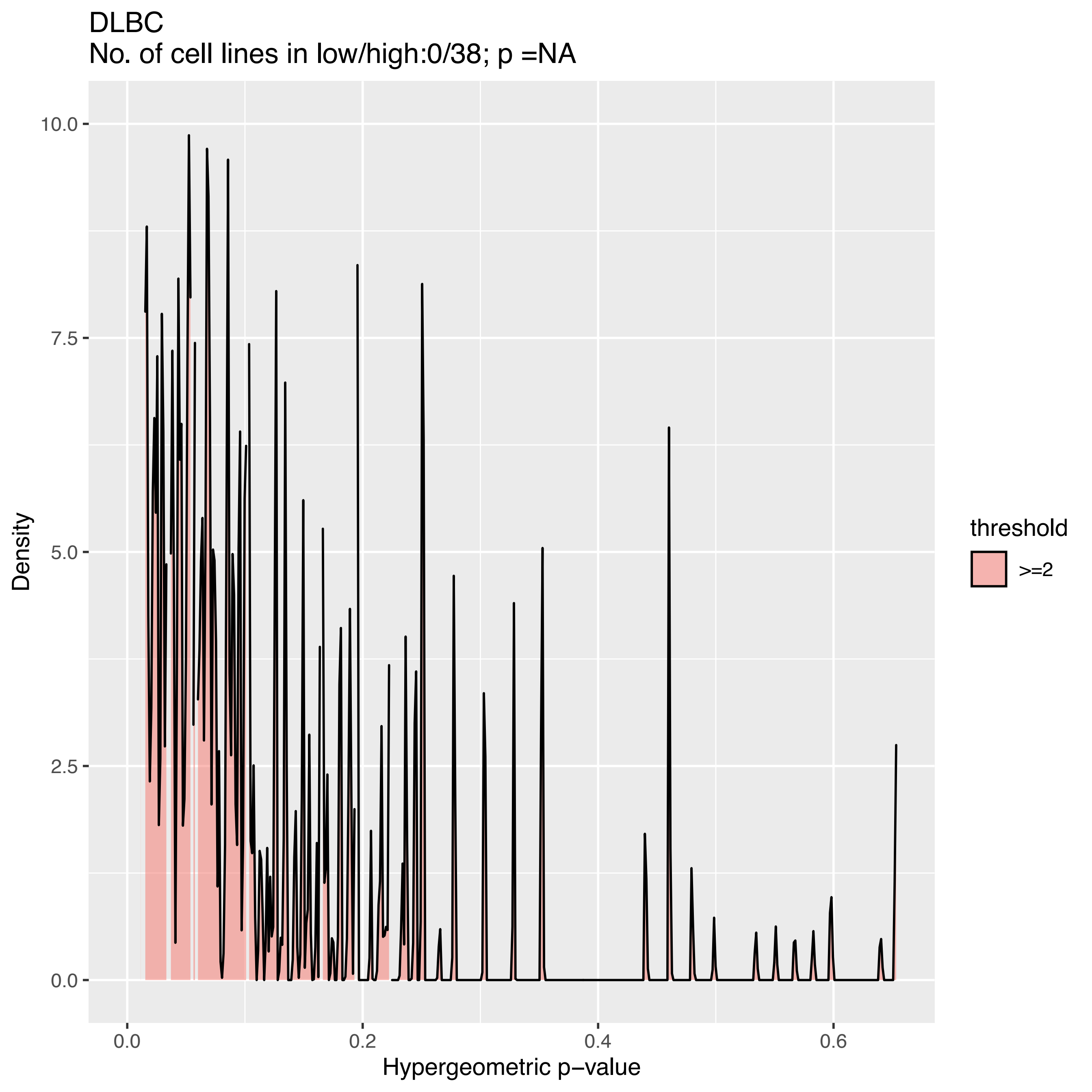

DLBC
No. of cell lines in low/high:0/38; p =NA
Density
Hypergeometric p−value
threshold
>=2
10.0
7.5
5.0
2.5
0.0
0.0
0.2
0.4
0.6

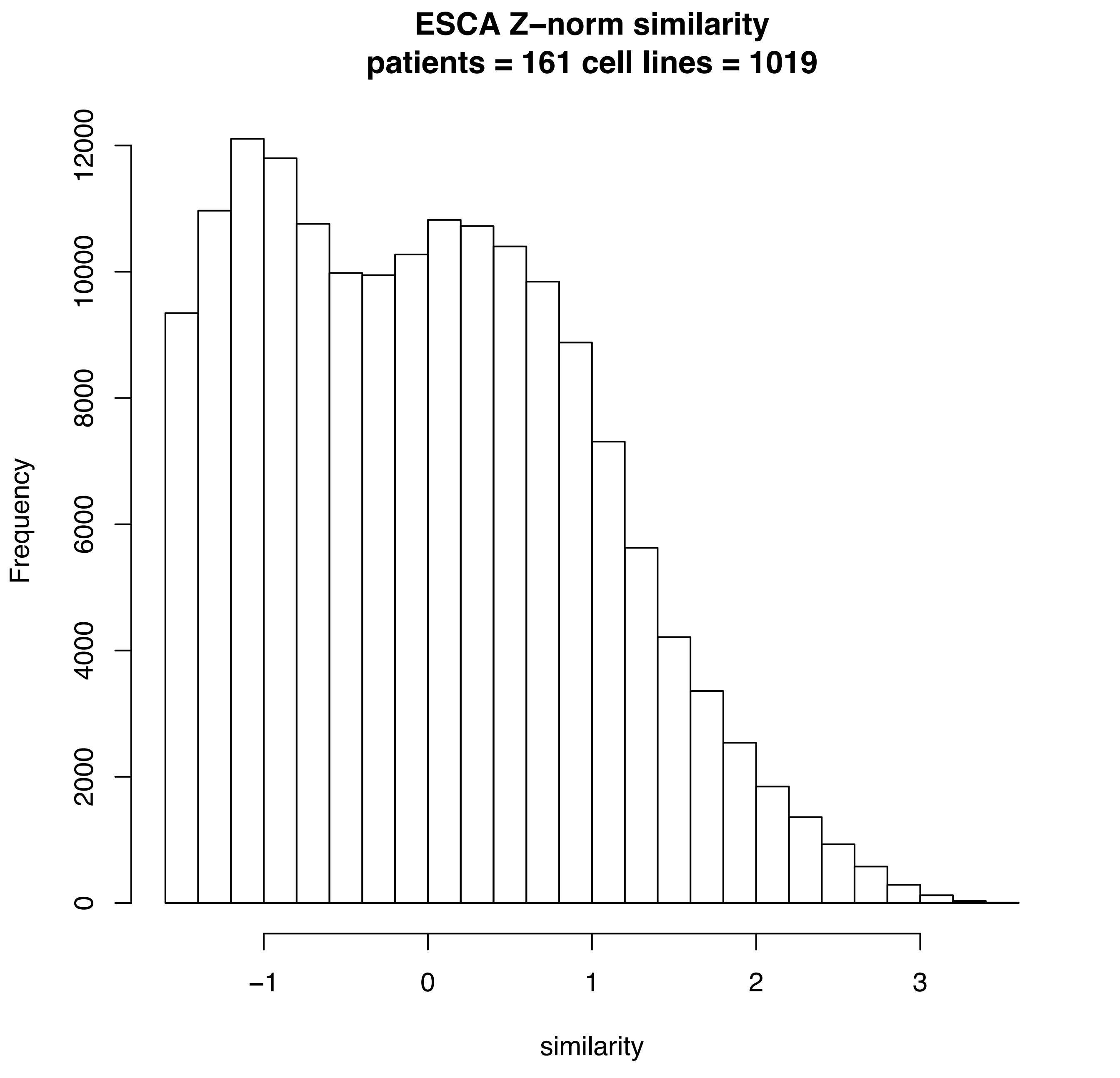
ESCA Z–norm similarity
patients = 161 cell lines = 1019
Frequency
0
2000
4000
6000
8000
10000
12000
–1
0
1
2
3
similarity

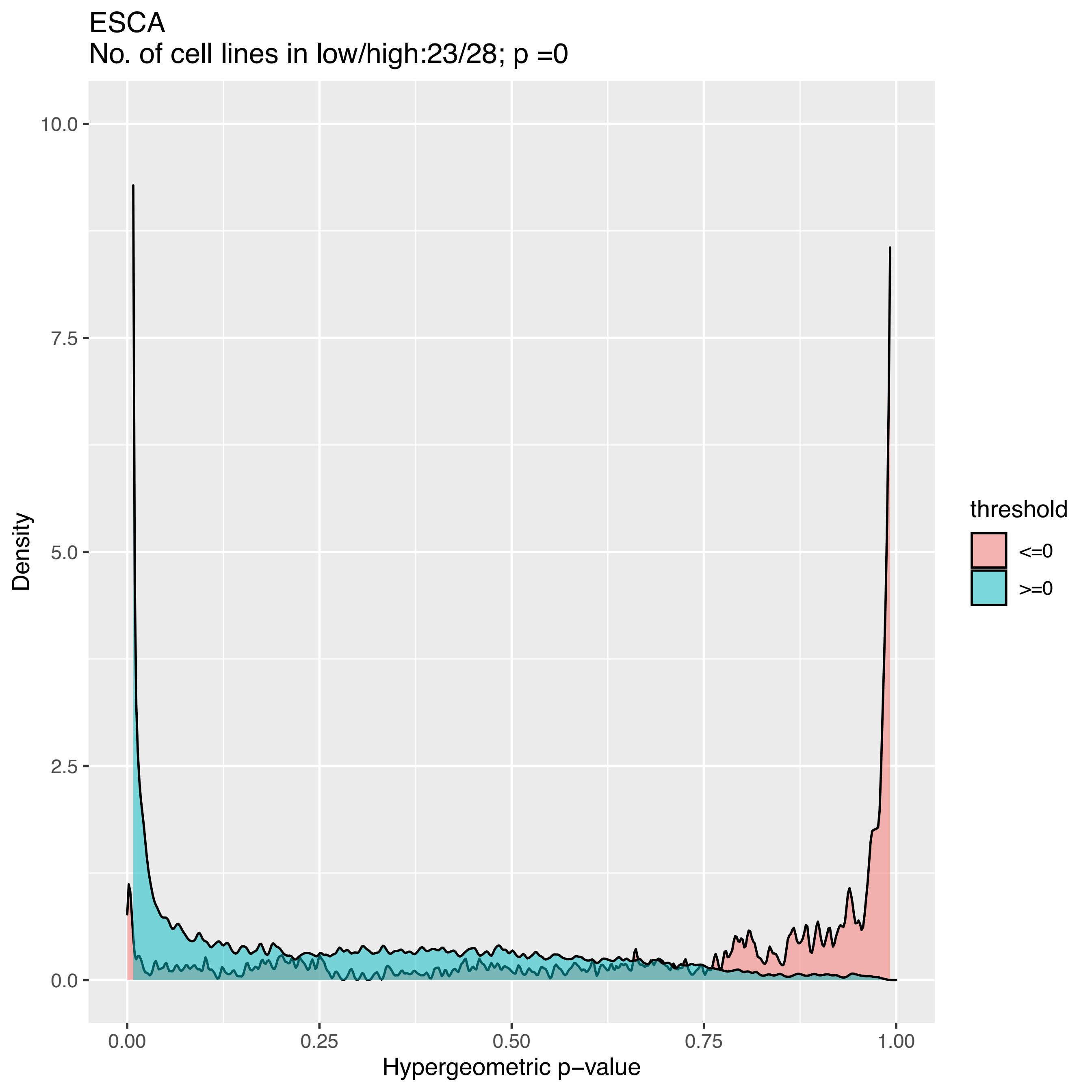

ESCA
No. of cell lines in low/high:23/28; p =0
Density
10.0
7.5
5.0
2.5
0.0
0.00
0.25
0.50
0.75
1.00
Hypergeometric p−value
threshold
<=0
>=0

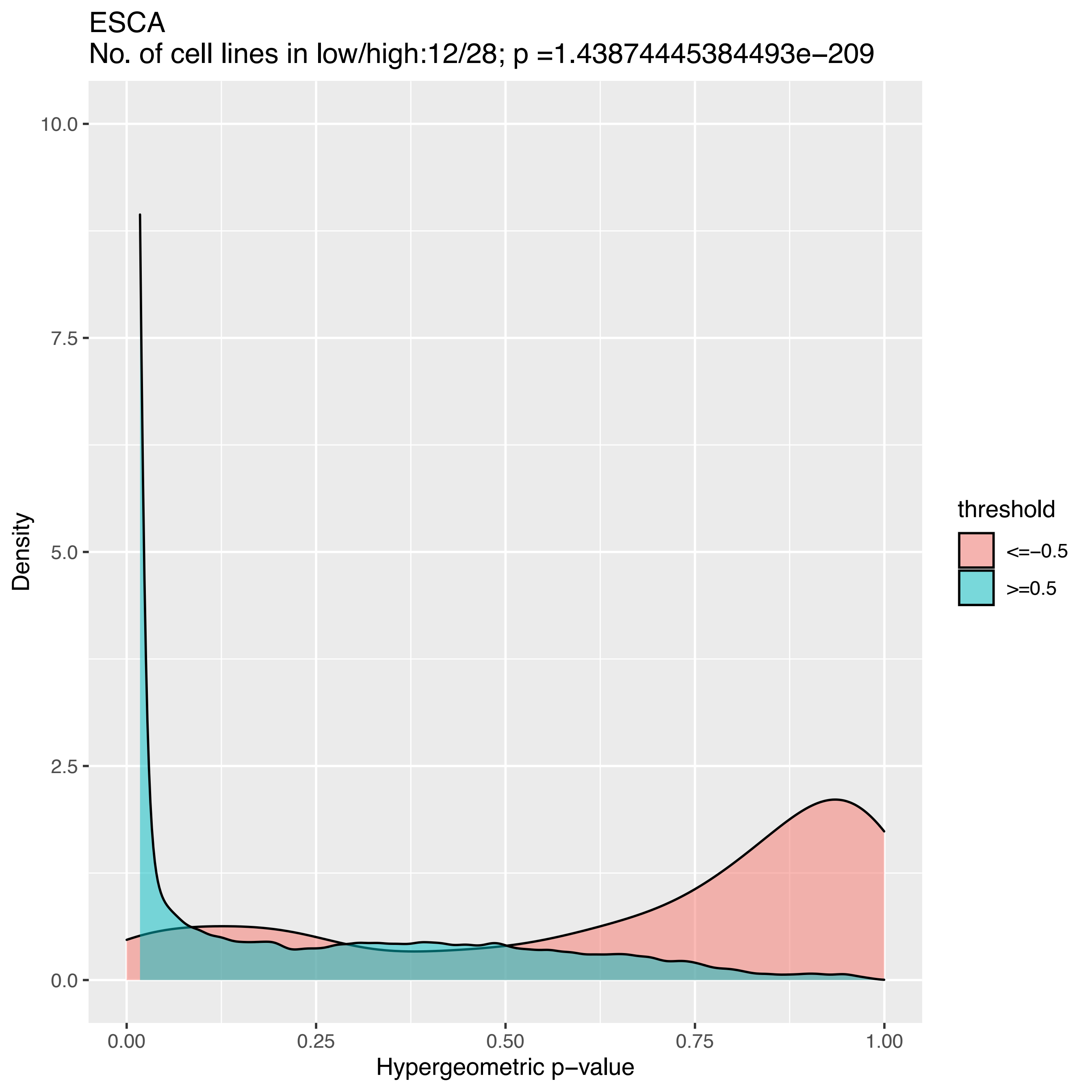
ESCA
No. of cell lines in low/high:12/28; p =1.43874445384493e−209
10.0
7.5
5.0
2.5
0.0
Density
0.00
0.25
0.50
0.75
1.00
Hypergeometric p−value
threshold
<=−0.5
>=0.5

ESCA
No. of cell lines in low/high:3/28; p =0.0227496267724688

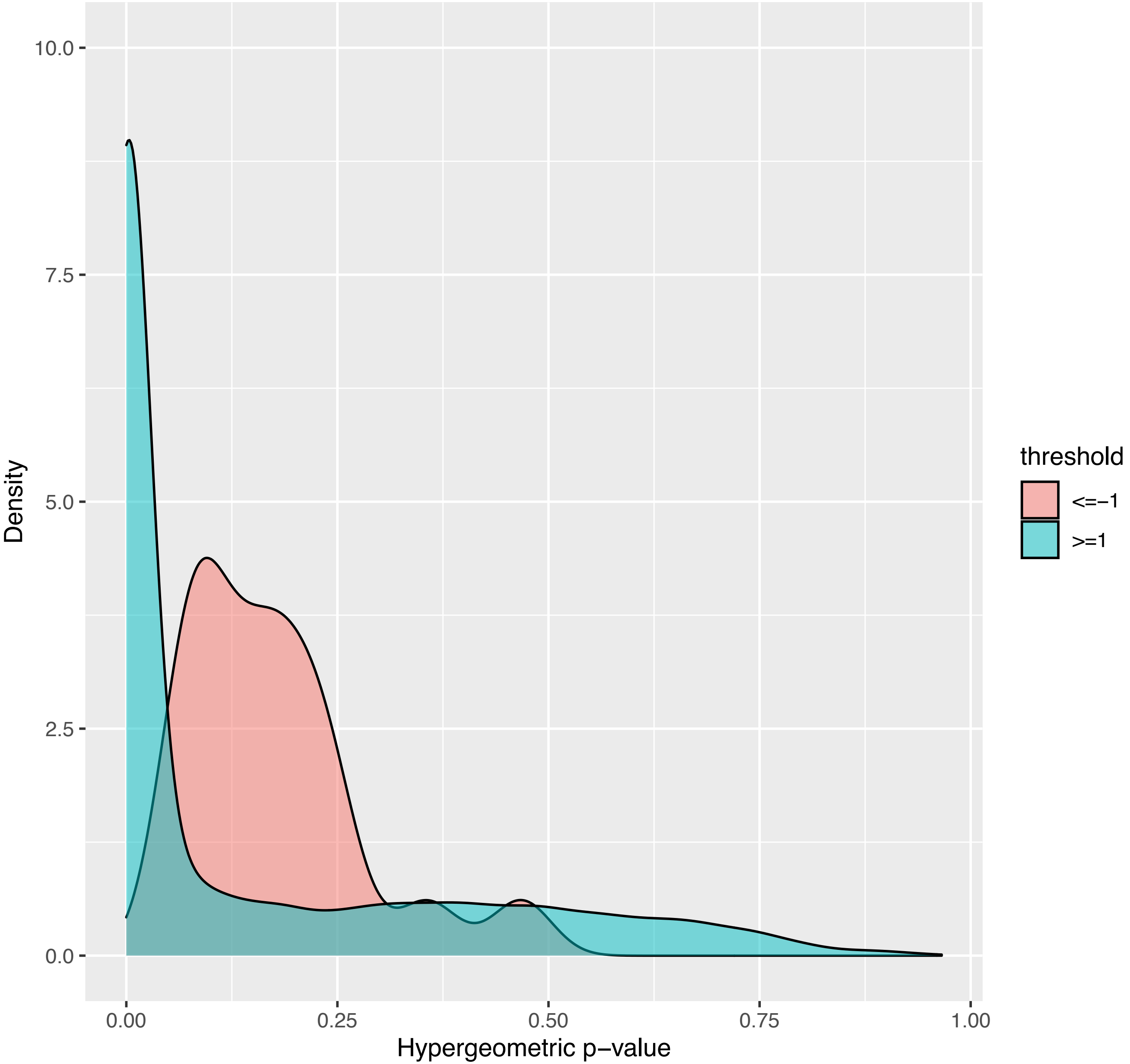

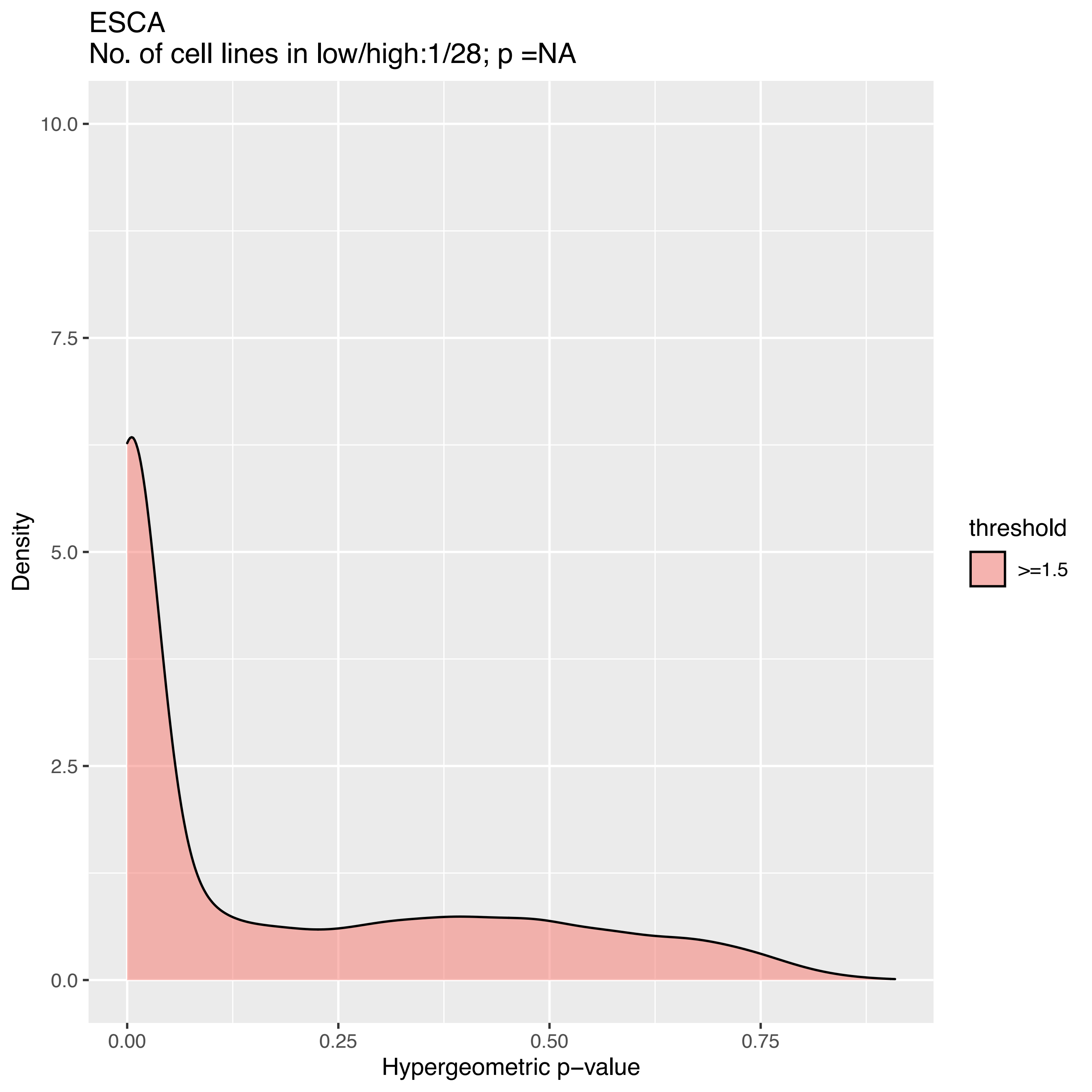

ESCA
No. of cell lines in low/high:1/28; p =NA
Density
10.0
7.5
5.0
2.5
0.0
0.00
0.25
0.50
0.75
Hypergeometric p−value
threshold
>=1.5

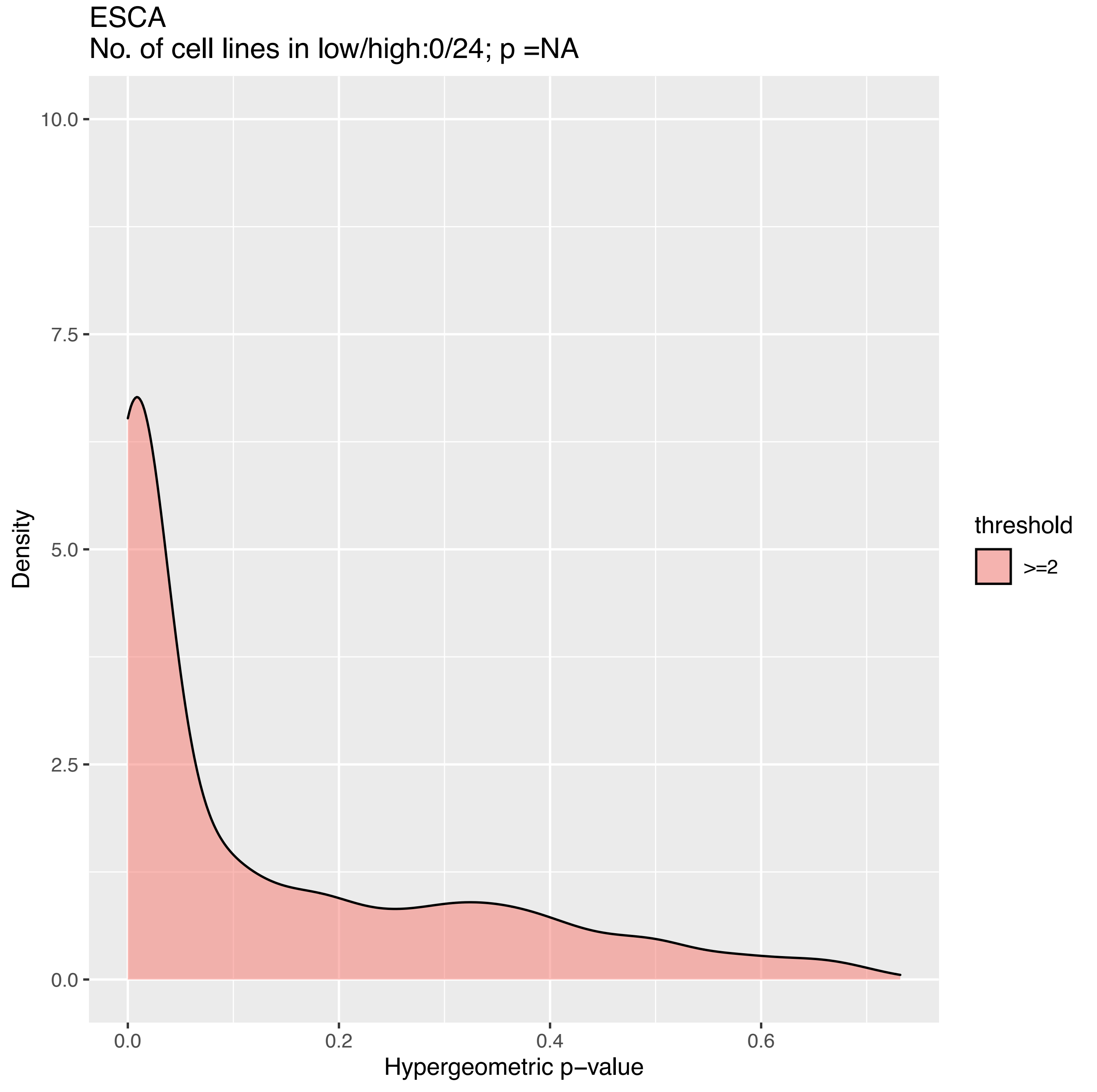

ESCA
No. of cell lines in low/high:0/24; p =NA
Density
10.0
7.5
5.0
2.5
0.0
0.0
0.2
0.4
0.6
Hypergeometric p−value
threshold
>=2

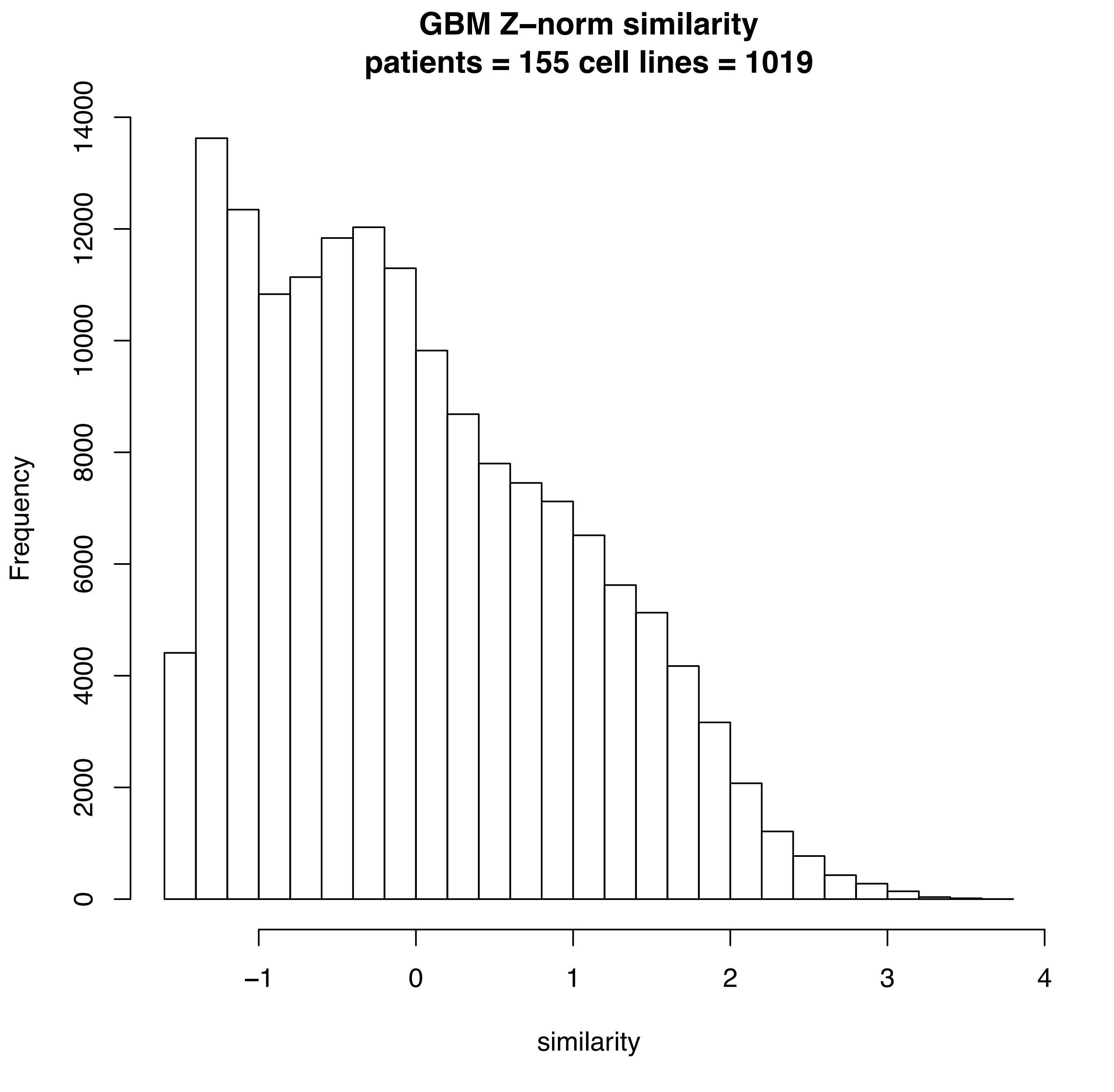

GBM Z–norm similarity
patients = 155 cell lines = 1019
Frequency
0
2000
4000
6000
8000
10000
12000
14000
−1
0
1
2
3
4
similarity

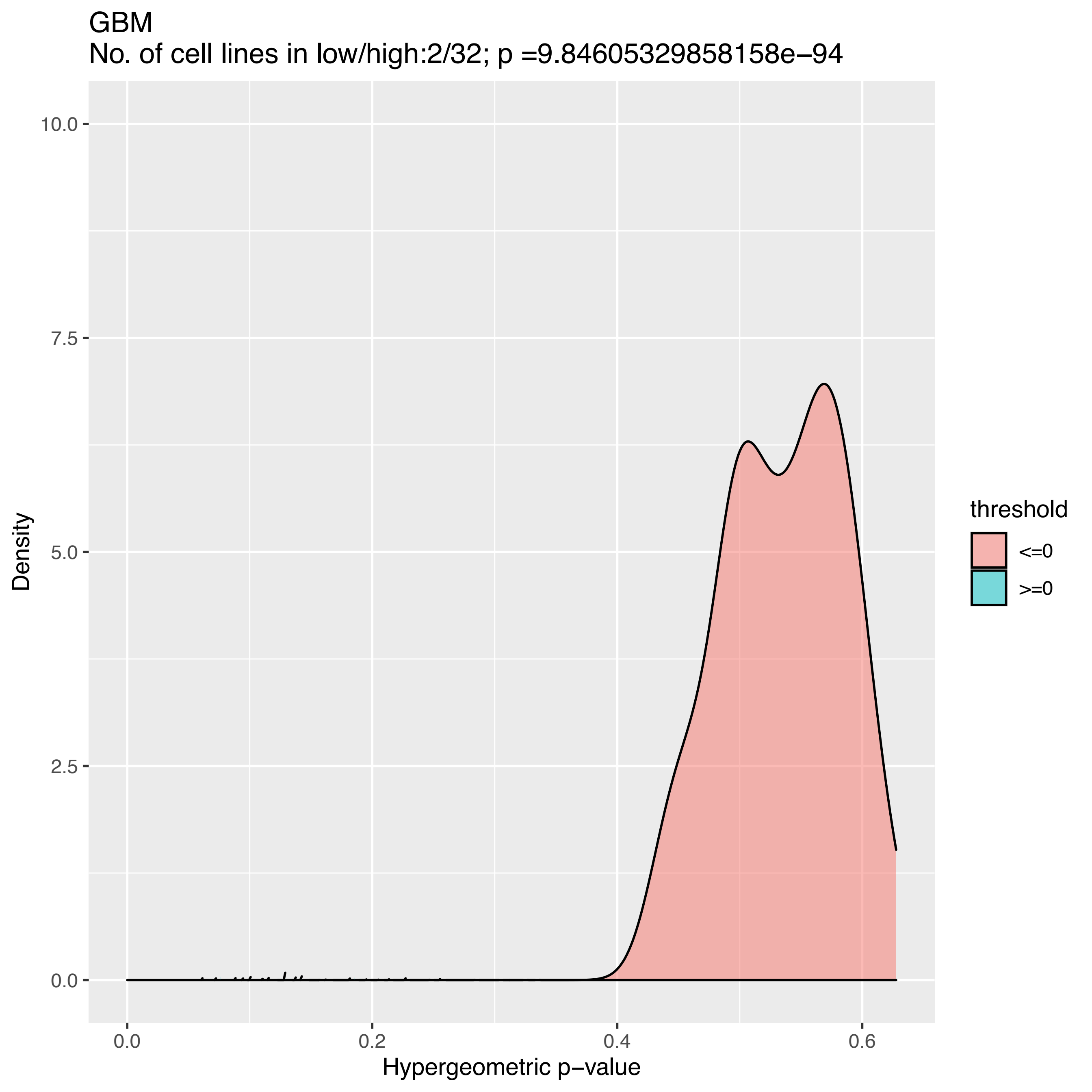
GBM
No. of cell lines in low/high:2/32; p =9.84605329858158e−94
Density
10.0
7.5
5.0
2.5
0.0
0.0
0.2
0.4
0.6
Hypergeometric p−value
threshold
<=0
>=0

GBM
No. of cell lines in low/high:1/32; p =NA

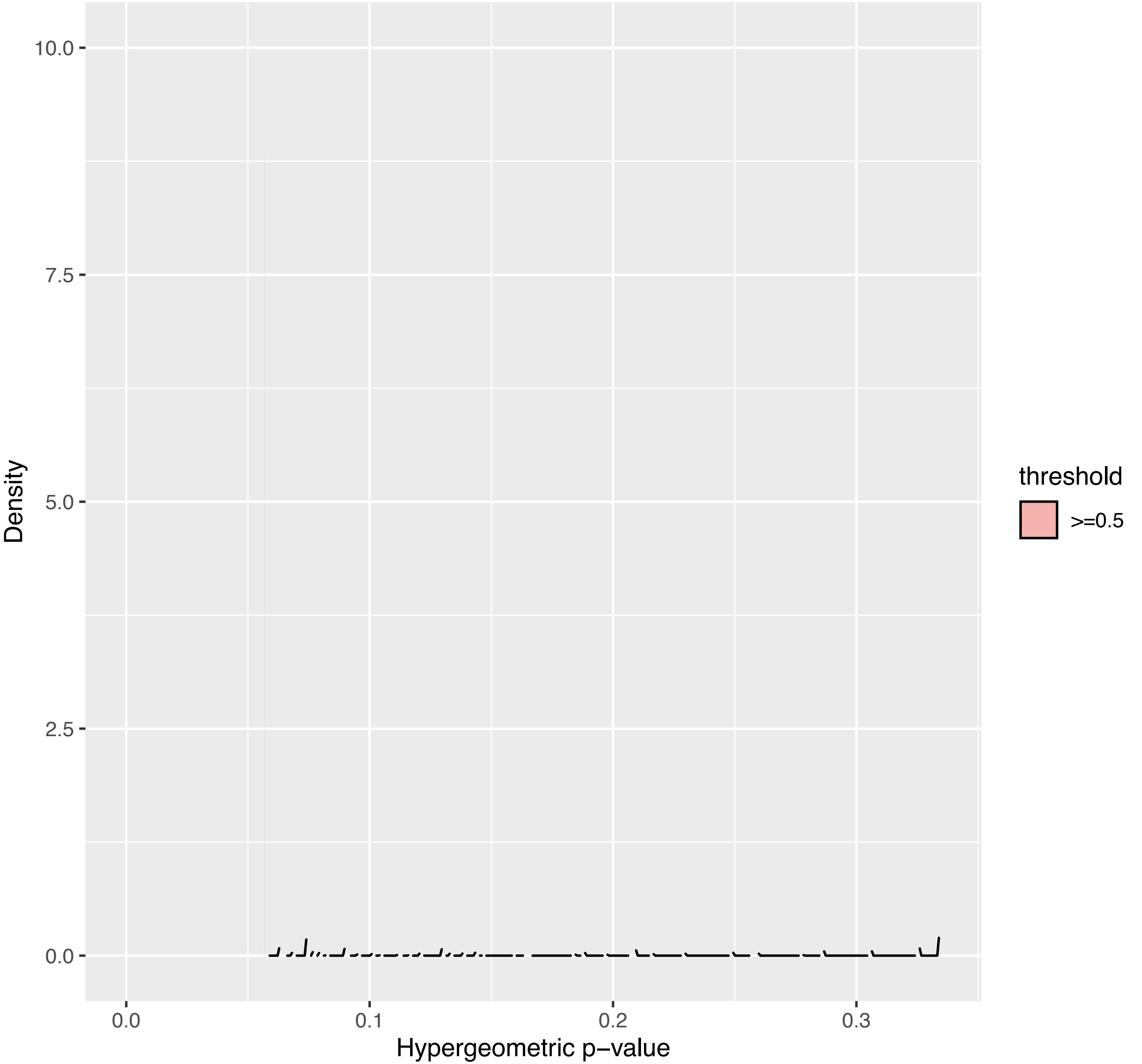

GBM
No. of cell lines in low/high:1/32; p =NA

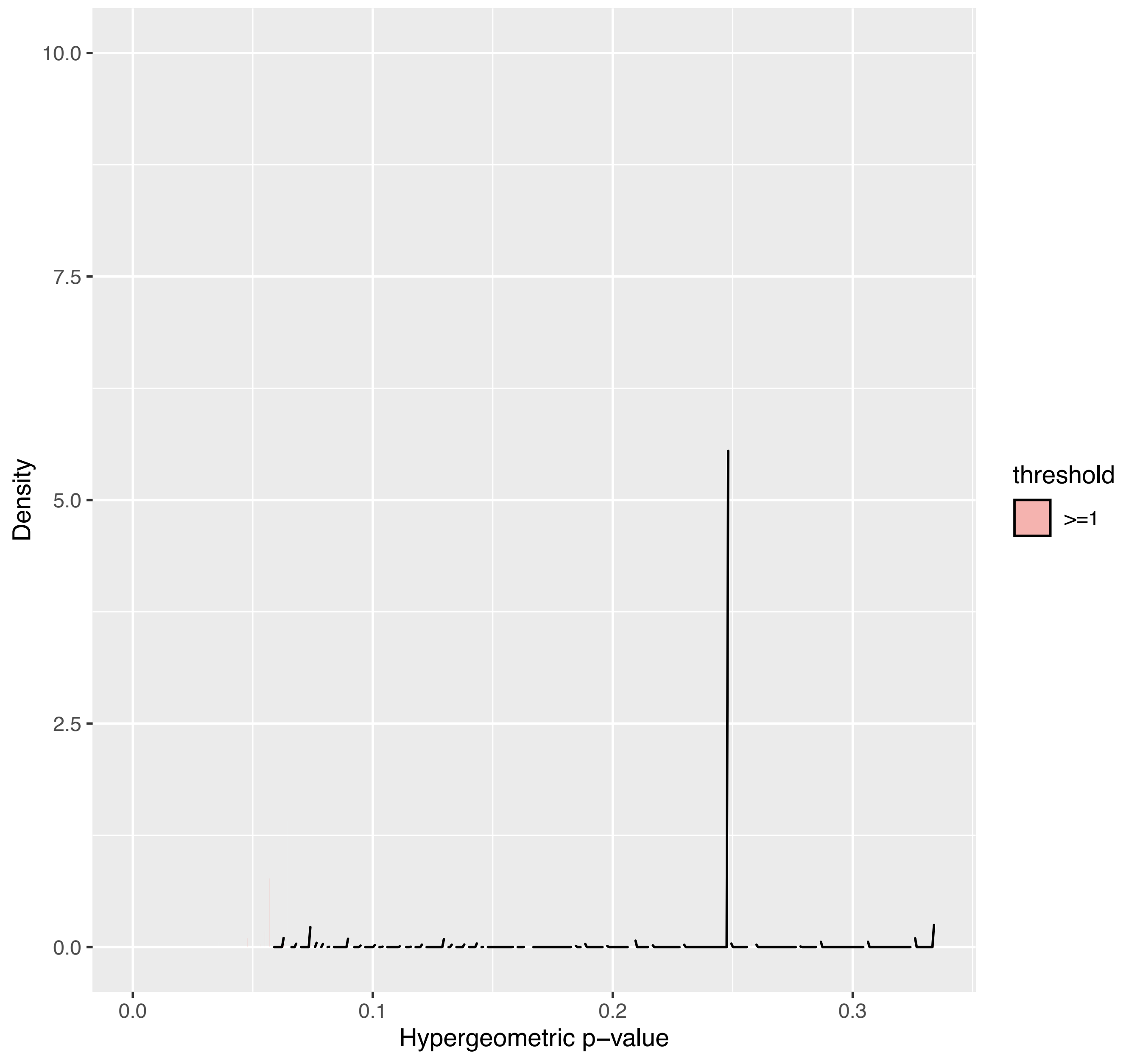

GBM
No. of cell lines in low/high:0/32; p =NA

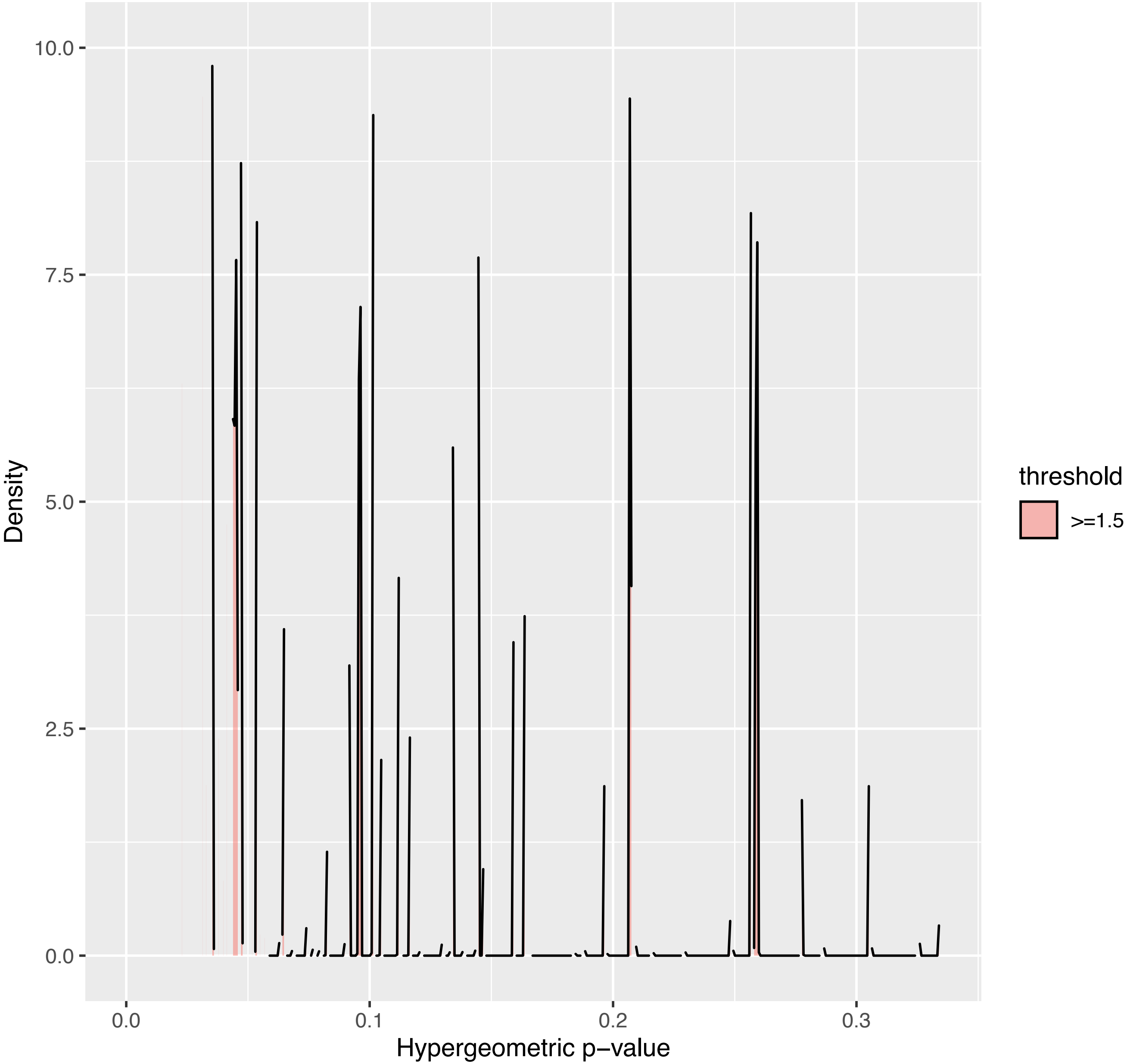

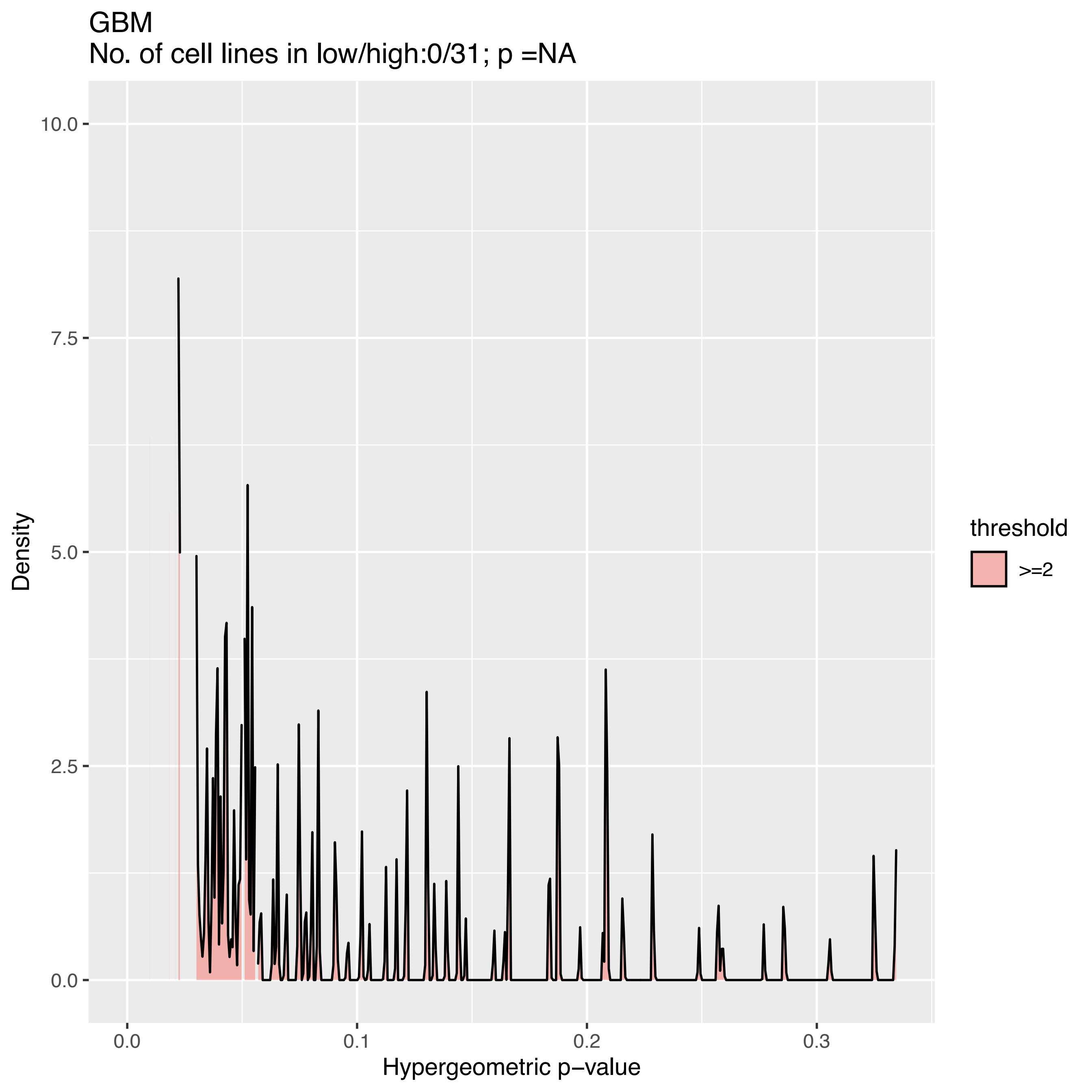
GBM
No. of cell lines in low/high:0/31; p =NA
Density
10.0
7.5
5.0
2.5
0.0
0.0
0.1
0.2
0.3
Hypergeometric p−value
threshold
>=2

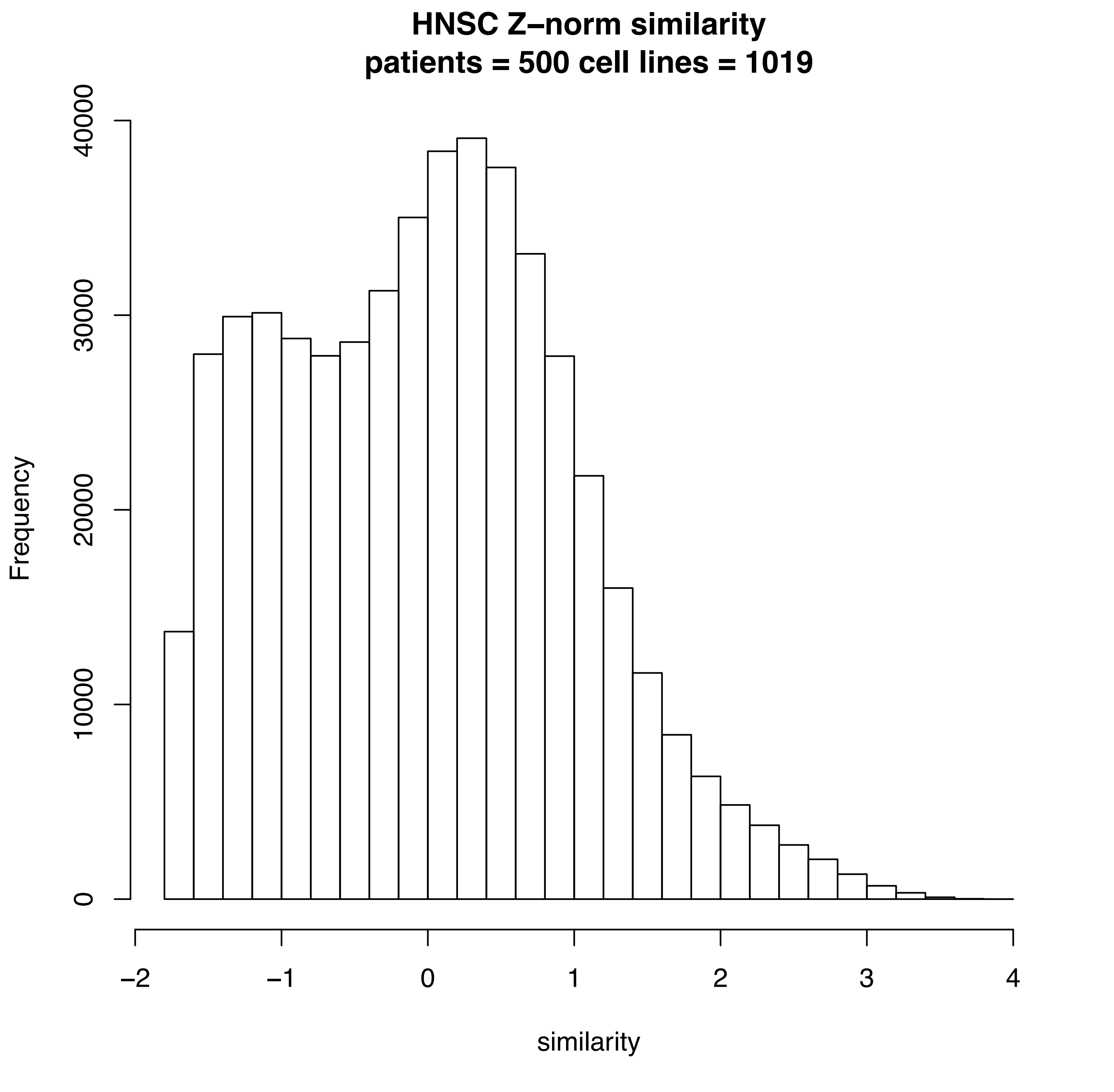

HNSC Z–norm similarity
patients = 500 cell lines = 1019
Frequency
0
10000
20000
30000
40000
−2
−1
0
1
2
3
4
similarity

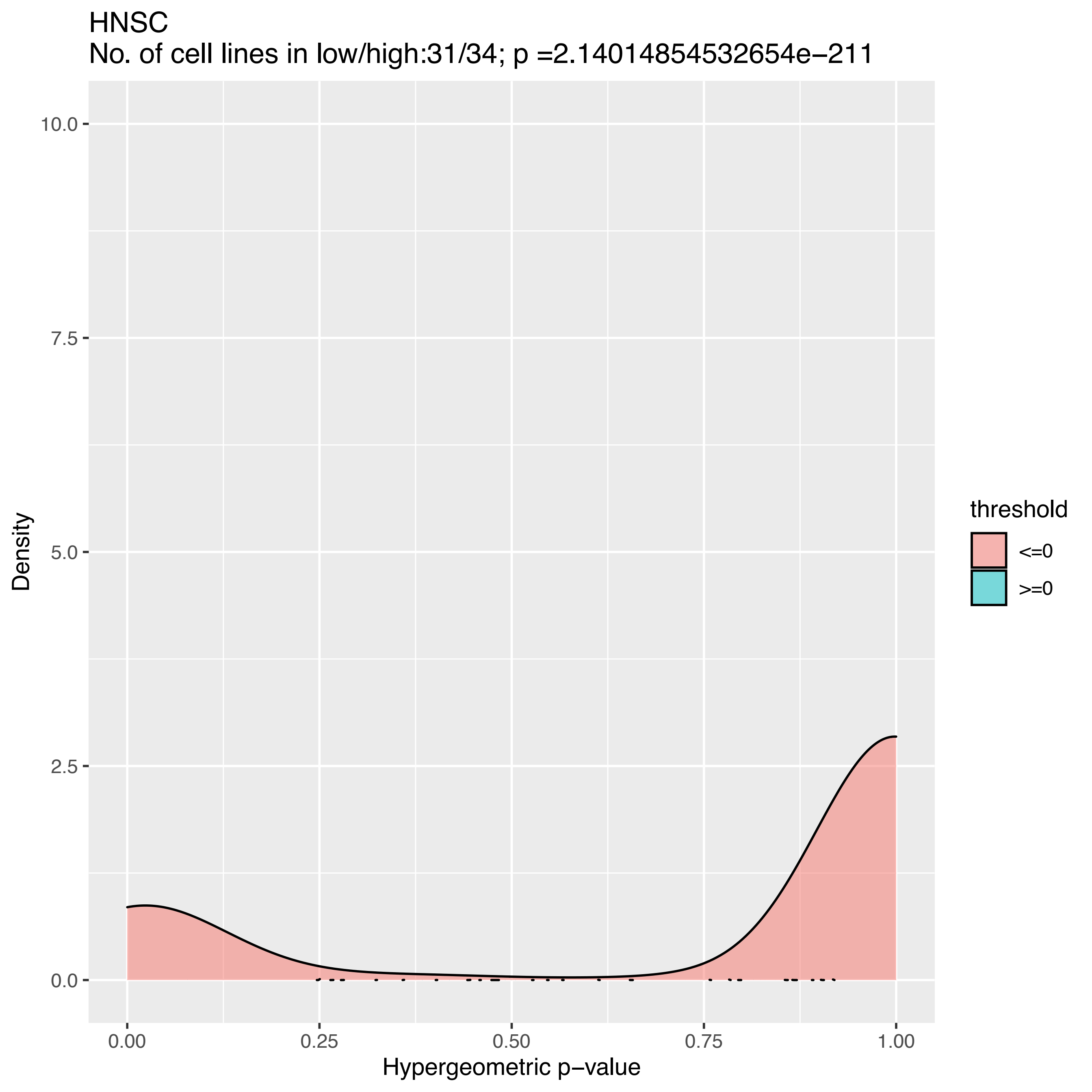
HNSC
No. of cell lines in low/high:31/34; p =2.14014854532654e−211
10.0
7.5
5.0
2.5
0.0
Density
0.00
0.25
0.50
0.75
1.00
Hypergeometric p−value
threshold
<=0
>=0

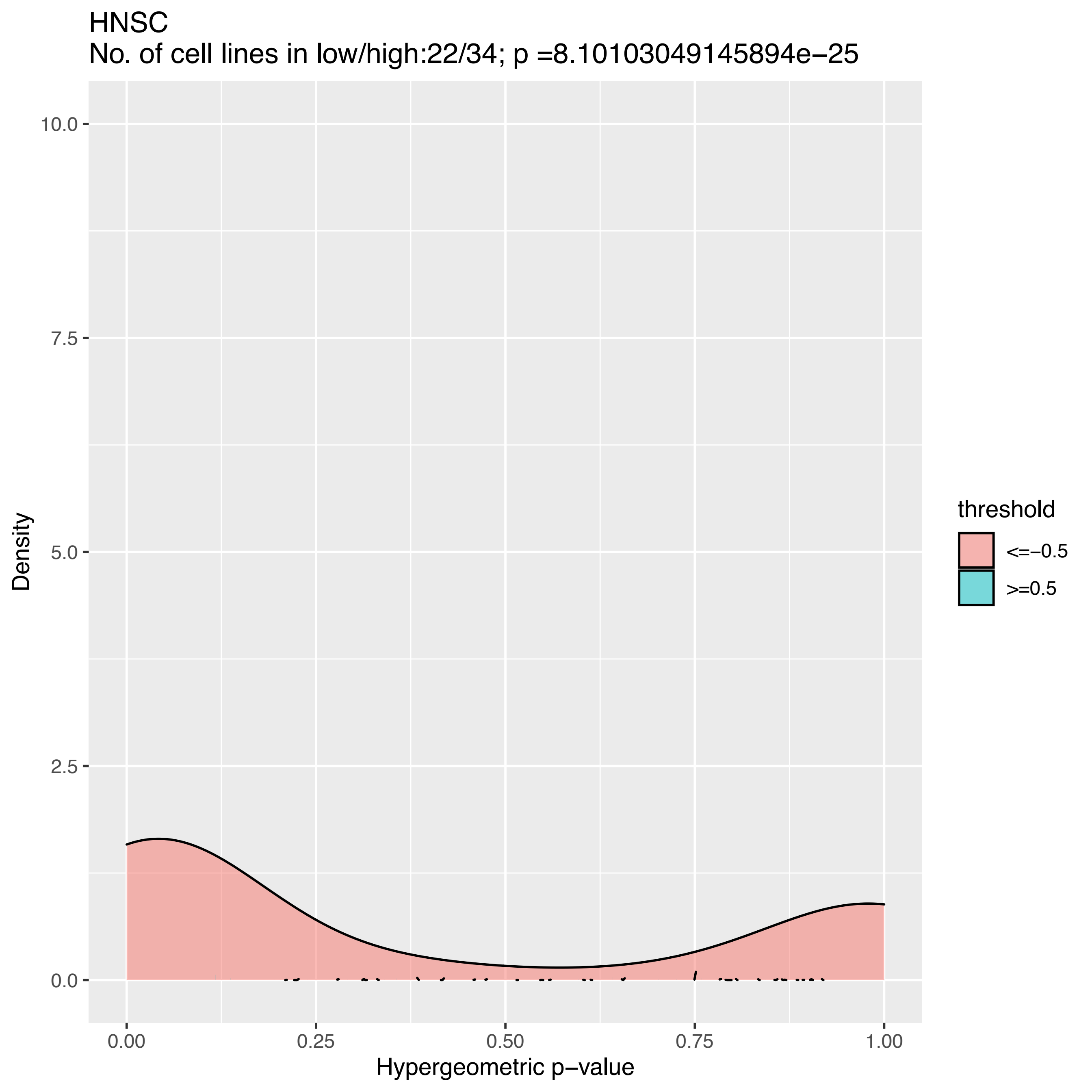
HNSC
No. of cell lines in low/high:22/34; p =8.10103049145894e−25
Density
10.0
7.5
5.0
2.5
0.0
0.00
0.25
0.50
0.75
1.00
Hypergeometric p−value
threshold
<=−0.5
>=0.5

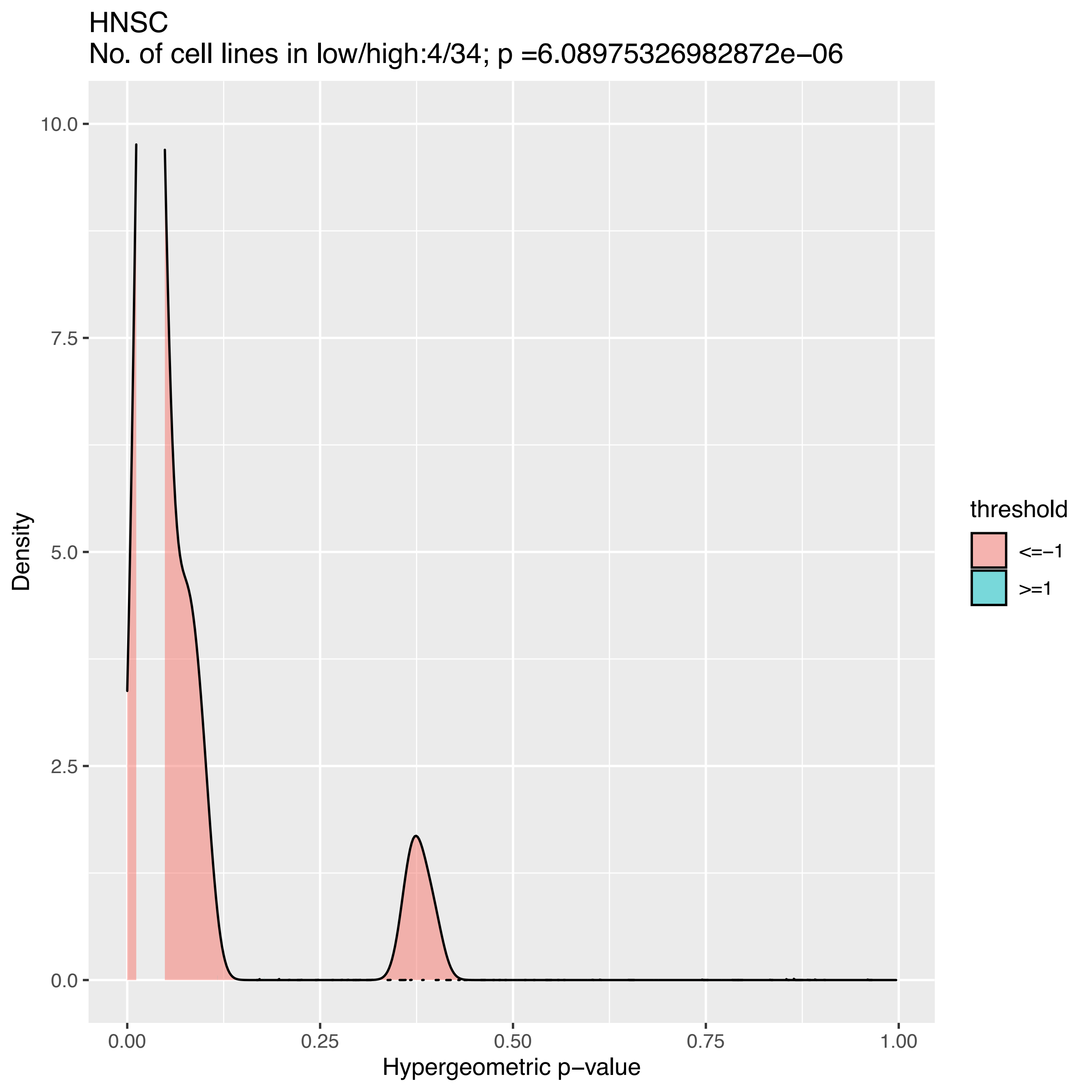
HNSC
No. of cell lines in low/high:4/34; p =6.08975326982872e−06
Density
10.0
7.5
5.0
2.5
0.0
0.00
0.25
0.50
0.75
1.00
Hypergeometric p−value
threshold
<=−1
>=1

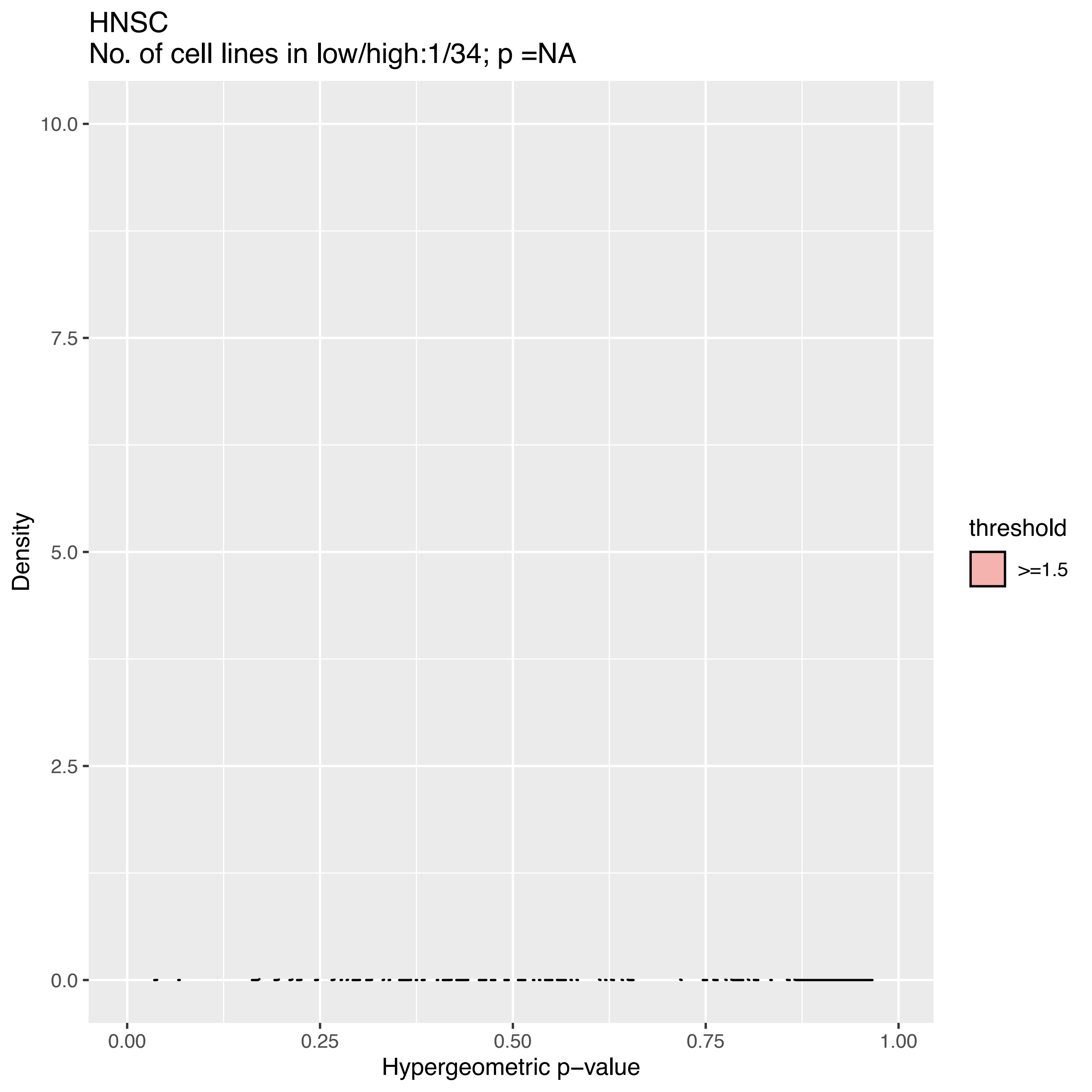

HNSC
No. of cell lines in low/high:1/34; p =NA
10.0
7.5
5.0
2.5
0.0
Density
0.00
0.25
0.50
0.75
1.00
Hypergeometric p−value
threshold
>=1.5

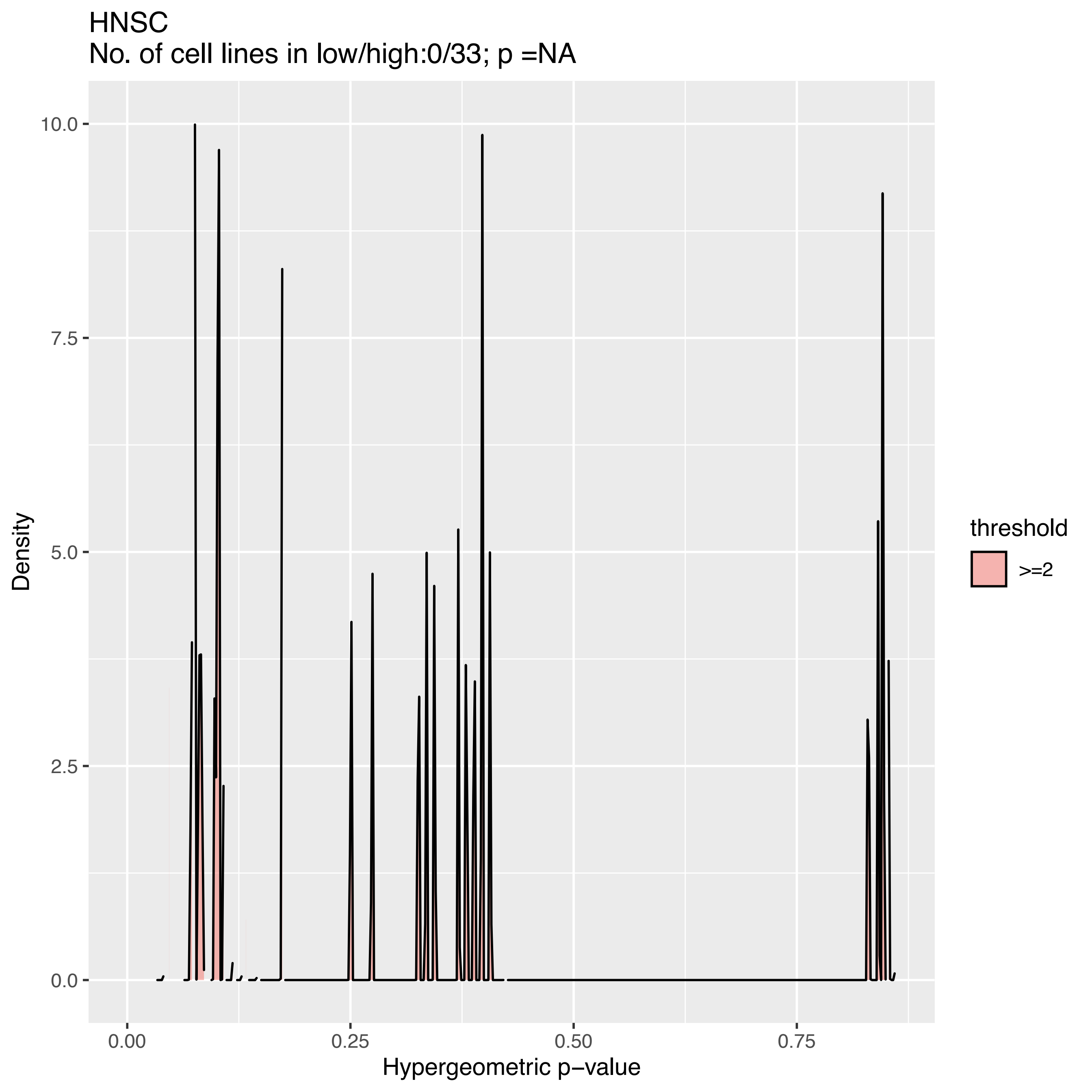
HNSC
No. of cell lines in low/high:0/33; p =NA
Density
10.0
7.5
5.0
2.5
0.0
0.00
0.25
0.50
0.75
Hypergeometric p−value
threshold
>=2

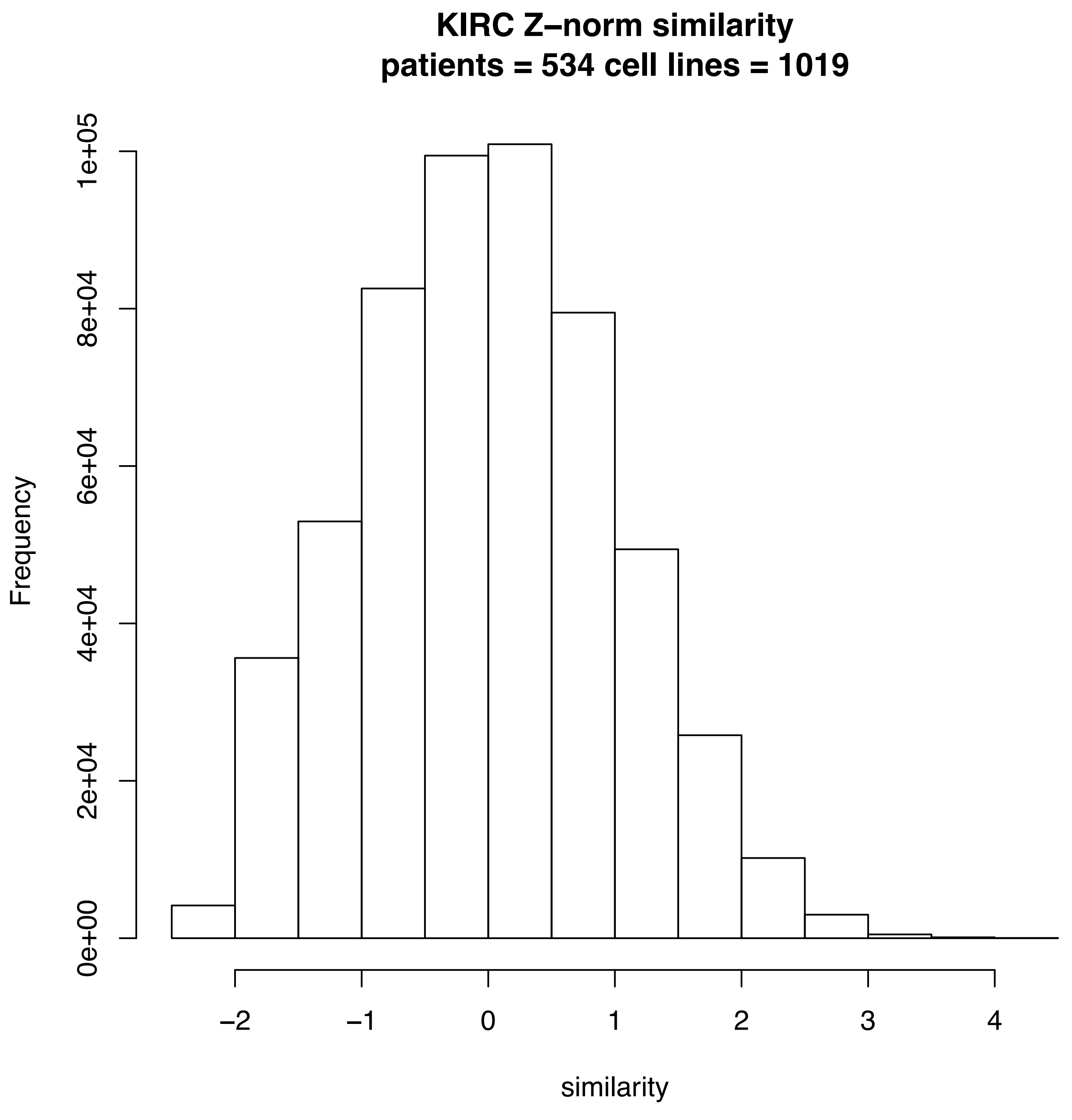
KIRC Z−norm similarity
patients = 534 cell lines = 1019
Frequency
0e+00
2e+04
4e+04
6e+04
8e+04
1e+05
−2
−1
0
1
2
3
4
similarity

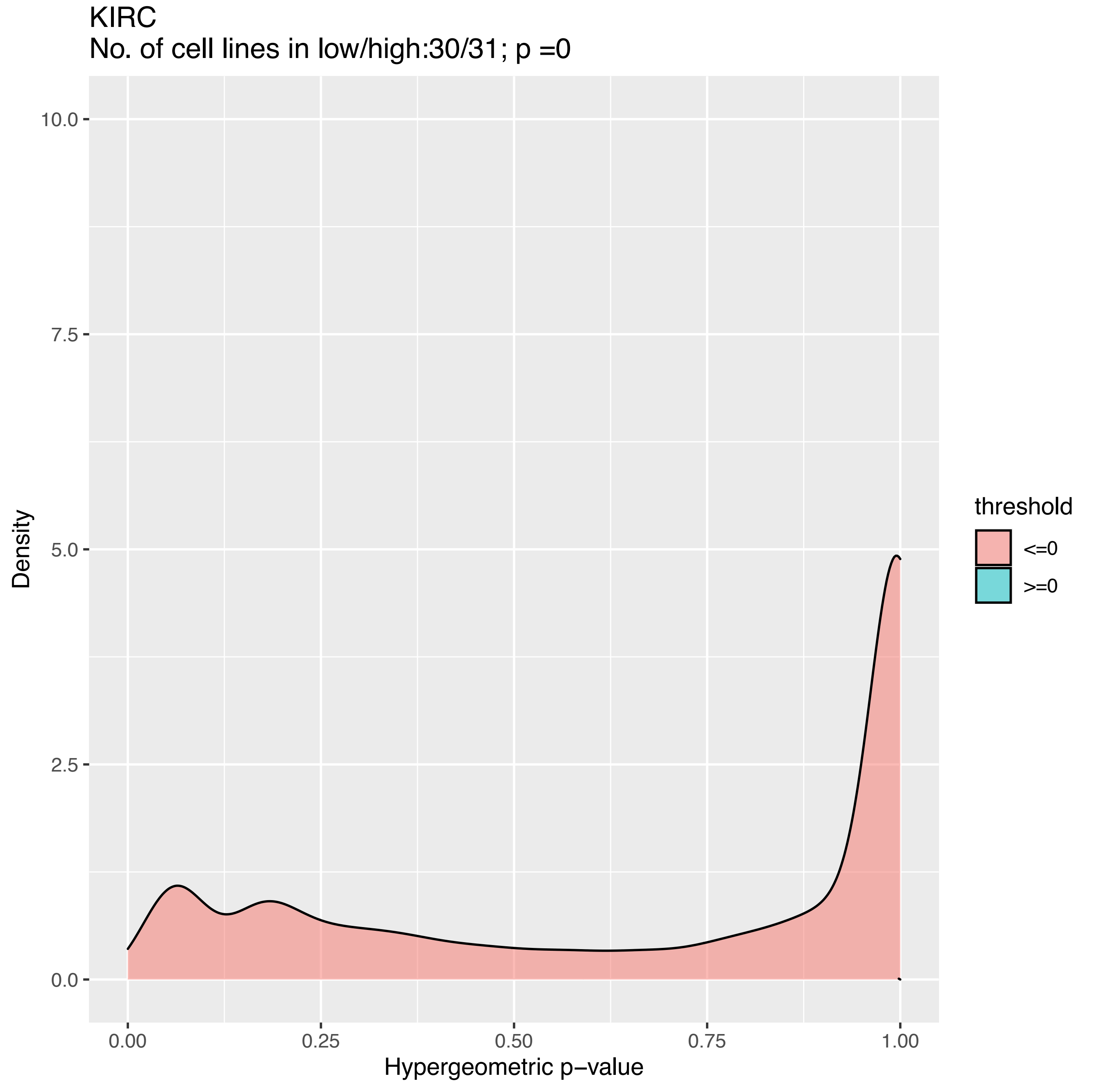
KIRC
No. of cell lines in low/high:30/31; p =0
Density
10.0
7.5
5.0
2.5
0.0
0.00
0.25
0.50
0.75
1.00
Hypergeometric p−value
threshold
<=0
>=0

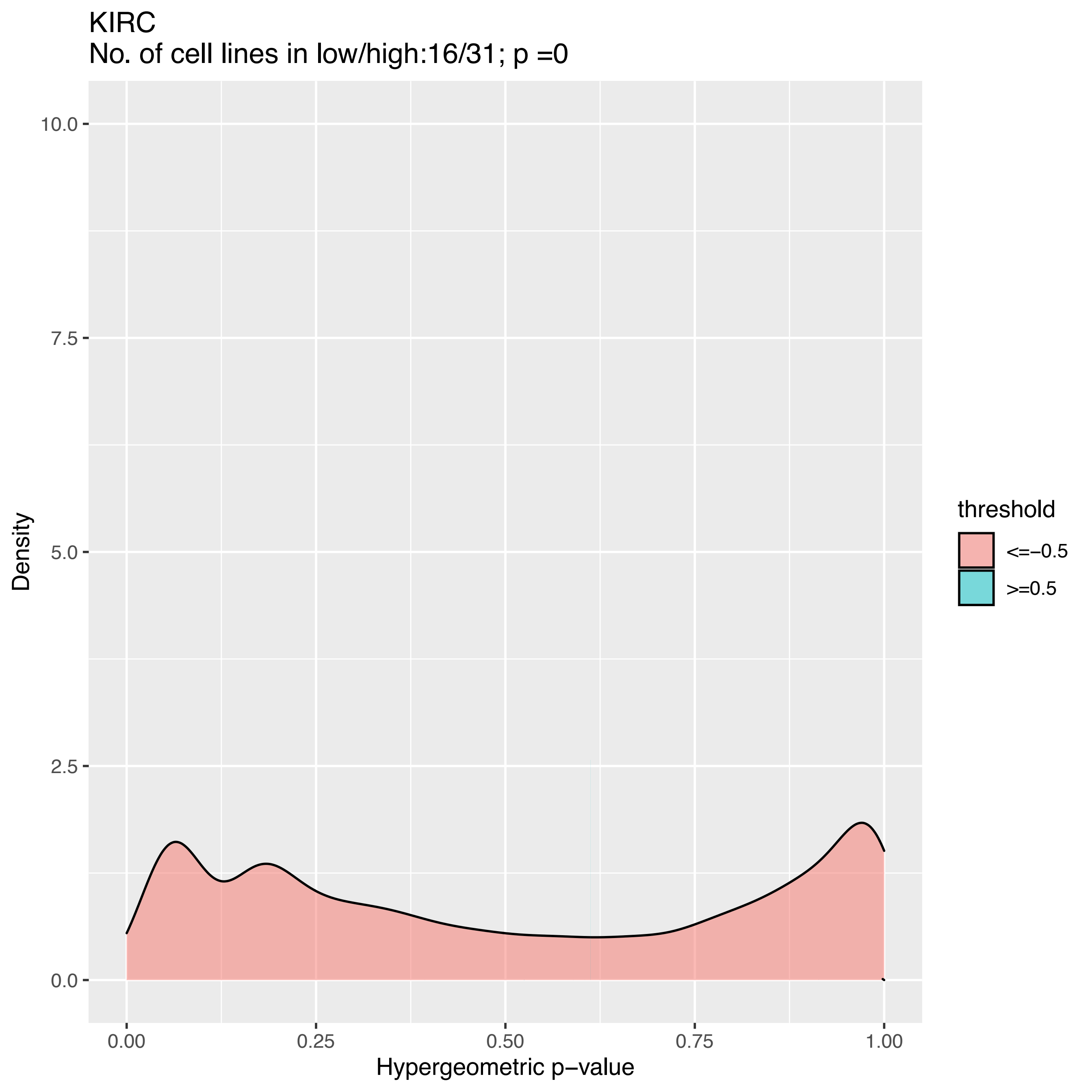

KIRC
No. of cell lines in low/high:16/31; p =0
Density
10.0
7.5
5.0
2.5
0.0
0.00
0.25
0.50
0.75
1.00
Hypergeometric p−value
threshold
<=−0.5
>=0.5

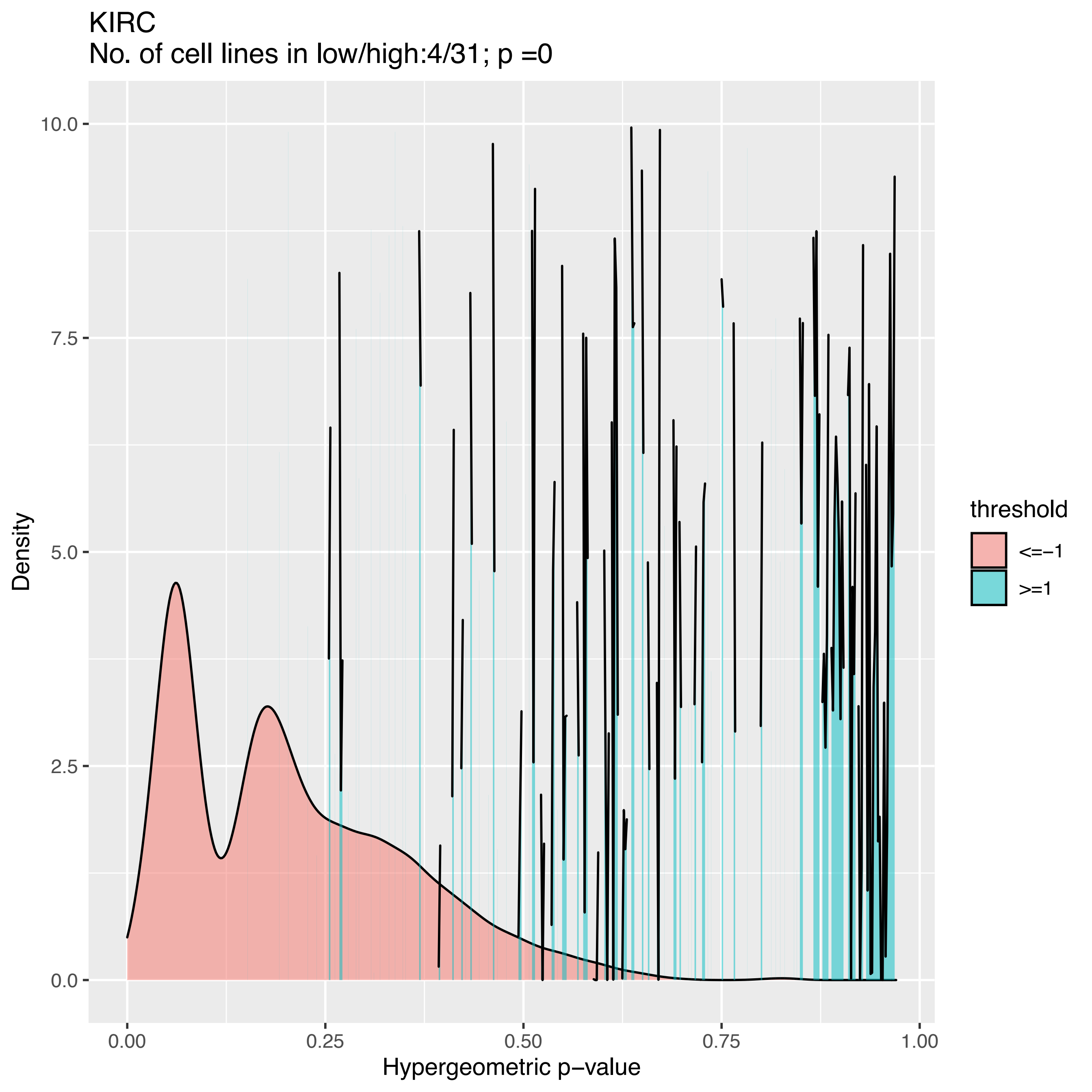

KIRC
No. of cell lines in low/high:4/31; p =0
Density
Hypergeometric p−value
10.0
7.5
5.0
2.5
0.0
0.00
0.25
0.50
0.75
1.00
threshold
<=−1
>=1

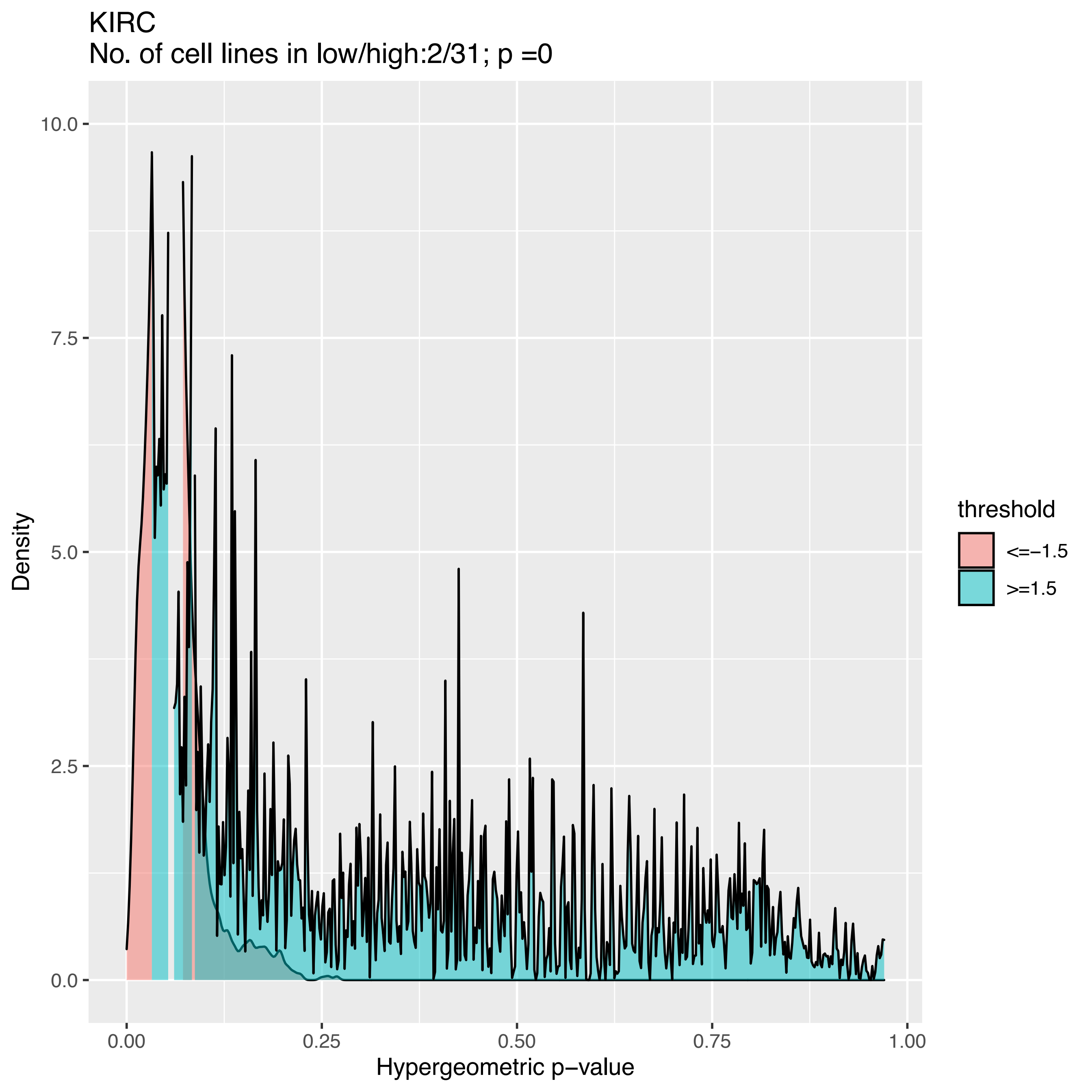

KIRC
No. of cell lines in low/high:2/31; p =0
Density
10.0
7.5
5.0
2.5
0.0
0.00
0.25
0.50
0.75
1.00
Hypergeometric p−value
threshold
<=−1.5
>=1.5

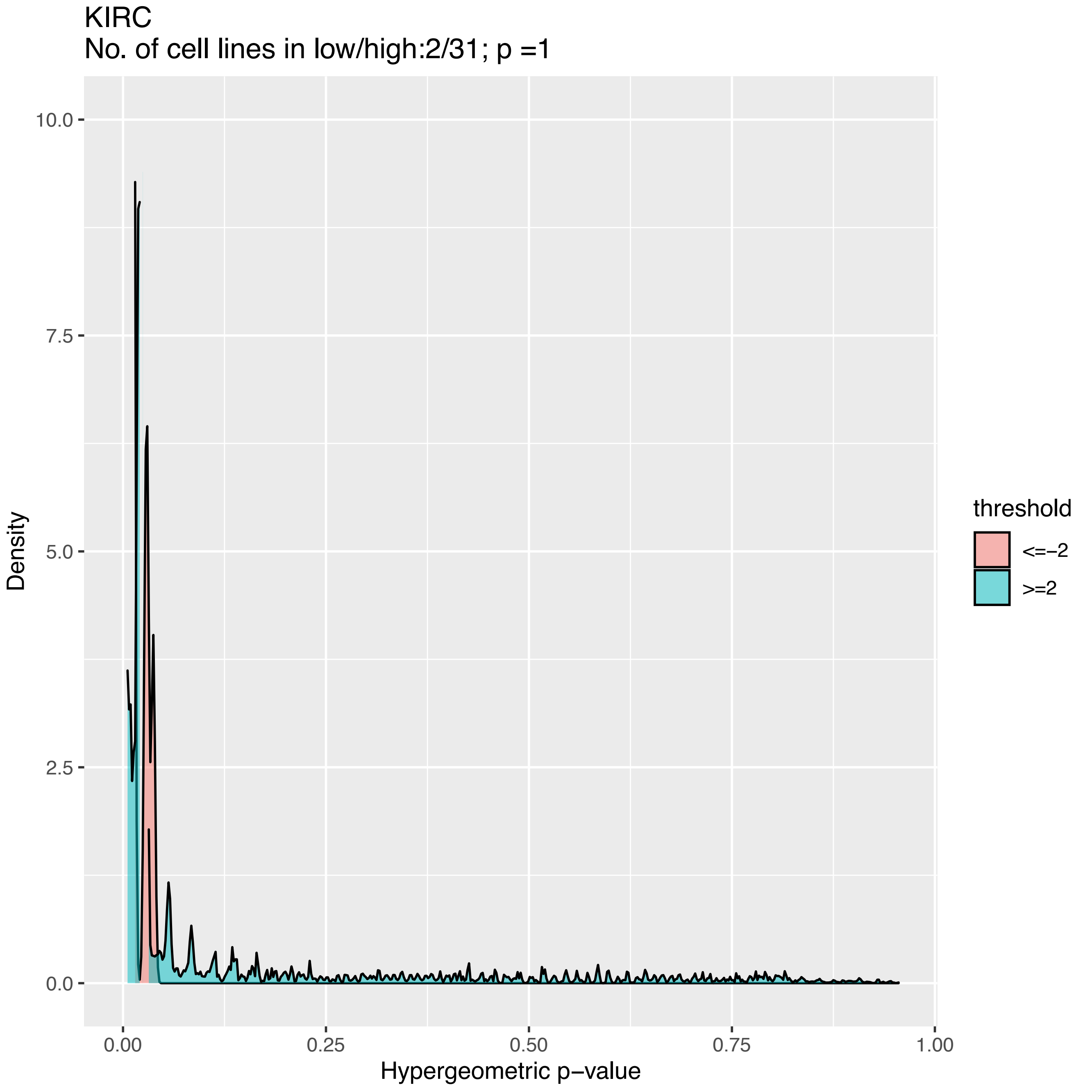
KIRC
No. of cell lines in low/high:2/31; p =1
Density
10.0
7.5
5.0
2.5
0.0
0.00
0.25
0.50
0.75
1.00
Hypergeometric p−value
threshold
<=−2
>=2

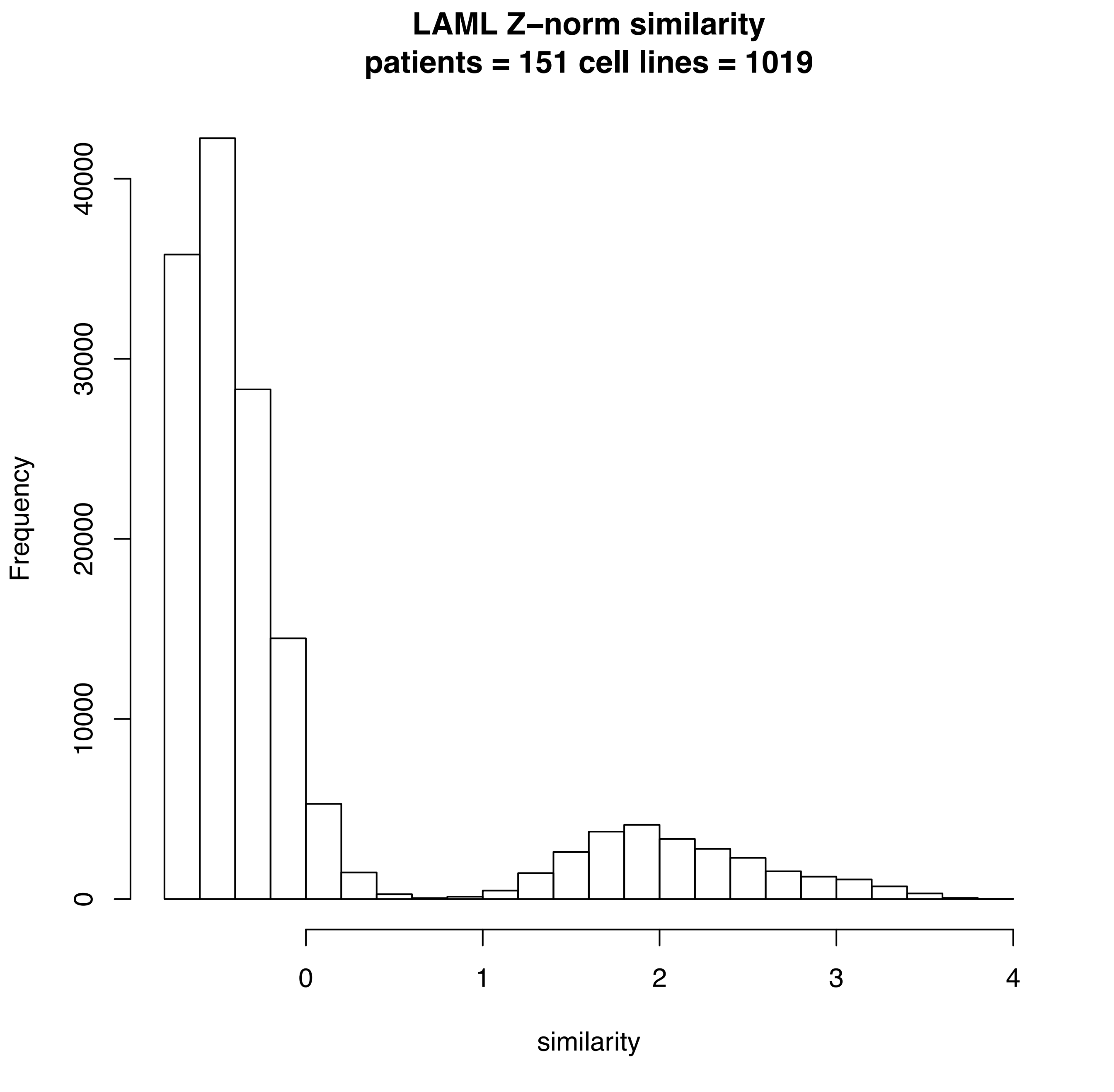

LAML Z–norm similarity
patients = 151 cell lines = 1019
Frequency
0
10000
20000
30000
40000
0
1
2
3
4
similarity

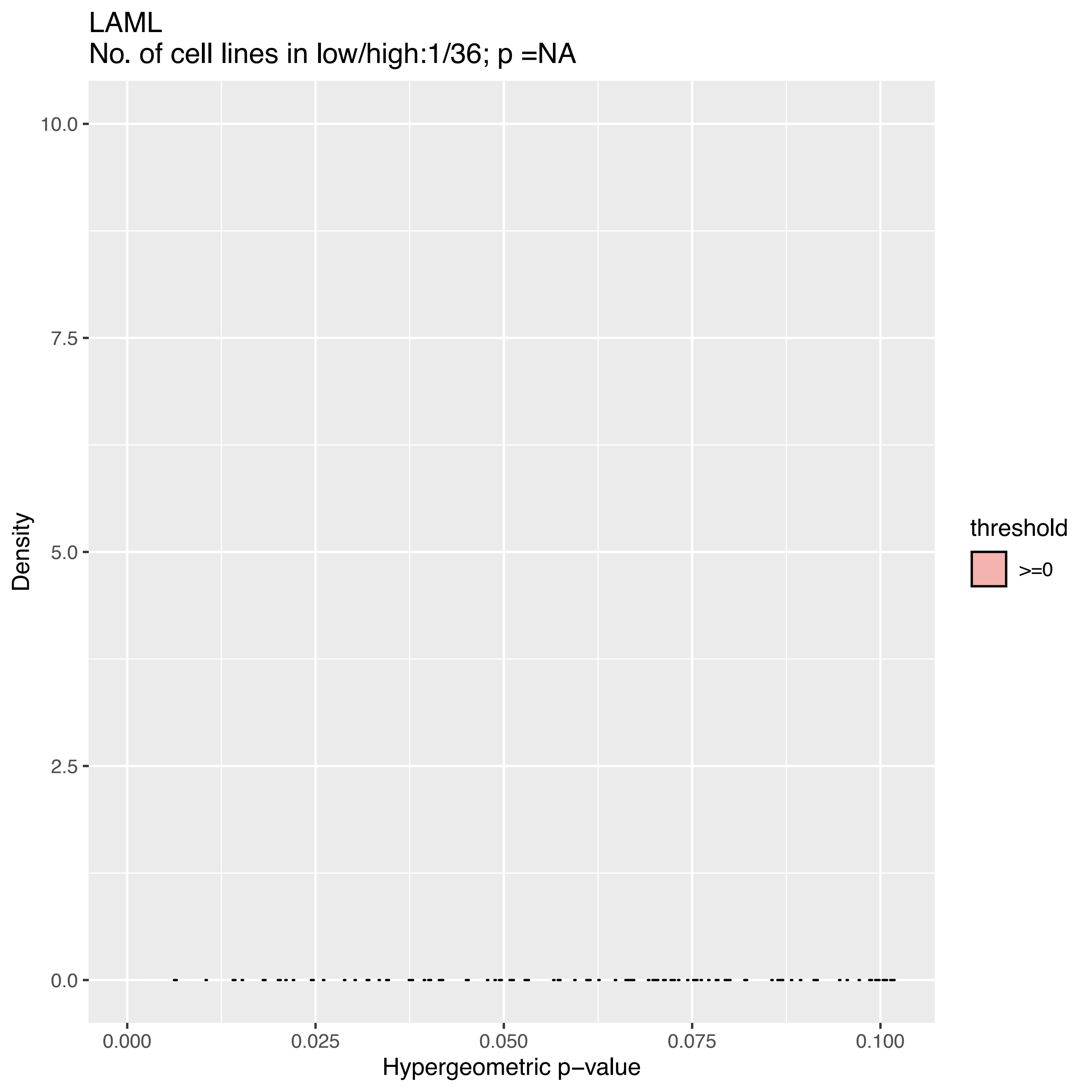

LAML
No. of cell lines in low/high:1/36; p =NA
10.0
7.5
5.0
2.5
0.0
Density
0.000
0.025
0.050
0.075
0.100
Hypergeometric p−value
threshold
>=0

LAML
No. of cell lines in low/high:1/36; p =NA

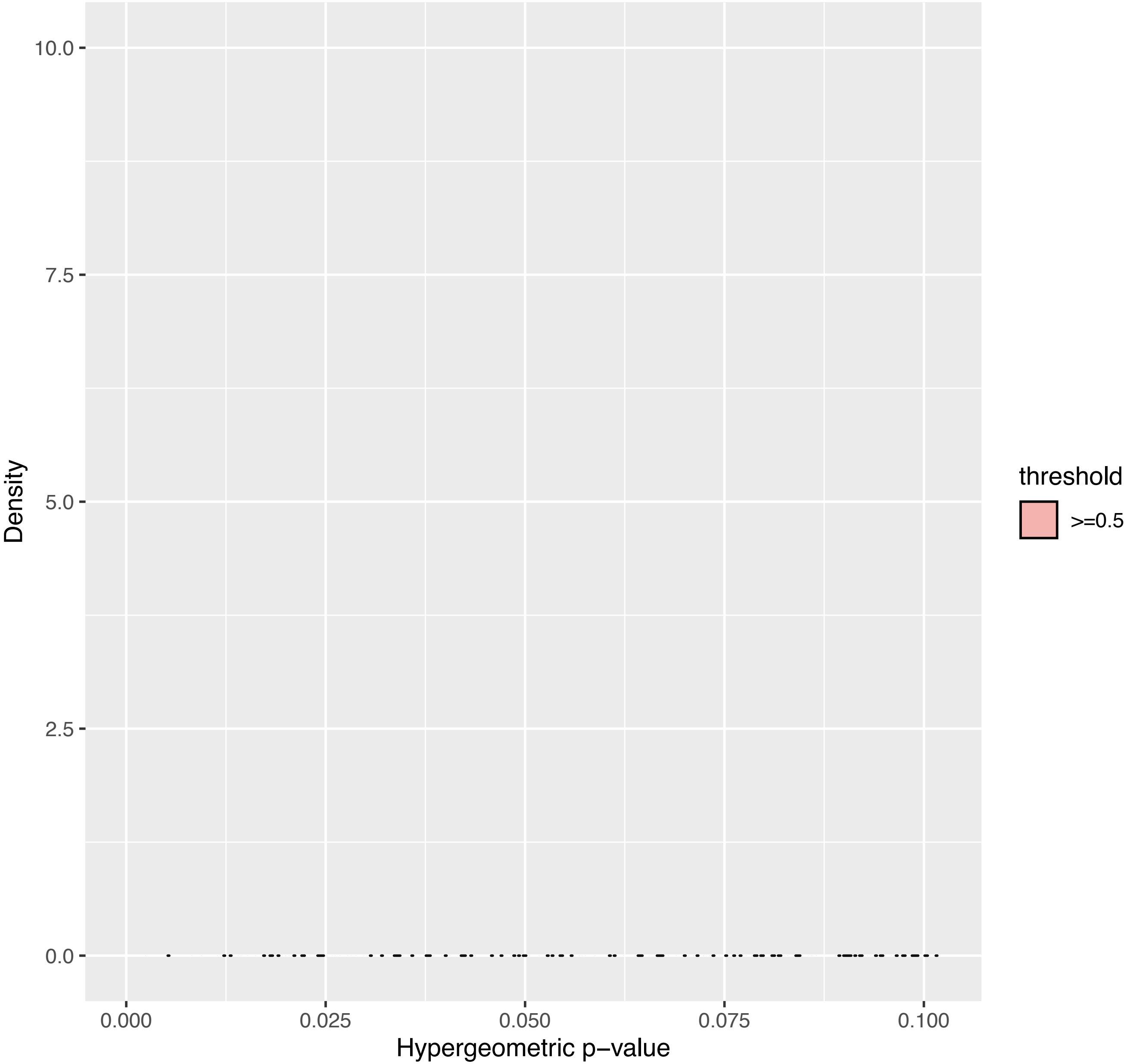

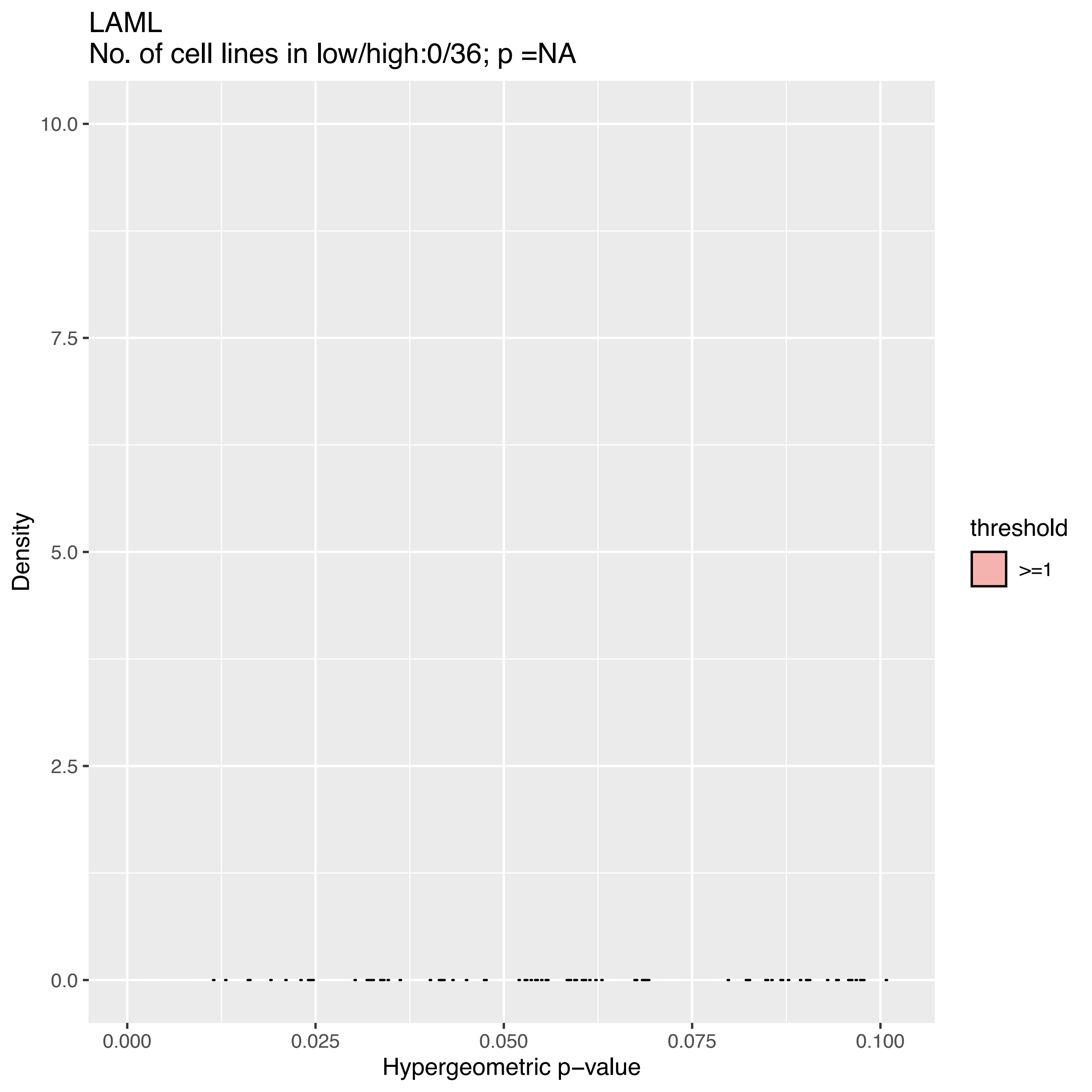

LAML
No. of cell lines in low/high:0/36; p =NA
10.0
7.5
5.0
2.5
0.0
Density
0.000
0.025
0.050
0.075
0.100
Hypergeometric p−value
threshold
>=1

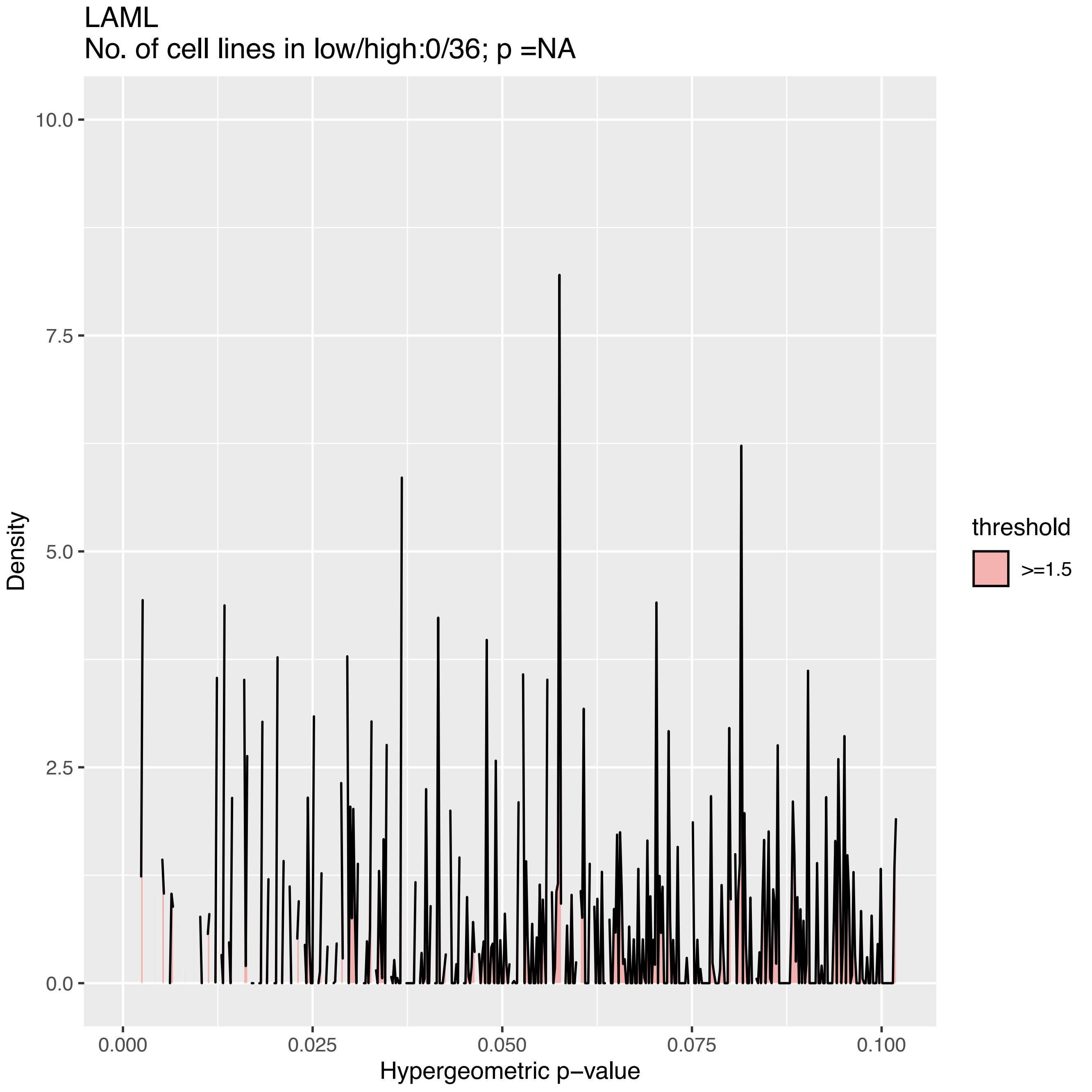
LAML
No. of cell lines in low/high:0/36; p =NA
Density
10.0
7.5
5.0
2.5
0.0
0.000
0.025
0.050
0.075
0.100
Hypergeometric p−value
threshold
>=1.5

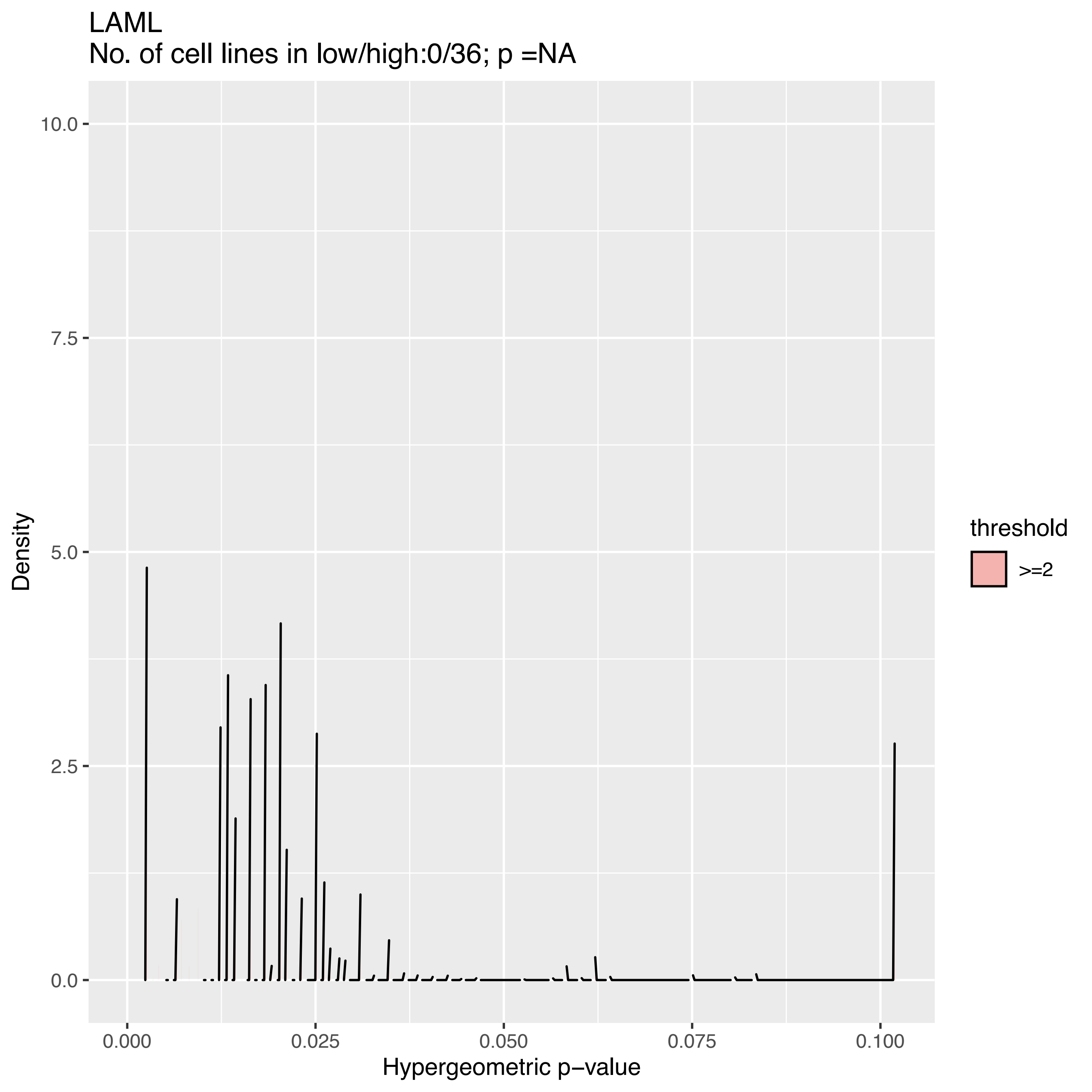

LAML
No. of cell lines in low/high:0/36; p =NA
Density
10.0
7.5
5.0
2.5
0.0
0.000
0.025
0.050
0.075
0.100
Hypergeometric p−value
threshold
>=2

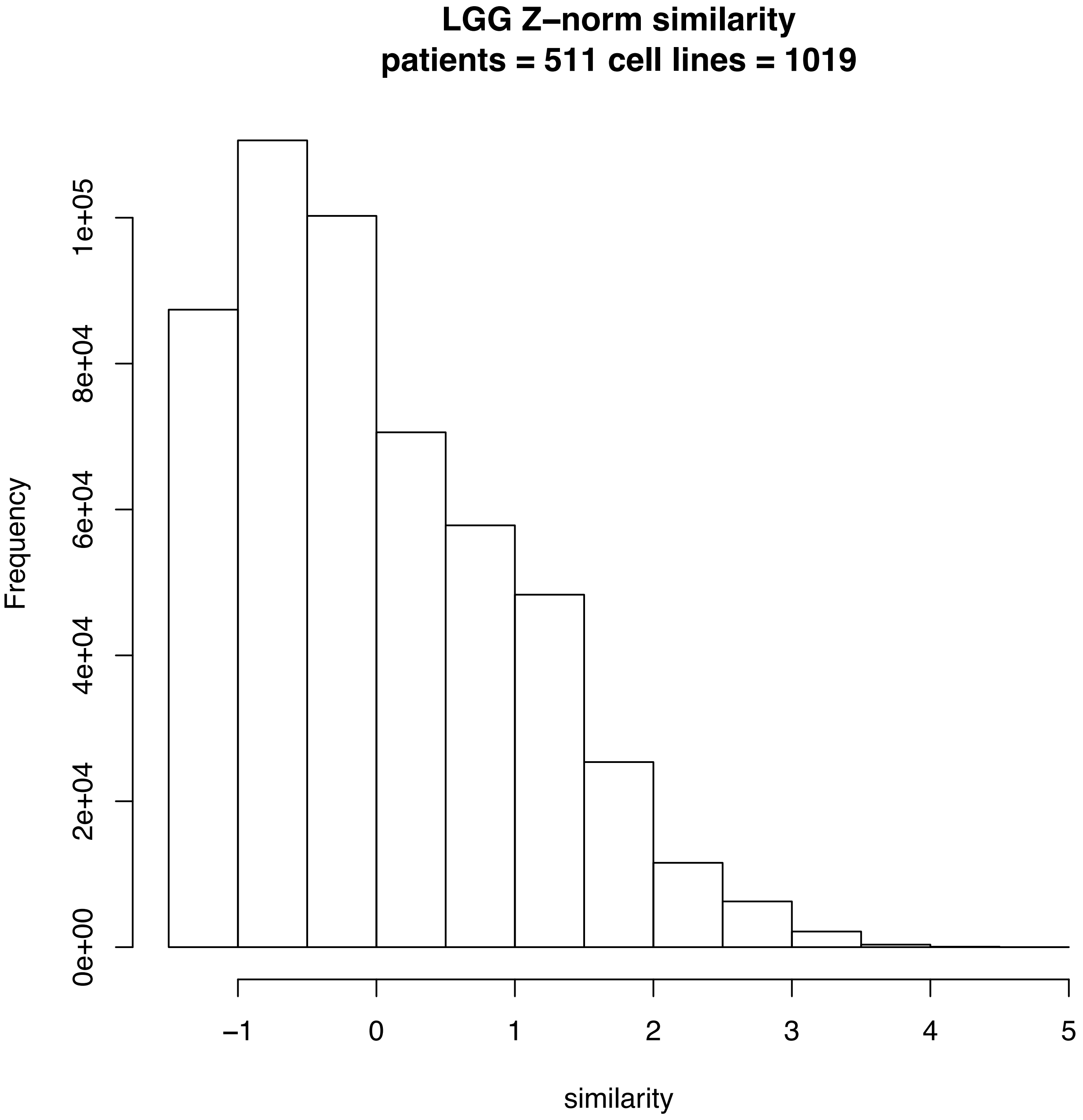
LGG Z–norm similarity
patients = 511 cell lines = 1019
Frequency
0e+00
2e+04
4e+04
6e+04
8e+04
1e+05
–1
0
1
2
3
4
5
similarity

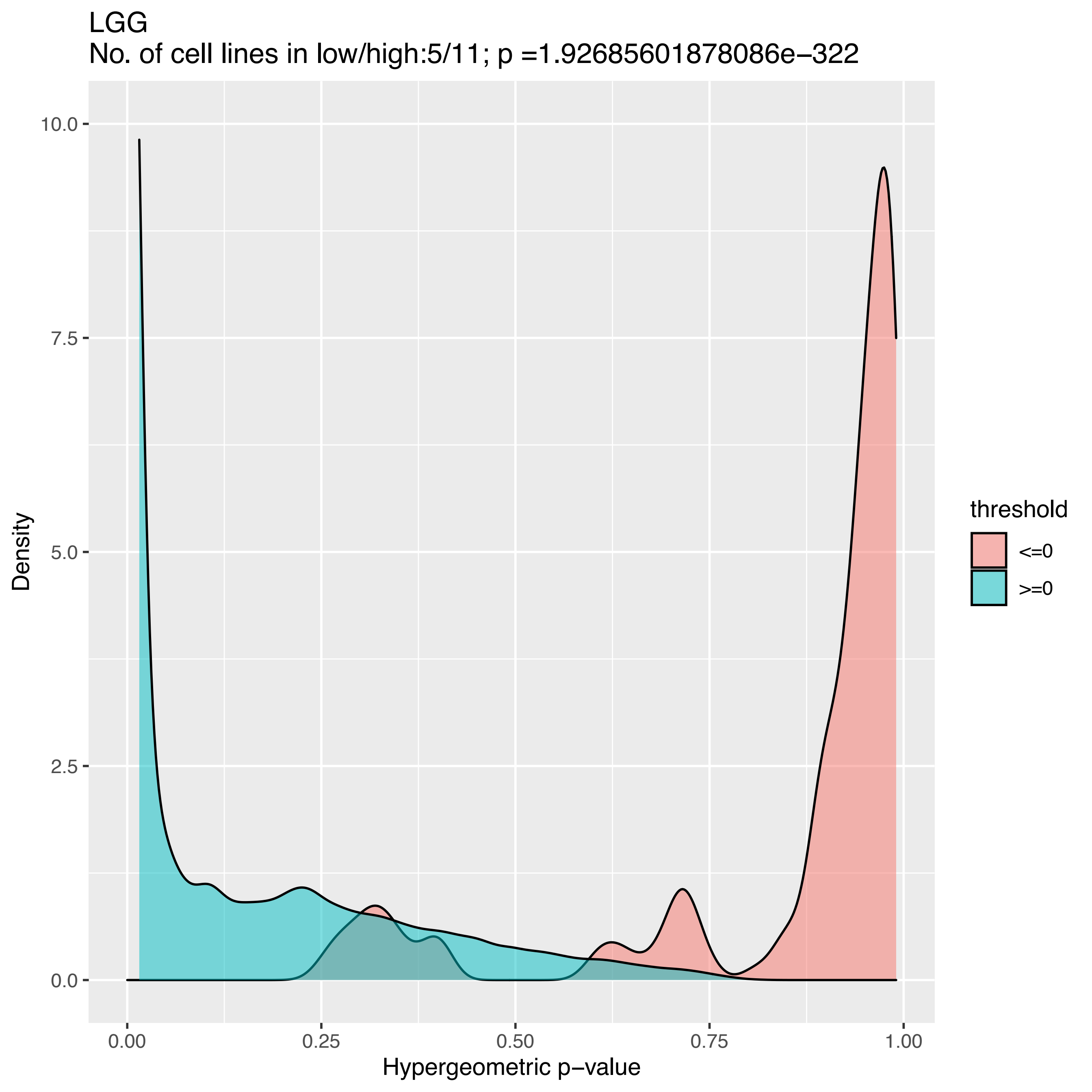
LGG
No. of cell lines in low/high:5/11; p =1.92685601878086e−322
Density
Hypergeometric p−value
10.0
7.5
5.0
2.5
0.0
0.00
0.25
0.50
0.75
1.00
threshold
<=0
>=0

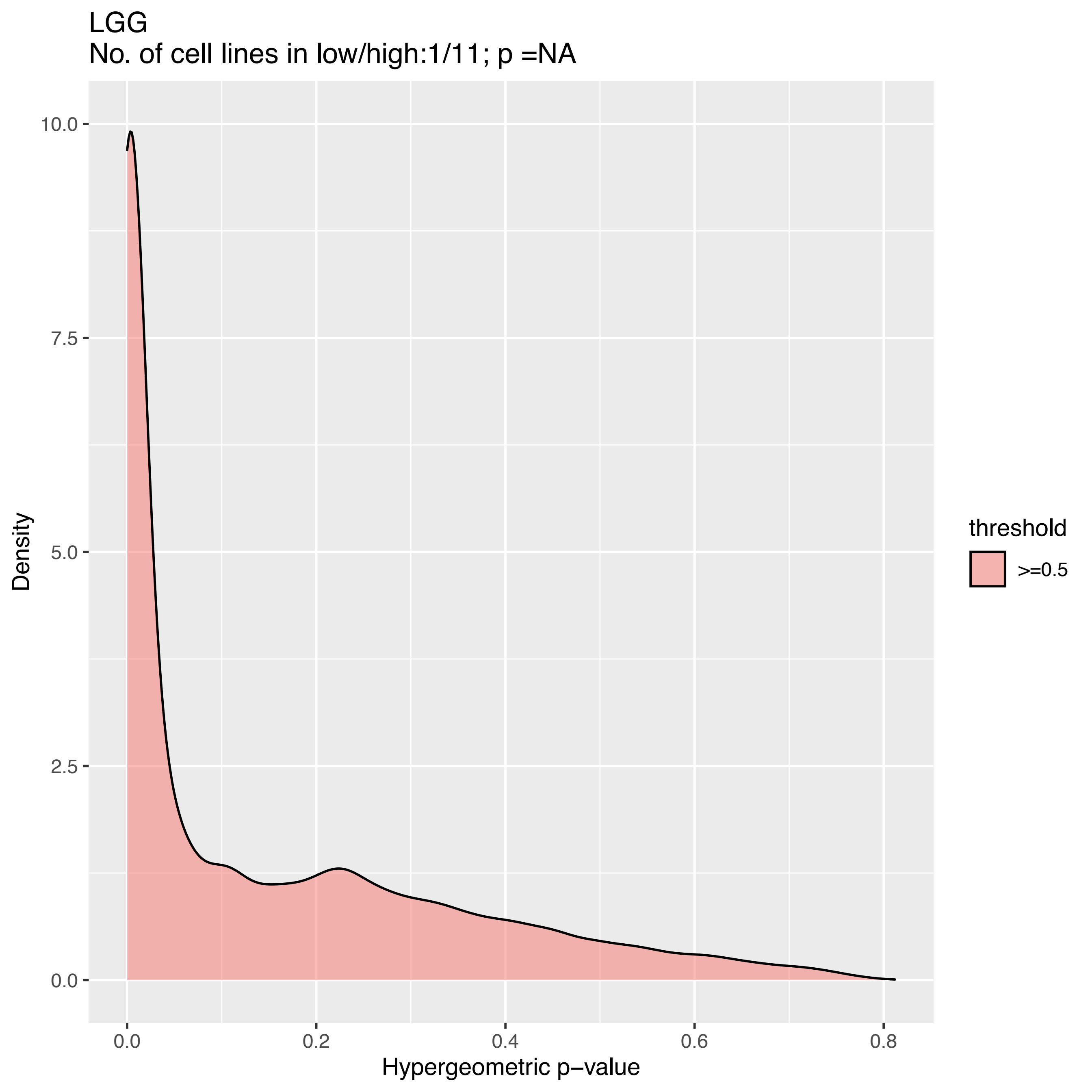
LGG
No. of cell lines in low/high:1/11; p =NA
Density
10.0
7.5
5.0
2.5
0.0
0.0
0.2
0.4
0.6
0.8
Hypergeometric p−value
threshold
>=0.5

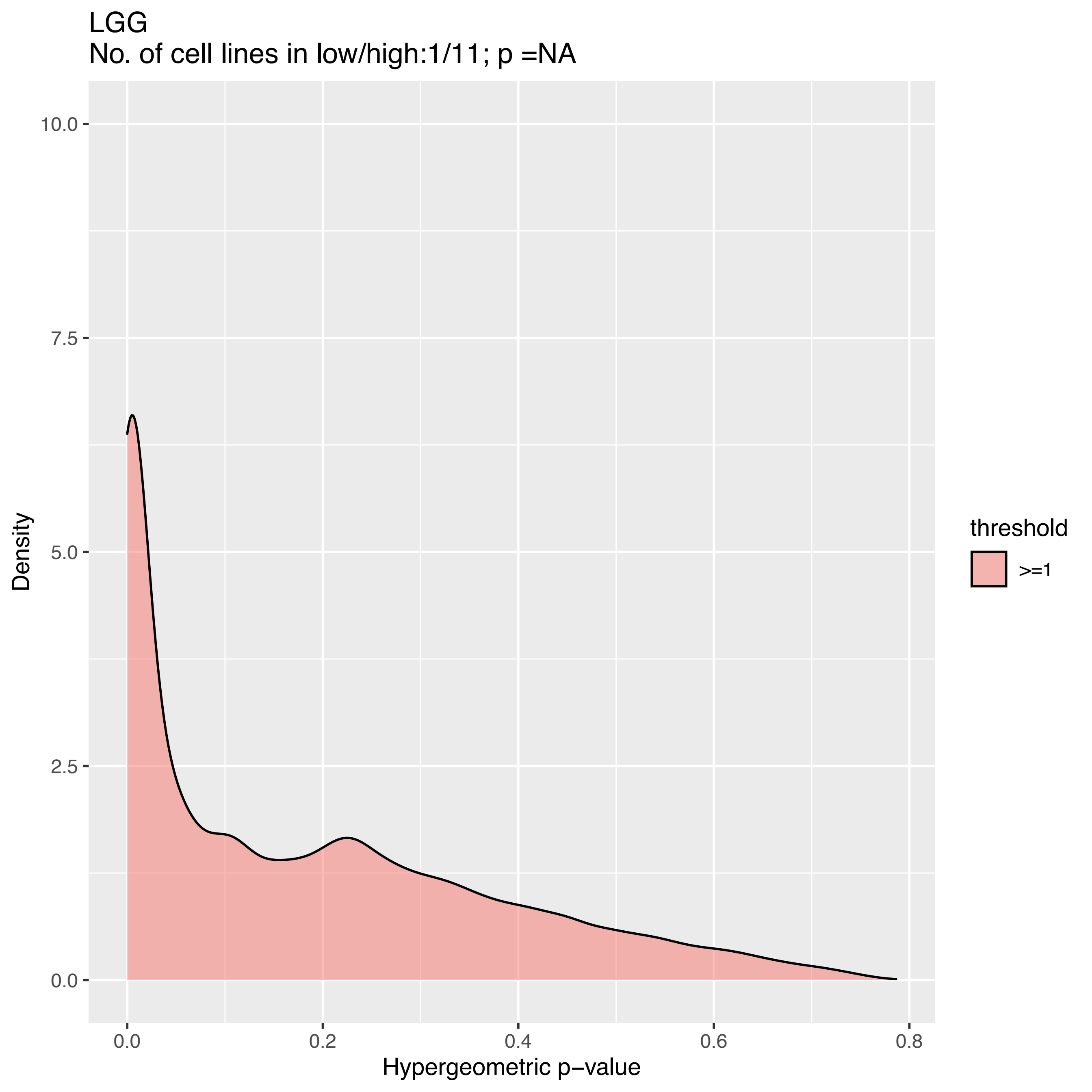

LGG
No. of cell lines in low/high:1/11; p =NA
10.0
7.5
5.0
2.5
0.0
Density
0.0
0.2
0.4
0.6
0.8
Hypergeometric p−value
threshold
>=1

LGG
No. of cell lines in low/high:0/11; p =NA

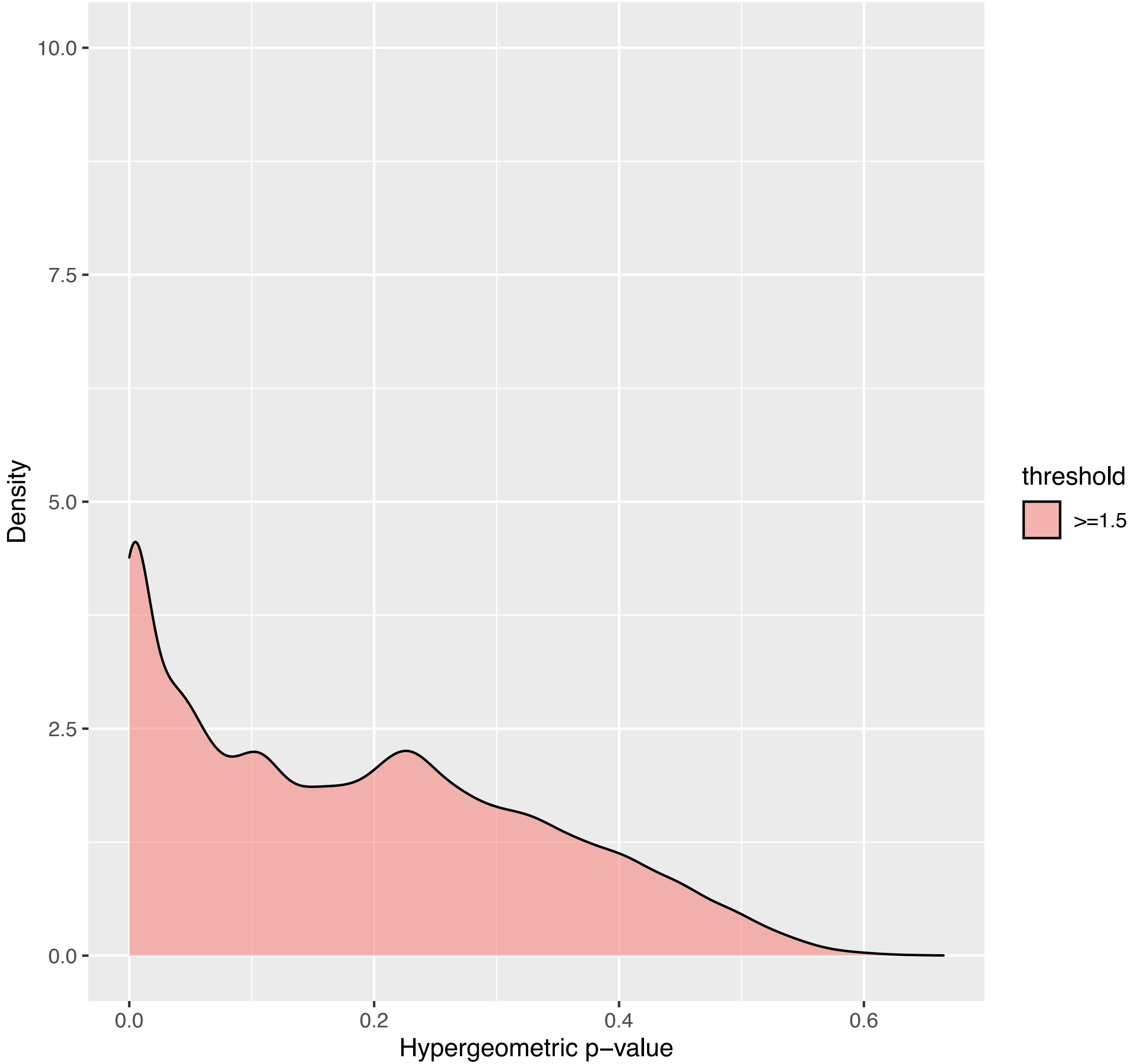

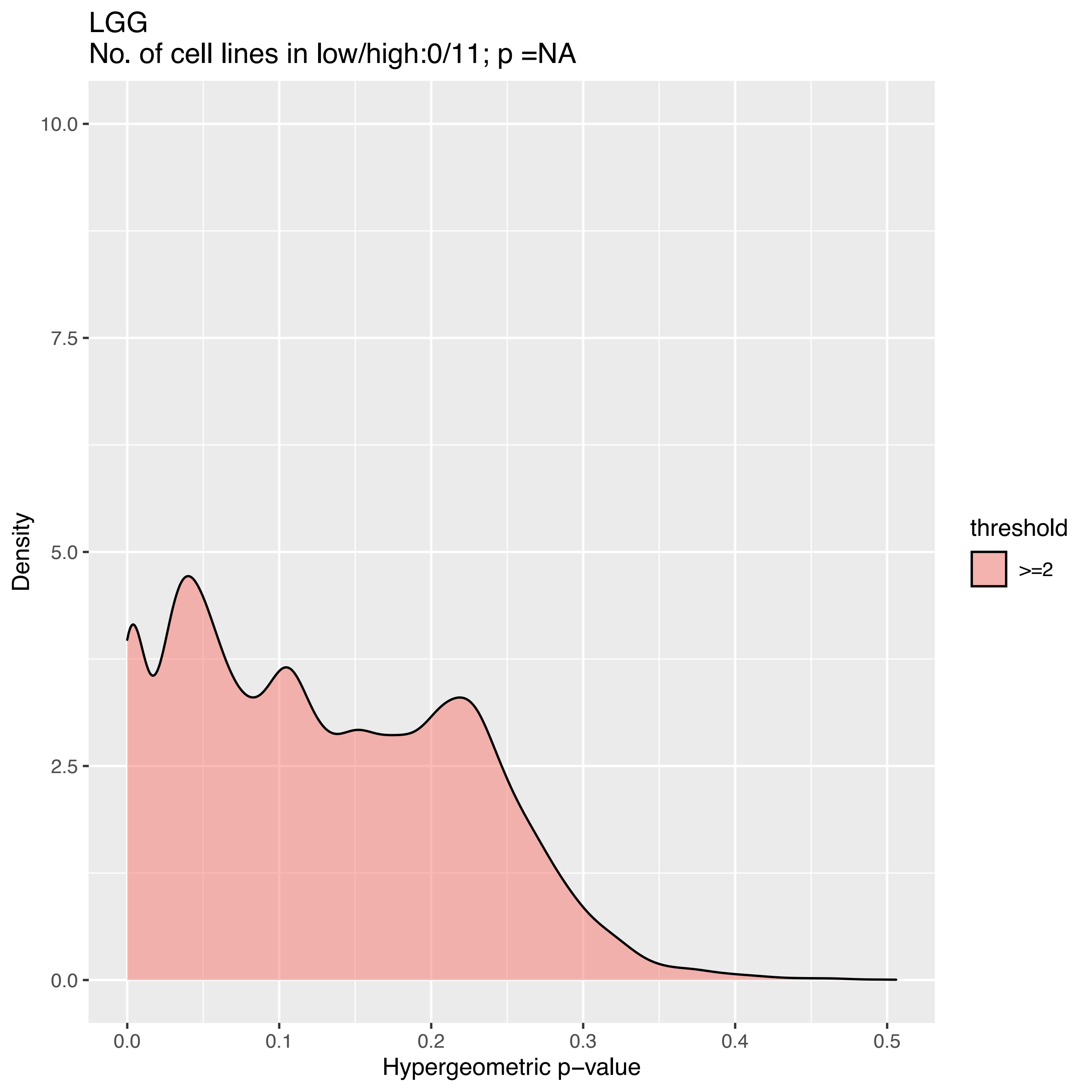
LGG
No. of cell lines in low/high:0/11; p =NA
10.0
7.5
5.0
2.5
0.0
Density
0.0
0.1
0.2
0.3
0.4
0.5
Hypergeometric p−value
threshold
>=2

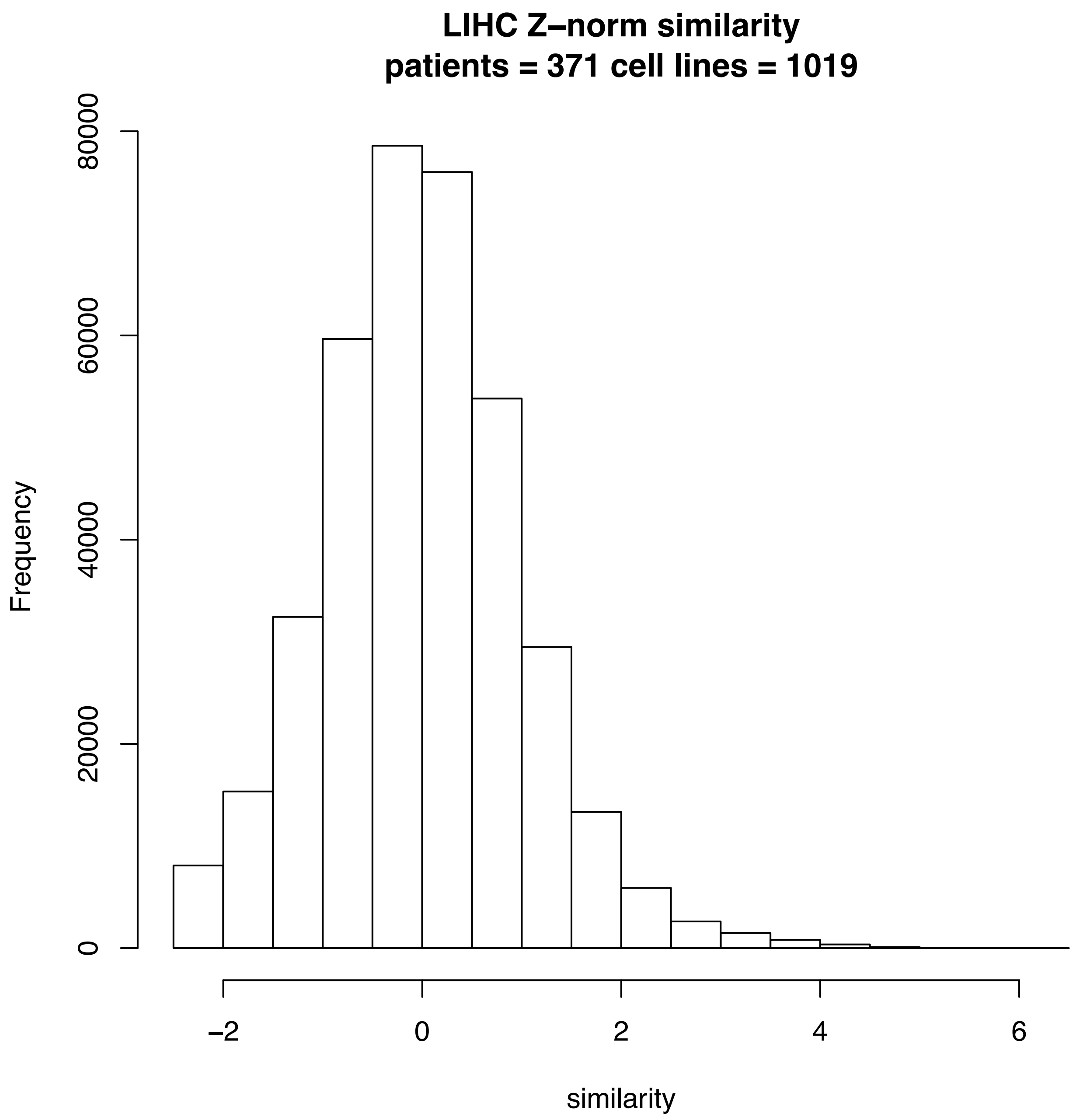
LIHC Z–norm similarity
patients = 371 cell lines = 1019
Frequency
0
20000
40000
60000
80000
–2
0
2
4
6
similarity

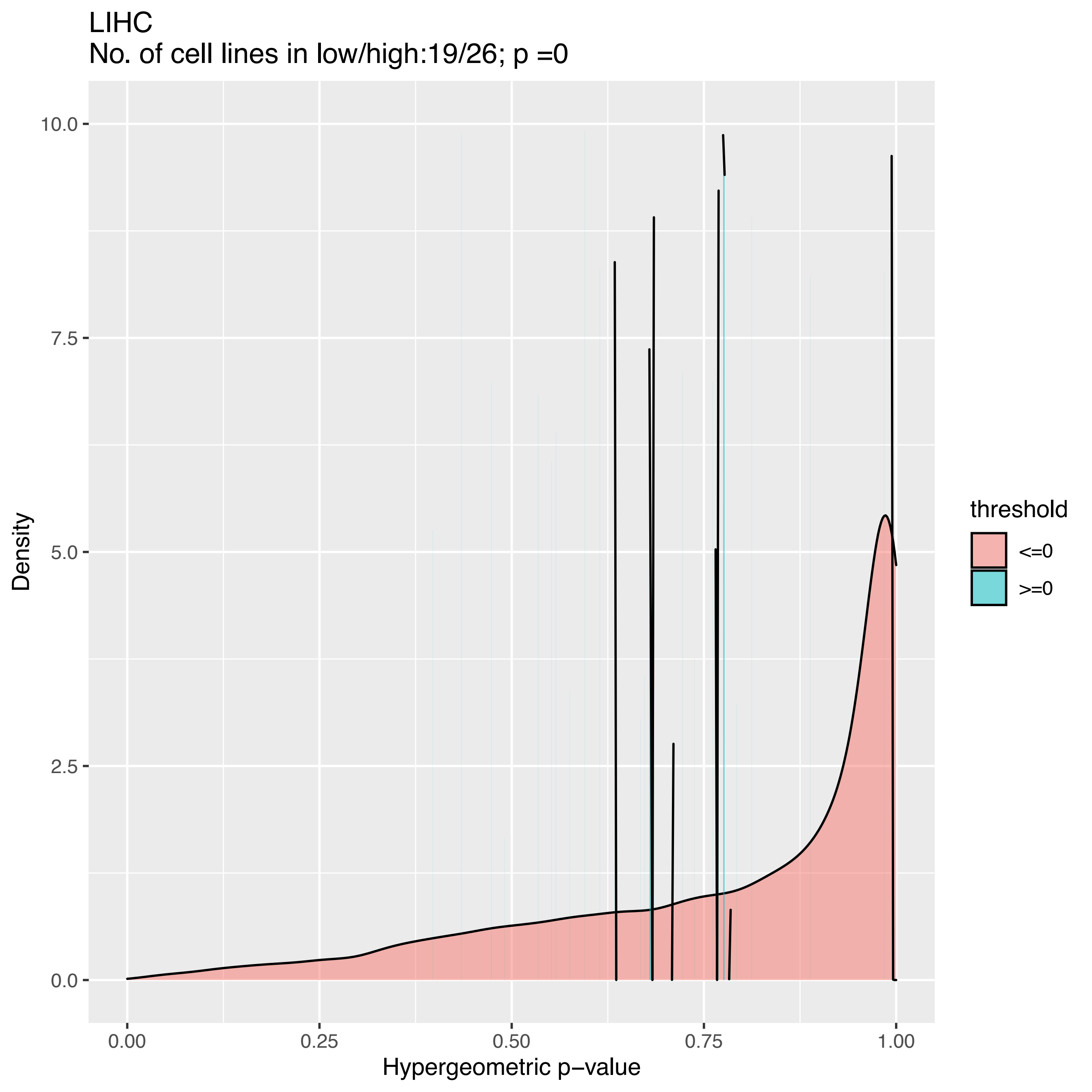

LIHC
No. of cell lines in low/high:19/26; p =0
Density
10.0
7.5
5.0
2.5
0.0
0.00
0.25
0.50
0.75
1.00
Hypergeometric p−value
threshold
<=0
>=0

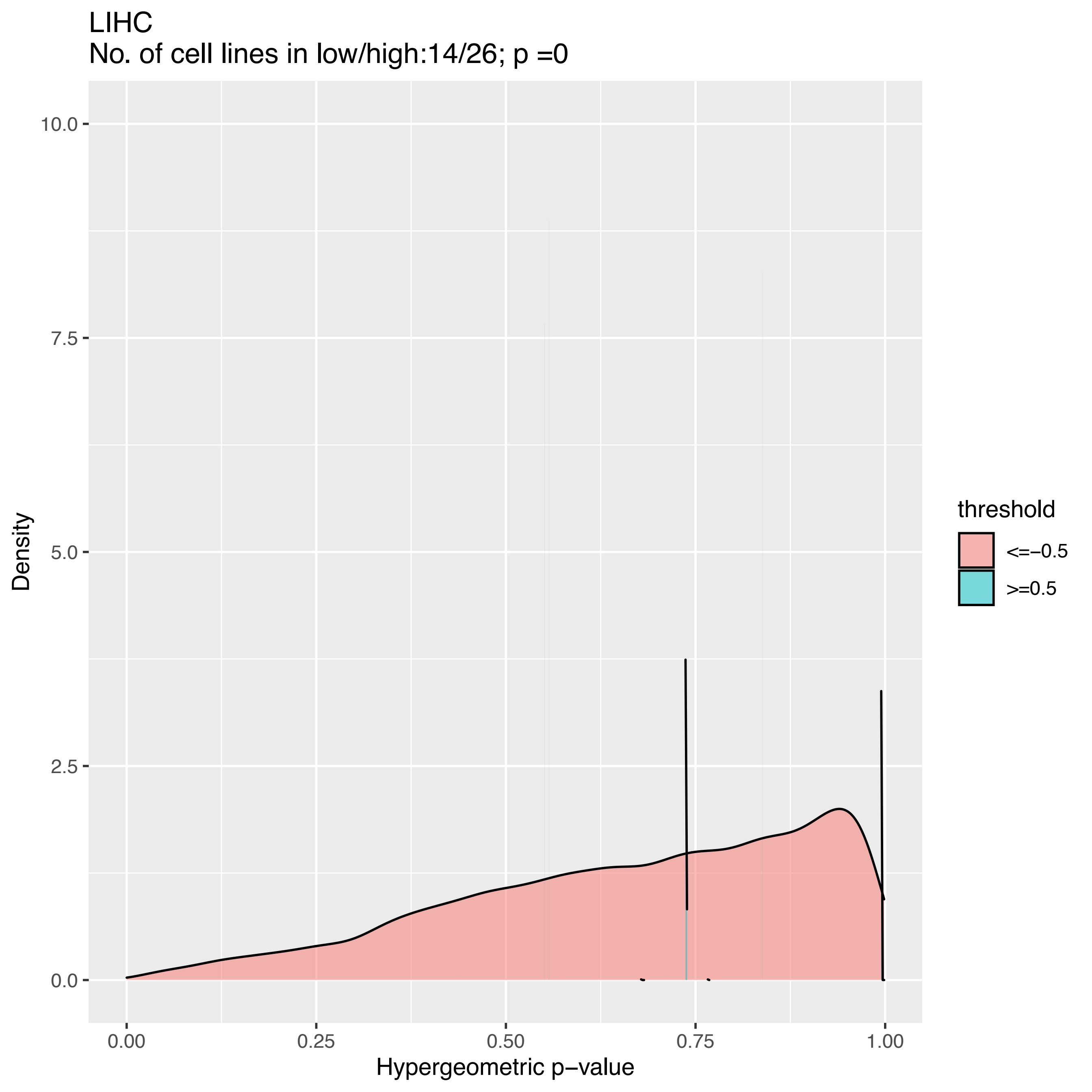
LIHC
No. of cell lines in low/high:14/26; p =0
10.0
7.5
5.0
2.5
0.0
Density
0.00
0.25
0.50
0.75
1.00
Hypergeometric p−value
threshold
<=−0.5
>=0.5

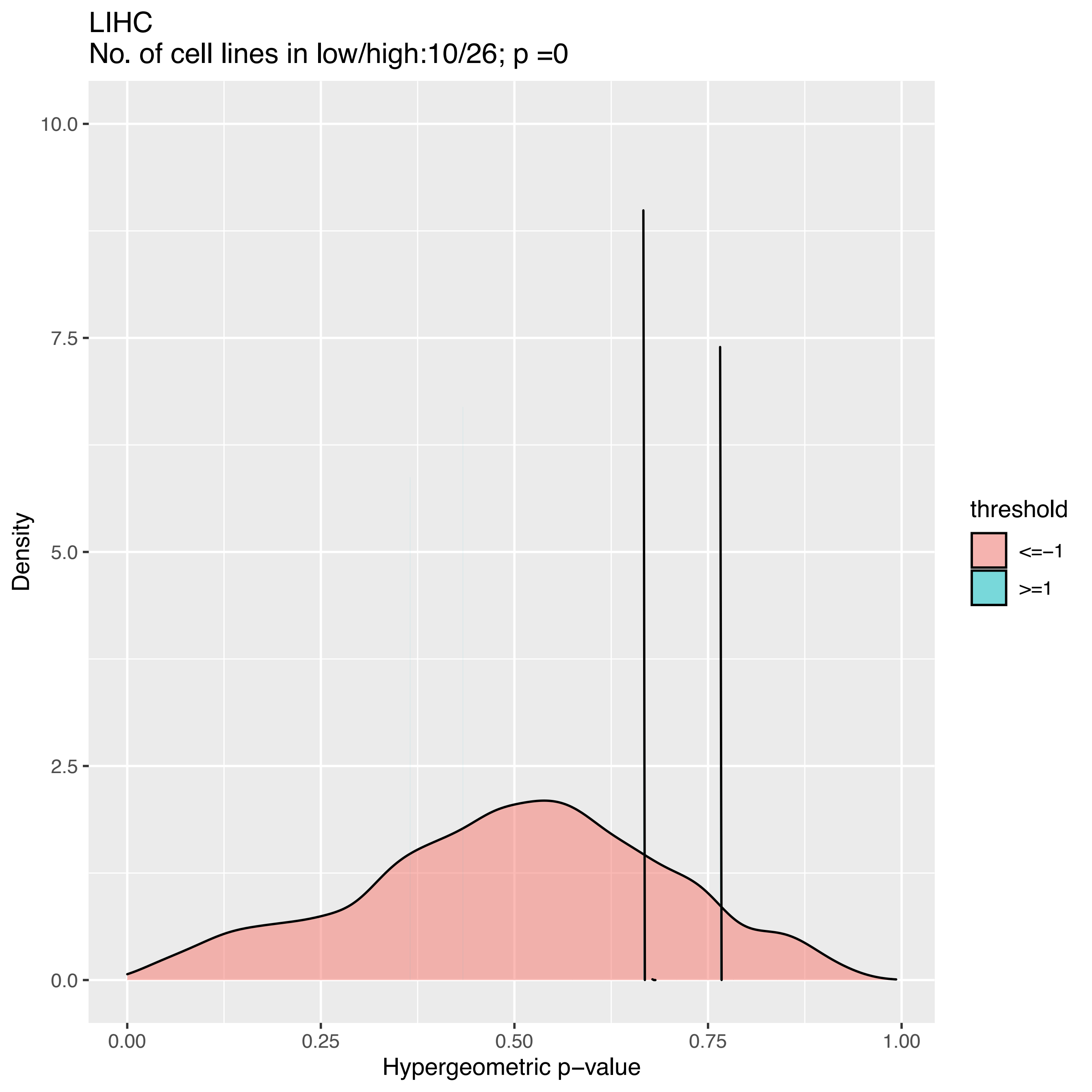

LIHC
No. of cell lines in low/high:10/26; p =0
Density
10.0
7.5
5.0
2.5
0.0
0.00
0.25
0.50
0.75
1.00
Hypergeometric p−value
threshold
<=−1
>=1

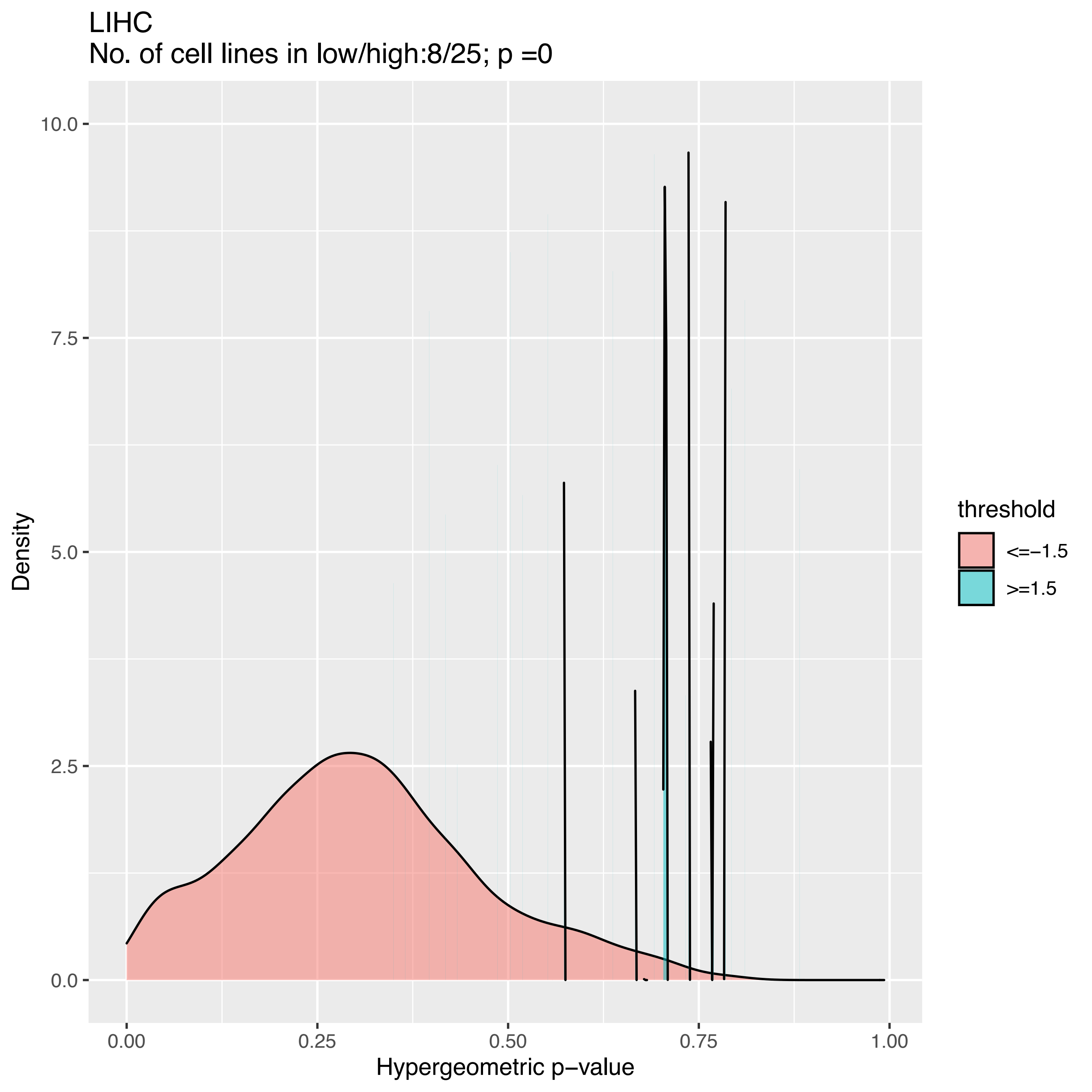
LIHC
No. of cell lines in low/high:8/25; p =0
Density
10.0
7.5
5.0
2.5
0.0
0.00
0.25
0.50
0.75
1.00
Hypergeometric p−value
threshold
<=−1.5
>=1.5

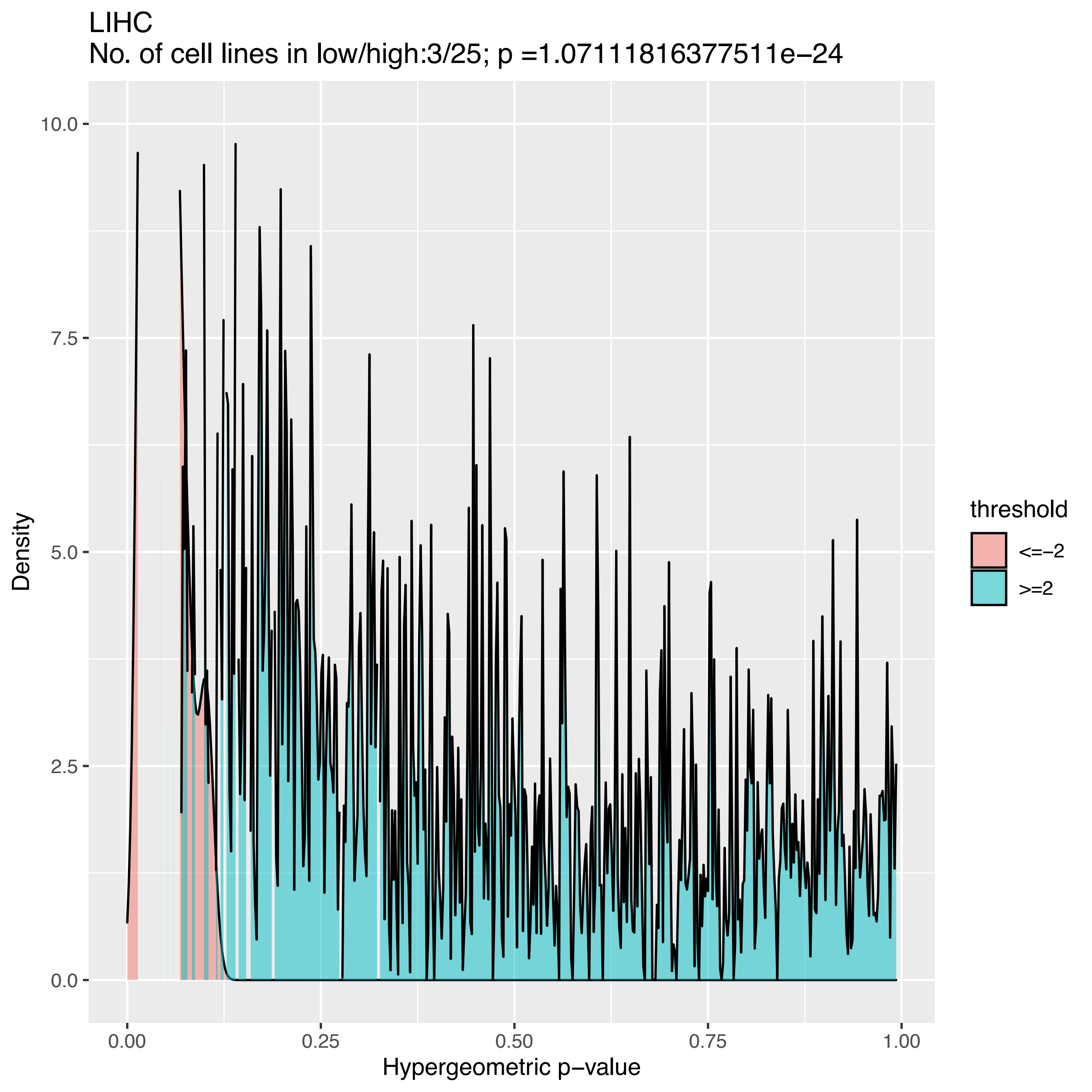
LIHC
No. of cell lines in low/high:3/25; p =1.07111816377511e−24
Density
Hypergeometric p−value
0.00
0.25
0.50
0.75
1.00
0.0
2.5
5.0
7.5
10.0
threshold
<=−2
>=2

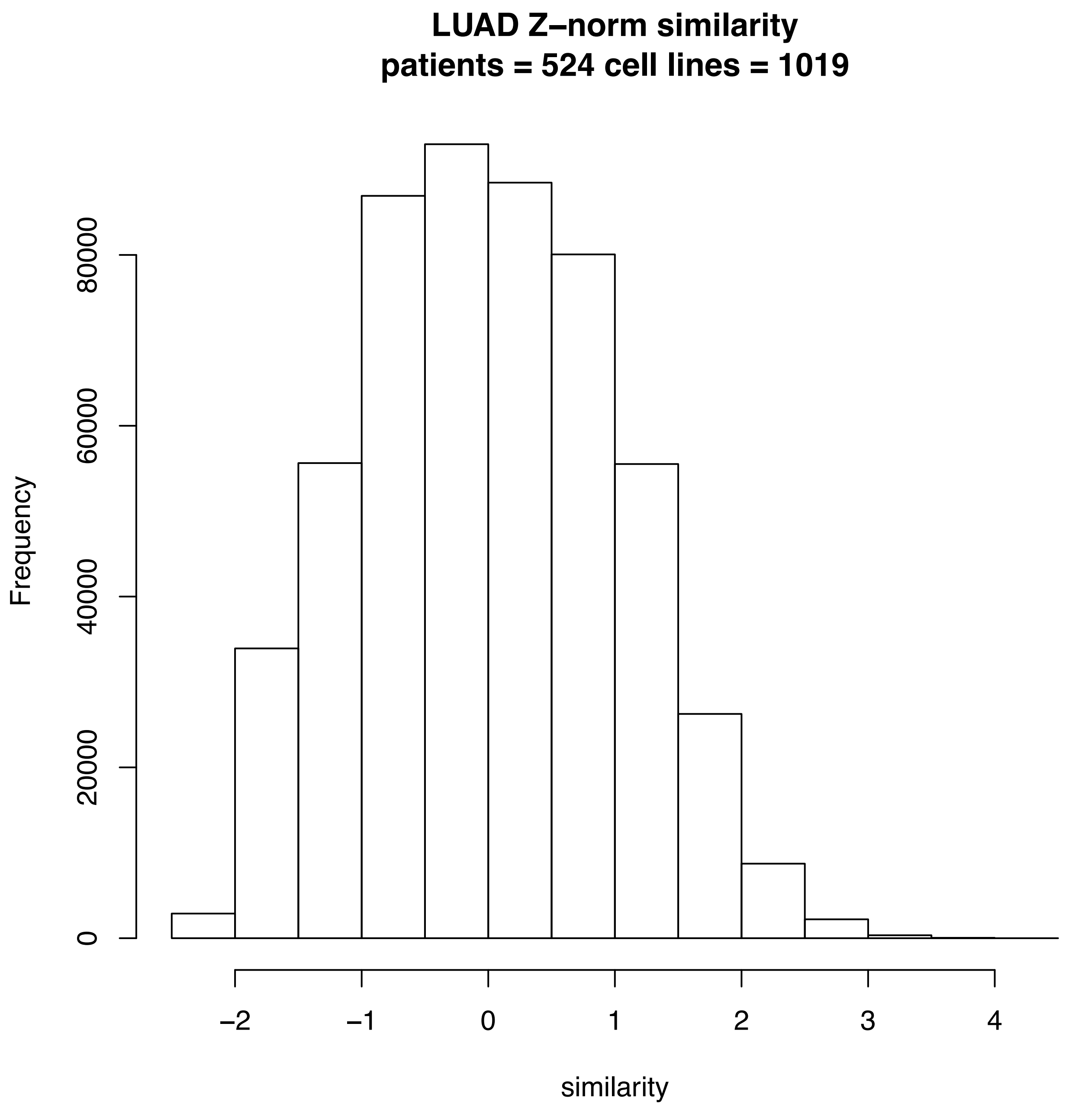
LUAD Z–norm similarity
patients = 524 cell lines = 1019
Frequency
0
20000
40000
60000
80000
−2
−1
0
1
2
3
4
similarity

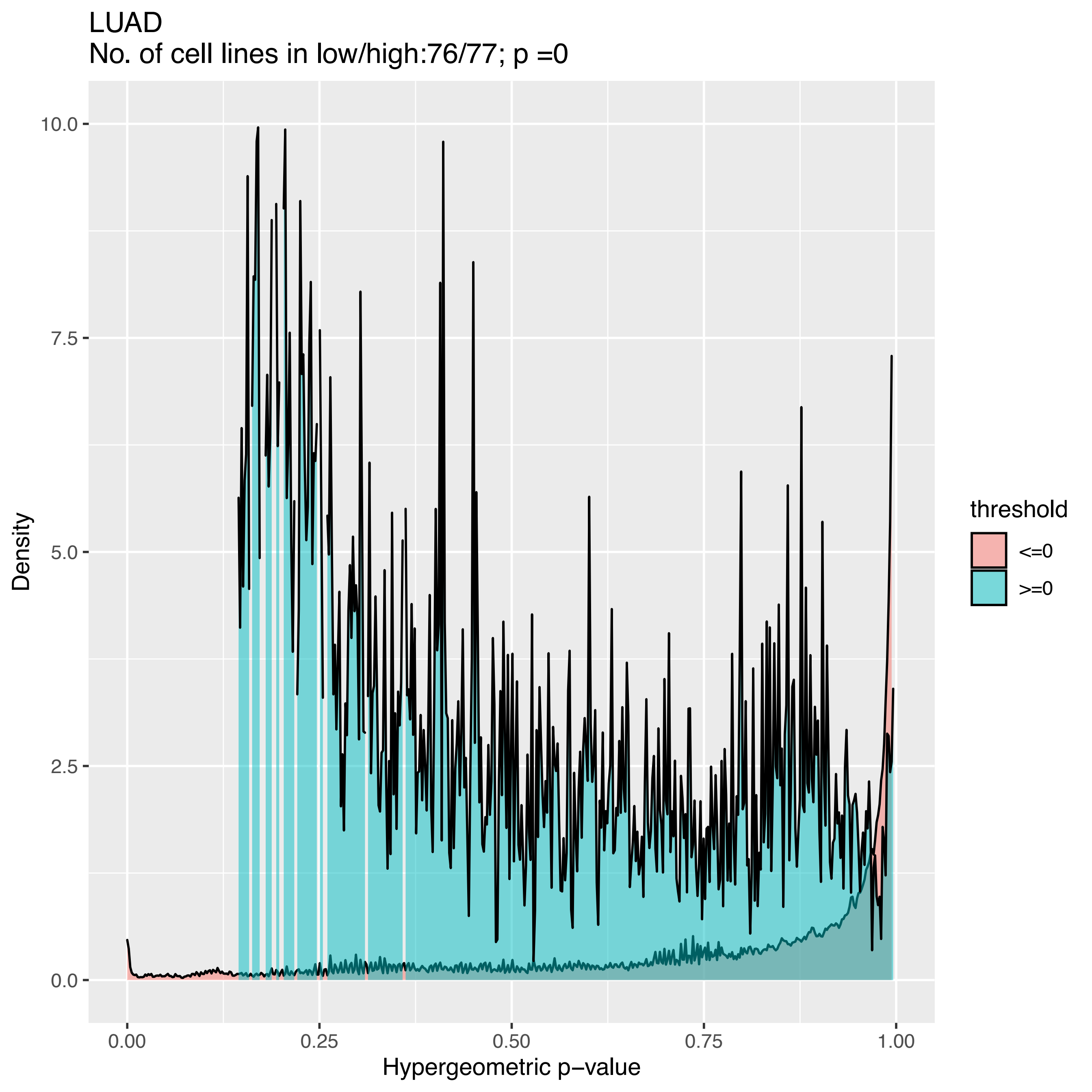

LUAD
No. of cell lines in low/high:76/77; p =0
Density
Hypergeometric p−value
0.0
2.5
5.0
7.5
10.0
0.00
0.25
0.50
0.75
1.00
threshold
<=0
>=0

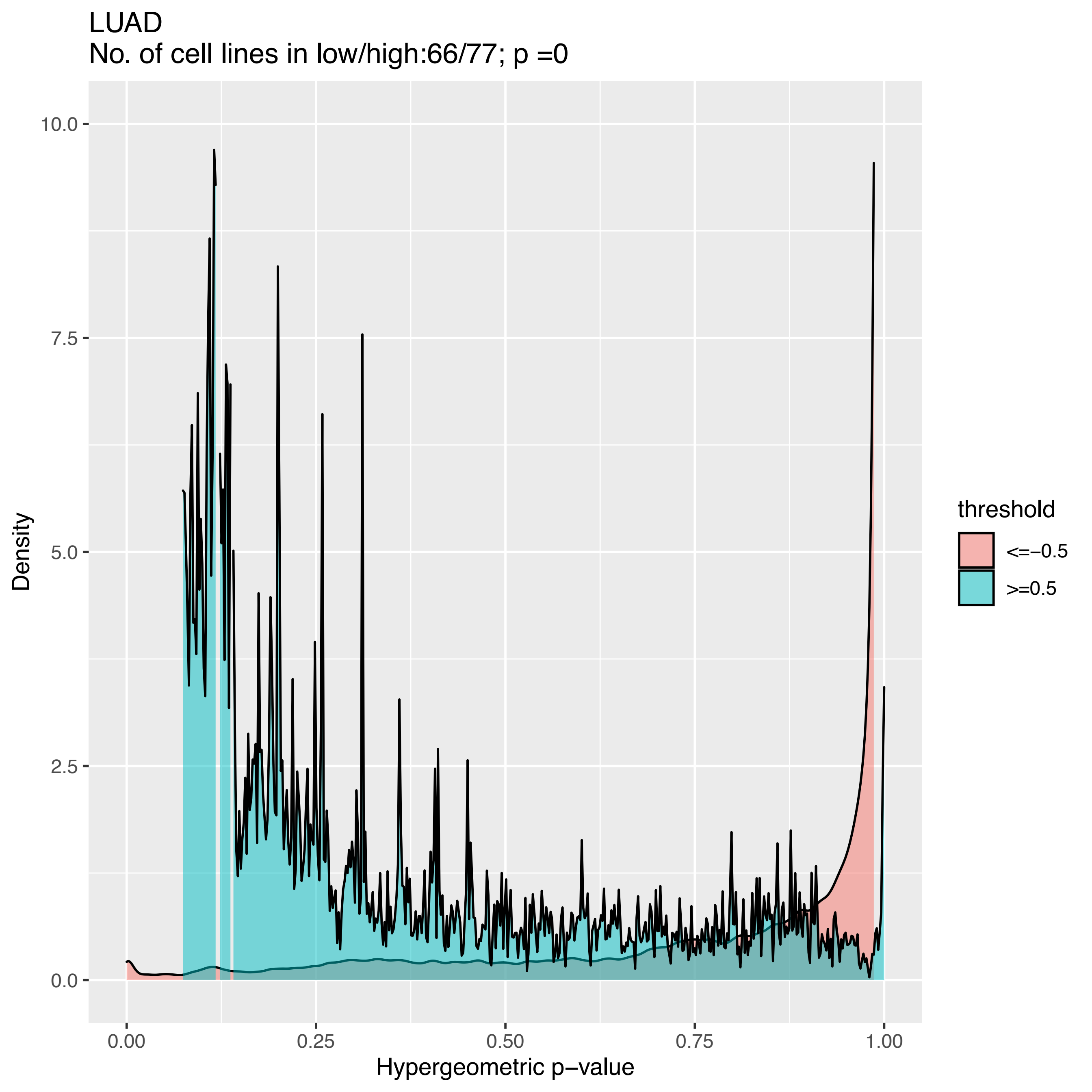

LUAD
No. of cell lines in low/high:66/77; p =0
Density
10.0
7.5
5.0
2.5
0.0
0.00
0.25
0.50
0.75
1.00
Hypergeometric p−value
threshold
<=−0.5
>=0.5

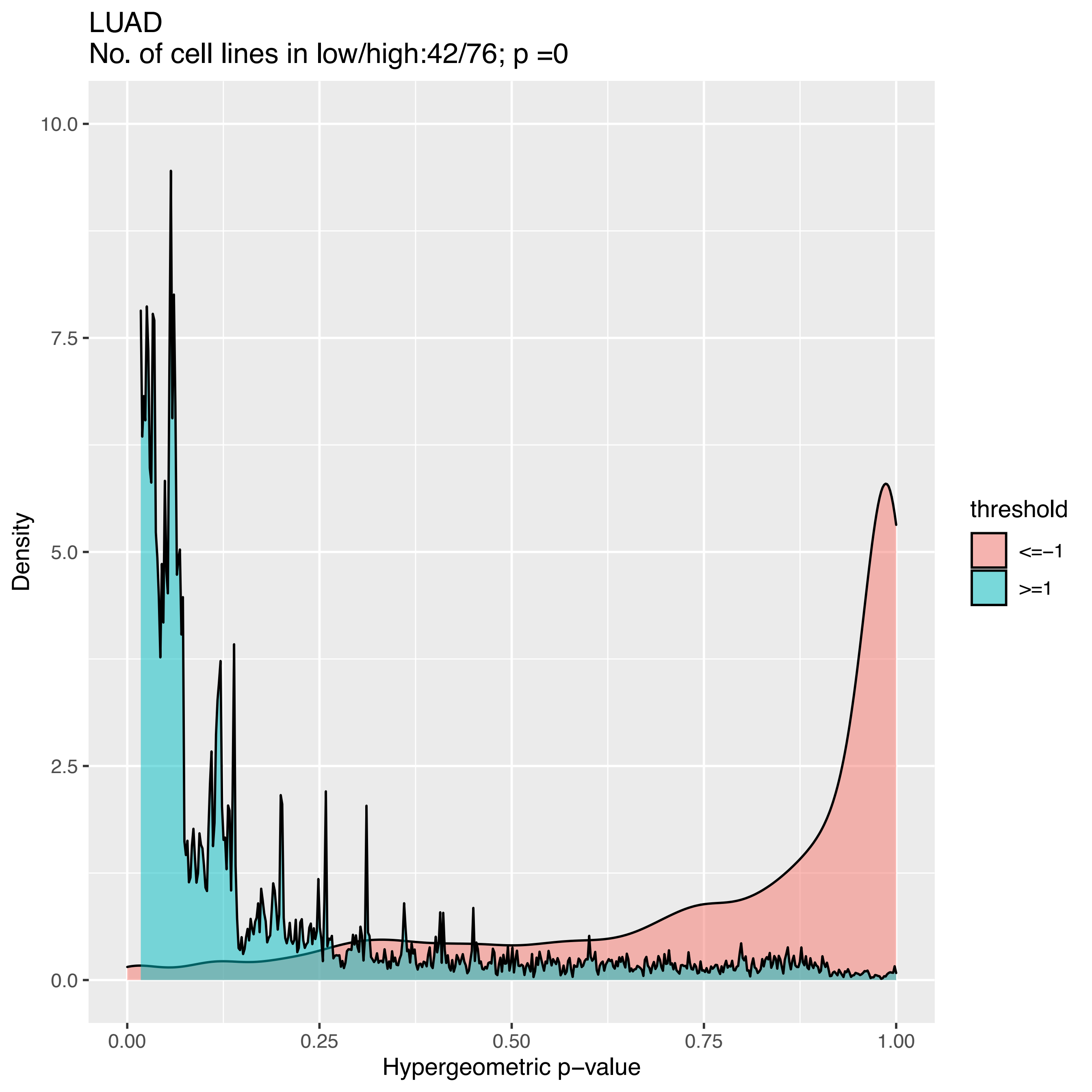

LUAD
No. of cell lines in low/high:42/76; p =0
Density
10.0
7.5
5.0
2.5
0.0
0.00
0.25
0.50
0.75
1.00
Hypergeometric p−value
threshold
<=−1
>=1

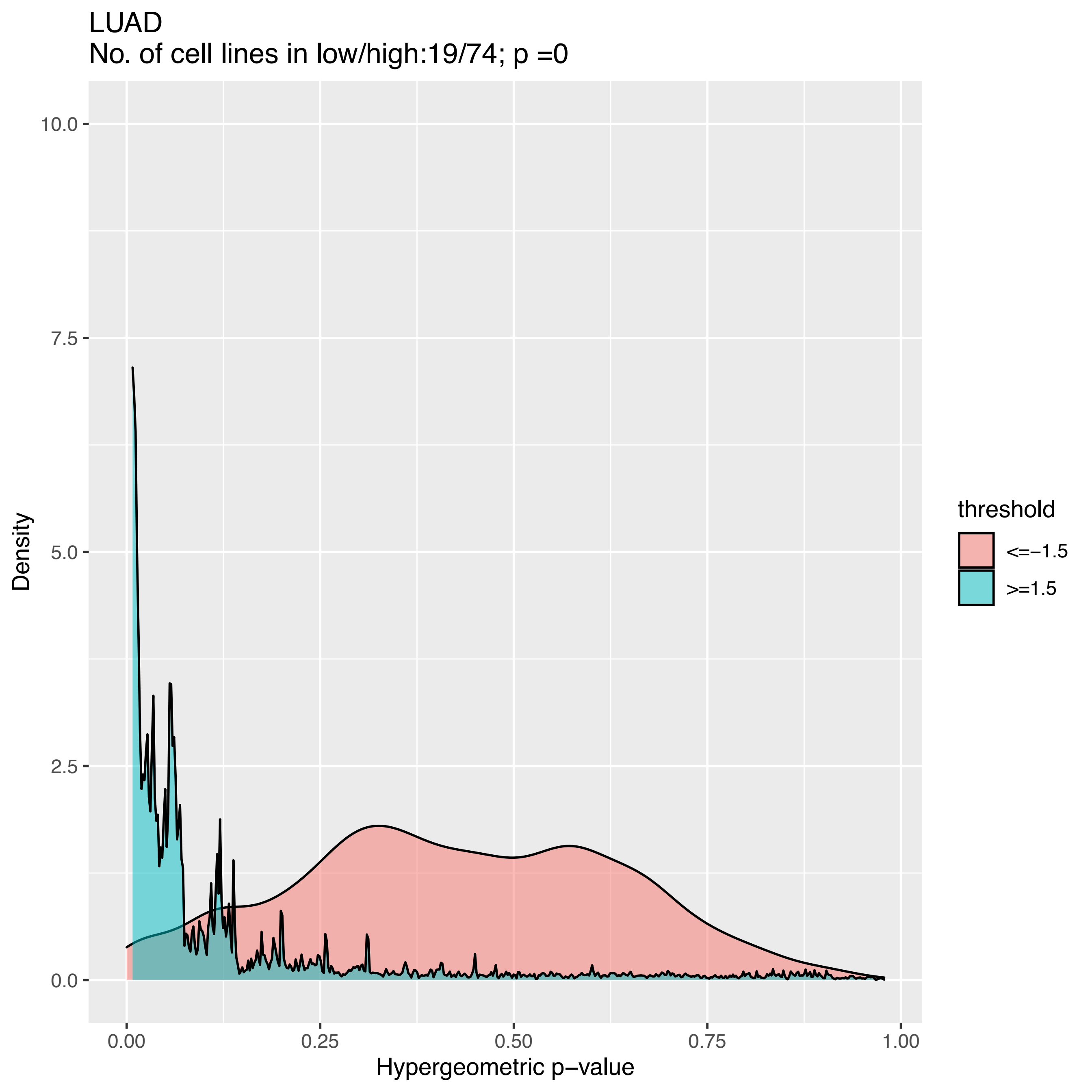
LUAD
No. of cell lines in low/high:19/74; p =0
Density
10.0
7.5
5.0
2.5
0.0
0.00
0.25
0.50
0.75
1.00
Hypergeometric p−value
threshold
<=−1.5
>=1.5

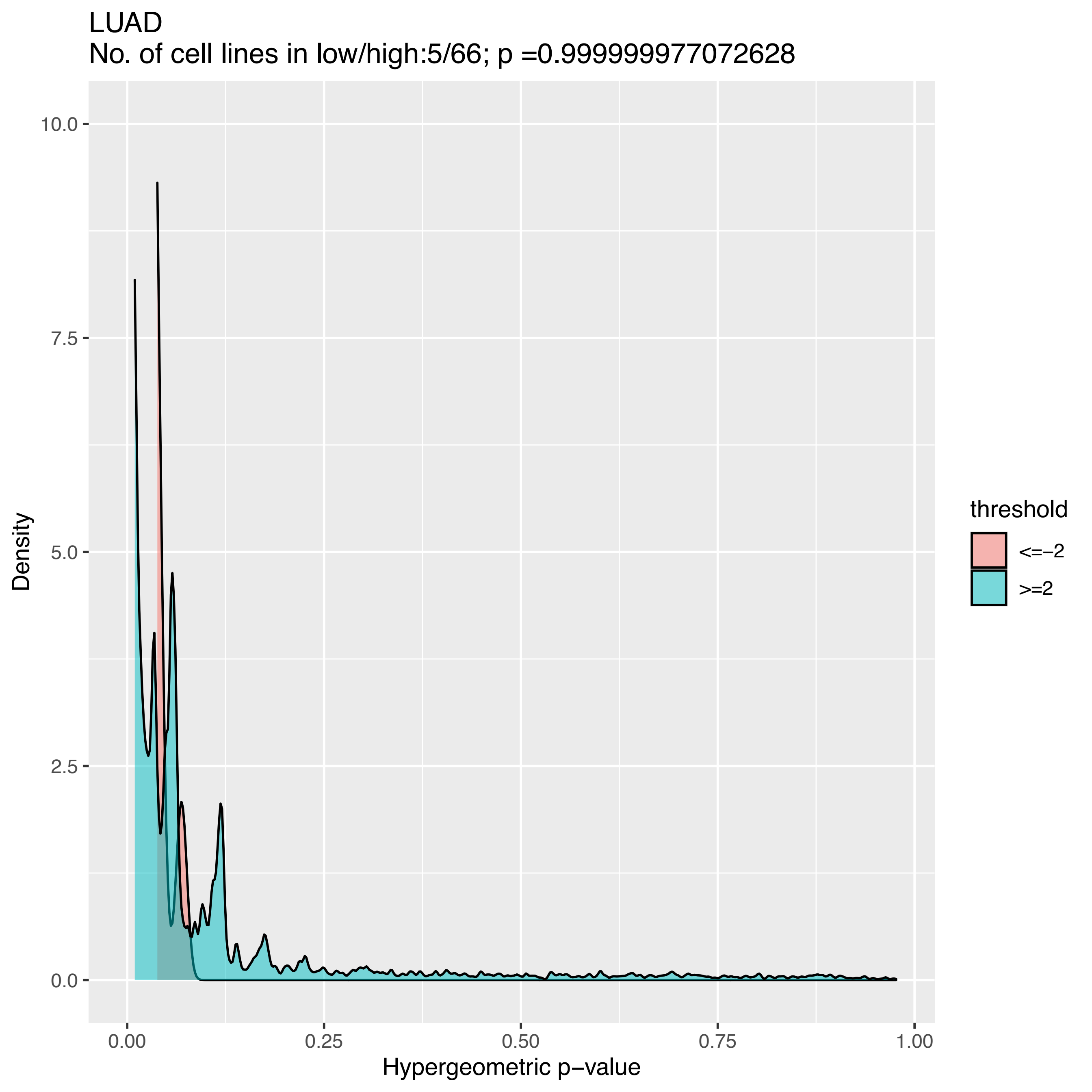
LUAD
No. of cell lines in low/high:5/66; p =0.999999977072628
10.0
7.5
5.0
2.5
0.0
Density
0.00
0.25
0.50
0.75
1.00
Hypergeometric p−value
threshold
<=−2
>=2

**LUSC Z–norm similarity**
**patients = 501 cell lines = 1019**

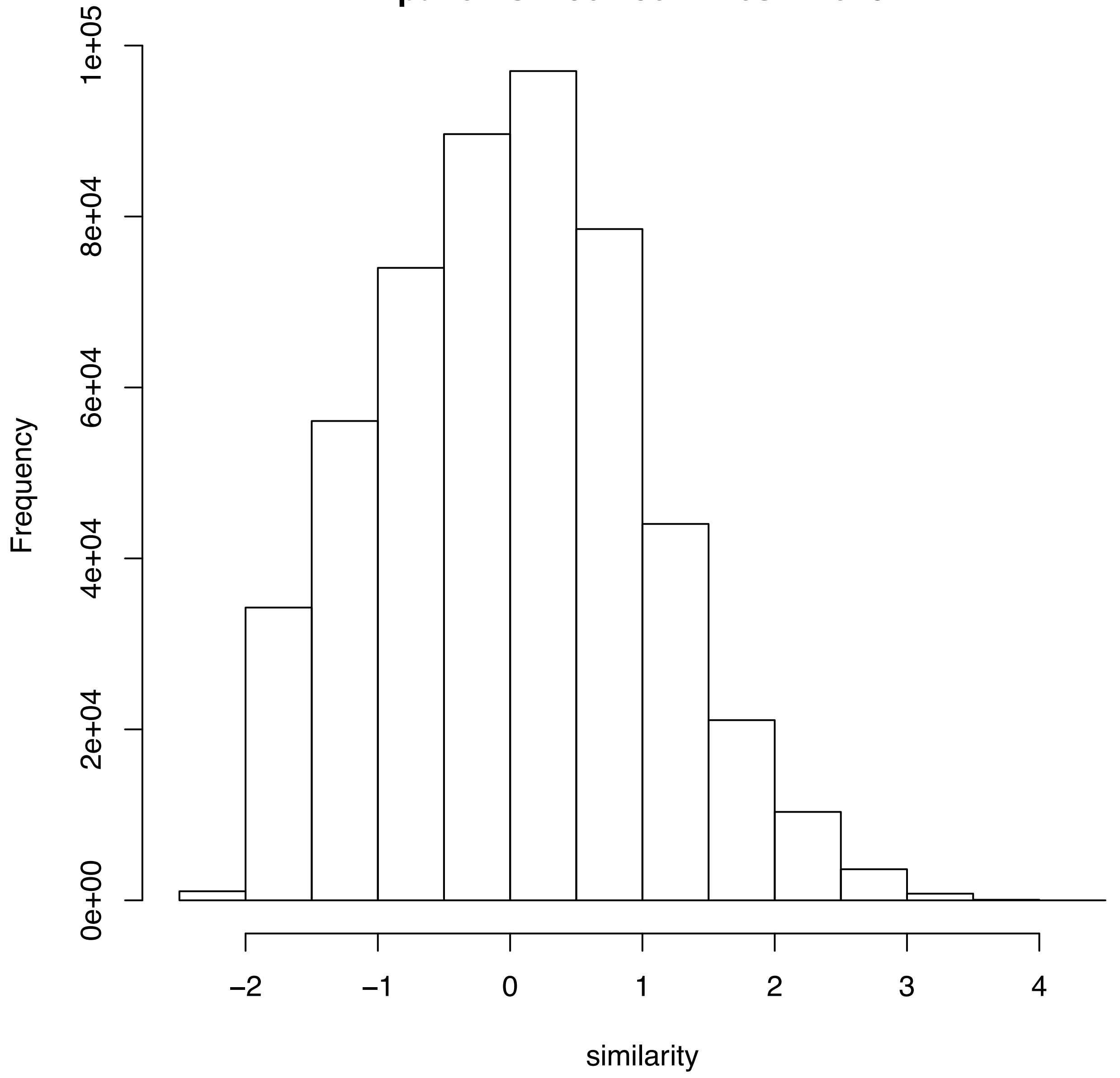

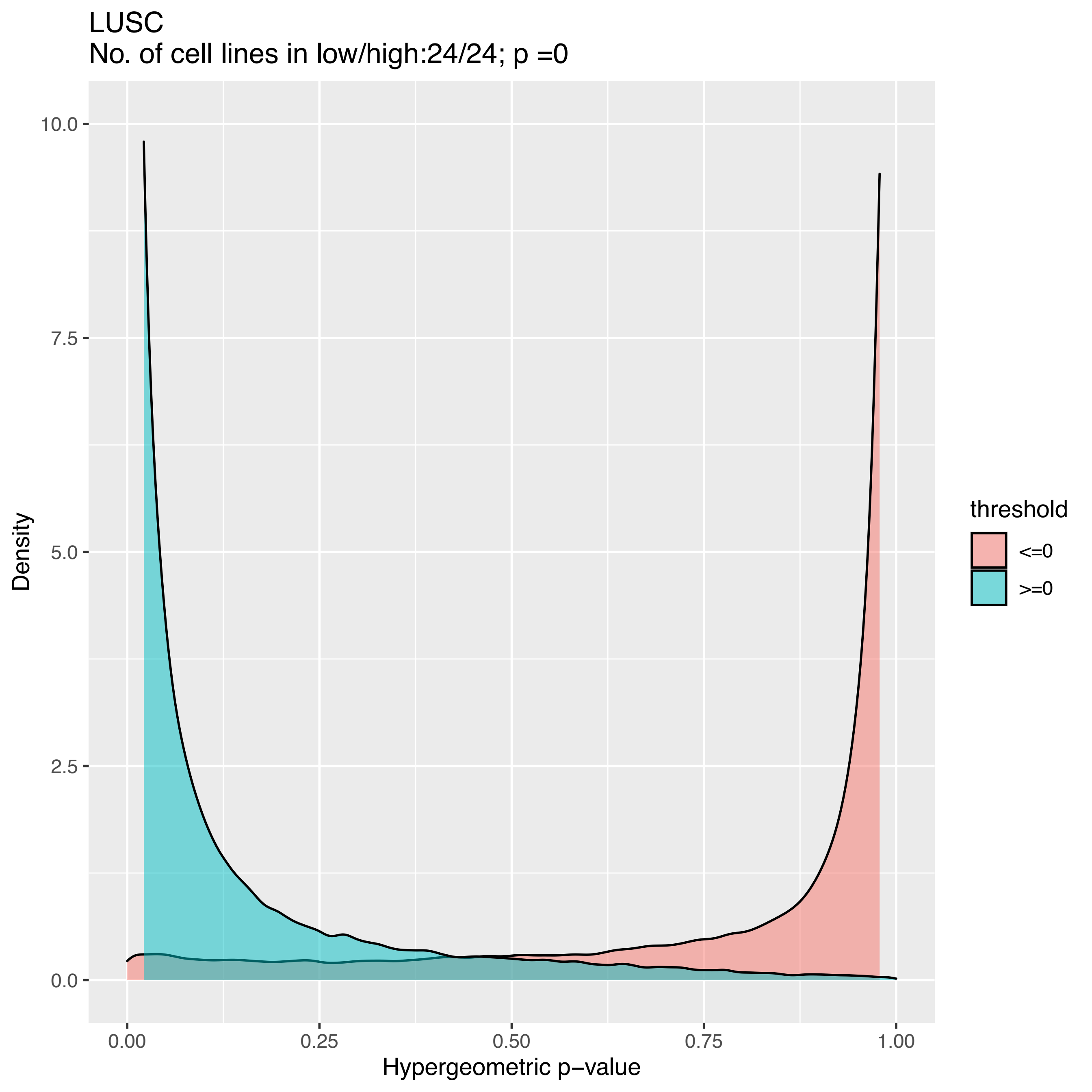

LUSC
No. of cell lines in low/high:24/24; p =0
10.0
7.5
5.0
2.5
0.0
Density
0.00
0.25
0.50
0.75
1.00
Hypergeometric p−value
threshold
<=0
>=0

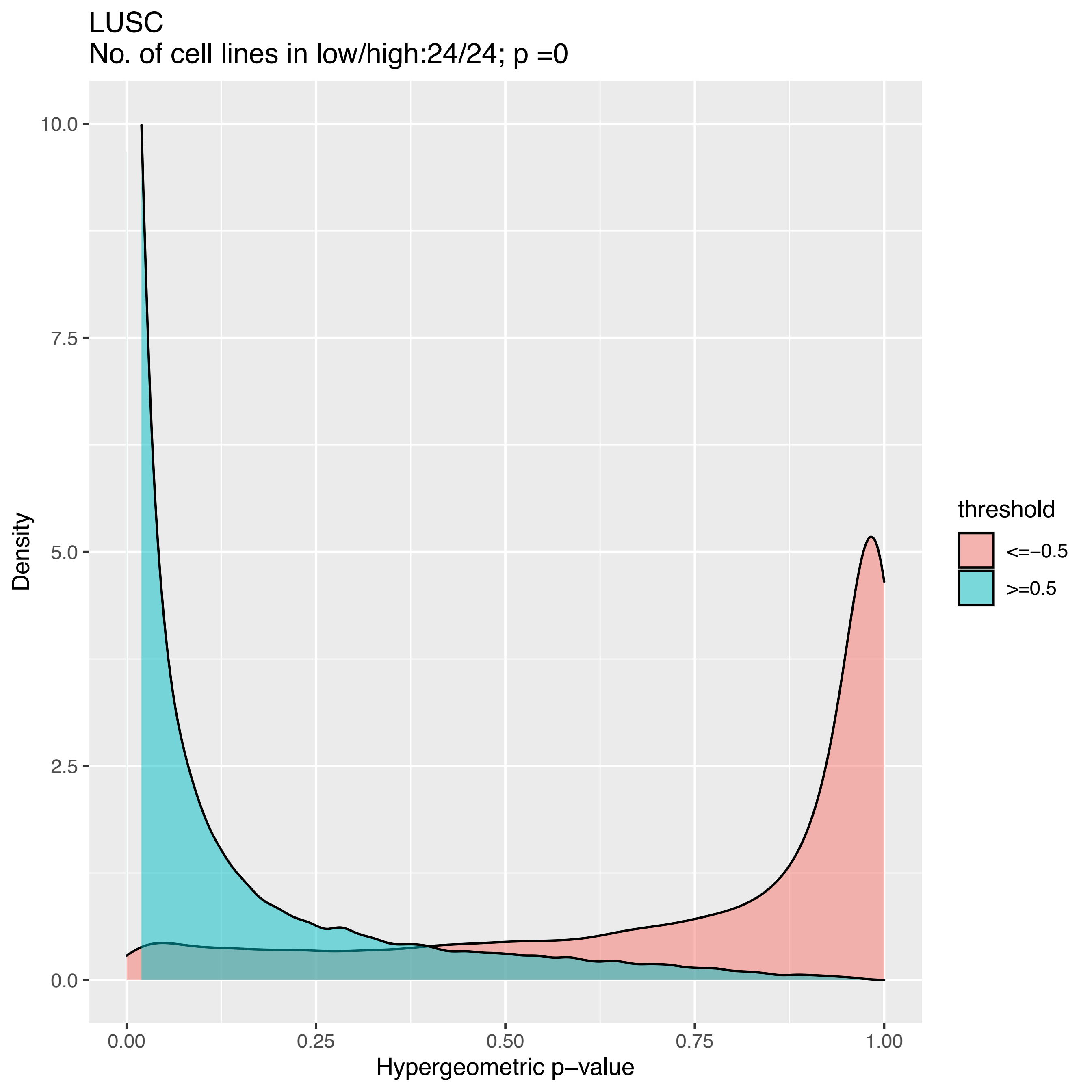

LUSC
No. of cell lines in low/high:24/24; p =0
Density
10.0
7.5
5.0
2.5
0.0
0.00
0.25
0.50
0.75
1.00
Hypergeometric p−value
threshold
<=−0.5
>=0.5

LUSC
No. of cell lines in low/high:21/23; p =0

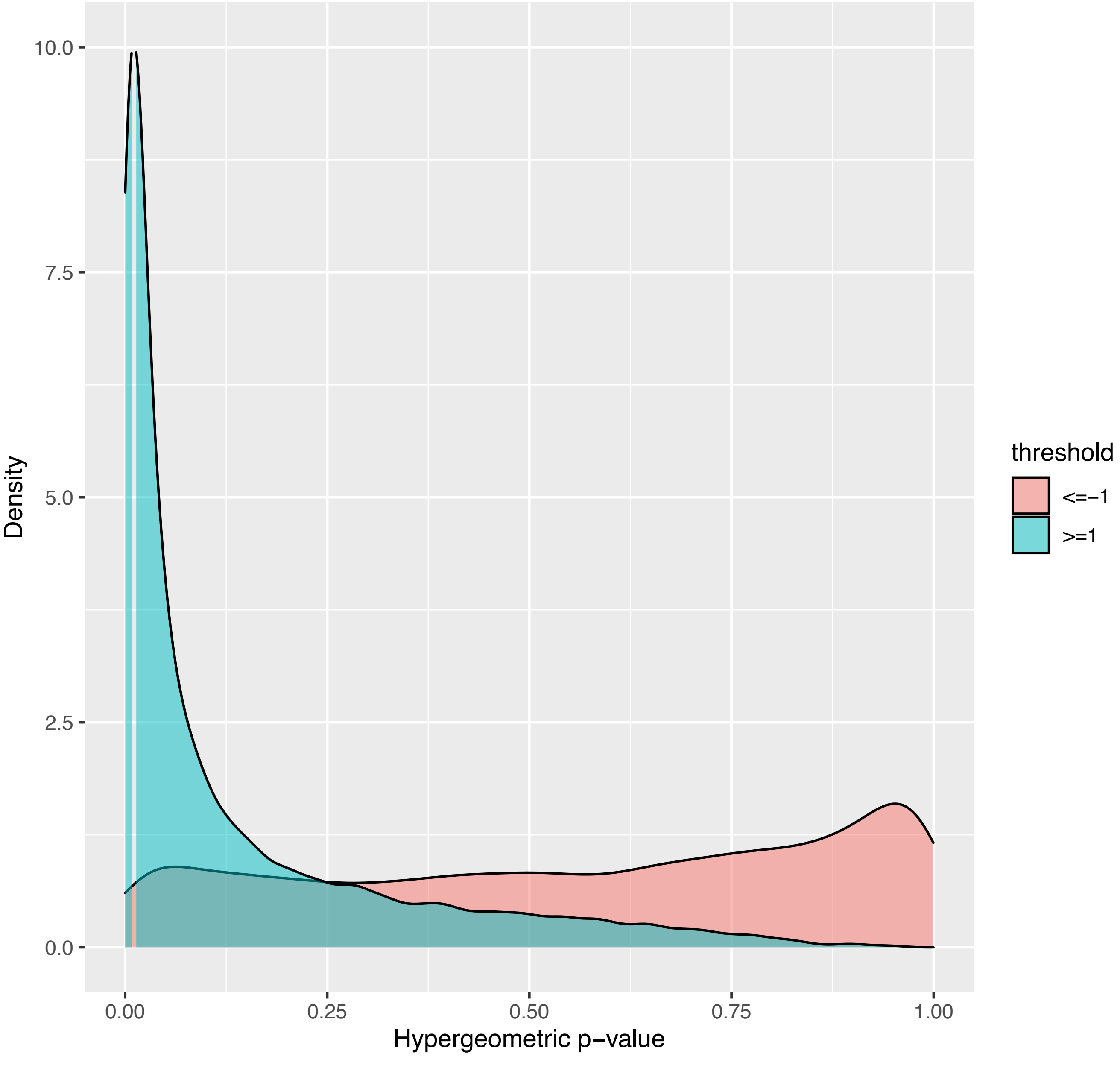

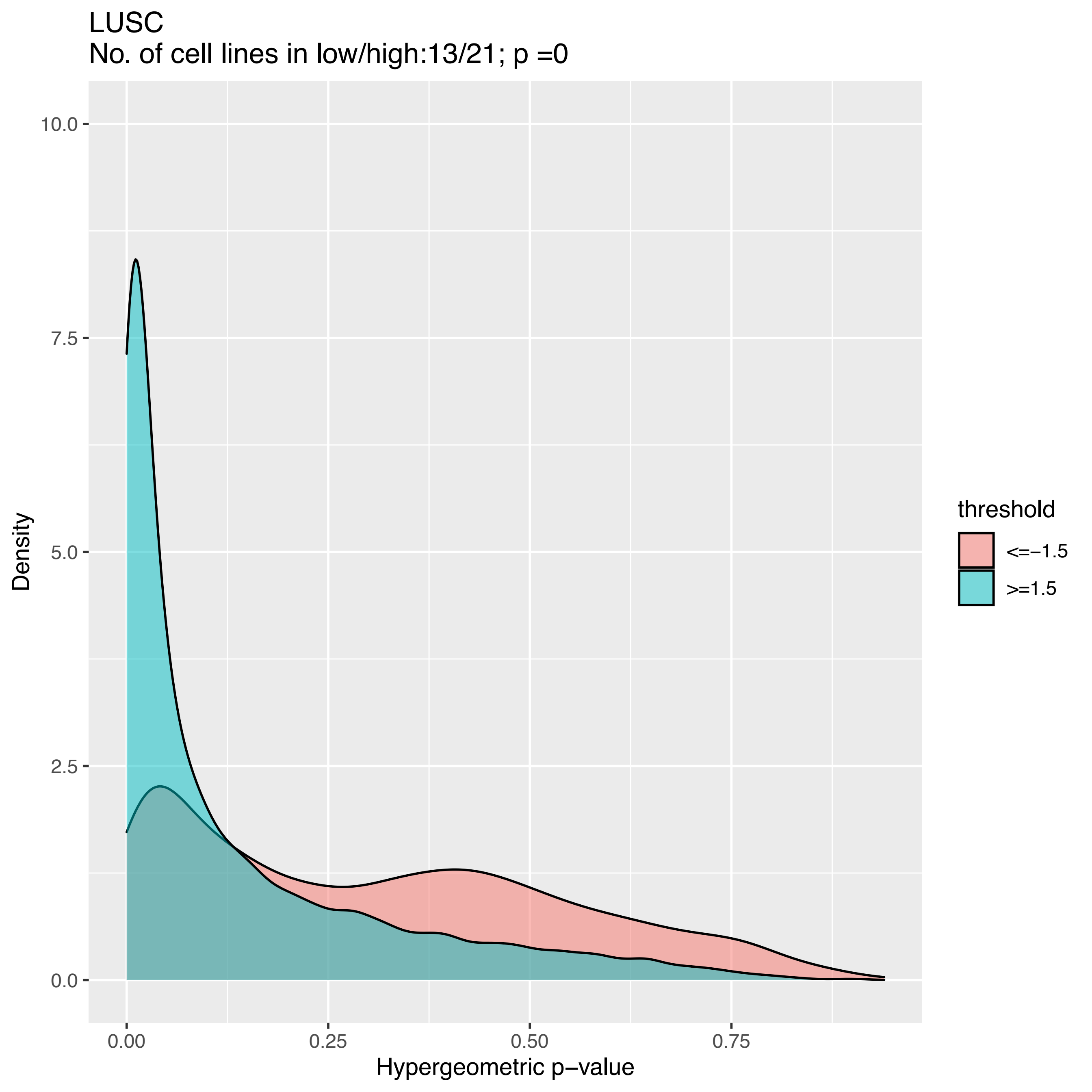
LUSC
No. of cell lines in low/high:13/21; p =0
Density
Hypergeometric p−value
threshold
<=−1.5
>=1.5
10.0
7.5
5.0
2.5
0.0
0.00
0.25
0.50
0.75

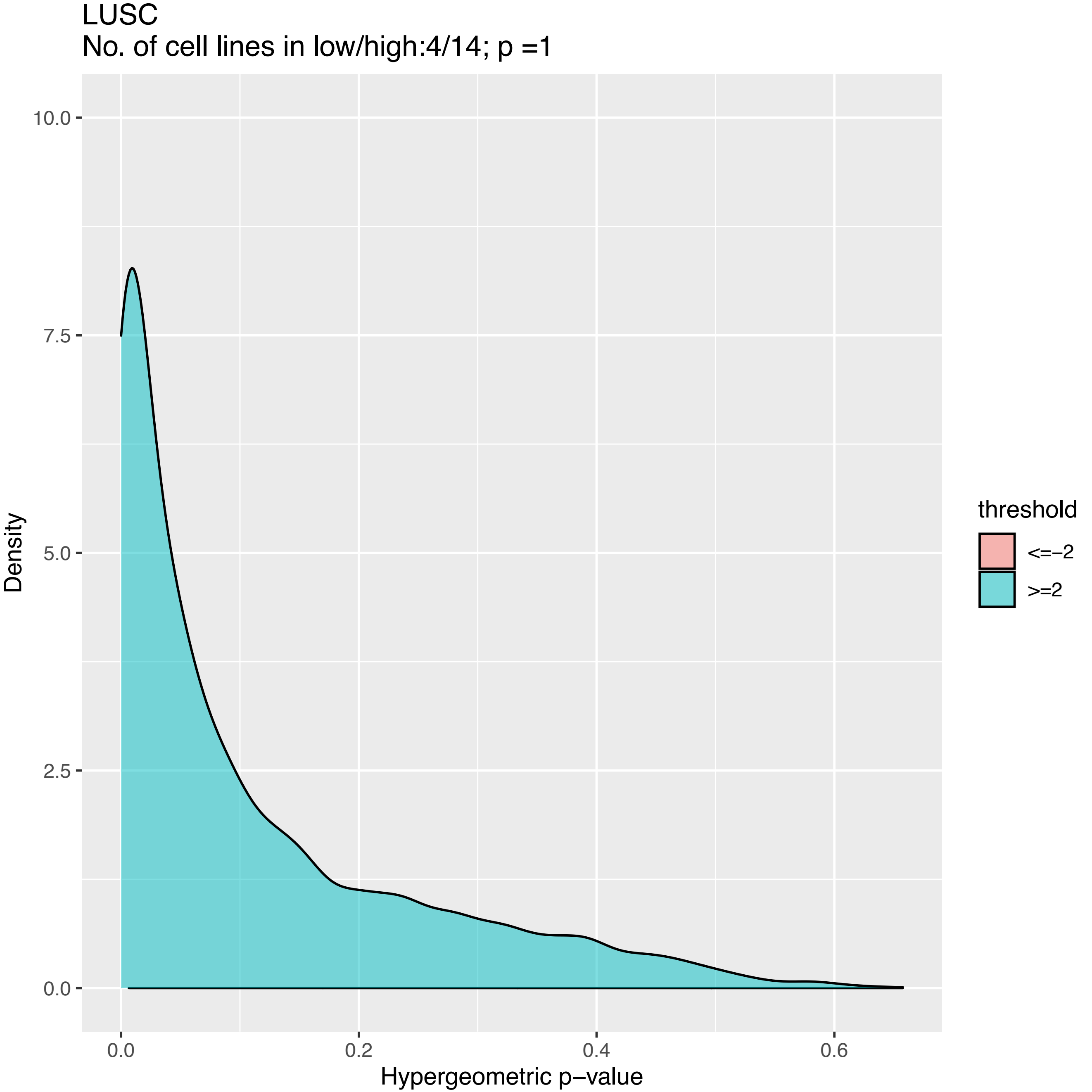
LUSC
No. of cell lines in low/high:4/14; p =1
10.0
7.5
5.0
2.5
0.0
Density
0.0
0.2
0.4
0.6
Hypergeometric p−value
threshold
<=−2
>=2

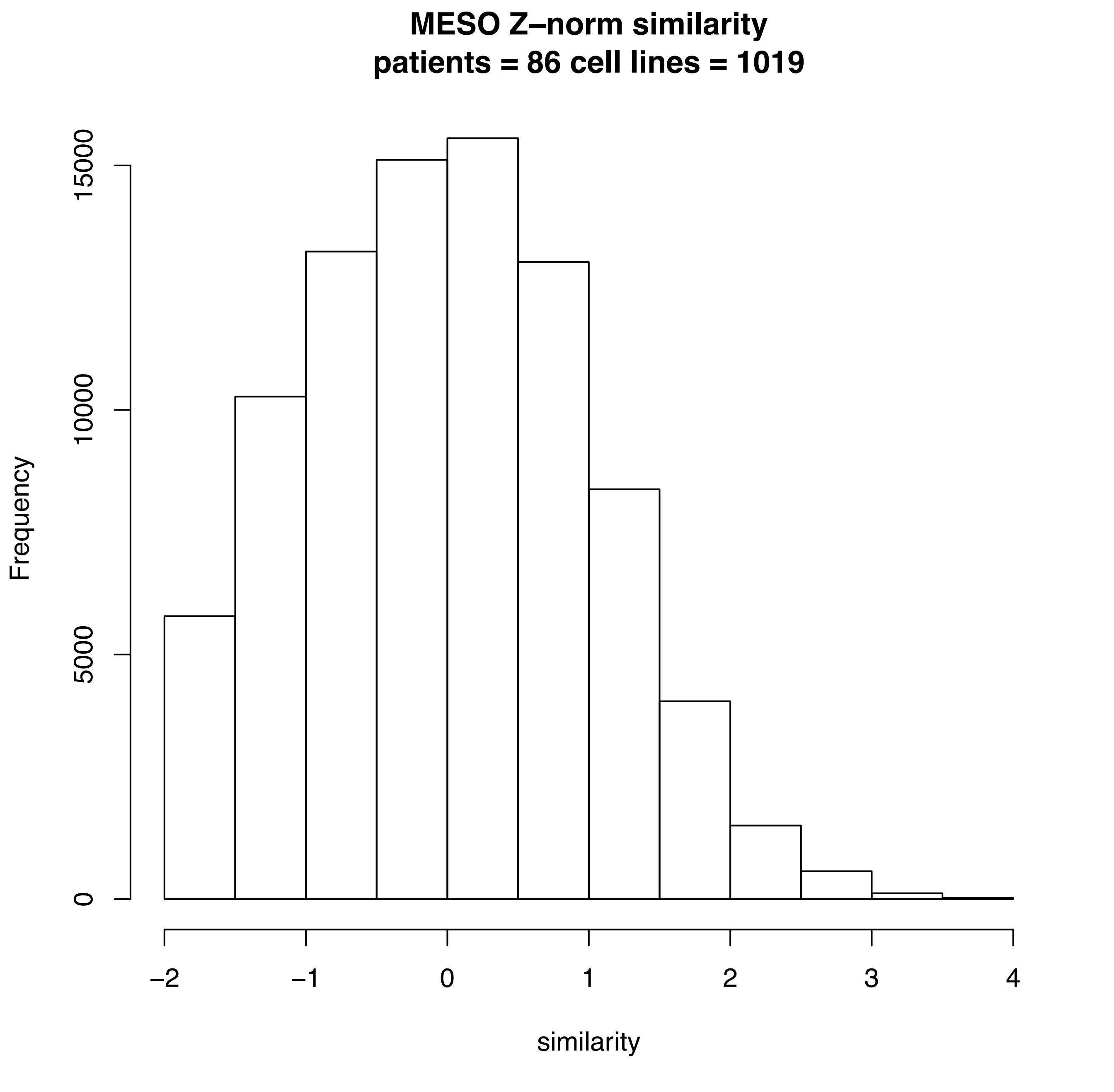
MESO Z–norm similarity
patients = 86 cell lines = 1019
Frequency
0
5000
10000
15000
−2
−1
0
1
2
3
4
similarity

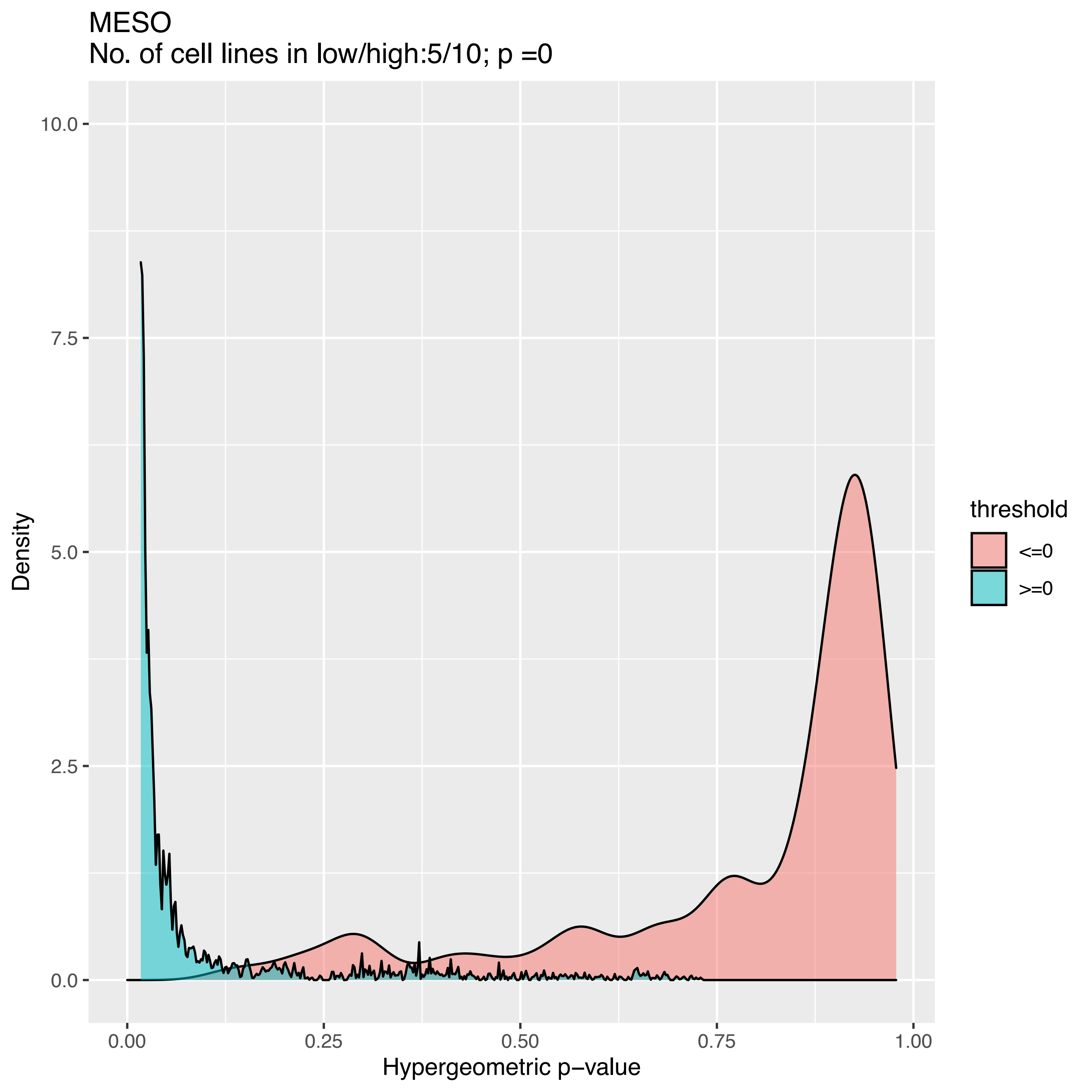

MESO
No. of cell lines in low/high:5/10; p =0
10.0
7.5
5.0
2.5
0.0
Density
0.00
0.25
0.50
0.75
1.00
Hypergeometric p−value
threshold
<=0
>=0

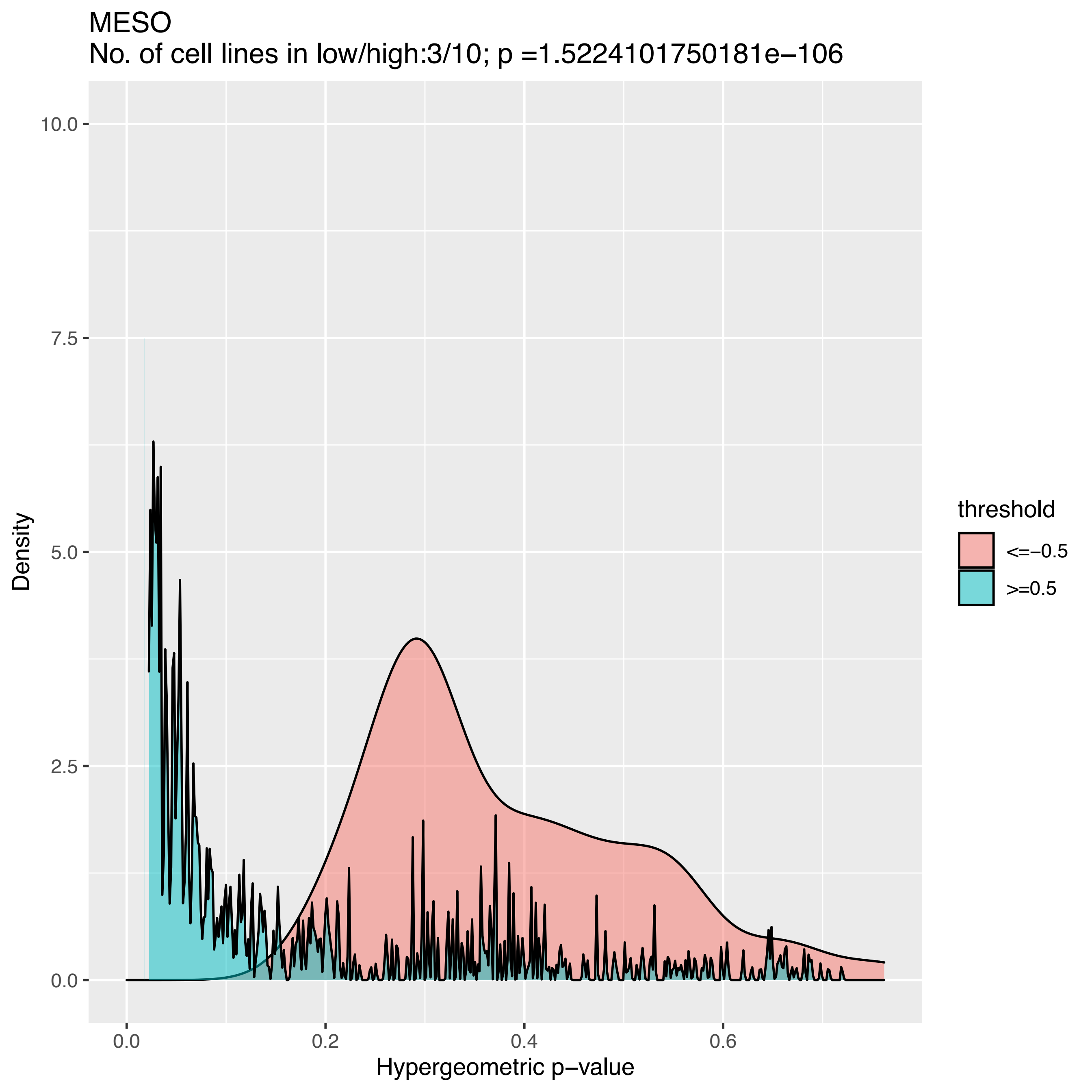
MESO
No. of cell lines in low/high:3/10; p =1.5224101750181e−106
Density
Hypergeometric p−value
0.0
2.5
5.0
7.5
10.0
0.0
0.2
0.4
0.6
threshold
<=−0.5
>=0.5

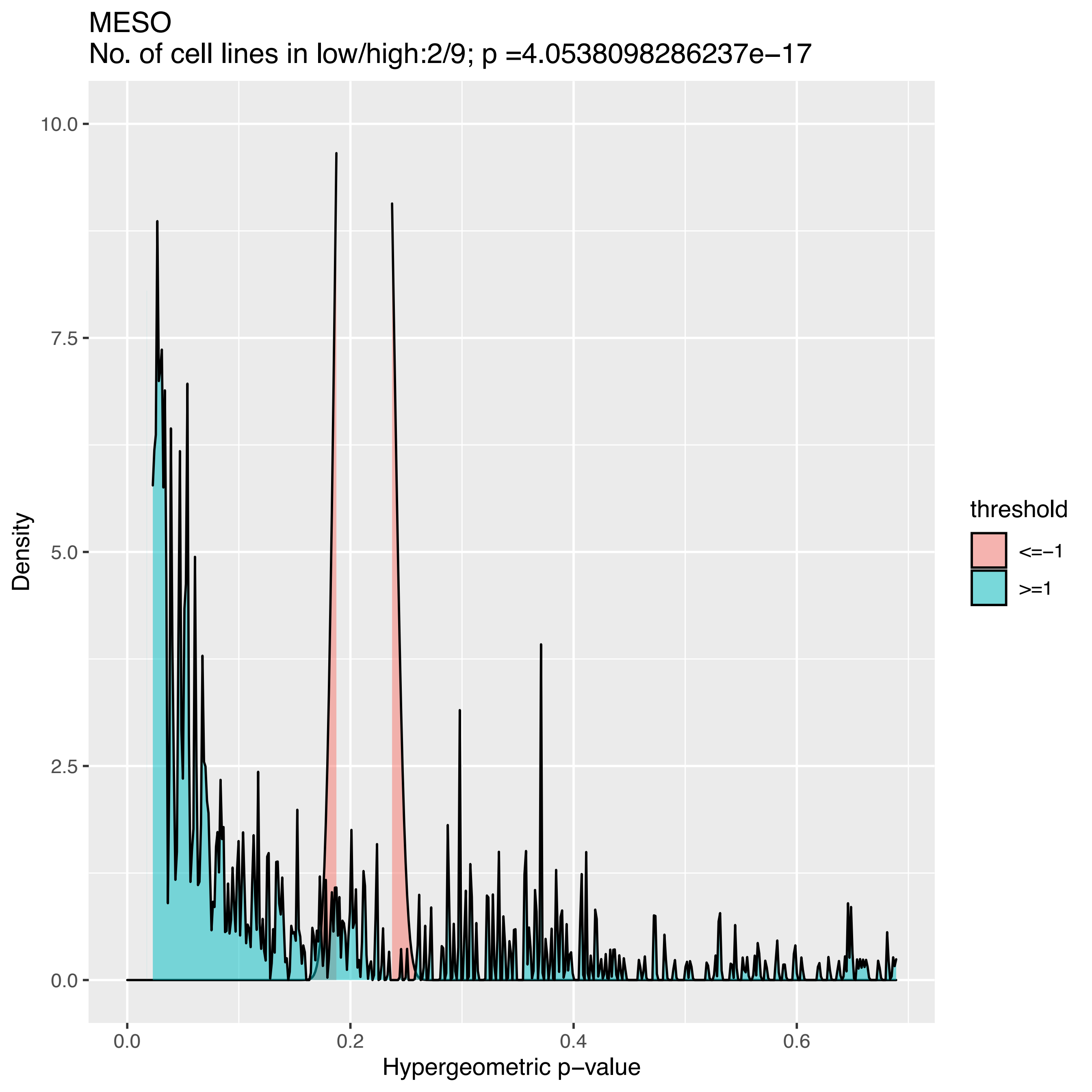
MESO
No. of cell lines in low/high:2/9; p =4.0538098286237e−17
10.0
7.5
5.0
2.5
0.0
Density
0.0
0.2
0.4
0.6
Hypergeometric p−value
threshold
<=−1
>=1

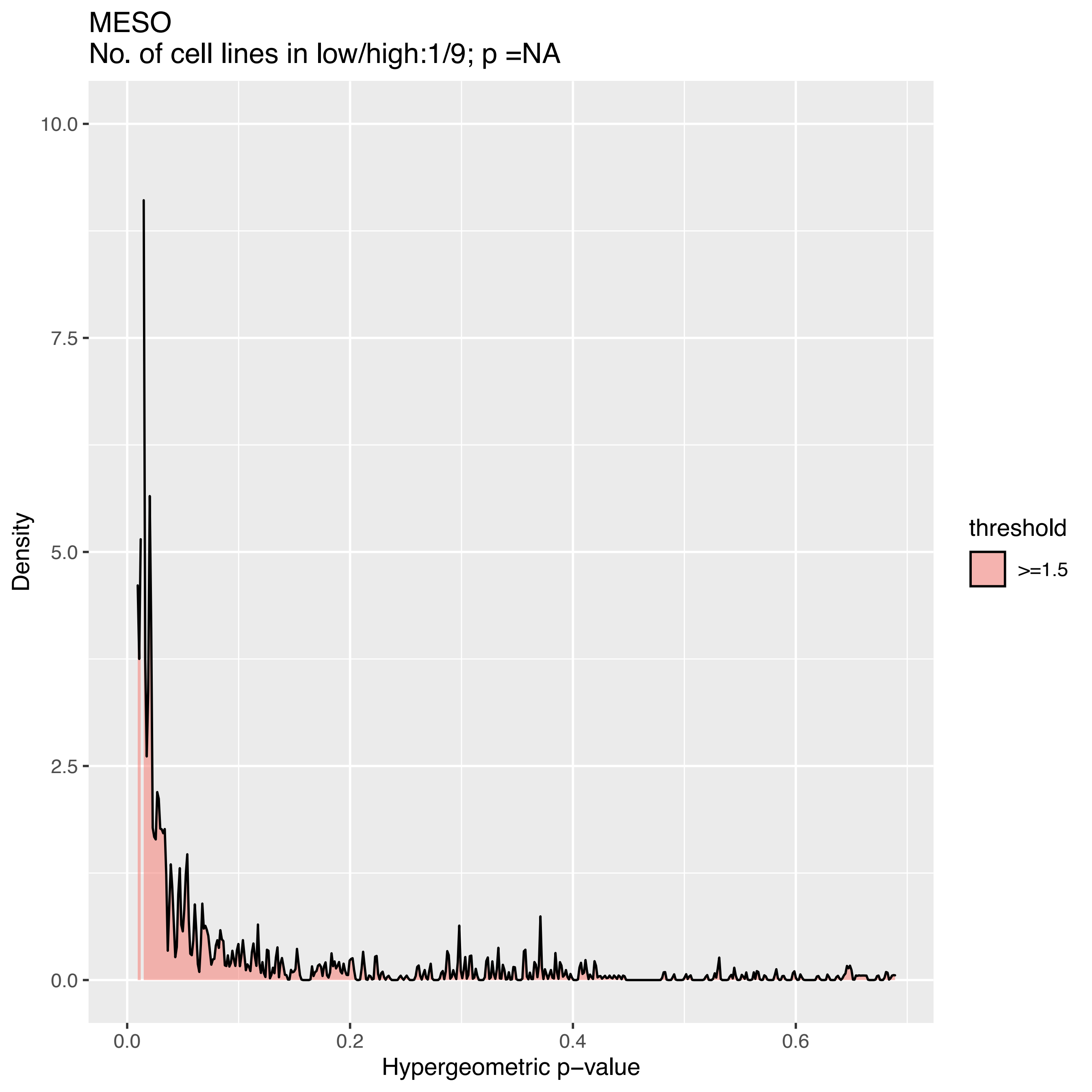
MESO
No. of cell lines in low/high:1/9; p =NA
10.0
7.5
5.0
2.5
0.0
Density
0.0
0.2
0.4
0.6
Hypergeometric p−value
threshold
>=1.5

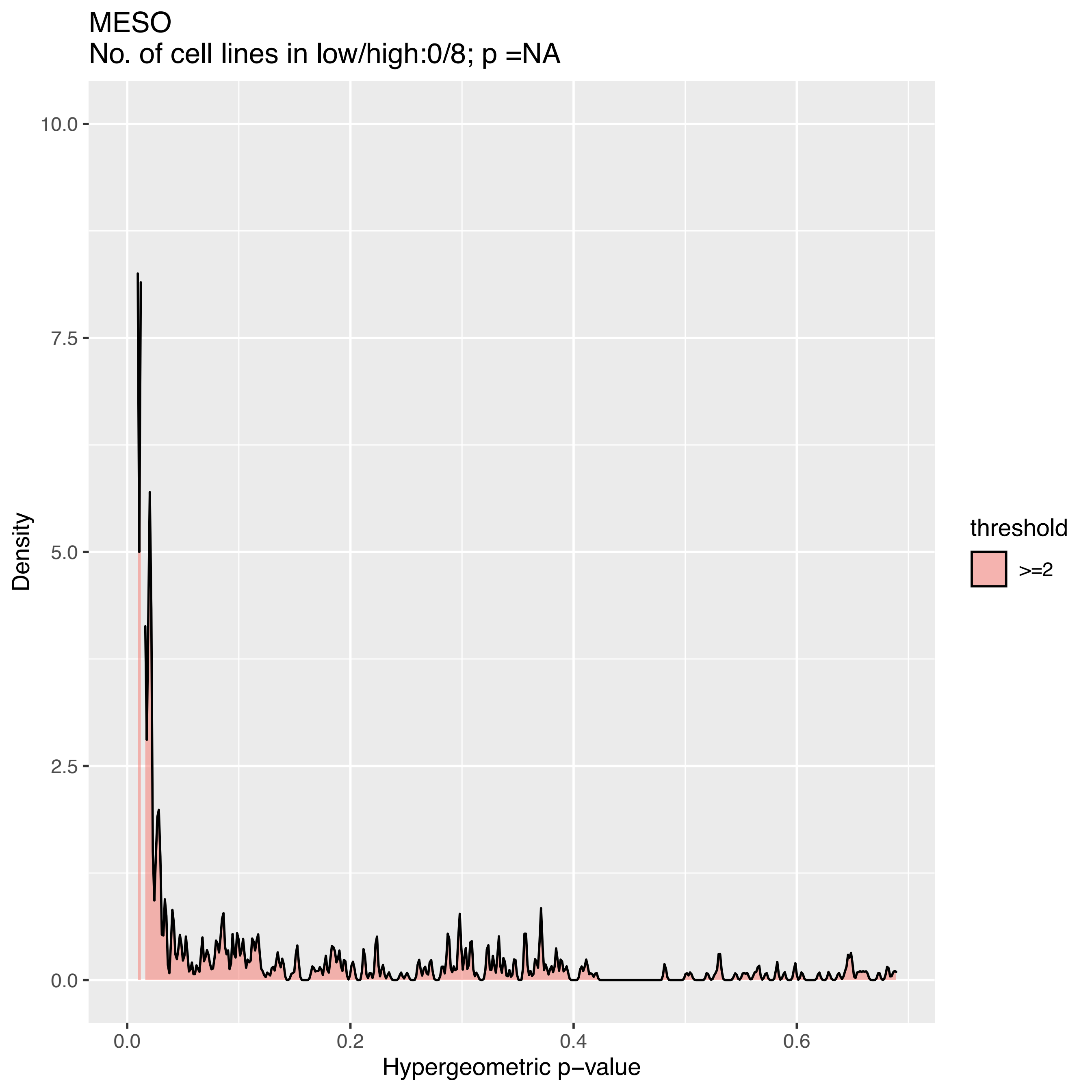
MESO
No. of cell lines in low/high:0/8; p =NA
Density
10.0
7.5
5.0
2.5
0.0
0.0
0.2
0.4
0.6
Hypergeometric p−value
threshold
>=2

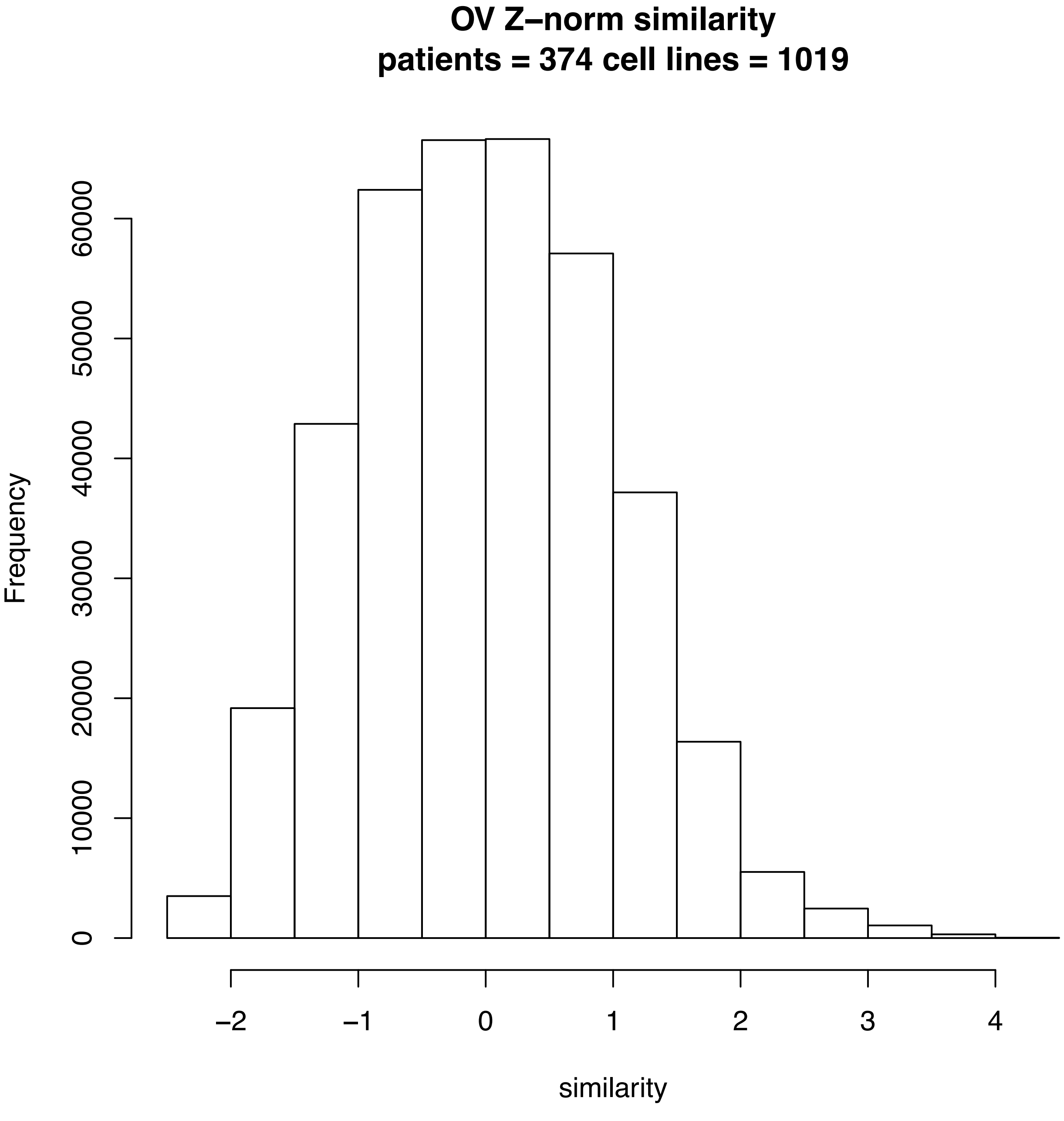

OV Z−norm similarity
patients = 374 cell lines = 1019
Frequency
0
10000
20000
30000
40000
50000
60000
−2
−1
0
1
2
3
4
similarity

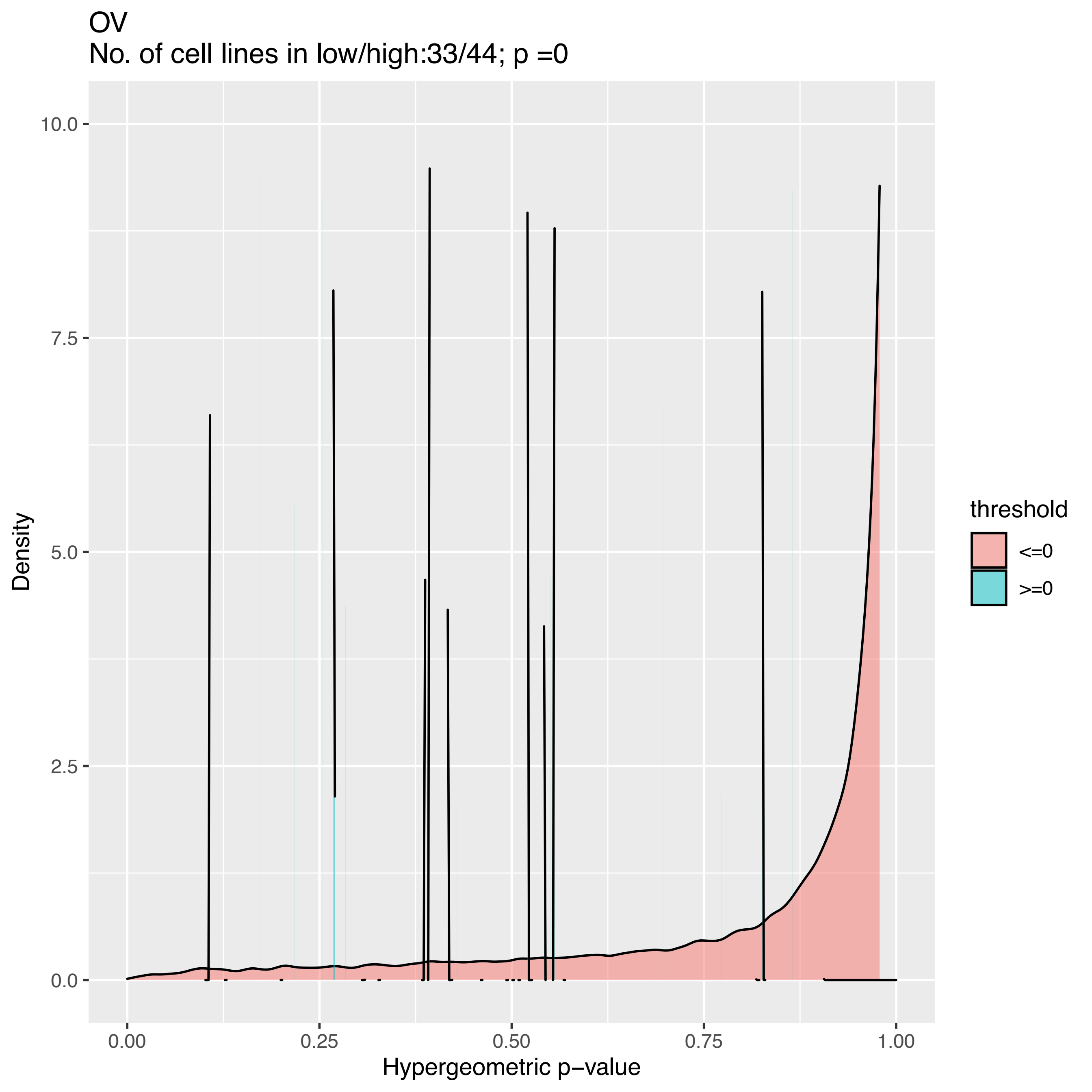
OV
No. of cell lines in low/high:33/44; p =0
Density
Hypergeometric p−value
0.0
2.5
5.0
7.5
10.0
0.00
0.25
0.50
0.75
1.00
threshold
<=0
>=0

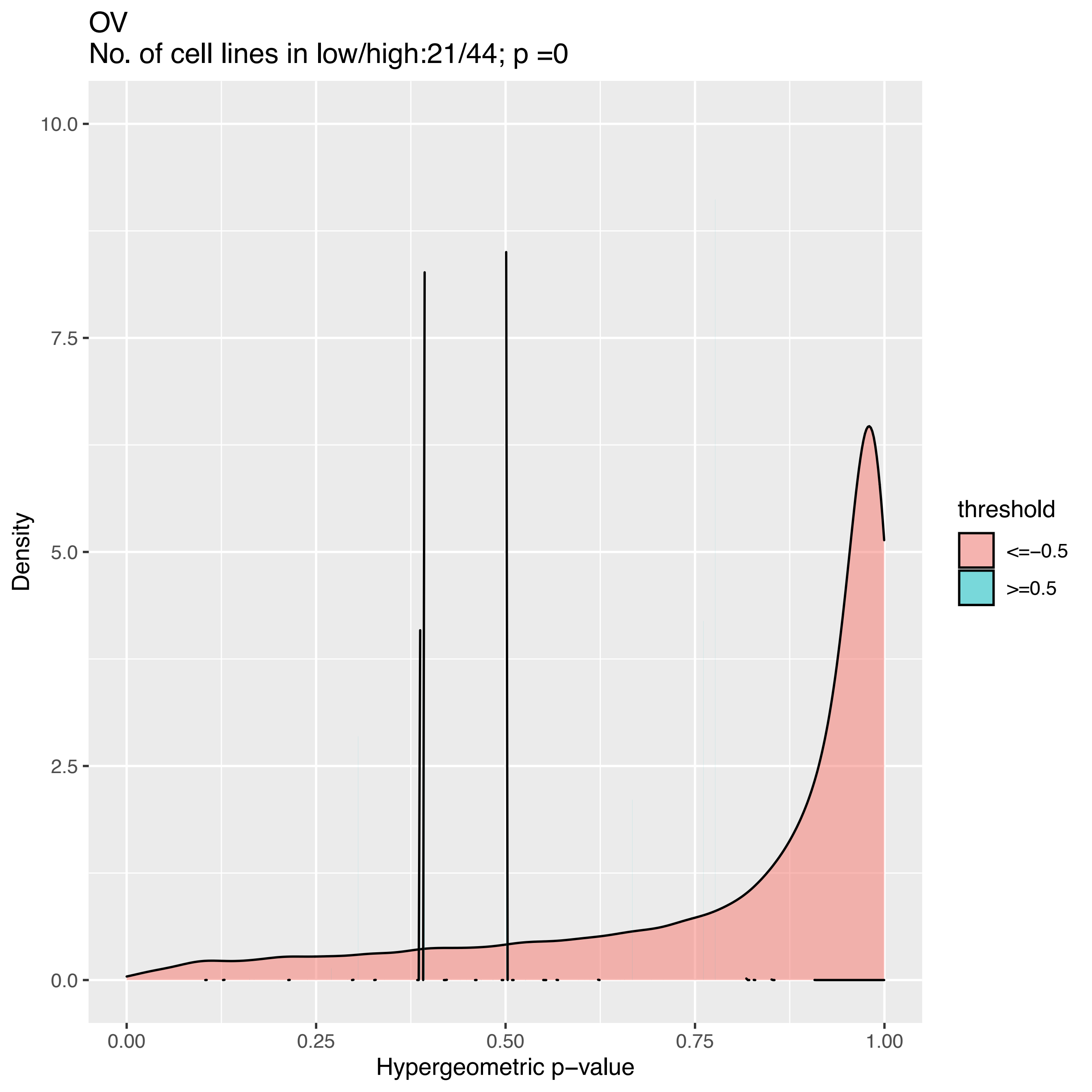
OV
No. of cell lines in low/high:21/44; p =0
Density
10.0
7.5
5.0
2.5
0.0
0.00
0.25
0.50
0.75
1.00
Hypergeometric p−value
threshold
<=−0.5
>=0.5

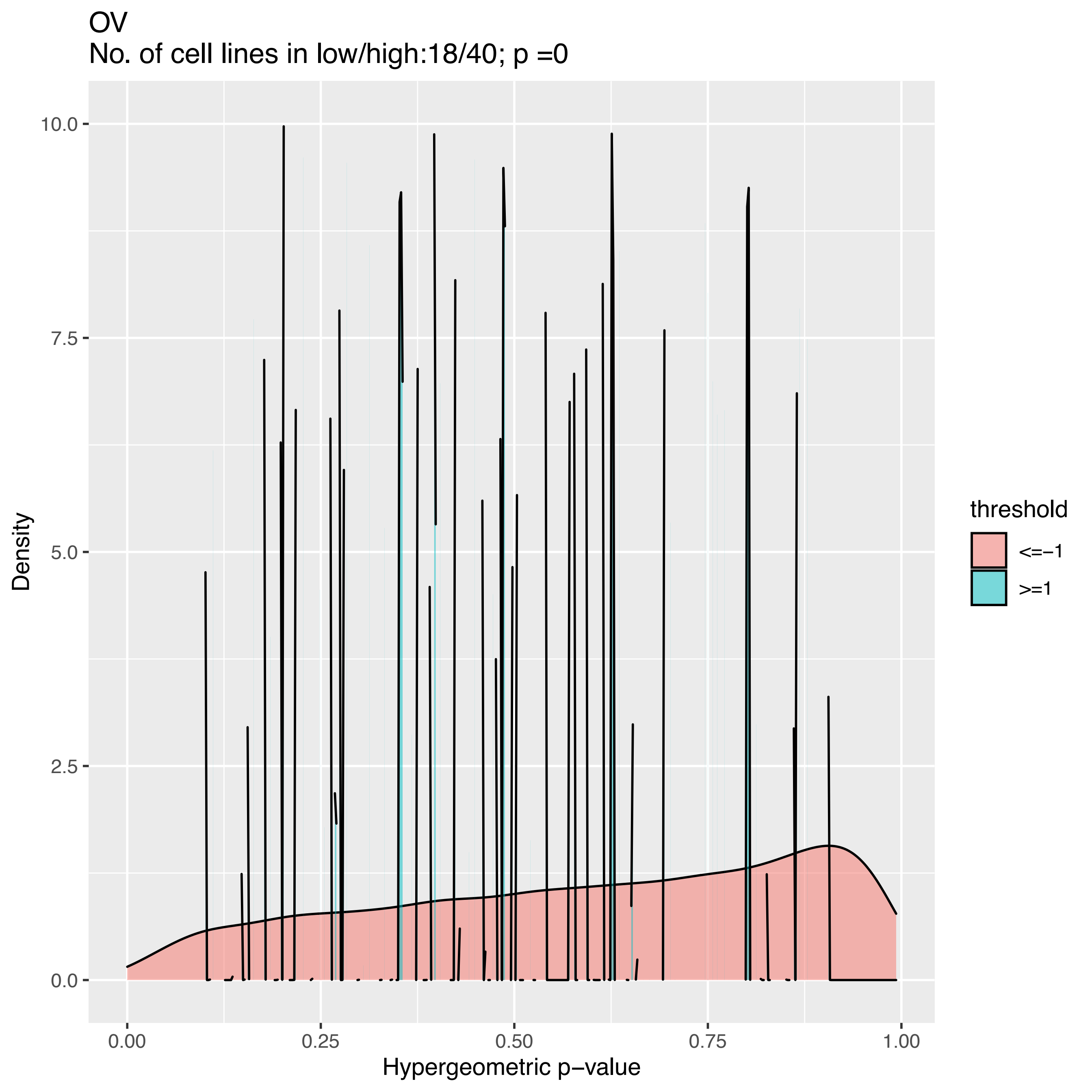
OV
No. of cell lines in low/high:18/40; p =0
10.0
7.5
5.0
2.5
0.0
Density
0.00
0.25
0.50
0.75
1.00
Hypergeometric p−value
threshold
<=−1
>=1

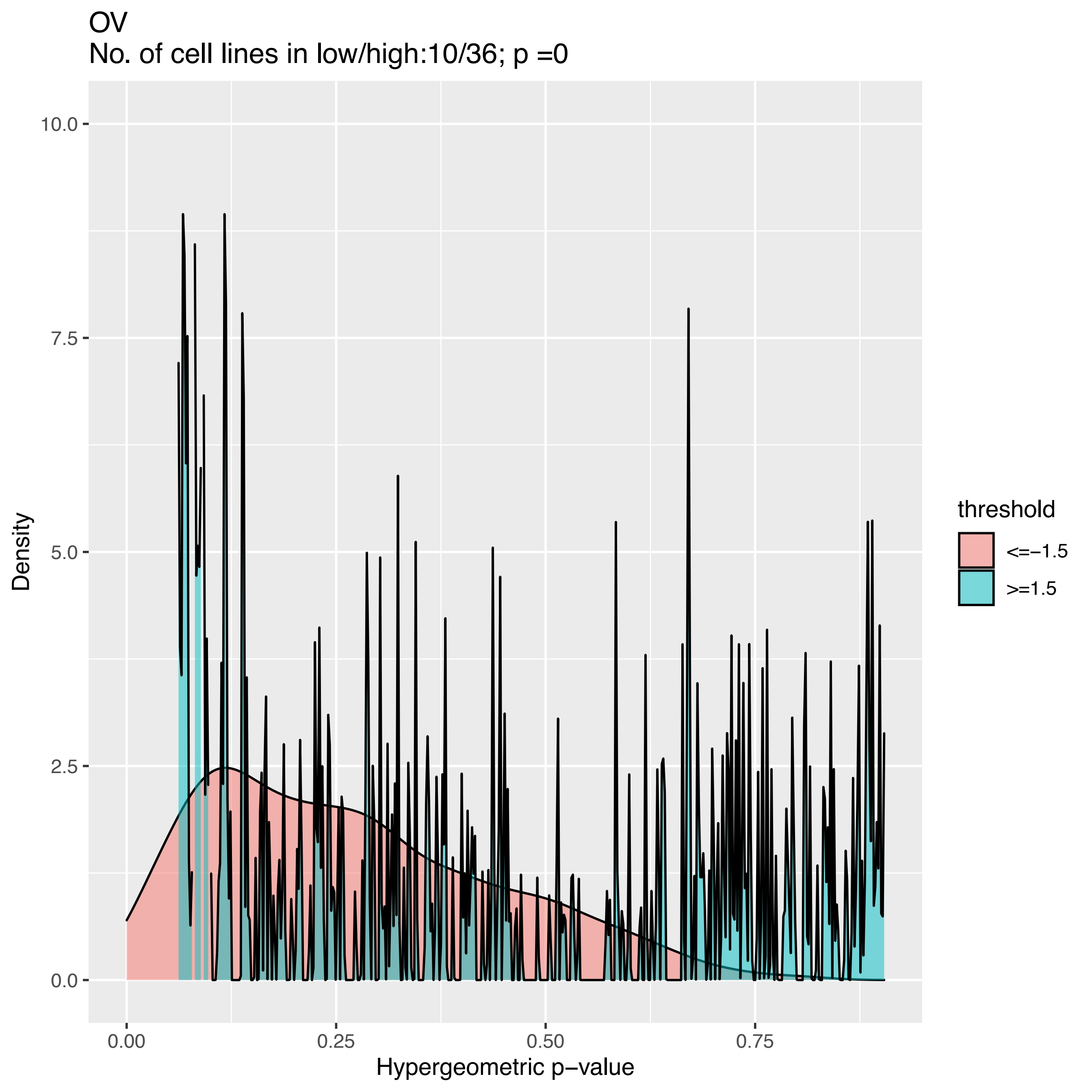

OV
No. of cell lines in low/high:10/36; p =0
Density
10.0
7.5
5.0
2.5
0.0
0.00
0.25
0.50
0.75
Hypergeometric p−value
threshold
<=−1.5
>=1.5

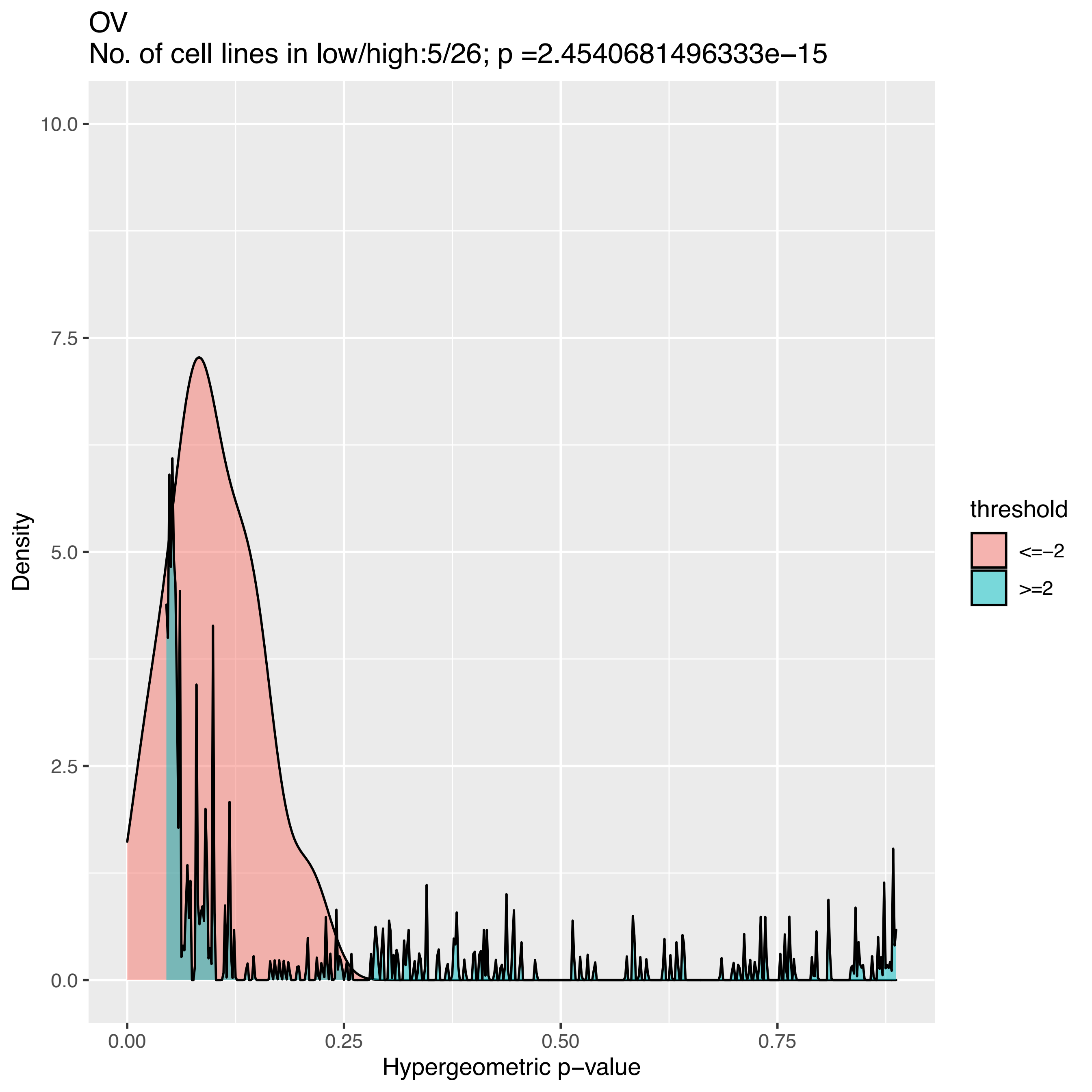
OV
No. of cell lines in low/high:5/26; p =2.4540681496333e−15
Density
10.0
7.5
5.0
2.5
0.0
0.00
0.25
0.50
0.75
Hypergeometric p−value
threshold
<=−2
>=2

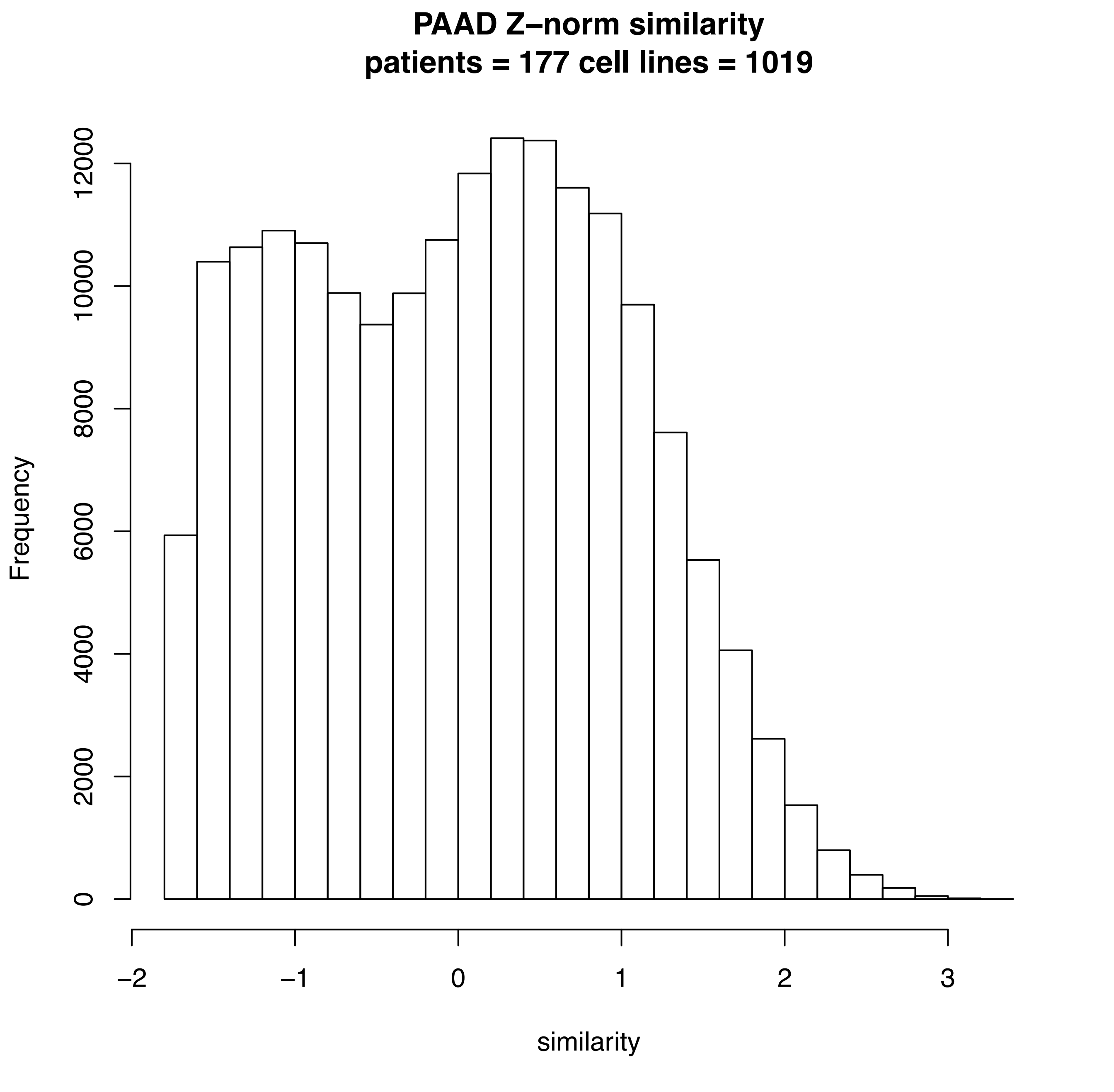
PAAD Z–norm similarity
patients = 177 cell lines = 1019
Frequency
0
2000
4000
6000
8000
10000
12000
–2
–1
0
1
2
3
similarity

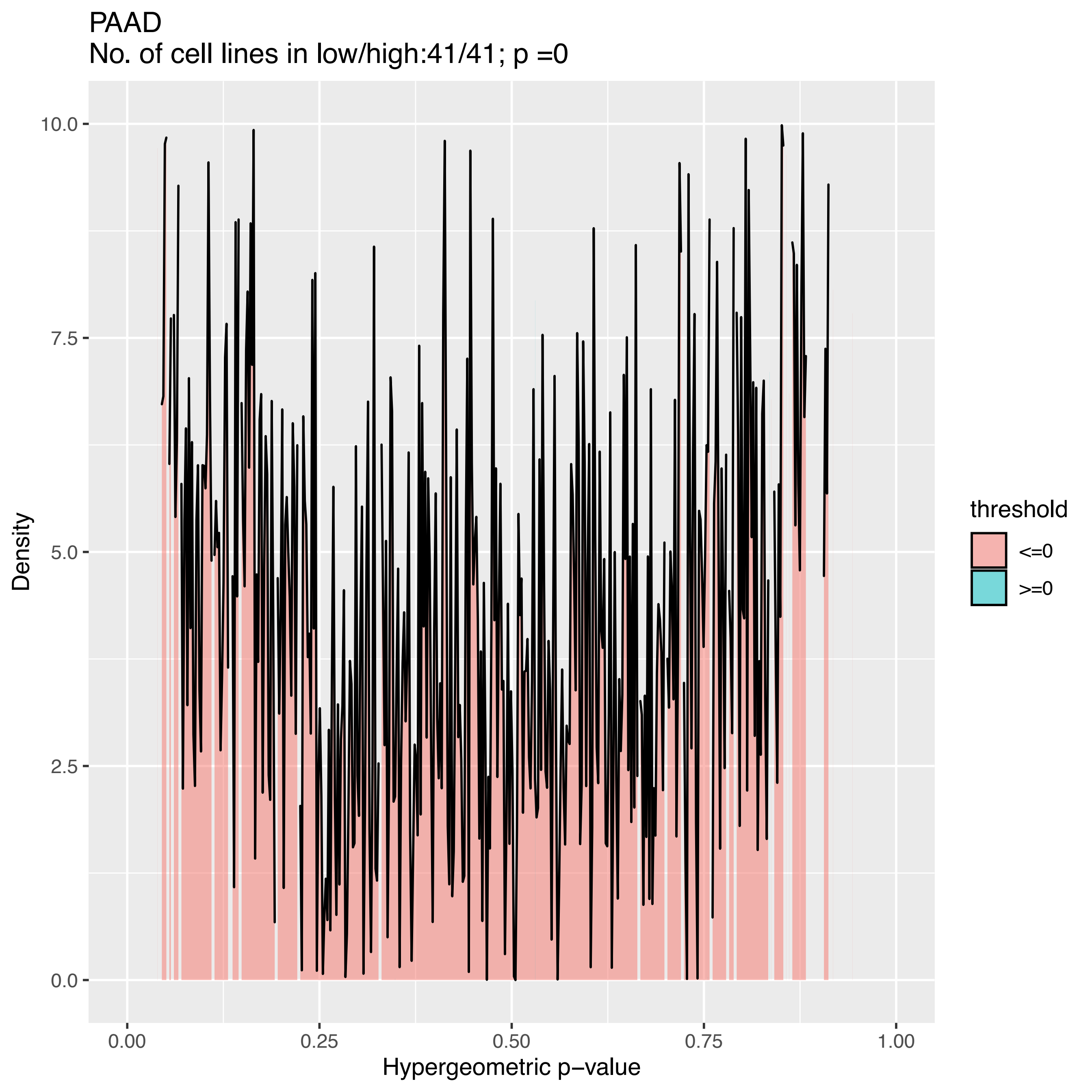

PAAD
No. of cell lines in low/high:41/41; p =0
Density
10.0
7.5
5.0
2.5
0.0
0.00
0.25
0.50
0.75
1.00
Hypergeometric p−value
threshold
<=0
>=0

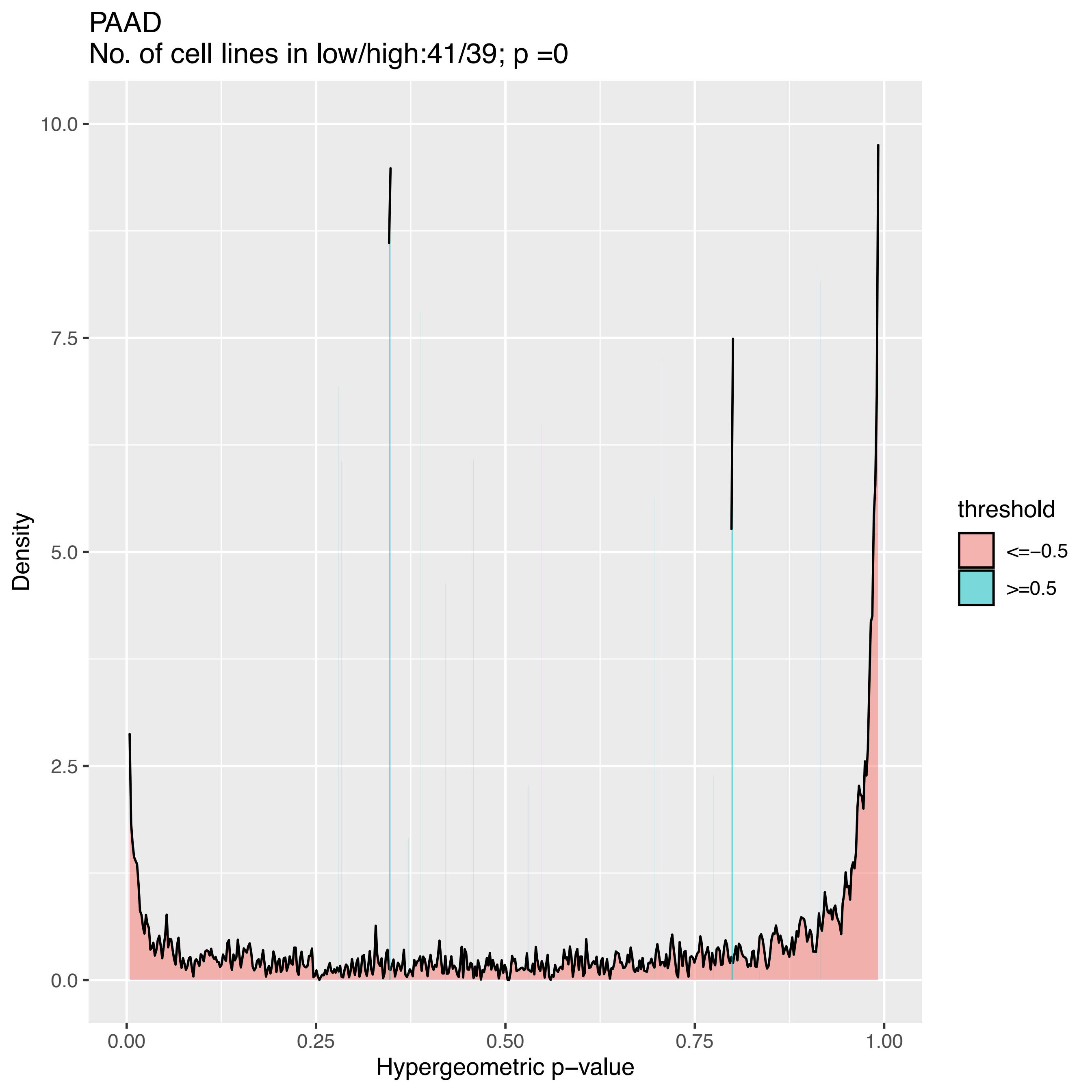

PAAD
No. of cell lines in low/high:41/39; p =0
Density
Hypergeometric p−value
10.0
7.5
5.0
2.5
0.0
0.00
0.25
0.50
0.75
1.00
threshold
<=−0.5
>=0.5

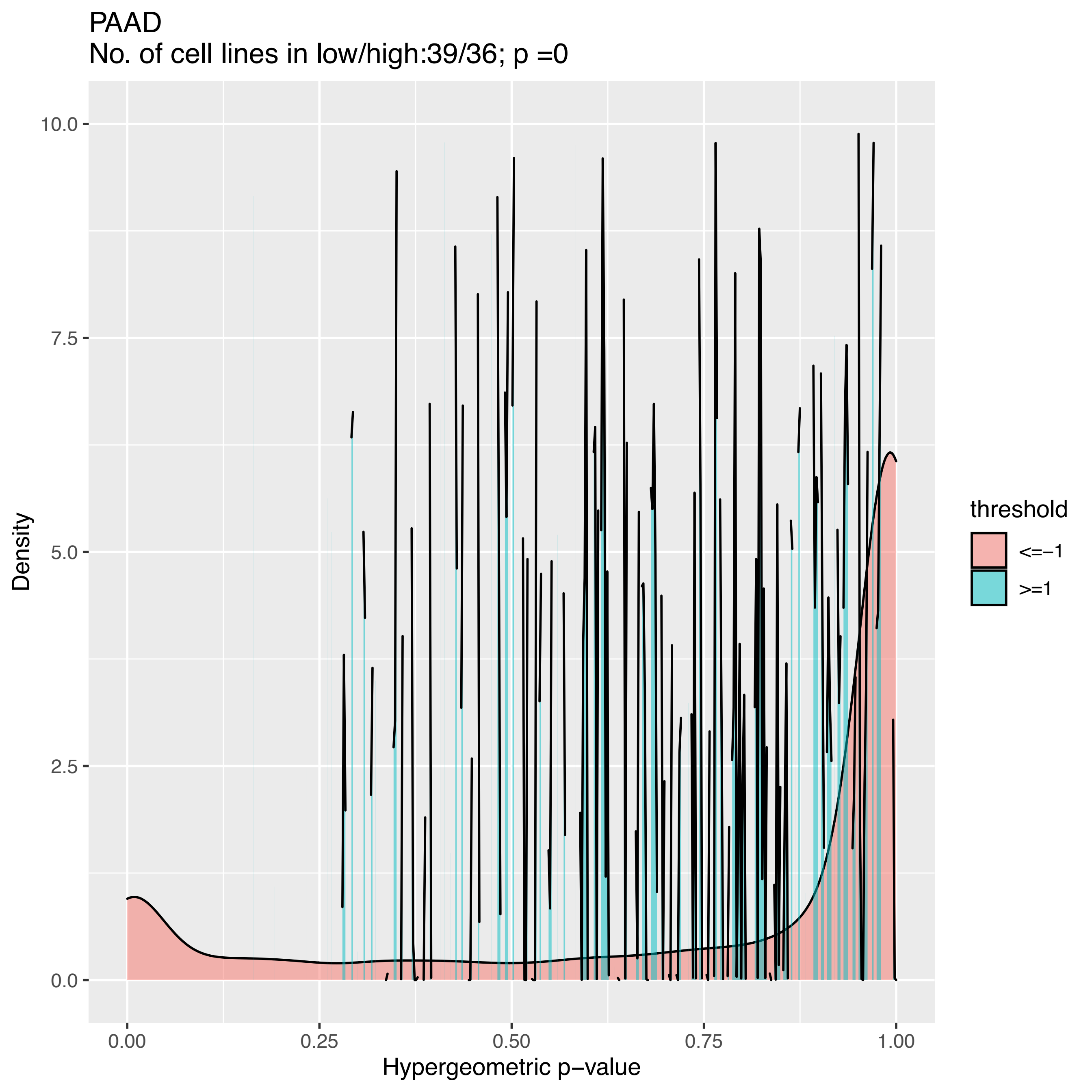

PAAD
No. of cell lines in low/high:39/36; p =0
Density
10.0
7.5
5.0
2.5
0.0
0.00
0.25
0.50
0.75
1.00
Hypergeometric p−value
threshold
<=−1
>=1

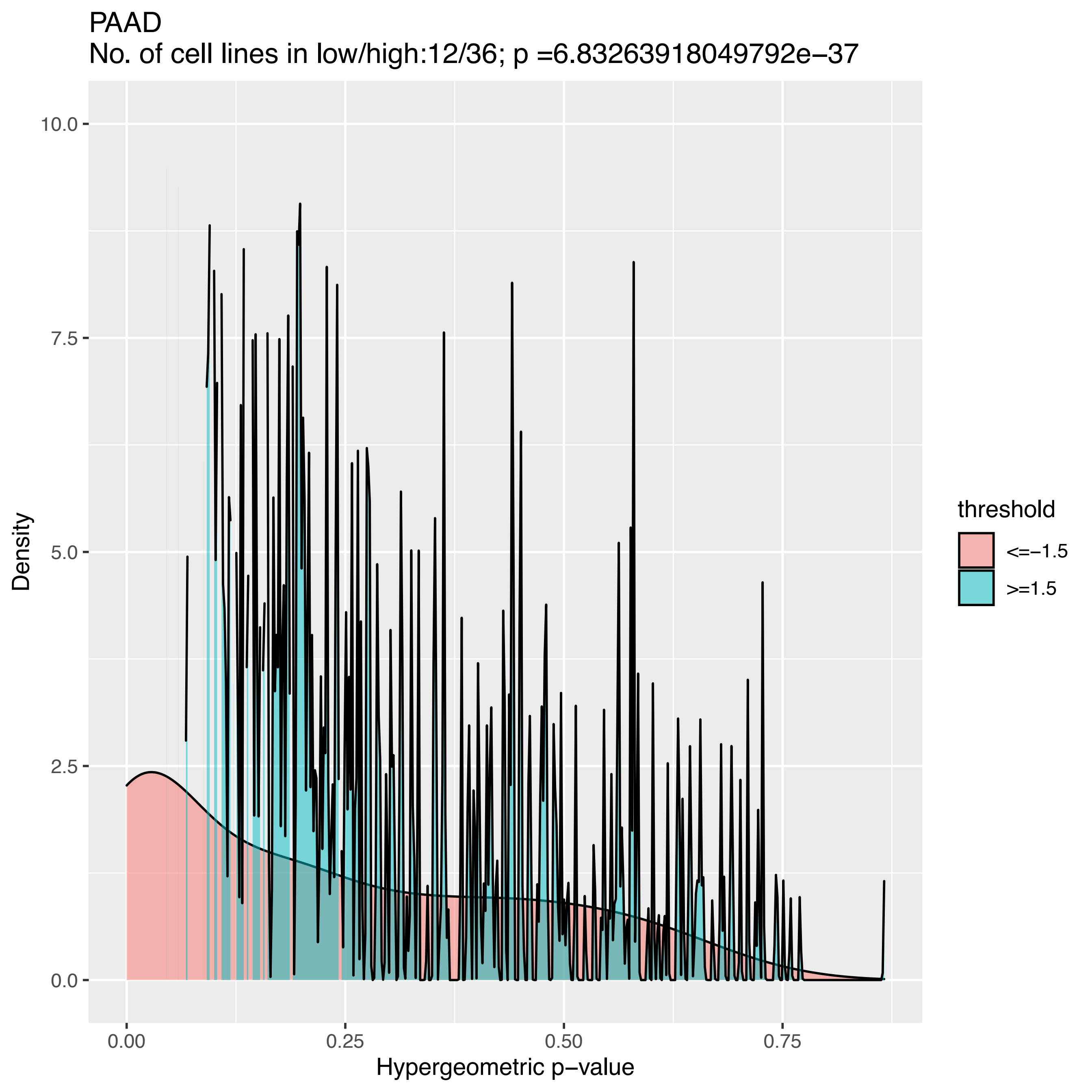

PAAD
No. of cell lines in low/high:12/36; p =6.83263918049792e−37
Density
10.0
7.5
5.0
2.5
0.0
0.00
0.25
0.50
0.75
Hypergeometric p−value
threshold
<=−1.5
>=1.5

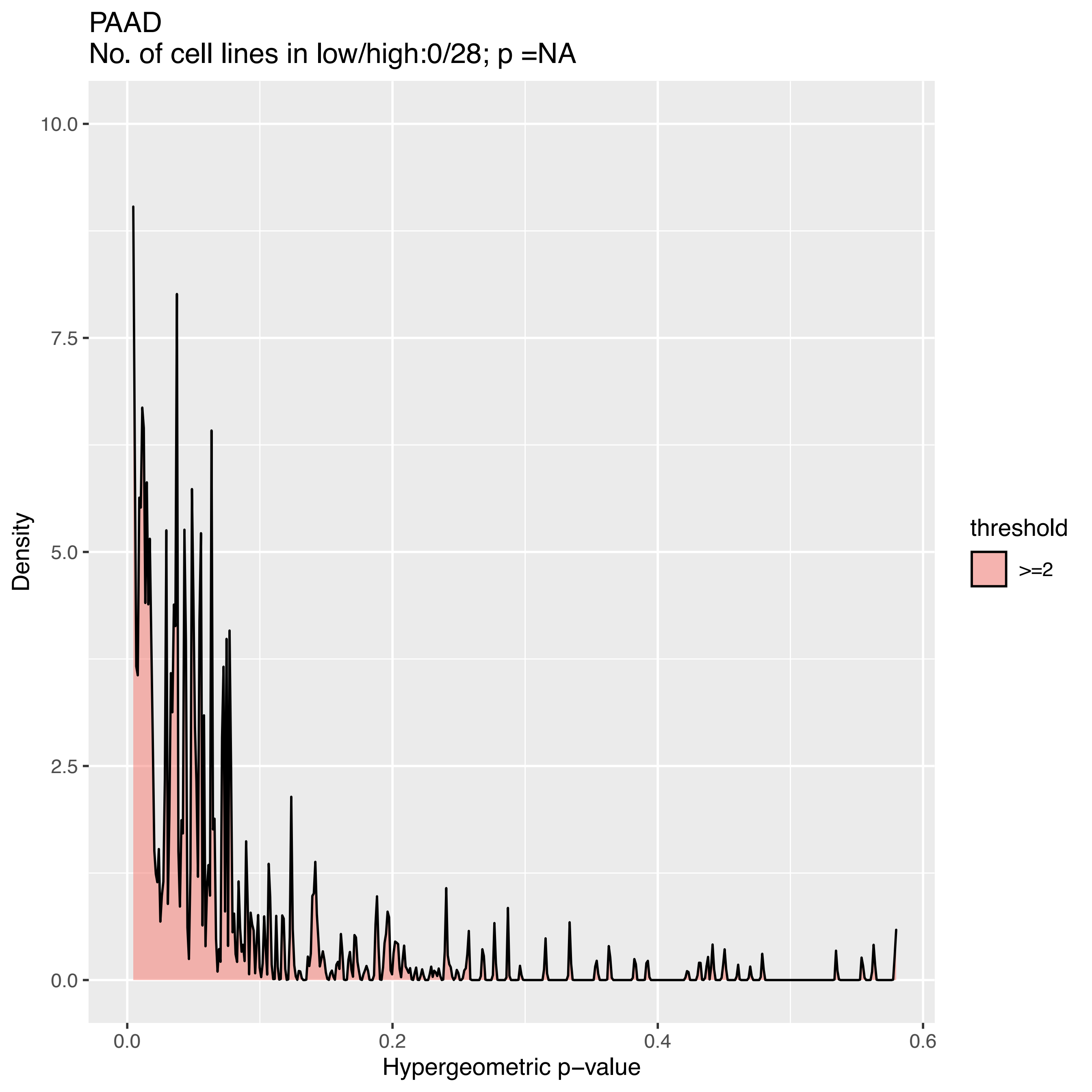
PAAD
No. of cell lines in low/high:0/28; p =NA
Density
10.0
7.5
5.0
2.5
0.0
0.0
0.2
0.4
0.6
Hypergeometric p−value
threshold
>=2

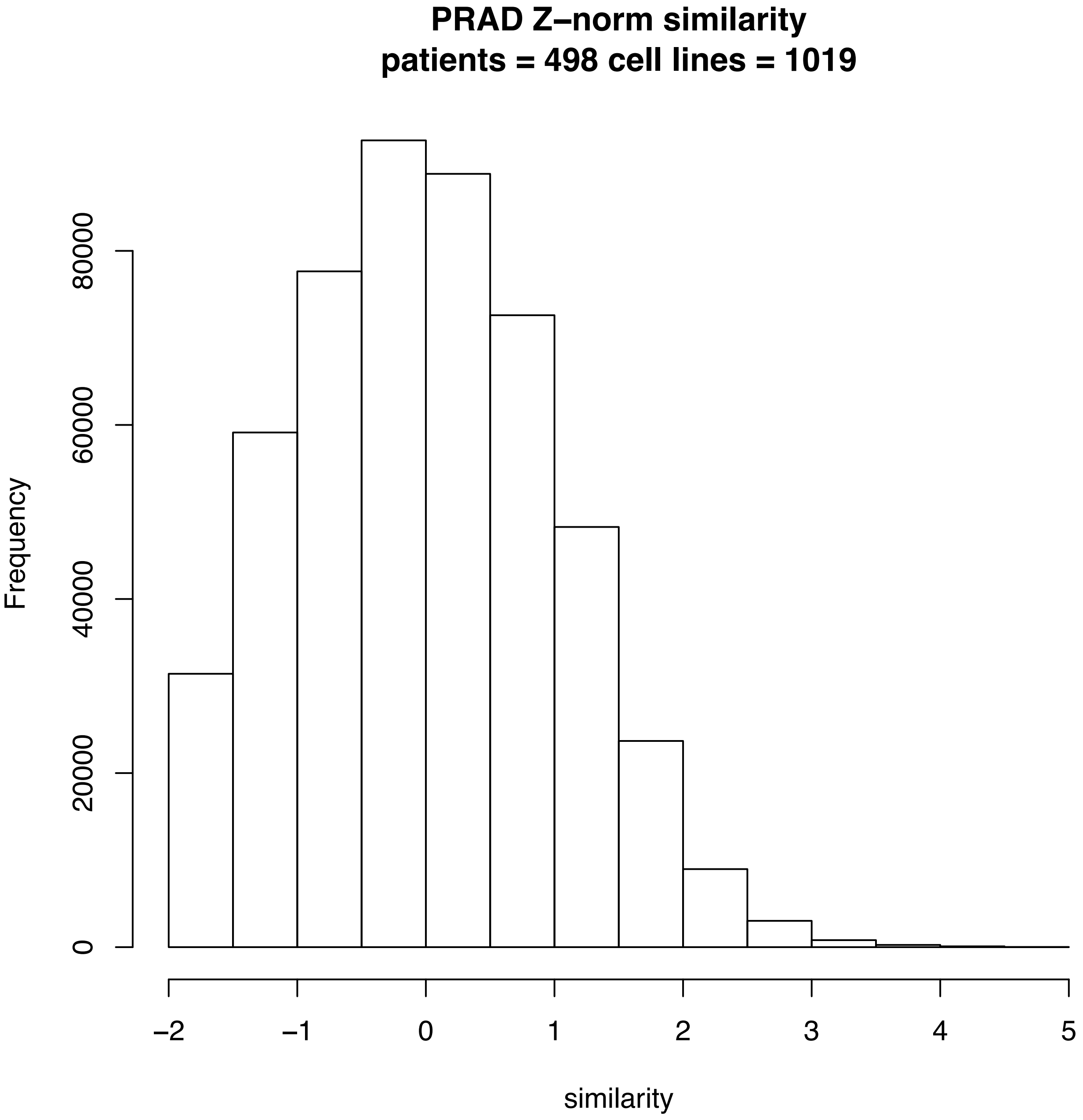
PRAD Z–norm similarity
patients = 498 cell lines = 1019
Frequency
0
20000
40000
60000
80000
−2
−1
0
1
2
3
4
5
similarity

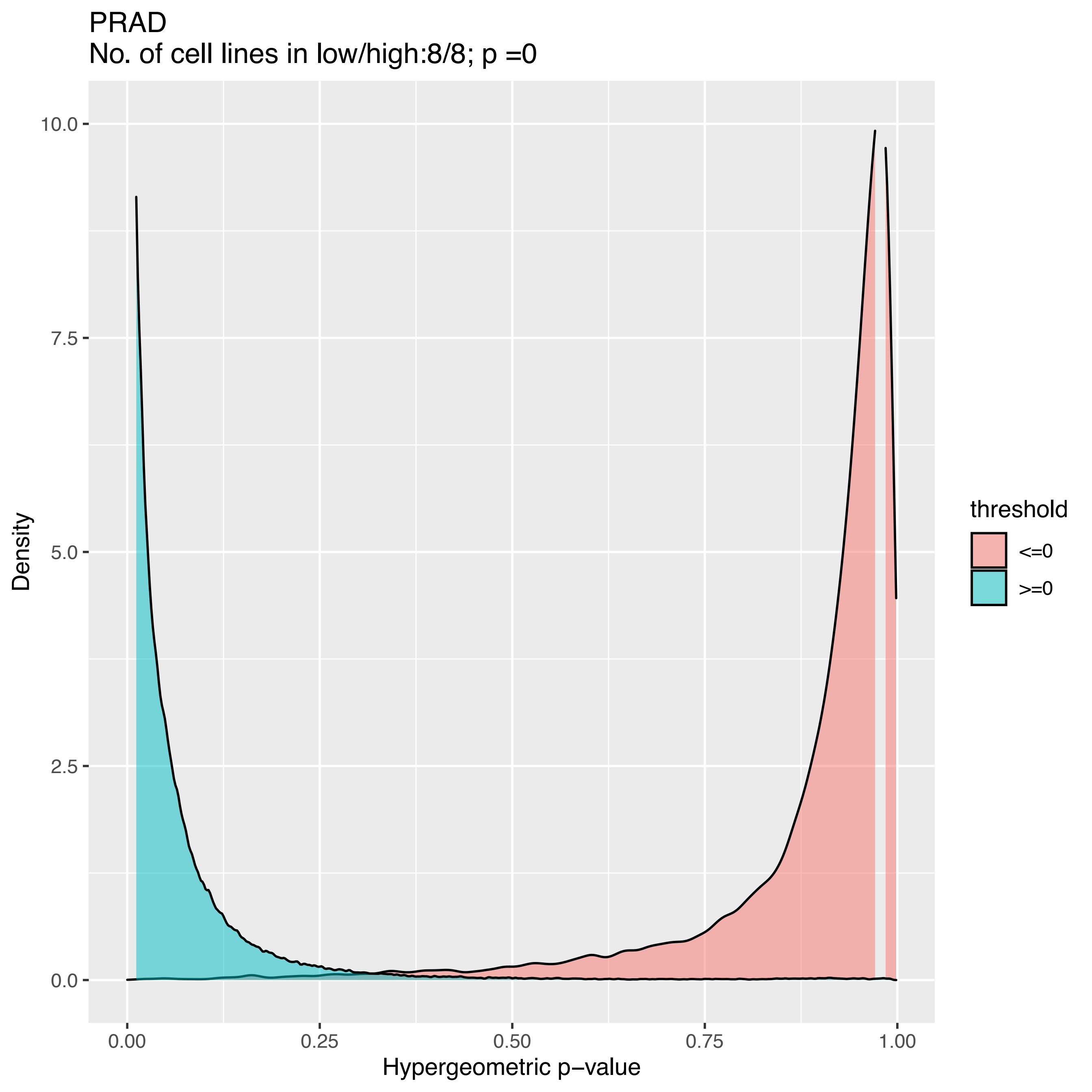
PRAD
No. of cell lines in low/high:8/8; p =0
Density
10.0
7.5
5.0
2.5
0.0
0.00
0.25
0.50
0.75
1.00
Hypergeometric p−value
threshold
<=0
>=0

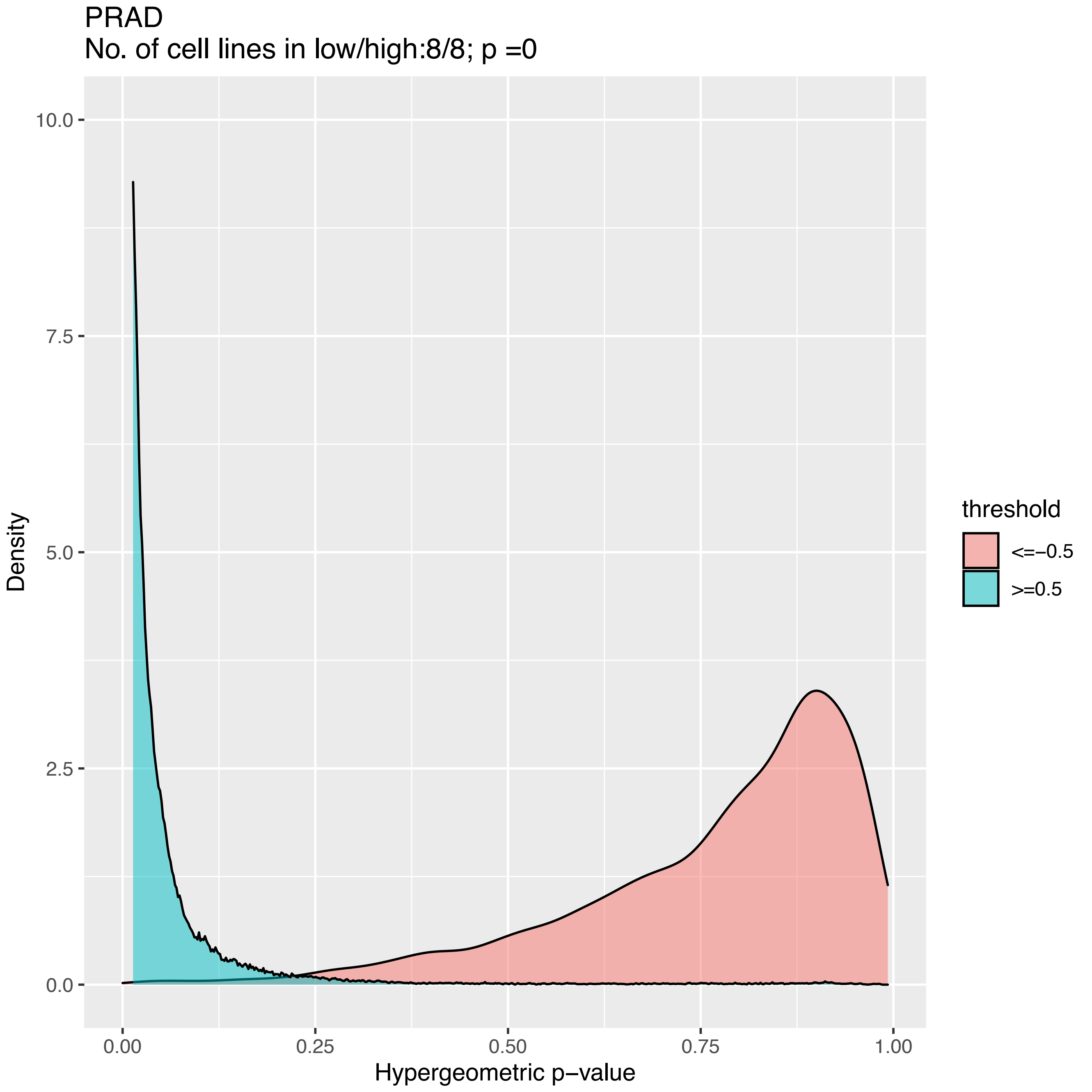
PRAD
No. of cell lines in low/high:8/8; p =0
10.0
7.5
5.0
2.5
0.0
Density
0.00
0.25
0.50
0.75
1.00
Hypergeometric p−value
threshold
<=−0.5
>=0.5

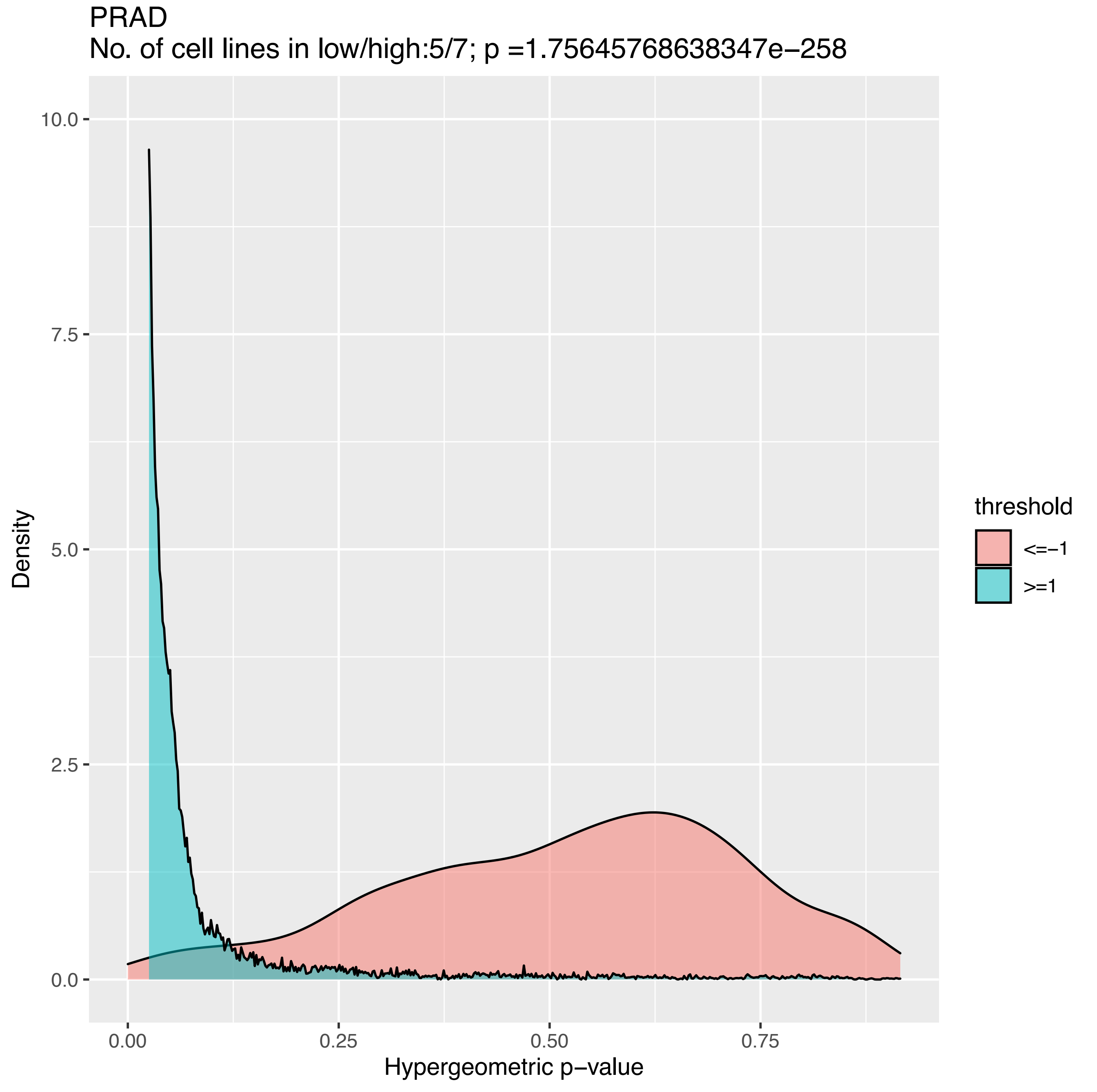
PRAD
No. of cell lines in low/high:5/7; p =1.75645768638347e−258
Density
10.0
7.5
5.0
2.5
0.0
0.00
0.25
0.50
0.75
Hypergeometric p−value
threshold
<=−1
>=1

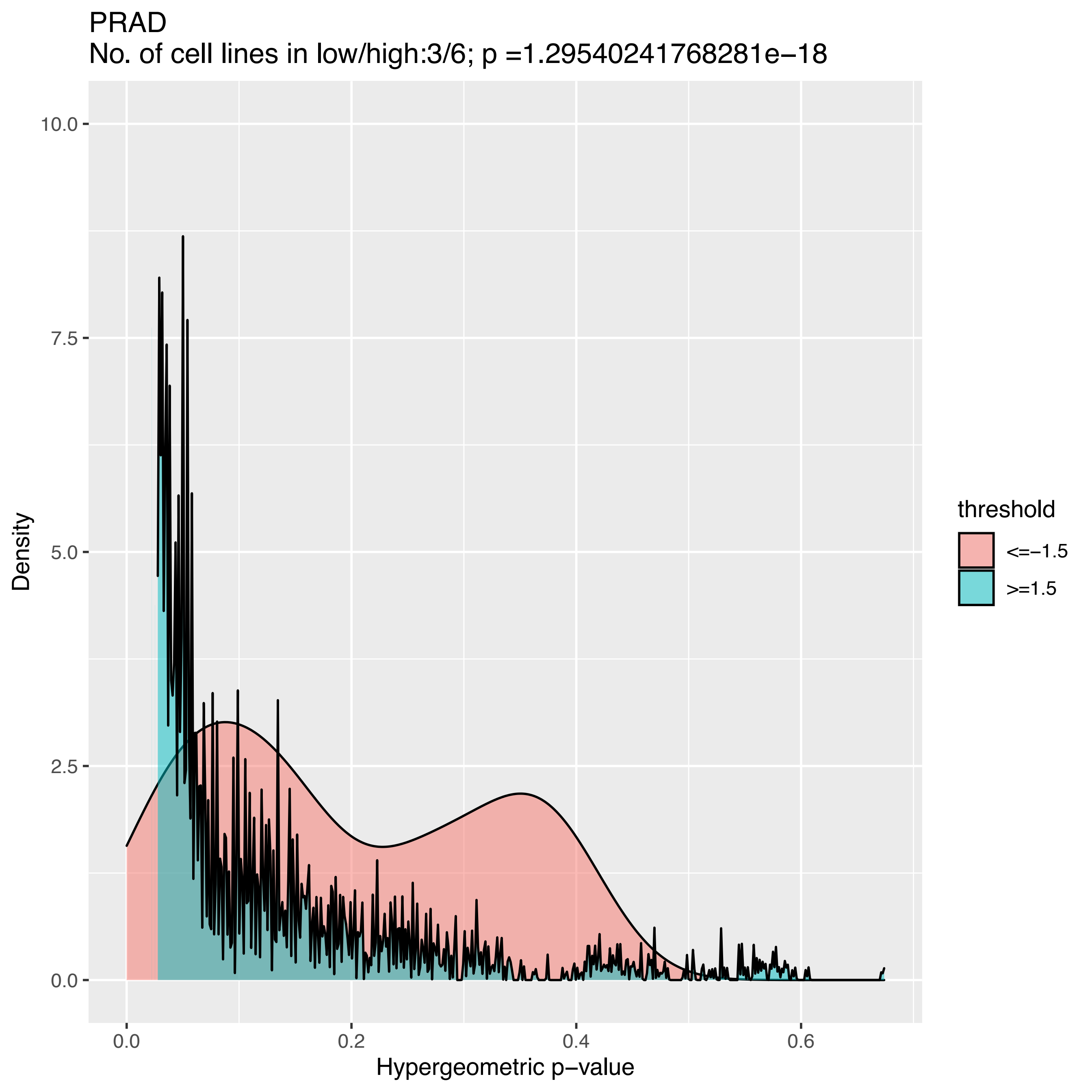
PRAD
No. of cell lines in low/high:3/6; p =1.29540241768281e−18
Density
10.0
7.5
5.0
2.5
0.0
0.0
0.2
0.4
0.6
Hypergeometric p−value
threshold
<=−1.5
>=1.5

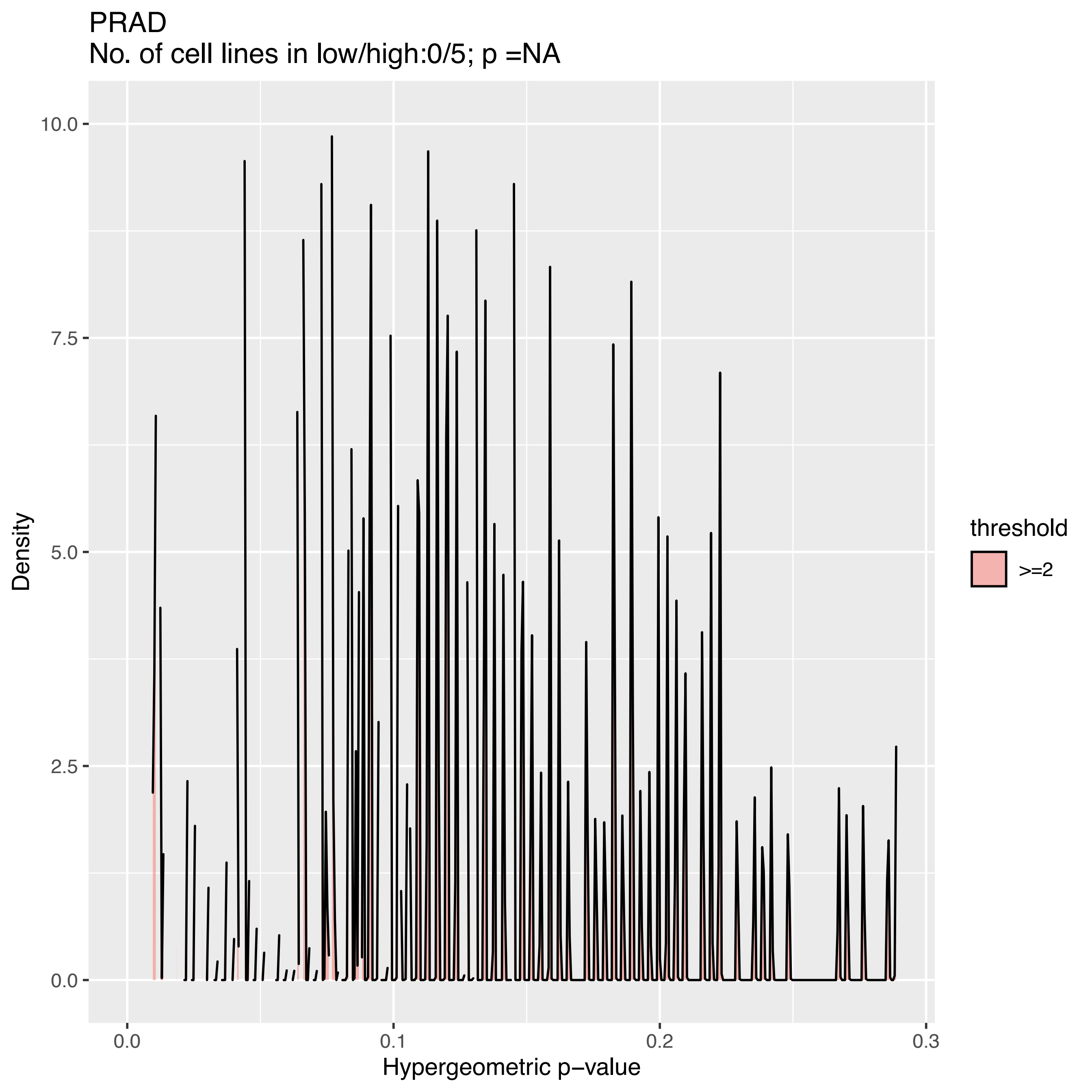
PRAD
No. of cell lines in low/high:0/5; p =NA
Density
10.0
7.5
5.0
2.5
0.0
0.0
0.1
0.2
0.3
Hypergeometric p−value
threshold
>=2

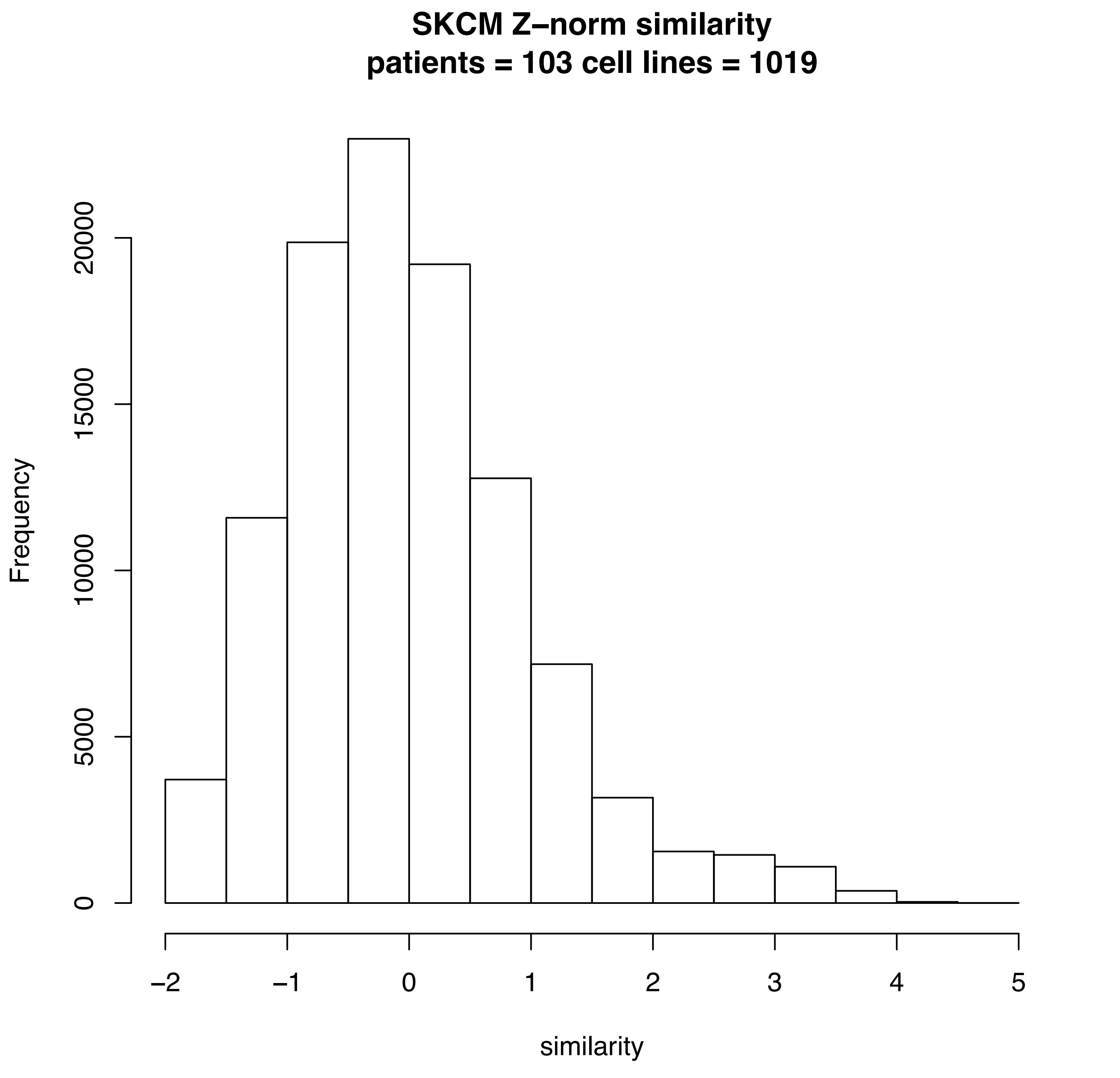
SKCM Z–norm similarity
patients = 103 cell lines = 1019
Frequency
0
5000
10000
15000
20000
–2
–1
0
1
2
3
4
5
similarity

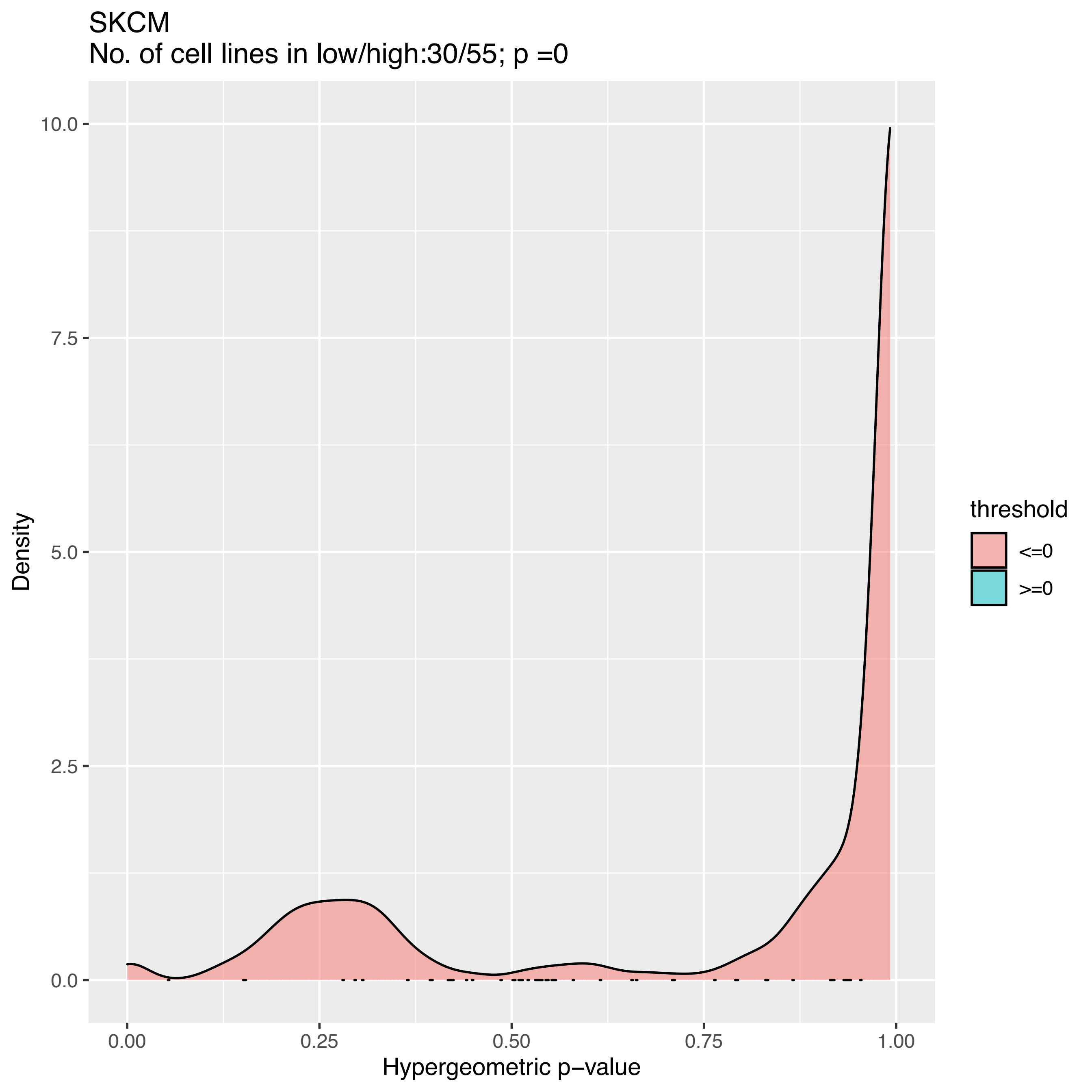
SKCM
No. of cell lines in low/high:30/55; p =0
Density
Hypergeometric p−value
0.0
2.5
5.0
7.5
10.0
0.00
0.25
0.50
0.75
1.00
threshold
<=0
>=0

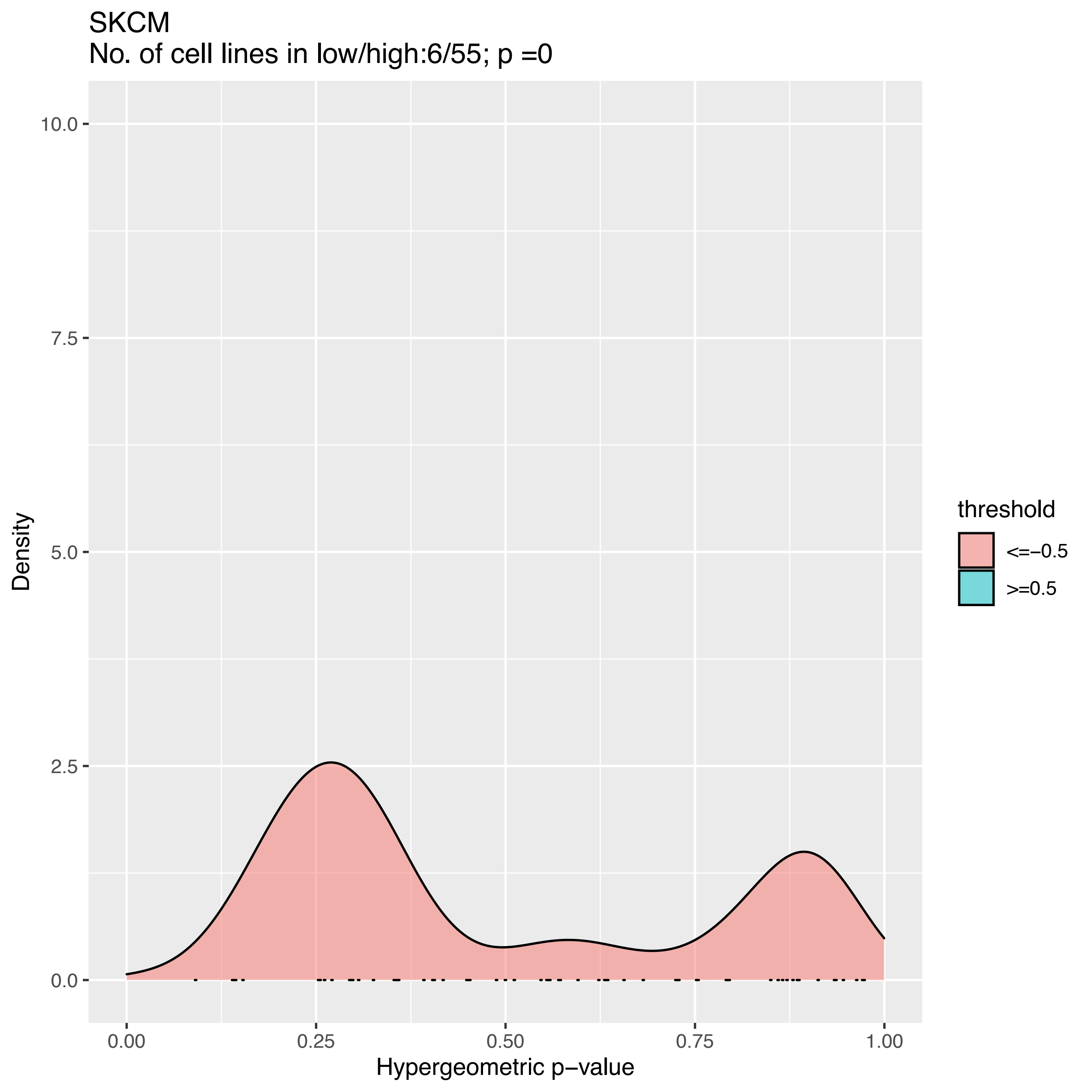
SKCM
No. of cell lines in low/high:6/55; p =0
10.0
7.5
5.0
2.5
0.0
Density
0.00
0.25
0.50
0.75
1.00
Hypergeometric p−value
threshold
<=−0.5
>=0.5

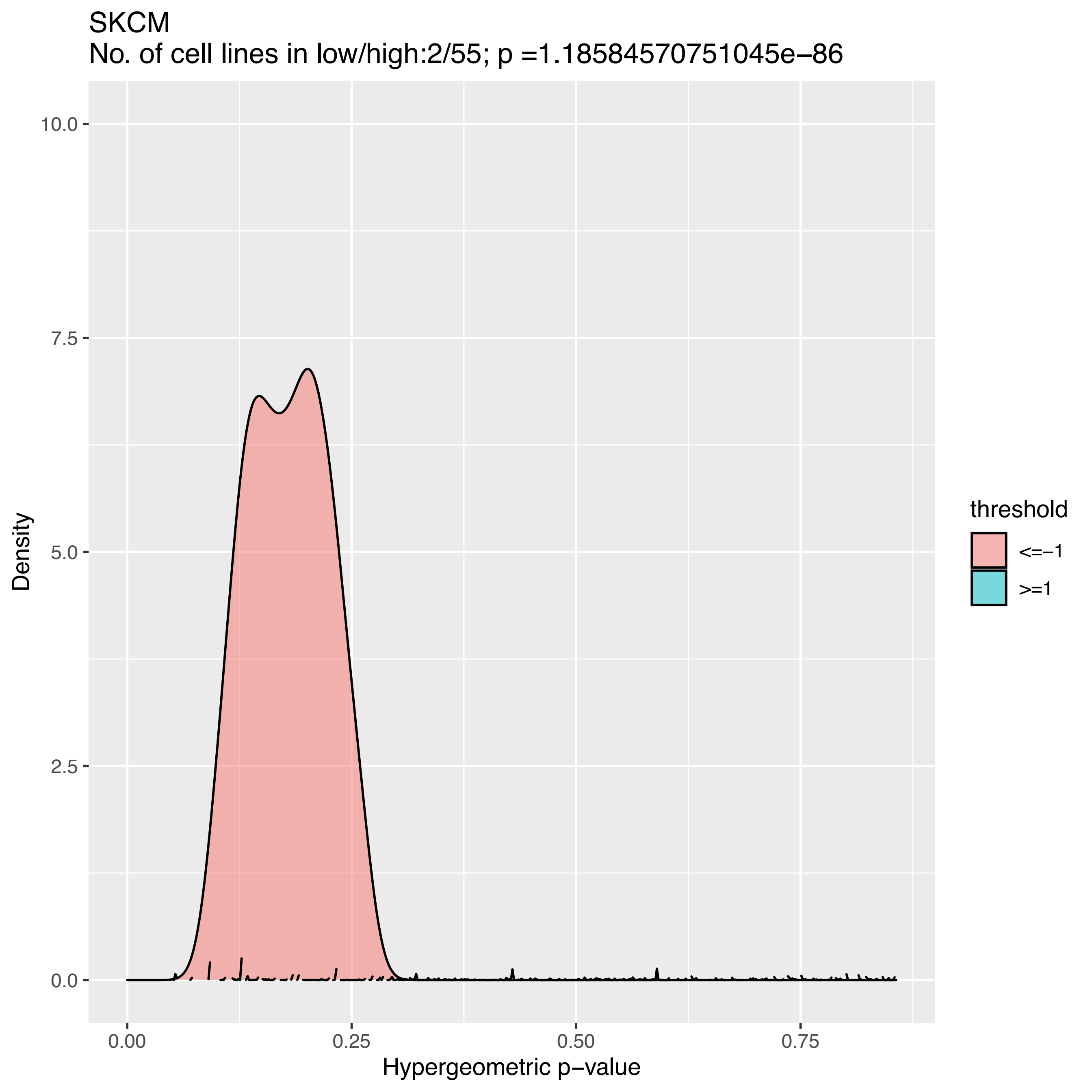

SKCM
No. of cell lines in low/high:2/55; p =1.18584570751045e−86
Density
10.0
7.5
5.0
2.5
0.0
0.00
0.25
0.50
0.75
Hypergeometric p−value
threshold
<=−1
>=1

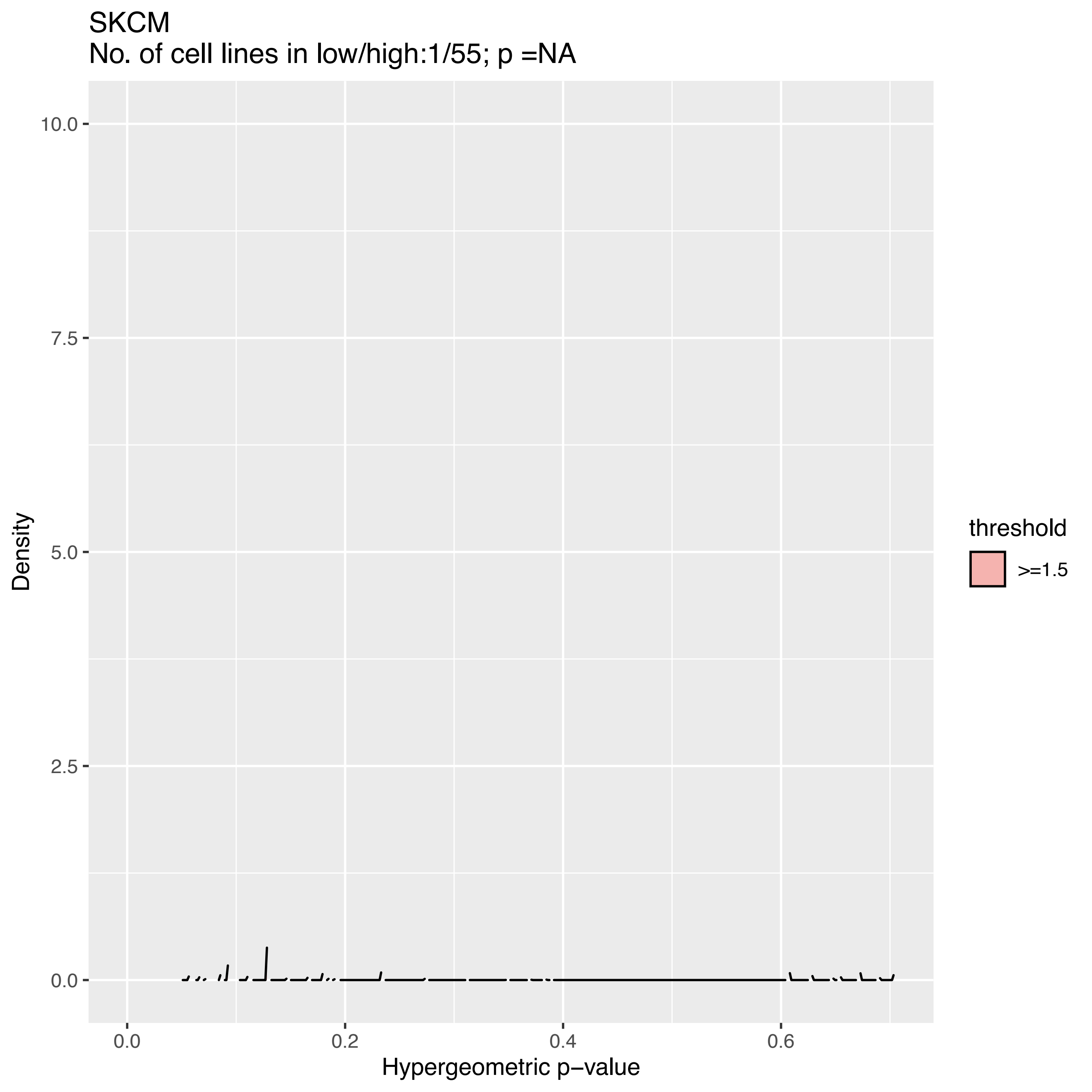

SKCM
No. of cell lines in low/high:1/55; p =NA
10.0
7.5
5.0
2.5
0.0
Density
0.0
0.2
0.4
0.6
Hypergeometric p−value
threshold
>=1.5

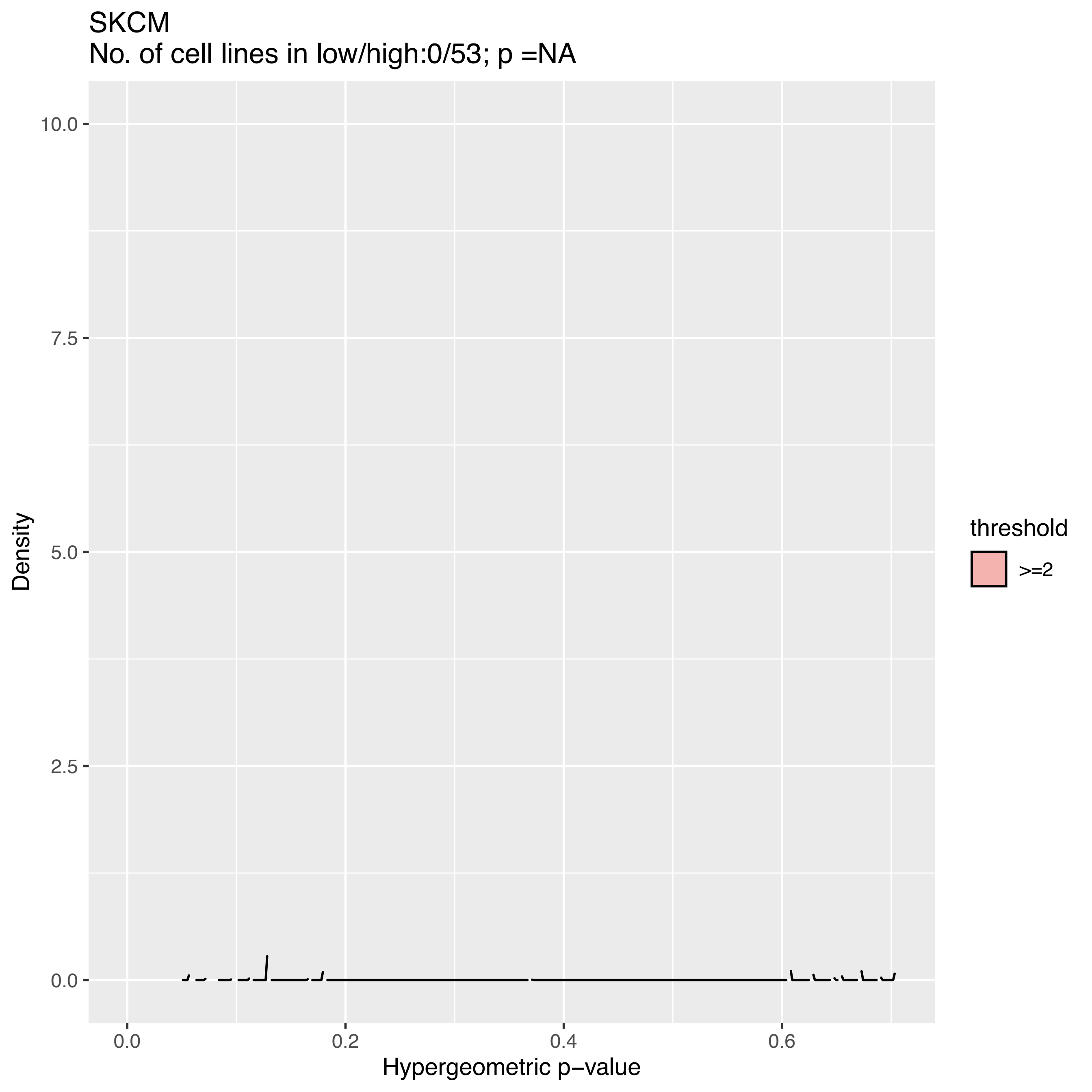
SKCM
No. of cell lines in low/high:0/53; p =NA
10.0
7.5
5.0
2.5
0.0
0.0
0.2
0.4
0.6
Density
Hypergeometric p−value
threshold
>=2

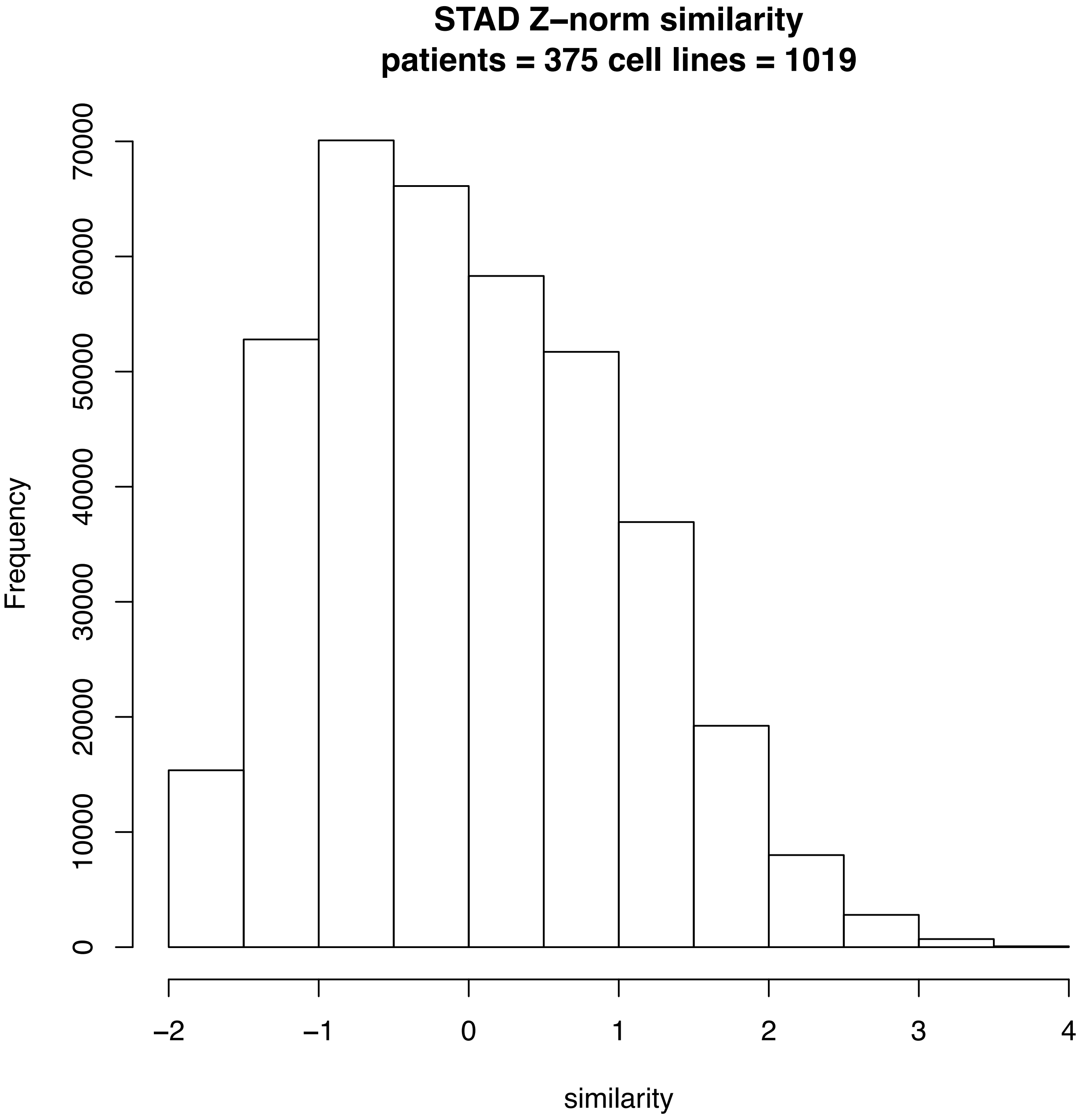
STAD Z–norm similarity
patients = 375 cell lines = 1019
Frequency
0
10000
20000
30000
40000
50000
60000
70000
–2
–1
0
1
2
3
4
similarity

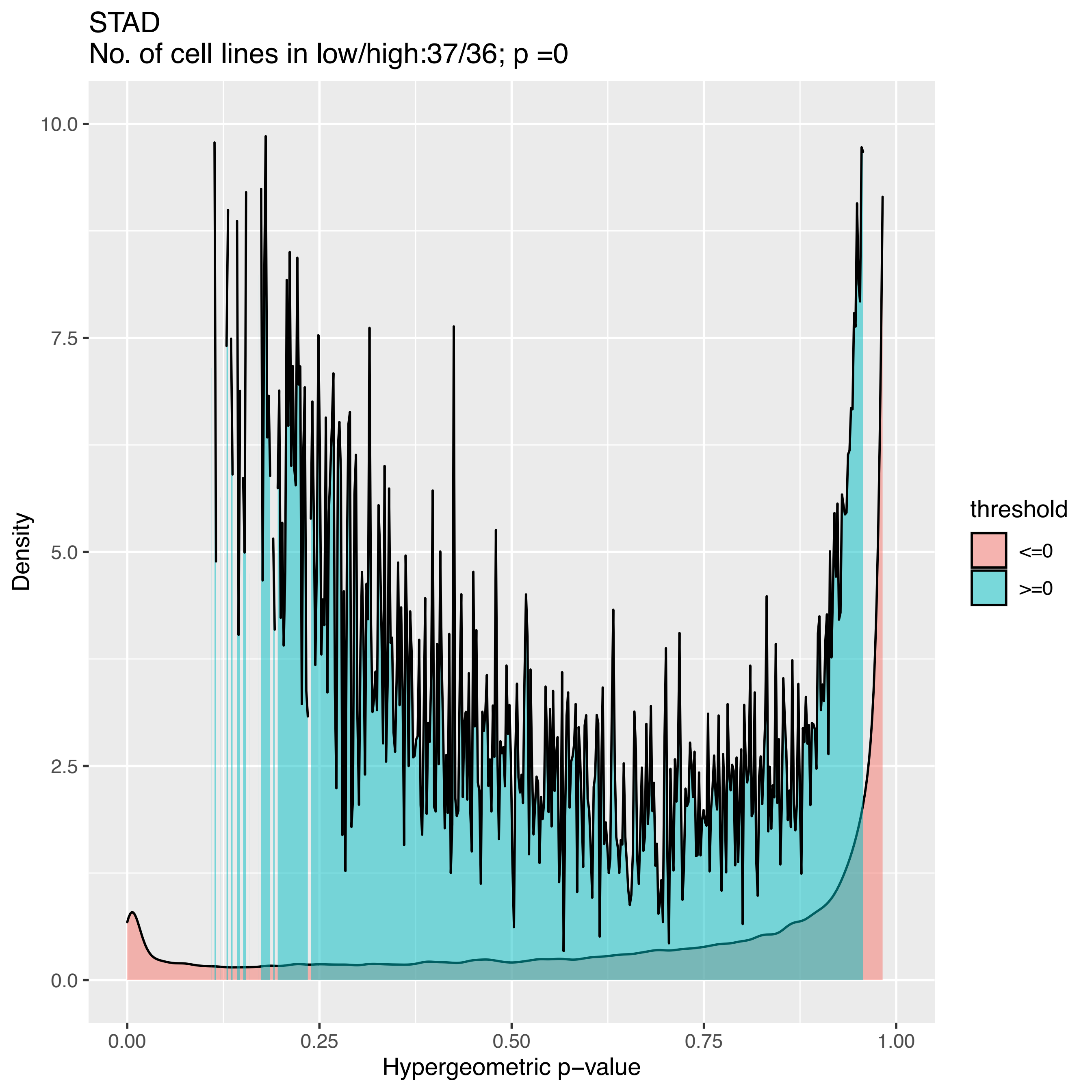
STAD
No. of cell lines in low/high:37/36; p =0
Density
Hypergeometric p−value
threshold
<=0
>=0
0.0
2.5
5.0
7.5
10.0
0.00
0.25
0.50
0.75
1.00

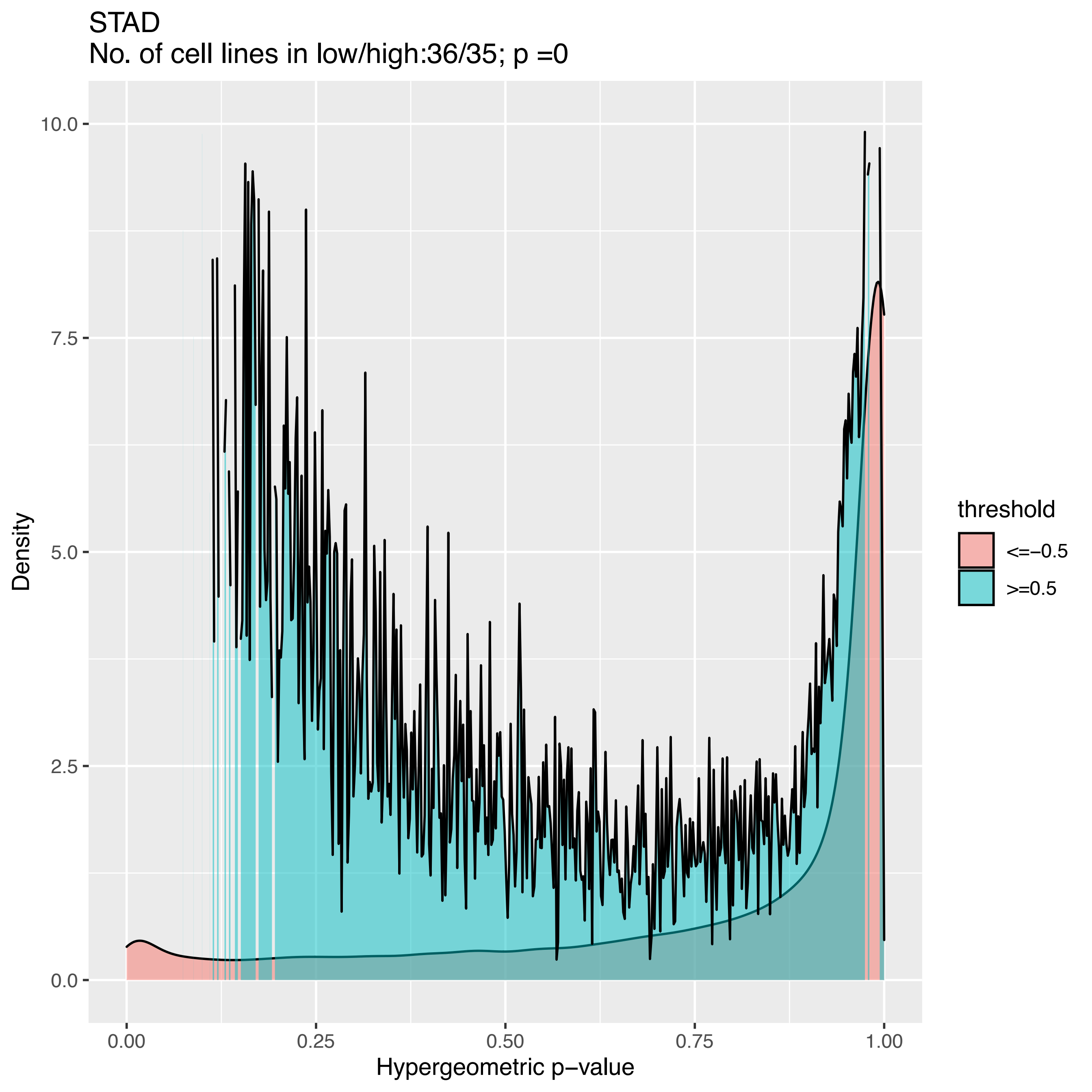
STAD
No. of cell lines in low/high:36/35; p =0
Density
Hypergeometric p−value
0.0
2.5
5.0
7.5
10.0
0.00
0.25
0.50
0.75
1.00
threshold
<=−0.5
>=0.5

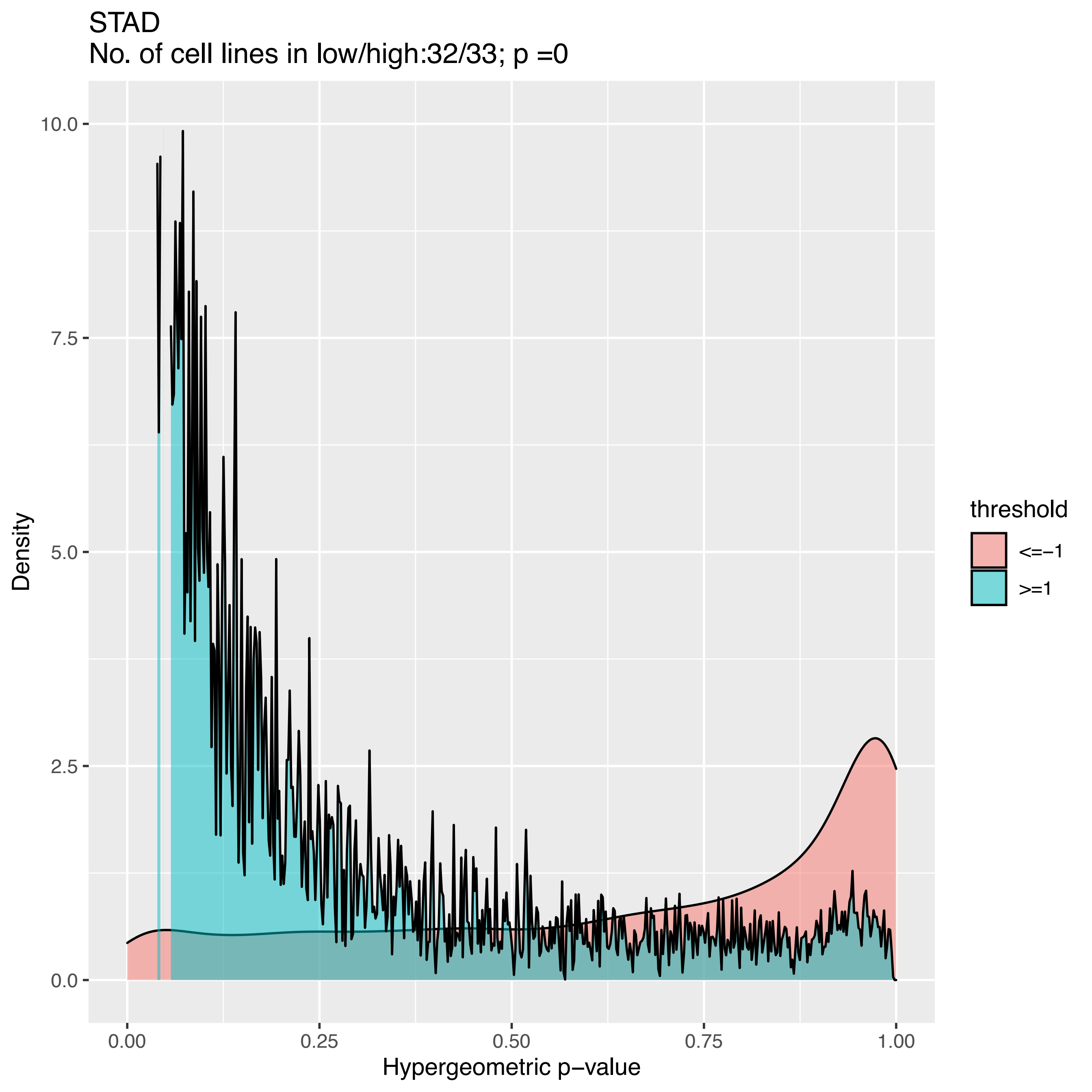

STAD
No. of cell lines in low/high:32/33; p =0
Density
10.0
7.5
5.0
2.5
0.0
0.00
0.25
0.50
0.75
1.00
Hypergeometric p−value
threshold
<=−1
>=1

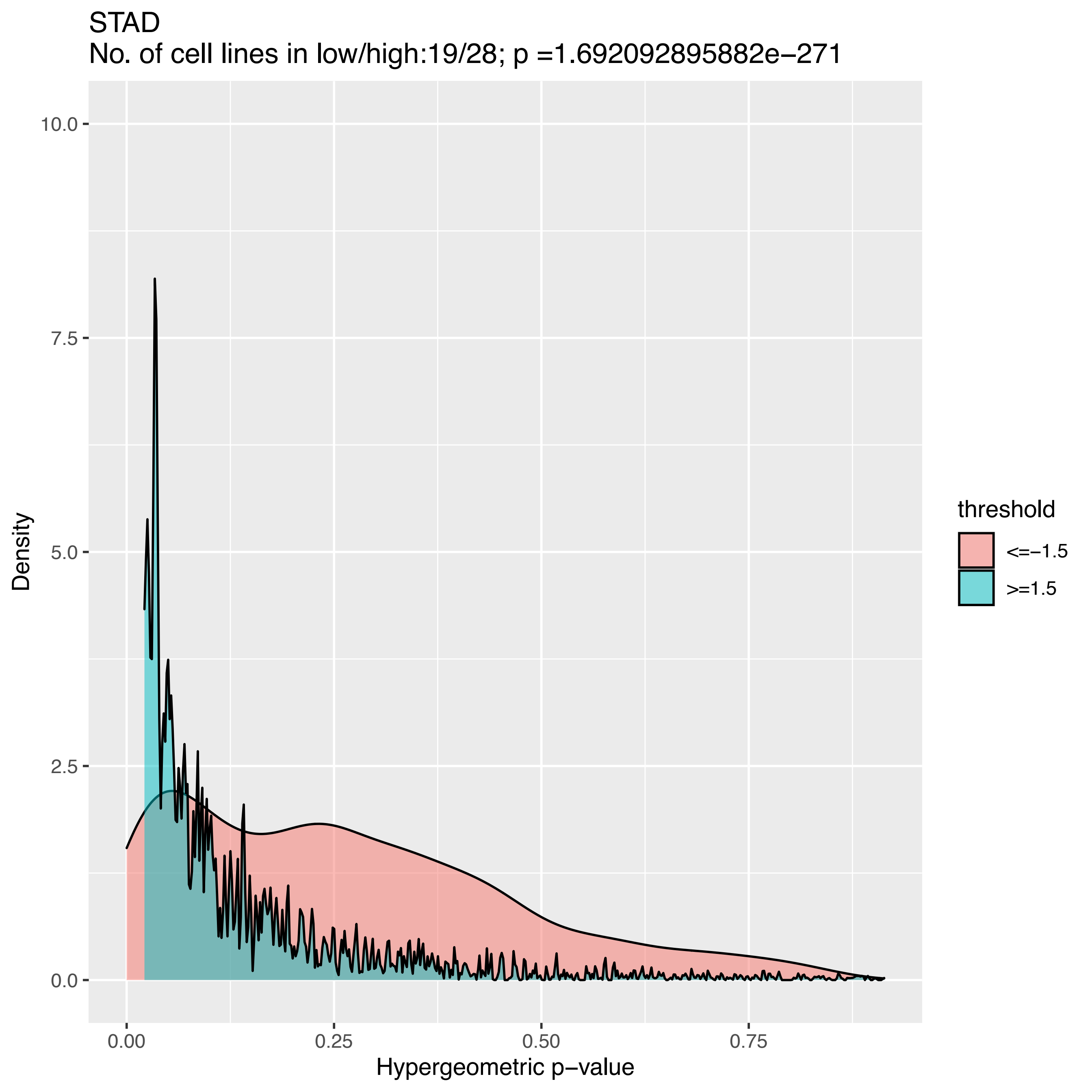
STAD
No. of cell lines in low/high:19/28; p =1.692092895882e−271
Density
10.0
7.5
5.0
2.5
0.0
0.00
0.25
0.50
0.75
Hypergeometric p−value
threshold
<=−1.5
>=1.5

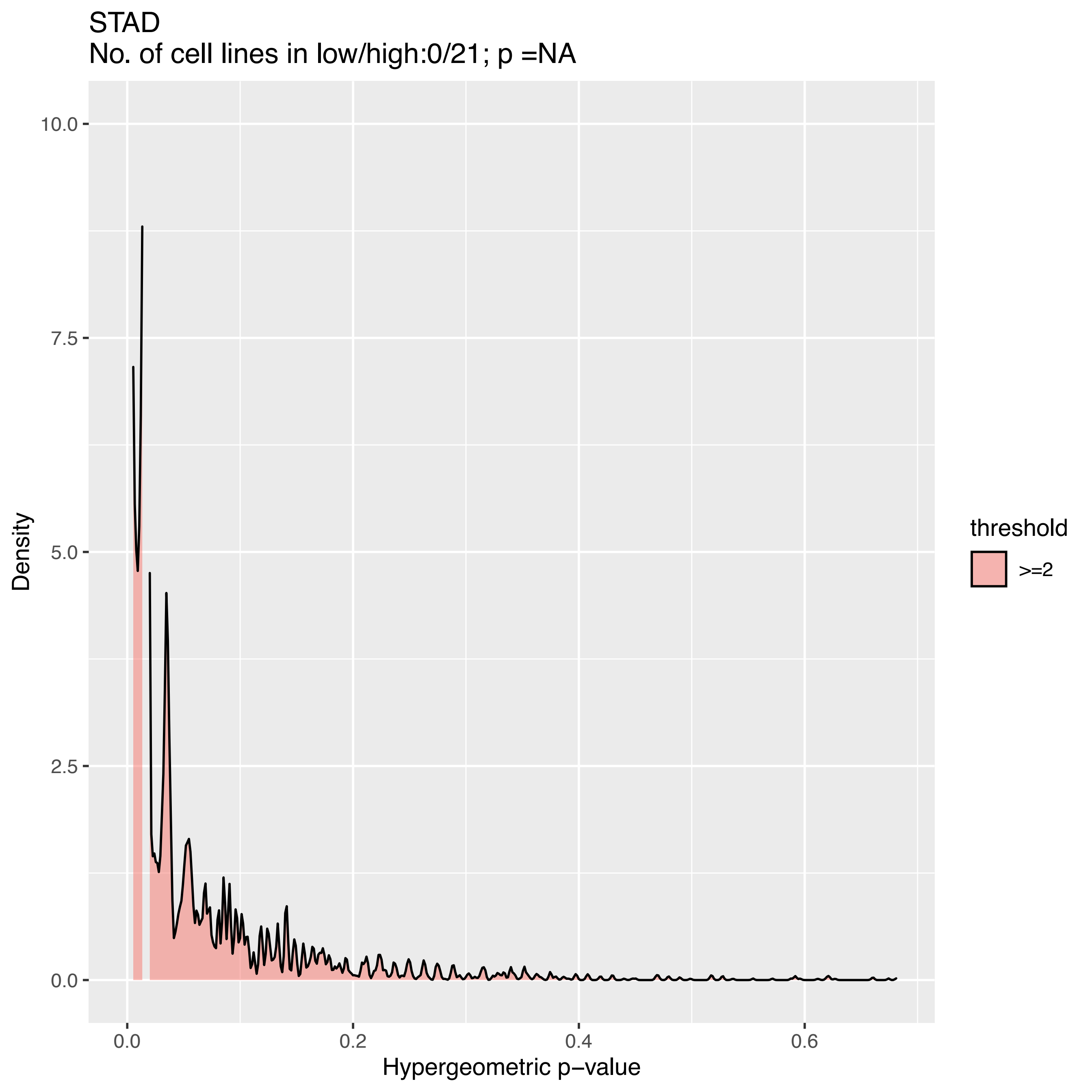

STAD
No. of cell lines in low/high:0/21; p =NA
Density
10.0
7.5
5.0
2.5
0.0
0.0
0.2
0.4
0.6
Hypergeometric p−value
threshold
>=2

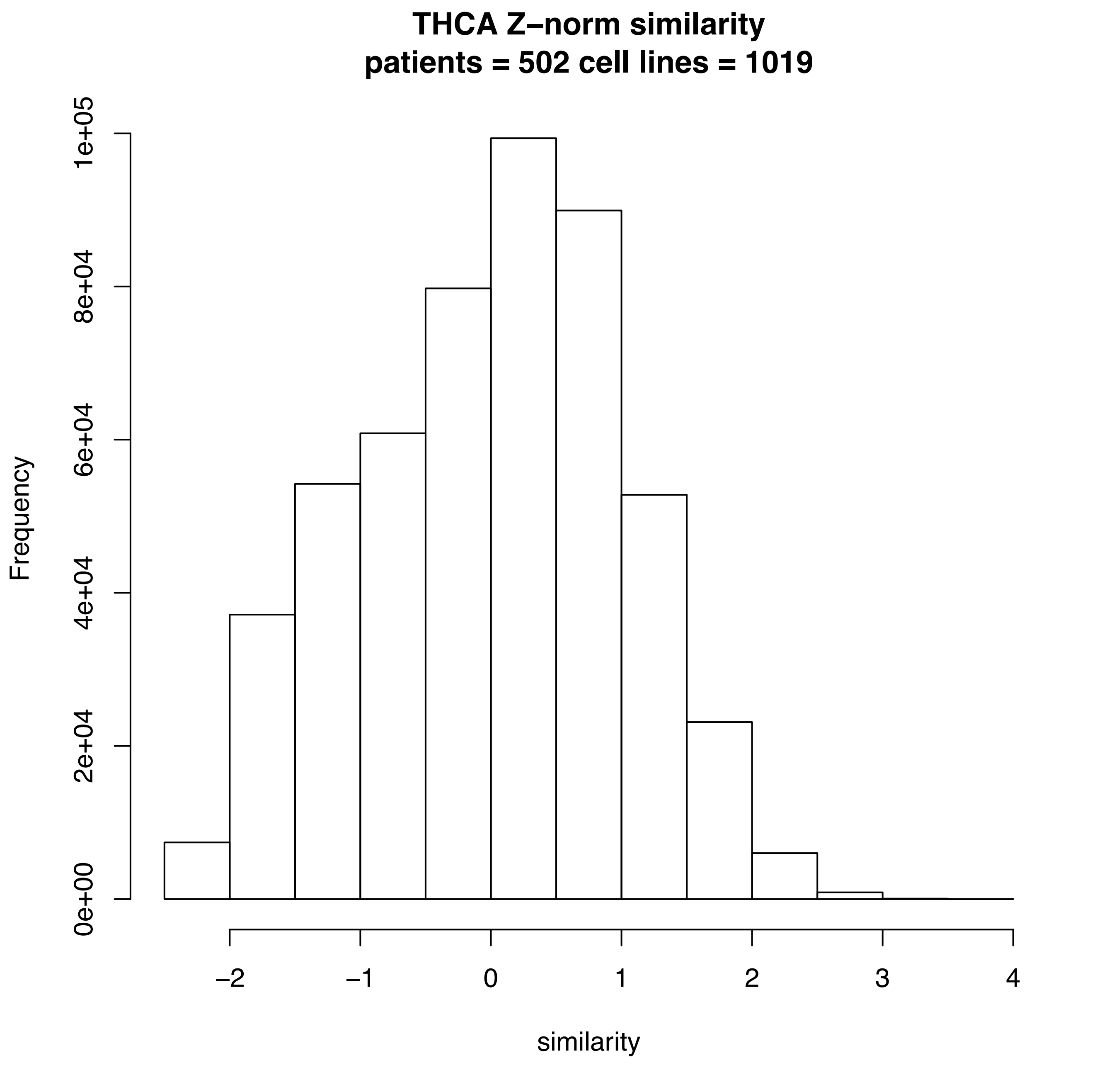
THCA Z–norm similarity
patients = 502 cell lines = 1019
Frequency
0e+00
2e+04
4e+04
6e+04
8e+04
1e+05
–2
–1
0
1
2
3
4
similarity

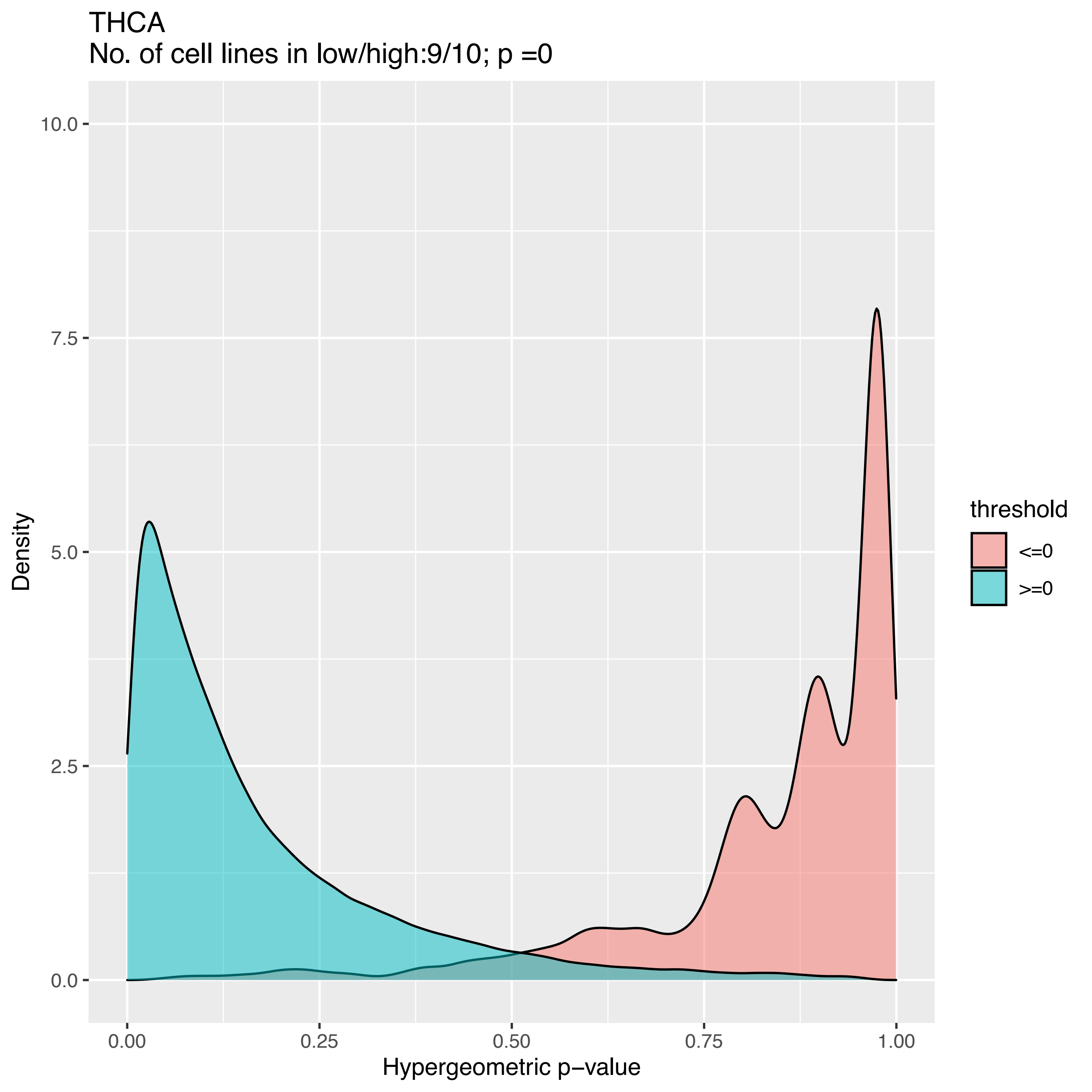
THCA
No. of cell lines in low/high:9/10; p =0
Density
10.0
7.5
5.0
2.5
0.0
0.00
0.25
0.50
0.75
1.00
Hypergeometric p−value
threshold
<=0
>=0

THCA
No. of cell lines in low/high:5/10; p =0

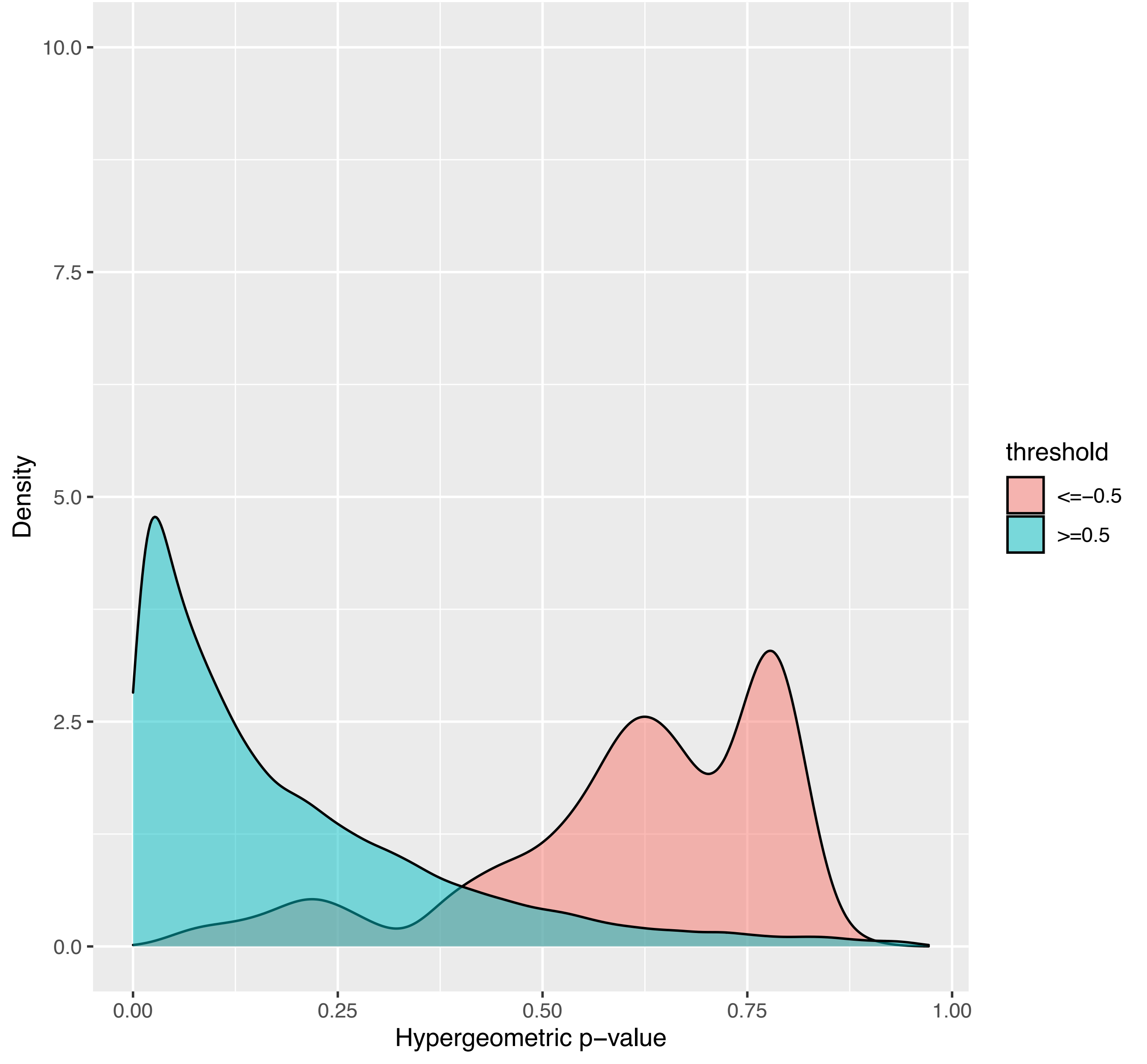

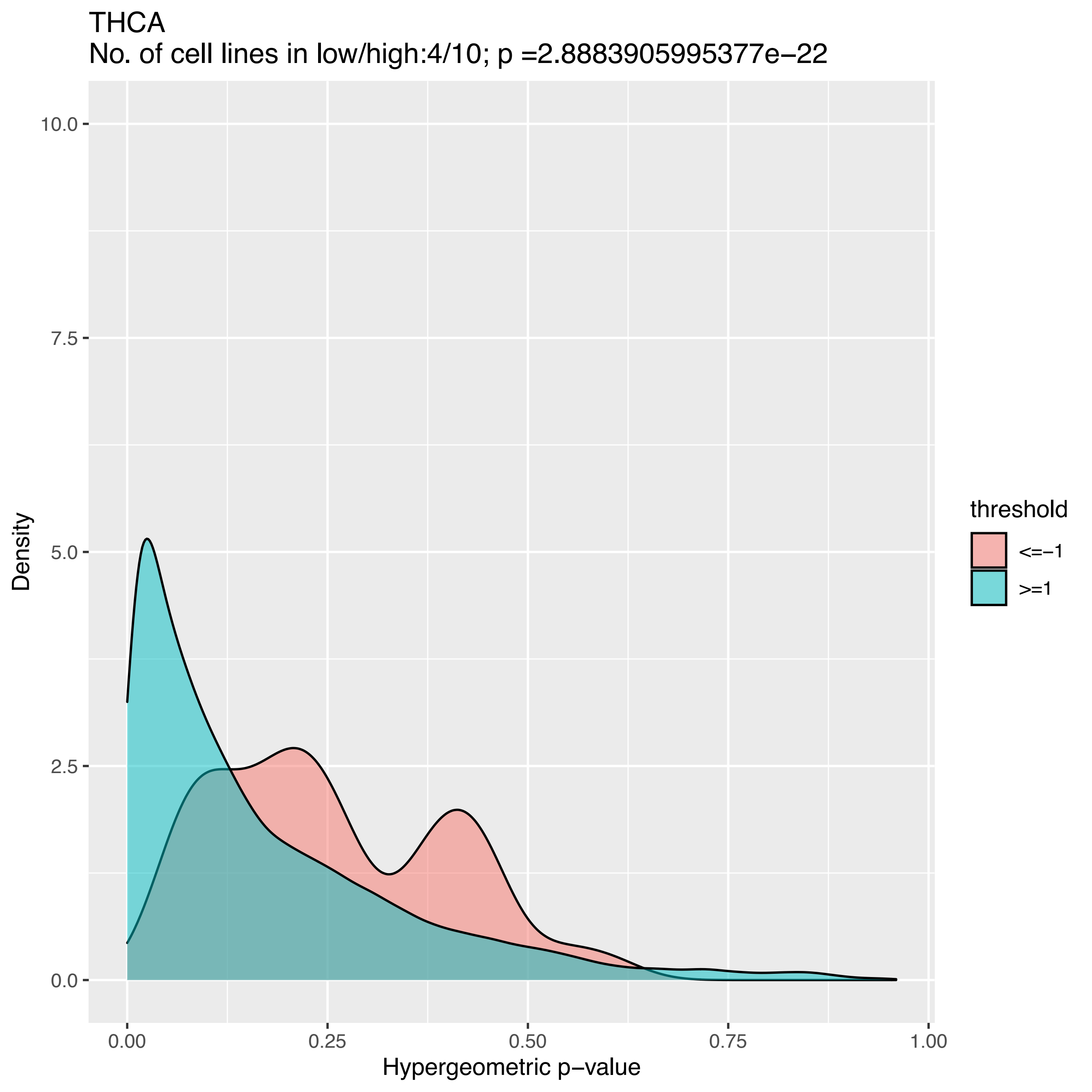

THCA
No. of cell lines in low/high:4/10; p =2.8883905995377e−22
10.0
7.5
5.0
2.5
0.0
Density
0.00
0.25
0.50
0.75
1.00
Hypergeometric p−value
threshold
<=−1
>=1

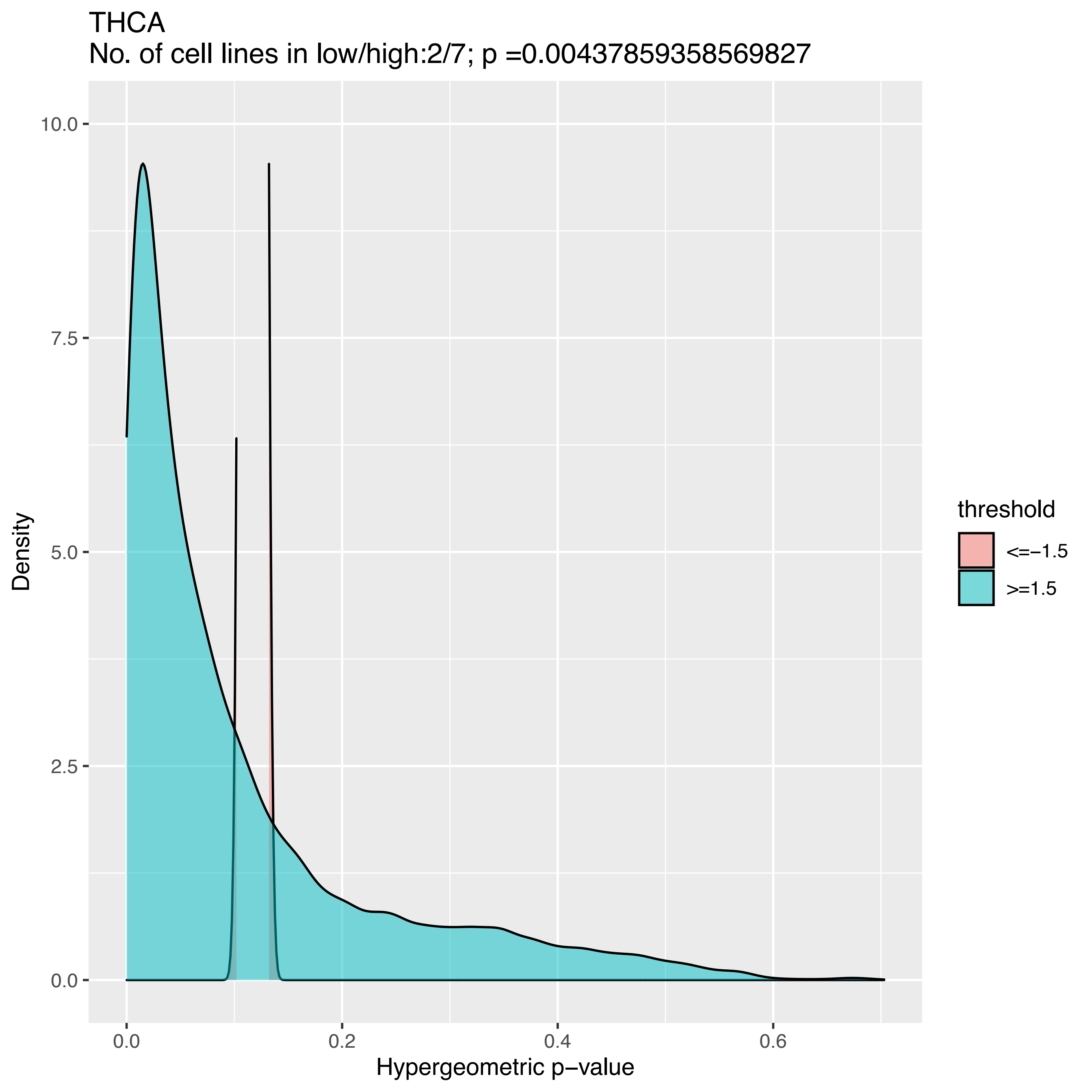
THCA
No. of cell lines in low/high:2/7; p =0.00437859358569827
10.0
7.5
5.0
2.5
0.0
Density
0.0
0.2
0.4
0.6
Hypergeometric p−value
threshold
<=−1.5
>=1.5

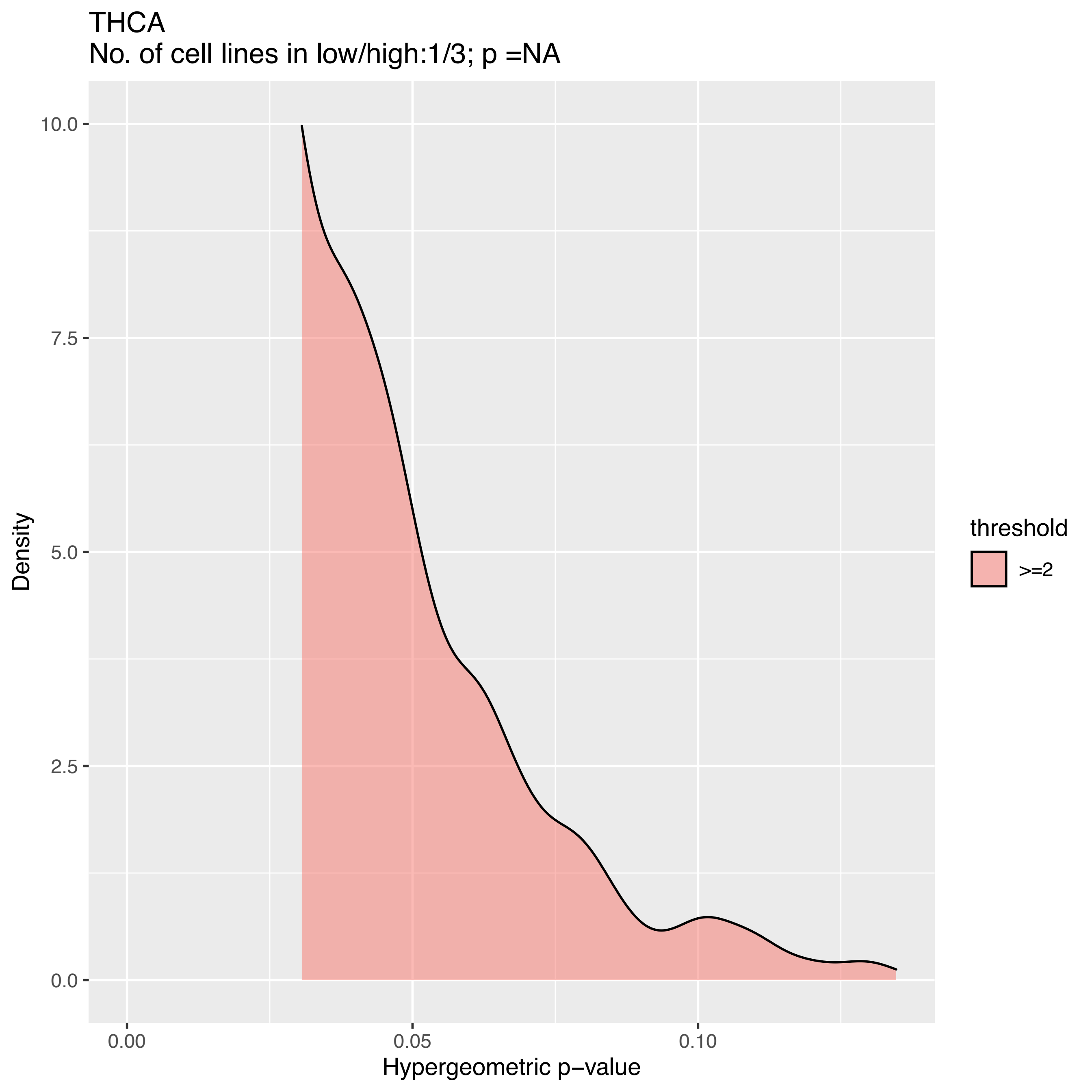
THCA
No. of cell lines in low/high:1/3; p =NA
10.0
7.5
5.0
2.5
0.0
Density
0.00
0.05
0.10
Hypergeometric p−value
threshold
>=2

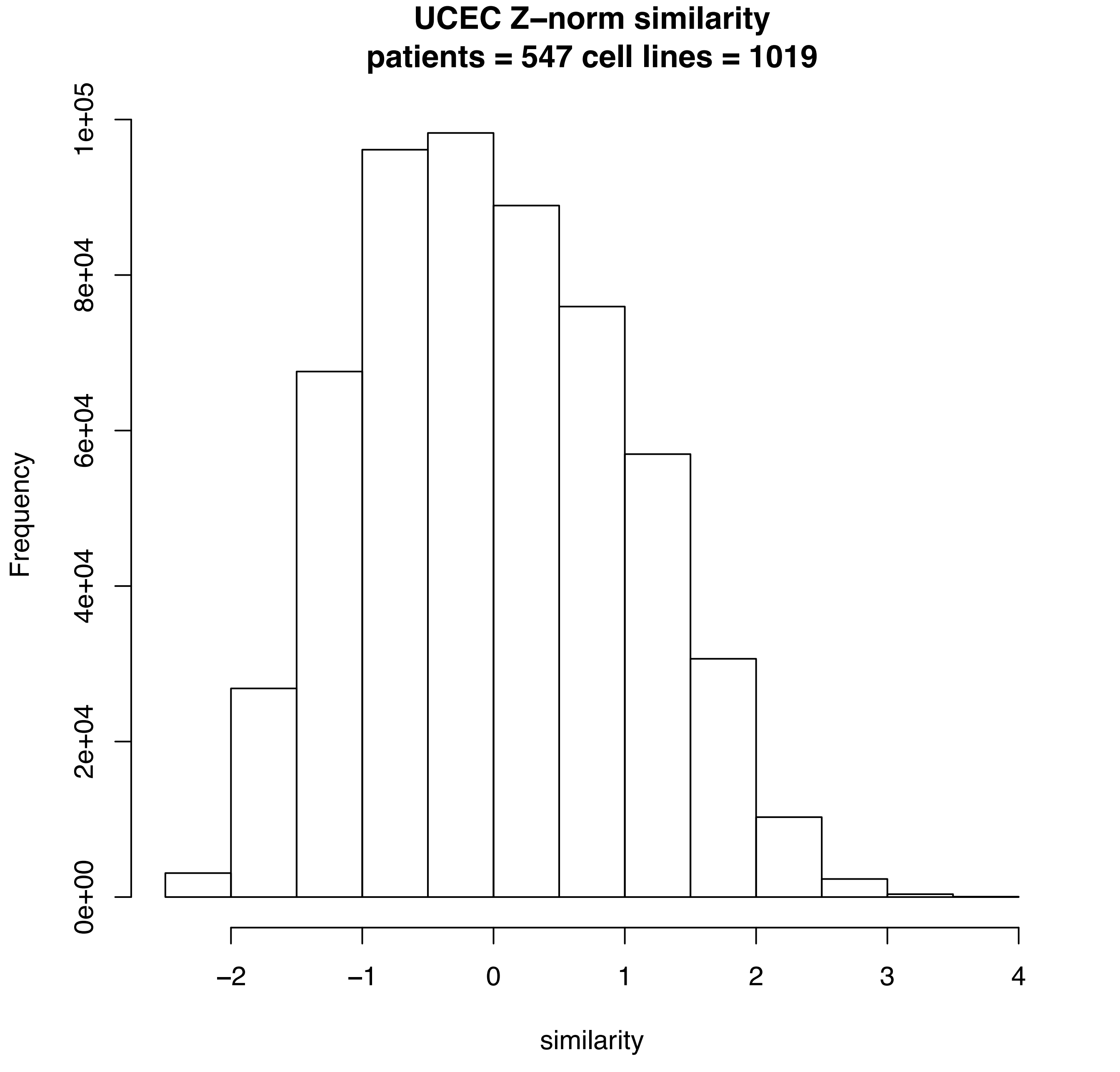
UCEC Z–norm similarity
patients = 547 cell lines = 1019
Frequency
0e+00
2e+04
4e+04
6e+04
8e+04
1e+05
–2
–1
0
1
2
3
4
similarity

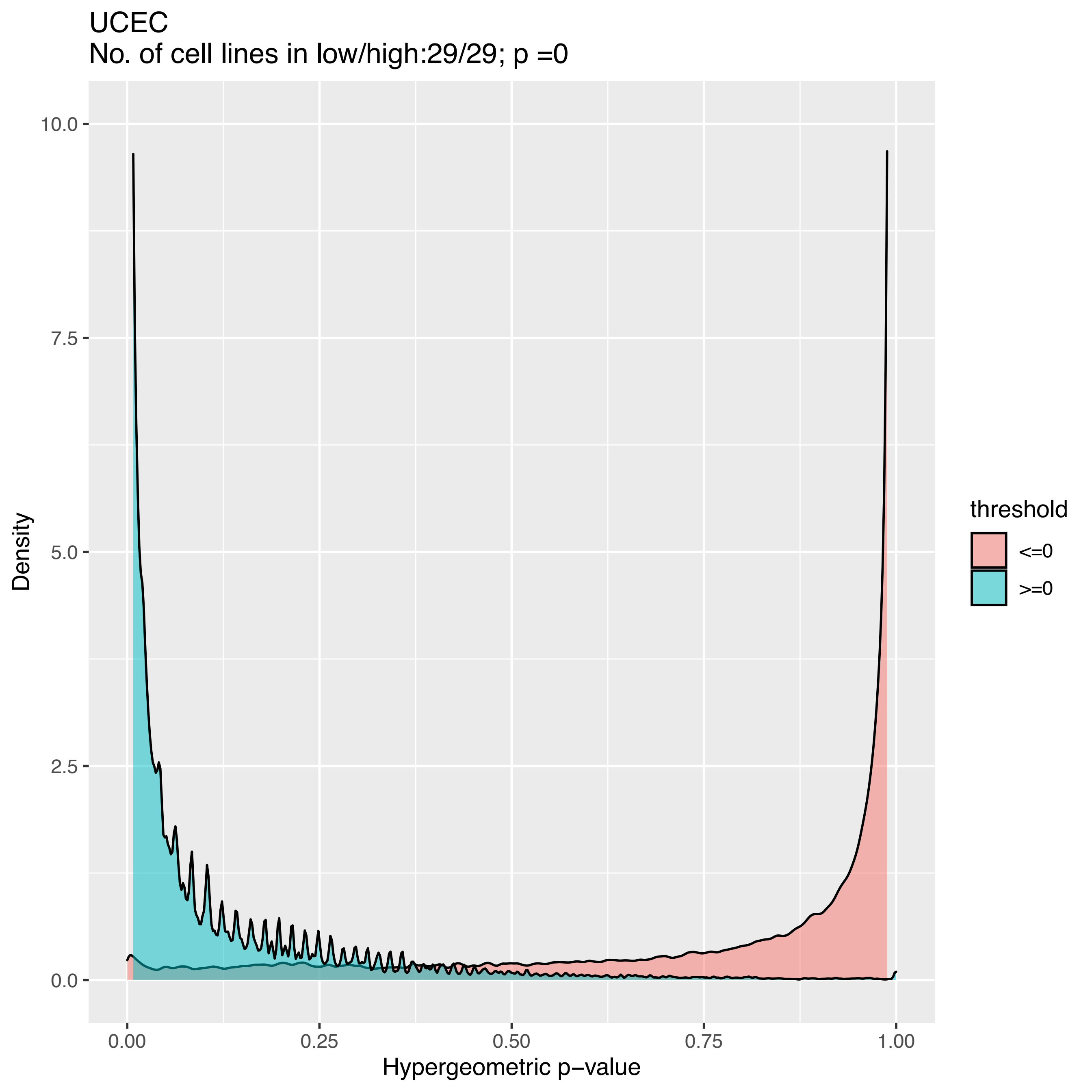

UCEC
No. of cell lines in low/high:29/29; p =0
Density
10.0
7.5
5.0
2.5
0.0
0.00
0.25
0.50
0.75
1.00
Hypergeometric p−value
threshold
<=0
>=0

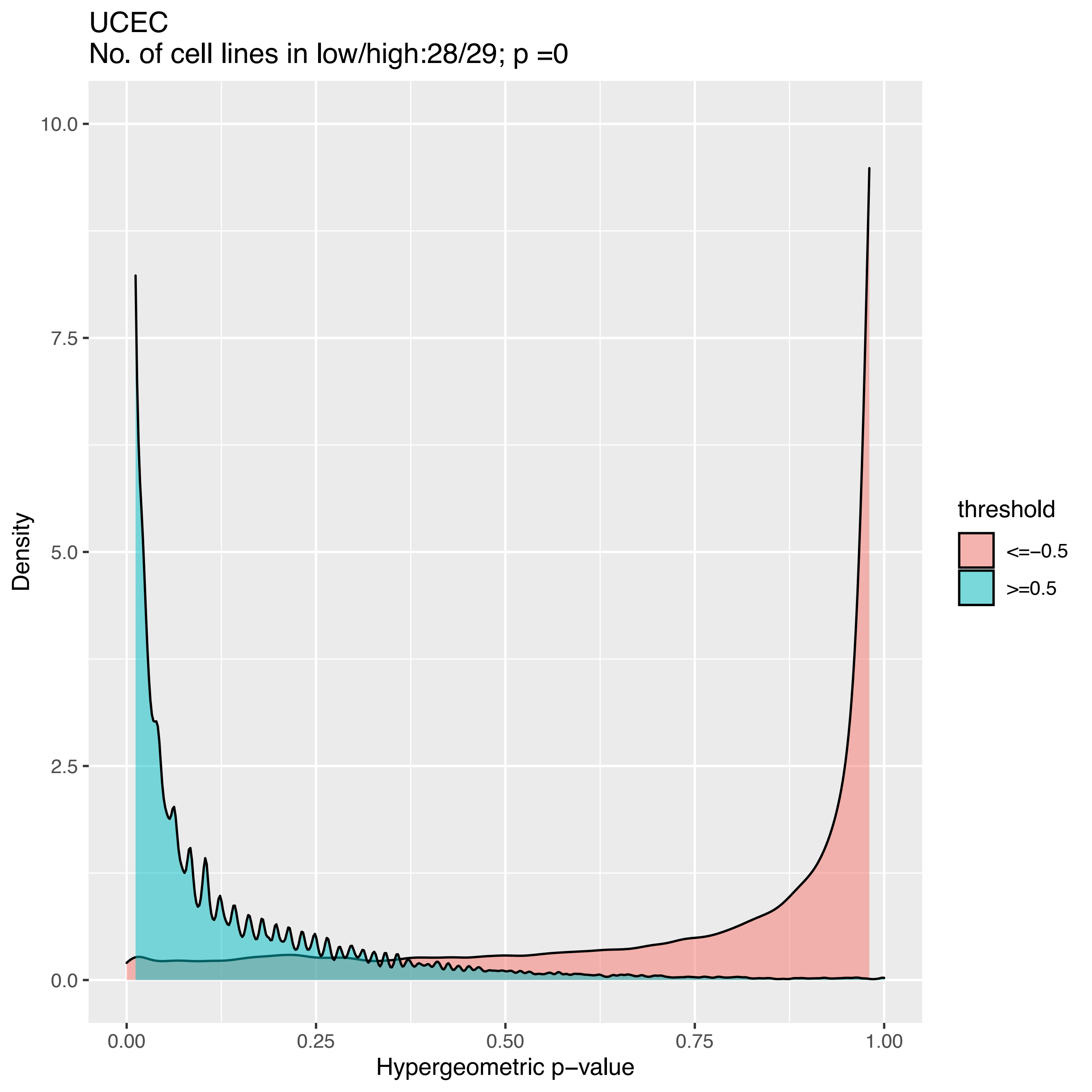
UCEC
No. of cell lines in low/high:28/29; p =0
Density
10.0
7.5
5.0
2.5
0.0
0.00
0.25
0.50
0.75
1.00
Hypergeometric p−value
threshold
<=−0.5
>=0.5

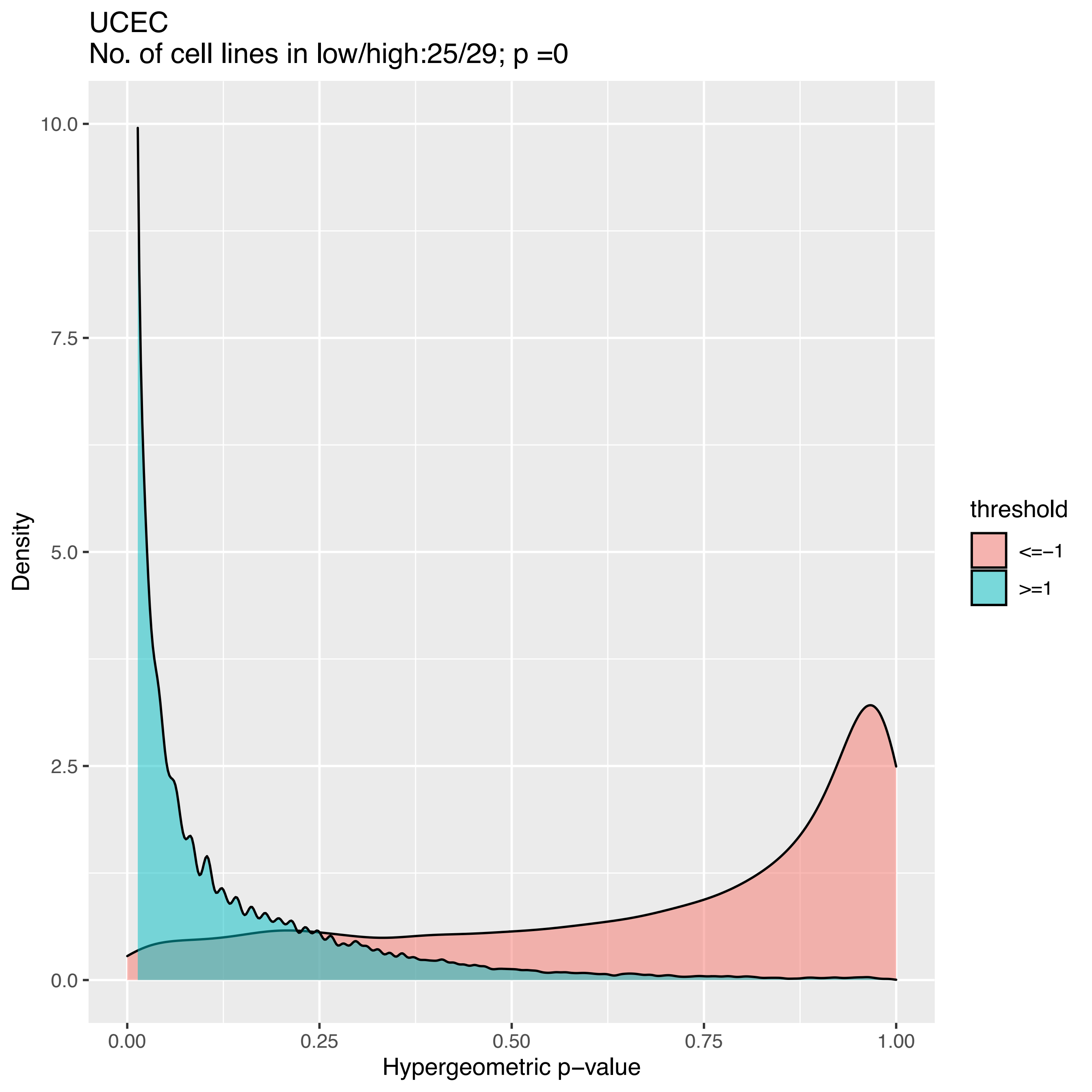
UCEC
No. of cell lines in low/high:25/29; p =0
Density
10.0
7.5
5.0
2.5
0.0
0.00
0.25
0.50
0.75
1.00
Hypergeometric p−value
threshold
<=−1
>=1

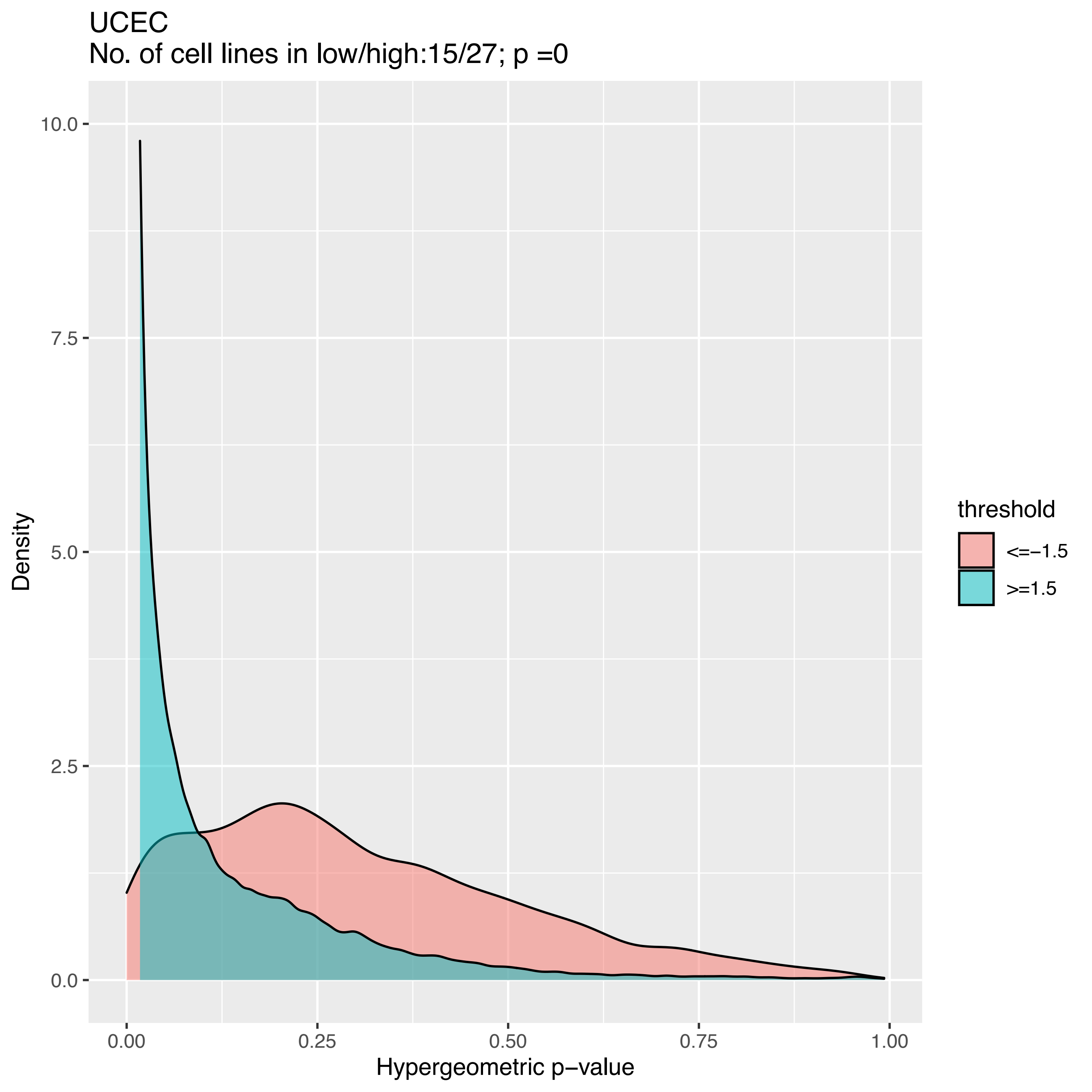
UCEC
No. of cell lines in low/high:15/27; p =0
Density
Hypergeometric p−value
0.00
0.25
0.50
0.75
1.00
0.0
2.5
5.0
7.5
10.0
threshold
<=−1.5
>=1.5

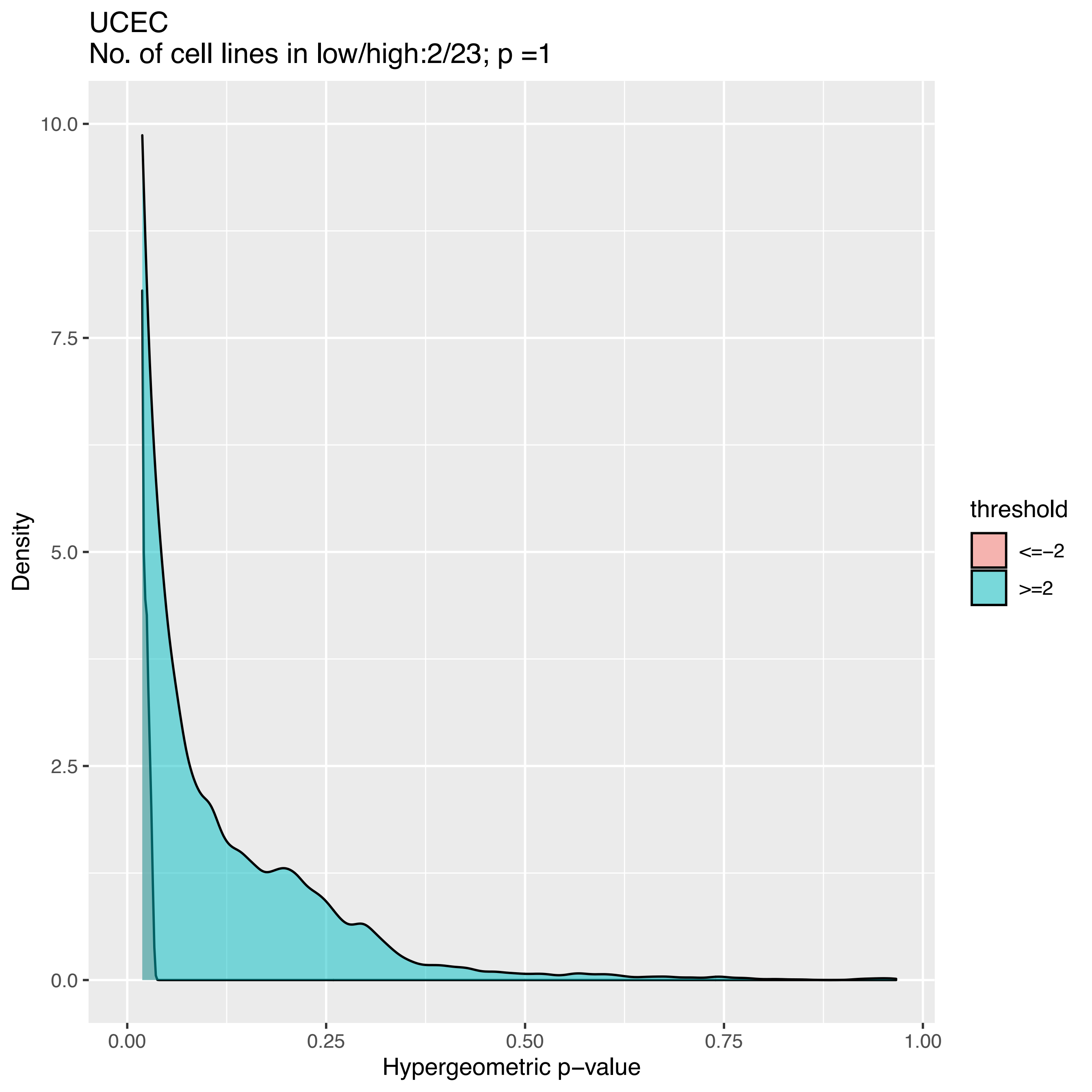

UCEC
No. of cell lines in low/high:2/23; p =1
10.0
7.5
5.0
2.5
0.0
Density
0.00
0.25
0.50
0.75
1.00
Hypergeometric p−value
threshold
<=−2
>=2

### Celligner

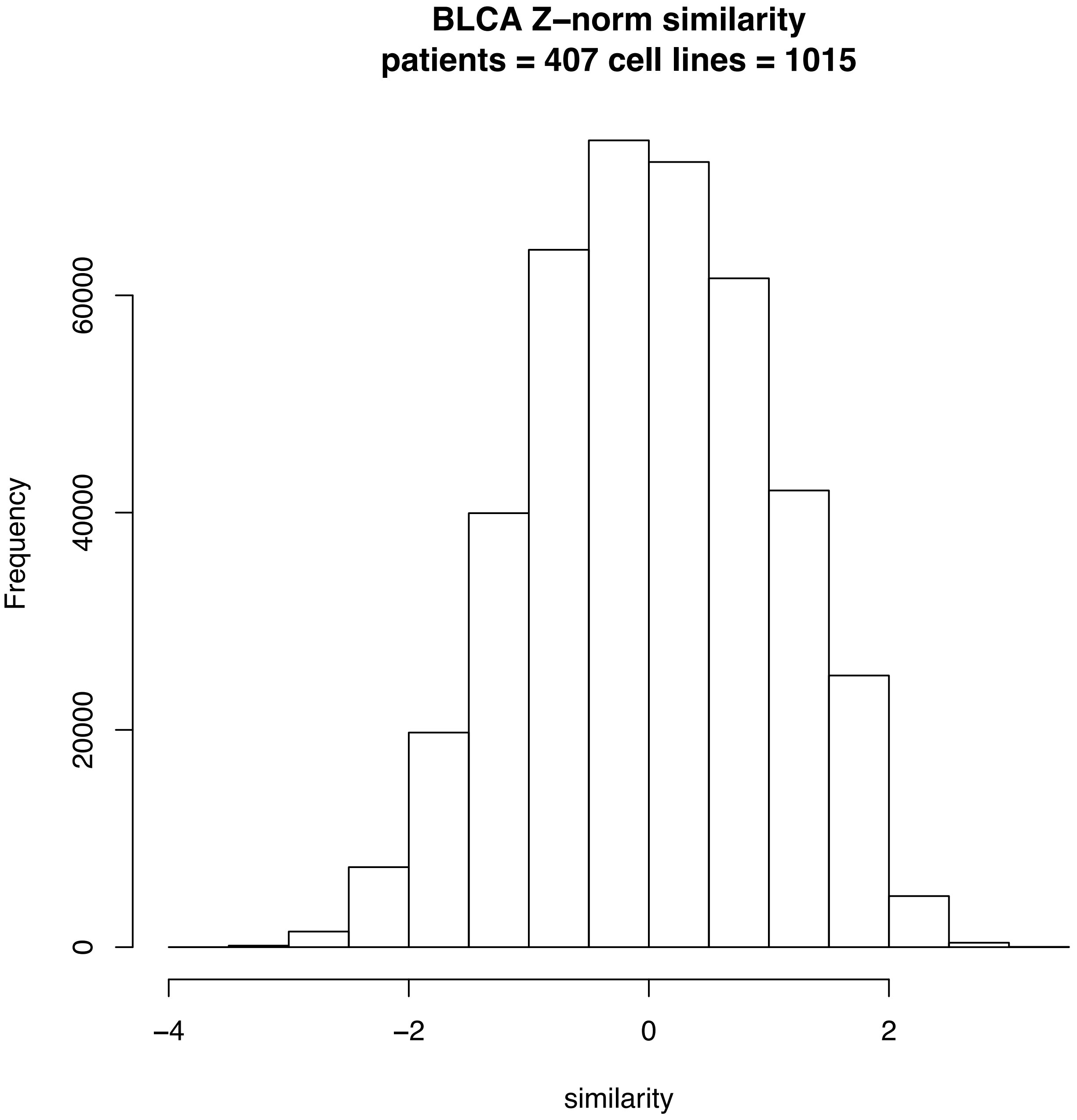

BLCA Z–norm similarity
patients = 407 cell lines = 1015
Frequency
0
20000
40000
60000
–4
–2
0
2
similarity

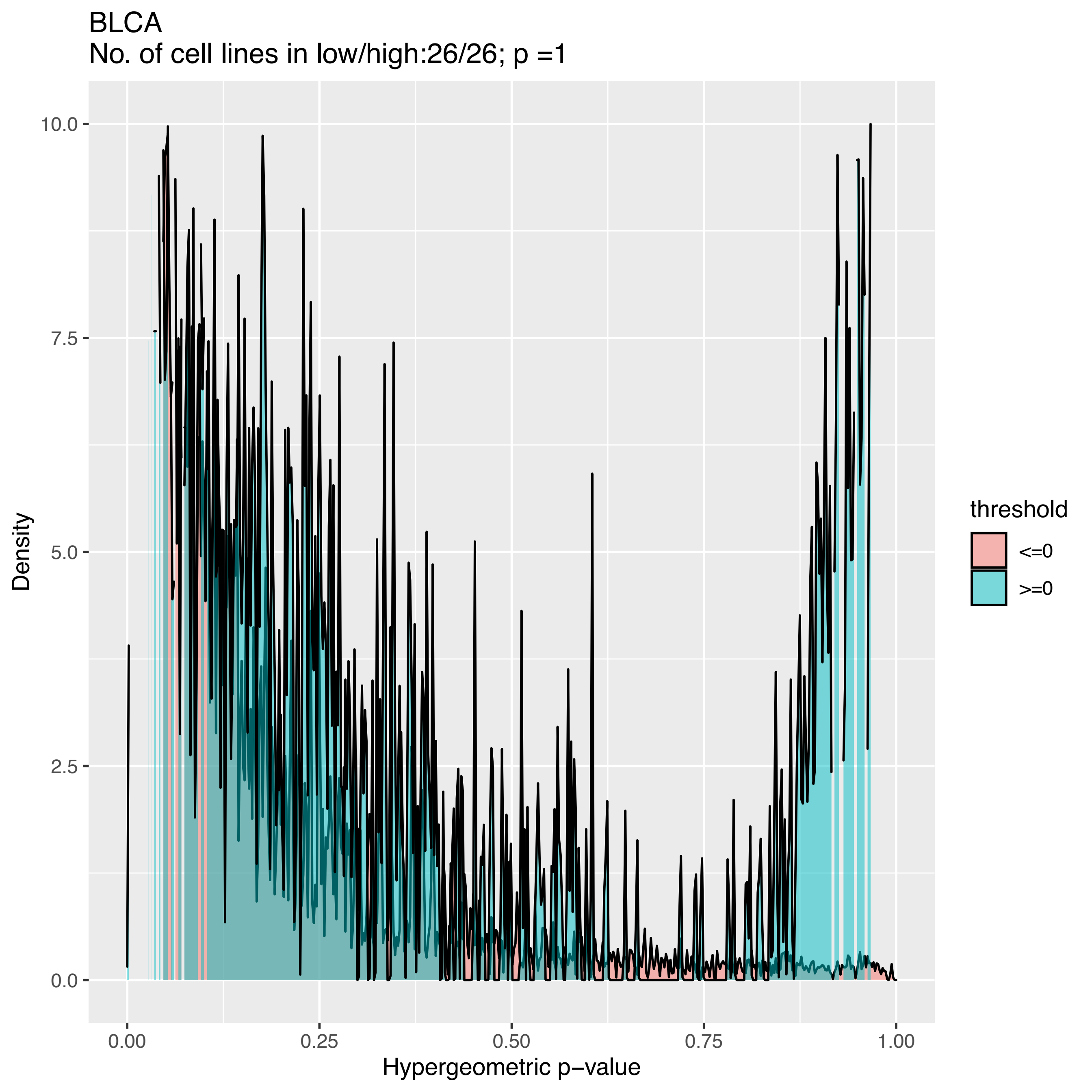

BLCA
No. of cell lines in low/high:26/26; p =1
Density
10.0
7.5
5.0
2.5
0.0
0.00
0.25
0.50
0.75
1.00
Hypergeometric p−value
threshold
<=0
>=0

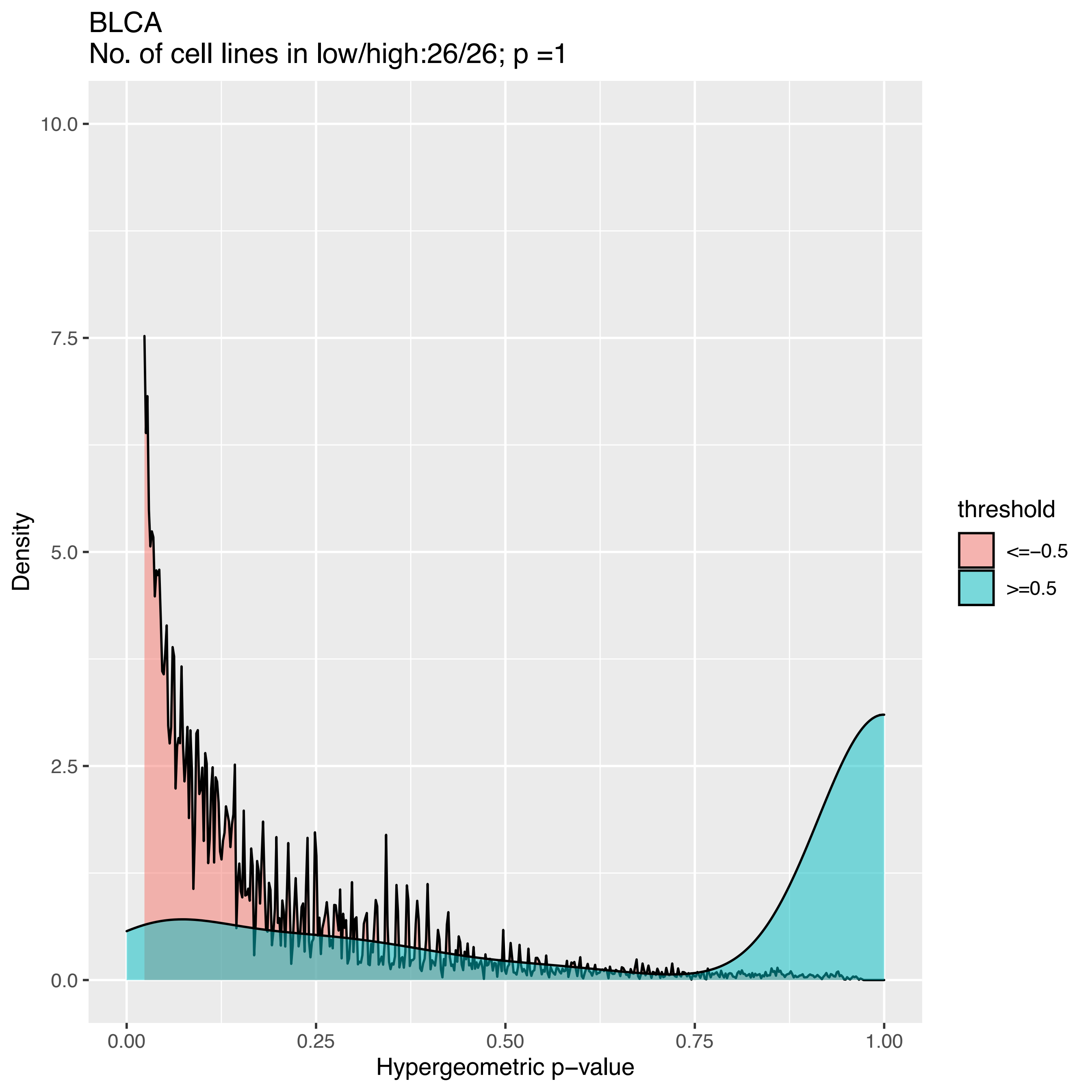
BLCA
No. of cell lines in low/high:26/26; p =1
Density
10.0
7.5
5.0
2.5
0.0
0.00
0.25
0.50
0.75
1.00
Hypergeometric p−value
threshold
<=−0.5
>=0.5

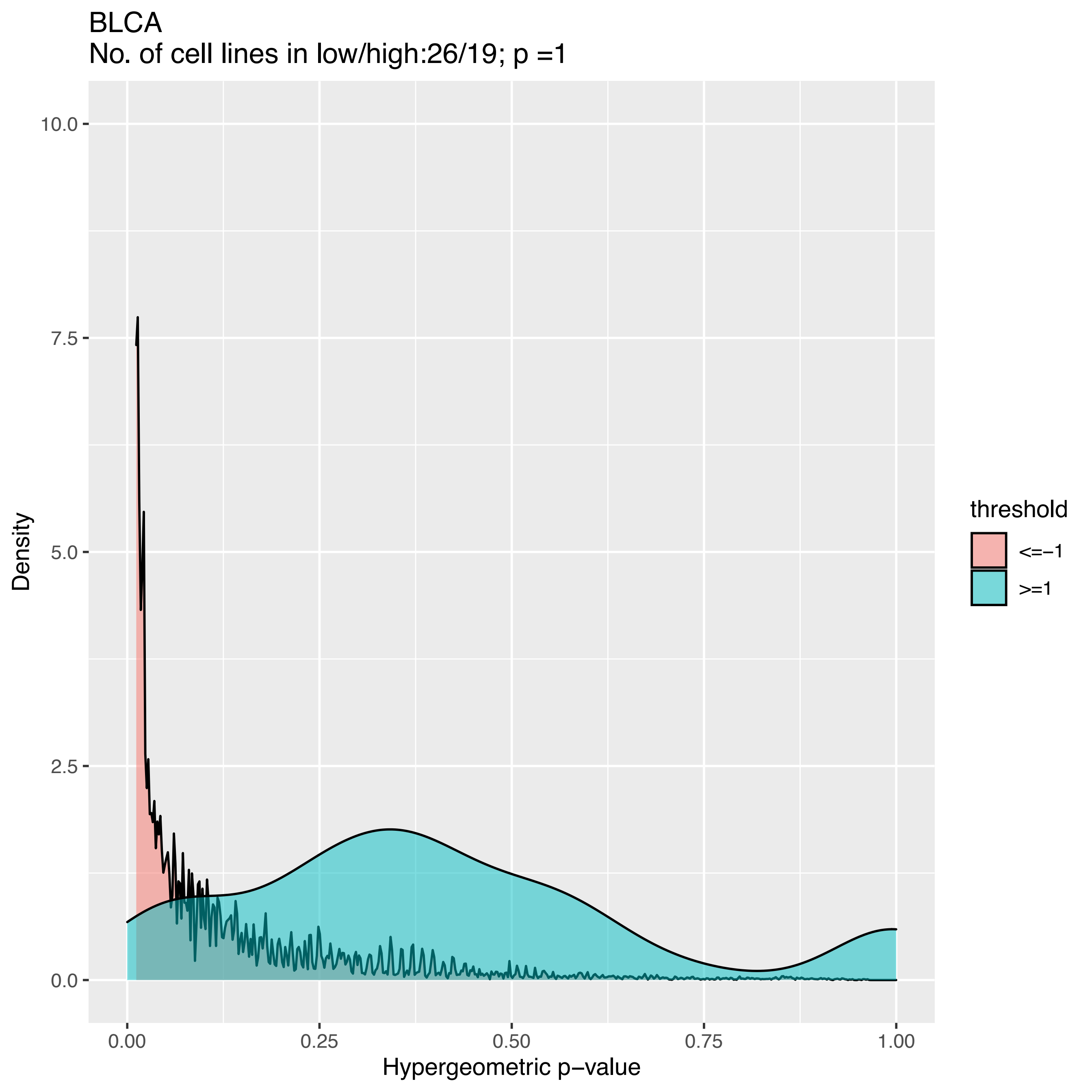
BLCA
No. of cell lines in low/high:26/19; p =1
10.0
7.5
5.0
2.5
0.0
Density
0.00
0.25
0.50
0.75
1.00
Hypergeometric p−value
threshold
<=−1
>=1

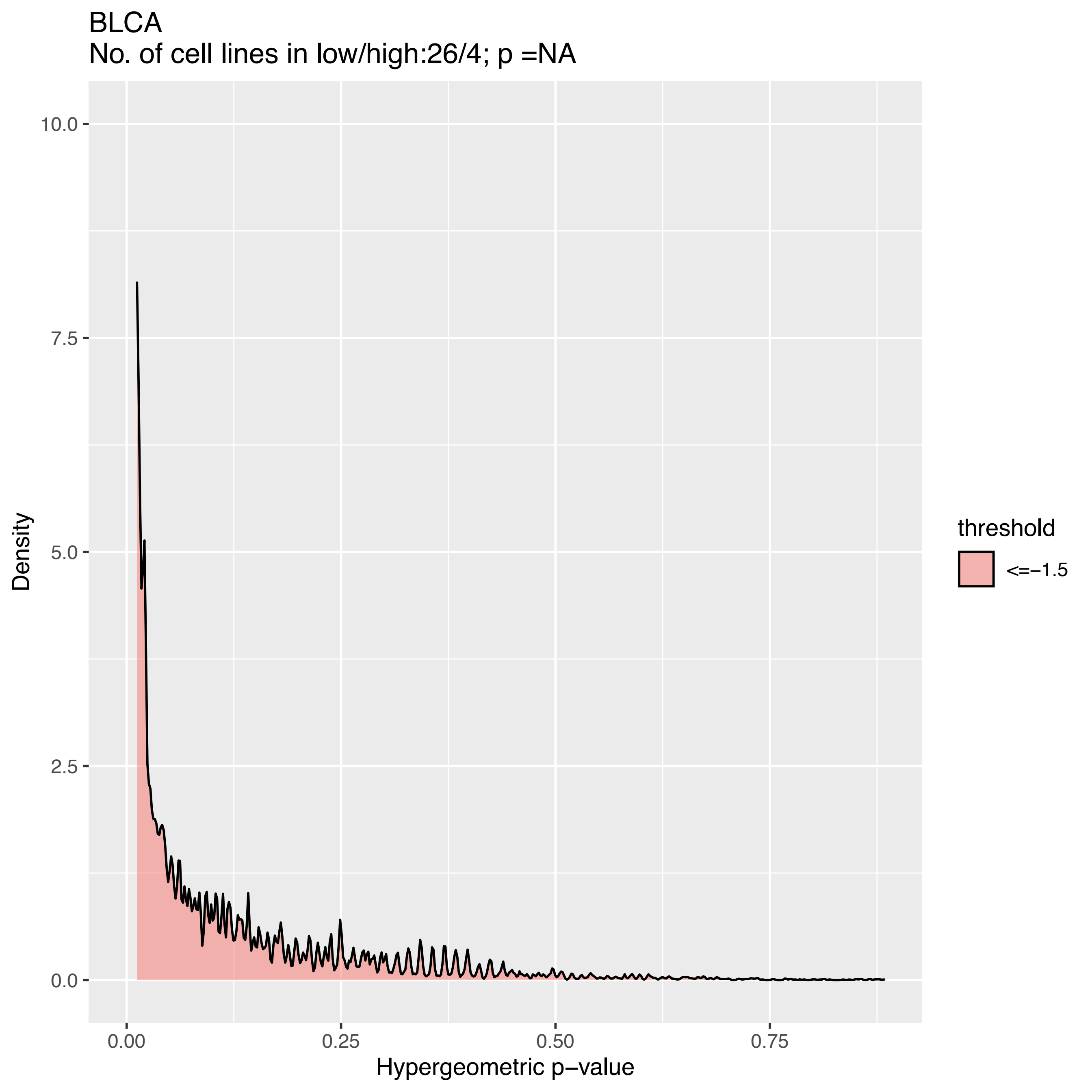

BLCA
No. of cell lines in low/high:26/4; p =NA
10.0
7.5
5.0
2.5
0.0
Density
0.00
0.25
0.50
0.75
Hypergeometric p−value
threshold
<=−1.5

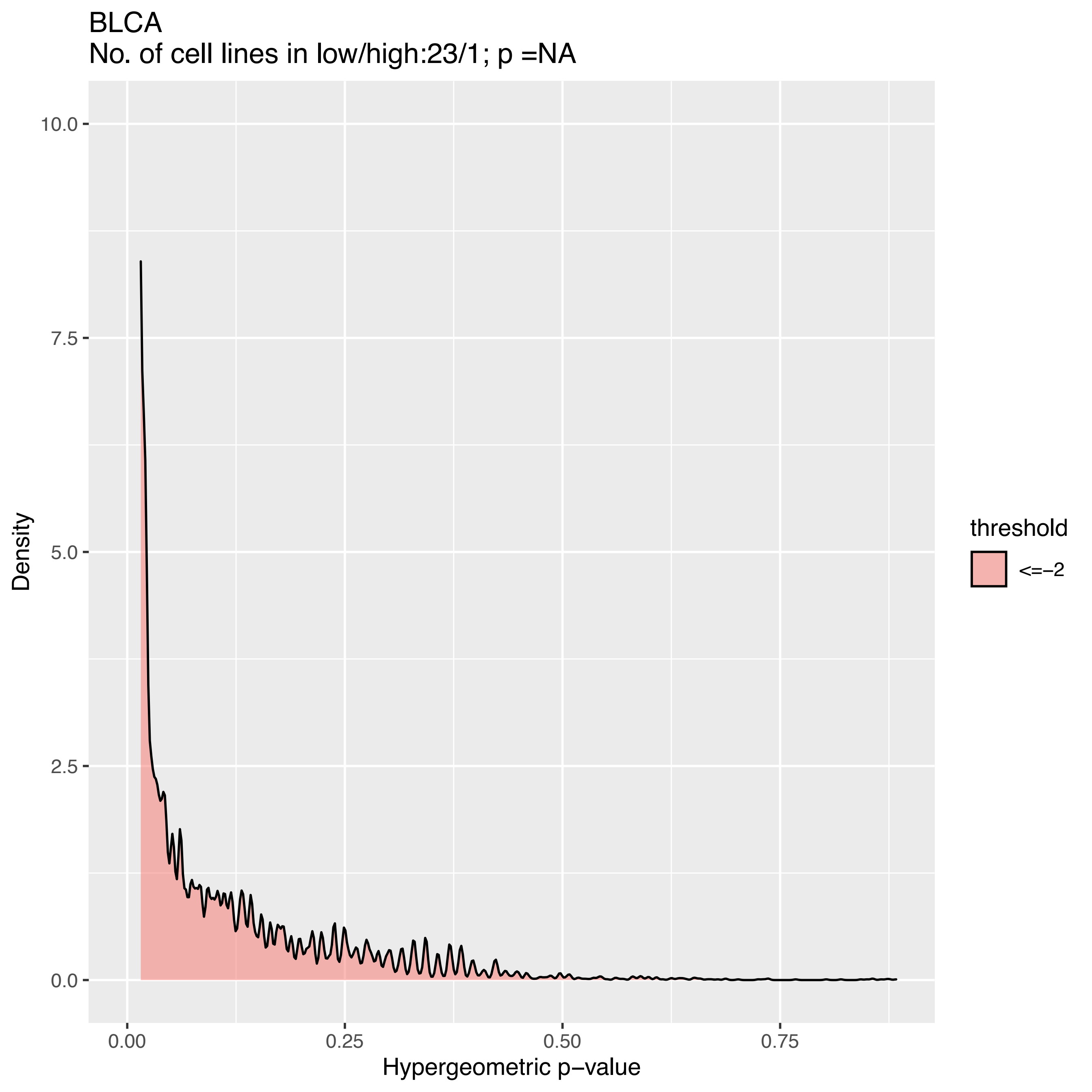
BLCA
No. of cell lines in low/high:23/1; p =NA
Density
10.0
7.5
5.0
2.5
0.0
0.00
0.25
0.50
0.75
Hypergeometric p−value
threshold
<=−2

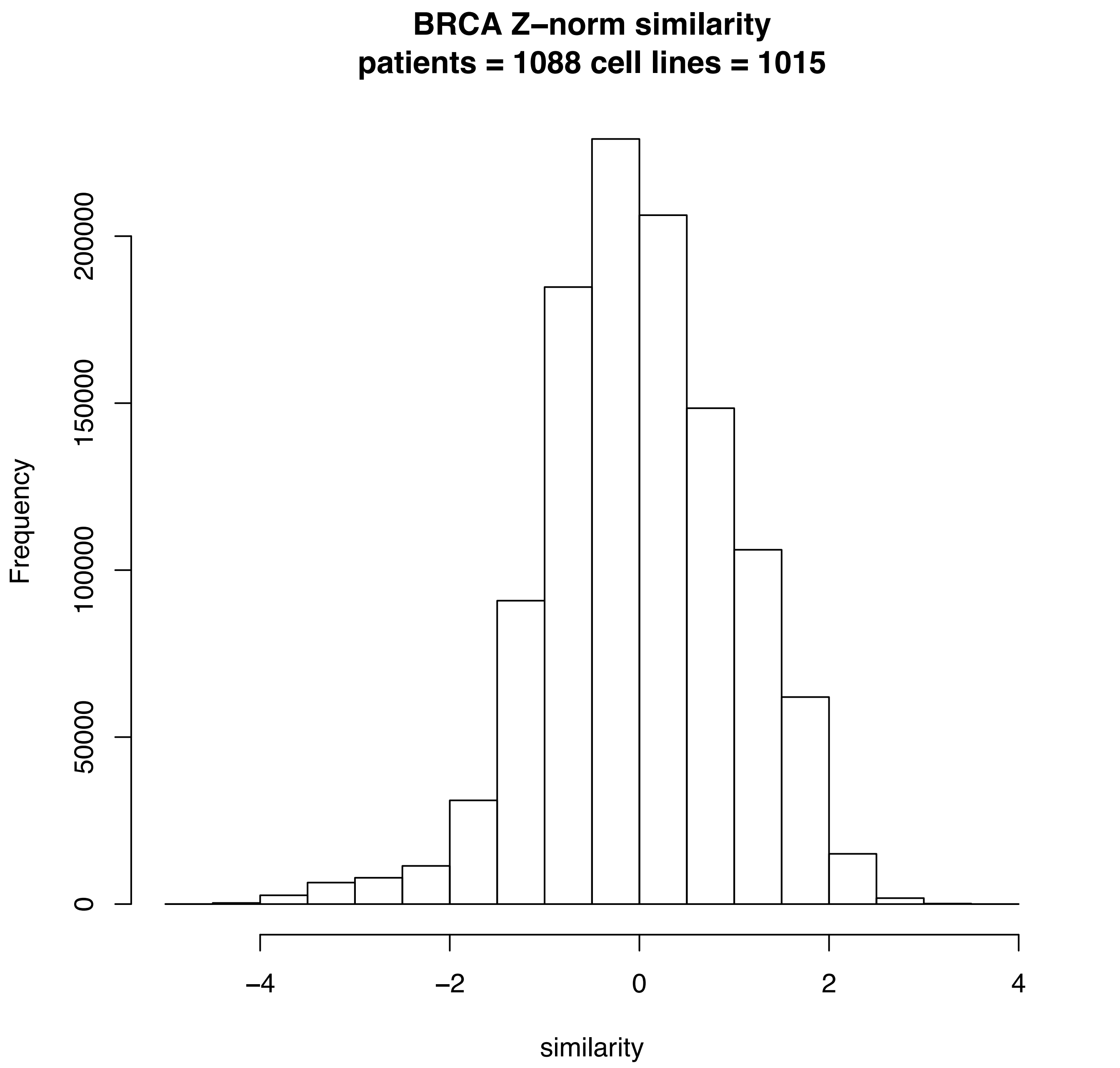
BRCA Z–norm similarity
patients = 1088 cell lines = 1015
Frequency
0
50000
100000
150000
200000
–4
–2
0
2
4
similarity

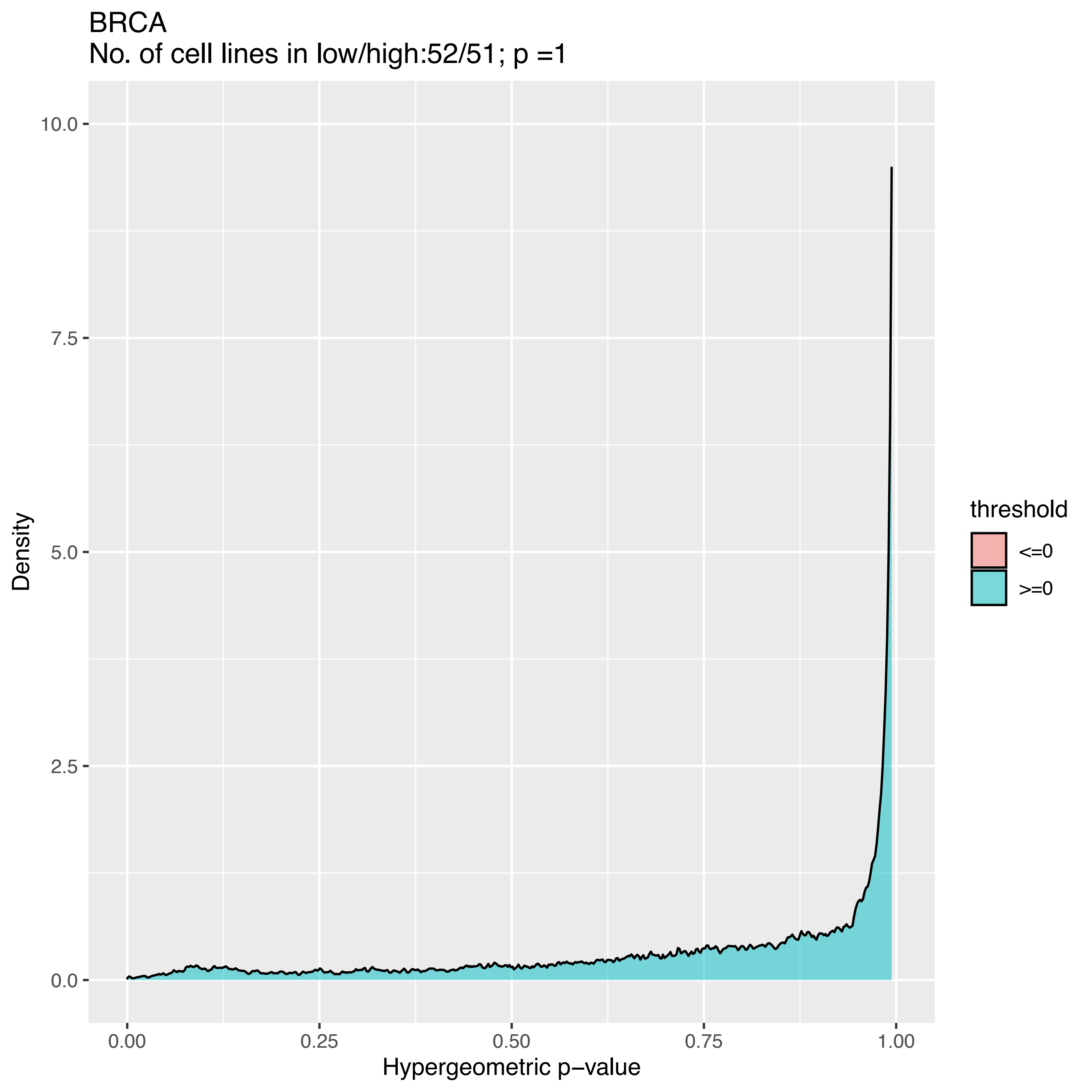
BRCA
No. of cell lines in low/high:52/51; p =1
10.0
7.5
5.0
2.5
0.0
Density
0.00
0.25
0.50
0.75
1.00
Hypergeometric p−value
threshold
<=0
>=0

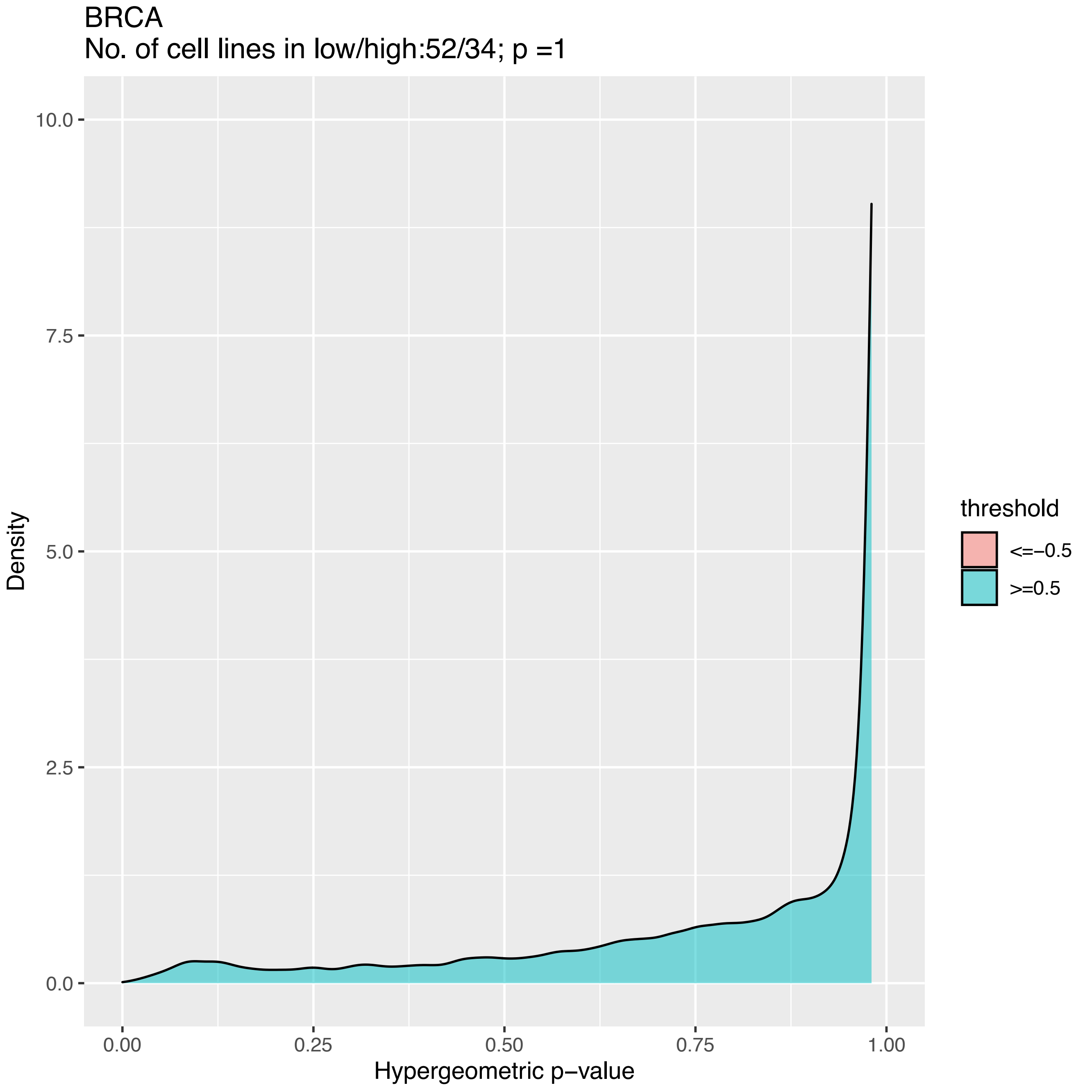
BRCA
No. of cell lines in low/high:52/34; p =1
Density
10.0
7.5
5.0
2.5
0.0
0.00
0.25
0.50
0.75
1.00
Hypergeometric p−value
threshold
<=−0.5
>=0.5

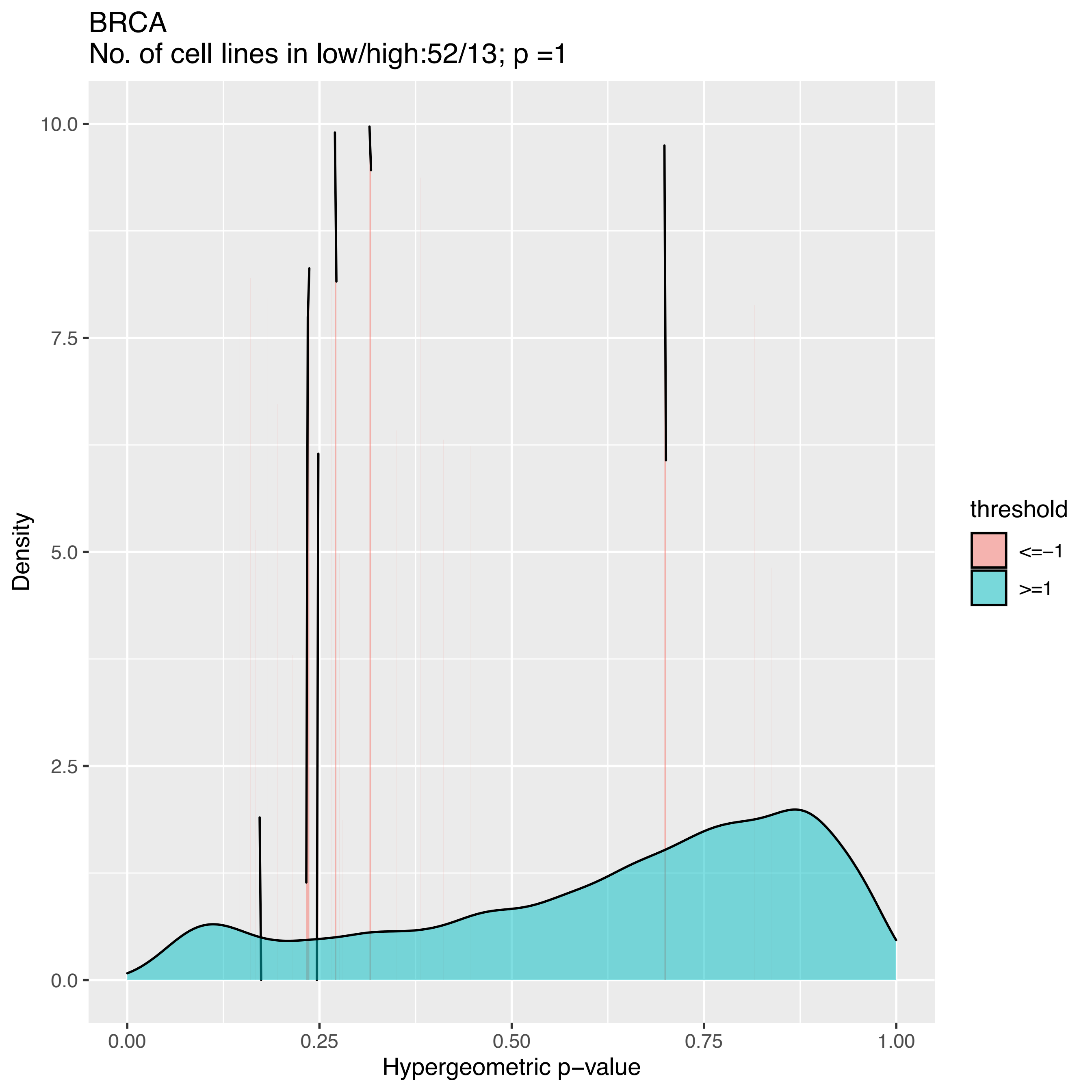

BRCA
No. of cell lines in low/high:52/13; p =1
Density
Hypergeometric p−value
0.0
2.5
5.0
7.5
10.0
0.00
0.25
0.50
0.75
1.00
threshold
<=−1
>=1

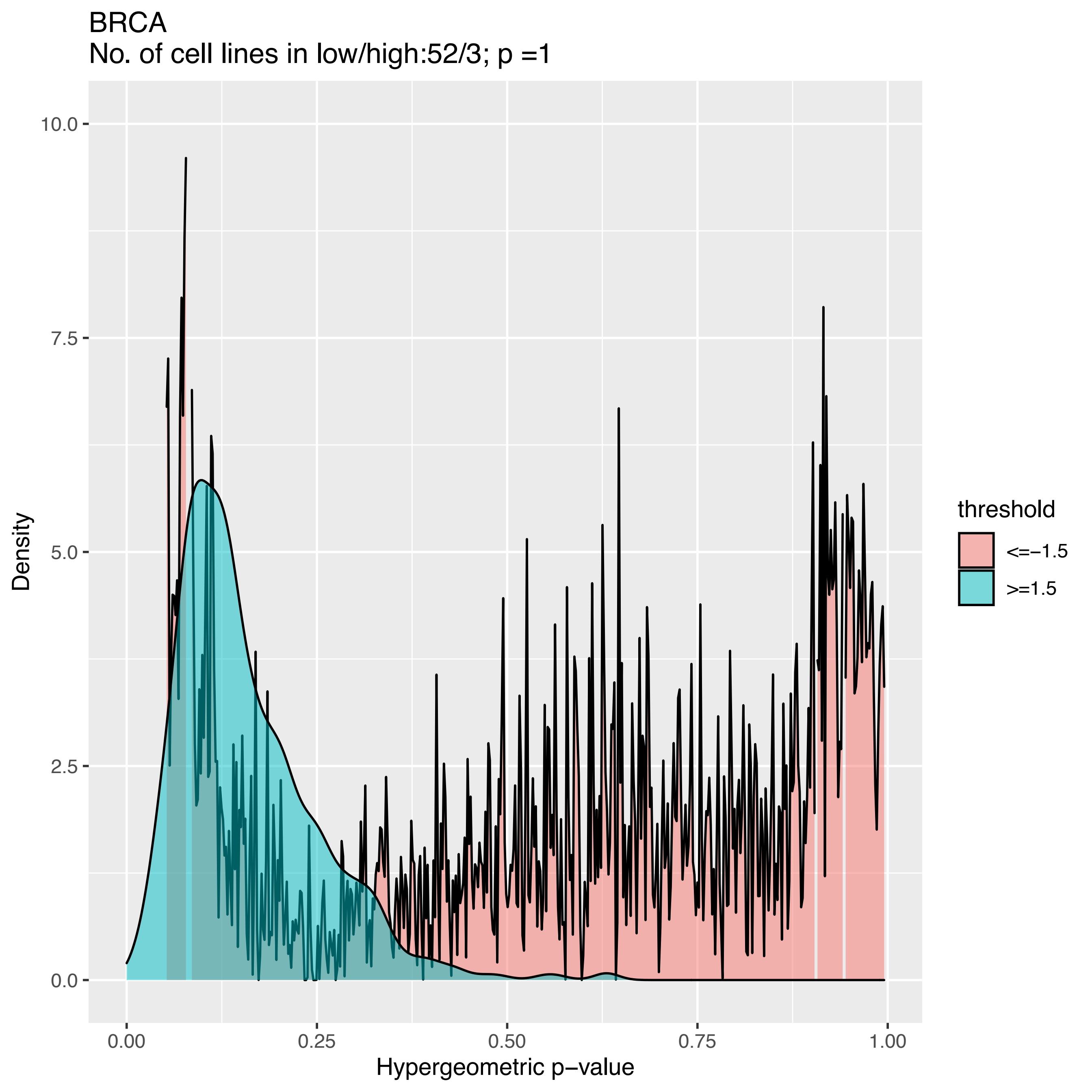

BRCA
No. of cell lines in low/high:52/3; p =1
Density
10.0
7.5
5.0
2.5
0.0
0.00
0.25
0.50
0.75
1.00
Hypergeometric p−value
threshold
<=−1.5
>=1.5

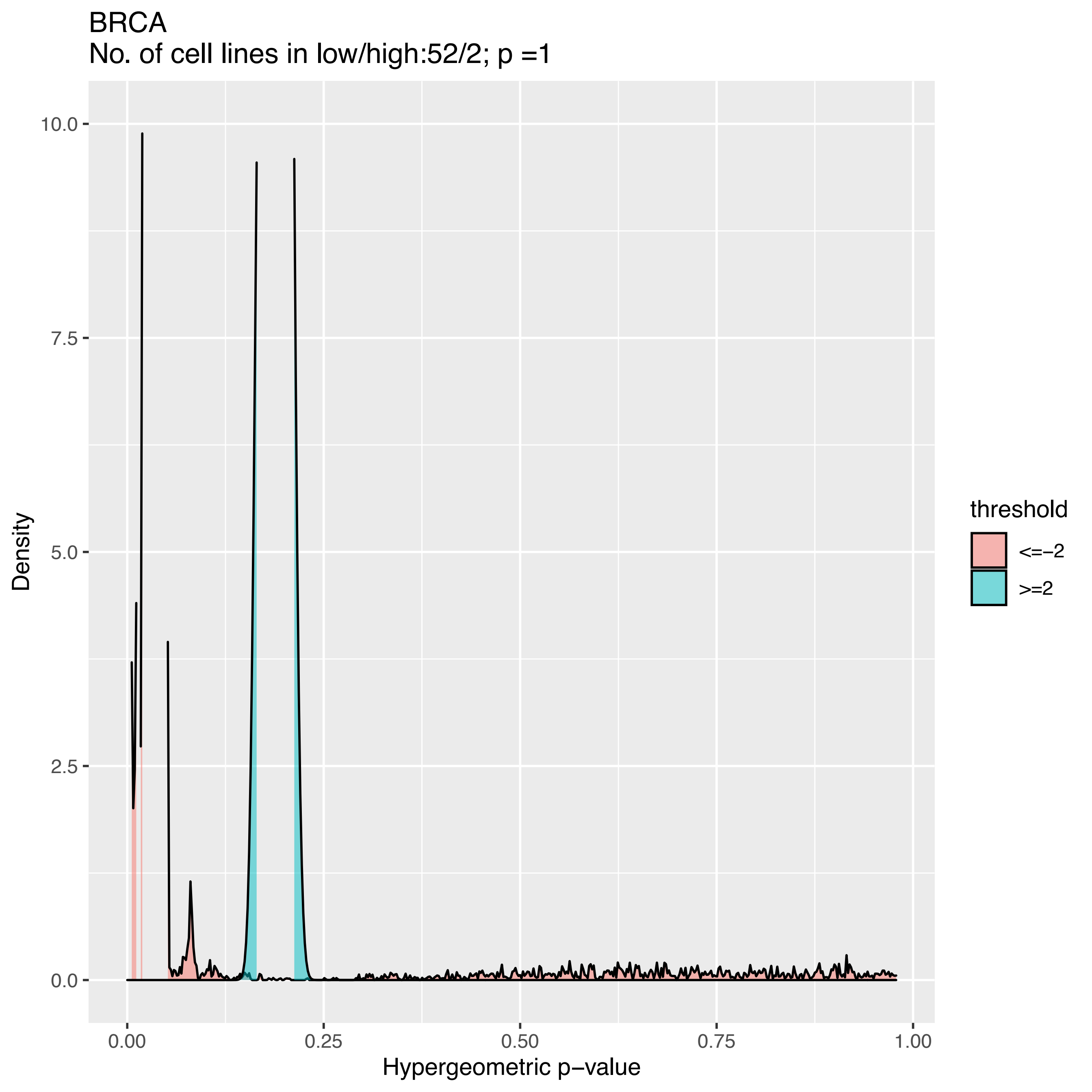
BRCA
No. of cell lines in low/high:52/2; p =1
Density
10.0
7.5
5.0
2.5
0.0
0.00
0.25
0.50
0.75
1.00
Hypergeometric p−value
threshold
<=−2
>=2

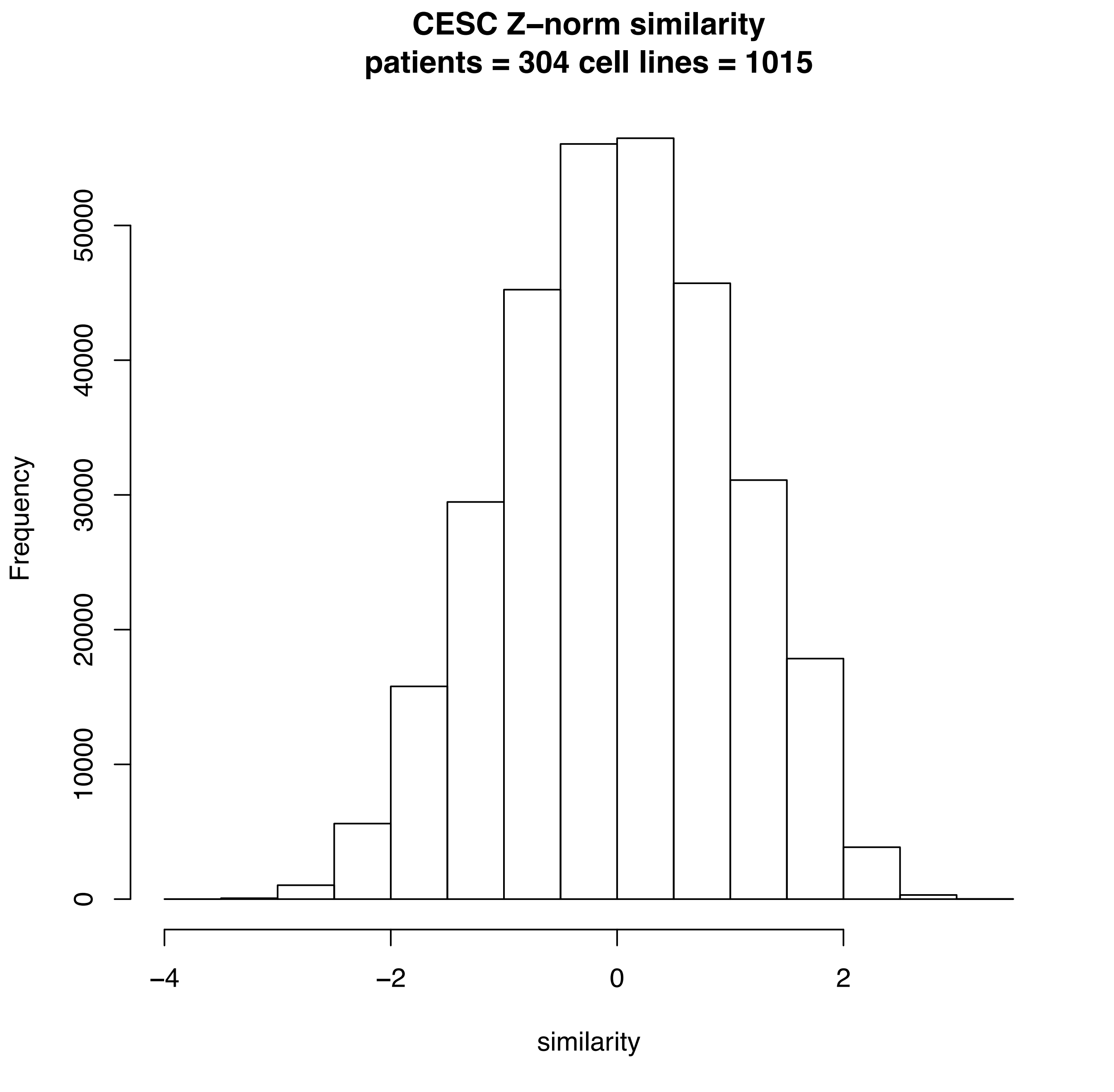

CESC Z–norm similarity
patients = 304 cell lines = 1015
Frequency
0
10000
20000
30000
40000
50000
–4
–2
0
2
similarity

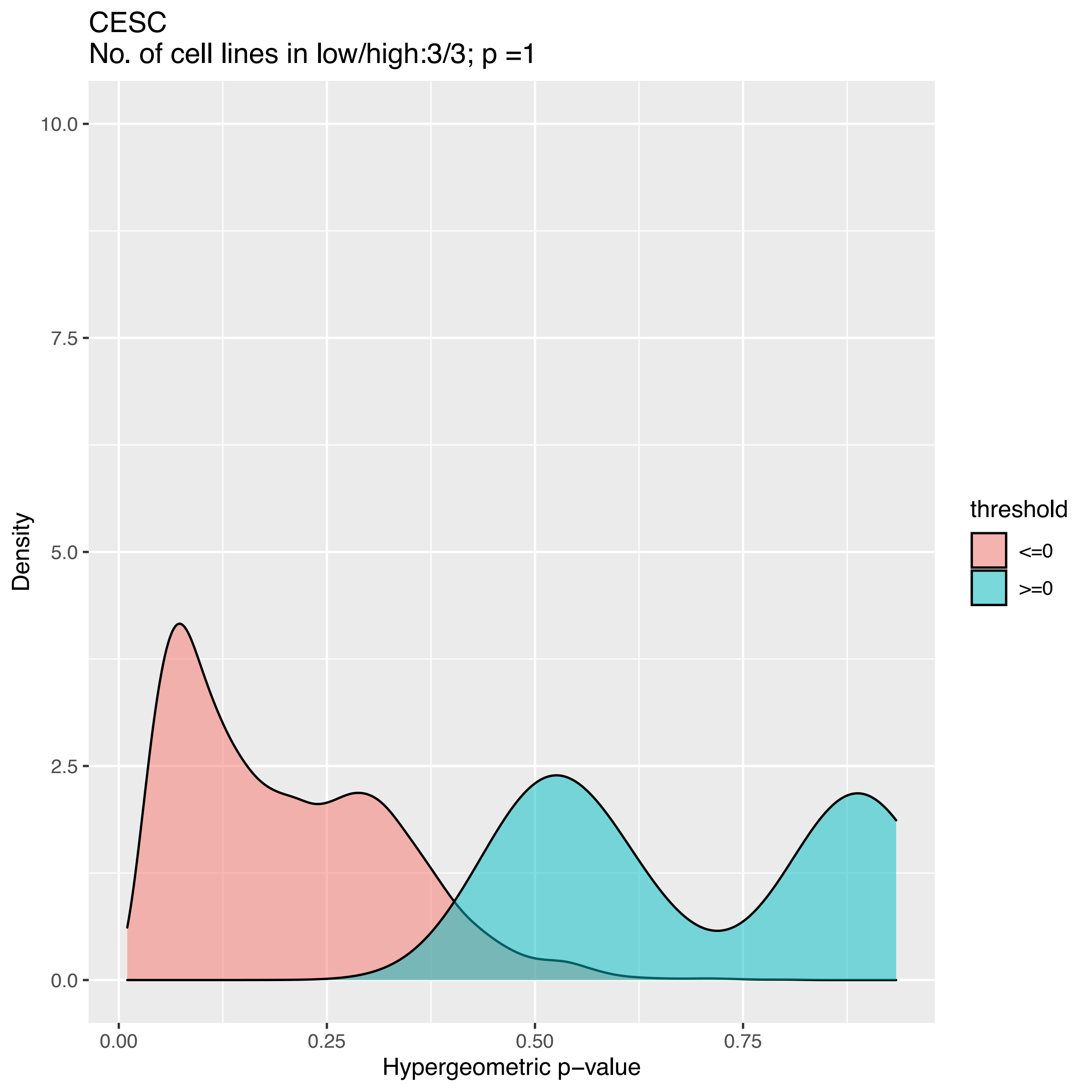
CESC
No. of cell lines in low/high:3/3; p =1
10.0
7.5
5.0
2.5
0.0
Density
0.00
0.25
0.50
0.75
Hypergeometric p−value
threshold
<=0
>=0

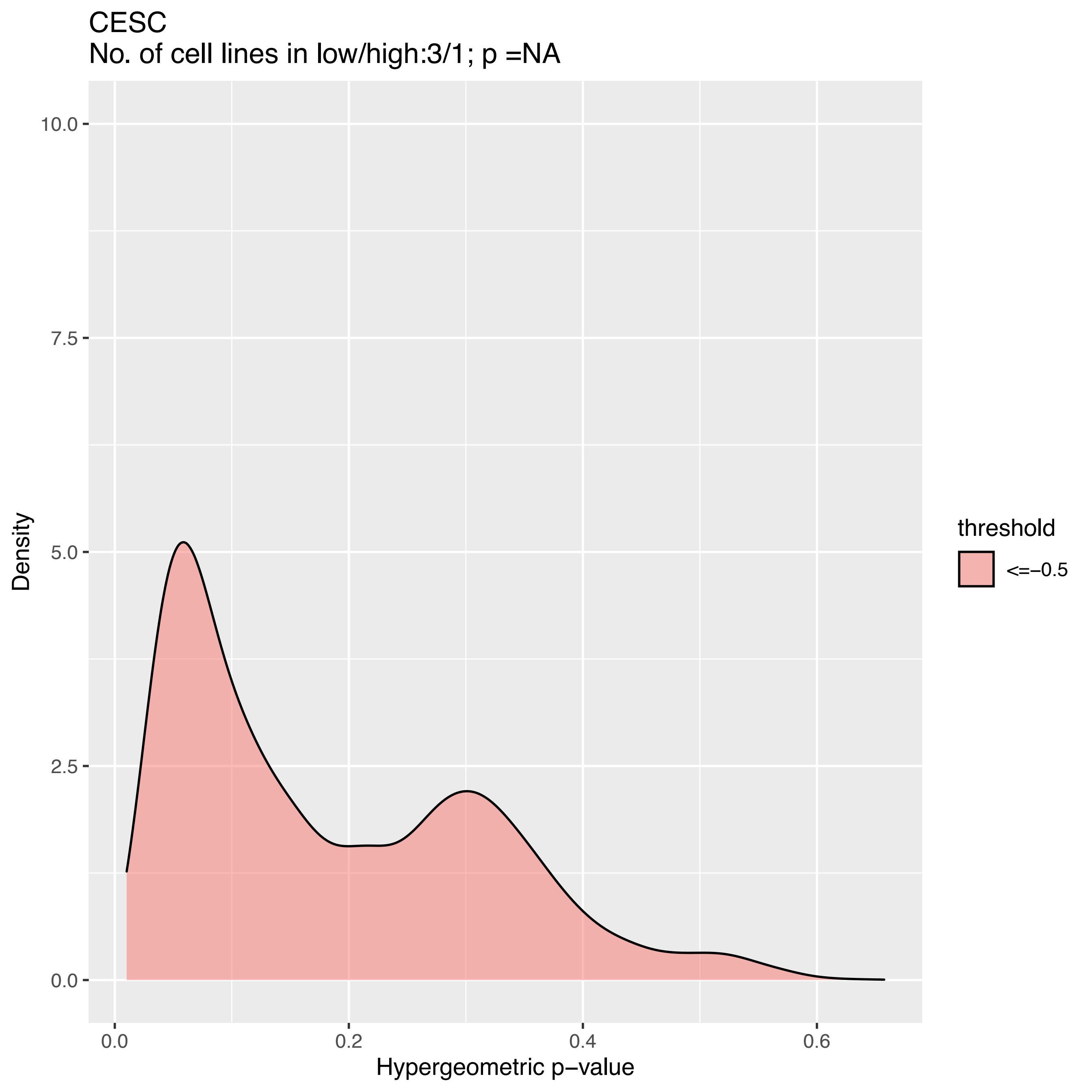
CESC
No. of cell lines in low/high:3/1; p =NA
Density
Hypergeometric p−value
10.0
7.5
5.0
2.5
0.0
0.0
0.2
0.4
0.6
threshold
<=−0.5

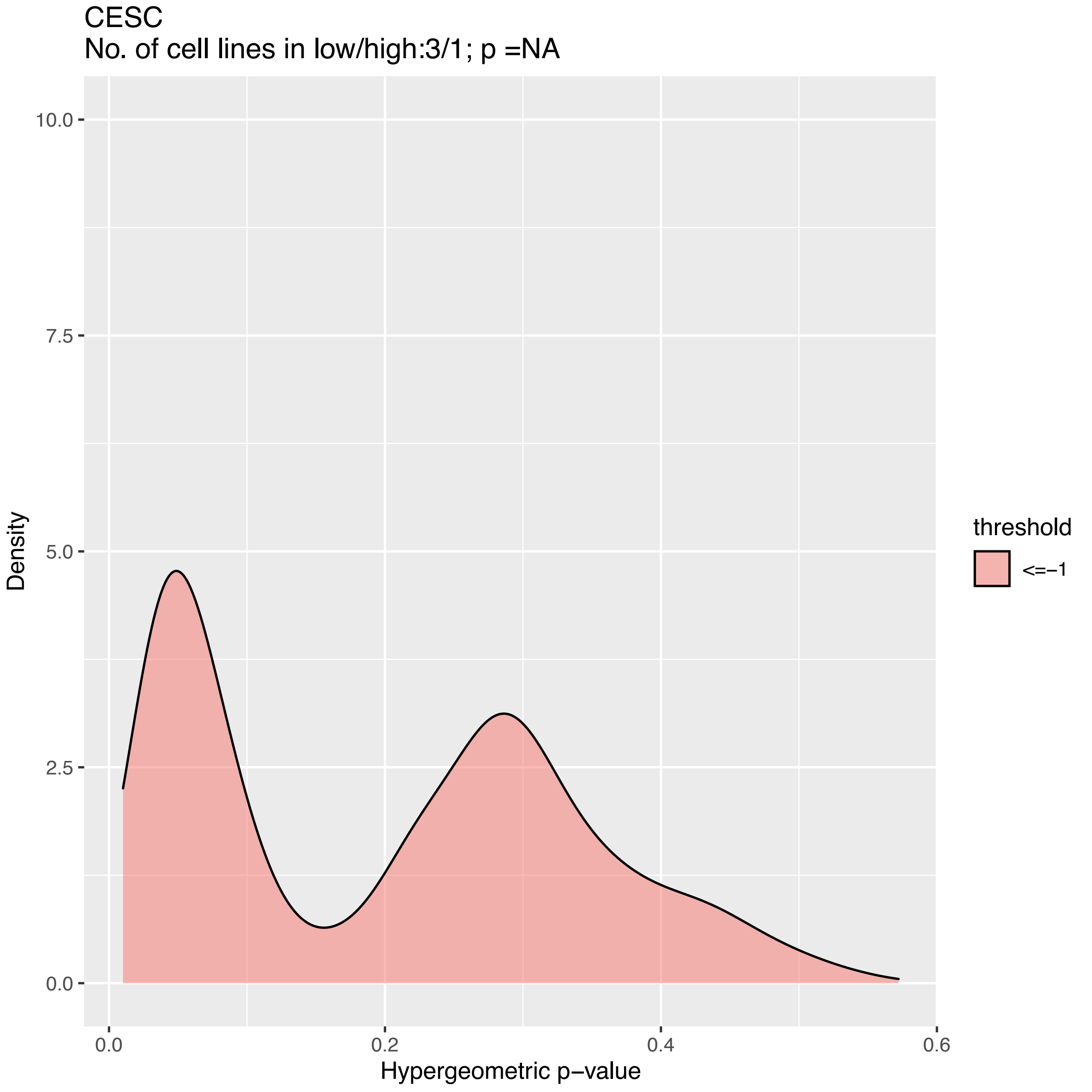
CESC
No. of cell lines in low/high:3/1; p =NA
10.0
7.5
5.0
2.5
0.0
Density
0.0
0.2
0.4
0.6
Hypergeometric p−value
threshold
<=−1

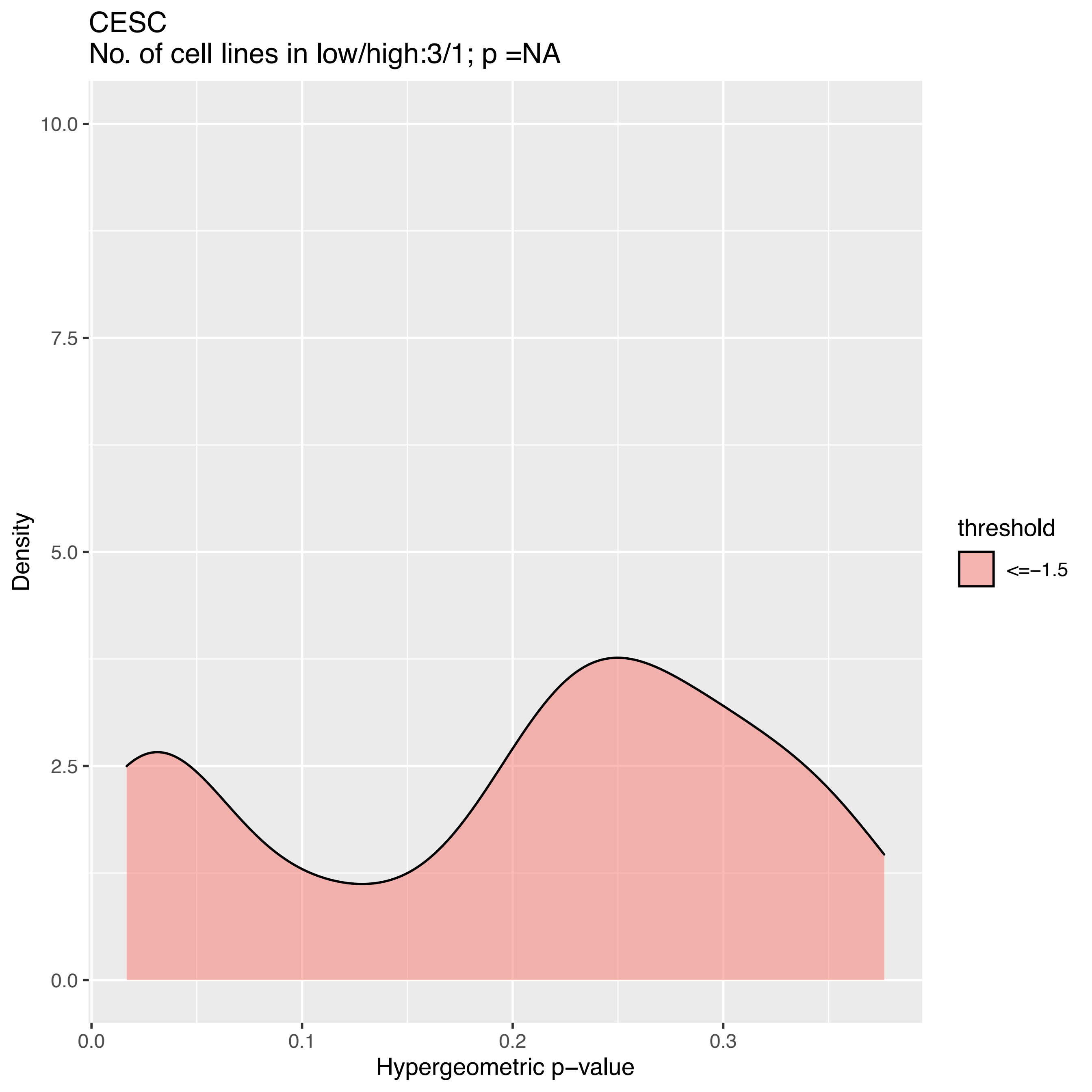

CESC
No. of cell lines in low/high:3/1; p =NA
Density
10.0
7.5
5.0
2.5
0.0
0.0
0.1
0.2
0.3
Hypergeometric p−value
threshold
<=−1.5

CESC
No. of cell lines in low/high:1/1; p =NA

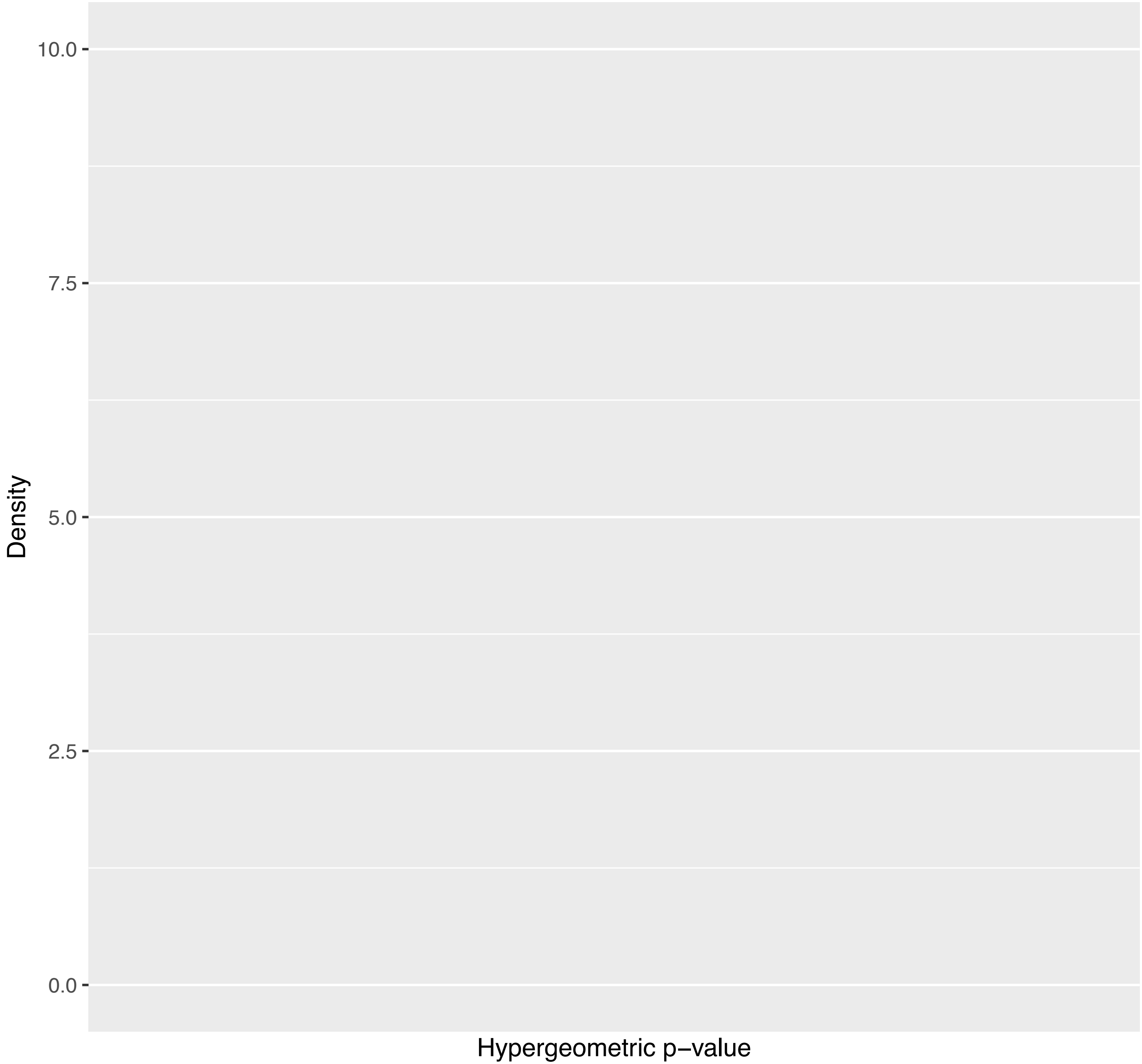

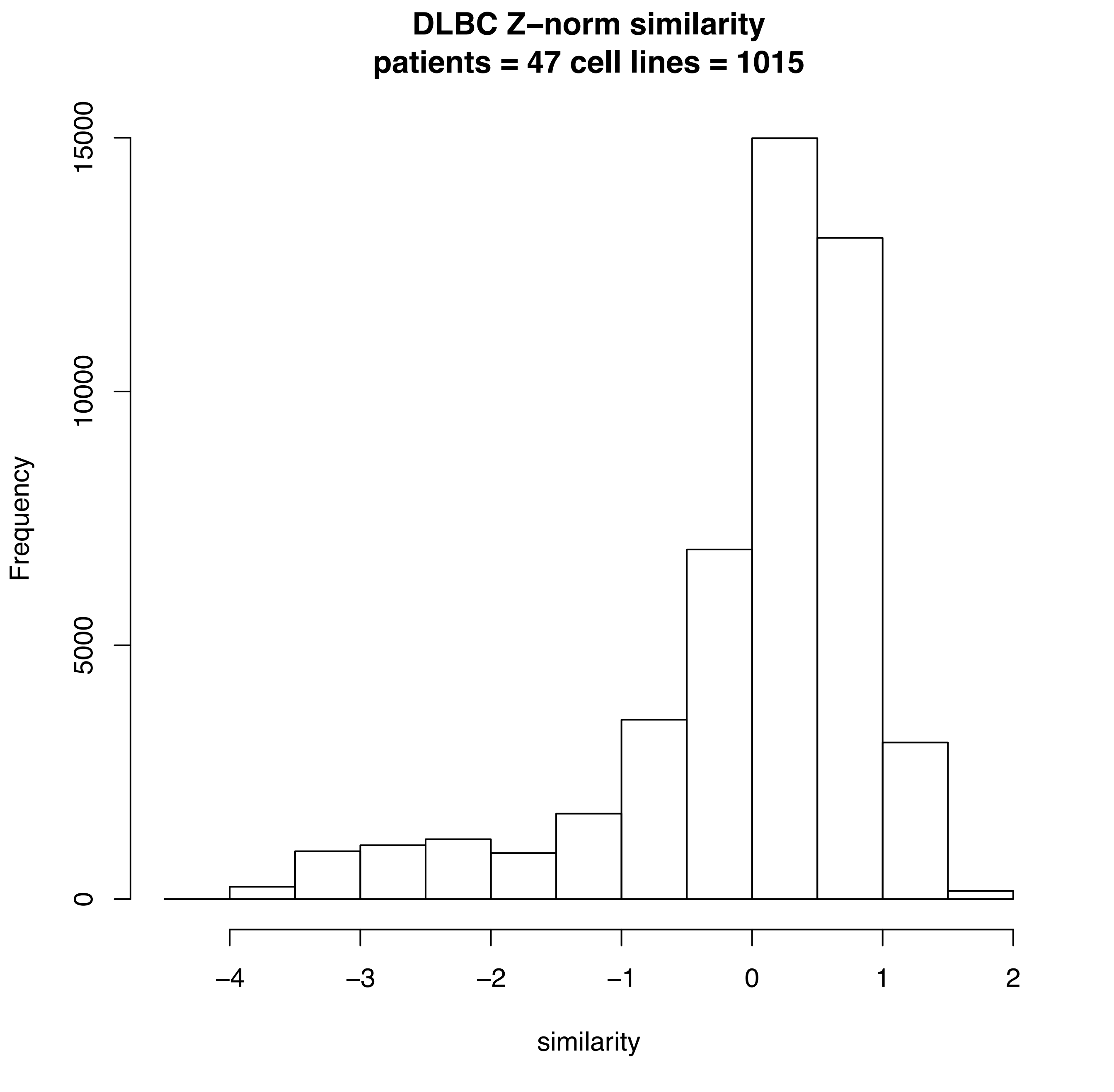
DLBC Z–norm similarity
patients = 47 cell lines = 1015
Frequency
0
5000
10000
15000
−4
−3
−2
−1
0
1
2
similarity

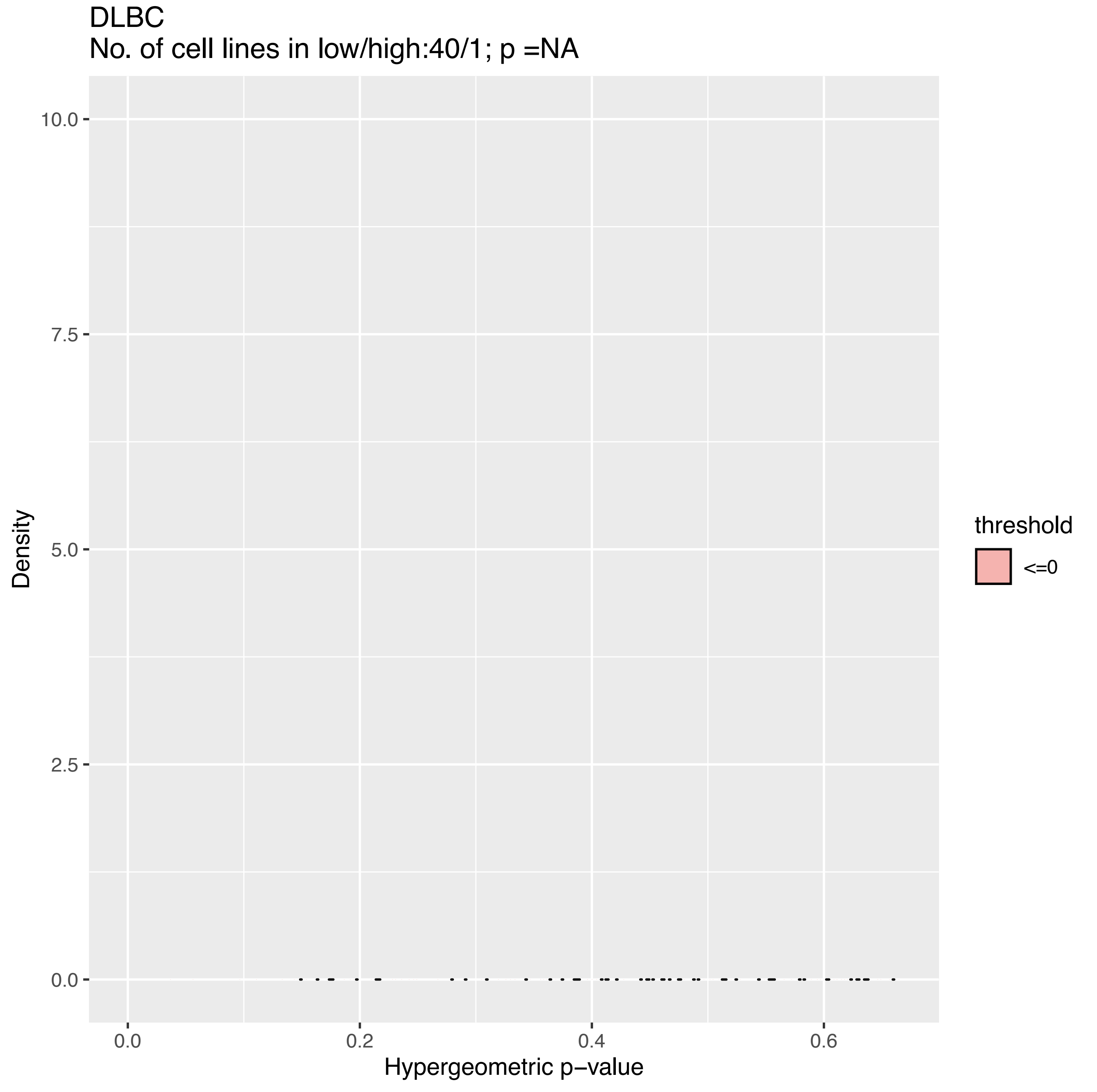
DLBC
No. of cell lines in low/high:40/1; p =NA
10.0
7.5
5.0
2.5
0.0
Density
0.0
0.2
0.4
0.6
Hypergeometric p−value
threshold
<=0

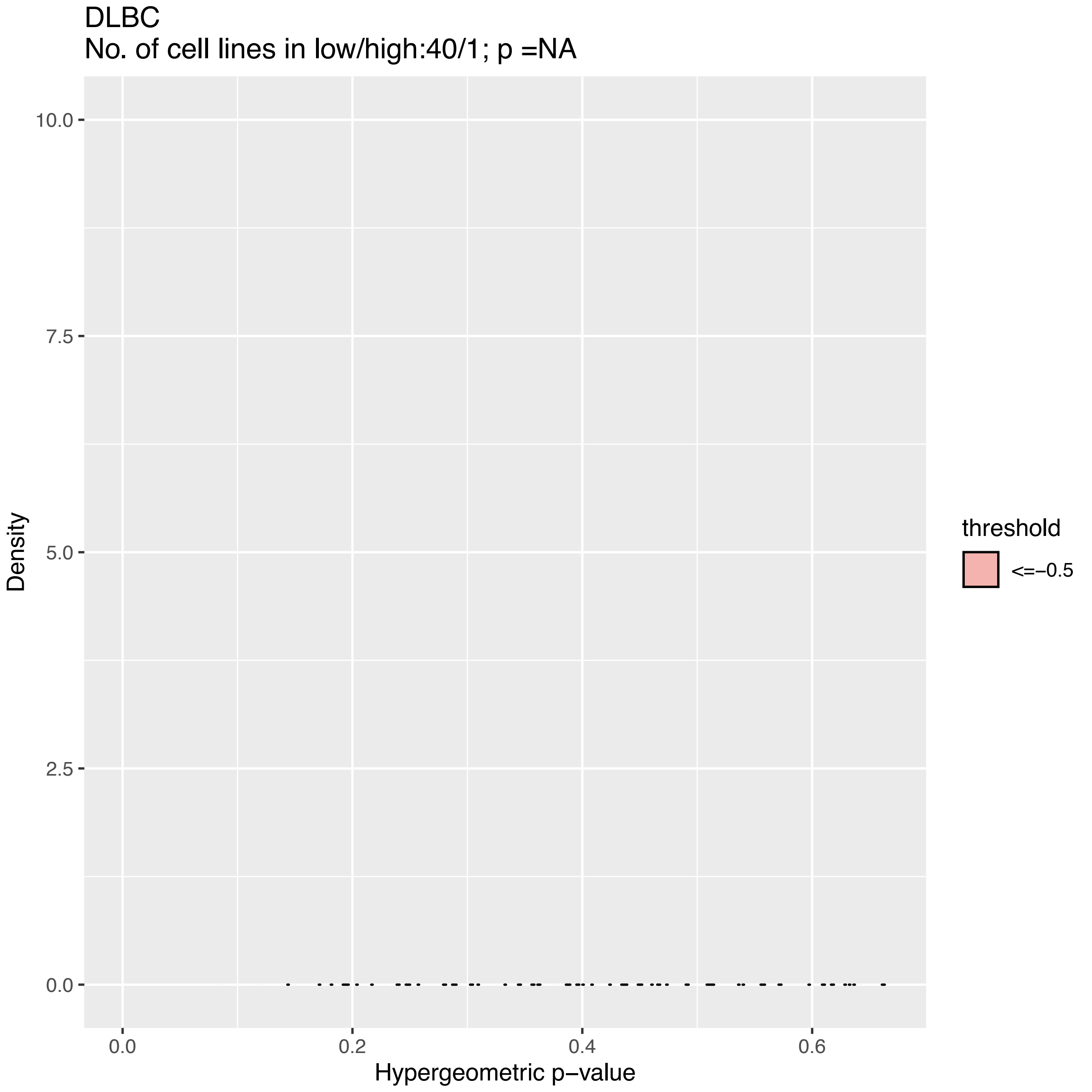
DLBC
No. of cell lines in low/high:40/1; p =NA
10.0
7.5
5.0
2.5
0.0
Density
0.0
0.2
0.4
0.6
Hypergeometric p−value
threshold
<=−0.5

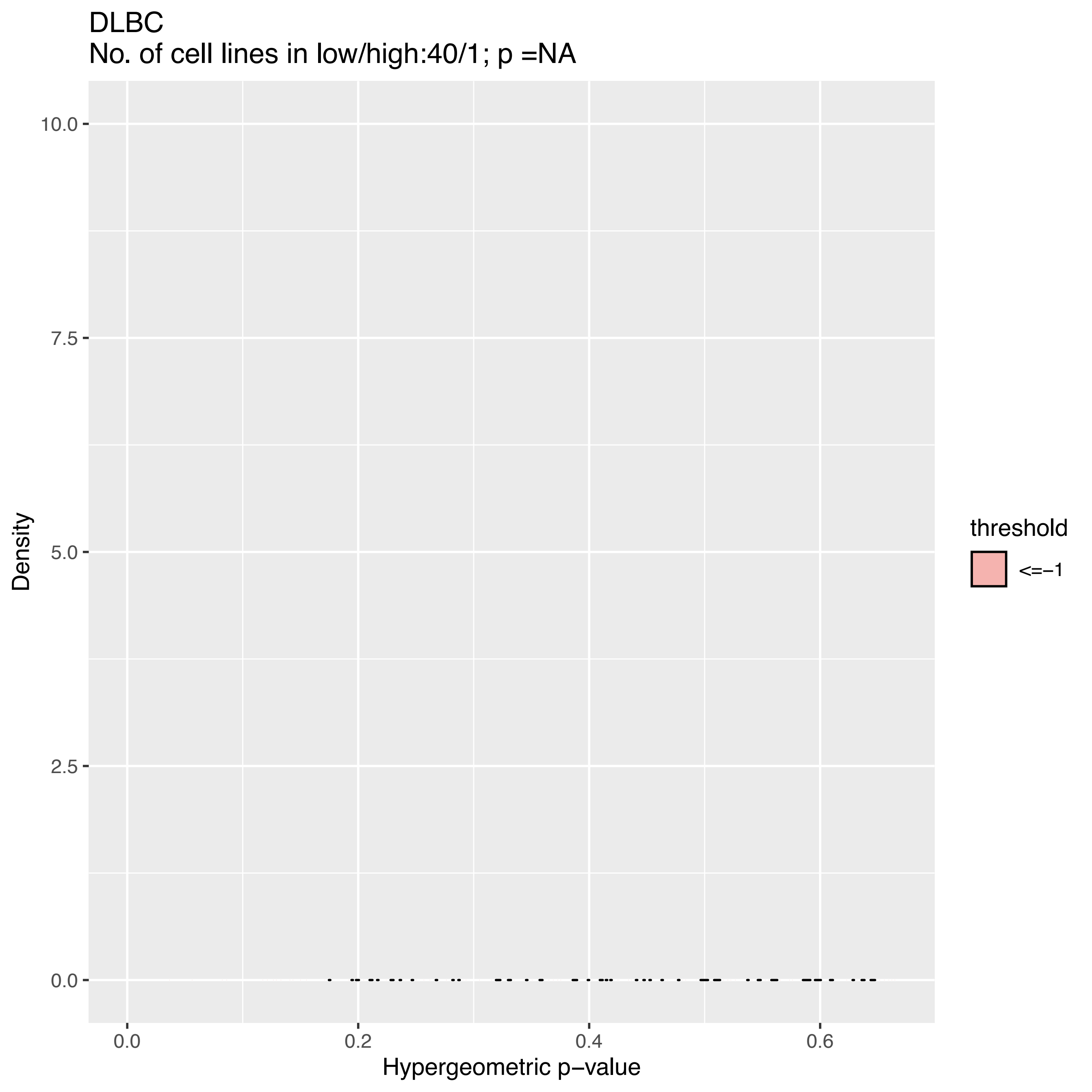
DLBC
No. of cell lines in low/high:40/1; p =NA
10.0
7.5
5.0
2.5
0.0
0.0
0.2
0.4
0.6
Density
Hypergeometric p−value
threshold
<=−1

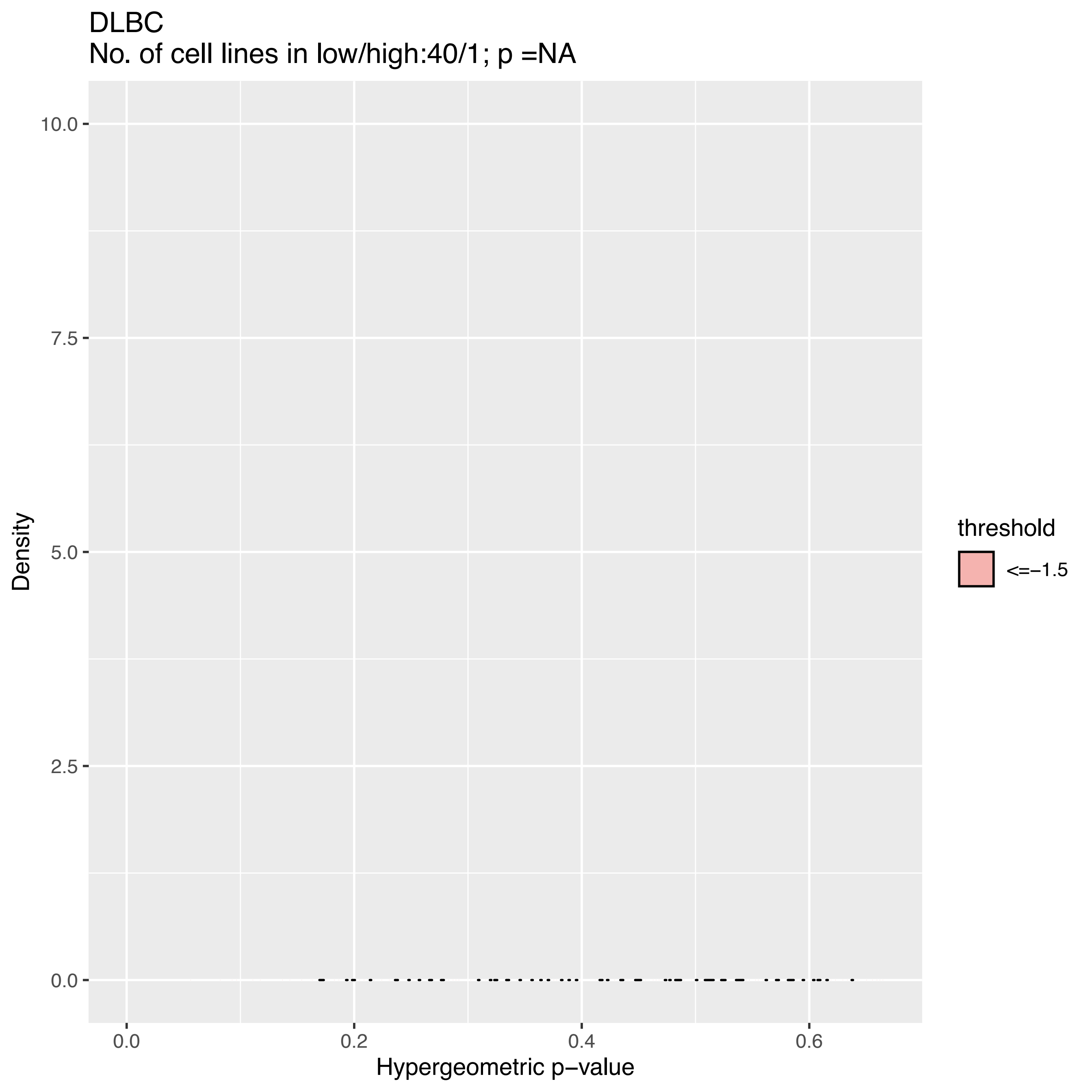
DLBC
No. of cell lines in low/high:40/1; p =NA
Density
10.0
7.5
5.0
2.5
0.0
0.0
0.2
0.4
0.6
Hypergeometric p−value
threshold
<=−1.5

DLBC
No. of cell lines in low/high:39/0; p =NA

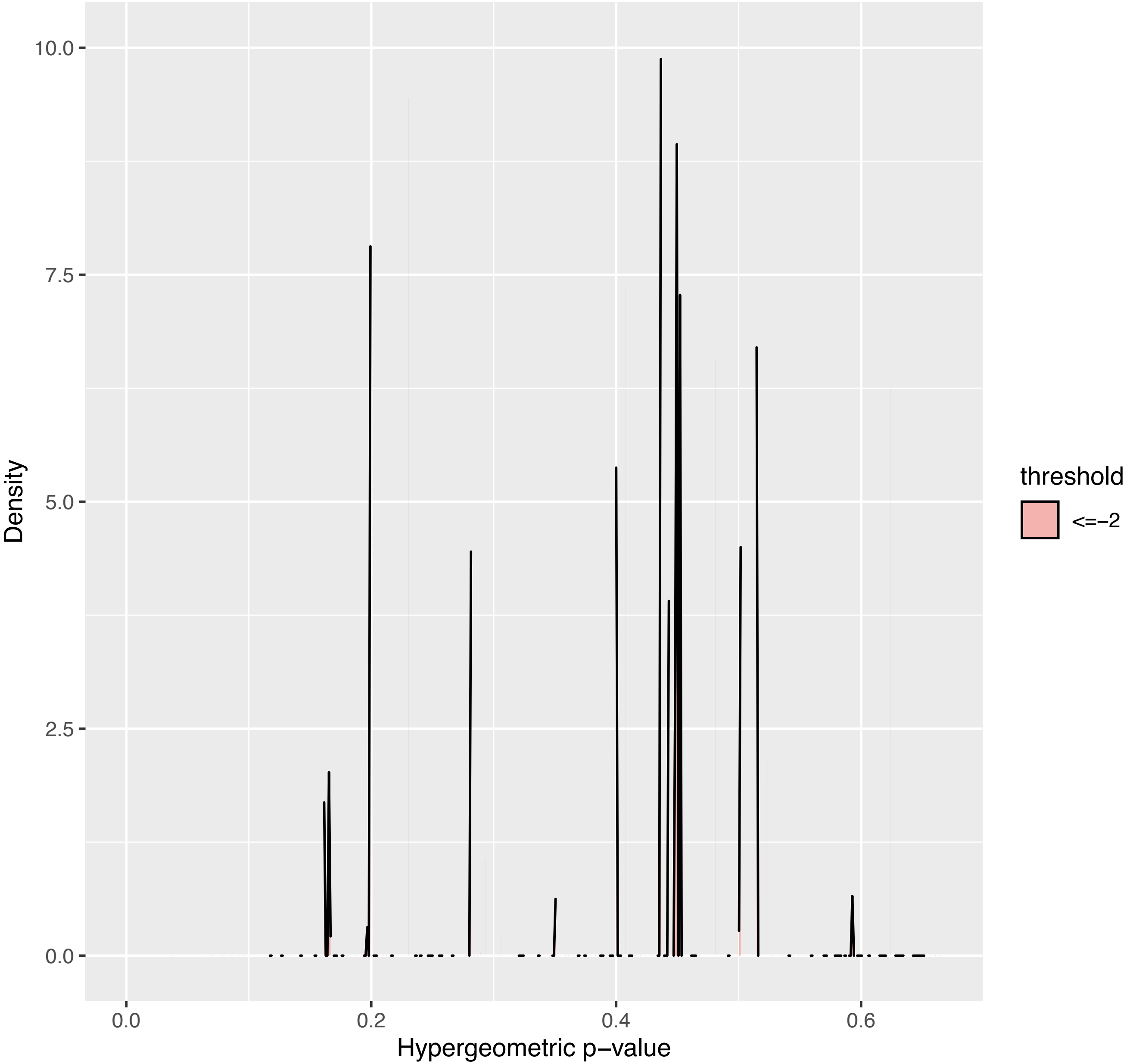

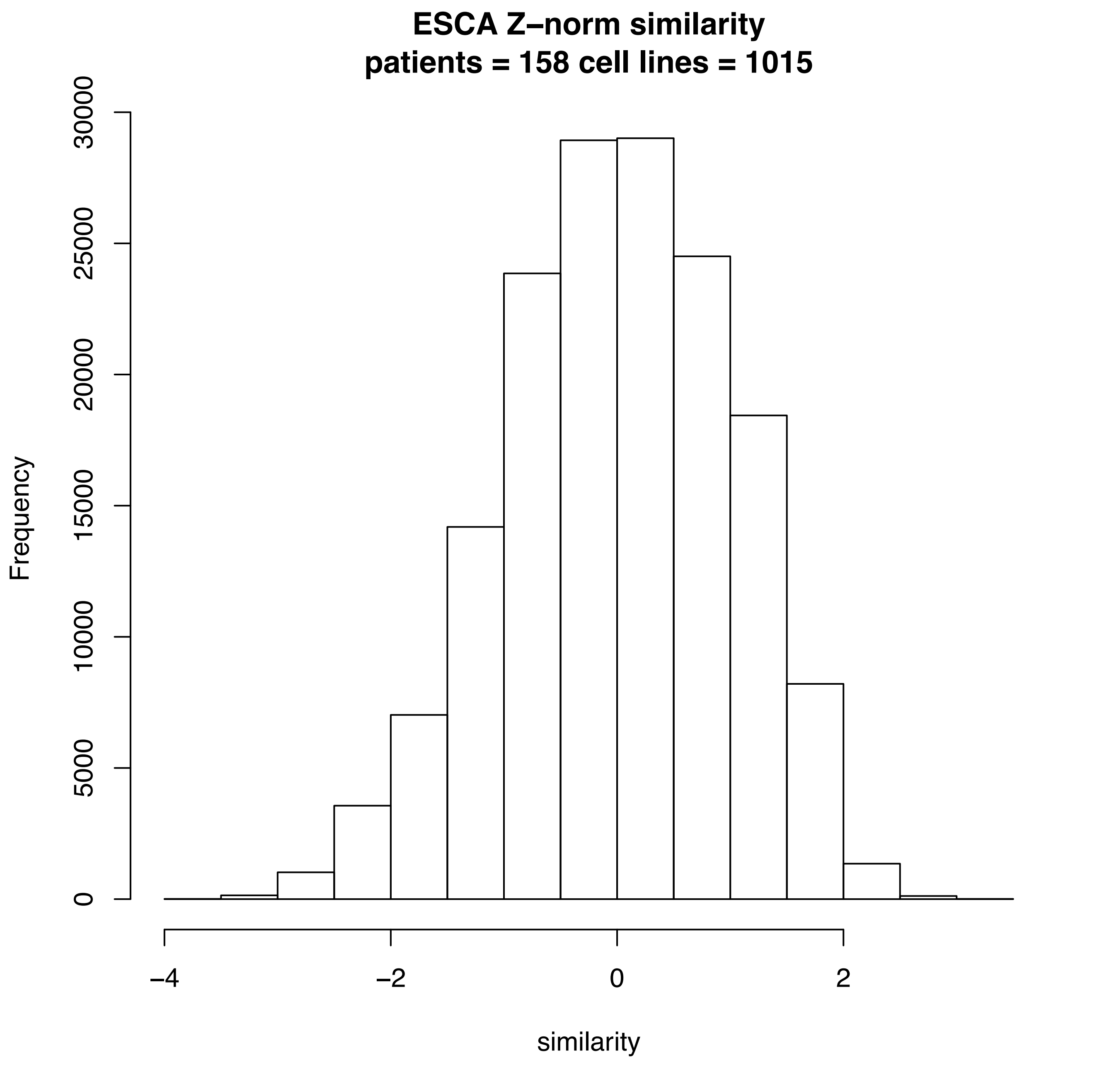
ESCA Z–norm similarity
patients = 158 cell lines = 1015
Frequency
0
5000
10000
15000
20000
25000
30000
−4
−2
0
2
similarity

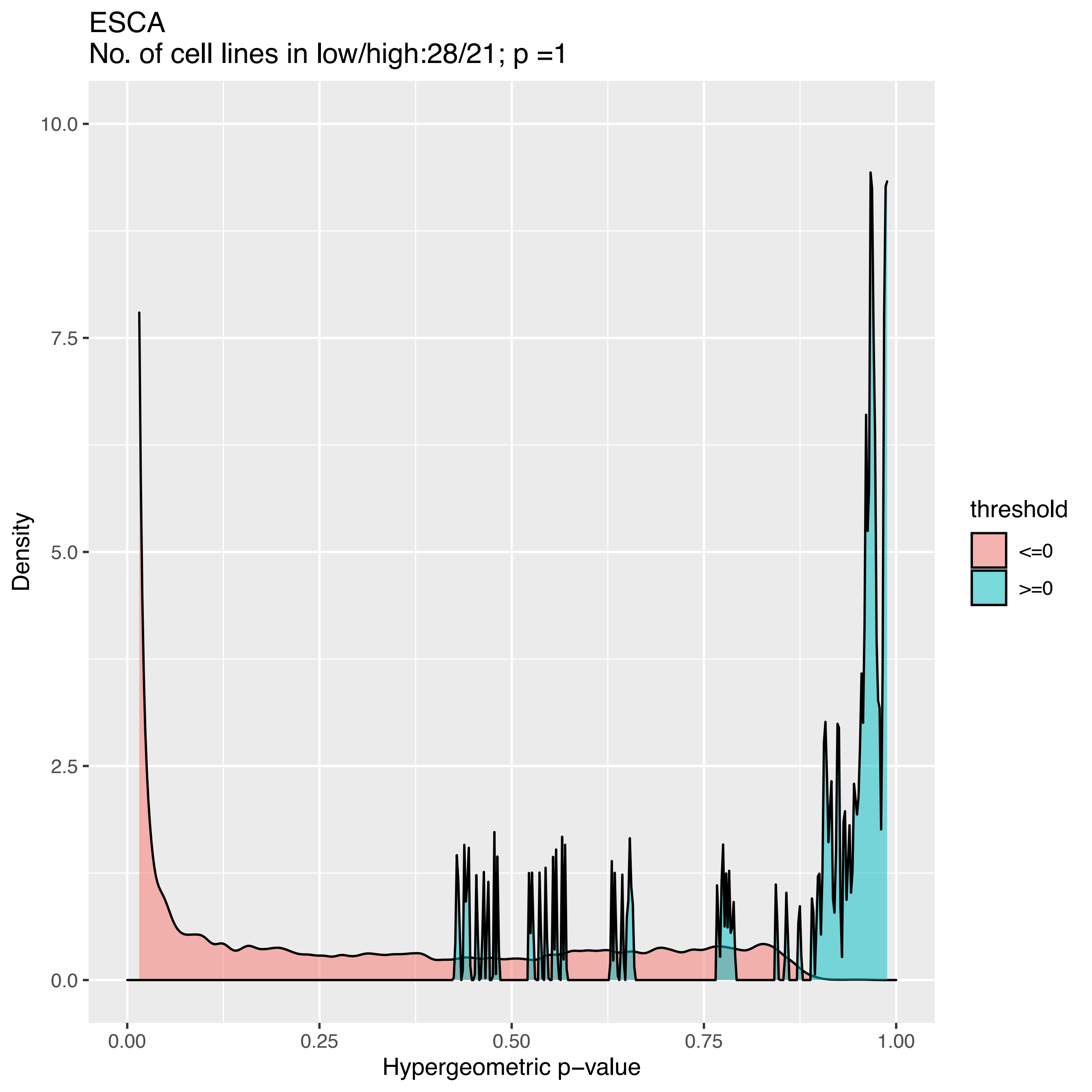
ESCA
No. of cell lines in low/high:28/21; p =1
Density
10.0
7.5
5.0
2.5
0.0
0.00
0.25
0.50
0.75
1.00
Hypergeometric p−value
threshold
<=0
>=0

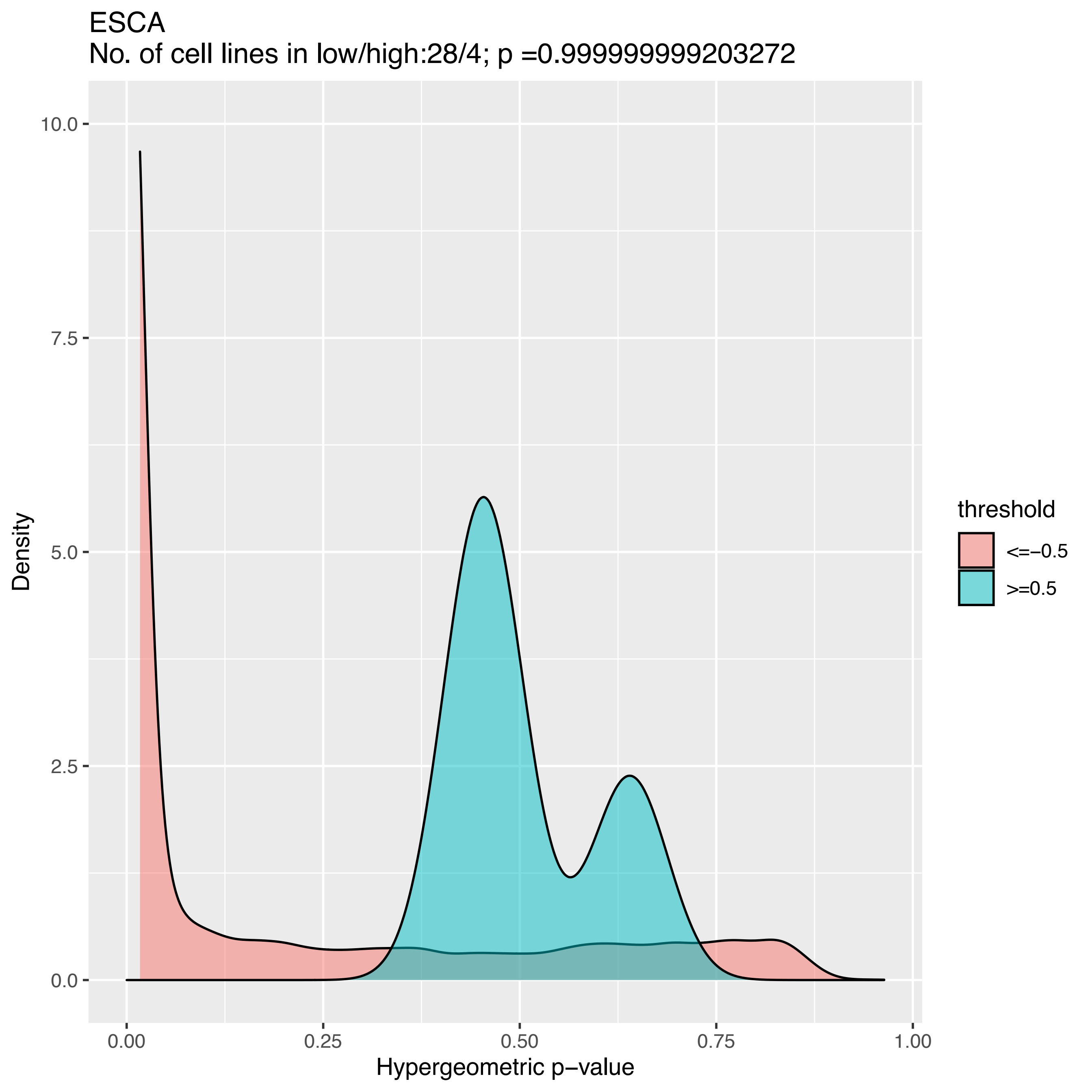

ESCA
No. of cell lines in low/high:28/4; p =0.999999999203272
Density
10.0
7.5
5.0
2.5
0.0
0.00
0.25
0.50
0.75
1.00
Hypergeometric p−value
threshold
<=−0.5
>=0.5

ESCA
No. of cell lines in low/high:28/1; p =NA

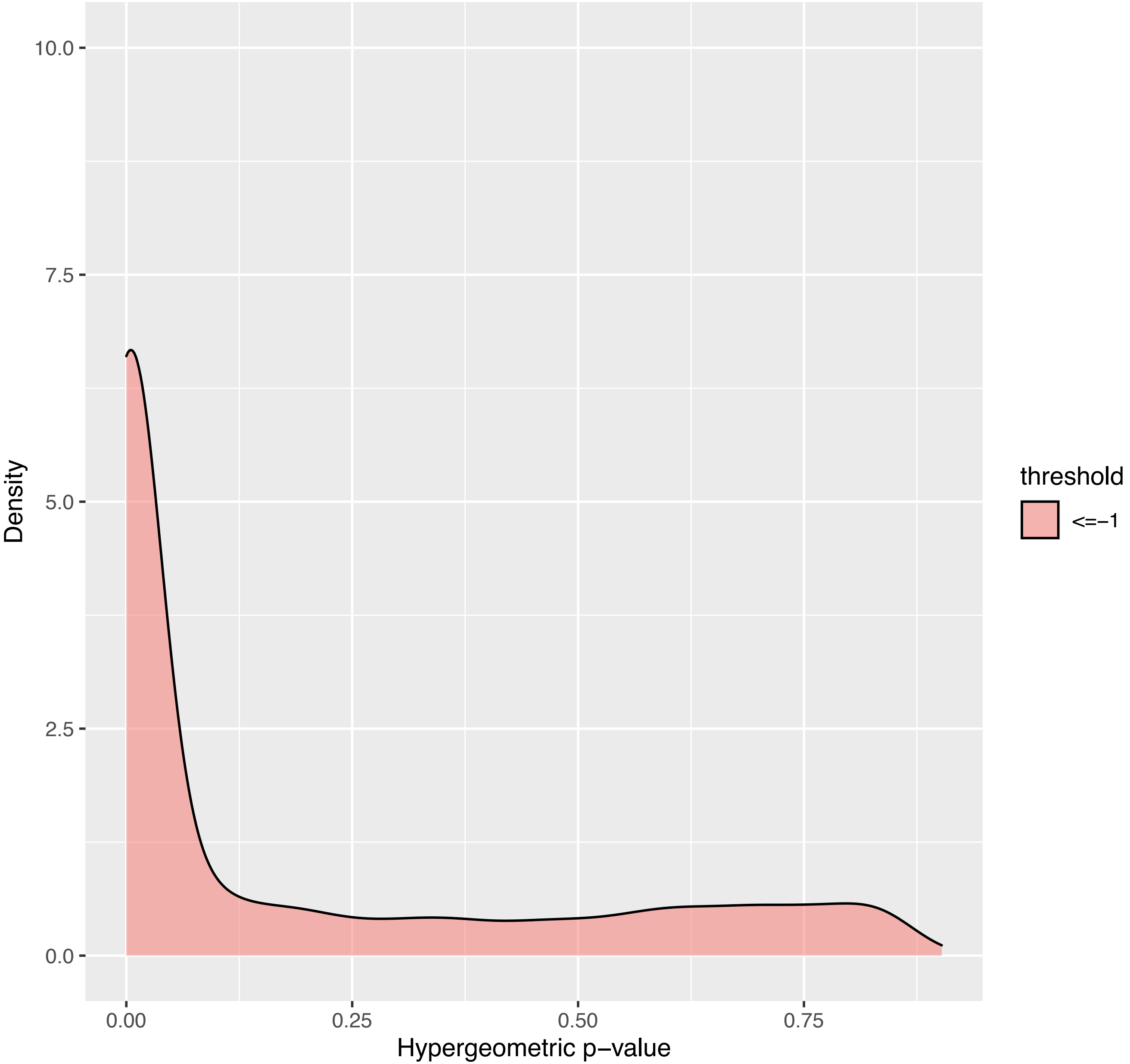

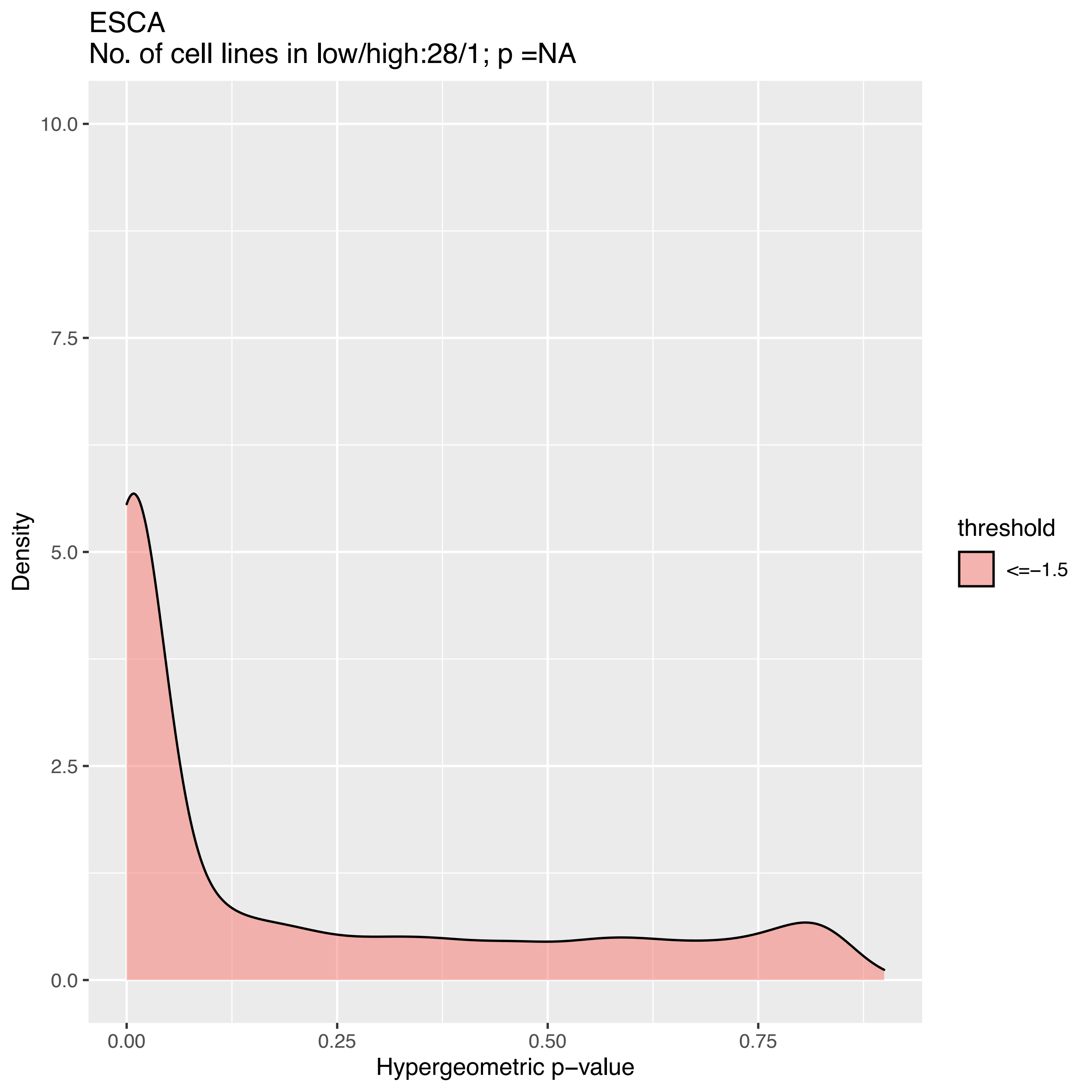
ESCA
No. of cell lines in low/high:28/1; p =NA
10.0
7.5
5.0
2.5
0.0
Density
0.00
0.25
0.50
0.75
Hypergeometric p−value
threshold
<=−1.5

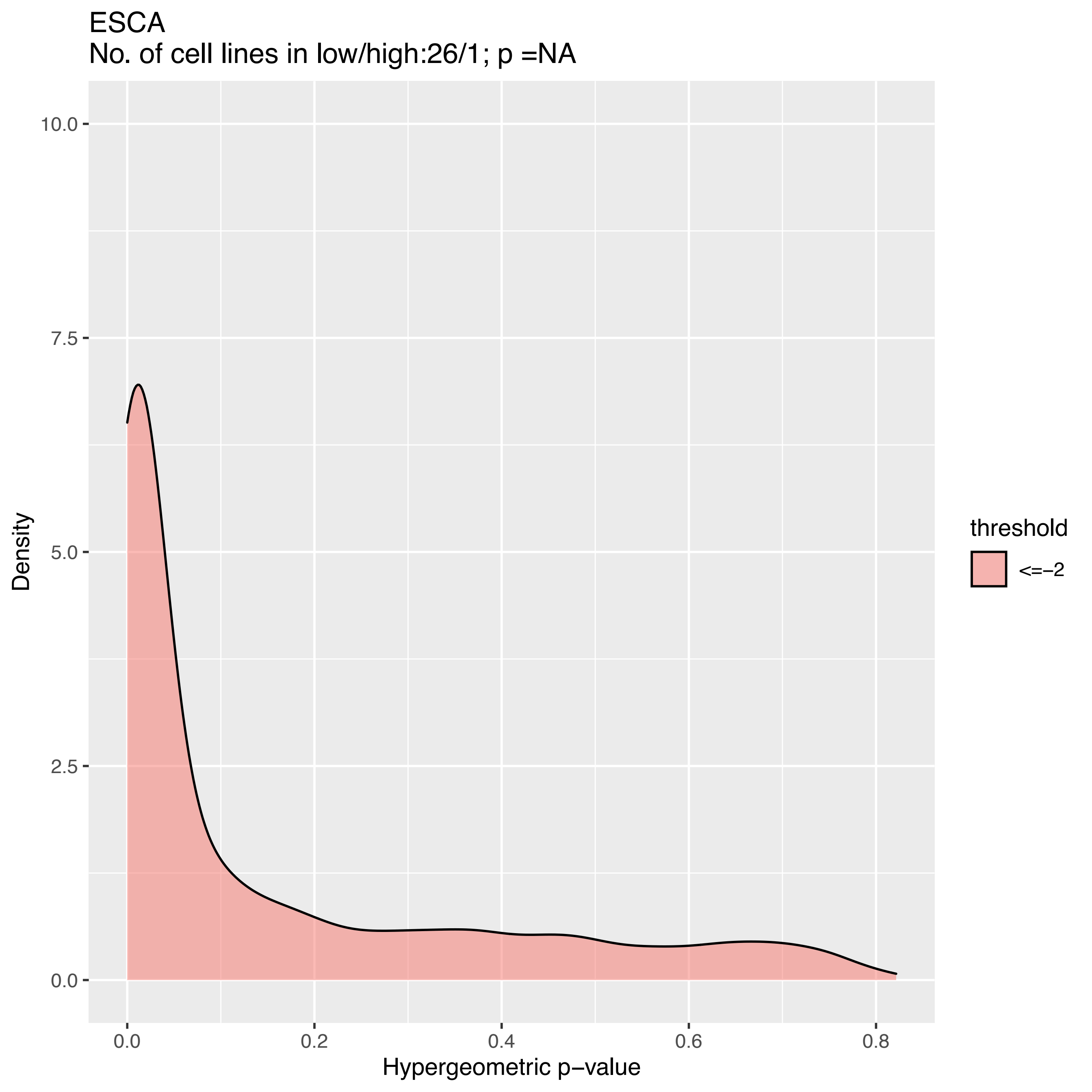
ESCA
No. of cell lines in low/high:26/1; p =NA
Density
10.0
7.5
5.0
2.5
0.0
0.0
0.2
0.4
0.6
0.8
Hypergeometric p−value
threshold
<=−2

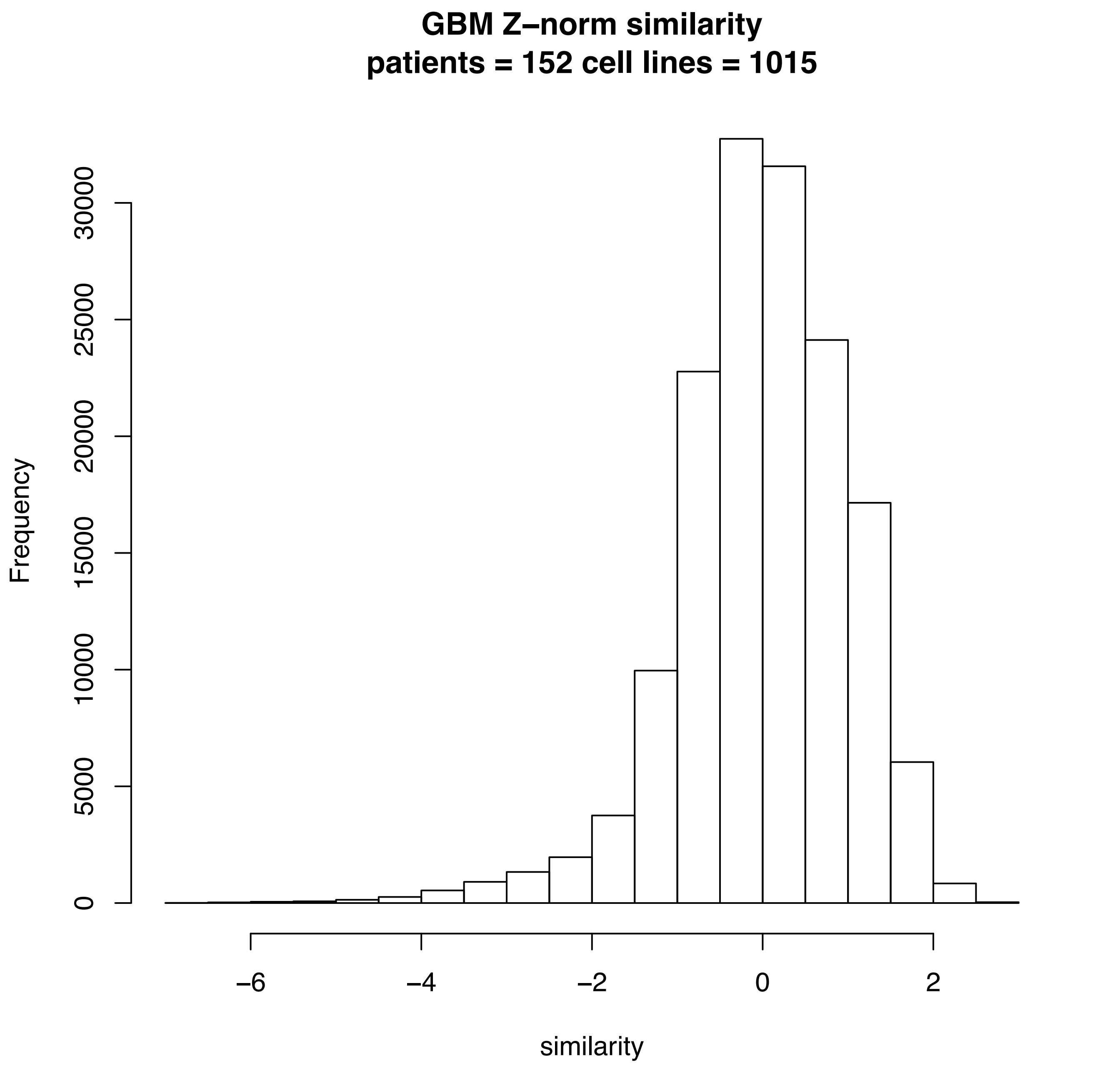
GBM Z–norm similarity
patients = 152 cell lines = 1015
Frequency
0
5000
10000
15000
20000
25000
30000
–6
–4
–2
0
2
similarity

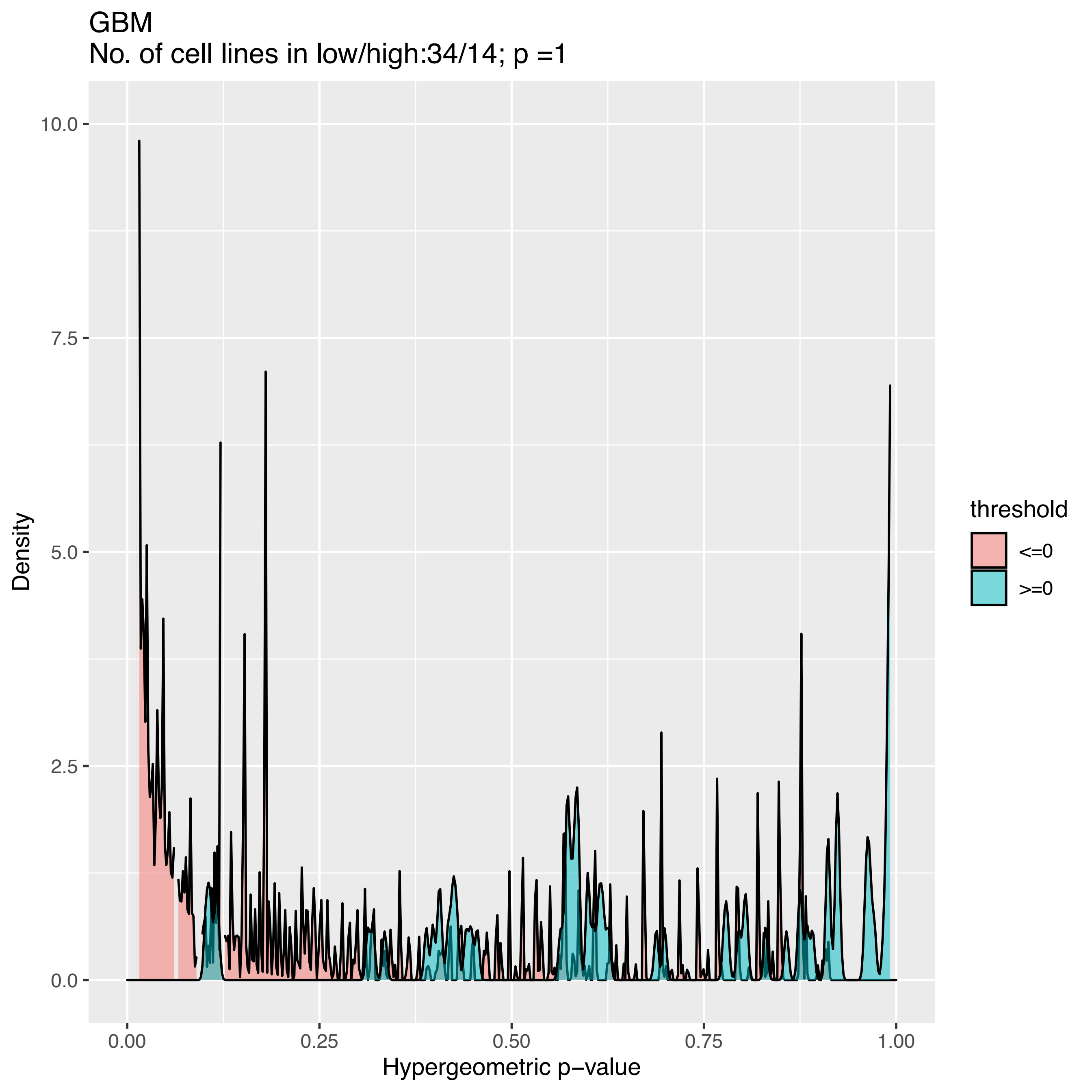
GBM
No. of cell lines in low/high:34/14; p =1
Density
10.0
7.5
5.0
2.5
0.0
0.00
0.25
0.50
0.75
1.00
Hypergeometric p−value
threshold
<=0
>=0

GBM
No. of cell lines in low/high:34/1; p =NA

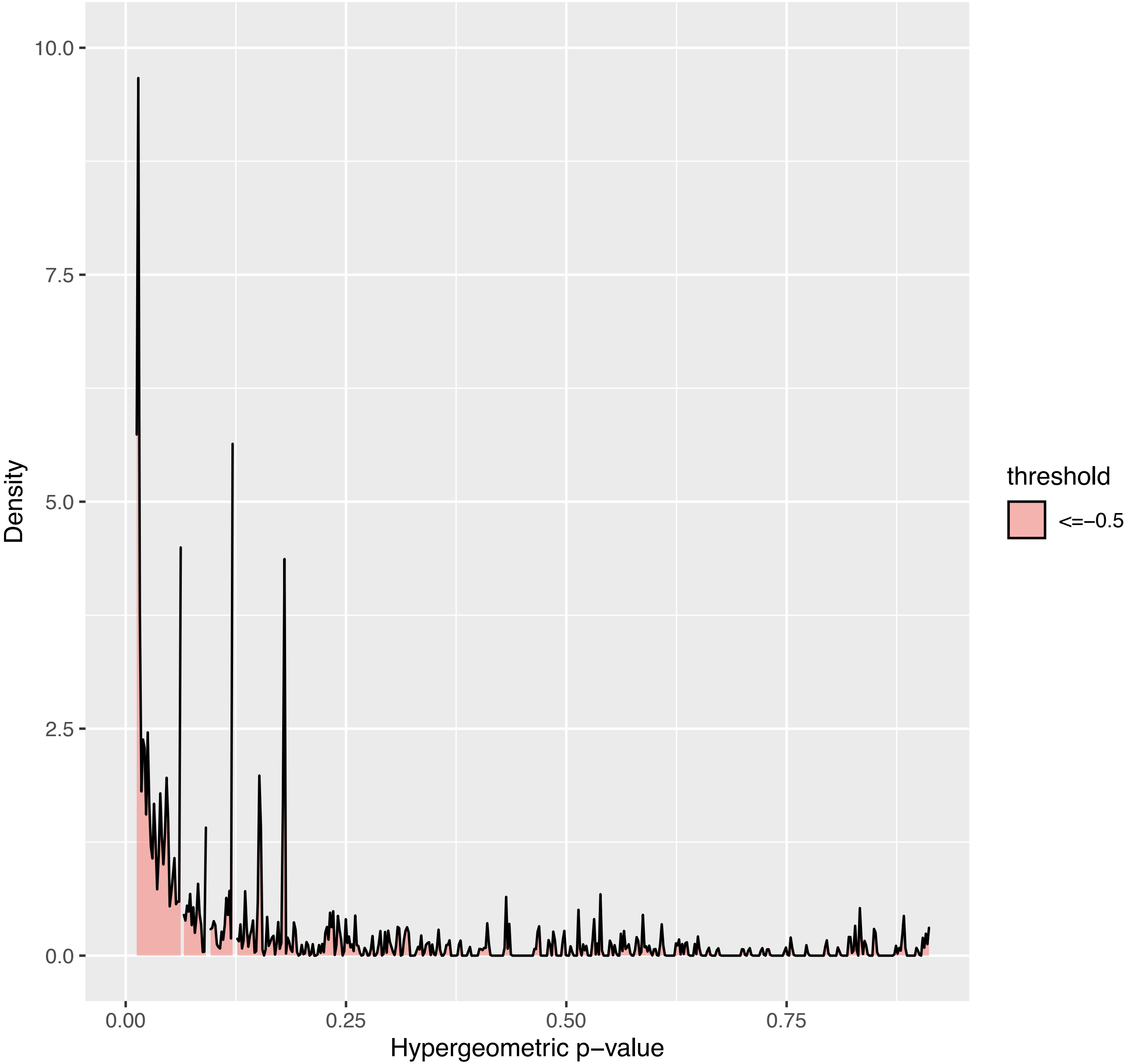

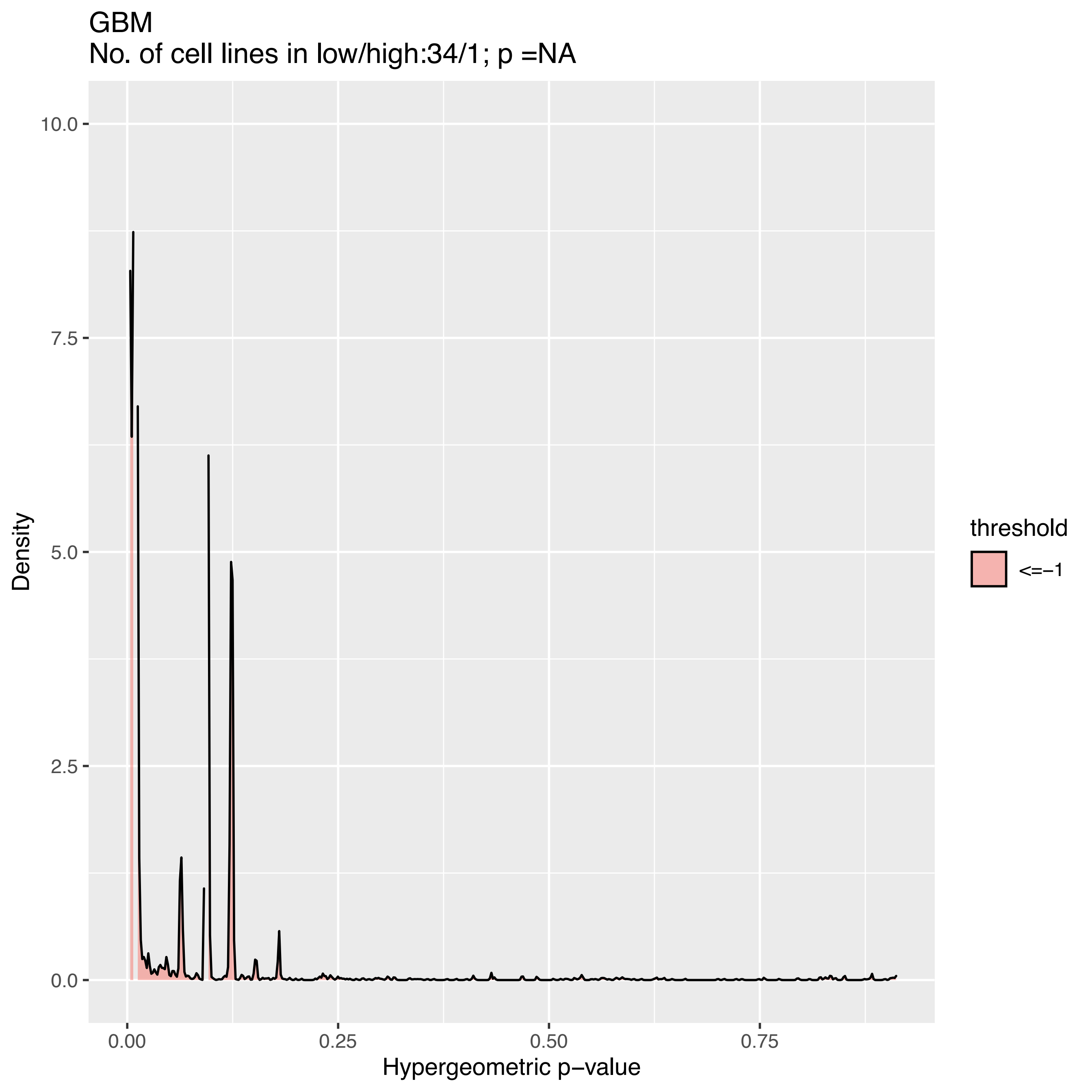
GBM
No. of cell lines in low/high:34/1; p =NA
Density
10.0
7.5
5.0
2.5
0.0
0.00
0.25
0.50
0.75
Hypergeometric p−value
threshold
<=−1

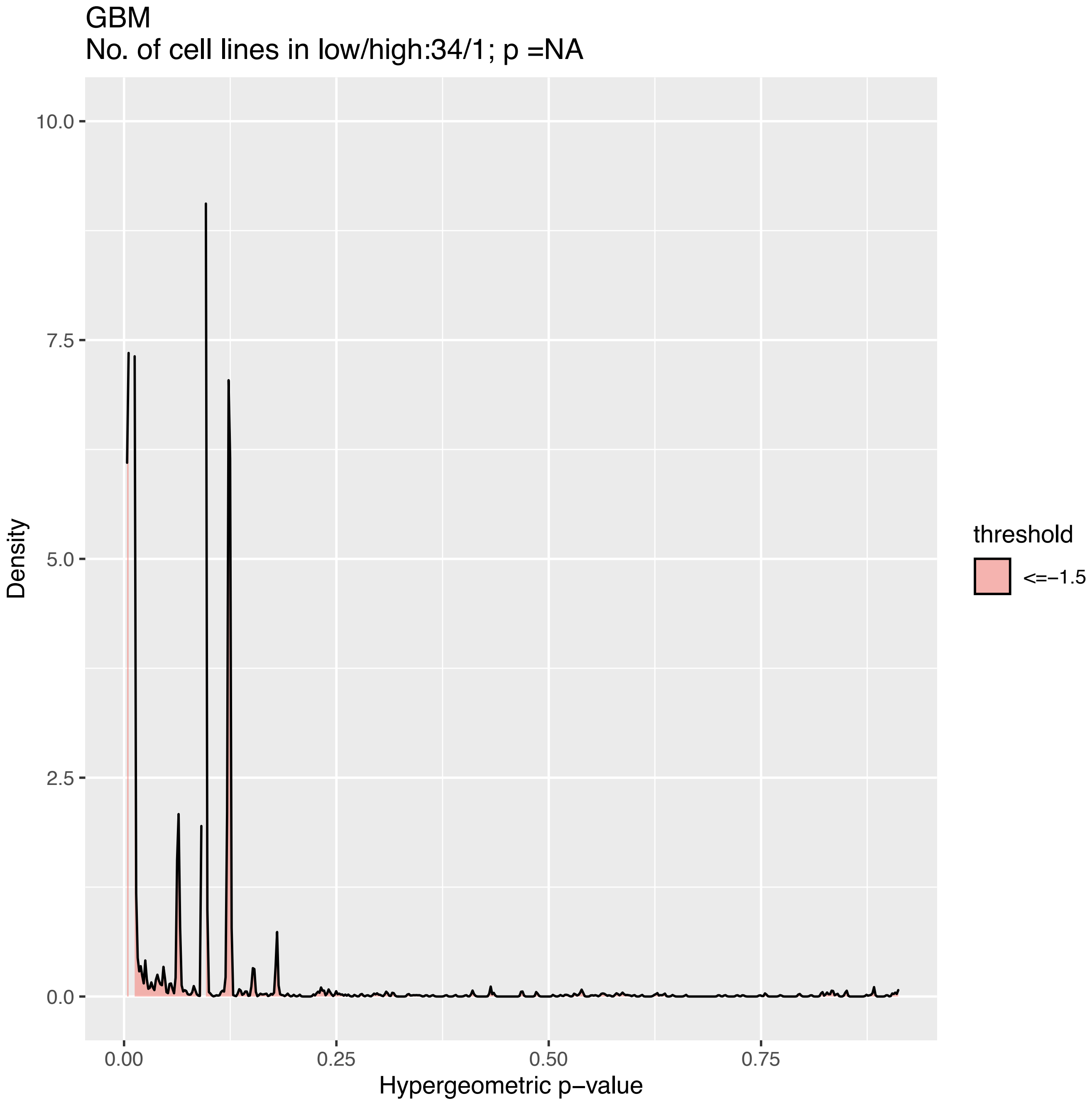
GBM
No. of cell lines in low/high:34/1; p =NA
10.0
7.5
5.0
2.5
0.0
Density
0.00
0.25
0.50
0.75
Hypergeometric p−value
threshold
<=−1.5

GBM
No. of cell lines in low/high:34/1; p =NA

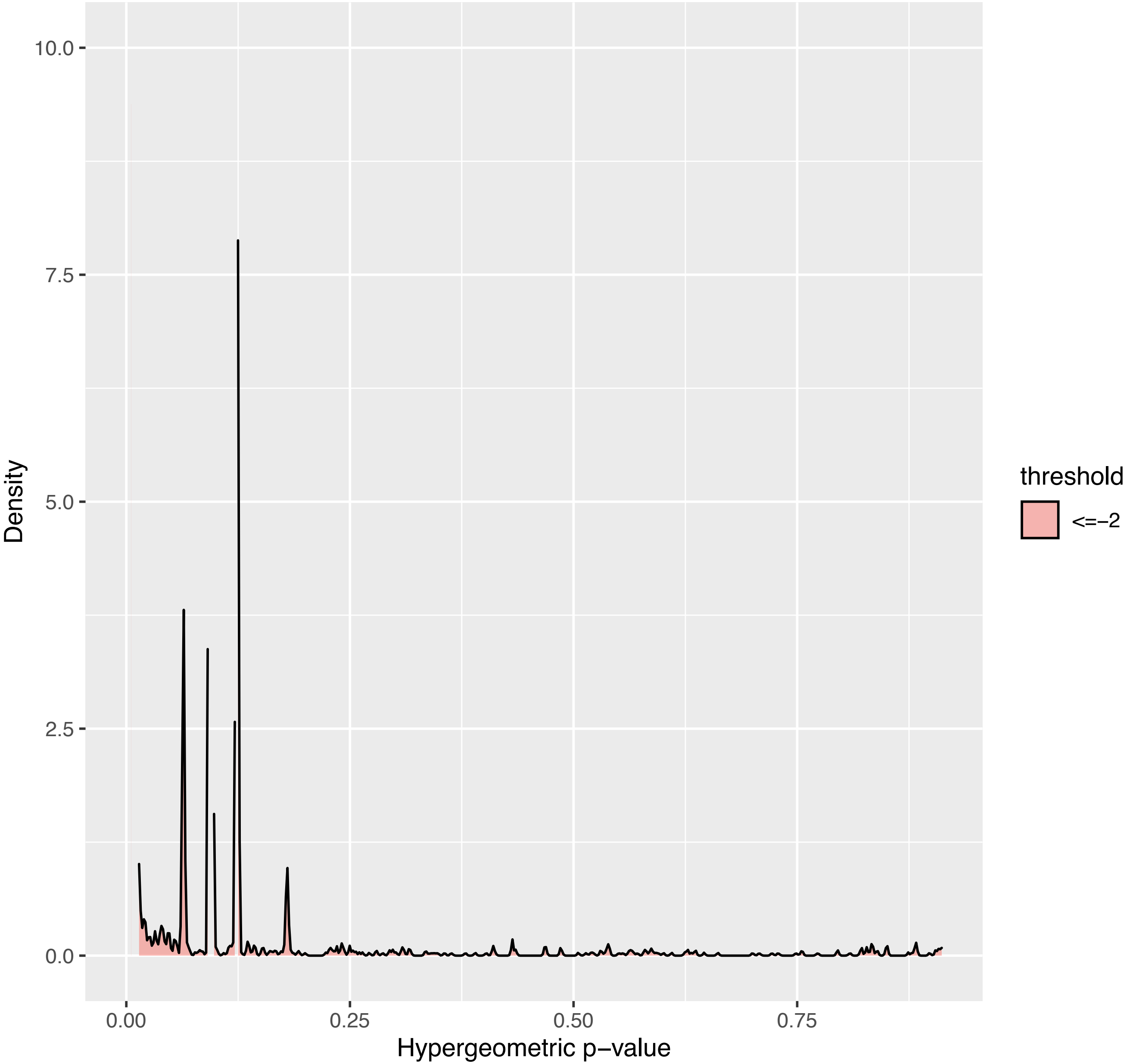

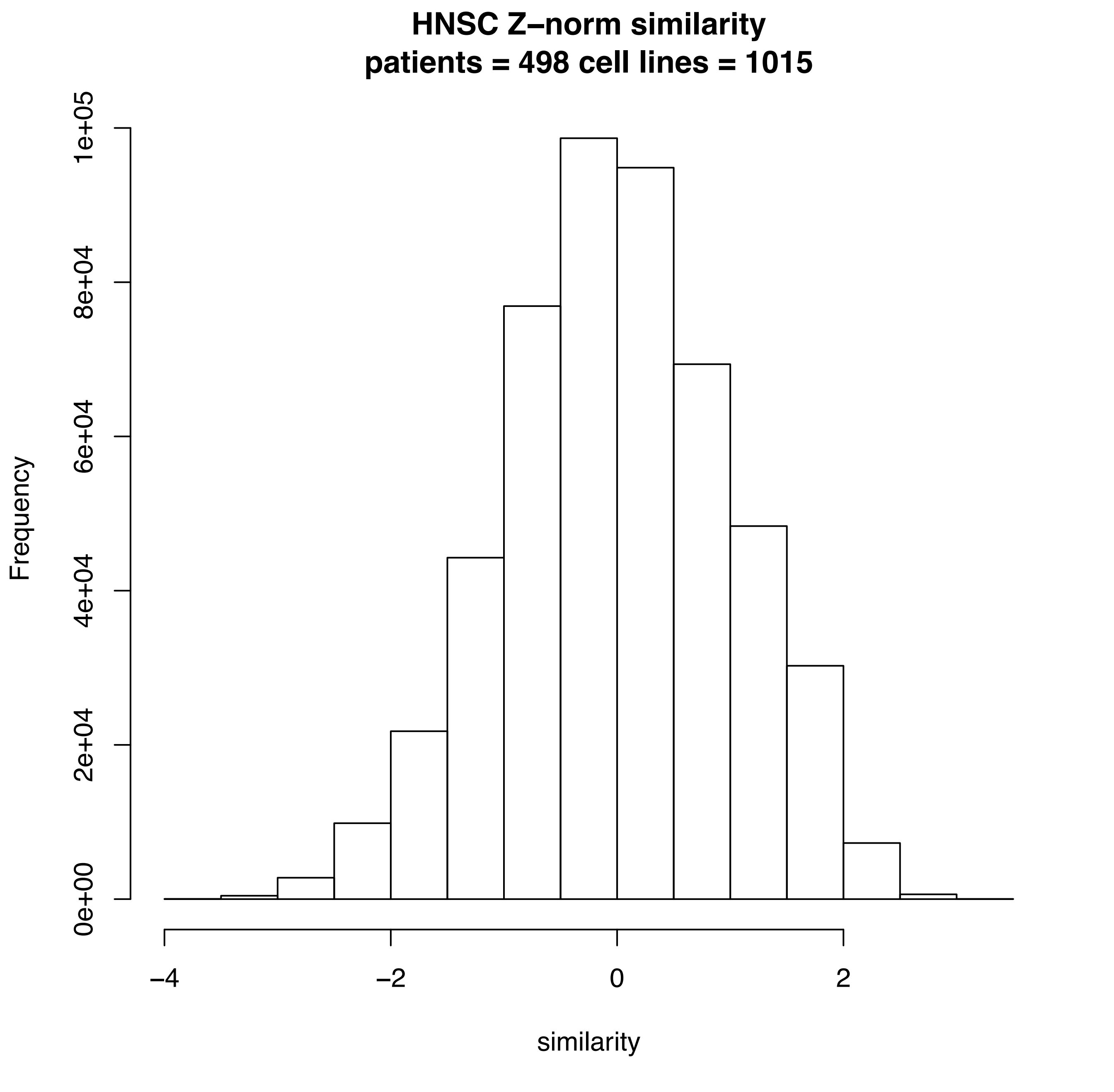
HNSC Z–norm similarity
patients = 498 cell lines = 1015
Frequency
0e+00
2e+04
4e+04
6e+04
8e+04
1e+05
−4
−2
0
2
similarity

HNSC
No. of cell lines in low/high:34/16; p =1

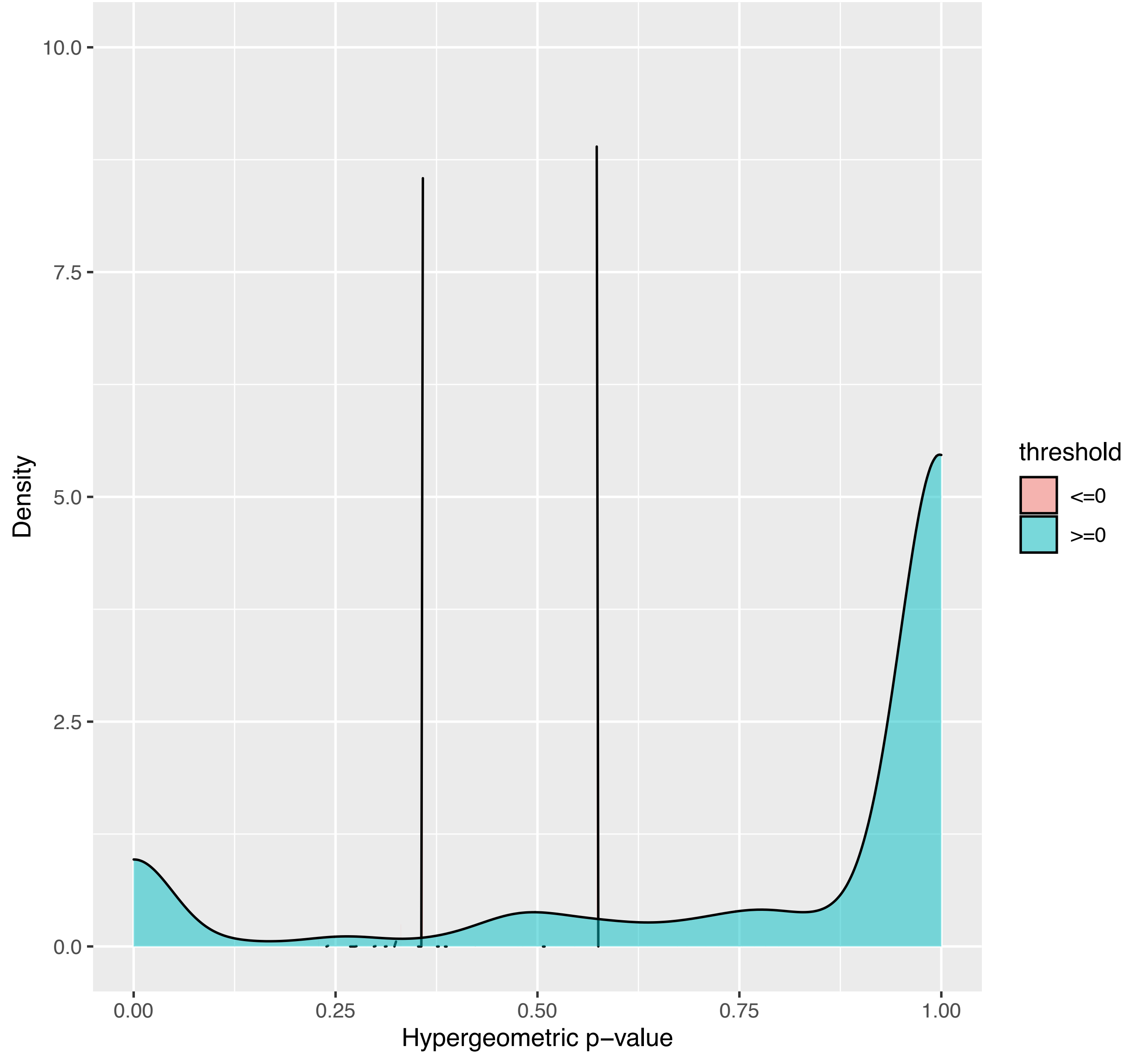

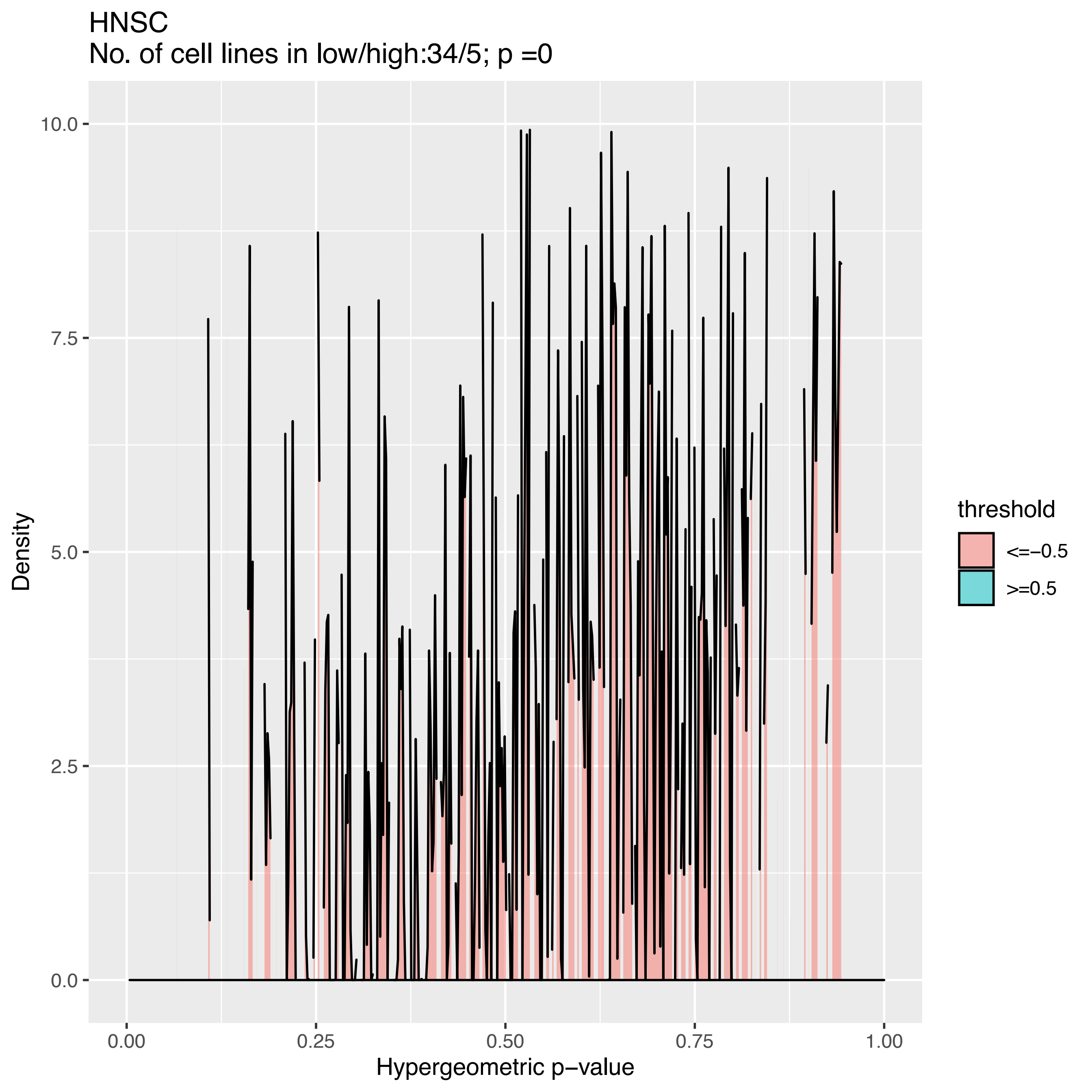
HNSC
No. of cell lines in low/high:34/5; p =0
Density
10.0
7.5
5.0
2.5
0.0
0.00
0.25
0.50
0.75
1.00
Hypergeometric p−value
threshold
<=−0.5
>=0.5

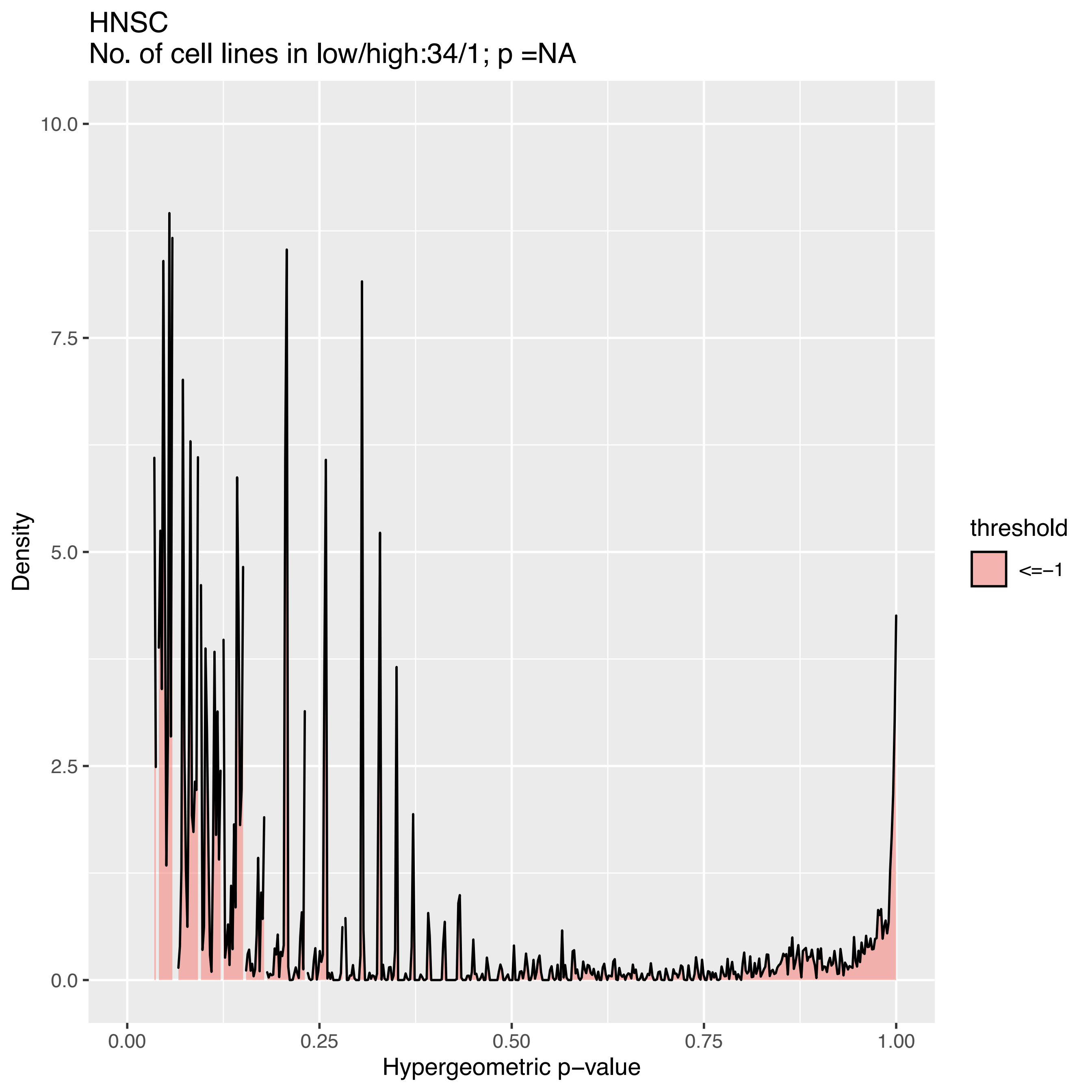
HNSC
No. of cell lines in low/high:34/1; p =NA
Density
Hypergeometric p−value
threshold
<=−1
10.0
7.5
5.0
2.5
0.0
0.00
0.25
0.50
0.75
1.00

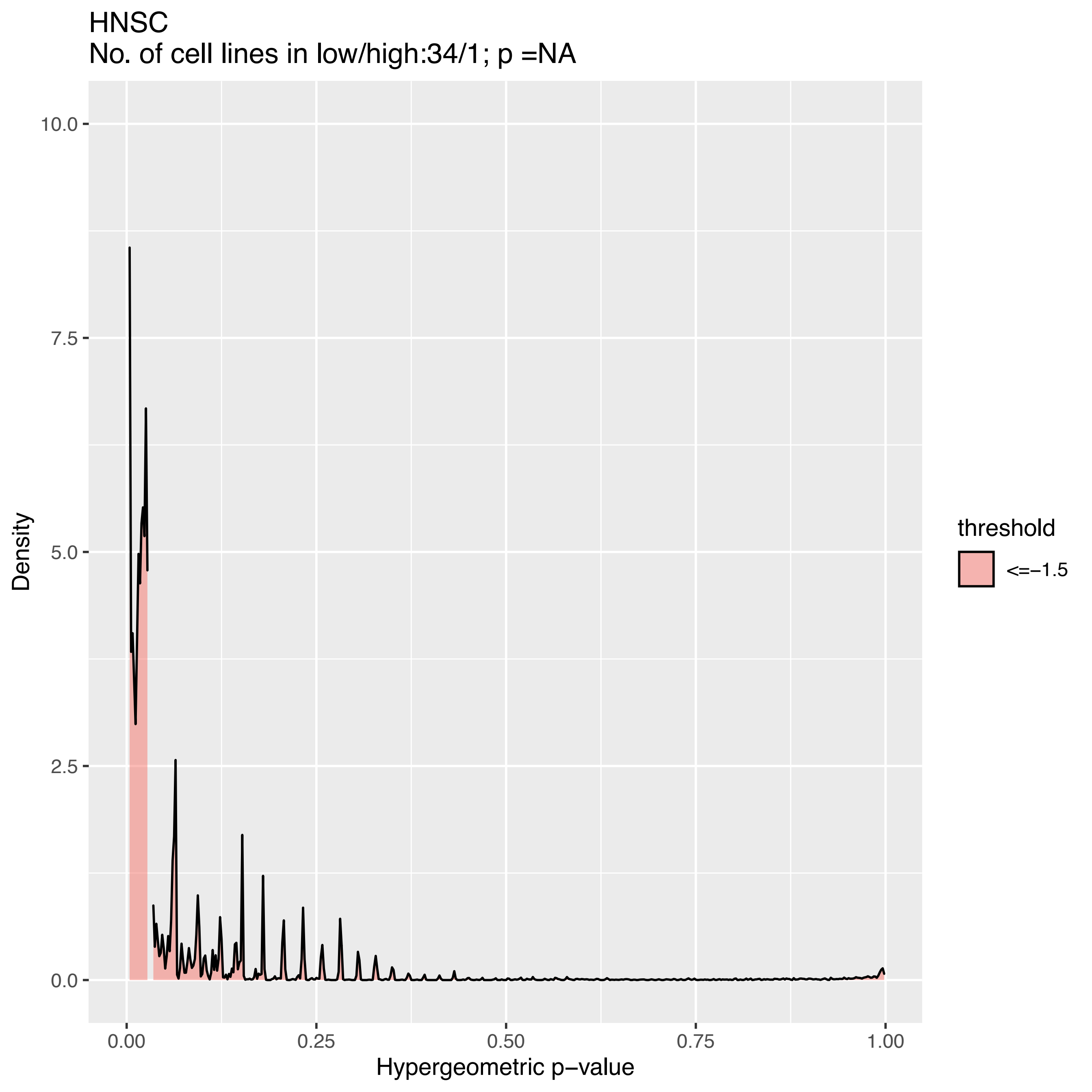
HNSC
No. of cell lines in low/high:34/1; p =NA
Density
10.0
7.5
5.0
2.5
0.0
0.00
0.25
0.50
0.75
1.00
Hypergeometric p−value
threshold
<=−1.5

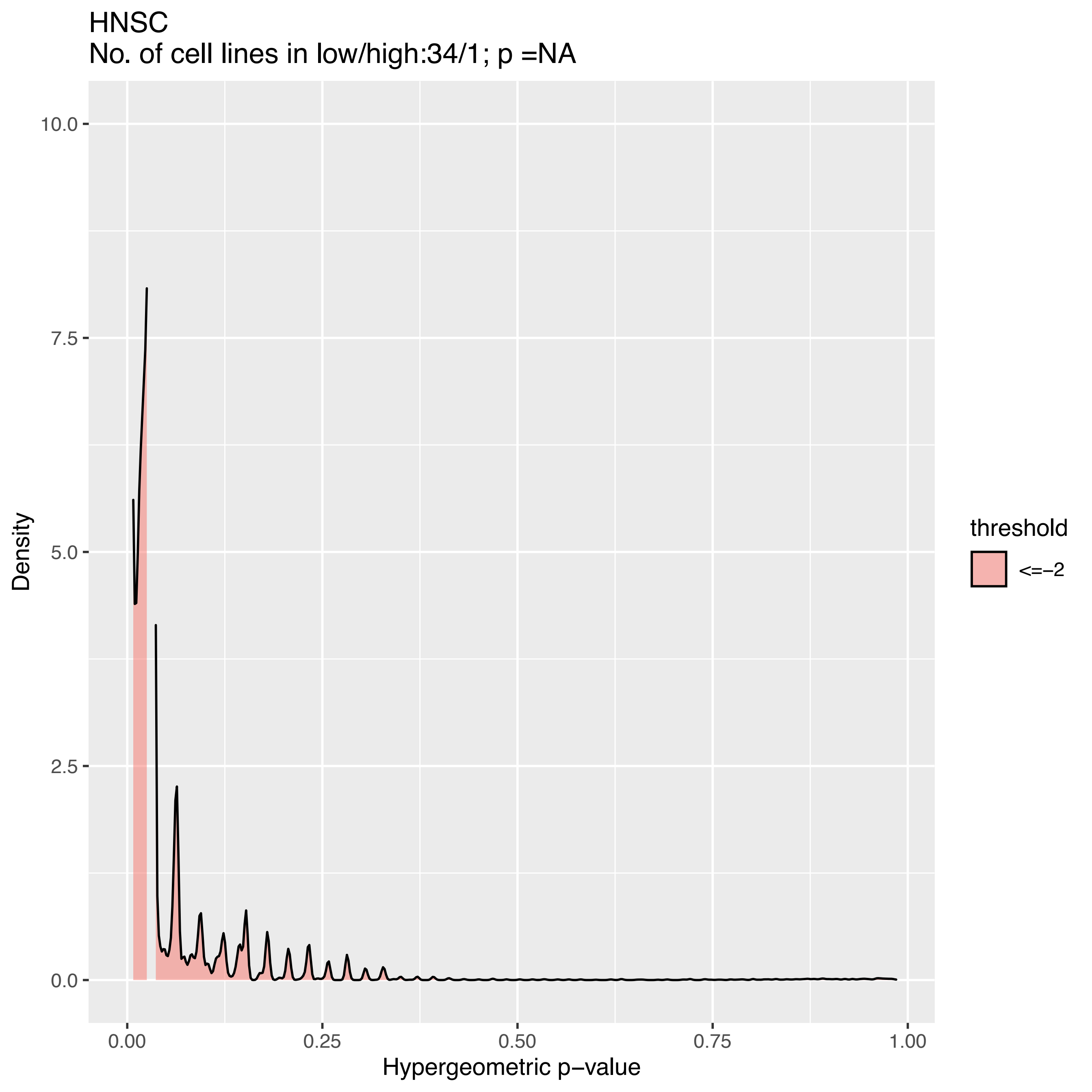
HNSC
No. of cell lines in low/high:34/1; p =NA
Density
10.0
7.5
5.0
2.5
0.0
0.00
0.25
0.50
0.75
1.00
Hypergeometric p−value
threshold
<=−2

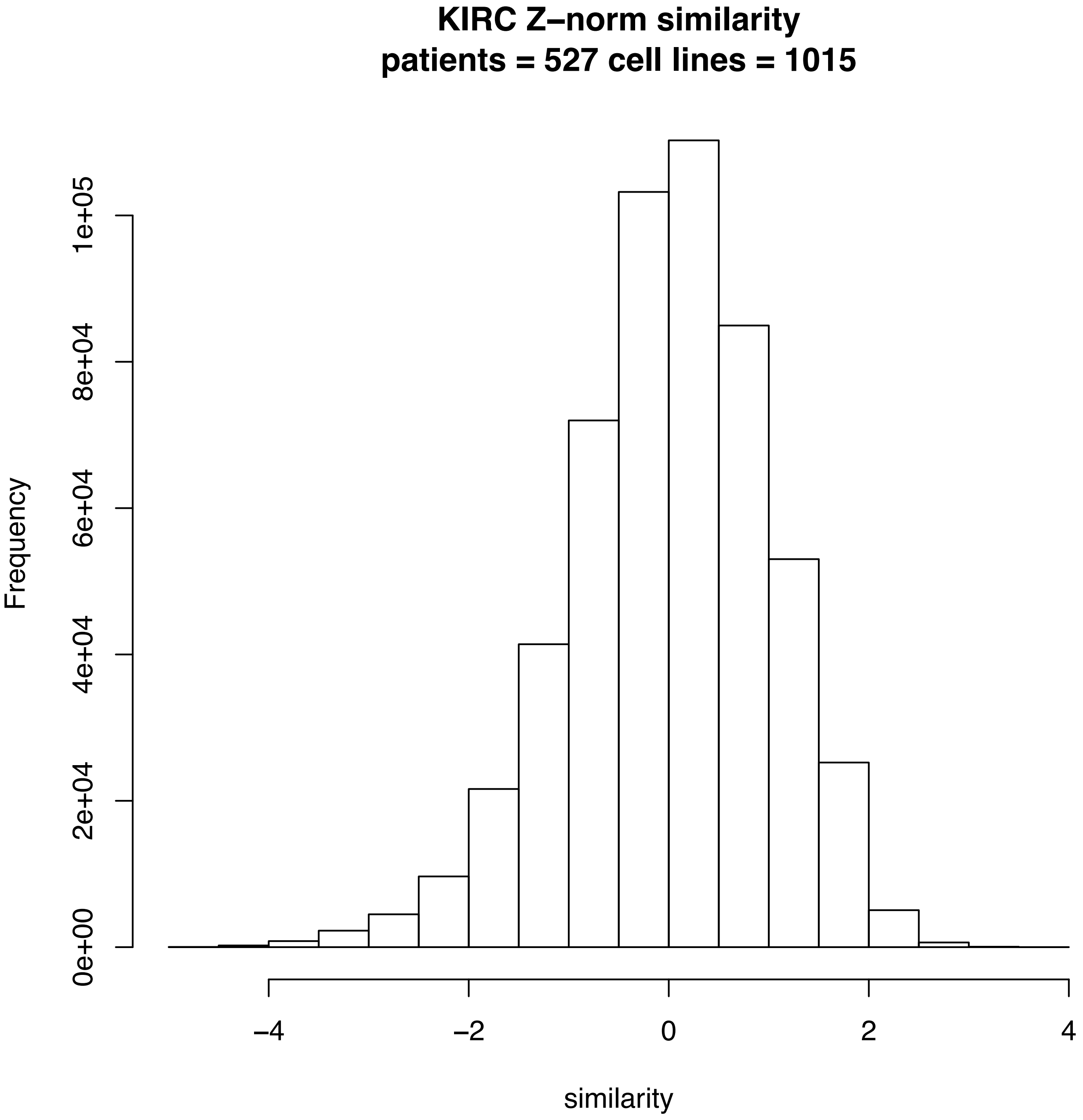

KIRC Z−norm similarity
patients = 527 cell lines = 1015
Frequency
0e+00
2e+04
4e+04
6e+04
8e+04
1e+05
−4
−2
0
2
4
similarity

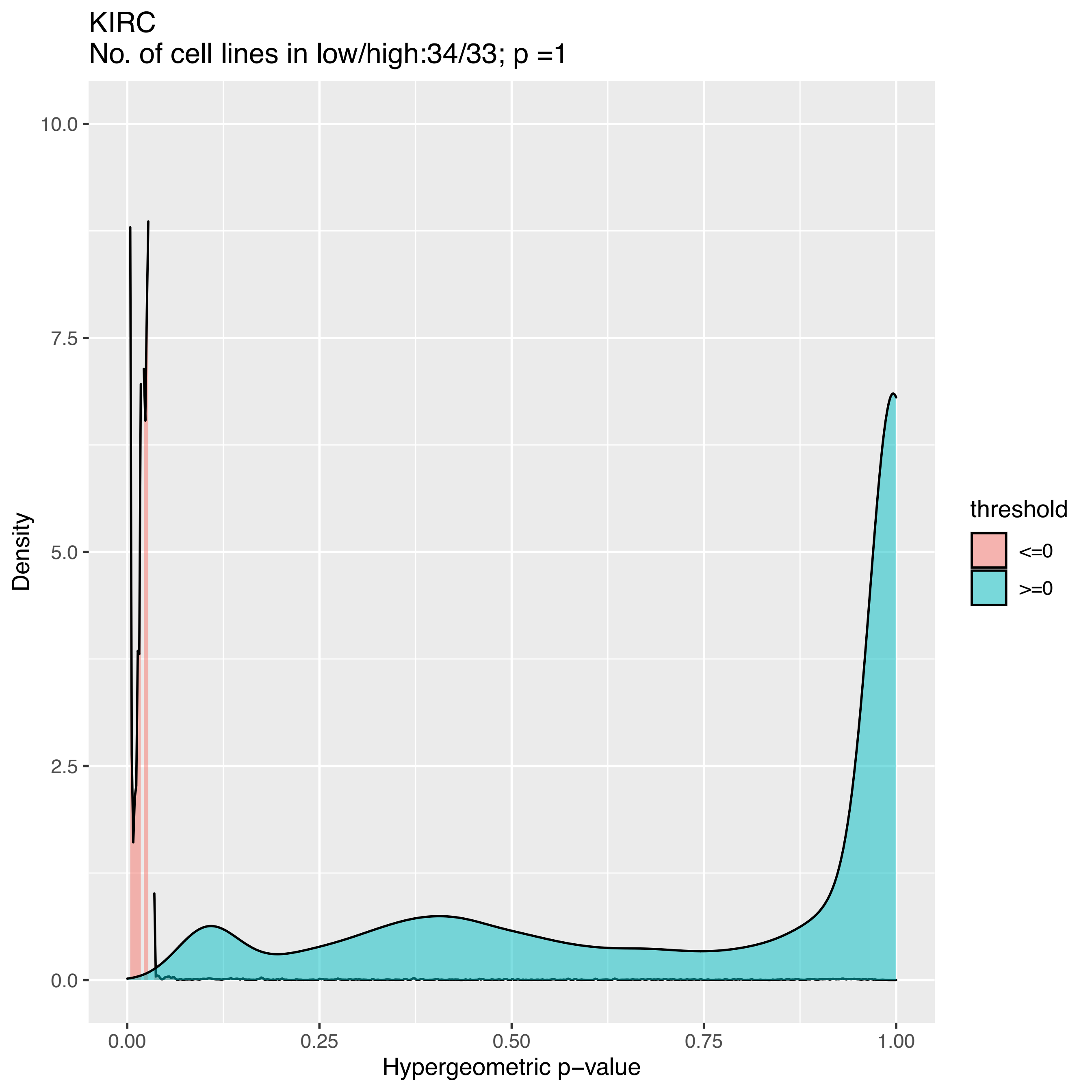
KIRC
No. of cell lines in low/high:34/33; p =1
10.0
7.5
5.0
2.5
0.0
Density
0.00
0.25
0.50
0.75
1.00
Hypergeometric p−value
threshold
<=0
>=0

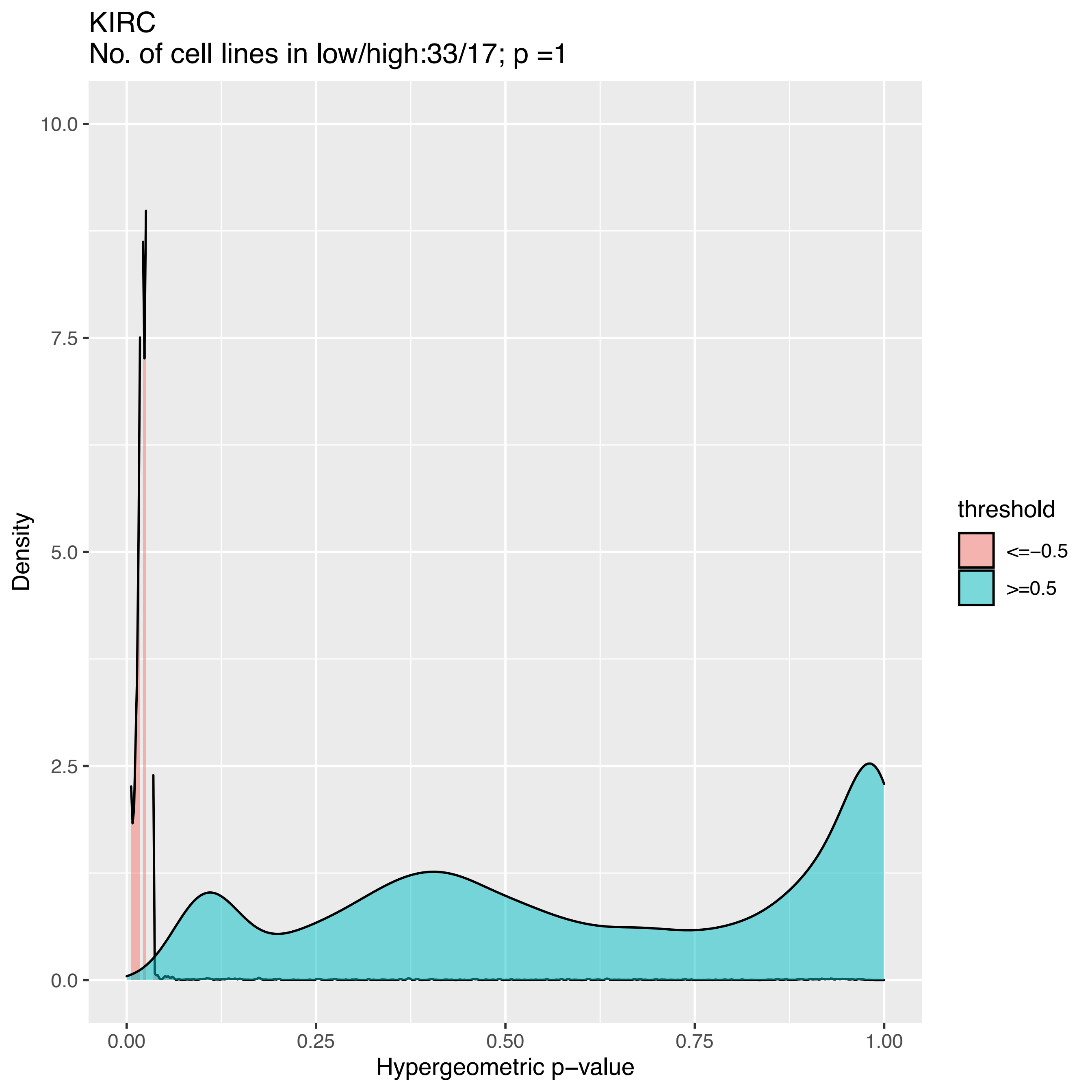
KIRC
No. of cell lines in low/high:33/17; p =1
Density
10.0
7.5
5.0
2.5
0.0
0.00
0.25
0.50
0.75
1.00
Hypergeometric p−value
threshold
<=−0.5
>=0.5

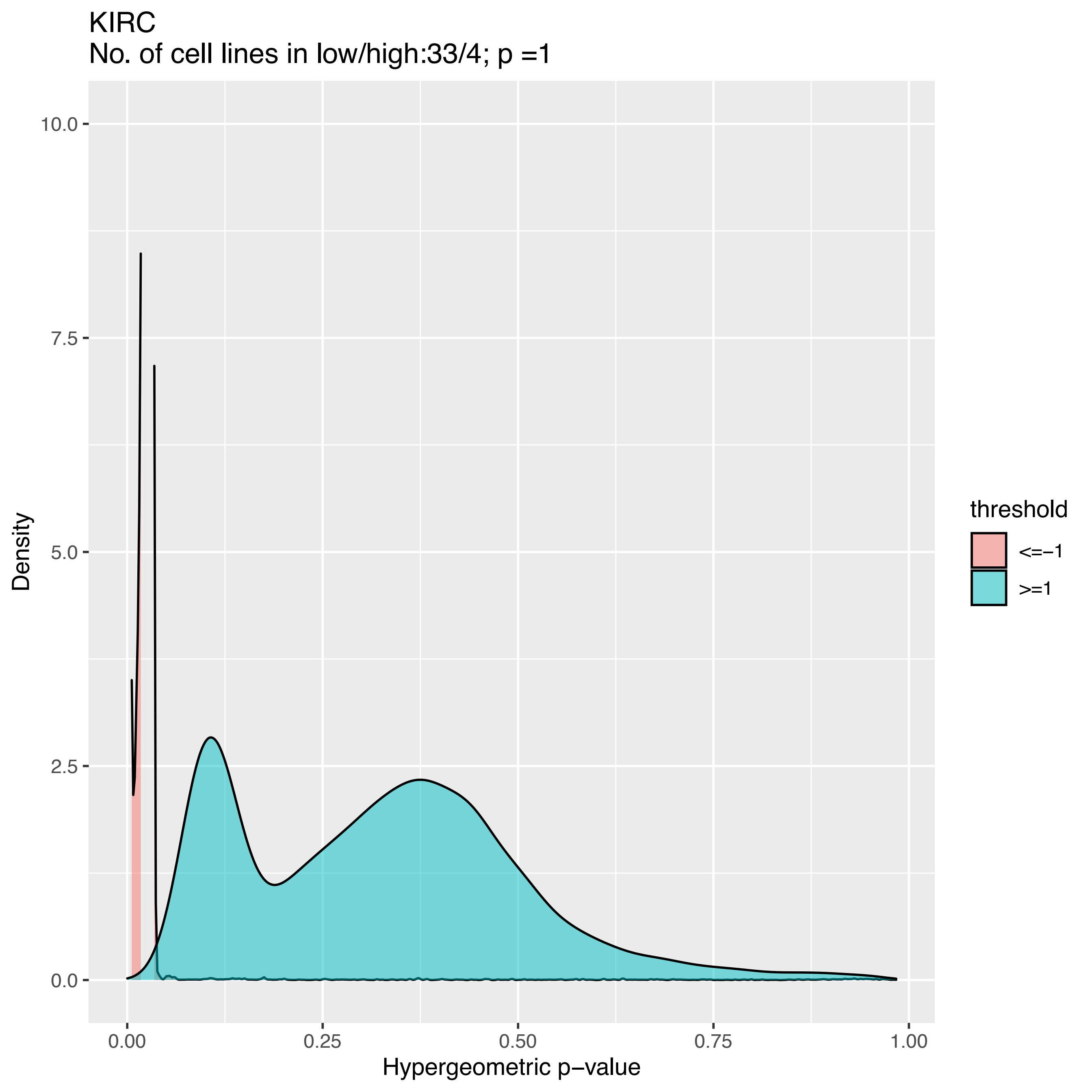
KIRC
No. of cell lines in low/high:33/4; p =1
Density
10.0
7.5
5.0
2.5
0.0
0.00
0.25
0.50
0.75
1.00
Hypergeometric p−value
threshold
<=−1
>=1

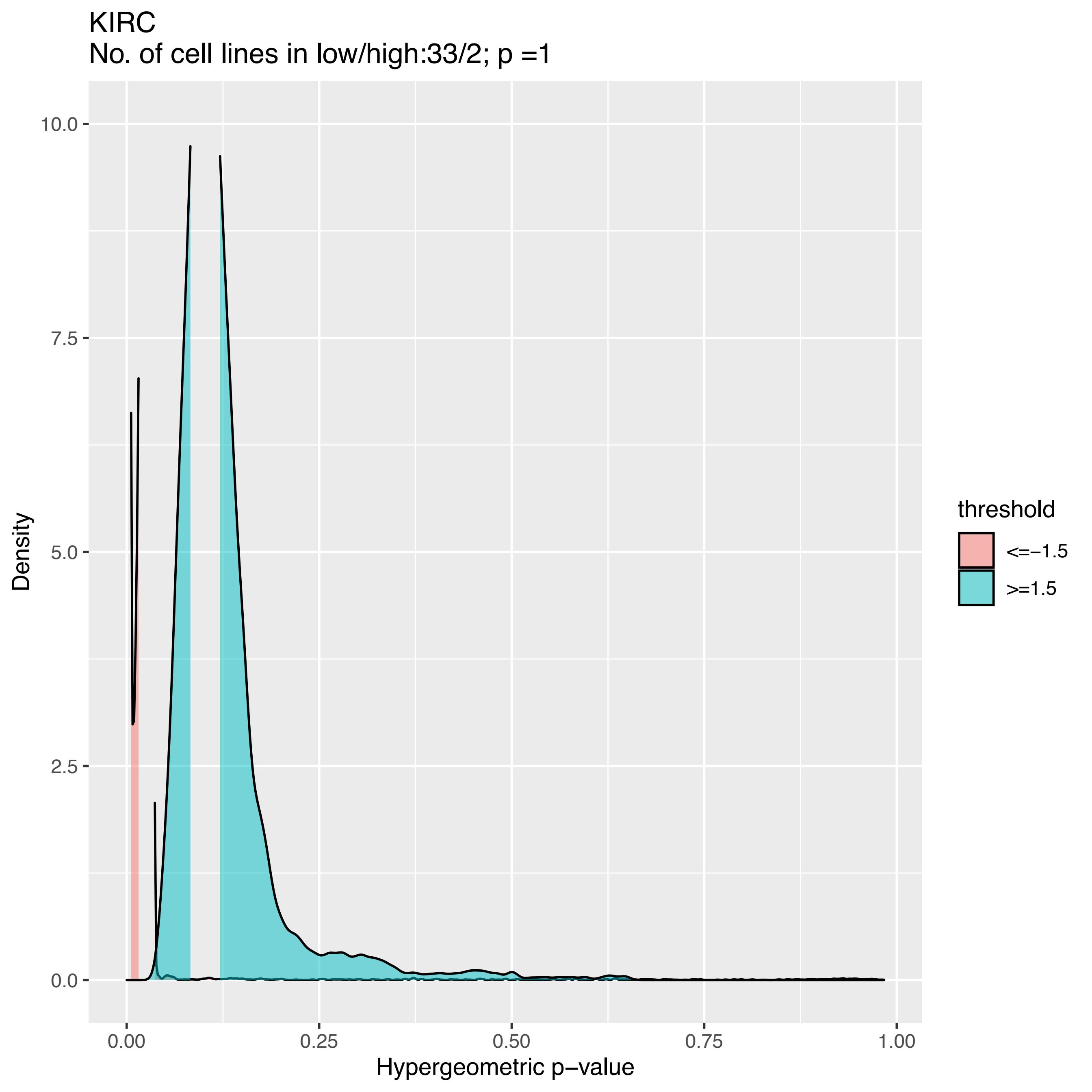
KIRC
No. of cell lines in low/high:33/2; p =1
10.0
7.5
5.0
2.5
0.0
Density
0.00
0.25
0.50
0.75
1.00
Hypergeometric p−value
threshold
<=−1.5
>=1.5

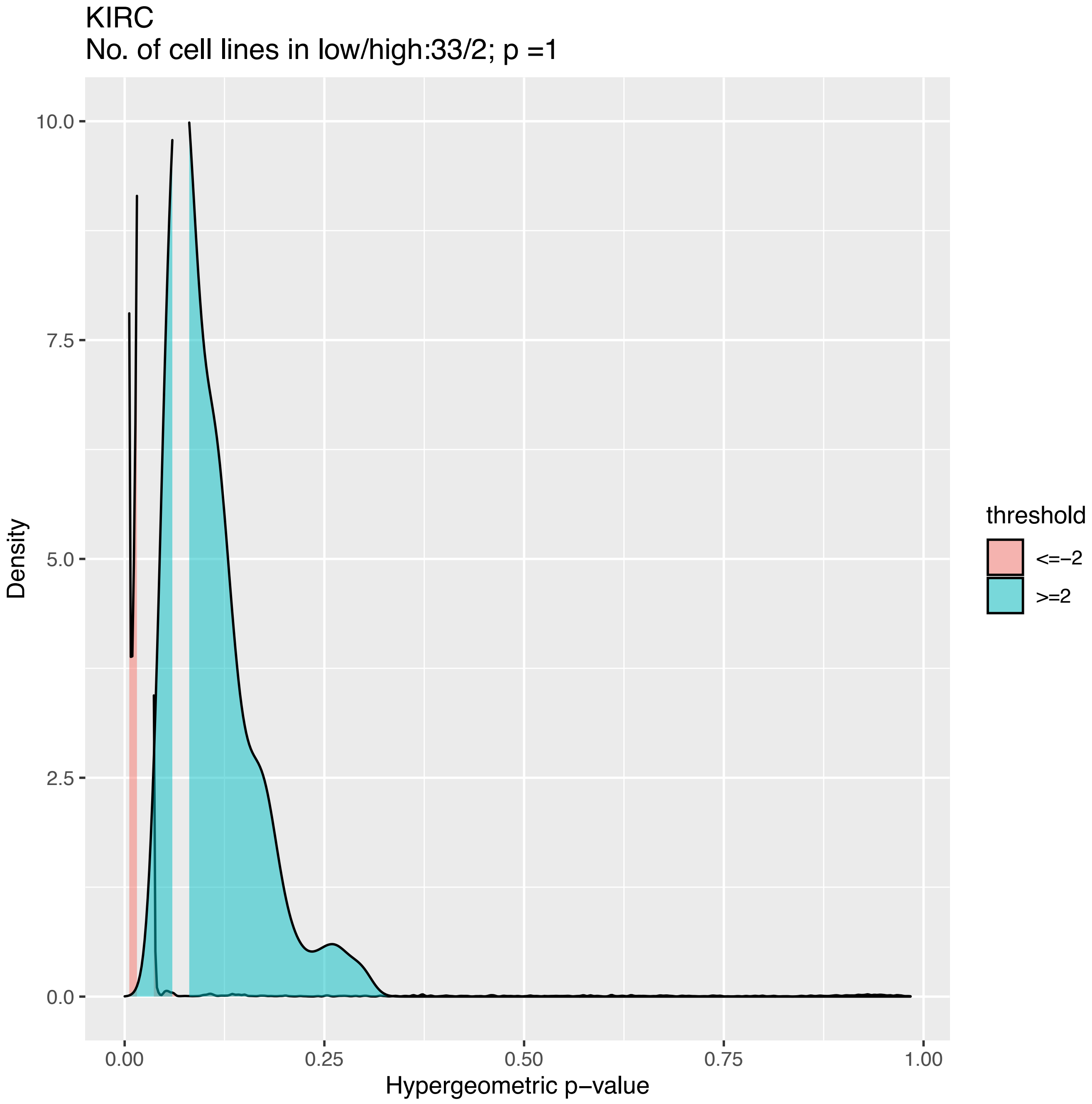
KIRC
No. of cell lines in low/high:33/2; p =1
Density
10.0
7.5
5.0
2.5
0.0
0.00
0.25
0.50
0.75
1.00
Hypergeometric p−value
threshold
<=−2
>=2

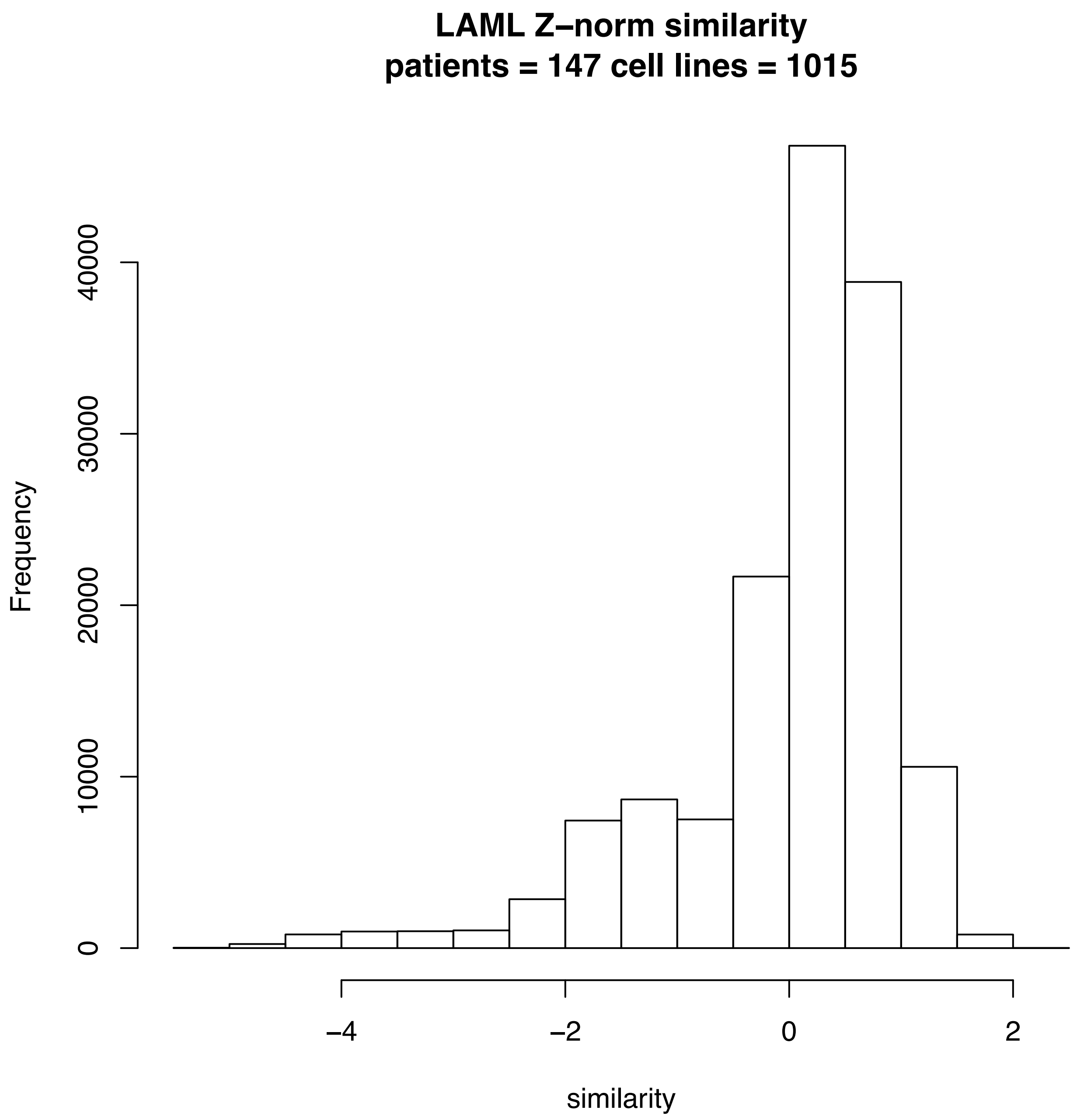
LAML Z–norm similarity
patients = 147 cell lines = 1015
Frequency
0
10000
20000
30000
40000
−4
−2
0
2
similarity

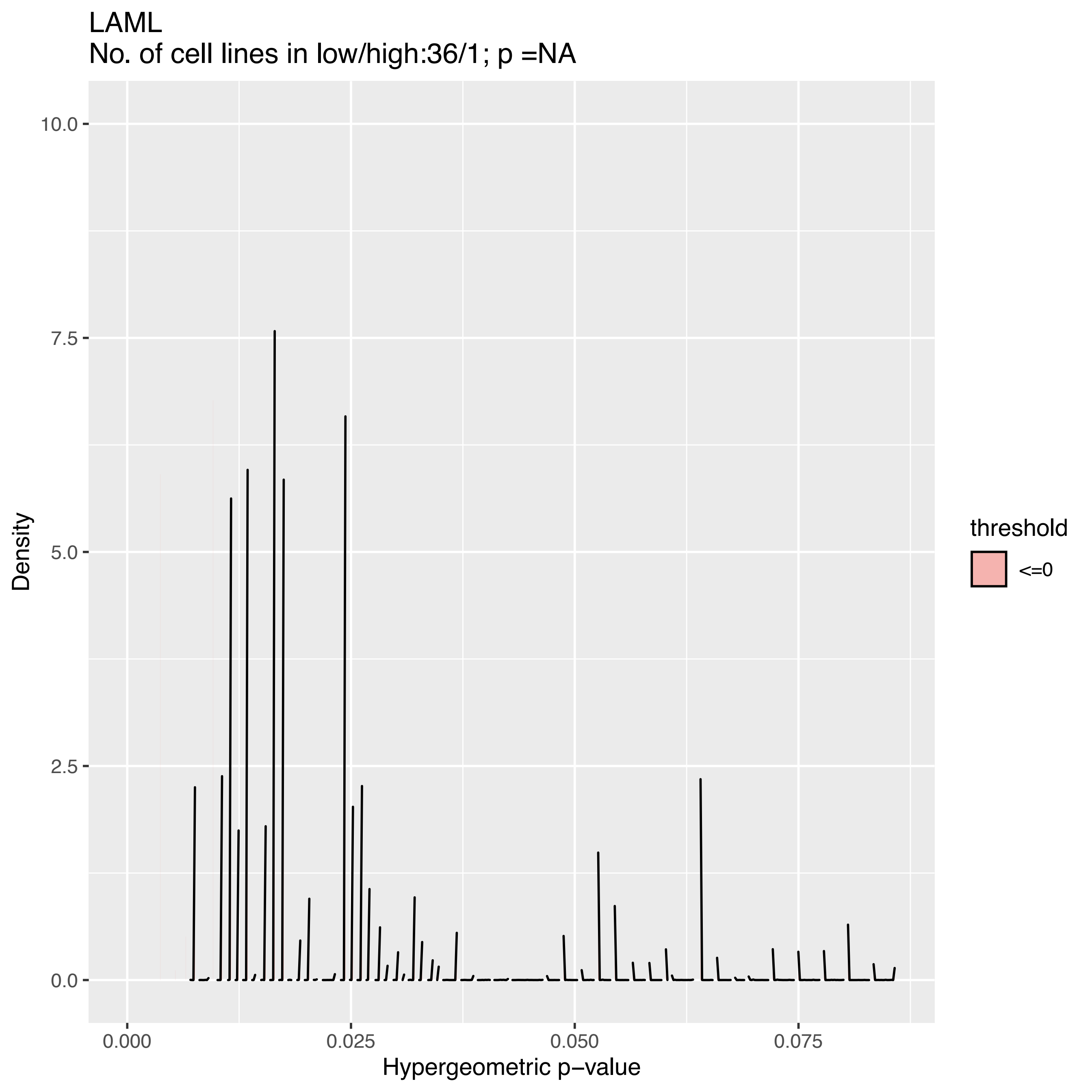
LAML
No. of cell lines in low/high:36/1; p =NA
Density
10.0
7.5
5.0
2.5
0.0
0.000
0.025
0.050
0.075
Hypergeometric p−value
threshold
<=0

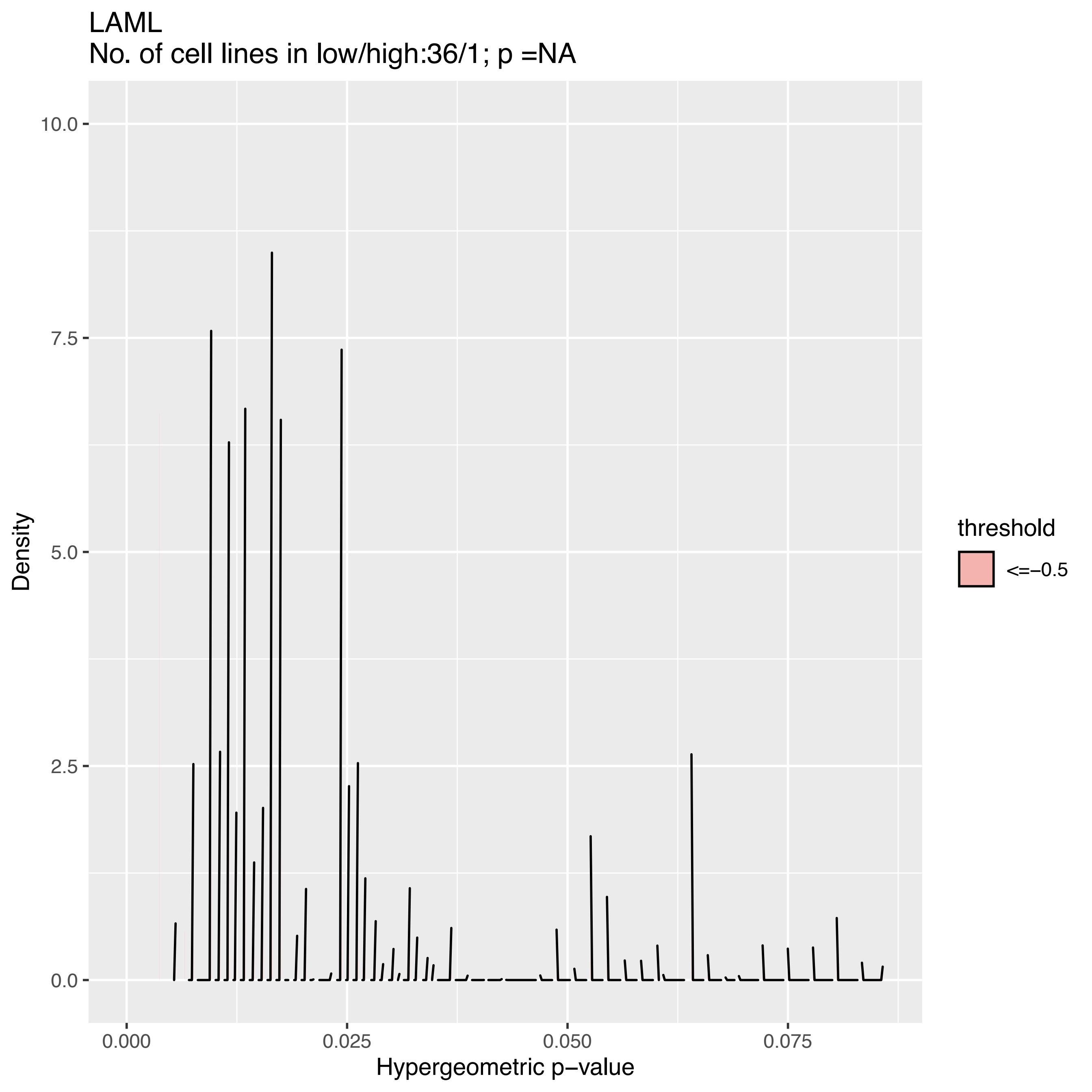
LAML
No. of cell lines in low/high:36/1; p =NA
Density
10.0
7.5
5.0
2.5
0.0
0.000
0.025
0.050
0.075
Hypergeometric p−value
threshold
<=−0.5

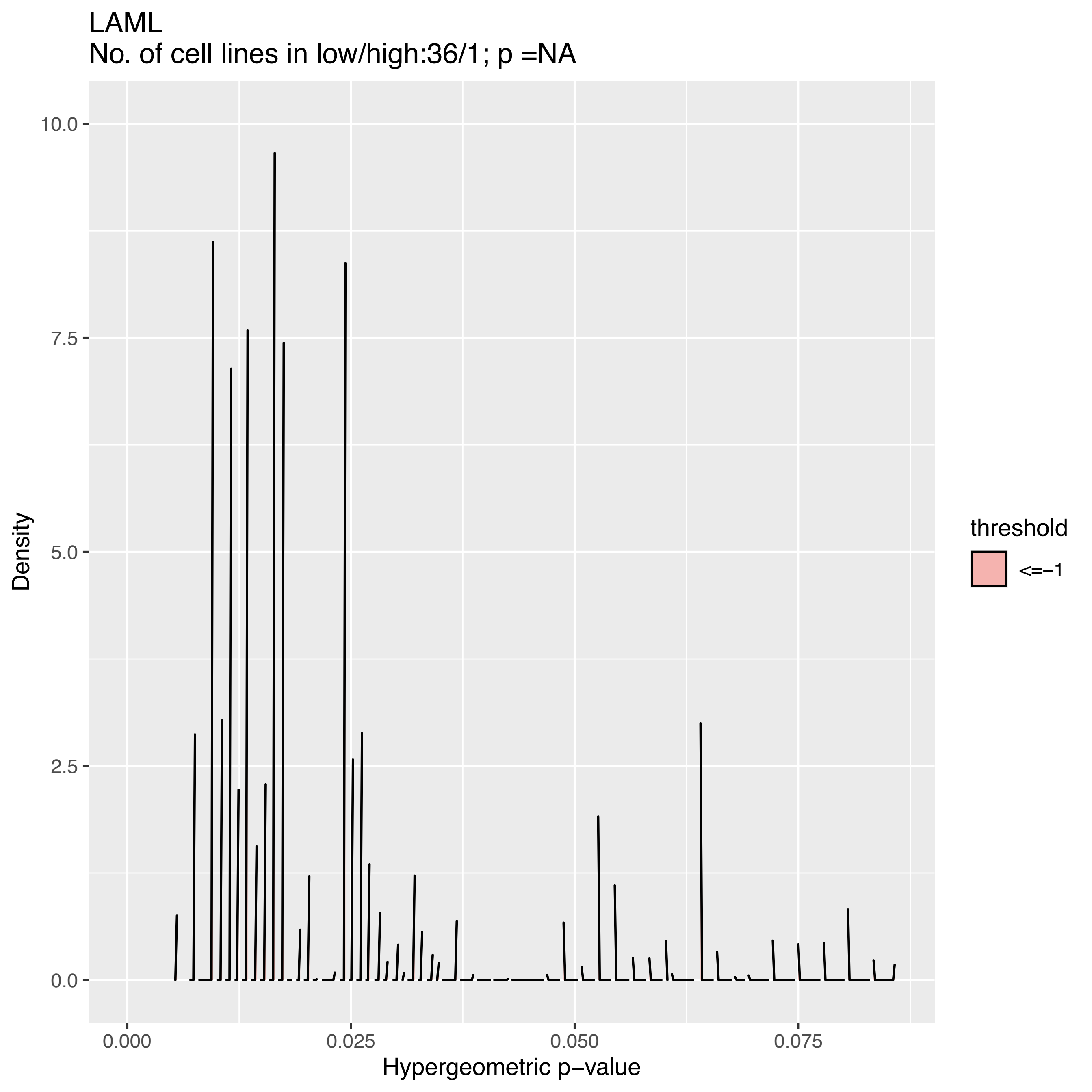

LAML
No. of cell lines in low/high:36/1; p =NA
Density
Hypergeometric p−value
threshold
<=−1
0.0
2.5
5.0
7.5
10.0
0.000
0.025
0.050
0.075

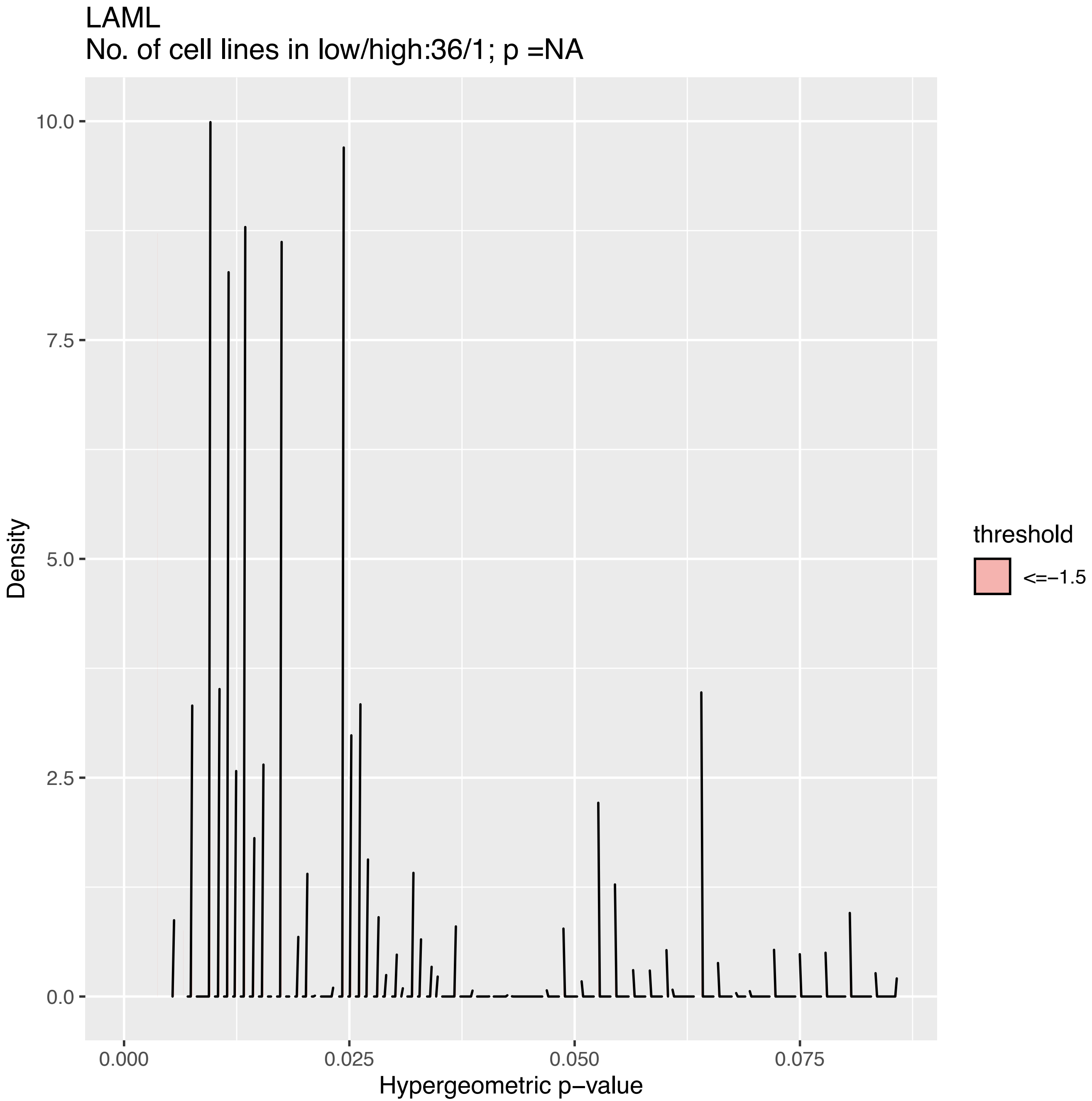

LAML
No. of cell lines in low/high:36/1; p =NA
Density
10.0
7.5
5.0
2.5
0.0
0.000
0.025
0.050
0.075
Hypergeometric p−value
threshold
<=−1.5

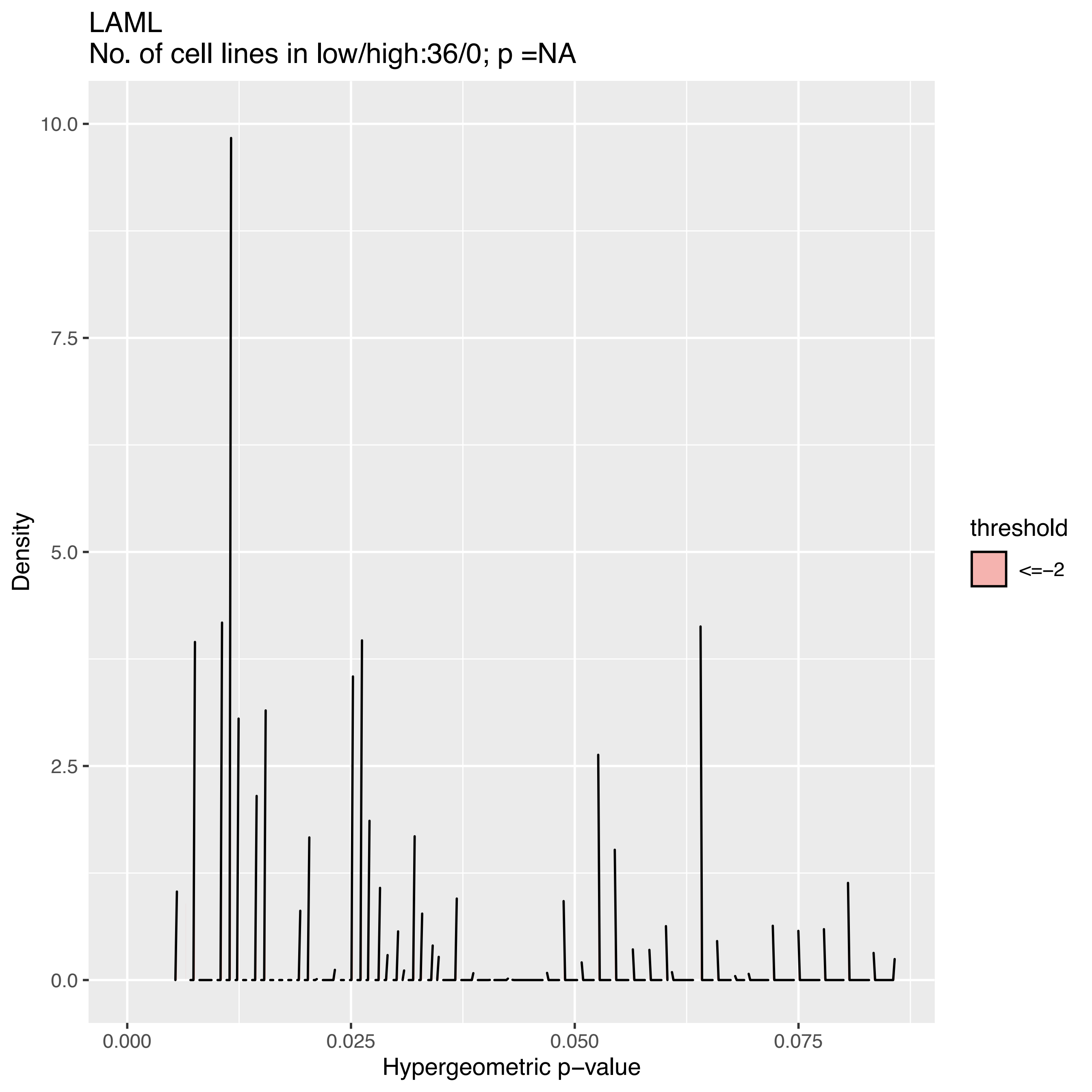
LAML
No. of cell lines in low/high:36/0; p =NA
Density
10.0
7.5
5.0
2.5
0.0
0.000
0.025
0.050
0.075
Hypergeometric p−value
threshold
<=−2

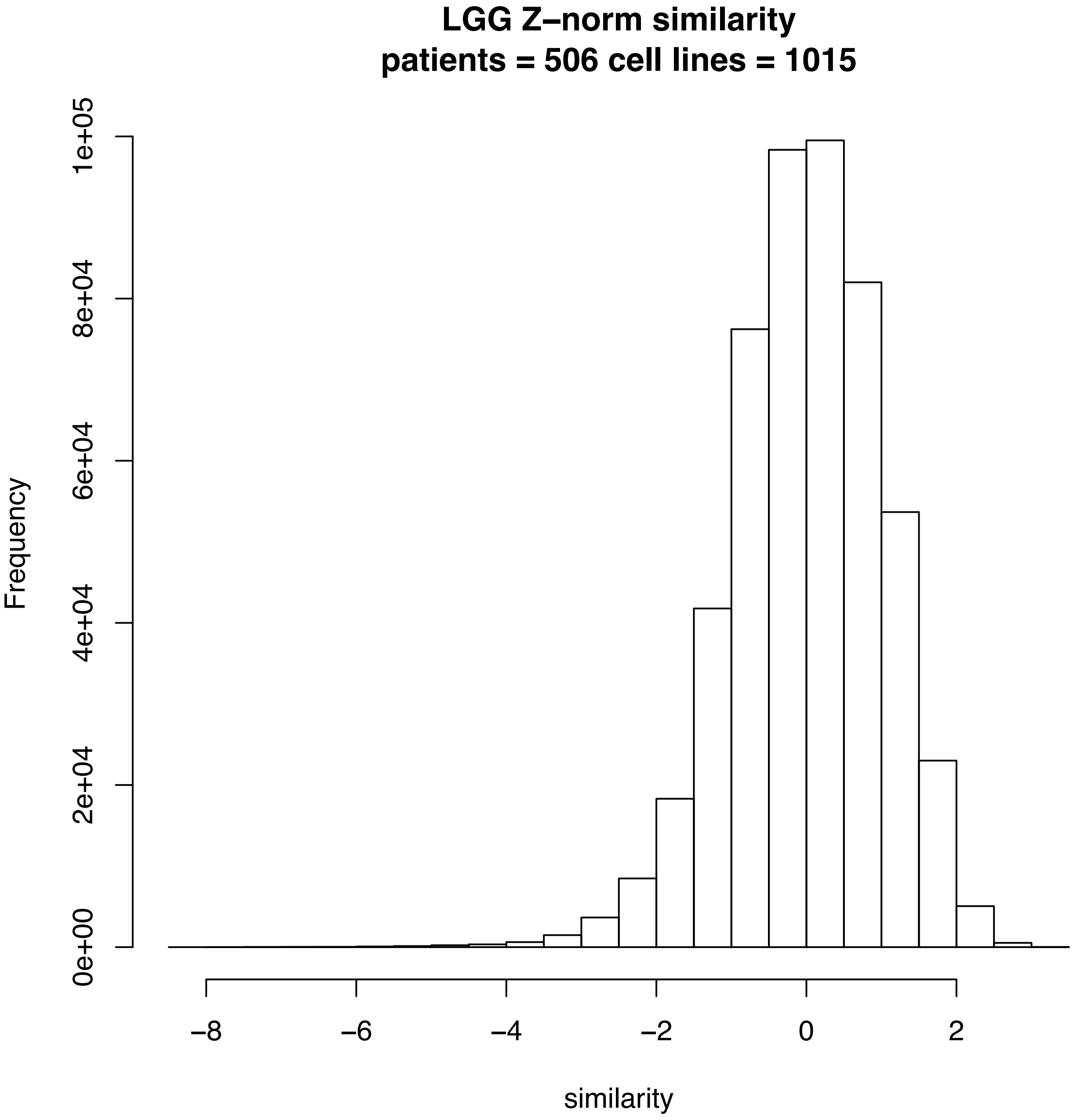

LGG Z–norm similarity
patients = 506 cell lines = 1015
Frequency
0e+00
2e+04
4e+04
6e+04
8e+04
1e+05
−8
−6
−4
−2
0
2
similarity

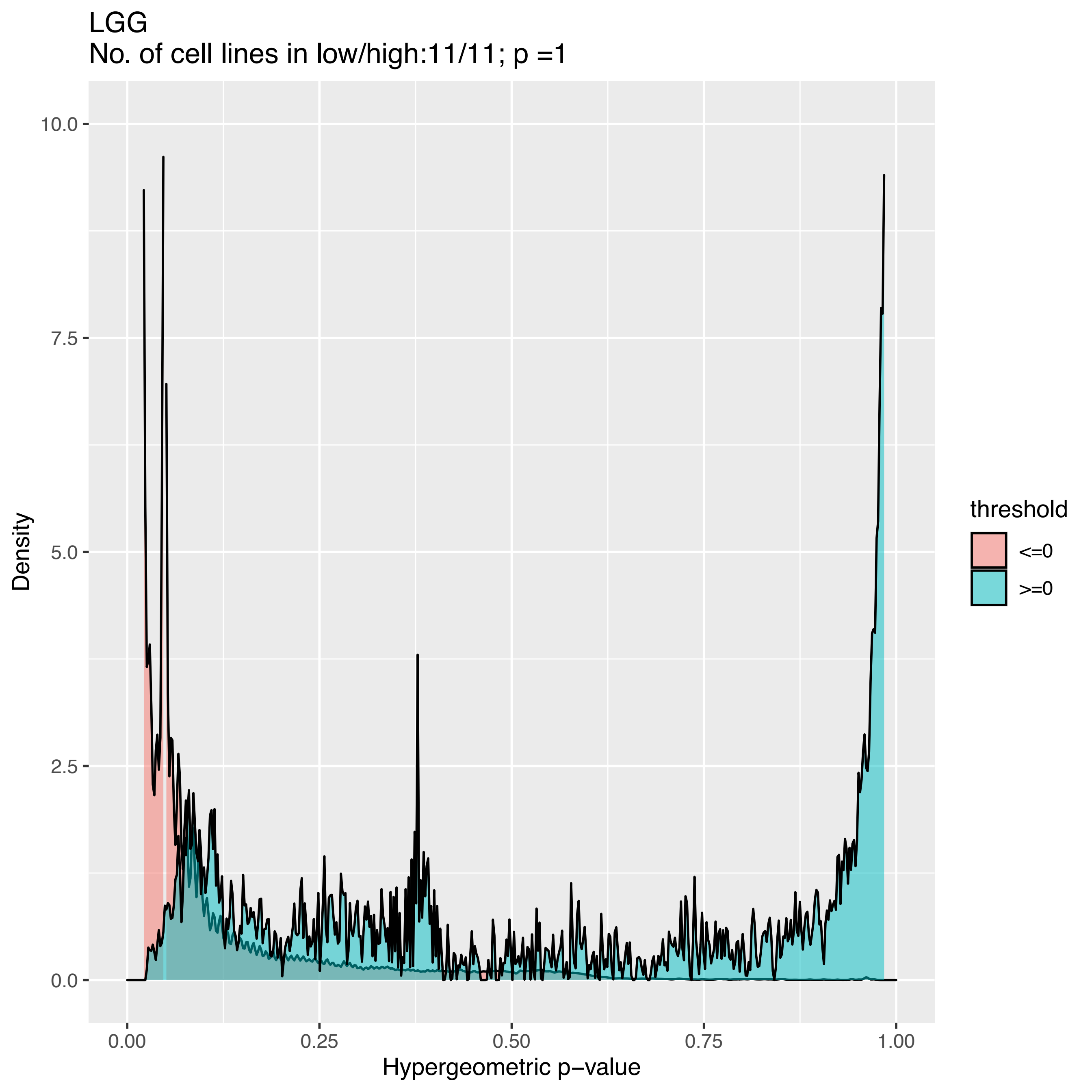
LGG
No. of cell lines in low/high:11/11; p =1
Density
10.0
7.5
5.0
2.5
0.0
0.00
0.25
0.50
0.75
1.00
Hypergeometric p−value
threshold
<=0
>=0

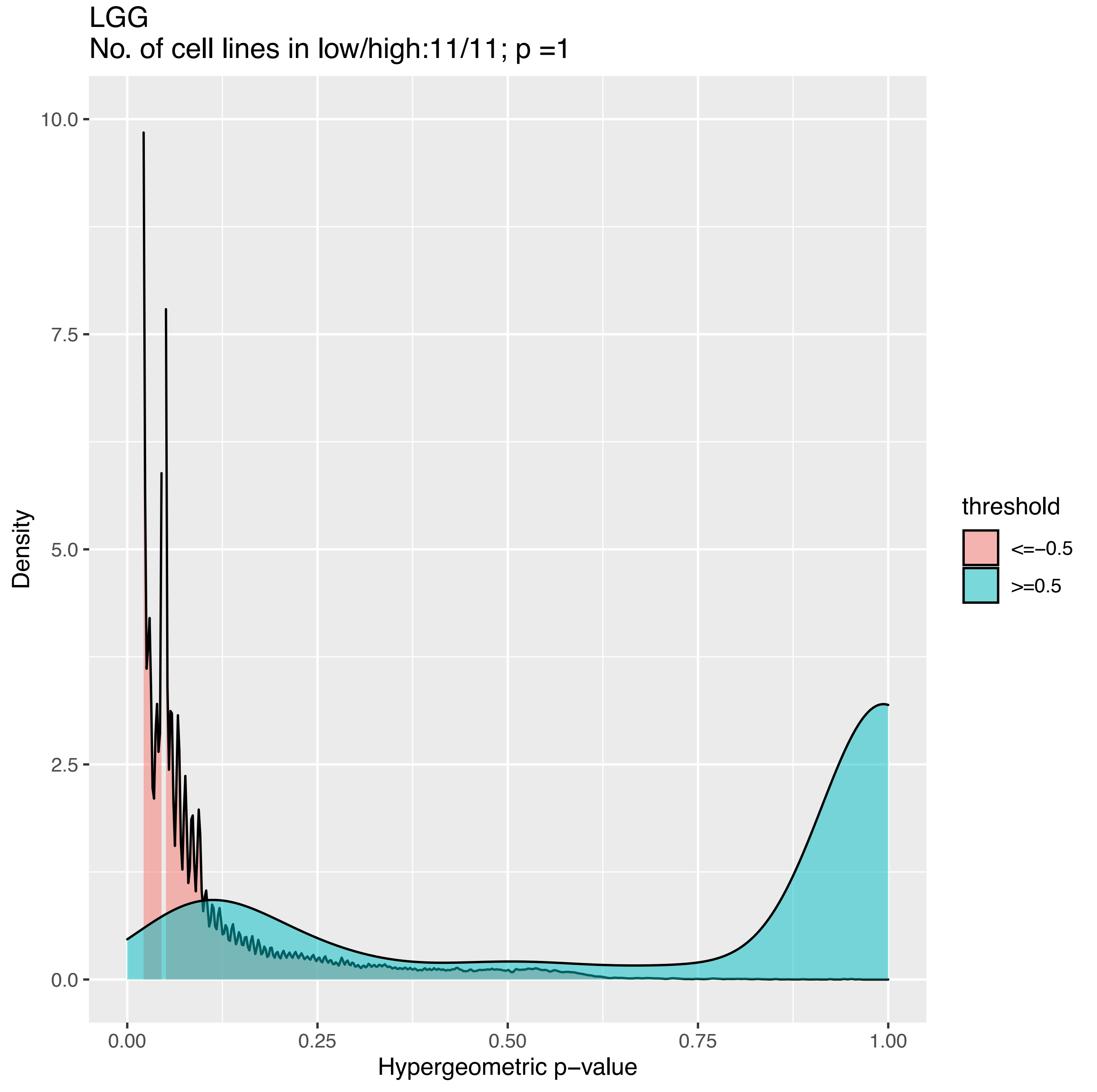
LGG
No. of cell lines in low/high:11/11; p =1
Density
10.0
7.5
5.0
2.5
0.0
0.00
0.25
0.50
0.75
1.00
Hypergeometric p−value
threshold
<=−0.5
>=0.5

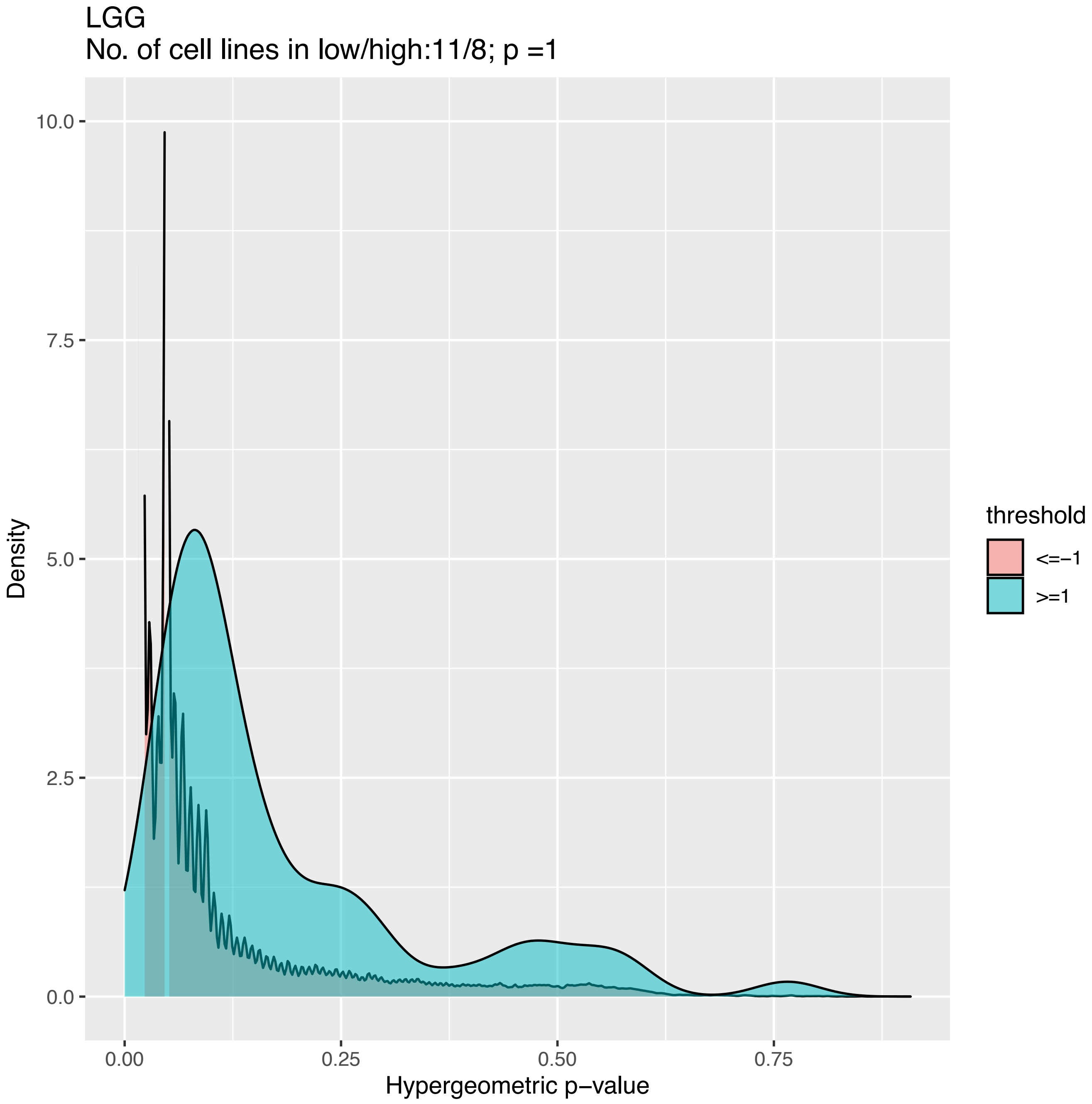
LGG
No. of cell lines in low/high:11/8; p =1
Density
10.0
7.5
5.0
2.5
0.0
0.00
0.25
0.50
0.75
Hypergeometric p−value
threshold
<=−1
>=1

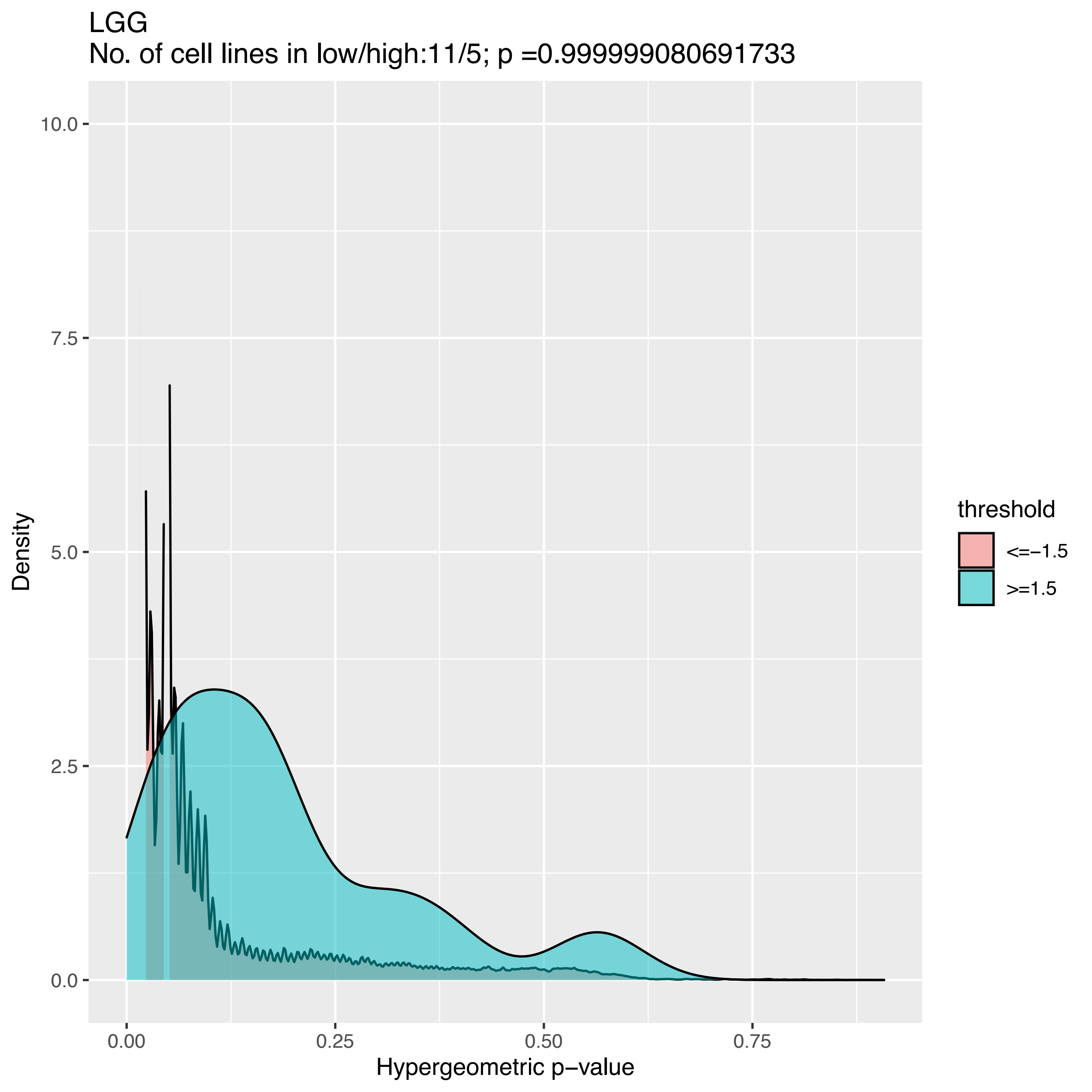

LGG
No. of cell lines in low/high:11/5; p =0.999999080691733
Density
10.0
7.5
5.0
2.5
0.0
0.00
0.25
0.50
0.75
Hypergeometric p−value
threshold
<=−1.5
>=1.5

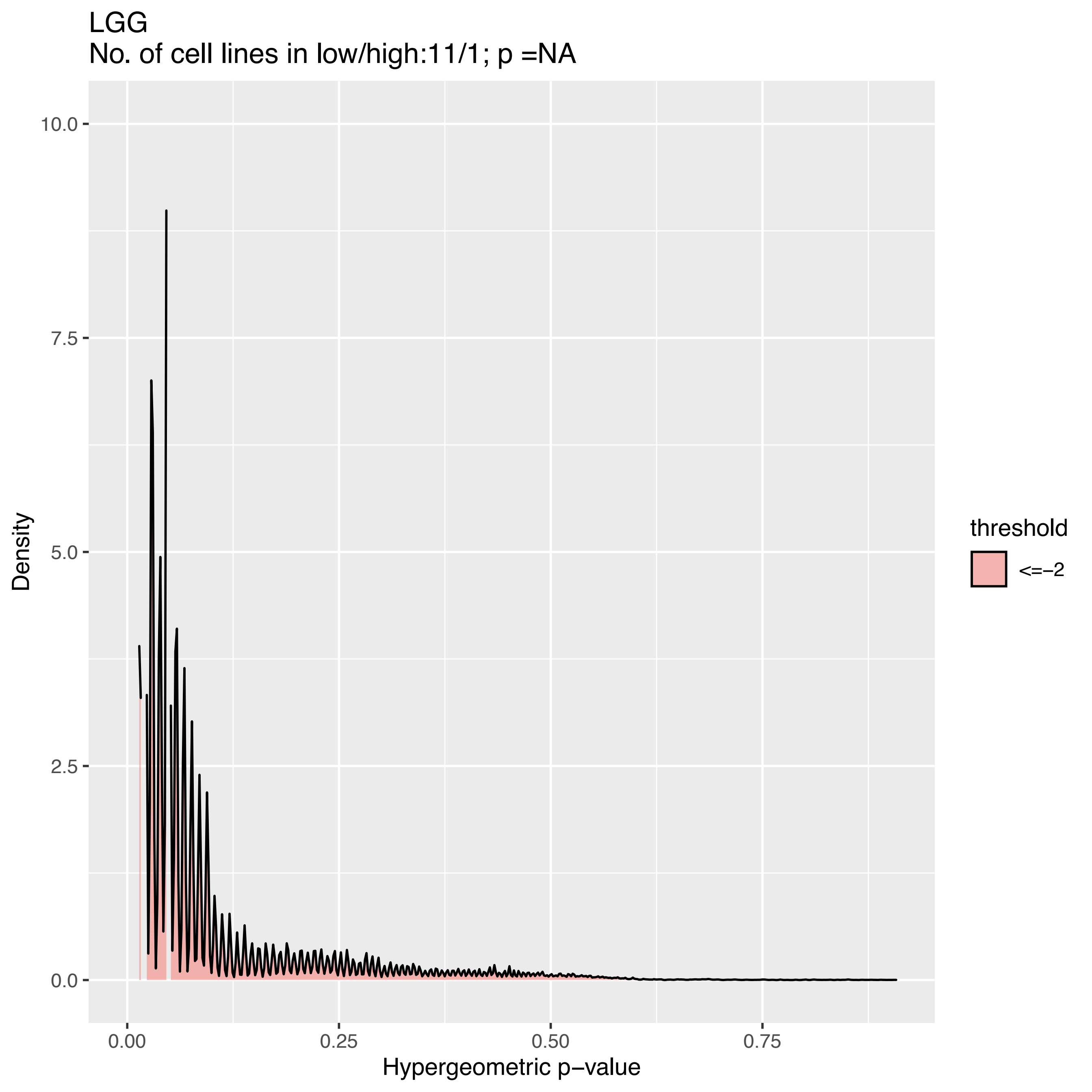

LGG
No. of cell lines in low/high:11/1; p =NA
Density
10.0
7.5
5.0
2.5
0.0
0.00
0.25
0.50
0.75
Hypergeometric p−value
threshold
<=−2

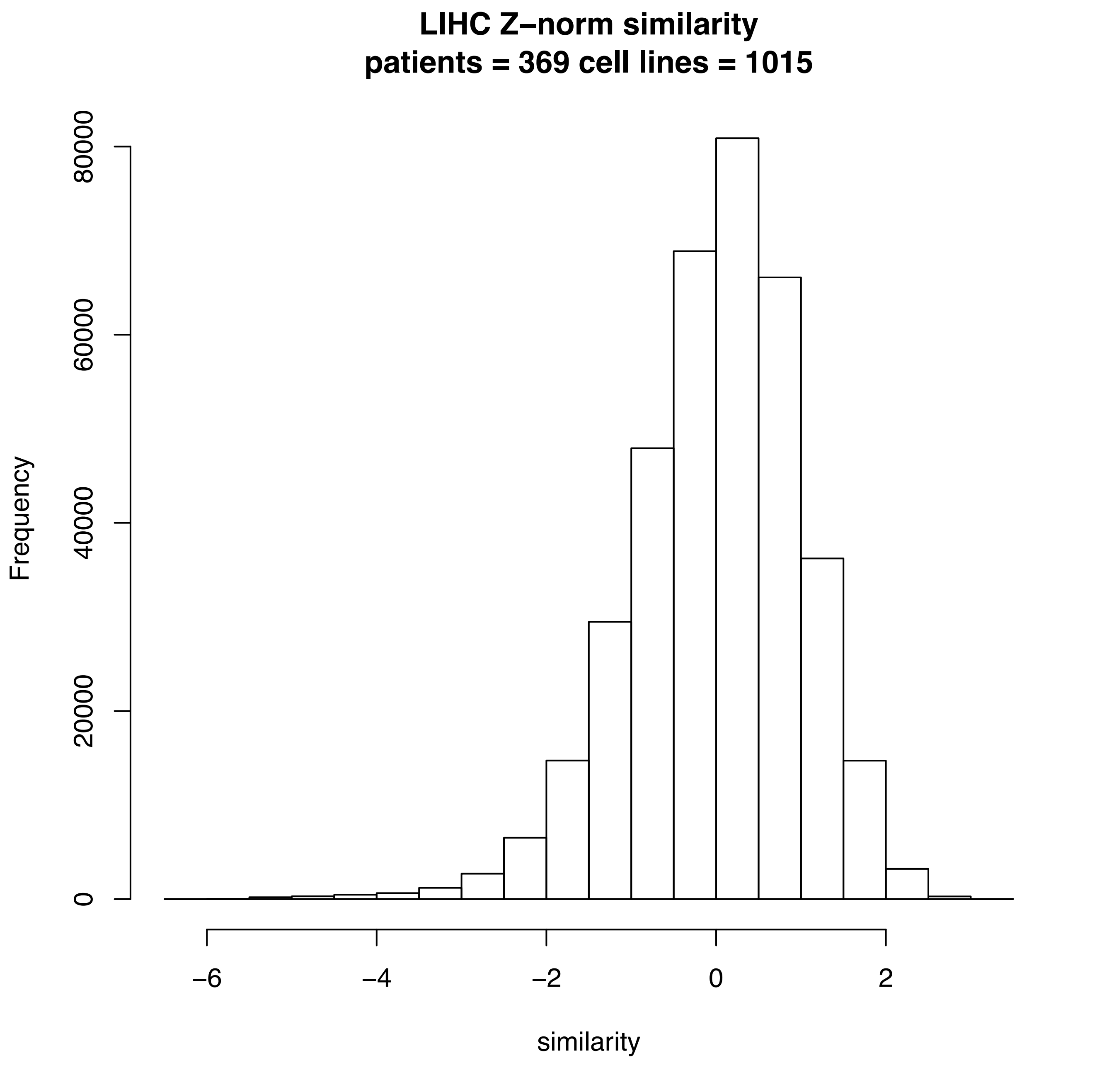
LIHC Z–norm similarity
patients = 369 cell lines = 1015
Frequency
0
20000
40000
60000
80000
−6
−4
−2
0
2
similarity

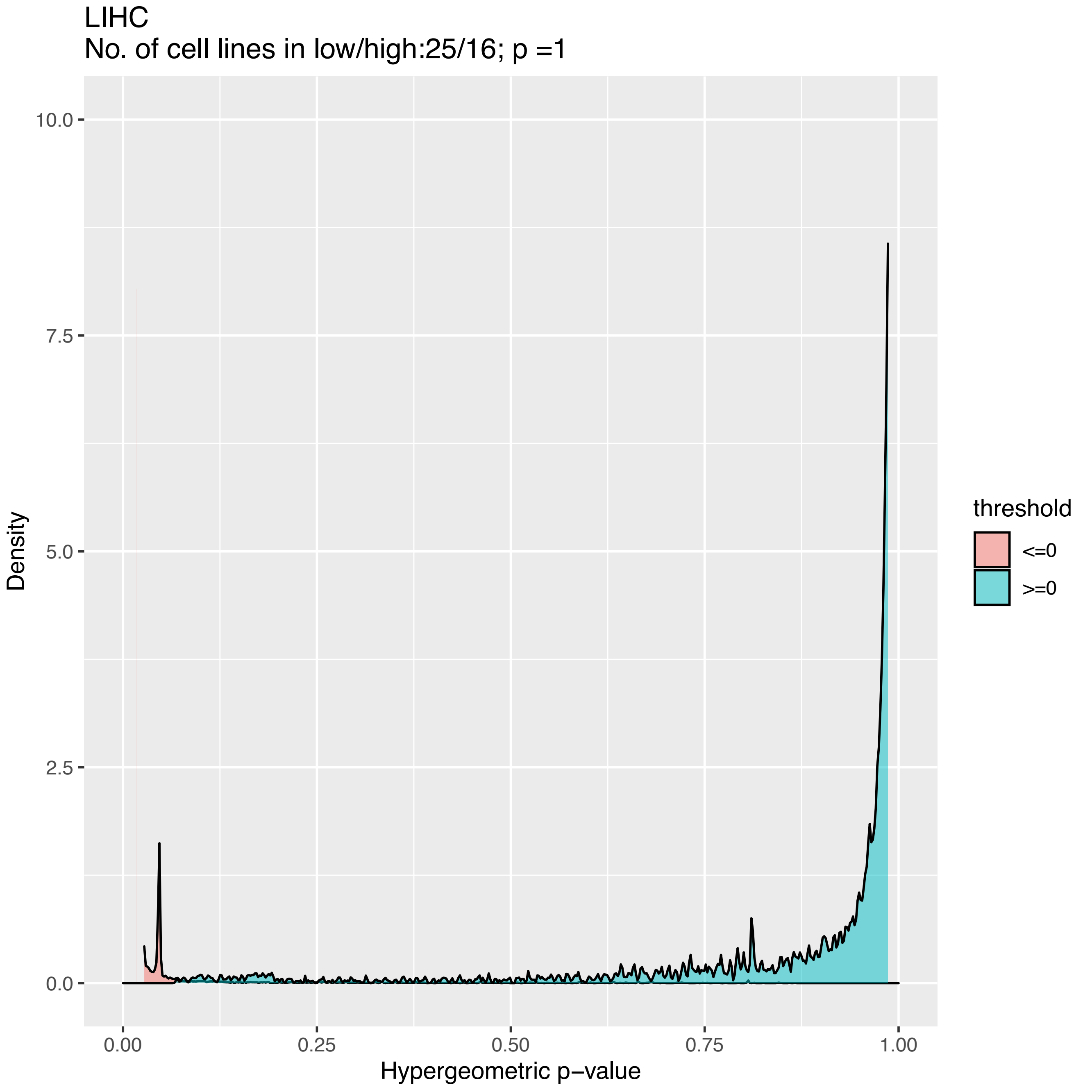
LIHC
No. of cell lines in low/high:25/16; p =1
Density
10.0
7.5
5.0
2.5
0.0
0.00
0.25
0.50
0.75
1.00
Hypergeometric p−value
threshold
<=0
>=0

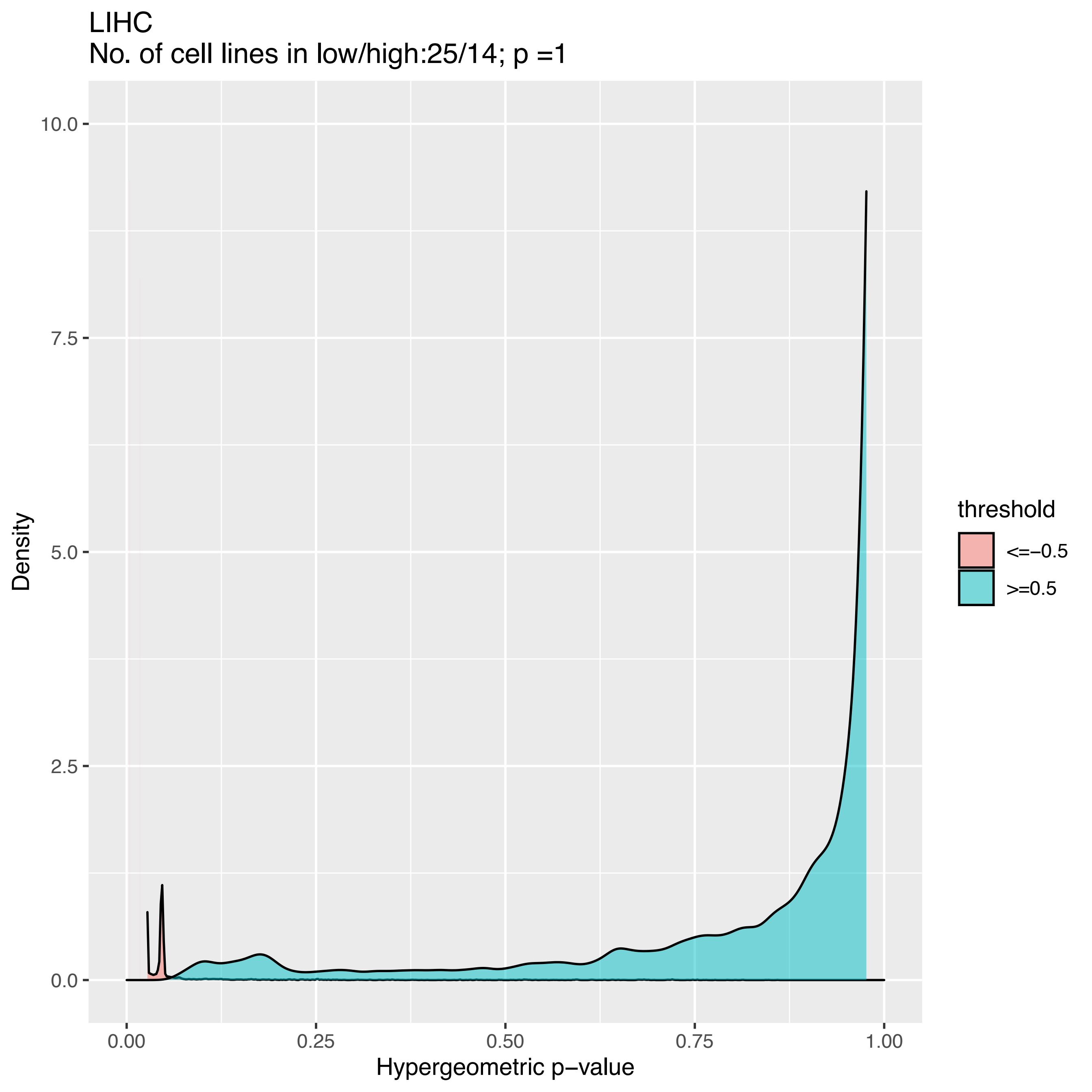
LIHC
No. of cell lines in low/high:25/14; p =1
Density
10.0
7.5
5.0
2.5
0.0
0.00
0.25
0.50
0.75
1.00
Hypergeometric p−value
threshold
<=−0.5
>=0.5

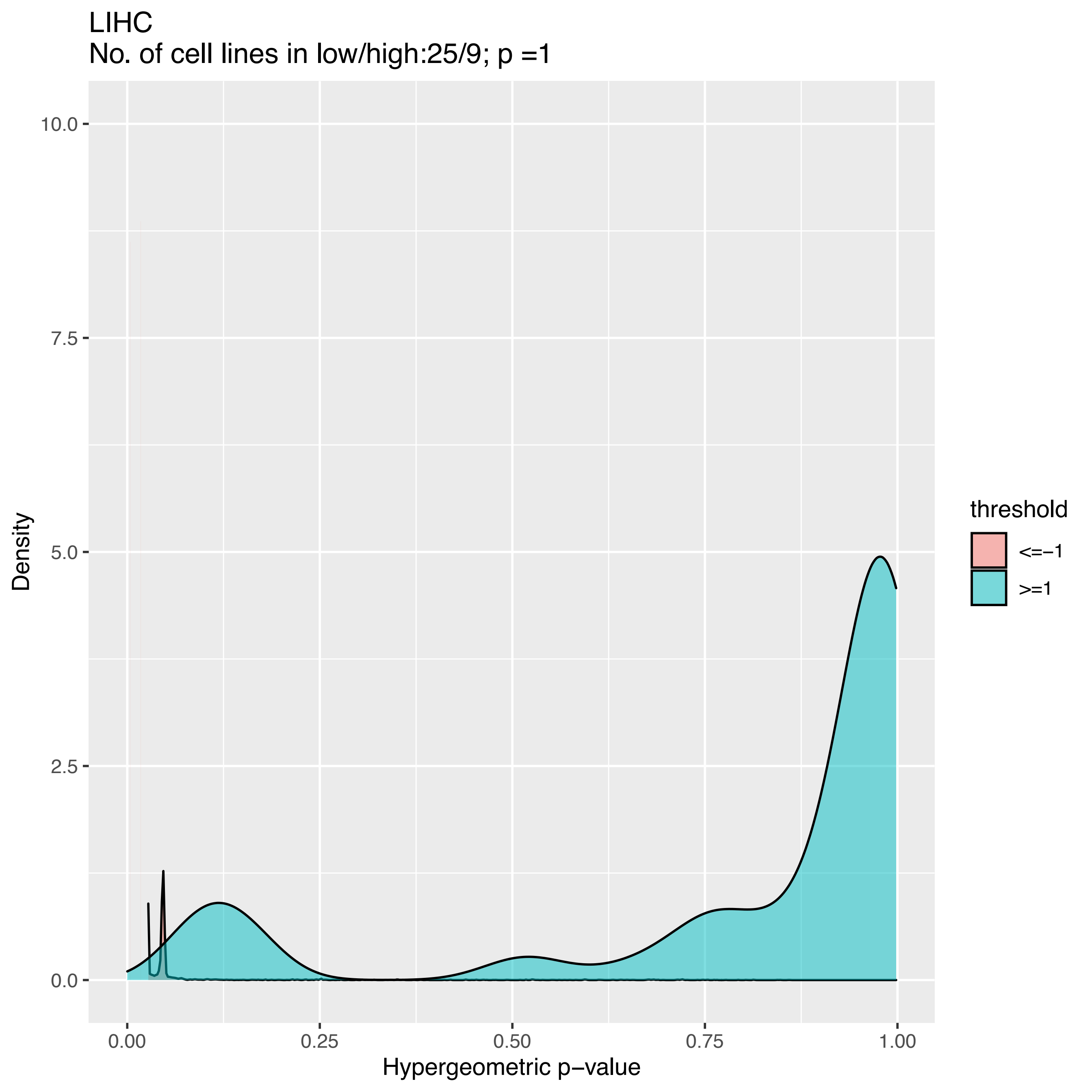
LIHC
No. of cell lines in low/high:25/9; p =1
10.0
7.5
5.0
2.5
0.0
Density
0.00
0.25
0.50
0.75
1.00
Hypergeometric p−value
threshold
<=−1
>=1

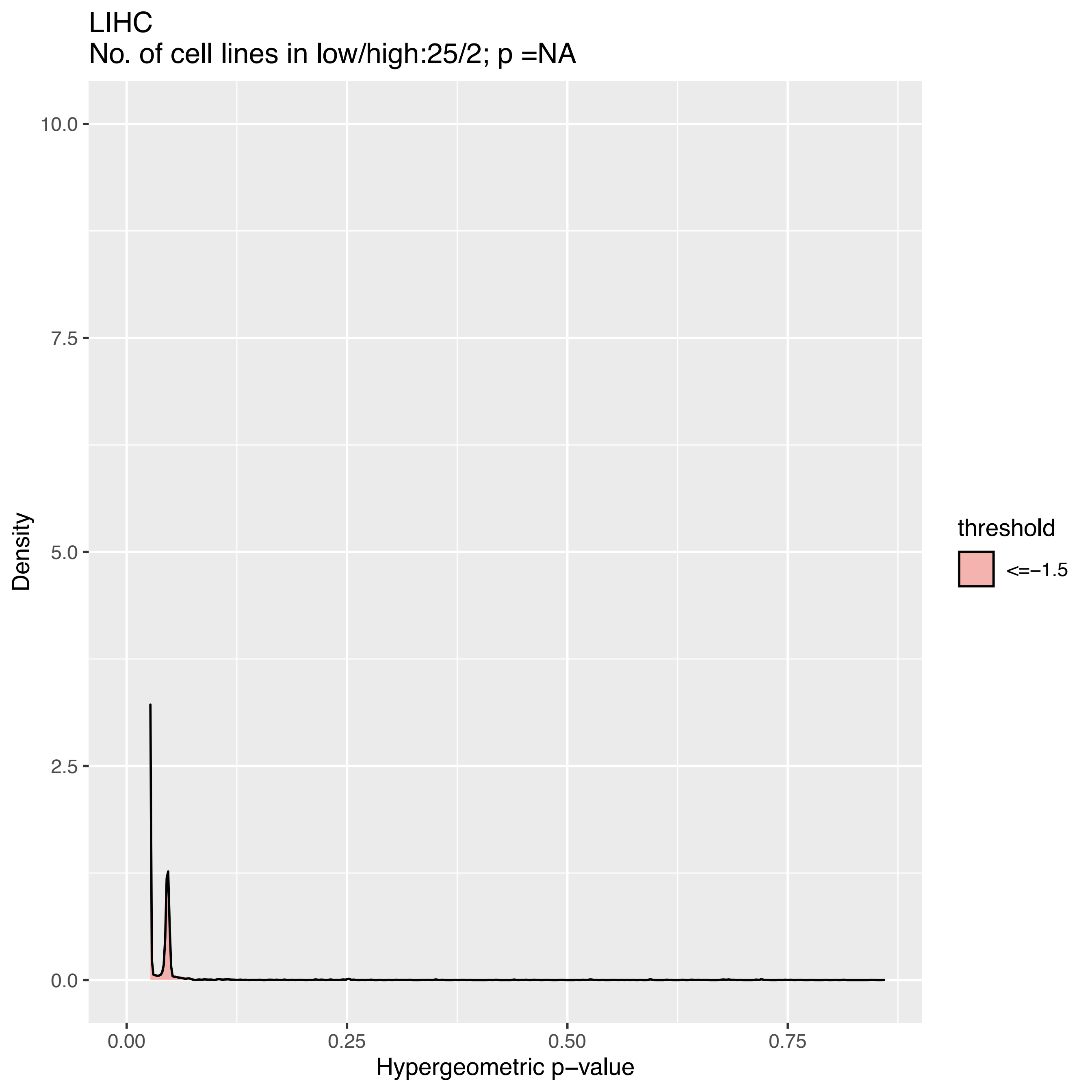

LIHC
No. of cell lines in low/high:25/2; p =NA
10.0
7.5
5.0
2.5
0.0
Density
0.00
0.25
0.50
0.75
Hypergeometric p−value
threshold
<=−1.5

LIHC
No. of cell lines in low/high:25/1; p =NA

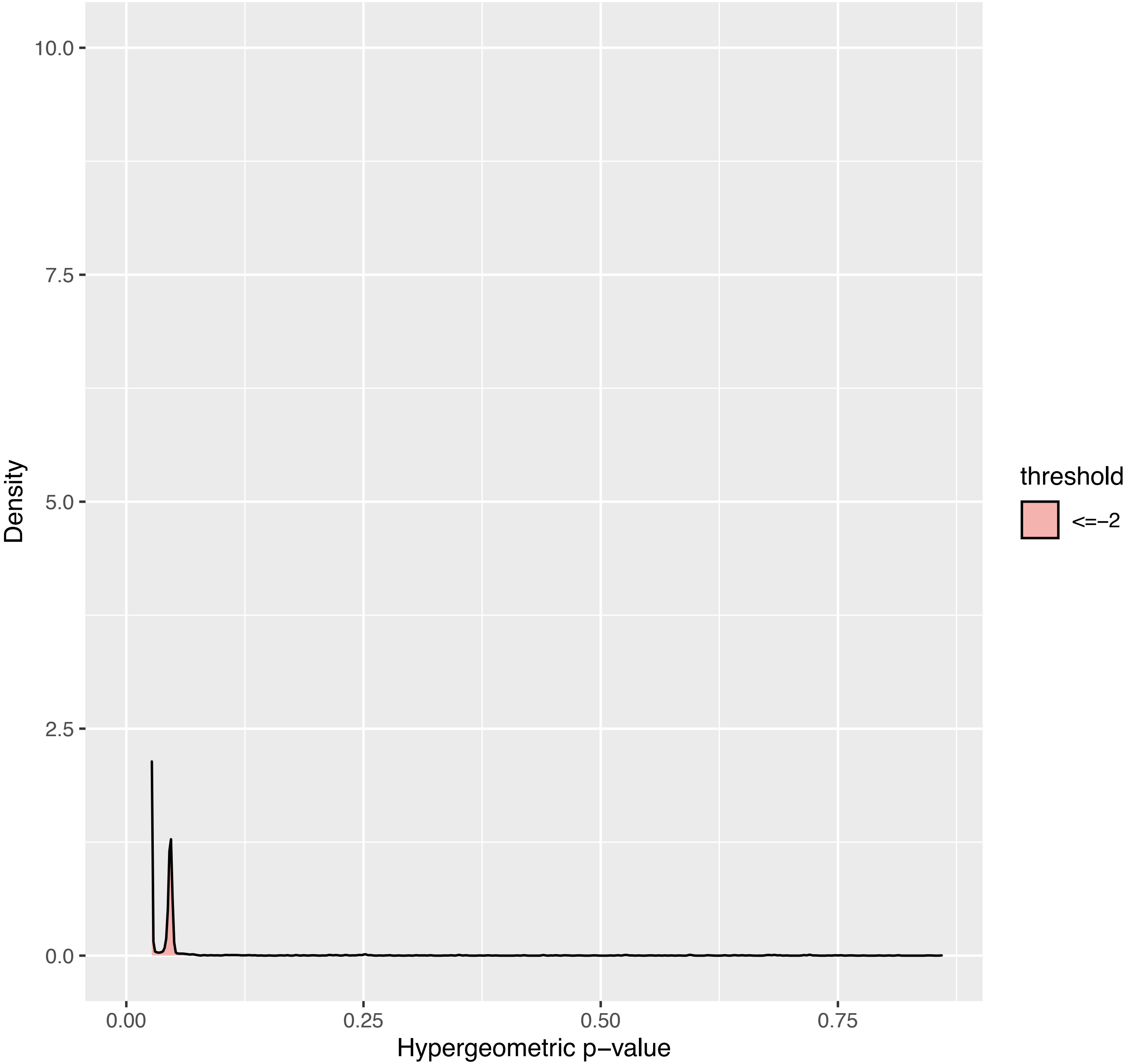

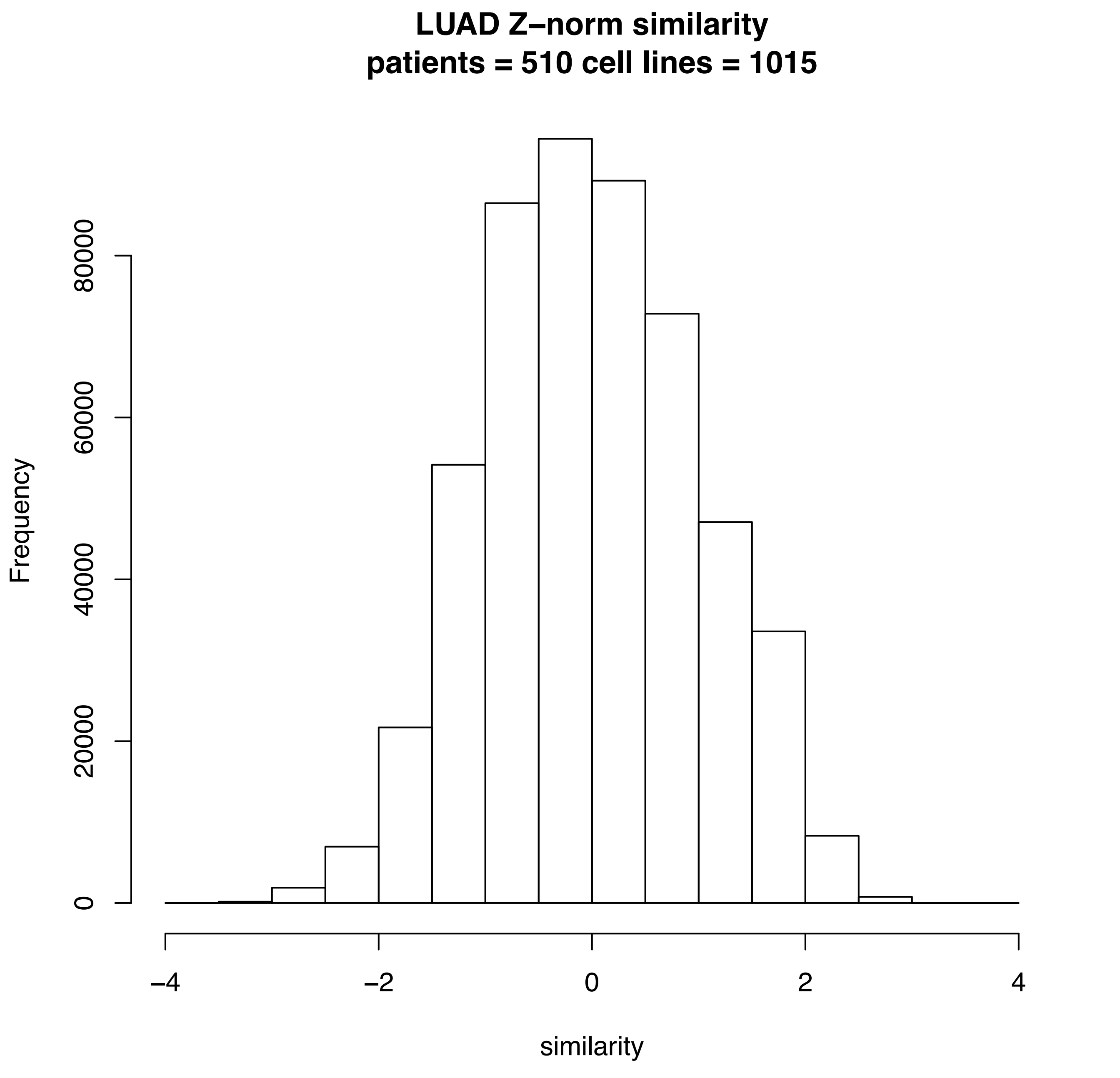

LUAD Z–norm similarity
patients = 510 cell lines = 1015
Frequency
0
20000
40000
60000
80000
–4
–2
0
2
4
similarity

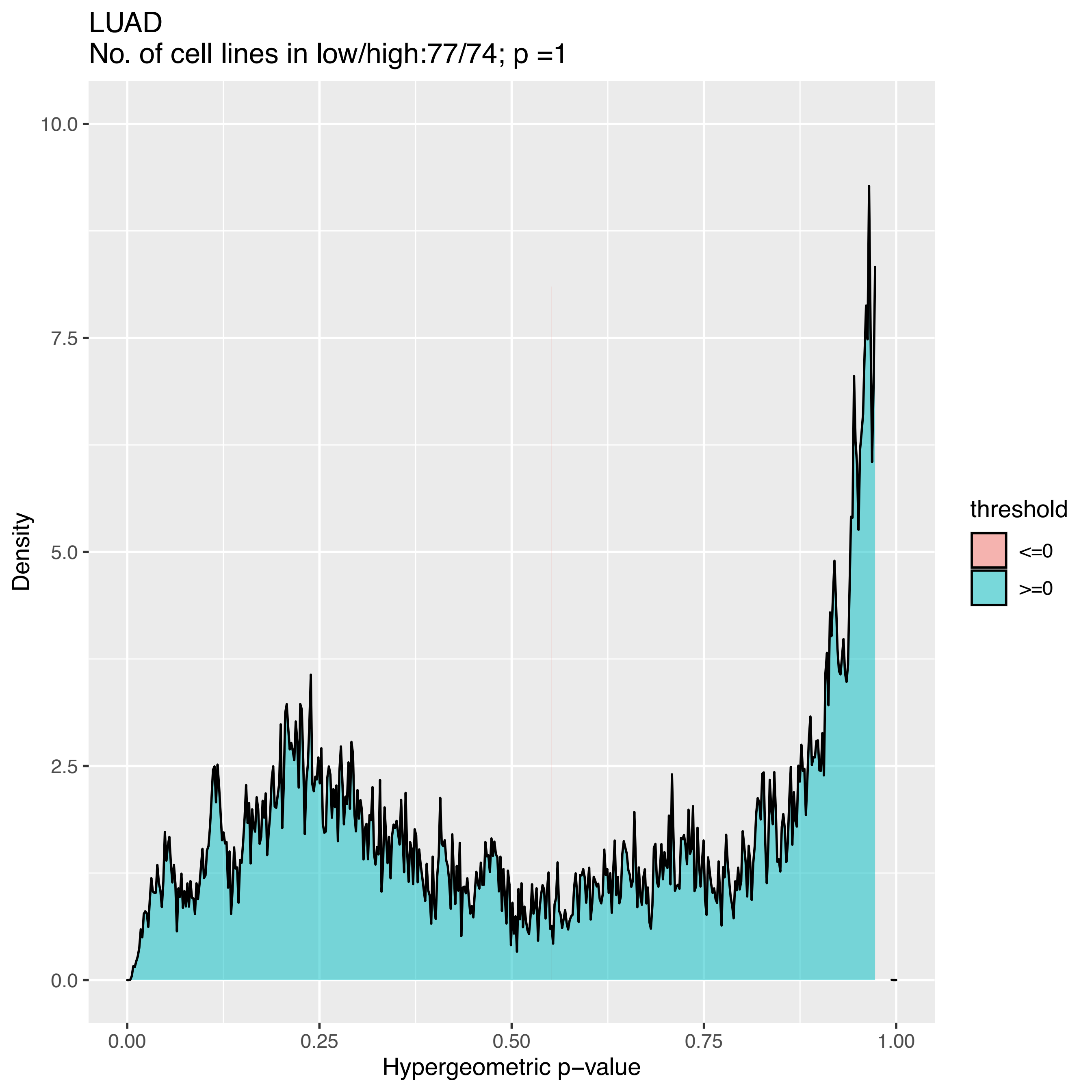
LUAD
No. of cell lines in low/high:77/74; p =1
Density
Hypergeometric p−value
0.0
2.5
5.0
7.5
10.0
0.00
0.25
0.50
0.75
1.00
threshold
<=0
>=0

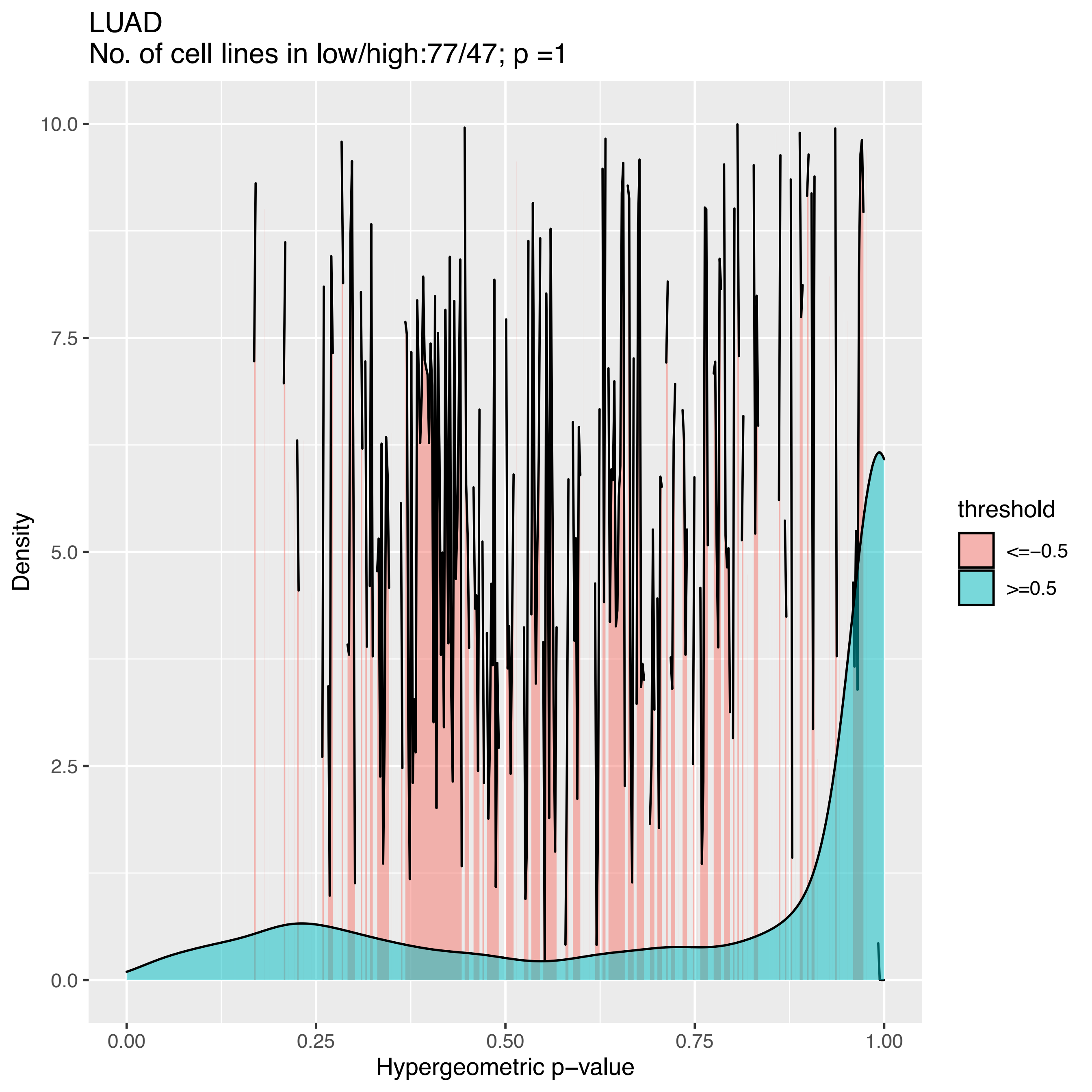

LUAD
No. of cell lines in low/high:77/47; p =1
Density
Hypergeometric p−value
0.0
2.5
5.0
7.5
10.0
0.00
0.25
0.50
0.75
1.00
threshold
<=−0.5
>=0.5

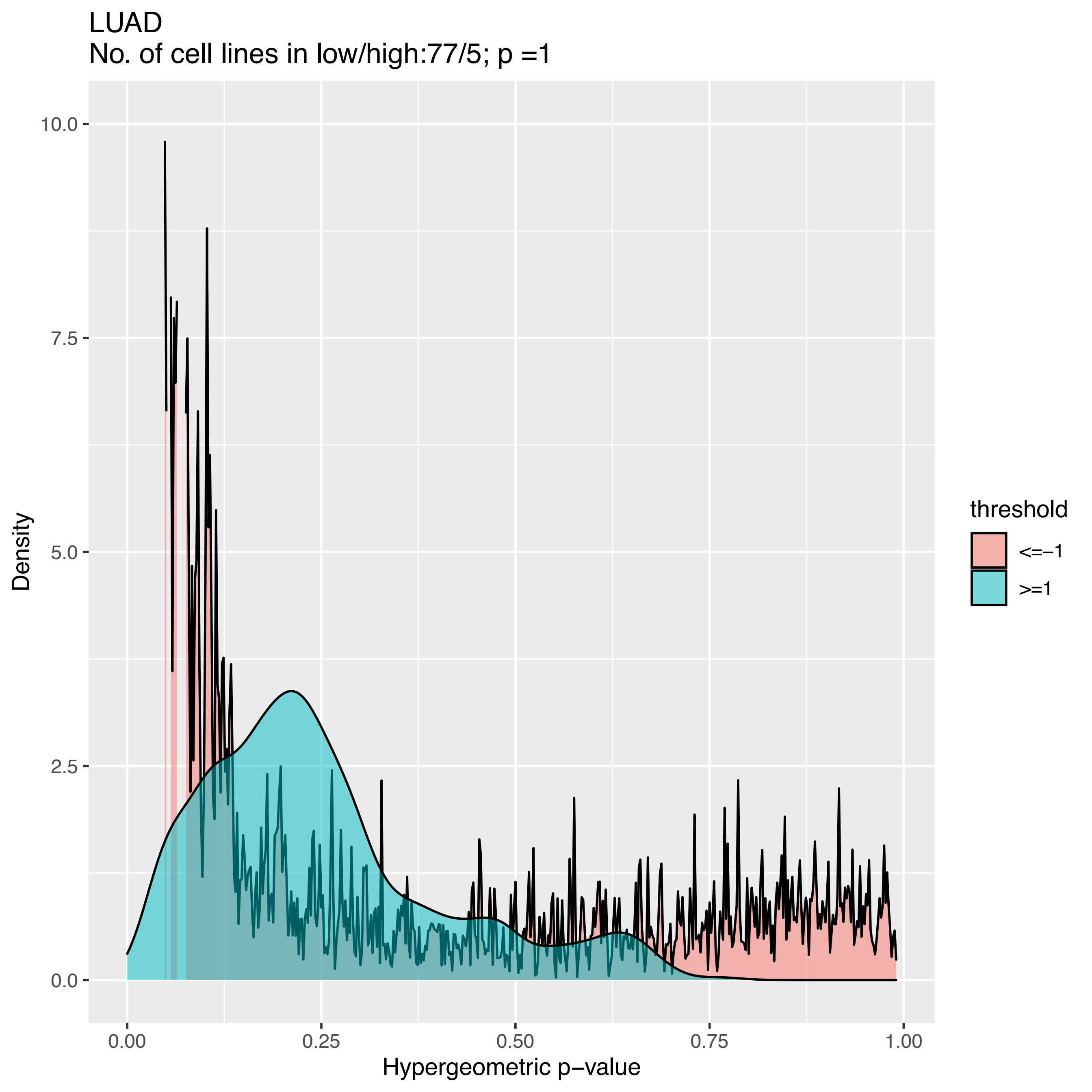
LUAD
No. of cell lines in low/high:77/5; p =1
10.0
7.5
5.0
2.5
0.0
Density
0.00
0.25
0.50
0.75
1.00
Hypergeometric p−value
threshold
<=−1
>=1

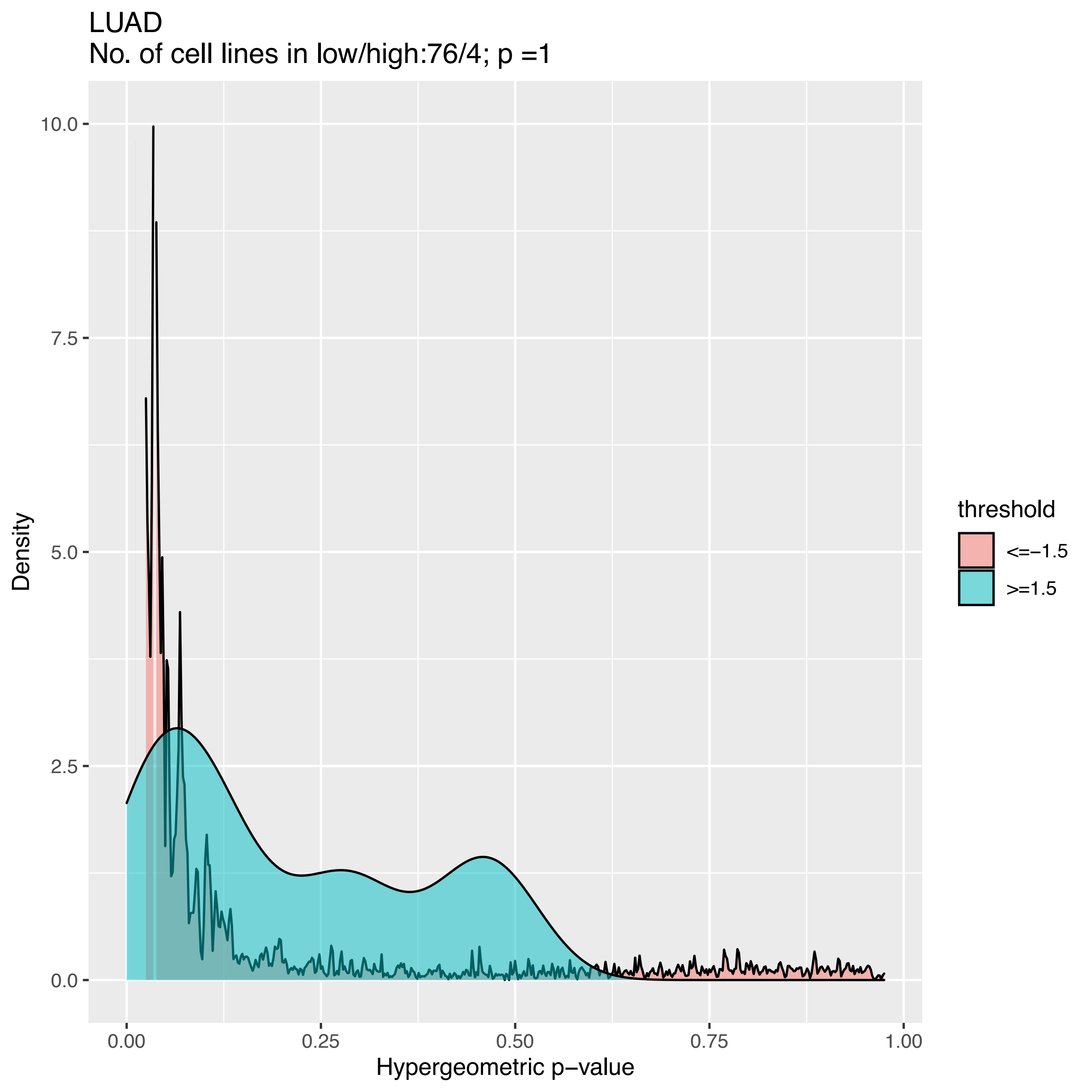
LUAD
No. of cell lines in low/high:76/4; p =1
Density
10.0
7.5
5.0
2.5
0.0
0.00
0.25
0.50
0.75
1.00
Hypergeometric p−value
threshold
<=−1.5
>=1.5

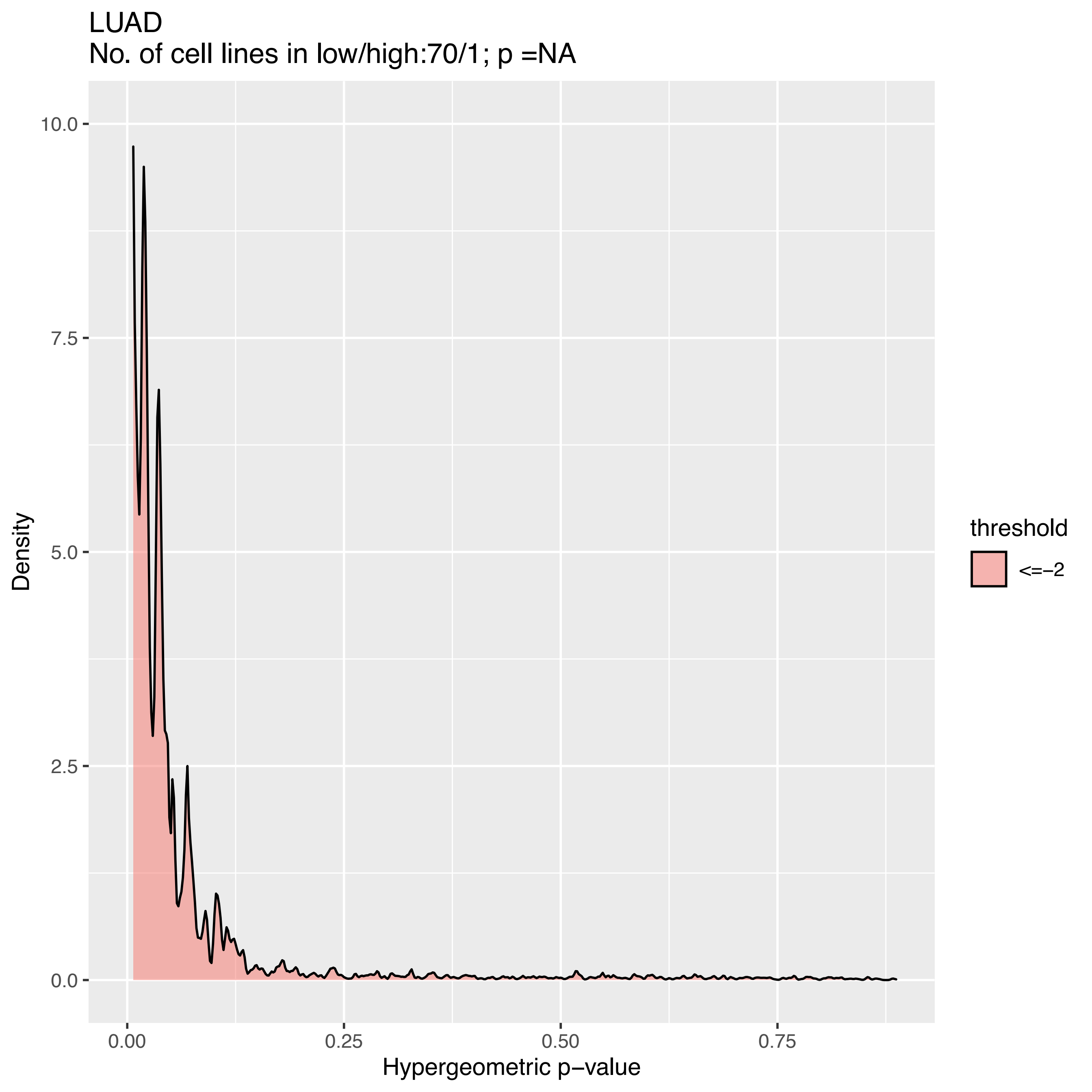

LUAD
No. of cell lines in low/high:70/1; p =NA
Density
10.0
7.5
5.0
2.5
0.0
0.00
0.25
0.50
0.75
Hypergeometric p−value
threshold
<=−2

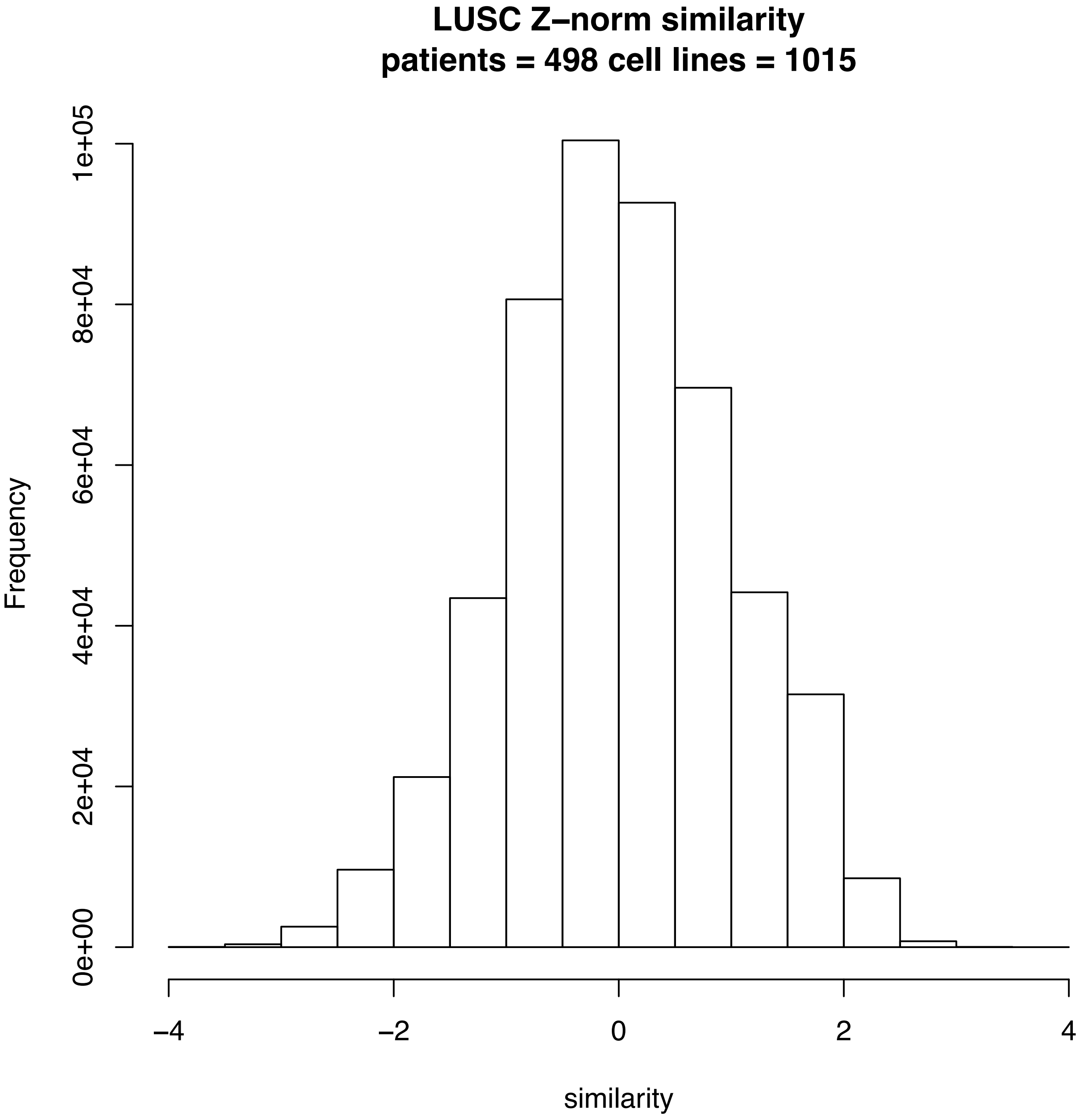
LUSC Z–norm similarity
patients = 498 cell lines = 1015
Frequency
0e+00
2e+04
4e+04
6e+04
8e+04
1e+05
–4
–2
0
2
4
similarity

LUSC
No. of cell lines in low/high:24/24; p =1

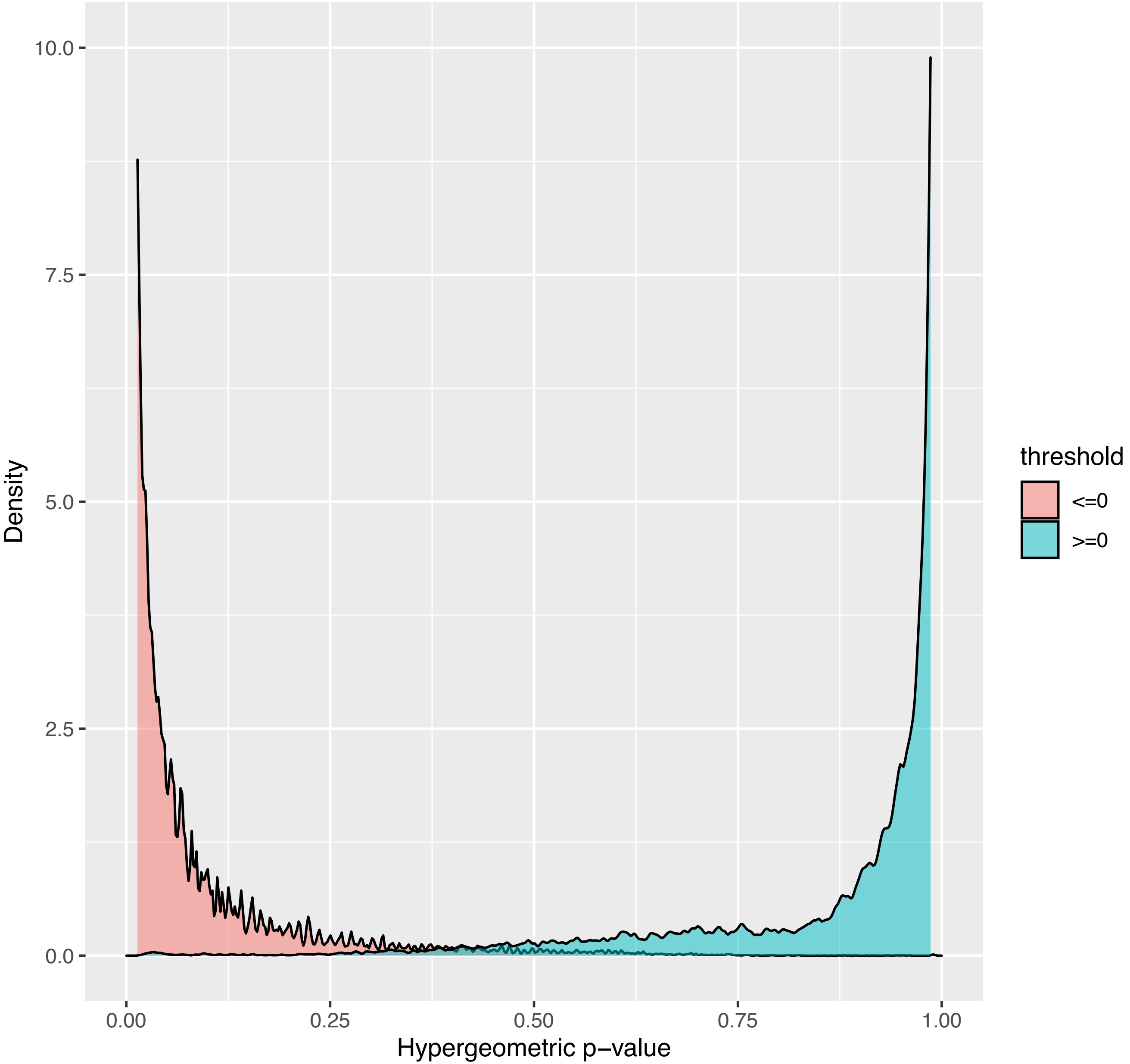

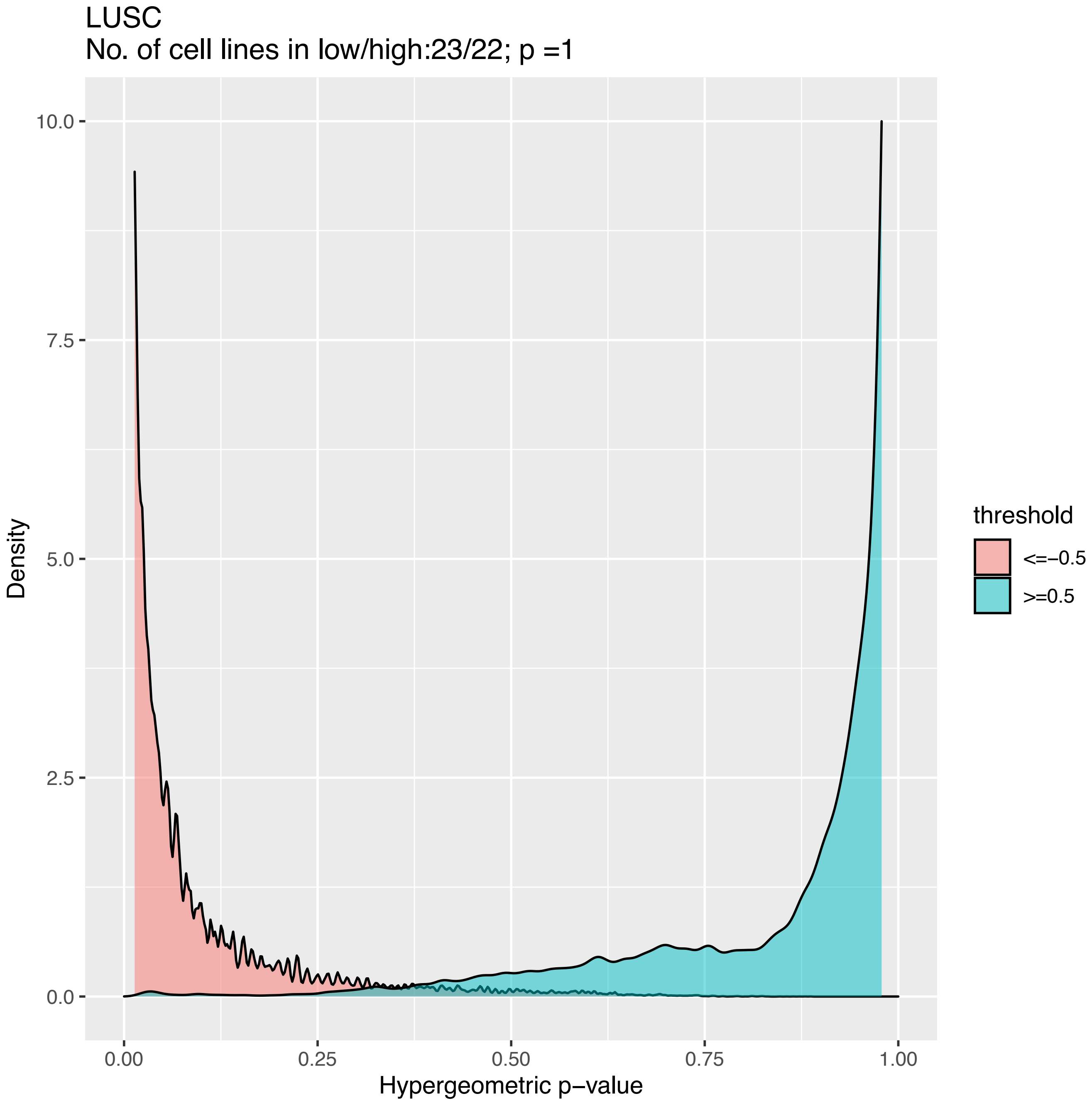
LUSC
No. of cell lines in low/high:23/22; p =1
Density
10.0
7.5
5.0
2.5
0.0
0.00
0.25
0.50
0.75
1.00
Hypergeometric p−value
threshold
<=−0.5
>=0.5

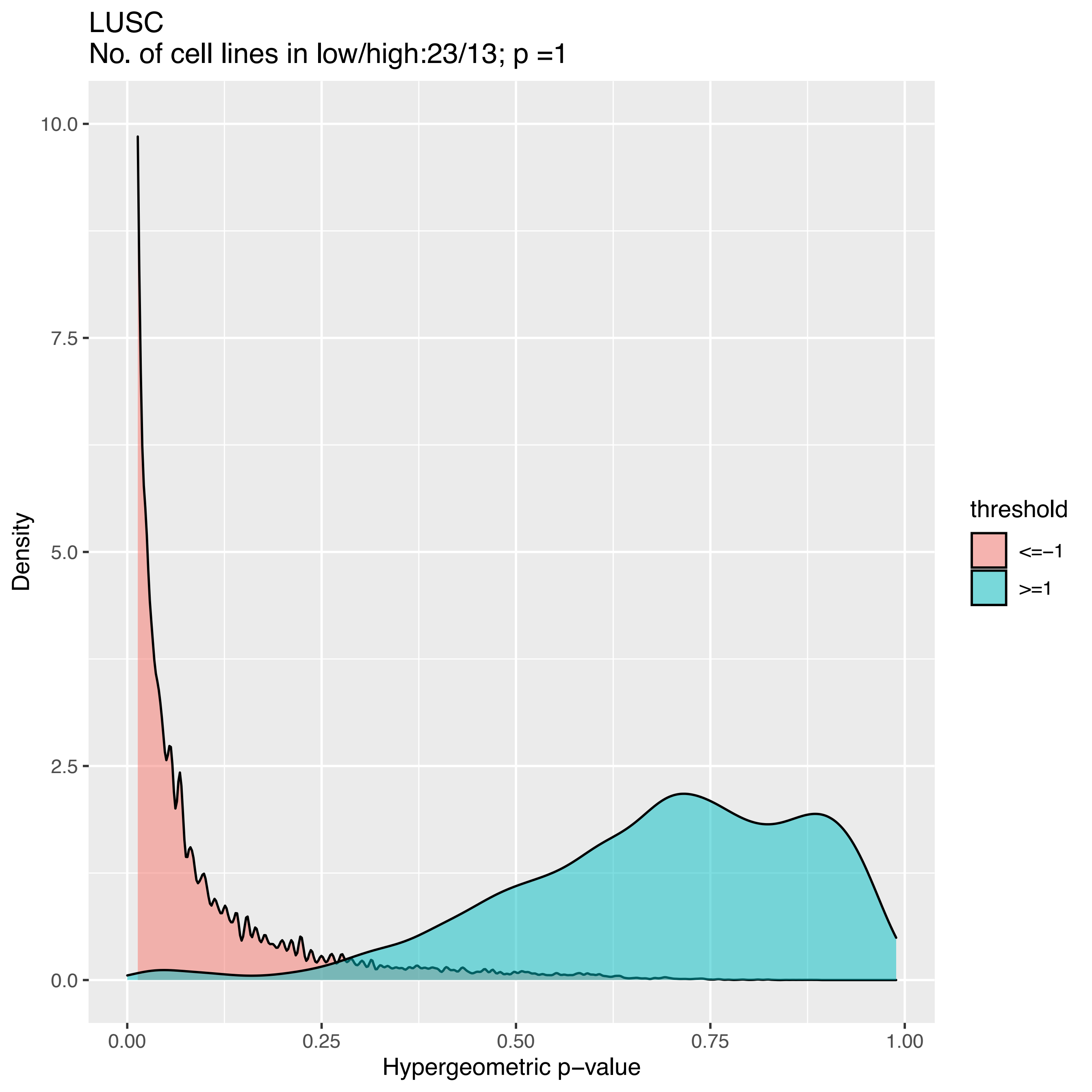

LUSC
No. of cell lines in low/high:23/13; p =1
Density
10.0
7.5
5.0
2.5
0.0
0.00
0.25
0.50
0.75
1.00
Hypergeometric p−value
threshold
<=−1
>=1

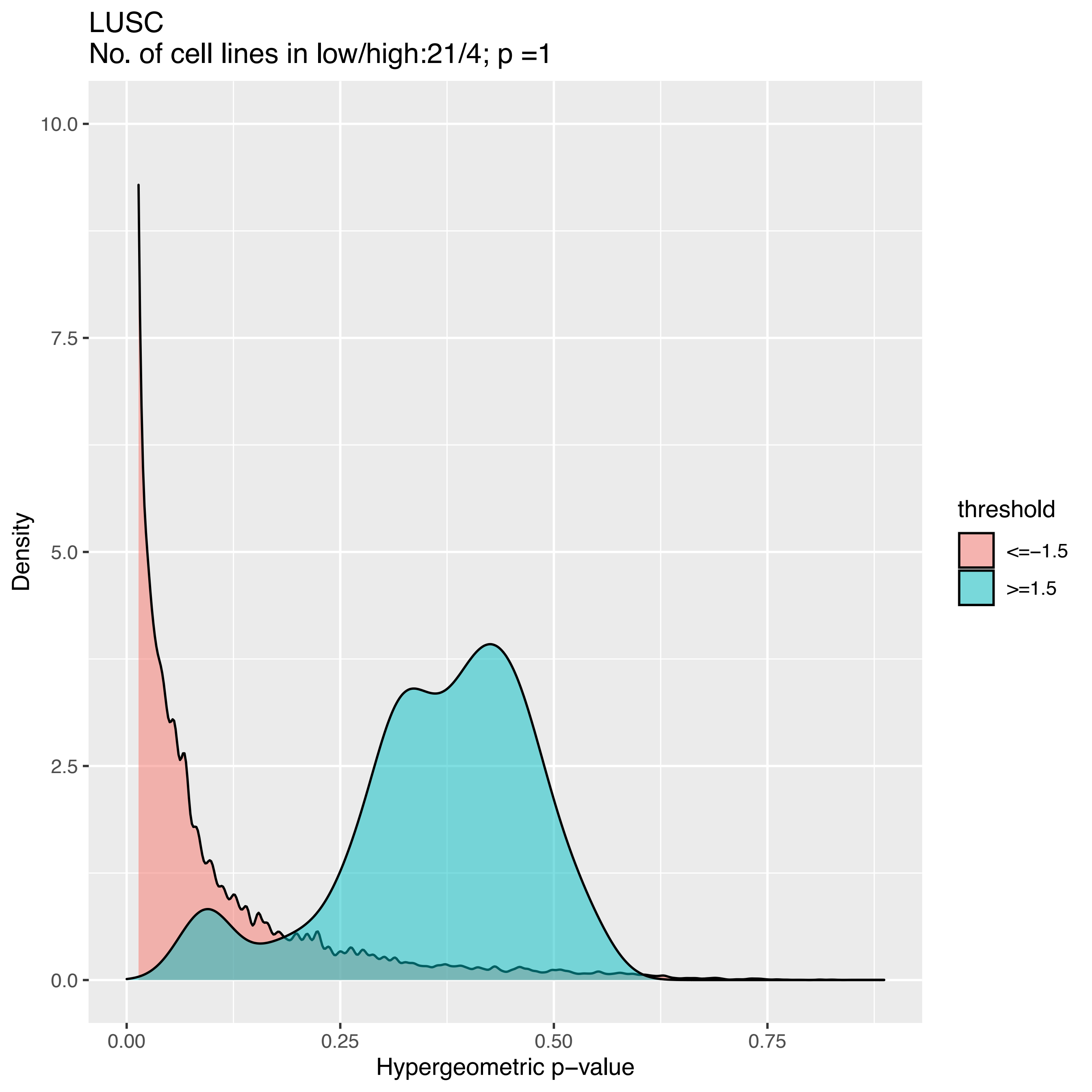

LUSC
No. of cell lines in low/high:21/4; p =1
Density
10.0
7.5
5.0
2.5
0.0
0.00
0.25
0.50
0.75
Hypergeometric p−value
threshold
<=−1.5
>=1.5

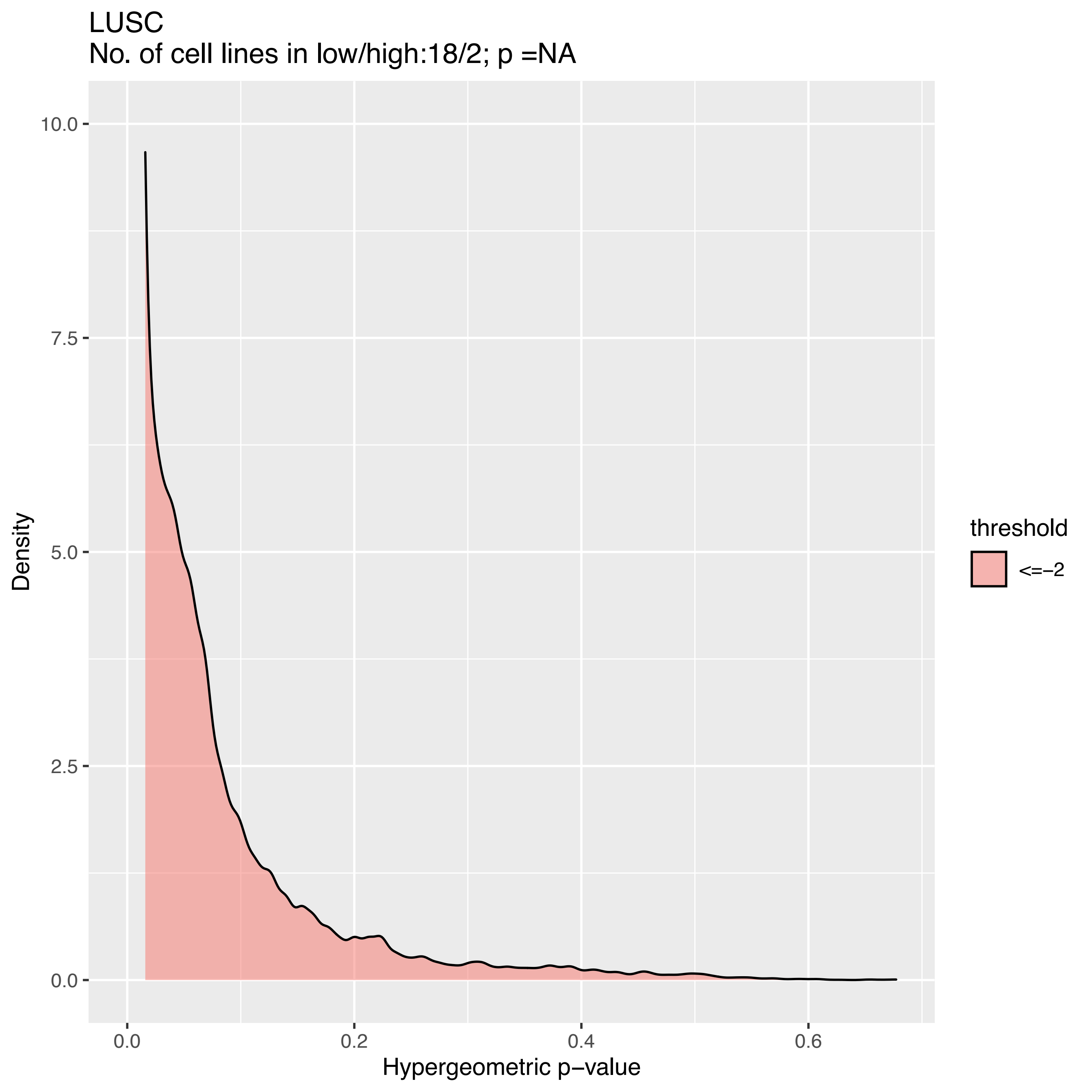
LUSC
No. of cell lines in low/high:18/2; p =NA
Density
Hypergeometric p−value
10.0
7.5
5.0
2.5
0.0
0.0
0.2
0.4
0.6
threshold
<=−2

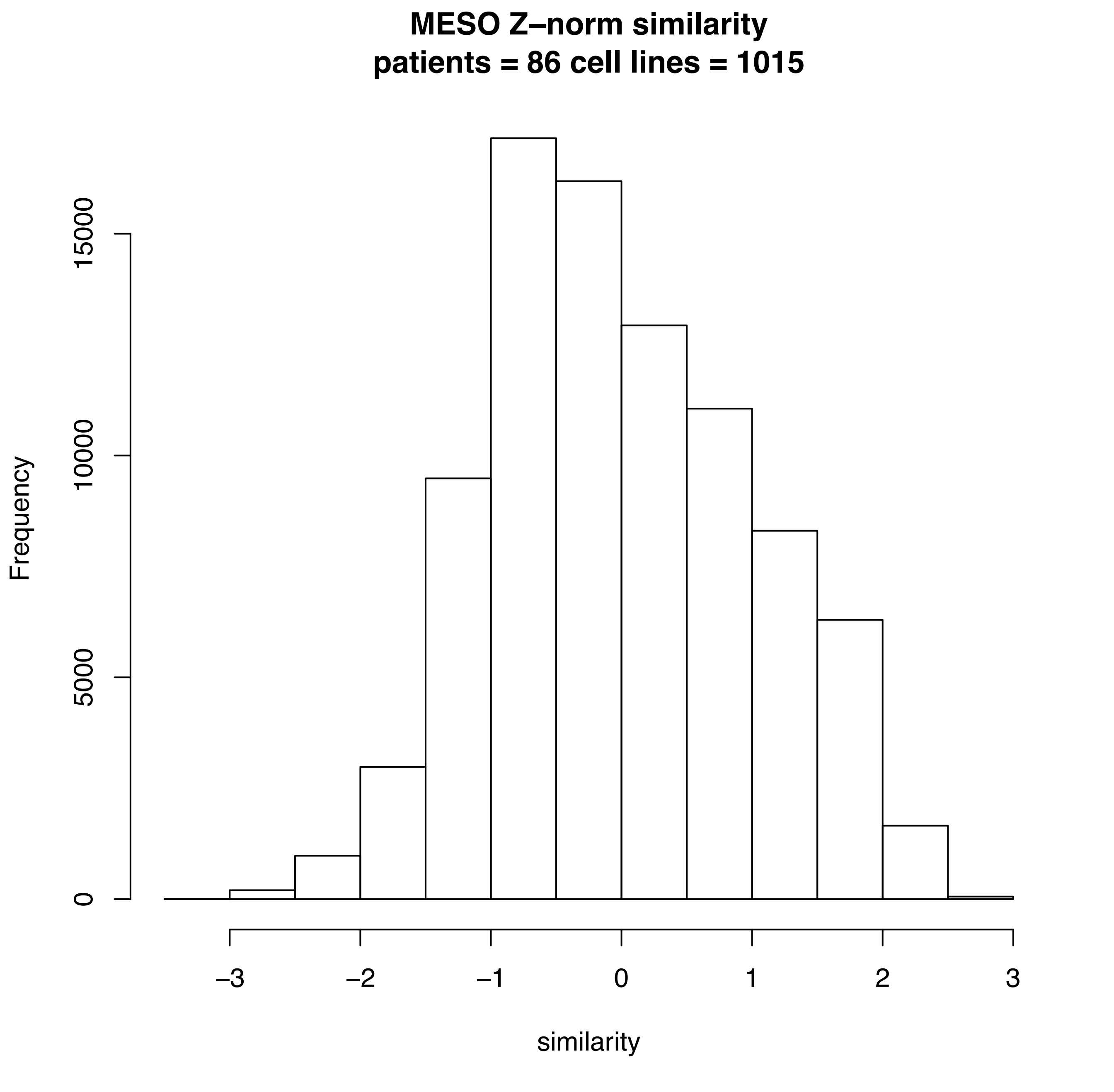
MESO Z–norm similarity
patients = 86 cell lines = 1015
Frequency
0
5000
10000
15000
–3
–2
–1
0
1
2
3
similarity

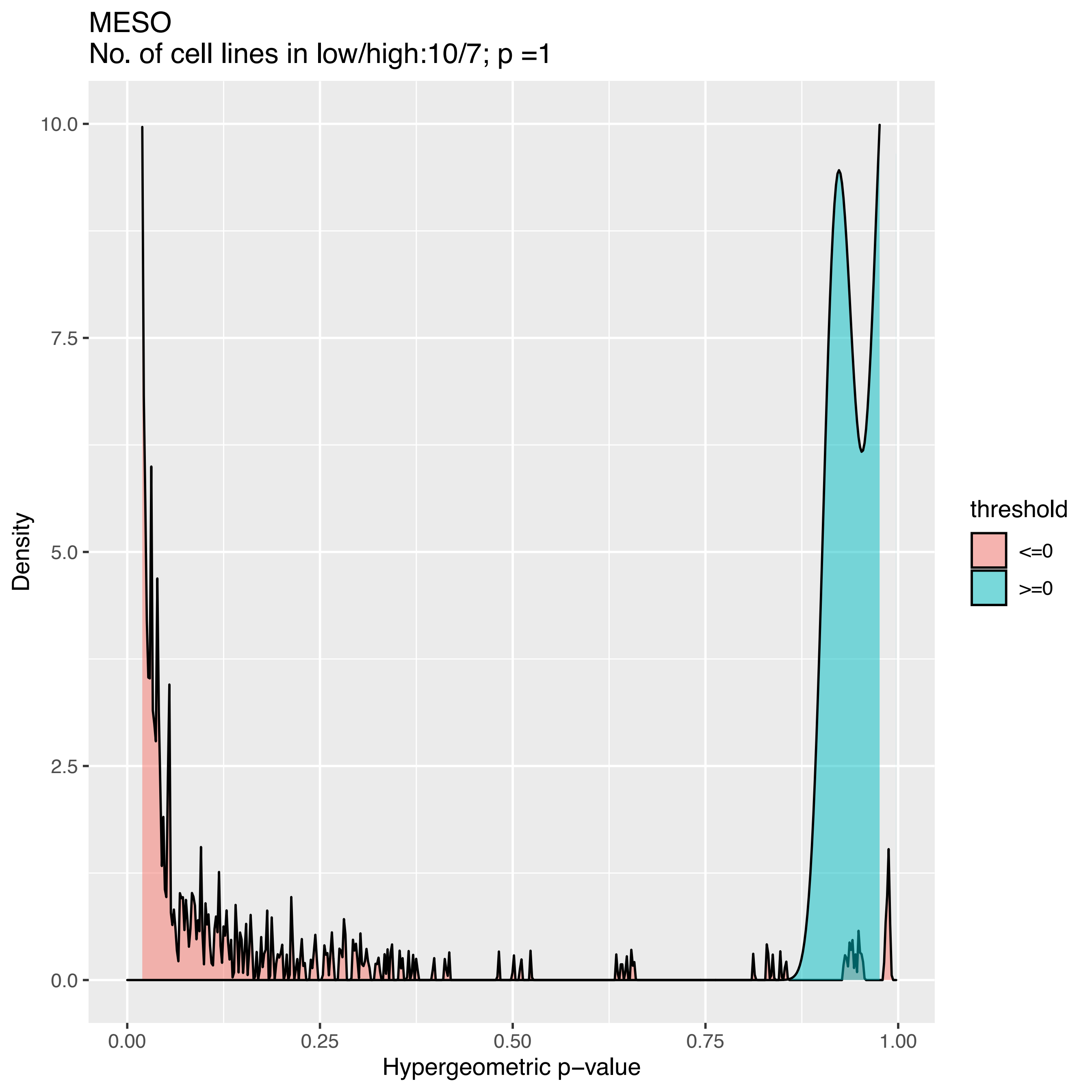

MESO
No. of cell lines in low/high:10/7; p =1
Density
10.0
7.5
5.0
2.5
0.0
0.00
0.25
0.50
0.75
1.00
Hypergeometric p−value
threshold
<=0
>=0

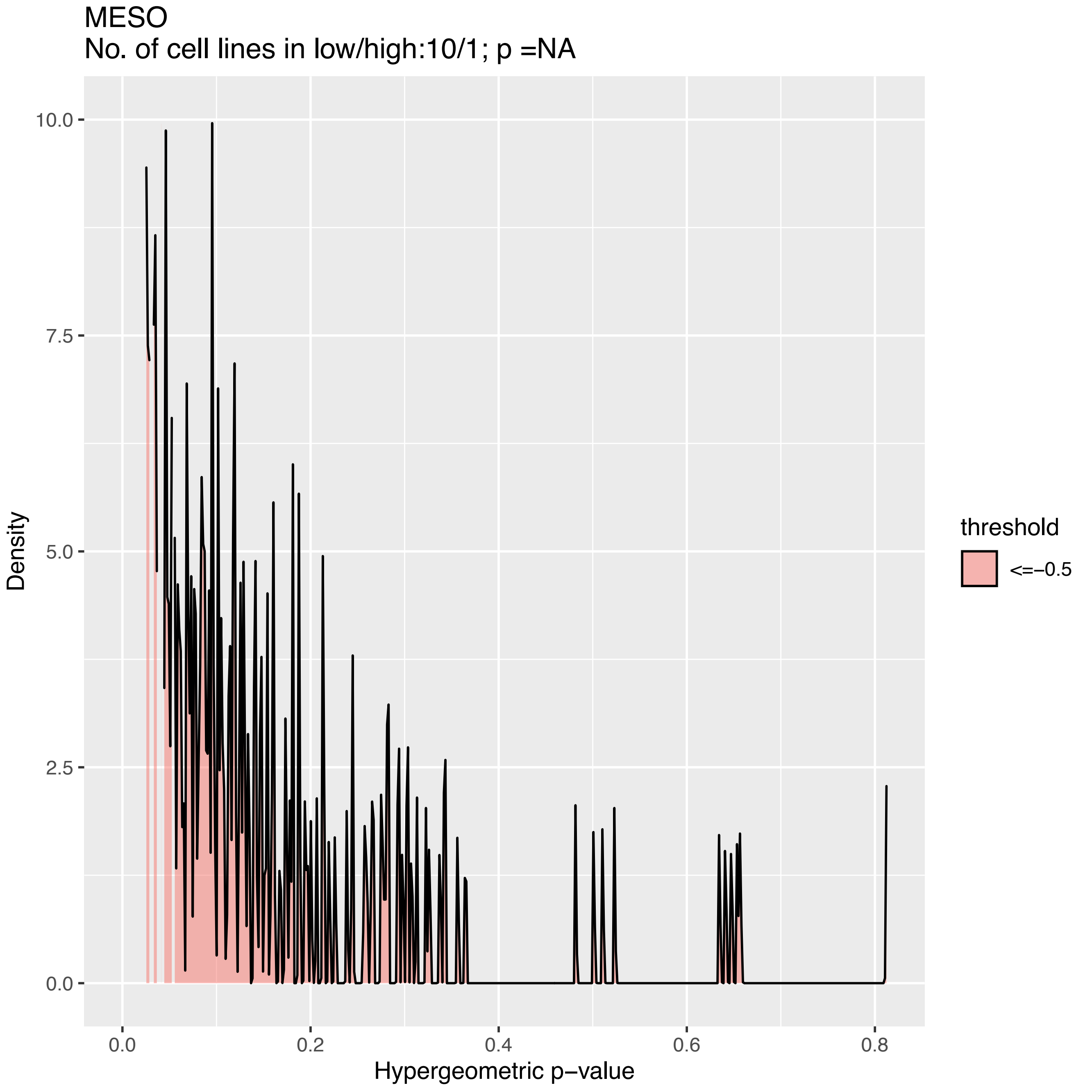

MESO
No. of cell lines in low/high:10/1; p =NA
Density
10.0
7.5
5.0
2.5
0.0
0.0
0.2
0.4
0.6
0.8
Hypergeometric p−value
threshold
<=−0.5

MESO
No. of cell lines in low/high:9/1; p =NA

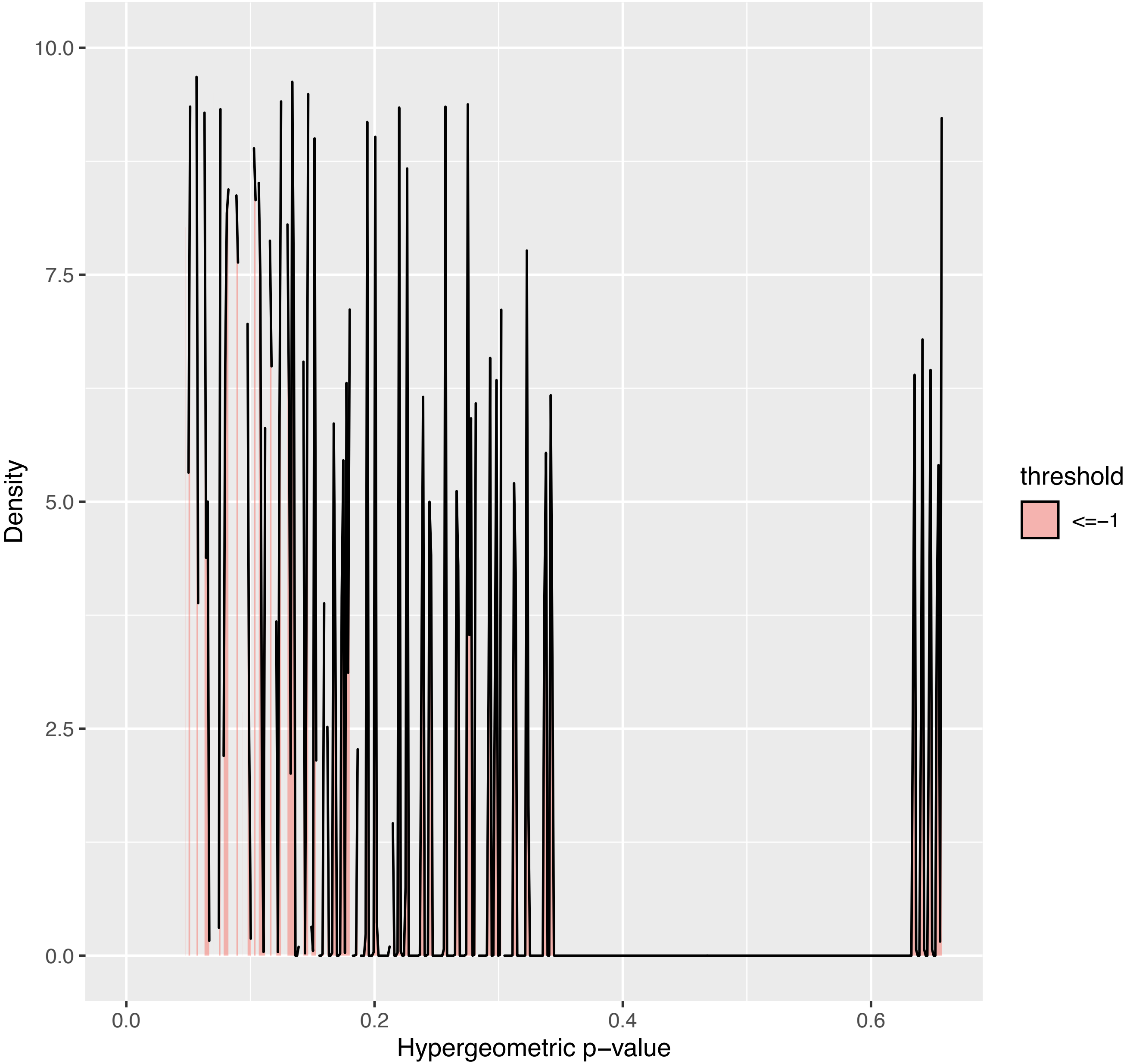

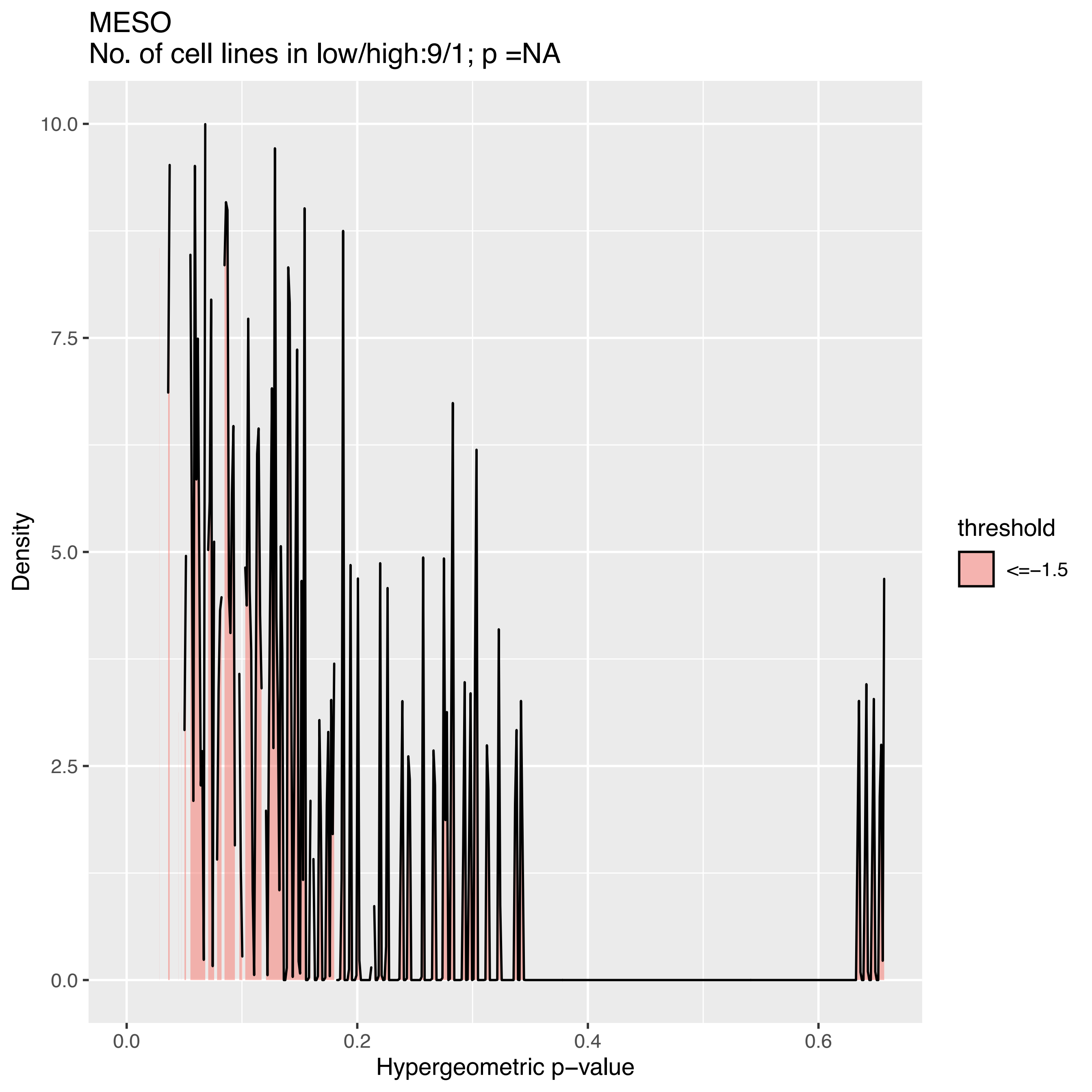
MESO
No. of cell lines in low/high:9/1; p =NA
10.0
7.5
5.0
2.5
0.0
Density
0.0
0.2
0.4
0.6
Hypergeometric p−value
threshold
<=−1.5

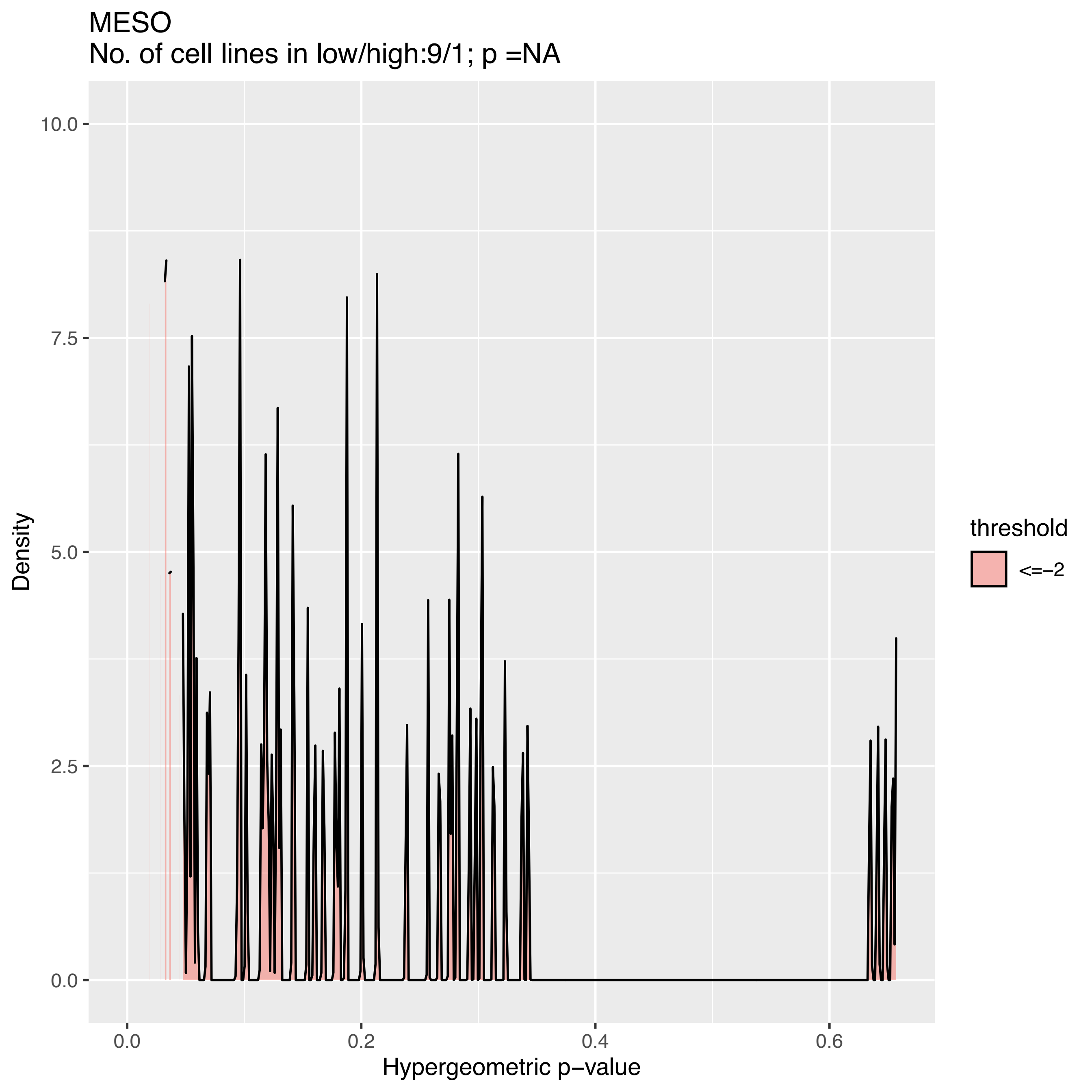

MESO
No. of cell lines in low/high:9/1; p =NA
Density
Hypergeometric p−value
0.0
2.5
5.0
7.5
10.0
0.0
0.2
0.4
0.6
threshold
<=−2

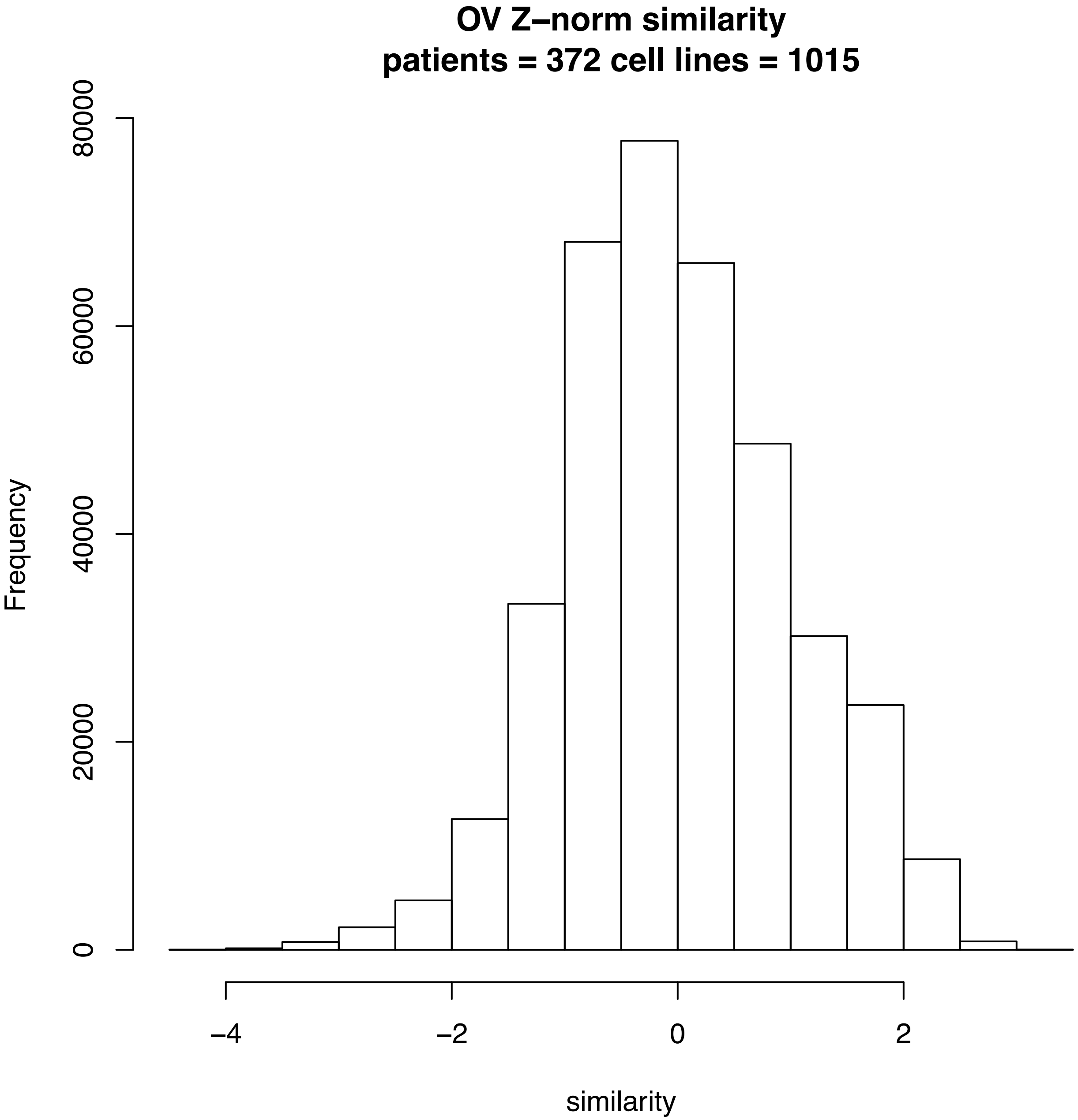

OV Z–norm similarity
patients = 372 cell lines = 1015
Frequency
0
20000
40000
60000
80000
similarity
–4
–2
0
2

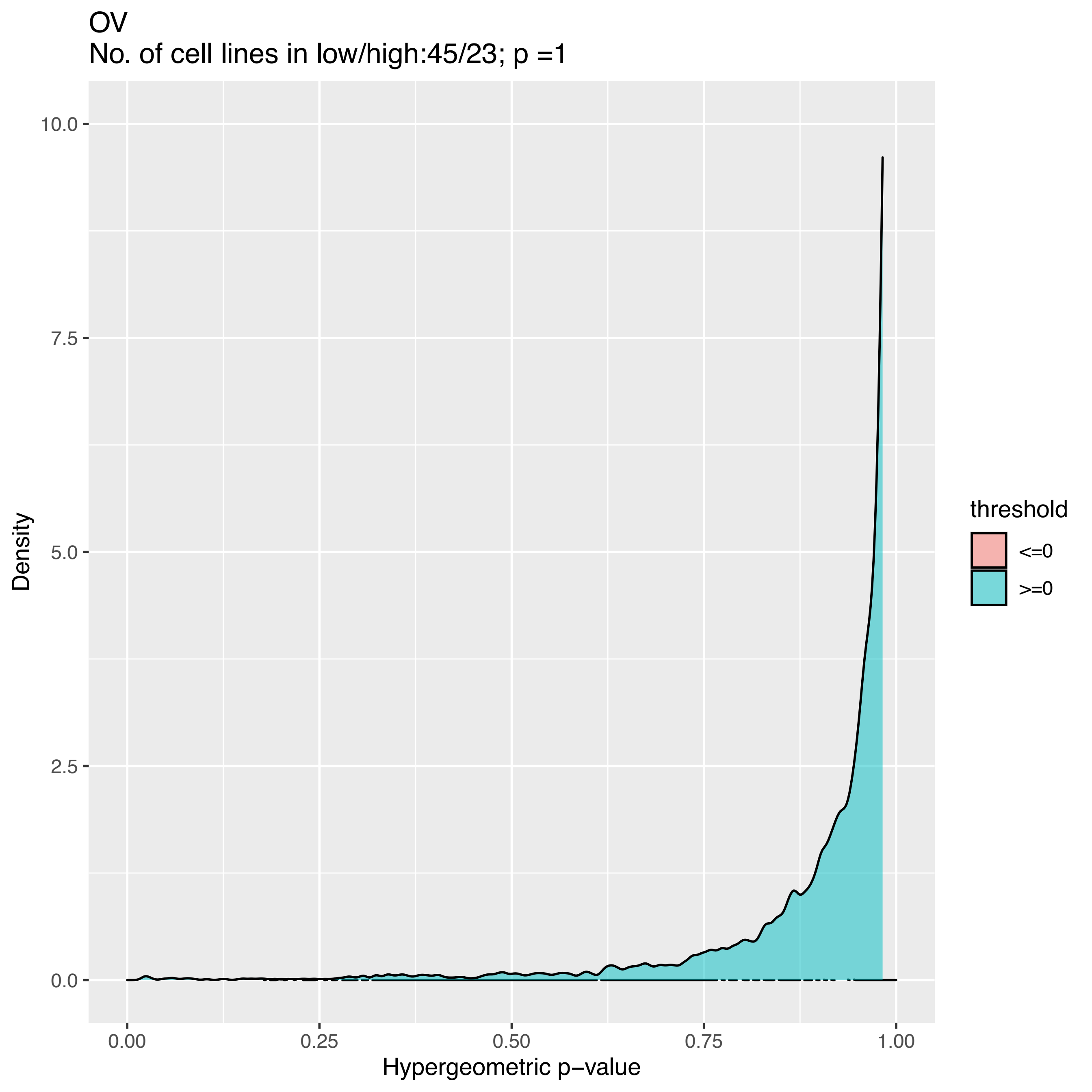

OV
No. of cell lines in low/high:45/23; p =1
Density
10.0
7.5
5.0
2.5
0.0
0.00
0.25
0.50
0.75
1.00
Hypergeometric p−value
threshold
<=0
>=0

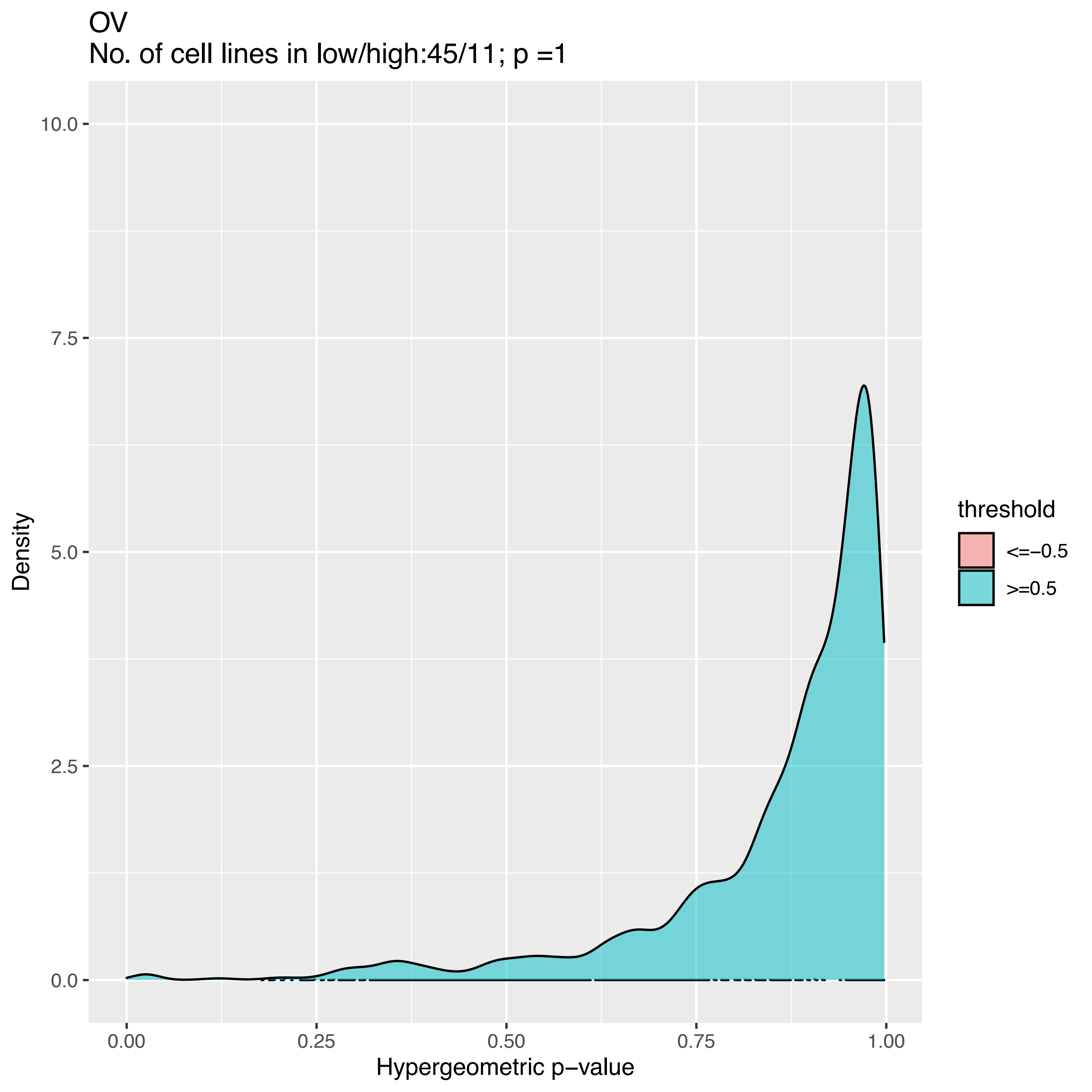
OV
No. of cell lines in low/high:45/11; p =1
10.0
7.5
5.0
2.5
0.0
Density
0.00
0.25
0.50
0.75
1.00
Hypergeometric p−value
threshold
<=−0.5
>=0.5

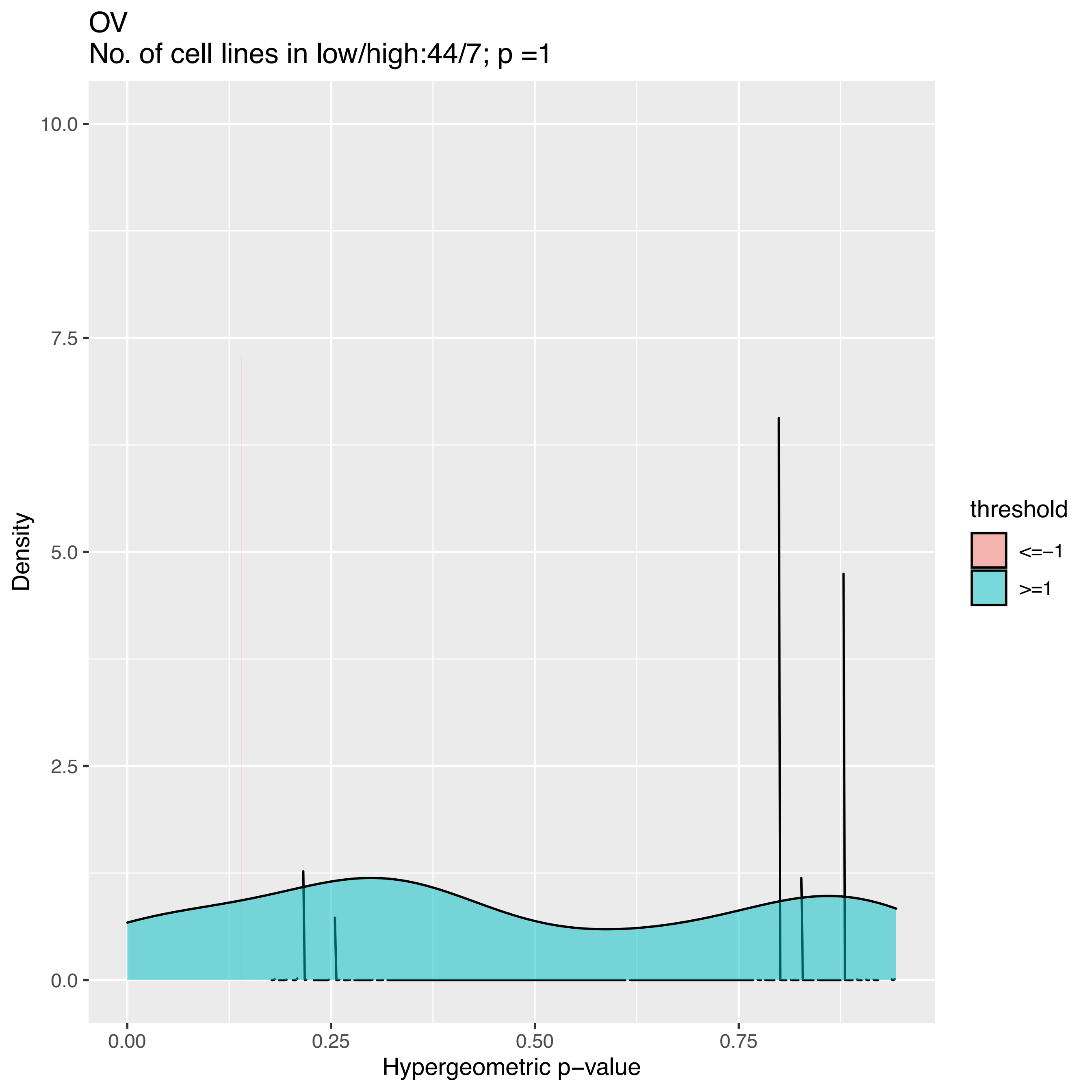

OV
No. of cell lines in low/high:44/7; p =1
10.0
7.5
5.0
2.5
0.0
Density
0.00
0.25
0.50
0.75
Hypergeometric p−value
threshold
<=−1
>=1

OV
No. of cell lines in low/high:43/2; p =NA

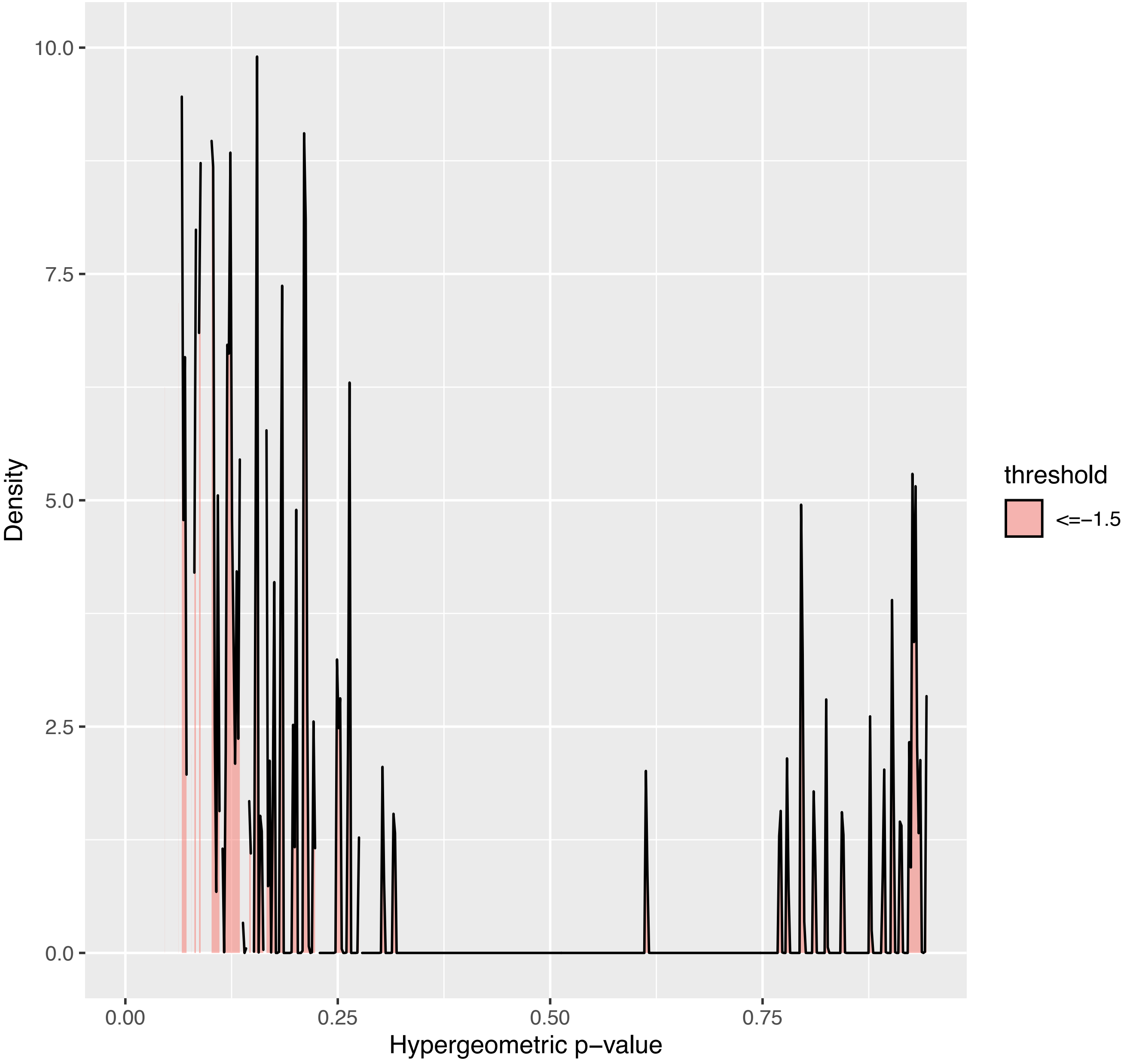

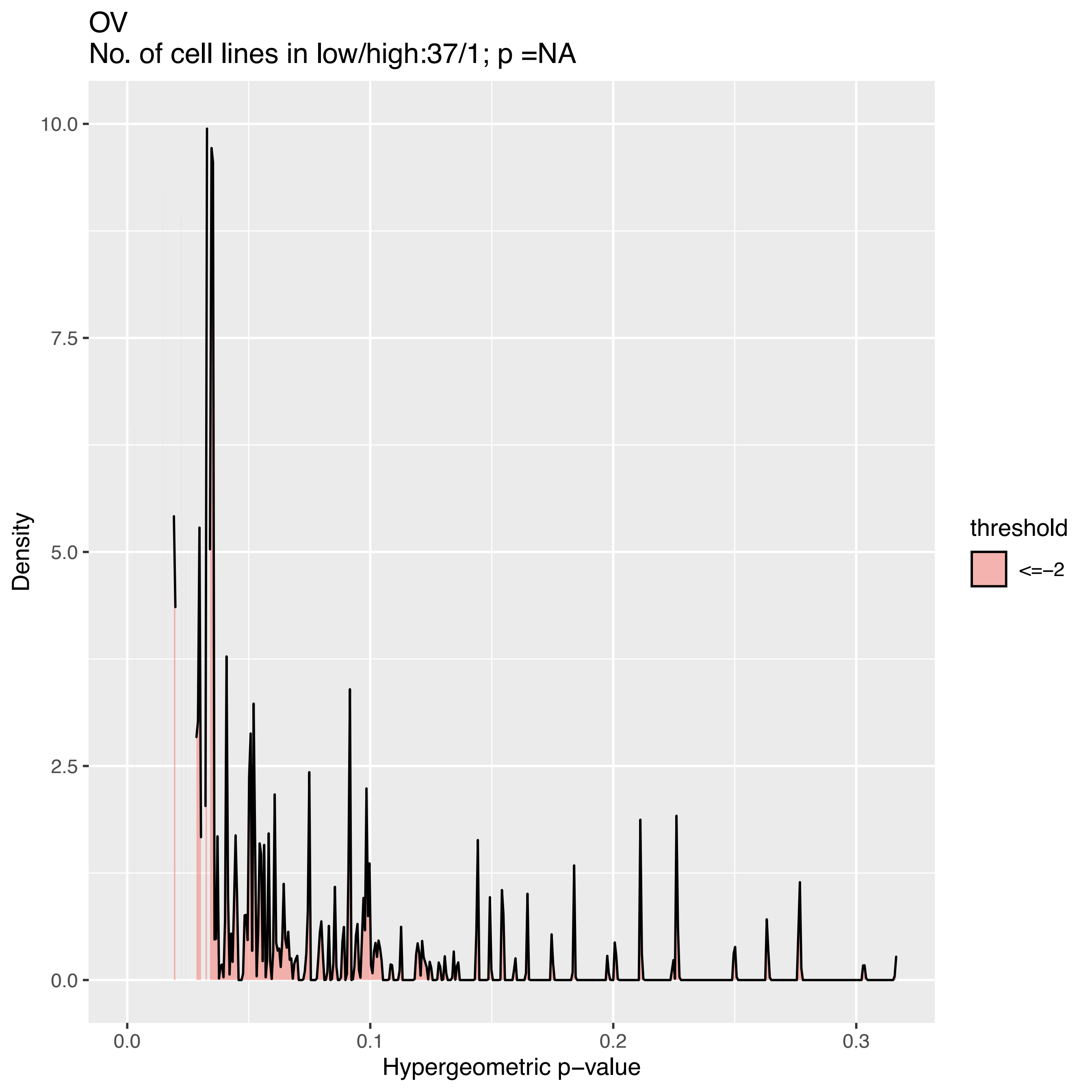
OV
No. of cell lines in low/high:37/1; p =NA
Density
10.0
7.5
5.0
2.5
0.0
0.0
0.1
0.2
0.3
Hypergeometric p−value
threshold
<=−2

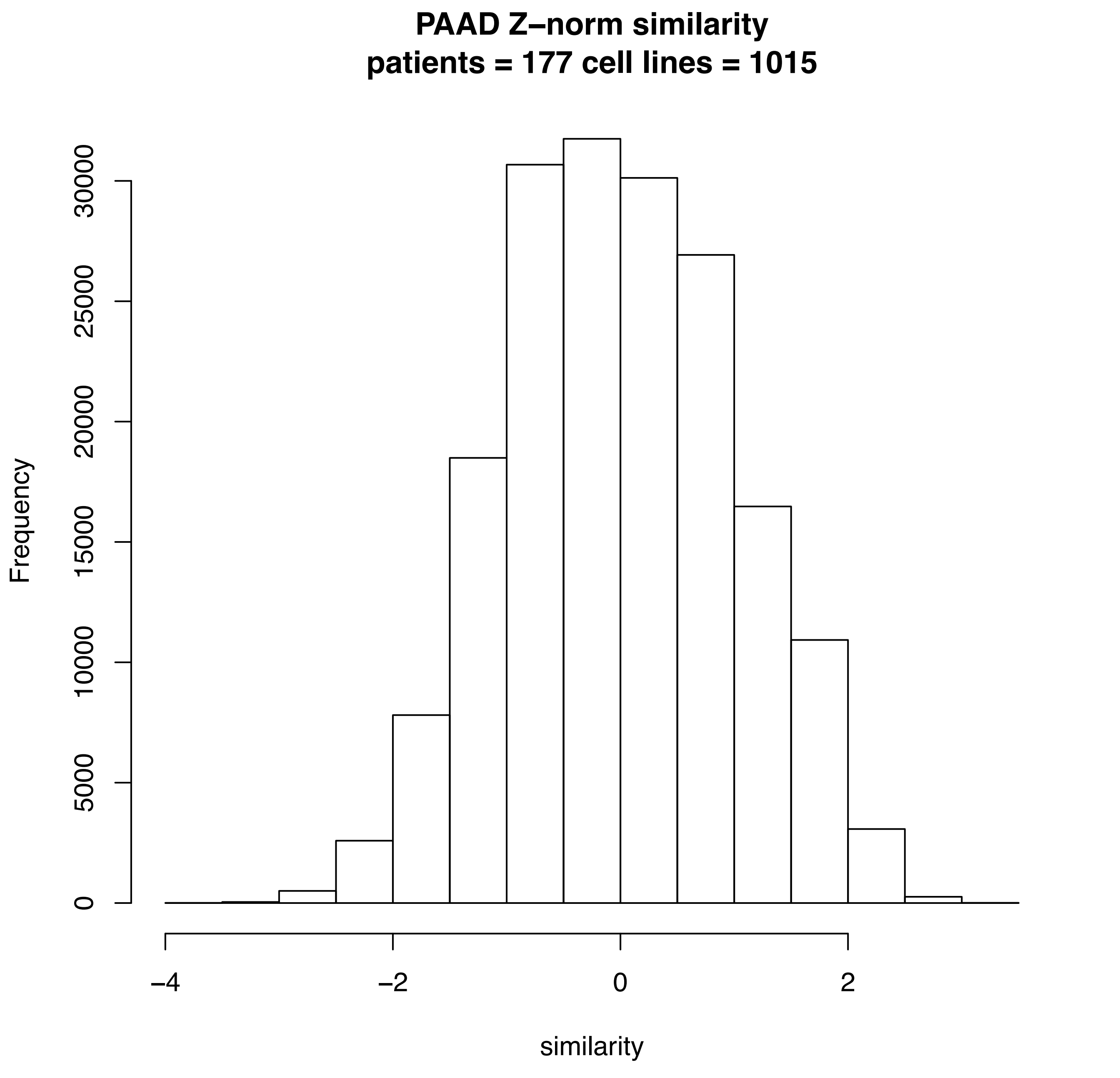
PAAD Z–norm similarity
patients = 177 cell lines = 1015
Frequency
0
5000
10000
15000
20000
25000
30000
−4
−2
0
2
similarity

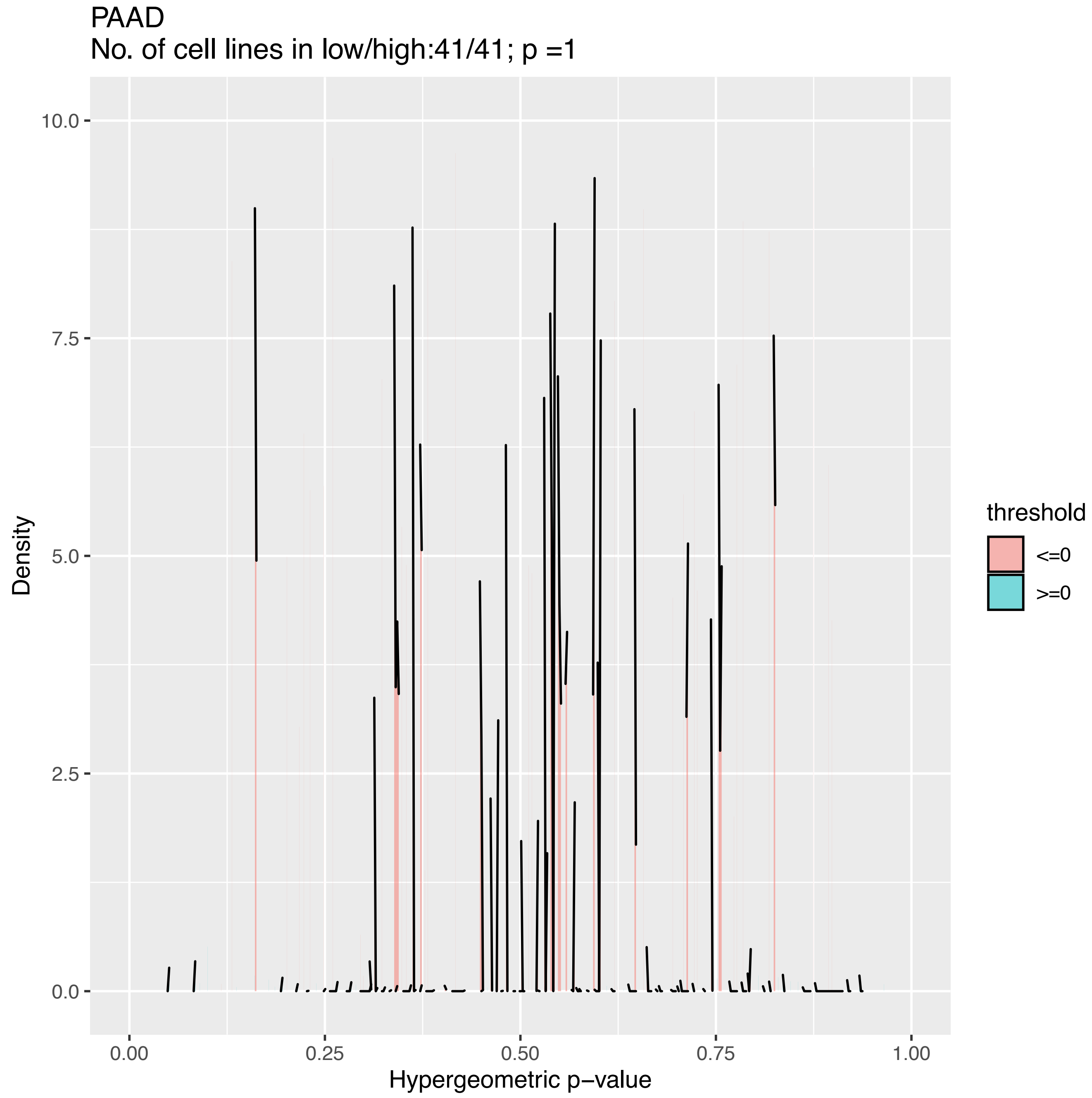

PAAD
No. of cell lines in low/high:41/41; p =1
10.0
7.5
5.0
2.5
0.0
Density
0.00
0.25
0.50
0.75
1.00
Hypergeometric p−value
threshold
<=0
>=0

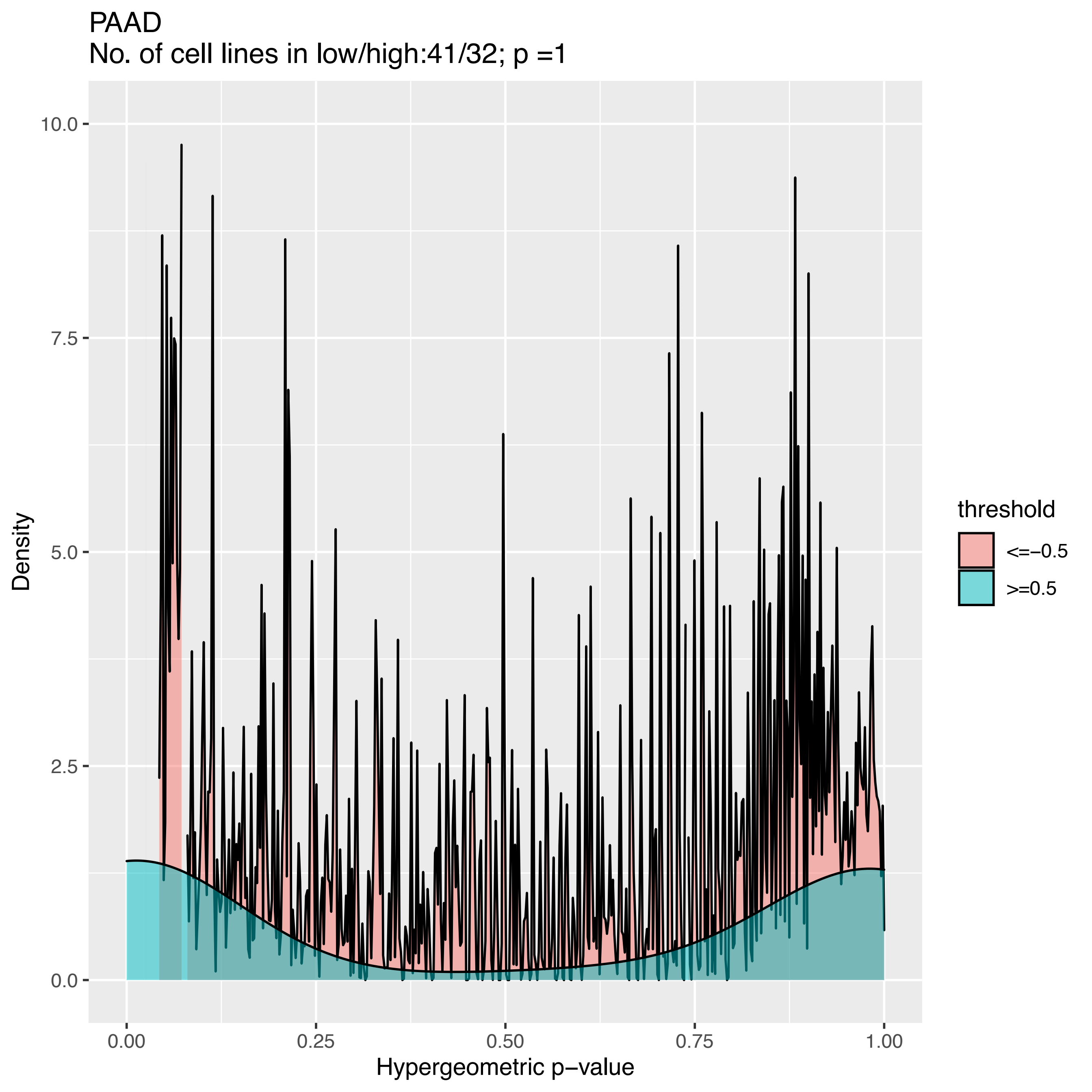
PAAD
No. of cell lines in low/high:41/32; p =1
Density
10.0
7.5
5.0
2.5
0.0
0.00
0.25
0.50
0.75
1.00
Hypergeometric p−value
threshold
<=−0.5
>=0.5

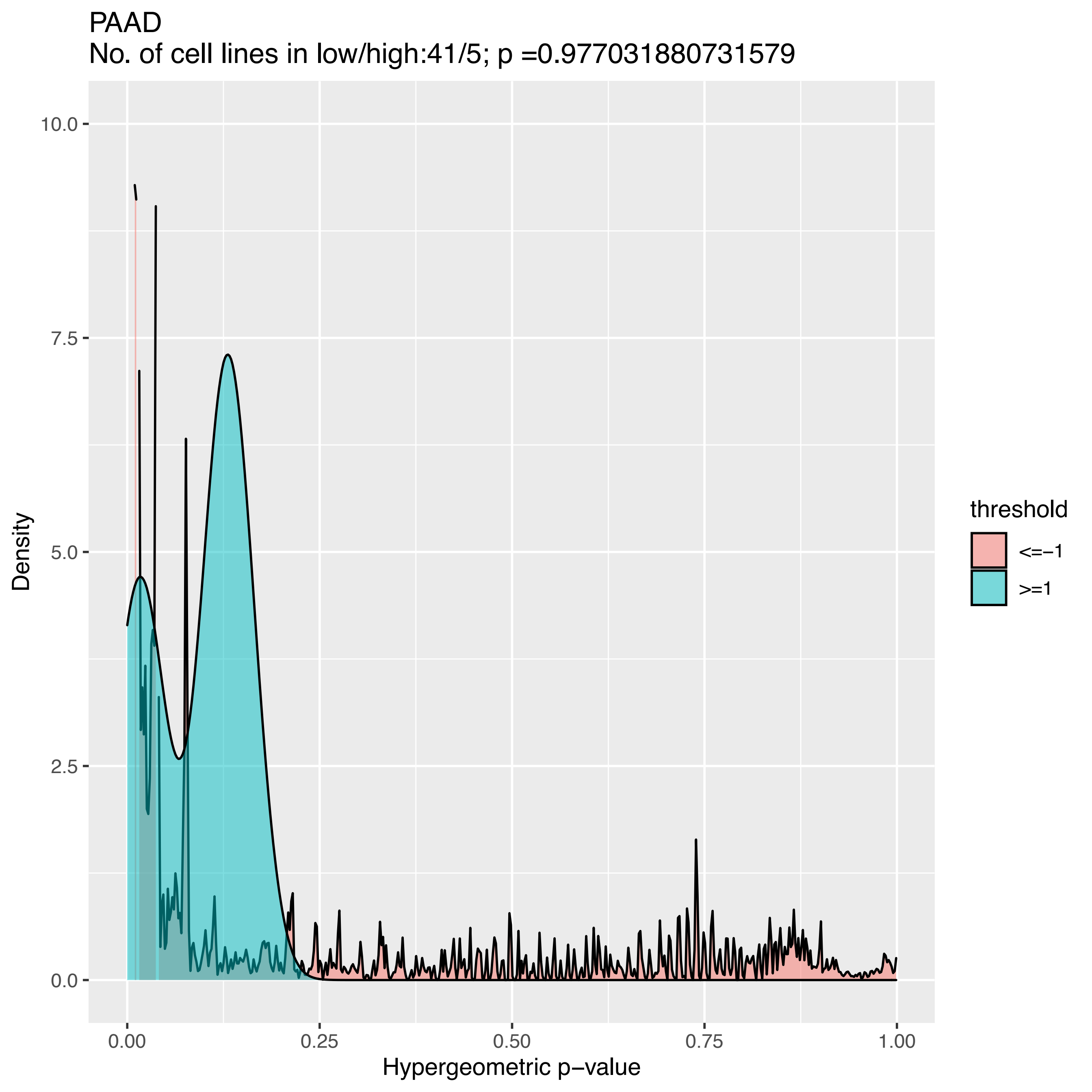
PAAD
No. of cell lines in low/high:41/5; p =0.977031880731579
Density
10.0
7.5
5.0
2.5
0.0
0.00
0.25
0.50
0.75
1.00
Hypergeometric p−value
threshold
<=−1
>=1

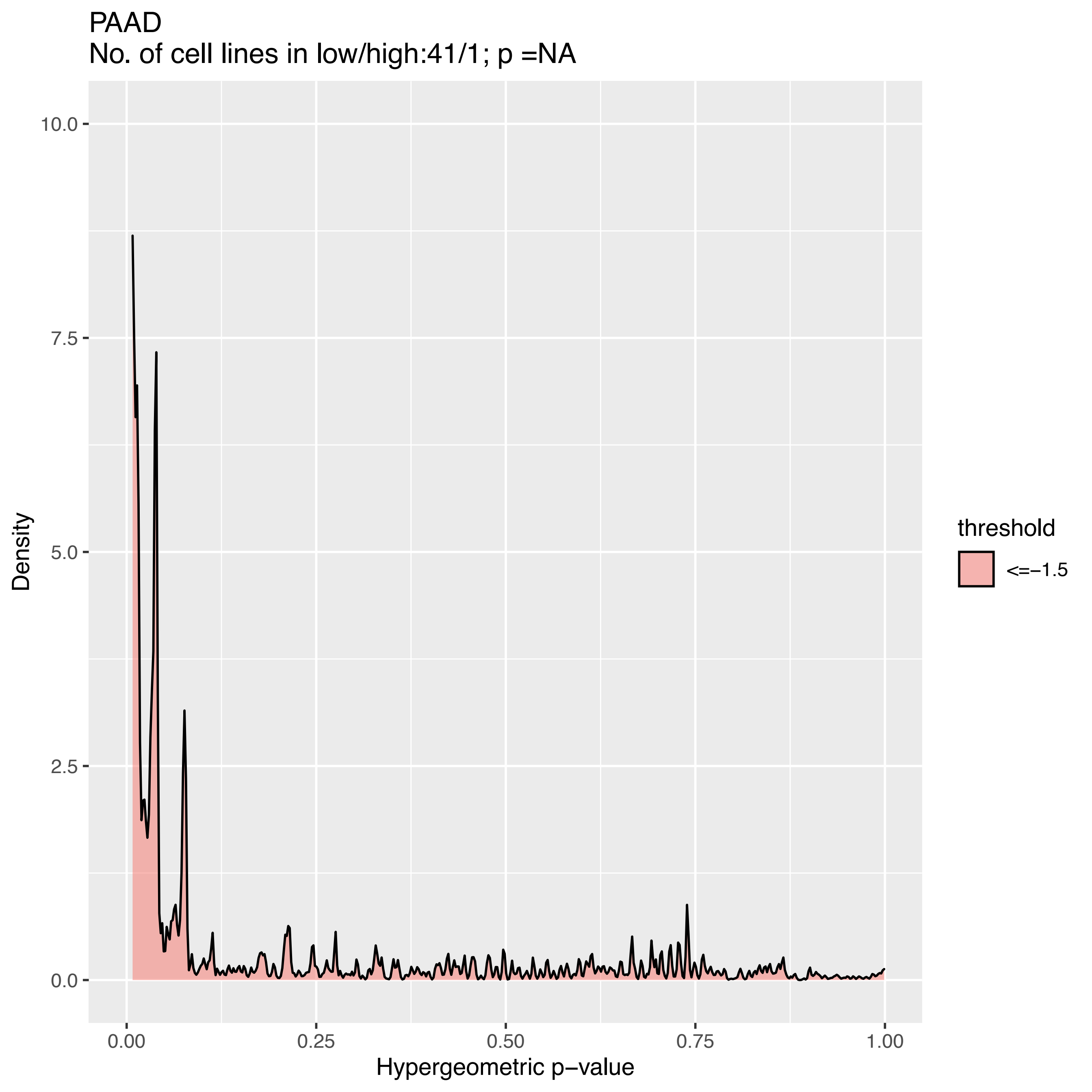

PAAD
No. of cell lines in low/high:41/1; p =NA
10.0
7.5
5.0
2.5
0.0
Density
0.00
0.25
0.50
0.75
1.00
Hypergeometric p−value
threshold
<=−1.5

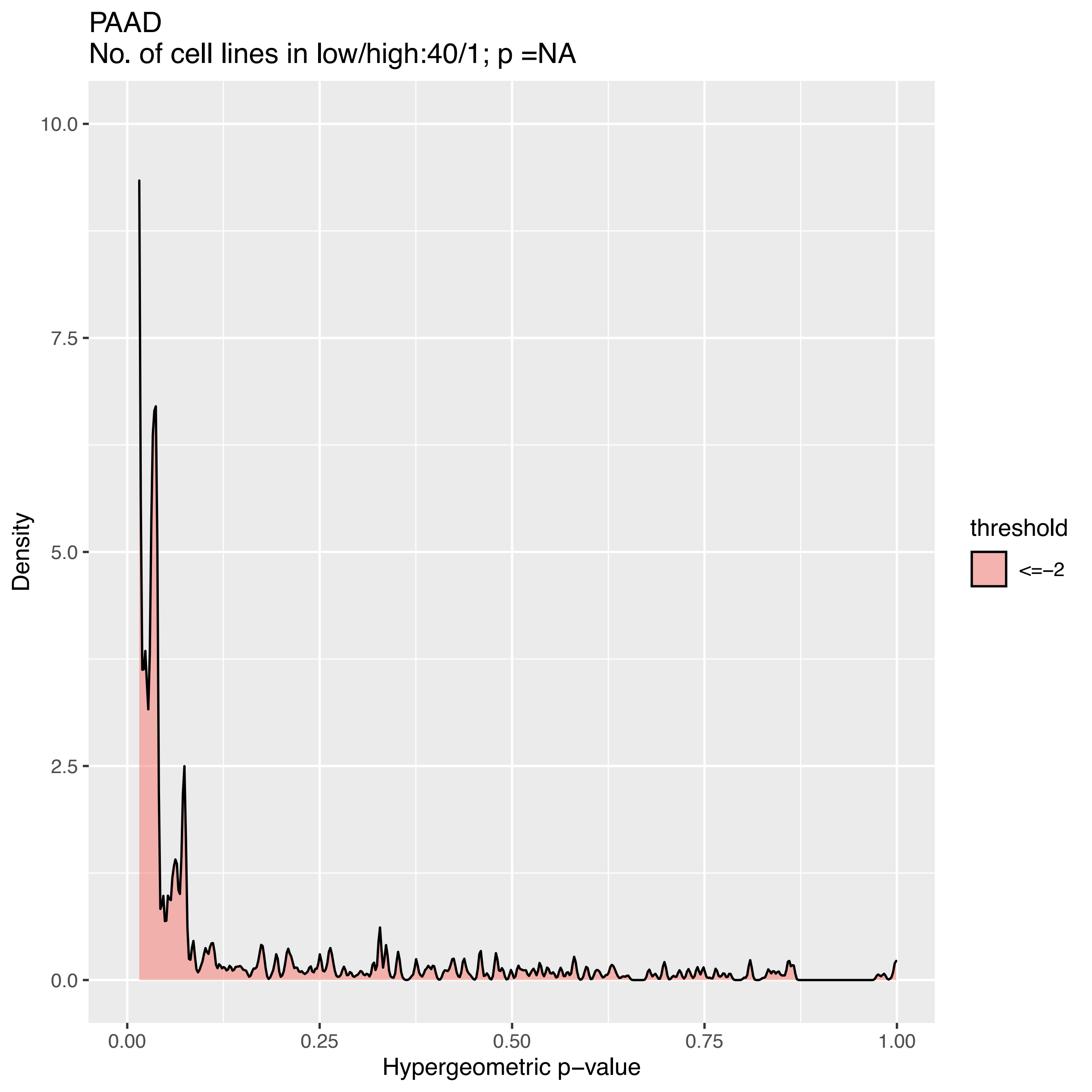

PAAD
No. of cell lines in low/high:40/1; p =NA
Density
10.0
7.5
5.0
2.5
0.0
0.00
0.25
0.50
0.75
1.00
Hypergeometric p−value
threshold
<=−2

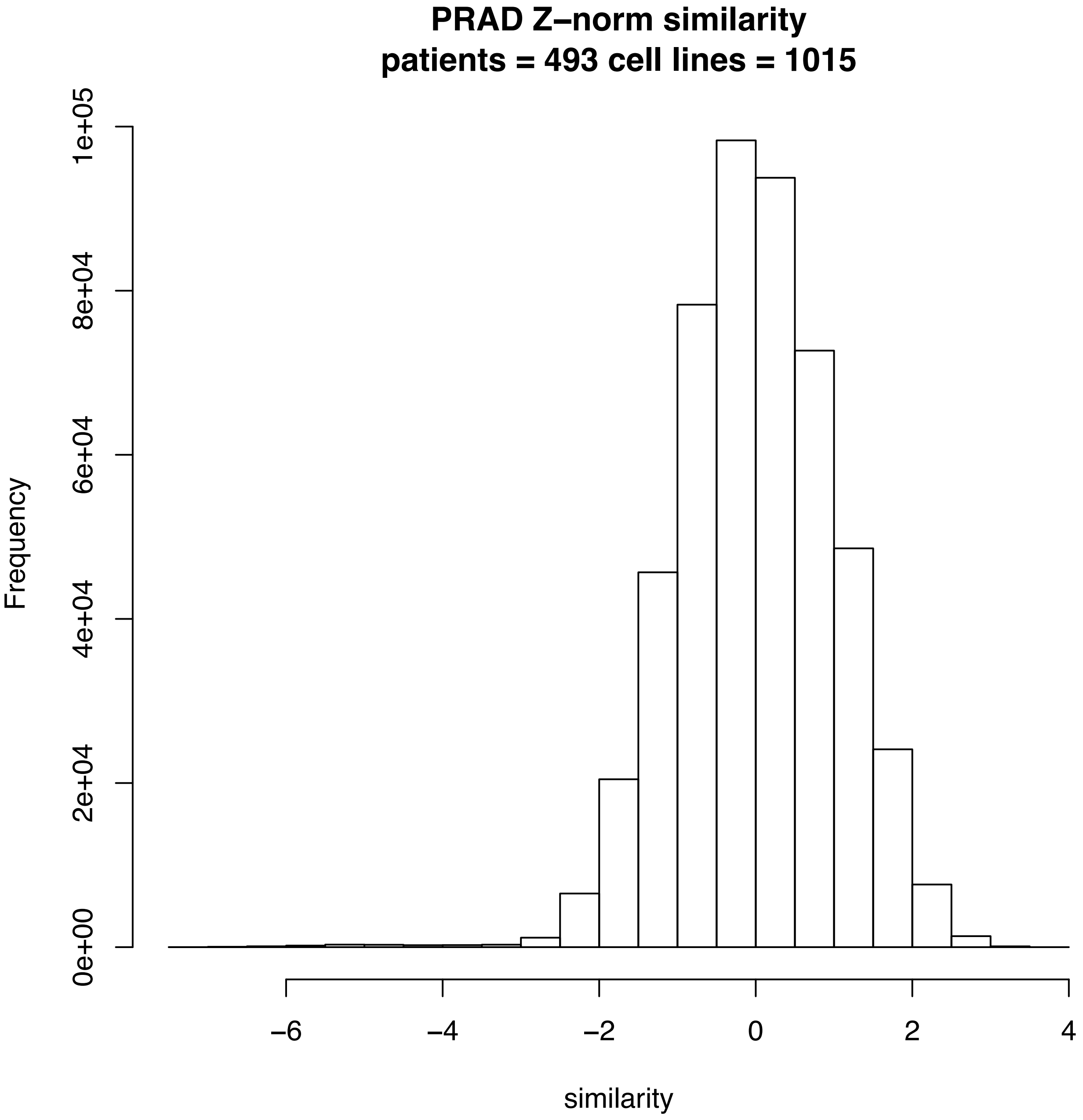

PRAD Z–norm similarity
patients = 493 cell lines = 1015
Frequency
0e+00
2e+04
4e+04
6e+04
8e+04
1e+05
−6
−4
−2
0
2
4
similarity

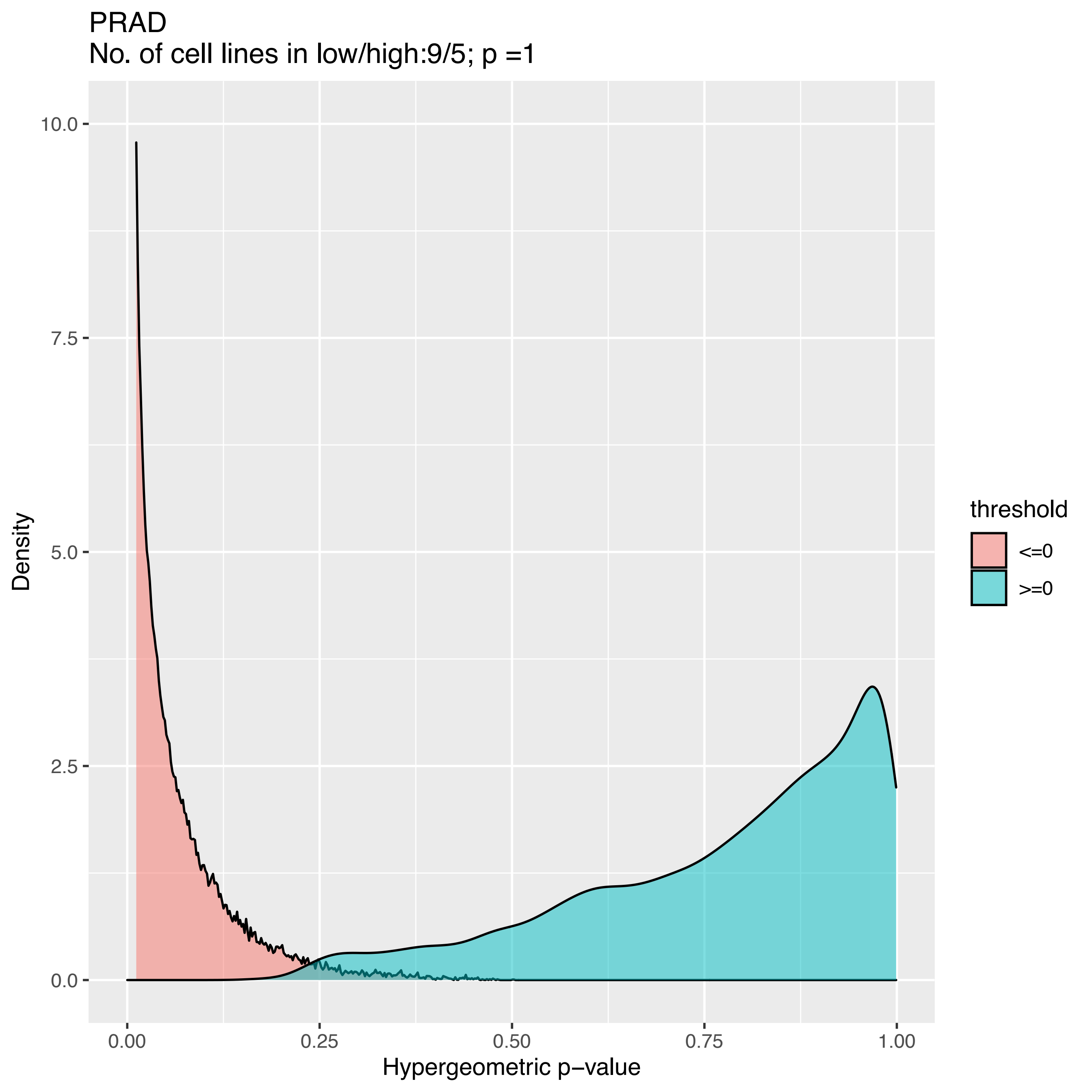

PRAD
No. of cell lines in low/high:9/5; p =1
Density
10.0
7.5
5.0
2.5
0.0
0.00
0.25
0.50
0.75
1.00
Hypergeometric p−value
threshold
<=0
>=0

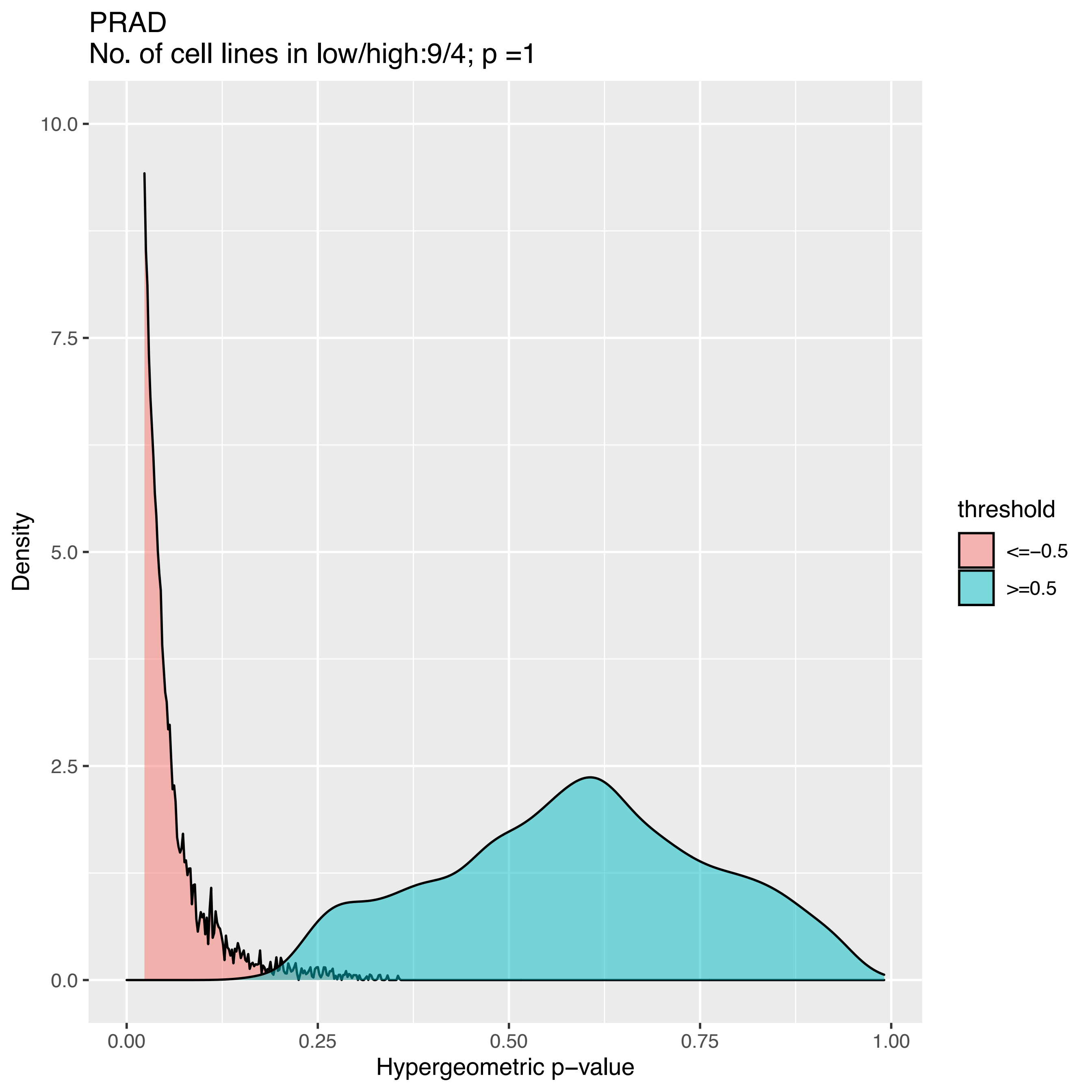

PRAD
No. of cell lines in low/high:9/4; p =1
Density
10.0
7.5
5.0
2.5
0.0
0.00
0.25
0.50
0.75
1.00
Hypergeometric p−value
threshold
<=−0.5
>=0.5

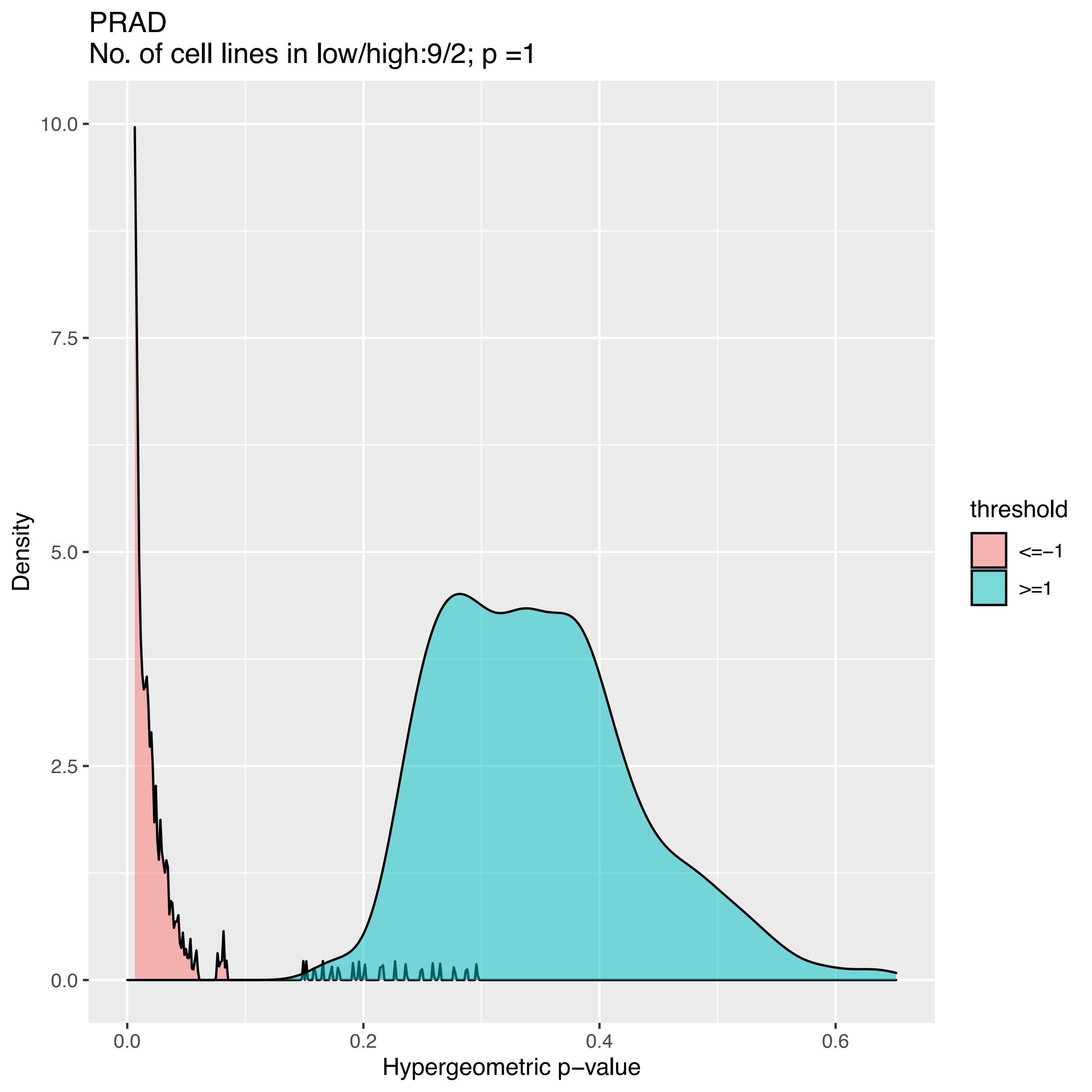

PRAD
No. of cell lines in low/high:9/2; p =1
Density
10.0
7.5
5.0
2.5
0.0
0.0
0.2
0.4
0.6
Hypergeometric p−value
threshold
<=−1
>=1

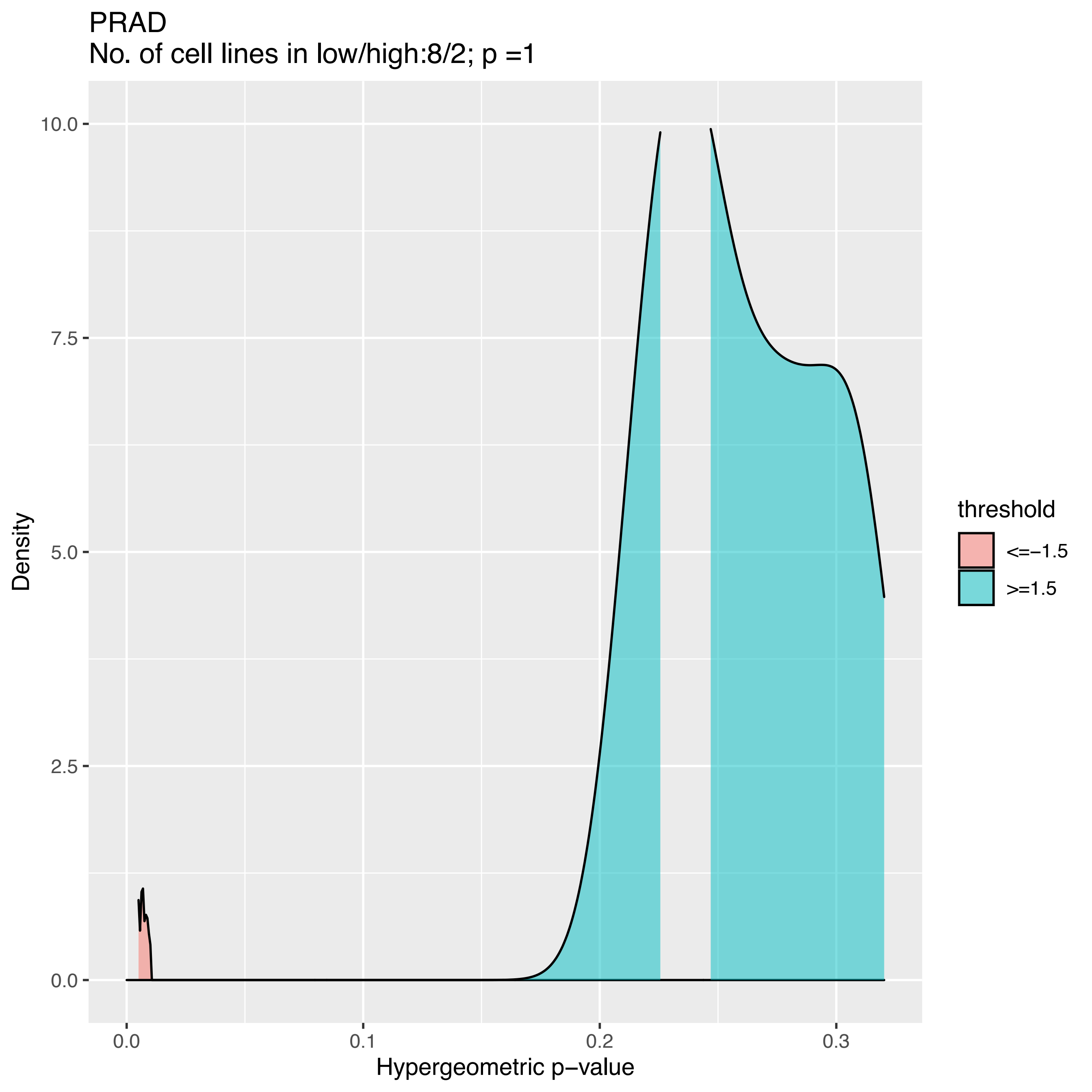
PRAD
No. of cell lines in low/high:8/2; p =1
10.0
7.5
5.0
2.5
0.0
Density
0.0
0.1
0.2
0.3
Hypergeometric p−value
threshold
<=−1.5
>=1.5

PRAD
No. of cell lines in low/high:8/1; p =NA

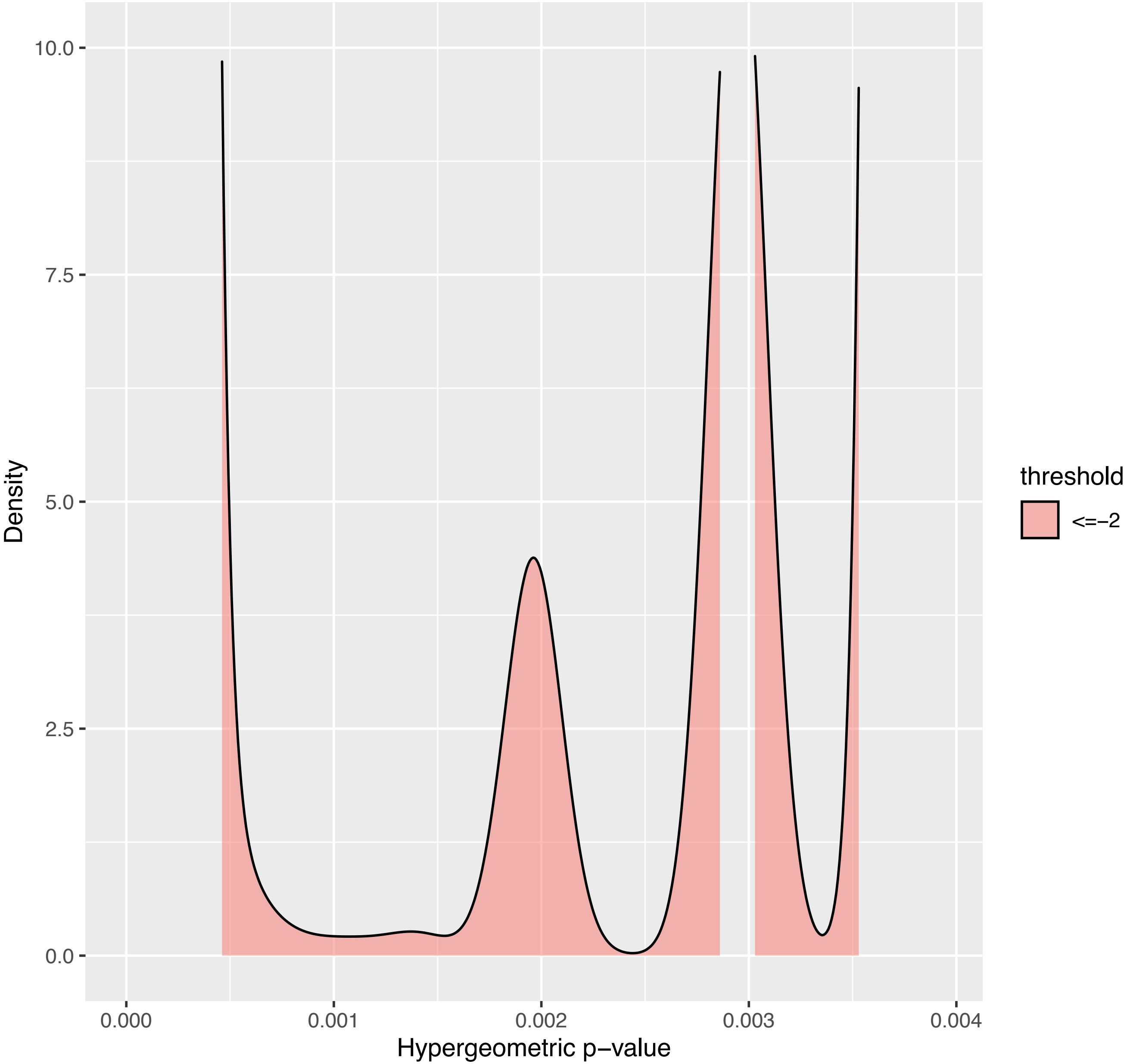

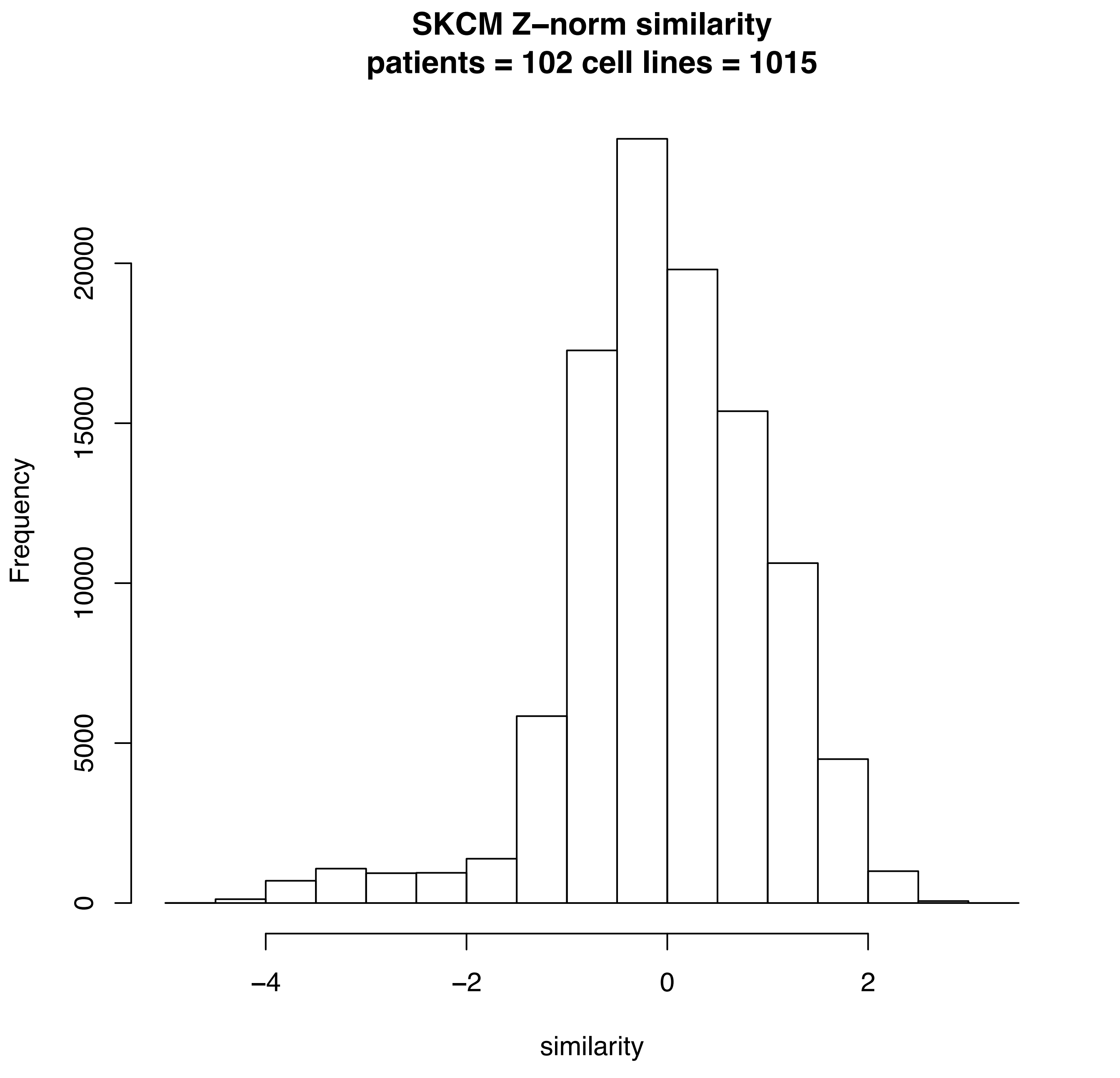
SKCM Z–norm similarity
patients = 102 cell lines = 1015
Frequency
0
5000
10000
15000
20000
−4
−2
0
2
similarity

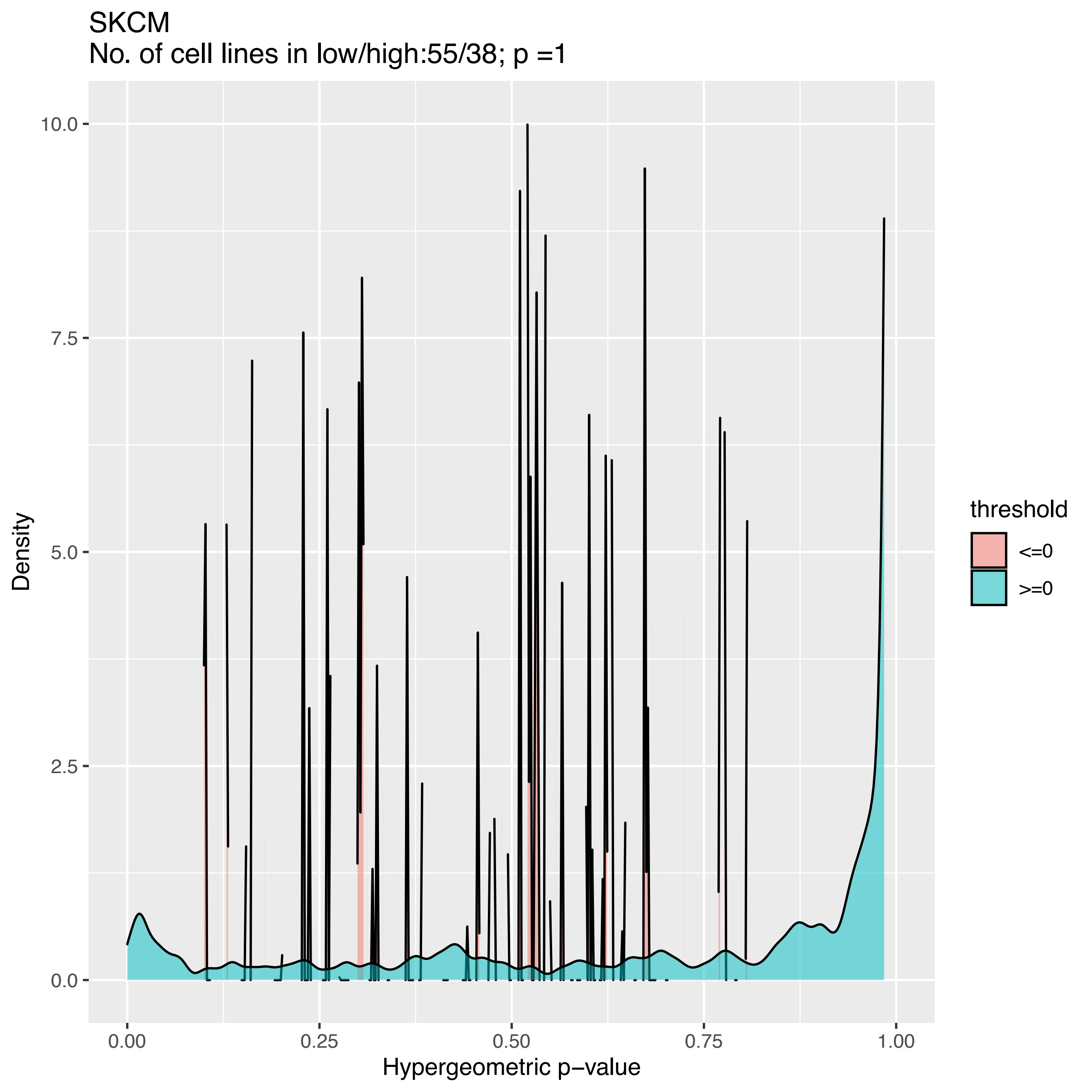
SKCM
No. of cell lines in low/high:55/38; p =1
Density
10.0
7.5
5.0
2.5
0.0
0.00
0.25
0.50
0.75
1.00
Hypergeometric p−value
threshold
<=0
>=0

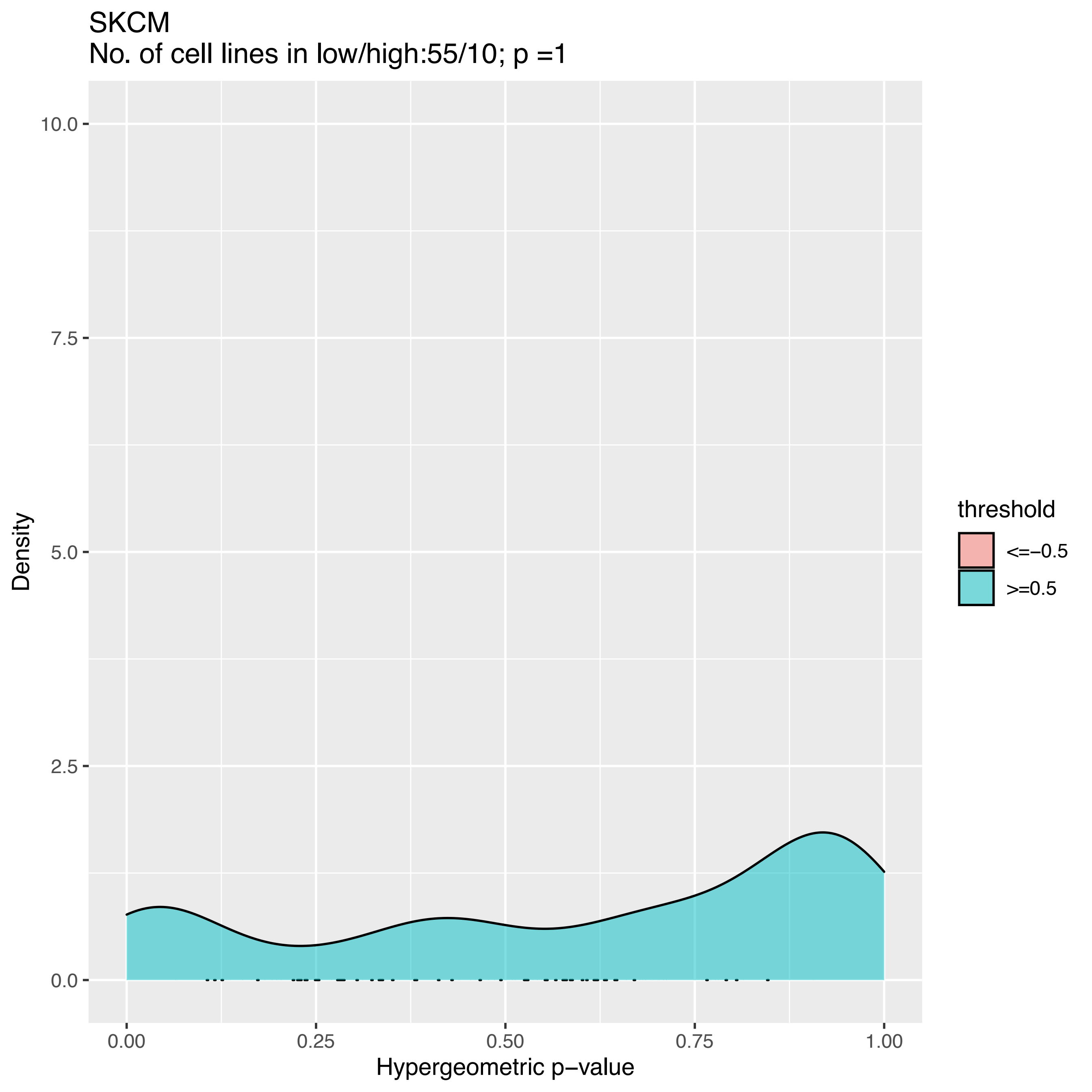

SKCM
No. of cell lines in low/high:55/10; p =1
Density
10.0
7.5
5.0
2.5
0.0
0.00
0.25
0.50
0.75
1.00
Hypergeometric p−value
threshold
<=−0.5
>=0.5

SKCM
No. of cell lines in low/high:55/3; p =1

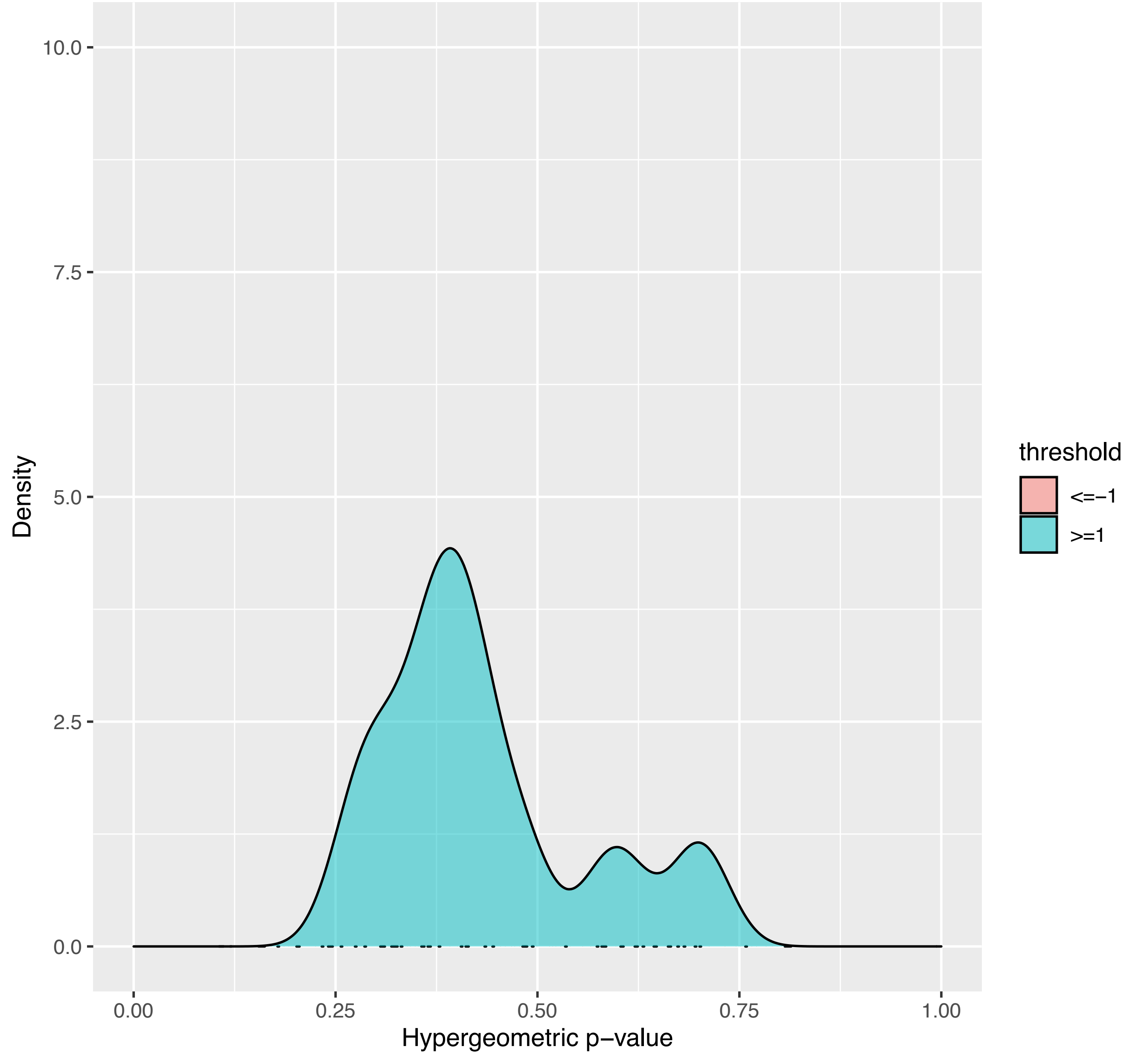

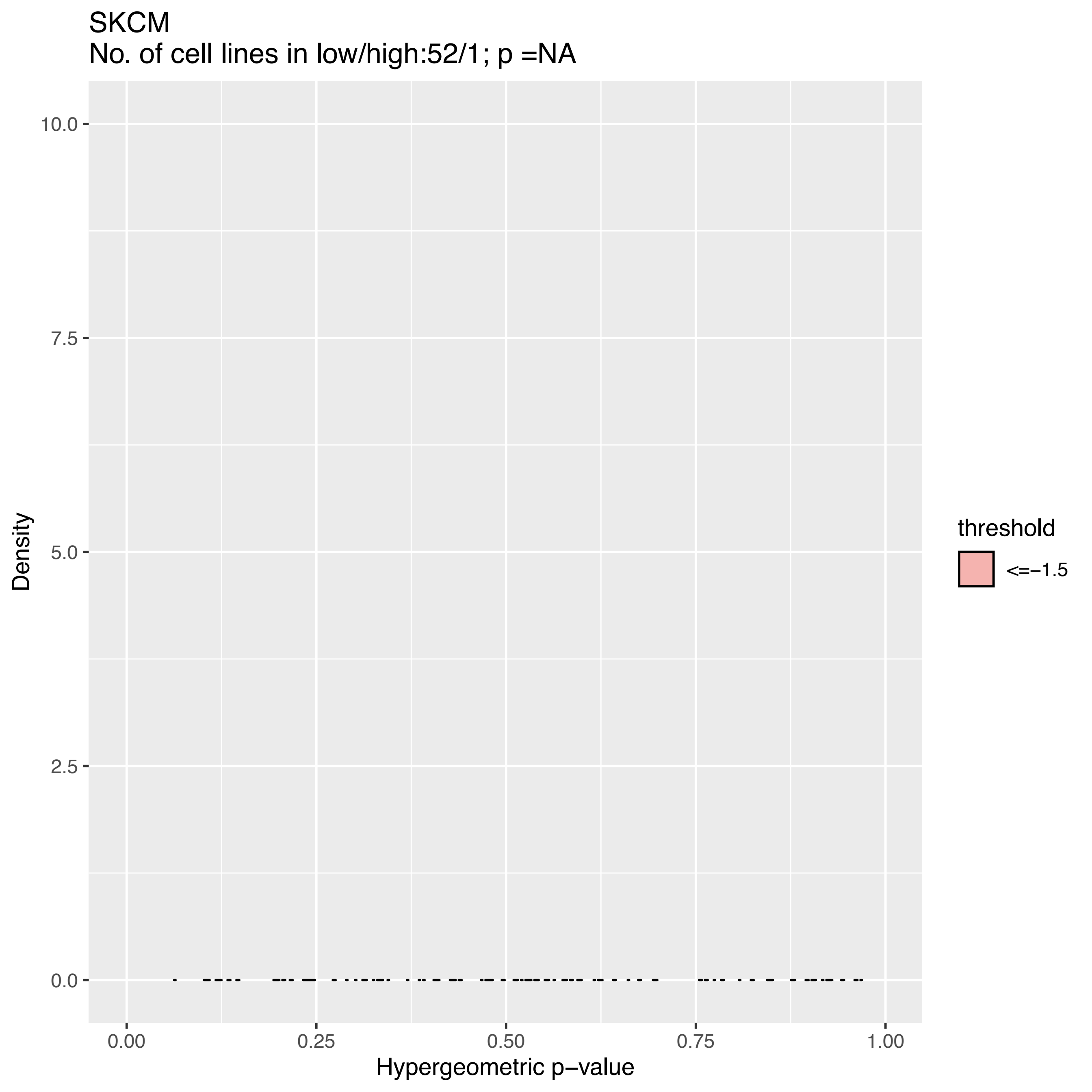

SKCM
No. of cell lines in low/high:52/1; p =NA
10.0
7.5
5.0
2.5
0.0
0.00
0.25
0.50
0.75
1.00
Density
Hypergeometric p−value
threshold
<=−1.5

SKCM
No. of cell lines in low/high:49/1; p =NA

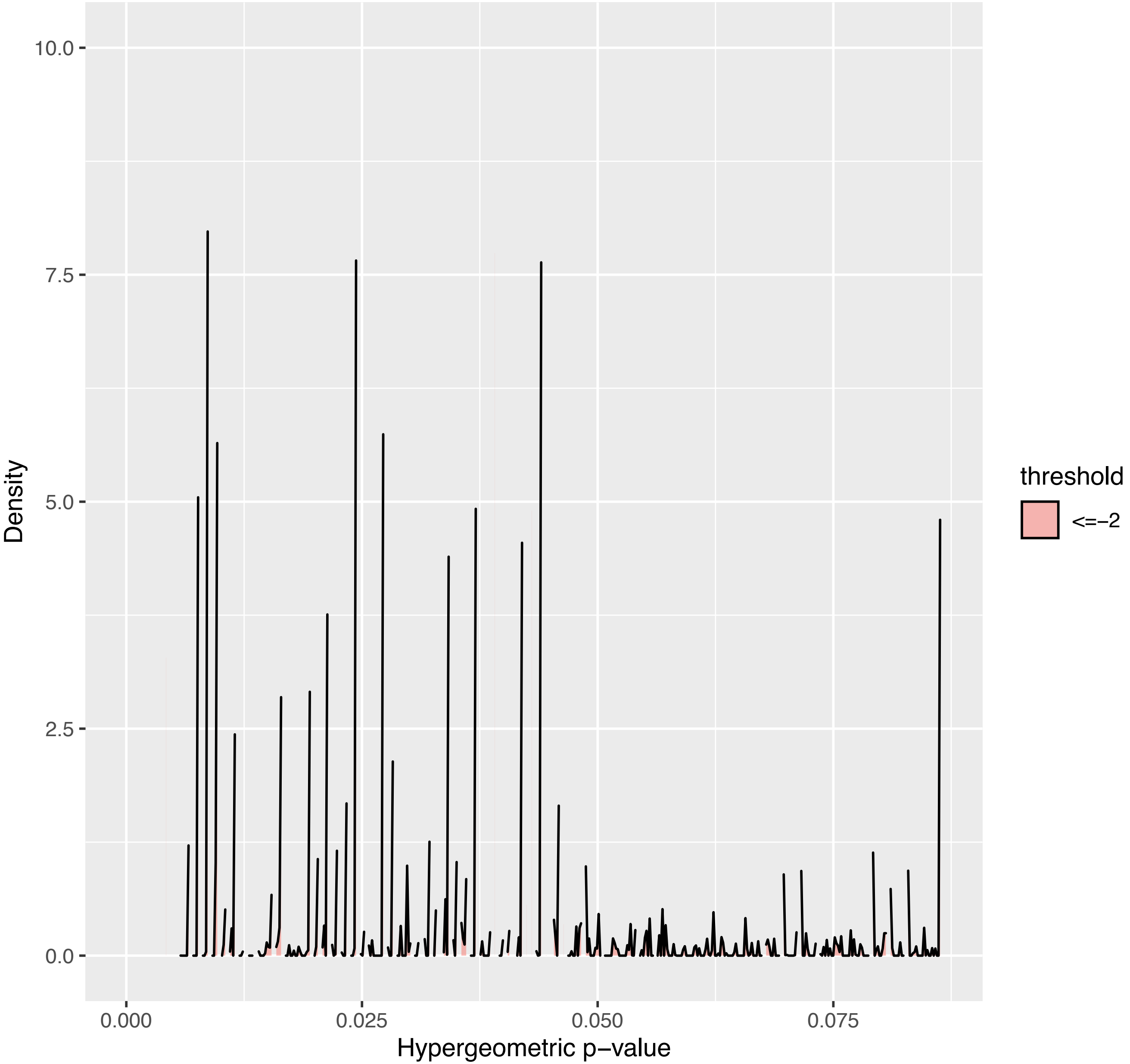

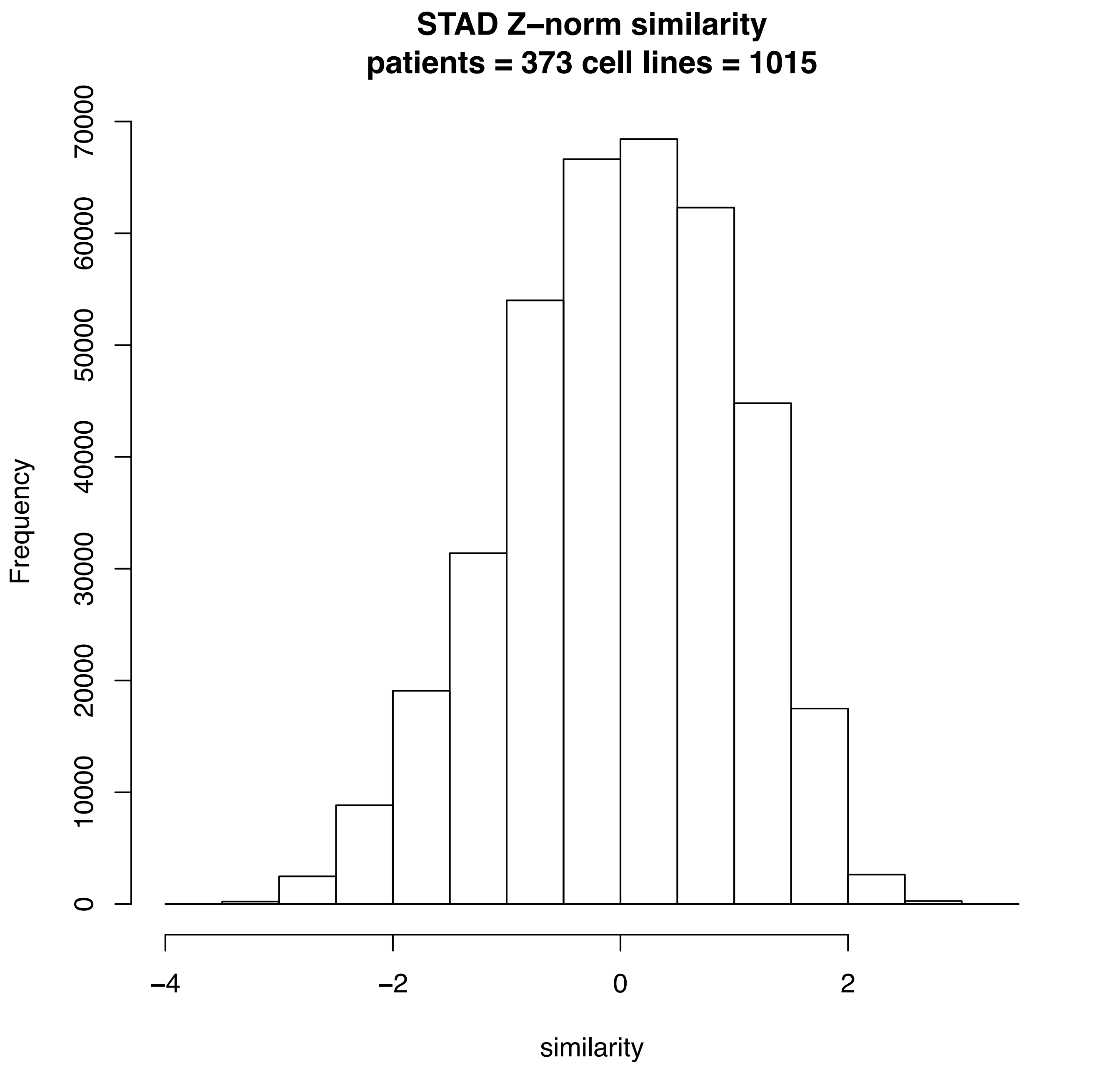

STAD Z–norm similarity
patients = 373 cell lines = 1015
Frequency
0
10000
20000
30000
40000
50000
60000
70000
–4
–2
0
2
similarity

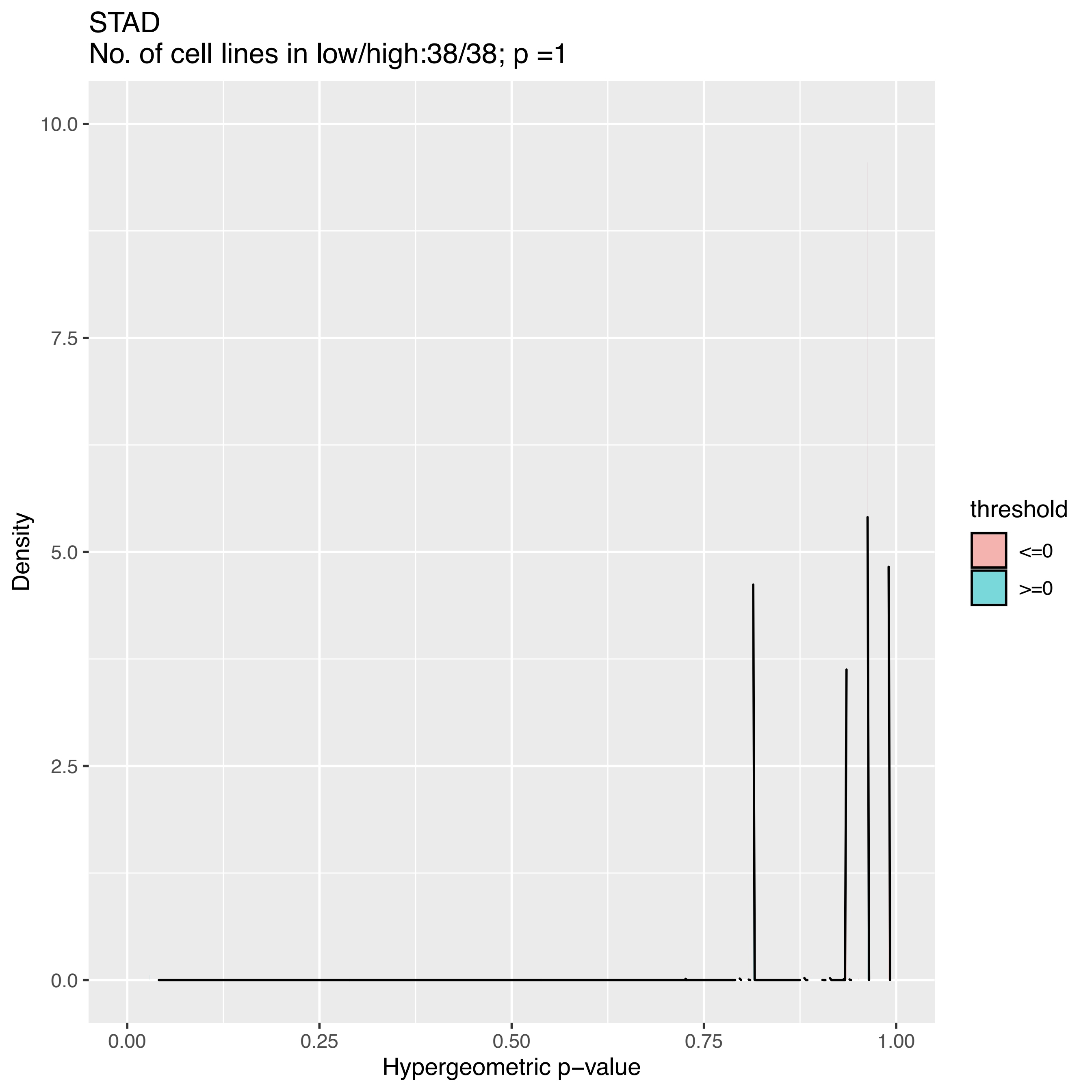
STAD
No. of cell lines in low/high:38/38; p =1
10.0
7.5
5.0
2.5
0.0
Density
0.00
0.25
0.50
0.75
1.00
Hypergeometric p−value
threshold
<=0
>=0

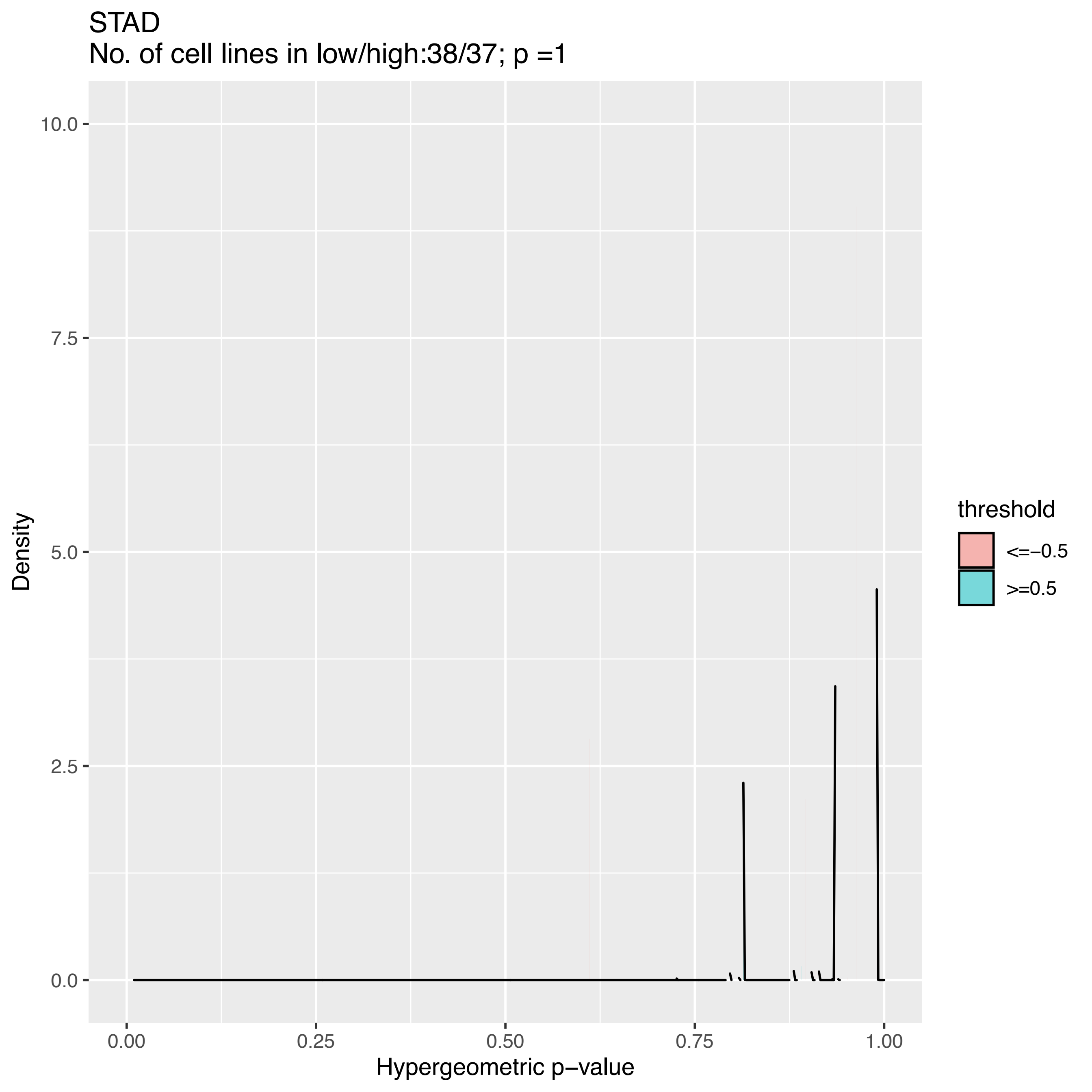
STAD
No. of cell lines in low/high:38/37; p =1
Density
10.0
7.5
5.0
2.5
0.0
0.00
0.25
0.50
0.75
1.00
Hypergeometric p−value
threshold
<=−0.5
>=0.5

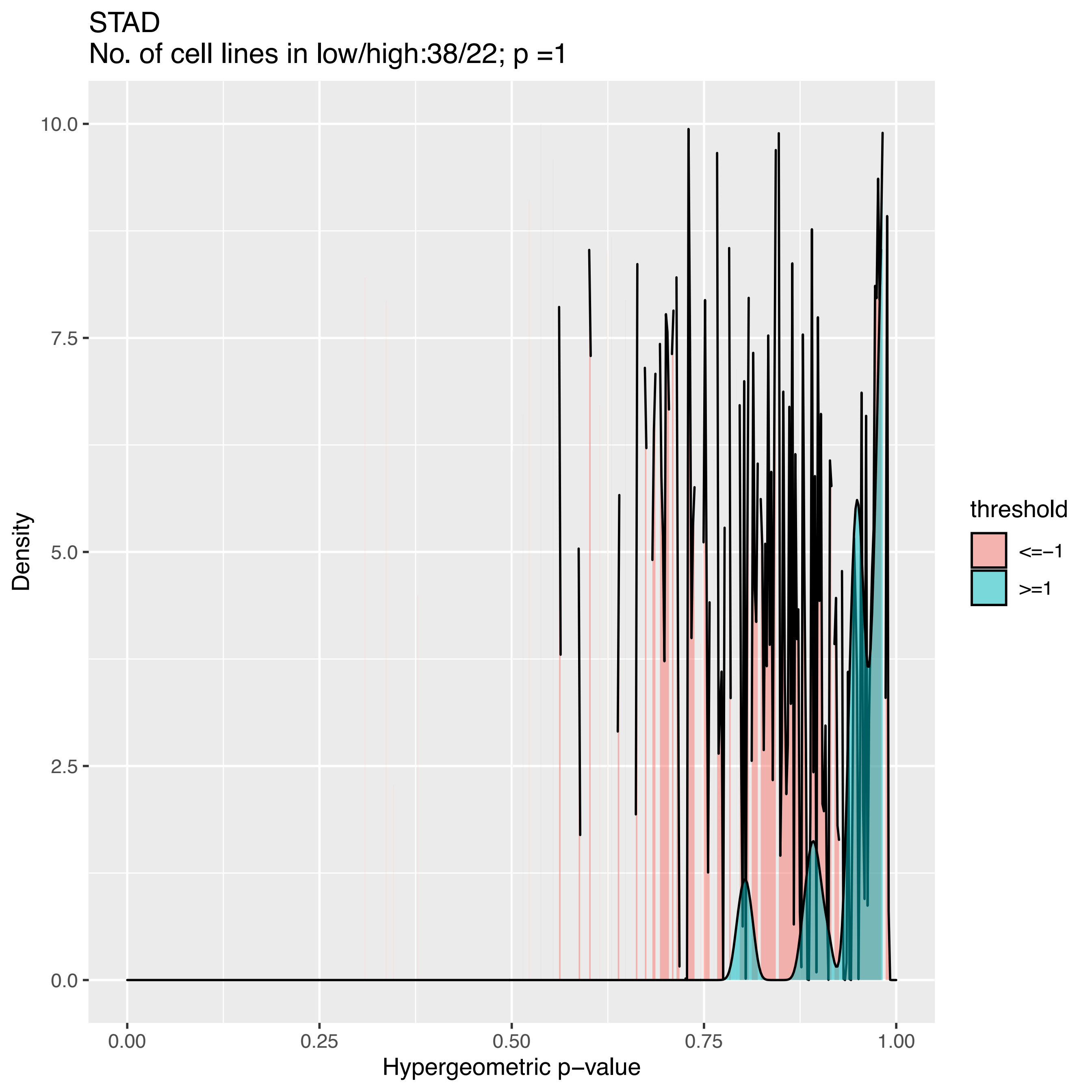

STAD
No. of cell lines in low/high:38/22; p =1
Density
10.0
7.5
5.0
2.5
0.0
0.00
0.25
0.50
0.75
1.00
Hypergeometric p−value
threshold
<=−1
>=1

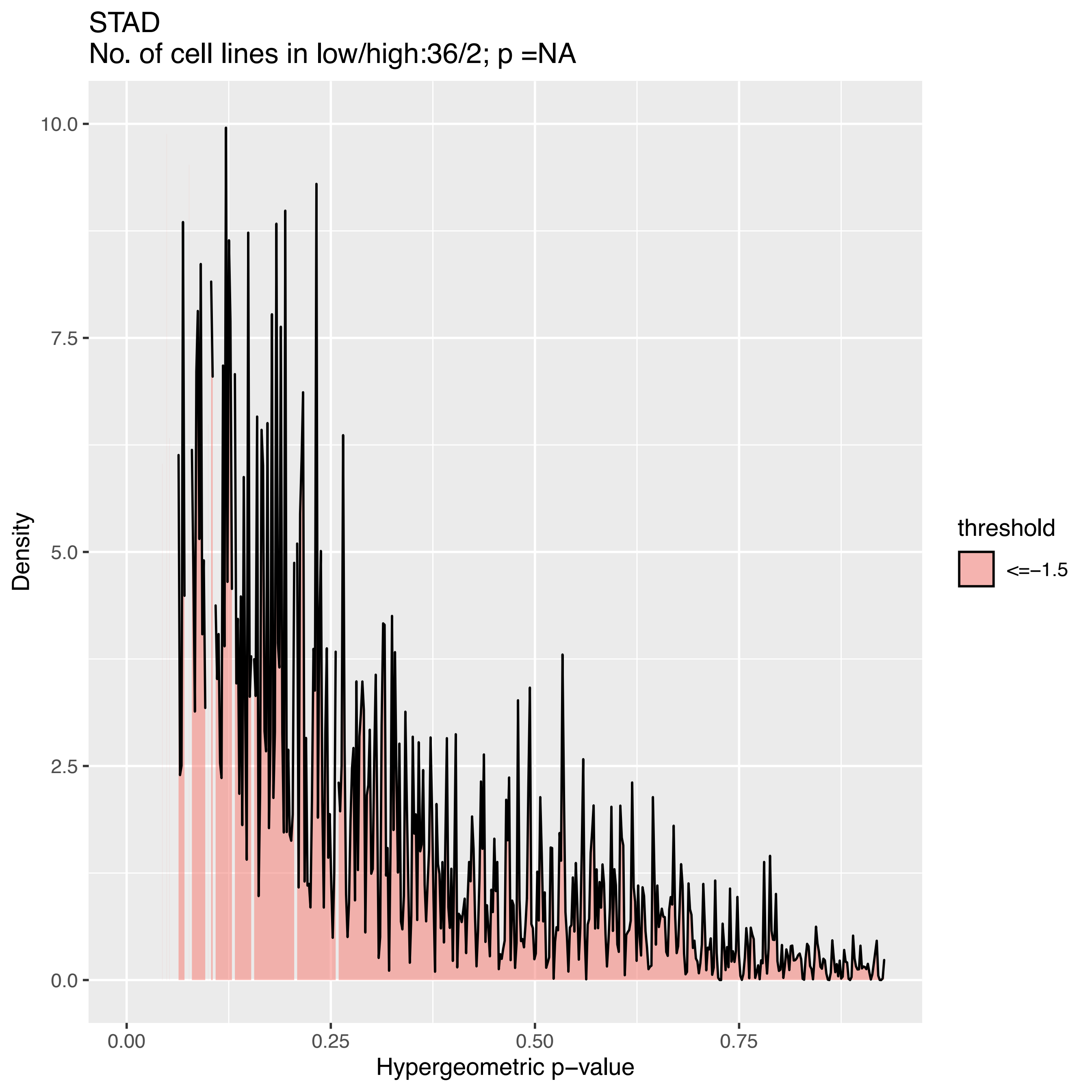

STAD
No. of cell lines in low/high:36/2; p =NA
Density
10.0
7.5
5.0
2.5
0.0
0.00
0.25
0.50
0.75
Hypergeometric p−value
threshold
<=−1.5

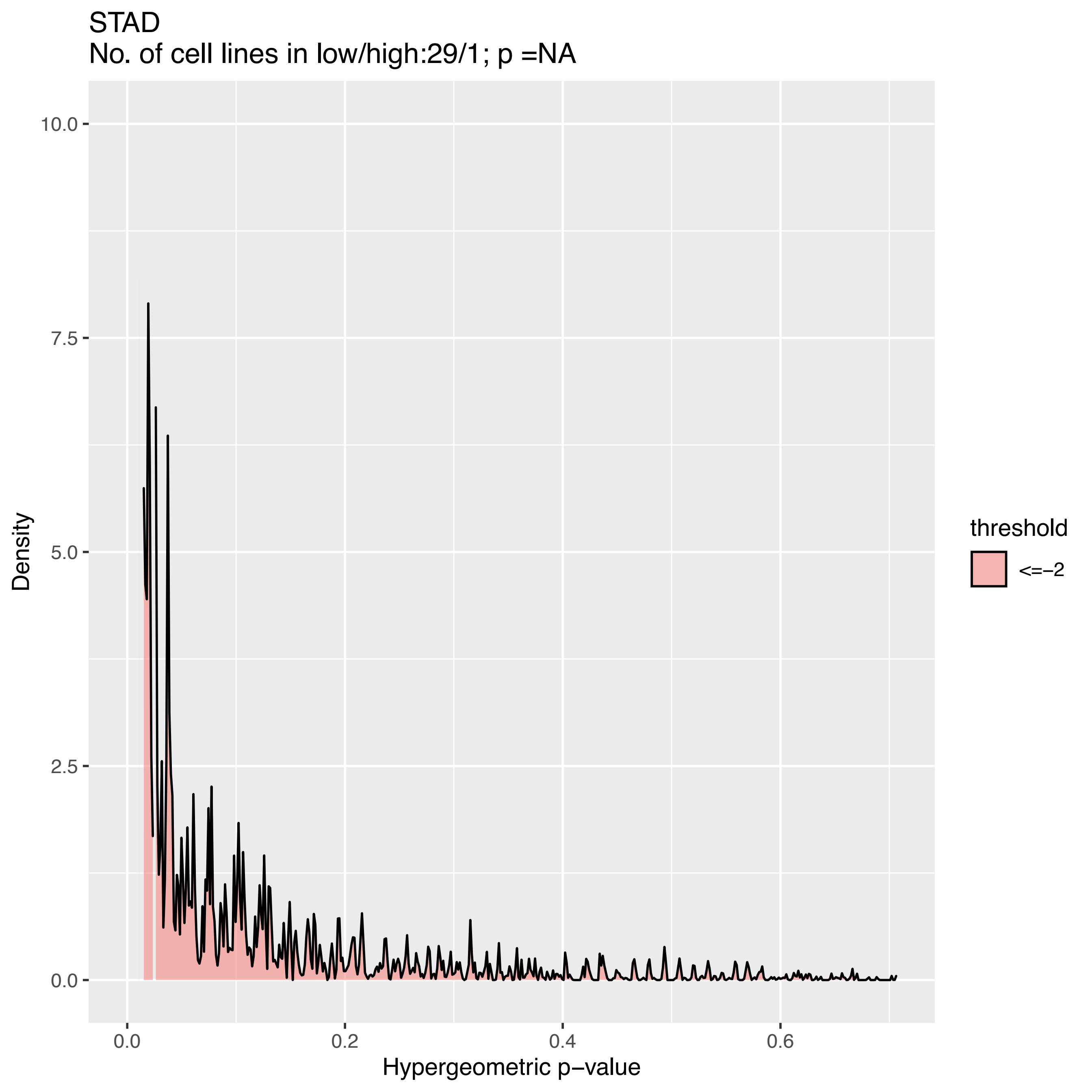
STAD
No. of cell lines in low/high:29/1; p =NA
10.0
7.5
5.0
2.5
0.0
Density
0.0
0.2
0.4
0.6
Hypergeometric p−value
threshold
<=−2

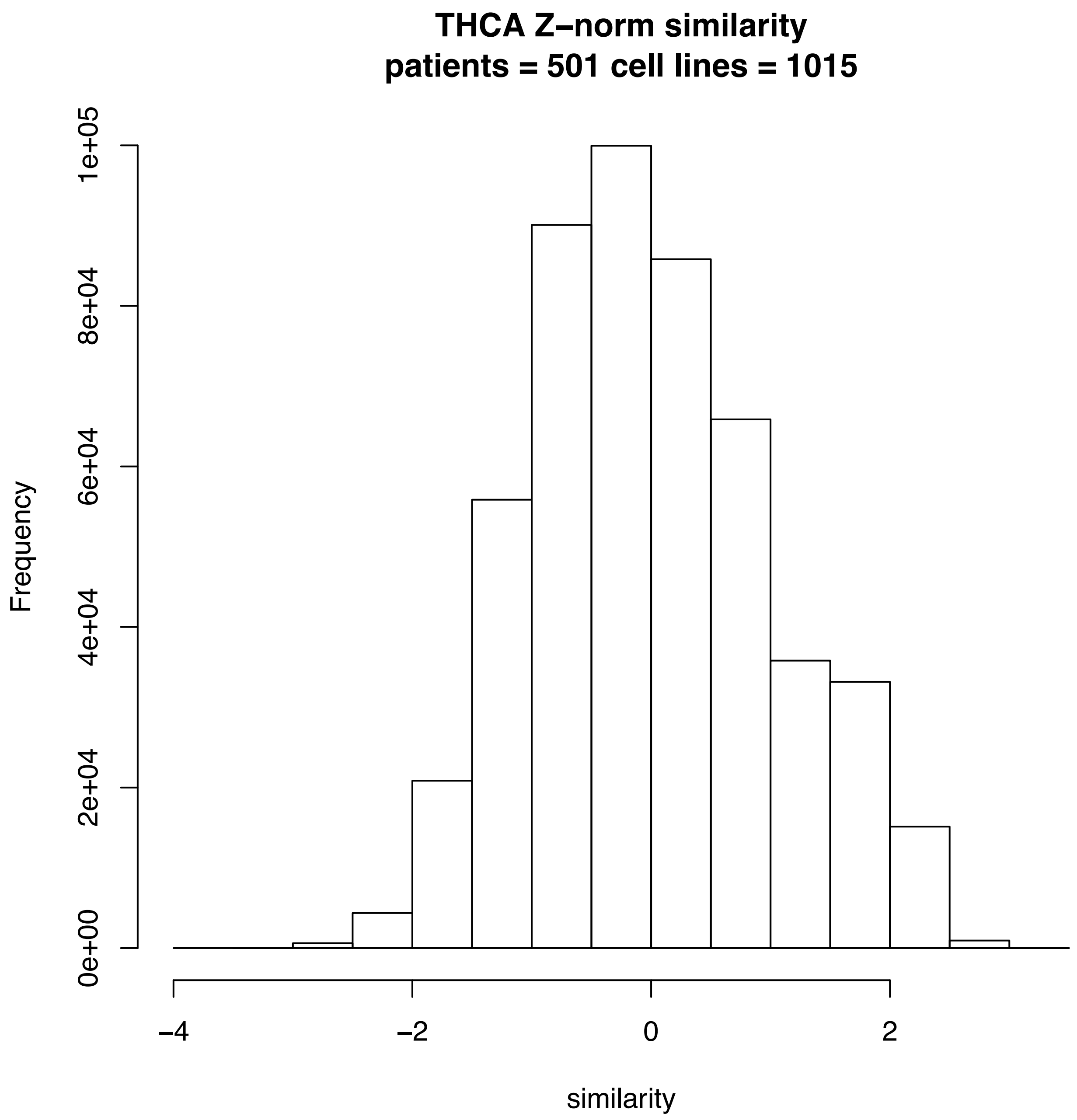
THCA Z–norm similarity
patients = 501 cell lines = 1015
Frequency
0e+00
2e+04
4e+04
6e+04
8e+04
1e+05
−4
−2
0
2
similarity

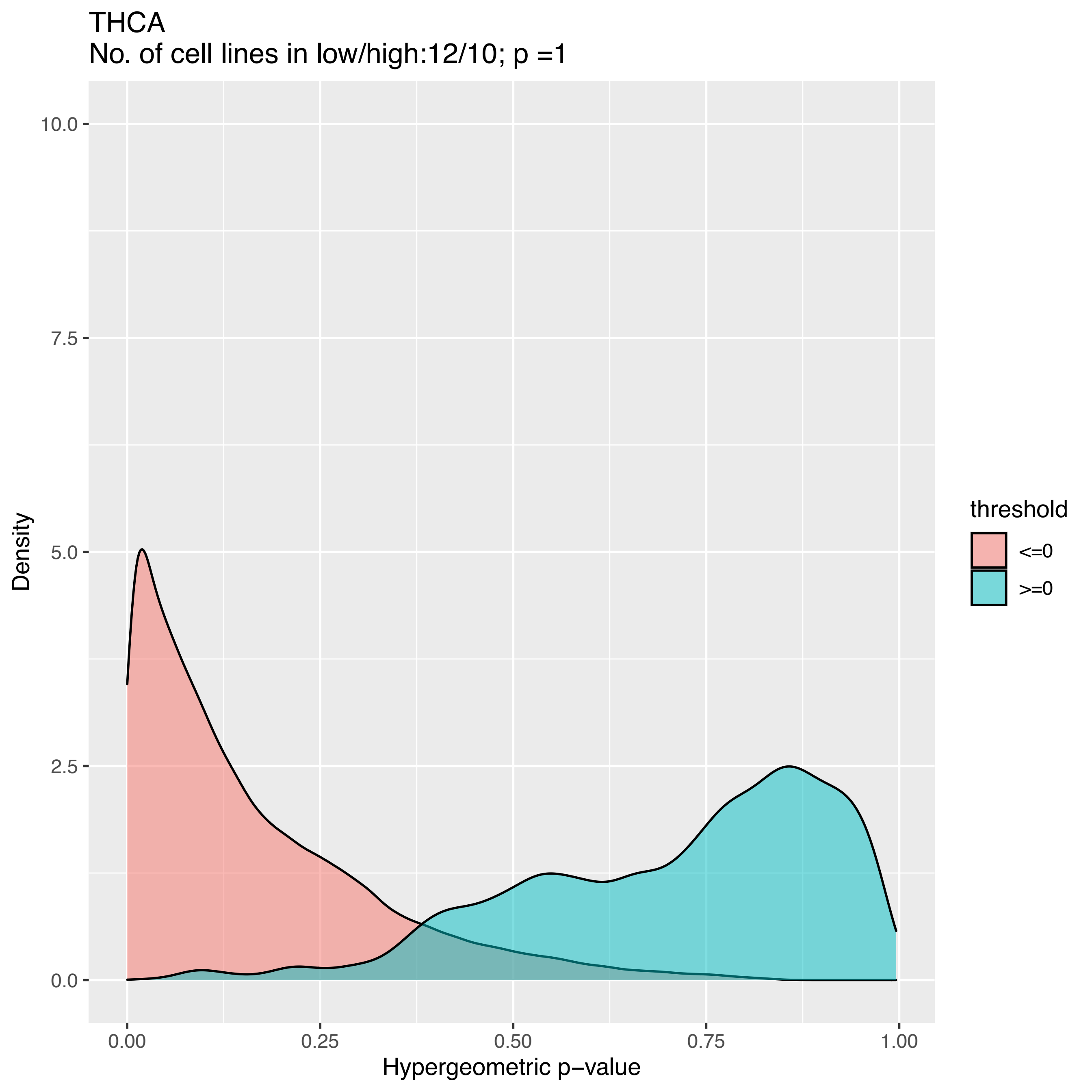

THCA
No. of cell lines in low/high:12/10; p =1
Density
10.0
7.5
5.0
2.5
0.0
0.00
0.25
0.50
0.75
1.00
Hypergeometric p−value
threshold
<=0
>=0

THCA
No. of cell lines in low/high:11/6; p =1

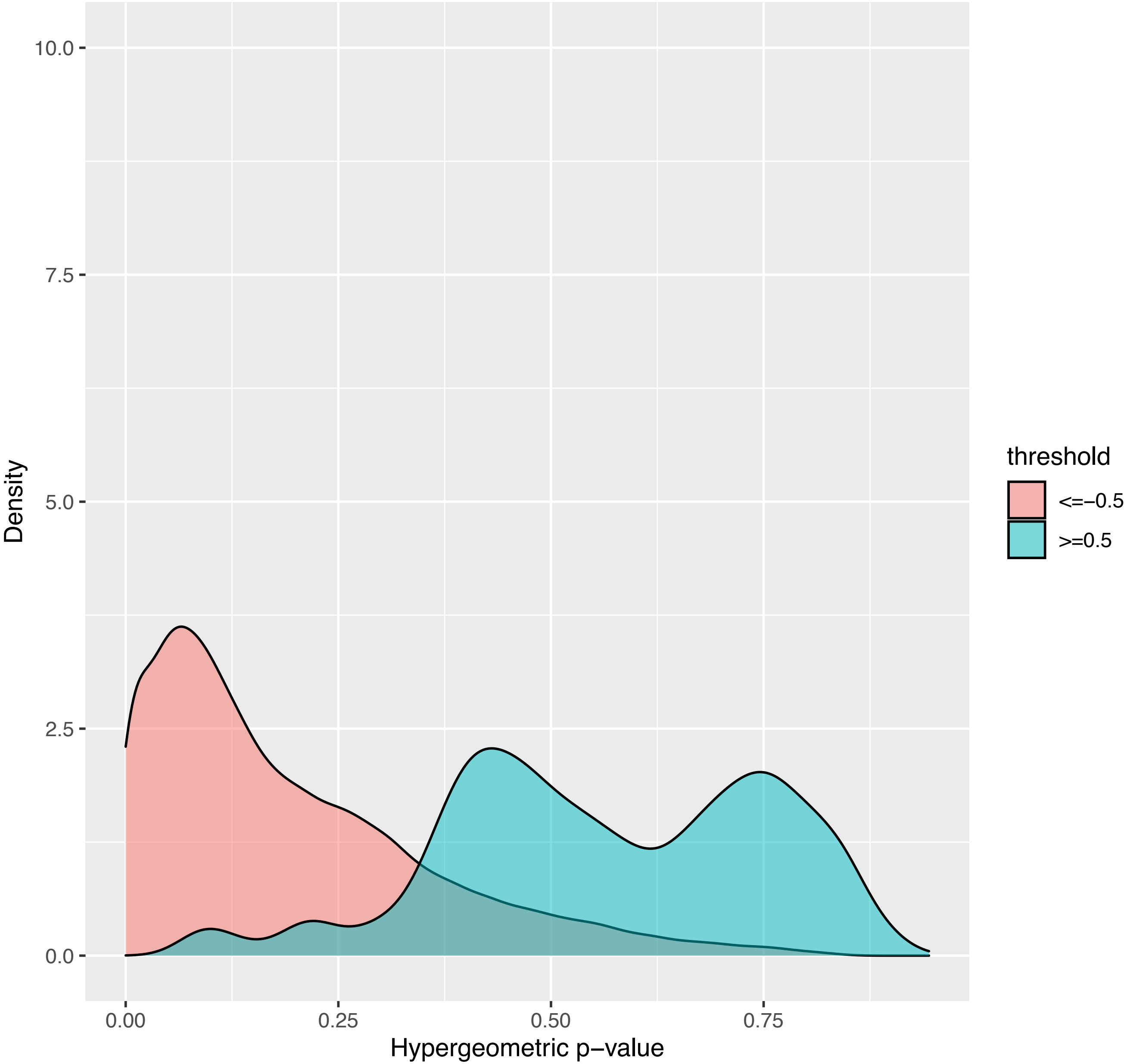

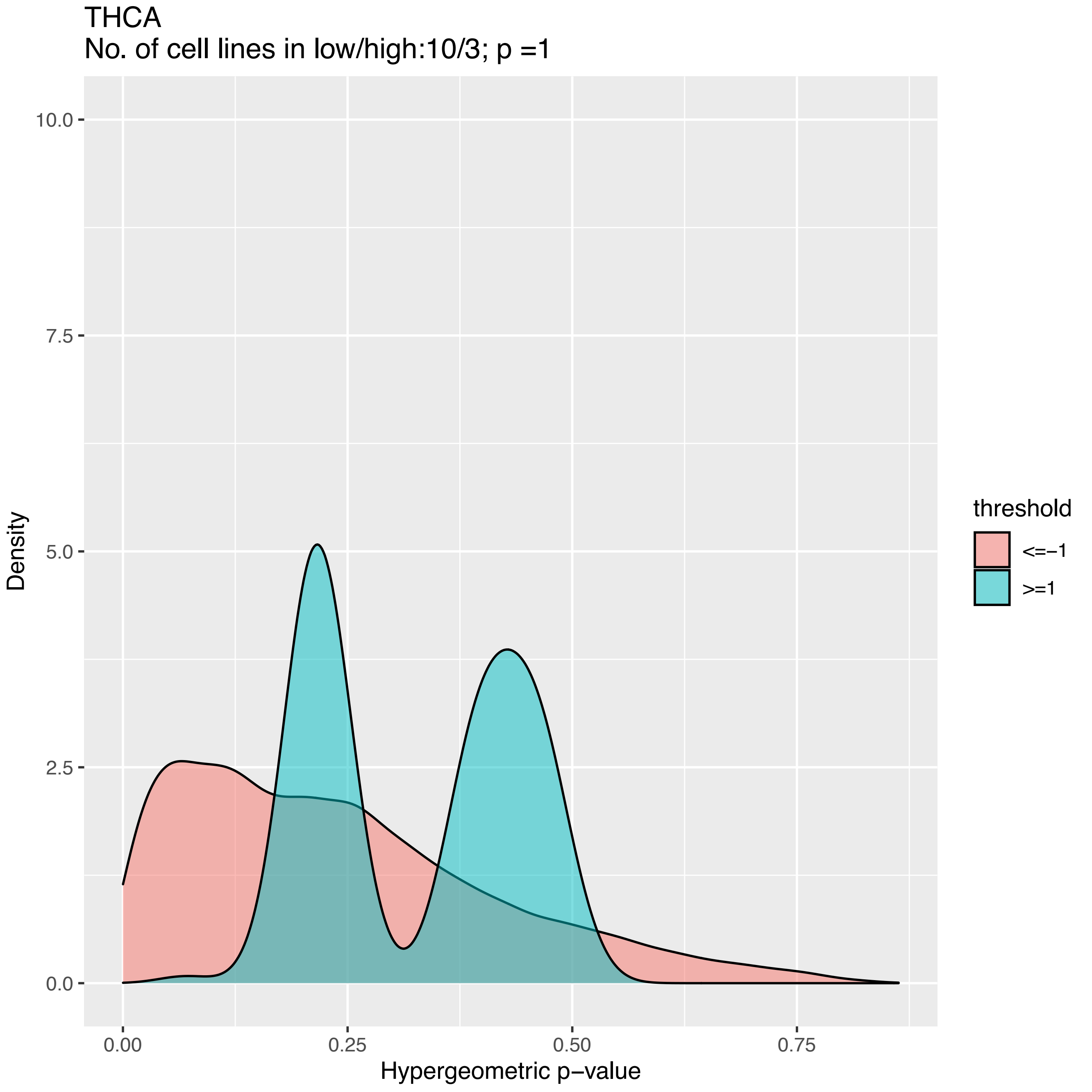
THCA
No. of cell lines in low/high:10/3; p =1
Density
10.0
7.5
5.0
2.5
0.0
0.00
0.25
0.50
0.75
Hypergeometric p−value
threshold
<=−1
>=1

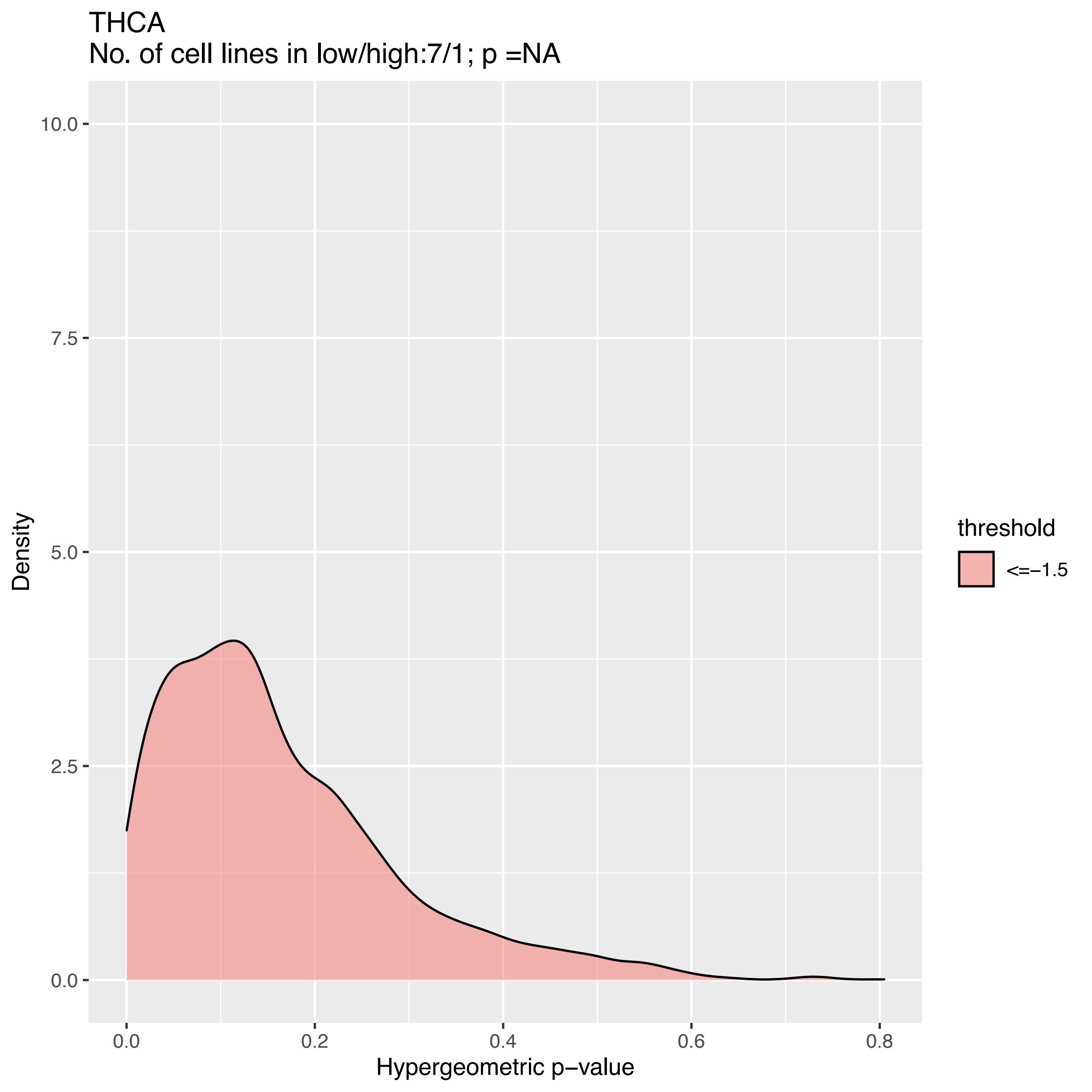

THCA
No. of cell lines in low/high:7/1; p =NA
Density
10.0
7.5
5.0
2.5
0.0
0.0
0.2
0.4
0.6
0.8
Hypergeometric p−value
threshold
<=−1.5

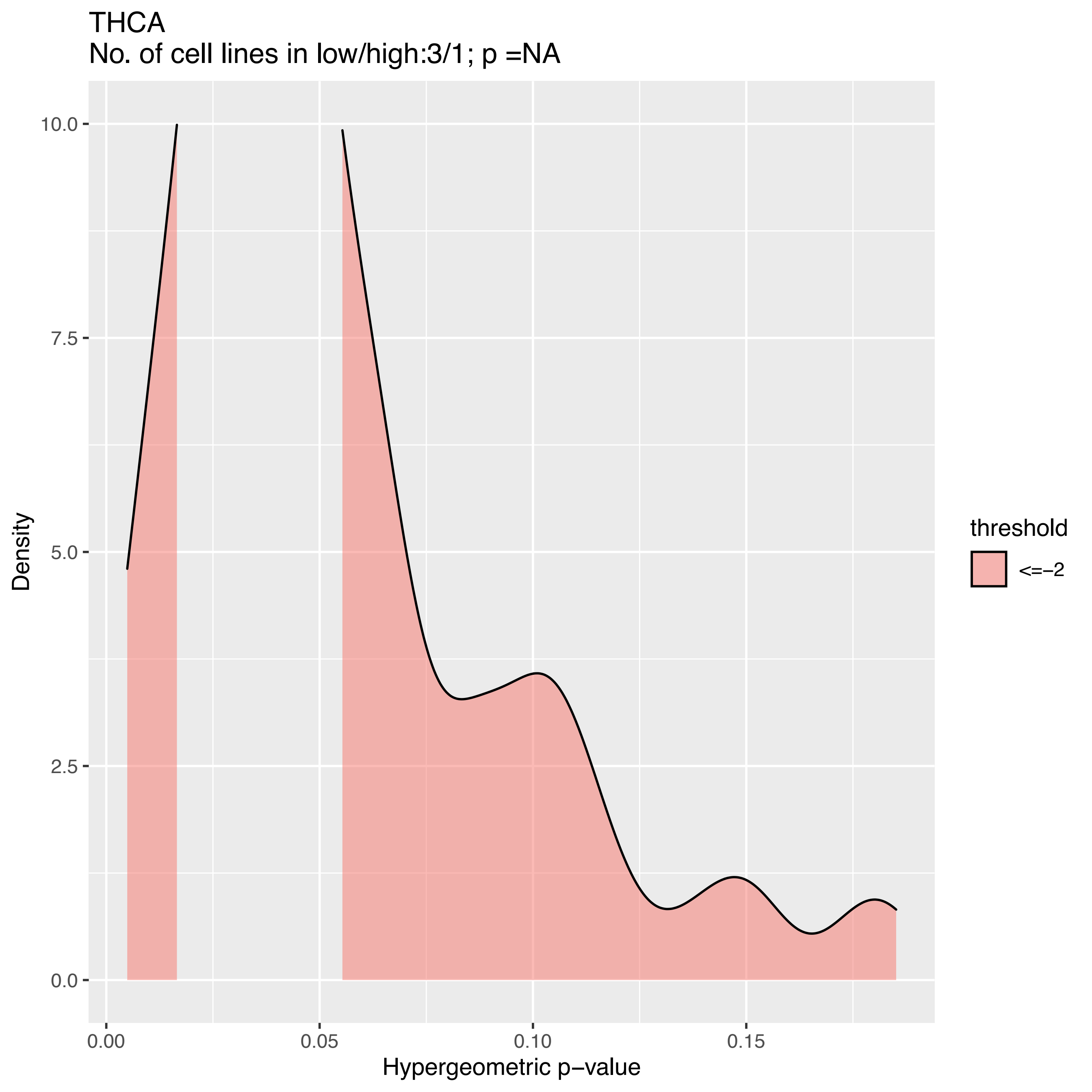
THCA
No. of cell lines in low/high:3/1; p =NA
Density
10.0
7.5
5.0
2.5
0.0
0.00
0.05
0.10
0.15
Hypergeometric p−value
threshold
<=−2

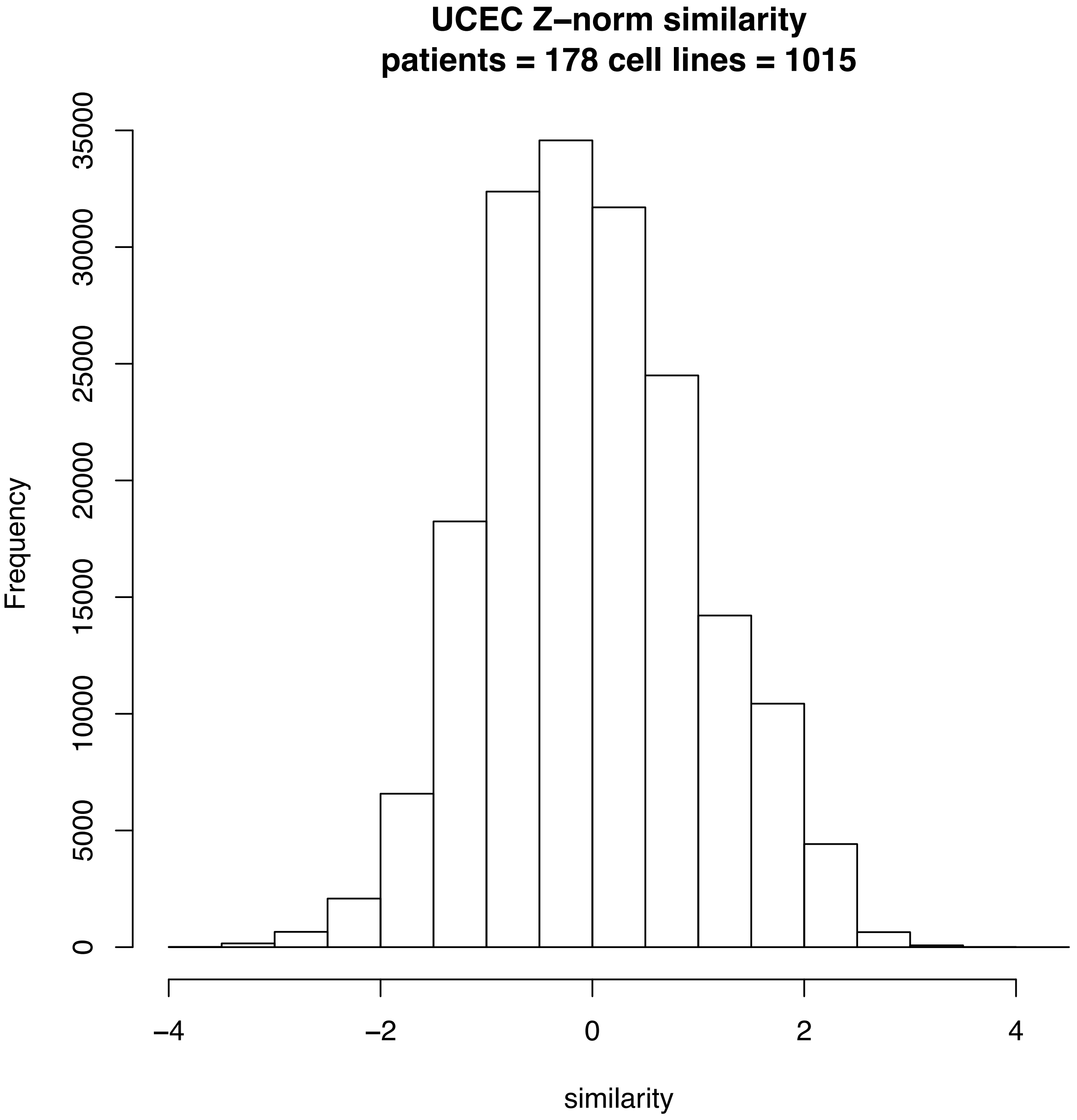

UCEC Z–norm similarity
patients = 178 cell lines = 1015
Frequency
0
5000
10000
15000
20000
25000
30000
35000
–4
–2
0
2
4
similarity

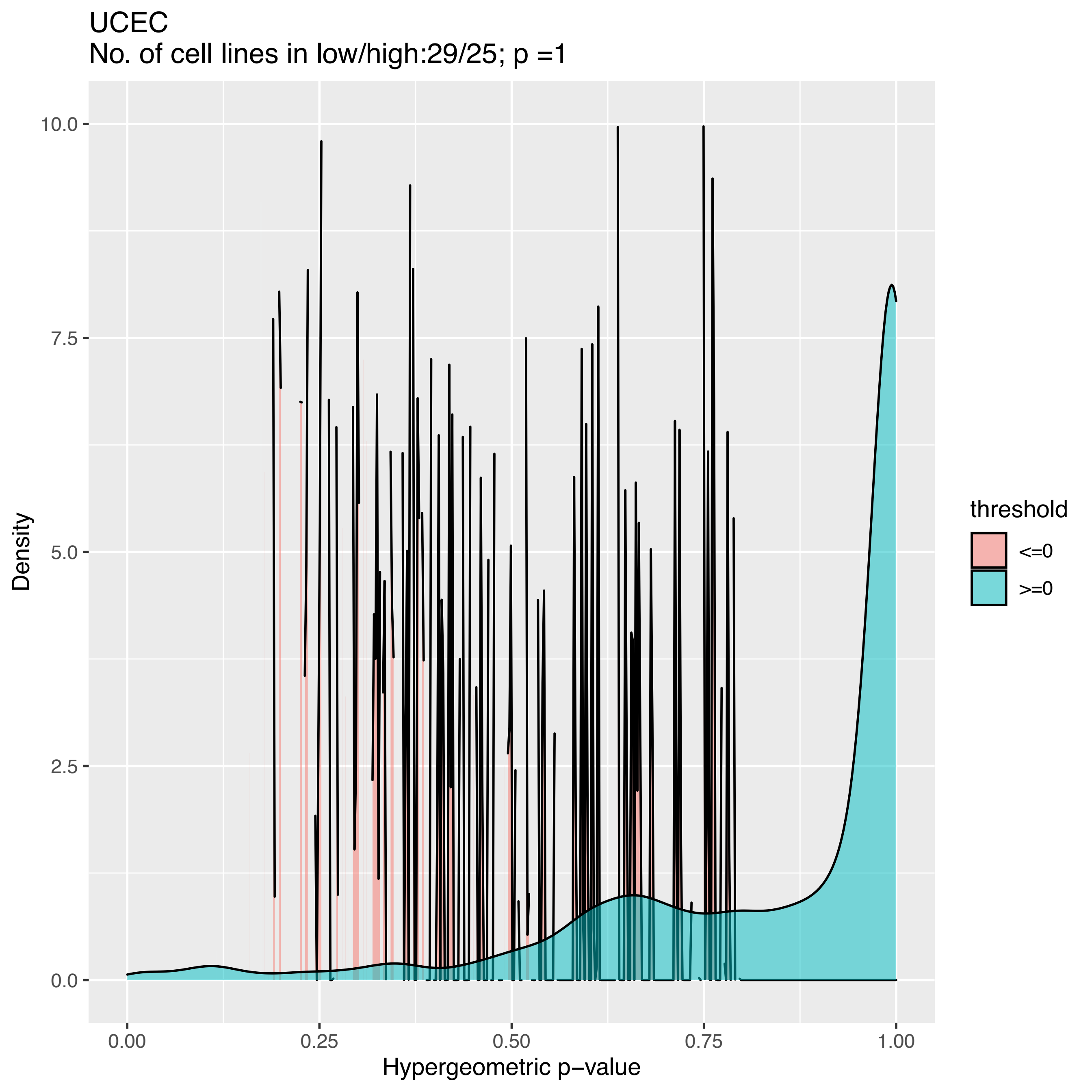
UCEC
No. of cell lines in low/high:29/25; p =1
Density
10.0
7.5
5.0
2.5
0.0
0.00
0.25
0.50
0.75
1.00
Hypergeometric p−value
threshold
<=0
>=0

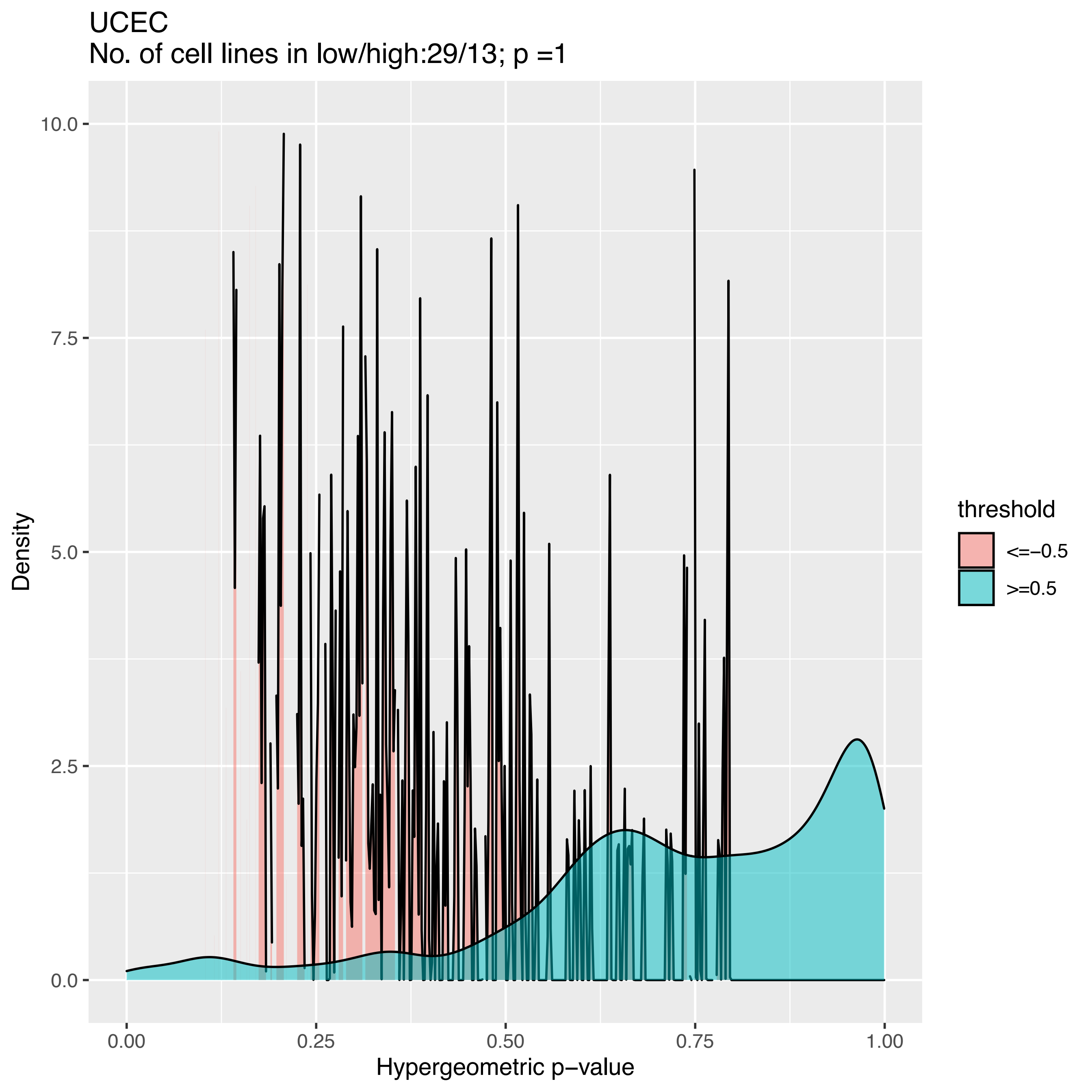

UCEC
No. of cell lines in low/high:29/13; p =1
Density
10.0
7.5
5.0
2.5
0.0
0.00
0.25
0.50
0.75
1.00
Hypergeometric p−value
threshold
<=−0.5
>=0.5

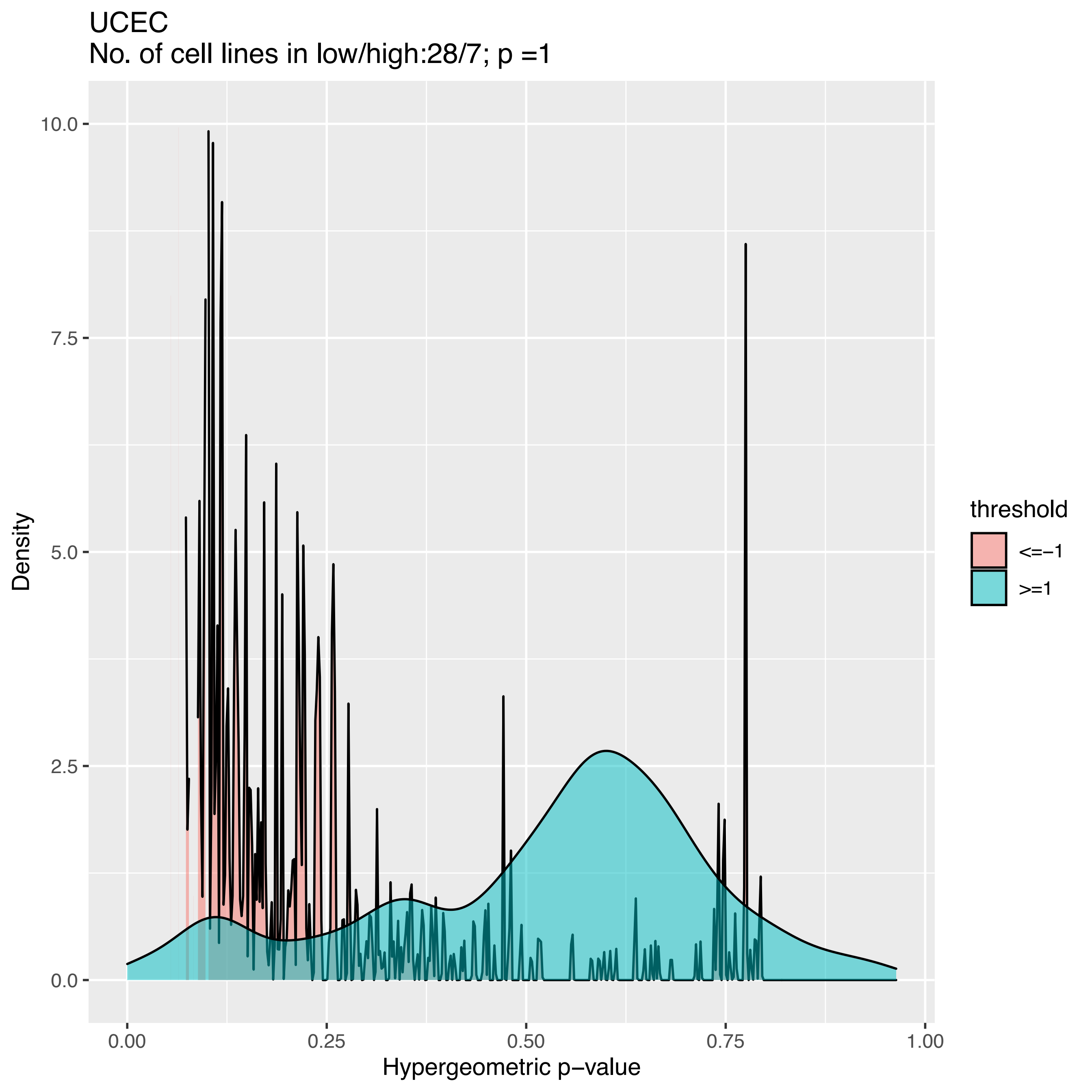

UCEC
No. of cell lines in low/high:28/7; p =1
Density
10.0
7.5
5.0
2.5
0.0
0.00
0.25
0.50
0.75
1.00
Hypergeometric p−value
threshold
<=−1
>=1

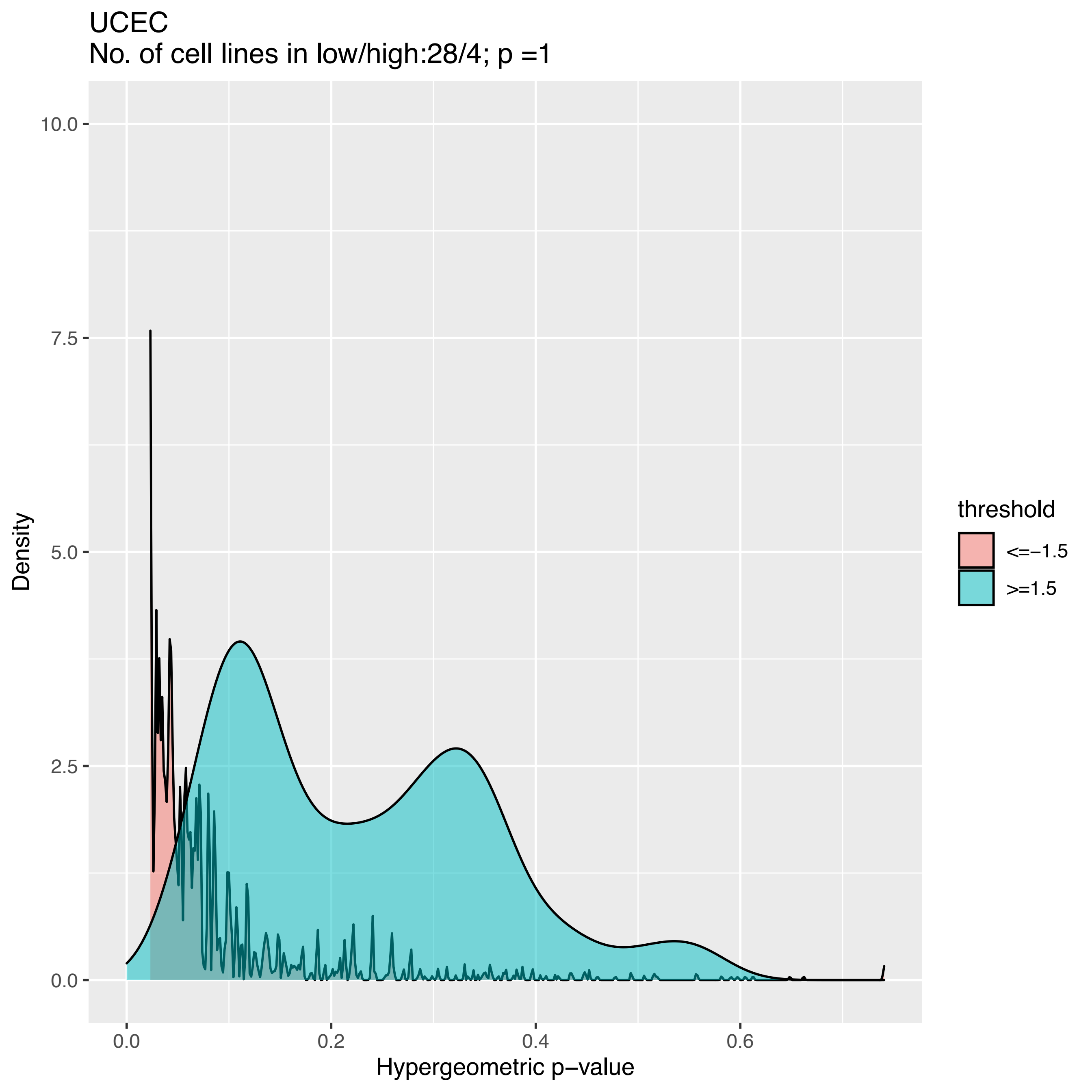
UCEC
No. of cell lines in low/high:28/4; p =1
Density
10.0
7.5
5.0
2.5
0.0
0.0
0.2
0.4
0.6
Hypergeometric p−value
threshold
<=−1.5
>=1.5

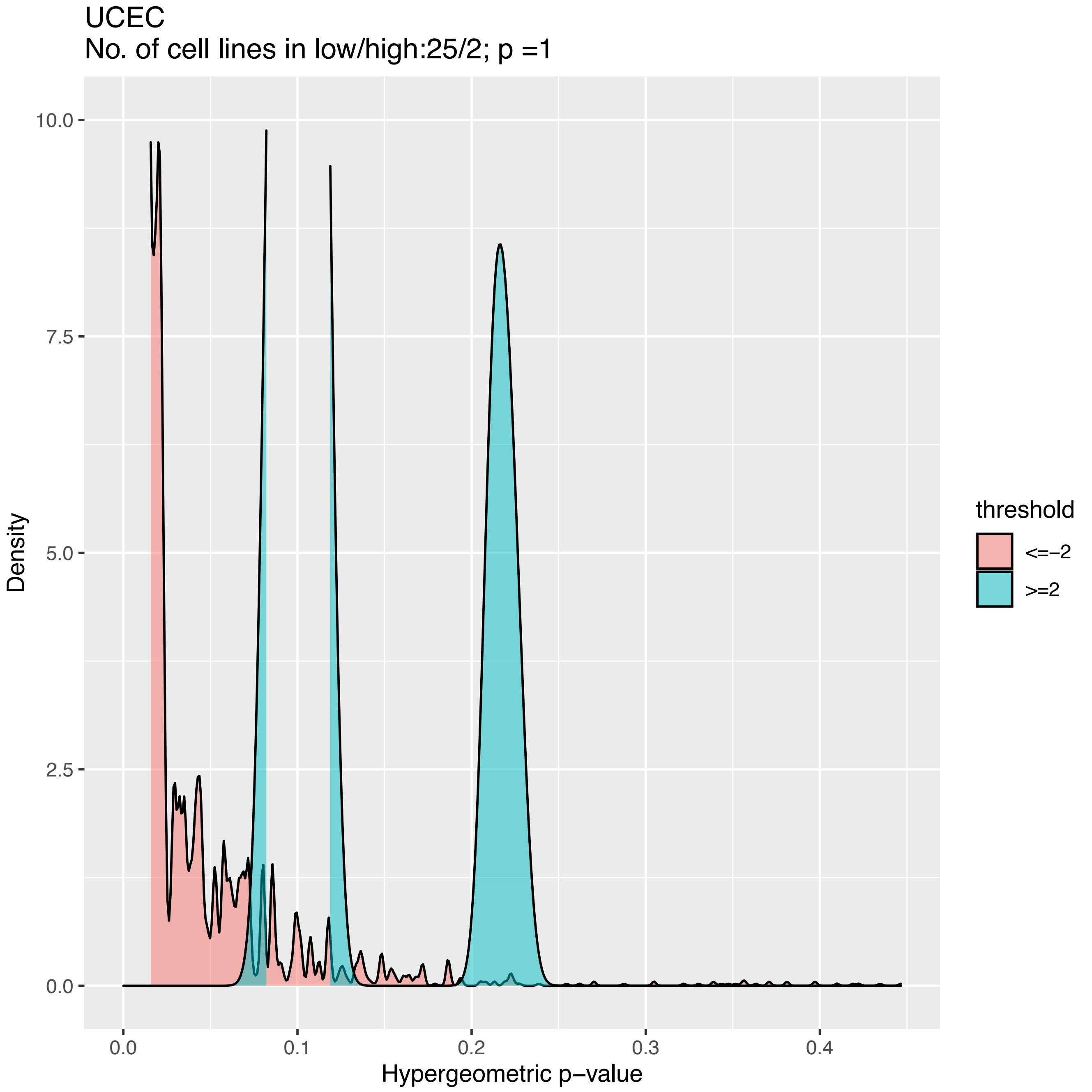
UCEC
No. of cell lines in low/high:25/2; p =1
Density
Hypergeometric p−value
threshold
<=−2
>=2
0.0
2.5
5.0
7.5
10.0
0.0
0.1
0.2
0.3
0.4
