## Supplemental Figure S3 for "Finding the best cell lines across pan-cancer to use in pre-clinical research as a proxy for patient tumor samples considering immune cells, multi-omics, and cancer pathways"

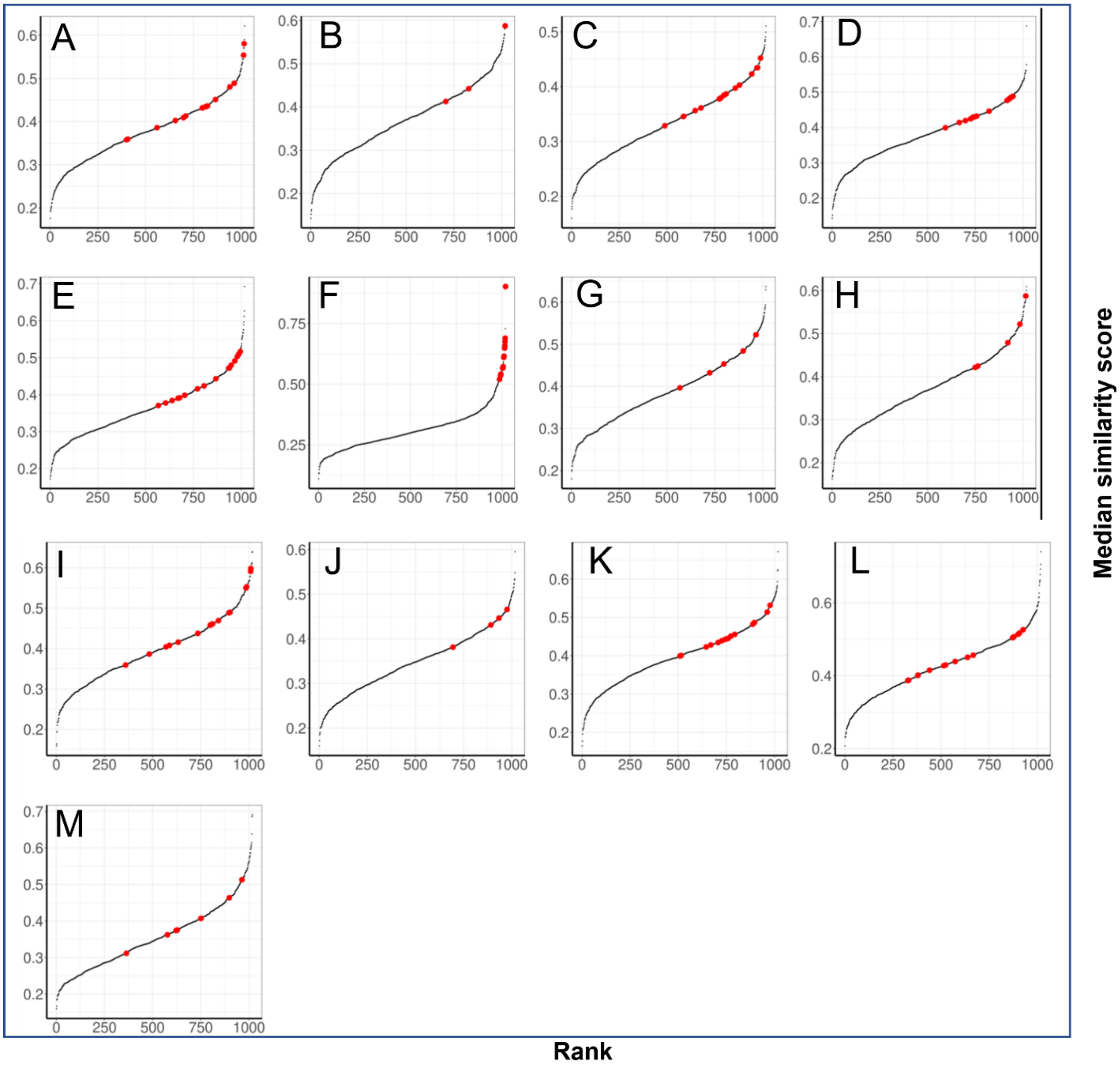


**The rank of all 1,018 CCLE cell lines based on median similarity scores with all samples in the corresponding cancer type computed by CTDPathSim2.0 in thirteen cancer types.** The rank of all cancer cell lines in A) BLCA, B) CESC, C) ESCA, D) HNSC, E) KIRC, F) LAML, G) LGG, H) MESO, I) PAAD, J) PRAD, K) SKCM, L) STAD and M) THCA. Each dot is a CCLE cell line and the few representatives of highly-ranked tissue-specific cell lines are marked by red.
