## Supplemental File S1 for "Finding the best cell lines across pan-cancer to use in pre-clinical research as a proxy for patient tumor samples considering immune cells, multi-omics, and cancer pathways"

**Explanation for the chosen thresholds for drug response concordance test.**

We examined drug response concordance using computed similarity scores in high and low similarities, using the top 20 and bottom 20 percentiles of the scores, respectively. We discovered that these thresholds were better at depicting conditional density plots of drug response matching and mismatch than other thresholds. Here, we present a comparison with different choices of thresholds (Top 30 & bottom 30; top 20 & bottom 20 and top 10 & bottom 10) in BRCA for the five different methods. We label a drug as "concordant" if the matching percentage of the response increases as the similarity score increases; we label it as "discordant" if the matching percentage of the response decreases as the similarity score increases; and all other trends are labeled as "undecided."

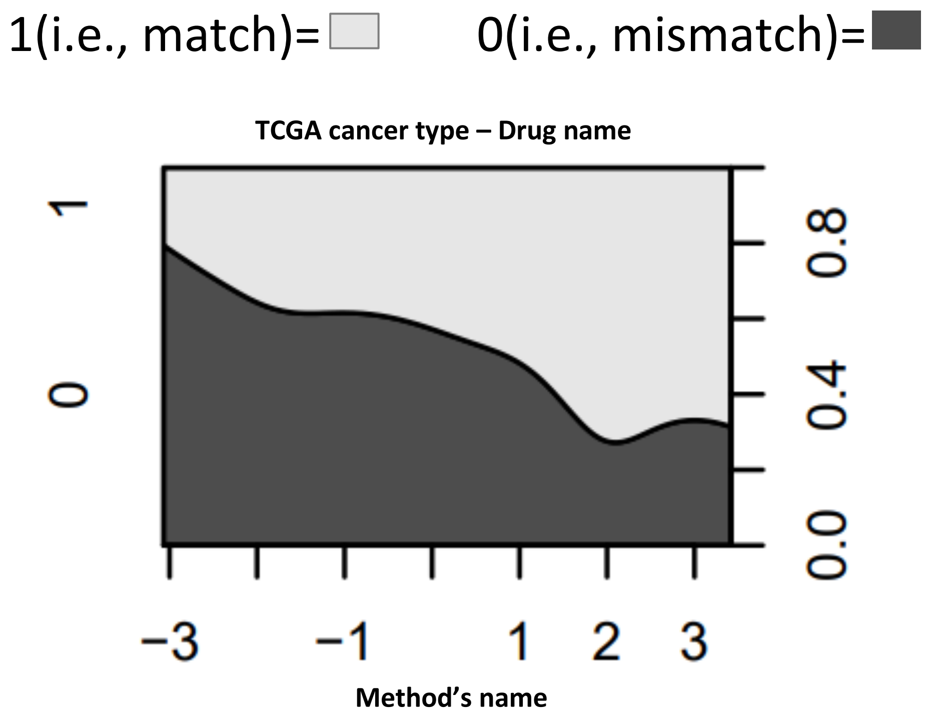


**Drug response concordance with sample-cell line similarity scores.** Conditional density plots of match and mismatch rate of drug response between TCGA samples and CCLE cell lines with computed z-normalized similarity scores by five different methods.

**Conditional density plots in BRCA for five different drugs**

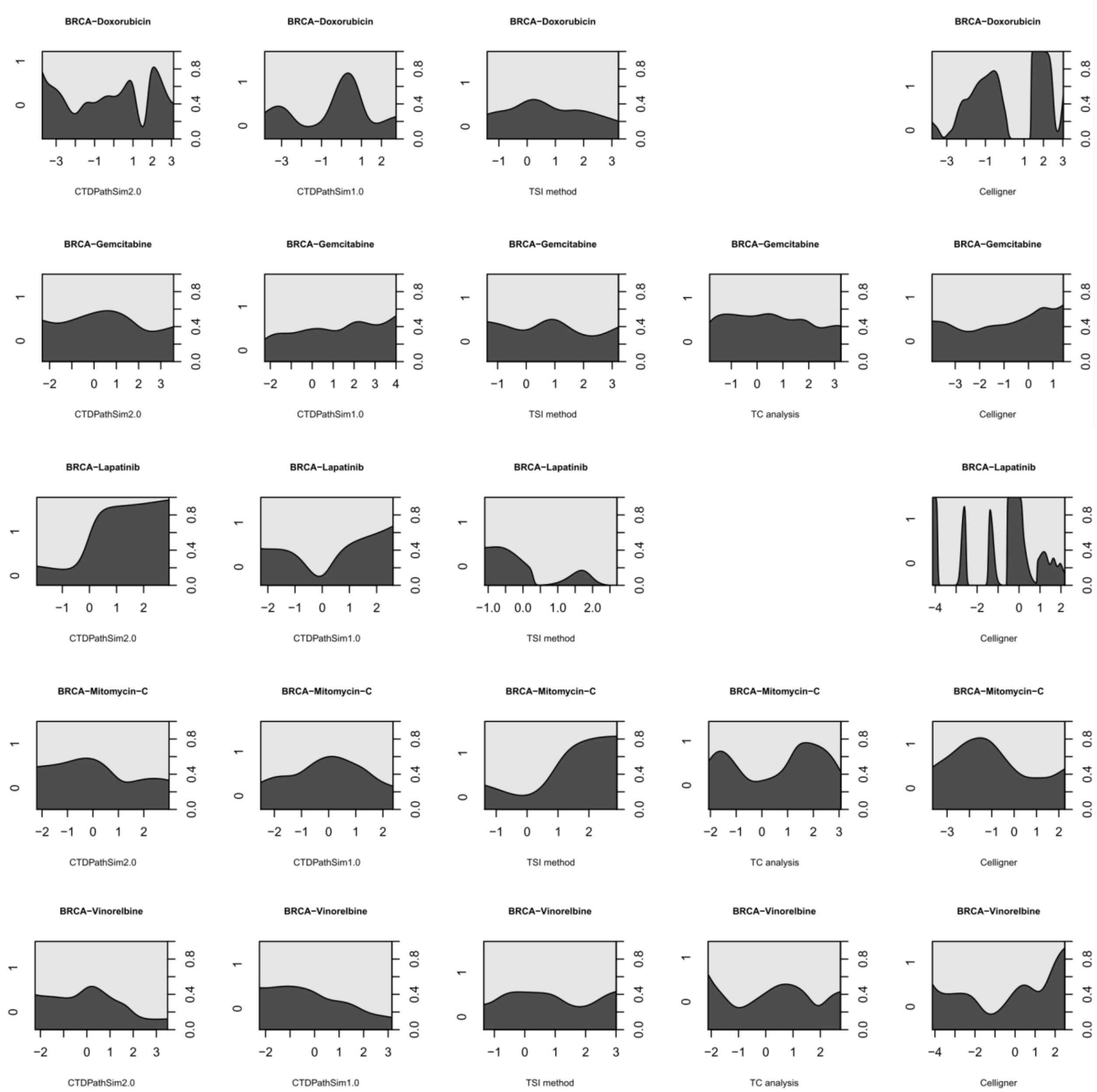


We created the three tables below using this parameters and varying top and bottom percentiles. There are labels for "concordant (i.e., c), "discordant (i.e., d)," and "undecided (i.e., u)" in each table, inside the parentheses, for each drug. The letter "Y" indicates that the label appears, while the letter "N" indicates that the label does not appear. In the case of missing conditional density plots for that drug due to filtering criteria supplied in the main manuscript's "Methods" section, NA stands for "not applicable." It appears in "bold" if the concordance pattern fits the visual assessment of the plots. The number of cells in bold is largest in the top 20 and bottom 20 threshold (Table 2) between three different thresholds. Hence, we estimated a drug response concordance score in each cancer type (i.e., micro concordance score) for each method to summarize its performance using computed similarity scores in high and low similarities, using the top 20 and bottom 20 percentiles of the scores, respectively.

**Table 1: Summary of conditional density plots using top 30 & bottom 30 percentiles in BRCA.**

| Drug name (Concodance label) | CTDPathSim2.0 | CTDPathSim1.0 | TSI method | TC analysis | Celligner |
|---|---|---|---|---|---|
| Doxorubicin (c, d, u) | **Y, N, N** | **Y,N,N** | **N,Y,N** | NA | **N,Y,N** |
| Gemcitabine (c, d, u) | Y, N, N | **N,Y,N** | N,N,Y | Y,N,N | **N,Y,N** |
| Lapatinib (c, d, u) | **N,Y,N** | **N,Y,N** | **Y, N, N** | NA | Y, N, N |
| Mitomycin-C (c, d, u) | N,Y,N | N,Y,N | **N,Y,N** | N,N,Y | **N,Y,N** |
| Vinorelbine (c, d, u) | **Y, N, N** | **Y, N, N** | Y, N, N | N,N,Y | **N,Y,N** |

**Table 2: Summary of conditional density plots using top 20 & bottom 20 percentiles in BRCA.**

| Drug name (Concodance label) | CTDPathSim2.0 | CTDPathSim1.0 | TSI method | TC analysis | Celligner |
|---|---|---|---|---|---|
| Doxorubicin (c, d, u) | **Y, N, N** | **Y,N,N** | **N,Y,N** | NA | **N,Y,N** |
| Gemcitabine (c, d, u) | **N,Y,N** | **N,Y,N** | **N,Y,N** | **N,Y,N** | **N,Y,N** |
| Lapatinib (c, d, u) | **N,Y,N** | **N,Y,N** | **Y, N, N** | NA | Y, N, N |
| Mitomycin-C (c, d, u) | **Y, N, N** | **N,N,Y** | **N,Y,N** | **N,Y,N** | **N,Y,N** |
| Vinorelbine (c, d, u) | **Y, N, N** | **Y, N, N** | Y, N, N | **Y, N, N** | **N,Y,N** |

**Table 3: Summary of conditional density plots using top 10 & bottom 10 percentiles in BRCA.**

| Drug name (Concodance label) | CTDPathSim2.0 | CTDPathSim1.0 | TSI method | TC analysis | Celligner |
|---|---|---|---|---|---|
| Doxorubicin (c, d, u) | **Y, N, N** | N,Y,N | **N,Y,N** | NA | **N,Y,N** |
| Gemcitabine (c, d, u) | **N,Y,N** | **N,Y,N** | **N,Y,N** | **N,Y,N** | **N,Y,N** |
| Lapatinib (c, d, u) | **N,Y,N** | **N,Y,N** | **Y, N, N** | NA | Y, N, N |
| Mitomycin-C (c, d, u) | **Y, N, N** | **N,N,Y** | Y, N, N | **N,Y,N** | **N,Y,N** |
| Vinorelbine (c, d, u) | **Y, N, N** | **Y, N, N** | Y, N, N | **Y, N, N** | **N,Y,N** |

**The explanation of the drug response mapping rule for TCGA patients.**

To compare the drug responses of patient tumor sample-cell line pairs, we compiled our ground truth drug response data for the patients from the TCGA database using Overall Survival (OS) and Progression-Free Survival (PFI) as clinical endpoints. These survival endpoints were considered representative of the drug response status of the patients. In OS, the patients who were dead from any cause were considered dead, otherwise censored. In PFI, the patients having new tumor event whether it was a progression of the disease, local recurrence, distant metastasis, new primary tumor event, or died with cancer without new tumor event, including cases with a new tumor event whose type is N/A were considered as the event occurred and all other patients were censored. We considered survival status 0 (i.e., censored), representative of sensitive drug response whereas survival status 1 (i.e., dead/event occurred), representative of resistant drug response as we mapped the patients' survival status to drug response. In order to determine if PFI or OS status can be used as a drug response metric, we looked at the time intervals between drug applications. The following five rules were used to map the patients' survival status to drug response.

1. **If PFI status = 0 and PFI.time = OS.time then use Drug response = OS status**
   In this case, if there is no progression of the disease but the time interval for OS and PFI are the same, then use OS status as drug response. Since, in this case if the patient has OS = 1 (dead) that will mean that the patient was resistant to the drug and if OS = 0, then that will mean that the patient is alive and there is no progression of the disease implying the sensitive drug response.

2. **If PFI status = 1 and PFI.time = OS.time then use Drug response = PFI status**
   In this case, if there is progression of the disease but the time interval for OS and PFI are the same, then use PFI status as drug response since patient may alive according to OS, but there is disease progression implying resistant drug response.

3. **If drug application end days ≤ PFI days and PFI status = 1 with PFI. time < OS.time then use Drug response = PFI status**
   In this case if the drug is applied within PFI interval days and there is disease progression and the time interval for PFI is less than OS, then use PFI status as drug response since patient may alive according to OS, but there is disease progression implying resistant drug response.

4. **If drug application end days ≤ PFI days and PFI status = 0 with PFI. time < OS.time then use Drug response as OS status**
   In this case if the drug is applied within PFI interval days and there is no disease progression and the time interval for PFI is less than OS, then use PFI status as drug response since patient may alive according to OS, but there is disease progression implying resistant drug response.

5. **If drug application start days ≥ PFI days then use Drug response = OS status**
   In this case if the drug is applied after PFI interval days then use drug response as OS status since progression status of the disease is not related with drug in this case.
