## Supplementary material for "Finding the best cell lines across pan-cancer to use in pre-clinical research as a proxy for patient tumor samples considering immune cells, multi-omics, and cancer pathways": CTDPathSim2.0 R Vignette

### CTDPathSim2.0 Vignette

**By Banabithi Bose**

**4/10/2022**

#### Contents

1 Introduction
2 Example Data Sets
2.1 Methylation.Sample
2.2 Reference.Methylation.Markers
2.3 Reference.Methylation.CellTypes
2.4 Annotation27K
2.5 Annotation450K
2.6 Expression.Sample
2.7 Expression.CellLine
2.8 CancerGeneList
2.9 GeneCentric.DNAmethylation.Sample
2.10 GeneCentric.DNAmethylation.CellLine
2.11 CNV.CellLine
3 CTDPathsim2.0 Functions
3.1 Step 1: Computing sample-specific deconvoluted methylation profile.
3.1.1 Function1: RunDeconvMethylFun
3.1.2 Function2: PlotDeconvMethylFun
3.1.3 Function3: ProbeToGeneFun
3.1.4 Function4: SampleMethylFun
3.2 Step 2: Computing sample-specific deconvoluted expression profile.
3.2.1 Function1: RunDeconvExprFun
3.2.2 Function2: SampleExprFun
3.3 Step 3: Computing sample and cell line-specific differentially expressed (DE) genes and enriched biological pathways.
3.3.1 Function1: GetSampleDEFun
3.3.2 Function2: GetCellLineDEFun
3.3.3 Function3: GetPathFun
3.4 Step 4: Computing sample and cell line-specific differentially methylated (DM) and differentially aberrated (DA) genes.
3.4.1 Function1: GetSampleDMFun
3.4.2 Function2: GetCellLineDMFun
3.4.3 Function3: GetCellLineDAFun
3.5 Step 5: Computing sample-cell line pathway activity-based similarity score.
3.5.1 Function1: FindSimFun

#### 1 Introduction

CTDPathSim2.0 is a computational tool, which utilizes a pathway activity-based approach to compute similarity scores between primary tumor samples and cell lines. CTDPathSim2.0 integrates DNA methylation, gene expression and copy number variation datasets. The package is able to run without copy number variation to produce the results based on previous version of the tool, CTDPathSim1.0. CTDPathSim2.0 has five main computational steps:

Step 1:Computing sample-specific deconvoluted DNA methylation profile

Step 2:Computing sample-specific deconvoluted expression profile

Step 3:Computing sample and cell line-specific differentially expressed (DE) genes and enriched biological pathways

Step 4:Computing sample and cell line-specific differentially methylated (DM) and differentially aberrated (DA) genes

Step 5:Computing sample-cell line pathway activity-based similarity score

```
library(CTDPathSim2.0)
library(printr)
## Warning: package 'printr' was built under R version 4.1
.3
## Registered S3 method overwritten by 'printr':
##   method                 from
##   knit_print.data.frame rmarkdown
data("CTDPathSim2")
```

### 2 Example Data Sets

#### 2.1 Methylation.Sample

```
knitr::kable(Methylation.Sample[1:5,1:3], digits = 2, capt
ion = 'Probe-centric DNA methylation values of the tumor s
amples.')
```

Table 1: **Probe-centric DNA methylation values of the tumor samples.**

| | TCGA-OU-A5PI-01A | TCGA-OR-A5K8-01A | TCGA-OR-A5JX-01A |
|---|---|---|---|
| cg00323915 | 0.93 | 0.55 | 0.32 |
| cg21830221 | 0.87 | 0.90 | 0.63 |
| cg11348106 | 0.91 | 0.55 | 0.77 |
| cg12232731 | 0.82 | 0.30 | 0.55 |
| cg07218880 | 0.16 | 0.13 | 0.07 |

#### 2.2 Reference.Methylation.Markers

```
knitr::kable(Reference.Methylation.Markers[1:5], caption =
'The marker loci (probes) of reference cell types.')
```

Table 2: **The marker loci (probes) of reference cell types.**

| x |
|---|
| cg00323915 |
| cg21830221 |
| cg11348106 |
| cg12232731 |
| cg07218880 |

#### 2.3 Reference.Methylation.CellTypes

```
knitr::kable(Reference.Methylation.CellTypes[1:3,1:3], cap
tion = 'R dataframe for the DNA methylation values for the
reference cell types and the probes.')
```

Table 3: **R dataframe for the DNA methylation values for the reference cell types and the probes.**

| | Neu | NK | Neu_1 |
|---|---|---|---|
| cg00323915 | 0.8715876 | 0.2367720 | 0.8266145 |
| cg21830221 | 0.8344109 | 0.0714415 | 0.8273859 |
| cg11348106 | 0.8878073 | 0.4177928 | 0.8717599 |

#### 2.4 Annotation27K

```
knitr::kable(Annotation27K[1:3,], caption = 'An R dataframe object with probe, symbol and synonym columns of 27K DNA methylation data annotation.')
```

Table 4: **An R dataframe object with probe, symbol and synonym columns of 27K DNA methylation data annotation.**

| ID | Symbol | Synonym |
|---|---|---|
| cg00000292 | ATP2A1 | ATP2A; SERCA1; |
| cg00002426 | SLMAP | SLAP; KIAA1601; |
| cg00003994 | MEOX2 | GAX; MOX2; |

#### 2.5 Annotation450K

```
knitr::kable(Annotation450K[c(6,7,10),], caption = 'An R dataframe object with probe ID, UCSC_REFGENE_NAME and UCSC_REFGENE_GROUP columns 450K DNA methylation data annotation.')
```

Table 5: **An R dataframe object with probe ID, UCSC_REFGENE_NAME and UCSC_REFGENE_GROUP columns 450K DNA methylation data annotation.**

| | ID | UCSC_REFGENE_NAME | UCSC_REFGENE_GROUP |
|---|---|---|---|
| 6 | cg00619207 | DENND2D | TSS200 |
| 7 | cg00705730 | NCK2;NCK2 | 5UTR;5UTR |
| 10 | cg00921266 | HOXA3;HOXA3;HOXA3 | 5UTR;5UTR;TSS200 |

#### 2.6 Expression.Sample

```
knitr::kable(Expression.Sample[1:5,1:3], caption = 'An R dataframe for the gene-centric gene expression (RNASeq) values of the tumor samples.')
```

Table 6: **An R dataframe for the gene-centric gene expression (RNASeq) values of the tumor samples.**

| | TCGA-OR-A5JT-01A | TCGA-OR-A5K6-01A | TCGA-OR-A5K3-01A |
|---|---|---|---|
| TSPAN6 | 10.0802526 | 13.3543602 | 10.9596161 |

Table 6: **An R dataframe for the gene-centric gene expression (RNASeq) values of the tumor samples.**

| | TCGA-OR-A5JT-01A | TCGA-OR-A5K6-01A | TCGA-OR-A5K3-01A |
|---|---|---|---|
| TNMD | 0.2315967 | 0.0536225 | 0.0652658 |
| DPM1 | 40.2218416 | 55.3076696 | 30.9633104 |
| SCYL3 | 1.3136879 | 2.2388920 | 0.5801193 |
| C1orf112 | 0.6405102 | 0.4268140 | 0.2524077 |

#### 2.7 Expression.CellLine

```
knitr::kable(Expression.CellLine[1:5,1:3], caption = 'An R
dataframe for the gene-centric gene expression (RNASeq) va
lues of the cell lines.')
```

Table 7: **An R dataframe for the gene-centric gene expression (RNASeq) values of the cell lines.**

| | LOUNH91_LUNG | T98G_CENTRAL_NERVOUS_SYSTEM | IPC298_SKIN |
|---|---|---|---|
| TSPAN6 | 10.0802526 | 13.3543602 | 10.9596161 |
| TNMD | 0.2315967 | 0.0536225 | 0.0652658 |
| DPM1 | 40.2218416 | 55.3076696 | 30.9633104 |
| SCYL3 | 1.3136879 | 2.2388920 | 0.5801193 |
| C1orf112 | 0.6405102 | 0.4268140 | 0.2524077 |

#### 2.8 CancerGeneList

```
knitr::kable(CancerGeneList[1:5,], caption = 'An R datafra
me object with a column of frequently mutated cancer drive
r genes.')
```

Table 8: **An R dataframe object with a column of frequently mutated cancer driver genes.**

| x |
|---|
| A1CF |
| ABI1 |
| ABL1 |
| ABL2 |
| ACKR3 |

#### 2.9 GeneCentric.DNAmethylation.Sample

```
knitr::kable(GeneCentric.DNAmethylation.Sample[1:5,1:3], c
aption = 'An R dataframe with gene-centric DNA methylation
beta values of bulk tumor samples. Rows are the genes, col
umns are the tumor samples.')
```

Table 9: **An R dataframe with gene-centric DNA methylation beta values of bulk tumor samples**
Rows are the genes, columns are the tumor samples.

| | TCGA-OU-A5PI-01A | TCGA-OR-A5K8-01A | TCGA-OR-A5JX-01A |
|---|---|---|---|
| A1BG | 0.5810136 | 0.2928524 | 0.5154303 |
| A1CF | 0.7602958 | 0.5895629 | 0.7548407 |
| A2BP1 | 0.6226549 | 0.5788986 | 0.3892227 |
| A2LD1 | 0.9217585 | 0.9184202 | 0.9097963 |
| A2M | 0.7231098 | 0.5392000 | 0.5754296 |

#### 2.10 GeneCentric.DNAmethylation.CellLine

```
knitr::kable(GeneCentric.DNAmethylation.CellLine[1:5,1:3],
caption = 'An R dataframe with gene-centric DNA methylatio
n beta values of cell lines. Rows are the genes, columns a
re the cell lines.')
```

Table 10: **An R dataframe with gene-centric DNA methylation beta values of cell lines**
Rows are the genes, columns are the cell lines.

| | LOUNH91_LUNG | T98G_CENTRAL_NERVOUS_SYSTEM | IPC298_SKIN |
|---|---|---|---|
| A1BG | 0.5810136 | 0.2928524 | 0.5154303 |
| A1CF | 0.7602958 | 0.5895629 | 0.7548407 |
| A2BP1 | 0.6226549 | 0.5788986 | 0.3892227 |
| A2LD1 | 0.9217585 | 0.9184202 | 0.9097963 |
| A2M | 0.7231098 | 0.5392000 | 0.5754296 |

#### 2.11 CNV.CellLine

```
knitr::kable(CNV.CellLine[1:5,1:3], caption = 'An R datafr
ame with gene-centric copy number values of cell lines. Ro
ws are the genes, columns are the cell lines.')
```

Table 11: **An R dataframe with gene-centric copy number values of cell lines**
Rows are the genes, columns are the cell lines.

| | LOUNH91_LUNG | T98G_CENTRAL_NERVOUS_SYSTEM | IPC298_SKIN |
|---|---|---|---|
| A1BG | 0.0259 | 0.1514 | 0.1511 |

Table 11: **An R dataframe with gene-centric copy number values of cell lines**
Rows are the genes, columns are the cell lines.

| | LOUNH91_LUNG | T98G_CENTRAL_NERVOUS_SYSTEM | IPC298_SKIN |
|---|---|---|---|
| NAT2 | 0.0325 | 0.0742 | 0.2180 |
| ADA | 0.7455 | 0.5280 | 0.1834 |
| CDH2 | 0.0136 | 0.4032 | 0.5538 |
| AKT3 | 0.0488 | 0.1349 | 0.1940 |

### 3 CTDPathsim2.0 Functions

#### 3.1 Step 1: Computing sample-specific deconvoluted methylation profile.

##### 3.1.1 Function1: RunDeconvMethylFun

```
?RunDeconvMethylFun
```

RunDeconvMethylFun R Documentation

#### Deconvolution of DNA methylation of bulk tumor samples

##### Description

To capture the accurate DNA methylation signal of tumor samples, this function utilizes a deconvolution-based algorithm (V. Onuchic et al., Cell Rep, 2016) to infer the samples deconvoluted DNA methylation profile with proportions of different cell types in the samples bulk tumor tissue. The deconvolution algorithm requires a reference DNA methylation profile of different cell types. Hence, we compiled a list of reference methylation profiles with eight different cell types, namely, B cell, natural killer, CD4T, CD8T, monocytes, adipocytes (AC), cortical neurons (CN), and vascular endothelial (VE) cells with replicates.

##### Usage

```
RunDeconvMethylFun(Methylation.Sample, Reference.Methylati
on.Probes)
```

##### Arguments

| | |
|---|---|
| `Methylation.Sample` | A R dataframe for the probe-centric DNA methylation values of the tumor samples. |
| `Reference.Methylation.Probes` | A vector of the marker loci (probes) based on comparisons of each class of reference against all other samples using t-test. |

##### Details

Created By: Banabithi Bose| Date Created: 4/10/2022 | Stage 1 | Function 1 |

##### Value

A list of seven items: Estimated DNA methylation profile of constituent cell types, estimated proportions of constituent cell types, iteration number, explained variance, residual sum of squares, Akaike information criterion (AIC), and the residual sum of squares per iteration.

```
# Running the function
```

```
stage1_result_ct <-
  RunDeconvMethylFun(Methylation.Sample,Reference.Methylat
ion.Markers)
# Outputs
knitr::kable(stage1_result_ct$methylation[1:5,], caption =
'Estimated DNA methylation profile of constituent cell typ
es.')
```

Table 12: **Estimated DNA methylation profile of constituent cell types.**

| | | | |
|---|---|---|---|
| cg00323915 | 0.1606656 | 0.5224483 | 0.1829575 |
| cg21830221 | 0.8006104 | 0.9129289 | 0.6576261 |
| cg11348106 | 0.6967357 | 0.8727030 | 0.7750149 |
| cg12232731 | 0.5240273 | 0.6487638 | 0.5502076 |
| cg07218880 | 0.0751444 | 0.5104532 | 0.3673889 |

```
knitr::kable(stage1_result_ct$proportions[1:5,], caption =
'Estimated proportions of constituent cell types.')
```

Table 12: **Estimated proportions of constituent cell types.**

| | | | |
|---|---|---|---|
| TCGA-OU-A5PI-01A | 0.31 | 0.69 | 0.00 |
| TCGA-OR-A5K8-01A | 0.55 | 0.45 | 0.00 |
| TCGA-OR-A5JX-01A | 0.86 | 0.07 | 0.07 |
| TCGA-OR-A5LT-01A | 0.58 | 0.19 | 0.23 |
| TCGA-PK-A5H9-01A | 0.73 | 0.07 | 0.20 |

##### 3.1.2 Function2: PlotDeconvMethylFun

```
?PlotDeconvMethylFun
```

PlotDeconvMethylFun R Documentation

#### Correlation plot to show the correlation between the reference cell types and estimated clusters DNA methylation deconvolution

##### Description

This function creates a correlation plot to show the correlation between the reference cell types used and estimated clusters after DNA methylation deconvolution. Users can use this plot to choose the appropriate labels for the cell types for the deconvoluted DNA methylation profiles.

##### Usage

```
PlotDeconvMethylFun(
  Methylation.Sample,
  Reference.Methylation.Probes,
```

```
  Reference.Methylation.CellTypes,
  stage1_result_ct,
  Reference.CellTypes.Names
)
```

#### Arguments

| | |
|---|---|
| `Methylation.Sample` | A R dataframe for the DNA methylation values for the samples and the probes. |
| `Reference.Methylation.Probes` | A vector of the marker loci (probes) based on comparisons of each class of reference against all other samples using t-test. |
| `Reference.Methylation.CellTypes` | A R dataframe for the DNA methylation values for the reference cell types and the probes. |
| `stage1_result_ct` | The output from RunDeconvMethylFun() |
| `Reference.CellTypes.Names` | A vector of the names of the reference cell types. |

#### Details

Created By: Banabithi Bose| Date Created: 4/10/2022 | Stage 1 | Function 2 |

#### Value

Correlation Plot.

```
# Running the function
print(
PlotDeconvMethylFun(Methylation.Sample,Reference.Methylati
on.Markers,Reference.Methylation.CellTypes,stage1_result_c
t,Reference.CellTypes.Names)
)
```

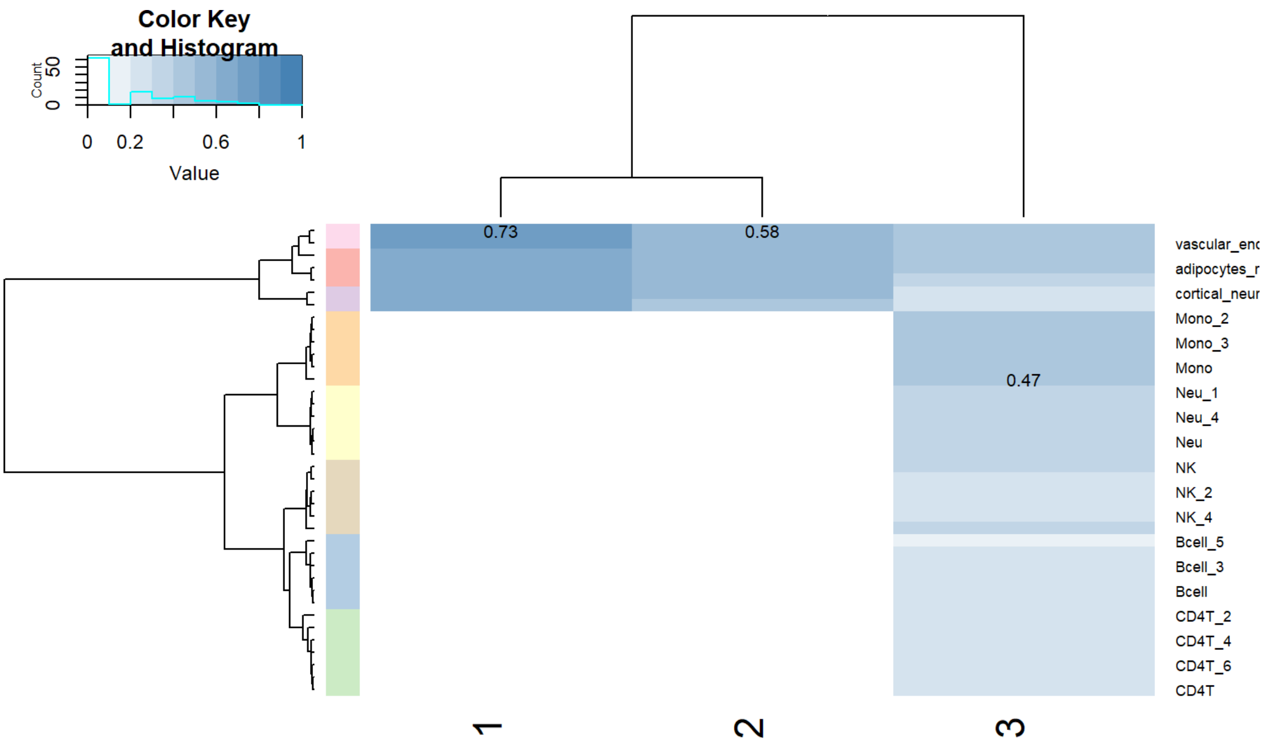


```
## $rowInd
```

```
##  [1]  7 28 30 19 21 11 12 23  4 17 20  8 27  5 26 24  6 
31  2  9  1 14 22 29  3
## [26] 13 10 25 16 18 15 32 33 37 38 36 34 35
##
## $colInd
## [1] 1 2 3
##
## $call
## gplots::heatmap.2(x = cors_deconv_refs_ct, breaks = seq
(0, 1,
##     0.1), col = color_gradient(10), cellnote = best_cor
_labels,
##     notecol = "black", trace = "none", margins = c(5, 5
), RowSideColors = ref_class_colors)
##
## $carpet
##          CD4T    CD4T_5   CD4T_6    CD4T_3    CD4T_4
CD4T_1    CD4T_2
## [1,] 0.0000000 0.0000000 0.000000 0.0000000 0.0000000 0
.0000000 0.0000000
## [2,] 0.0000000 0.0000000 0.000000 0.0000000 0.0000000 0
.0000000 0.0000000
## [3,] 0.2397522 0.2398797 0.239818 0.2356827 0.2318422 0
.2490813 0.2706636
##        Bcell_4     Bcell  Bcell_2   Bcell_3   Bcell_1
Bcell_5      NK_1
## [1,] 0.000000 0.0000000 0.000000 0.0000000 0.0000000 0.
0000000 0.0000000
## [2,] 0.000000 0.0000000 0.000000 0.0000000 0.0000000 0.
0000000 0.0000000
## [3,] 0.214539 0.2134911 0.209599 0.2038733 0.2077725 0.
1947822 0.3078597
##          NK_4      NK_3      NK_2      NK_5        NK
Neu_2       Neu
## [1,] 0.000000 0.0000000 0.0000000 0.0000000 0.0000000 0
.0000000 0.0000000
## [2,] 0.000000 0.0000000 0.0000000 0.0000000 0.0000000 0
.0000000 0.0000000
## [3,] 0.287379 0.2901608 0.2977594 0.2968749 0.3038183 0
.3504076 0.3539911
##          Neu_3    Neu_4     Neu_5     Neu_1       Mono_
1      Mono    Mono_5
## [1,] 0.0000000 0.000000 0.0000000 0.0000000 7.095989e-0
6 0.0000000 0.0000000
## [2,] 0.0000000 0.000000 0.0000000 0.0000000 0.000000e+0
0 0.0000000 0.0000000
## [3,] 0.3543929 0.359657 0.3568648 0.3614733 4.682052e-0
1 0.4500018 0.4544033
##         Mono_3    Mono_4    Mono_2 cortical_neurons_rep
2 cortical_neurons_rep3
## [1,] 0.0000000 0.0000000 0.0000000              0.65948
6             0.6693294
## [2,] 0.0000000 0.0000000 0.0000000              0.48771
1             0.5047249
## [3,] 0.4514434 0.4562636 0.4599182              0.26350
1             0.2799680
##      adipocytes_rep2 adipocytes_rep3 adipocytes_rep1
## [1,]       0.6732391       0.6731806       0.6855552
## [2,]       0.5325767       0.5374538       0.5485259
## [3,]       0.3983218       0.4063843       0.4621058
##      vascular_endothelial_cells_rep1 vascular_endotheli
al_cells_rep2
## [1,]                       0.7220792
0.7299444
## [2,]                       0.5682833
0.5753545
## [3,]                       0.4304132
0.4423359
##
```

```
## $rowDendrogram
## 'dendrogram' with 2 branches and 38 members total, at h
eight 1.196428
##
## $colDendrogram
## 'dendrogram' with 2 branches and 3 members total, at he
ight 2.550236
##
## $breaks
##  [1] 0.0 0.1 0.2 0.3 0.4 0.5 0.6 0.7 0.8 0.9 1.0
##
## $col
##  [1] "#FFFFFF" "#EAF1F6" "#D5E3EE" "#C1D5E6" "#ACC7DD"
"#98B9D5" "#83ABCD"
##  [8] "#6F9DC4" "#5A8FBC" "#4682B4"
##
## $colorTable
##    low high   color
## 1  0.0  0.1 #FFFFFF
## 2  0.1  0.2 #EAF1F6
## 3  0.2  0.3 #D5E3EE
## 4  0.3  0.4 #C1D5E6
## 5  0.4  0.5 #ACC7DD
## 6  0.5  0.6 #98B9D5
## 7  0.6  0.7 #83ABCD
## 8  0.7  0.8 #6F9DC4
## 9  0.8  0.9 #5A8FBC
## 10 0.9  1.0 #4682B4
##
## $layout
## $layout$lmat
##      [,1] [,2] [,3]
## [1,]    5    0    4
## [2,]    3    1    2
##
## $layout$lhei
## [1] 1.5 4.0
##
## $layout$lwid
## [1] 1.5 0.2 4.0
dev.off()
## null device
##           1
```

##### 3.1.3 Function3: ProbeToGeneFun

```
?ProbeToGeneFun
```

ProbeToGeneFun R Documentation

#### Converting DNA methylation data from probes to genes

##### Description

This function converts the DNA methylation data from probe-centric to gene-centric. This function works for both the 450K probe format and 27K probe format.

##### Usage

```
ProbeToGeneFun(
  Methylation.Probe.Annotation,
```

```
  Deconv_methylation,
  Annotation.skim
)
```

#### Arguments

| | |
|---|---|
| `Methylation.Probe.Annotation` | A R dataframe object with 'probe', 'symbol' and 'synonym' columns for 27K or 'ID', 'UCSC_REFGENE_NAME', 'UCSC_REFGENE_GROUP' columns for 450K DNA methylation data according to the convention of the annotation file in "Genome Browser" (https://genome.ucsc.edu/. |
| `Deconv_methylation` | A R dataframe with DNA methylation values for each probe and cell type. |
| `Annotation.skim` | A character string indicating if the user is using either 450K or 27K probes. |

#### Details

Created By: Banabithi Bose| Date Created: 4/10/2022 | Stage 1 | Function 3 |

#### Value

An R dataframe object with deconvoluted DNA methylation values for each gene in cell types. The rows are the genes and the columns are the cell types. This function labels the columns as V1, V2, V3,….etc.

```
# Running the function

Deconv_methylation <- stage1_result_ct$methylation

Deconv_meth_gene <-

  ProbeToGeneFun(Annotation450K,Deconv_methylation,"450K")

## Loading required package: gsubfn

## Loading required package: proto

## Loading required package: RSQLite

# Output

knitr::kable(Deconv_meth_gene[1:5,], caption = 'An R dataf
rame object with deconvoluted DNA methylation values for e
ach gene in cell types. The rows are the genes and the col
umns are the cell types. This function labels the columns
as V1, V2, V3,....etc.')
```

Table 13: **An R dataframe object with deconvoluted DNA methylation values for each gene in cell types**
The rows are the genes and the columns are the cell types. This function labels the columns as V1, V2, V3,….etc.

| | V1 | V2 | V3 |
|---|---|---|---|
| ABI3 | 0.9318309 | 0.9343204 | 0.8692456 |
| ADORA2A | 0.9693054 | 0.9688003 | 0.4833535 |
| ANKRD11 | 0.8464842 | 0.9703333 | 0.3610563 |
| BIN2 | 0.9823428 | 0.9621157 | 0.5049663 |

Table 13: **An R dataframe object with deconvoluted DNA methylation values for each gene in cell types**
The rows are the genes and the columns are the cell types. This function labels the columns as V1, V2, V3,….etc.

| | V1 | V2 | V3 |
|---|---|---|---|
| BLK | 0.5299996 | 0.8206106 | 0.2800671 |

##### 3.1.4 Function4: SampleMethylFun

```
?SampleMethylFun
```

SampleMethylFun R Documentation

#### Sample-specific deconvoluted DNA methylation profile.

##### Description

This function utilizes the estimated proportion of constituent cell types and the estimated gene-centric DNA methylation profile of constituent cell type to compute the sample-specific deconvoluted DNA methylation profile for each tumor sample.

##### Usage

```
SampleMethylFun(Deconv_meth_gene, Deconv_proportions, CTDD
irectory = "~")
```

##### Arguments

| | |
|---|---|
| `Deconv_meth_gene` | A R dataframe with the deconvoluted DNA methylation values for each gene in constituent cell types. |
| `Deconv_proportions` | A R dataframe with the estimated proportions of constituent cell types in samples. |
| `CTDDirectory` | A character string for the file path of the directory for the output to be stored. |

##### Value

R dataframes including the deconvoluted DNA methylation profile for each sample in the specified directory. The rows are the genes and the columns are the cell types.This function labels the columns as V1, V2, V3,….etc.

```
# Running the function
CTDDirectory <- tempdir()
SampleMethylFun(Deconv_meth_gene,Deconv_proportions, CTDDi
rectory)
## NULL
# Output
load("~/Sample_methylation/TCGA-OR-A5J1-01A.Rda")
knitr::kable(methylation[1:5,], caption = 'Deconvoluted DN
A methylation profile of the sample TCGA-OR-A5J1-01A')
```

Table 14: **Deconvoluted DNA methylation profile of the sample TCGA-OR-A5J1-01A**

| | V1 | V2 | V3 |
|---|---|---|---|
| ABI3 | 0.3075042 | 0.0086925 | 0.6166515 |

Table 14: **Deconvoluted DNA methylation profile of the sample TCGA-OR-A5J1-01A**

| | V1 | V2 | V3 |
|---|---|---|---|
| ADORA2A | 0.3198708 | 0.0048335 | 0.6394082 |
| ANKRD11 | 0.2793398 | 0.0036106 | 0.6404200 |
| BIN2 | 0.3241731 | 0.0050497 | 0.6349964 |
| BLK | 0.1748998 | 0.0028007 | 0.5416030 |

#### 3.2 Step 2: Computing sample-specific deconvoluted expression profile.

##### 3.2.1 Function1: RunDeconvExprFun

```
?RunDeconvExprFun
```

RunDeconvExprFun R Documentation

#### Deconvolution of gene expression of bulk tumor samples

##### Description

This function utilizes the estimated cell proportions from Step 1 as a fixed input and computes an average gene expression profiles of constituent cell types through a constrained least-squares using quadratic programming.

##### Usage

```
RunDeconvExprFun(Expression.Sample, Deconv_proportions)
```

##### Arguments

| | |
|---|---|
| `Expression.Sample` | A R dataframe for the gene-centric gene expression (RNASeq) values of the tumor samples. |
| `Deconv_proportions` | A R dataframe with estimated cell proportions in tumor samples from output of the RunDeconvMethylFun() function. |

##### Details

Created By: Banabithi Bose| Date Created: 4/10/2022 | Stage 2 | Function 1 |

##### Value

A R dataframe with gene expression values in constituent cell types. The rows are the genes and the columns are the cell types. This function labels the columns as ,1, ,2,,3,….etc.

```
# Running the function
Deconv_expression<-RunDeconvExprFun(Expression.Sample,Deco
nv_proportions)
# Output
knitr::kable(Deconv_expression[1:5,], caption = 'An R data
frame object with deconvoluted gene expression values for
each gene in cell types. The rows are the genes and the co
lumns are the cell types. This function labels the columns
as V1, V2, V3,....etc.')
```

Table 15: **An R dataframe object with deconvoluted gene expression values for each gene in cell types**
The rows are the genes and the columns are the cell types. This function labels the columns as V1, V2, V3,….etc.

| | | | |
|---|---|---|---|
| TSPAN6 | 10.2170293 | 7.662672 | 7.4049619 |
| TNMD | 0.1067874 | 0.244172 | 0.0321168 |
| DPM1 | 35.6648144 | 25.368440 | 42.1562712 |
| SCYL3 | 1.3917714 | 1.781208 | 1.2404976 |
| C1orf112 | 0.2324402 | 1.249619 | 0.9056265 |

##### 3.2.2 Function2: SampleExprFun

```
?SampleExprFun
```

SampleExprFun R Documentation

#### Sample-specific deconvoluted gene expression profile.

##### Description

This function computes the deconvoluted expression profile for each tumor sample.

##### Usage

```
SampleExprFun(Deconv_expression, Deconv_proportions, CTDDi
rectory = "~")
```

##### Arguments

| | |
|---|---|
| `Deconv_expression` | A R dataframe with the deconvoluted gene expression values for each gene in constituent cell types. |
| `Deconv_proportions` | A R dataframe with the estimated proportions of constituent cell types in samples. |
| `CTDDirectory` | A character string for the file path of the directory for the output to be stored. |

##### Details

Created By: Banabithi Bose| Date Created: 4/10/2022 | Stage 2 | Function 2 |

##### Value

R dataframes including the deconvoluted gene expression profile for each sample in the specified directory. The rows are the genes and the columns are the cell types. This function labels the columns as V1, V2, V3,….etc.

```
# Running the function
SampleExprFun(Deconv_expression,Deconv_proportions, CTDDir
ectory)
## NULL
# Output
load("~/Sample_expression/TCGA-OR-A5J1-01A.Rda")
knitr::kable(expression[1:5,], caption = 'Deconvoluted gen
e expression profile of the sample TCGA-OR-A5J1-01A')
```

Table 16: **Deconvoluted gene expression profile of the sample TCGA-OR-A5J1-01A**

| | | | |
|---|---|---|---|
| TSPAN6 | 3.3716197 | 0.0766267 | 4.8872748 |
| TNMD | 0.0352398 | 0.0024417 | 0.0211971 |
| DPM1 | 11.7693888 | 0.2536844 | 27.8231390 |
| SCYL3 | 0.4592846 | 0.0178121 | 0.8187284 |
| C1orf112 | 0.0767053 | 0.0124962 | 0.5977135 |

#### 3.3 Step 3: Computing sample and cell line-specific differentially expressed (DE) genes and enriched biological pathways.

##### 3.3.1Function1: GetSampleDEFun

```
?GetSampleDEFun
```

GetSampleDEFun R Documentation

#### Sample-specific differentially expressed (DE) genes

##### Description

This function computes the median expression of a gene across all samples and further computed the fold change of that gene to the computed median expression. A gene was considered up if the fold-change ≥ 4 and down if the fold-change ≤ -4.

##### Usage

```
GetSampleDEFun(
  RnaSeq_data,
  parallel = c("TRUE", "FALSE"),
  ncores = 2,
  CTDDirectory = "~"
)
```

##### Arguments

| | |
|---|---|
| `RnaSeq_data` | A R dataframe with gene-centric expression values of bulk tumor samples. Rows are the genes, columns are the tumor samples. |
| `parallel` | A boolean value (‘TRUE’ or ‘FALSE’) indicating if the users want to run this function in multi-core. |
| `ncores` | An integer value for the number of cores. |
| `CTDDirectory` | A character string for the file path of the directory for the output to be stored. |

##### Details

Created By: Banabithi Bose| Date Created: 4/10/2022 | Stage 3 | Function 1 |

##### Value

R dataframe objects each with a single column containing DE genes for each tumor sample. The column is labeled as gene.

```r
# Running the function
CTDDirectory <- tempdir()
GetSampleDEFun(RnaSeq_data=Expression.Sample,parallel= TRU
E,ncores=2, CTDDirectory)
## Warning: package 'pbapply' was built under R version 4.
1.3
## [1] "cluster making started"
## [1] "cluster export started"
## [1] "cluster making done"
## [1] "cluster stopped"
## NULL
# Output
load(paste0(CTDDirectory,"/Patient_DE_genes/DE_Gene/TCGA-O
R-A5J6-01A_pat_de_genes.Rda"))
knitr::kable(pat_de_genes, caption = 'DE genes of the samp
le TCGA-OR-A5J1-01A')
```

Table 17: **DE genes of the sample TCGA-OR-A5J1-01A**

| gene |
|---|
| TNMD |

##### 3.3.2 Function2: GetCellLineDEFun

```r
?GetCellLineDEFun
```

GetCellLineDEFun R Documentation

#### Getting cell line-specific differentially expressed (DE) genes

##### Description

This function computes the median expression of a gene across all cell lines and further computed the fold change of that gene to the computed median expression. A gene was considered up if the fold-change ≥ 4 and down if the fold-change ≤ -4.

##### Usage

```r
GetCellLineDEFun(
  RnaSeq_data,
  parallel = c("TRUE", "FALSE"),
  ncores = 2,
  CTDDirectory = "~"
)
```

##### Arguments

| | |
|---|---|
| `RnaSeq_data` | A R dataframe with gene-centric expression values of cell lines. Rows are the genes, columns are the tumor samples. |
| `parallel` | A boolean value (‘TRUE’ or ‘FALSE’) indicating if the users want to run this function in multi-core. |
| `ncores` | A integer value for the number of cores. |

`CTDDirectory`
A character string for the file path of the directory for the output to be stored.

#### Details

Created By: Banabithi Bose| Date Created: 4/10/2022 | Stage 3 | Function 2 |

#### Value

R dataframe objects each with a single column containing DE genes for each cell line. The column is labeled as gene.

```
# Running the function
GetCellLineDEFun(RnaSeq_data=Expression.CellLine,parallel=
TRUE,ncores=2, CTDDirectory)
## [1] "cluster making started"
## [1] "cluster export started"
## [1] "cluster making done"
## [1] "cluster stopped"
## NULL
# Output
load(paste0(CTDDirectory,"/CellLine_DE_genes/DE_Gene/LK2_L
UNG_cell_de_genes.Rda"))
knitr::kable(cell_de_genes[1:5,,drop=F], caption = 'DE gen
es of the LK2 lung cancer cell line')
```

Table 18: **DE genes of the LK2 lung cancer cell line**

| gene |
|---|
| A1CF |
| A2M |
| A2M-AS1 |
| A2ML1-AS1 |
| A2MP1 |

##### 3.3.3 Function3: GetPathFun

```
?GetPathFun
```

GetPathFun R Documentation

### Enriched biological pathways

#### Description

This function identifies the enriched biological pathways for each sample and cell line utilizing the DE genes. It finds enriched biological pathways listed in the REACTOME database via a pathway enrichment tool Pathfinder. To prioritize cancer-related biological pathways, specifically, it uses the known frequently mutated genes that were considered cancer-driving genes from the DE gene list for this analysis. We obtained the frequently mutated cancer-driving genes from the COSMIC database. GetPathFun() considers the pathways that were significantly enriched with FDR-corrected hypergeometric p-values < 0.05 in this DE cancer gene list.

#### Usage

```
GetPathFun(CancerGeneList, SampleCell_de_genes)
```

#### Arguments

`CancerGeneList` A R dataframe with a column of frequently mutated cancer driver genes.

`SampleCell_de_genes` A R dataframe with a column containing DE genes of a sample or cell line.

#### Details

Created By: Banabithi Bose| Date Created: 4/10/2022 | Stage 3 | Function 3 |

#### Value

An R dataframe object with three columns labeled as 'ID', 'reactome_pathway', and 'p.adjust.pat', respectively.

```
# Running the function
data("CTDPathSim2")
Enriched.pathways.sample<-GetPathFun(CancerGeneList,pat_de
_genes)
# Output
knitr::kable(Enriched.pathways.sample[1:5,], caption = 'Sa
mple(TCGA-OR-A5J1-01A)-specific enriched biological pathwa
ys')
```

Table 19: Sample(TCGA-OR-A5J1-01A)-specific enriched biological pathways

| ID | reactome_pathway | p.adjust.pat |
|---|---|---|
| R-HSA-162582 | Signal Transduction | 0.00e+00 |
| R-HSA-1266738 | Developmental Biology | 0.00e+00 |
| R-HSA-1643685 | Disease | 0.00e+00 |
| R-HSA-212436 | Generic Transcription Pathway | 4.60e-06 |
| R-HSA-73857 | RNA Polymerase II Transcription | 1.28e-05 |

```
knitr::kable(Enriched.pathways.cellLine[1:5,], caption = '
Cell line(LK2)-specific enriched biological pathways')
```

Table 19: Cell line(LK2)-specific enriched biological pathways

| ID | reactome_pathway | p.adjust.pat |
|---|---|---|
| R-HSA-162582 | Signal Transduction | 0 |
| R-HSA-168256 | Immune System | 0 |
| R-HSA-9006934 | Signaling by Receptor Tyrosine Kinases | 0 |

Table 19: **Cell line(LK2)-specific enriched biological pathways**

| ID | reactome_pathway | p.adjust.pat |
|---|---|---|
| R-HSA-1280215 | Cytokine Signaling in Immune system | 0 |
| R-HSA-5663202 | Diseases of signal transduction by growth factor receptors and second messengers | 0 |

#### 3.4 Step 4: Computing sample and cell line-specific differentially methylated (DM) and differentially aberrated (DA) genes.

##### 3.4.1 Function1: GetSampleDMFun

```
?GetSampleDMFun
```

GetSampleDMFun R Documentation

#### Sample-specific differentially methylated (DM) genes

##### Description

This function computes sample-specific DM genes for each sample. It computes M-values from gene-centric beta values and computes the median of gene-centric M-values across all the samples. It considers a gene hypermethylated if the M-value fold-change ≥ 4 and hypomethylated if fold-change ≤ -4.

##### Usage

```
GetSampleDMFun(
  DNAmethylation_data,
  parallel = c("TRUE", "FALSE"),
  ncores = 2,
  CTDDirectory = "~"
)
```

##### Arguments

| | |
|---|---|
| `DNAmethylation_data` | A R dataframe with gene-centric DNA methylation beta values of bulk tumor samples. Rows are the genes, columns are the tumor samples. |
| `parallel` | A boolean value (‘TRUE’ or ‘FALSE’) indicating if the users want to run this function in multi-core. |
| `ncores` | An integer value for the number of cores. |
| `CTDDirectory` | A character string for the file path of the directory for the output to be stored. |

##### Details

Created By: Banabithi Bose| Date Created: 4/10/2022 | Stage 4 | Function 1 |

##### Value

R dataframe objects each with a single column containing DM genes for each tumor sample. The column is labeled as gene.

```
# Running the function
GetSampleDMFun(GeneCentric.DNAmethylation.Sample,parallel=
TRUE,ncores=2, CTDDirectory)
## [1] "cluster making started"
## [1] "cluster export started"
## [1] "cluster making done"
## [1] "cluster stopped"
## NULL
# Output
load(paste0(CTDDirectory,"/Patient_DM_genes/DM_Gene/TCGA-O
R-A5J6-01A_pat_dm_genes.Rda"))
knitr::kable(pat_dm_genes, caption = 'DM genes of the samp
le TCGA-OR-A5J1-01A')
```

Table 20: DM genes of the sample TCGA-OR-A5J1-01A

| gene |
|---|
| A1CF |
| A2M |
| A4GALT |
| ABAT |
| ABCA1 |
| AADACL2 |
| ABCA10 |

##### 3.4.2Function2: GetCellLineDMFun

```
?GetCellLineDMFun
```

GetCellLineDMFun R Documentation

#### Cell line-specific differentially (DM) genes

##### Description

This function computes cell line-specific DM genes for each cell lines It computes M-values from gene-centric beta values and computes the median of gene-centric M-values across all the cell lines It considers a gene hypermethylated if the M-value fold-change ≥ 4 and hypomethylated if fold-change ≤ -4.

##### Usage

```
GetCellLineDMFun(
  DNAmethylation_data,
  parallel = c("TRUE", "FALSE"),
  ncores = 2,
  CTDDirectory = "~"
)
```

##### Arguments

| | |
|---|---|
| `DNAmethylation_data` | A R dataframe with gene-centric DNA methylation beta values of cell lines. Rows are the genes, columns are the cell lines. |
| `parallel` | A boolean value ('TRUE' or 'FALSE') indicating if the users want to run this function in multi-core. |
| `ncores` | A integer value for the number of cores. |
| `CTDDirectory` | A character string for the file path of the directory for the output to be stored. |

##### Details

Created By: Banabithi Bose| Date Created: 4/10/2022 | Stage 4 | Function 2 |

##### Value

R dataframe objects each with a single column containing DM genes for each cell line. The column is labeled as gene.

```
# Running the function
GetCellLineDMFun(GeneCentric.DNAmethylation.CellLine,parallel= TRUE,ncores=2, CTDDirectory)
## [1] "cluster making started"
## [1] "cluster export started"
## [1] "cluster making done"
## [1] "cluster stopped"
## NULL
# Output
load(paste0(CTDDirectory,"/CellLine_DM_genes/DM_Gene/LK2_LUNG_cell_dm_genes.Rda"))
knitr::kable(cell_dm_genes[1:5,], caption = 'DM genes of the LK2 cell line')
```

Table 21: **DM genes of the LK2 cell line**

| gene |
|---|
| ADCY9 |
| ADCYAP1 |
| ADCYAP1R1 |
| ADD3 |
| ADGRE1 |

#### 3.4.3 Function3: GetCellLineDAFun

```
?GetCellLineDAFun
```

GetCellLineDAFun R Documentation

### Cell line-specific differentially aberrated (DA) genes

##### Description

This function selects the highly amplified and deleted genes for each cell line compared with all other cell lines from various cancer types to capture the cancer-specific variation in copy number profiles in computing the similarity score between a patient sample and cell line. It selects the cell line-specific DA genes based on Tukey mean-difference curve based on gene-centric copy number data of cell lines. To determine whether a gene is DA for a specific cell line, it calculates the mean of that gene's copy number values across all cell lines and a difference between the copy number value of that gene in the cell line under consideration and the mean of the copy number values of that gene in the other cell lines. For each cell line, the highly amplified genes are chosen based on mean ≥ 0.5 and difference ≥1, and highly deleted genes are chosen based on mean ≥ 0.5 and difference ≤ -1.

##### Usage

```
GetCellLineDAFun(CNV.CellLine, ncores = 1, CTDDirectory =
"~")
```

##### Arguments

| | |
|---|---|
| `CNV.CellLine` | A R dataframe with gene-centric copy number values of cell lines. Rows are the genes, columns are the cell lines. |
| `ncores` | An integer value for the number of cores. |
| `CTDDirectory` | A character string for the file path of the directory for the output to be stored. |

##### Details

Created By: Banabithi Bose| Date Created: 4/10/2022 | Stage 4 | Function 1 |

##### Value

R dataframe objects each with a single column containing DA genes for each cell line. The column is labeled as gene.

```
# Running the function
GetCellLineDAFun(CNV.CellLine,ncores=2, CTDDirectory)
## [1] "cluster making started"
## NULL
# Output
load(paste0(CTDDirectory,"/CellLine_DA_genes/DA_Gene/LK2_L
UNG_cell_dm_genes.Rda"))
knitr::kable(cell_da_genes[1:2,], caption = 'DA genes of t
he LK2 lung cancer cell line')
```

Table 22: **DA genes of the LK2 lung cancer cell line**

| V1 |
|---|
| SIGLEC14 |
| CDKN2B-AS1 |

Computing sample-specific DA genes: To compute sample-specific DA genes, users need to compute highly amplified and deleted genes utilizing chromosomal copy number profiles of each cancer type cohort as inputs in the GISTIC 2.0 tool in GenePattern web server (https://www.genepattern.org/) using a confidence interval of 0.90. They should select the genes that exceed the high-level GISTIC thresholds for amplification and deletions as 2 and -2, respectively as DA genes.

#### 3.5 Step 5: Computing sample-cell line pathway activity-based similarity score.

##### 3.5.1 Function1: FindSimFun

```
?FindSimFun
```

FindSimFun R Documentation

#### Sample-cell line pathway activity-based similarity score

##### Description

This function computes Spearman rank correlation between each sample-cell line pair using the sample-specific deconvoluted expression, sample-specific deconvoluted DNA methylation, or gene-centric copy number values of samples. All DE, DM, and DA genes that occur in an enriched pathway of a sample or cell line are used to compute the Spearman rank correlation between each sample-cell line pair. Using DM and DE genes, it uses the deconvoluted profiles of samples to compute Spearman rank correlation whereas for DA genes, it uses the gene-centric copy number profile of samples to compute Spearman rank correlation.

##### Usage

```
FindSimFun(
  Patient.D.Genes,
  Patient.Reactome,
  Patient.Expr.Meth.CNV,
  Cell.D.Genes,
  Cell.Reactome,
  Cell.Expr.Meth.CNV
)
```

##### Arguments

| | |
|---|---|
| `Patient.D.Genes` | A vector of DE or DM or DA genes of a tumor sample |
| `Patient.Reactome` | A R dataframe for patient-specific enriched biological pathways with three columns labeled as ID, reactome_pathway, and p.adjust.pat. |
| `Patient.Expr.Meth.CNV` | A R dataframe with sample-specific deconvoluted gene expression or deconvoluted DNA methylation or copy number profiles. |
| `Cell.D.Genes` | A vector of DE or DM or DA genes of a cell line |
| `Cell.Reactome` | A R dataframe for cell line-specific enriched biological pathways with three columns labeled as ID, reactome_pathway, and p.adjust.pat. |
| `Cell.Expr.Meth.CNV` | A R dataframe with cell line-specific gene expression or DNA methylation or copy number profiles. |

##### Details

Created By: Banabithi Bose| Date Created: 4/10/2022 | Stage 5 | Function 1 |

##### Value

A numeric value (Spearman similarity score)

```
# Running the function to compute DNA methylation based Sp
earman similarity score of a patient-cell line pair
```

```
data("CTDPathSim2")

x<-FindSimFun(pat_dm_genes,pat_reactome,Deconv_meth_gene,c
ell_dm_genes,cell_reactome,ccle_methylation)

knitr::kable(x, caption = 'DNA methylation based Spearman
similarity score of a patient-cell line pair')
```

Table 23: DNA methylation based Spearman similarity score of a patient-cell line pair

| Spearman_Similarity_Score |
|---|
| 0.6796275 |

```
# Running the function to compute gene expression-based Sp
earman similarity score of a patient-cell line pair

y<-FindSimFun(pat_de_genes,pat_reactome,Deconv_expression,
cell_de_genes,cell_reactome,ccle_expr)

knitr::kable(y, caption = 'Gene expression-based Spearman
similarity score of a patient-cell line pair')
```

Table 23: Gene expression-based Spearman similarity score of a patient-cell line pair

| Spearman_Similarity_Score |
|---|
| 0.1575758 |

```
# Running the function to compute copy number value based
Spearman similarity score of a patient-cell line pair

z<-FindSimFun(pat_da_genes,pat_reactome,pat_cnv,cell_da_ge
nes,cell_reactome,ccle_cnv)

knitr::kable(z, caption = 'Copy number aberration-based Sp
earman similarity score of a patient-cell line pair')
```

Table 23: Copy number aberration-based Spearman similarity score of a patient-cell line pair

| Spearman_Similarity_Score |
|---|
| 0.0966884 |

For CTDPathSim2.0, after computing the three different scores for all the sample-cell line pairs, the users need to scale the Spearman similarities to the range of 0 to 1 using min-max normalization and should compute an average similarity score taking a mean of expression-based score, DNA methylation-based score, and copy number-based score for each sample-cell line pair.

For CTDPathSim1.0, the users should perform the above step with only expression-based score and DNA methylation-based score.
