## Supplementary material for "Finding the best cell lines across pan-cancer to use in pre-clinical research as a proxy for patient tumor samples considering immune cells, multi-omics, and cancer pathways": README

Running the pipeline: For running our pipeline, please use the CTDPathSim2.0 R package. 

For installation the package, use:

install.packages("remotes")
remotes::install_github("boseb/CTDPathSim2.0")

Check the vignette: browseVignettes("CTDPathSim2.0")

A copy of the vignette can also be found in the supplemental documents.

Running the evaluations: For evaluation of our method, the provided scripts in the supplemental documents can be used to generate different results in the figures and tables of the main manuscripts and supplemental tables and files. 


 
