## Supplementary material for "Finding the best cell lines across pan-cancer to use in pre-clinical research as a proxy for patient tumor samples considering immune cells, multi-omics, and cancer pathways": Code9_Table 6

## This R script is for five top-ranked tissue-specific cell lines for each cancer type in Table 6.
## Before running this script, we need to run "Code6_Table 5.R" for "cell_median_sim_based_rank.Rda" files.
## Author: Banabithi Bose
## Date: 04/01/2022
fileNames <-
  c(
    "BLCA",
    "BRCA",
    "DLBC",
    "ESCA",
    "GBM",
    "HNSC",
    "KIRC",
    "LAML",
    "LIHC",
    "LUAD",
    "LUSC",
    "OV",
    "PAAD",
    "SKCM",
    "STAD"
  )

high.rank.cells.in.cancer <- data.frame("", "", "", "", "", "")
colnames(high.rank.cells.in.cancer) <-
  c("cancer", "cell 1", "cell 2", "cell 3", "cell 4", "cell 5")

for (i in 1:length(fileNames1)) {
  f <- fileNames1[i]
  load(paste0(
    "~/EXTENDED_CTDPathSim/",
    f,
    "/cell_median_sim_based_rank.Rda"
  ))
  n <- nrow(df1) - 5
  high.rank.cells <-
    t(data.table(gsub("_.*", "", row.names(df1[df1$rank >= n, ])[1:5])))
  high.rank.cells.in.cancer1 <- cbind(f, high.rank.cells)
  colnames(high.rank.cells.in.cancer1) <-
    c("cancer", "cell 1", "cell 2", "cell 3", "cell 4", "cell 5")
  high.rank.cells.in.cancer <-
    rbind(high.rank.cells.in.cancer, high.rank.cells.in.cancer1)
  
}
high.rank.cells.in.cancer <-
  data.table(high.rank.cells.in.cancer[-1, ])
