## Supplementary material for "Finding the best cell lines across pan-cancer to use in pre-clinical research as a proxy for patient tumor samples considering immune cells, multi-omics, and cancer pathways": Code10_Supplemental Table S1

## This R script is for a list of the filtered drugs with the number of resistant and sensitive patients and cell lines
## provided in Supplemental Table S1.
## For running this script, we need "CCLE_GDSC_cellLine_response.xlsx"; "FINAL_CORRECTED_Gdisc_GDSC_DRUG_RESPONSE.Rda" and "_Drug_cdplot_Final4.rda" files under cancer folders.
## Author: Banabithi Bose
## Date: 01/01/2022

fileNames <-
  c(
    "ACC",
    "BLCA",
    "BRCA",
    "CESC",
    "DLBC",
    "ESCA",
    "GBM",
    "HNSC",
    "KIRC",
    "LGG",
    "LIHC",
    "LUAD",
    "LUSC",
    "MESO",
    "OV",
    "PAAD",
    "SKCM",
    "STAD",
    "THCA"
  )

## Reading cell lines' response data and getting response value
CCLE_cell_with_GDSC_drugs <-
  read_xlsx(path = "~/backup/EXTENDED_CTDPathSim/CCLE_GDSC_cellLine_response.xlsx")
CCLE_cell_with_GDSC_drugs$Response <- 0
CCLE_cell_with_GDSC_drugs$lnIC50 <-
  as.numeric(as.character(CCLE_cell_with_GDSC_drugs$lnIC50))
CCLE_cell_with_GDSC_drugs[CCLE_cell_with_GDSC_drugs$lnIC50 < (-2), 5] <-
  1
C1 <- CCLE_cell_with_GDSC_drugs[, c(2, 3, 5)]
colnames(C1) <- c("CellLine", "Drug", "CellLine_Response")

Cancer.Sample.CellLine <- data.frame("", "", "", "", "", "")
colnames(Cancer.Sample.CellLine) <-
  c(
    "Cancer.Type",
    "Drug",
    "No.of.Sensitive.Samples",
    "No.of.Resistant.Samples",
    "No.of.Sensitive.CellLines",
    "No.of.Resistant.CellLines"
  )

for (i in 1:19) {
  f <- str_sub(fileNames1[i], 1,-17)
  print(f)
  load(
    paste0(
      "~/backup/EXTENDED_CTDPathSim/",
      f,
      "/FINAL_CORRECTED_Gdisc_GDSC_DRUG_RESPONSE.Rda"
    )
  )
  S1 <-
    unique(FINAL_CORRECTED_Gdisc_GDSC_DRUG_RESPONSE[, c(3, 1, 6)])
  colnames(S1) <- c("Sample", "Drug", "Sample_Response")
  
  ## Getting the same drugs in TCGA samples and CCLE cell lines
  load(paste0(
    "~/backup/EXTENDED_CTDPathSim/",
    f,
    "/",
    f,
    "_Drug_cdplot_Final4.rda"
  ))
  Drug <- data.table(unique(as.character(M$drug)))
  colnames(Drug) <- "Drug"
  C2 <- merge(C1, Drug, by = "Drug")
  S2 <- merge(S1, Drug, by = "Drug")
  
  ## No. of senitive and resistant samples
  S <- S2[S2$Sample_Response == 1,-2, drop = F]
  if (nrow(S) == 0) {
    S = data.frame("", "")
    colnames(S) <- c("Drug", "Sensitive.Samples")
  }
  S.sens <- data.table(table(S$Drug))
  colnames(S.sens) <- c("Drug", "Sensitive.Samples")
  
  S <- S2[S2$Sample_Response == 0,-2, drop = F]
  if (nrow(S) == 0) {
    S = data.frame("", "")
    colnames(S) <- c("Drug", "Resistant.Samples")
  }
  S.res <- data.table(table(S$Drug))
  colnames(S.res) <- c("Drug", "Resistant.Samples")
  
  S.sens.res <- merge(S.sens, S.res, by = "Drug", all = T)
  
  ## No. of senitive and resistant cell lines
  C <- C2[C2$CellLine_Response == 1,-2, drop = F]
  if (nrow(C) == 0) {
    C = data.frame("", "")
    colnames(C) <- c("Drug", "Sensitive.CellLines")
  }
  C.sens <- data.table(table(C$Drug))
  colnames(C.sens) <- c("Drug", "Sensitive.CellLines")
  
  C <- C2[C2$CellLine_Response == 0,-2, drop = F]
  if (nrow(C) == 0) {
    C = data.frame("", "")
    colnames(C) <- c("Drug", "Resistant.CellLines")
  }
  C.res <- data.table(table(C$Drug))
  colnames(C.res) <- c("Drug", "Resistant.CellLines")
  
  ## Getting the number of sensitive and resistant sample and cell lines per cancer type per drug
  S.sens.res <- merge(S.sens, S.res, by = "Drug", all = T)
  C.sens.res <- merge(C.sens, C.res, by = "Drug", all = T)
  SC <- merge(S.sens.res, C.sens.res, by = "Drug", all = T)
  Cancer.Sample.CellLine1 <- data.table(cbind(f, SC))
  colnames(Cancer.Sample.CellLine1) <-
    c(
      "Cancer.Type",
      "Drug",
      "No.of.Sensitive.Samples",
      "No.of.Resistant.Samples",
      "No.of.Sensitive.CellLines",
      "No.of.Resistant.CellLines"
    )
  Cancer.Sample.CellLine <-
    rbind(Cancer.Sample.CellLine, Cancer.Sample.CellLine1)
}
Cancer.Sample.CellLine$Drug <-
  as.character(Cancer.Sample.CellLine$Drug)
Cancer.Sample.CellLine$No.of.Sensitive.Samples <-
  as.character(Cancer.Sample.CellLine$No.of.Sensitive.Samples)
Cancer.Sample.CellLine$No.of.Resistant.Samples <-
  as.character(Cancer.Sample.CellLine$No.of.Resistant.Samples)
Cancer.Sample.CellLine$No.of.Sensitive.CellLines <-
  as.character(Cancer.Sample.CellLine$No.of.Sensitive.CellLines)
Cancer.Sample.CellLine$No.of.Resistant.CellLines <-
  as.character(Cancer.Sample.CellLine$No.of.Resistant.CellLines)

Cancer.Sample.CellLine <-
  Cancer.Sample.CellLine[!Cancer.Sample.CellLine$Drug == "",]
Cancer.Sample.CellLine[is.na(Cancer.Sample.CellLine)] = 0

write.xlsx(Cancer.Sample.CellLine, file = "~/Supplemental Table S1.xlsx")
