## Supplementary material for "Finding the best cell lines across pan-cancer to use in pre-clinical research as a proxy for patient tumor samples considering immune cells, multi-omics, and cancer pathways": Code15_Supplemental Table S3

## This R script is for computing the classification matrices with precision, recall, and F1 scores for Supplemental Table S3
## Author: Banabithi Bose
## Date: 04/03/2022

## For classification of disease types
library(ggplot2)
######################
subtype_info <- read_xlsx(path = "~/Disease_subtype_annotation")
S1 <- subtype_info[, c(1, 2, 5, 6, 8)]

## Reading all patients and cell lines from 22 different cancer types with CTDPathSim2.0 computed score
P1 <-
  read_xlsx(path = "~/backup/EXTENDED_CTDPathSim/CTDPathSim2.0_patient_cellLine_score.xlsx", col_names = T)
colnames(P1) <- c("patient", "cellLine", "score")

## Getting the disease info for the patients
P1$patient <- str_sub(P1$patient, 1, 15)
P1.S1 <- merge(P1, S1, by.y = "sampleID", by.x = "patient")
P1.S1 <- P1.S1[, -4]
length(unique(P1.S1$patient))#7318
length(unique(P1.S1$disease))##19 different diseases among 7318 patients
P1.S2 <- P1.S1[, 1:3]
P1.S3 <- dcast(P1.S2, patient ~ cellLine, fun.aggregate = mean)
P1.S4 <- na.omit(P1.S3)
sample <- data.table(P1.S4$patient)
colnames(sample) <- "patient"
sample.disease <- unique(merge(sample, P1.S1, by = "patient")[, c(1, 6)])
colnames(sample.disease) <- c("patient", "disease")
length(unique(sample.disease$disease))#19
D <- P1.S4
x <- data.table(table(sample.disease$disease))
colnames(x) <- c("class", "frequency")
D$class <- as.factor(as.character(sample.disease$disease))
D <- merge(x, D, by = "class")
## Removing class labels having less than 10 tumor samples
D <- D[D$frequency >= 10, ]
library(caret)
D <- D[, -c(2:3)]
length(unique(D$class))#26 class
D$class <- gsub(" ", "_", D$class)

# create a list of 80% of the rows in the original dataset we can use for training
validation_index <- createDataPartition(D$class, p = 0.75, list = FALSE)
# select 20% of the data for validation
validation <- D[-validation_index, ]
Dtesttrain <- D[validation_index, ]
percentage <- prop.table(table(D$class)) * 100
cbind(freq = table(D$class), percentage = percentage)
control <-
  trainControl(method = "cv", number = 10)## Testing and traning are done by 10-fold cross validation
metric <- "Accuracy"
Dtesttrain$class <- as.factor(as.character(Dtesttrain$class))
fit.rf <-
  train(
    class ~ .,
    data = Dtesttrain,
    method = "rf",
    metric = metric,
    trControl = control
  )
results.summary <- resamples(list(fit.rf, fit.rf))
summary(results.summary)
# summarize the best model
print(fit.rf)
R.ran.forest <- as.data.frame(fit.rf[["byClass"]][, "F1"])

validation$class <- as.factor(as.character(validation$class))
predictions <- predict(fit.rf, validation)
result <-
  confusionMatrix(predictions, as.factor(as.character(validation$class)))
## Confusion matrix
result.confusion.matrix <- as.matrix(result)
# View confusion matrix overall with F1 score,precision-recall etc.
result.confusion.matrix1 <- as.data.frame(result$byClass)
mean(result.confusion.matrix1$F1)# [1] 0.96249

cm <- fit.rf[["finalModel"]][["confusion"]][, -20]
n = sum(cm) # number of instances
nc = nrow(cm) # number of classes
diag = diag(cm) # number of correctly classified instances per class
rowsums = apply(cm, 1, sum) # number of instances per class
colsums = apply(cm, 2, sum) # number of predictions per class
p = rowsums / n # distribution of instances over the actual classes
q = colsums / n # distribution of instances over the predicted classes
accuracy = sum(diag) / n
precision = diag / colsums
recall = diag / rowsums
f1 = 2 * precision * recall / (precision + recall)
F1.per.class <- data.frame(precision, recall, f1)
mean(F1.per.class$f1)#0.9612072


## For classification of subtypes
sample.subtype <- unique(merge(sample, P1.S1, by = "patient")[, c(1, 5)])
length(unique(sample.subtype$subtype))#29
D <- P1.S4
x <- data.table(table(sample.subtype$subtype))
colnames(x) <- c("class", "frequency")
D$class <- as.factor(as.character(sample.subtype$subtype))
D <- merge(x, D, by = "class")
## Removing class labels having less than 10 tumor samples
D <- D[D$frequency >= 10, ]
length(unique(D$class))#26

library(caret)
D <- D[, -c(2:3)]
length(unique(D$class))#26 class
D$class <- gsub(" ", "_", D$class)
D$class <- gsub("-", "_", D$class)
D[D$class == "normal", 1] <- "normal_sample"
D[D$class == "melanocytic", 1] <- "melanocytic_tumor"
# create a list of 75% of the rows in the original dataset we can use for training
validation_index <- createDataPartition(D$class, p = 0.75, list = FALSE)
# select 25% of the data for validation
validation <- D[-validation_index, ]
#length(unique(validation$class)) #26
# use the remaining 80% of data to training and testing the models
Dtesttrain <- D[validation_index, ]
#length(unique(Dtesttrain$class))#26
# summarize the class distribution
percentage <- prop.table(table(D$class)) * 100
cbind(freq = table(D$class), percentage = percentage)
control <- trainControl(method = "cv", number = 10)
metric <- "Accuracy"
Dtesttrain$class <- as.factor(as.character(Dtesttrain$class))
fit.rf <-
  train(
    class ~ .,
    data = Dtesttrain,
    method = "rf",
    metric = metric,
    trControl = control
  )
validation$class <- as.factor(as.character(validation$class))
predictions <- predict(fit.rf, validation)
result <-
  confusionMatrix(predictions, as.factor(as.character(validation$class)))
result.confusion.matrix <- as.matrix(result)
# View confusion matrix overall
result.confusion.matrix1 <- as.data.frame(result$byClass)
mean(na.omit(result.confusion.matrix1$F1))#[1] 0.8773842


cm <- as.data.frame(fit.rf[["finalModel"]][["confusion"]])
save(cm, file = "~/EXTENDED_CTDPathSim/RCODES/FIGURES/subtype.RF.cm.rda")
cm <- cm[, -27]
n = sum(cm) # number of instances
nc = nrow(cm) # number of classes
diag = diag(cm) # number of correctly classified instances per class ##THIS SHOWS her2 wasn't correctly predicted
##her2 samples were identified as luminal A mostly also basal
##lumi A and basal are her2 -ve but in https://www.ncbi.nlm.nih.gov/pmc/articles/PMC6828840/
##A clear example of this is the HER2-positive breast cancer. In these tumors, we can find all of the transcriptional
#subtypes of breast cancer, even the basal like or luminal A subtypes.
## **HER2-enriched subtype can have a distinctive transcriptional landscape independent of HER2 amplification.
#In this review, we present an extensive revision about the last **highlights

##melanocytic as transitory mostly
##normal samples as luminal A mostly
#The transitory and melanocytic subtypes are a refinement
#of the previously reported differentiated proliferative phenotype
#https://www.ncbi.nlm.nih.gov/pmc/articles/PMC5953834/
rowsums = apply(cm, 1, sum) # number of instances per class
colsums = apply(cm, 2, sum) # number of predictions per class
p = rowsums / n # distribution of instances over the actual classes
q = colsums / n # distribution of instances over the predicted classes
accuracy = sum(diag) / n
precision = diag / colsums
recall = diag / rowsums
f1 = 2 * precision * recall / (precision + recall)
F1.per.class <- data.frame(precision, recall, f1)
mean(F1.per.class$f1)#0.9612072
